# Recurrent erosion of *COA1/MITRAC15* demonstrates gene dispensability in oxidative phosphorylation

**DOI:** 10.1101/2021.06.09.447812

**Authors:** Sagar Sharad Shinde, Sandhya Sharma, Lokdeep Teekas, Ashutosh Sharma, Nagarjun Vijay

**Affiliations:** Computational Evolutionary Genomics Lab, Department of Biological Sciences, IISER Bhopal, Bhauri, Madhya Pradesh, India

**Keywords:** Cytochrome C Oxidase Assembly Factor 1, *COA1*, *MITRAC15*, Chicken, gene loss, rodent

## Abstract

Skeletal muscle fibers rely upon either oxidative phosphorylation or glycolytic pathway to achieve muscular contractions that power mechanical movements. Species with energy-intensive adaptive traits that require sudden bursts of energy have a greater dependency on fibers that use the glycolytic pathway. Glycolytic fibers have decreased reliance on OXPHOS and lower mitochondrial content compared to oxidative fibers. Hence, we hypothesized that adaptive gene loss might have occurred within the OXPHOS pathway in lineages that largely depend on glycolytic fibers. The protein encoded by the *COA1/MITRAC15* gene with conserved orthologs found in budding yeast to humans promotes mitochondrial translation. We show that gene disrupting mutations have accumulated within the *COA1/MITRAC15* gene in the cheetah, several species of galliforms, and rodents. The genomic region containing *COA1/MITRAC15* is a well-established evolutionary breakpoint region in mammals. Careful inspection of genome assemblies of closely related species of rodents and marsupials suggests two independent *COA1/MITRAC15* gene loss events co-occurring with chromosomal rearrangements. Besides recurrent gene loss events, we document changes in *COA1/MITRAC15* exon structure in primates and felids. The detailed evolutionary history presented in this study reveals the intricate link between skeletal muscle fiber composition and dispensability of the chaperone-like role of the *COA1/MITRAC15* gene.

## 1. Introduction

Skeletal muscles control numerous locomotor functions in vertebrates (Weeks, 1989). The hundreds of different muscles in the body consist of highly organized heterogeneous bundles of fibers. These muscle fibers are classified based on contractile properties, power source, and myosin component into type-1, 2A, 2B, and 2X (Talbot and Maves, 2016). Muscles with type-1 and type-2A fibers rely on the oxidative phosphorylation (OXPHOS) pathway, the primary source of ATP needed for locomotion and other energy-intensive tasks (Shen et al., 2010). The energy releasing electron transport chain (ETC) coupled with the energy-requiring chemiosmosis is known as (OXPHOS) (Hatefi, 1985; Mitchell, 1961). A chain of mitochondrial inner membrane-embedded proteins encoded by both mitochondrial and nuclear genes form four large complexes that transport electrons through redox reactions. The energy released during these reactions results in a proton gradient, which uses a fifth membrane-embedded complex to generate ATP through chemiosmosis. Optimization of crucial steps in the OXPHOS pathway leads to improved locomotor performance (Conley, 2016). Origin of novel energetically demanding phenotypes has been possible through adaptations in the OXPHOS pathway (Doan et al., 2004; Garvin et al., 2015; Wu et al., 2000; Zhang and Broughton, 2015). Multiple genes of the OXPHOS pathway are under positive selection in mammalian species with high energy demanding adaptations such as powered flight in bats (Shen et al., 2010), survival of polar bears in cold Arctic environment (Welch et al., 2014), high altitude adaptation in yak (Qiu et al., 2012), hypoxia tolerance in cetaceans (Tian et al., 2018), ecotype specific divergence in killer whales (Foote et al., 2011) and evolution of large brains in anthropoid primates (Grossman et al., 2004).

The loss of energetically demanding phenotypes reduces the strength of purifying selection acting on the OXPHOS pathway. For instance, the domestication of dogs (Björnerfeldt et al., 2006) and degeneration of locomotor abilities in birds (Shen et al., 2009) resulted in relaxed selective constraint on the OXPHOS pathway proteins. Among other carnivores and rodents, the great diversity of functionally important locomotor habits have variations in energy requirements and corresponding differences in the magnitude of purifying selection (Samuels and Van Valkenburgh, 2008; Taylor, 1989). Even within the same species, mitochondrially encoded protein components of the OXPHOS pathway are under stronger purifying selection than those protein components encoded by the nuclear genome (Popadin et al., 2013). These differing purifying selection levels are due to gene expression level differences between nuclear and mitochondrial OXPHOS genes (Nabholz et al., 2013). Despite the movement of most genes from the ancestral mitochondria to the nucleus in eukaryotes, a separate mitochondrial organelle is well conserved with scarce exceptions (Karnkowska et al., 2016; Sloan et al., 2018). Turnover in the content of mitochondrial protein complexes has mainly occurred before the emergence of eukaryotes with few gene gain/loss events reported in vertebrates (Adams and Palmer, 2003; Cardol, 2011; Gabaldón et al., 2005; Gabaldón and Huynen, 2007; Huynen et al., 2013; van Esveld and Huynen, 2018). However, lineage-specific gene loss from the mitochondria has occurred in nonbilaterian organisms (Lavrov and Pett, 2016), other metazoan lineages (Gissi et al., 2008), and plants (Depamphilis et al., 1997; Palmer et al., 2000). The duplication of mitochondrial genes in bird lineages followed by gene loss and genomic rearrangement events is relatively unique (Akiyama et al., 2017; Mackiewicz et al., 2019; San Mauro et al., 2006; Urantówka et al., 2020).

The proton gradient established by the ETC also powers the generation of heat in mammalian Non-Shivering Thermogenesis (NST) (Nedergaard et al., 2001). Thermogenin or uncoupling protein 1 (*UCP1*) expressed in the inner mitochondrial membrane facilitates the regulated leakage of protons to generate heat in brown adipose tissue (Krauss et al., 2005). The *UCP1* gene is absent in all birds (Newman et al., 2013) and some mammals (Emre et al., 2007; Mcgaugh and Schwartz, 2017) despite its presence in fish (Jastroch et al., 2005), amphibians (Hughes et al., 2009), and marsupials (Polymeropoulos et al., 2012). The integration of *UCP1* in the thermogenic pathway is considered a eutherian-mammal-specific adaptation unrelated to its ancestral innate immune functions (Jastroch, 2017). The exceptional repeated loss of this mitochondrial membrane protein in vertebrate lineages appears to result from its changing functional roles (Gaudry et al., 2017; Mcgaugh and Schwartz, 2017). In contrast to *UCP1*, most OXPHOS pathway genes are highly conserved, and defective protein components generally result in clinical phenotypes (Hock et al., 2020). The proteins *TMEM186* and *COA1/MITRAC15* are chaperones interacting with the Mitochondrial Complex I Assembly (MCIA) complex, and defects in these genes do not result in any clinical phenotypes (Hock et al., 2020; Signes and Fernandez-Vizarra, 2018).

Functional studies have implicated a role for *COA1/MITRAC15* in promoting mitochondrial translation and complex I and IV biogenesis (Wang et al., 2020). However, overexpression of other genes easily compensates for the mild effect of *COA1/MITRAC15* gene knockout (Hess et al., 2009; Pierrel et al., 2007). Notably, the *COA1/MITRAC15* gene was also identified as a positively selected gene in a genome-wide screen in primates (Van Der Lee et al., 2017) and suggests that despite its mild phenotype, *COA1/MITRAC15* can contribute to fitness increases through its role as a chaperone. *COA1/MITRAC15* resembles *TIMM21*, a subunit of the *TIM23* complex (Mick et al., 2012). Such *TIMM21* gene duplicates interacting with the mitochondrial import apparatus and respiratory chain complexes occur in Arabidopsis (Murcha et al., 2014). Diversification of the mitochondrial import system has benefitted from gene duplication events that have contributed new members to the Translocase of the Inner Membrane (TIM) and Translocase of the Outer Membrane (TOM) protein complexes (Fukasawa et al., 2017). Hence, it is plausible that *COA1/MITRAC15* results from a duplication of the *TIMM21* gene followed by divergence.

Divergence of conventional or class-2 myosin genes after duplication has led to the diversification of the *MYH* gene family (Moore et al., 1993; Weiss and Leinwand, 1996). These myosin genes have distinct functions defined by their contractile properties and ATPase activity (Resnicow et al., 2010). While *MYH7* and *MYH2* expressing fibers rely upon the OXPHOS pathway, *MYH1* and *MYH4* expressing fibers are dependent on the glycolytic pathway. The protein encoded by the *MYH7* gene occurs in both cardiac muscles and the slow-contracting type-1 fibers (Schiaffino and Reggiani, 2011). However, the *MYH* genes expressed in type-2 fibers are restricted mainly to skeletal muscles. The fast-contracting type-2 fibers power explosive movements like jumping and sprinting. Such rapid movements form an essential component of hunting strategies used by terrestrial predators and the escape strategy of the prey (Kohn, 2014; J. W. Wilson et al., 2013). Felids, small-bodied rodents, marsupials, certain cervids, and galliform birds have exceptional adaptations for rapid locomotion.

The world’s fastest mammal, the cheetah (*Acinonyx jubatus*), epitomizes the relevance of speed and acceleration (A. M. Wilson et al., 2013). In general, felids are adept at sprinting and can accelerate more rapidly than canids but cannot sustain it for a prolonged period (Bailey et al., 2013). The predominance of type-2X fibers in felid species provides the ability to achieve rapid acceleration (Hyatt et al., 2010; Kohn et al., 2011; Williams et al., 1997). Compared to canids, felids have a greater reliance on glycolytic fibers. Glycolytic fibers have decreased reliance on OXPHOS and lower mitochondrial content than oxidative fibers (Mishra et al., 2015; Picard et al., 2012). Hence, the OXPHOS pathway might be under stronger selective constraint in canids than felids. Like these predators, prey species like antelopes are fast sprinters but have the added advantage of resistance to fatigue. The high speed of these species relies on type-2X fibers with high glycolytic capacities, and the added resistance to fatigue is possible due to the remarkable oxidative ability of these fibers (Curry et al., 2012). The use of both glycolytic and oxidative pathways suggests the OXPHOS pathway in these antelope species and other cervids would be under strong purifying selection.

Despite drastic variation in body size within mammals, the relative speed of locomotion is thought to be largely independent of body mass, at least in small mammals (Iriarte-Díaz, 2002). The higher relative speed of small mammals results from faster constriction made possible by the higher proportion of fast fibers (mostly 2X and 2B) in each muscle (Schiaffino and Reggiani, 2011). For instance, rodent limb muscles are known to have more abundant type 2B fibers compared to larger mammals (including humans, which have no type 2B fibers in the limb muscles) (Kohn, 2014; Kohn and Myburgh, 2007). Marsupial species also have high relative speeds and possess muscle fibers equivalent to eutherian mammals (Zhong et al., 2001). The smaller marsupial species have type-2B and 2X muscle fibers in several important muscles (Zhong et al., 2008). The higher proportion of fast glycolytic fibers in rodents and marsupials potentially results in relaxed selection on the OXPHOS pathway genes in these species.

The ability to fly is a distinctive feature of birds except for lineages that have become entirely flightless or retain only a limited flying capacity (Harshman et al., 2008; Pan et al., 2019; Sackton et al., 2019; Sayol et al., 2020). The large amount of energy required for flight has necessitated a high metabolic rate in birds (Holmes and Austad, 1995). Increased ATP generation fulfills these energy demands through metabolic adaptations in the OXPHOS pathway (Das, 2006). The set of flight muscles possessed by a bird species determine several aspects of flight performance and strongly influences life history and ecology (DuBay et al., 2020). Avian flight is possible through a combination of flight muscles that consist of white (fast glycolytic), intermediate/red-pink (fast oxidative), and red (slow oxidative) muscle fibers (Barnard et al., 1982; Butler, 2016; Ogata and Yamasaki, 1997). Birds with strong flight abilities, such as long-distant migrants and small passerines, contain mostly fast oxidative fibers (Welch and Altshuler, 2009). In contrast to this, Galliformes contain mostly glycolytic fibers that only allow short bursts of activity (Dial, 2003). Hence, the OXPHOS pathway is under stronger selective constraint in non-Galliform bird species than Galliform birds due to the functional specialization of mitochondria to different muscle fibers (Picard et al., 2012).

This study evaluates whether the protein encoded by the *COA1/MITRAC15* gene, a mitochondrial complex I translation factor with a chaperone-like role, is dispensable when the OXPHOS pathway is under relaxed selective constraints. We hypothesized that the OXPHOS pathway might have experienced reduced purifying selection in felids, rodents, marsupials, and galliform birds based on increased glycolytic muscle fibers in these species. Duplicate copies or alternative metabolic pathways compensate for gene function and decide gene dispensability (Gu et al., 2003). Hence, to evaluate our hypothesis, we aim to (1) investigate whether *COA1/MITRAC15* has any homologs that could compensate its function, (2) screen the genomes of vertebrate species to identify and track the evolutionary history of *COA1/MITRAC15* orthologs, (3) identify evidence of gene disruptive changes within the *COA1/MITRAC15* locus and (4) reconstruct the sequence of events associated with the potential erosion of the *COA1/MITRAC15* locus due to chromosomal rearrangement events at the evolutionary breakpoint region spanning the *COA1/MITRAC15* gene. We extensively screened publicly available genomes and transcriptomes of more than 200 vertebrate species to establish recurrent loss of the widely conserved *COA1/MITRAC15* gene.

## 2. Materials and methods

### 2.1 Finding homologs of *COA1/MITRAC15*

The amino acid sequence of the human *COA1/MITRAC15* gene was used as a query in PSI-BLAST (Altschul et al., 1997) against the nr database with eight iterations to identify homologs. Similarly, the human *COA1/MITRAC15* amino acid sequence was the query in the program HHblits of HHsuite (Remmert et al., 2012; Steinegger et al., 2019) with the flags “-e 1e-3 -n 8 -p 20 -Z 5000 -z 1 -b 1 -B 5000 -d UniRef30_2020_06”. The output from HHblits was used as input to the CLANS program (Frickey and Lupas, 2004) with an e-value cut-off of 1e-4 to cluster the blast hits using the MPI Bioinformatics Toolkit (Gabler et al., 2020; Zimmermann et al., 2018). We ran the CLANS java application for more than 50,000 rounds on the webserver output to ensure stable clusters. Manually inspection of gene annotations allowed identification of each of the groups. Subsequently, we performed the HHblits search again with different settings such as “-glob” to perform global alignments and “-loc” to conduct local alignments. The PFAM database was the alternative to the Uniclust 30 database. Manually curated multiple sequence alignment of *COA1/MITRAC15* open reading frames from 24 primate species was also separately used to query for better sensitivity. The protein *TIMM21* provides a consistent hit with different search settings and databases.

To further verify whether the database matches are homologous, we evaluated the biological function, secondary structure similarity, relationship among top hits, and occurrence of conserved motifs. To obtain secondary structure predictions for the proteins *COA1/MITRAC15* and *TIMM21*, we used the PROTEUS2 webserver (Montgomerie et al., 2008). The HeliQuest webserver (Gautier et al., 2008) provided each predicted helix’s physicochemical properties and amino acid compositions. While the three-dimensional (3-D) structure of the *COA1/MITRAC15* protein is not available yet, multiple structures of the *TIMM21* protein are available in the Protein Data Bank (PDB). It is possible to use comparative/homology modeling to predict the 3-D structure based on the protein structure of a related protein (Webb and Sali, 2016). Hence, we used the comparative modeling approach implemented in Modeller (v10.0) software to model the structure of *COA1/MITRAC15* based on the homologous structures available in PDB. The Phyre2 (Kelley et al., 2015) and Expasy Swiss-Model (Waterhouse et al., 2018) webservers also predicted homologous 3-D structures of *COA1/MITRAC15*. All the top hits were from 3-D structures of the IMS (Inter Membrane Space) domain of *TIMM21* protein. The IMS domain of *TIMM21,* whose 3-D structures are available on PDB, contains only the part of the protein that occurs after the membrane-spanning helix. To model the structure of *COA1/MITRAC15* using these existing 3-D structures, we used the *COA1/MITRAC15* amino acid sequence that occurs after the membrane-spanning domain. We visualized the structure of *TIMM21* and the predicted *COA1/MITRAC15* structure using (UCSF Chimera v1.15) ChimeraX (Pettersen et al., 2021).

### 2.2 Validation of *COA1/MITRAC15* annotation

Despite being a fast-evolving gene, orthologs of *COA1/MITRAC15* can be identified based on gene synteny and sequence identity. However, identifying *COA1/MITRAC15* orthologs between distantly related species is challenging (Szklarczyk et al., 2012). We screened the genome assemblies and annotations available on NCBI and Ensembl for *COA1* (C7orf44 or *MITRAC15*) protein-coding transcripts. The *COA1/MITRAC15* gene orthologs have been annotated in almost 300 vertebrate species (see **Supplementary Table S1**). However, the number of exons and the length of the ORF is highly variable between species. We validated the annotation of the *COA1/MITRAC15* gene relying upon gene synteny in the genomic vicinity of the *COA1/MITRAC15* gene, multiple sequence alignments, and RNA-seq data. Annotation across most species endorses the existence of four coding exons that produce a ∼130 to 140 amino acid (aa) protein. The *COA1/MITRAC15* annotation in the human genome (see **Supplementary Figure S1**) has multiple isoforms with seven exons. The additional three exons annotated in the human genome upstream from the widely conserved four exons need further investigation. Bird species such as *Nipponia nippon*, *Cuculus canorus*, *Pterocles gutturalis*, *Gavia stellate*, *Buceros rhinoceros silvestris*, *Anser cygnoides domesticus*, *Anas platyrhynchos* (corrected in XM_027451320.2), and *Fulmarus glacialis* have annotation for a fifth exon upstream from the widely conserved four exons. Annotation for multiple isoforms of the *COA1/MITRAC15* gene also exists in *Athene cunicularia*, *Tyto alba*, *Calidris pugnax*, *Serinus canaria*, *Corvus moneduloides*, *Corvus brachyrhynchos*, *Egretta garzetta*, *Aquila chrysaetos*, *Pipra filicauda*, *Corvus cornix*, *Cygnus atratus,* and *Parus major*. We examined RNA-seq datasets of multiple species to evaluate the expression of the isoforms. RNA-seq data in *Colius striatus* and *Eurypyga helias* (which had partial sequences annotated) allowed reconstruction of full-length ORFs. In addition to bird genomes, the *COA1/MITRAC15* gene ortholog is annotated in lizards (*Zootoca vivipara*, *Podarcis muralis*, *Lacerta agilis*, *Anolis carolinensis*, *Gekko japonicus*, *Thamnophis sirtalis*, *Pantherophis guttatus*, *Notechis scutatus*, *Pseudonaja textilis* and *Python bivittatus*), turtles (*Trachemys scripta elegans*, *Chelonia mydas*, *Chelonoidis abingdonii*, *Chrysemys picta*, *Gopherus evgoodei* and *Pelodiscus sinensis*), alligators (*Gavialis gangeticus*, *Alligator sinensis*, *Alligator mississippiensis* and *Crocodylus porosus*), Even-toed ungulates (*Bos taurus*, *Sus scrofa*, *Odocoileus virginianus texanus*, *Bison bison bison*, *Bos indicus x Bos taurus*, *Bos mutus*, *Bubalus bubalis*, *Capra hircus*, *Ovis aries*, *Vicugna pacos*, *Camelus ferus*, *Camelus bactrianus*, *Camelus dromedarius*, *Neophocaena asiaeorientalis asiaeorientalis*, *Balaenoptera acutorostrata scammoni*, *Lipotes vexillifer*, *Lagenorhynchus obliquidens*, *Globicephala melas*, *Orcinus orca*, *Tursiops truncatus*, *Phocoena sinus*, *Monodon monoceros*, *Delphinapterus leucas*, *Physeter catodon* and *Balaenoptera musculus*), Odd-toed ungulates (*Equus caballus*, *Equus asinus*, *Equus przewalskii* and *Ceratotherium simum simum*), Pangolins (Manis pentadactyla and Manis javanica), *Galeopterus variegatus*, *Tupaia chinensis* and Primates (*Homo sapiens*, *Macaca mulatta*, *Pan troglodytes*, *Chlorocebus sabaeus*, *Callithrix jacchus*, *Colobus angolensis palliatus*, *Cercocebus atys*, *Macaca fascicularis*, *Macaca nemestrina*, *Papio anubis*, *Theropithecus gelada*, *Mandrillus leucophaeus*, *Trachypithecus francoisi*, *Rhinopithecus bieti*, *Rhinopithecus roxellana*, *Piliocolobus tephrosceles*, *Gorilla gorilla*, *Pan paniscus*, *Pongo abelii*, *Nomascus leucogenys*, *Hylobates moloch*, *Saimiri boliviensis*, *Sapajus apella*, *Cebus imitator*, *Aotus nancymaae*, *Carlito syrichta*, *Propithecus coquereli*, *Microcebus murinus* and *Otolemur garnettii*).

We screened the synteny pattern of the candidate *COA1/MITRAC15* gene in Galliformes and Anseriformes using five upstream genes (*STK17A*, *HECW1*, *TNS3*, *PSMA2*, *MRPL32*) and the five downstream genes (*BLVRA*, *VOPP1*, *LANCL2*, *EGFR*, *SEC61G*). The chicken (*Gallus gallus*) has a chromosome level assembly, and the gene occurs on Chromosome 2, and its region is syntenic with human (*Homo sapiens*) chromosome 2 (**Supplementary Figure S2-S3**). The gene synteny is mostly conserved in these species and is present on the same scaffold/chromosome. The blast search of the genome using the query gene sequence of closely related species identified genes missing in the annotation. *Anas platyrhynchos* has chromosome level assembly with the same gene order as *Gallus gallus* (**Supplementary Figure S4**). *Anser cygnoides* and *Anseranas semipalmata* also contain this conserved gene order. *Anas platyrhynchos*, *Numida meleagris*, *Coturnix japonica*, *Meleagris gallopavo* show syntenic blocks aligning with the human chromosome 7 (**Supplementary Figure S5-S8**). Synteny-based verification was done clade-wise in birds (see **Supplementary Table S2**), rodents (**Supplementary Table S3**), carnivores (**Supplementary Table S4**), and primates (**Supplementary Table S5**). Gene order and synteny relationships for representative species from each of the clades are in **Supplementary Figure S9-S230**.

Vertebrate species have a conserved *COA1/MITRAC15* gene intron/exon organizational structure. However, two lineages (primates and carnivores) with evidence of intron/exon organization changes have also had *COA1/MITRAC15* gene duplication events. To ensure that the observed differences were not a result of incorrect annotation, alignment artifacts, or duplicated copies, we compared the *COA1/MITRAC15* gene organization across diverse vertebrate species. Subsequently, we validated the annotations from NCBI and Ensembl using RNA-seq datasets. Sequencing read haplotypes from the functional and pseudogenised copy can be distinguished as their sequences have diverged.

### 2.3 Verification of *COA1/MITRAC15* gene disrupting changes in raw read data

We used a previously published 5-pass strategy to verify gene loss events (S. Sharma et al., 2020). Briefly, to verify the correctness of the genome assembly nucleotide sequence, we used the *COA1/MITRAC15* gene sequence of multiple species as a query for a blastn search of the raw short-read database. The details of short-read datasets (both DNA and RNA) used to validate gene sequence are in **Supplementary Table S6**. Manual inspection of the blast search results ensured concordance between gene sequence and raw read data. All the blast output files are in **Supplementary File S1**. In the chicken genome, we also verified the correctness of genome assembly in the vicinity of the *COA1/MITRAC15* gene by evaluating Pacbio long-read data (see **Supplementary Figure S231-S234**).

### 2.4 Assessing the transcriptional status of *COA1/MITRAC15*

We analyzed transcriptomic datasets for evidence of transcription of *COA1/MITRAC15* locus. The RNA-seq reads were mapped to the genome assemblies using the STAR read mapper (Dobin et al., 2013). We visualized the resulting bam files using the IGV browser (Robinson et al., 2011; Thorvaldsdottir et al., 2013). For consistent representation across tissues and species, we used three different views: (1) Positions of all four exons of *COA1/MITRAC15* identified using blast search are shown as a bed record below the RNA-seq bam files, (2) Zoomed-in views of each of the four exons are presented in four panels within a single screenshot and (3) Zoomed-in view of the first and last exons of *COA1/MITRAC15* are shown along with the adjacent genes on both sides. The adjacent genes in the IGV screenshot act as positive controls.

No evidence for transcription of *COA1/MITRAC15* gene in chicken exists in the RNA-seq data from 23 tissues consisting of blood, bone marrow, breast muscle, bursa, cerebellum, cerebrum, comb, eye, fascia, gallbladder, gizzard, gonad, heart, immature egg, kidney, liver, lung, mature egg, pancreas, shank, skin, spleen, uterus (**Supplementary Figure S235-S304**). Among other Galliformes species, we found no evidence for expression of the *COA1/MITRAC15* gene. (The spleen and gonad of the peacock, the skin of golden pheasant, gonad, spleen, brain, muscle, liver, and heart of ring-necked pheasant, bursa, gonad spleen, blood and uterus of helmeted guineafowl, breast muscle, gonad, spleen, brain, liver, heart, and bursa of turkey, kidney, liver, muscle, lung, and heart of Japanese quail, the blood of *Colinus virginianus* and blood of *Syrmaticus Mikado*, see **Supplementary Figure S305-S373**). The only Galliform species to have a transcribed *COA1/MITRAC15* gene was *Alectura lathami* (blood tissue: **Supplementary Figure S374-S376**).

In contrast to Galliformes, the *COA1/MITRAC15* gene is intact in Anseriformes species. However, the *COA1/MITRAC15* gene annotation in duck (*Anas platyrhynchos platyrhynchos*) contains two isoforms. The more extended isoform codes for a 265 amino acid protein and consists of five exons. The shorter isoform (139 amino acid) is orthologous to the Galliform ORF. Upon closer inspection of the first exon, only 24 of the 372 bases have RNA-seq read support **(Supplementary Figure S377)**. Hence, this additional exon might be an annotation artifact or part of the untranslated region. The last four annotated exons, which correspond to the intact 139 amino acid encoding sequence, were found to be robustly expressed in the gonad, spleen, liver, brain, and skin (**Supplementary figure S378-S385**). A similar annotation of the fifth exon in *Anser cygnoides domesticus* appears to be an artifact. The gonad, liver, and spleen express the last four exons (see **Supplementary Figure S386-S392**). The RNA-seq data from blood tissue for magpie goose (*Anseranas semipalmata*) and southern screamer (*Chauna torquata*) also supported the transcription of the *COA1/MITRAC15* gene (**Supplementary Figure S393-S396**).

Having verified the expression of the *COA1/MITRAC15* gene in multiple Anseriformes species, we screened additional bird RNA-seq datasets to evaluate the transcriptional activity of the intact ORF found in these species. Many other bird genomes have annotations for multiple isoforms of the *COA1*/*MITRAC15* gene, like the duck genomes. These isoforms range in length from 136 to 265 amino acids and 4 to 7 exons. Based on careful examination of multiple RNA-seq datasets across several closely related species and sequence homology, we found that in most cases, the four-exon transcript coding for a 139 amino acid protein was the only correct annotation. However, in some rare cases, additional exons have robust expression and require further investigation. In the Corvidae group, annotation exists for transcripts of lengths 170 and 139 aa. The first exon of the longer transcript lacked expression.

In comparison, all four transcripts of the shorter transcript are present in the blood tissue of western Jackdaw (*Corvus monedula*) as well as gonad, brain, spleen, and liver of hooded crow (*Corvus cornix*) **(Supplementary Figure S397-S402)**. The common canary (*Serinus canaria*) has three transcripts with 177, 154, and 139 aa **(Supplementary Figure S403-S404)**. We checked the expression using liver and skin tissue and found support for all three transcripts. However, upon closer inspection, the transcript with 139 aa was strongly expressed, and the other two transcripts are potentially artifacts. Great tit (*Parus major*) has two transcripts of lengths 169, 139 aa. While the kidney and liver express both transcripts, the first exon has feeble expression and appears artefactual **(Supplementary Figure S405-S406)**.

The golden eagle (*Aquila chrysaetos*) has four annotated transcripts with lengths of 219, 180, 159, and 139 aa. Transcript of 219 aa length contains six exons, transcripts of length 180 aa, and 159 aa have five exons, and 139 aa transcript contains four exons. We found that exon 1 showed negligible expression, and exons 2 to 6 have high expression levels. However, exon 1 and 2 both have an in-frame stop codon **(Supplementary Figure S407-S411)**. Hence, we consider that the 139 aa long transcript expressed in the liver and muscle is correct. Red-throated loon (*Gavia stellata*) has a single five exon transcript of length 155 aa annotated. We discovered a lack of expression in the first exon compared to the last four exons that are orthologous to the transcript of length 139 aa **(Supplementary Figure S412-S413)**.

The ruff (*Calidris pugnax*) genome annotates three transcripts with lengths of 233, 229, and 139 aa. Transcript one and two contain seven exons each, and the third transcript contains four exons. Exons 1 and 2 lack expression in the first two transcripts, and the third exon did not have any start codon explaining the transcript. The last four exons have transcripts and are orthologous to other species’ *COA1*/*MITRAC15* gene **(Supplementary Figure S414-S418)**. In the little egret (*Egretta garzetta*), transcripts of lengths 212 and 203 are annotated and contain five exons. We found evidence of expression of *COA1/MITRAC15* in blood tissue **(Supplementary Figure S419-S420)**. Although the first exon has a lower level of expression than the last four exons, the consistent occurrence of the fifth exon across many species suggests it might be part of the untranslated region. We annotated and verified the expression of *COA1/MITRAC15* in *Phalacrocorax carbo, Phaethon lepturus, Opisthocomus hoazin, Leptosomus discolor* (**Supplementary Figure S421-S428)**. *Eurypyga helias* has an unverified transcript length of 121 aa. Hence, we screened the genome and RNA-seq data and found its transcript length is 139 aa **(Supplementary Figure S429-S431)**. We verified the *COA1*/*MITRAC15* gene expression using RNA-seq data in *Strigops habroptilus* as it had less than 100 percent RNA-seq coverage **(Supplementary Figure S432-S433)**. We also examined the RNA-seq data from few other bird species to verify the *COA1/MITRAC15* gene (see **Supplementary Figure S434-S481**). Bird species share this conserved gene order (**Supplementary Figure S482**). The Anolis lizard (*Anolis carolinensis*) liver also expresses the *COA1/MITRAC15* gene (**Supplementary Figure S483-S485**).

RNA-seq datasets from the European rabbit (*Oryctolagus cuniculus*) heart and liver showed no evidence of transcription of *COA1/MITRAC15* (see **Supplementary Figure S486-S489**). In contrast to the rabbit, intact *COA1/MITRAC15* gene is present in the Royle’s pika (*Ochotona roylei*) and Daurian pika (*Ochotona dauurica*) with blood RNA-seq datasets showing robust expression (see **Supplementary Figure S490**). Screening of RNA-seq datasets from the root ganglion, spinal cord, ovary, liver, spleen, and testis in the naked mole-rat (*Heterocephalus glaber*) revealed no transcription of *COA1/MITRAC15* locus (see **Supplementary Figure S491**). The closely related Damaraland mole-rat (*Fukomys damarensis*) has robust *COA1/MITRAC15* expression in the brain, liver, and testis (see **Supplementary Figure S492-S497**). The Brazilian guinea pig (*Cavia aperea*), the guinea pig (*Cavia porcellus*), and the long-tailed chinchilla (*Chinchilla lanigera*) were all found to express the *COA1/MITRAC15* gene robustly (see **Supplementary Figure S498-S505**). The thirteen-lined ground squirrel (*Ictidomys tridecemlineatus*), the Arctic ground squirrel (*Urocitellus parryii*), the groundhog (*Marmota monax*), and the Himalayan marmot (*Marmota himalayana*) do not express the *COA1/MITRAC15* locus (see **Supplementary Figure S506-S520**). In contrast to these species, the Eurasian red squirrel (*Sciurus vulgaris*) has an intact *COA1/MITRAC15* expressed in the skin (see **Supplementary Figure S521-S522**). Despite gene disrupting mutations, the North American beaver (*Castor canadensis*) *COA1/MITRAC15* locus is expressed in the blood and spleen (see **Supplementary Figure S523-S524**). Other tissues such as the brain, liver, stomach, ovarian follicle, skeletal muscle, and kidney do not show any expression at the *COA1/MITRAC15* locus (see **Supplementary Figure S525-S530**). The expressed transcript might represent a new long non-coding RNA that cannot produce a functional *COA1/MITRAC15* protein due to the presence of premature stop codons.

Chromosomal rearrangement in rodent species has resulted in the movement of genes flanking *COA1*/*MITRAC15* to new locations. The *BLVRA* gene is transcriptionally active in the mouse (*Mus musculus*) liver and heart even though it has translocated to an entirely different location between *AP4E1* and *NCAPH* (see **Supplementary Figure S531**). Genes on the left flank consisting of *HECW1*, *PSMA2,* and *MRPL32* are now located beside *ARID4B* and are expressed in the mouse (see **Supplementary Figure S532-S533**). The genes from the right flank (*MRPS24* and *URGCP*) are also transcriptionally active in the mouse at their new location beside *ANKRD36* (see **Supplementary Figure S534**). Remnants of *COA1*/*MITRAC15* occur between the *PTPRF* and *HYI* genes. However, no transcriptionally activity is seen in the mouse in the region between *PTPRF* and *HYI* genes (see **Supplementary Figure S535**). The new gene order and gene expression patterns are shared by rat (*Rattus norvegicus*) (see **Supplementary Figure S536-S540**), steppe mouse (*Mus spicilegus*) (see **Supplementary Figure S541-S545**), Gairdner’s shrewmouse (*Mus pahari*) (see **Supplementary Figure S546-S550**), Ryukyu mouse (*Mus caroli*) (see **Supplementary Figure S551-S555**), Algerian mouse (*Mus spretus*) (see **Supplementary Figure S556-S560**), deer mouse (*Peromyscus maniculatus*) (see **Supplementary Figure S561-S565**), prairie vole (*Microtus ochrogaster*) (see **Supplementary Figure S566-S570**), golden hamster (*Mesocricetus auratus*) (see **Supplementary Figure S571-S575**), Mongolian gerbil or Mongolian jird (*Meriones unguiculatus*) (see **Supplementary Figure S576-S579**), Chinese hamster (Cricetulus griseus) (see **Supplementary Figure S580-S584**), Northern Israeli blind subterranean mole rat (*Nannospalax galili*) (see **Supplementary Figure S585-S589**), white-footed mouse (*Peromyscus leucopus*) (see **Supplementary Figure S590-S594**) and fat sand rat (*Psammomys obesus*) (see **Supplementary Figure S595-S599**). The banner-tailed kangaroo rat (*Dipodomys spectabilis*) (see **Supplementary Figure S600-S601**) has a different gene order and appears to represent one of the pre-EBR species. However, we cannot rule out the possibility of genome assembly errors.

The genome assemblies of rodents such as the mouse and rat are well-curated and represent some of the highest-quality reference genomes (Rhie et al., 2021). To ensure that the chromosomal rearrangements identified are correct, we evaluated the correctness of genome assemblies of the mouse (see **Supplementary Figure S602-S608**) and white-footed mouse (*Peromyscus leucopus*) (see **Supplementary Figure S609-S616**) using PacBio long-read sequencing datasets. The mouse genome assembly has been finished to a very high quality using artificial clones of genome fragments (Osoegawa et al., 2000). We further verified the mouse genome assembly by visualizing the coverage of assembly fragments across the genomic regions of interest (see **Supplementary Figure S618-S623**). Repeat regions occur at the boundaries of the evolutionary breakpoint regions (see the last row of screenshots). Although repeat regions are a major contributing factor for the misassembly of genomes, the conserved gene orders across several species and concordance in the timing of the chromosomal rearrangement and support from long-read data support the presence of a genuine change in gene order.

The *COA1*/*MITRAC15* gene is intact and robustly expressed in the platypus (*Ornithorhynchus anatinus*) heart and brain (see **Supplementary Figure S624-S627**). Gene order in the short-beaked echidna (*Tachyglossus aculeatus*) matches the platypus and other outgroup species (see **Supplementary Figure S628)**. In contrast to the monotreme species, all marsupial genomes analyzed have a different gene order following chromosomal rearrangements. The gray short-tailed opossum (*Monodelphis domestica*) has the gene *ACVR2B* beside the new location of right flank genes of *COA1*/*MITRAC15*. The left flank genes are beside *GPR141B*. No traces of the *COA1*/*MITRAC15* gene are found either in the genome assembly or raw read datasets. The opossum brain expresses these adjacent genes with no transcripts in the intergenic regions (see **Supplementary Figure S629-S631**). The gene order and transcriptional activity were the same in the tammar wallaby (*Notamacropus eugenii*) (Uterus: see **Supplementary Figure S632-S633**), koala (*Phascolarctos cinereus*) (Liver and PBMC: see **Supplementary Figure S634-S636**), the Tasmanian devil (*Sarcophilus harrisii*) (Lung and Spleen: see **Supplementary Figure S637-S639**), and the common brushtail (*Trichosurus vulpecula*) (Liver: see **Supplementary Figure S640-S642**). Long-read sequencing data in the koala supports the correctness of genome assembly (see **Supplementary Figure S643-S645**).

The NCBI annotation documents the presence of transcripts, and the *COA1/MITRAC15* gene is remarkably well conserved in ungulate species (see **Supplementary Table S1**). Within ungulate species, certain Cervid species have remarkable sprinting abilities that allow them to escape from predators. However, in addition to sprinting ability, these species are resistant to fatigue. Hence, the prediction from our hypothesis is that gene loss would not occur in Cervid species. The white-tailed deer (*Odocoileus virginianus*) liver and retropharyngeal lymph node and the red deer (*Cervus elaphus*) blood transcriptomes express *COA1/MITRAC15* (see **Supplementary Figure S646-S649**).

The *COA1/MITRAC15* gene has undergone duplication within the primate lineage. We screened the genomes of 27 primate species to track down when the gene duplication event occurred. Based on the presence of the duplicate copies, the duplication event is estimated to have happened in the last 43 million years (see **Supplementary Figure S650-S651**). Subsequent duplications have also occurred in Nancy Ma’s night monkey (*Aotus nancymaae*) and a shared duplication in the black-capped squirrel monkey (*Saimiri boliviensis*) and the Panamanian white-faced capuchin (*Cebus imitator*). Concurrent with the gene duplication, the intron-exon structure of the *COA1/MITRAC15* gene has also changed (see **Supplementary Figure S652**). The functional copy of the *COA1*/*MITRAC15* gene is transcriptionally active in the gray mouse lemur (*Microcebus murinus*) (Kidney and Lung: see **Supplementary Figure S653-S654**), the northern greater galago (*Otolemur garnettii*) (Liver: see **Supplementary Figure S655**), Coquerel’s sifaka (*Propithecus coquereli*) (see **Supplementary Figure S656**), Nancy Ma’s night monkey (*Aotus nancymaae*) (Liver, Heart, and Kidney: see **Supplementary Figure S657-S659**), the common marmoset (*Callithrix jacchus*) (Lung, Liver, and Kidney: see **Supplementary Figure S660-S661**), the Panamanian white-faced capuchin (*Cebus imitator*) (Blood: see **Supplementary Figure S662-S664**), the black-capped squirrel monkey (*Saimiri boliviensis*) (Ovary and Heart: see **Supplementary Figure S665-S668**), the sooty mangabey (*Cercocebus atys*) (Liver: see **Supplementary Figure S669-S670**), the olive baboon (*Papio anubis*) (Kidney and Heart: see **Supplementary Figure S671-672**), the crab-eating macaque (*Macaca fascicularis*) (Blood and Liver: see **Supplementary Figure S673-S674**), the golden snub-nosed monkey (*Rhinopithecus roxellana*) (Heart and Blood: see **Supplementary Figure S675-S676**), human (*Homo sapiens*) (Liver : see **Supplementary Figure S677-S682**) and the Philippine tarsier (*Carlito syrichta*) (see **Supplementary Figure S683**).

The intron/exon structure of the *COA1/MITRAC15* gene has undergone several changes in the carnivore lineage (see **Supplementary Figure S684-S685**). However, outgroup species such as the horse (*Equus caballus*) and pangolin (*Manis javanica*) lack intron/exon structure (see **Supplementary Figure S686-S687**). We screened the RNA-seq dataset of multiple carnivore species to validate the annotation and evaluate the intron/exon structure changes. Alternative exon usage was also carefully analyzed to quantify the transcriptional status of *COA1/MITRAC15* in different carnivore species. The *COA1/MITRAC15* gene is transcriptionally active in the meerkat (*Suricata suricatta*) (testis and liver: see **Supplementary Figure S688-S690**), dog (*Canis lupus familiaris*) (spleen and skeletal muscle: see **Supplementary Figure S691-S702**), ferret (*Mustela putorius furo*) (heart and kidney: see **Supplementary Figure S703-S704**), Giant panda (*Ailuropoda melanoleuca*) (heart and liver: see **Supplementary Figure S705-S706**), American black bear (*Ursus americanus*) (liver, kidney, and the brain: see **Supplementary Figure S707-S708**), and Weddell seal (*Leptonychotes weddellii*) (lung and muscle: see **Supplementary Figure S709-S712**). Detailed investigation of the splice junctions and actual positions of splice sites in dog transcriptome also supports the *COA1/MITRAC15* gene annotation.

Skipping of the dog-like-exon-3 occurs in the transcriptomes of tiger (*Panthera tigris altaica*), lion (*Panthera leo persica*), cat (*Felis catus*), and puma (*Puma concolor*) (see **Supplementary Figure S713-S738**). Although annotation for the *COA1*/*MITRAC15* locus exists in the cheetah (*Acinonyx jubatus*), we found no transcripts in the skin RNA-seq data (see **Supplementary Figure S739-S740**). Close inspection of the *COA1*/*MITRAC15* locus in cheetah suggests gene loss. We further compared the splice isoforms found in canine and felid species through sashimi plots of the *COA1*/*MITRAC15* locus. The sashimi plot shows the links between the splice sites and the number of reads that are splice mapped between these sites (see **Supplementary Figure S741-S745**). Changes in the splice enhancers and splice silencer elements were also compared between cat and dog (see **Supplementary Figure S746**).

Co-expressed genes tend to perform related functions and are lost together. Hence, to identify the loss of genes related to *COA1/MITRAC15,* we identified the top 50 genes co-expressed with human ortholog based on the correlation values in COXPRESdb ver. 7.3 (Obayashi et al., 2019). The presence of orthologs of these co-expressed genes in the high-quality genomes of chicken and mouse using ENSEMBL BioMart (**Supplementary Table S7**). None of these co-expressed genes appear lost in Galliformes or rodents.

### 2.5 Molecular evolutionary analyses

#### 2.5.1 Relaxed selection signatures

Molecular signatures of relaxation in the degree of purifying selection generally accompany the loss of gene functionality and have been used as evidence of gene loss (Hecker et al., 2017; Sharma and Hiller, 2018; Shinde et al., 2019). Based on the gene sequence of *COA1/MITRAC15*, we could identify eleven Galliform species with gene-disrupting mutations (see **Supplementary Table S8** and **S9**). Two other Galliform species (*Chrysolophus pictus* and *Phasianus colchicus*) do not express the *COA1/MITRAC15* gene. Hence, we looked for signatures of relaxed selection in each of the terminal branches leading to each Galliform species. We quantified branch-specific selection patterns using the program RELAX (Wertheim et al., 2015) from the HyPhy package and the codeml program from the PAML (Yang, 2007) package. To test for relaxed selection in the terminal branches, we labeled the focal species as the foreground and used the Anseriformes species as the background species. We downloaded the phylogenetic tree with branch lengths from the TimeTree website. Although we found some evidence of relaxed selection in some of the Galliform species, the RELAX program also reported intensification of selection (see **Supplementary Table S10**). None of the internal branches were under relaxed selection.

We used the same phylogenetic tree and multiple sequence alignment to obtain branch-specific estimates of ω using the codeml program. The branch-specific estimates of ω are all greater than 1 in *Odontophorus gujanensis*, *Coturnix japonica*, *Meleagris gallopavo*, *Tympanuchus cupido*, *Pavo cristatus*, *Chrysolophus pictus*, *Phasianus colchicus,* and *Numida meleagris*. In the case of Galliform species (*Alectura lathami*, *Callipepla squamata,* and *Penelope pileata*) with intact *COA1/MITRAC15* gene, the values of ω are all less than 1. Except for chicken (*Gallus gallus*), species with gene-disrupting changes are not under purifying selection (see **Supplementary Table S10** and **S11**). We evaluated the internal nodes leading to the terminal branches for signatures of relaxed selection to ascertain whether gene loss had occurred in the common ancestor of the Galliform species with gene-disrupting mutations. However, all the ancestral branches appear to be under purifying selection and support the idea of recurrent lineage-specific gene loss suggested by the lineage-specific gene disrupting mutations seen in the Galliform species. Based on this branch-by-branch analysis of selection signatures, we could identify the approximate time frame in which gene loss might have occurred. To get a more accurate estimate of the gene loss timing, we used the method described by (Meredith et al., 2009).

We relied upon multiple sequence alignments of carnivores (see **Supplementary Table S12**), rodents (see **Supplementary Table S13**), and primates (see **Supplementary Table S14**) to identify gene disrupting mutations and changes in intron-exon structure. We evaluated each taxonomic group for lineage-specific relaxed selection (see **Supplementary Table S15**). Based on previous reports (Van Der Lee et al., 2017) of positive selection in primates, we additionally identified positively selected sites among primate species (see **Supplementary Table S16**).

#### 2.5.2 Time of gene loss

Different ω values were estimated for both of these labels (see **Supplementary Table S17**). The ω values for mixed(ω_m_) and functional(ω_f_) branches were estimated using two different codon substitution models (F1X4 and F3X4) to ensure the robustness of the estimates. The calculation of gene loss timing relies upon estimates of T_p_ (time for which the gene has been pseudogenic) using the method proposed by Meredith et al. (2009) by considering ω_p_ as 1. Based on the assumptions of 1ds and 2ds, we could get a confidence interval for the estimated time of gene loss (see **Supplementary Table S17**). Gene loss timing was estimated separately in rodents and carnivores (see **Supplementary Table S17**).

#### 2.5.3 GC content range and kmer abundance

The GC content range (minimum and maximum possible values of GC% for a given amino acid sequence) was calculated (see **Supplementary Table S18**) for *COA1/MITRAC15* and *PDX1* amino acid sequences in rodent and primate species using the window-based tool CodSeqGen (Al-Ssulami et al., 2020). The ContMap function in the R package phytools extrapolates the evolution of GC content along the phylogeny for both genes (see **Supplementary Figure S747-S749**). The program jellyfish (v2.2.8) (Marçais and Kingsford, 2011) was used to get the kmers (count command with the flags -C -m 21 -s 1000M and –t 16) and their abundance (dump command). The seqkit fx2tab (v0.10.1) (Shen et al., 2016) option calculated the abundance of kmers at different GC content bins and the GC content of each of the *COA1*/*MITRAC15* gene exons (see **Supplementary Table S19**).

#### 2.5.4 Quantification of gBGC

We calculated the (gBGC) for *COA1/MITRAC15* gene sequences of more than 200 species using the program mapNH(v1.3.0) implemented in the testNH package (Dutheil, 2008). In mapNH, we used multiple sequence alignments of the *COA1/MITRAC15* gene and species tree as input with the flag model=K80. A single gene-wide estimate of gBGC termed GC* is obtained for each species (see **Supplementary Table S20**). These estimates of GC* (GC* > 0.9 is significant) help understand the evolution of gBGC along the phylogeny using the ContMap function of the phytools package. Additionally, we also calculated the gBGC for taxonomic group-wise alignments using the programs phastBias and phyloFit implemented in the PHAST (v1.3) package (Capra et al., 2013; Hubisz et al., 2011). In the first step, we use the phyloFit program to fit phylogenetic models to multiple sequence alignments using the specified tree topology (--tree flag with species tree as argument) and substitution model (-- subst-mod flag with HKY85 model as argument). Next, the phastBias program with the –bgc flag identified gBGC tracts using the “.mod” file output from phyloFit (see **Supplementary Table S21**, see **Supplementary Figure S750-S778**). The gBGC tracts are positions along the gene with posterior probability >0.5.

#### 2.5.5 Computational prediction of RNA binding sites

The regulation of gene expression and splicing tends to be determined by the RNA binding sites present within the exons or introns of a gene (Fu and Ares, 2014). A combination of such splice enhancers and splice silencer elements work in concert to facilitate the expression of different isoforms (Dassi, 2017). The *COA1*/*MITRAC15* gene has changed the intron-exon organization and has acquired novel splice isoforms in felid species. These changes in splicing could result from changes in the RNA binding motifs present within the exons or introns of the gene. In contrast to felids, the splicing pattern in canid species matches the ancestral state. Hence, we compared the *COA1*/*MITRAC15* gene sequences of canid and felid species to identify differences in the RNA binding motifs. We used the RBPmap (Paz et al., 2014) webserver to predict the RNA binding sites in each exon and intron separately (see **Supplementary Table S22**).

## 3. Results

### 3.1 *COA1/MITRAC15* is a distant homolog of *TIMM21*

We identified that the *TIMM21* gene is a distant homolog of *COA1/MITRAC15* based on PSI-Blast and HHblits iterative profile-profile search of the uniport database. Of the 500 top search results from HHblits, 59 have annotation as “Cytochrome C oxidase assembly factor” or “Cytochrome C oxidase assembly protein” or “*COA1*”, and 120 as “*TIMM21*” homologs. The annotation of 13 proteins are “hypothetical”, nine are “membrane” proteins, eight are “DUF1783 domain-containing” proteins, and 27 proteins are from diverse proteins. The remaining 264 of the 500 hits are “Uncharacterized”. The large number of “Uncharacterized” proteins identified are challenging to interpret. Hence, to trace the relationships between the proteins identified as homologs of *COA1/MITRAC15*, we investigated the sequence identity-based clusters established by CLANS (see **Fig. 1A**). The large group of red dots consists of proteins annotated as *TIMM21,* and the collection of blue dots contains proteins annotated as *COA1/MITRAC15*. Homologs of *COA1/MITRAC15* from bacterial species form two clusters, a distinct light blue cluster consisting of predominantly Planctomycetes bacteria and a diffuse bunch of brown dots that consists of largely proteobacterial species. The group of orange dots consists of proteins annotated as *COA1/MITRAC15* in fungal genomes. The *COA1/MITRAC15* homologs in plants consist of a yellow cluster consisting of *Arabidopsis thaliana* homolog At2g20390 and the magenta cluster of *TIMM21*-like proteins containing *Arabidopsis thaliana* homolog At2g37940. The distinct *COA1/MITRAC15* and *TIMM21* groups found by the cluster analysis suggest that *TIMM21* is a very distant homolog of *COA1/MITRAC15*.

**Figure 1:**
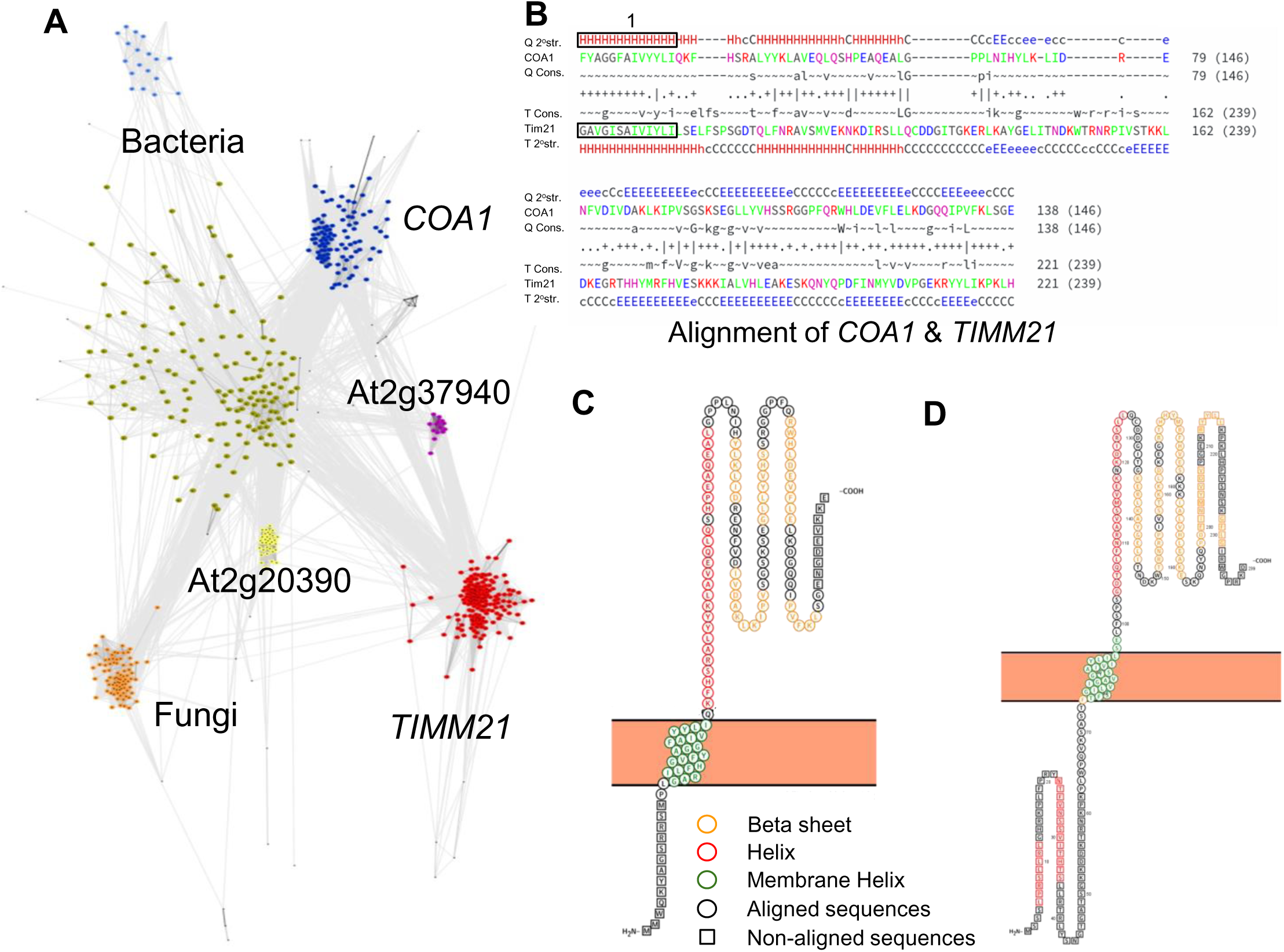
*COA1/MITRAC15* and *TIMM21* are distant homologs with similar amino acid sequence profiles and secondary structures. (**A**) Cluster map of *COA1/MITRAC15* homologs identified using profile-profile search implemented in HHblits. The cluster of *COA1/MITRAC15*: blue, *TIMM21*: red, homologs of *COA1/MITRAC15* from species of fungi: orange, homologs of *COA1/MITRAC15* from bacterial species: light blue cluster and diffuse brown cluster, *COA1/MITRAC15* homologs in plants, represented by *Arabidopsis thaliana* homolog At2g20390: yellow, *TIMM21*-like proteins that exist as duplicated copies in plants, represented by *Arabidopsis thaliana* homolog At2g37940: magenta. (**B**) The output of HHpred showing the alignment of human *COA1/MITRAC15* with yeast *TIMM21*. The region in the box highlights the predicted transmembrane helix. (**C**) The predicted secondary structure of human (*Homo sapiens) COA1/MITRAC15*. (**D**) The predicted secondary structure of yeast (*Saccharomyces cerevisiae*) *TIMM21*.

The list of proteins identified as homologs of human *COA1/MITRAC15* (**Supplementary File S2-S3**) and primate *COA1/MITRAC15* orthologs (**Supplementary File S4**) contain several *TIMM21* like proteins. Iterative PSI-BLAST search identified *TIMM21* homologs from the second iteration onwards and found an increasing number of *TIMM21* hits in each subsequent iteration (see **Supplementary File S5**). The pairwise alignment of the human *COA1/MITRAC15* protein sequence with the *TIMM21* protein with the best alignment (i.e., *TIMM21* from *Amblyomma cajennenseis*) shows that regions with the most substantial homology include the membrane-spanning domain and covers >100 residues (see **Fig. 1B**). In addition to the primary sequence-homology detected, both *TIMM21* and *COA1/MITRAC15* are known to play prominent roles in the mitochondria and have comparable secondary structures (see **Fig. 1C, 1D**). The strong homology between these proteins also allows for homology-based modeling of the tertiary structure of the *COA1/MITRAC15* protein using *TIMM21* as a model (see **Supplementary Figure S779-S783**). Despite the lack of well-conserved motifs, we found three well-matching columns (marked with a ‘|’ sign in **Fig. 1B**) between residues 91 to 95 in *COA1/MITRAC15*. Two consecutive conserved residues occur at residues 57-58, 64-65, and 67-68 of *COA1/MITRAC15*. The similar sequence, structure, and function of *COA1/MITRAC15* and *TIMM21* strongly support that these genes are homologs.

### 3.2 *COA1/MITRAC15* gene duplication, pseudogenisation, and exon reorganization

The sequence divergence between *COA1/MITRAC15* and *TIMM21* appears to result from changes in the *COA1/MITRAC15* gene intron/exon organization. The *COA1/MITRAC15* gene has undergone independent gene duplications followed by pseudogenisation and degeneration of the duplicated copy in both primates and carnivores. Consequently, the functional and pseudogene copies of *COA1/MITRAC15* have diverged considerably and formed distinct haplotypes. For example, the blast search of sequencing raw read data from the human genome with *COA1/MITRAC15* gene sequence as a query results in two distinct haplotypes. One set of reads correspond to the intact *COA1/MITRAC15* gene in humans, and the other set of reads are from the pseudogenic copy (see **Fig. 2A**). Comparative analysis of primate genome assemblies suggests that the pseudogenic copy results from a duplication of *COA1/MITRAC15* within the primate lineage (see **Supplementary Figure S651**). After duplication of the *COA1/MITRAC15* gene in primates, an extension of the N-terminal region has occurred in Cercopithecidae and Catarrhini and is transcriptionally active (see **Supplementary Figure S652**). However, new world monkeys do not have this N-terminal extension denoted as exon-1a. Both Cercopithecidae and Catarrhini have an additional start codon in exon-1a upstream from the original start codon in the ancestral exon-1 denoted as exon-1b in species with N-terminal extension. A striking difference between Cercopithecidae and Catarrhini is the lack of the internal start codon in Cercopithecidae, where Catarrhini has a start codon. Since proteome level data is not available for these species, we rely solely on the RNA-seq datasets and start and stop codons within the expressed transcripts to evaluate the exon/intron structure changes. Using these carefully annotated primate sequences of *COA1/MITRAC15*, we verified (see **Supplementary Table S16**) a previous report (Van Der Lee et al., 2017) of positive selection in this gene among primates.

**Figure 2:**
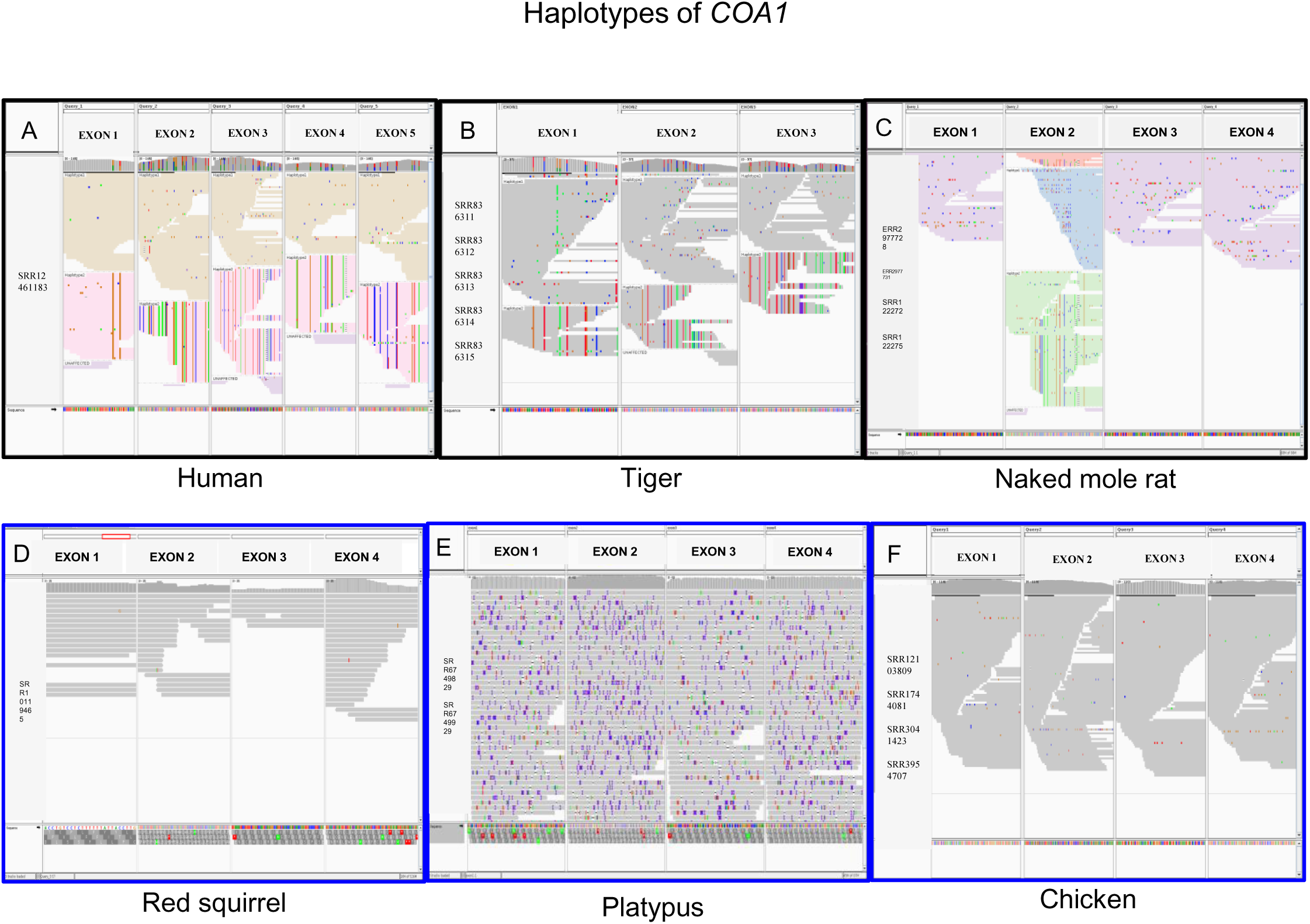
Comparison of haplotypes of *COA1/MITRAC15* gene inferred based on sequencing reads in different species visualized in IGV browser. (**A**) Two haplotypes of *COA1/MITRAC15* in humans (*Homo sapiens*) corresponding to the functional and pseudogene copies. (**B**) Two haplotypes of exon 1 to exon 4 of *COA1/MITRAC15* in tiger (*Panthera tigris*). (**C**) Two haplotypes in exon two and one haplotype of remaining exons of *COA1/MITRAC15* in naked mole-rat (*Heterocephalus glaber*). (**D**) The single haplotype of *COA1/MITRAC15* gene in chicken (*Gallus gallus*). (**E**) The single haplotype of *COA1/MITRAC15* in the platypus (*Ornithorhynchus anatinus*). (**F**) The single haplotype of *COA1/MITRAC15* in red squirrel (*Sciurus vulgaris*).

Independent duplication of *COA1/MITRAC15* has occurred in carnivores (see **Supplementary Figure S685**). However, similar to primates, the duplicated copy has undergone pseudogenization and diverged from the functional gene sequence. For example, sequencing raw read data in the tiger consist of two distinct haplotypes corresponding to the intact and pseudogene copies (see **Fig. 2B**). While the intact copy is located at a genomic region (*STK17A* & *HECW1* upstream and *BLVRA* & *VOPP1* downstream) with conserved synteny across other mammals, the pseudogene copy occurs adjacent to the *PRR32* gene. Outgroup species such as horse (*Equus caballus*) and pangolin (*Manis javanica*) have a single copy of the *COA1/MITRAC15* gene with all raw reads supporting a single haplotype (see **Supplementary Figure S686**). Both sub-orders (Caniformia and Feliformia) within Carnivora share this duplication of the *COA1/MITRAC15* gene (see **Supplementary Figure S685**).

The intact *COA1/MITRAC15* copy is expressed in diverse transcriptomes among Caniformia species, while the pseudogene copy lacks expression. The first and second exons are orthologous; however, the genomic location of the transcribed third exon is different between Feliformia (cat-like-exon-3) and Caniformia species (dog-like-exon-3) (see **Fig. 3**). The final exon of the *COA1/MITRAC15* gene in Feliformia extends to 163 base pairs (*Panthera tigris altaica, Panthera leo, Panthera pardus,* and *Lynx lynx*) and 160 base pairs (*Puma concolor* and *Felis catus*) compared to the 100 base pairs in Caniformia species. A single deletion event causes the difference of three base pairs between these two groups of Feliformia at the 24^th^ base of exon-4. The extended final exon shared by all Feliformia species results from a two-base frameshift deletion before the erstwhile stop codon in exon-4. Despite the extended last exon in Feliformia species, the full-length open reading frames of Feliformia (130/131 amino acids) and Caniformia (135 amino acids) are comparable.

**Figure 3:**
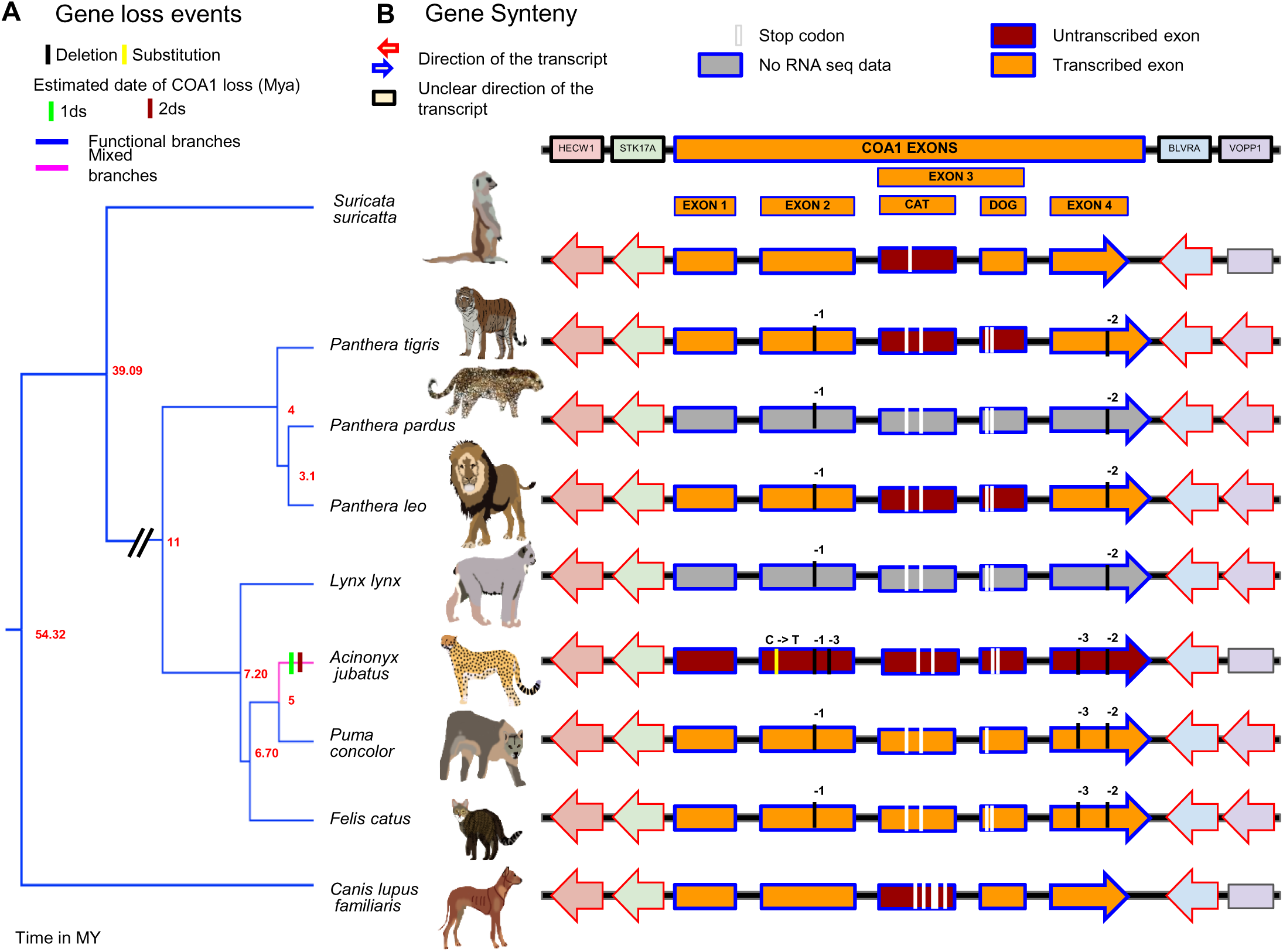
Loss of *COA1/MITRAC15* gene in Feliform. (A) Gene loss event in *Acinonyx jubatus* besides a time-calibrated phylogenetic tree downloaded from the time tree website. (B) Gene order in the genomic region flanking the *COA1/MITRAC15* gene and its exons in genomes. Red and blue arrows depict the direction of gene transcription relative to the *COA1/MITRAC15* gene for consistency across species. Gray boxes represent the genes located on short scaffolds with unknown orientation.

The shorter reading frame in Feliformia results from the majority of *COA1/MITRAC15* transcripts skipping the dog-like-exon-3, whose inclusion results in premature stop codons in all the seven Feliformia species. The dog-like-exon-3 is present in all *COA1/MITRAC15* transcripts of Caniformia species and does not contain gene-disrupting changes. A single base deletion in all Feliformia species changes the end phase of exon-2 to maintain an intact reading frame while skipping the dog-like-exon-3. Transcriptomes of the cat (*Felis catus*) from the spleen (see **Supplementary Figure S744**) and puma (*Puma concolor*) from blood (see **Supplementary Figure S745**) exhibit expression of a proto cat-like-exon-3 which gets spliced into some of the *COA1/MITRAC15* transcripts. However, the majority of transcripts skip this proto cat-like-exon-3 which contains premature stop codons. These changes in exon splicing patterns between Caniformia and Feliformia species appear to result from changes in splice factor binding sites at the *COA1/MITRAC15* locus (see **Supplementary Figure S746**).

Except for the cheetah (*Acinonyx jubatus*), intact transcribed open reading frames are discernible in all carnivore species at the *COA1/MITRAC15* locus identified based on conserved synteny across mammals (see **Fig. 3**). The gene disrupting premature stop codon in the cheetah is due to a single base C->T substitution at the 27^th^ base of exon-2 assembled at the *COA1/MITRAC15* locus. The duplicated copy of *COA1/MITRAC15* also contains a premature stop codon at the 49^th^ base of exon-2 caused by a single base insertion at the 11^th^ base of exon-2. The *COA1/MITRAC15* gene transcripts are missing in the skin transcriptome of the cheetah. Hence, multiple lines of evidence support *COA1/MITRAC15* gene loss in the cheetah. Gene loss in the cheetah occurred between 2.98-3 MYA (**Supplementary Table S17**).

In contrast to primates and carnivores, reads support multiple haplotypes of *COA1/MITRAC15* only in the second exon of naked mole-rat (see **Fig. 2C**). Hence, the duplicated copy of *COA1/MITRAC15* in naked mole-rat appears to have mostly degraded. However, we cannot rule out the possibility that the reads from other haplotypes spanning the remaining three exons are missing due to high GC content. The sequencing reads support the presence of a single intact open reading frame in the red squirrel (see **Fig. 2D**) and platypus (see **Fig. 2E**). Although a single haplotype occurs in the raw read dataset of chicken, this haplotype has gene-disrupting changes (see **Fig. 2F**). The gene-disrupting modifications identified in the chicken *COA1/MITRAC15* gene were investigated further by screening long-read datasets, transcriptomes, and genomes of various Galliform species.

### 3.3 *COA1/MITRAC15* gene loss in Galliform species

We found evidence of eight independent gene-disruption events in the *COA1/MITRAC15* gene in the galliform group (see **Fig. 4A**). The chicken (*Gallus gallus*) and Amazonian wood quail (*Odontophorus gujanensis*) have single-base G to T substitutions at the 69^th^ base of exon-2 and the 72^nd^ base of exon-4 in the *COA1/MITRAC15* gene, respectively (see **Supplementary Table S9**). These substitutions lead to (**G**AA→**T**AA) premature stop codons. Gene loss of *COA1/MITRAC15* is estimated to have occur between 23 MYA and 29 MYA in chicken and between 17 MYA and 18 MYA in the Amazonian wood quail (see **Supplementary Table S9** and **S17**). In the Indian peafowl (*Pavo cristatus*), two single-base deletions, one at 37^th^ base of exon-1 and another at 31^st^ base of exon-4, result in premature stop codons in exons 2, 3, and 4. The gene disrupting mutations identified in the Indian peafowl (*Pavo cristatus*) also occur in the green peafowl (*Pavo muticus*). Loss of the *COA1/MITRAC15* gene is estimated to have occurred between 20 MYA and 29 MYA in the peafowls (see **Supplementary Table S17**). The exon-2 of Pinnated grouse (*Tympanuchus cupido*) and Helmeted guineafowl (*Numida Meleagris*) have independent 13 and 17 base deletions. Changes in the reading frame resulting from these deletions lead to several premature stop codons (see **Supplementary Table S9**). The 13-base deletion in the exon-2 of the Pinnated Grouse (*Tympanuchus cupido*) also occurs in Gunnison grouse (*Centrocercus minimus*), Rock ptarmigan (*Lagopus muta*), and the black grouse (*Lyrurus tetrix*). The estimated time of gene loss in these four species is between 18 MYA and 20 MYA, and for Helmeted guineafowl, it is between 39 MYA and 40 MYA (see **Supplementary Table S17**).

**Figure 4:**
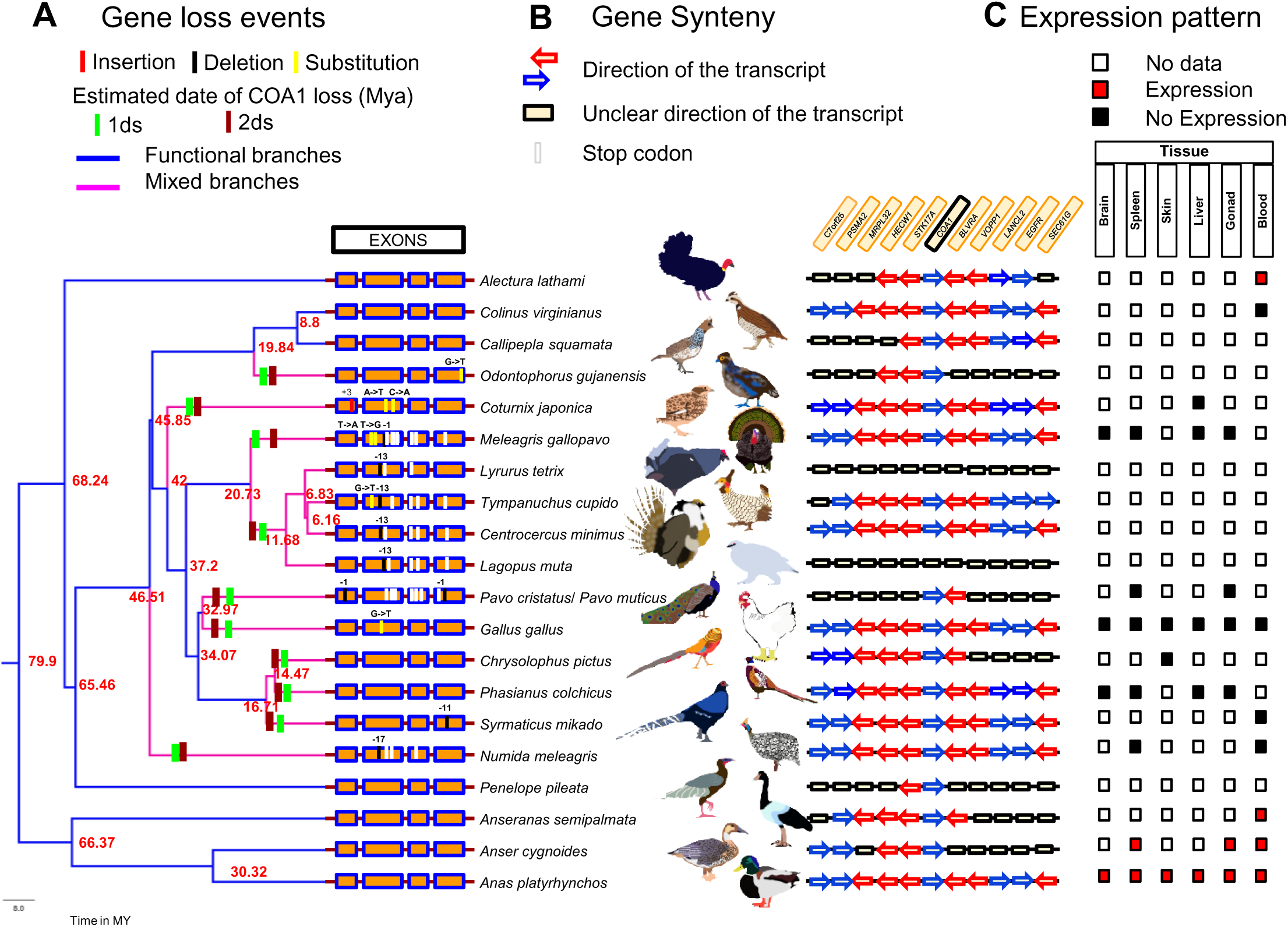
Recurrent loss of *COA1/MITRAC15* gene in Galliform species. (**A**) Gene loss events in ten Galliform species besides a time-calibrated phylogenetic tree downloaded from the time tree website. Blue branches in the tree represent functional branches, and the magenta-colored branches represent mixed (functional + pseudogenic) branches. The method proposed by Meredith et al., 2009 was used to estimate the time of gene loss using two different substitution rates (1ds and 2ds). Short colored bars depict the locations of the gene disrupting mutations on the four exons of *COA1/MITRAC15*. (**B**) Gene order in the genomic region flanking the *COA1/MITRAC15* gene in bird genomes. Red and blue arrows depict the direction of gene transcription relative to the *COA1/MITRAC15* gene for consistency across species. Gray boxes represent the genes located on short scaffolds with unknown orientation. (**C**) The gene expression pattern of the *COA1/MITRAC15* gene in six tissues (brain, spleen, skin, liver, gonad, and blood) was assessed by screening RNA-seq datasets. The red-colored blocks depict the robust expression of the *COA1/MITRAC15* gene, the black-colored blocks depict a lack of *COA1/MITRAC15* gene expression in that particular tissue, and the white-colored blocks represent a lack of data for that tissue.

In Turkey (*Meleagris gallopavo*), a two-base substitution at bases 7 & 8 and a single base deletion at the 37^th^ base of exon-2 result in a frameshift in the *COA1/MITRAC15* gene leading to premature stop codons. Gene loss in Turkey is estimated to have occurred between 14 MYA and 18 MYA. Two closely spaced single base substitutions (**<u>A</u>**A**<u>C</u>**→**<u>T</u>**A**<u>A</u>**) at 48th and 50th positions of exon-2 result in a premature stop codon in the Japanese quail (*Coturnix japonica*). The time of gene loss in the Japanese quail is estimated between 35 MYA and 36 MYA (see **Supplementary Table S17**). The Mikado pheasant (*Syrmaticus mikado*) has an 11-base deletion in exon-4, and the time of gene loss is between 14 MYA and 16 MYA. Other Galliform species such as Australian brushturkey (*Alectura lathami*), Blue quail (*Callipepla squamata*), Ring-necked pheasant (*Phasianus colchicus*), Golden pheasant (*Chrysolophus pictus*), and White-crested guan (*Penelope pileata*) have intact *COA1/MITRAC15* coding sequences. The coding region is intact in outgroup species such as Swan goose (*Anser cygnoides*), Duck (*Anas platyrhynchos*), and Magpie goose (*Anseranas semipalmata*). Five genes upstream (*BLVRA*, *VOPP1*, *LANCL2*, *EGFR,* and *SEC61G*) and downstream (*STK17A*, *HECW1*, *MRPL32*, *PSMA2,* and *C7orf25*) from *COA1/MITRAC15* retain a conserved order in birds. We relied upon this conserved order to verify the 1 to 1 orthology of the *COA1/MITRAC15* gene across species (see **Fig. 4B**).

Signatures of relaxed selection in Galliform species with gene disrupting changes further support the loss of *COA1/MITRAC15* in these lineages (see **Supplementary Table S10**). Despite intact coding regions, the Ring-necked pheasant (*Phasianus colchicus*) and Golden pheasant (*Chrysolophus pictus*) *COA1/MITRAC15* sequences also have signatures of relaxed selection (see **Supplementary Table S10**). None of the four tissues (Brain, Spleen, Liver, and Gonad) for which RNA-seq data is available from the Ring-necked pheasant shows any *COA1/MITRAC15* transcripts. Similarly, the one tissue (Skin) for which RNA-seq data is available in the Golden pheasant (*Chrysolophus pictus*) does not show *COA1/MITRAC15* expression. To evaluate the relevance of the gene disrupting mutations and signatures of relaxed selection identified in galliform species, the transcriptomes of Galloanserae species were screened to assess the transcriptional status of *COA1/MITRAC15*. We evaluated RNA-seq datasets of several comparable tissues across species and found the *COA1/MITRAC15* gene is not transcribed in chicken despite screening more than 20 tissues (see **Fig. 4C**). Other Galloanserae species have RNA-seq data available for very few tissues. We evaluated the RNA-seq datasets from six tissues (Brain, Spleen, Skin, Liver, Gonad, and Blood) available in several species for the presence of *COA1/MITRAC15* transcripts. Our search consistently revealed transcription of *COA1/MITRAC15* gene in Anseriformes species in contrast to lack of transcription in Galliform species except for Australian brushturkey (*Alectura lathami*) and northern bobwhite (*Colinus virginianus*), which have intact *COA1/MITRAC15* gene that is under strong purifying selection (see **Fig. 4C**). The lack of gene expression and signatures of relaxed selection in the Ring-necked pheasant (*Phasianus colchicus*) and Golden pheasant (*Chrysolophus pictus*) suggests gene loss. The putative gene loss in both these species occurred between 12 MYA and 13 MYA.

### 3.4 Complete erosion of *COA1/MITRAC15* locus is challenging to prove

Search for the *COA1/MITRAC15* gene in the mammoth genome demonstrated striking heterogeneity in coverage of the four exons based on the Illumina ancient DNA sequencing datasets analyzed (see **Fig. 5A-F**). Despite having comparable genome-wide coverage, we could see that not all exons occur in all the datasets. For instance, the re-sequencing dataset from PRJEB29510 (162 Gb) does not have reads from any of the four *COA1/MITRAC15* exons. However, the datasets from PRJEB7929 (88.34 Gb) and PRJNA397140 (155 Gb) have reads covering three exons each despite having much lower genome-wide coverage. The third exon of *COA1/MITRAC15* was also missing or had fewer reads than the other three exons in most of the datasets. The dataset from PRJEB42269 had no reads from the first exon but had a few reads from exons three and four. We reasoned that this heterogeneity in the coverage of various *COA1/MITRAC15* exons was mainly a result of the well-established sequencing bias of Illumina that results in inadequate coverage of GC-rich regions (Chen et al., 2013). Quantification of GC content in each of the four *COA1/MITRAC15* exons and kmer abundance in different GC content bins in each of the mammoth Illumina re-sequencing datasets explains most of the heterogeneity in coverage between datasets as well as exons (see **Fig. 5G**). In contrast to the *COA1/MITRAC15* gene, we did not see heterogeneity in the sequencing coverage of *TIMM21* exons despite comparable GC content for some of the exons (see **Fig. 5G** and **Supplementary Figure S784-S791**).

**Figure 5:**
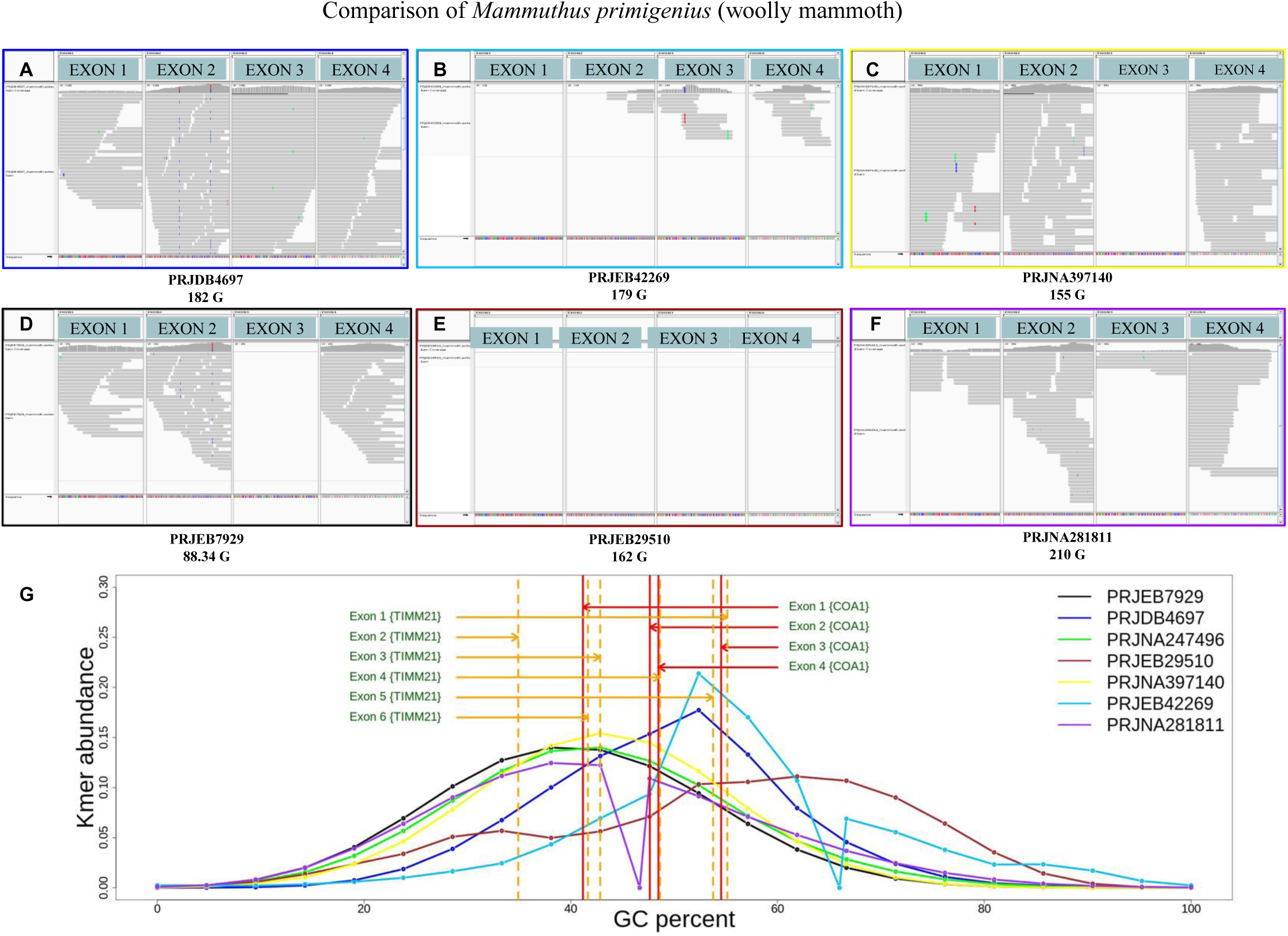
Comparison of different sequencing datasets of woolly mammoth (*Mammuthus primigenius*) for *COA1/MITRAC15* gene exons. Gray rectangles show the reads mapped to each exon. Panels **A** to **F** shows the reads with sequence supporting each exon from different woolly mammoth SRA bio projects [PRJDB4697 (182 Gb), PRJEB42269 (179 Gb), PRJNA397140 (155 Gb), PRJEB7929 (88.34 Gb), PRJEB29510 (162 Gb), and PRJNA281811 (210 Gb)]. Panel **G** indicates the GC percentage vs. K-mer abundance of different project IDs mentioned in different colors. The vertical dotted lines in orange denote the GC percentage of *TIMM21* exons, and vertical solid lines in red indicate the GC percentage of *COA1/MITRAC15* exons.

The heterogeneity in sequencing coverage of *COA1/MITRAC15* exons demonstrates the challenges of detecting its presence in Illumina sequencing datasets. GC-biased gene conversion (gBGC) plays a defining role in the base composition for any particular gene or genomic region. It preferentially fixes GC in AT/GC heterozygotes and increases the GC content. The GC content of the *COA1/MITRAC15* exons can be driven to extreme values by gBGC. The magnitude of gBGC also varies across the genome within a species as well as between species. Therefore, *COA1/MITRAC15* orthologs from closely related species or even duplicated copies of *COA1/MITRAC15* within the same species can have very different GC content. Such differences in GC content can result in correspondingly different coverage of the gene sequence in Illumina data and masquerade as a gene loss event (Botero-Castro et al., 2017; Hargreaves et al., 2017).

A well-known example for high GC content impeding sequencing is the gene *PDX1*, which has striking differences in GC content between closely related rodent species and requires dedicated GC-rich DNA enrichment protocols for sequencing. We contrasted *COA1/MITRAC15* with the *PDX1* genes of rodents by comparing the minimum (see **Supplementary Table S20**) and maximum (see **Supplementary Table S20**) GC contents possible given their amino acid sequence. Although *COA1/MITRAC15* had lower GC content levels than *PDX1*, we could not rule out the possibility of gBGC affecting some of the exons. The values of GC* across more than 200 vertebrate species with intact *COA1/MITRAC15* reading frames suggested considerable heterogeneity between taxa (see **Supplementary Figure S749)**. In each taxonomic group, the prevalence of gBGC was separately quantified (see **Supplementary Figure S750-S772**). Strong patterns of gBGC occur in the *COA1/MITRAC15* sequence of several species (see **Supplementary Figure S750-S772**: elephant (*Loxodonta africana*), kagu (*Rhynochetus jubatus*), blue-crowned manakin (*Lepidothrix coronata*), Chilean tinamou (*Nothoprocta perdicaria*), American black bear (*Ursus americanus*), North American river otter *(Lontra canadensis)*, meerkat (*Suricata suricatta*), California sea lion (*Zalophus californianus*), little brown bat (*Myotis lucifugus*), large flying fox (*Pteropus vampyrus*), southern pig-tailed macaque (*Macaca nemestrina*), Brazilian guinea pig (*Cavia aperea*), sheep (*Ovis aries*), eastern brown snake (*Pseudonaja textilis*) and the Goode’s thornscrub tortoise (*Gopherus evgoodei*)). However, none of the rodent species with intact *COA1/MITRAC15* show any striking gBGC patterns. The GC content vs. kmer abundance plots of Pacbio, BGI-seq, and Illumina datasets spans the entire range of GC contents seen in *COA1/MITRAC15* exons (see **Supplementary Figure S775**). Since the GC content of individual *COA1/MITRAC15* exons differs between species groups (see **Supplementary Figure S775-S778**), the high GC content of certain regions might result in inadequate sequencing coverage of the *COA1/MITRAC15* gene in some species. Hence, the lack of sequencing reads covering *COA1/MITRAC15* cannot serve as definitive evidence of gene loss.

### 3.5 *COA1/MITRAC15* occurs in an evolutionary breakpoint region

We find evidence of *COA1/MITRAC15* gene disrupting mutations and lack of gene expression in multiple RNA-seq datasets despite a conserved gene order in the rabbit (*Oryctolagus cuniculus*), naked mole-rat (*Heterocephalus glaber*), and four Sciuridae species (*Urocitellus parryii*, *Spermophilus dauricus*, *Ictidomys tridecemlineatus*, *Marmota marmota marmota*). The gene disrupting mutations identified in the rabbit *COA1/MITRAC15* gene includes a two-base pair deletion at the 22^nd^ codon of exon-1 and single base pair deletions in exon-2 at the 13^th^ and 37^th^ codons. Gene disrupting changes in the third exon consist of a five-base insertion between the 11^th^ and 12^th^ codon, one base insertion at the 17^th^ codon, and one base deletion in the 23^rd^ codon (see **Fig. 6** and **Supplementary Table S13**). These six gene-disrupting changes result in premature stop codons in exon-2 and exon-4. Gene loss in the rabbit is estimated to have occurred between 12 MYA and 17 MYA (see **Fig. 6** and **Supplementary Table S17**). The lack of *COA1/MITRAC15* RNA-seq reads in tissues such as the brain, liver, and testis that express *COA1/MITRAC15* in closely related species supports the loss of the *COA1/MITRAC15* gene in the naked mole-rat. Besides the lack of a start codon, a single gene disrupting mutation is found in the naked mole-rat *COA1/MITRAC15* gene and consists of a single base deletion at the 21^st^ codon of exon-1. Gene loss in the naked mole-rat is estimated between 7 MYA and 11 MYA (see **Supplementary Table S17** and **Fig. 6**).

**Figure 6:**
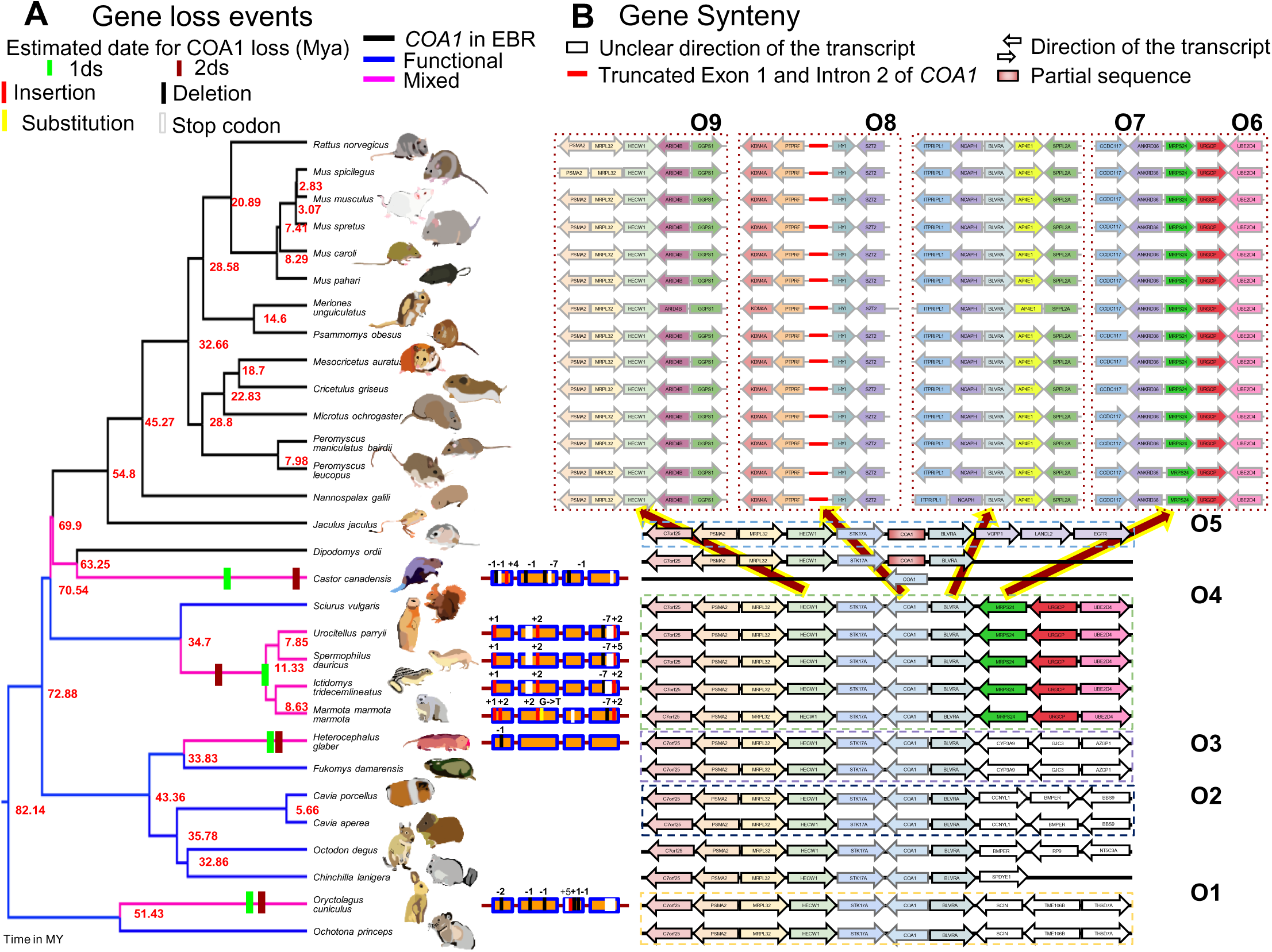
Recurrent loss of *COA1/MITRAC15* gene in rodent species. (**A**) Gene loss events in seven rodent species through four events are represented exon-wise beside the pink-colored branches of the time-calibrated phylogenetic tree obtained from the time tree website. Blue branches correspond to functional copies of *COA1/MITRAC15,* and black branches correspond to the Evolutionary Breakpoint Region (EBR) (**B**). Gene order in the genomic region flanking the *COA1/MITRAC15* gene in rodent genomes. Arrows depict the direction of gene transcription relative to the *COA1/MITRAC15* gene for consistency across species. Boxes represent the genes located on short scaffolds with unknown orientation. Each dotted box contains one type of gene order, and the brown arrows highlighted in yellow emerging from gene order O5 depict the EBR event that leads to gene orders O6, O7, O8, and O9. Gene order O8 and O7 contain partial remains of the *COA1/MITRAC15* gene and a functional *BLVRA* gene, respectively. A solid red line within gene order O8 depicts the partial exon one and intron 2 of *COA1/MITRAC15* located between the *PTPRF* and *HYI* genes. The gene order O6 and O9 correspond to the regions on the left and right flanks of the region containing *STK17A*, *COA1/MITRAC15,* and *BLVRA*.

The presence of common gene disrupting changes such as a one base pair insertion at second codon of exon-1, two base pair insertion at 25^th^ codon of exon-2, seven base pair deletion between 25^th^ and 26^th^ codon of exon-4, and a 2-base insertion at 33^rd^ codon of exon-4 supports a shared gene loss in four Sciuridae species (*Urocitellus parryii*, *Spermophilus dauricus*, *Ictidomys tridecemlineatus*, *Marmota marmota marmota*). The *COA1/MITRAC15* gene of alpine marmot has additional gene disrupting changes consisting of a 2-base insertion between the 8^th^ and 9^th^ codon of exon-1 and a single nucleotide substitution at the 26^th^ codon of exon-2. The 2-base insertion at the 33^rd^ codon of exon-4 has extended to a five-base pair insertion in the Daurian ground squirrel (*Spermophilus dauricus*). The estimated time of gene loss for this shared event is between 10 MYA and 30 MYA (see **Supplementary Table S17** and **Fig. 6**).

The presence of intact open reading frames robustly expressed at syntenic locations in closely related (∼30 to 50 million years) species strongly supports at least three independent *COA1/MITRAC15* gene loss events (see **Fig. 6**). Multiple gene-disrupting mutations in the *COA1/MITRAC15* gene of the North American beaver (*Castor canadensis*) suggest a fourth independent gene loss event. Gene-disrupting mutations in the beaver result in at least two premature stop codons. In the first exon, single-base deletions occur in the 3^rd^ and 20^th^ codon, a four-base insertion occurs between 33^rd^ and 34^th^ codon. The second exon has a single-base deletion in the 33^rd^ codon and a seven-base pair deletion between 29^th^ and 30^th^ codons. A single base deletion occurs at the 12^th^ codon of exon-3 (see **Supplementary Table S13** and **Fig. 6**). The genome assembly of the North American beaver is fragmented, and the synteny of the flanking regions cannot be verified. The Illumina sequencing raw reads support the gene disrupting mutations identified in the genome assembly (**Supplementary File S1**), and duplicate copies don’t occur. The loss of the *COA1/MITRAC15* gene in the beaver is estimated to have occurred sometime between 3 MYA and 23 MYA (see **Supplementary Table S17** and **Fig. 6**).

The North American beaver is phylogenetically closely related to the Ord’s kangaroo rat (*Dipodomys ordii*) and the lesser Egyptian jerboa (*Jaculus jaculus*). The more contiguous genome assemblies of the jerboa and kangaroo rat allow verification of a conserved gene order likely to be shared by the North American beaver (see **Fig. 6**). The presence of repetitive elements and lack of long-read sequencing data in most rodent species prevents genome assembly verification. Hence, we have screened the genomes of several closely related rodent species and verified the genome assemblies using long-read sequencing data or cloned fragments that cover parts of the genome. Gaps in the genome assembly also hamper the identification of the correct gene order. Previous reports that examined genome assemblies and EST data have claimed loss of the *STK17A* gene in mice due to a chromosomal rearrangement spanning this genomic region (Fitzgerald and Bateman, 2004). Detailed examination of gene order flanking the *COA1/MITRAC15* locus in several rodent genomes revealed the occurrence of this previously reported chromosomal rearrangement event (see **Fig. 6**).

Identifying gene loss events coinciding with EBRs is notoriously challenging and has motivated nuanced inferences in both bird (Botero-Castro et al., 2017) and rodent species (Hargreaves et al., 2017). Nonetheless, more than a dozen rodent species share the putative combined loss of *STK17A* and *COA1/MITRAC15* (see **Fig. 6**). Based on the presence of adjacent genes, the rearranged regions could be tracked down to two different chromosomes (see **Fig. 6, O6,** and **O9**). Genes on the left flank of *STK17A*-*COA1*-*BLVRA* consist of *PSMA2*, *MRPL32,* and *HECW1* in gene orders O1 to O5. After the chromosomal rearrangement, the same sequence of genes can be found in gene order O9 and occur adjacent to *ARID4B* and *GGPS1*. Genes on the right flank of *STK17A*-*COA1*-*BLVRA* consist of *MRPS24*, *URGCP,* and *UBE2D4* in gene order O4. Several other gene orders (O1 to O5) occur on the right flank in various species. The sequence of genes found on the right flank in gene order O4 is also found sequentially in gene order O6 and occurs adjacent to *ANKRD36* and *CCDC117* after the chromosomal rearrangement.

We found that the *BLVRA* gene has translocated to an entirely new location and does not co-occur with either the left or right flank. However, the new location of the *BLVRA* gene between the *NCAPH* and *ITPRIPL1* genes on the left flank and *AP4E1* and *SPPL2A* genes on the right flank is consistently conserved across all 14 post-EBR species and corresponds to gene order O7. Both *COA1/MITRAC15* and *STK17A* are missing in the post-EBR rodent genome assemblies. The search of the genome assemblies, sequencing raw read datasets, and RNA-seq datasets also failed to find any evidence of an intact *COA1/MITRAC15* or *STK17A* gene. All raw read and genome assembly hits for *STK17A* while using queries from pre-EBR rodent genomes could be traced back to the *STK17B* gene that matches with the *STK17A* gene at a short sequence stretch. The *STK17A* gene is lost or has sequence properties that prevent it from being sequenced with currently available technologies. The exon-1 region of *COA1/MITRAC15* occurs in a gene desert region between *PTPRF* and *HYI* genes in post-EBR species. Using blast search of *COA1/MITRAC15* introns, we found strong support for the existence of *COA1/MITRAC15* intron-2 close to the exon-1 hit. Pairwise genome alignments provide support for the presence of *COA1/MITRAC15* gene remains at this location (see **Supplementary File S6**). Notably, the *COA1/MITRAC15* remnants of a truncated exon-1 and intron-2 occur in the gene desert located between *PTPRF* and *HYI* genes only in post-EBR species. None of the pre-EBR species had any such remains. Hence, the *COA1/MITRAC15* remnants between *PTPRF* and *HYI* genes are unlikely to have resulted from duplicated copies of *COA1/MITRAC15*. The synteny of this region is well conserved with *KDM4A* and *PTPRF* on the left flank and *HYI* and *SZT2* on the right side and corresponds to gene order O8. Careful examination of this region in RNA-seq datasets found no evidence of transcripts.

Comparison of gene order in marsupial species with various outgroup species (including the platypus and short-beaked echidna from the order Monotremata) identified the presence of an independent chromosomal rearrangement event spanning the *COA1/MITRAC15* locus (see **Fig. 7**). In contrast to the rodent-specific EBR, we found that the *STK17A* gene is intact in post-EBR (gene order O2 and O3 in **Fig. 7**) marsupial species. However, an extensive search of marsupial genomes, transcriptomes, and raw sequencing read datasets (including high coverage Pacbio datasets for the Koala) failed to find any evidence of *COA1/MITRAC15* orthologs or its remnants. Lack of sequencing reads from *COA1/MITRAC15* in marsupial species suggests either complete erosion of the gene or drastic change in sequence composition that eludes sequencing with currently available technologies.

**Figure 7:**
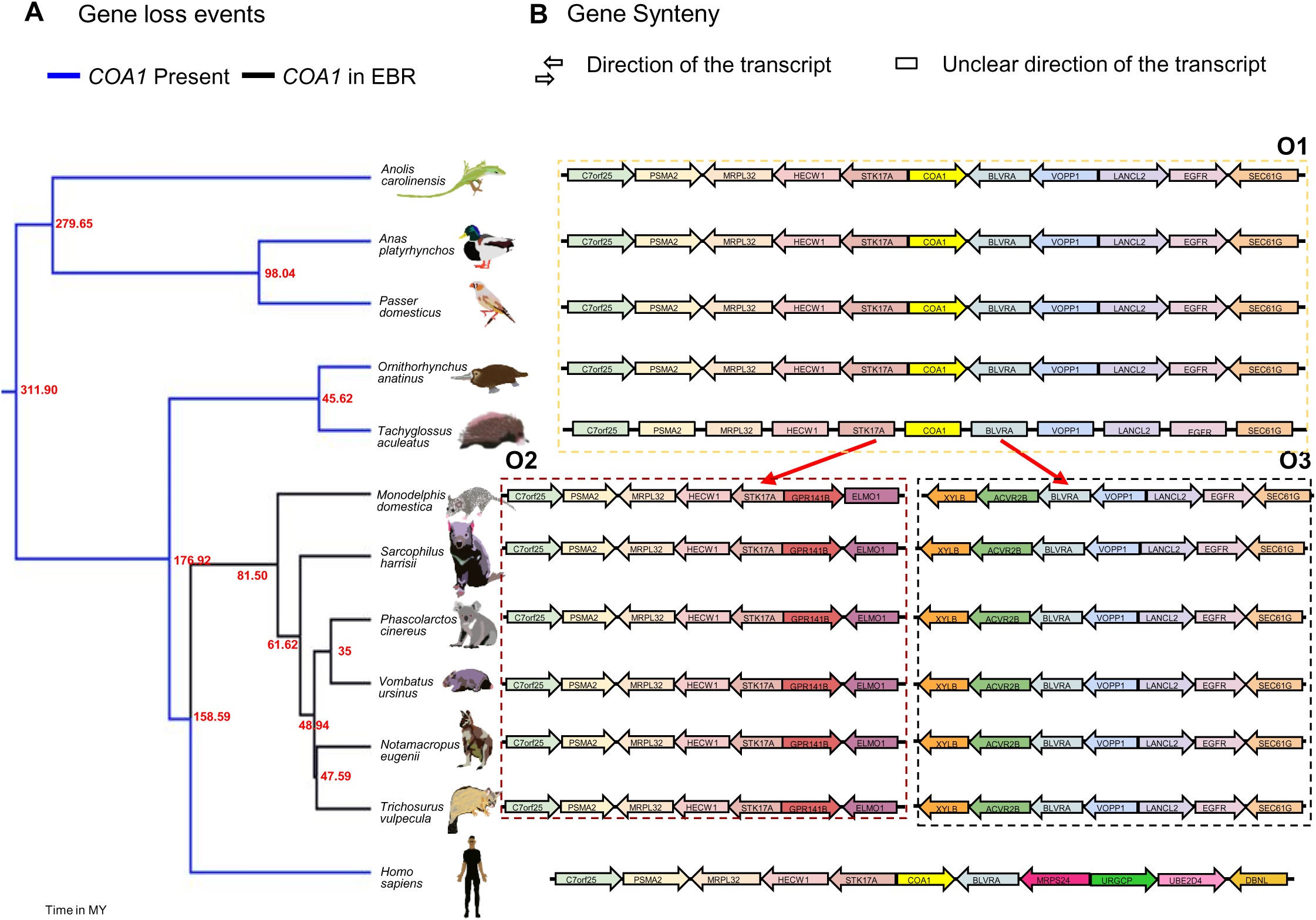
The genomic region spanning the *COA1/MITRAC15* gene coincides with an evolutionary breakpoint (EBR). (**A**) The phylogenetic relationship between marsupial species along with few outgroup species. The phylogenetic tree is from the time tree website. (**B**) The gene order in the region flanking the *COA1/MITRAC15* gene. The arrows show the direction of gene transcription relative to the *COA1/MITRAC15* gene for consistency across species. Each dotted box contains one type of gene order, and the red arrows from gene order O1 depict the EBR event that leads to gene orders O2 and O3 in the six marsupial species. The outgroup species have the Pre-EBR gene order O1. In the post-EBR gene orders O2 and O3, the *COA1/MITRAC15* gene occurs in the EBR, and the gene order of flanking genes is changed. A functional *COA1/MITRAC15* can be identified in the outgroup species but is presumably lost in marsupial species as it is missing in the genome assembly and raw read datasets.

## 4. Discussion

Our search of the sequence databases identified that *COA1/MITRAC15* and *TIMM21* are distant homologs with representative genes found in animals, plants, and fungi. The occurrence of *COA1/MITRAC15* homologs in α-proteobacteria supports an ancestral role for these genes (Kurland and Andersson, 2000). Endosymbiotic theories explain the origin of eukaryotes and their mitochondria (Martin et al., 2015). Cells that lacked mitochondria never attained the complexity seen in eukaryotes. Hence, true intermediates to this transition from prokaryotes to eukaryotes are not available. The number of genes within mitochondria varies from five to over a hundred in different eukaryotes (Bevan and Lang, 2004). Species with a higher number of genes in the mitochondria provide a snapshot of the endocytosed bacteria-like ancestral entity. The gene-rich mitochondrial genomes of Jakobid protists are models to study the evolution of mitochondria (Burger et al., 2013). Although *TIMM21* homologs are present in the genome of the Jakobid *Andalucia godoyi*, the *COA1/MITRAC15* gene is missing (Gray et al., 2020). The single-copy homologs of *TIMM21* and *COA1/MITRAC15* in bacterial species and Jakobid protist mitochondria suggest that duplication of *TIMM21* might have occurred during the movement of *TIMM21* homologs from the mitochondria to the nucleus. A sampling of more Jakobid genomes might resolve the timing of duplication of *TIMM21* to *COA1/MITRAC15*.

The *COA1/MITRAC15* gene has undergone subsequent duplication events in carnivores and primates. The prevalence of such duplication events suggests that either a higher *COA1*/*MITRAC15* protein dosage is not harmful or sophisticated regulatory machinery to maintain the correct dosage exists. Genes with duplicated copies have greater flexibility for subfunctionalization or neofunctionalisation (Taylor and Raes, 2004). In contrast to gene duplication, the origin of new splice-isoforms increases the transcriptome complexity without increasing the gene count. The evolution of phenotypic novelty through alternative splicing has received greater attention thanks to the availability of large-scale transcriptomic and proteomic datasets in diverse species (Bush et al., 2017). While positive selection has a role in specific examples of alternative splicing (Parker et al., 2014; Ramensky et al., 2008), the vast majority of splicing is probably noisy, and neutral processes may explain its evolution (Pickrell et al., 2010). Alternative splicing also reduces premature protein truncation due to purifying selection (Xing and Lee, 2004). In the case of felid species, the alternative splicing of the third exon (see **Fig. 2**) may have evolved in response to the gene-disrupting changes. Verifying the relevance of the alternative splicing observed at the transcriptional level would require further scrutiny of the protein level isoforms of the *COA1/MITRAC15* gene in felid and canid species. In primate species, the potential addition of the extra coding-exon occurs by a shift of the start codon into the untranslated region. Such changes at the reading frame termini occur when the gene is under relaxed selective constraints (Shinde et al., 2019). Acquisition of novel protein-coding sequences through changes in the exon length is also known to occur (Kishida et al., 2018). We speculate that drastic lineage-specific changes in purifying selection have allowed for changes in intron-exon structure resulting in the evolution of new splice-isoforms of *COA1/MITRAC15*.

Gene loss can be dealt with through compensation from another gene (Xiong and Lei, 2020) or is associated with a biological pathway rewiring (Vijay, 2020). Large-scale changes in gene content are associated with major evolutionary transitions that drastically alter the fitness landscape. Prominent examples of such shifts are the origin of flight in birds (Meredith et al., 2014) and the movement of mammals from land to water seen in cetaceans (Huelsmann et al., 2019). Recurrent gene loss events following relaxed selective constraint in various other lineages have also been documented (Schneider et al., 2019; Sharma and Hiller, 2018; Valente et al., 2020). Loss of genes in the Galliform lineage while being intact in the Anseriformes lineage has been linked to differences in the immune response of these clades (Barber et al., 2010; S. Sharma et al., 2020). The *COA1/MITRAC15* gene is not known to have any obvious immune functions, and its loss in Galliform birds appears to be a consequence of relaxed selection on the OXPHOS pathway. Our computational analysis of more than 200 vertebrate genomes has found that the *COA1/MITRAC15* gene is intact and transcribed in most species, except for the Cheetah, Galliformes, rodents, and marsupial species. Notably, the detailed investigation of the *COA1/MITRAC15* gene in other bird species that are flightless or have a limited ability to fly has found an intact transcribed gene. Therefore, the loss of the *COA1/MITRAC15* gene appears to be associated with changes in skeletal muscle fiber composition. The prominent role of mitochondria in skeletal muscles is evident from diseases of the muscle tissue caused by defects in mitochondrial genes (Gan et al., 2018).

The correlation between recurrent gene loss and the presence of specific phenotypes has provided crucial insights into the evolution of these traits. Stomach loss in gnathostomes co-occurs with the loss of several genes that code for digestive enzymes (Castro et al., 2013). The loss of ketogenesis has occurred through the recurrent loss of the *HMGCS2* gene (Jebb and Hiller, 2018). Gene losses associated with dietary composition, the patterns of feeding, and gut microbiomes have also been identified (Hecker et al., 2019). Recurrent loss of Toll-like receptors (TLRs), which play prominent roles in the innate immune system, is associated with impaired ability to detect extracellular flagellin (V. Sharma et al., 2020). The repeated loss of the cortistatin gene is related to modifications in the circadian pathway (Valente et al., 2020). In the *COA1/MITRAC15* gene, we record the independent occurrence of gene disrupting changes in closely related species of Galliformes and rodents. However, we can rule out the possibility of a common regulatory mutation that initially resulted in the loss of gene expression followed by the independent accumulation of the gene disrupting changes that we observe. Our hypothesis predicts gene loss following changes in skeletal muscle fiber composition. The *COA1/MITRAC15* gene does not directly alter the muscle fiber composition and might have subsequently experienced relaxed selective constraint due to increased glycolytic muscle fibers. Hence, it is tempting to speculate that the independent gene disrupting changes reflect recurrent gene loss events. However, the mechanistic basis of changes in muscle fiber composition between species is yet to be understood. Identifying the genetic changes that determine muscle fiber composition and the sequence of events would provide greater clarity regarding when and why the *COA1/MITRAC15* gene loss occurred.

Evolutionary Breakpoint Regions (EBRs) are genomic regions that have undergone one or more structural changes resulting in altered karyotypes between lineages (Lemaitre et al., 2009). Recurrent non-random structural changes at the same regions in multiple lineages potentially occur due to the presence of repeat elements (Farré et al., 2016; Schibler et al., 2006), chromosome fragile sites (Durkin and Glover, 2007; Ruiz-Herrera et al., 2006, 2005), nucleotide composition, methylation level (Carbone et al., 2009) and chromatin state (Boteva et al., 2020; Huvet et al., 2007). However, the prevalence of EBRs and their relevance to evolutionary processes has been the focus of considerable debate (Alekseyev and Pevzner, 2007; Peng et al., 2006; Trinh et al., 2004). Several lineage-specific gene loss events near EBRs in rodents are due to chromosomal rearrangements (Capilla et al., 2016; Fitzgerald and Bateman, 2004). Notably, one of these lost genes, *STK17A,* is located adjacent to the *COA1/MITRAC15* gene. The co-occurrence of an EBR with putative *COA1/MITRAC15* gene loss in rodents and marsupials is very intriguing. However, rodent genomes have mutational hotspots with high lineage-specific gBGC resulting in a substantial gene sequence divergence (Hargreaves et al., 2017). Such highly diverged orthologs can be challenging to identify due to difficulties in sequencing high GC regions. In *COA1/MITRAC15*, the magnitude of gBGC is relatively low, especially in rodents. Moreover, we find remnants of *COA1/MITRAC15* in several post-EBR species that suggest actual gene loss, at least in rodents. Several pre-EBR rodent species have also independently accumulated gene disrupting mutations in the *COA1/MITRAC15* gene. Hence, the *COA1/MITRAC15* gene appears to be under relaxed selective constraint even before the occurrence of the EBR.

Species with exceptionally large body sizes or extremely long lifespans have a greater number of cell divisions. An increment in the number of cell divisions enhances cancer risk. However, paradoxically, large-bodied animals like elephants and whales do not have a higher incidence of cancer (Peto et al., 1975; Tollis et al., 2017). Cancer resistance due to lineage-specific changes in gene content may explain this paradox (Caulin et al., 2015; Caulin and Maley, 2011; DeGregori, 2011). While specific genetic changes in mammalian species lead to cancer resistance (Tollis et al., 2019; Vazquez et al., 2018), the reasons for lower cancer incidence in birds compared to mammals are mostly unexplored (Møller et al., 2017). Interestingly, the *COA1/MITRAC15* gene is an oncogene with a role in colorectal cancer (Xue et al., 2020), and its loss could reduce cancer risk. Silencing of *COA1/MITRAC15* by miRNAs strongly suppresses Giant cell tumors of the bone (Fellenberg et al., 2016; Herr et al., 2017). Our discovery of *COA1/MITRAC15* gene loss in Galliformes sets a precedent for the indisputable identification of gene loss events in birds and might reveal other oncogenes which are lost. We also identify *COA1/MITRAC15* gene loss in the beaver and naked mole-rat genomes, species that are models to study longevity (Zhou et al., 2020). High-quality near-complete vertebrate genomes with very few errors will further aid in the large-scale identification of gene loss events across the vertebrate phylogeny (Rhie et al., 2021).

## 5. Conclusions

*COA1/MITRAC15* is a distant homolog of the *TIMM21* gene that has undergone recurrent gene loss in several Galliform and rodent species. Gene loss events occurred between 15 MYA and 46 MYA in Galliform species and between 2 MYA and 30 MYA in rodents. The gene loss event occurs in species that rely primarily on glycolytic muscle fibers to achieve short bursts of activity. We show that *COA1/MITRAC15* and the adjacent *STK17A* gene are located at an Evolutionary Breakpoint Region (EBR) and are missing from the genomes of several rodent species following chromosomal rearrangement events. Pseudogenic and functional copies of *COA1/MITRAC15* are present in carnivores and primates, with the functional copy diverging in its intron-exon structure. Prevalence of repeated gene loss and duplication events in the history of *COA1/MITRAC15* not only demonstrates the dispensability of this gene but also hints at its ability to provide fitness increases in a context-dependent manner.

## Acknowledgment

We thank the Council of Scientific & Industrial Research for fellowship to SSS and the Ministry of Human Resource Development for fellowship to LT and SS and University Grants commission to AS. NV has been awarded the Innovative Young Biotechnologist Award 2018 from the Department of Biotechnology (Government of India).

## Availability of data

All data associated with this study are available in the Supplementary Materials.

