## Supplementary Figures for "Recurrent erosion of *COA1/MITRAC15* demonstrates gene dispensability in oxidative phosphorylation": Supplementary_Figures.pptx

### Slide 1
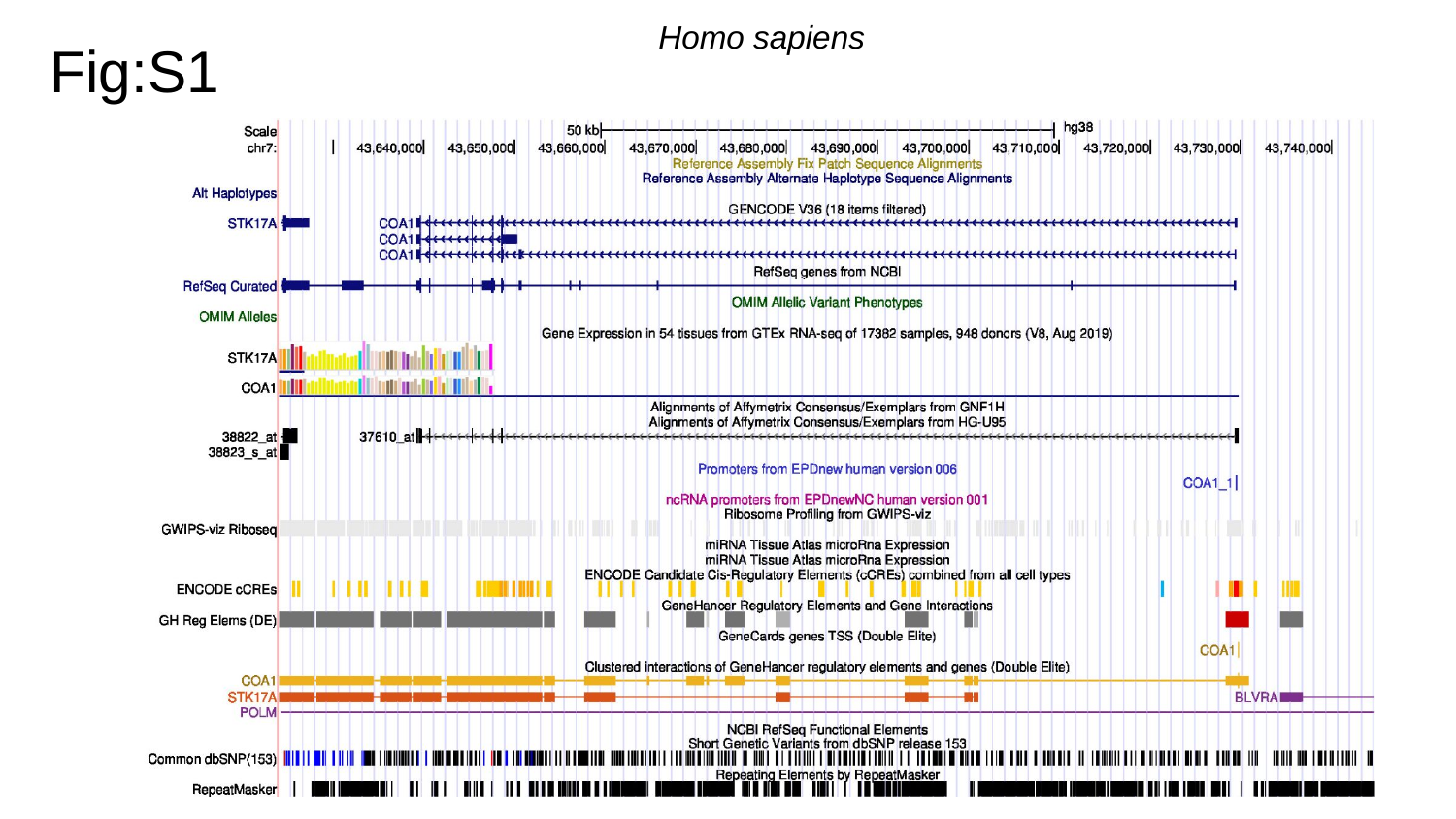

Homo sapiens
Fig:S1

### Slide 2
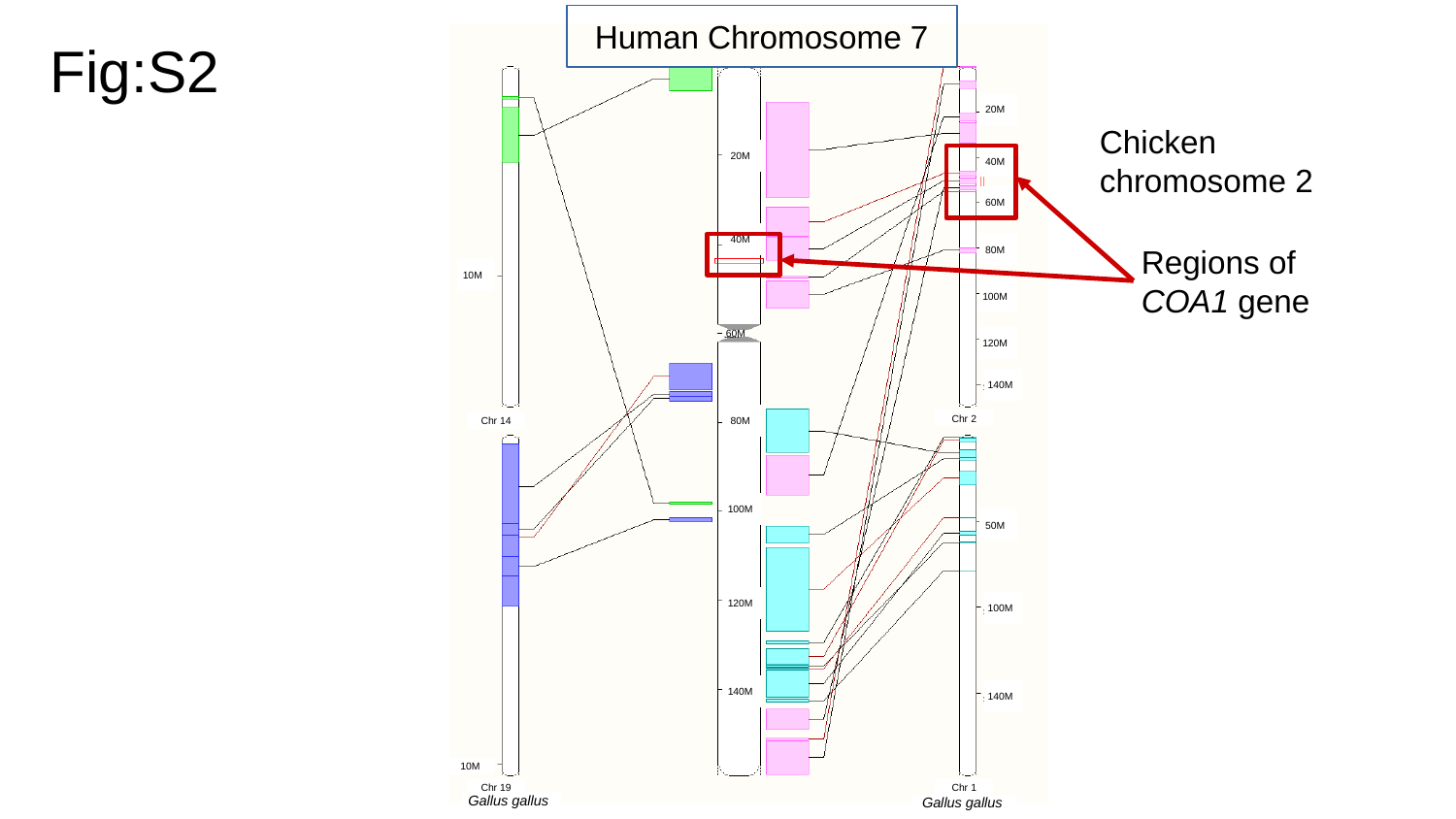

Human Chromosome 7
Fig:S2
20M
Chicken chromosome 2
20M
40M
60M
Regions of COA1 gene
40M
80M
10M
100M
120M
60M
140M
80M
Chr 2
Chr 14
100M
50M
120M
100M
140M
140M
10M
Chr 19
Chr 1
Gallus gallus
Gallus gallus

### Slide 3
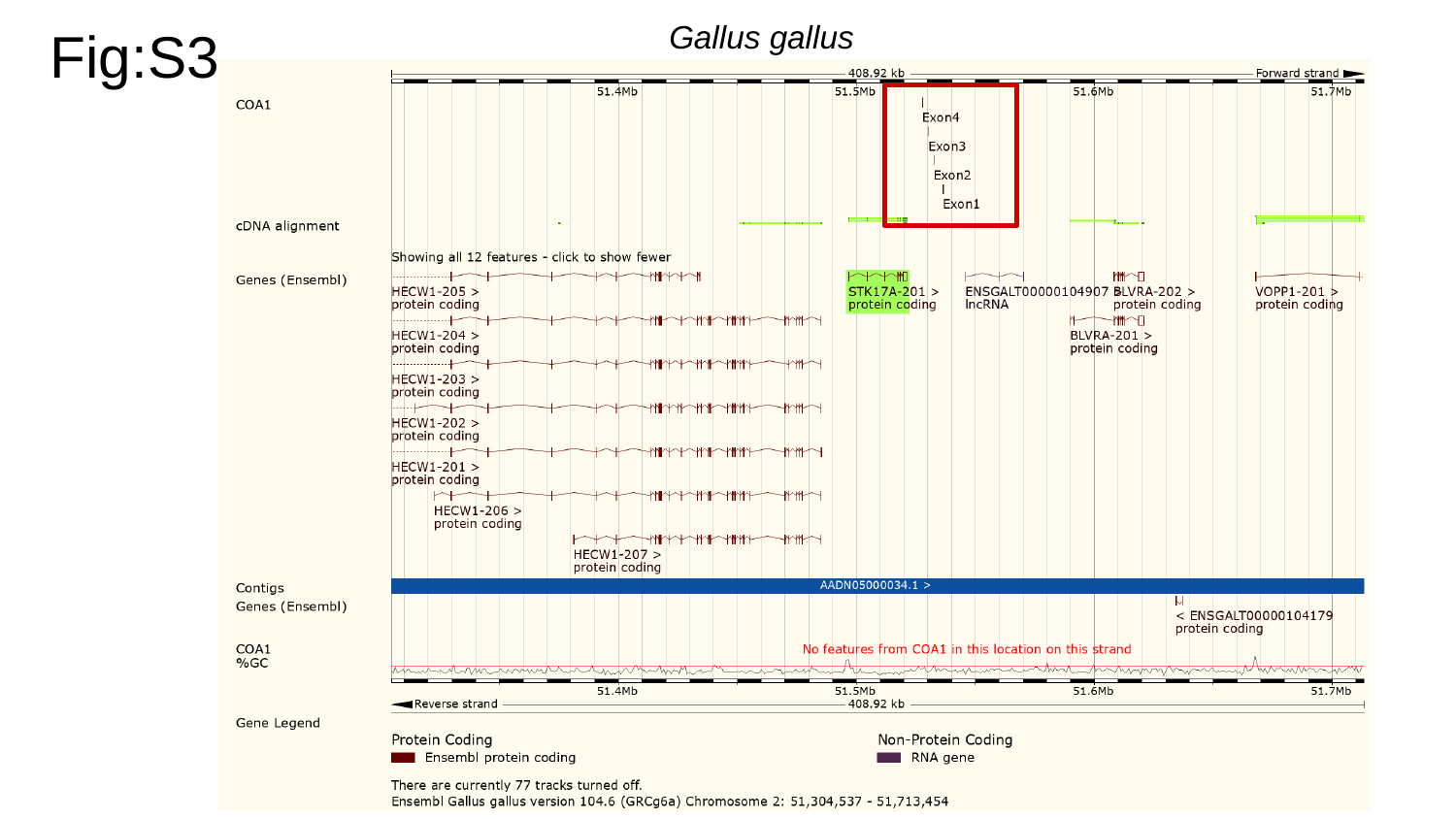

Fig:S3
Gallus gallus

### Slide 4
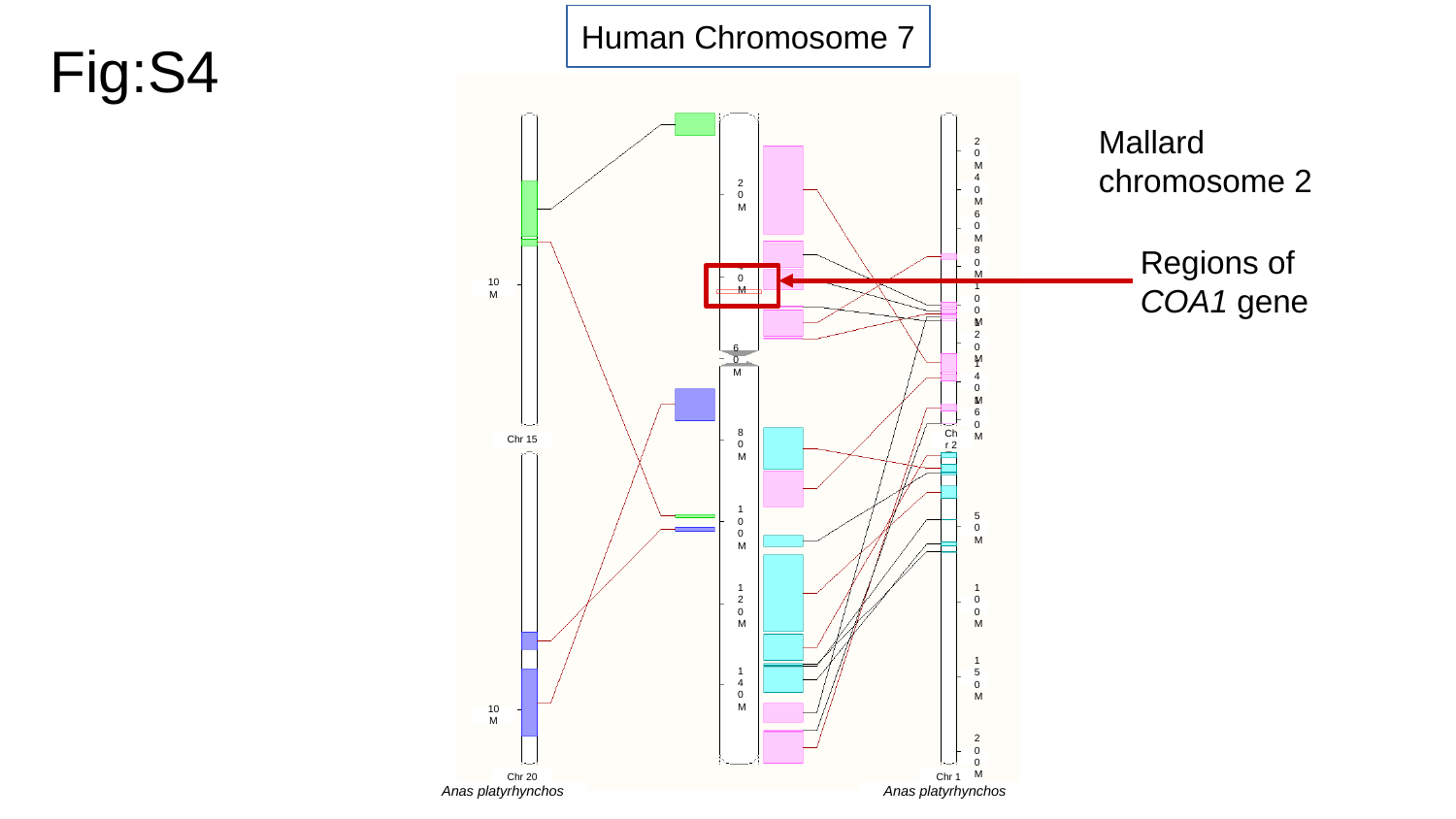

Human Chromosome 7
Fig:S4
Mallard chromosome 2
20M
40M
20M
60M
Regions of COA1 gene
80M
40M
10M
100M
120M
60M
140M
160M
Chr 15
Chr 2
80M
100M
50M
120M
100M
150M
140M
10M
200M
Chr 20
Chr 1
Anas platyrhynchos
Anas platyrhynchos

### Slide 5
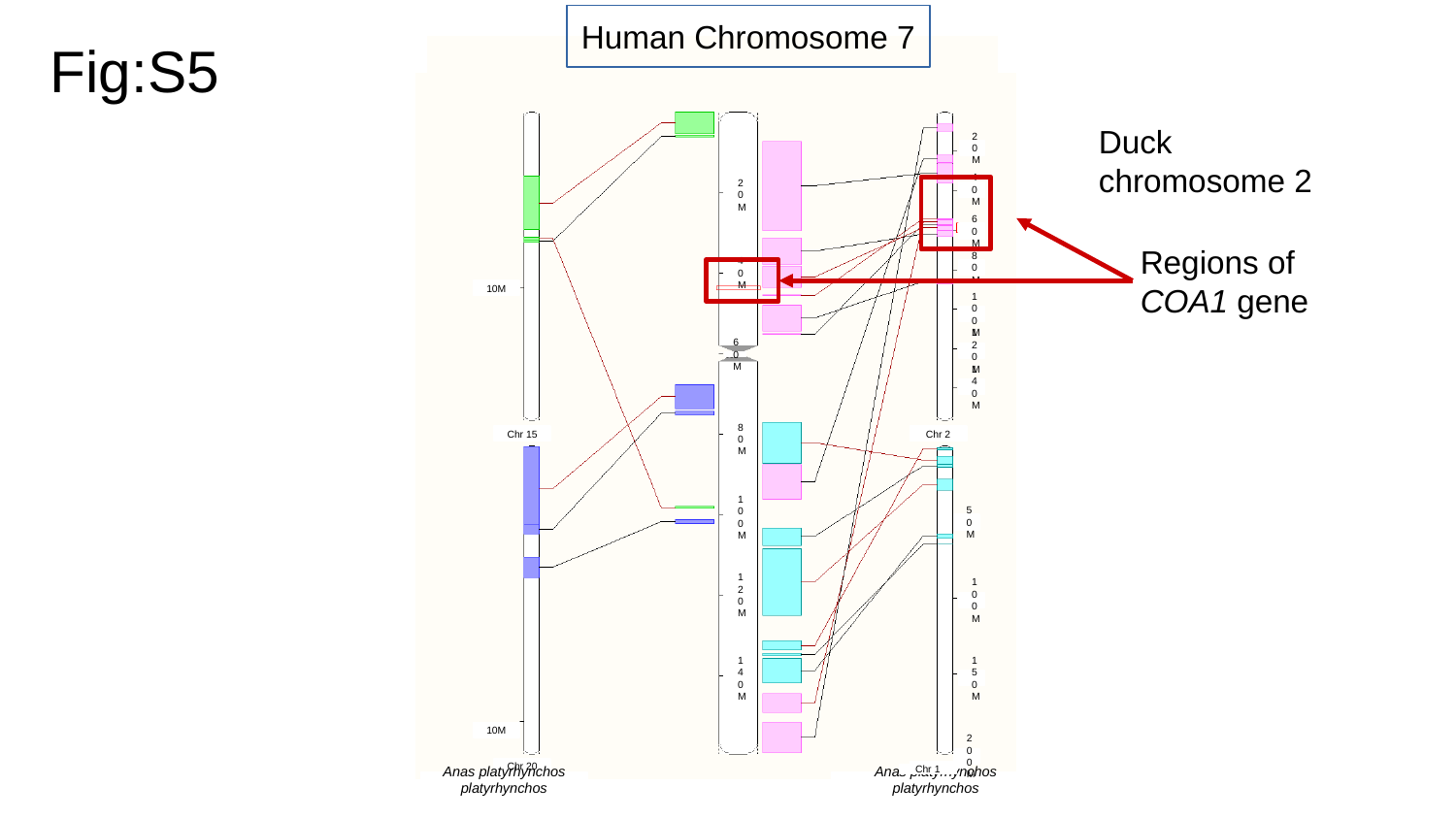

Human Chromosome 7
Fig:S5
Duck chromosome 2
20M
40M
20M
Regions of COA1 gene
60M
80M
40M
10M
100M
120M
60M
140M
Chr 15
Chr 2
80M
100M
50M
120M
100M
140M
150M
10M
200M
Chr 20
Chr 1
Chr 1
Anas platyrhynchos platyrhynchos
Anas platyrhynchos platyrhynchos

### Slide 6
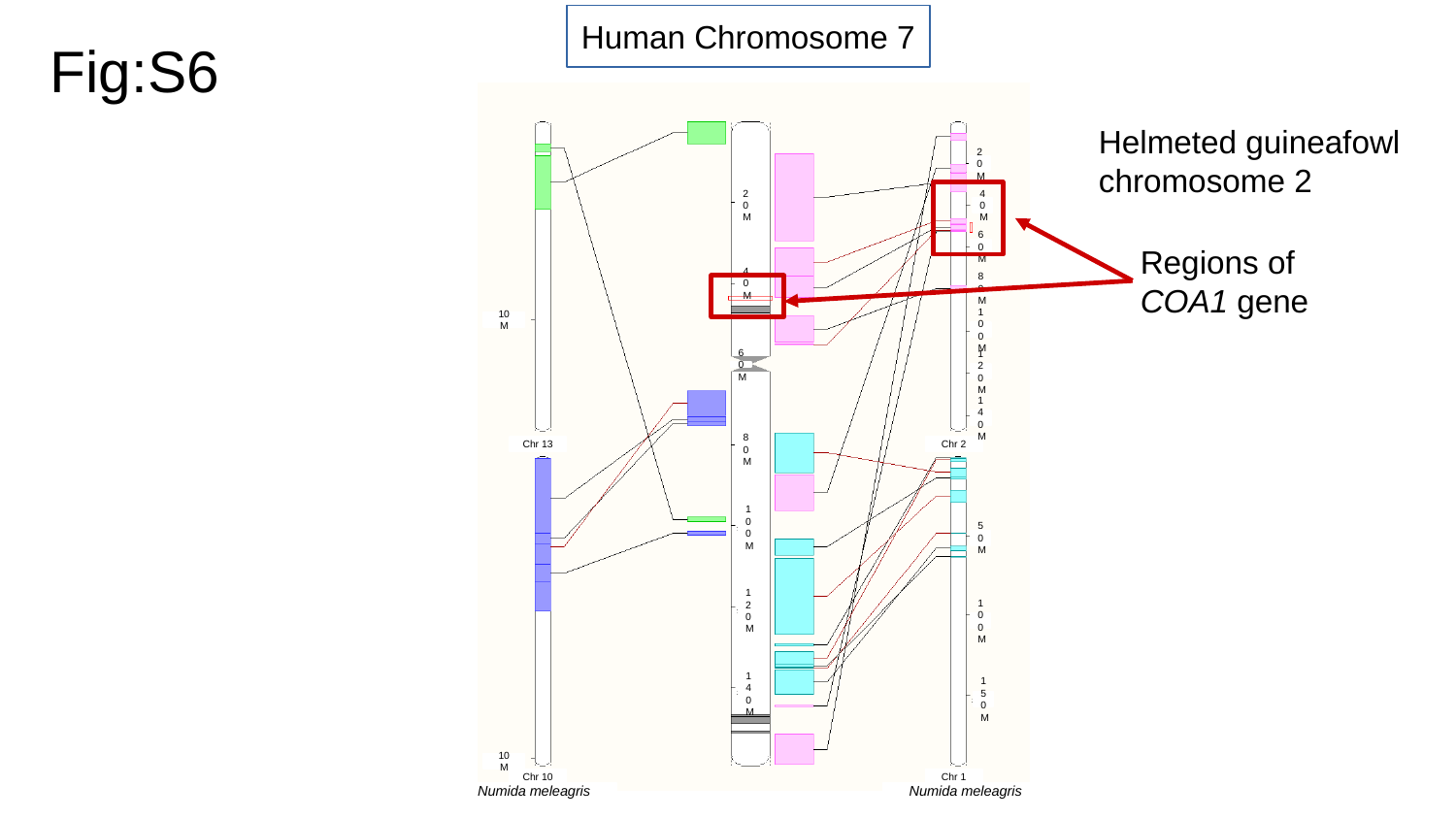

Human Chromosome 7
Fig:S6
Helmeted guineafowl chromosome 2
20M
20M
40M
Regions of COA1 gene
60M
40M
80M
10M
100M
60M
120M
140M
Chr 13
Chr 2
80M
100M
50M
120M
100M
140M
150M
10M
Chr 10
Chr 1
Numida meleagris
Numida meleagris

### Slide 7
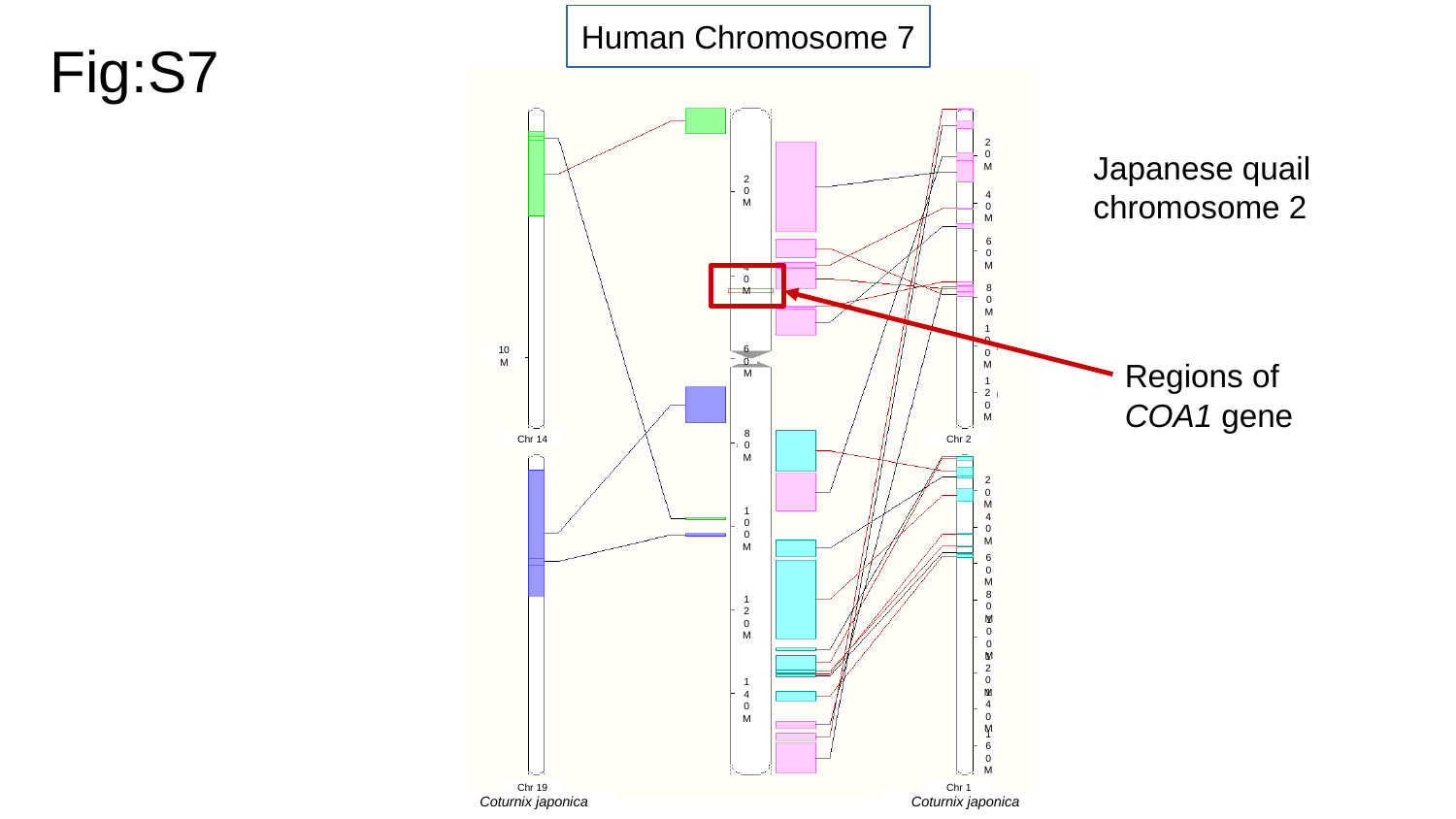

Human Chromosome 7
Fig:S7
Japanese quail chromosome 2
20M
20M
40M
60M
40M
80M
Regions of COA1 gene
100M
10M
60M
120M
Chr 14
Chr 2
80M
20M
100M
40M
60M
80M
120M
100M
120M
140M
140M
160M
Chr 19
Chr 1
Coturnix japonica
Coturnix japonica

### Slide 8
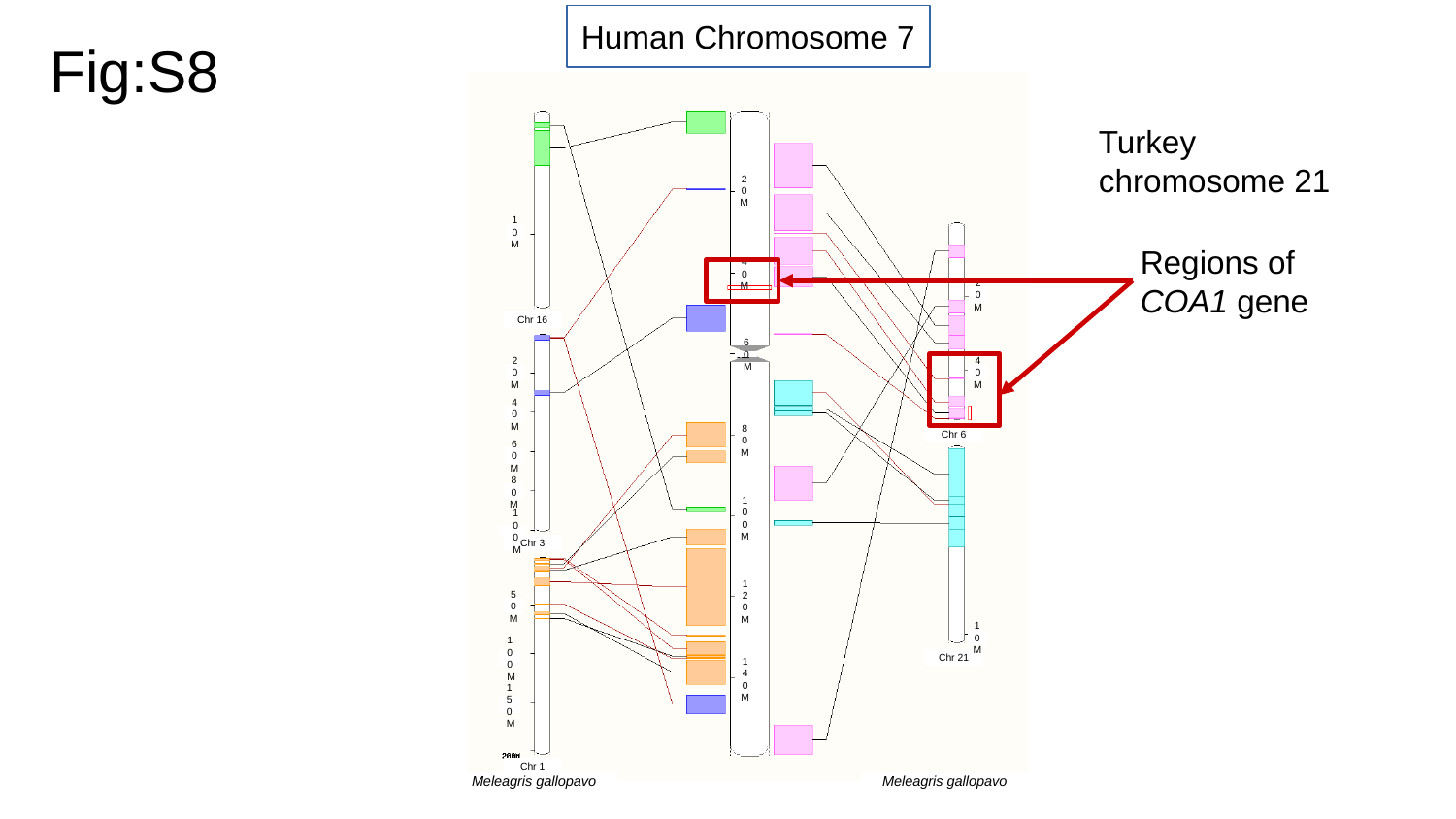

Human Chromosome 7
Fig:S8
Turkey chromosome 21
20M
Regions of COA1 gene
10M
40M
20M
Chr 16
60M
20M
40M
40M
Chr 6
80M
60M
80M
100M
100M
Chr 3
120M
50M
10M
Chr 21
100M
140M
150M
Chr 1
Meleagris gallopavo
Meleagris gallopavo

### Slide 9
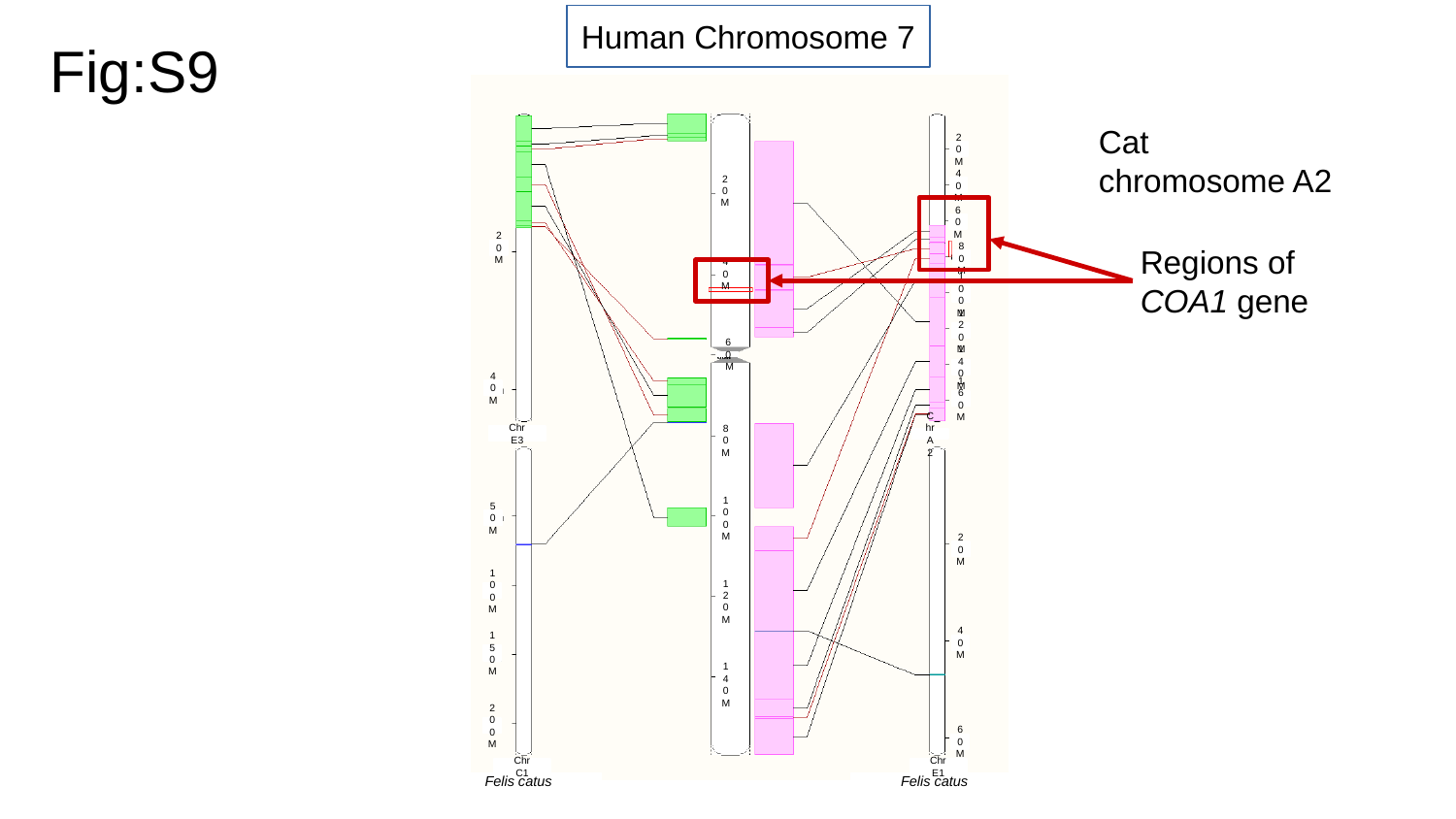

Human Chromosome 7
Fig:S9
Cat chromosome A2
20M
40M
20M
60M
Regions of COA1 gene
20M
80M
40M
100M
120M
60M
140M
40M
160M
Chr E3
Chr A2
80M
50M
100M
20M
100M
120M
40M
150M
140M
200M
60M
Chr C1
Chr E1
Felis catus
Felis catus

### Slide 10
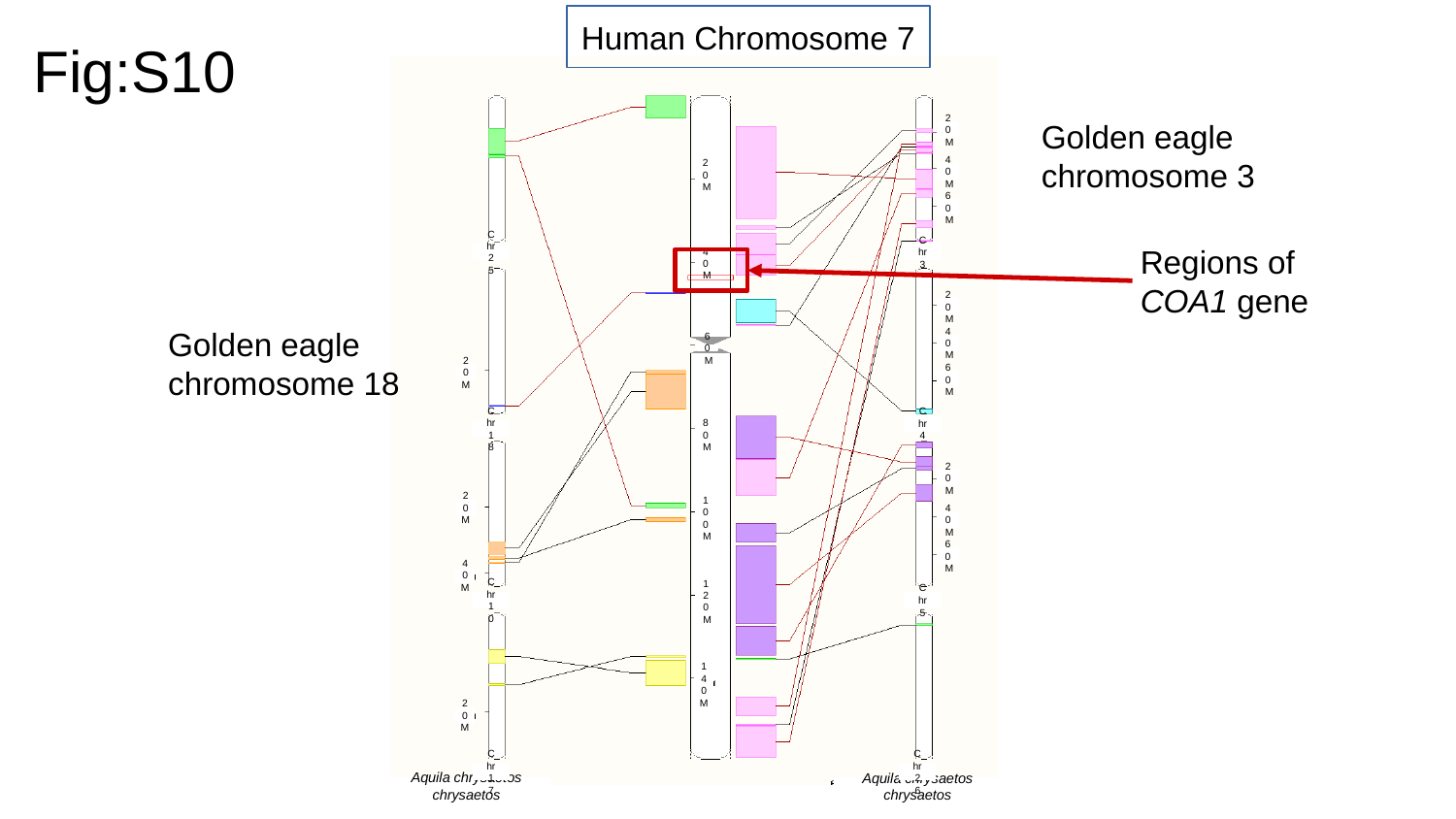

Human Chromosome 7
Fig:S10
Golden eagle chromosome 3
20M
40M
20M
60M
Regions of COA1 gene
Chr 25
Chr 3
40M
20M
Golden eagle chromosome 18
40M
60M
20M
60M
Chr 4
Chr 18
80M
20M
20M
100M
40M
60M
40M
Chr 10
Chr 5
120M
140M
20M
Chr 17
Chr 26
Aquila chrysaetos chrysaetos
Aquila chrysaetos chrysaetos

### Slide 11
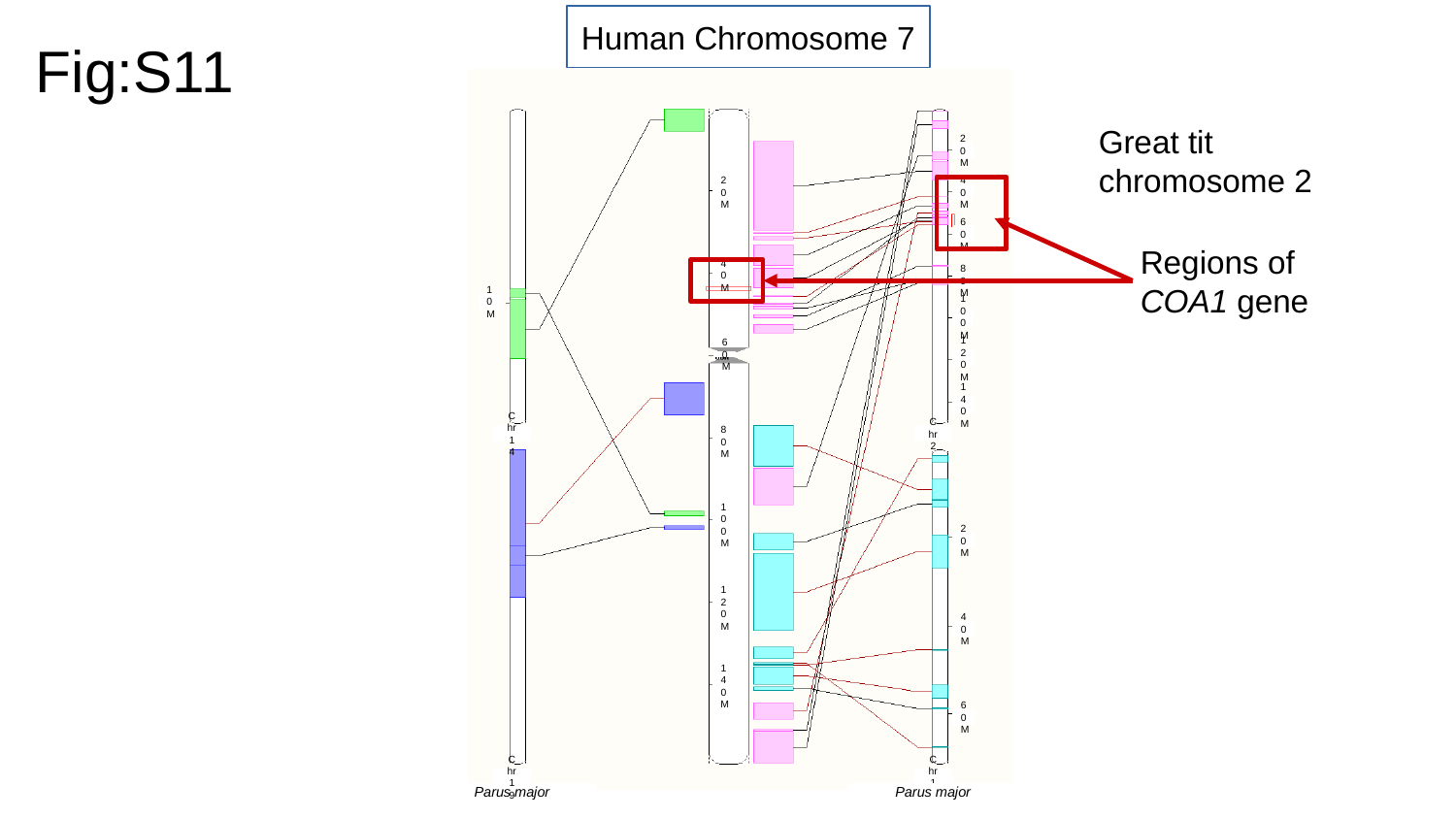

Human Chromosome 7
Fig:S11
Great tit chromosome 2
20M
20M
40M
Regions of COA1 gene
60M
40M
80M
10M
100M
120M
60M
140M
Chr 14
Chr 2
80M
100M
20M
120M
40M
140M
60M
Chr 19
Chr 18
Parus major
Parus major

### Slide 12
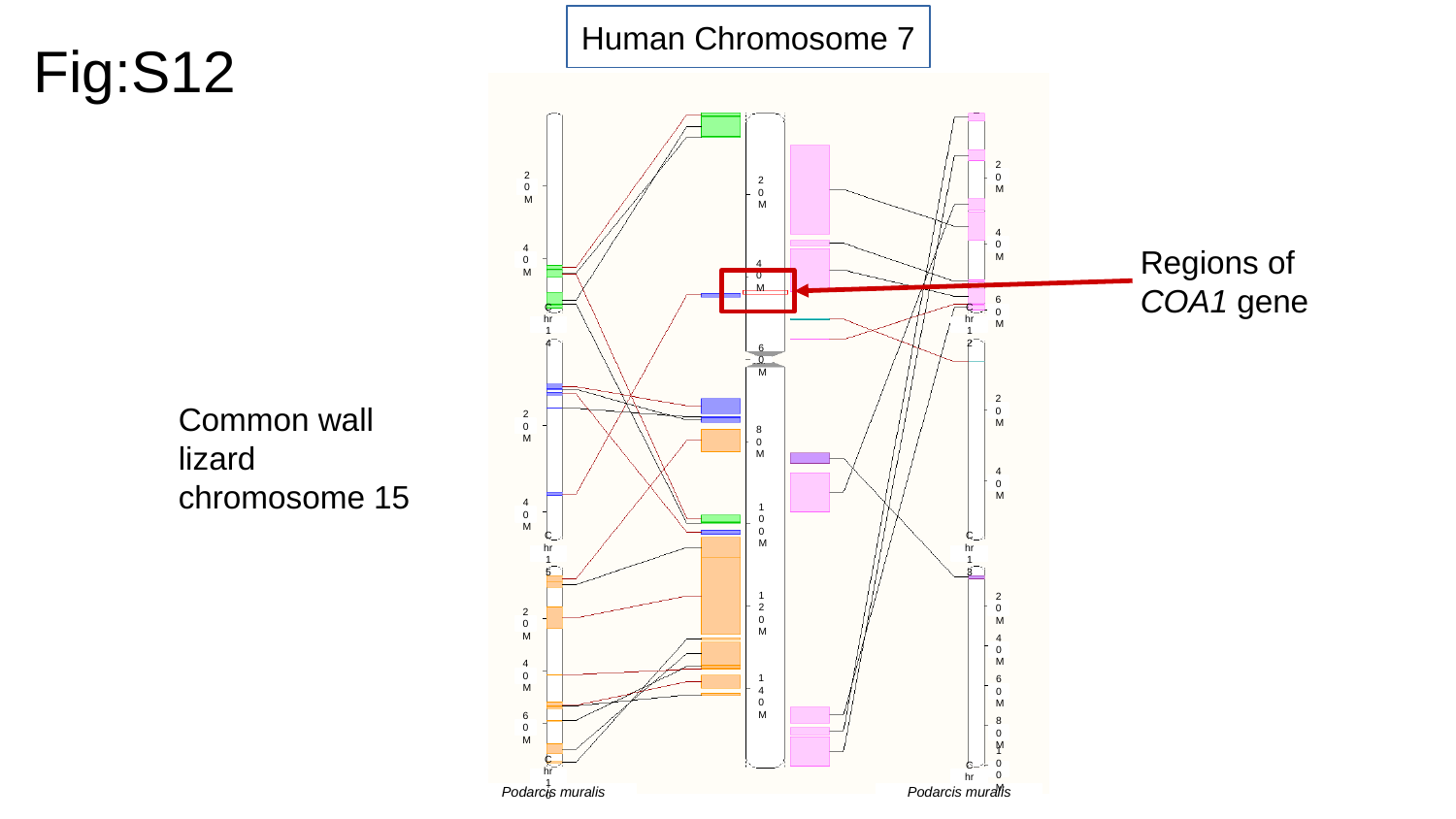

Human Chromosome 7
Fig:S12
20M
20M
20M
Regions of COA1 gene
40M
40M
40M
60M
Chr 14
Chr 12
60M
Common wall lizard chromosome 15
20M
20M
80M
40M
40M
100M
Chr 15
Chr 13
20M
120M
20M
40M
40M
60M
140M
60M
80M
100M
Chr 10
Chr 5
Podarcis muralis
Podarcis muralis

### Slide 13
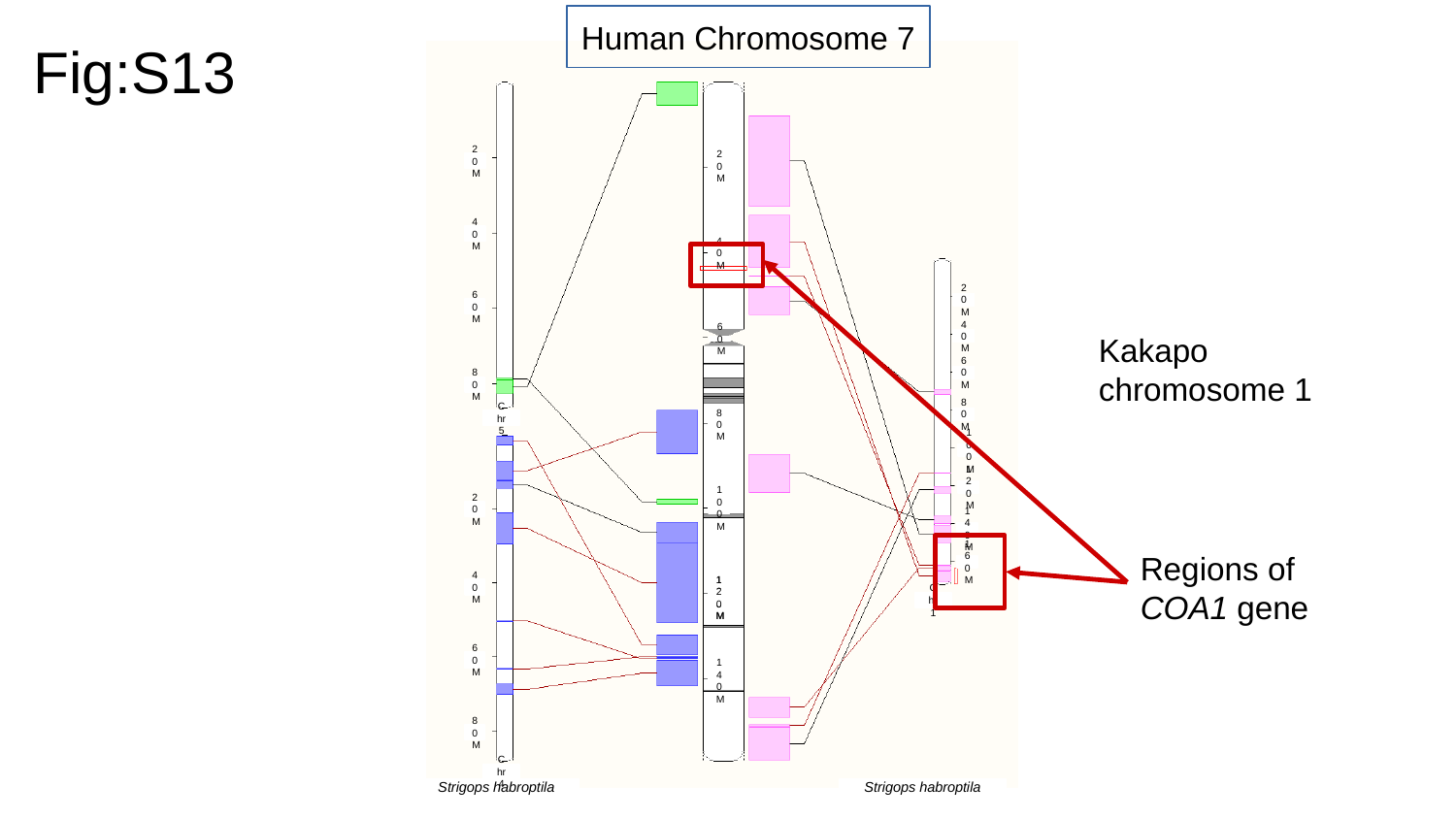

Human Chromosome 7
Fig:S13
20M
20M
40M
40M
20M
60M
Kakapo chromosome 1
40M
60M
60M
80M
80M
Chr 5
80M
100M
120M
20M
100M
140M
Regions of COA1 gene
160M
40M
120M
120M
Chr 1
60M
140M
80M
Chr 4
Strigops habroptila
Strigops habroptila

### Slide 14
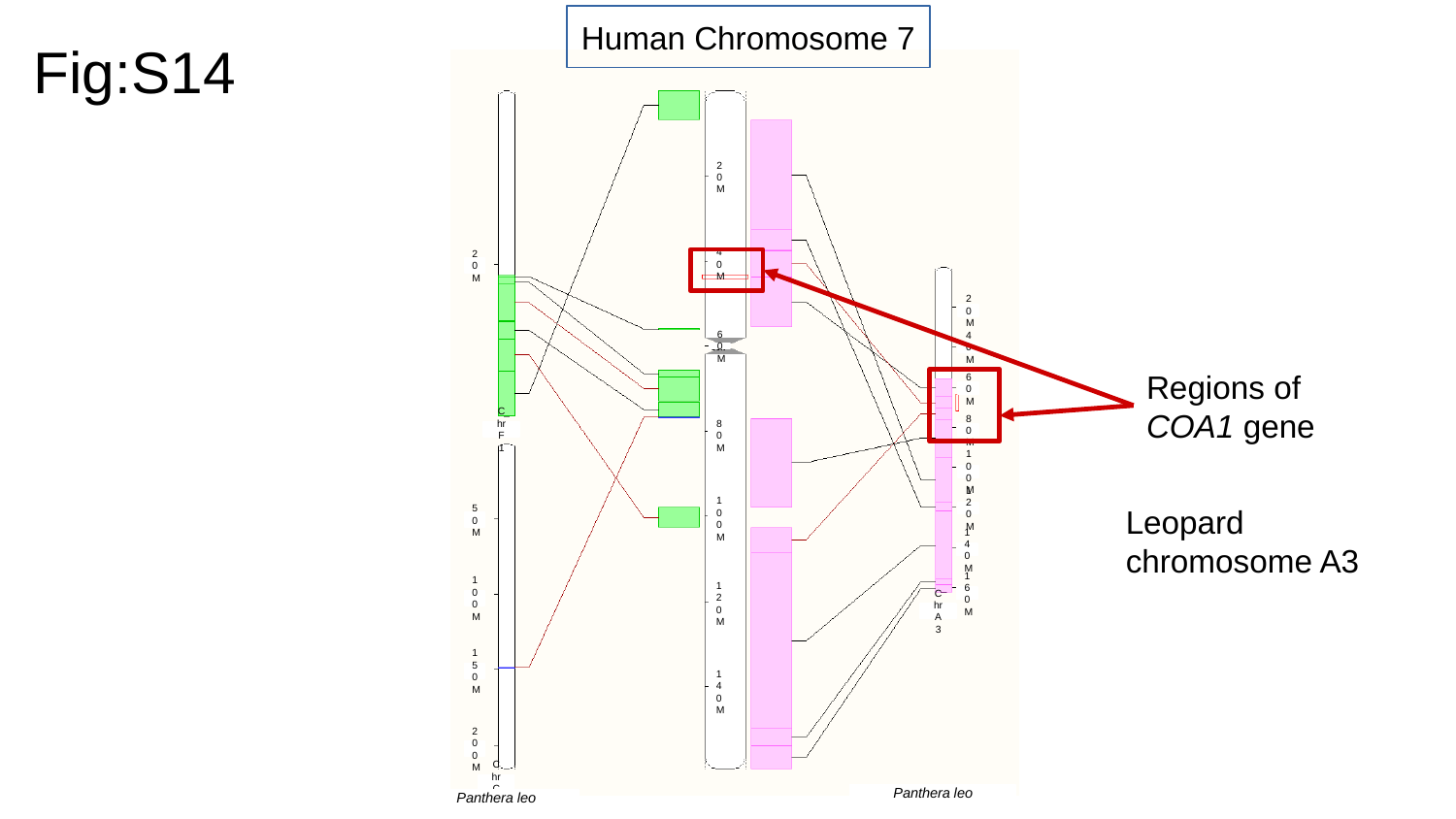

Human Chromosome 7
Fig:S14
20M
20M
40M
20M
40M
60M
Regions of COA1 gene
60M
Chr F1
80M
80M
100M
Leopard chromosome A3
120M
50M
100M
140M
160M
100M
120M
Chr A3
150M
140M
200M
Chr C2
Panthera leo
Panthera leo

### Slide 15
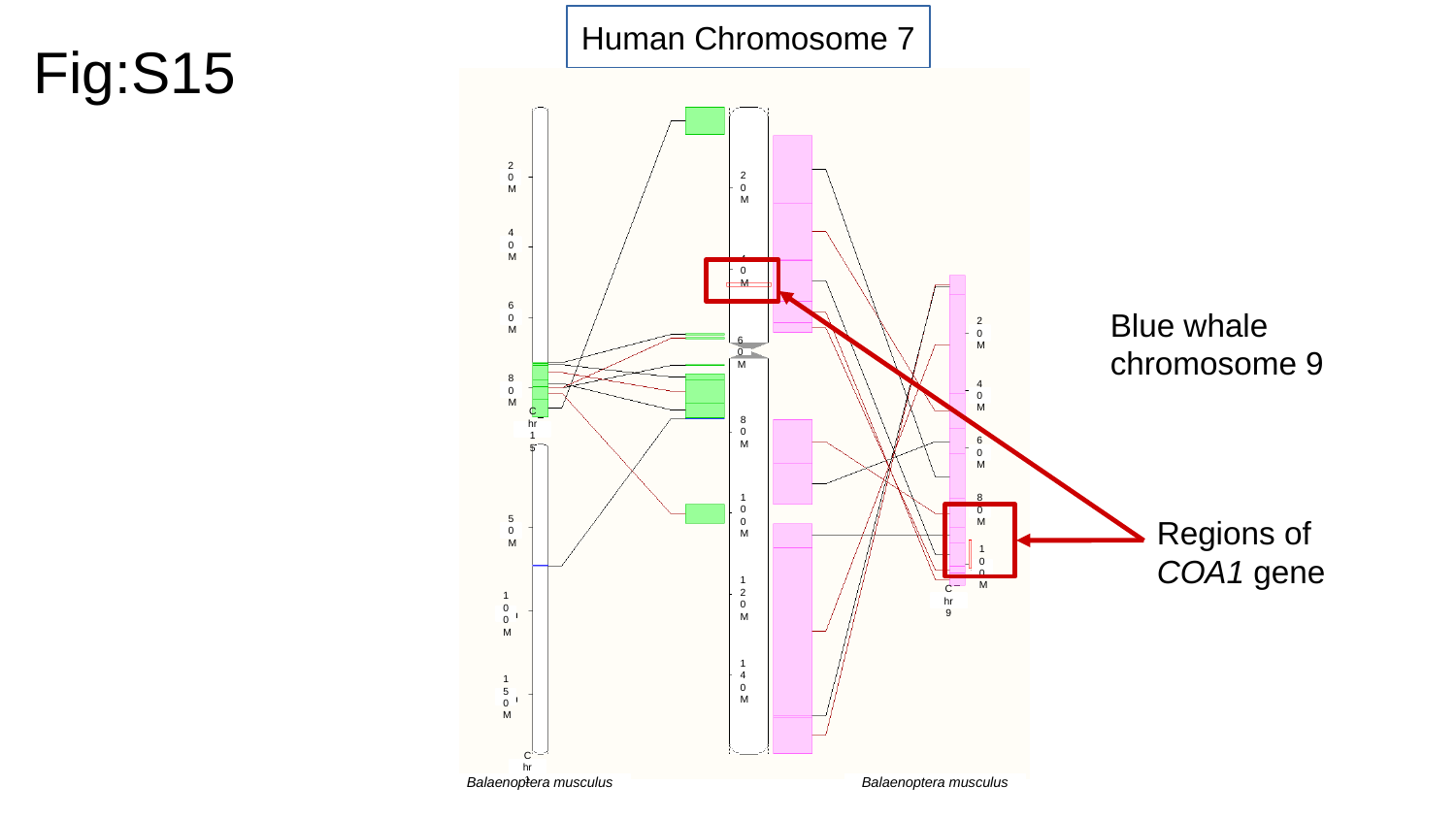

Human Chromosome 7
Fig:S15
20M
20M
40M
40M
Blue whale chromosome 9
60M
20M
60M
80M
40M
Chr 15
80M
60M
Regions of COA1 gene
80M
100M
50M
100M
120M
Chr 9
100M
140M
150M
Chr 1
Balaenoptera musculus
Balaenoptera musculus

### Slide 16
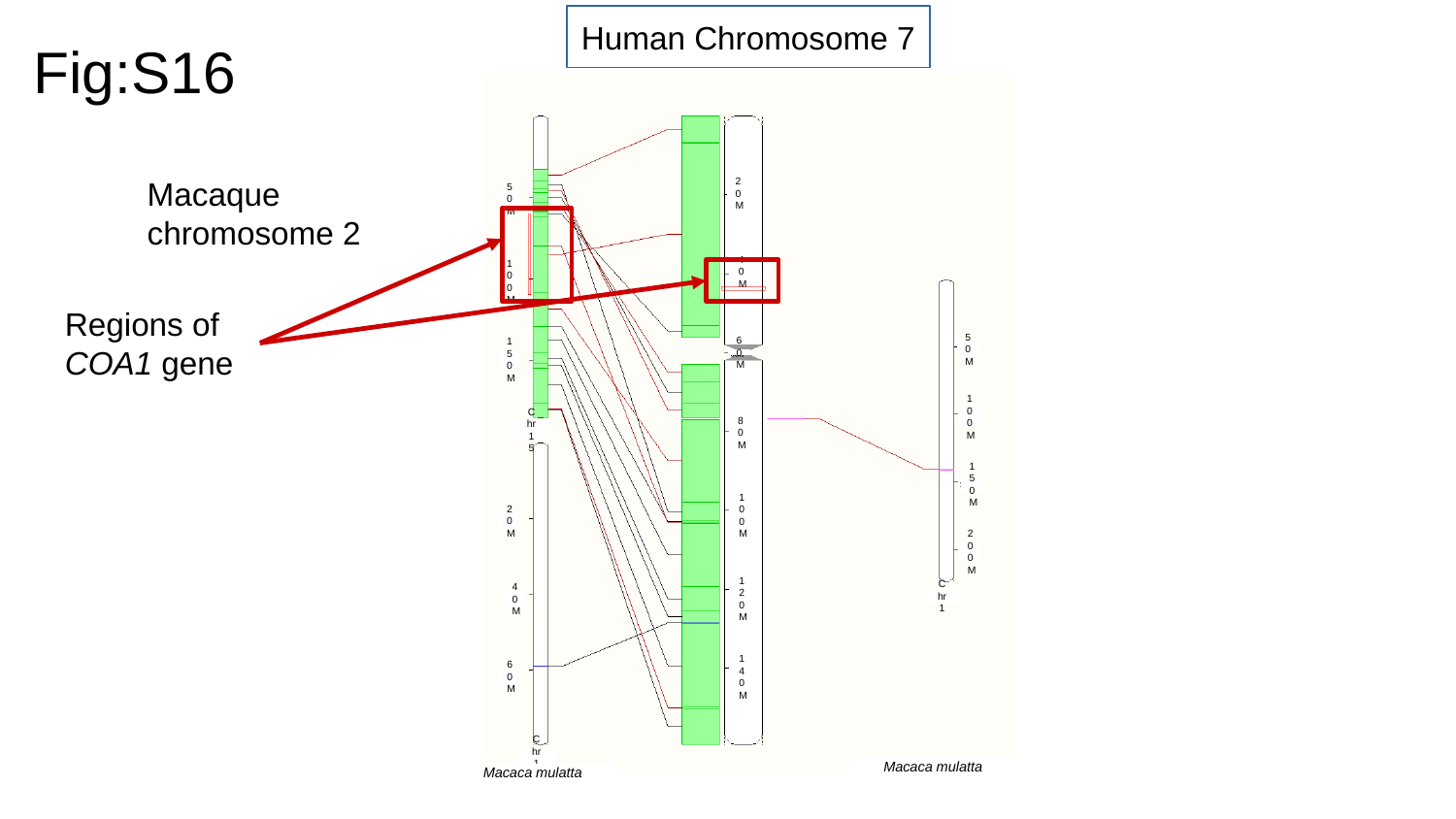

Human Chromosome 7
Fig:S16
Macaque chromosome 2
20M
50M
40M
100M
Regions of COA1 gene
50M
60M
150M
100M
Chr 15
80M
150M
100M
20M
200M
Chr 1
40M
120M
60M
140M
Chr 16
Macaca mulatta
Macaca mulatta

### Slide 17
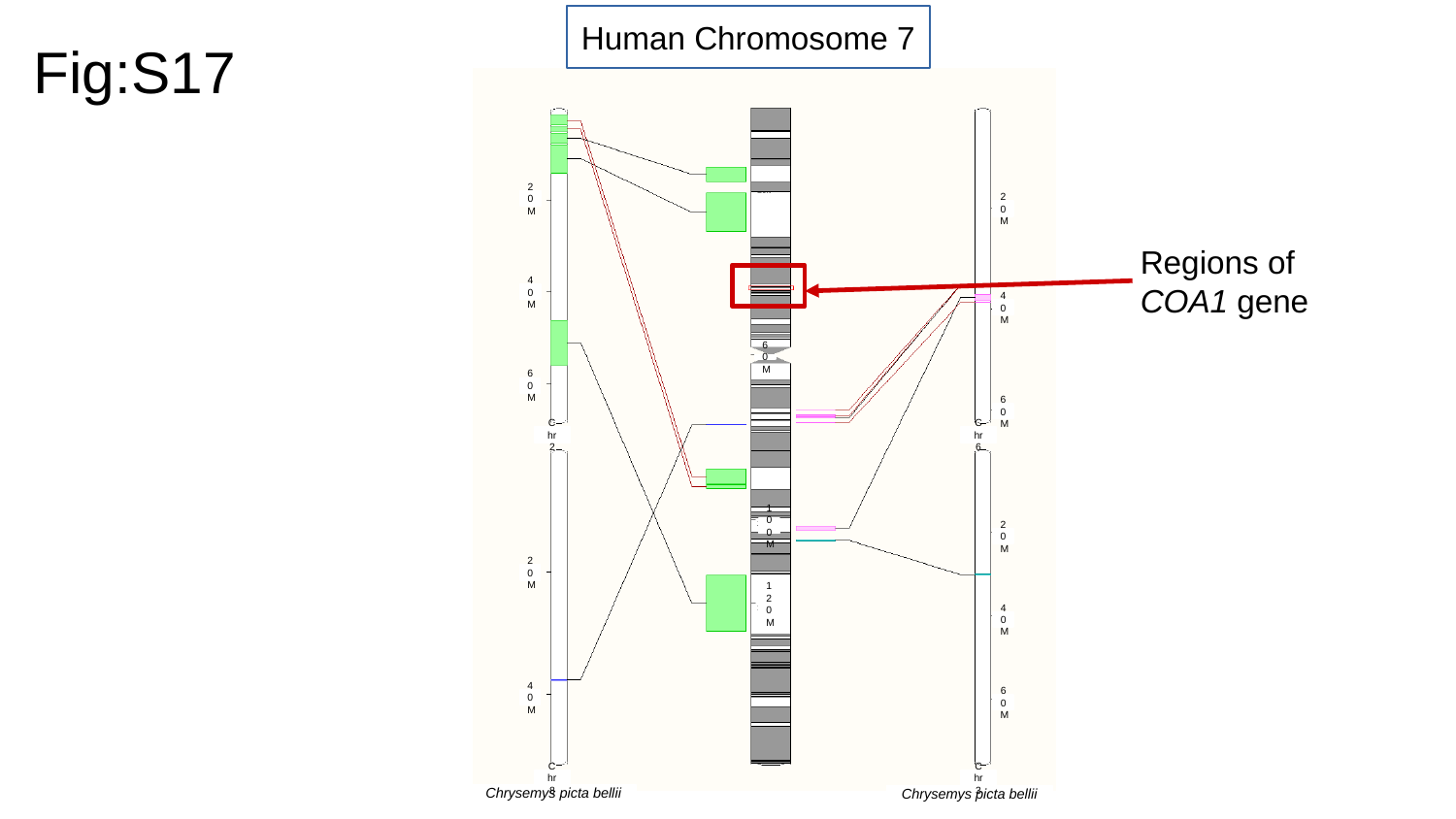

Human Chromosome 7
Fig:S17
20M
20M
Regions of COA1 gene
40M
40M
60M
60M
60M
Chr 2
Chr 6
100M
20M
20M
120M
40M
40M
60M
Chr 8
Chr 3
Chrysemys picta bellii
Chrysemys picta bellii

### Slide 18
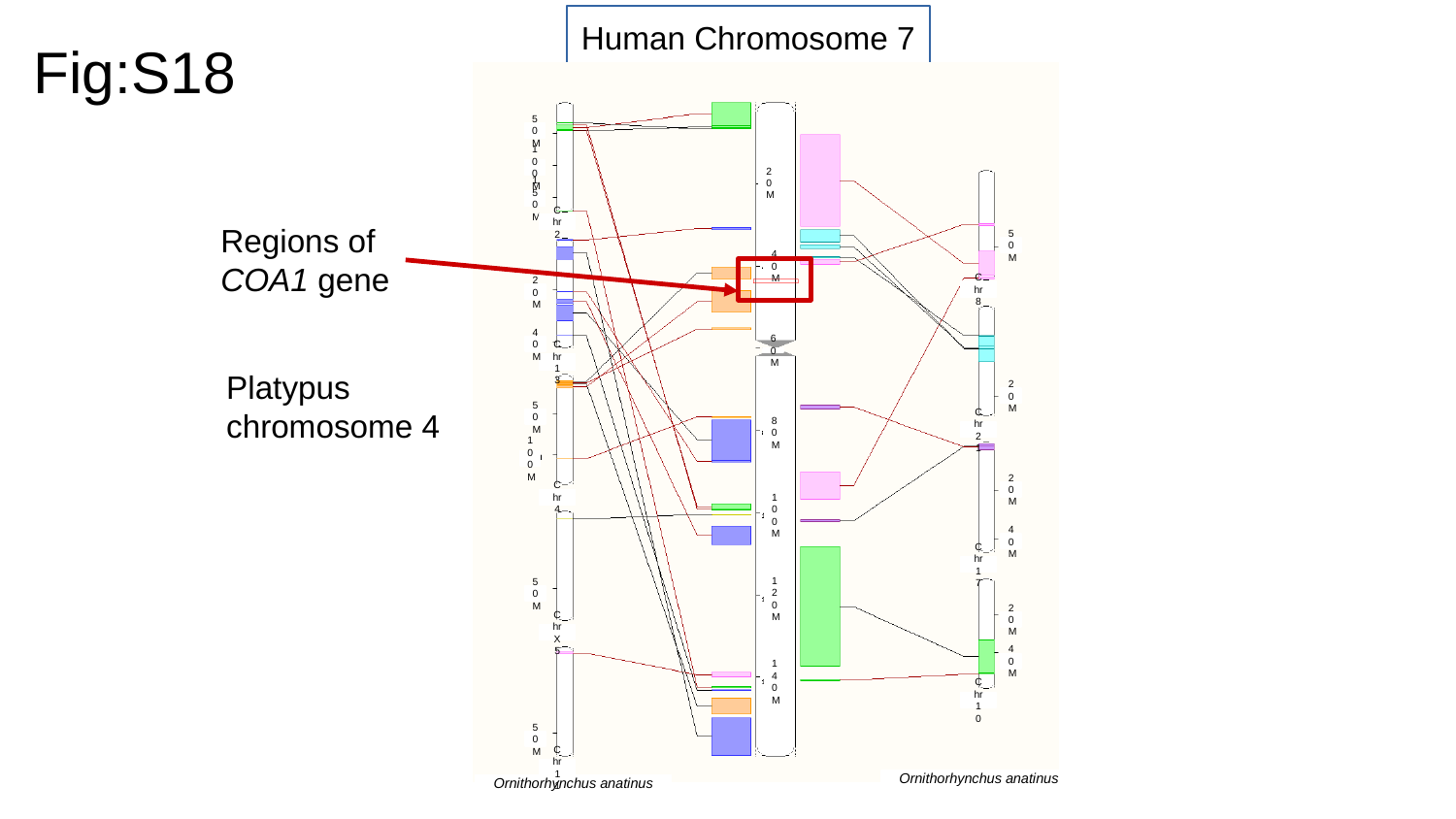

Human Chromosome 7
Fig:S18
50M
100M
20M
150M
Regions of COA1 gene
Chr 2
50M
40M
Chr 8
20M
40M
Platypus chromosome 4
60M
Chr 13
20M
50M
Chr 21
80M
100M
20M
Chr 4
100M
40M
Chr 17
50M
120M
20M
Chr X5
40M
140M
Chr 10
50M
Chr 11
Ornithorhynchus anatinus
Ornithorhynchus anatinus

### Slide 19
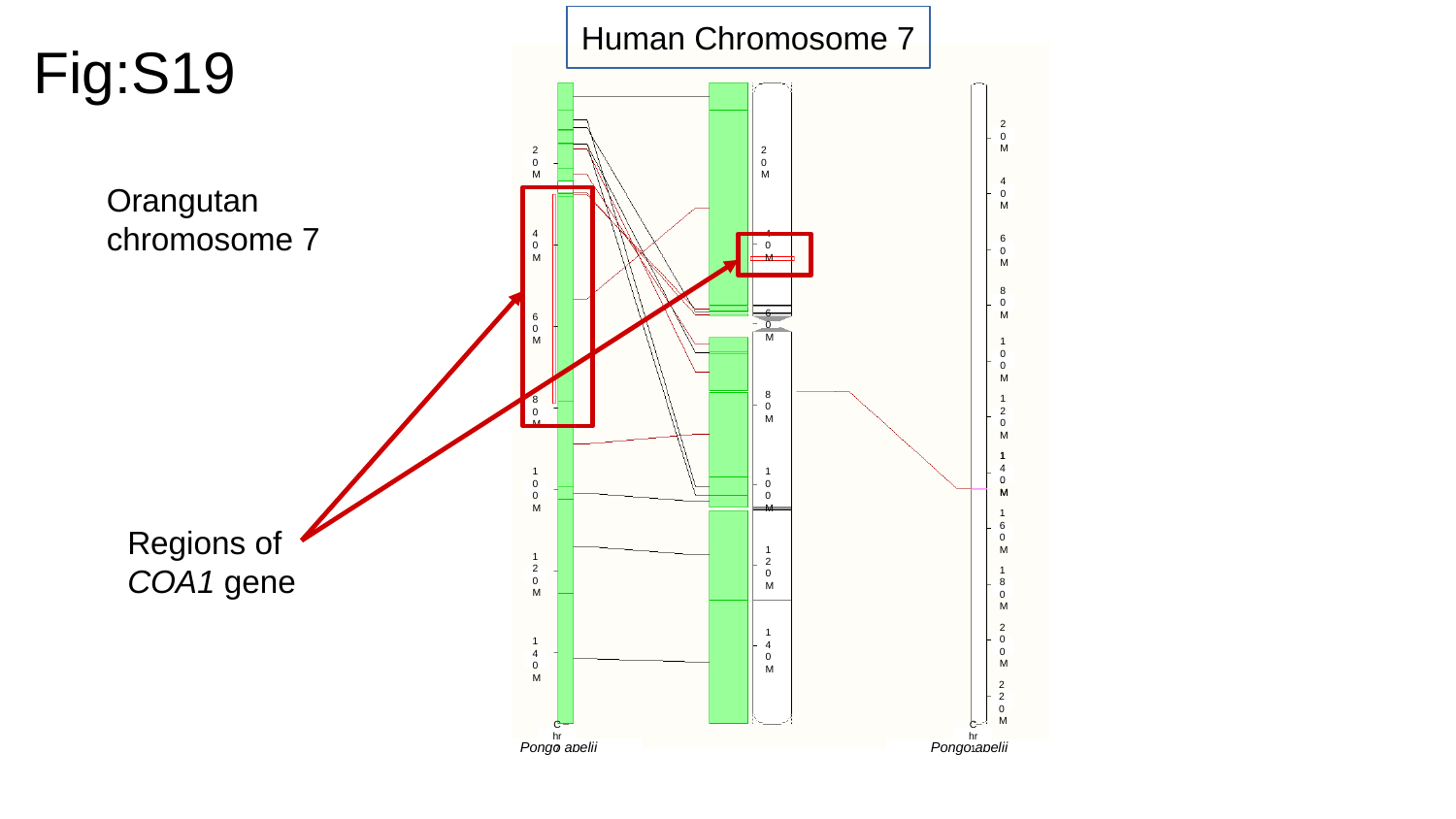

Human Chromosome 7
Fig:S19
20M
20M
20M
Orangutan chromosome 7
40M
40M
40M
60M
80M
60M
60M
100M
80M
80M
120M
140M
140M
100M
100M
Regions of COA1 gene
160M
120M
120M
180M
200M
140M
140M
220M
Chr 7
Chr 1
Pongo abelii
Pongo abelii

### Slide 20
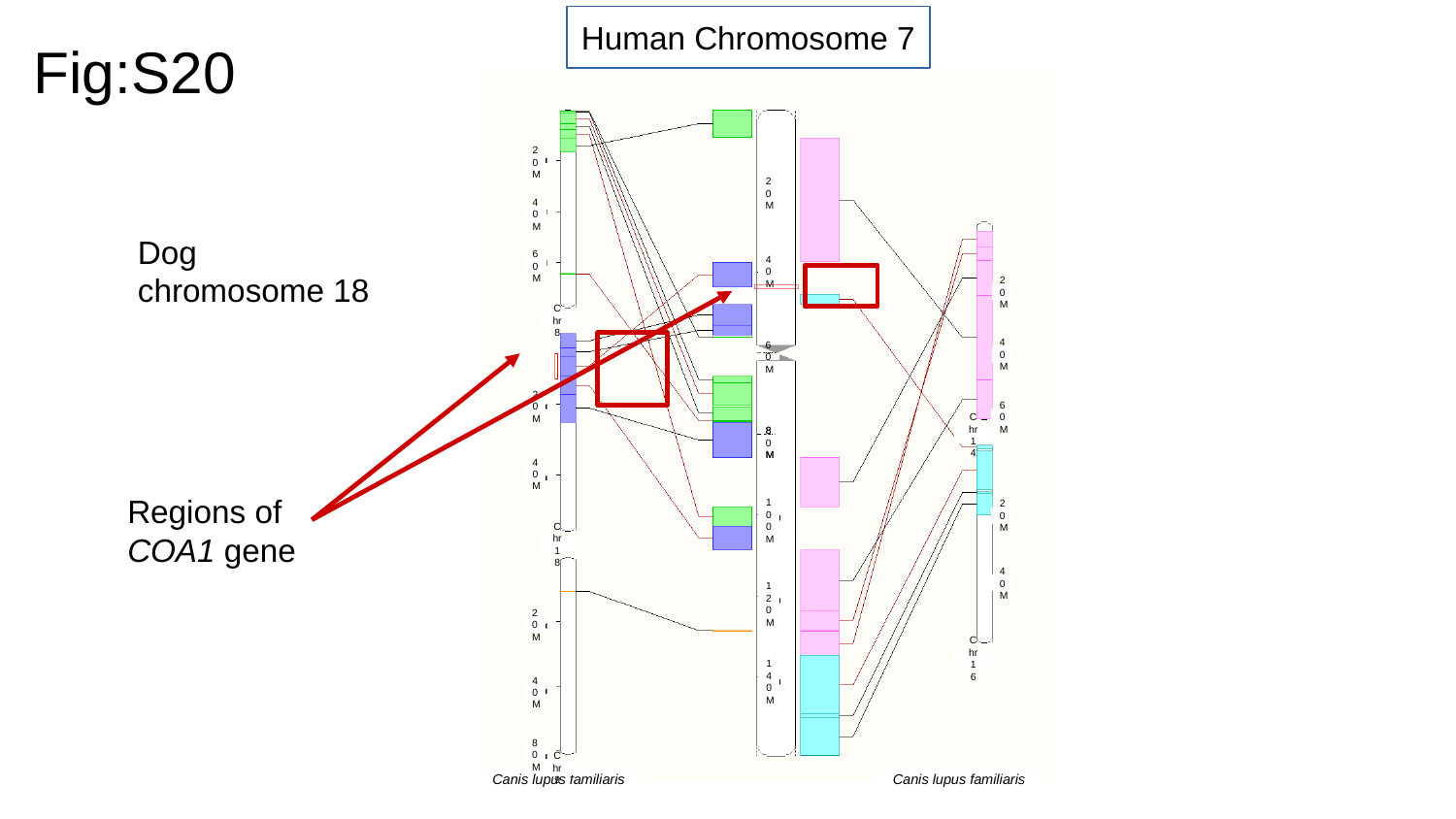

Human Chromosome 7
Fig:S20
20M
20M
40M
Dog chromosome 18
60M
40M
20M
Chr 8
40M
60M
20M
60M
Chr 14
80M
80M
40M
Regions of COA1 gene
20M
100M
Chr 18
40M
120M
20M
Chr 16
140M
40M
80M
Chr 9
Canis lupus familiaris
Canis lupus familiaris

### Slide 21
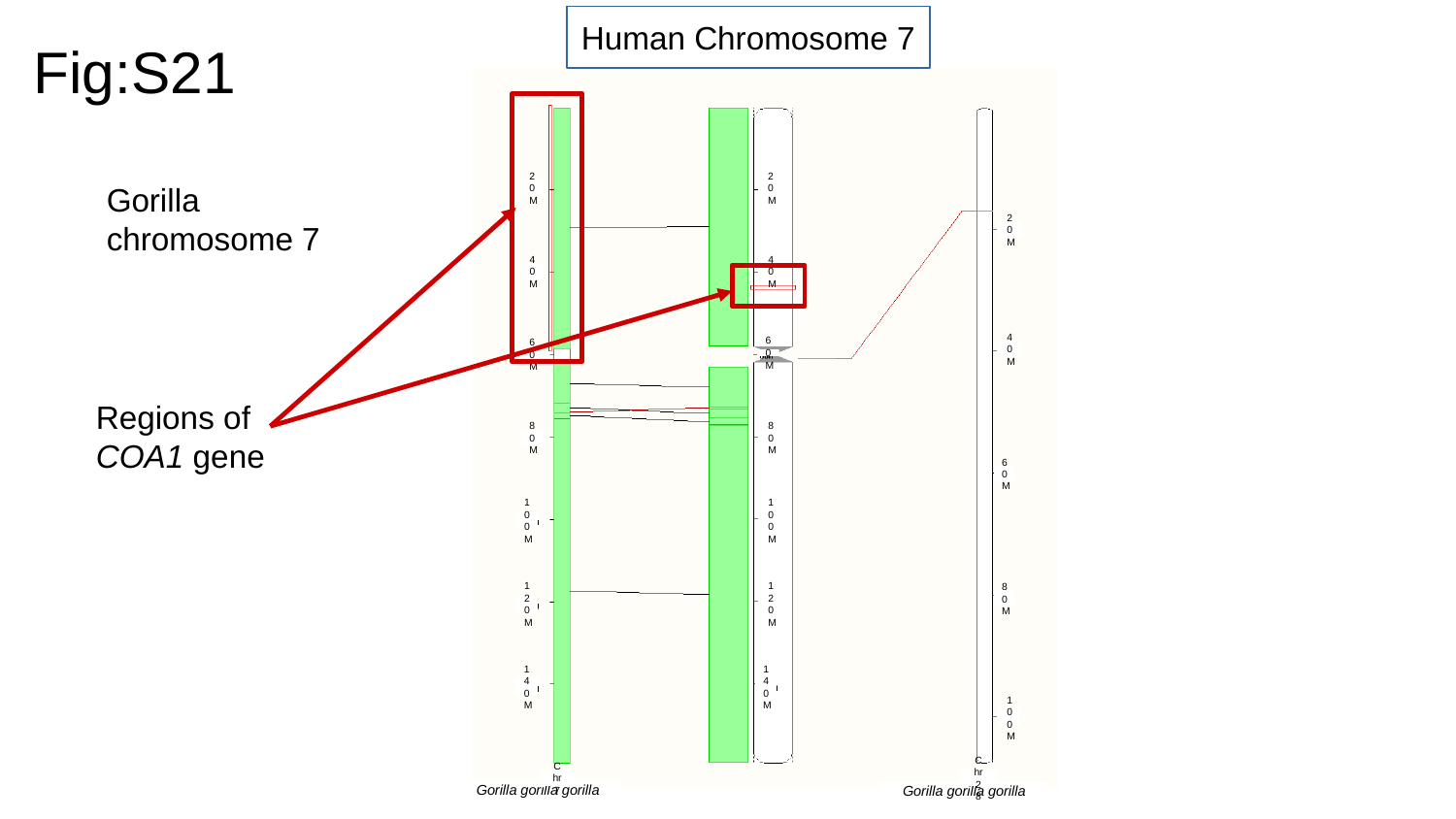

Human Chromosome 7
Fig:S21
Gorilla chromosome 7
20M
20M
20M
40M
40M
40M
60M
60M
Regions of COA1 gene
80M
80M
60M
100M
100M
80M
120M
120M
140M
140M
100M
Chr 7
Chr 28
Gorilla gorilla gorilla
Gorilla gorilla gorilla

### Slide 22
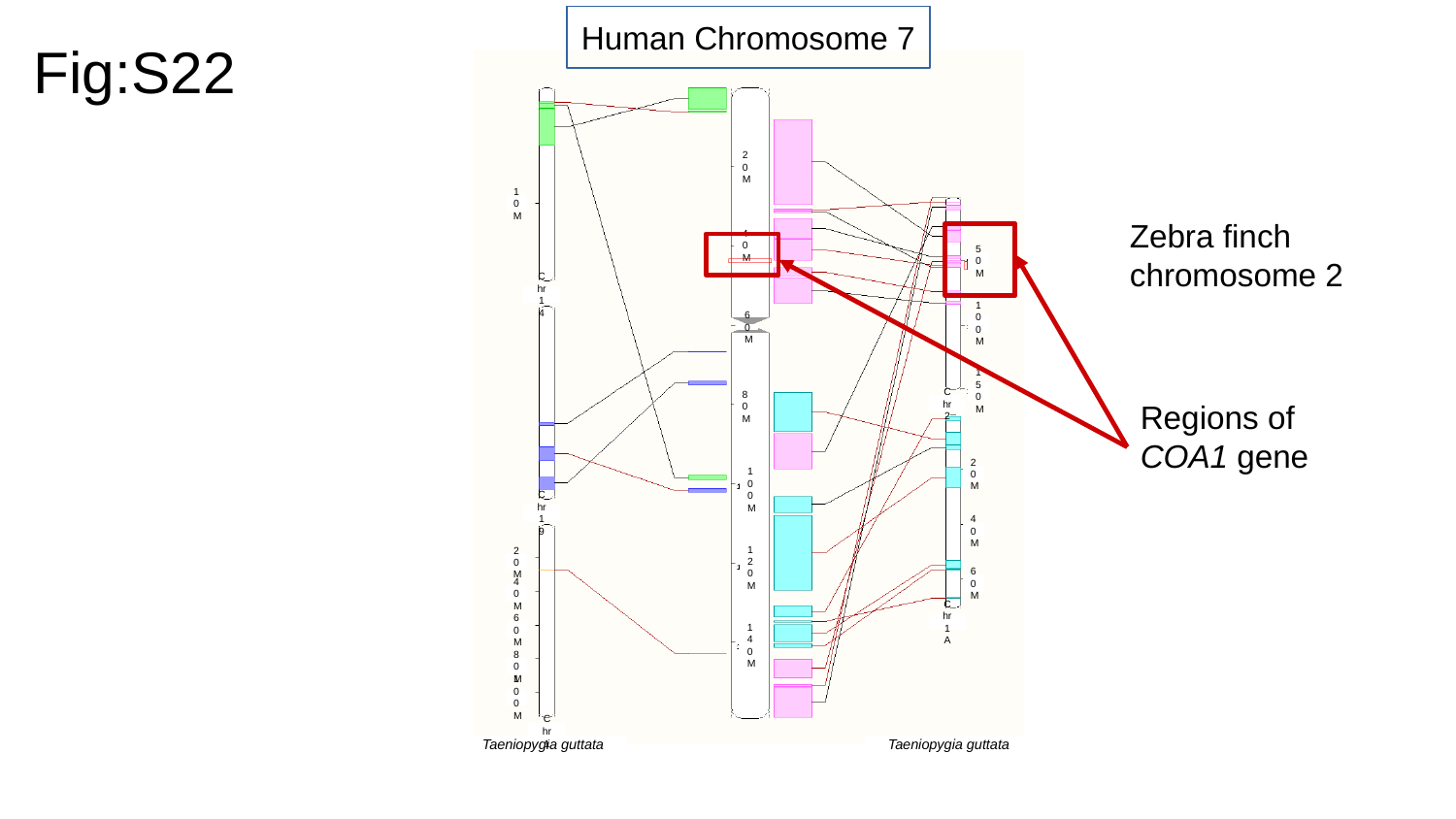

Human Chromosome 7
Fig:S22
20M
Zebra finch chromosome 2
10M
40M
50M
Chr 14
100M
60M
Regions of COA1 gene
150M
Chr 2
80M
20M
100M
Chr 19
40M
20M
120M
60M
40M
Chr 1A
60M
140M
80M
100M
Chr 1
Taeniopygia guttata
Taeniopygia guttata

### Slide 23
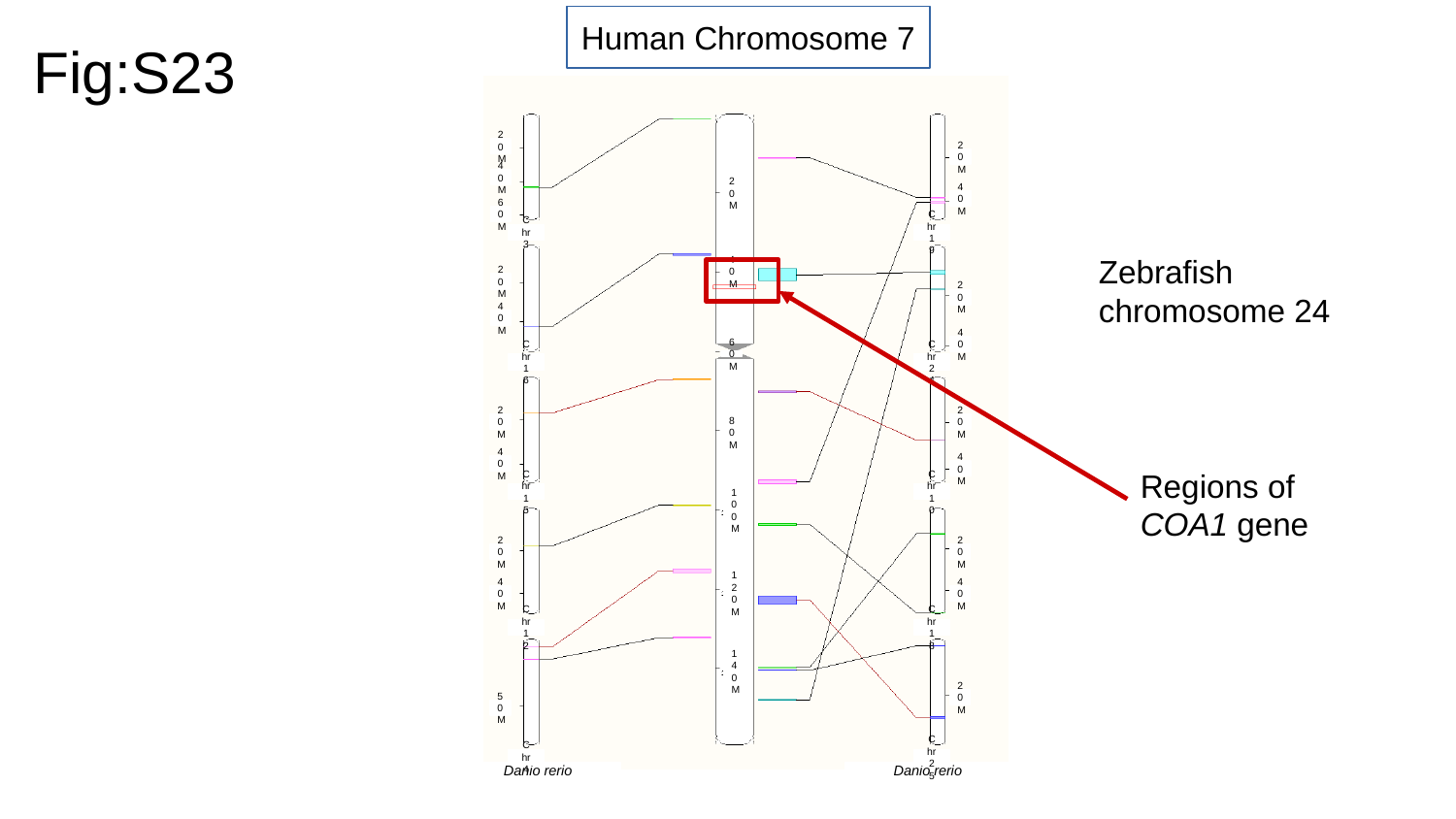

Human Chromosome 7
Fig:S23
20M
20M
40M
20M
40M
60M
Chr 3
Chr 19
Zebrafish chromosome 24
40M
20M
20M
40M
40M
60M
Chr 16
Chr 24
20M
20M
80M
Regions of COA1 gene
40M
40M
Chr 15
Chr 10
100M
20M
20M
40M
120M
40M
Chr 12
Chr 18
140M
20M
50M
Chr 4
Chr 25
Danio rerio
Danio rerio

### Slide 24
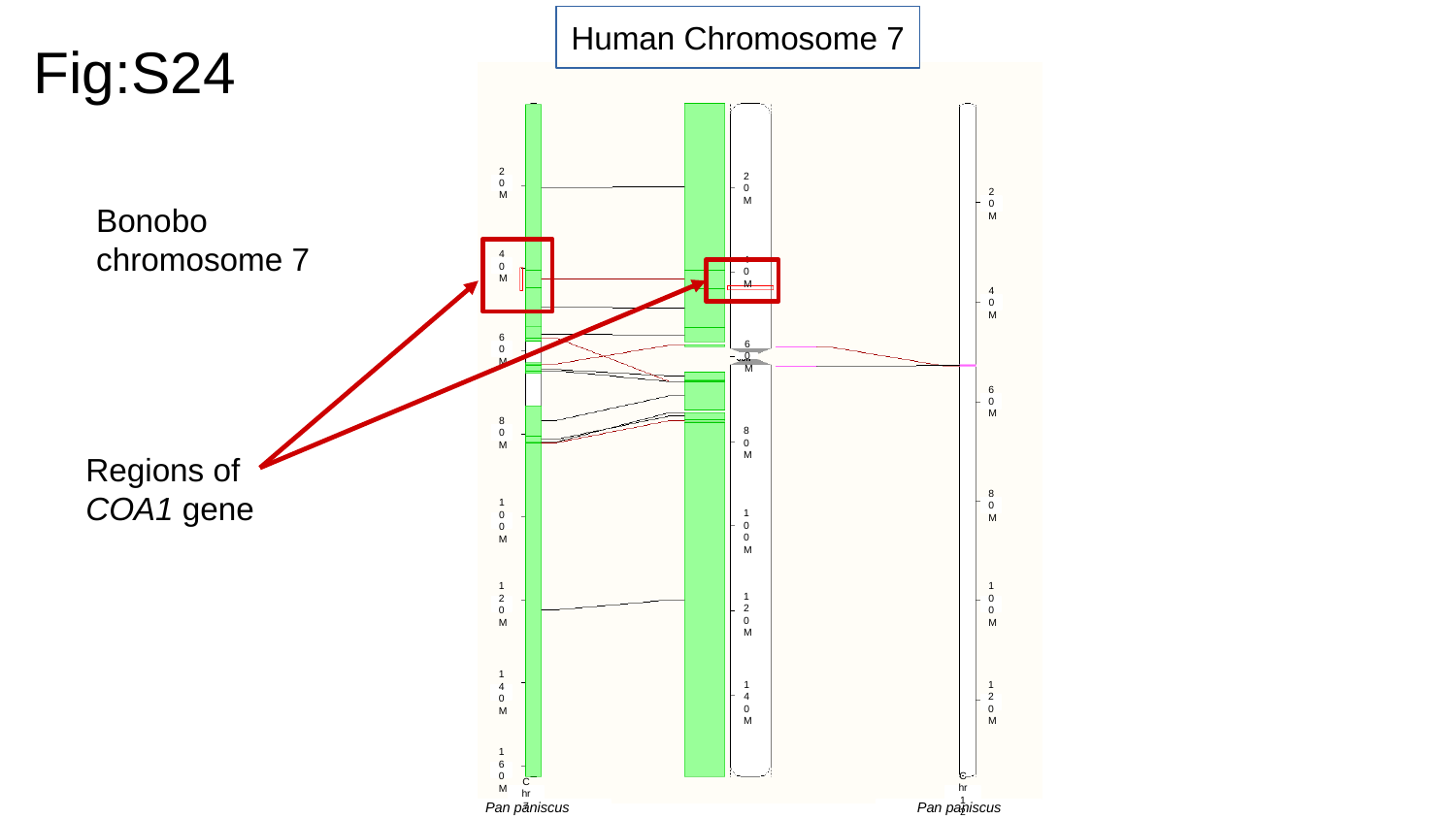

Human Chromosome 7
Fig:S24
20M
Bonobo chromosome 7
20M
20M
40M
40M
40M
60M
60M
60M
80M
Regions of COA1 gene
80M
80M
100M
100M
120M
100M
120M
140M
140M
120M
160M
Chr 7
Chr 12
Pan paniscus
Pan paniscus

### Slide 25
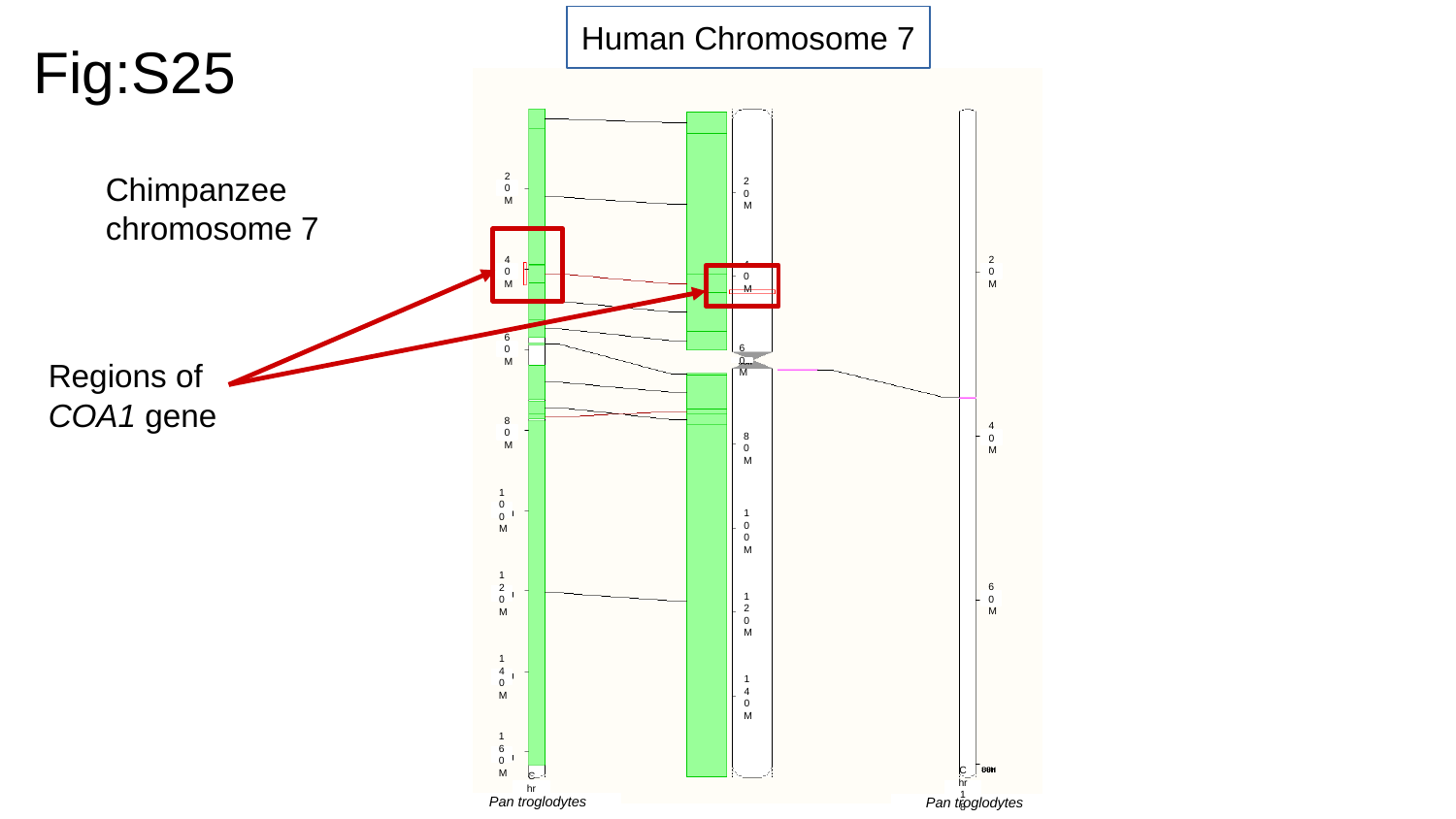

Human Chromosome 7
Fig:S25
Chimpanzee chromosome 7
20M
20M
40M
20M
40M
Regions of COA1 gene
60M
60M
80M
40M
80M
100M
100M
120M
60M
120M
140M
140M
160M
Chr 7
Chr 18
Pan troglodytes
Pan troglodytes

### Slide 26
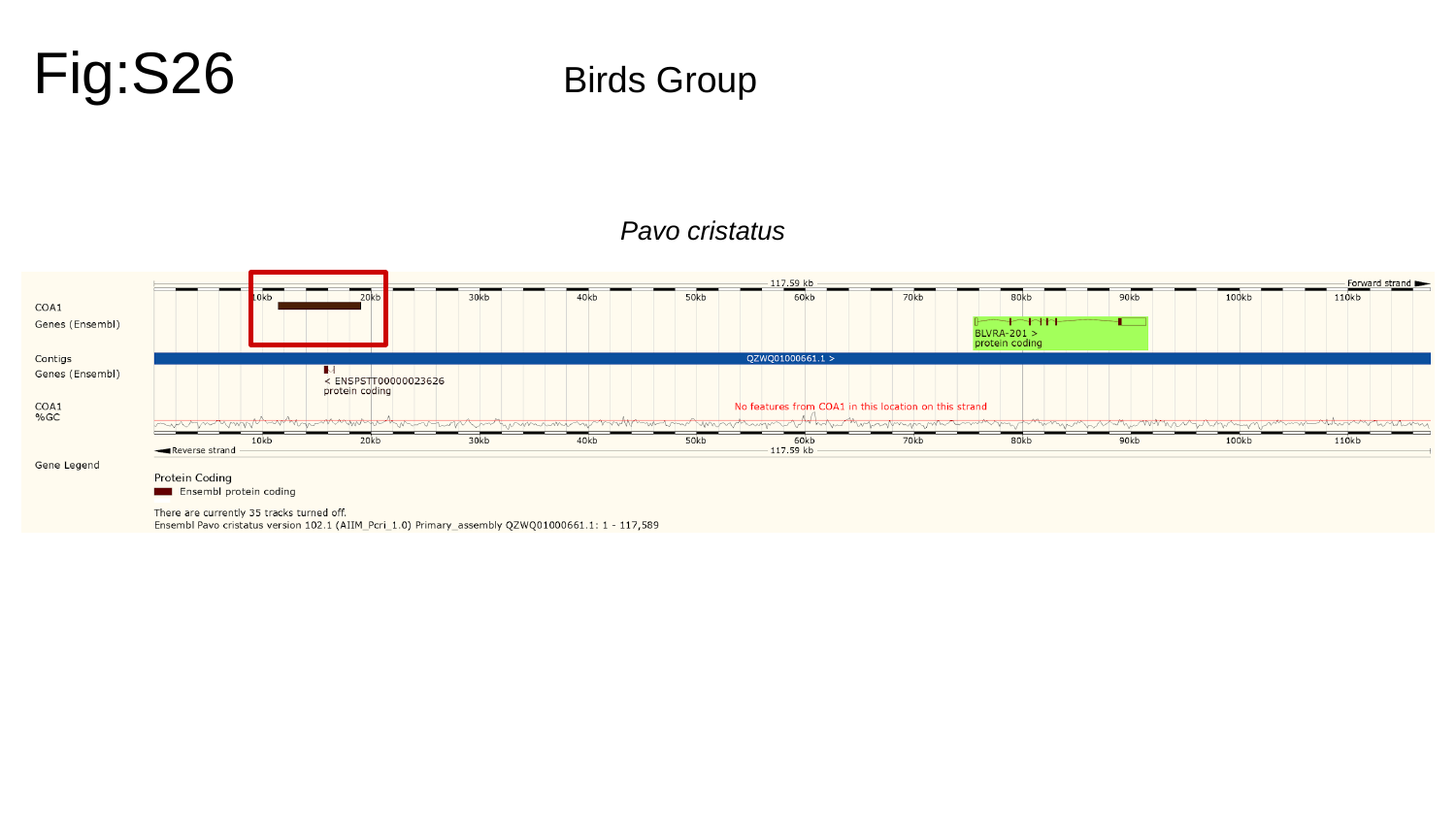

Fig:S26
Birds Group
Pavo cristatus

### Slide 27
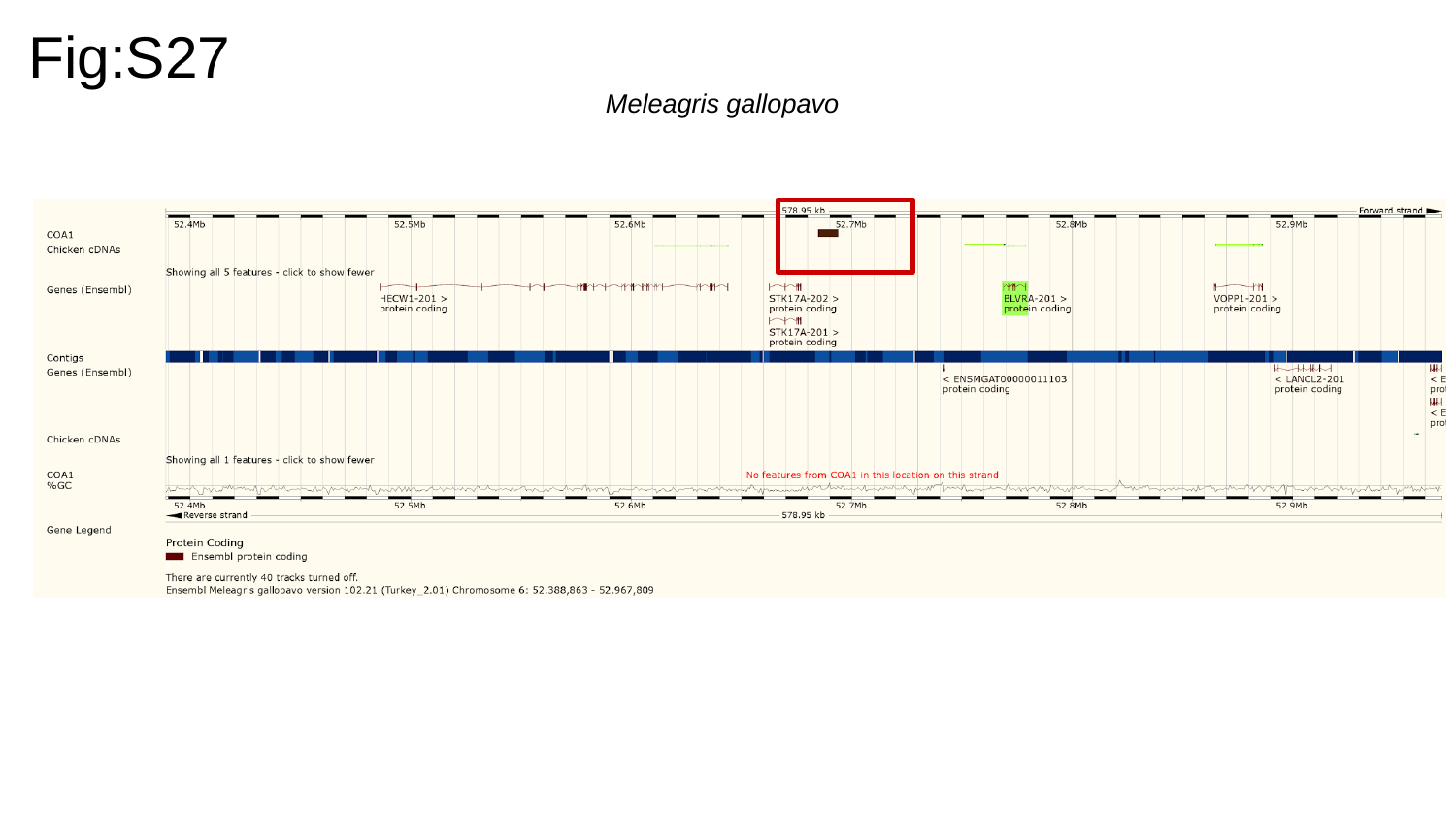

Fig:S27
Meleagris gallopavo

### Slide 28
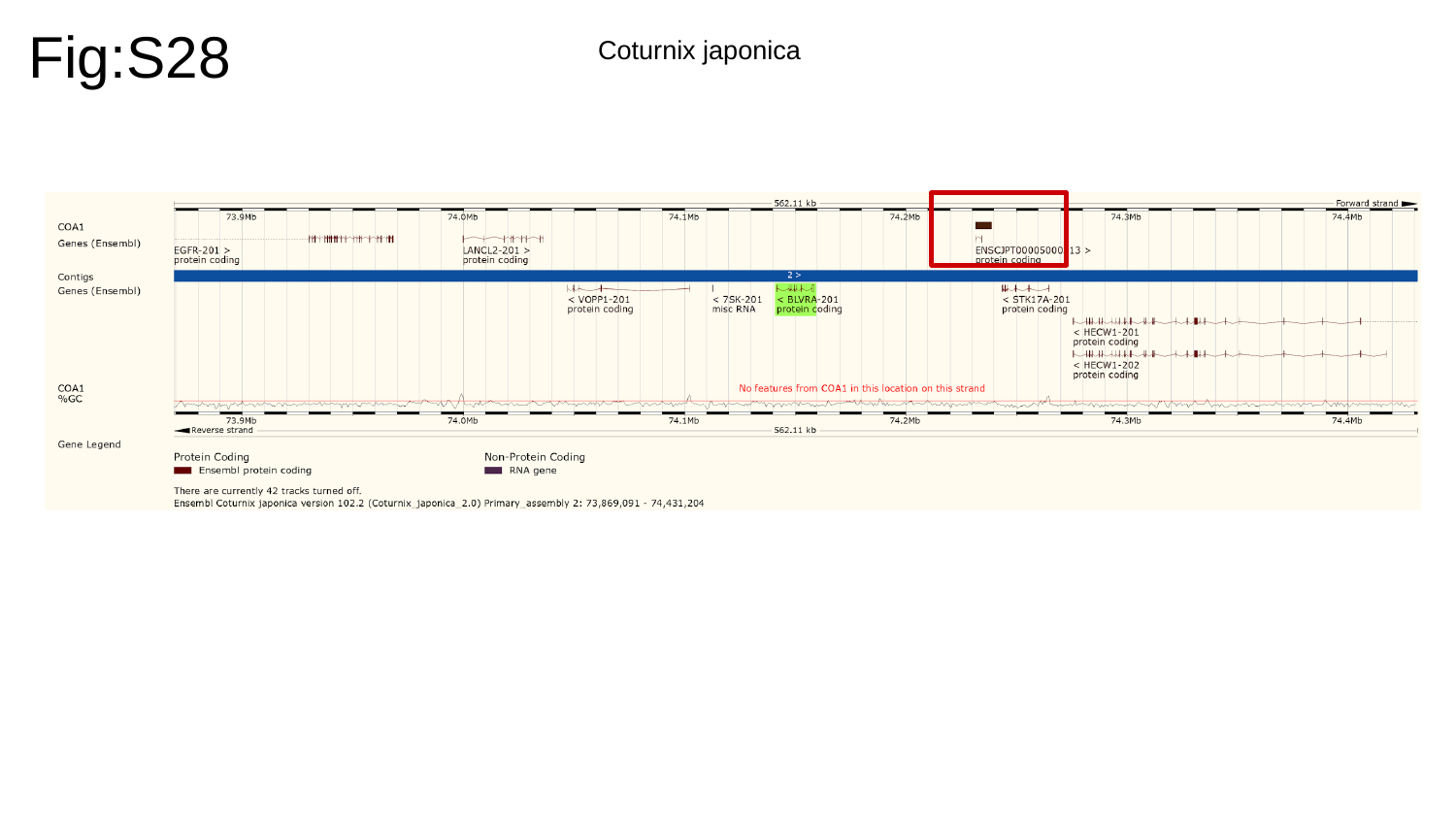

Fig:S28
Coturnix japonica

### Slide 29
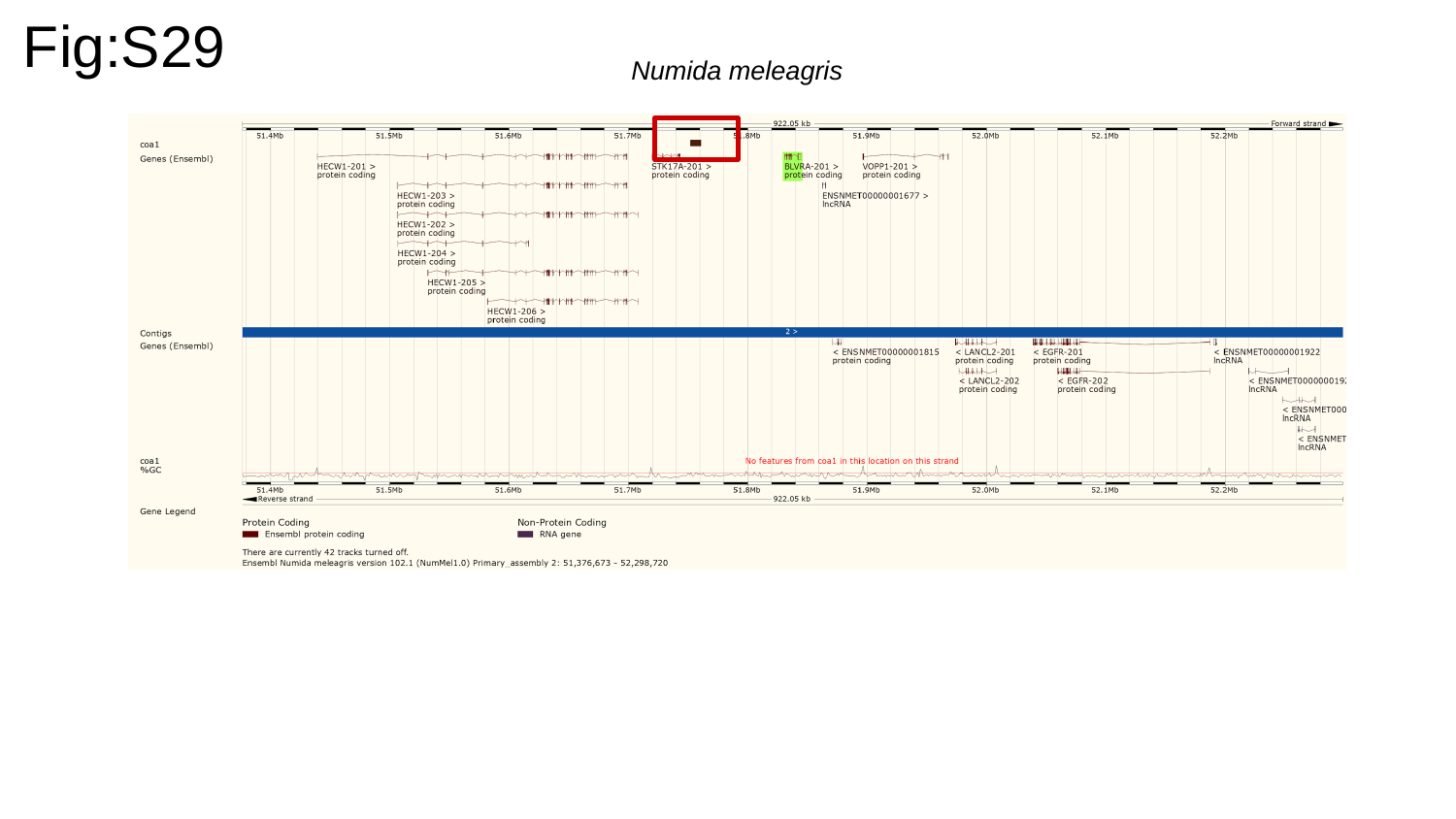

Fig:S29
Numida meleagris

### Slide 30
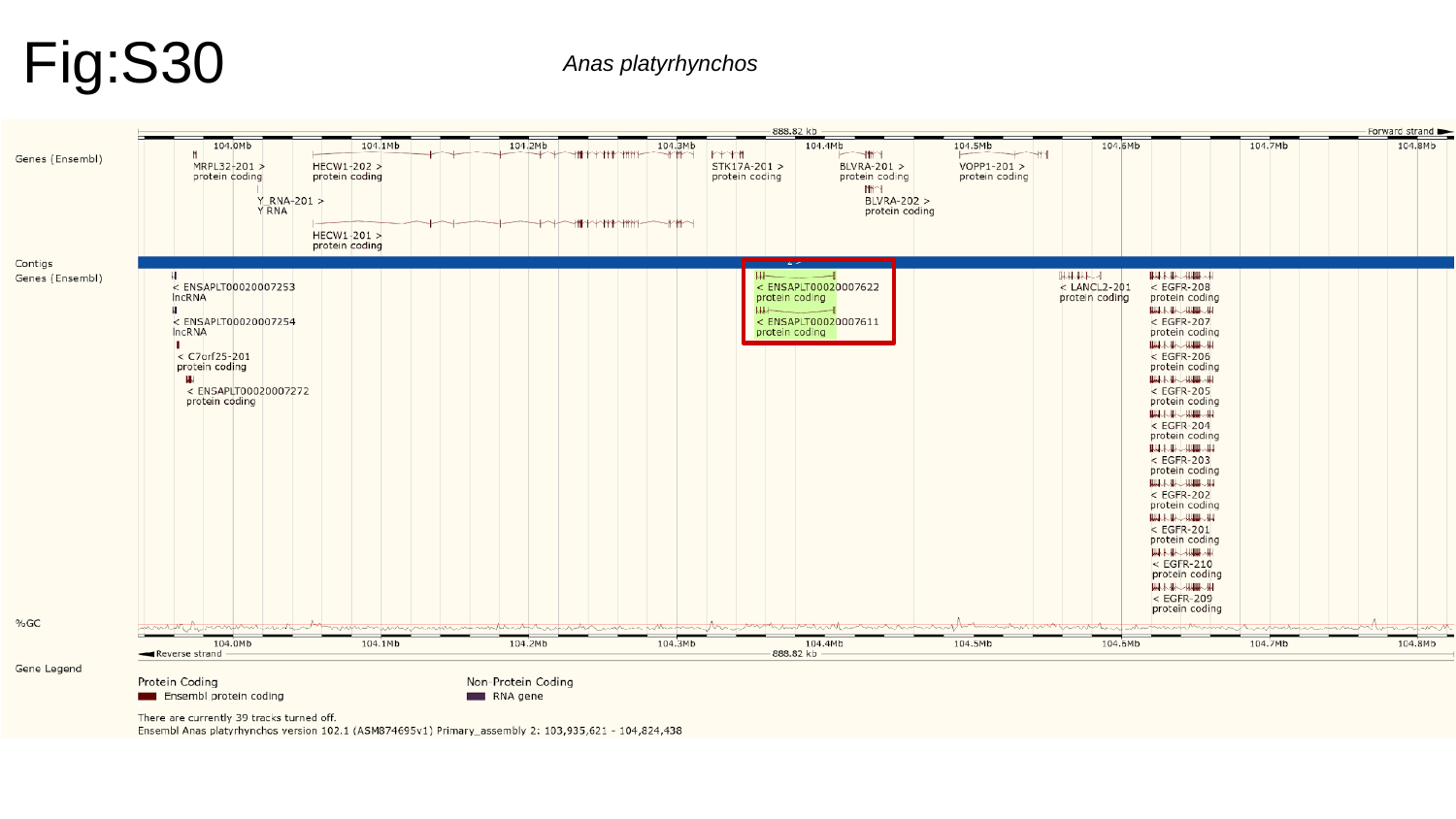

Fig:S30
Anas platyrhynchos

### Slide 31
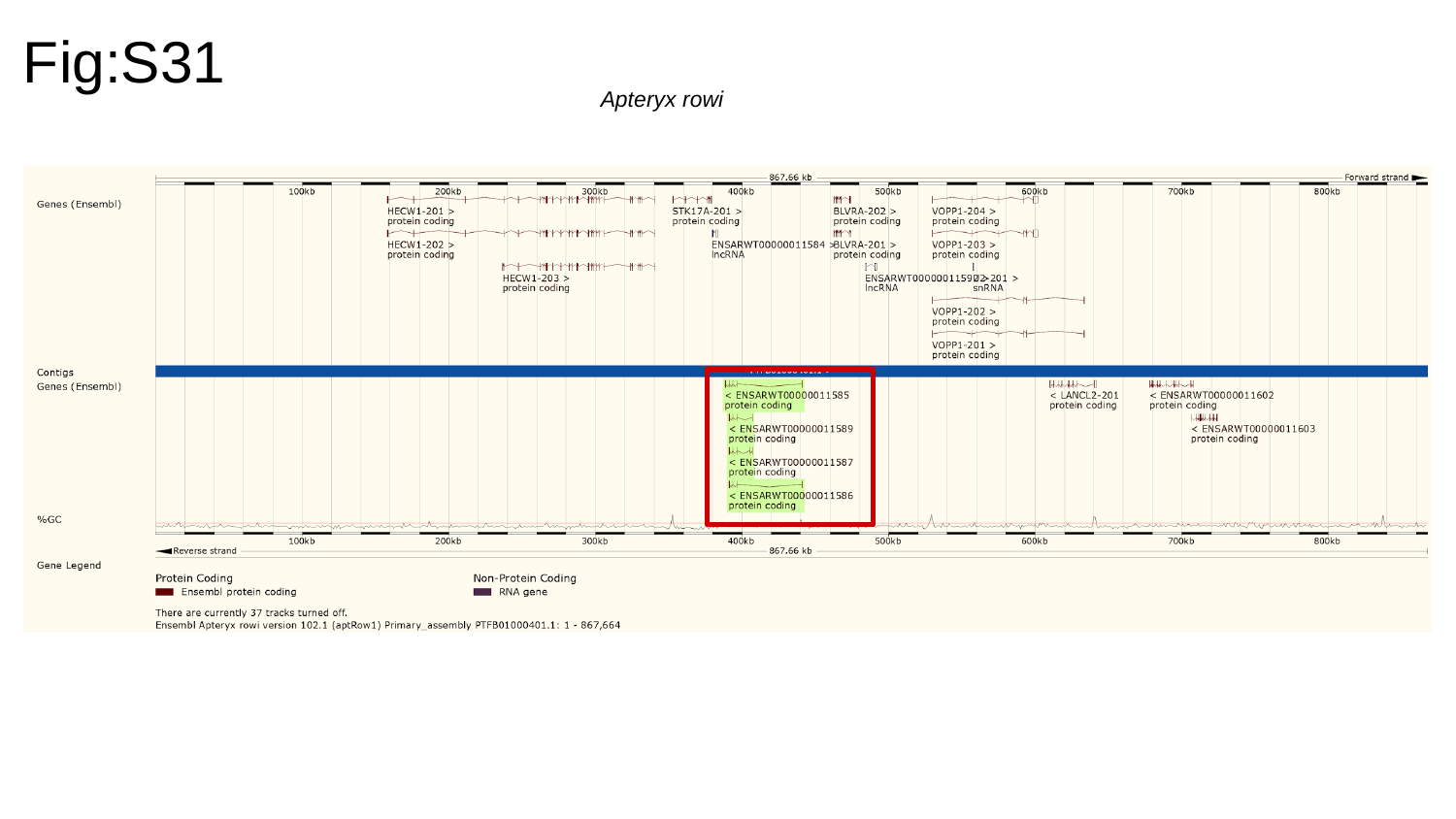

Fig:S31
Apteryx rowi

### Slide 32
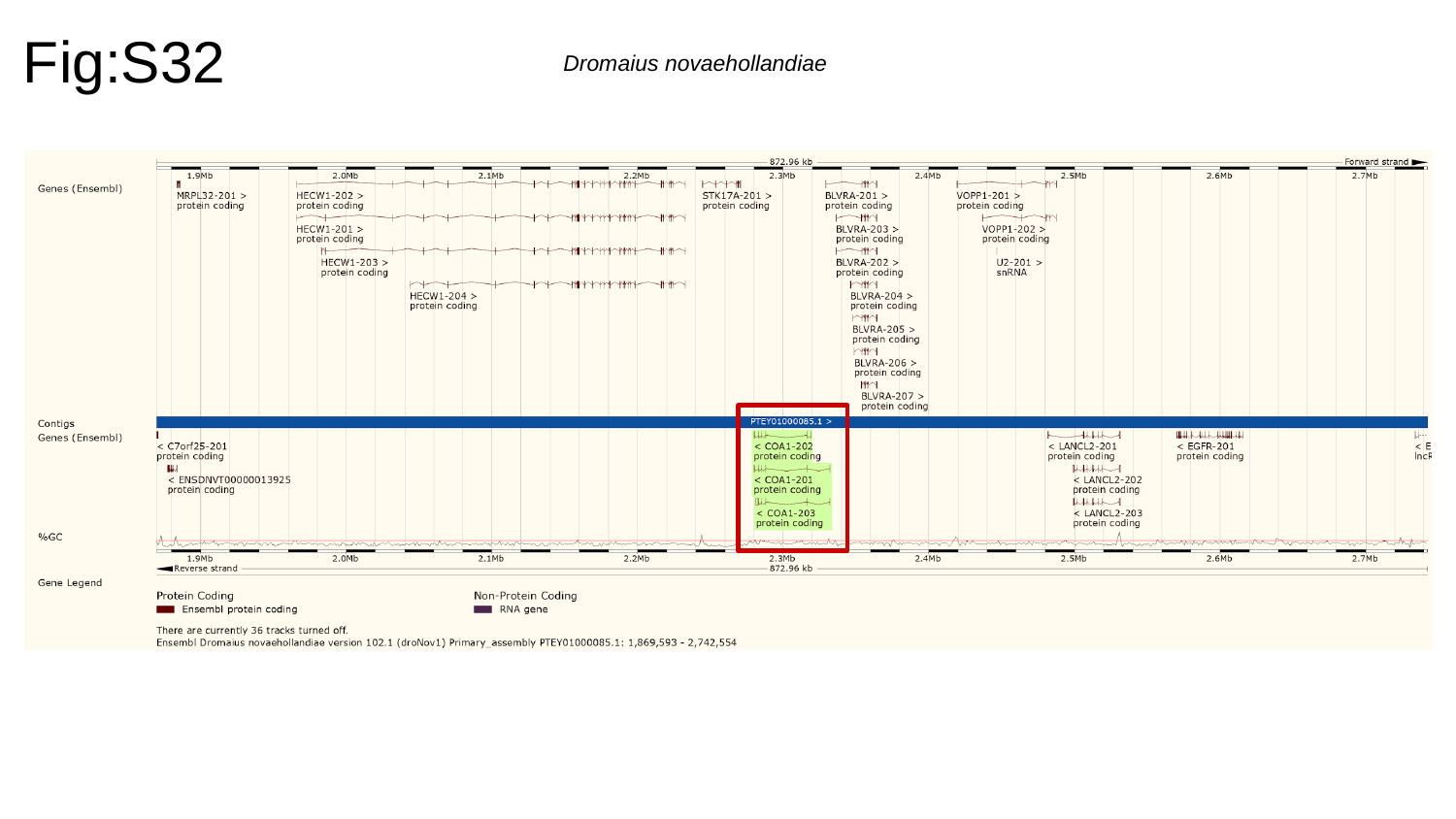

Fig:S32
Dromaius novaehollandiae

### Slide 33
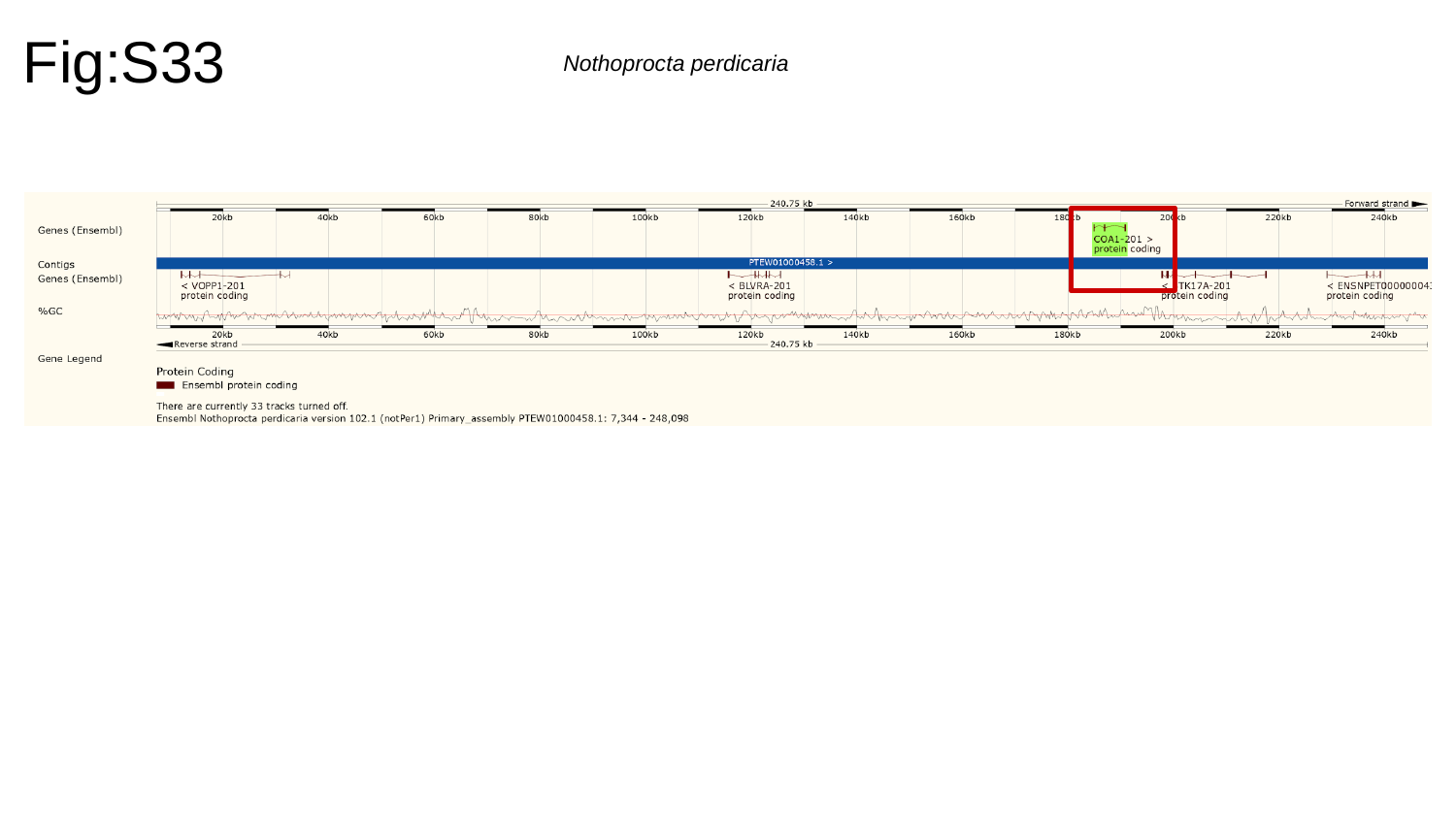

Fig:S33
Nothoprocta perdicaria

### Slide 34
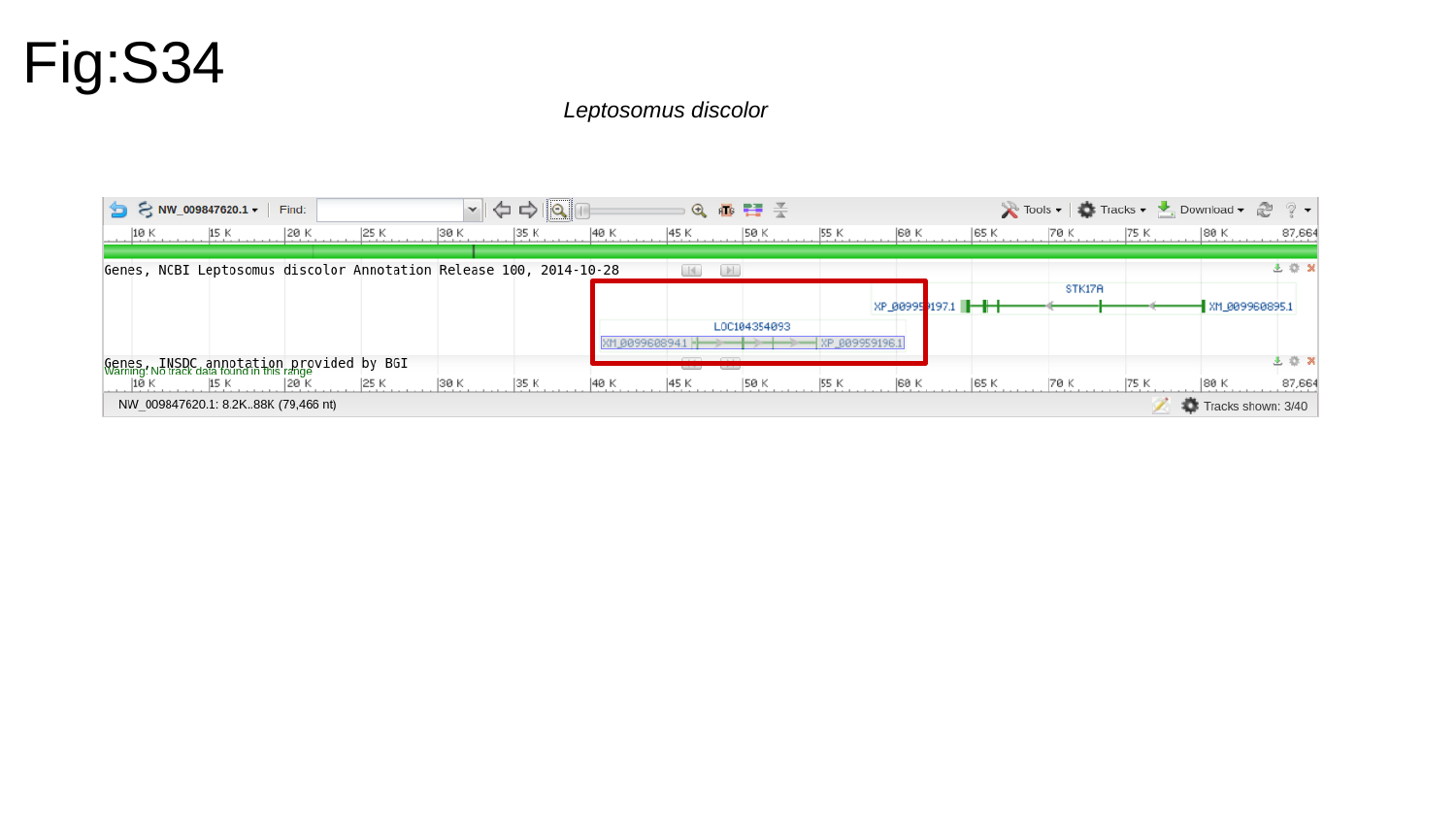

Fig:S34
Leptosomus discolor

### Slide 35
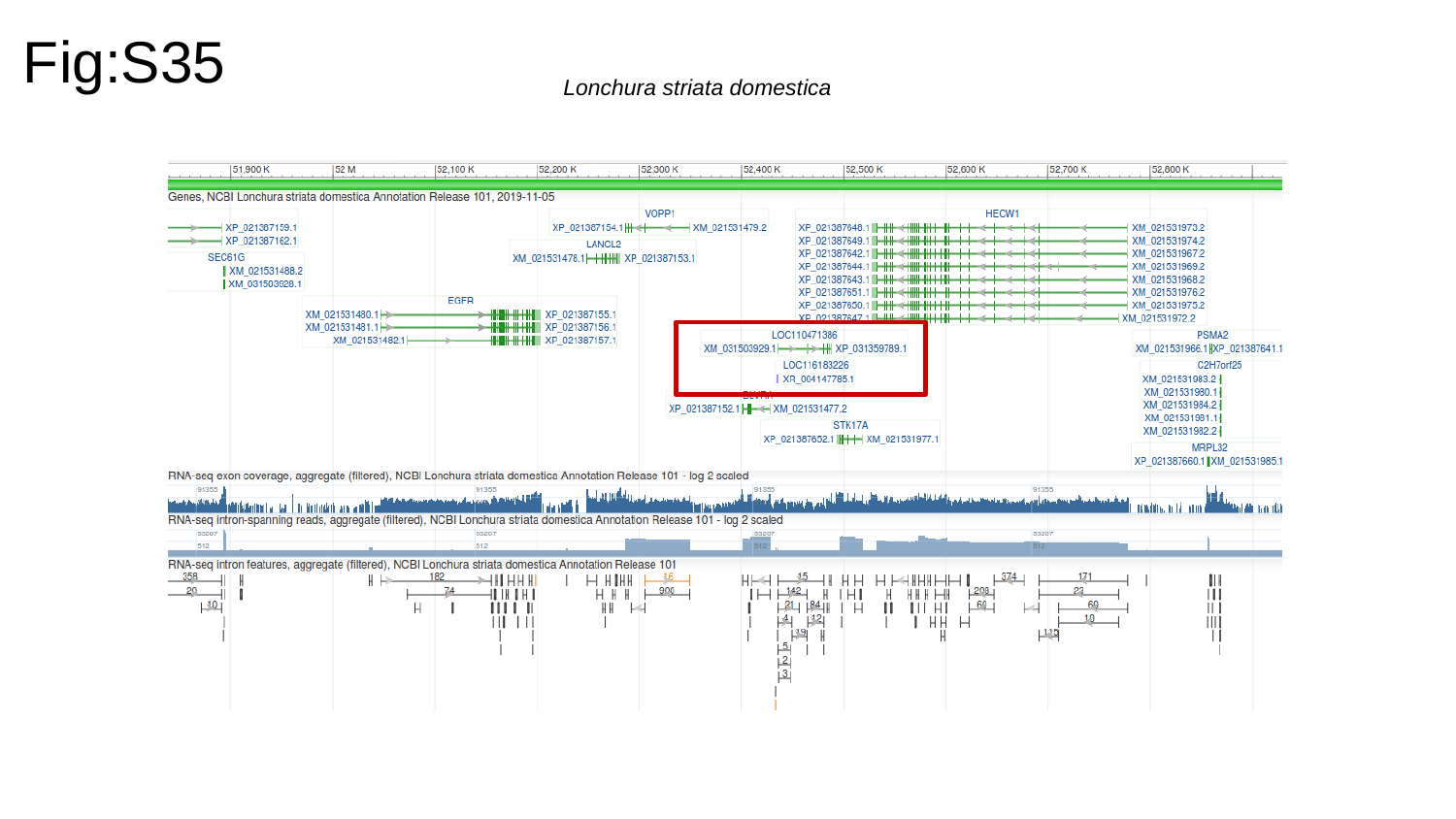

Fig:S35
Lonchura striata domestica

### Slide 36
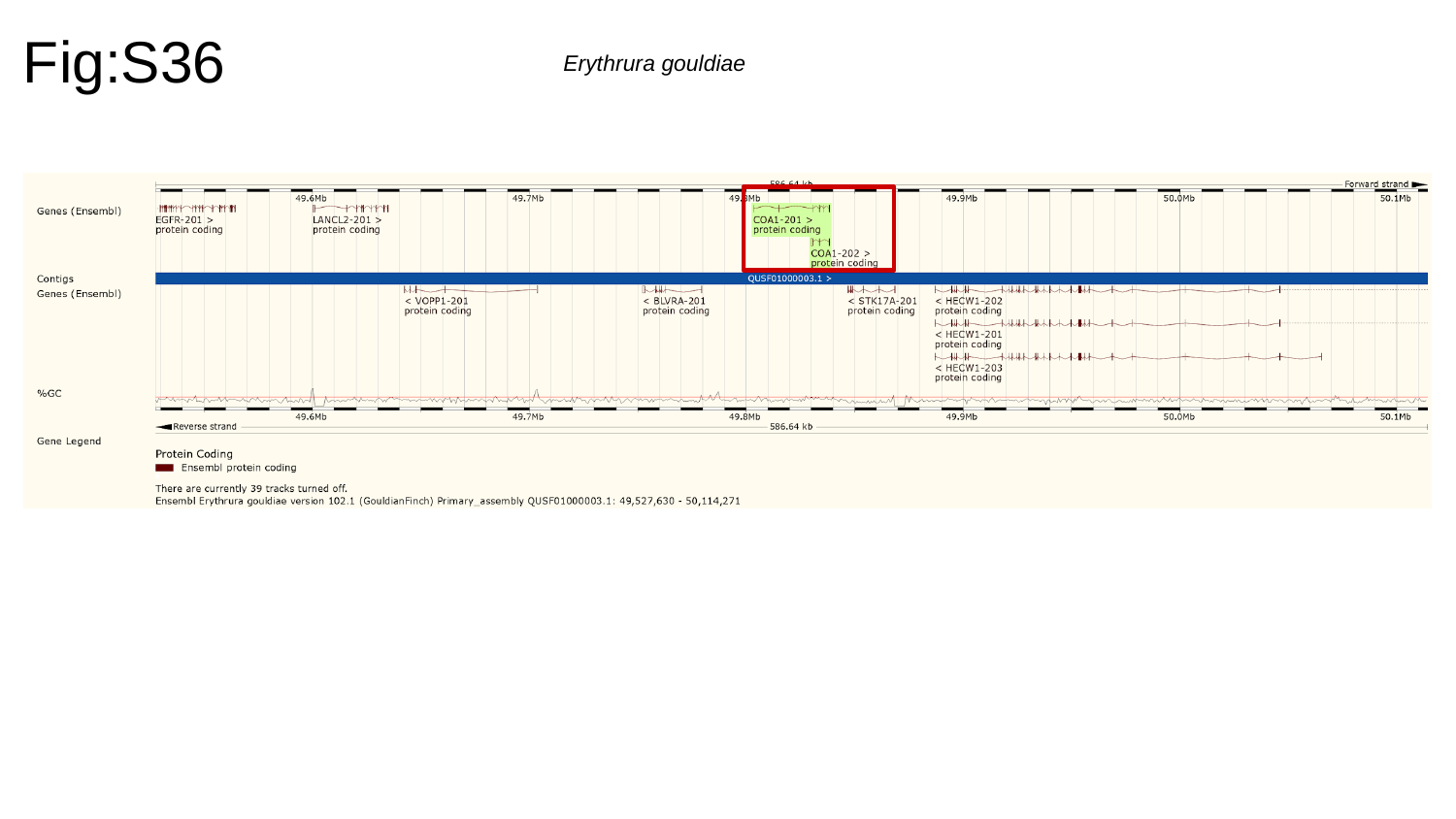

Fig:S36
Erythrura gouldiae

### Slide 37
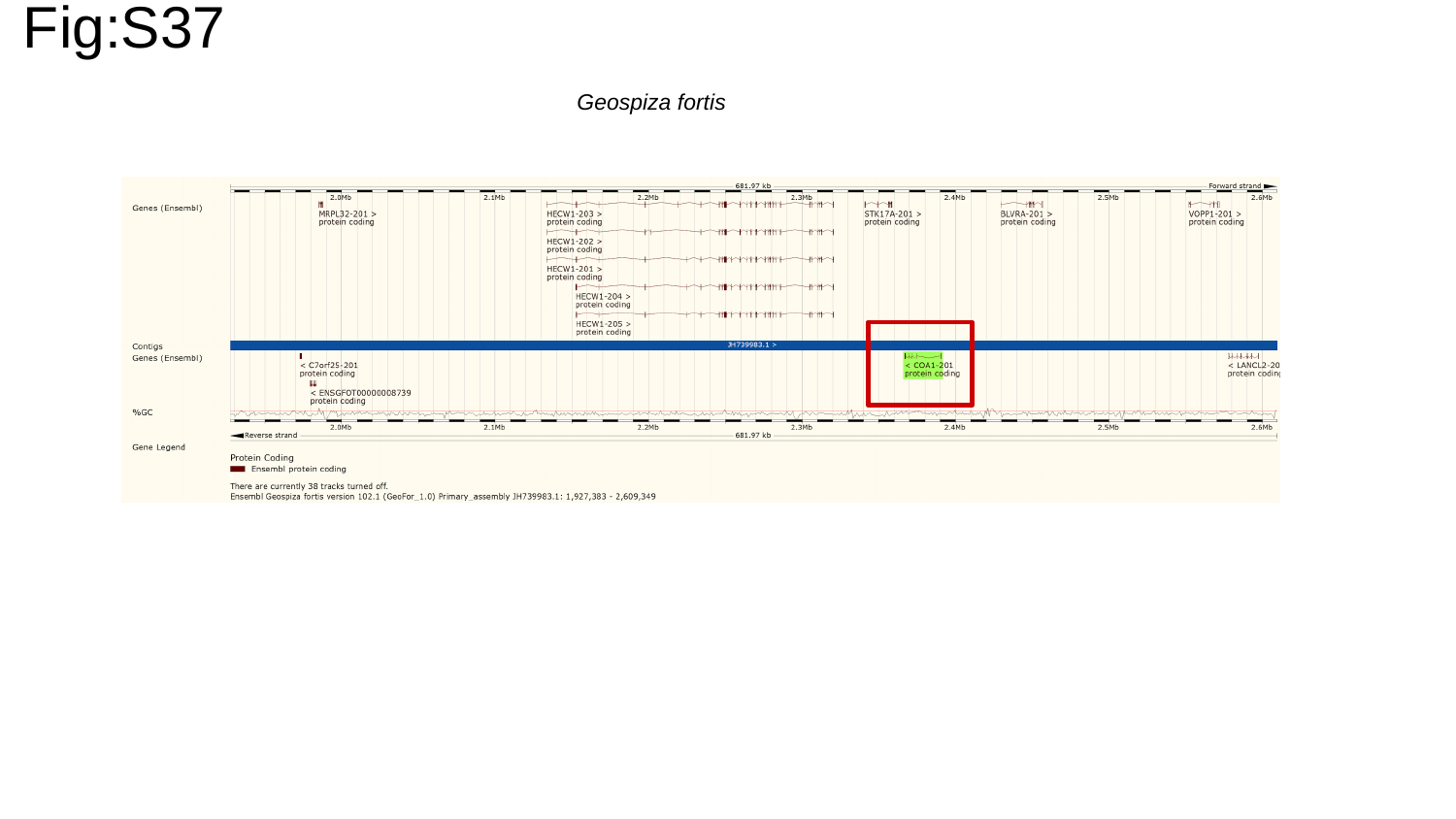

Fig:S37
Geospiza fortis

### Slide 38
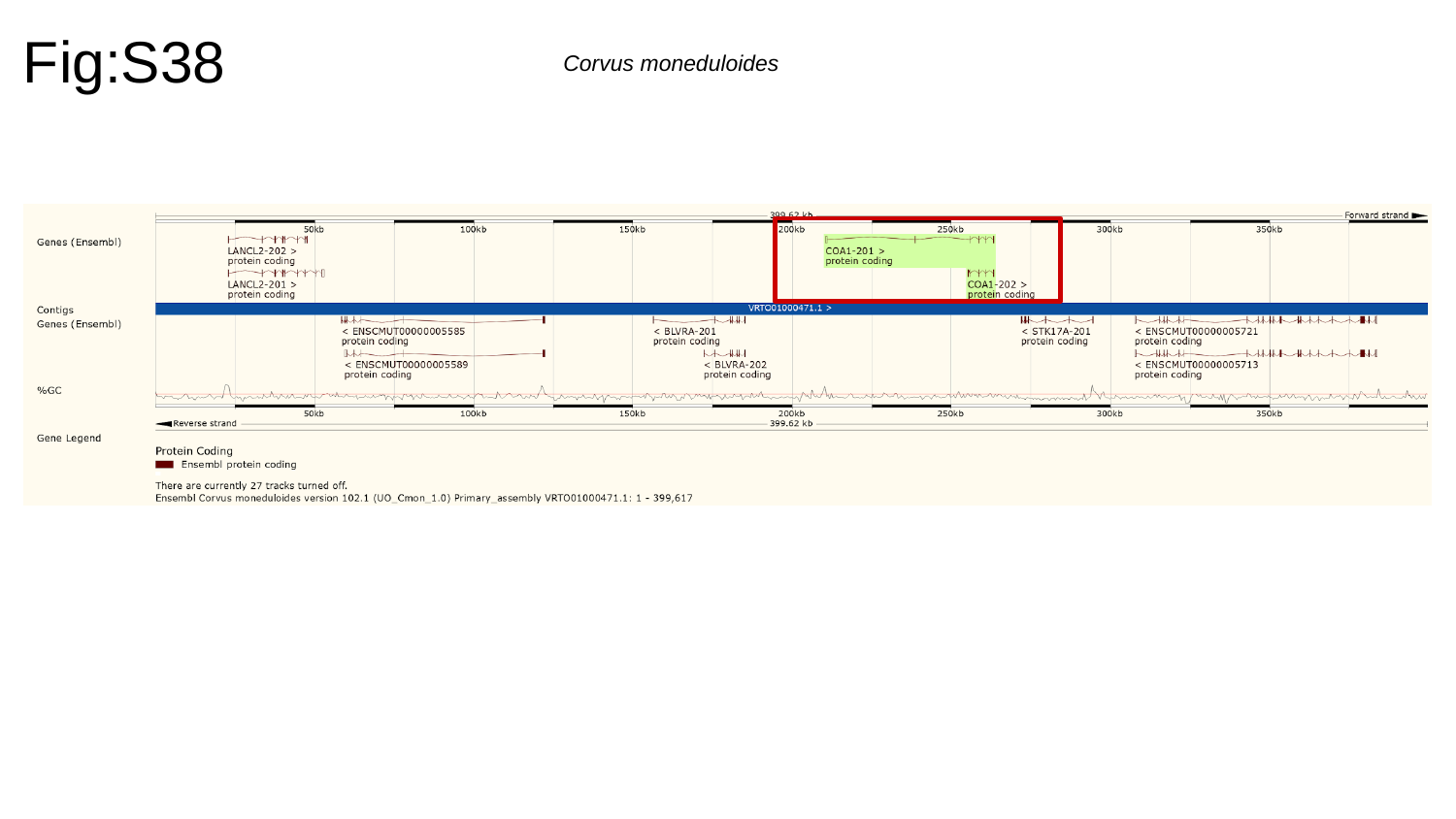

Fig:S38
Corvus moneduloides

### Slide 39
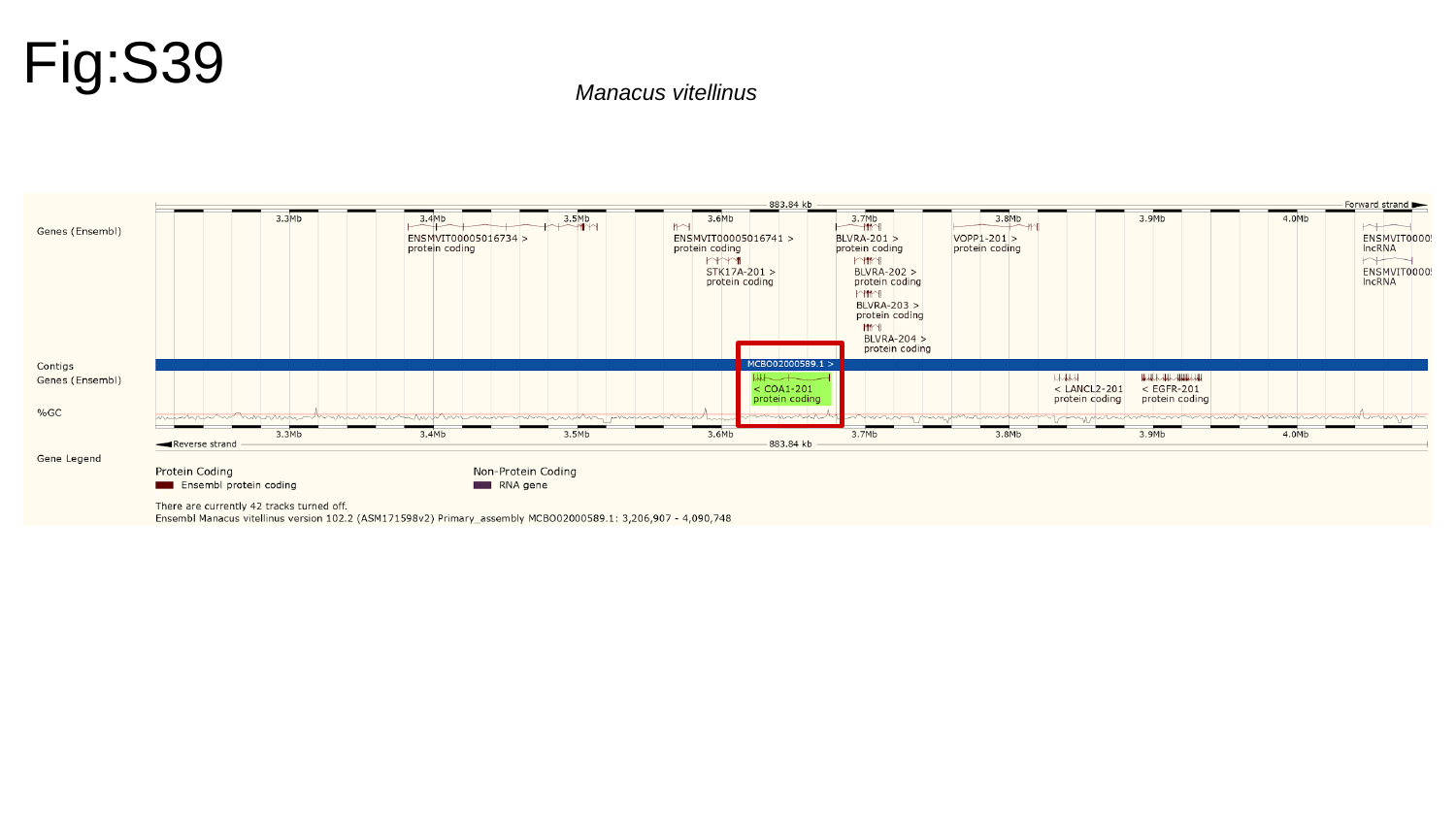

Fig:S39
Manacus vitellinus

### Slide 40
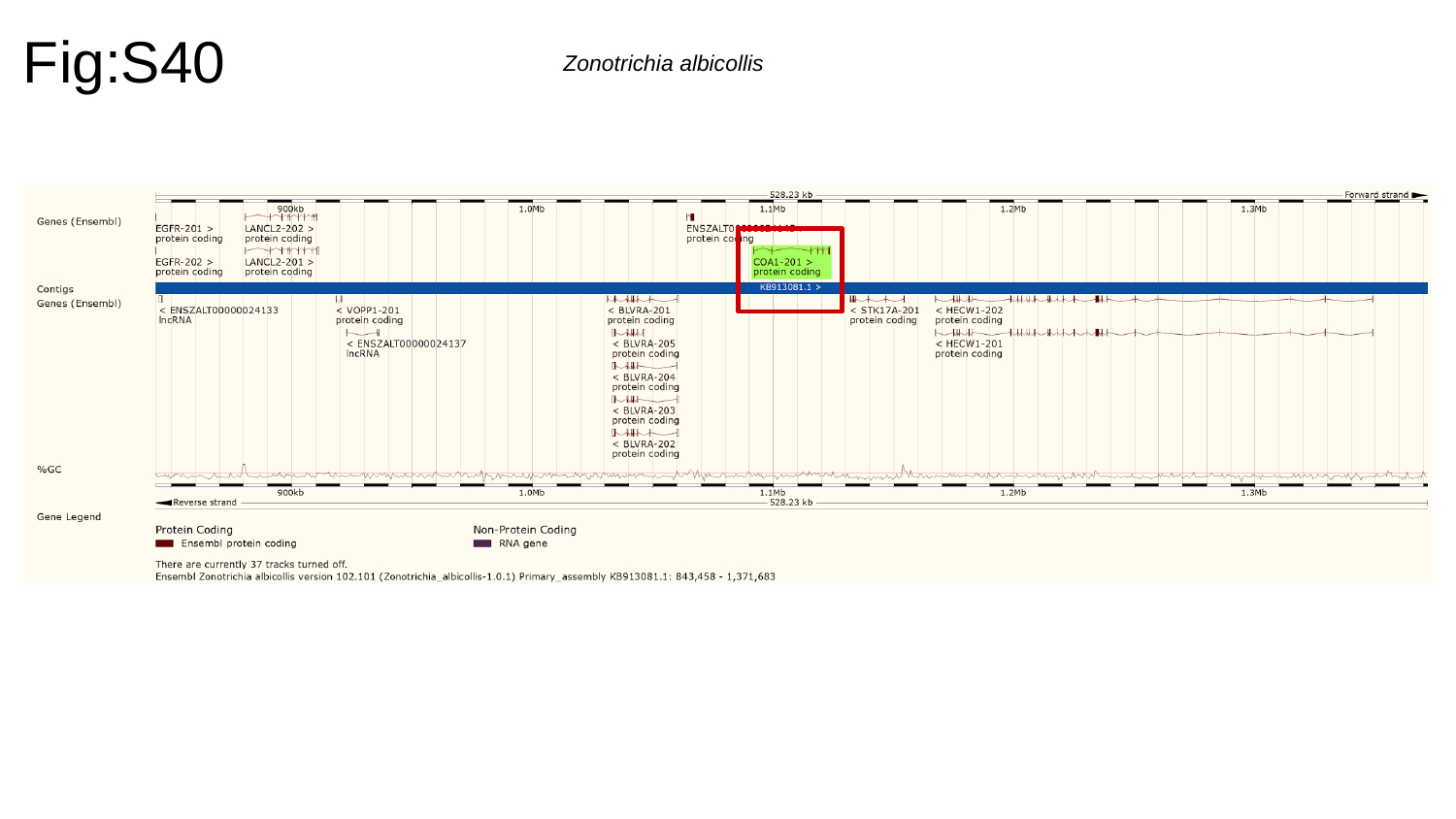

Fig:S40
Zonotrichia albicollis

### Slide 41
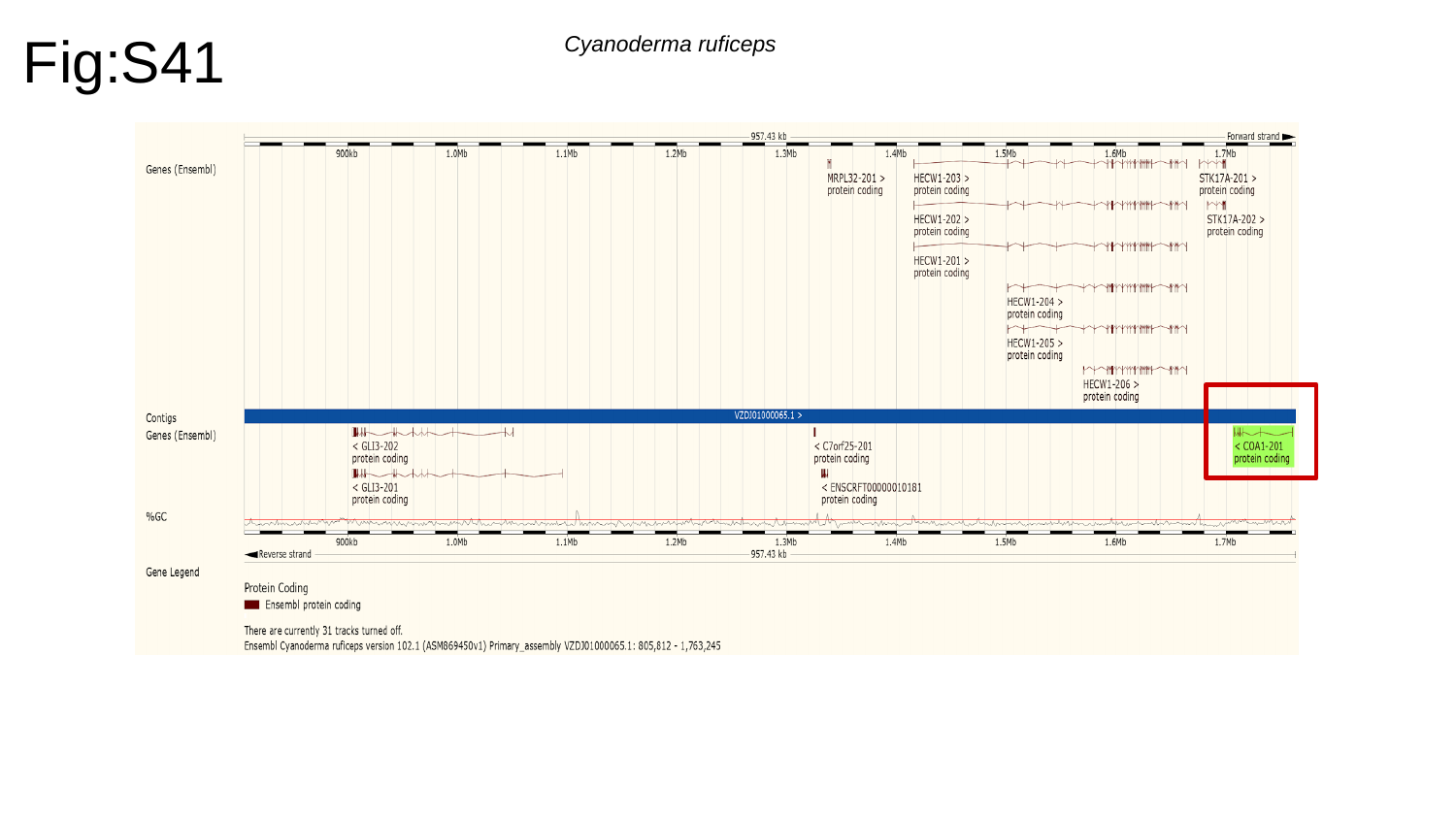

Fig:S41
Cyanoderma ruficeps

### Slide 42
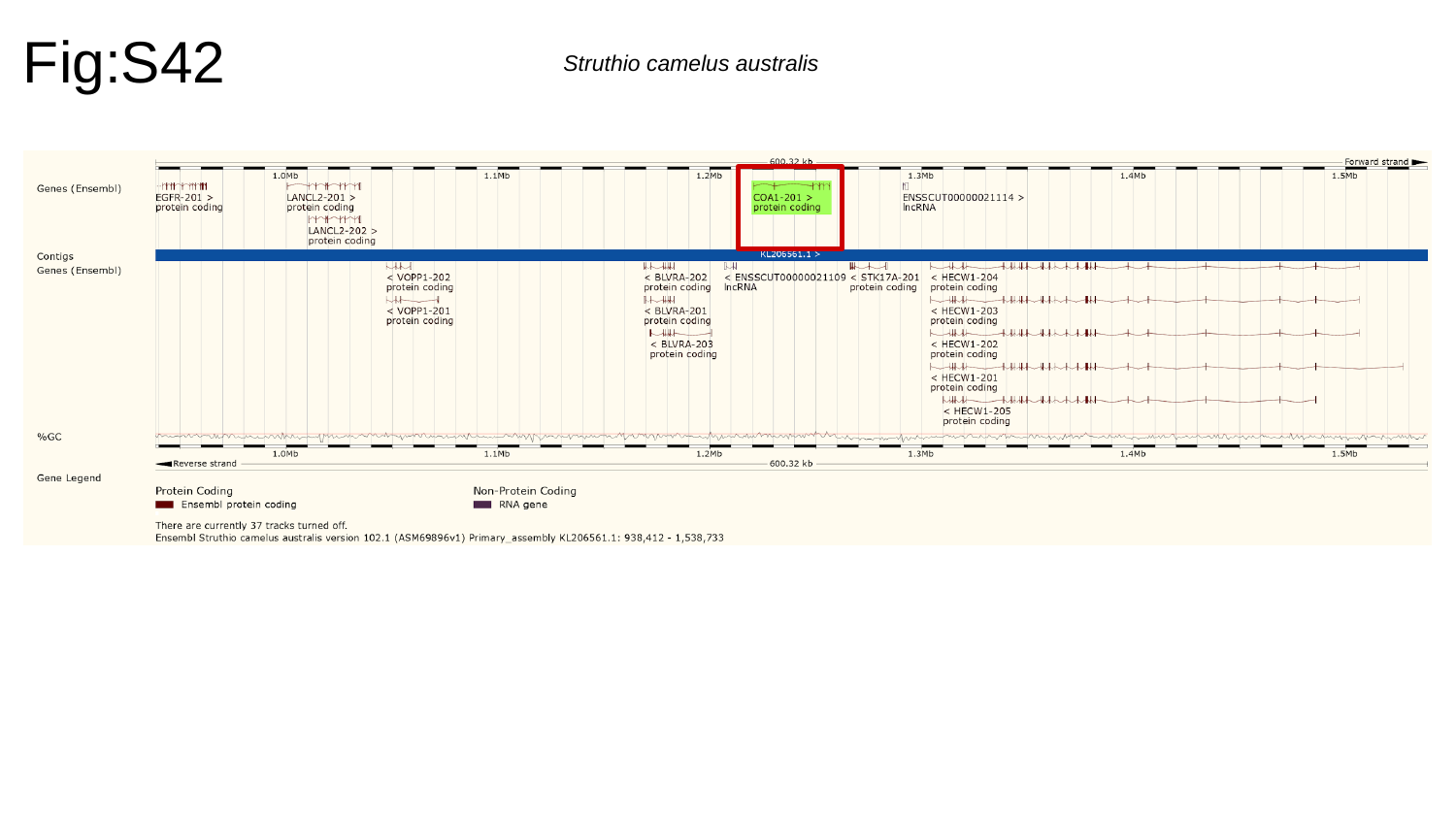

Fig:S42
Struthio camelus australis

### Slide 43
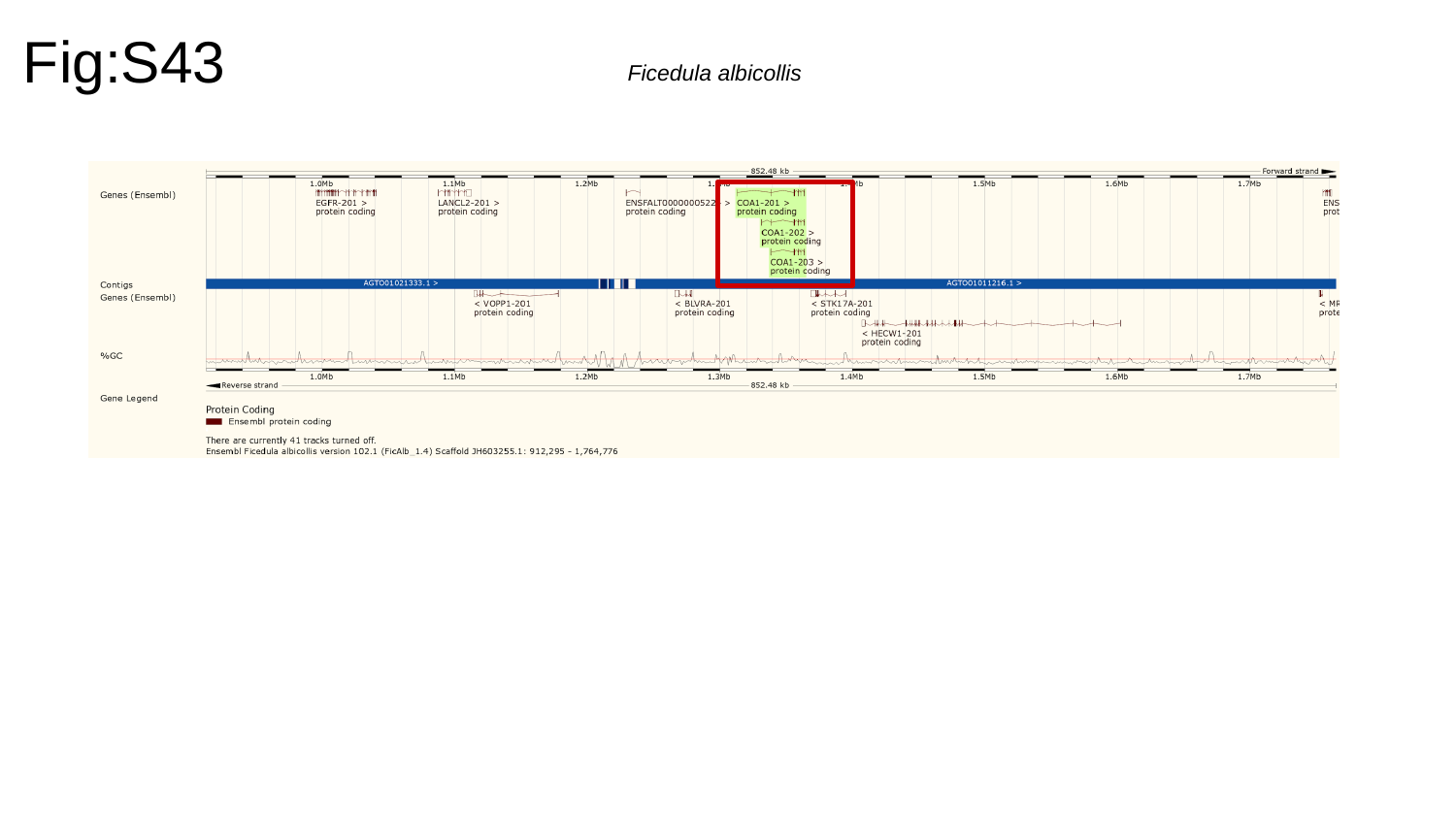

Fig:S43
Ficedula albicollis

### Slide 44
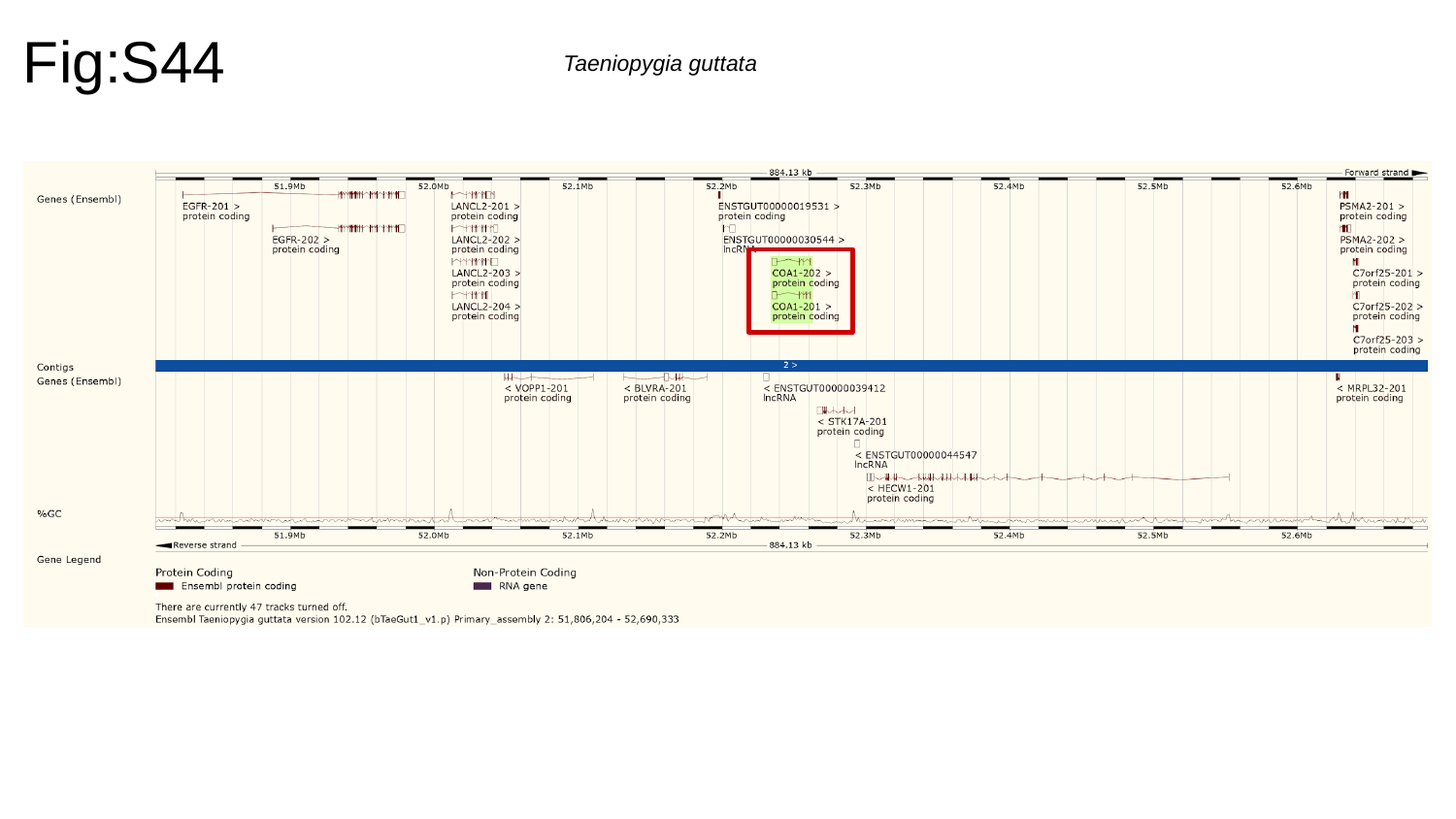

Fig:S44
Taeniopygia guttata

### Slide 45
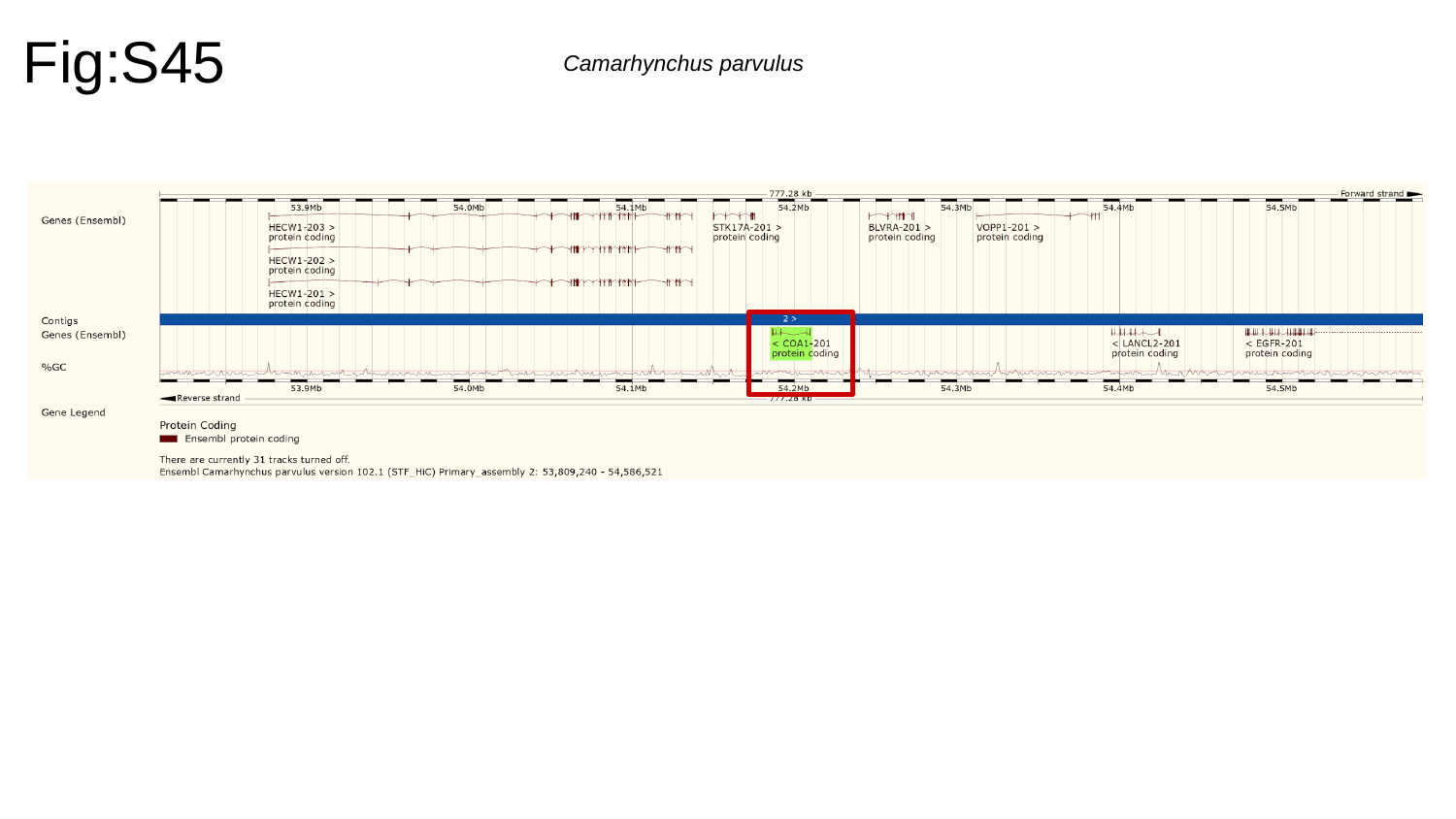

Fig:S45
Camarhynchus parvulus

### Slide 46
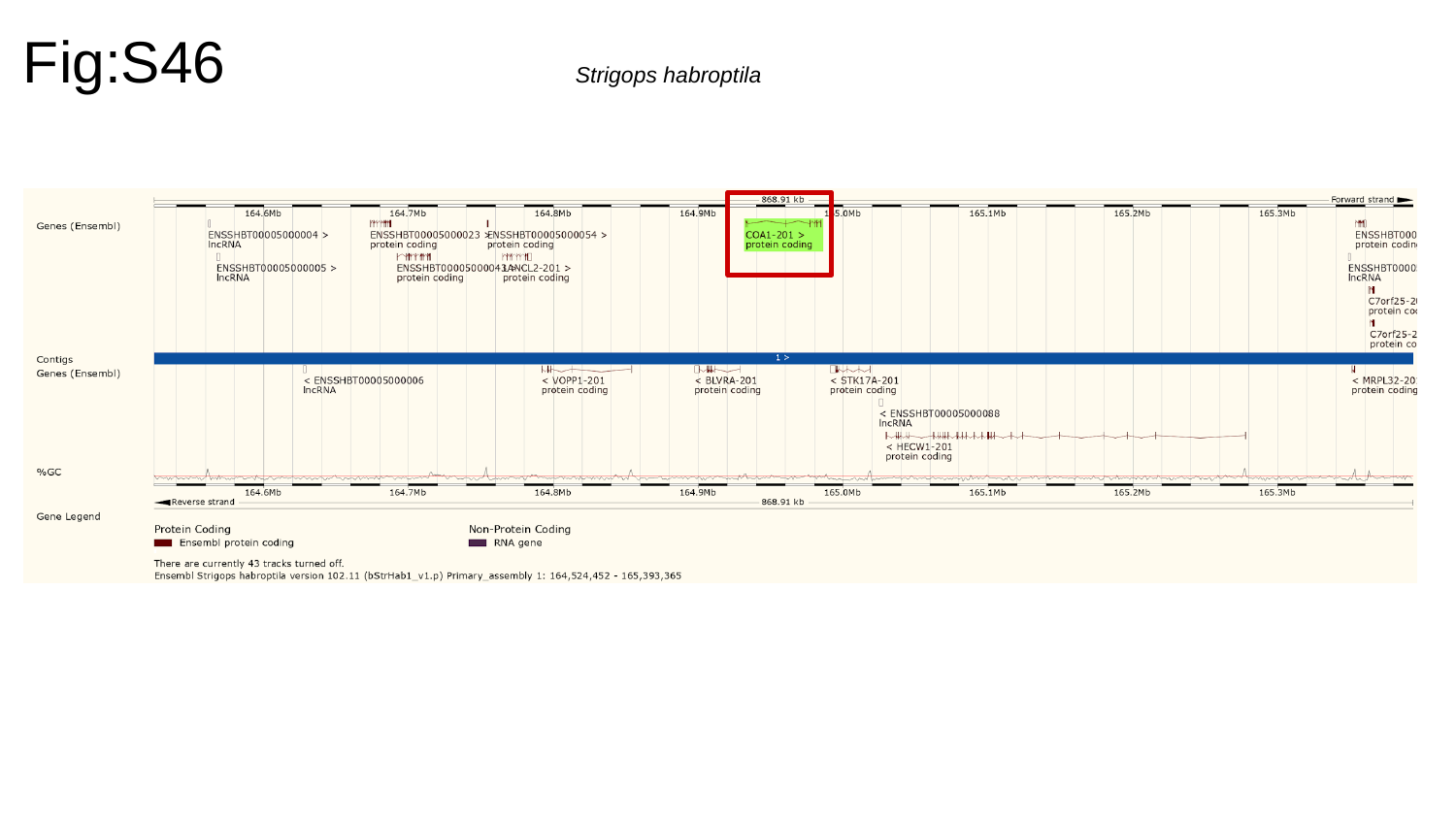

Fig:S46
Strigops habroptila

### Slide 47
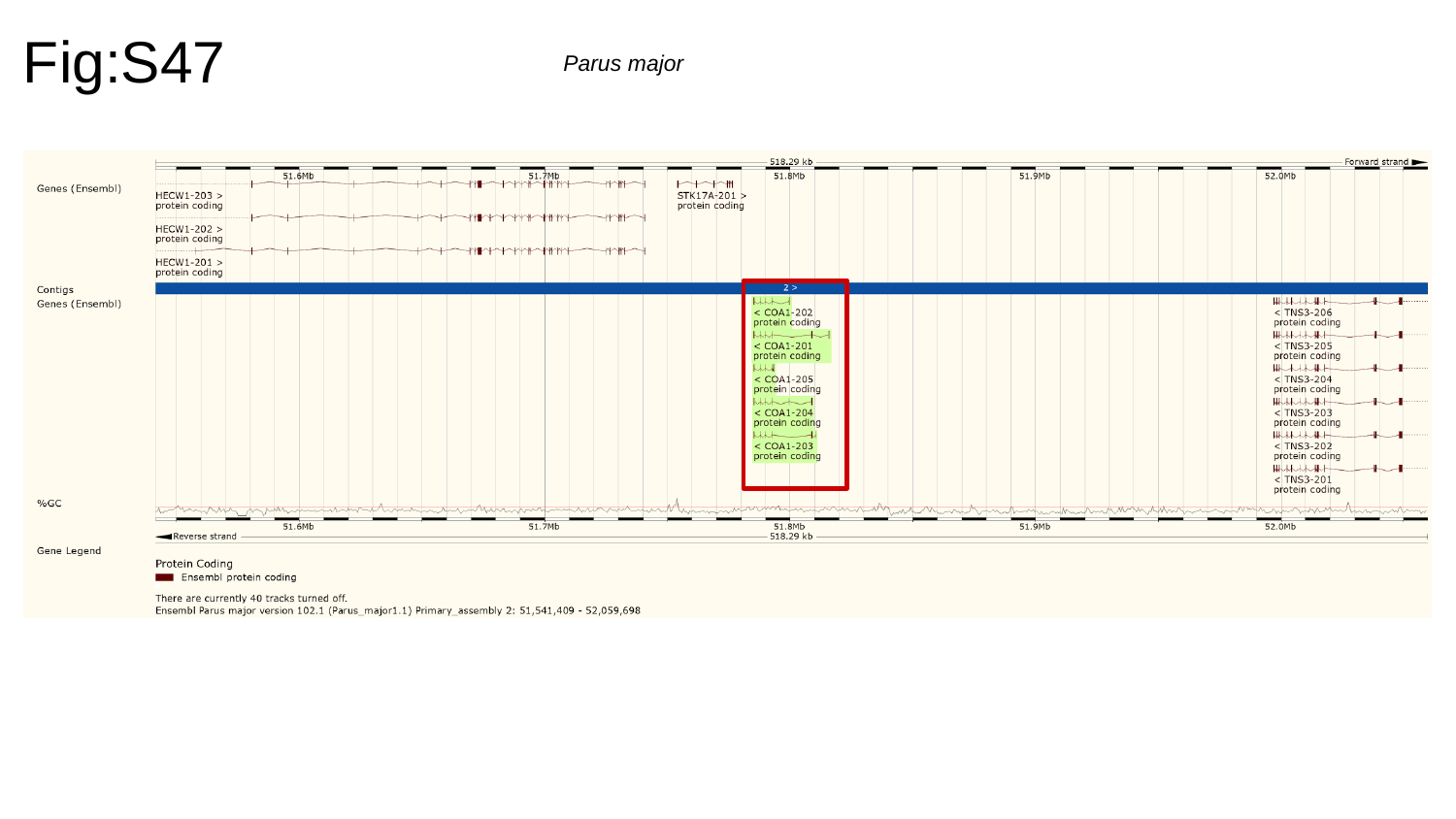

Fig:S47
Parus major

### Slide 48
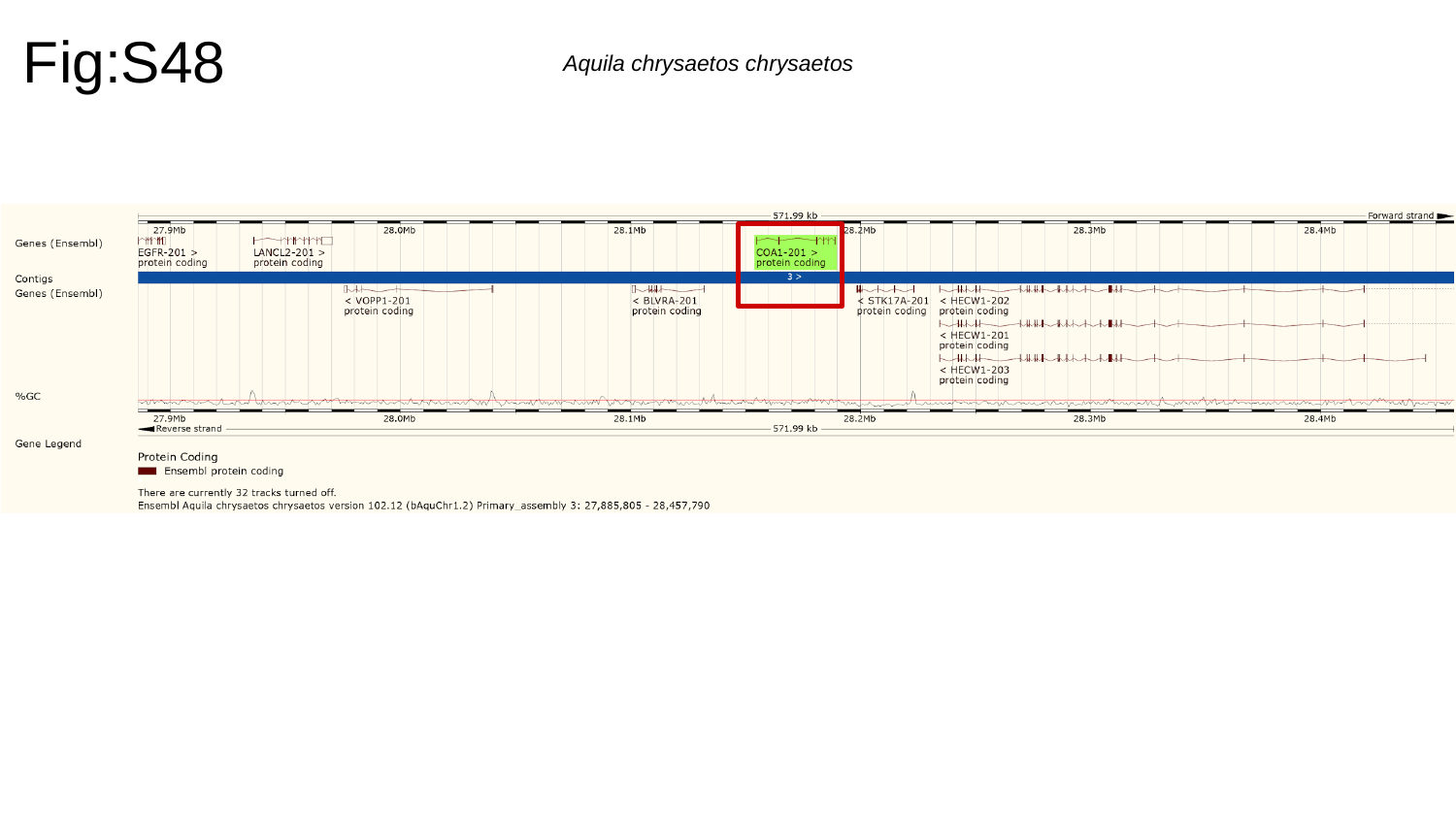

Fig:S48
Aquila chrysaetos chrysaetos

### Slide 49
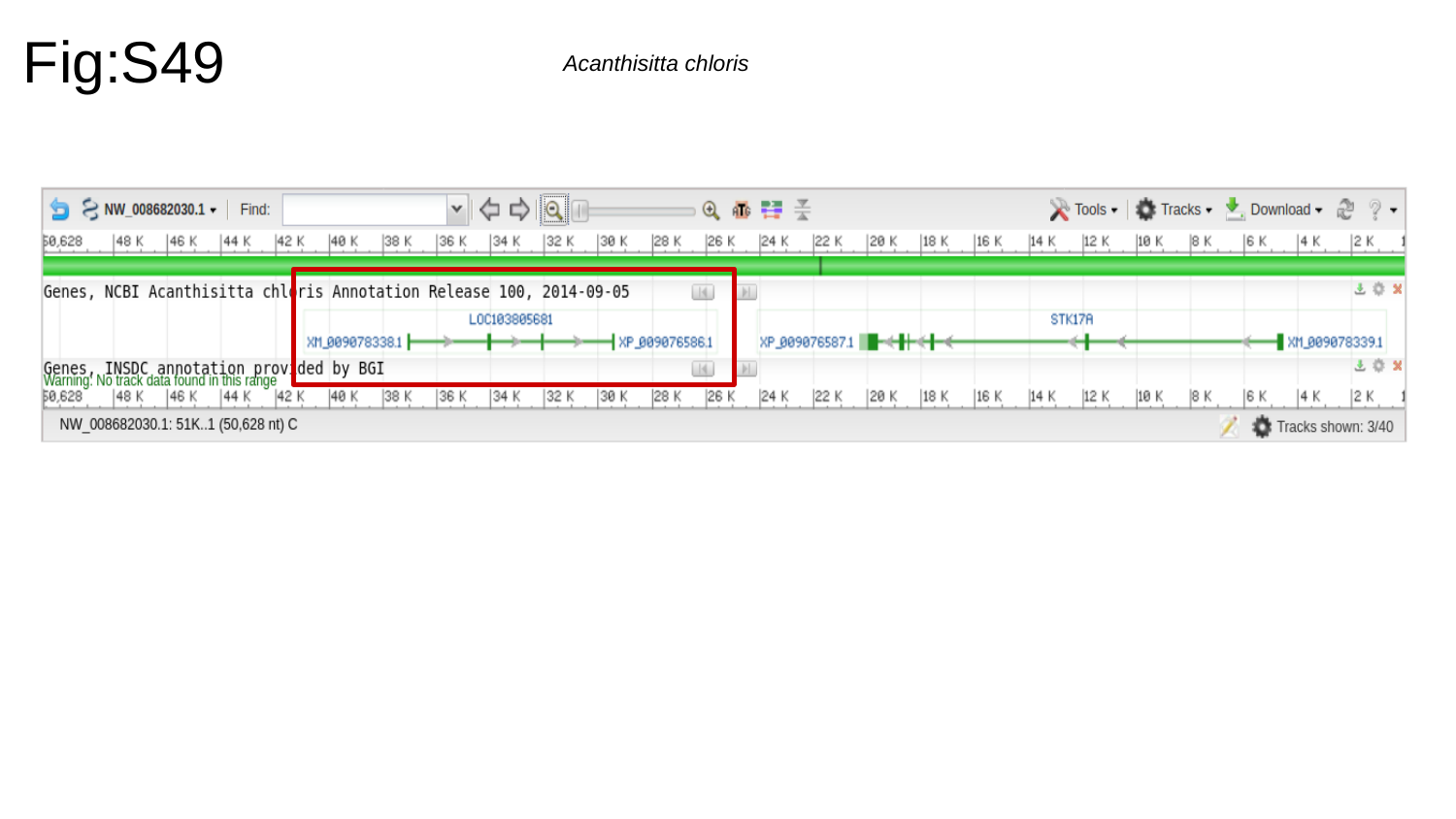

Fig:S49
Acanthisitta chloris

### Slide 50
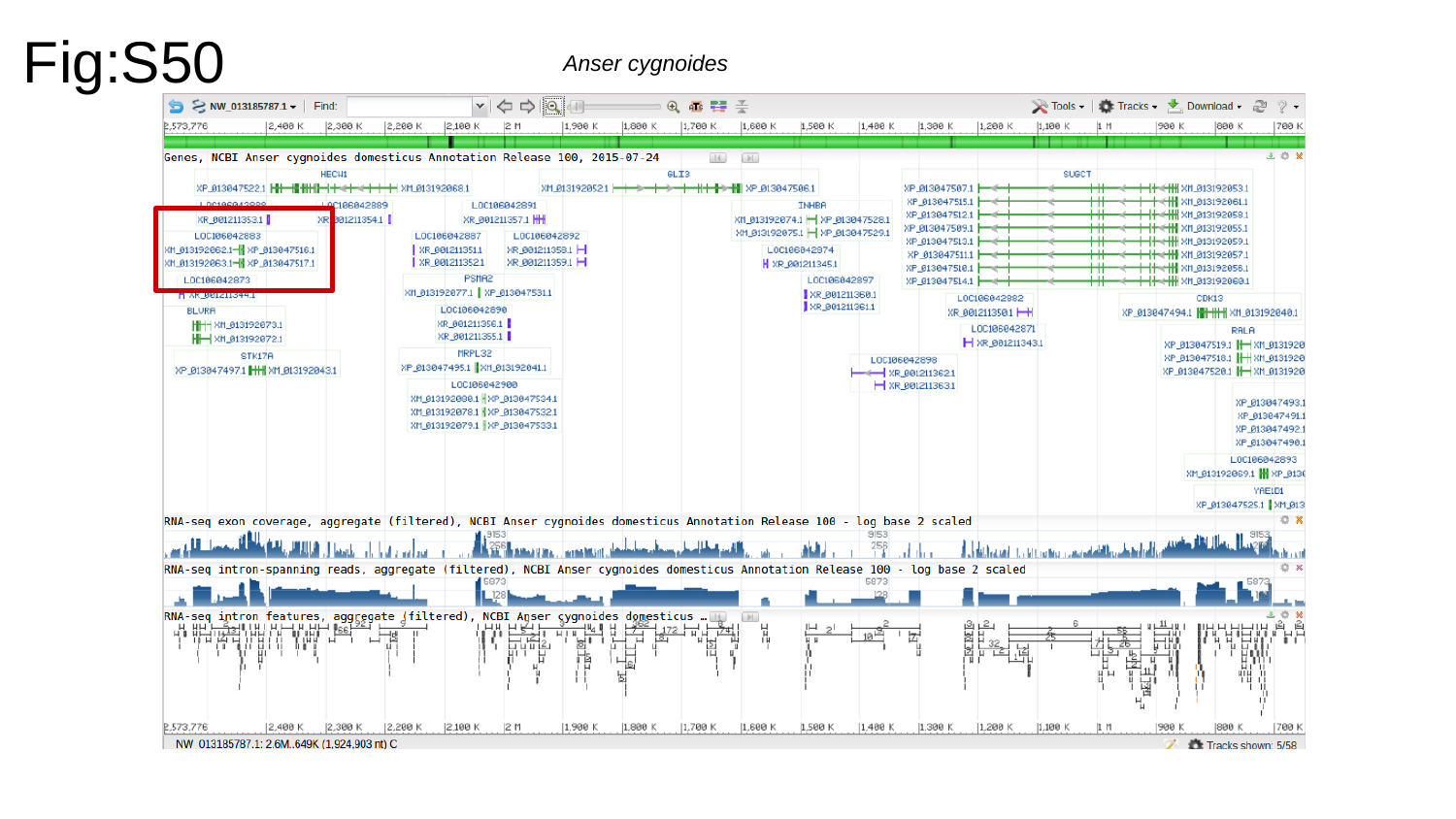

Fig:S50
Anser cygnoides

### Slide 51
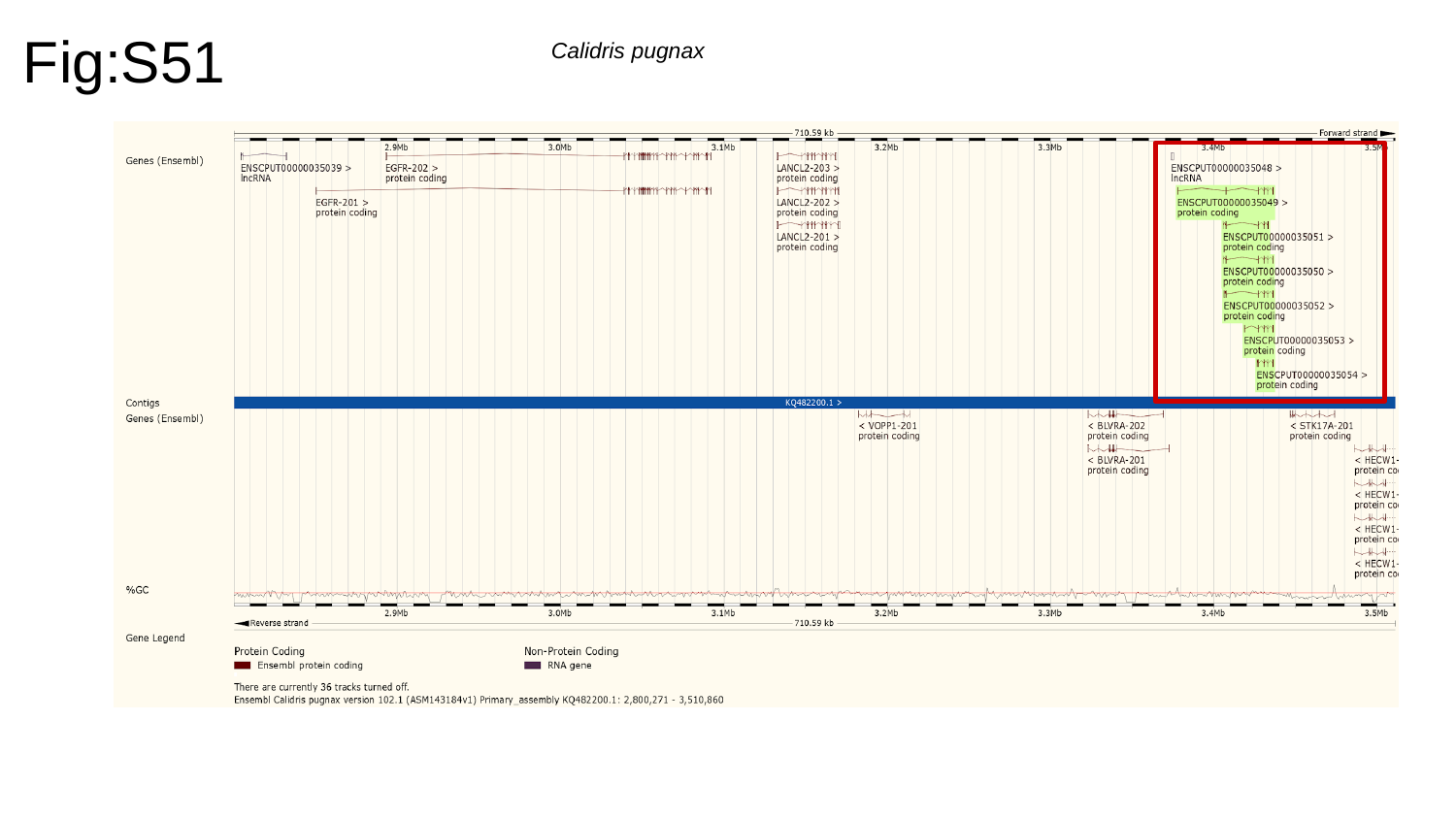

Fig:S51
Calidris pugnax

### Slide 52
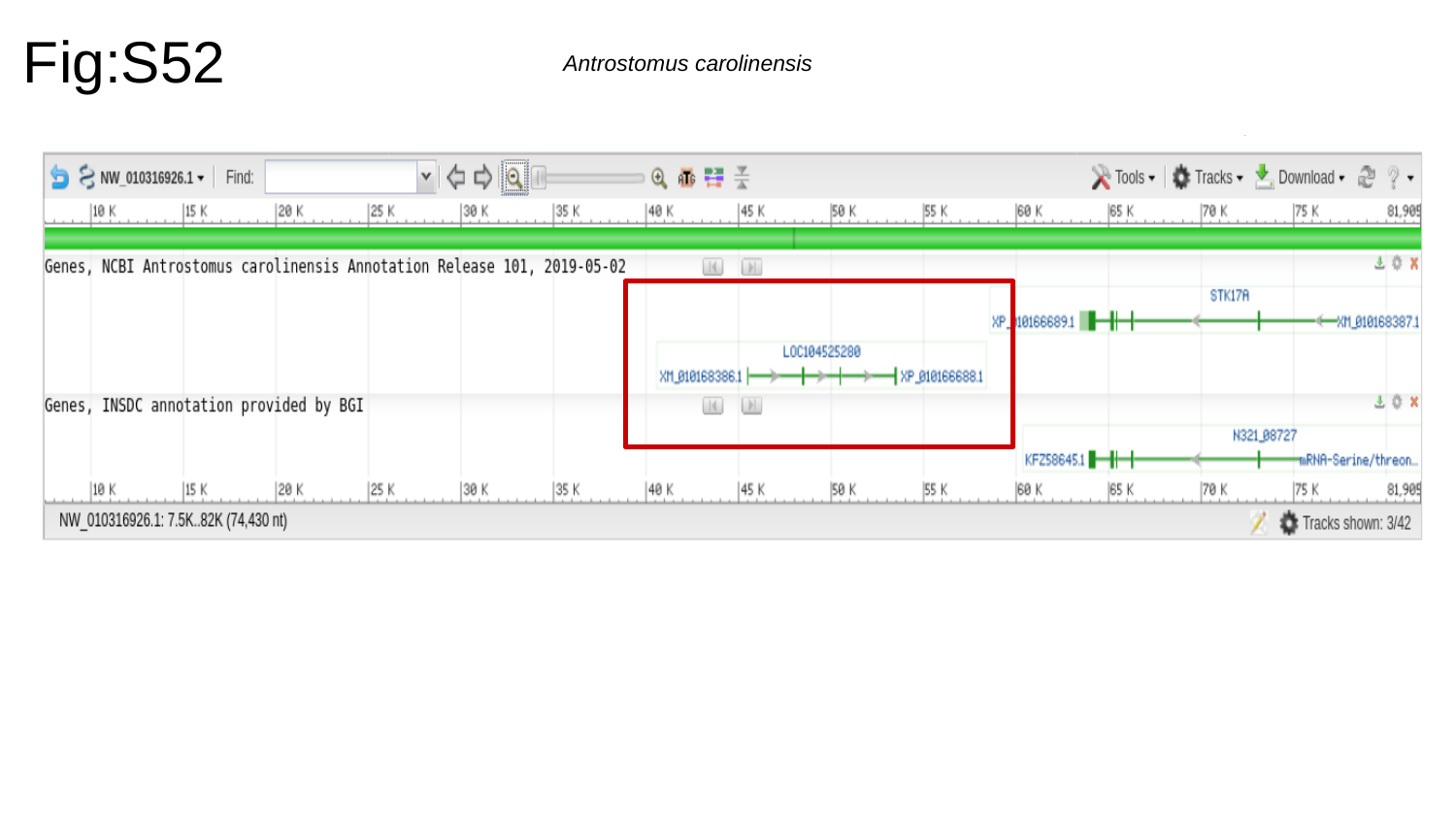

Fig:S52
Antrostomus carolinensis

### Slide 53
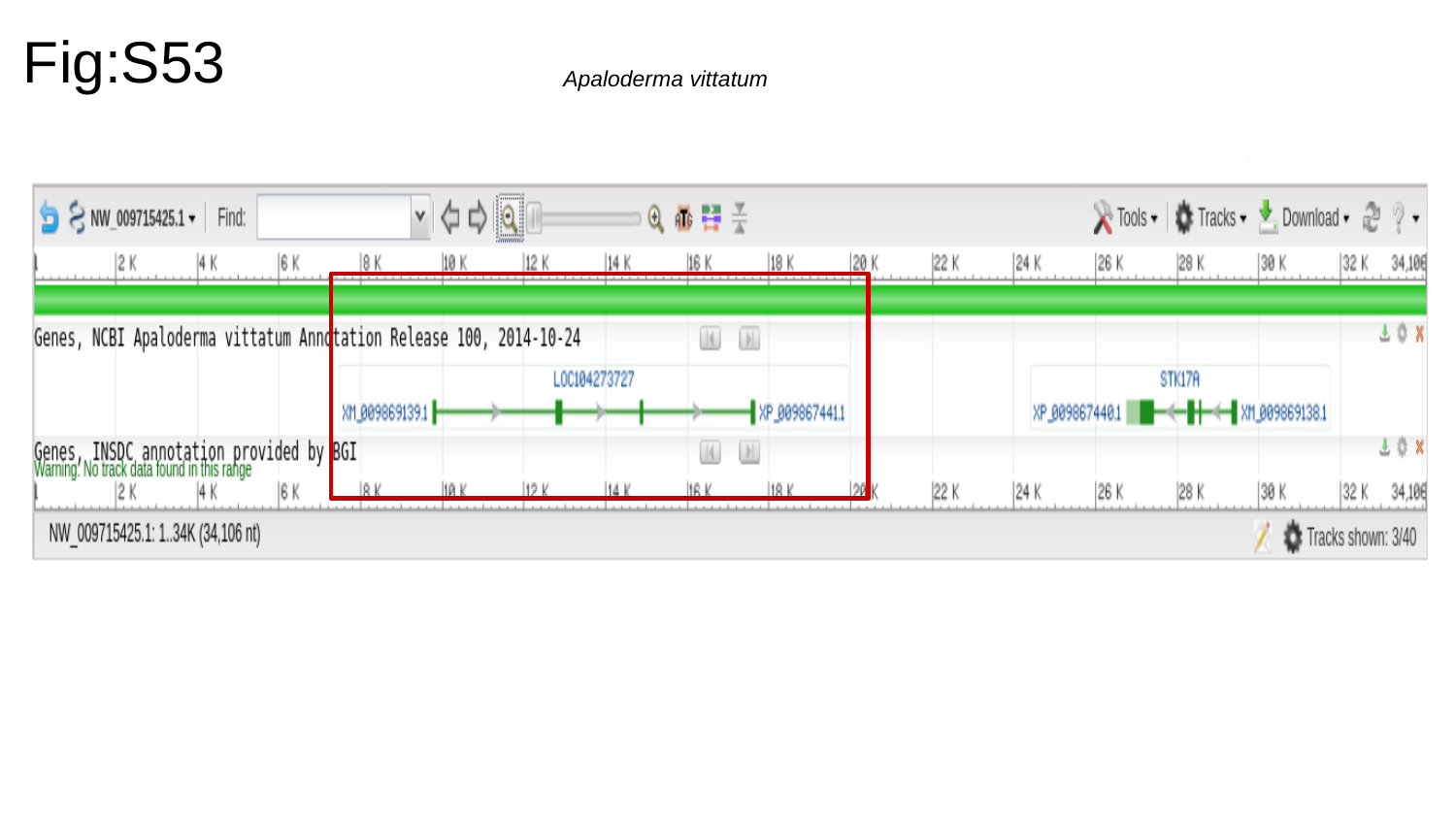

Fig:S53
Apaloderma vittatum

### Slide 54
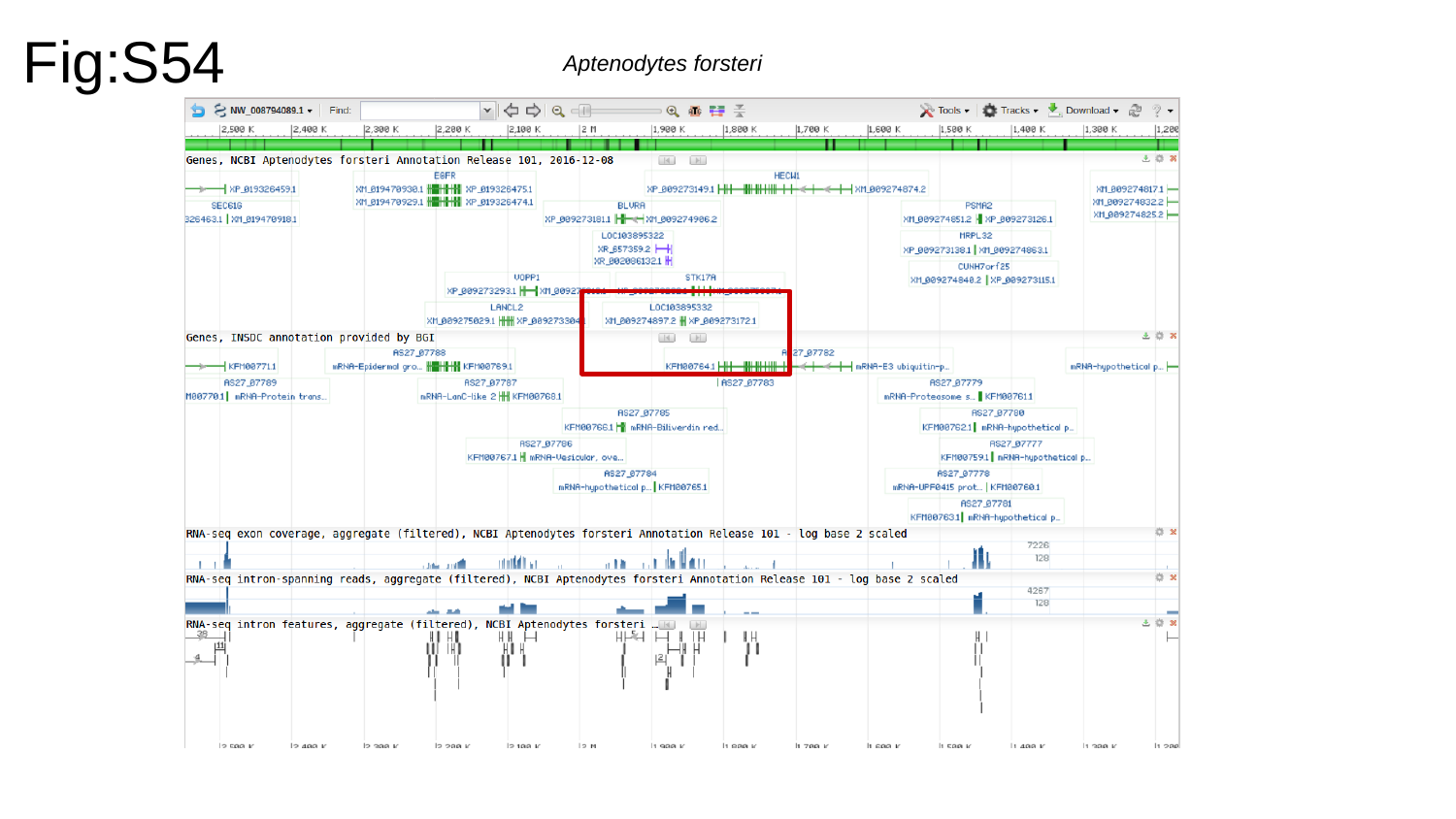

Fig:S54
Aptenodytes forsteri

### Slide 55
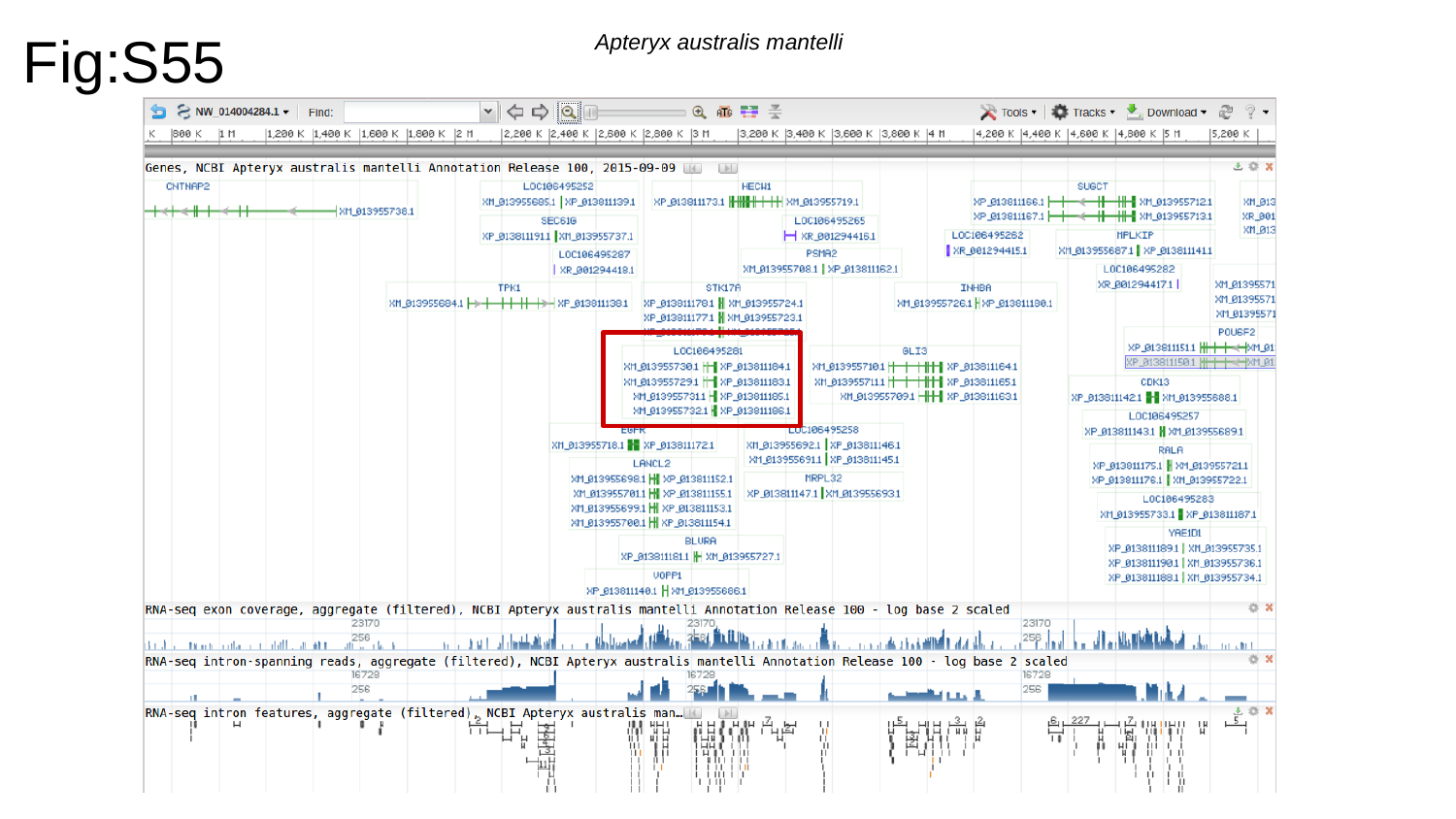

Fig:S55
Apteryx australis mantelli

### Slide 56
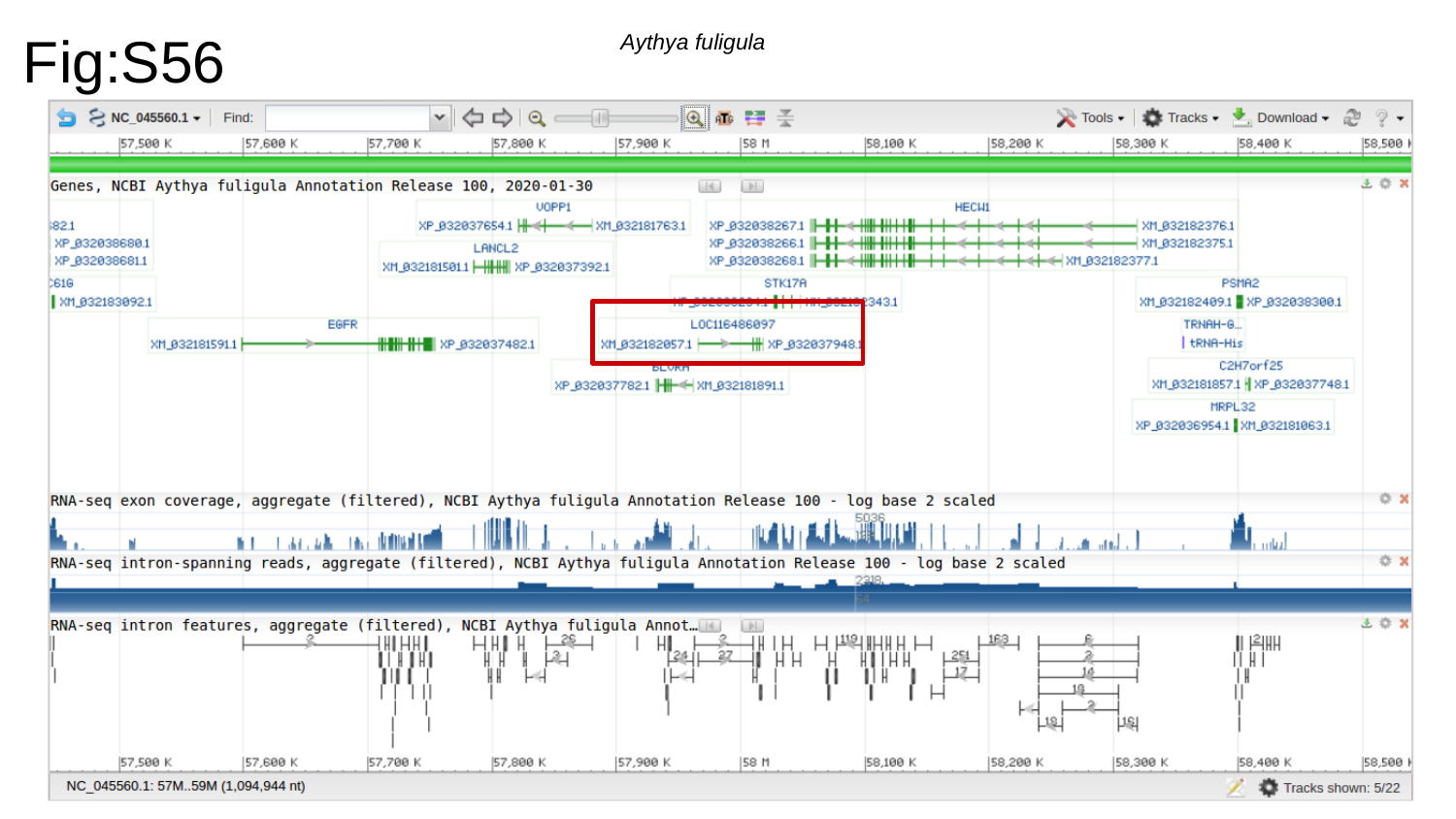

Fig:S56
Aythya fuligula

### Slide 57
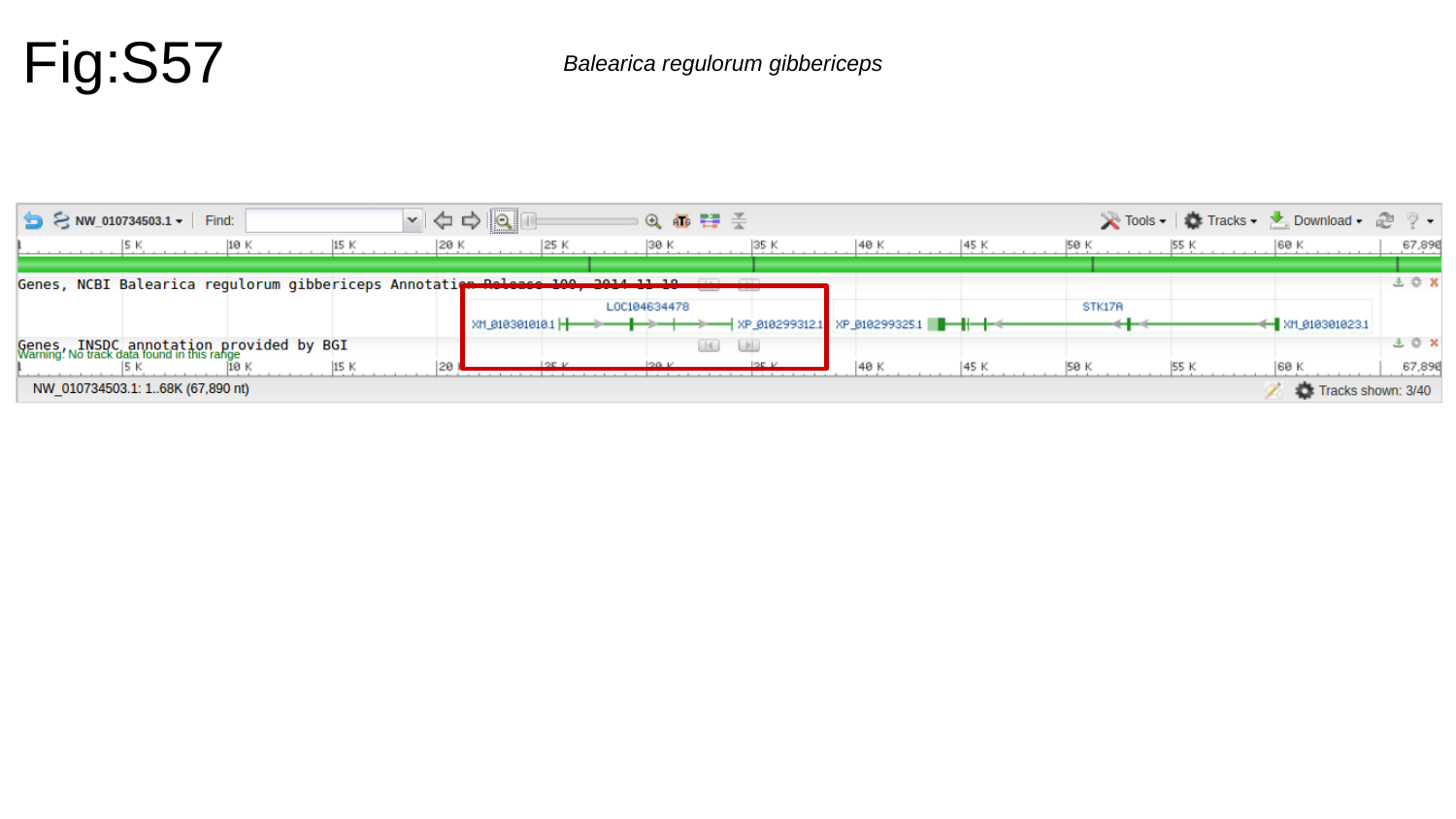

Fig:S57
Balearica regulorum gibbericeps

### Slide 58
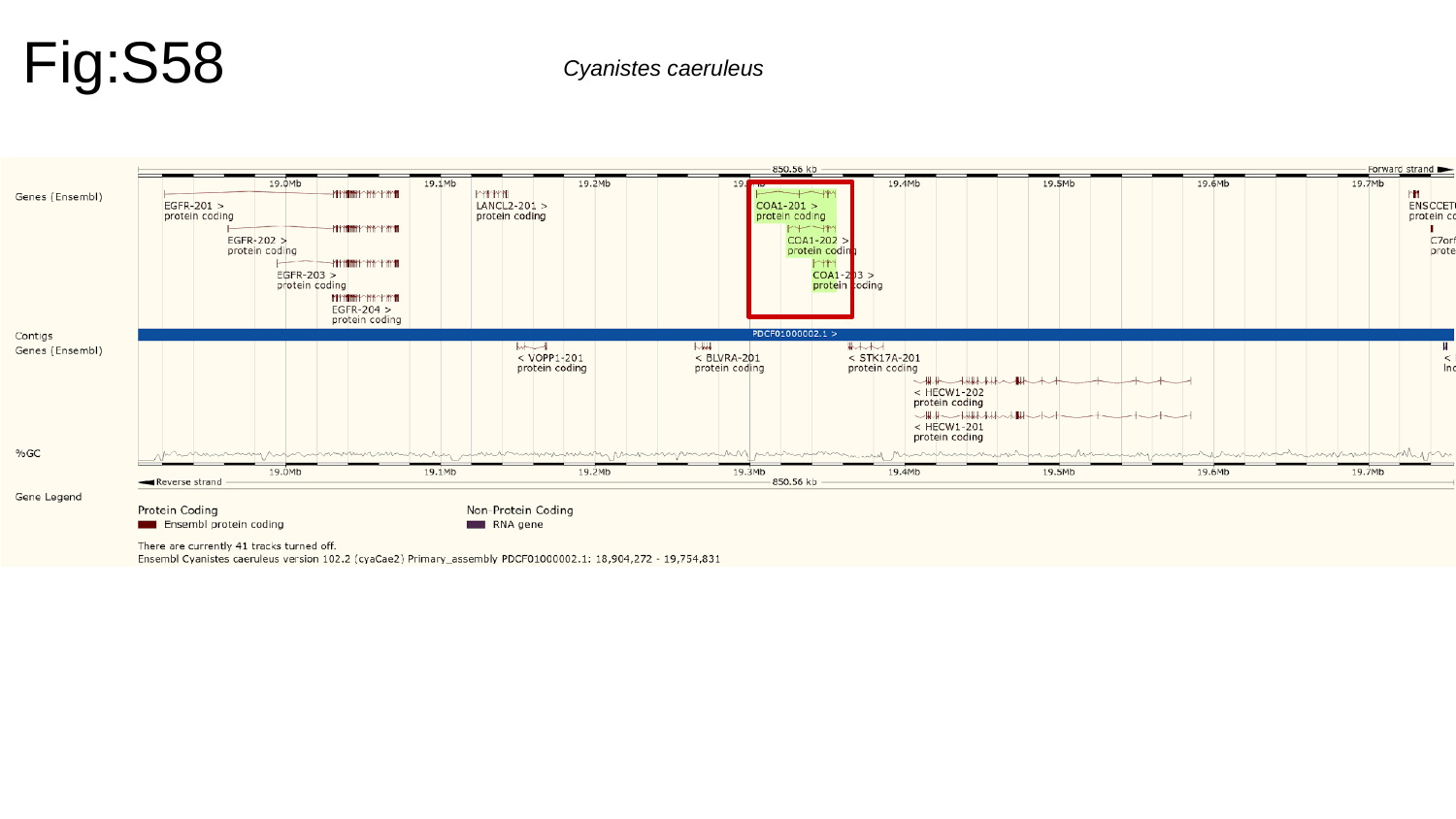

Fig:S58
Cyanistes caeruleus

### Slide 59
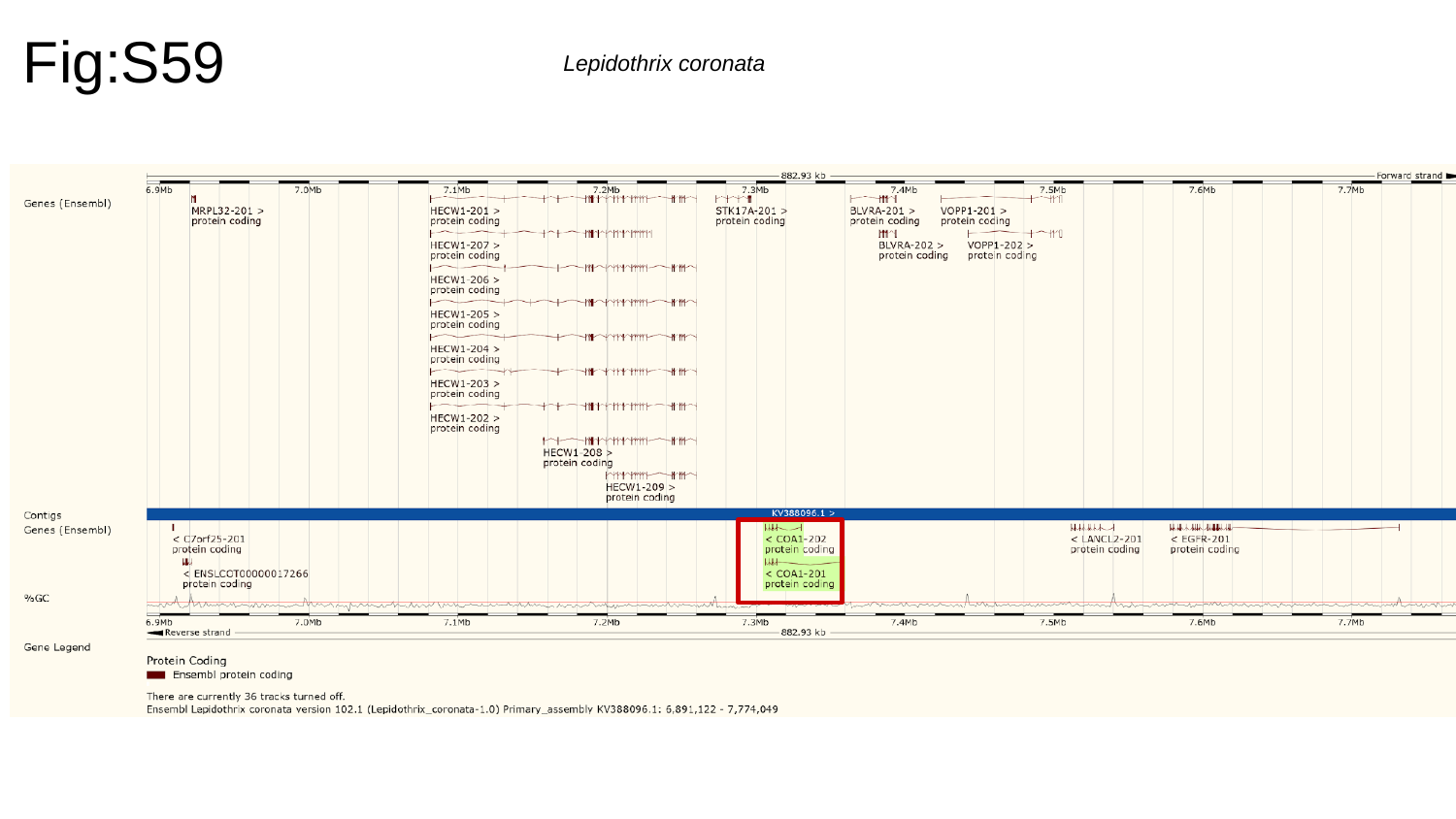

Fig:S59
Lepidothrix coronata

### Slide 60
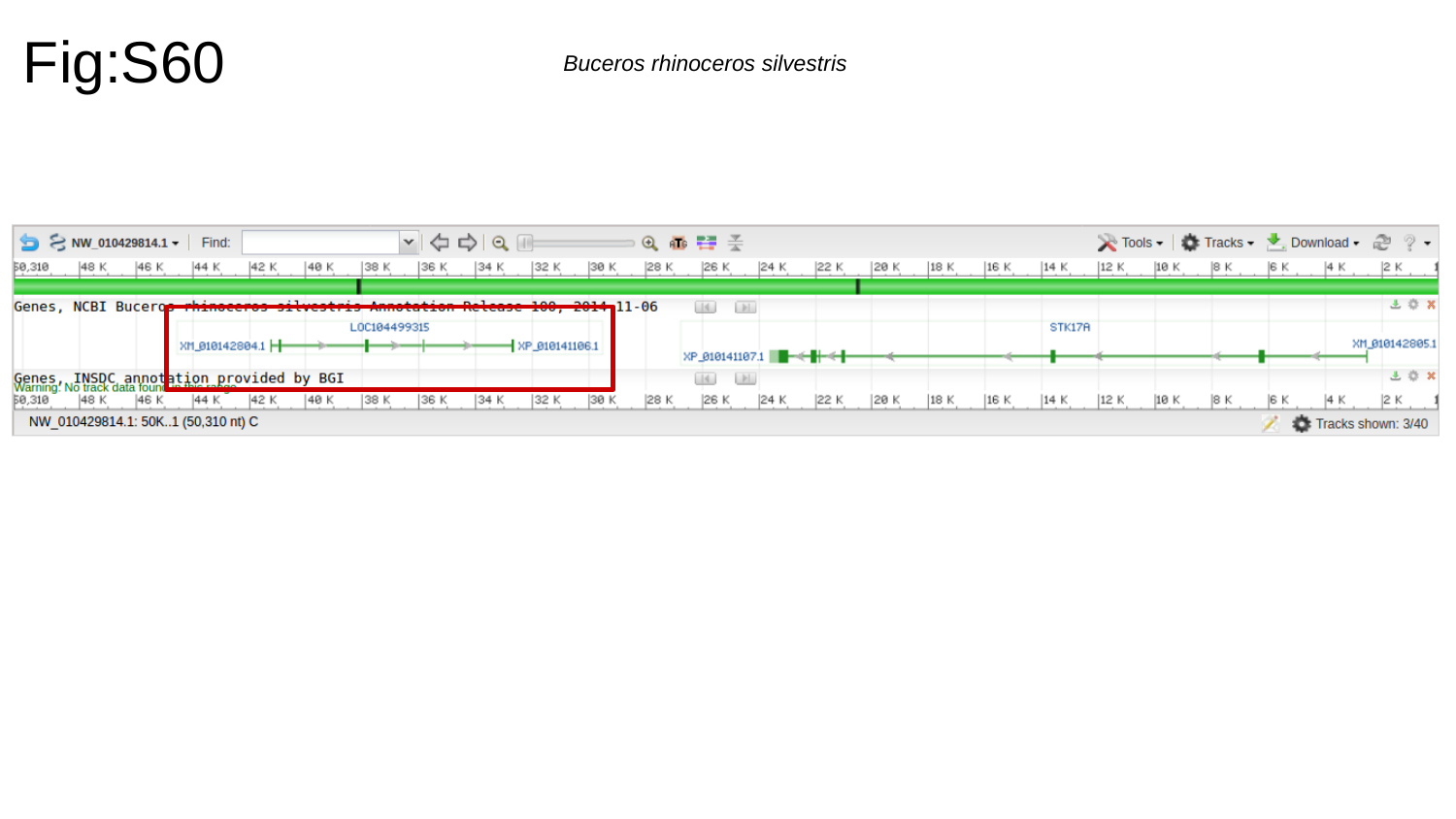

Fig:S60
Buceros rhinoceros silvestris

### Slide 61
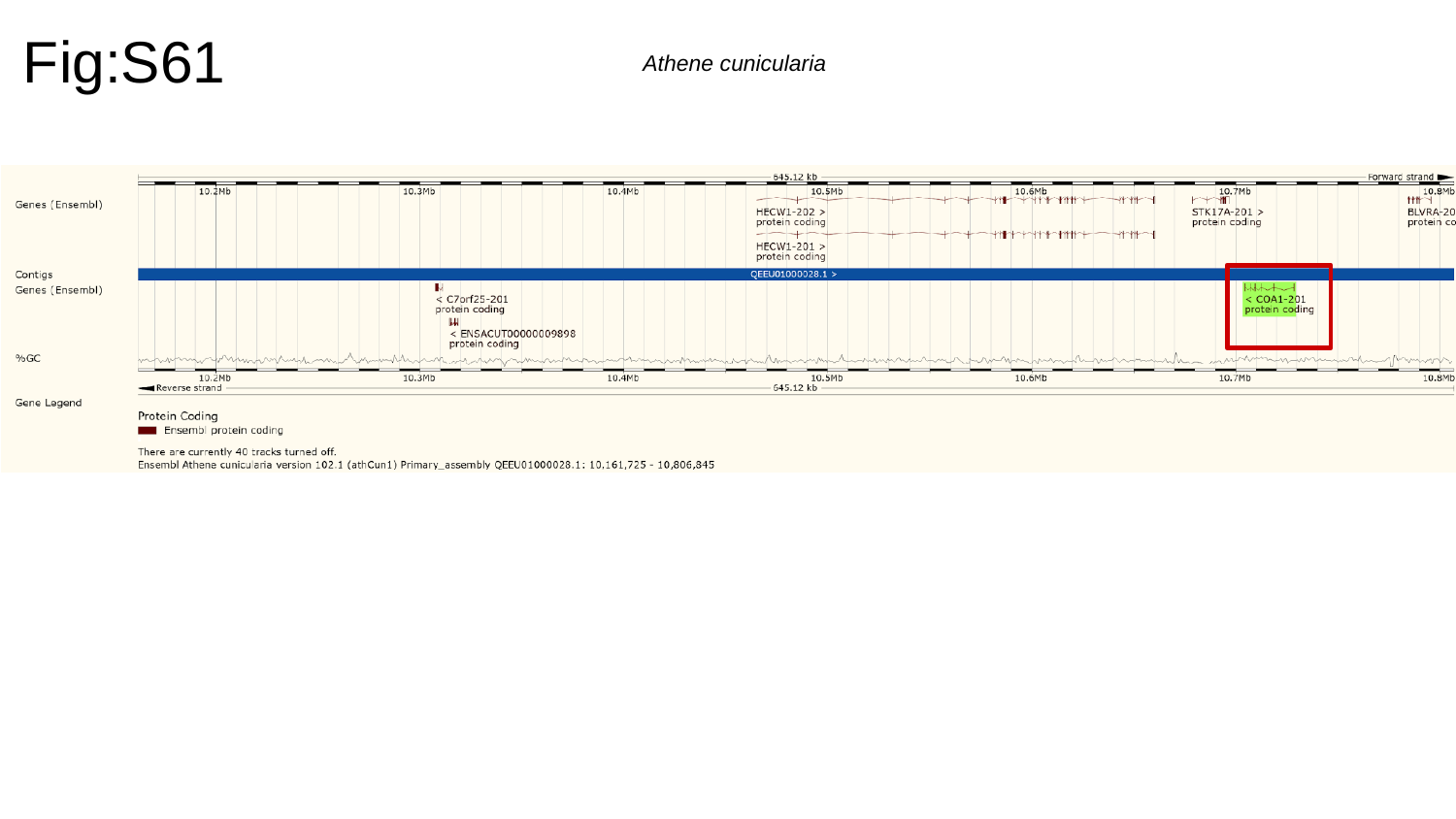

Fig:S61
Athene cunicularia

### Slide 62
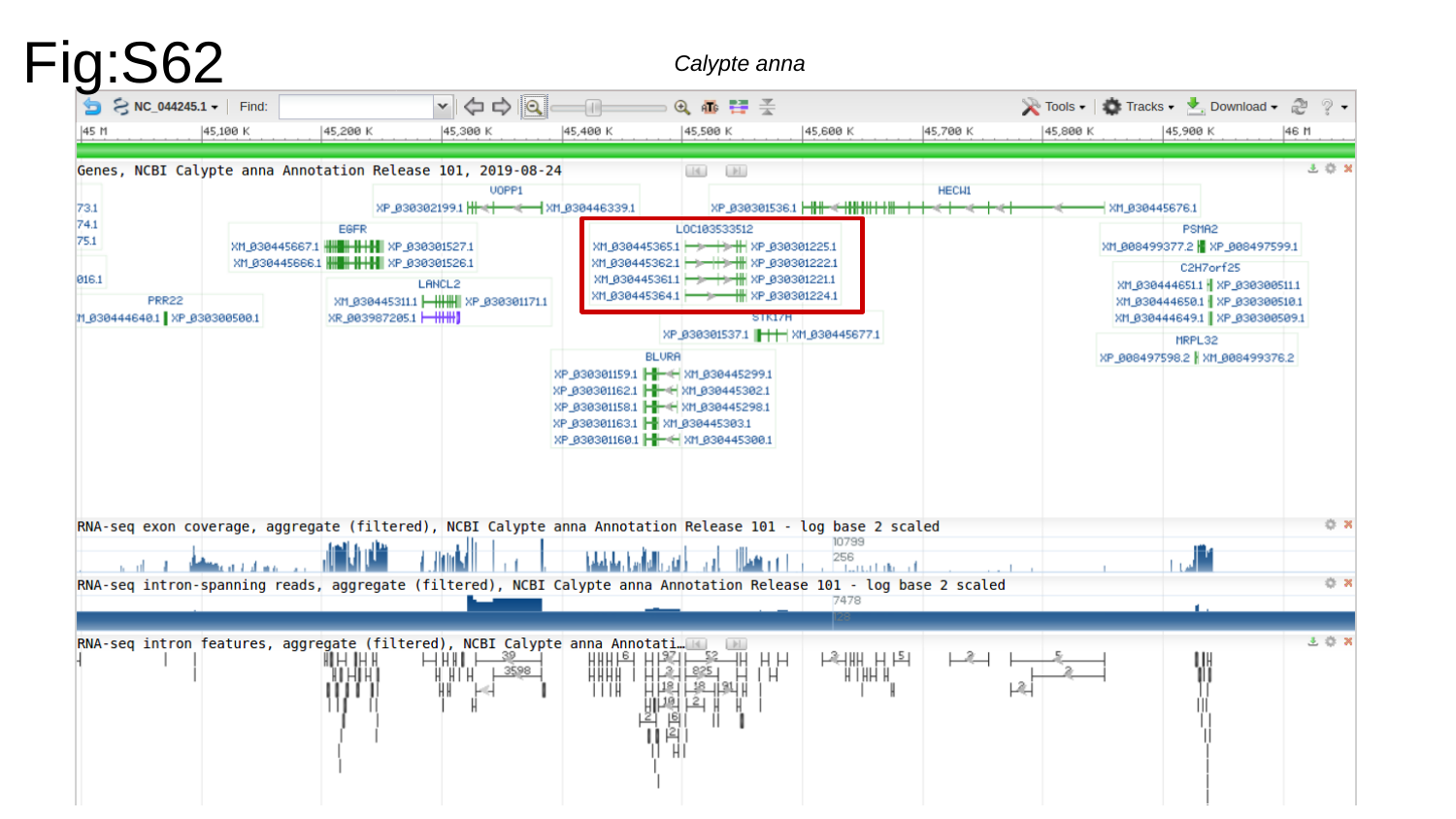

Fig:S62
Calypte anna

### Slide 63
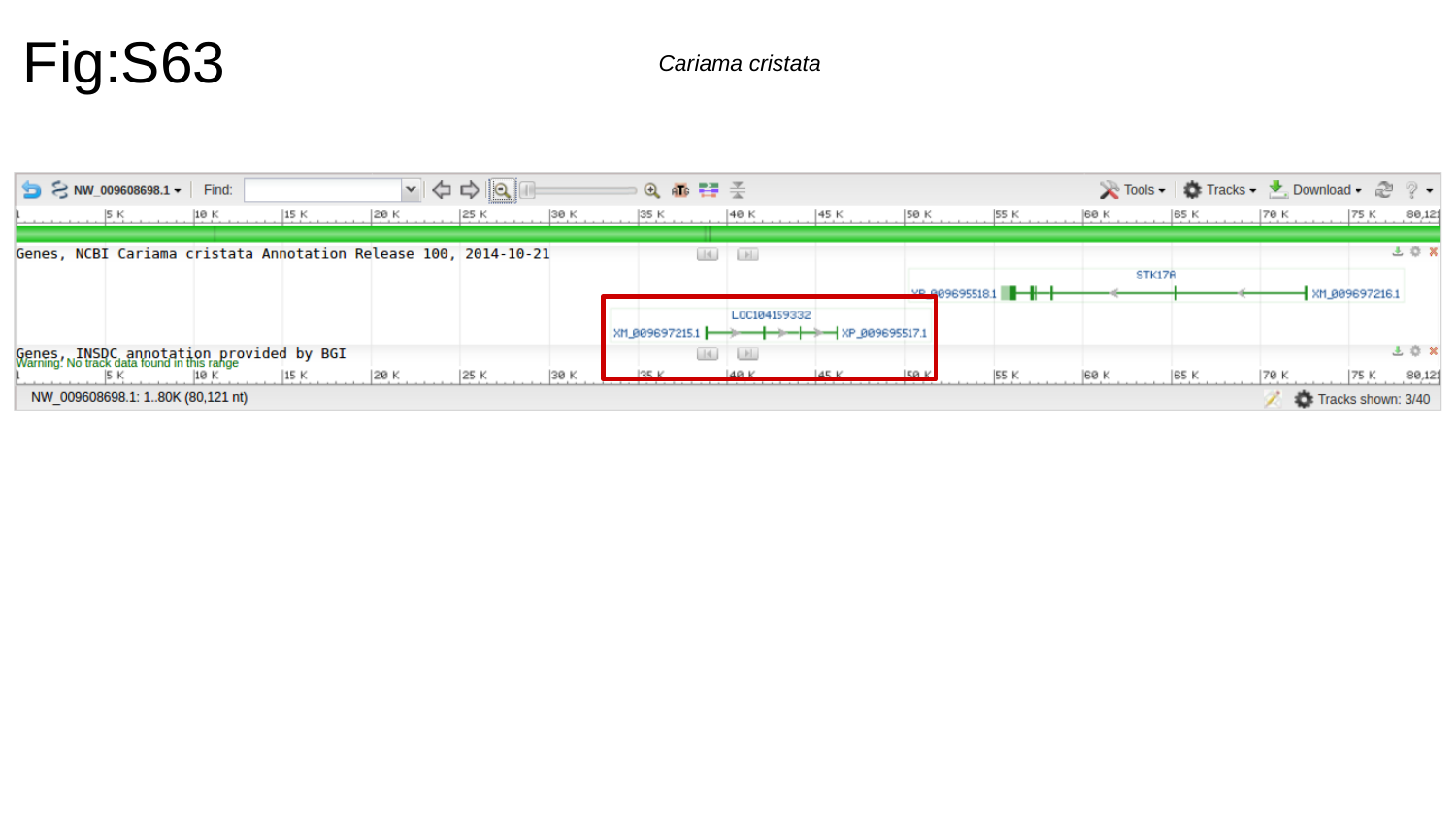

Fig:S63
Cariama cristata

### Slide 64
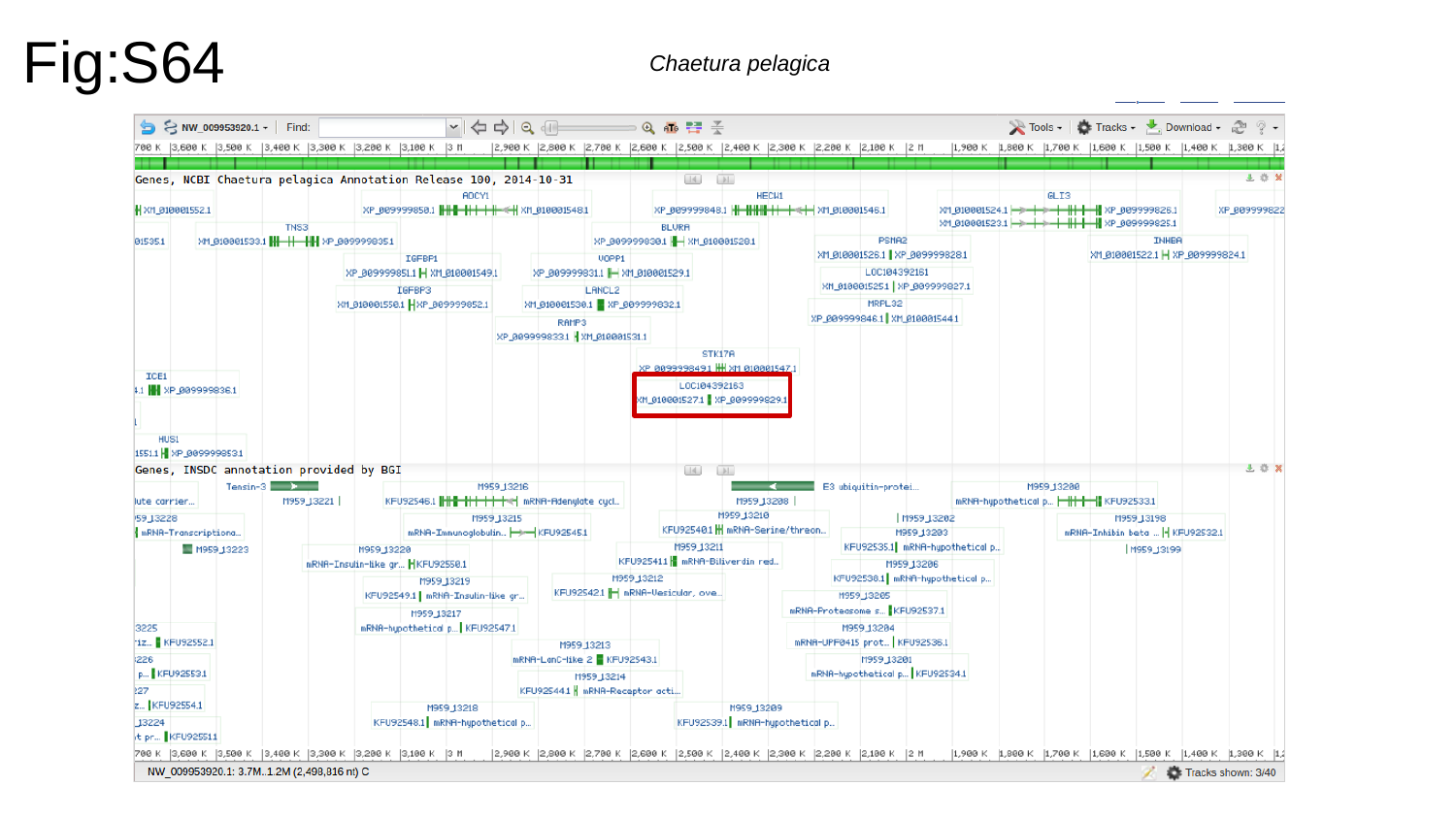

Fig:S64
Chaetura pelagica

### Slide 65
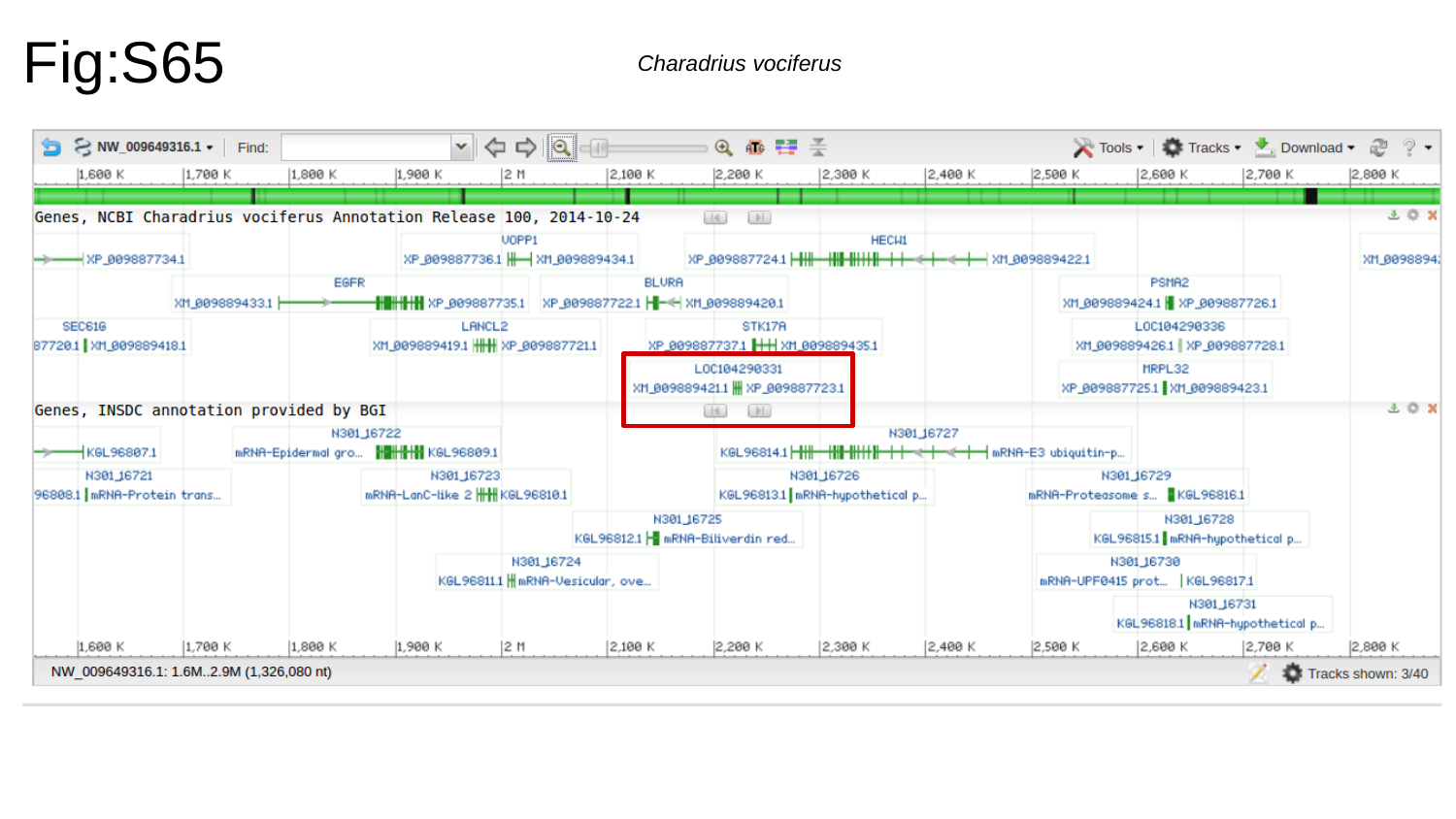

Fig:S65
Charadrius vociferus

### Slide 66
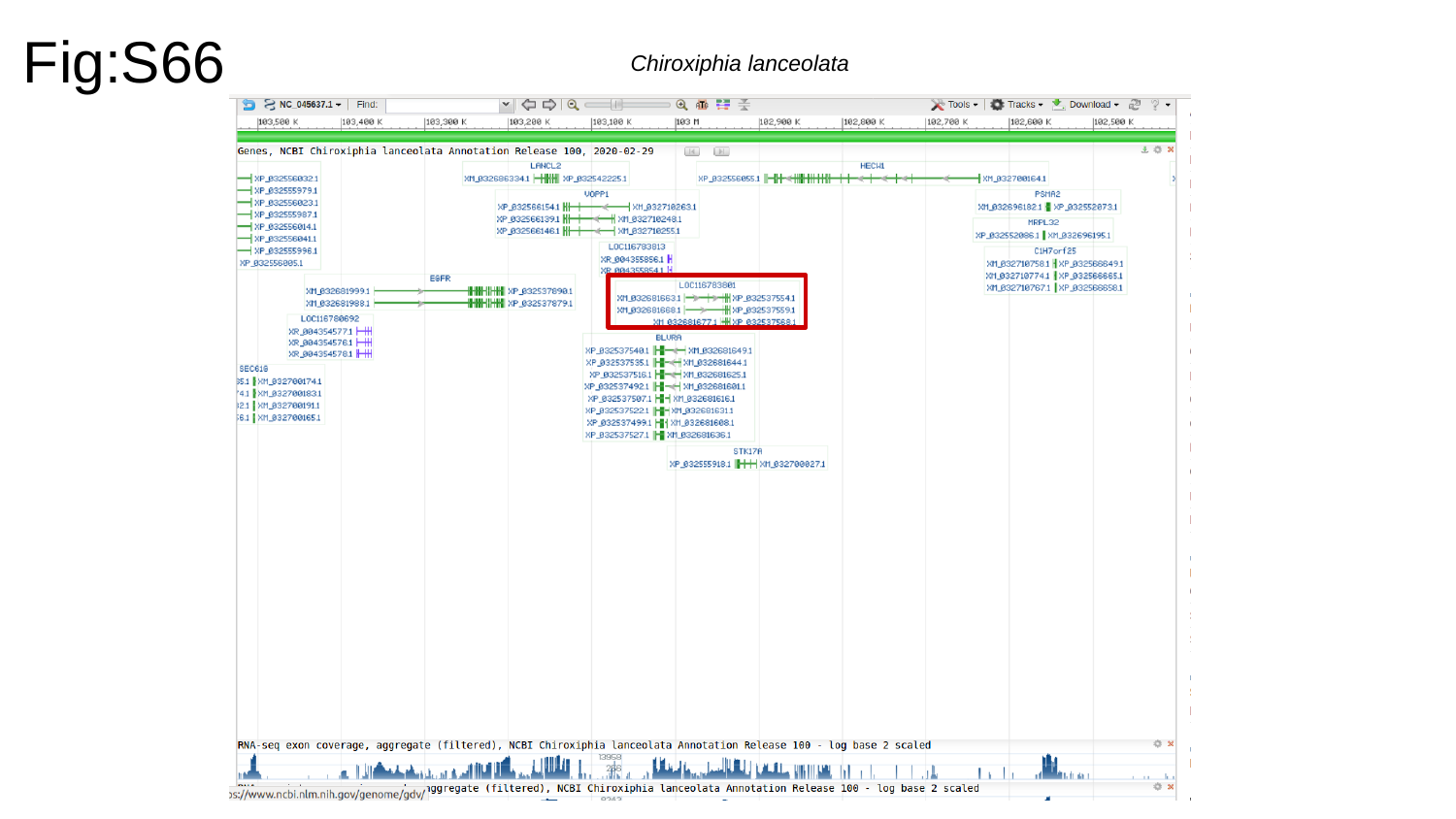

Fig:S66
Chiroxiphia lanceolata

### Slide 67
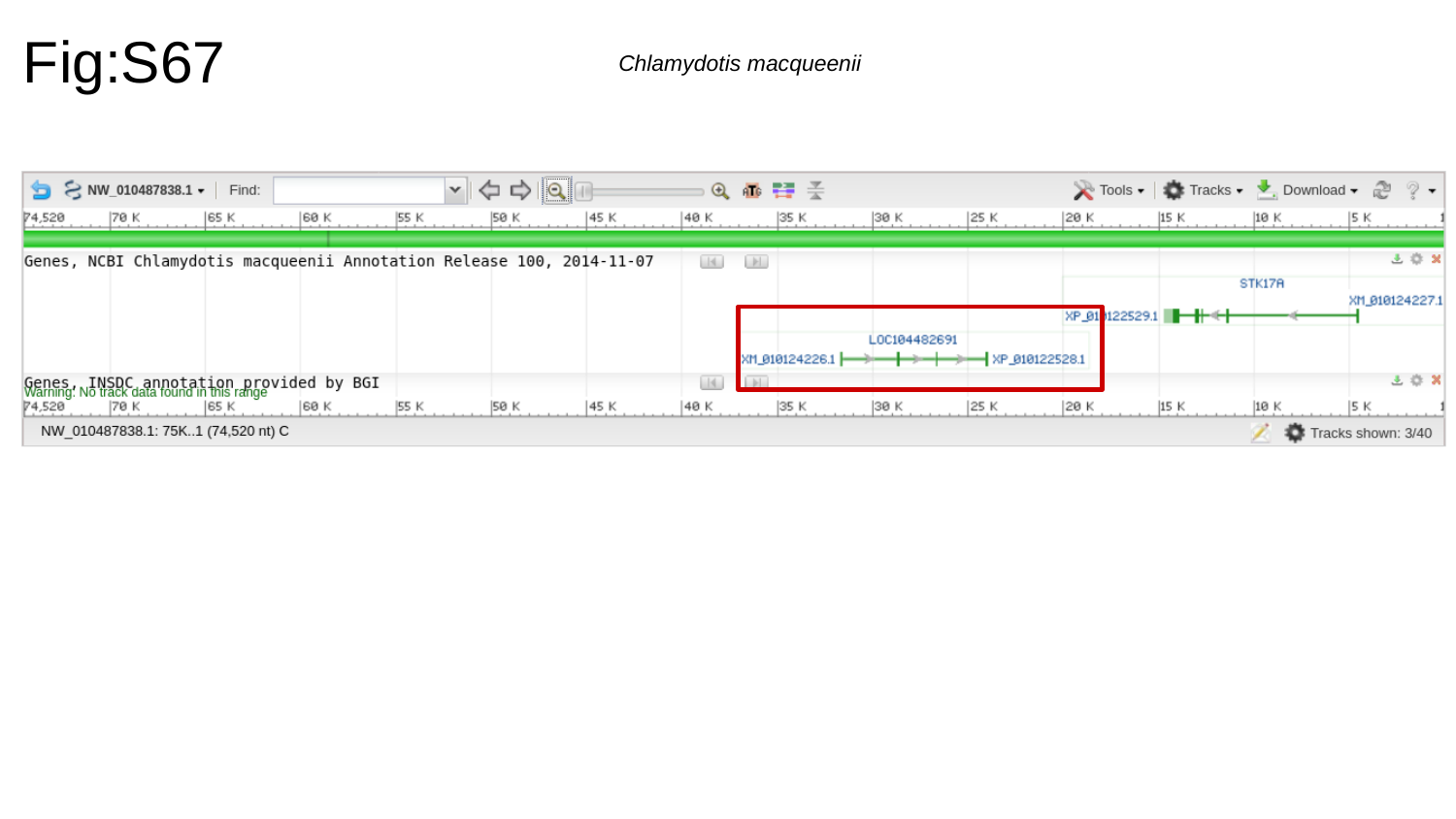

Fig:S67
Chlamydotis macqueenii

### Slide 68
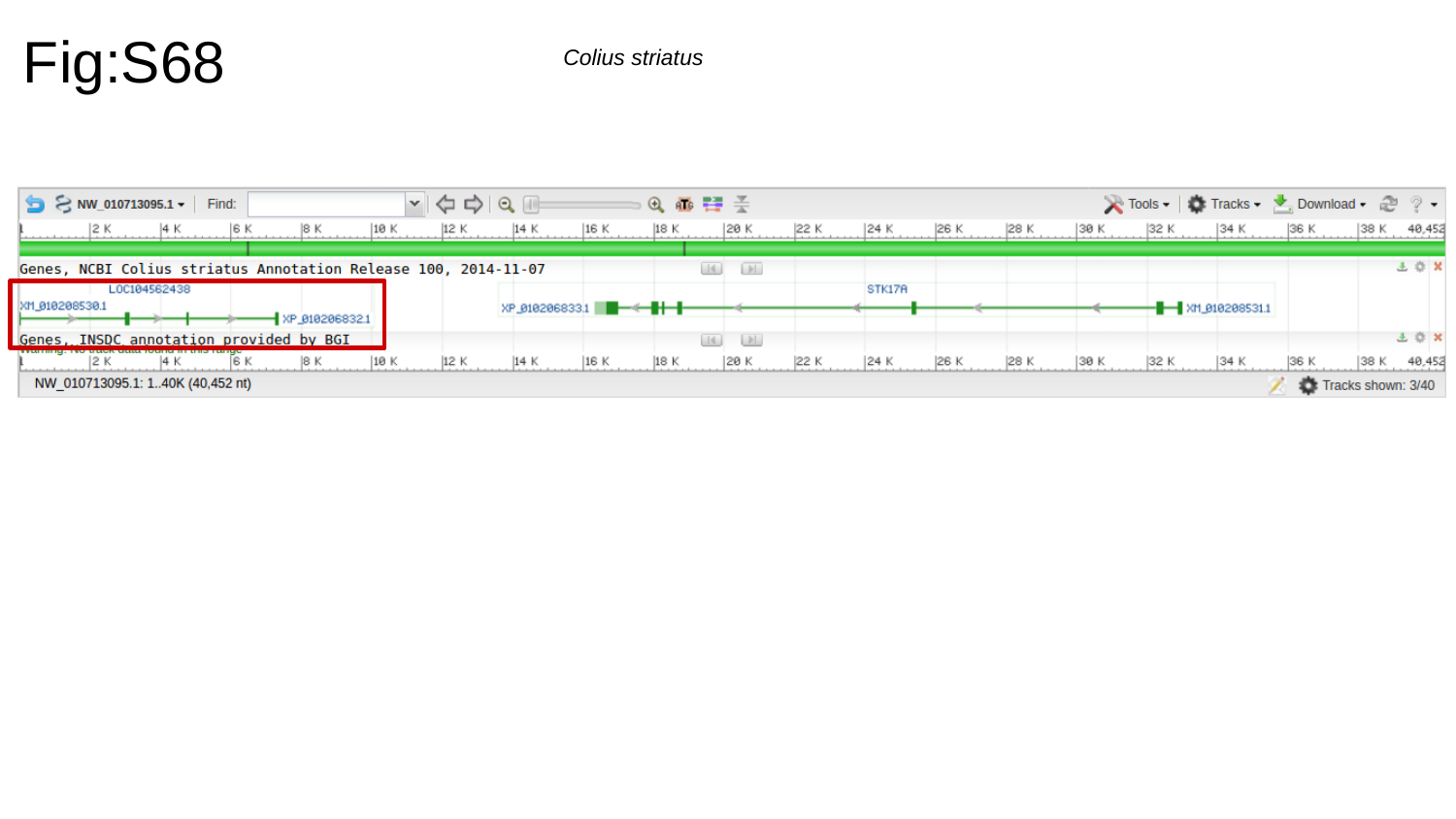

Fig:S68
Colius striatus

### Slide 69
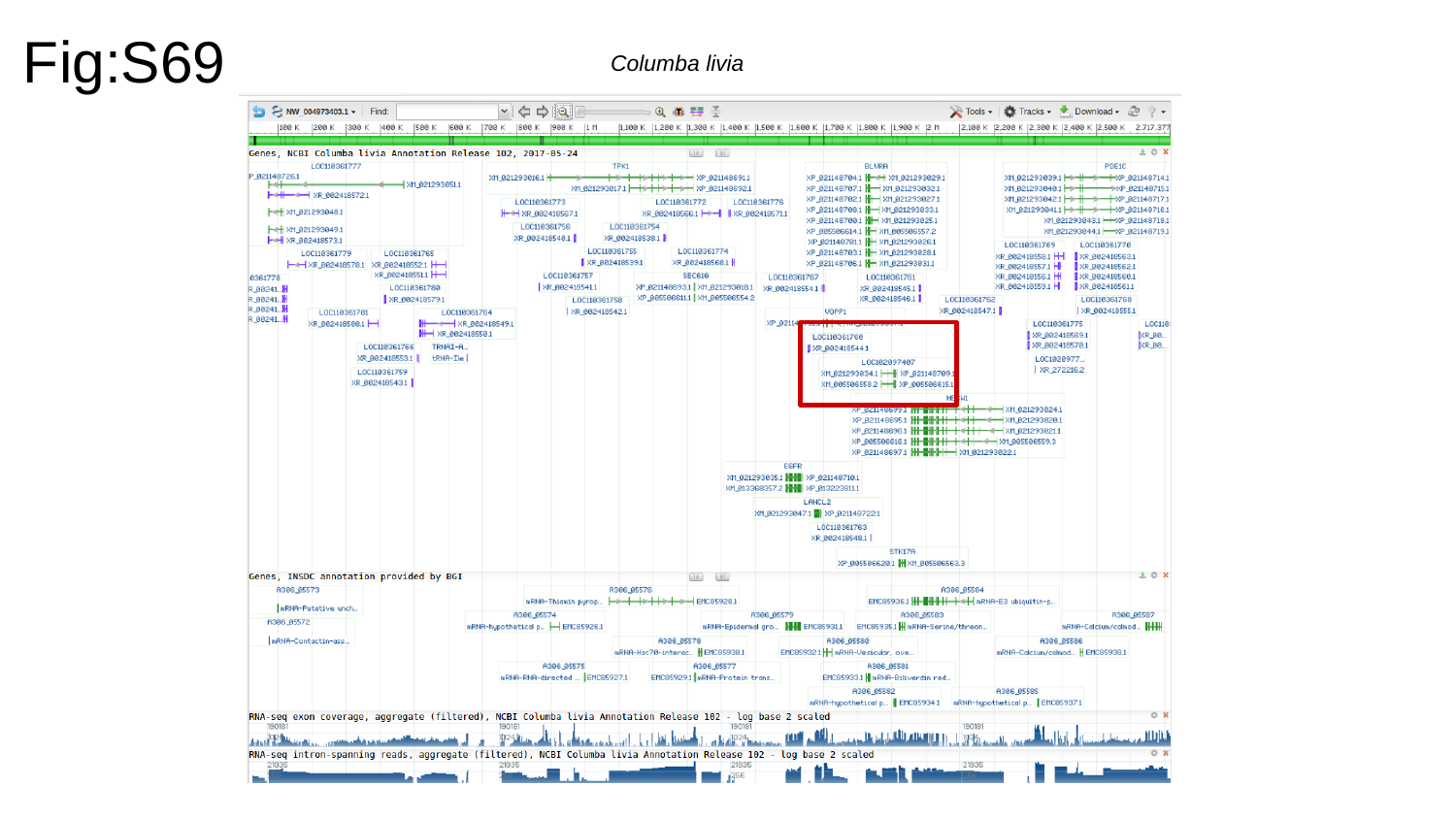

Fig:S69
Columba livia

### Slide 70
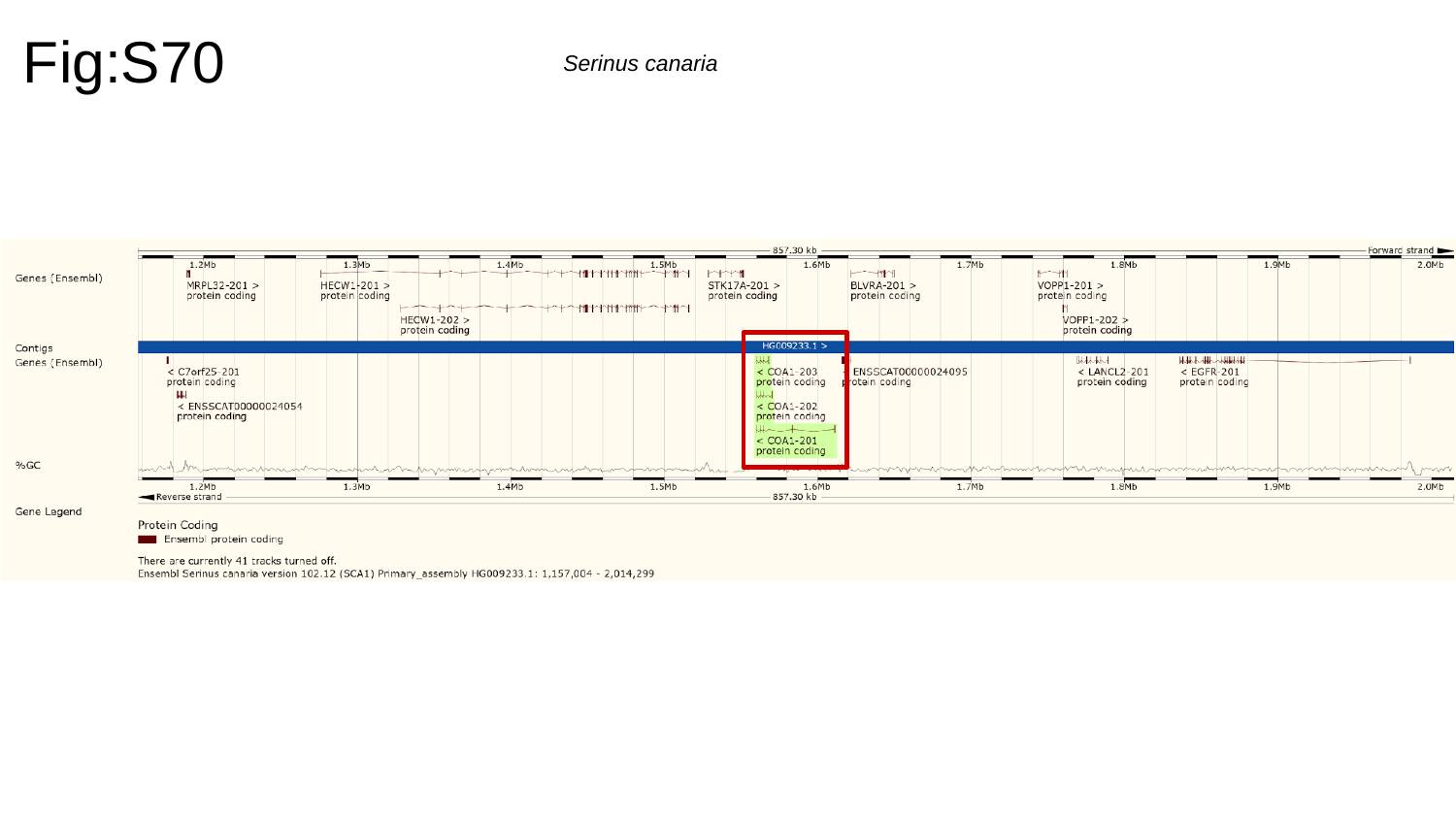

Fig:S70
Serinus canaria

### Slide 71
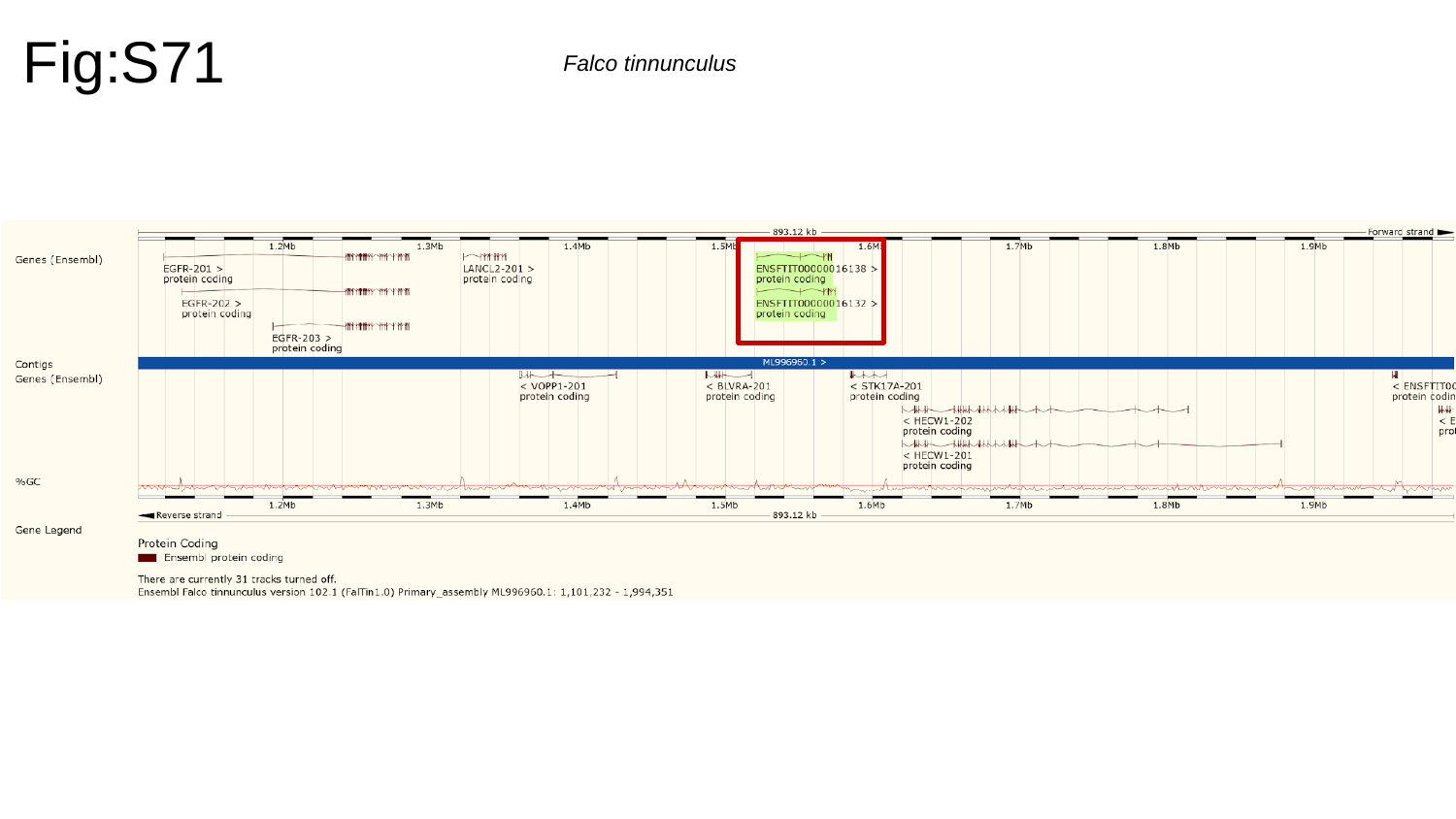

Fig:S71
Falco tinnunculus

### Slide 72
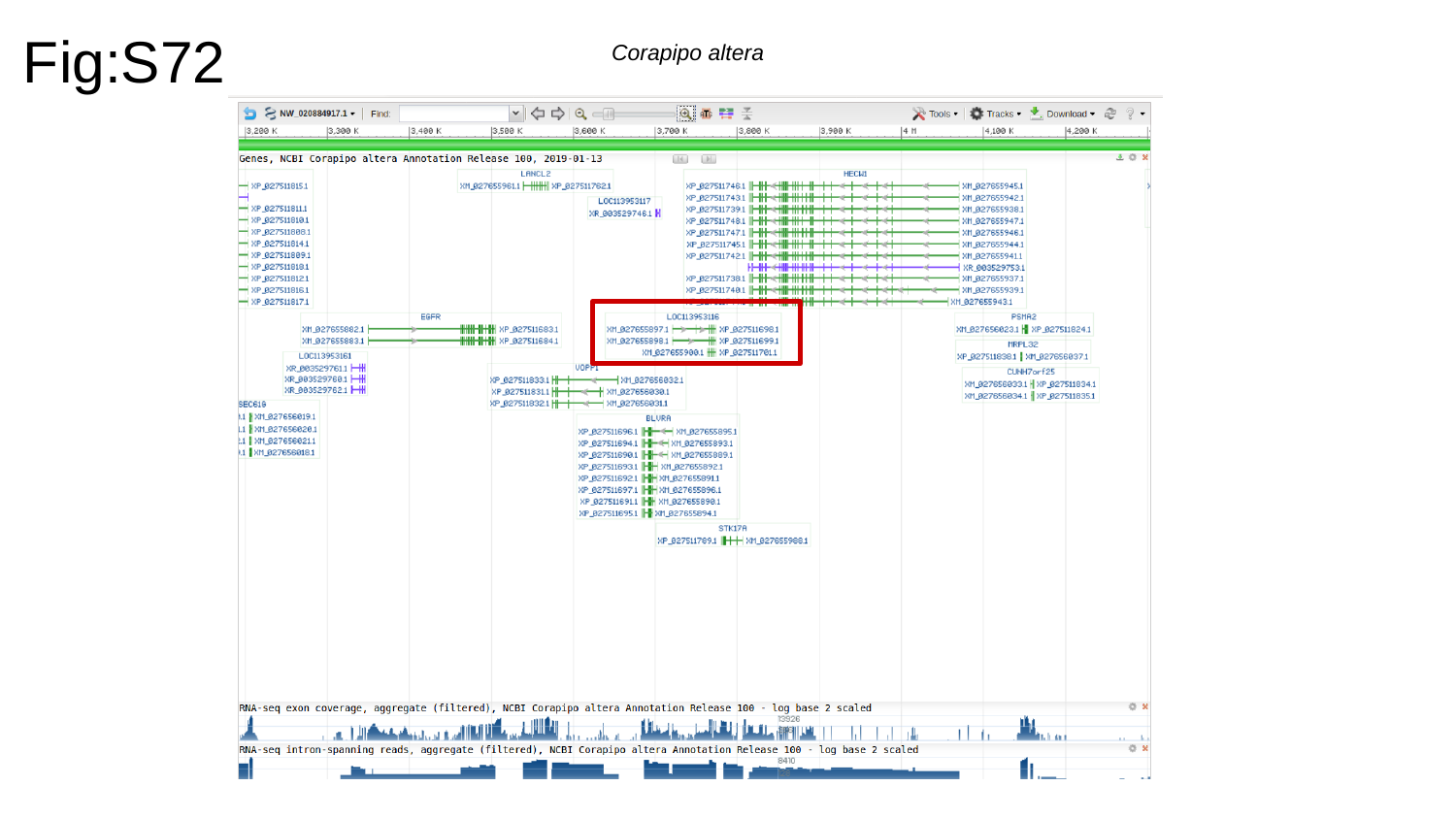

Fig:S72
Corapipo altera

### Slide 73
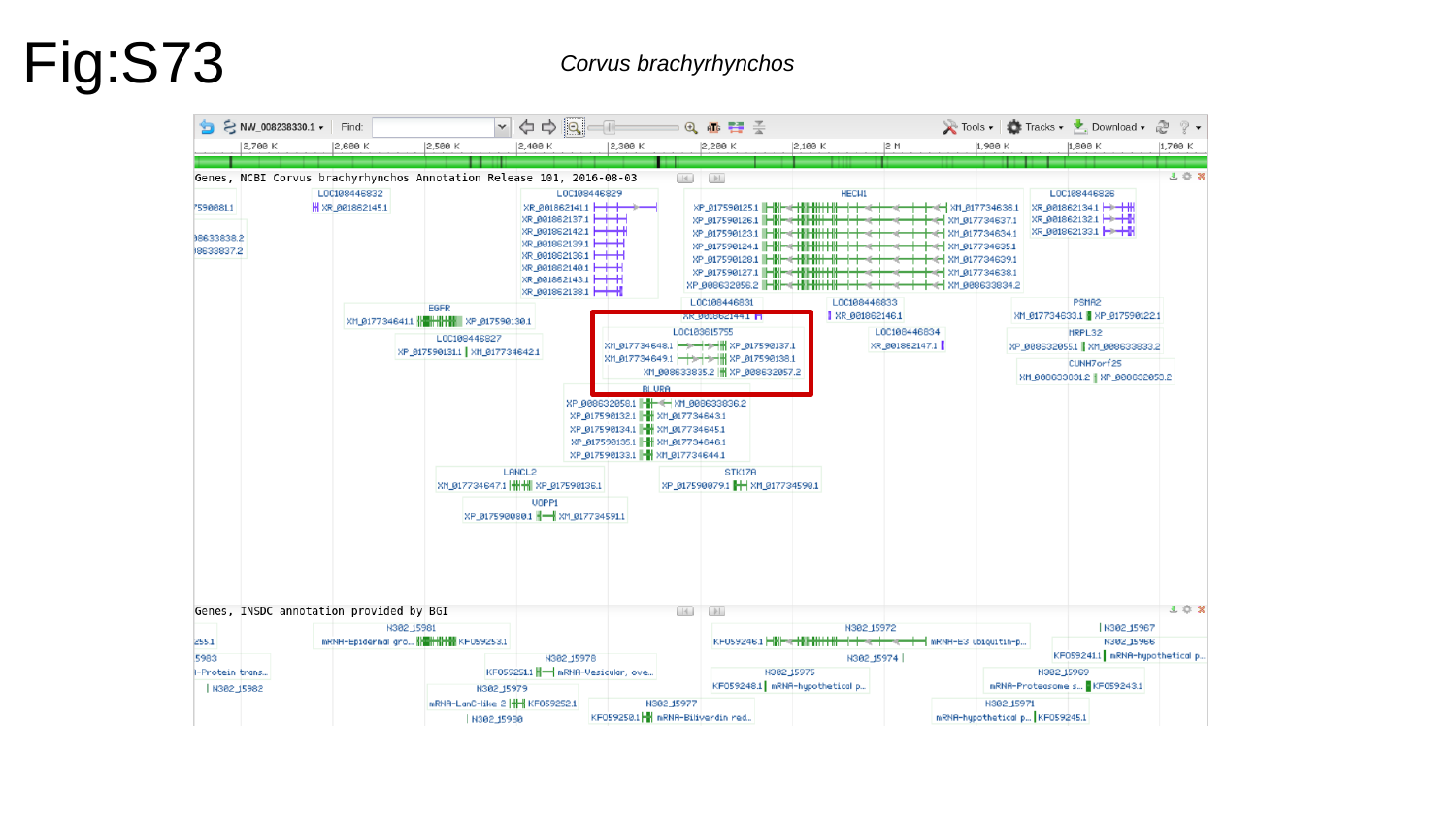

Fig:S73
Corvus brachyrhynchos

### Slide 74
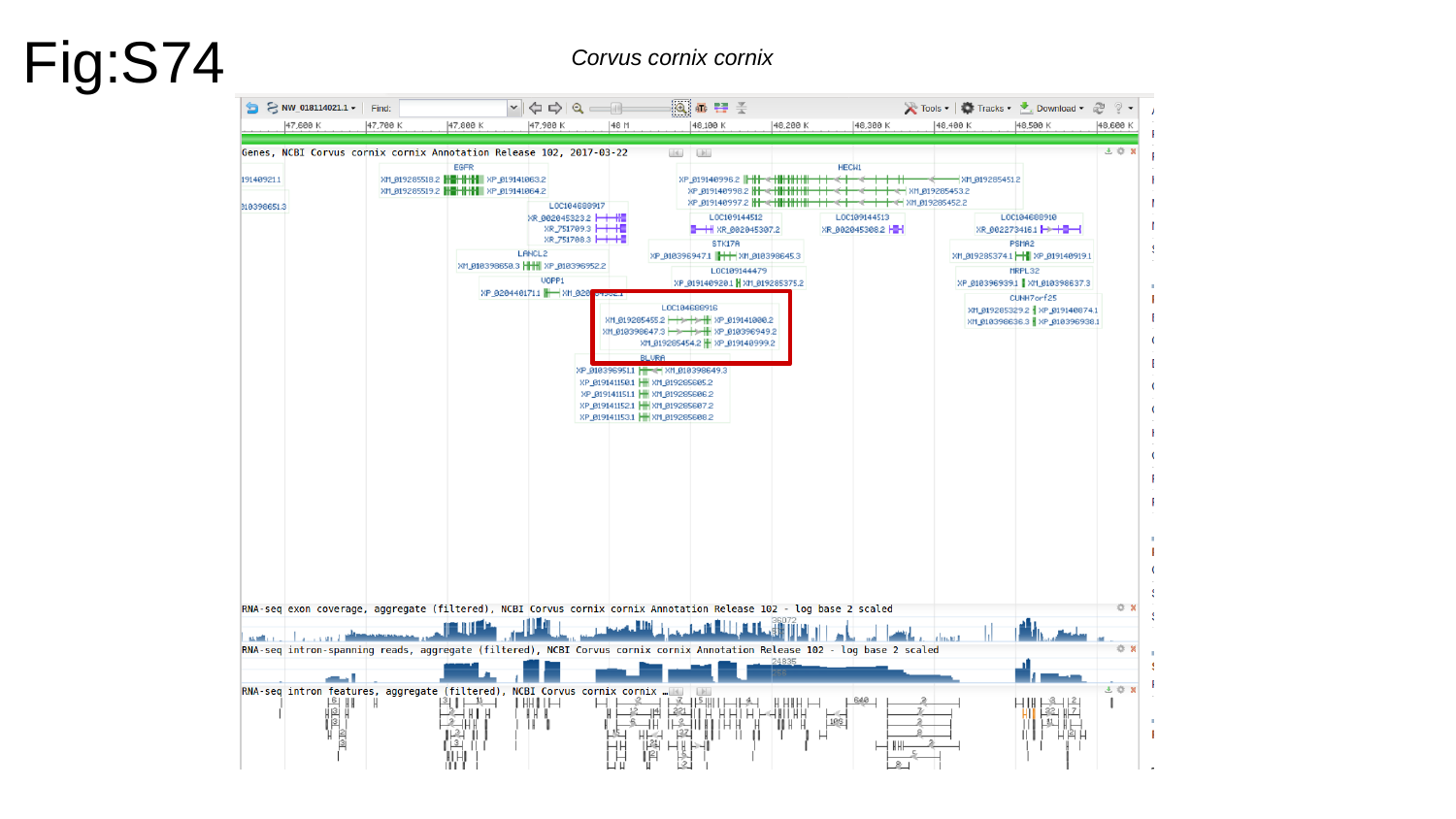

Fig:S74
Corvus cornix cornix

### Slide 75
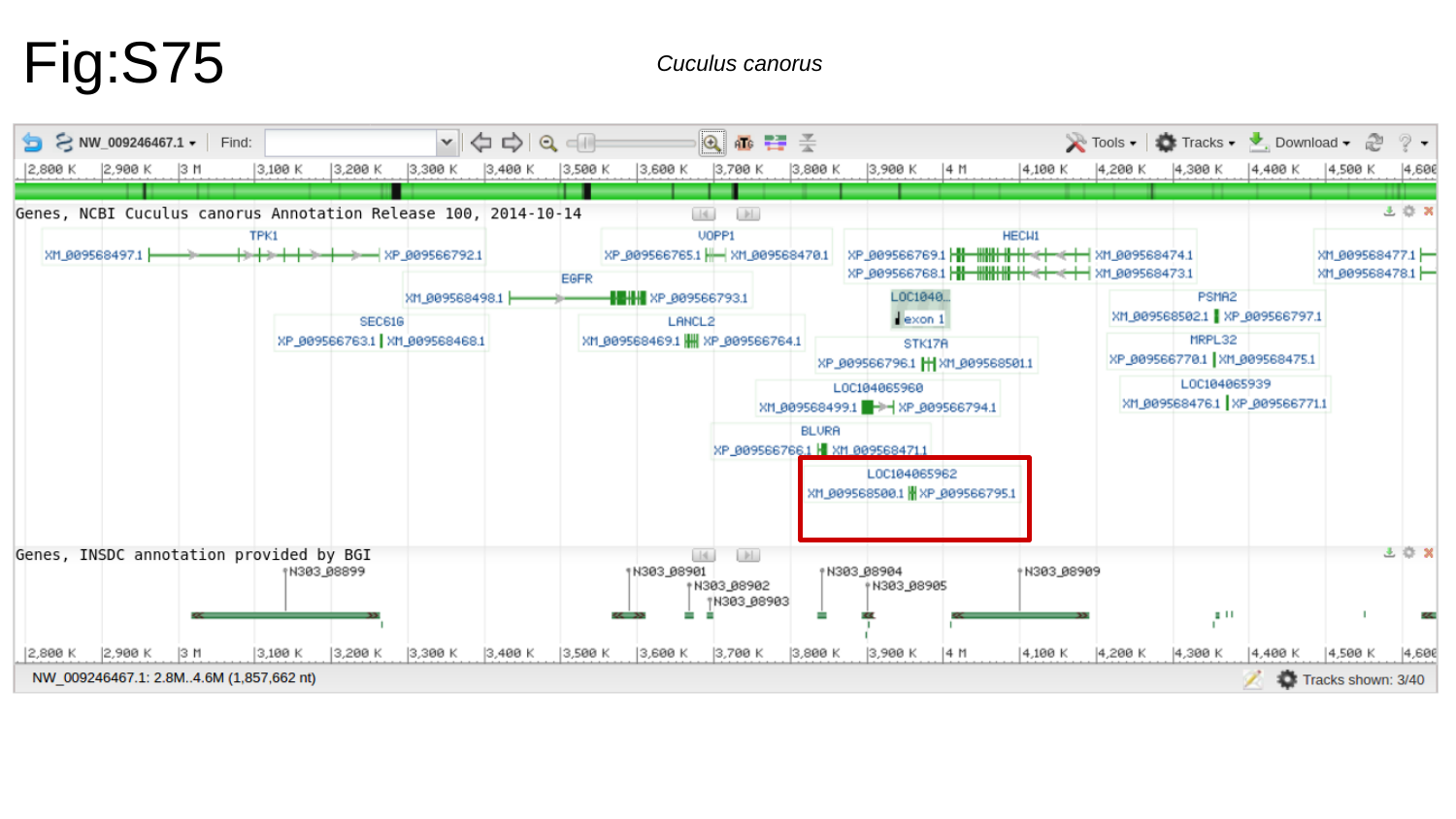

Fig:S75
Cuculus canorus

### Slide 76
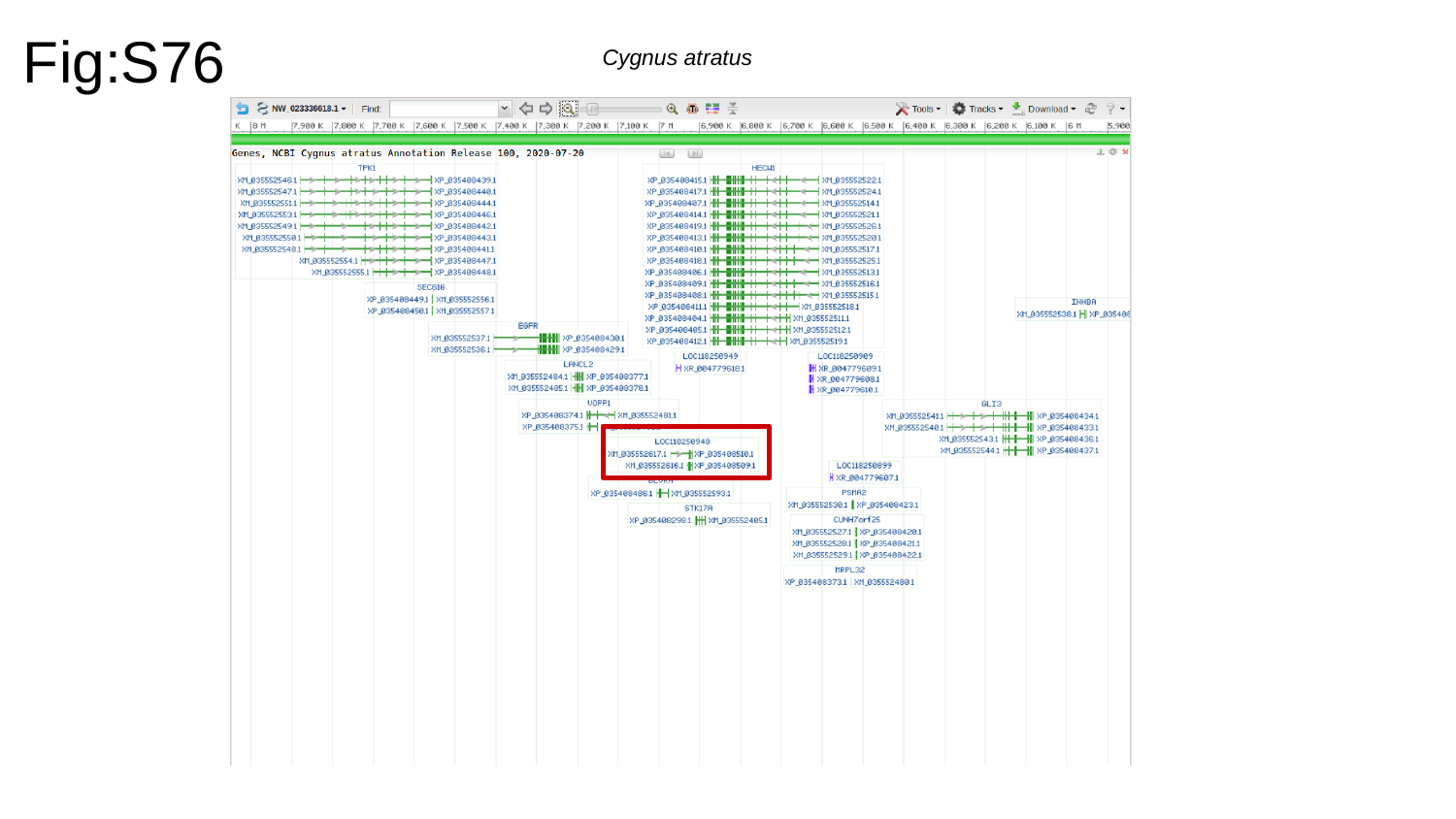

Fig:S76
Cygnus atratus

### Slide 77
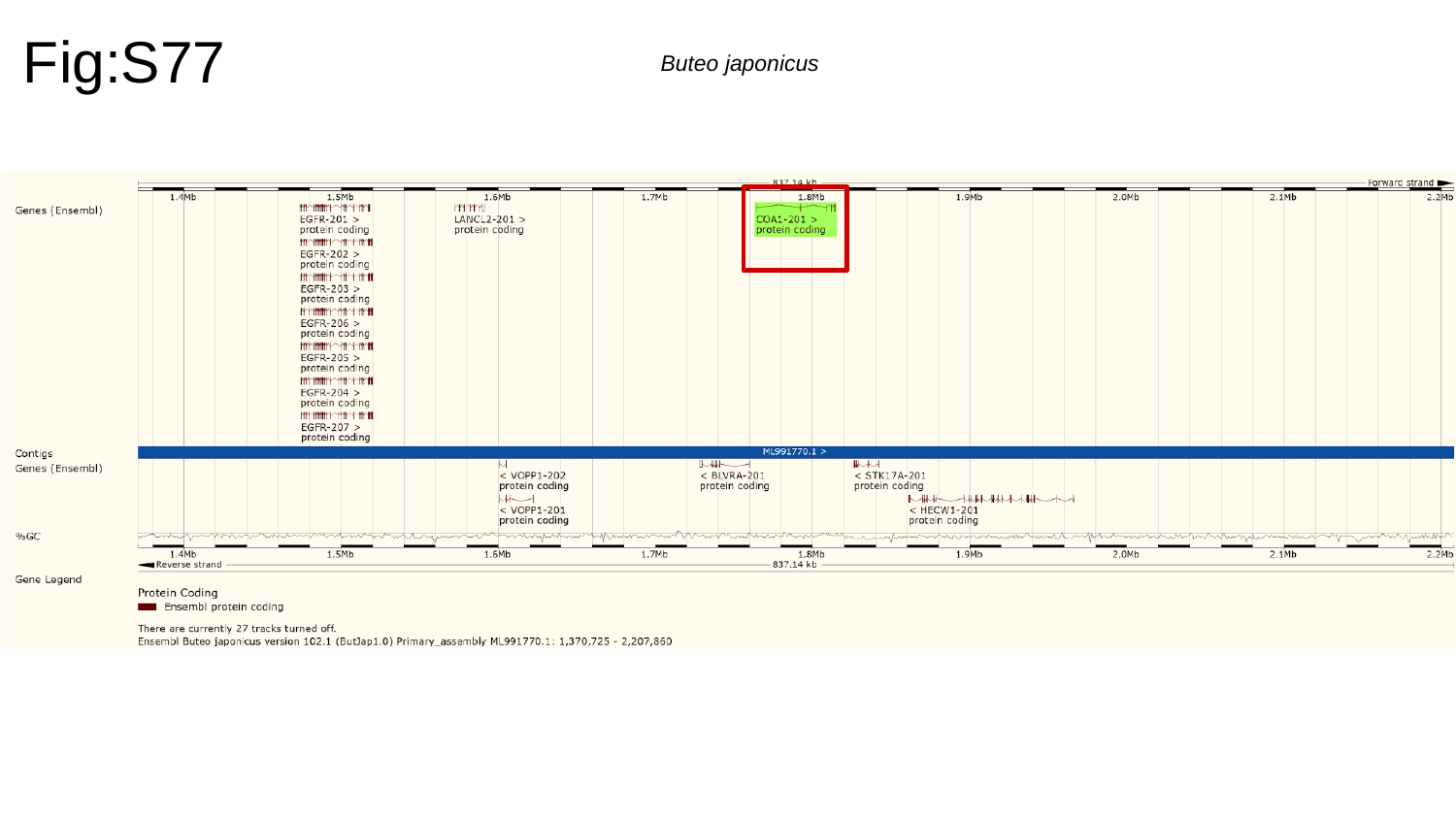

Fig:S77
Buteo japonicus

### Slide 78
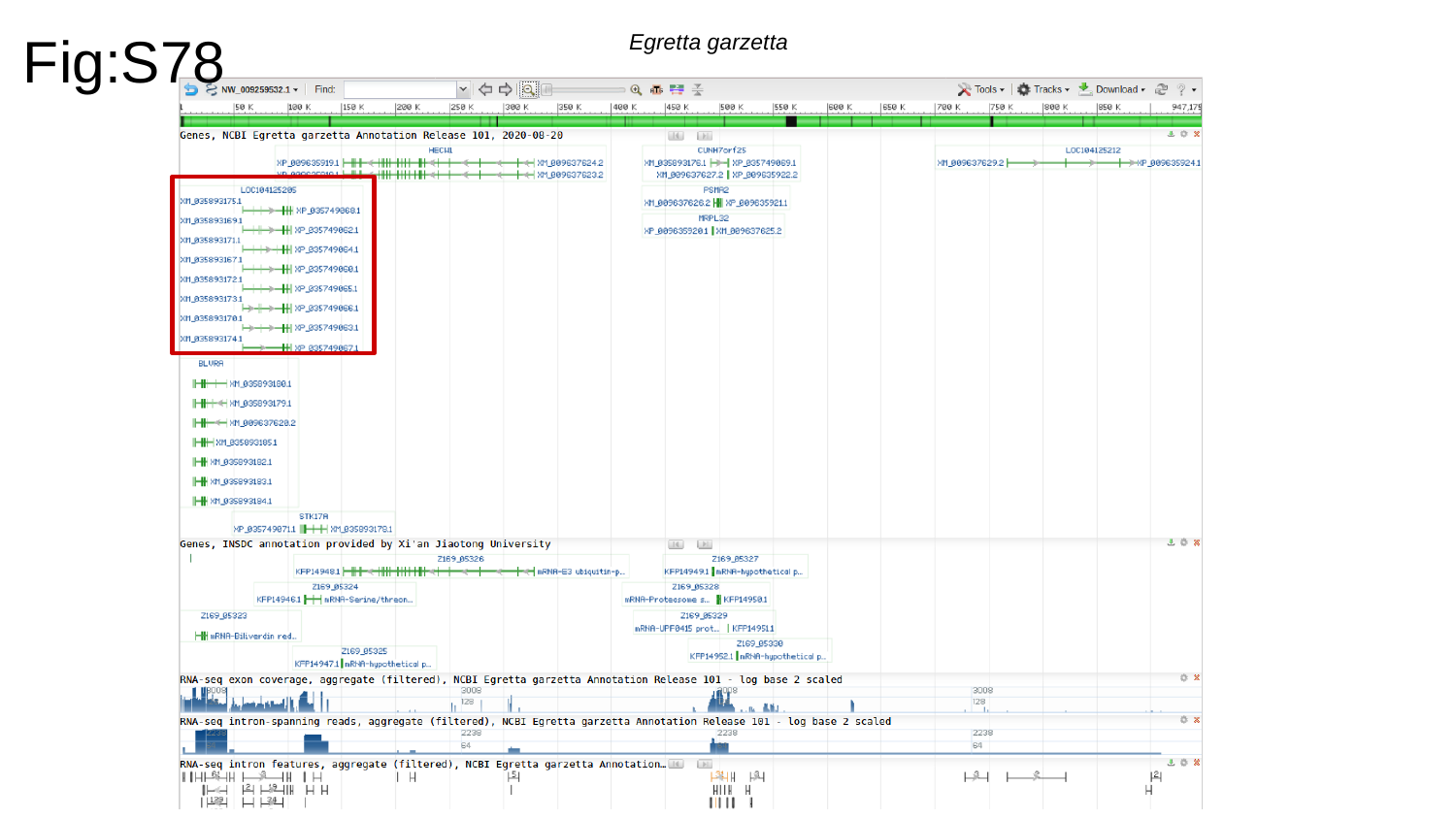

Fig:S78
Egretta garzetta

### Slide 79
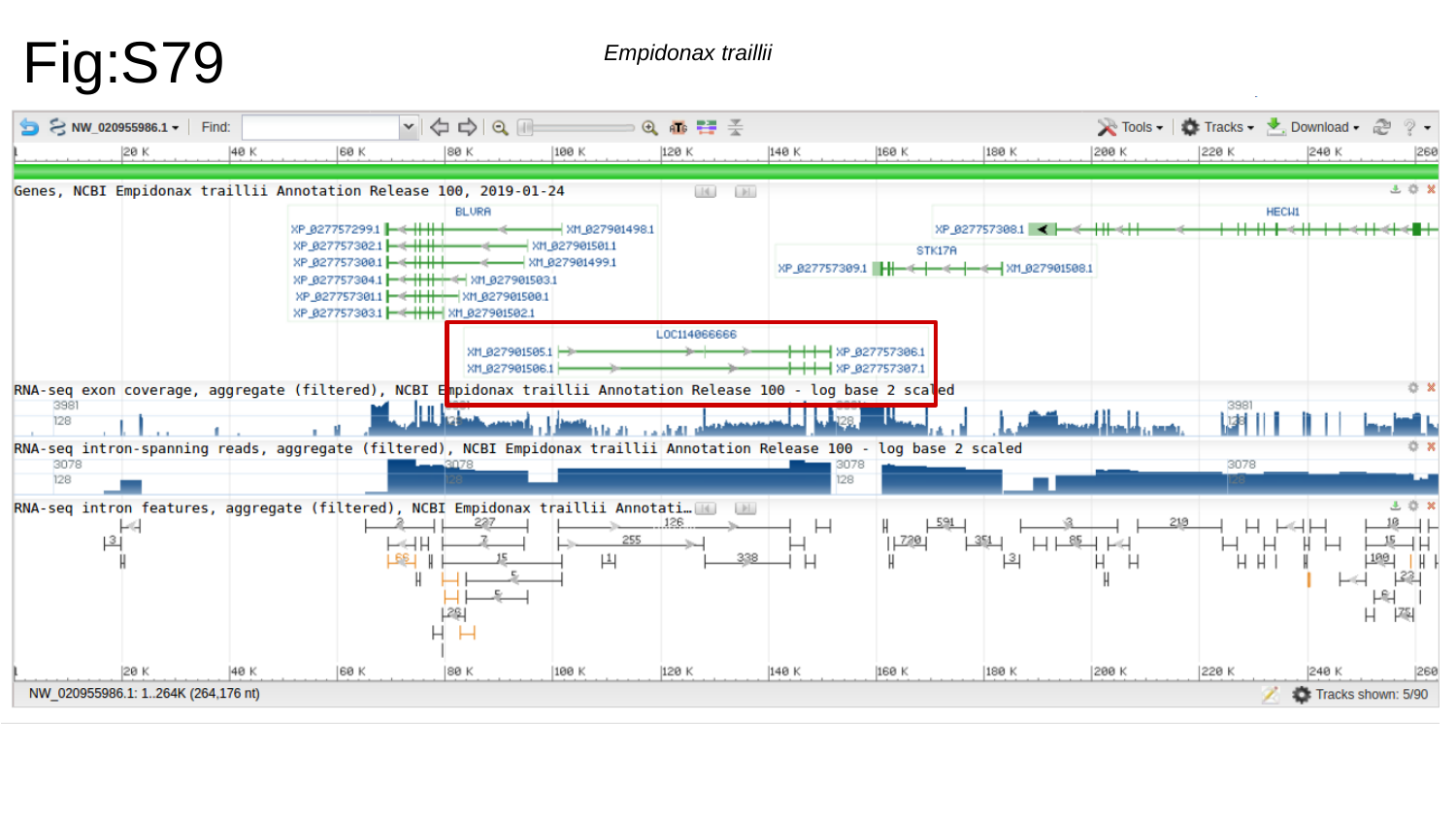

Fig:S79
Empidonax traillii

### Slide 80
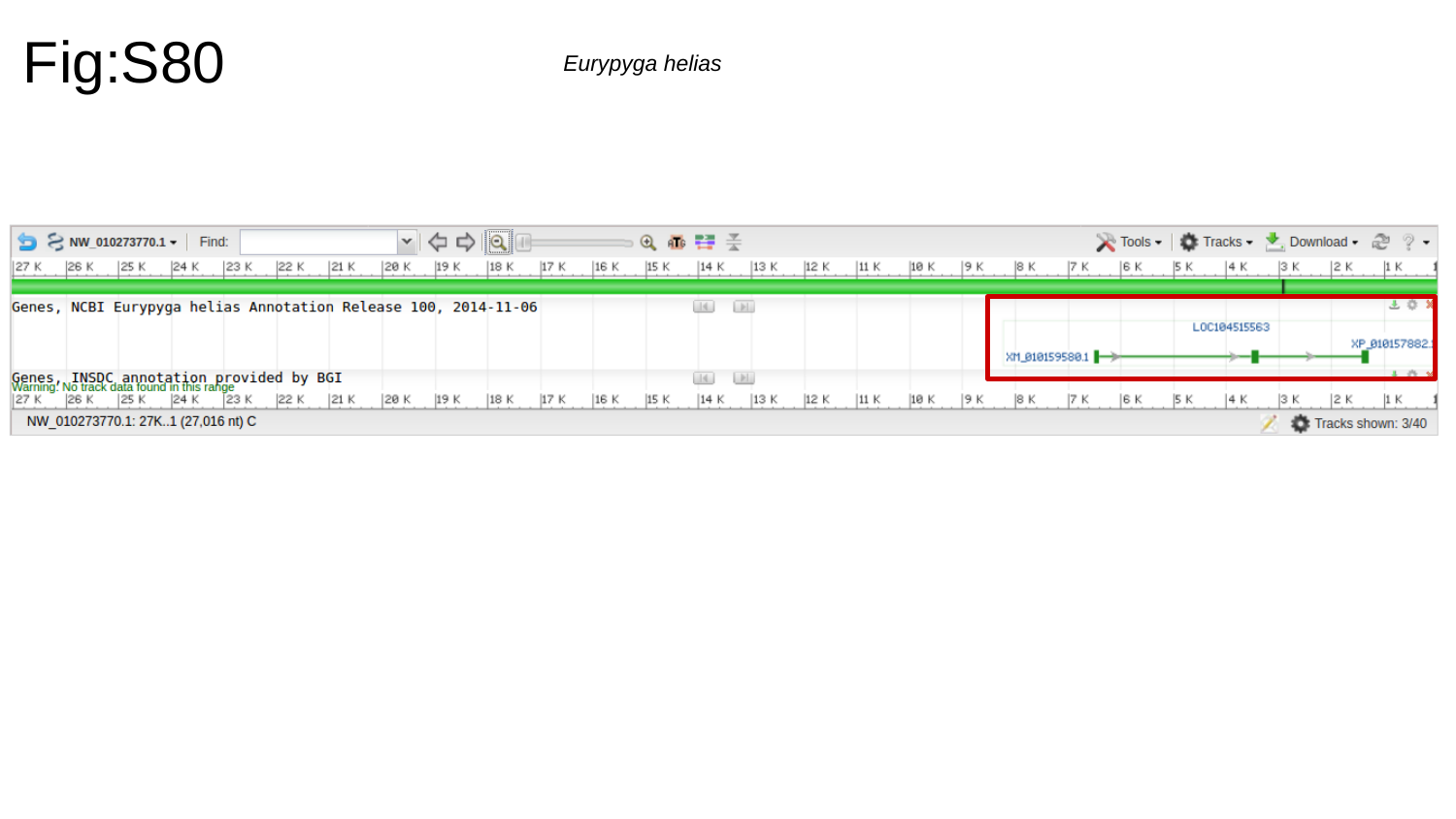

Fig:S80
Eurypyga helias

### Slide 81
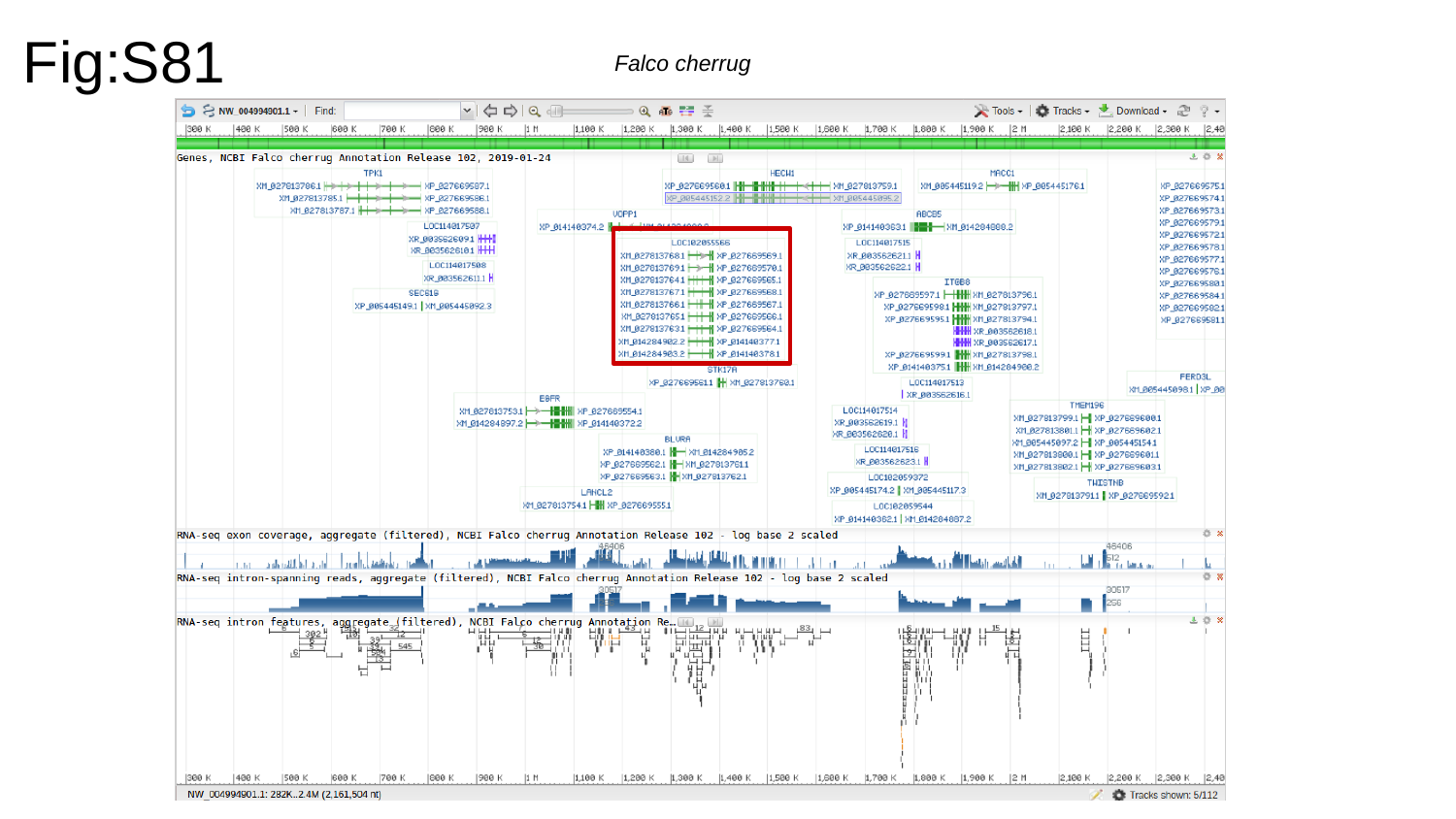

Fig:S81
Falco cherrug

### Slide 82
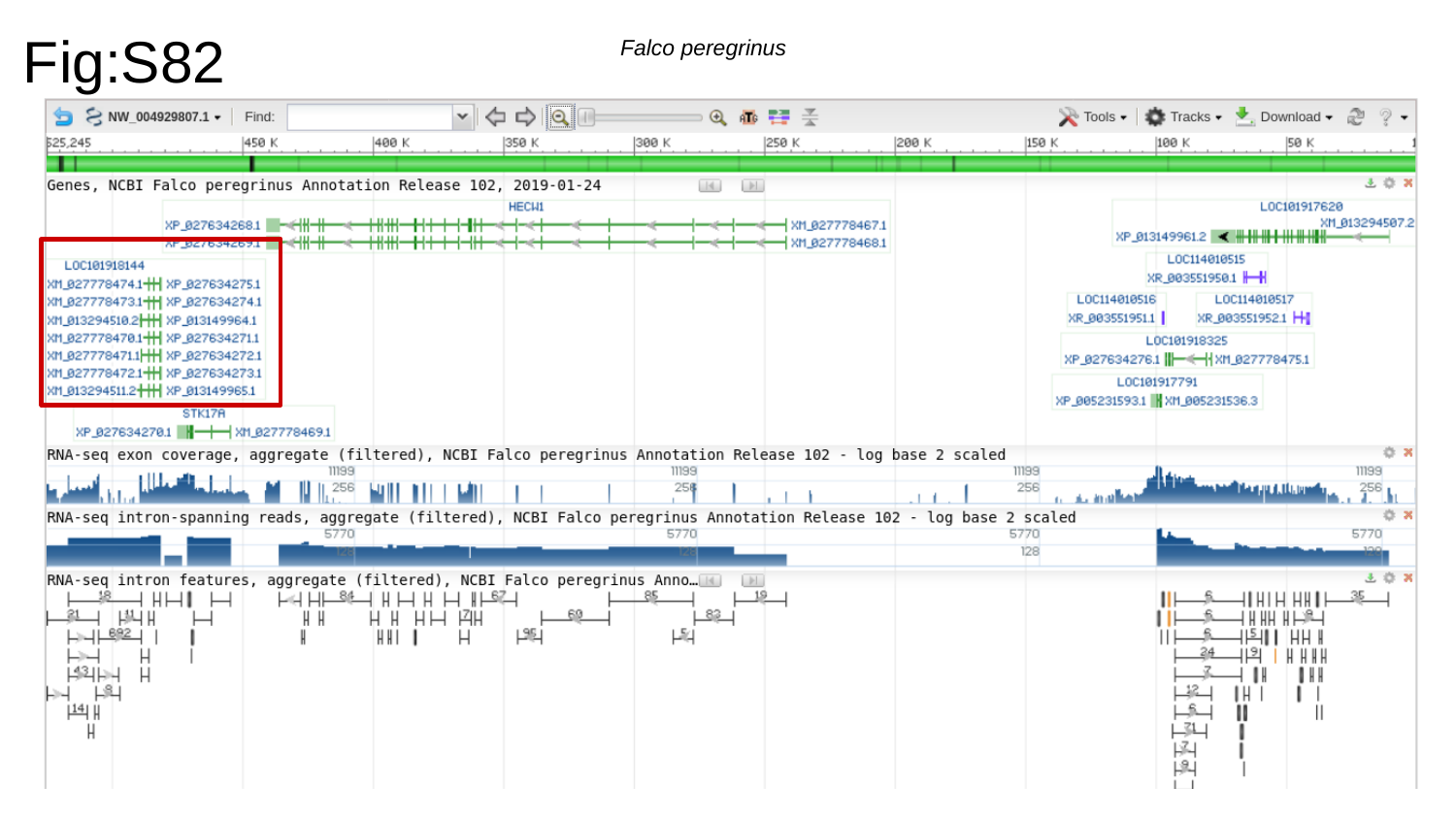

Fig:S82
Falco peregrinus

### Slide 83
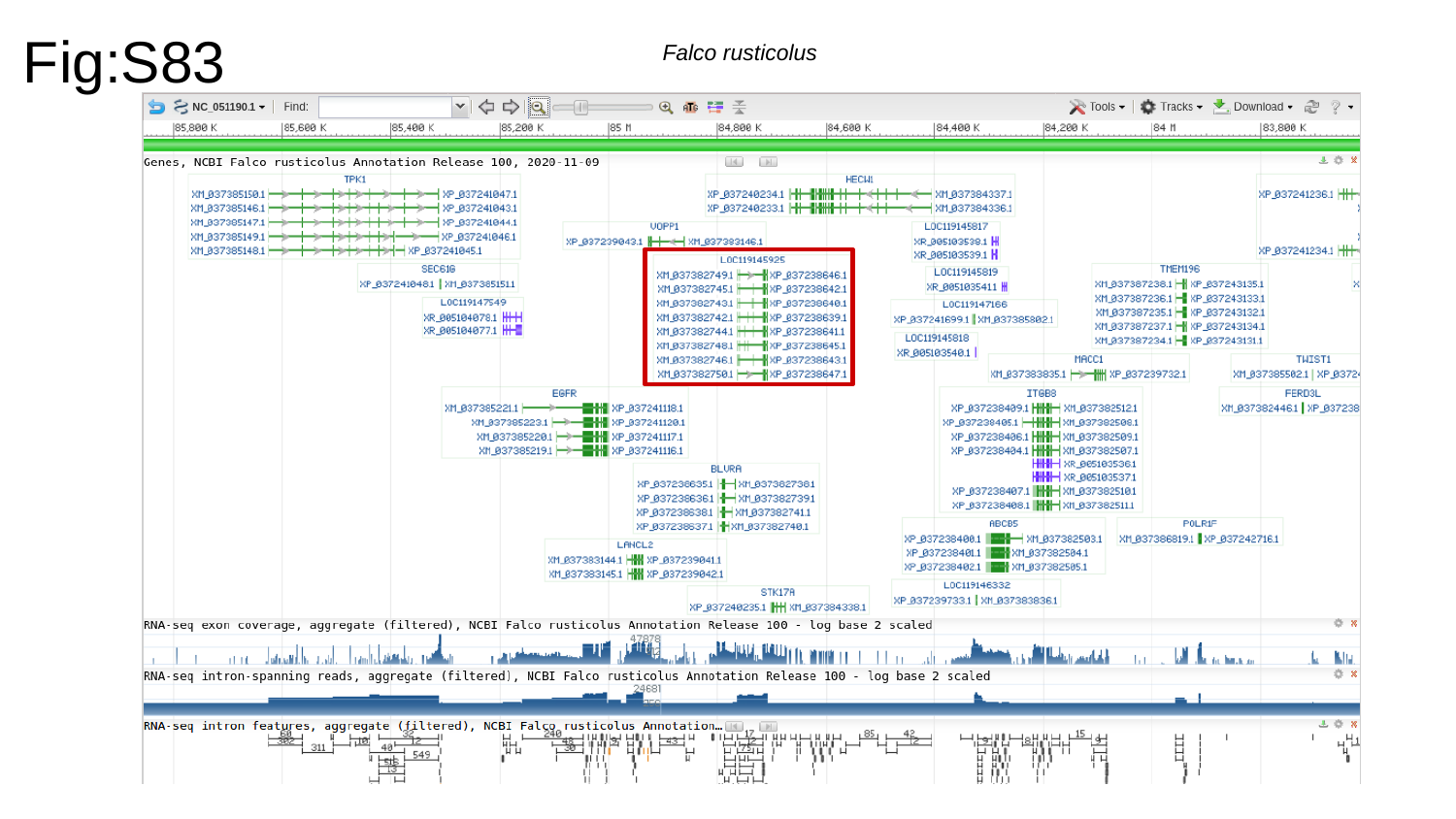

Fig:S83
Falco rusticolus

### Slide 84
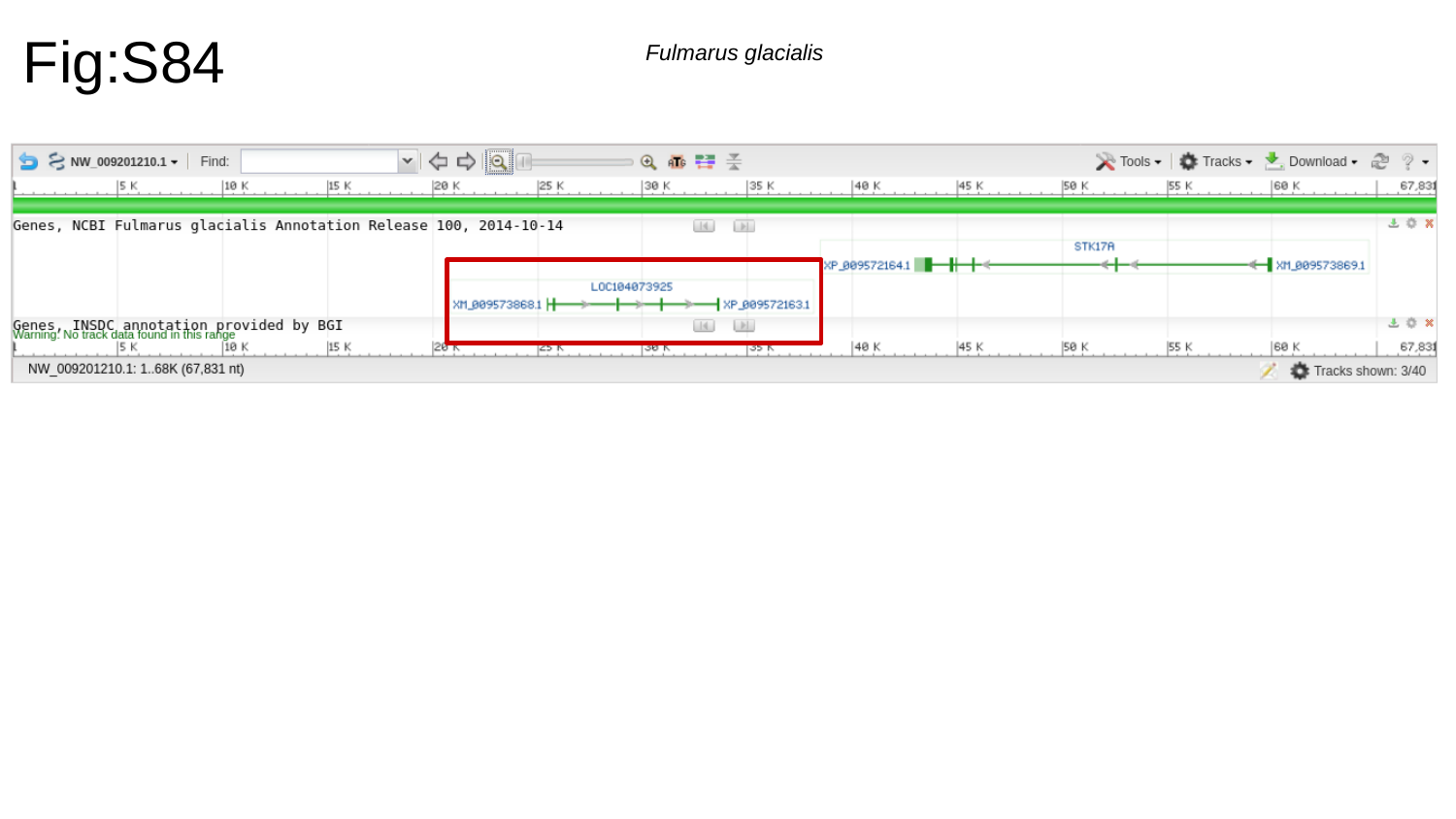

Fig:S84
Fulmarus glacialis

### Slide 85
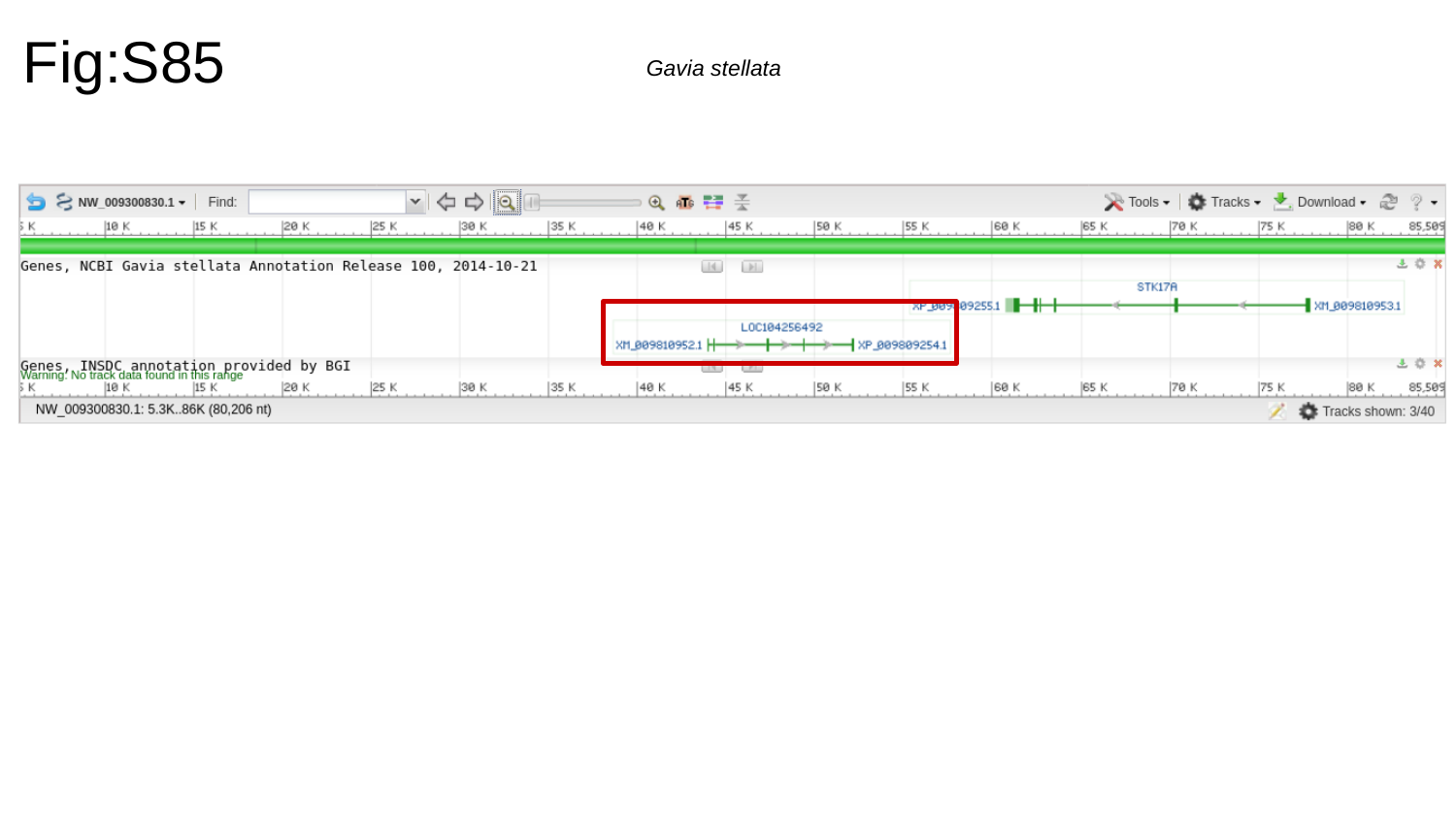

Fig:S85
Gavia stellata

### Slide 86
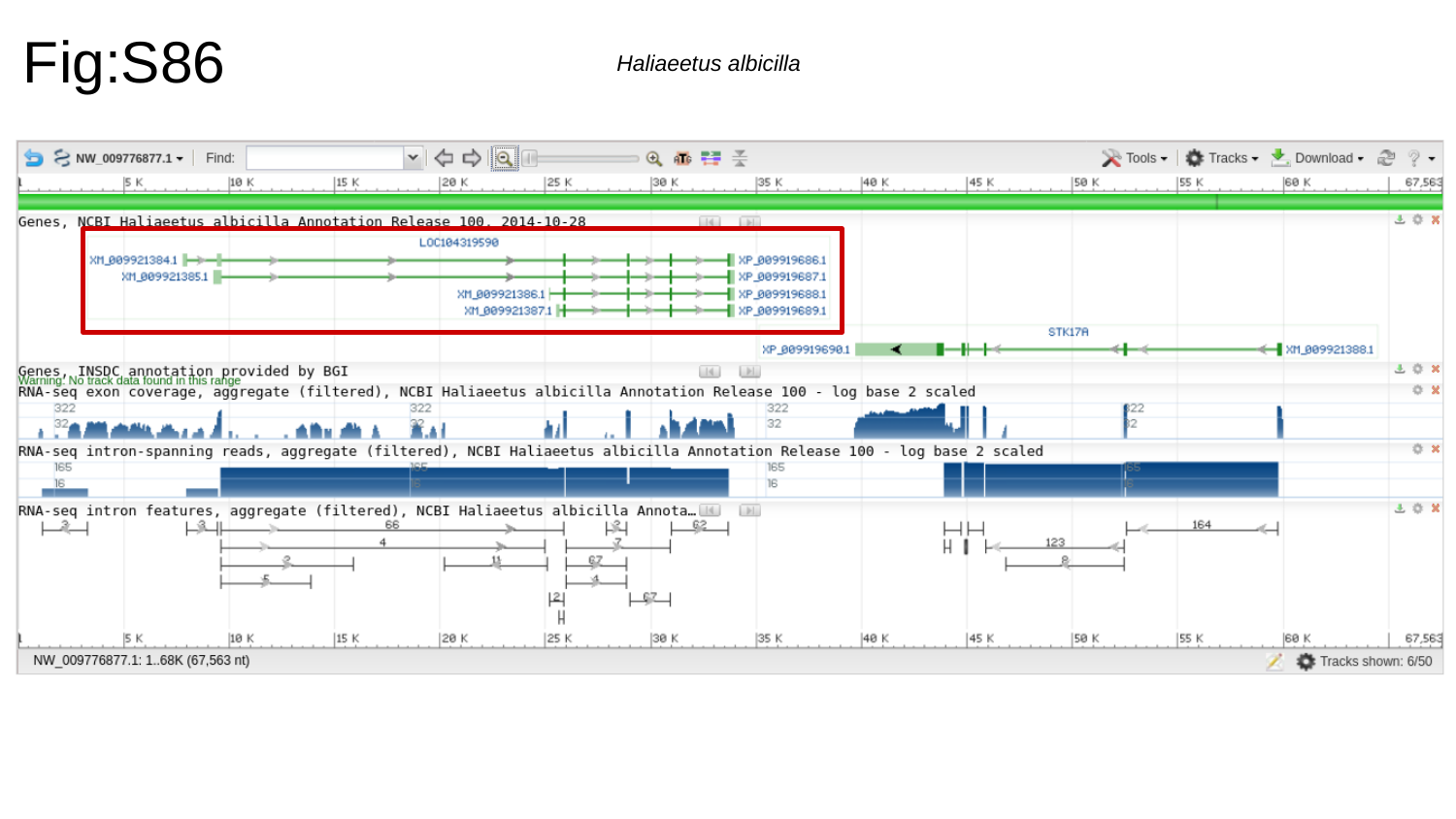

Fig:S86
Haliaeetus albicilla

### Slide 87
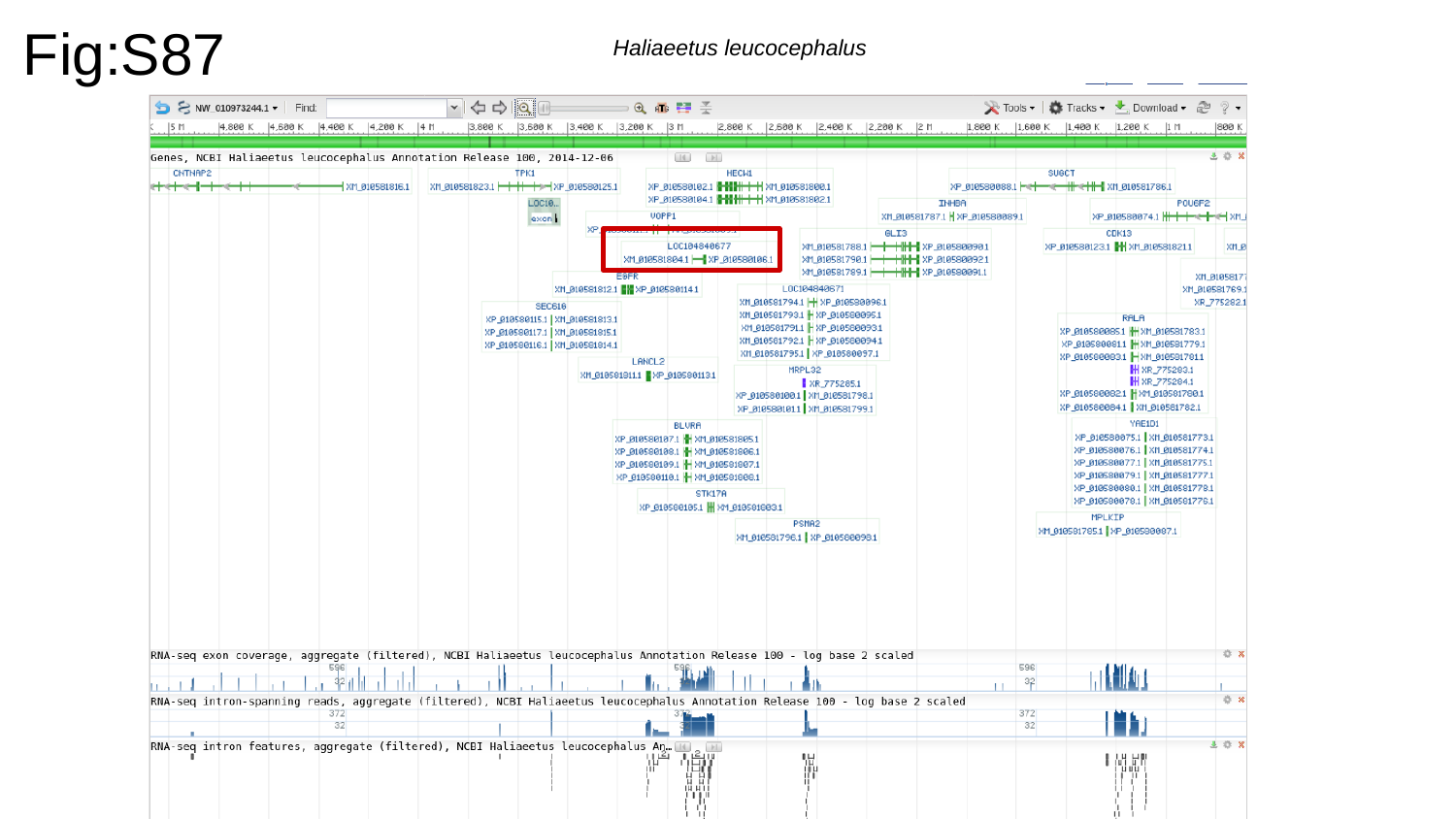

Fig:S87
Haliaeetus leucocephalus

### Slide 88
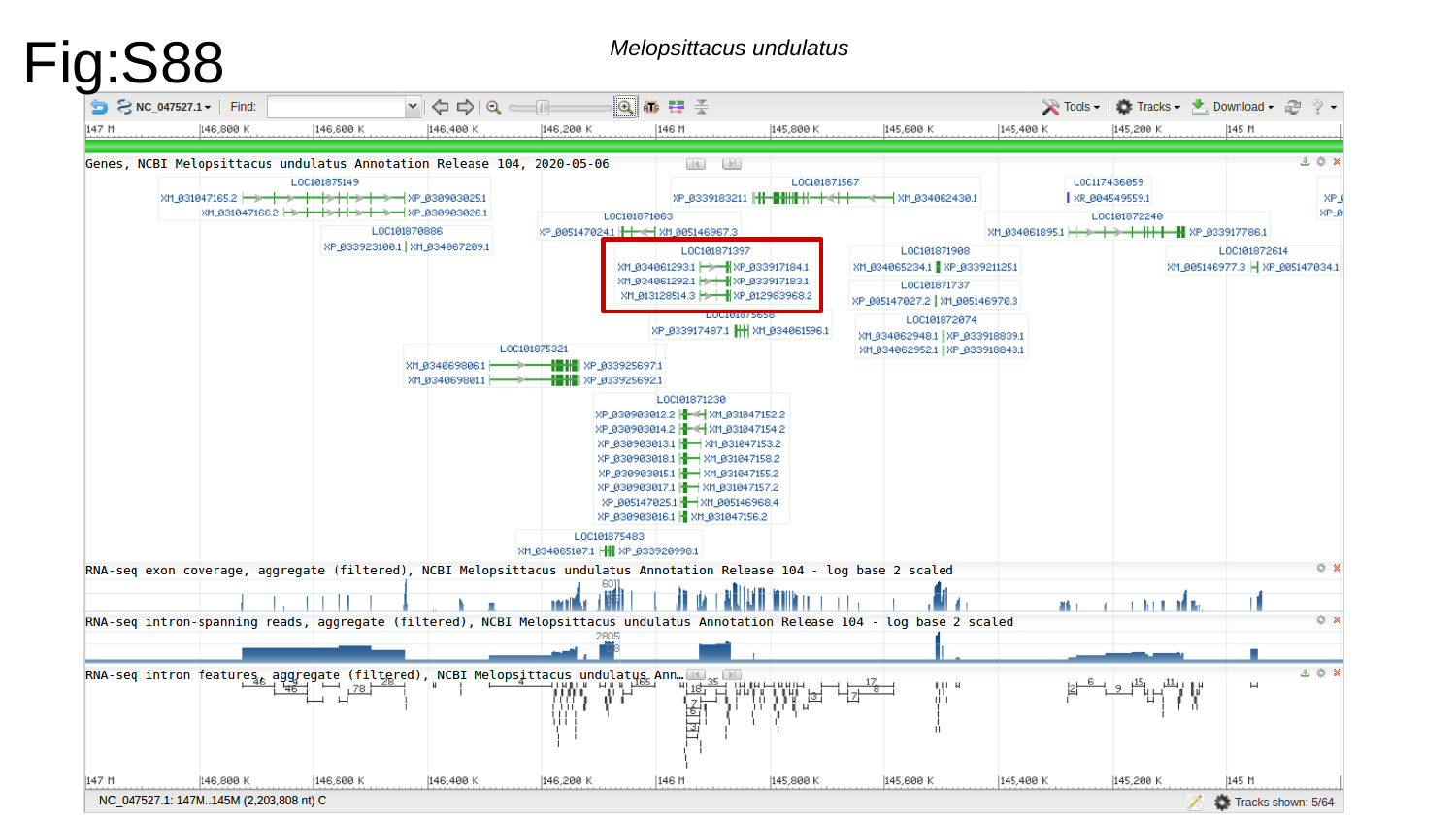

Fig:S88
Melopsittacus undulatus

### Slide 89
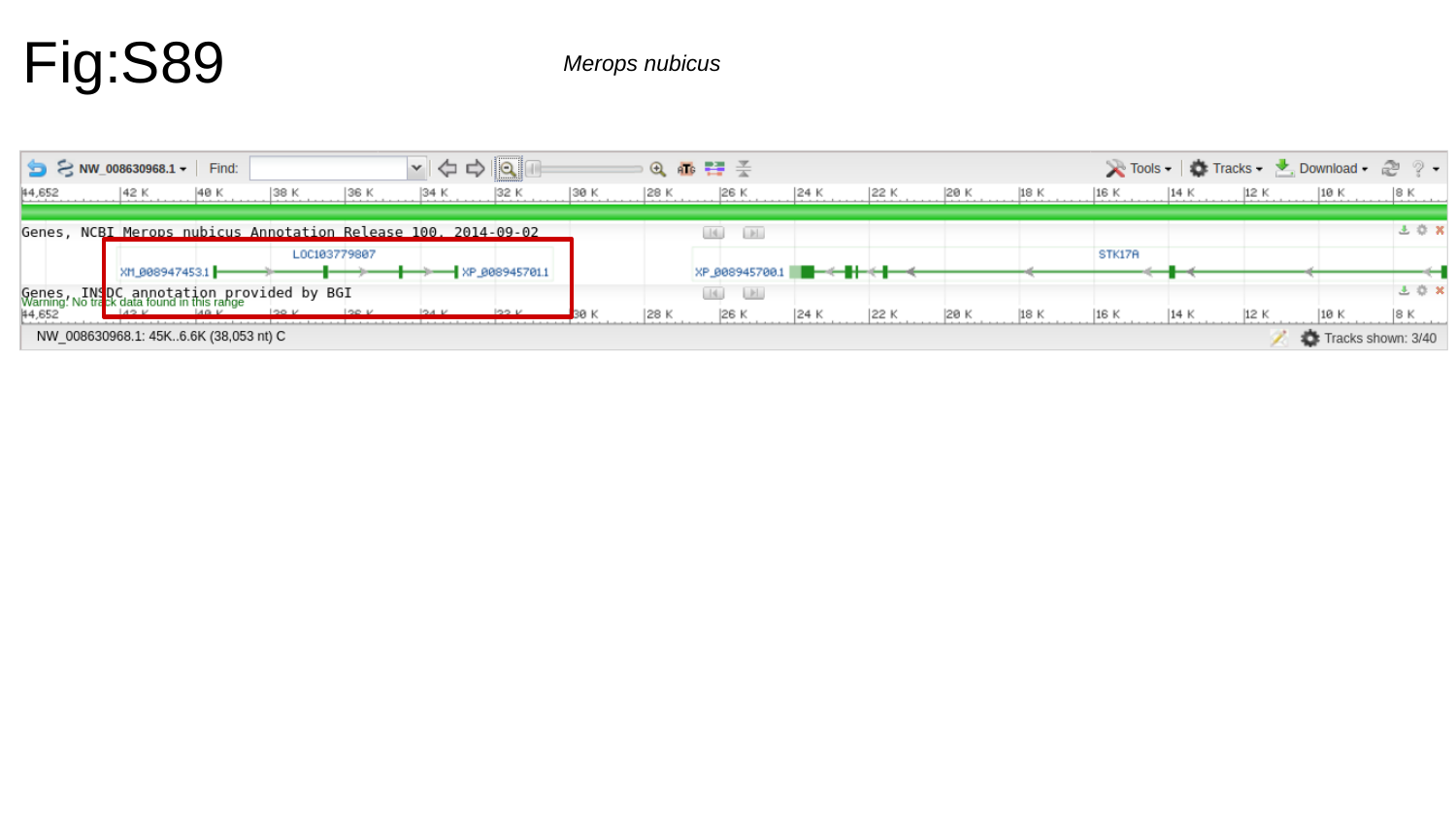

Fig:S89
Merops nubicus

### Slide 90
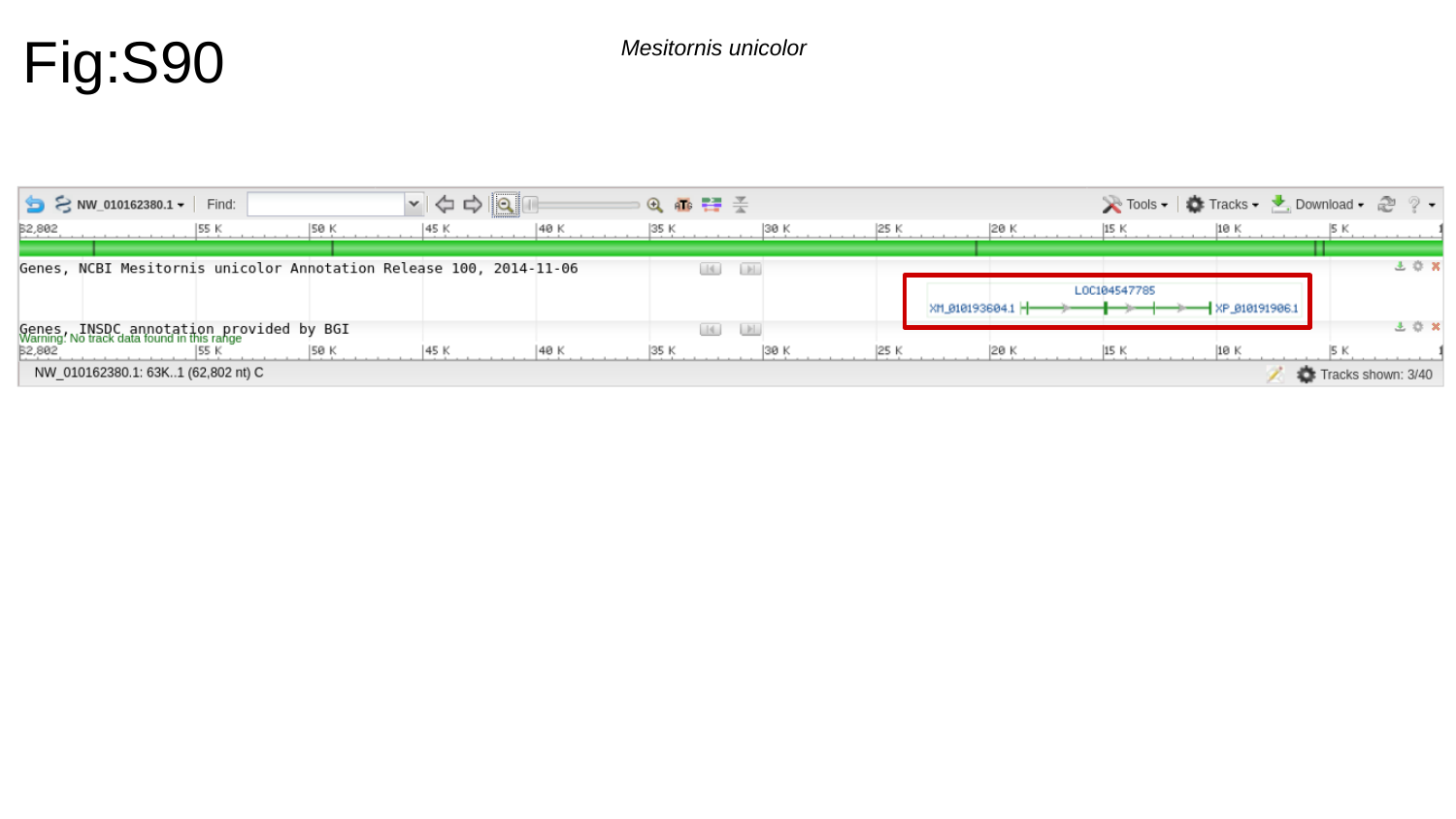

Fig:S90
Mesitornis unicolor

### Slide 91
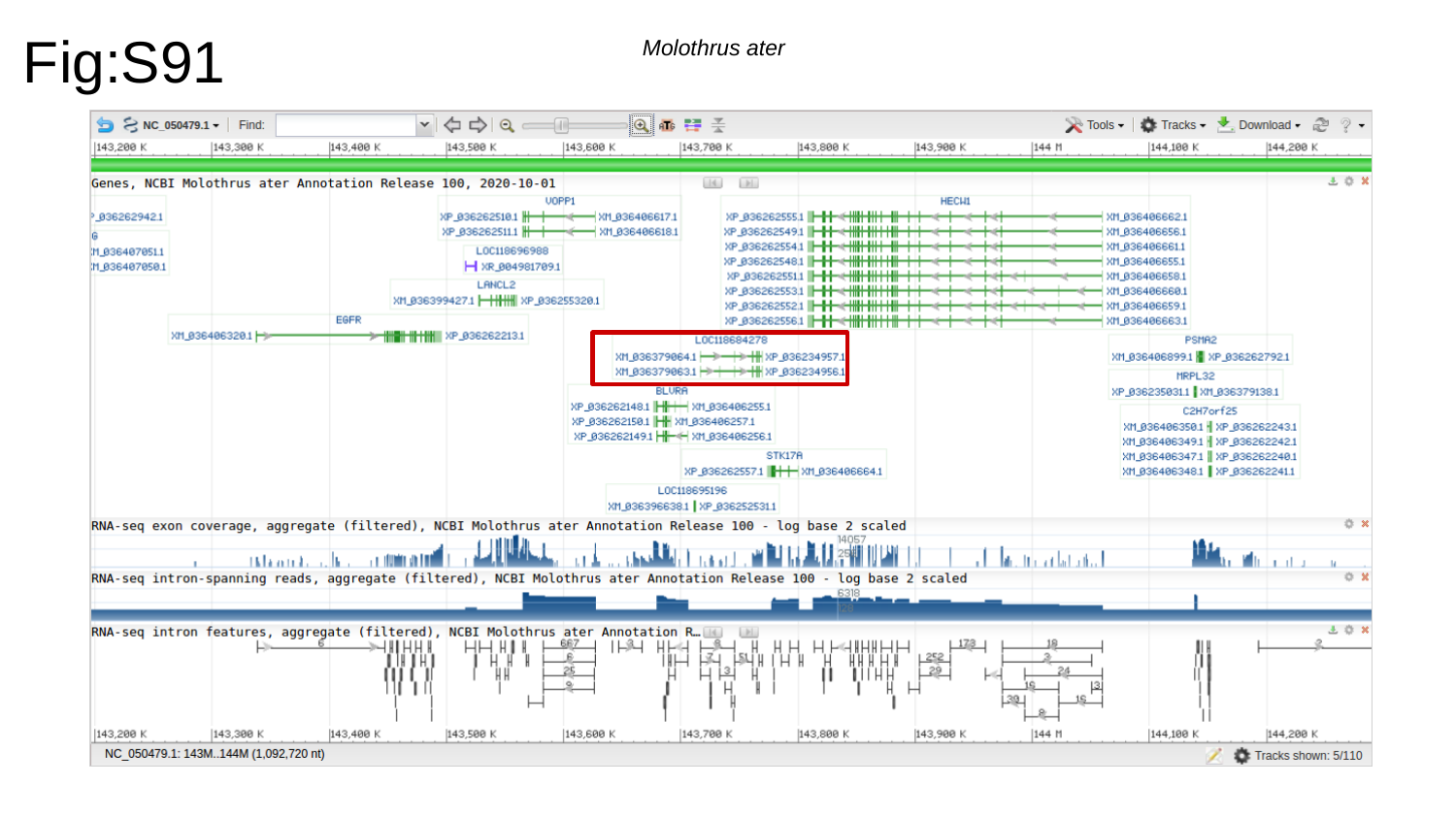

Fig:S91
Molothrus ater

### Slide 92
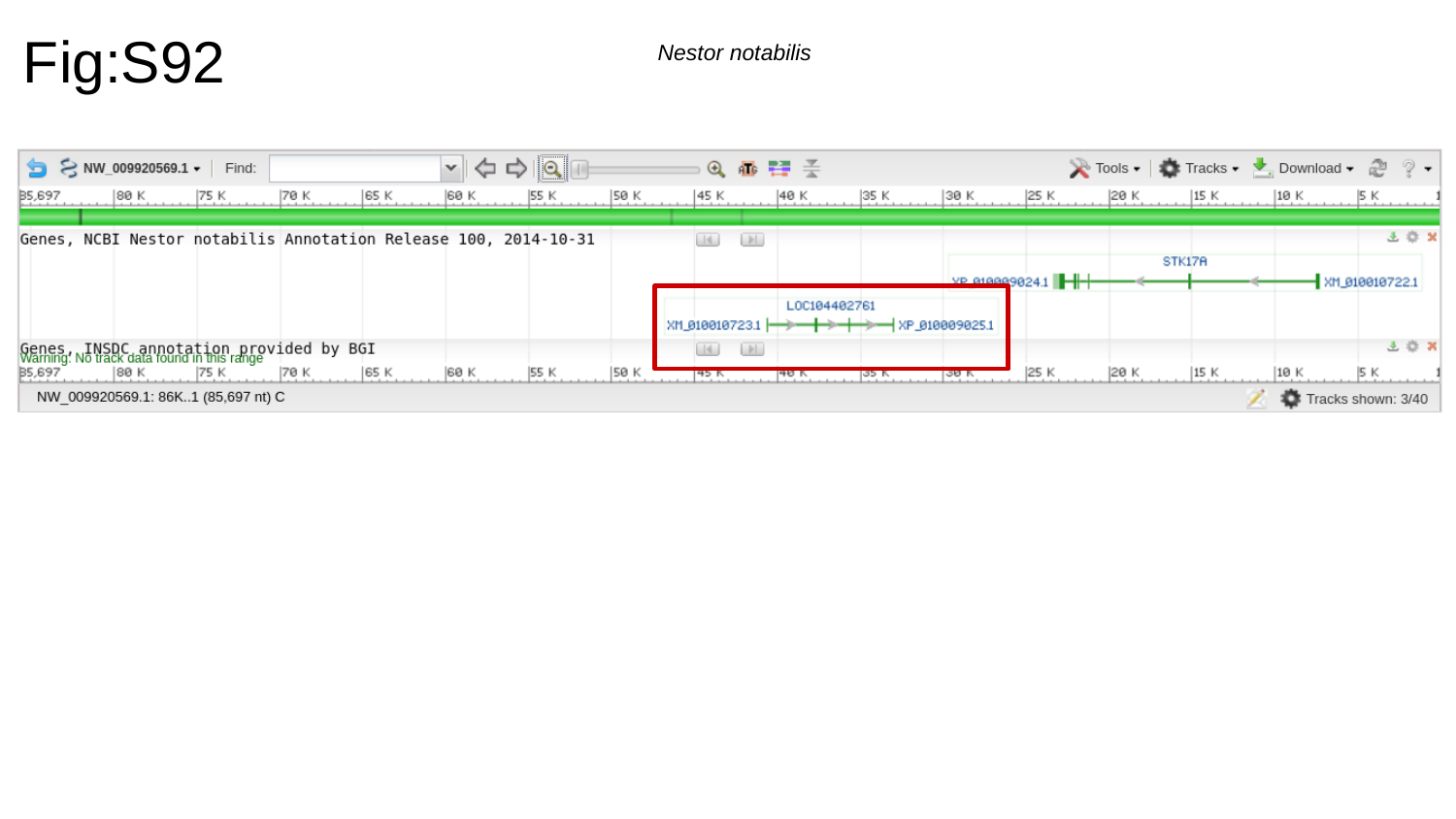

Fig:S92
Nestor notabilis

### Slide 93
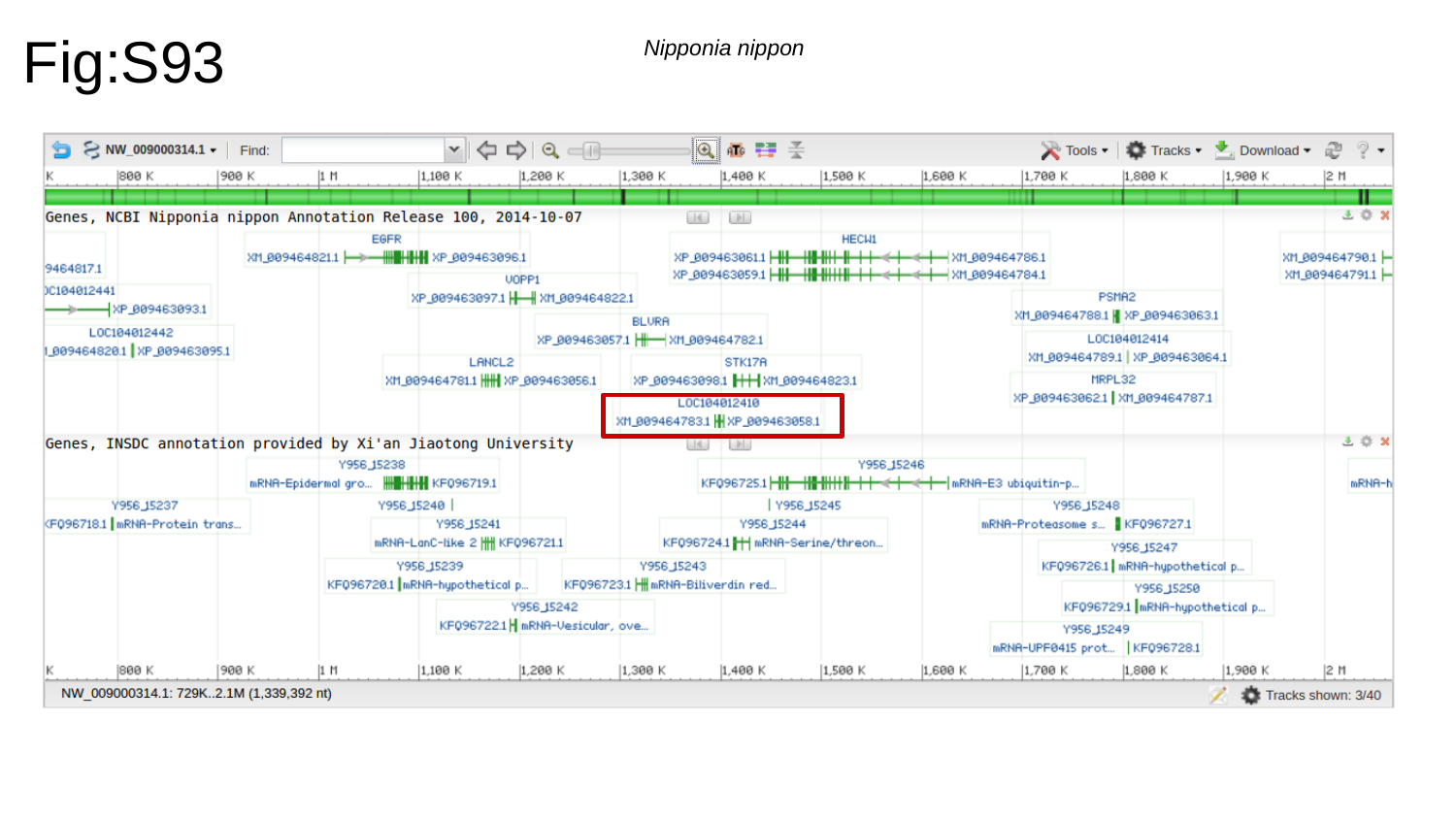

Fig:S93
Nipponia nippon

### Slide 94
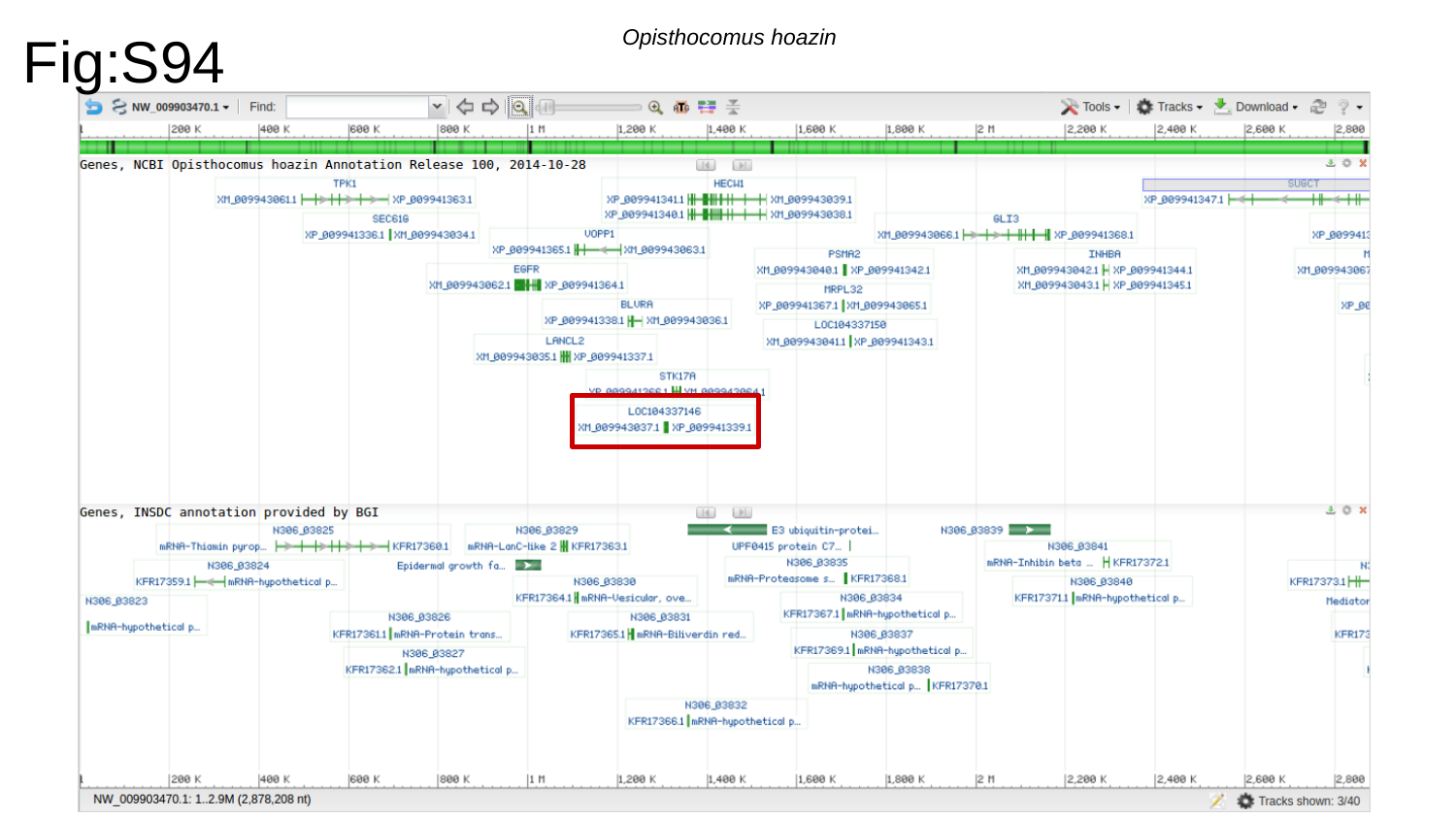

Fig:S94
Opisthocomus hoazin

### Slide 95
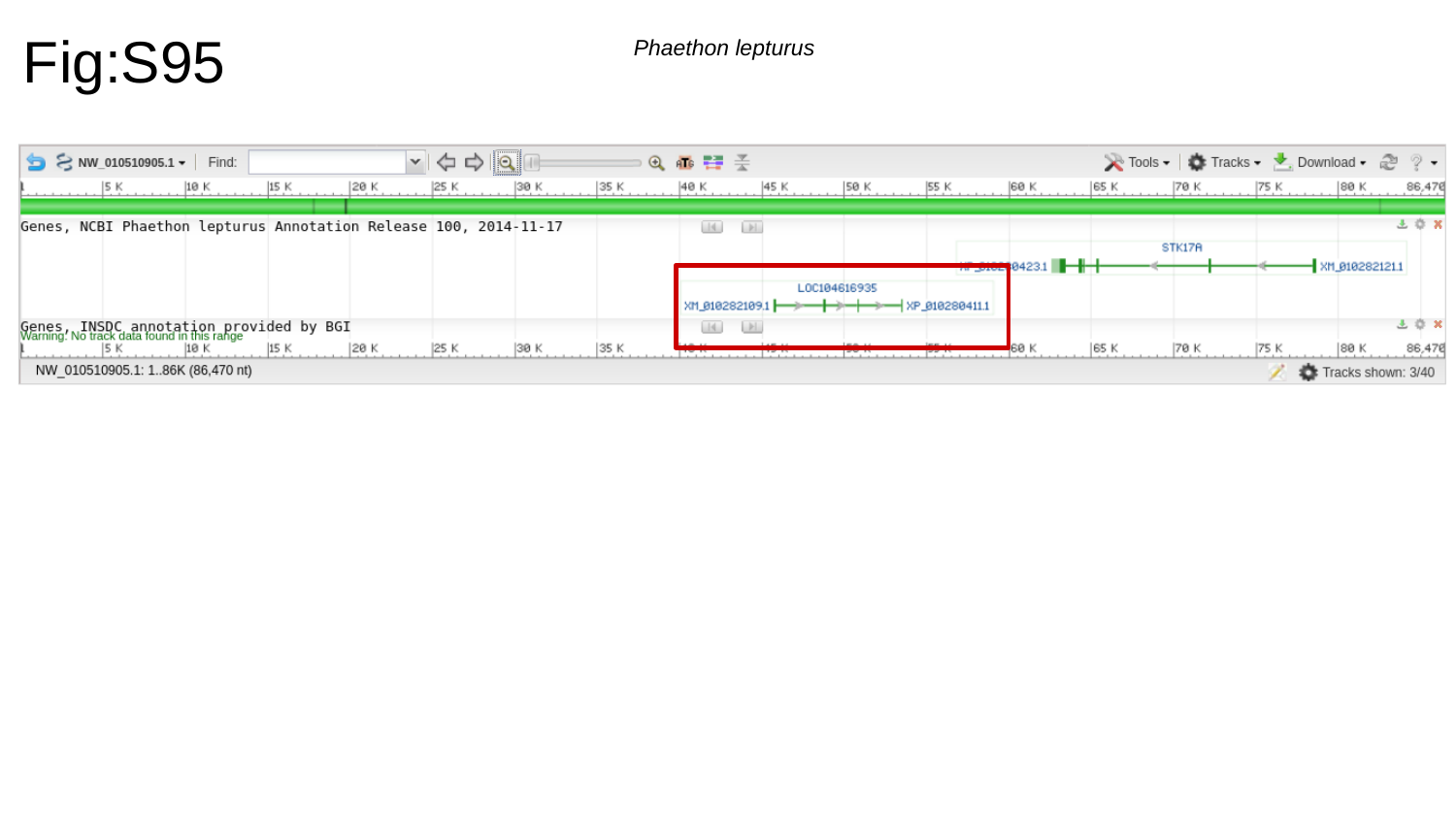

Fig:S95
Phaethon lepturus

### Slide 96
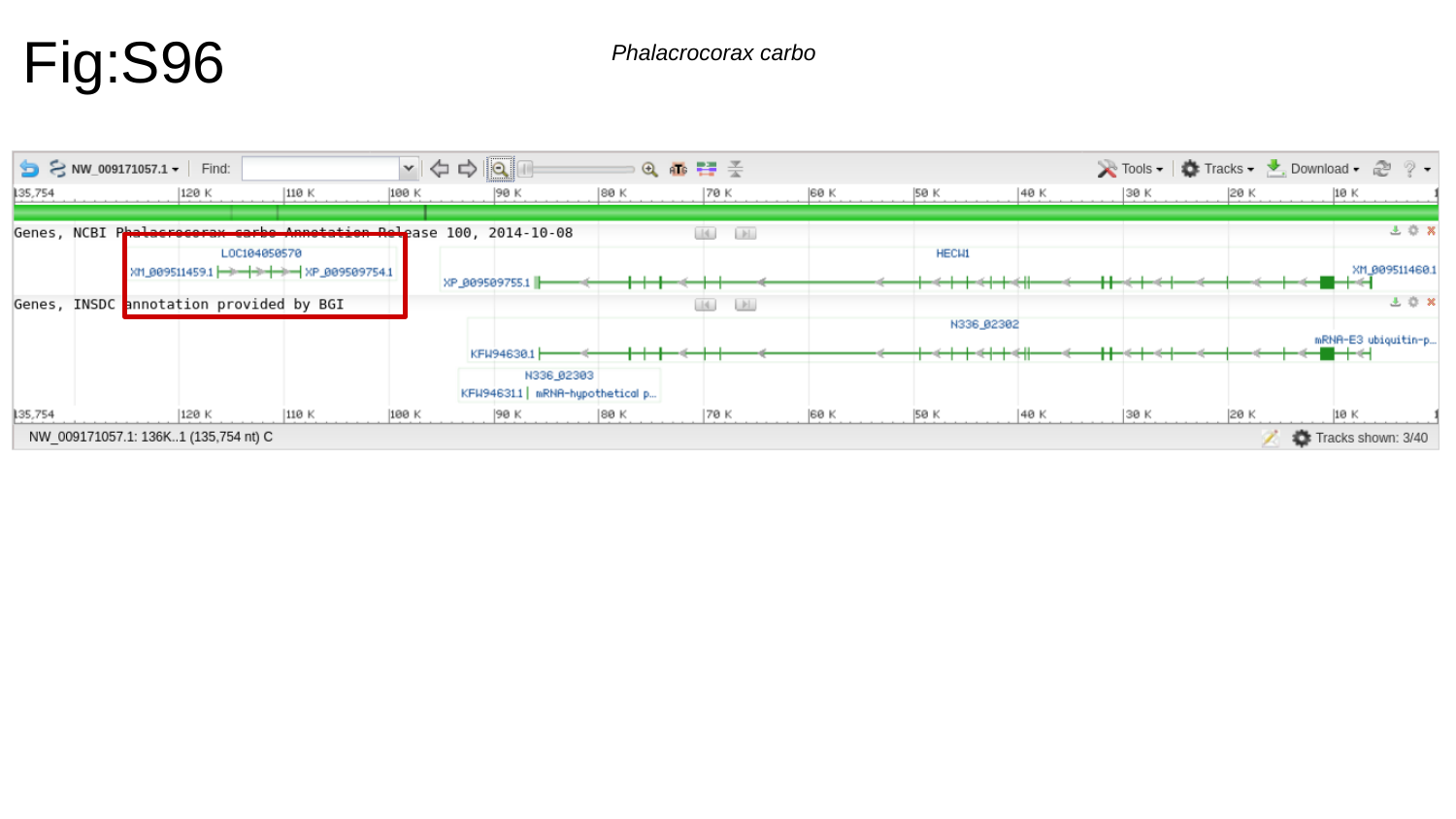

Fig:S96
Phalacrocorax carbo

### Slide 97
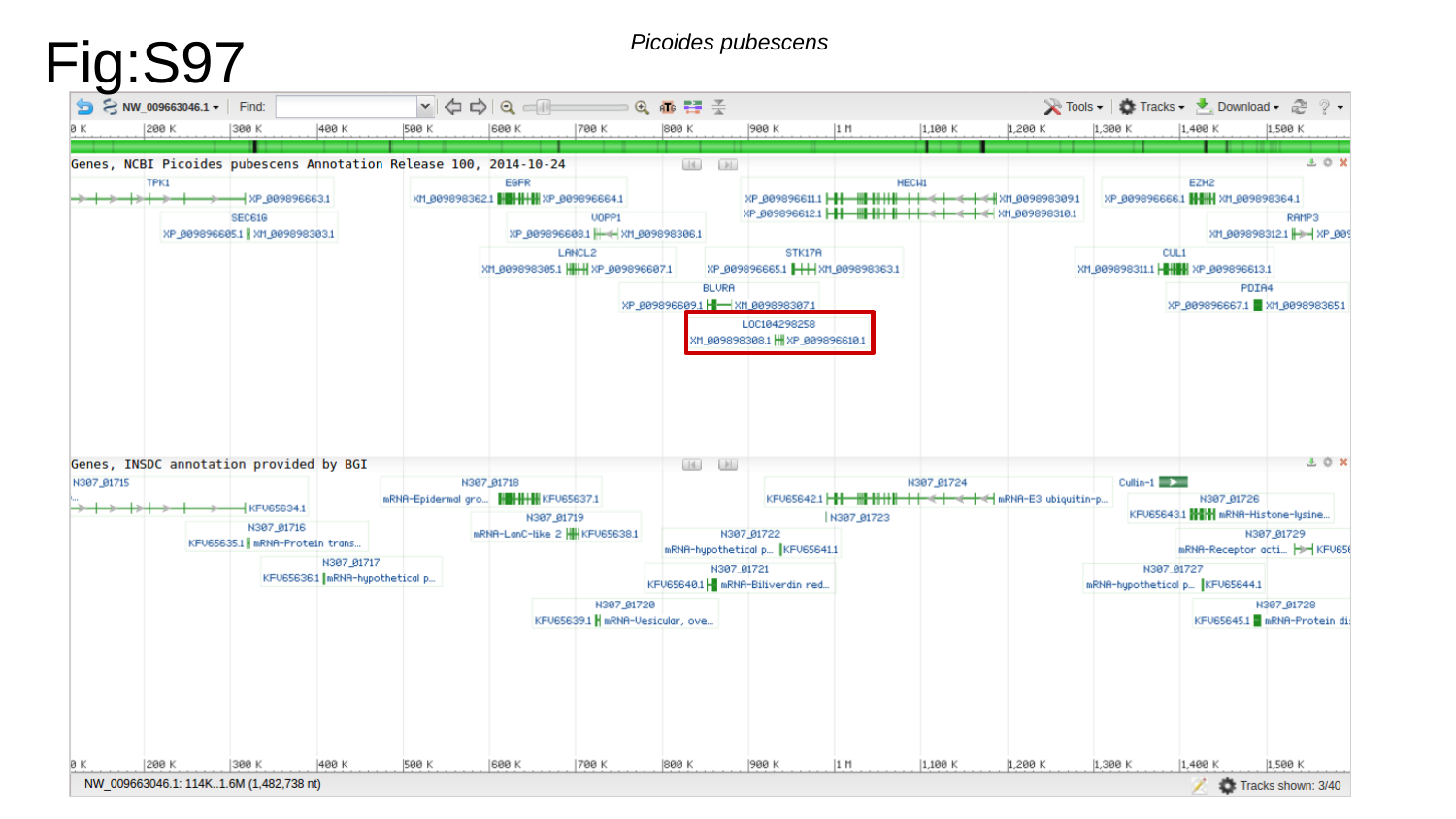

Fig:S97
Picoides pubescens

### Slide 98
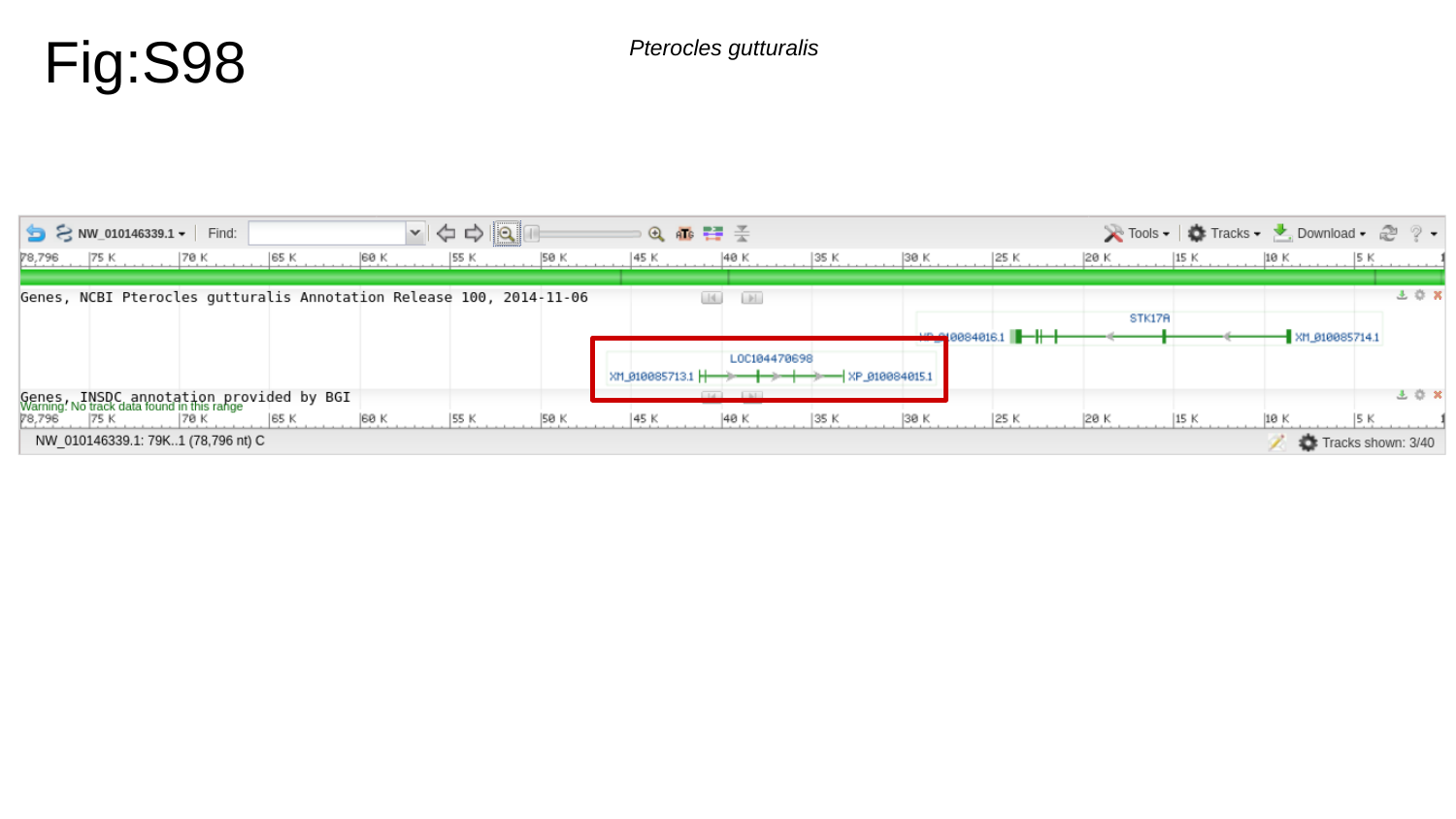

Fig:S98
Pterocles gutturalis

### Slide 99
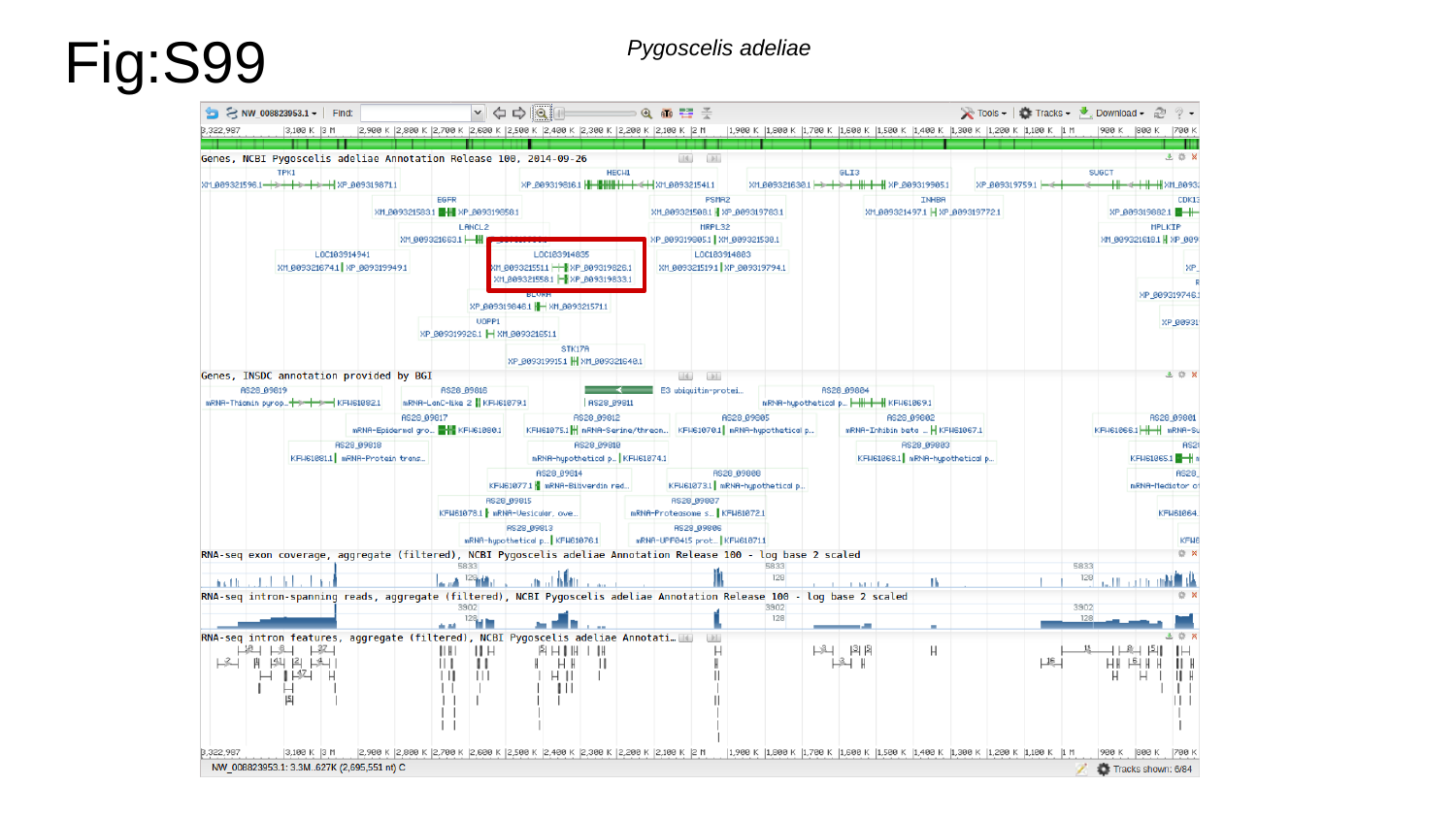

Fig:S99
Pygoscelis adeliae

### Slide 100
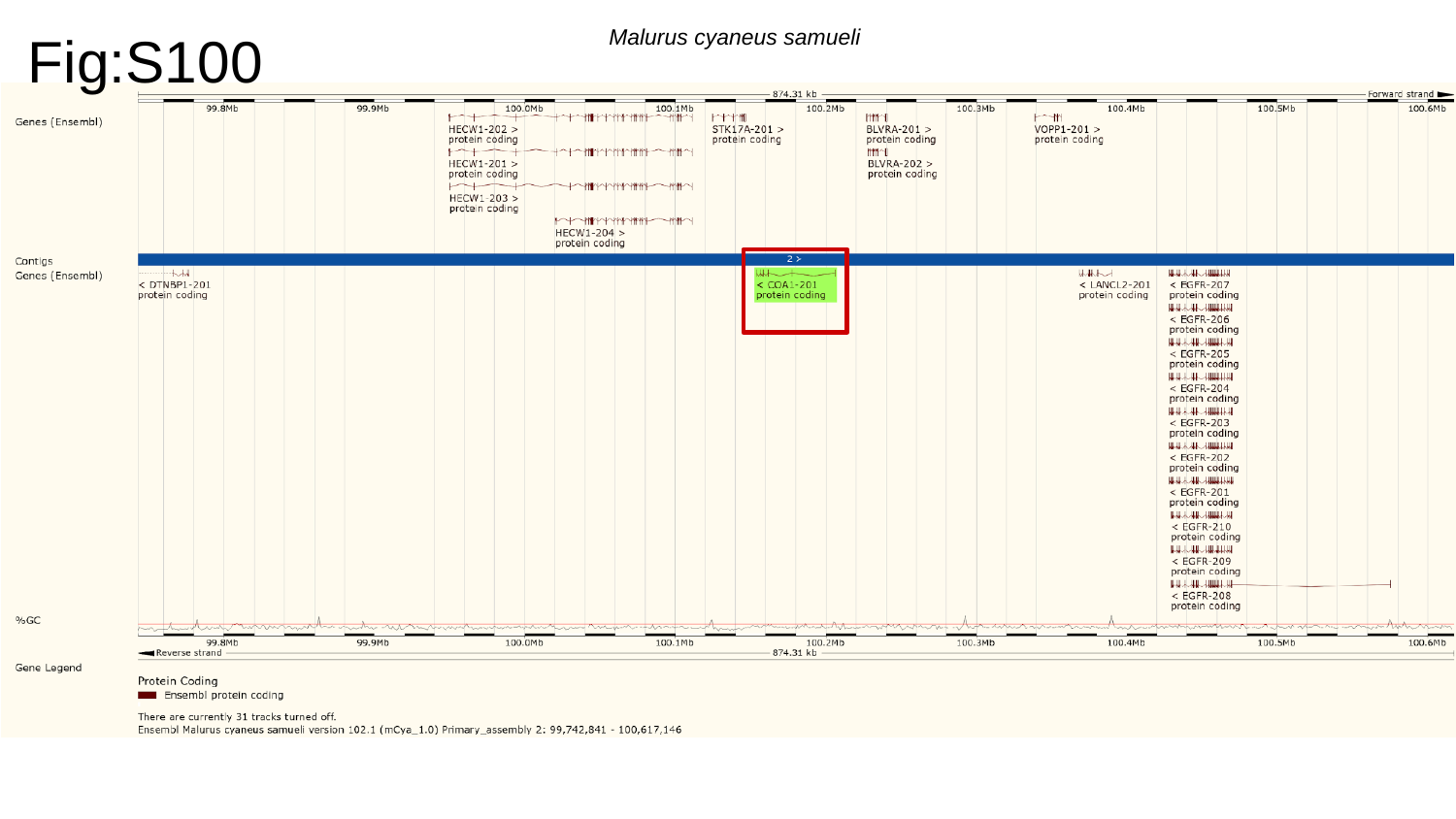

Fig:S100
Malurus cyaneus samueli

### Slide 101
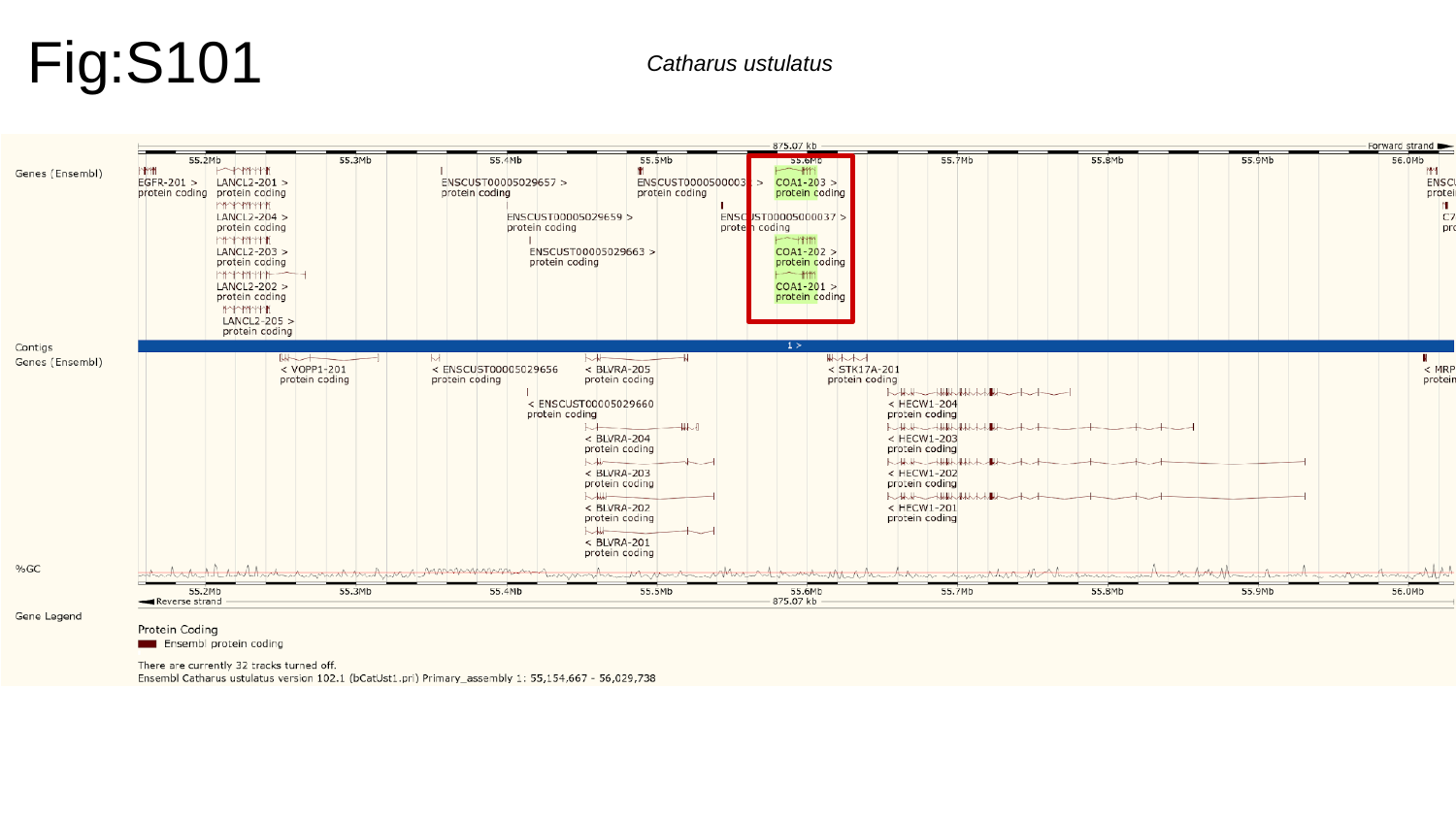

Fig:S101
Catharus ustulatus

### Slide 102
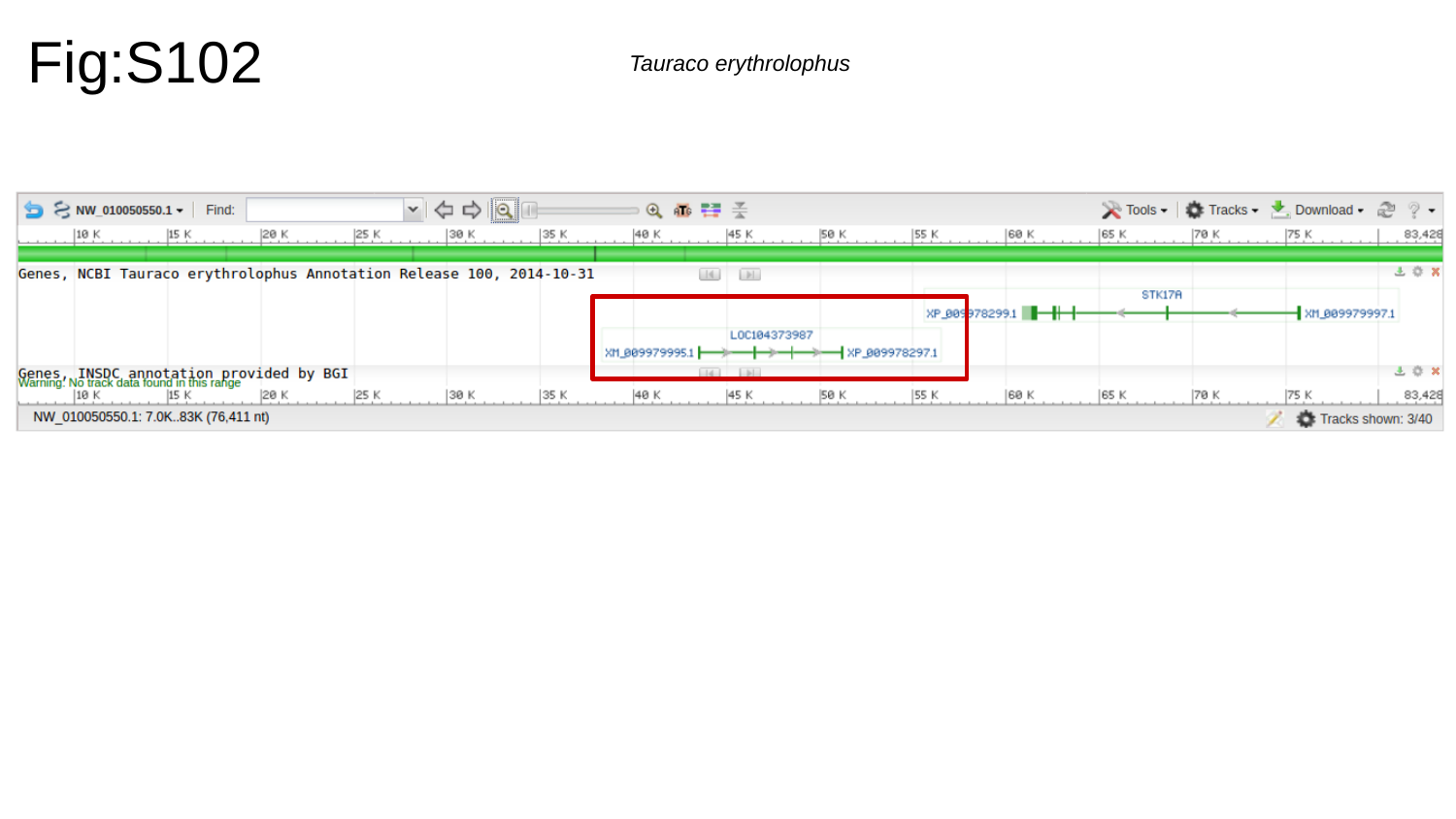

Fig:S102
Tauraco erythrolophus

### Slide 103
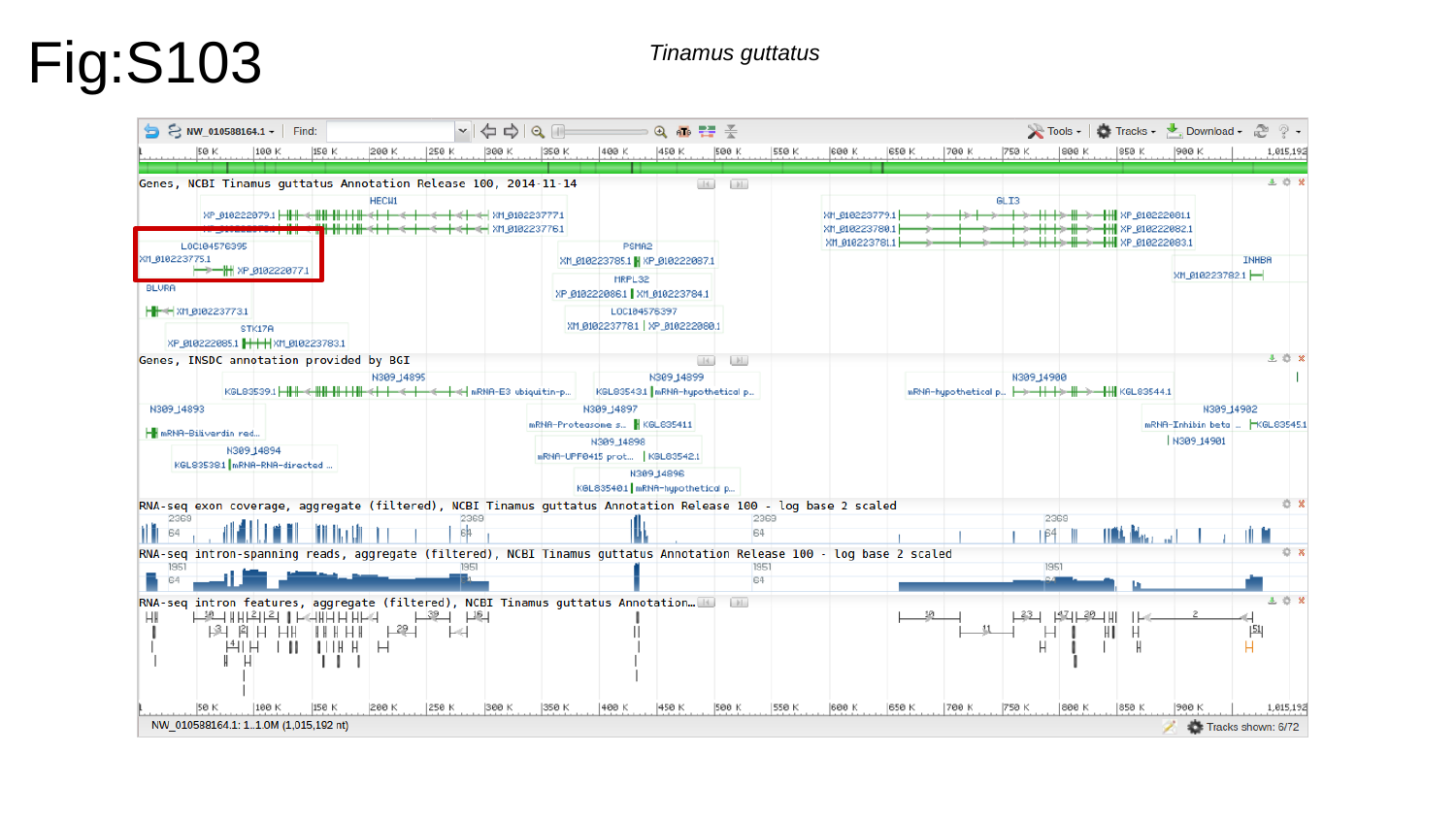

Fig:S103
Tinamus guttatus

### Slide 104
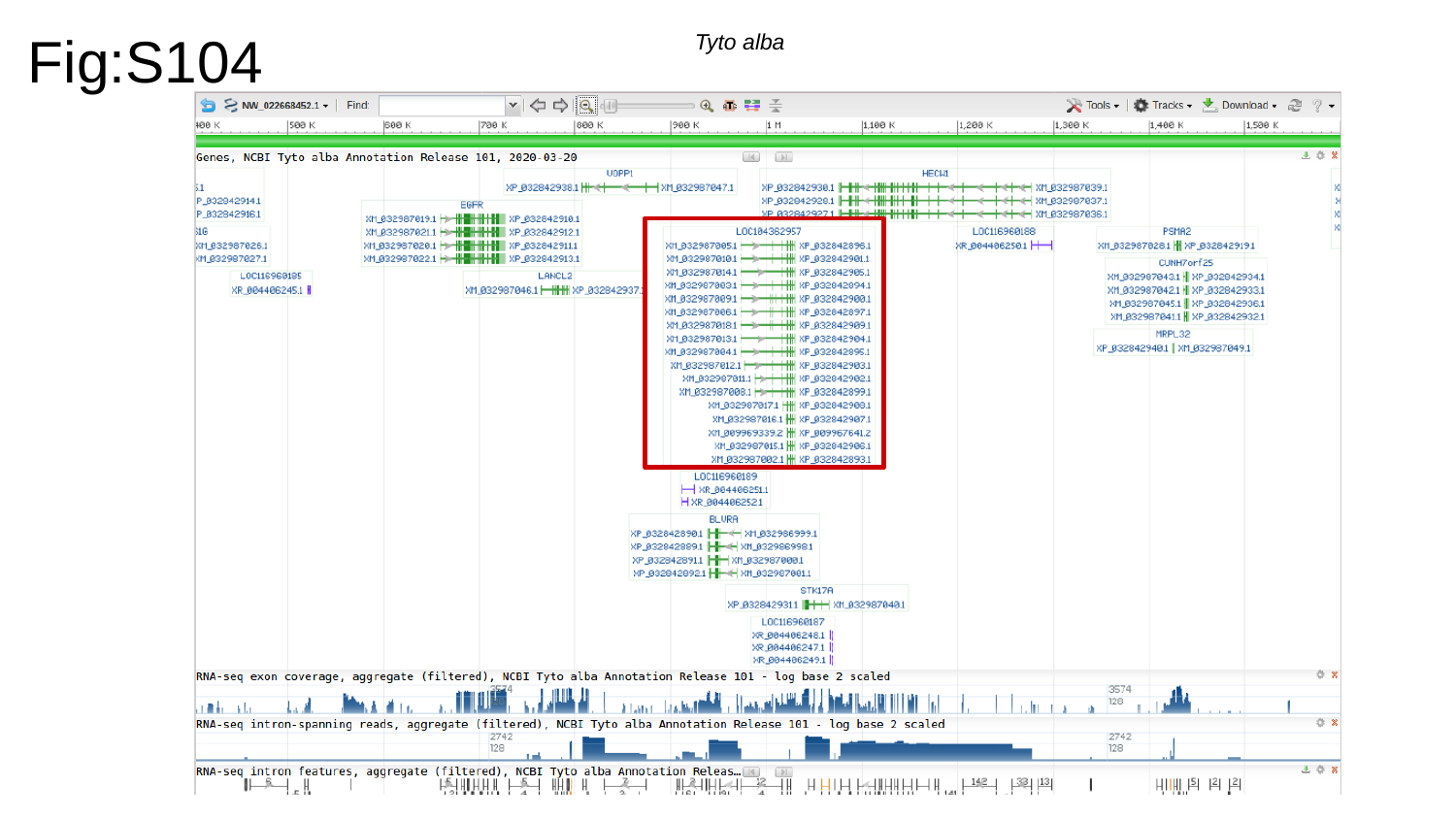

Fig:S104
Tyto alba

### Slide 105
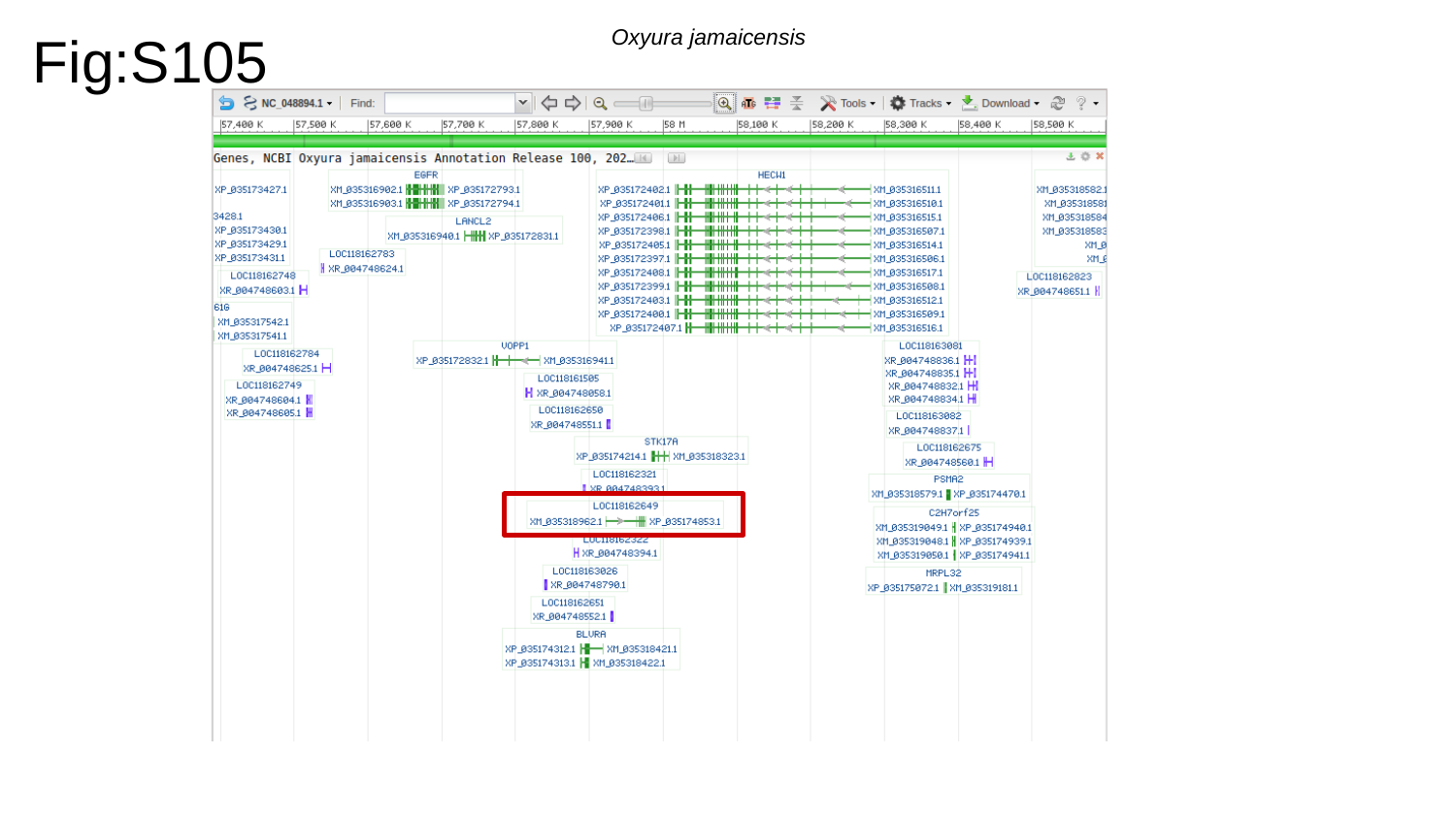

Fig:S105
Oxyura jamaicensis

### Slide 106
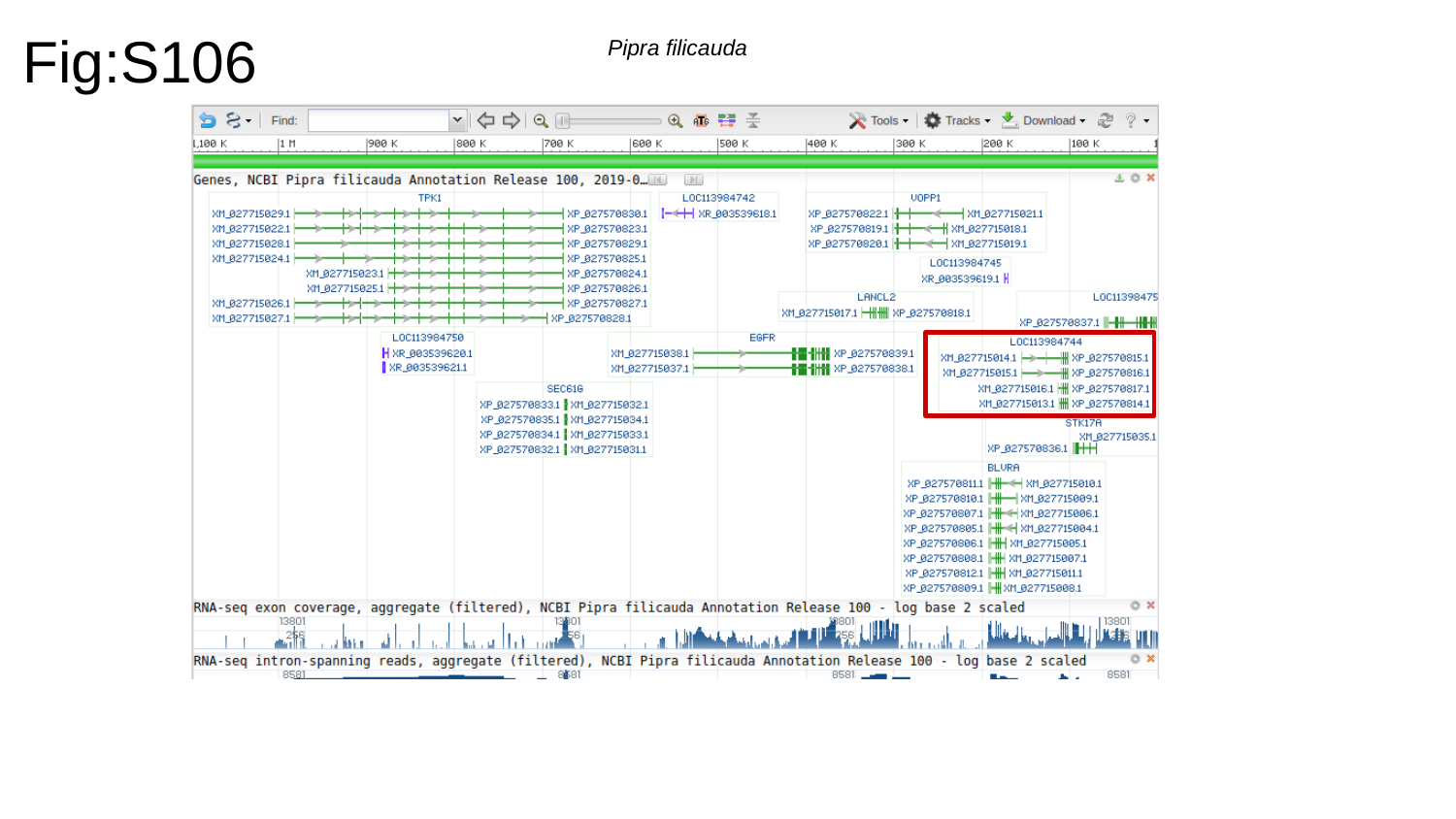

Fig:S106
Pipra filicauda

### Slide 107
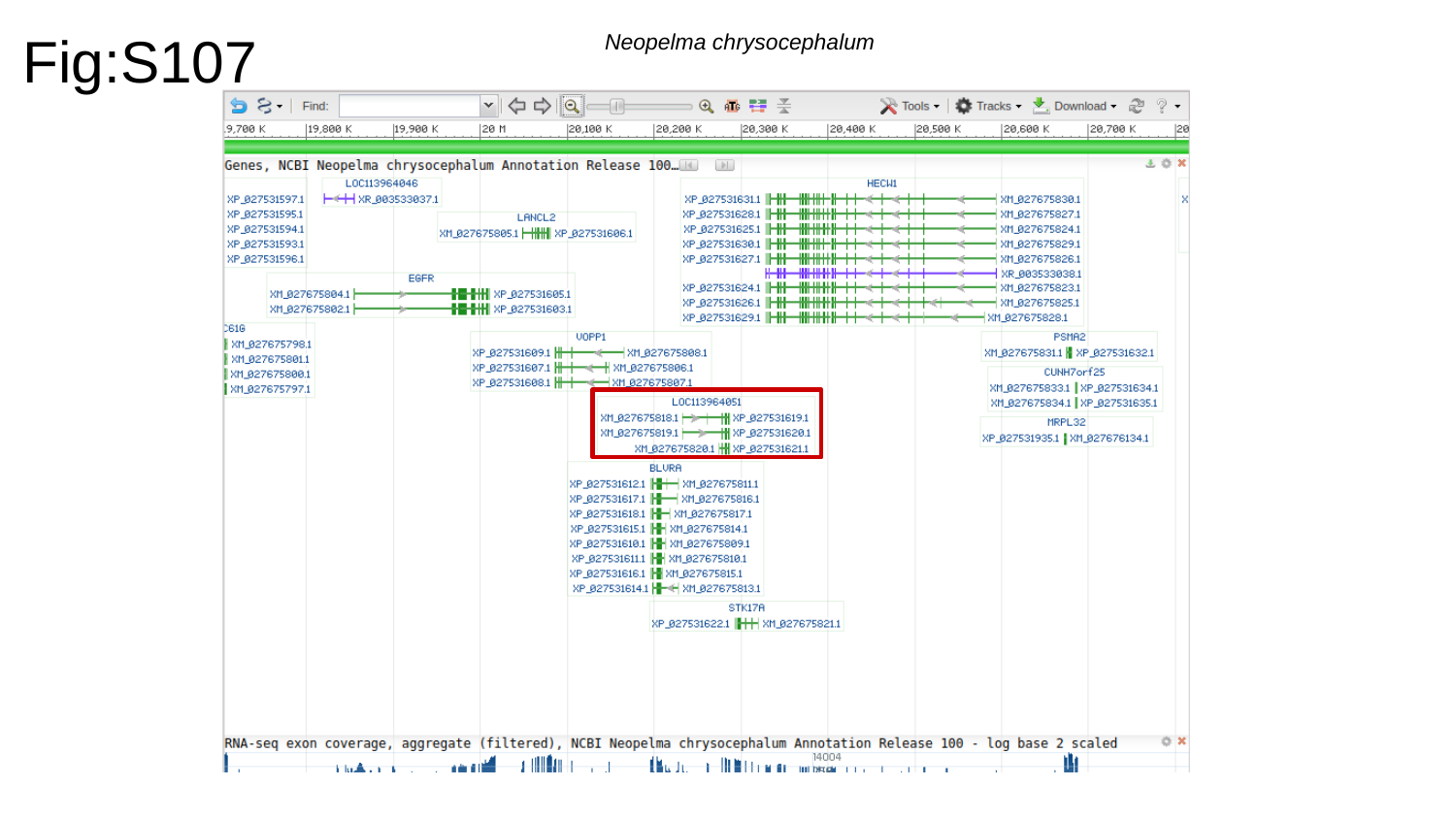

Fig:S107
Neopelma chrysocephalum

### Slide 108
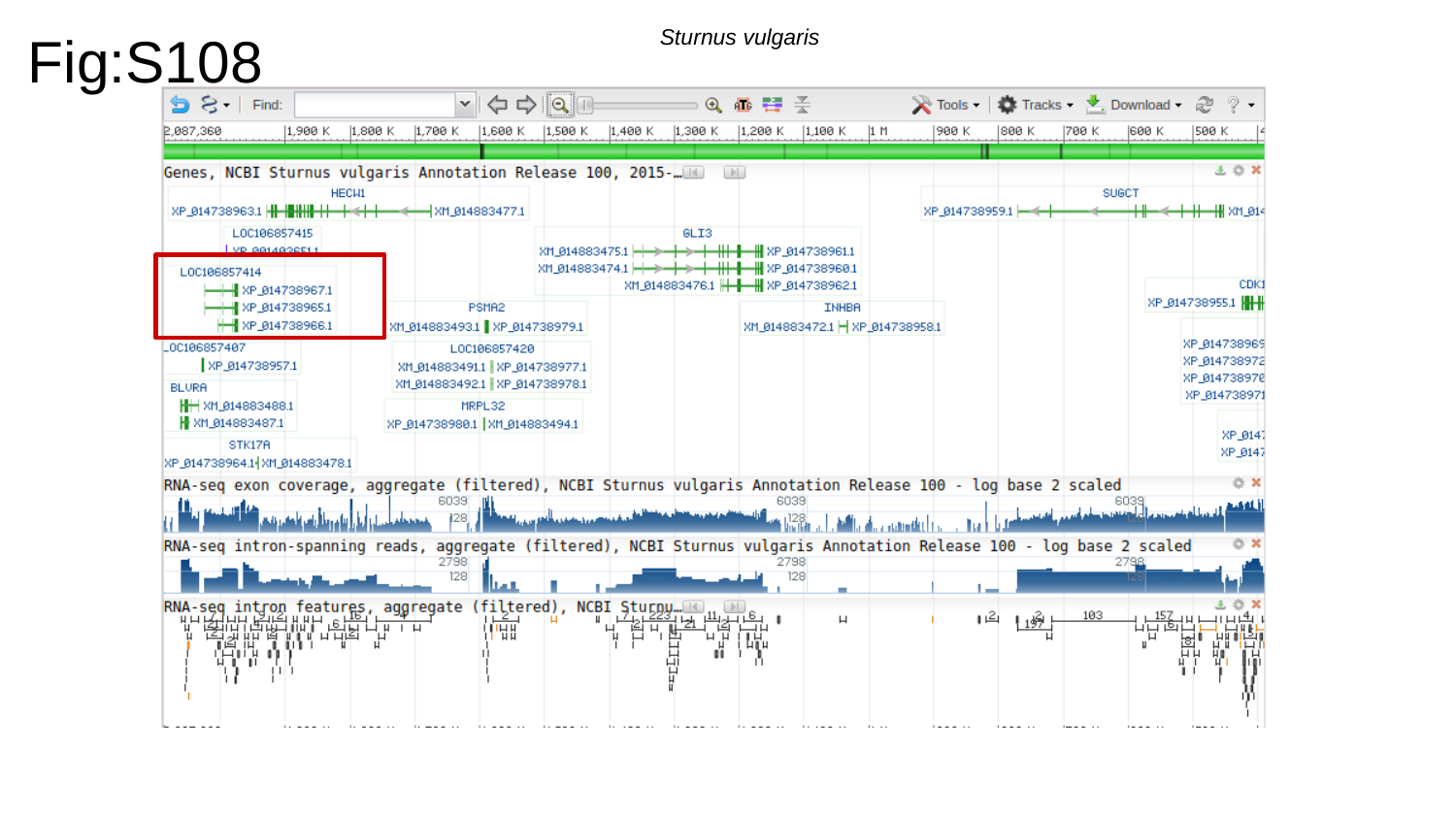

Fig:S108
Sturnus vulgaris

### Slide 109
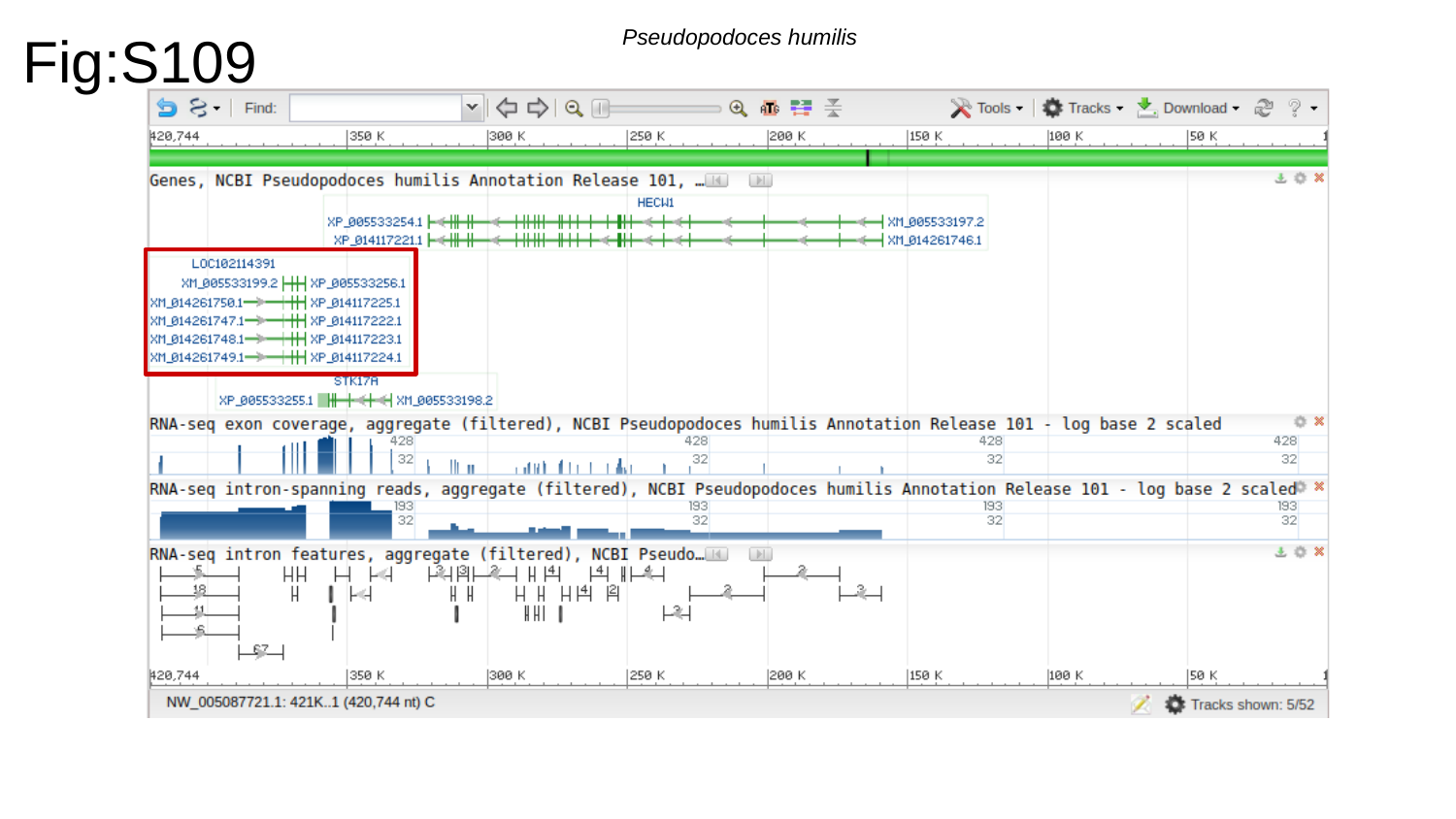

Fig:S109
Pseudopodoces humilis

### Slide 110
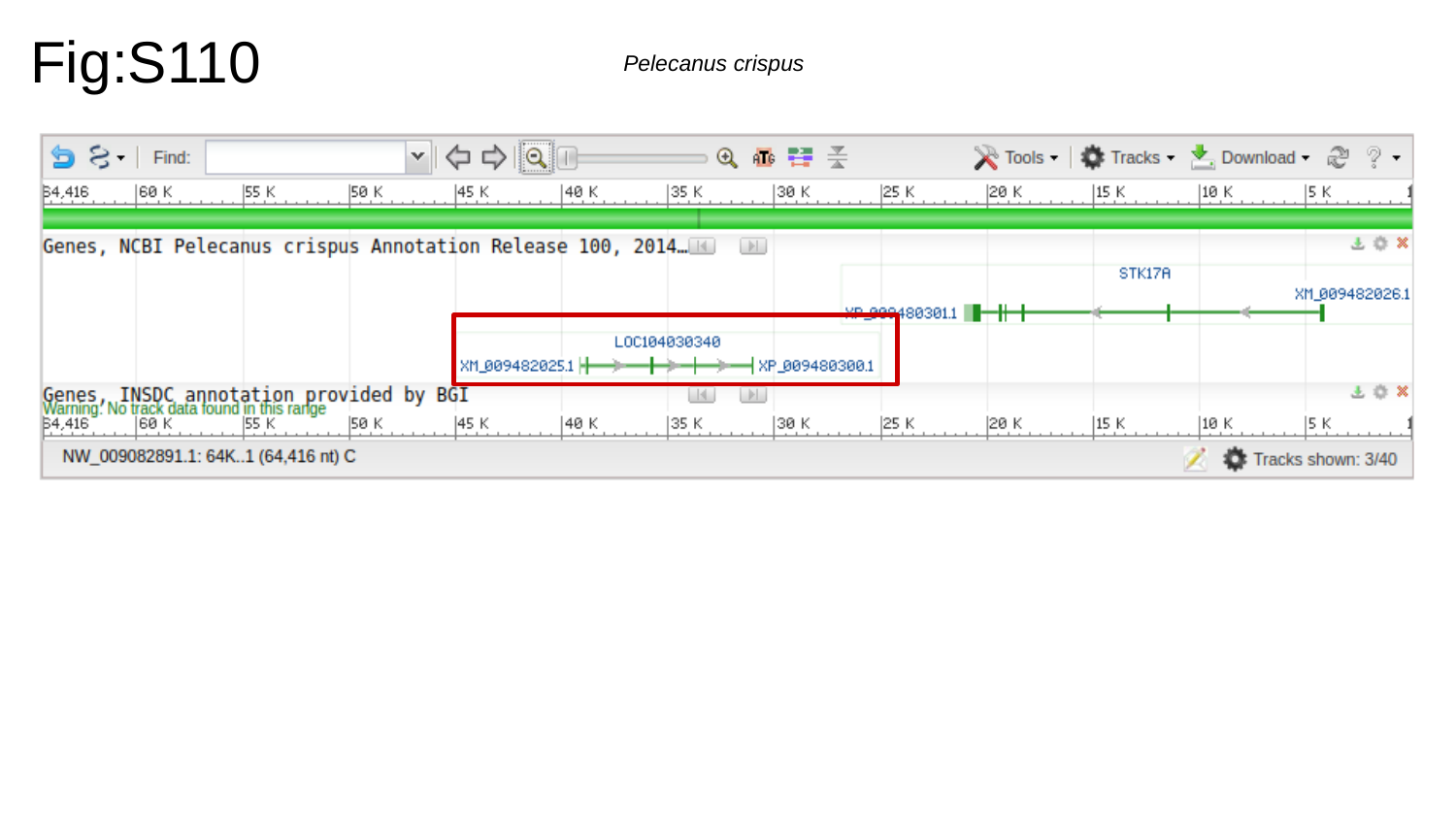

Fig:S110
Pelecanus crispus

### Slide 111
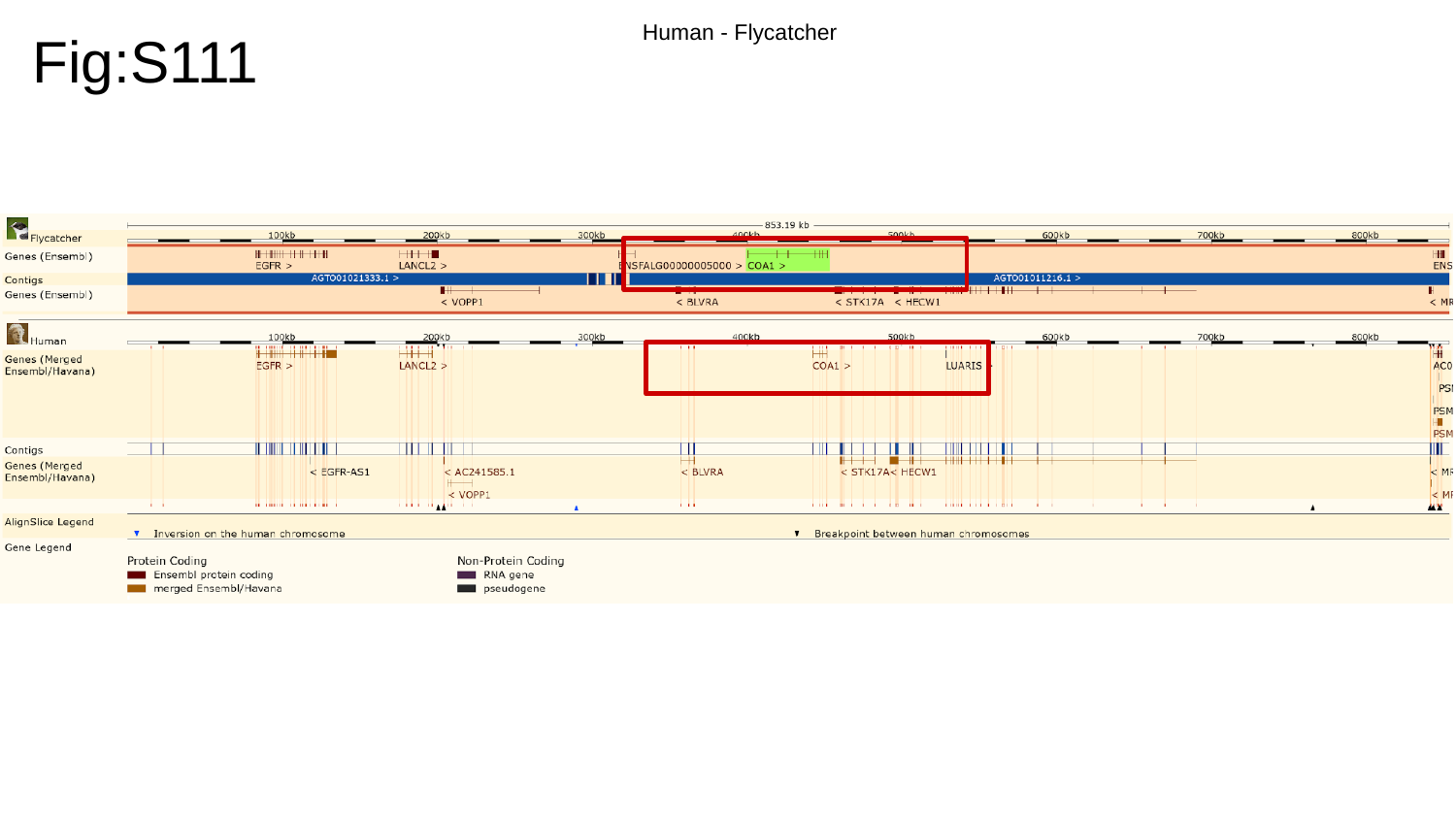

Fig:S111
Human - Flycatcher

### Slide 112
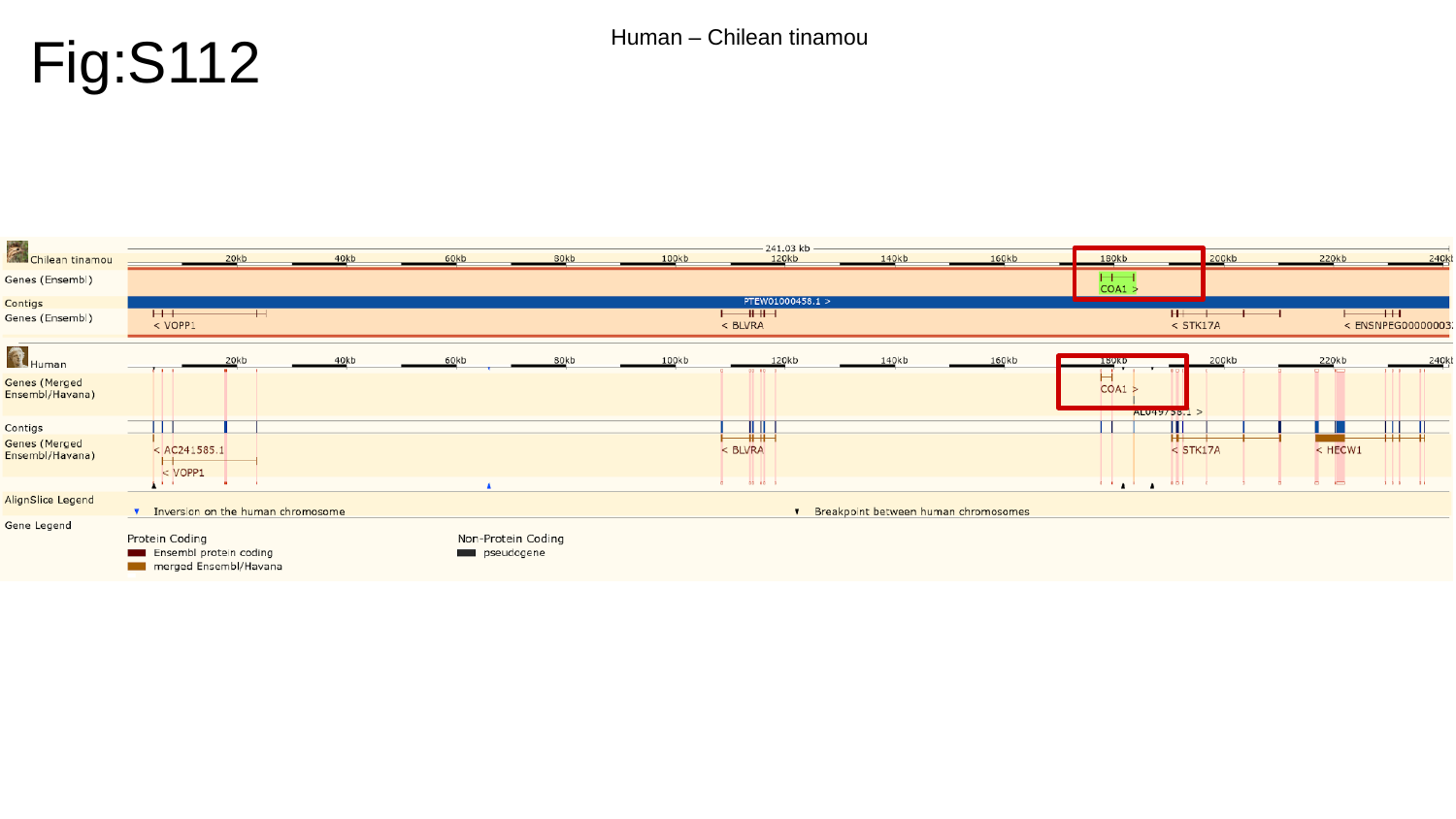

Fig:S112
Human – Chilean tinamou

### Slide 113
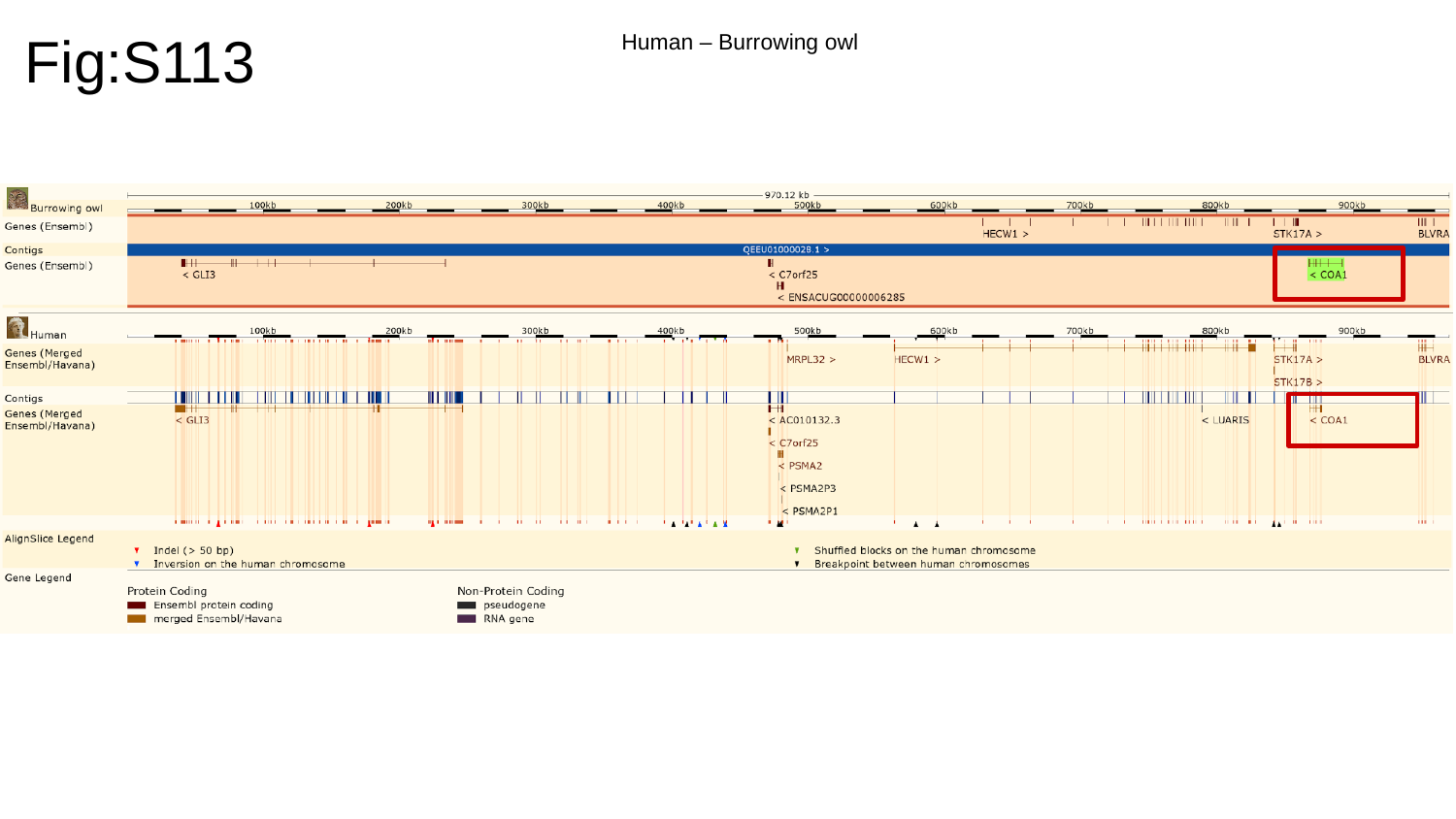

Fig:S113
Human – Burrowing owl

### Slide 114
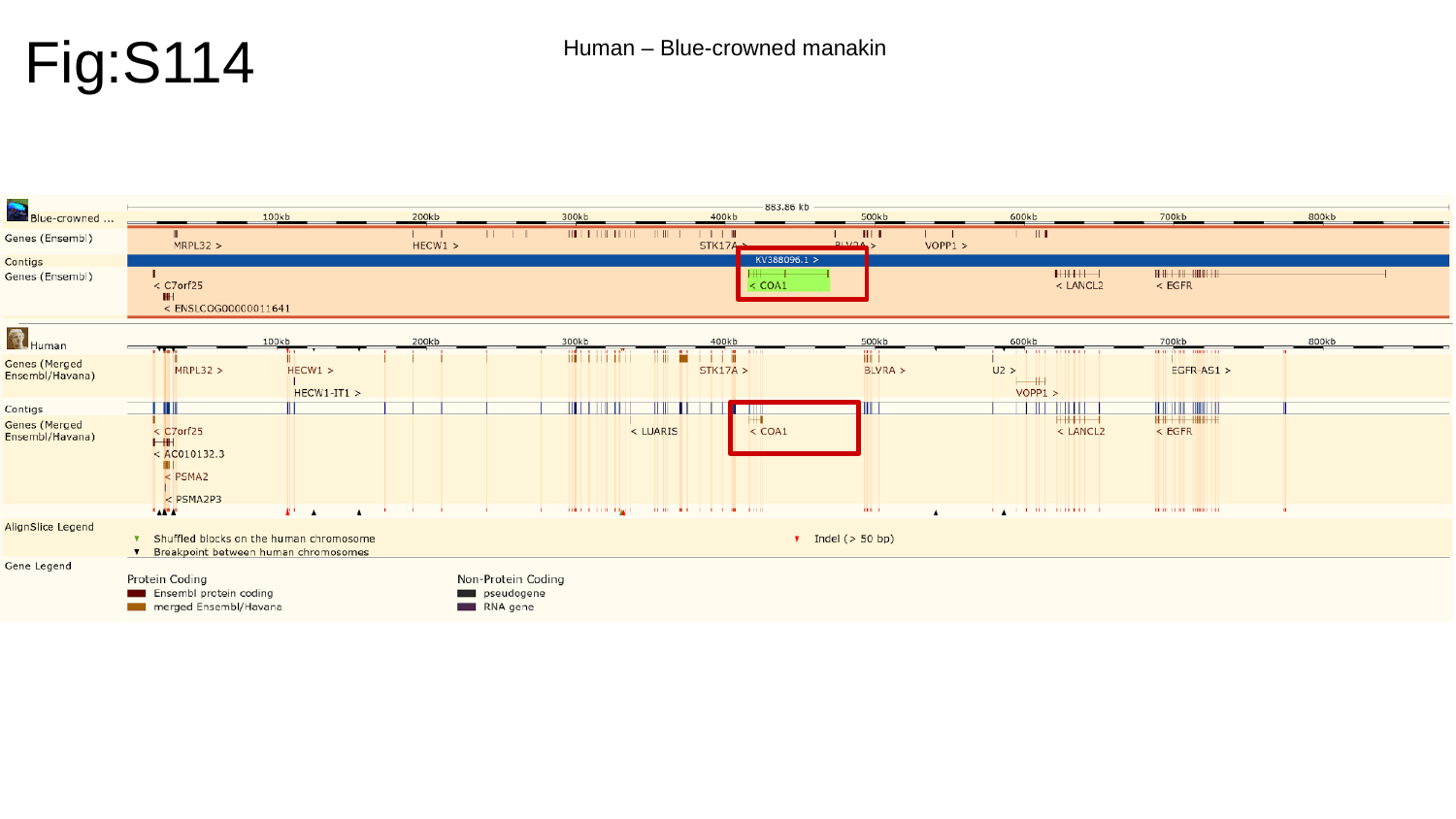

Fig:S114
Human – Blue-crowned manakin

### Slide 115
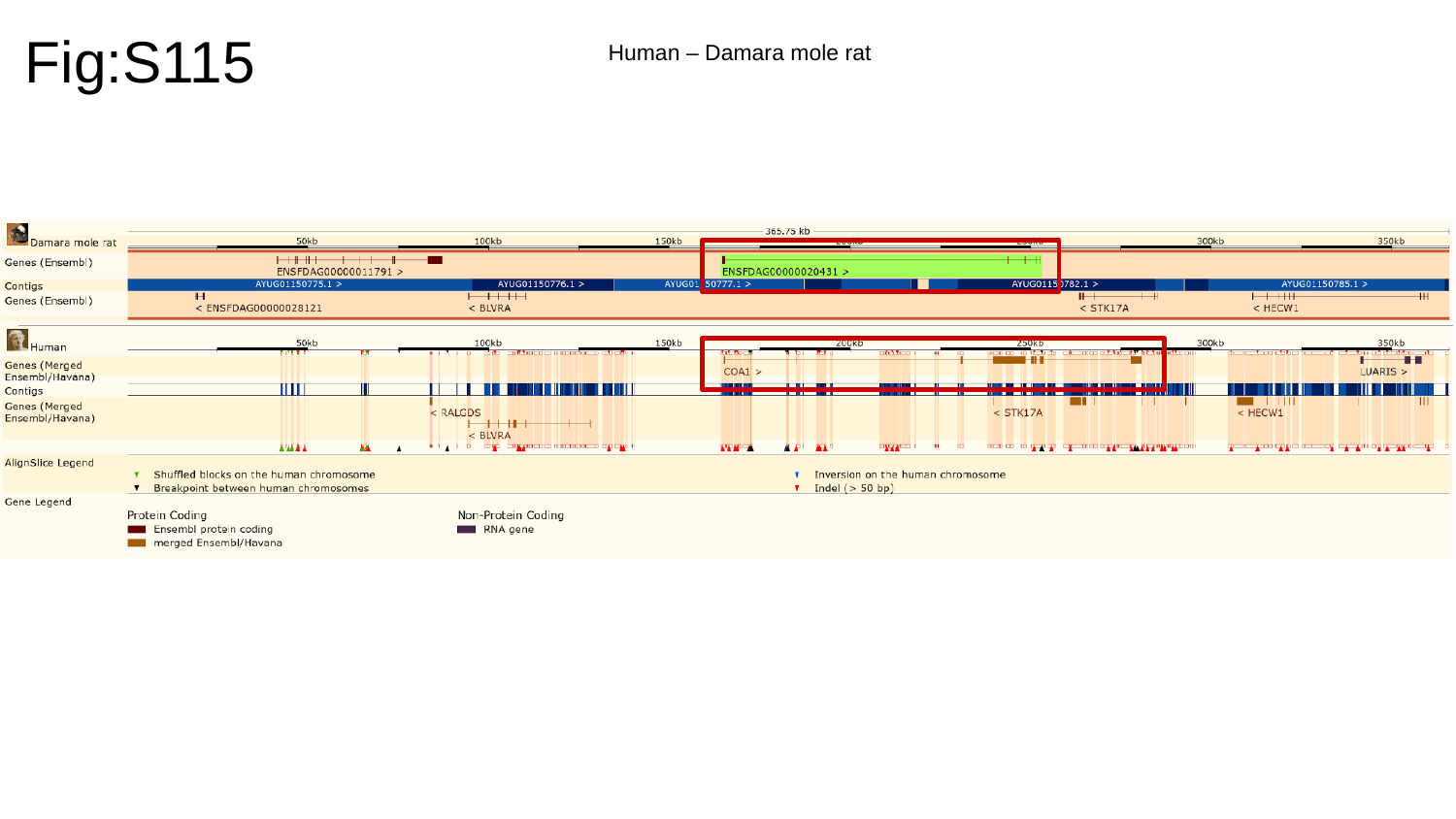

Fig:S115
Human – Damara mole rat

### Slide 116
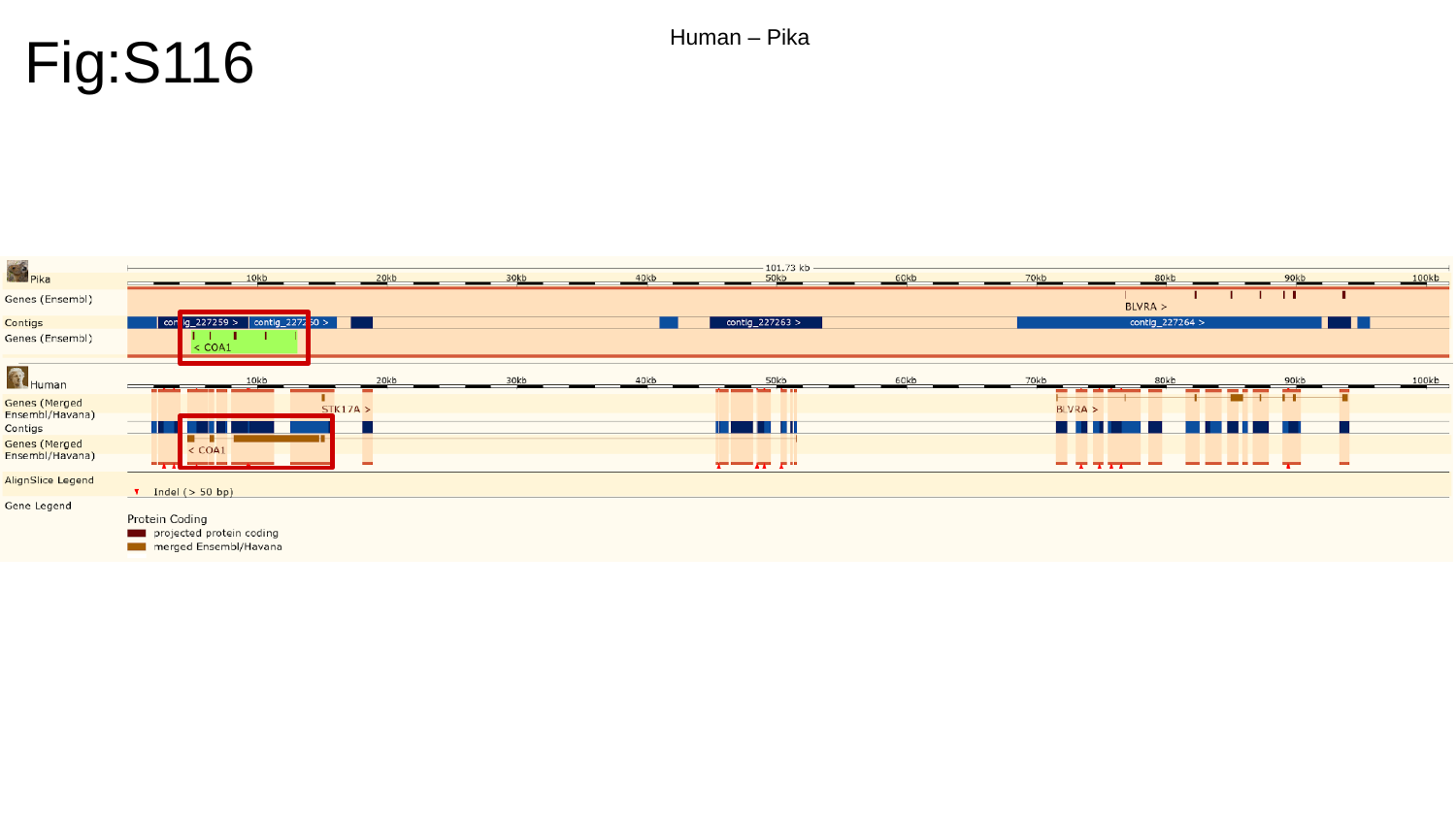

Fig:S116
Human – Pika

### Slide 117
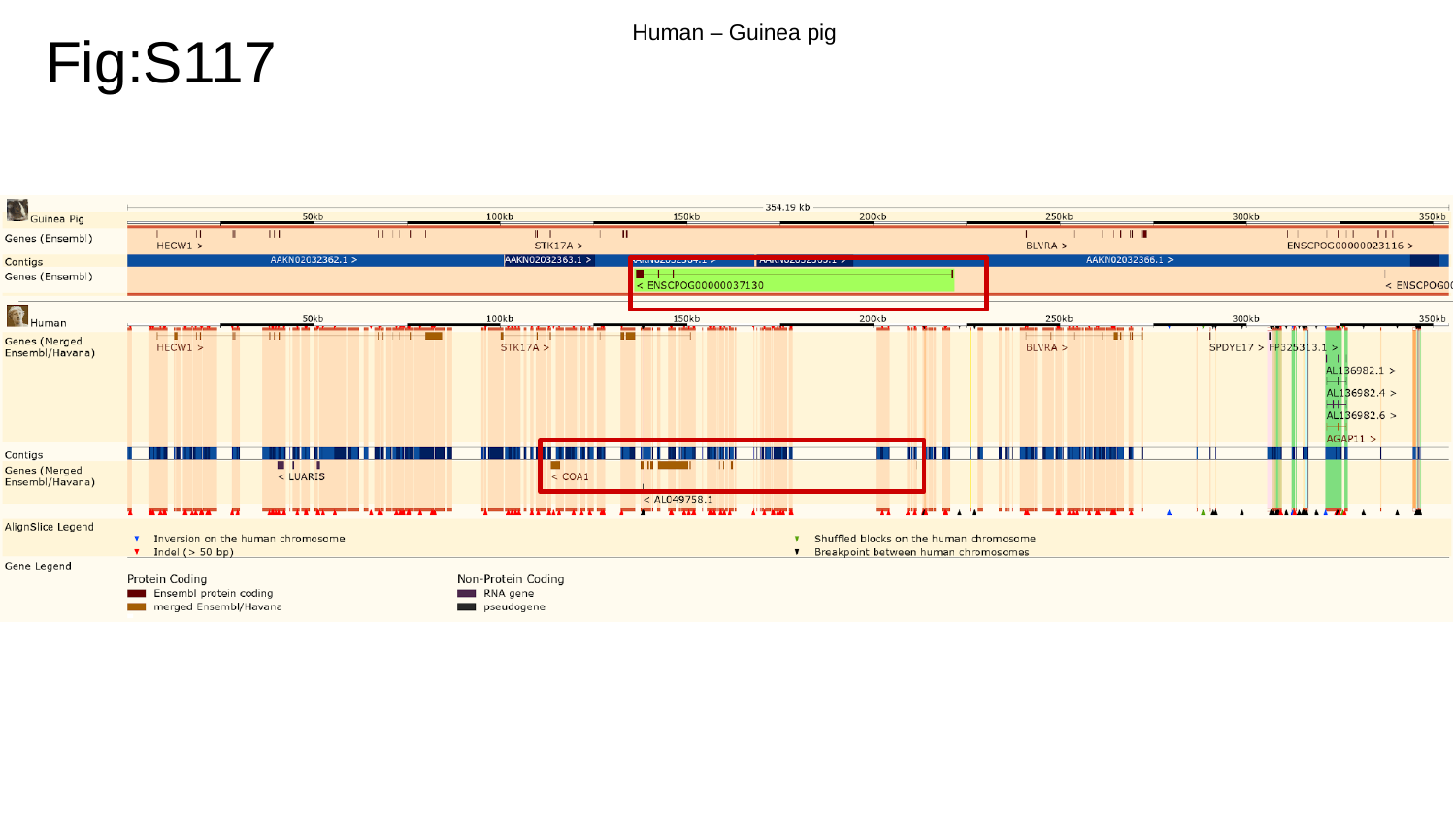

Fig:S117
Human – Guinea pig

### Slide 118
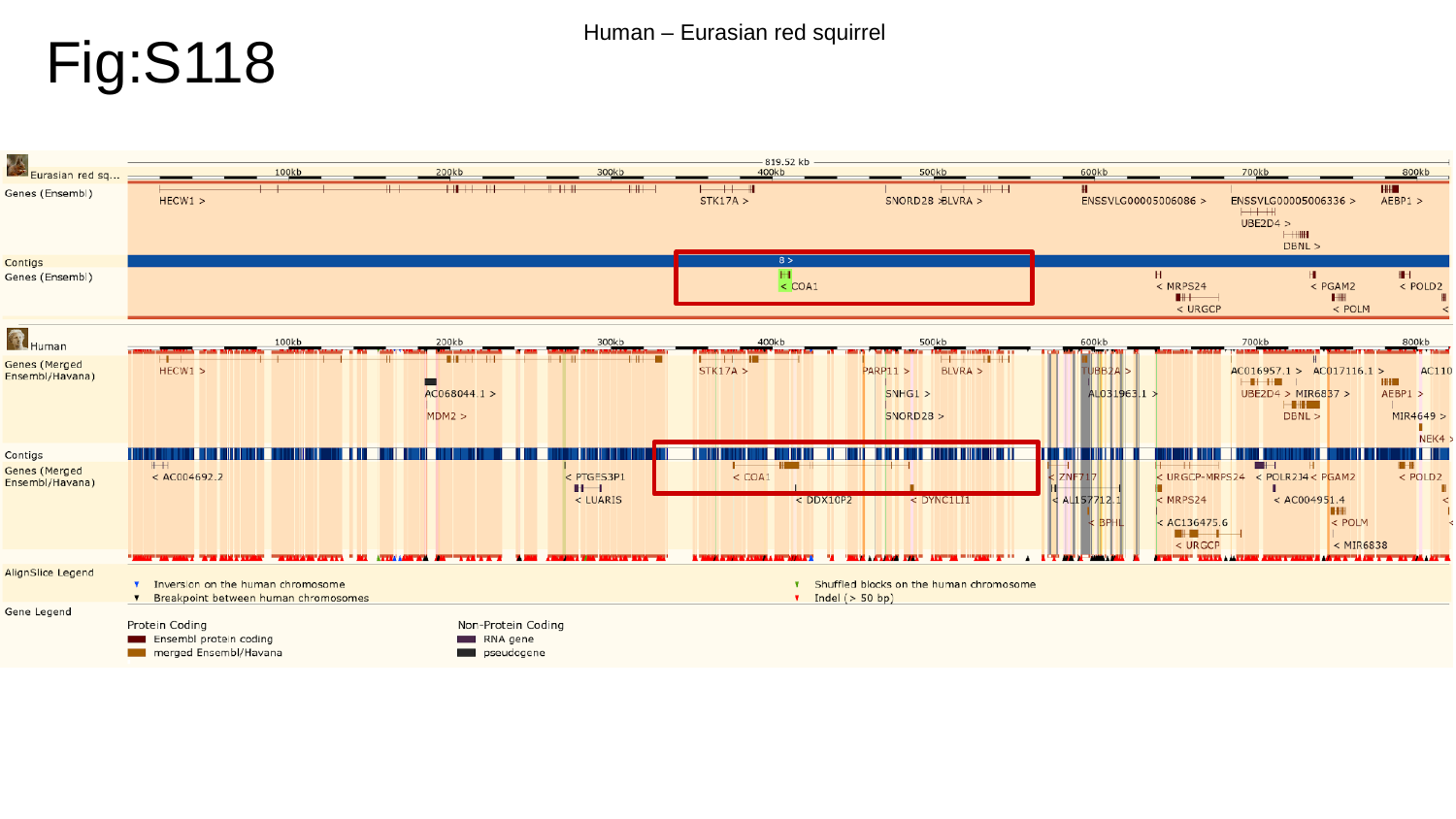

Fig:S118
Human – Eurasian red squirrel

### Slide 119
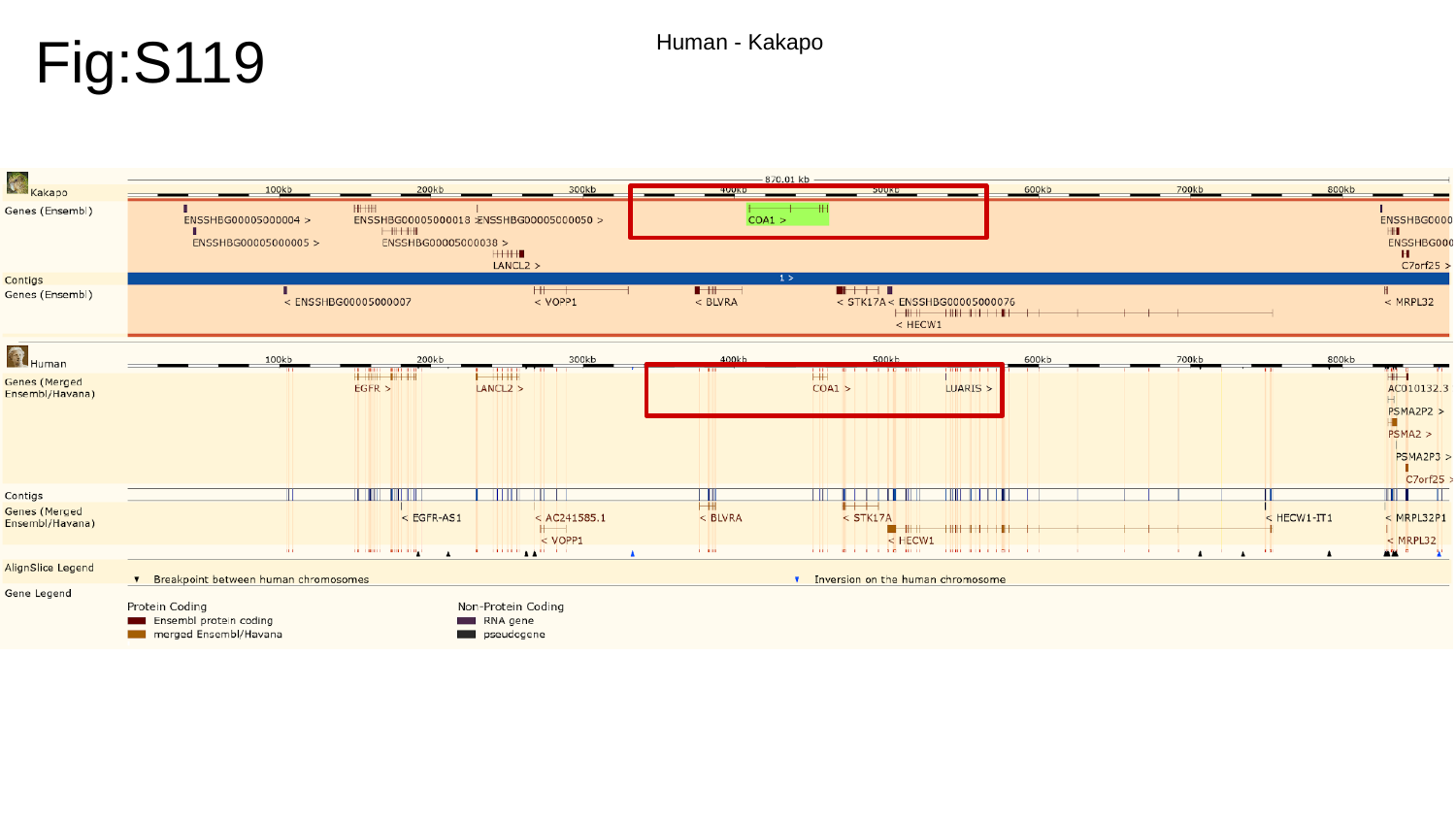

Fig:S119
Human - Kakapo

### Slide 120
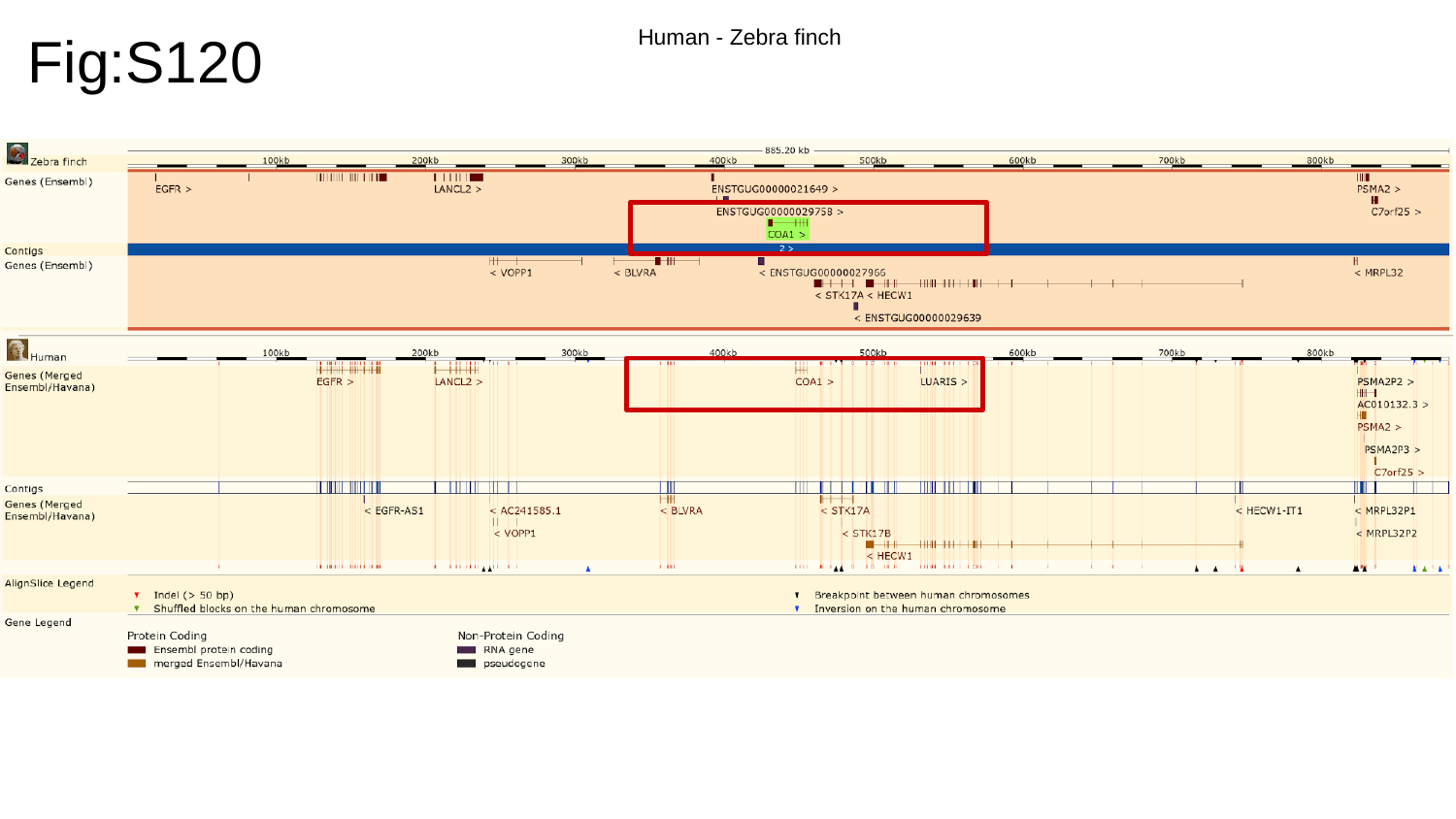

Fig:S120
Human - Zebra finch

### Slide 121
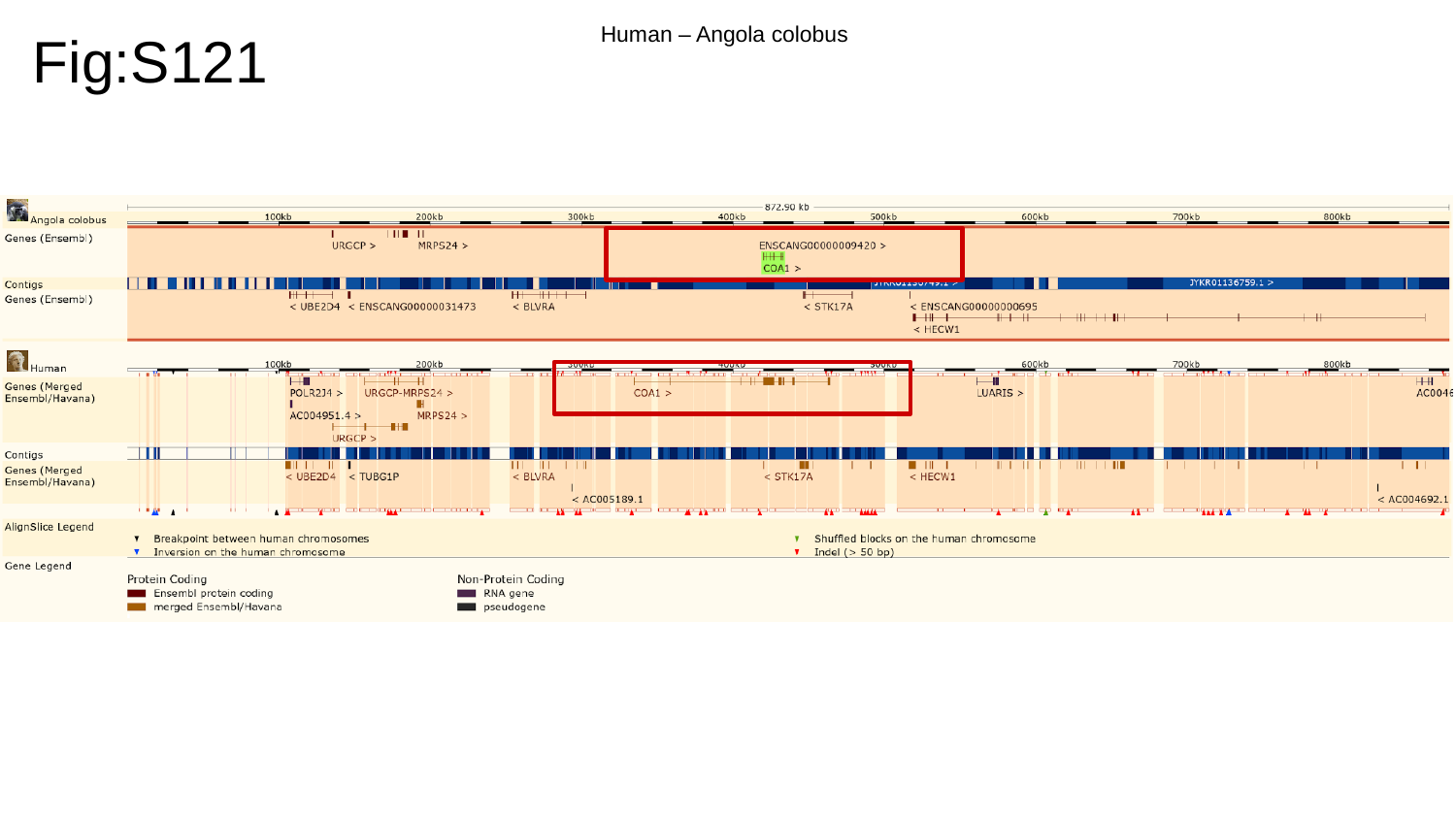

Fig:S121
Human – Angola colobus

### Slide 122
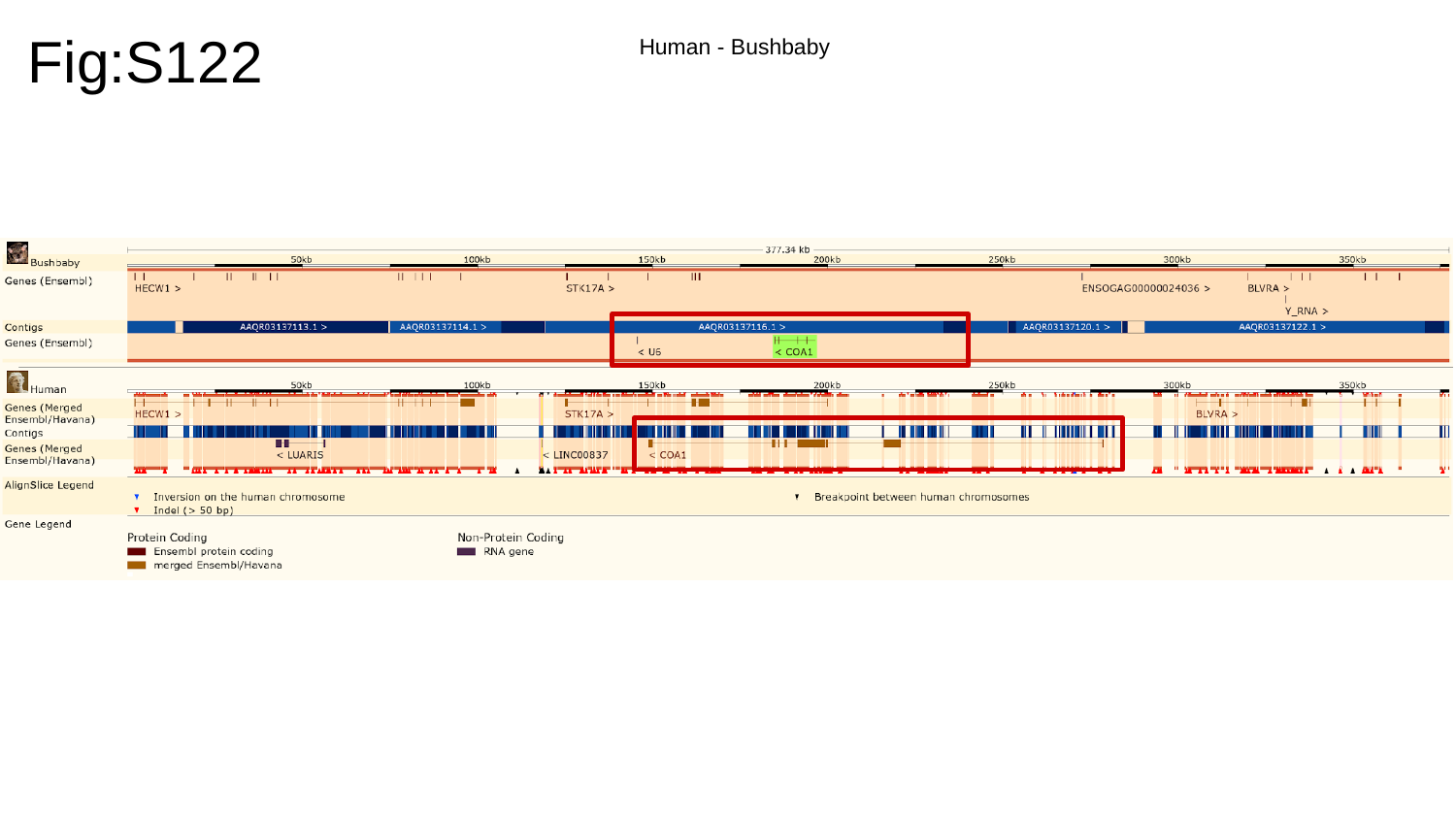

Fig:S122
Human - Bushbaby

### Slide 123
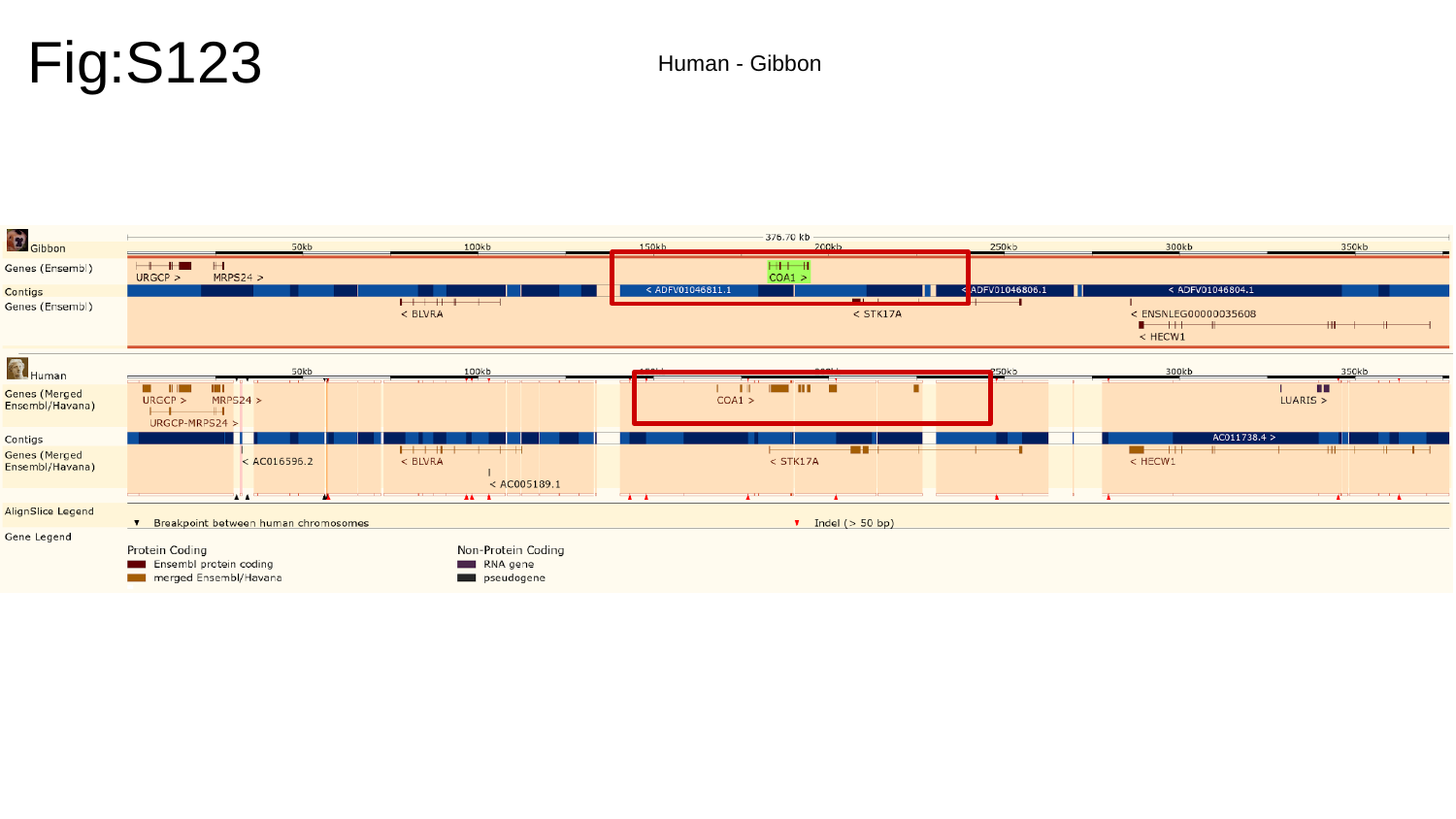

Fig:S123
Human - Gibbon

### Slide 124
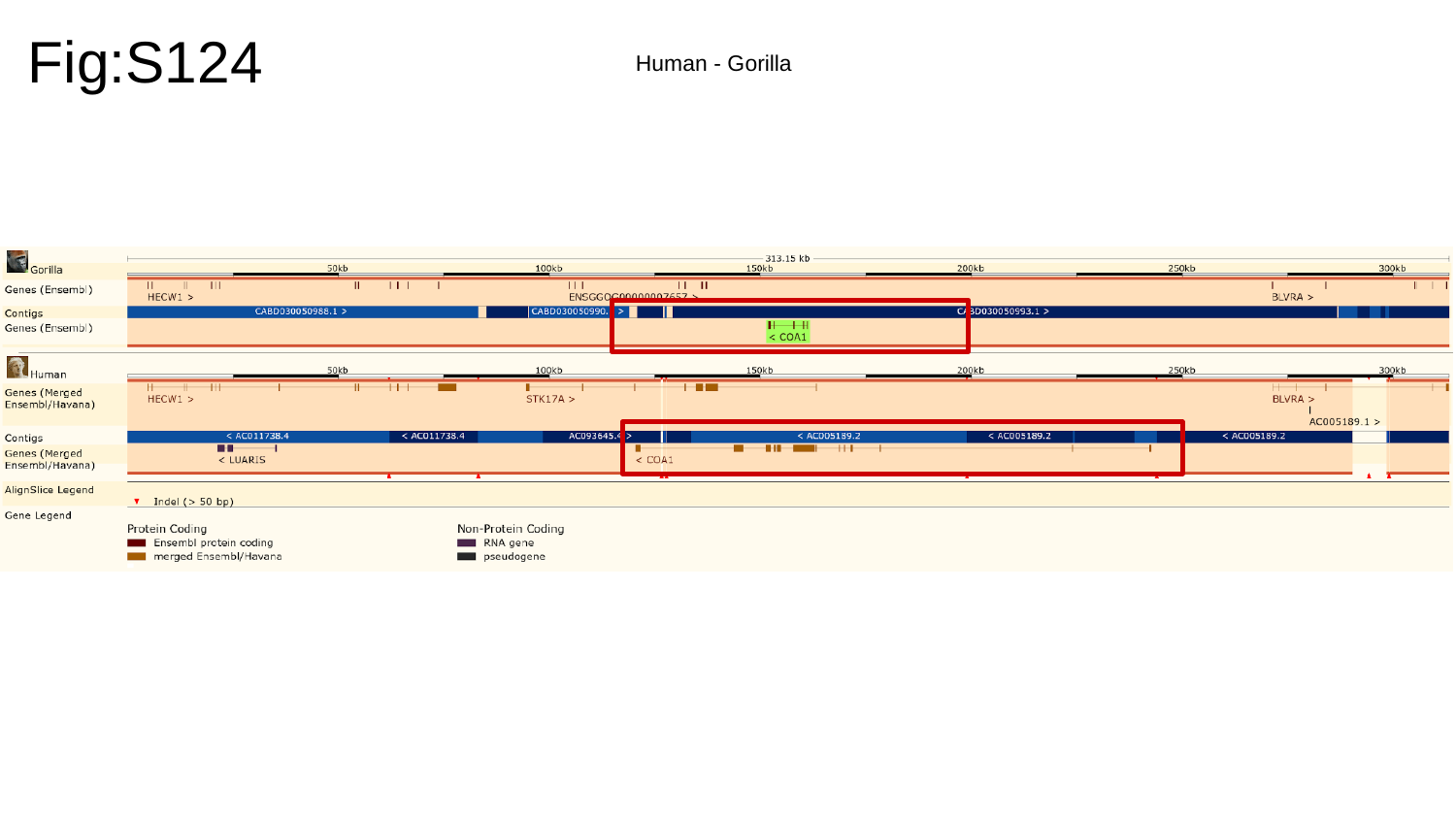

Fig:S124
Human - Gorilla

### Slide 125
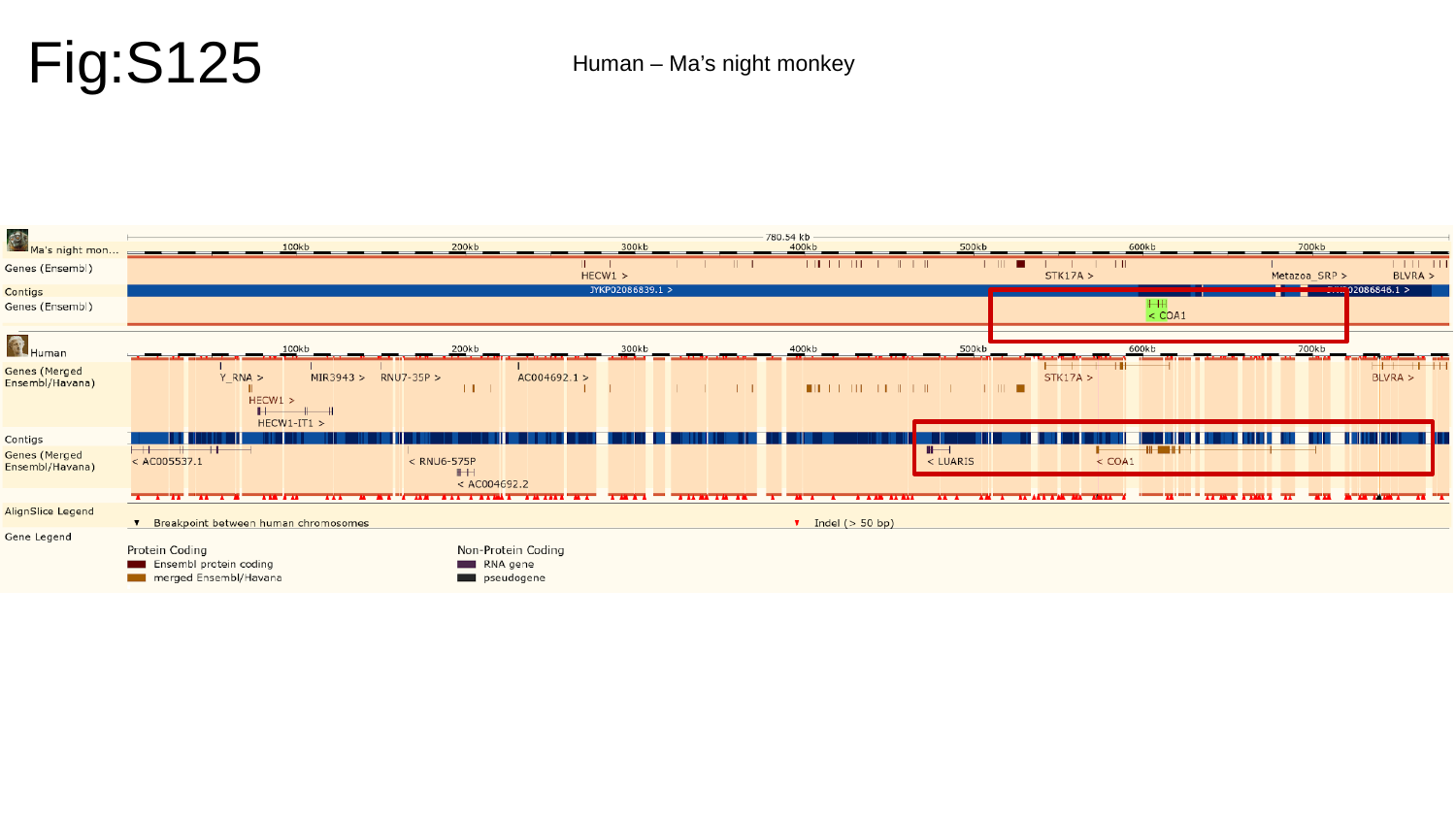

Fig:S125
Human – Ma’s night monkey

### Slide 126
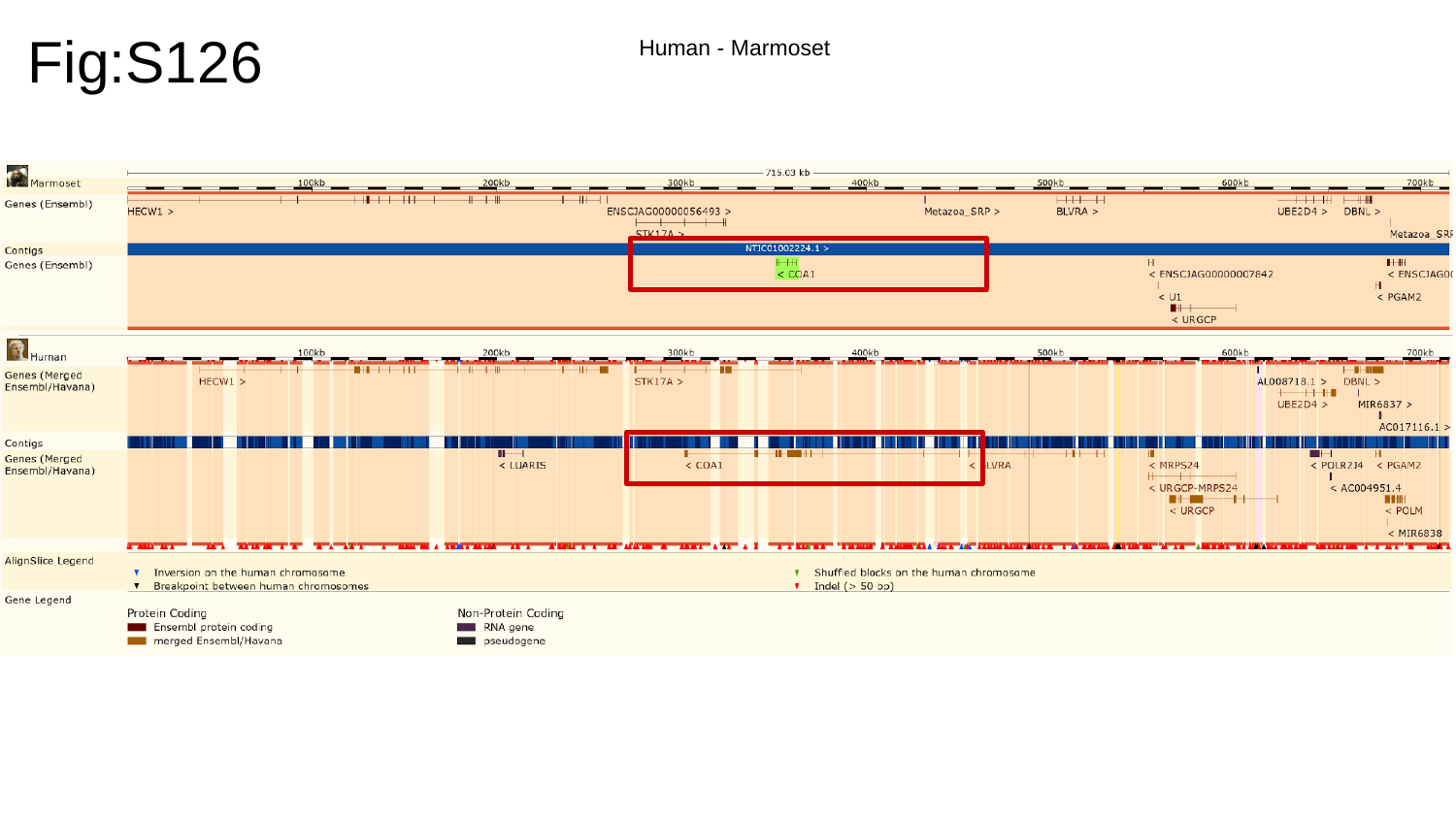

Fig:S126
Human - Marmoset

### Slide 127
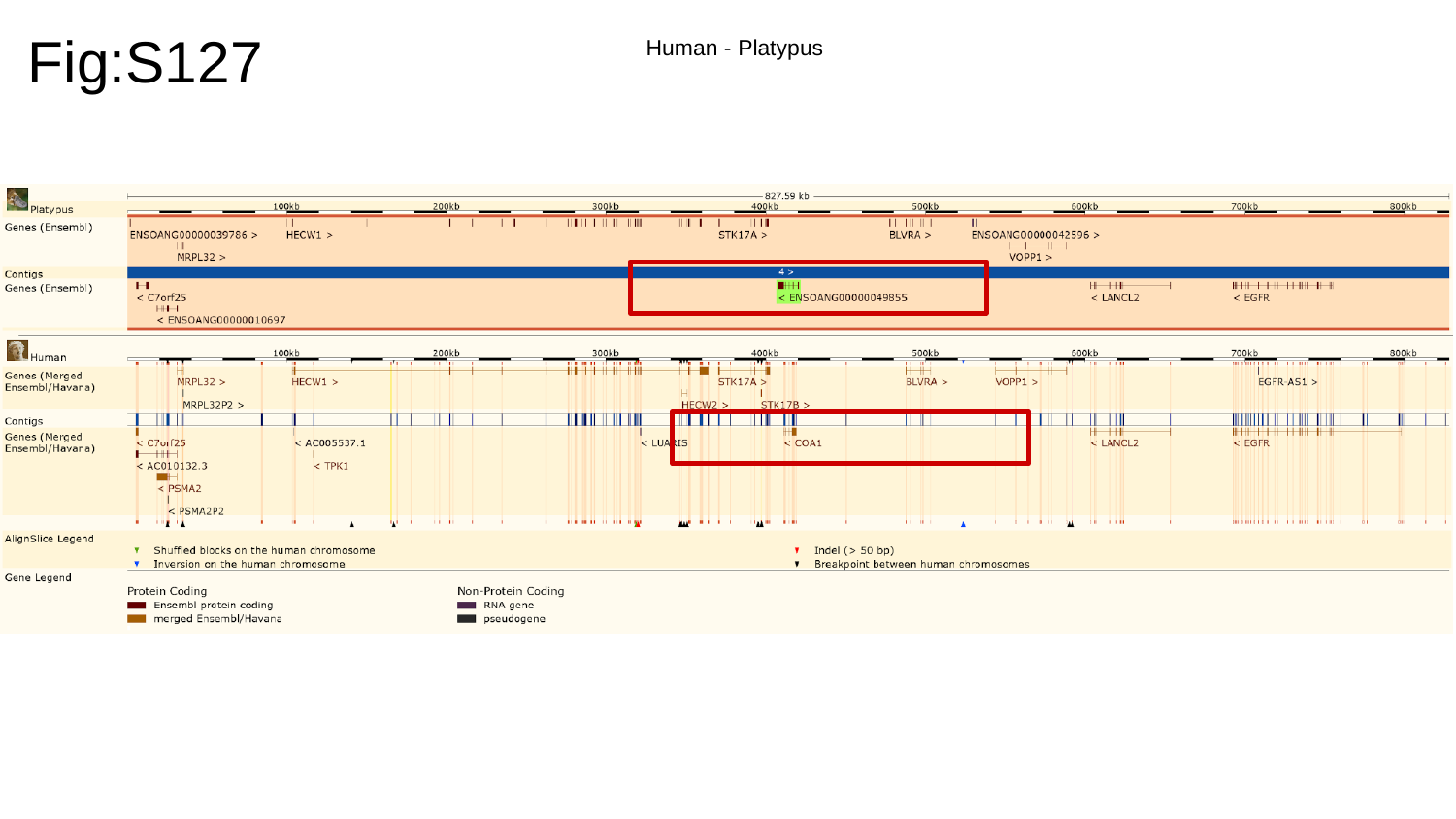

Fig:S127
Human - Platypus

### Slide 128
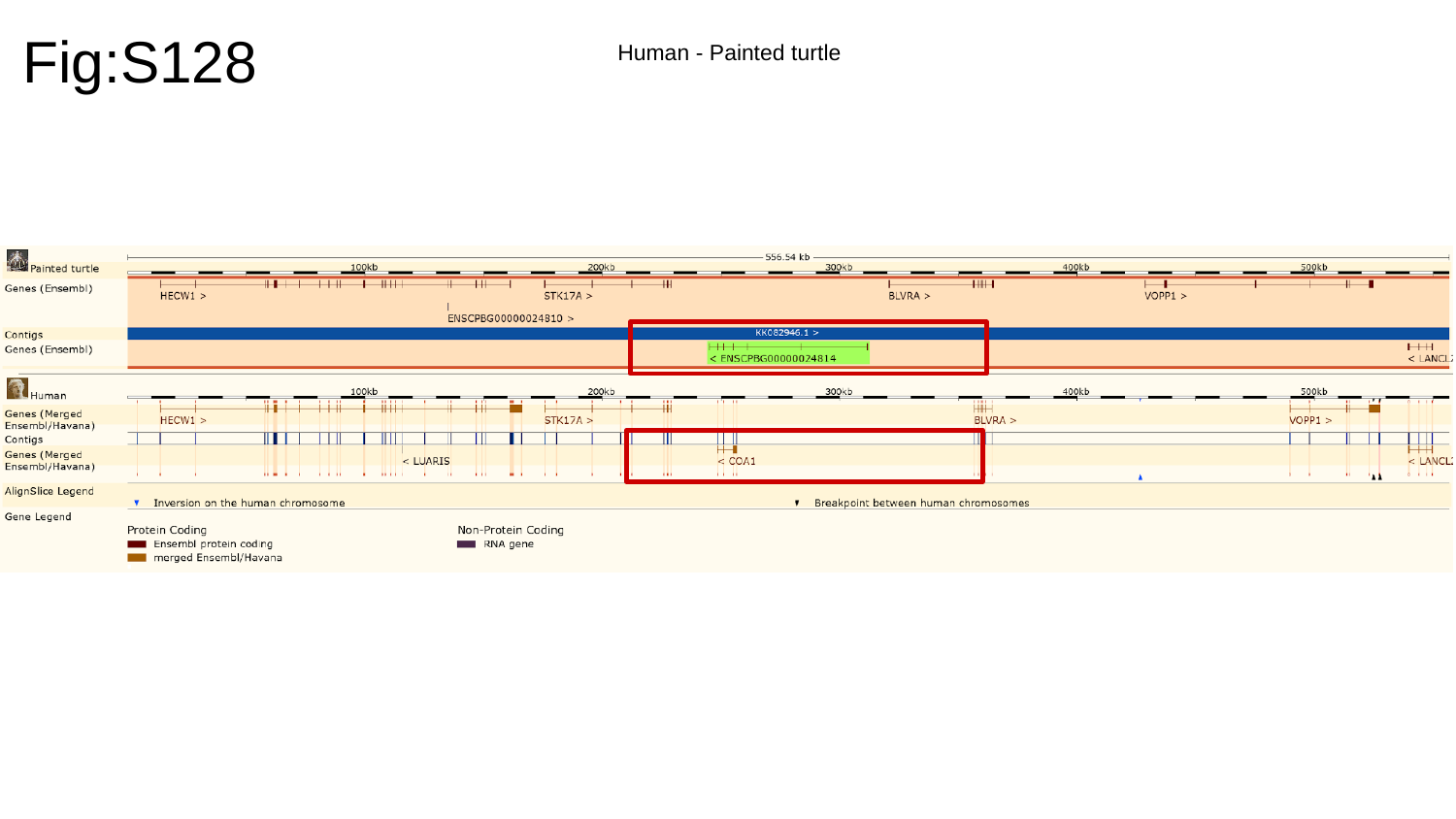

Fig:S128
Human - Painted turtle

### Slide 129
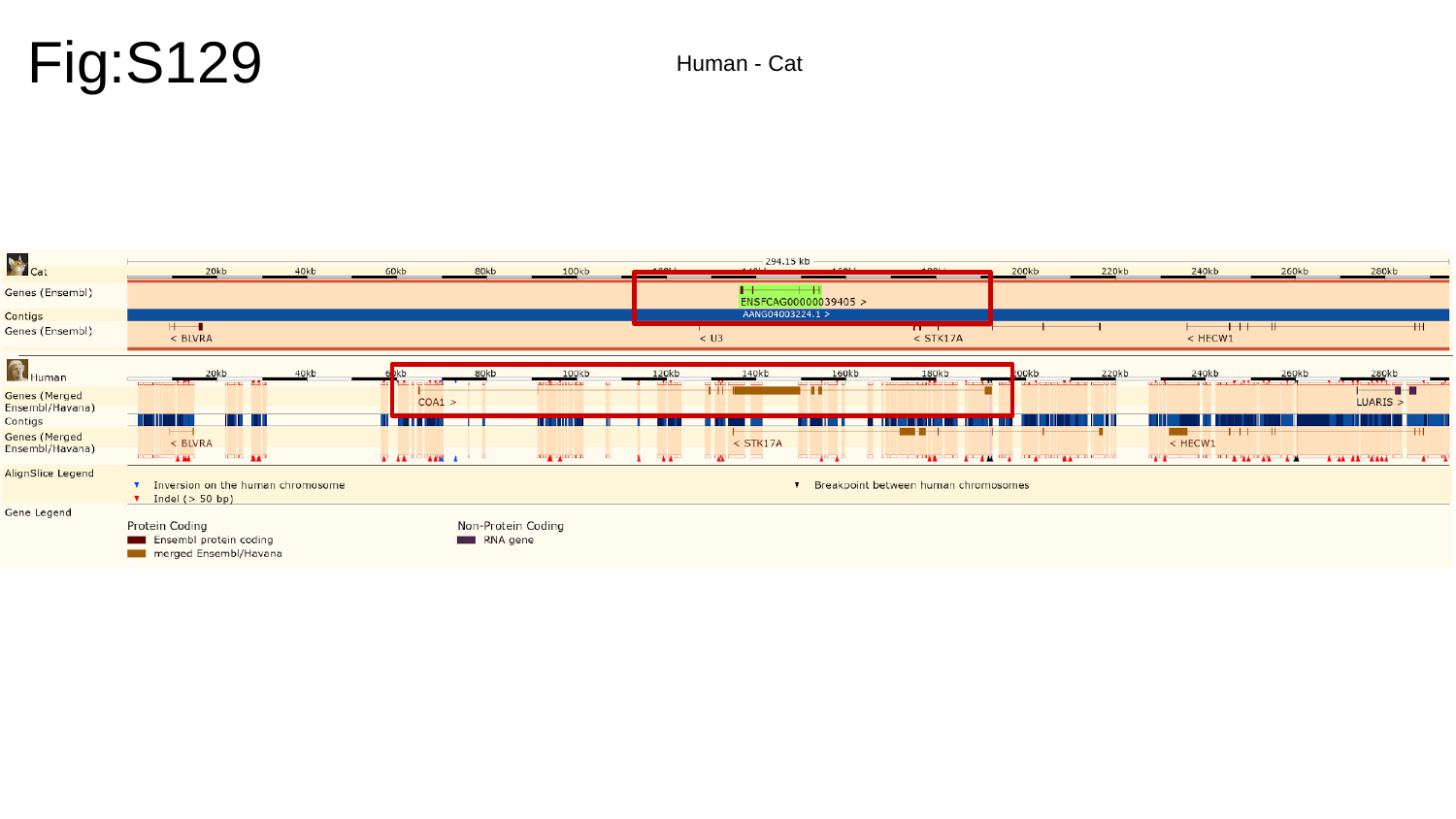

Fig:S129
Human - Cat

### Slide 130
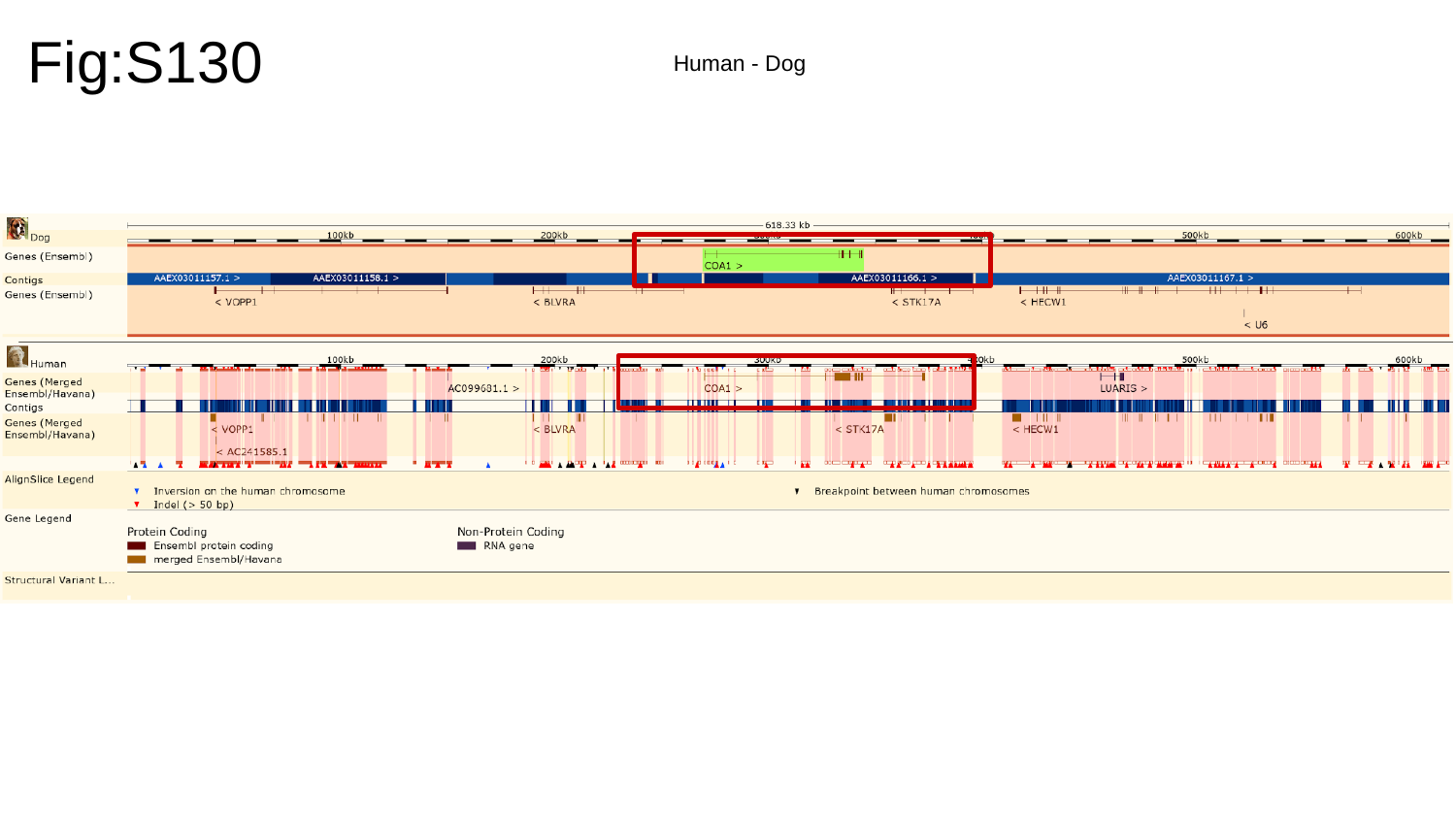

Fig:S130
Human - Dog

### Slide 131
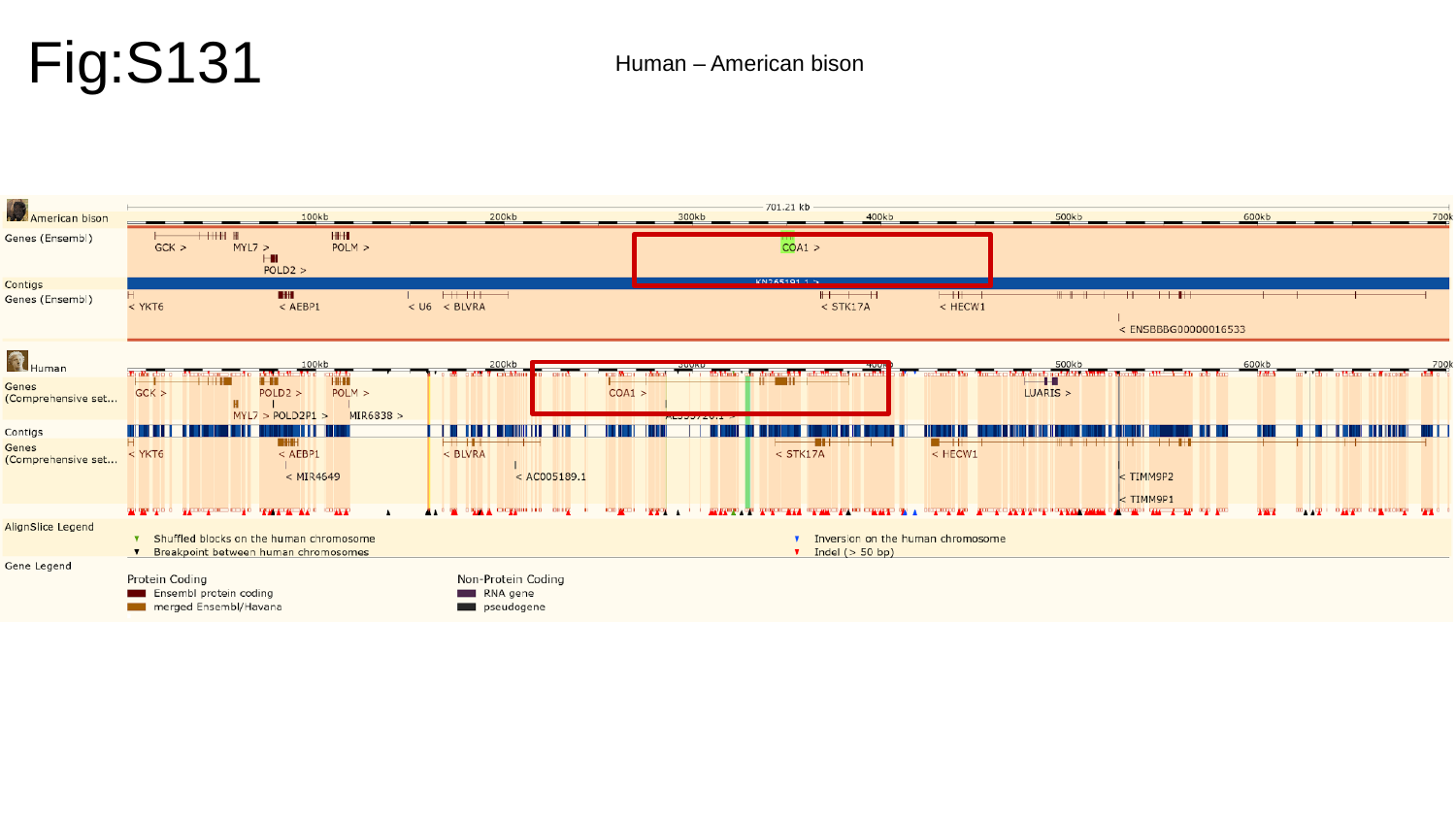

Fig:S131
Human – American bison

### Slide 132
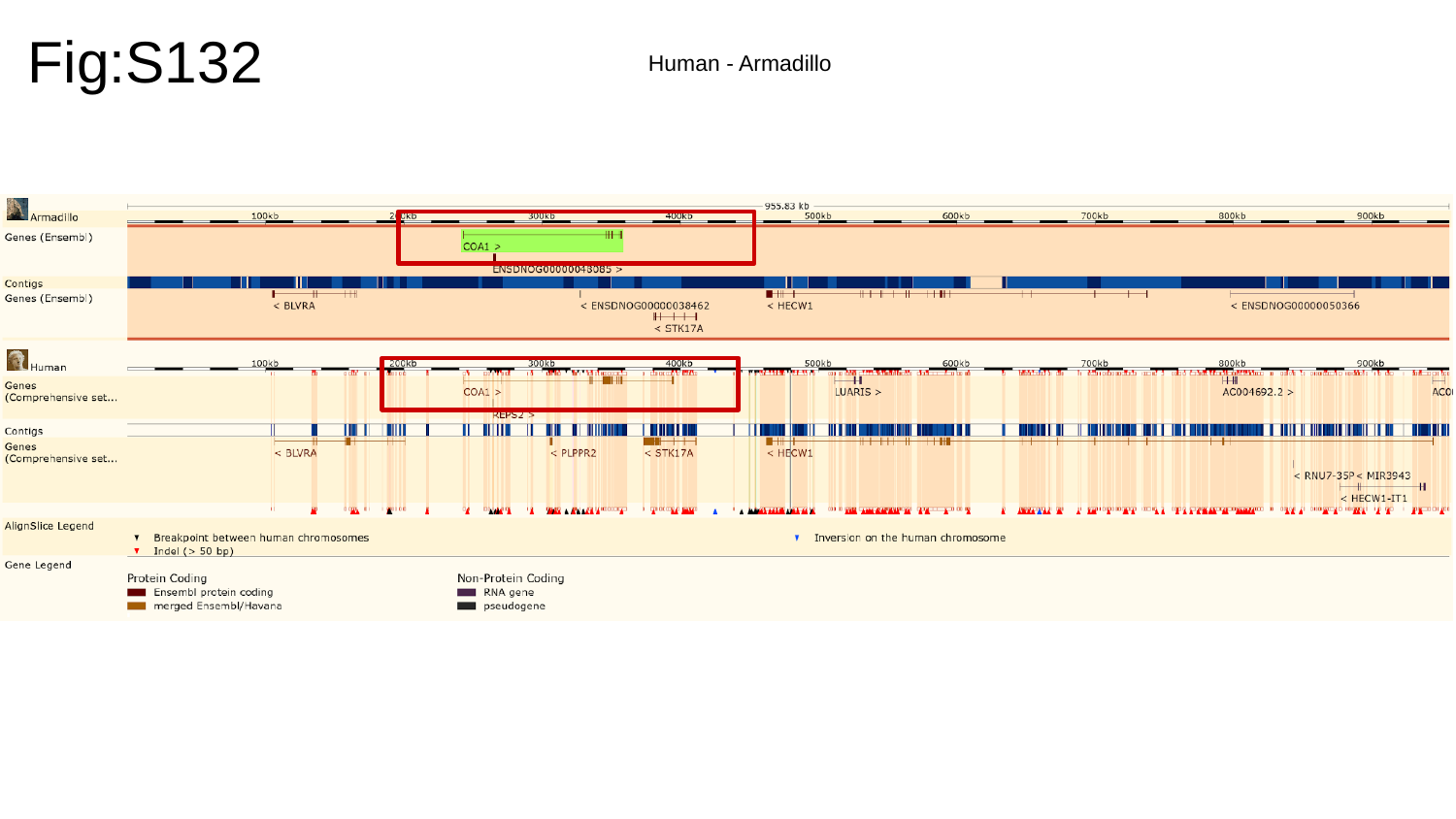

Fig:S132
Human - Armadillo

### Slide 133
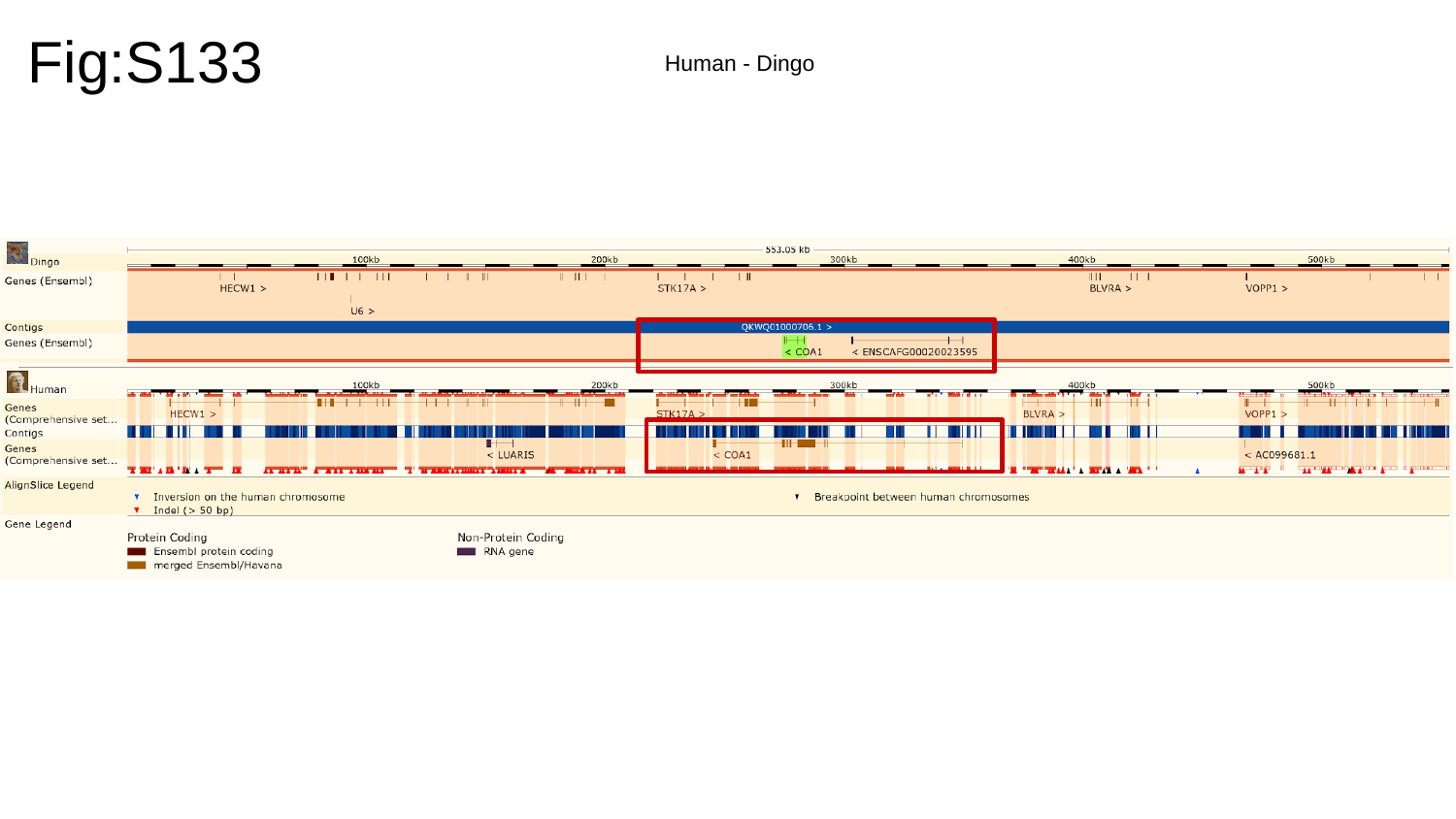

Fig:S133
Human - Dingo

### Slide 134
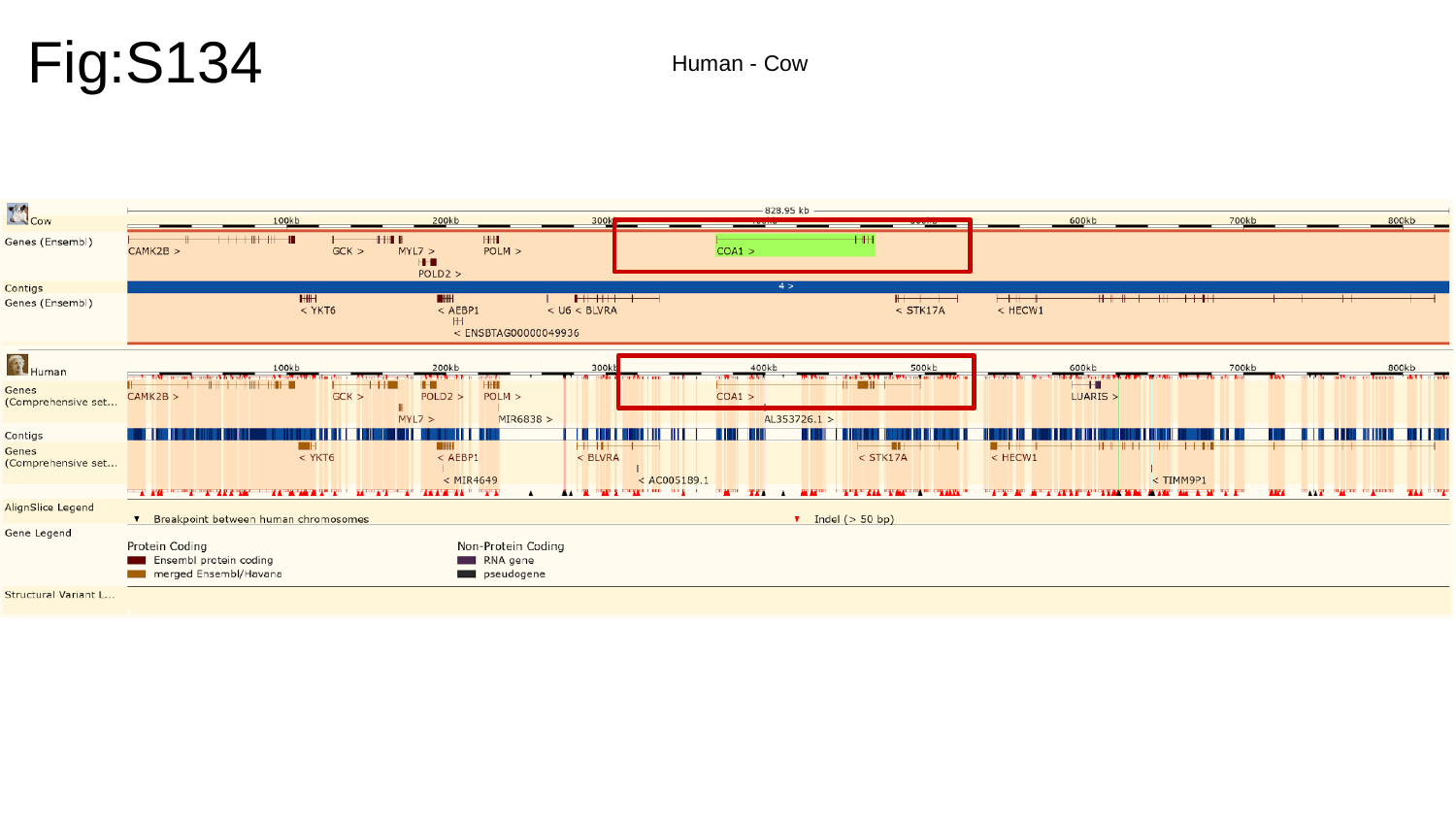

Fig:S134
Human - Cow

### Slide 135
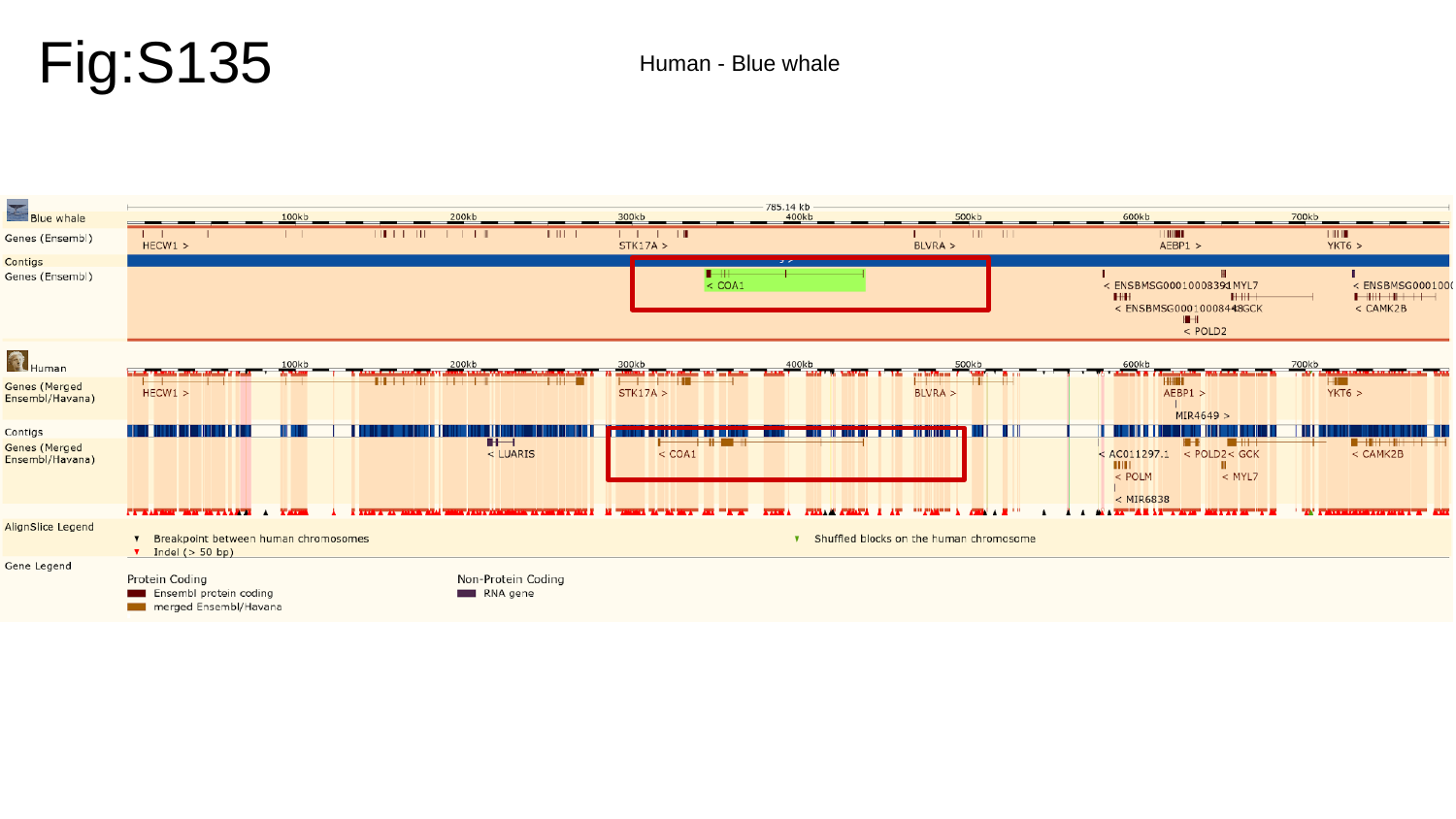

Fig:S135
Human - Blue whale

### Slide 136
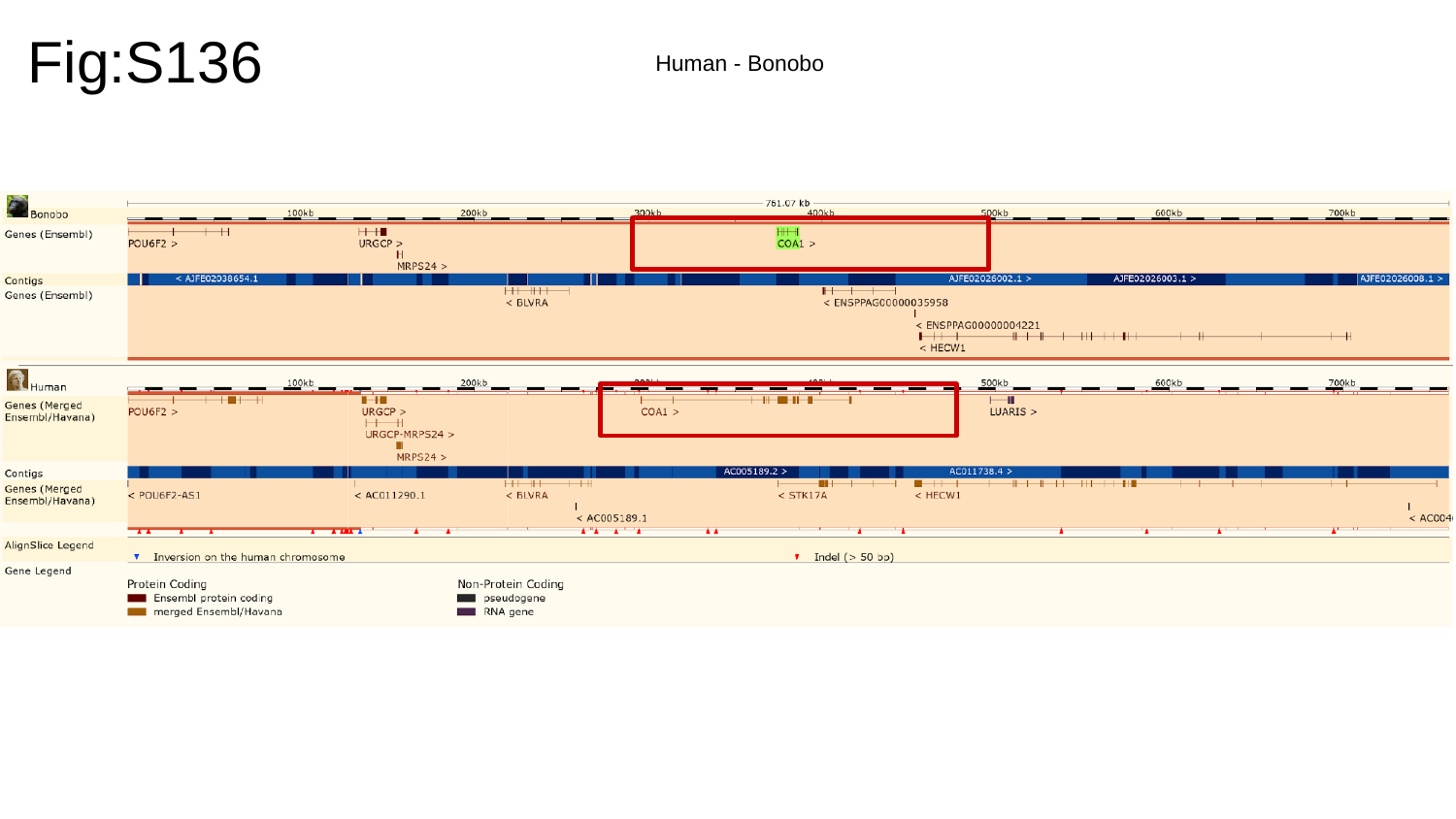

Fig:S136
Human - Bonobo

### Slide 137
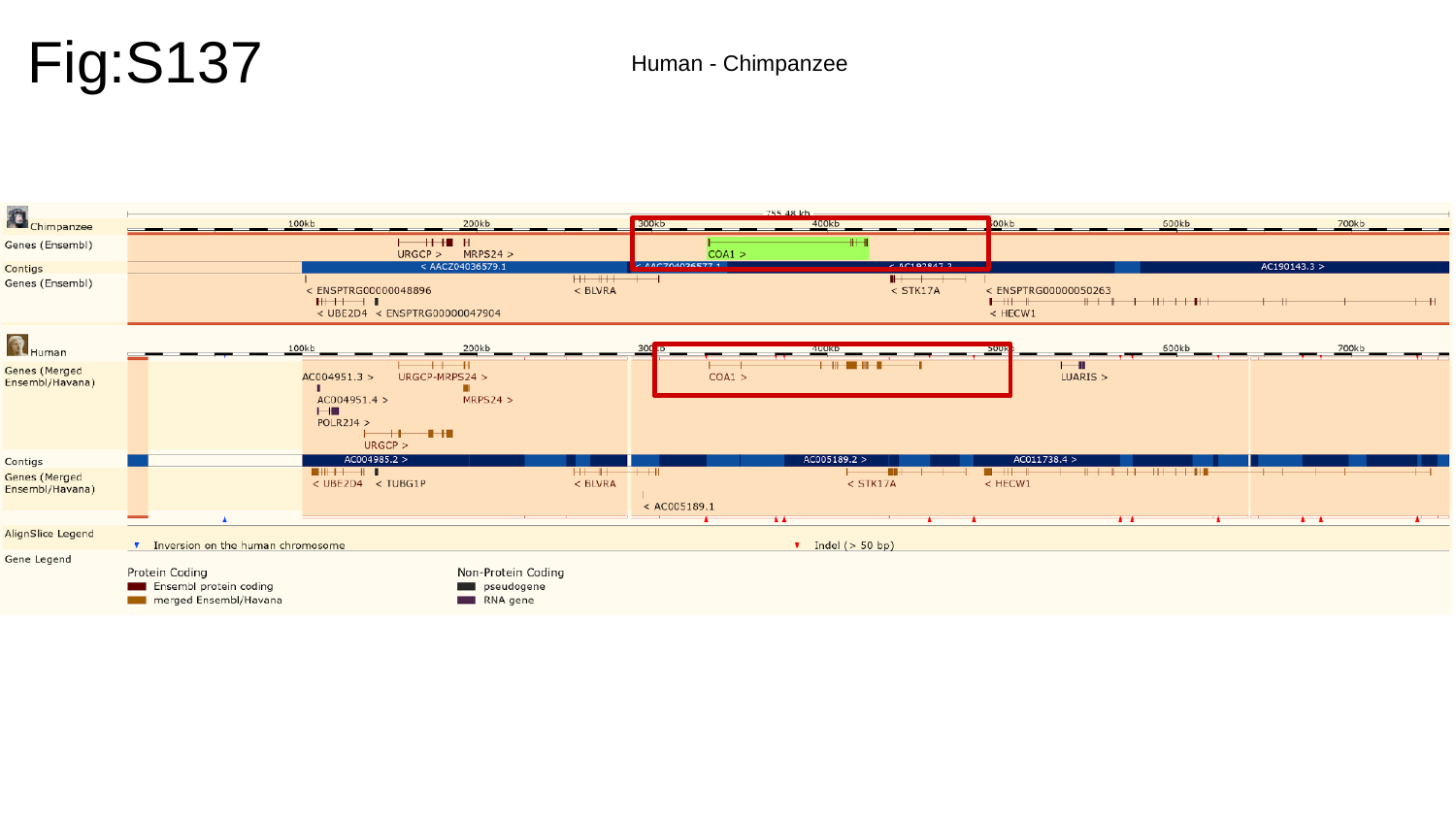

Fig:S137
Human - Chimpanzee

### Slide 138
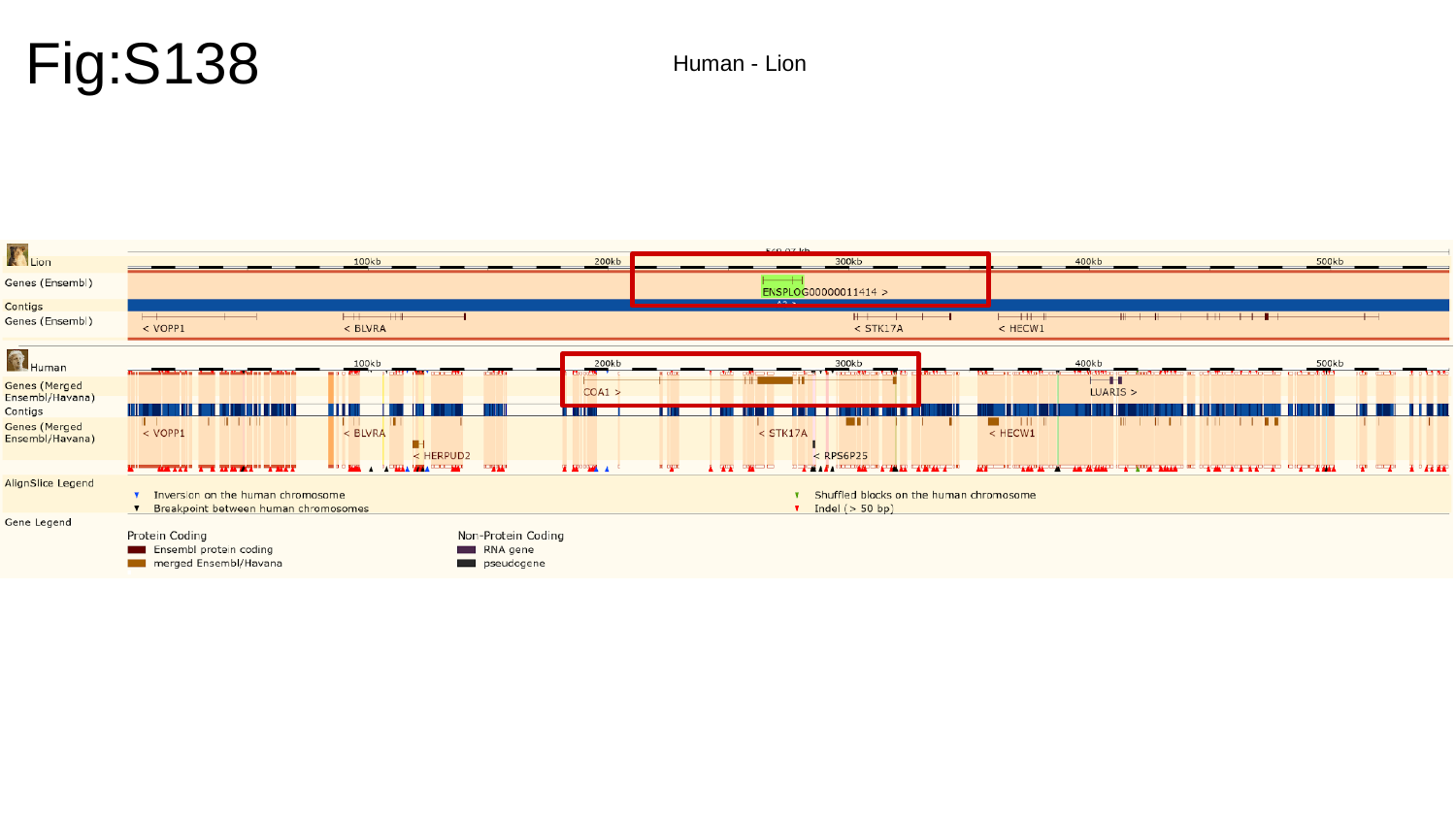

Fig:S138
Human - Lion

### Slide 139
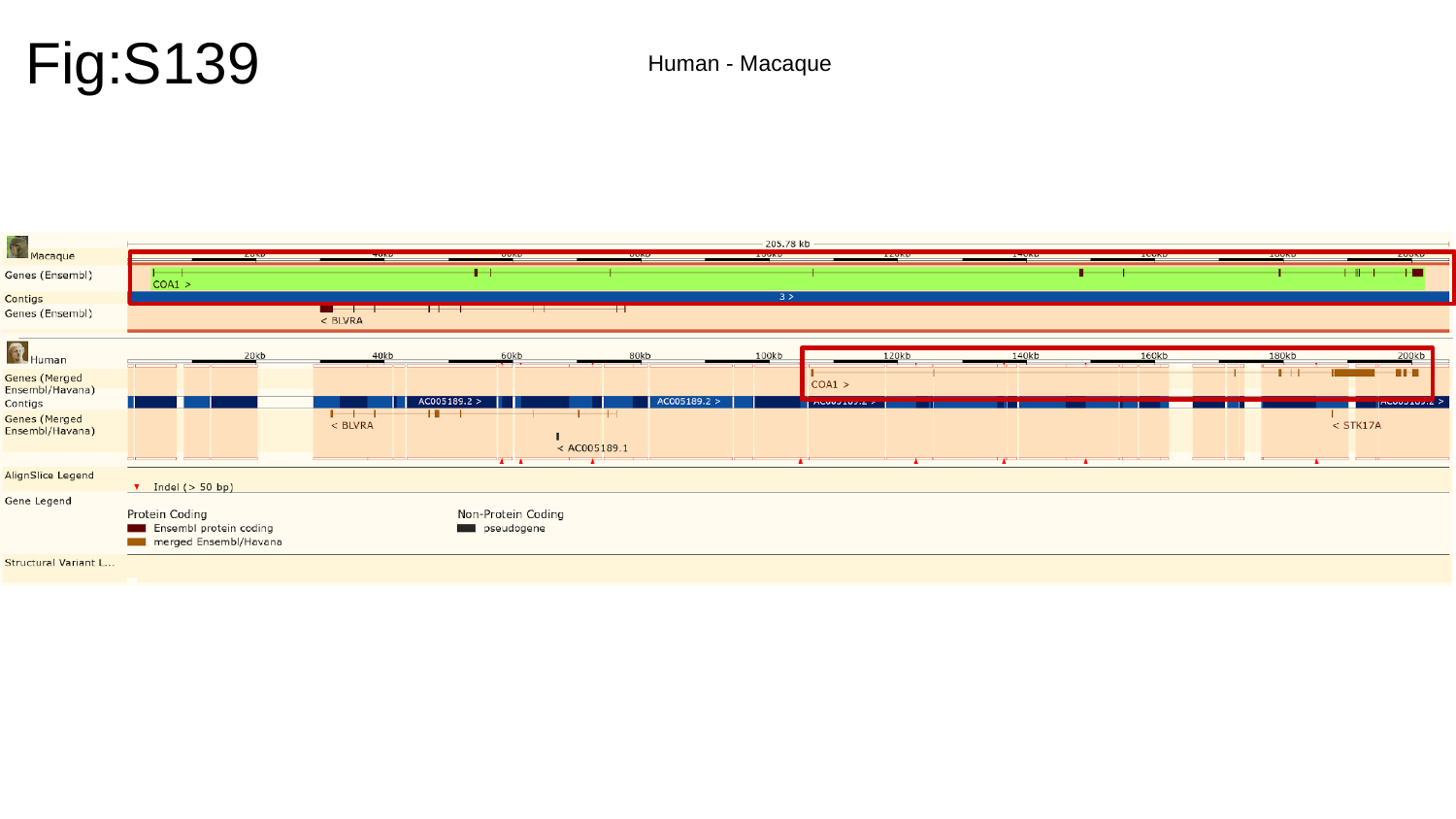

Fig:S139
Human - Macaque

### Slide 140
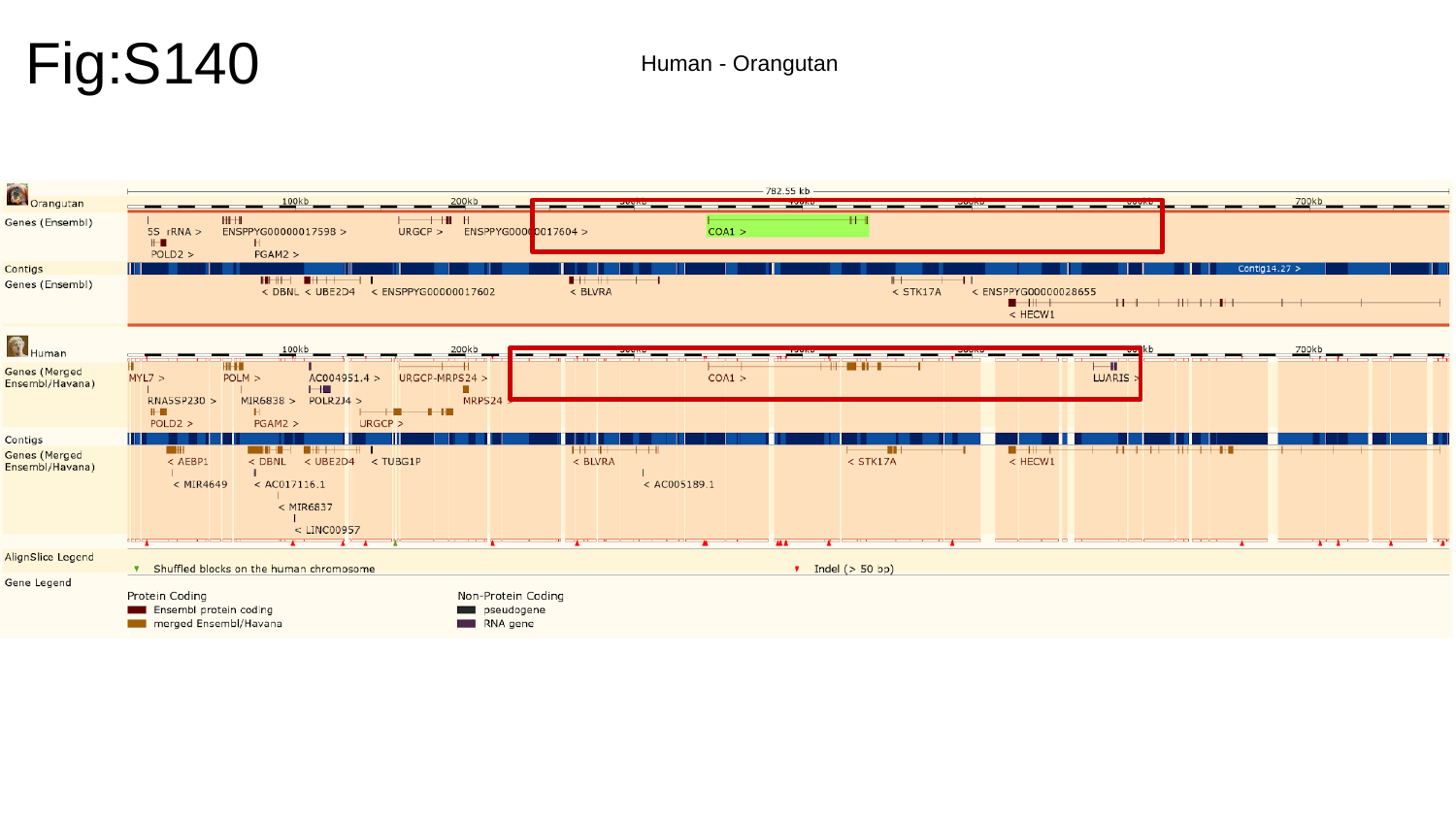

Fig:S140
Human - Orangutan

### Slide 141
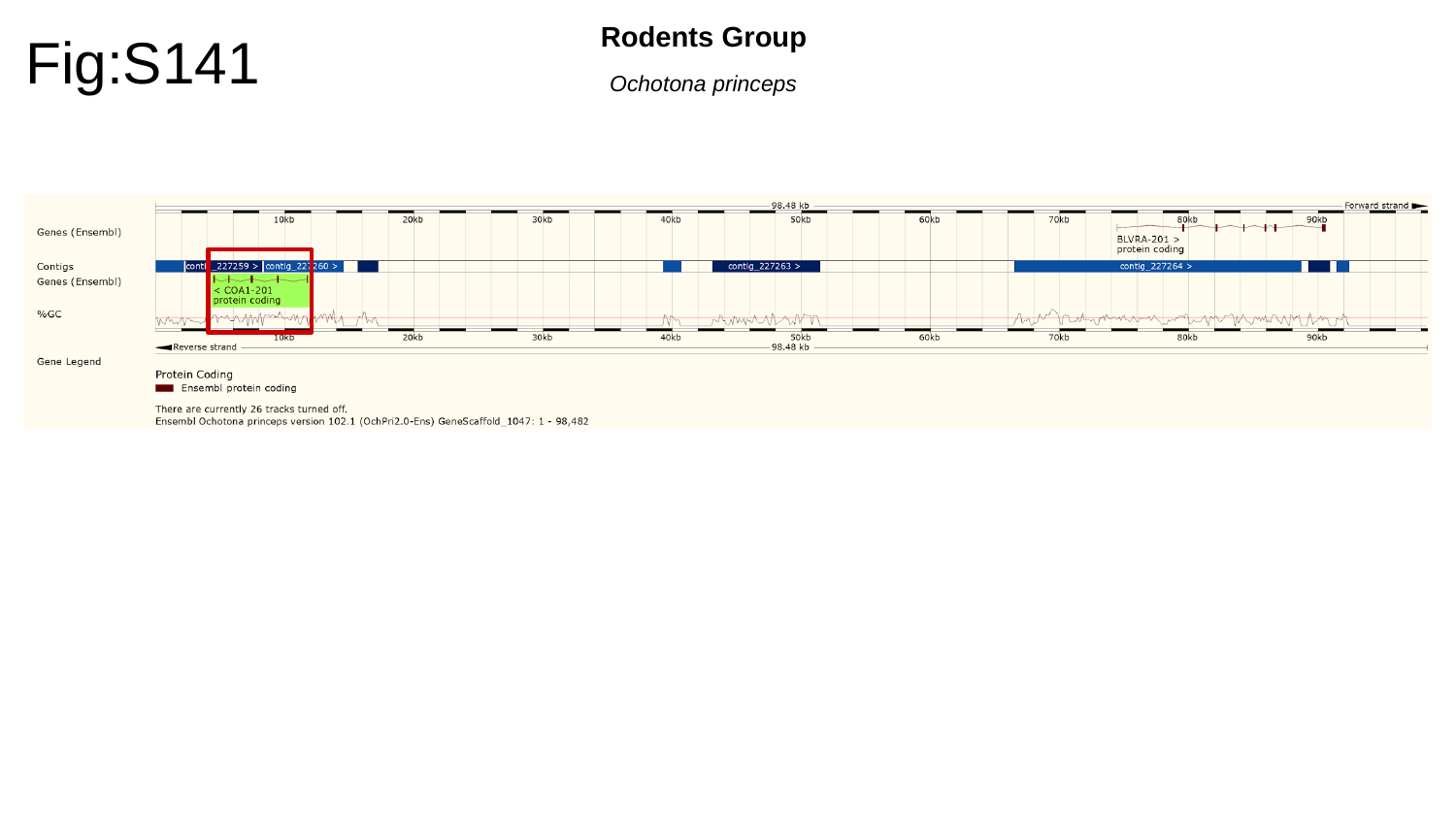

Fig:S141
Rodents Group
Ochotona princeps

### Slide 142
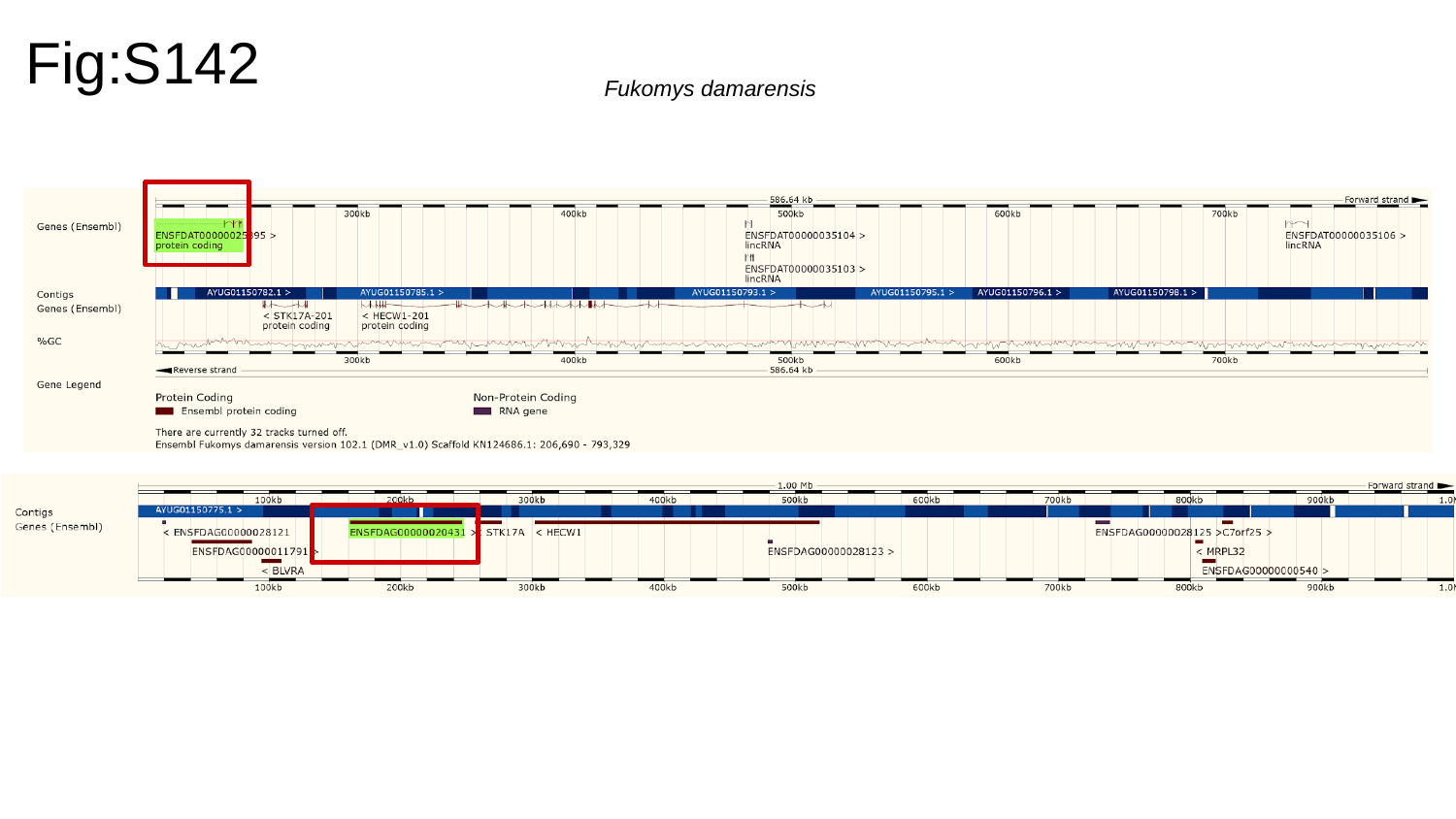

Fig:S142
Fukomys damarensis

### Slide 143
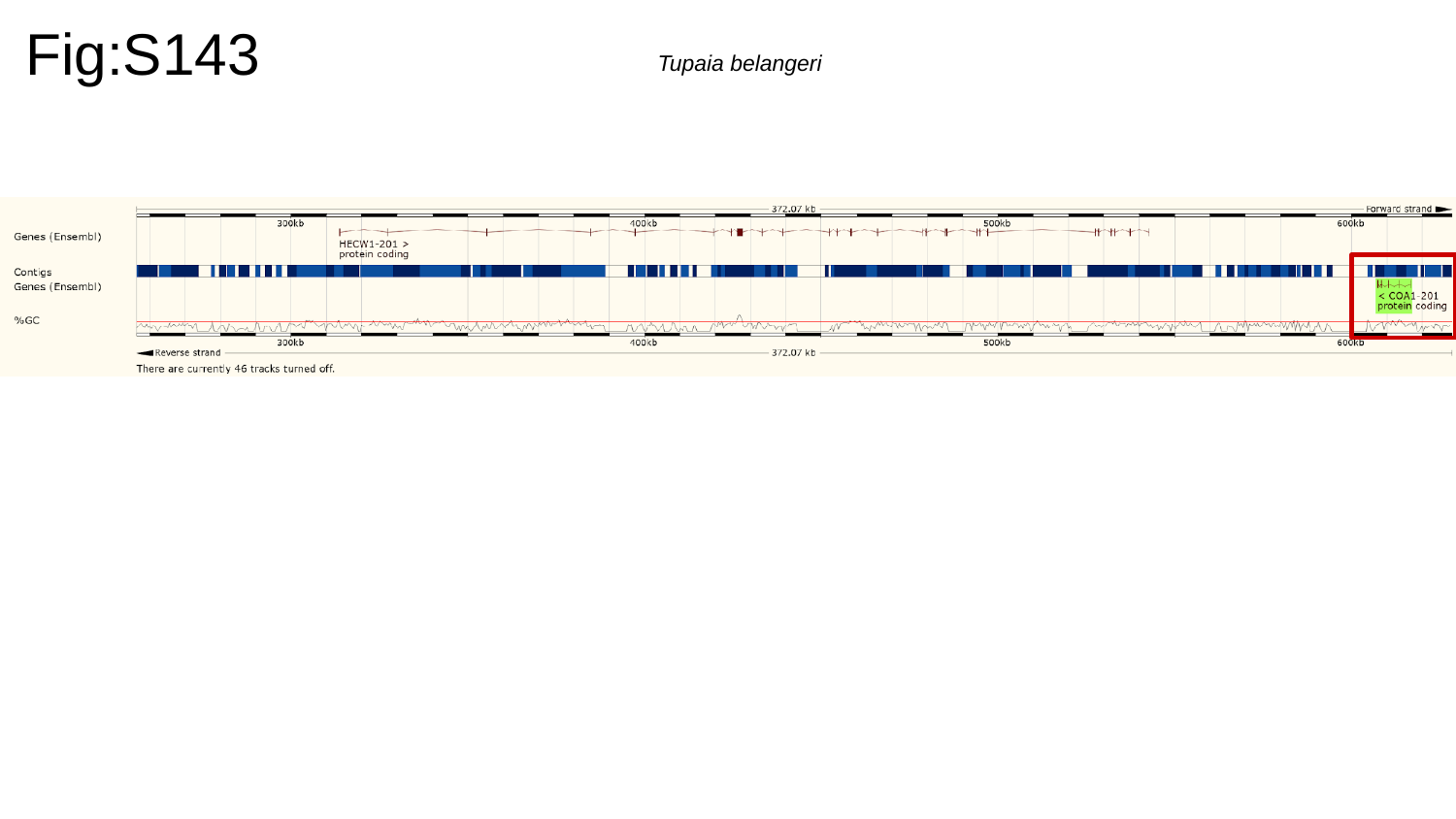

Fig:S143
Tupaia belangeri

### Slide 144
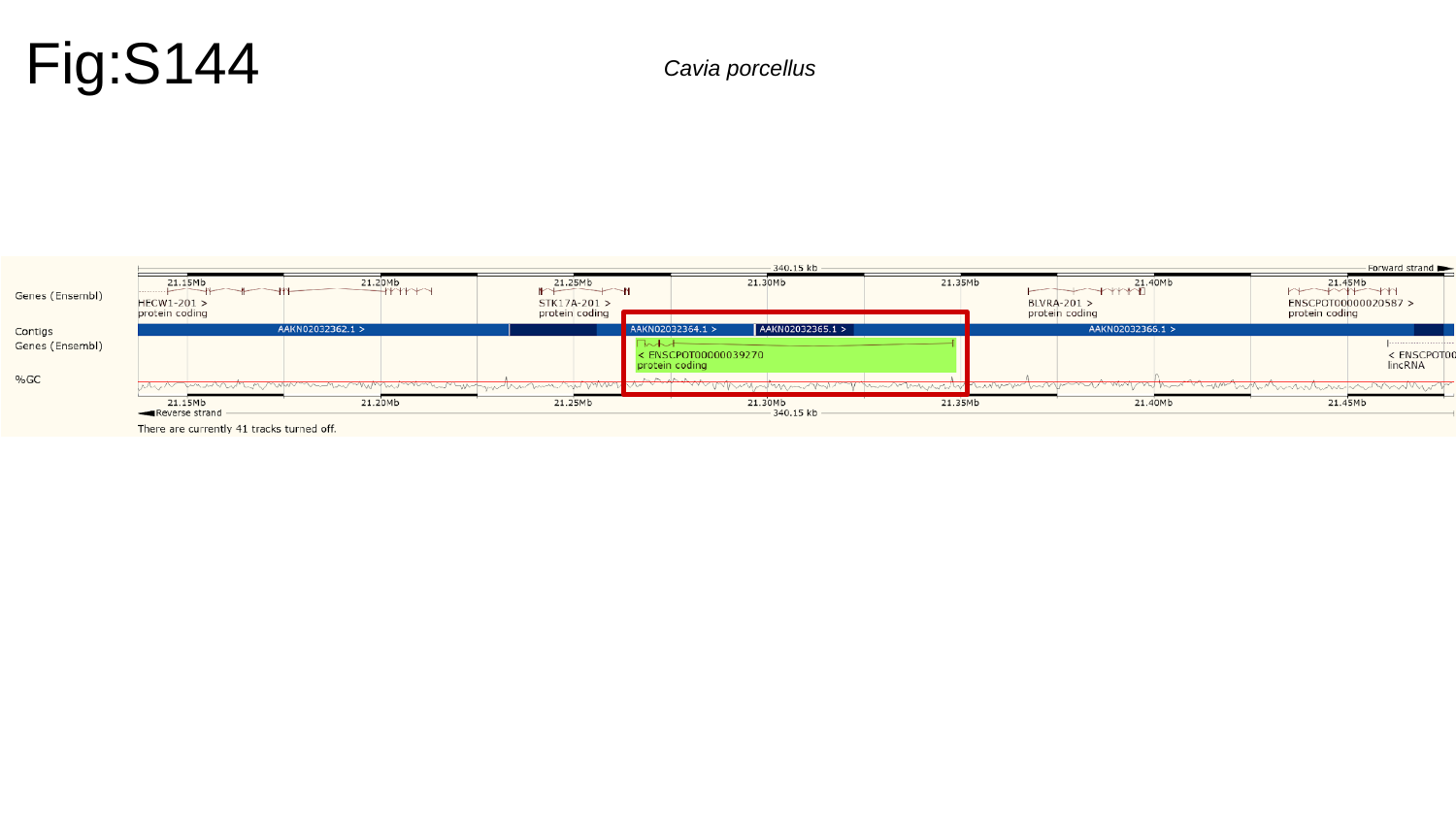

Fig:S144
Cavia porcellus

### Slide 145
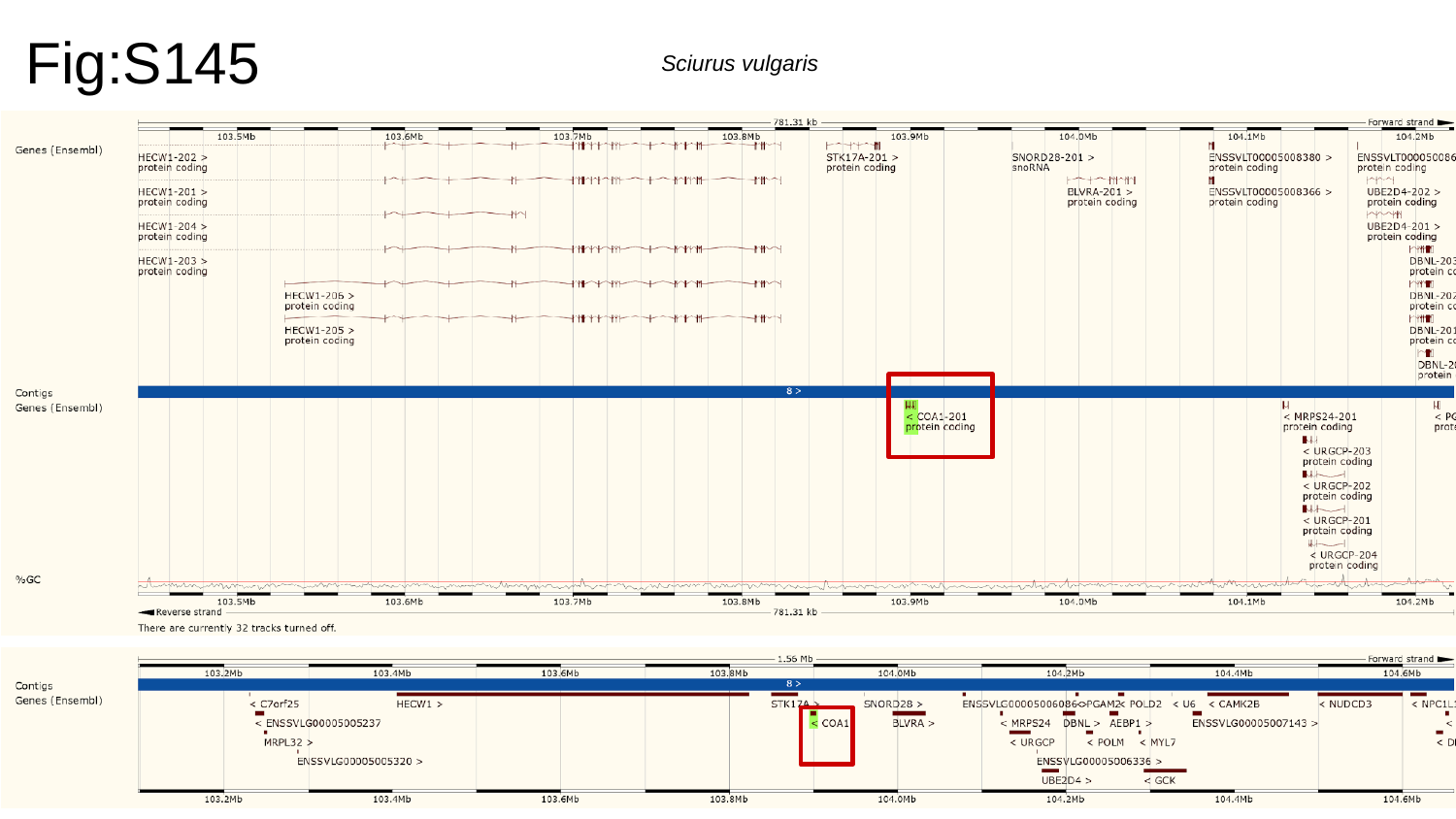

Fig:S145
Sciurus vulgaris

### Slide 146
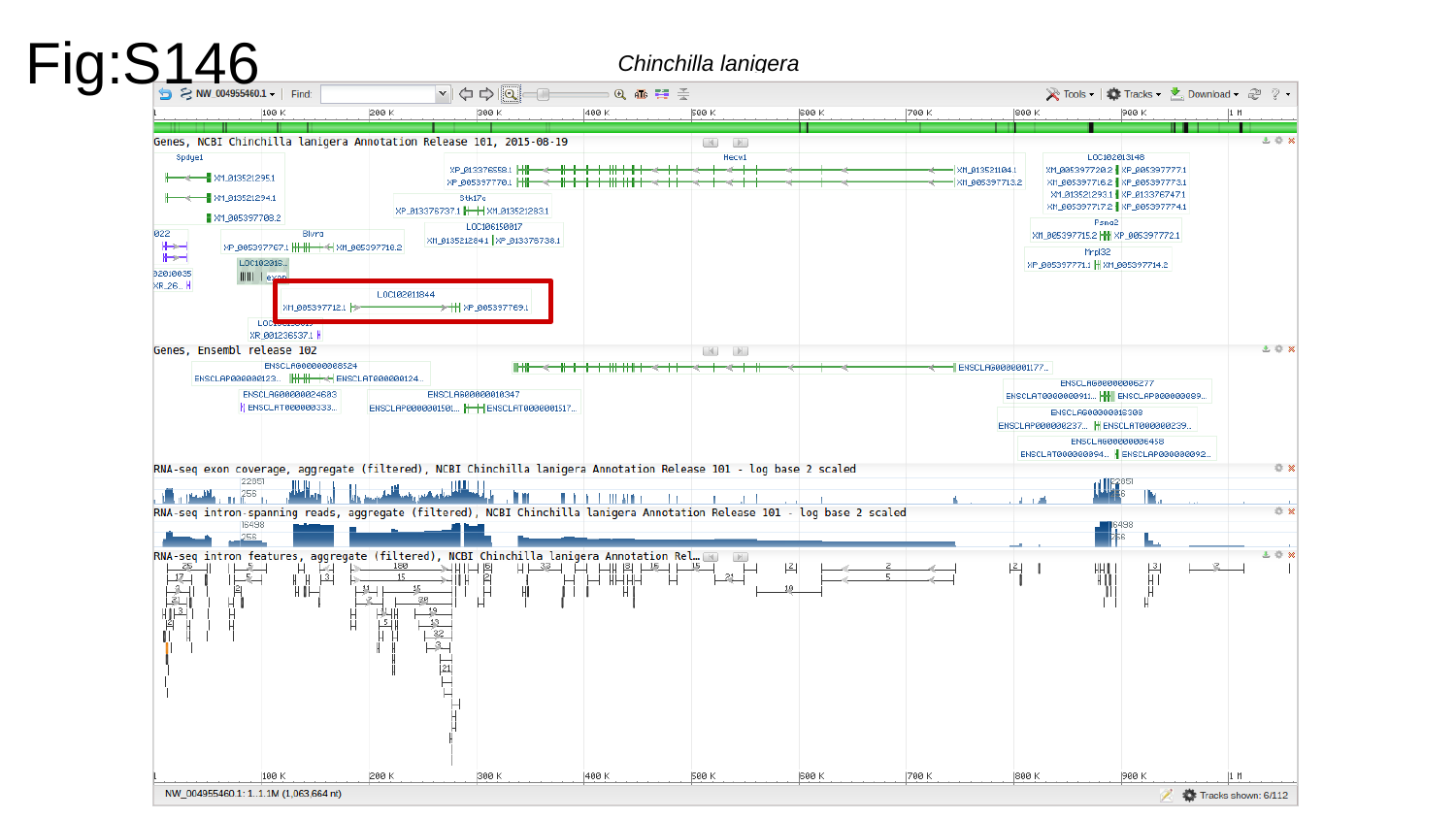

Fig:S146
Chinchilla lanigera

### Slide 147
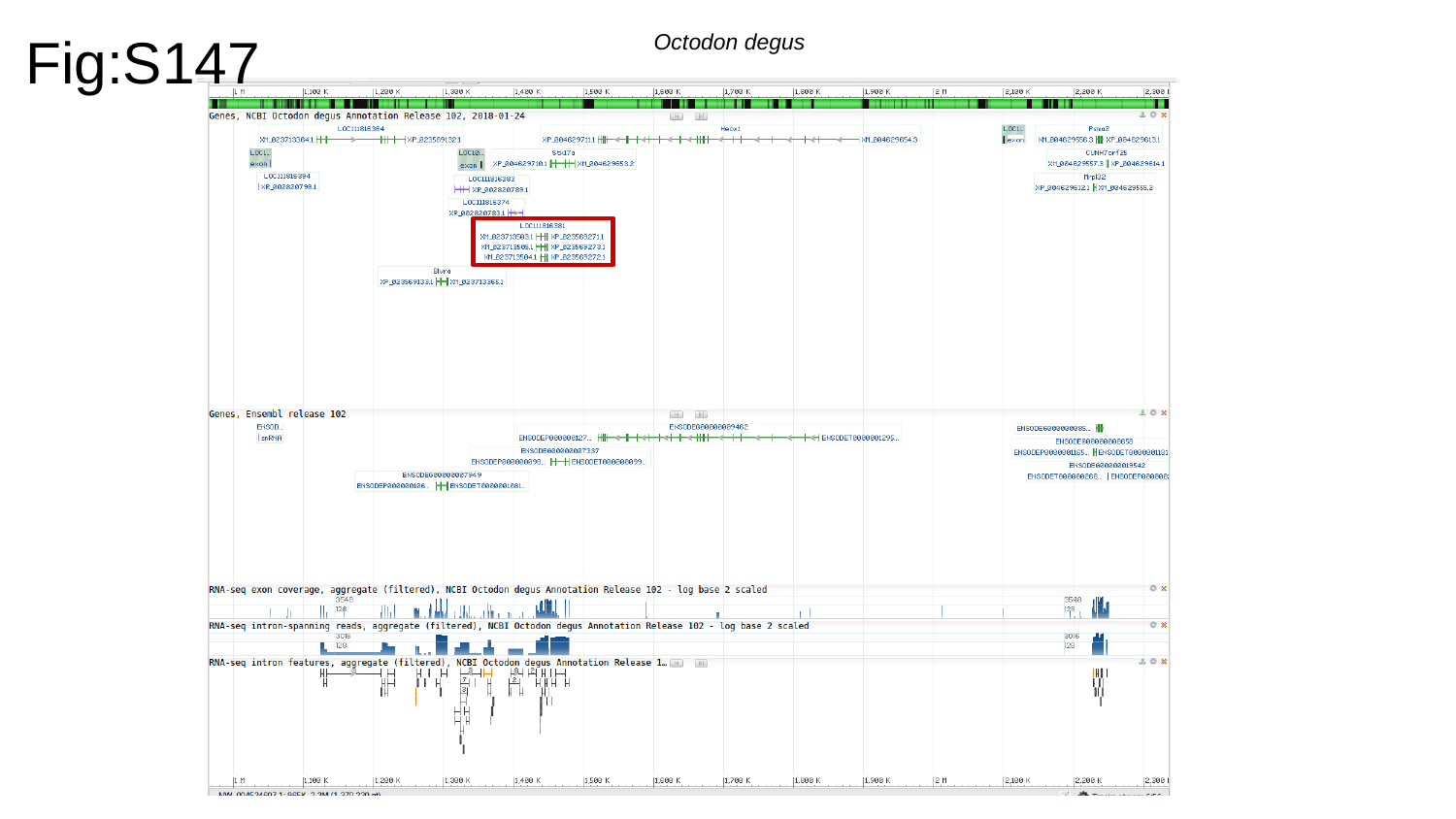

Fig:S147
Octodon degus

### Slide 148
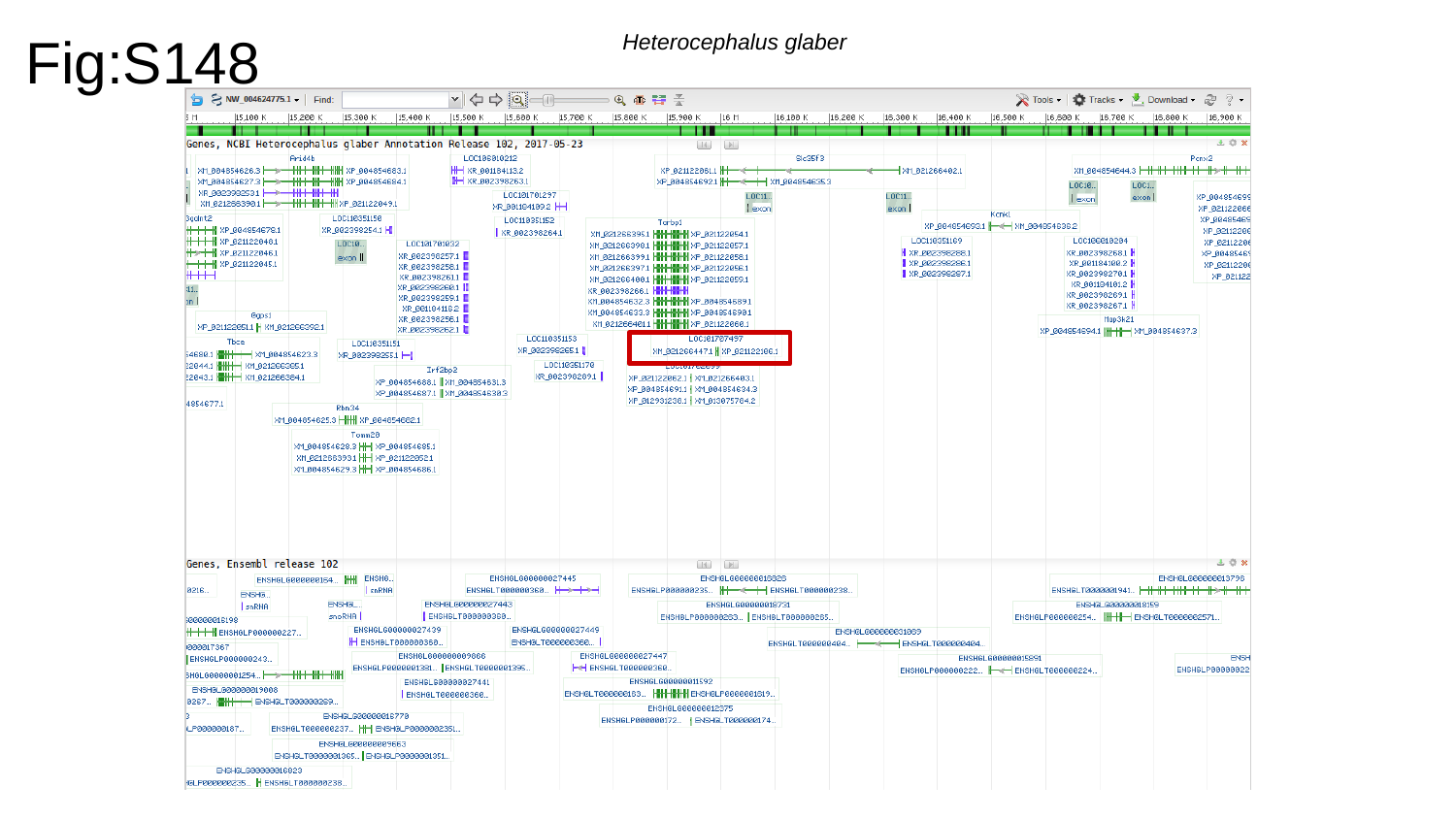

Fig:S148
Heterocephalus glaber

### Slide 149
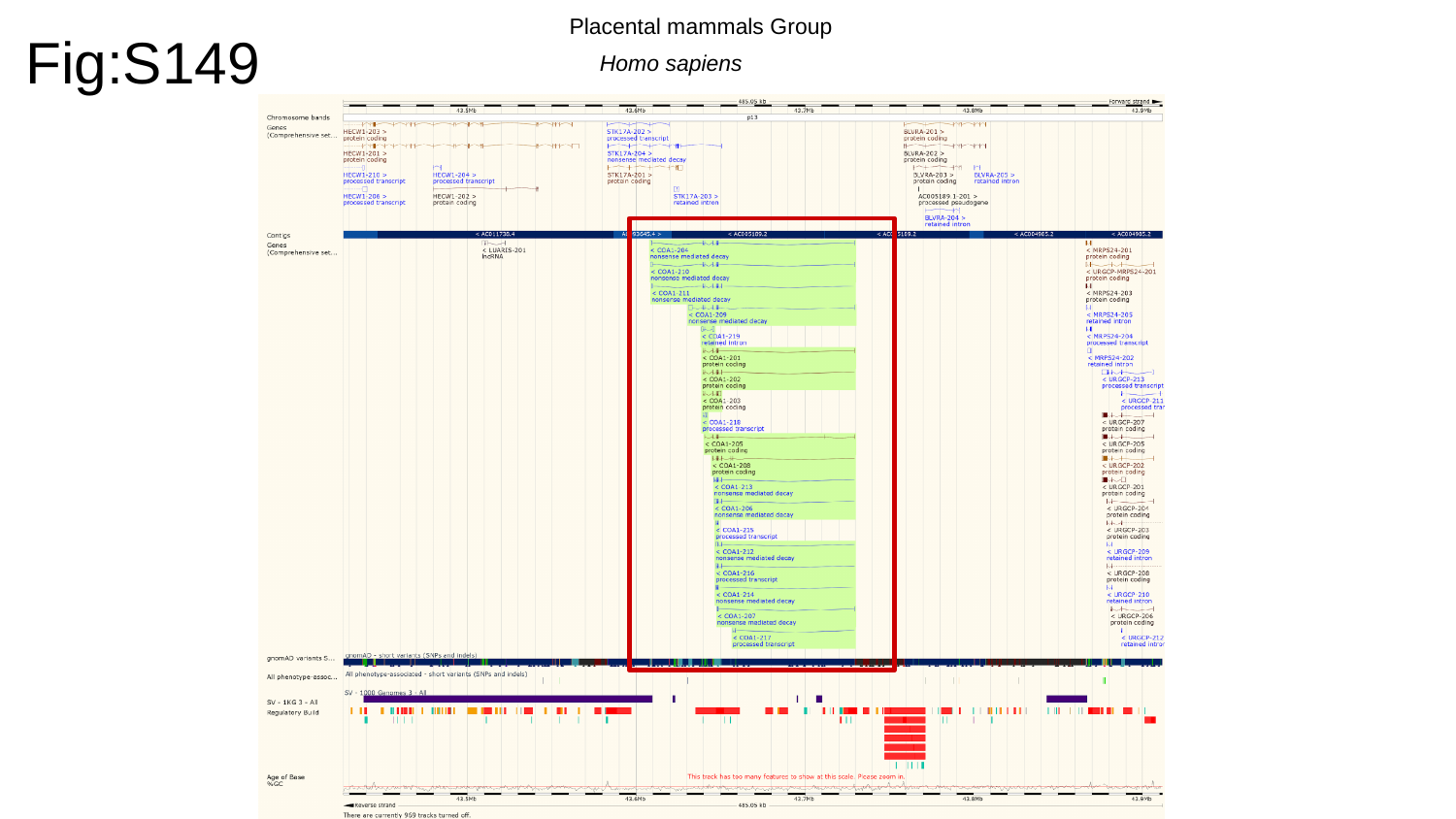

Fig:S149
Placental mammals Group
Homo sapiens

### Slide 150
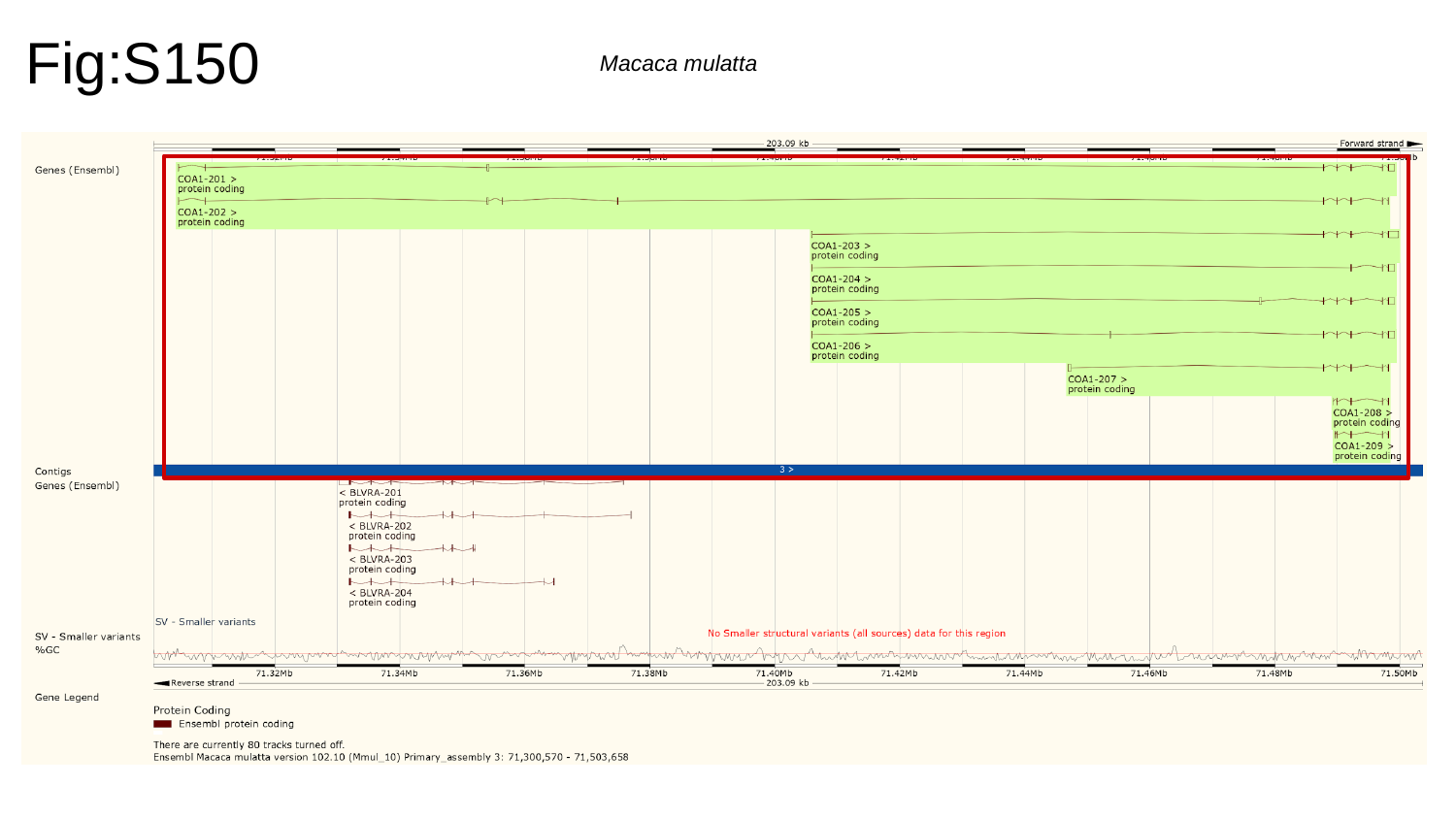

Fig:S150
Macaca mulatta

### Slide 151
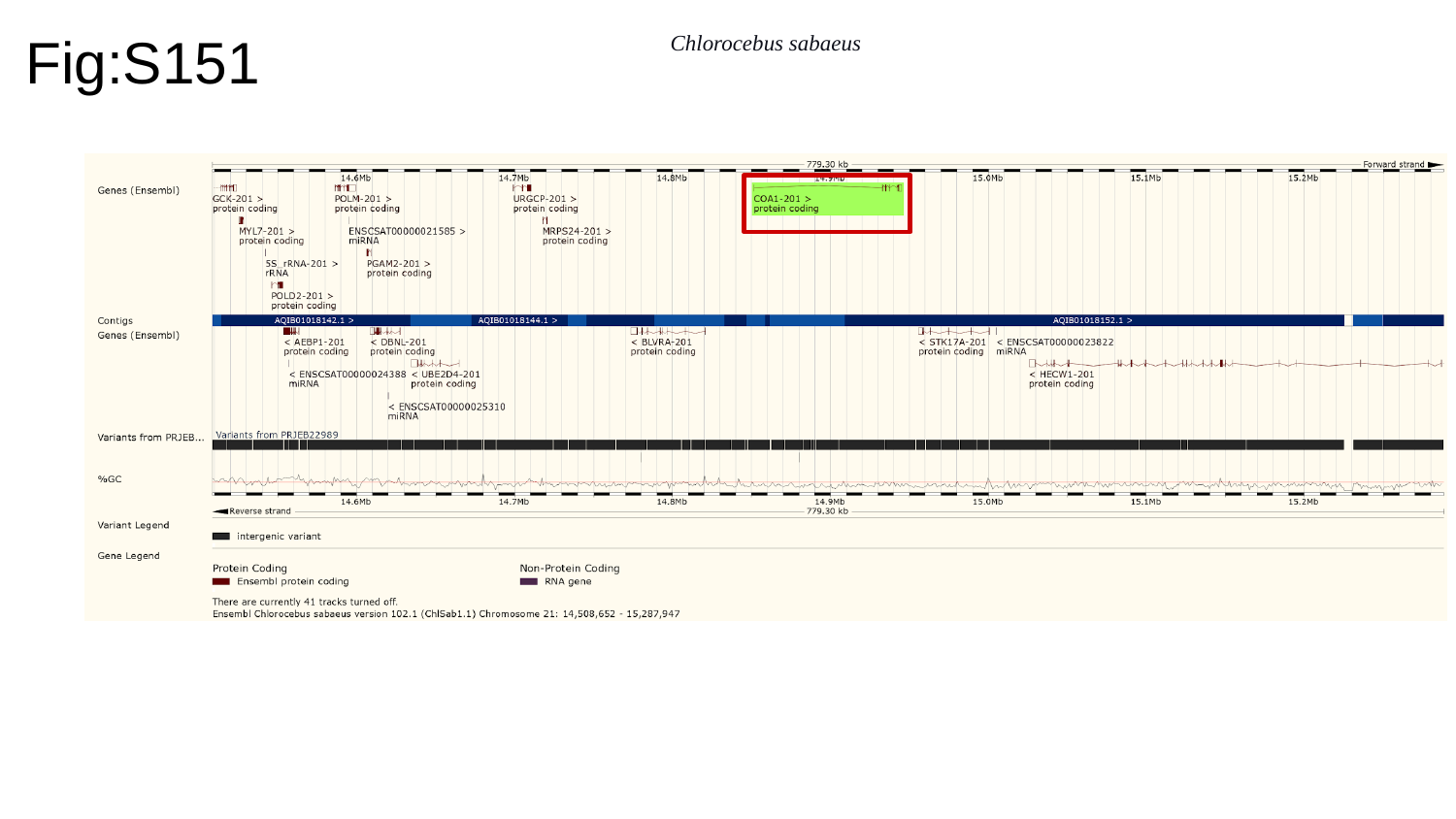

Fig:S151
Chlorocebus sabaeus

### Slide 152
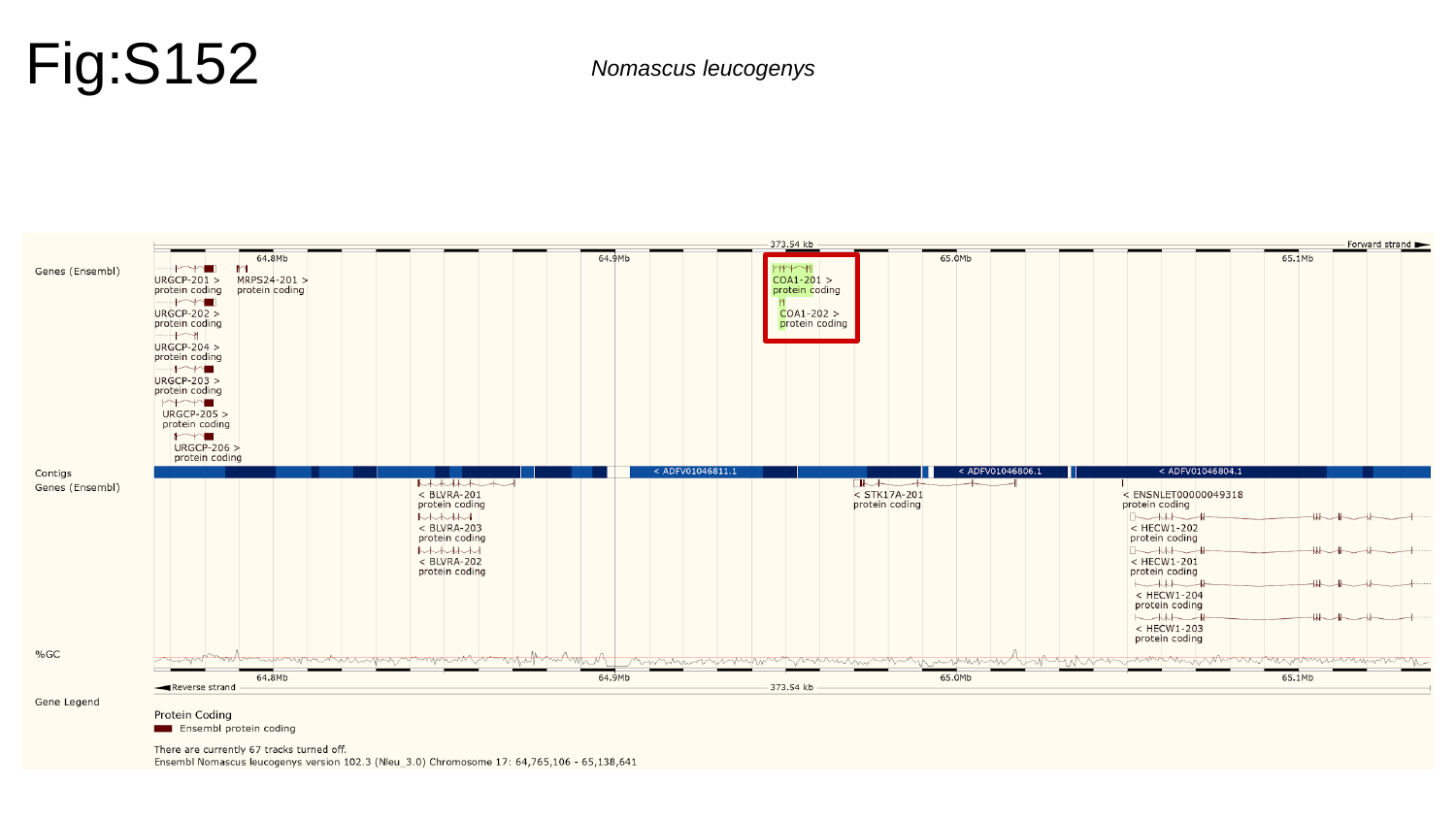

Fig:S152
Nomascus leucogenys

### Slide 153
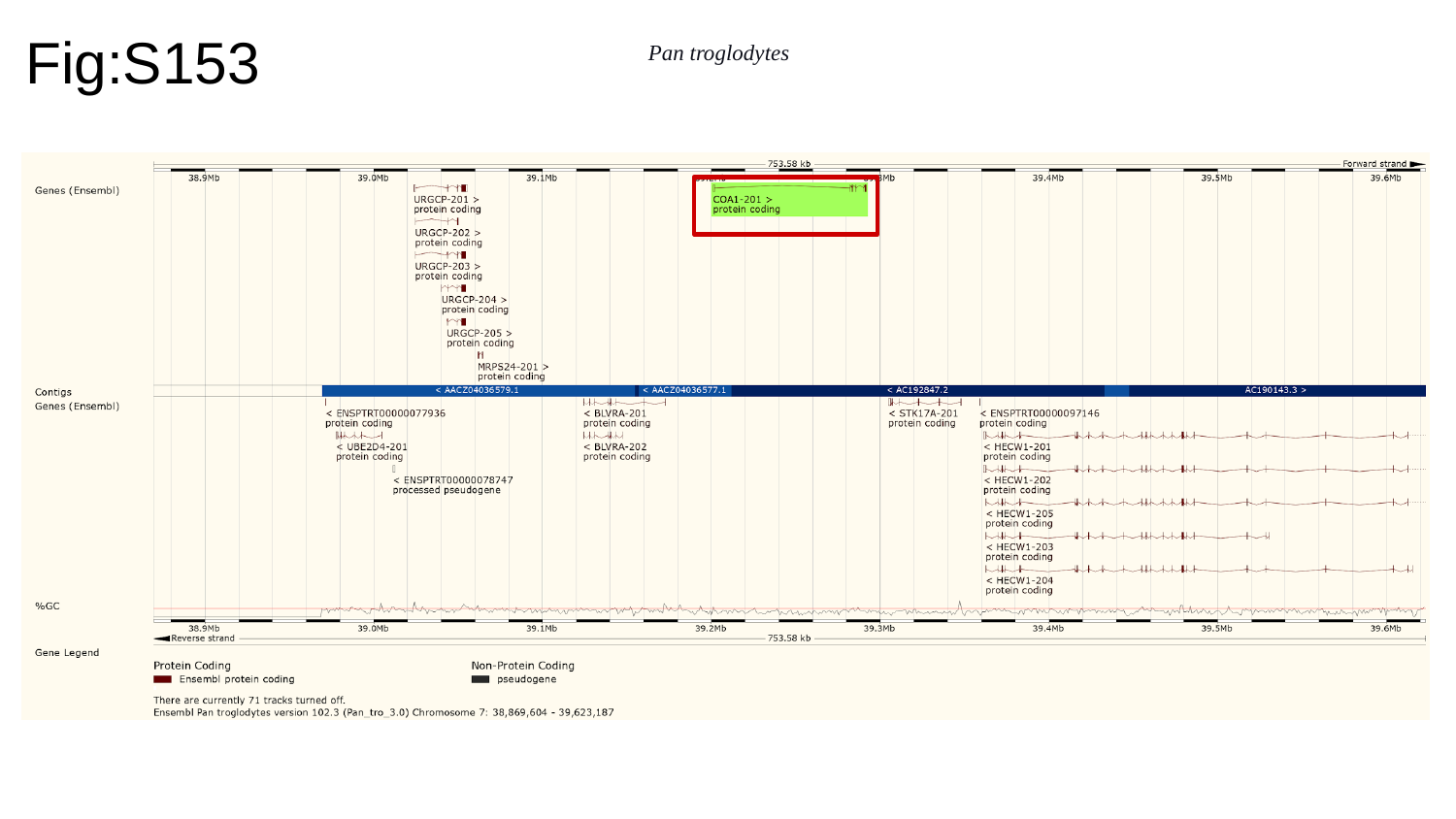

Fig:S153
Pan troglodytes

### Slide 154
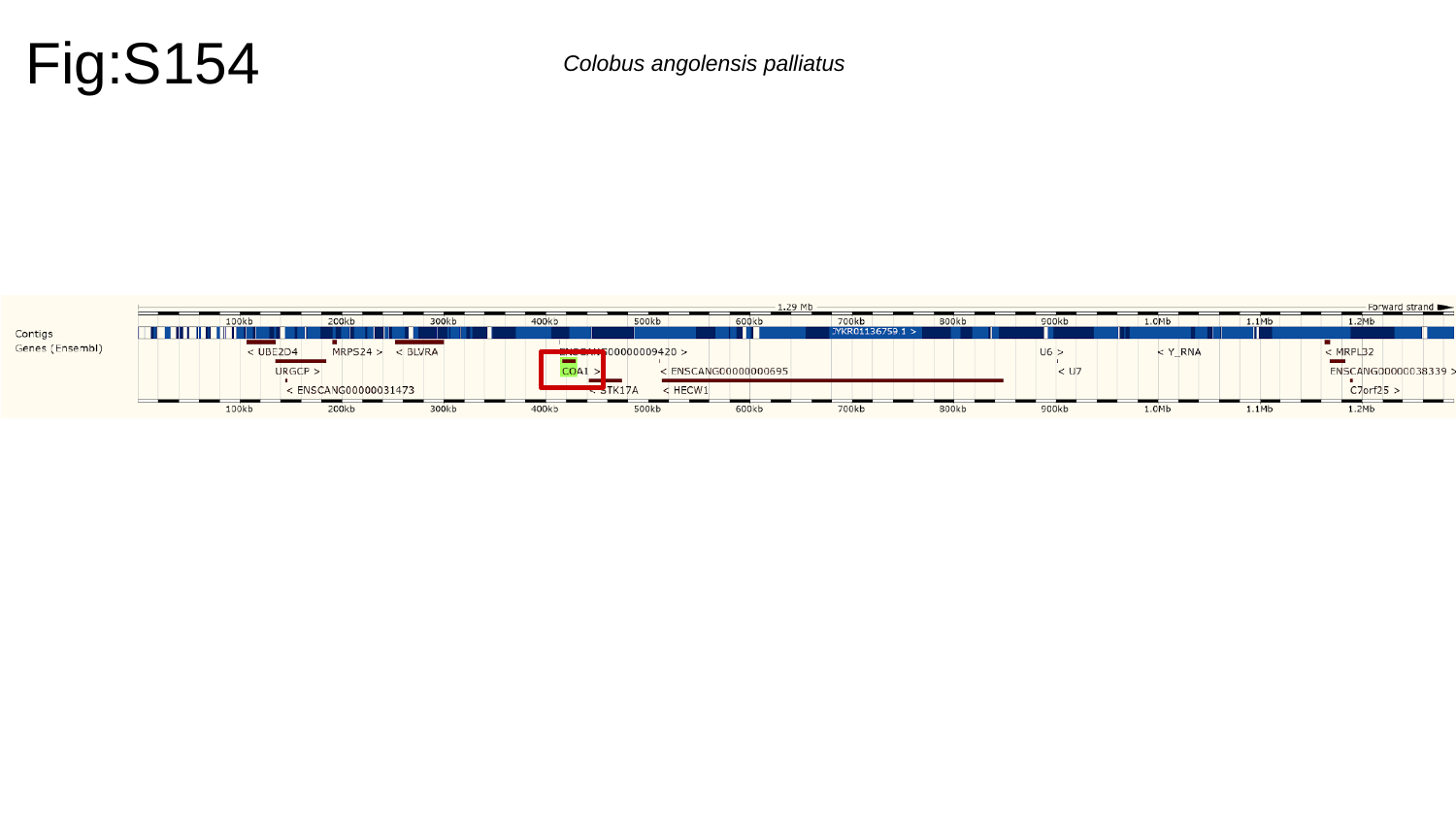

Fig:S154
Colobus angolensis palliatus

### Slide 155
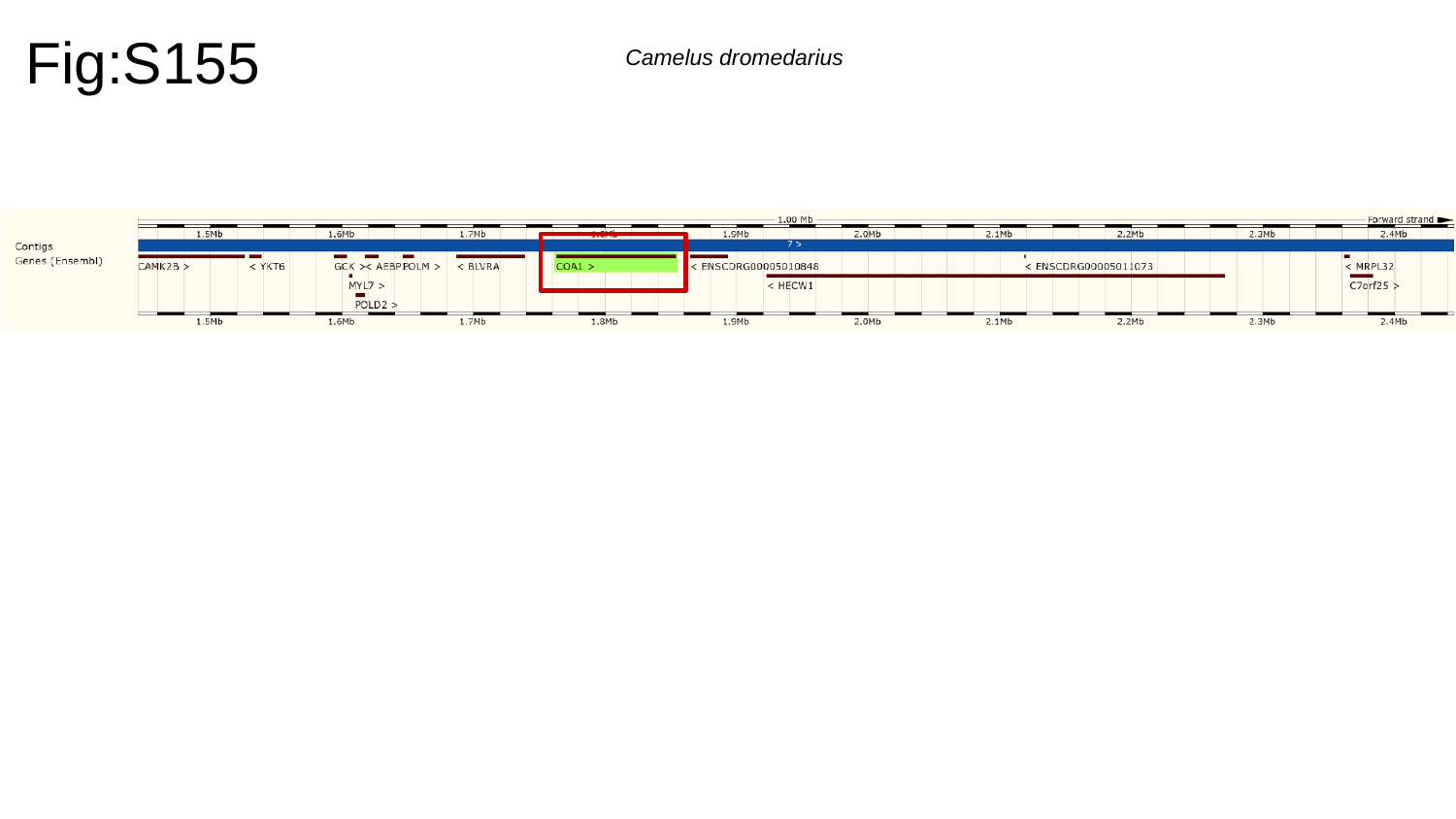

Fig:S155
Camelus dromedarius

### Slide 156
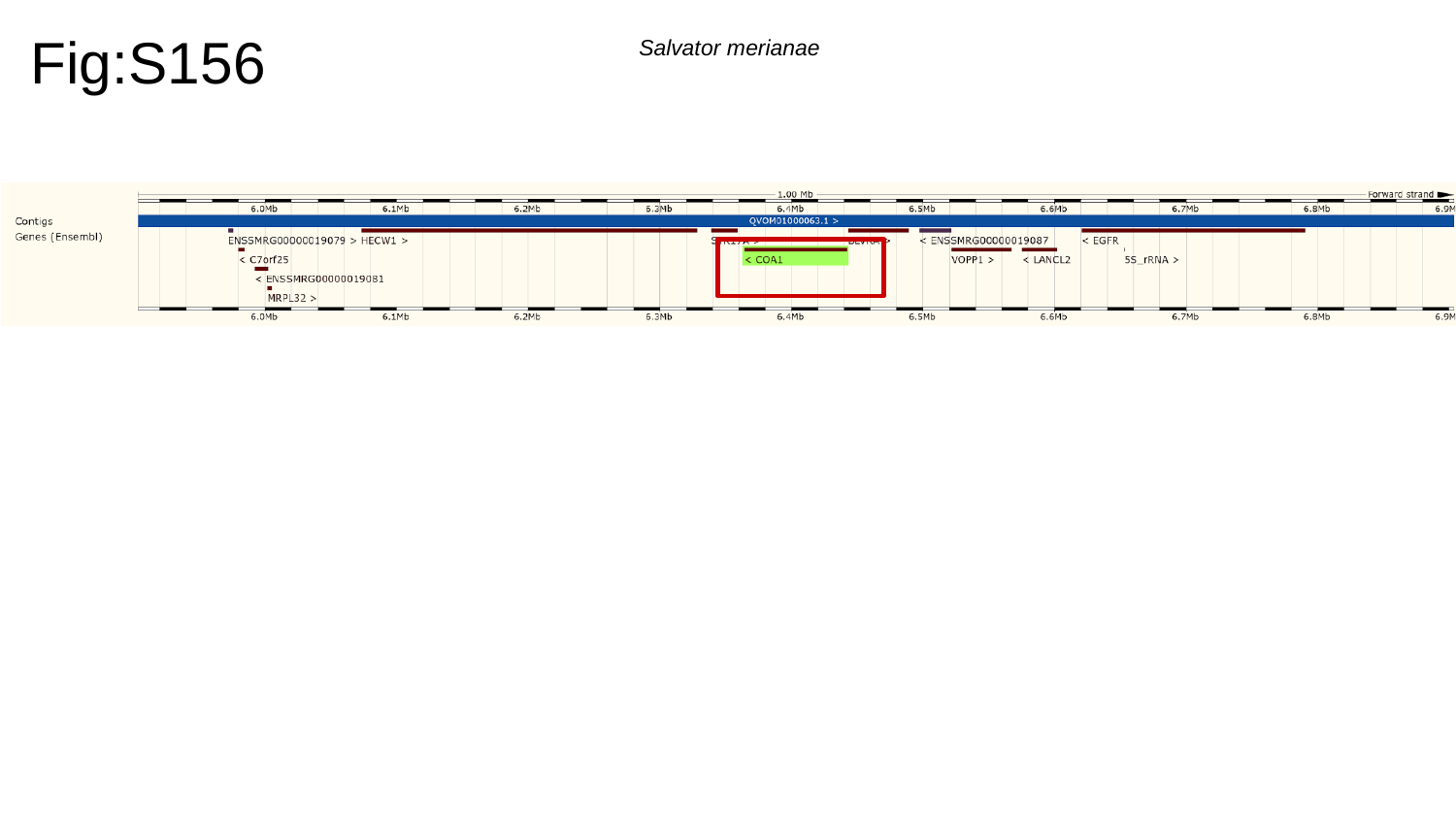

Fig:S156
Salvator merianae

### Slide 157
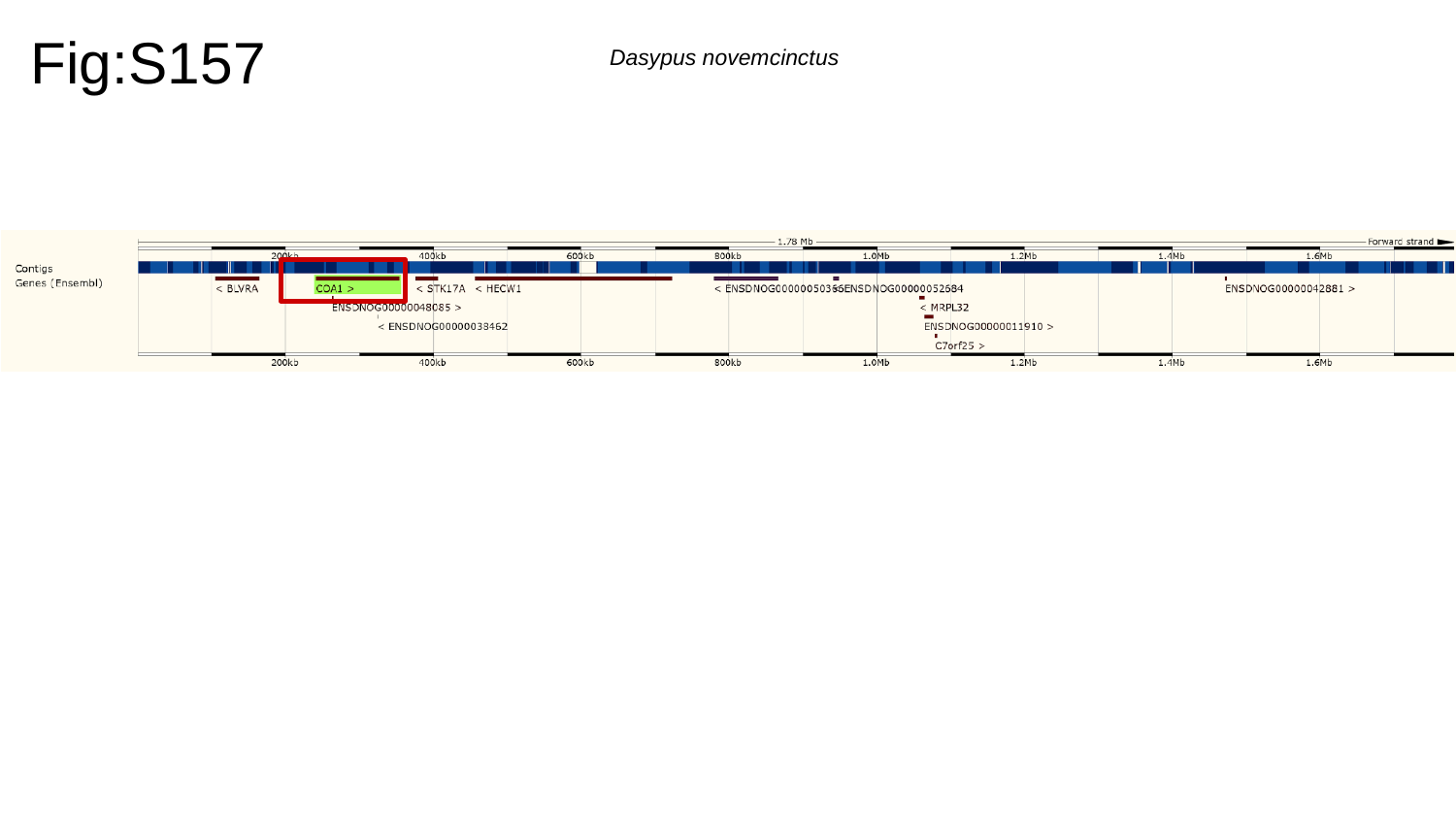

Fig:S157
Dasypus novemcinctus

### Slide 158
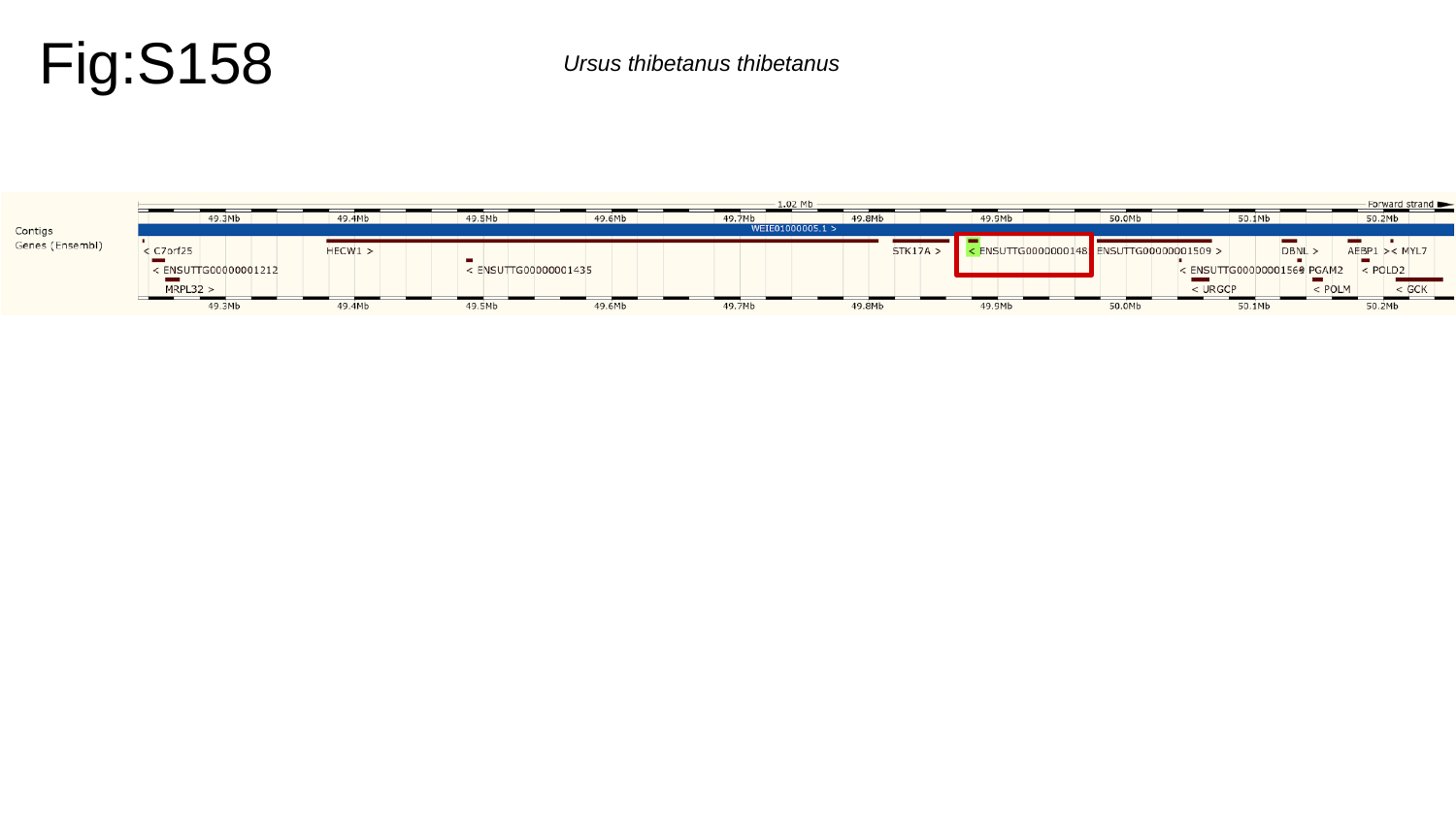

Fig:S158
Ursus thibetanus thibetanus

### Slide 159
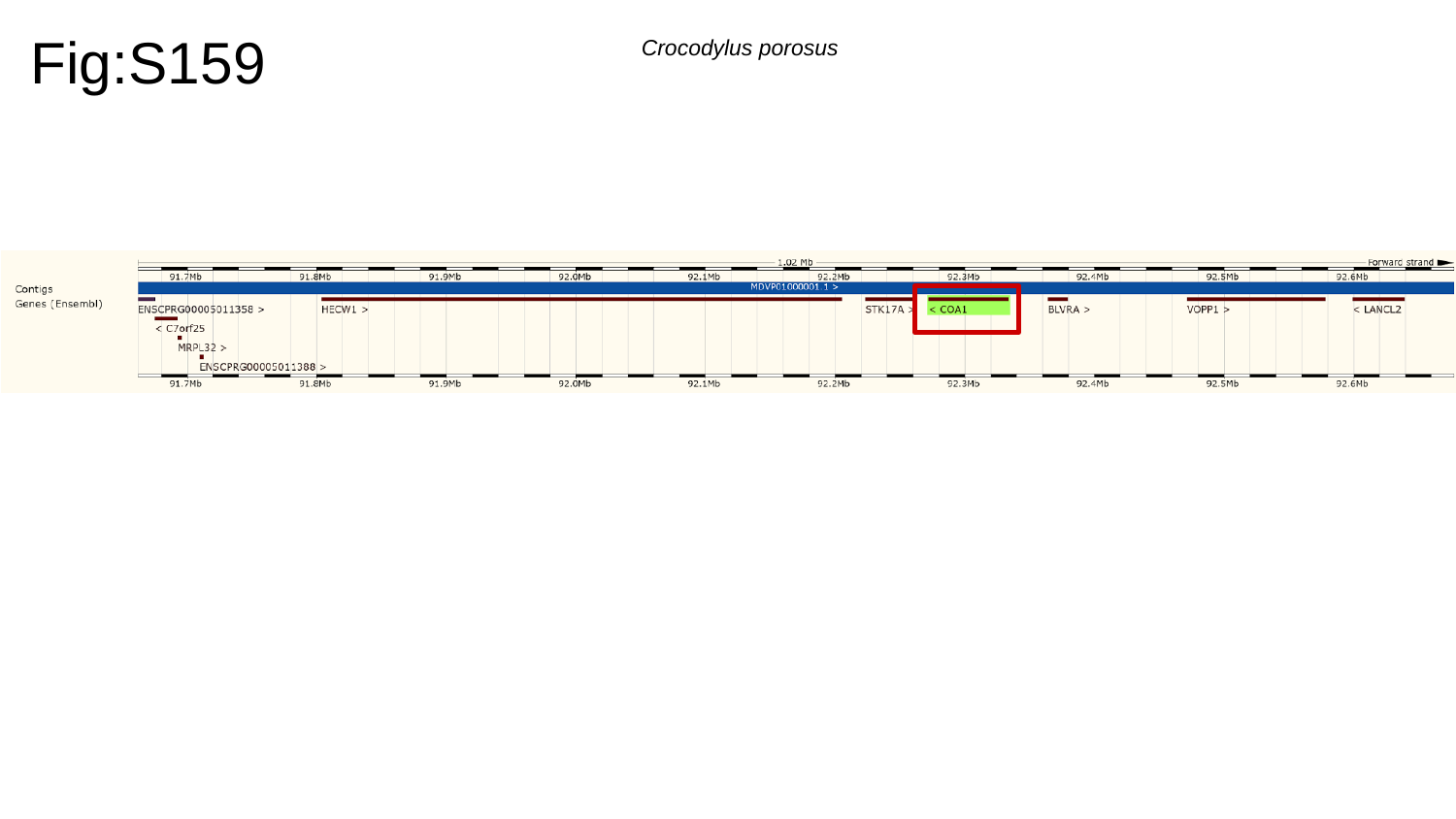

Fig:S159
Crocodylus porosus

### Slide 160
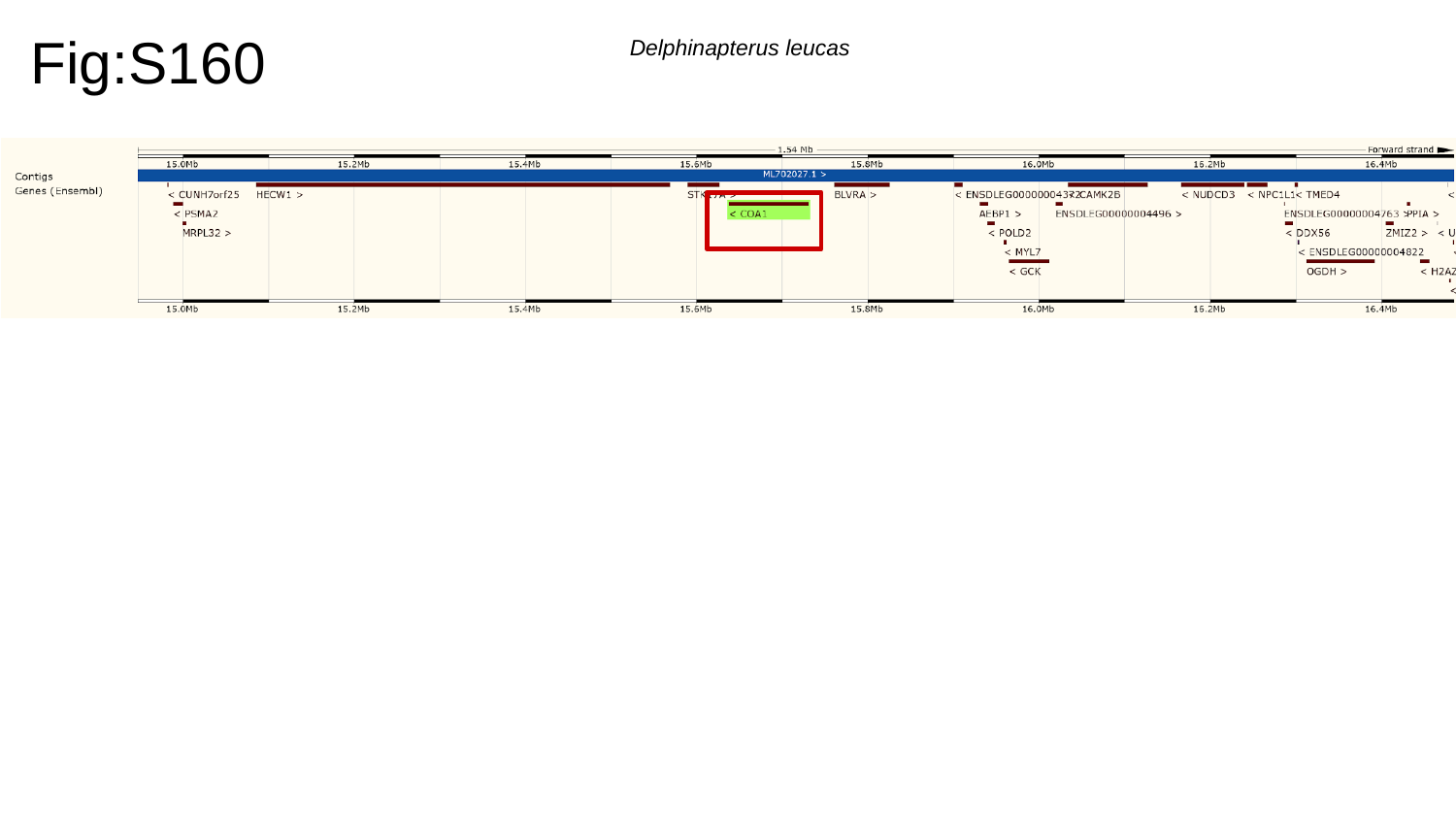

Fig:S160
Delphinapterus leucas

### Slide 161
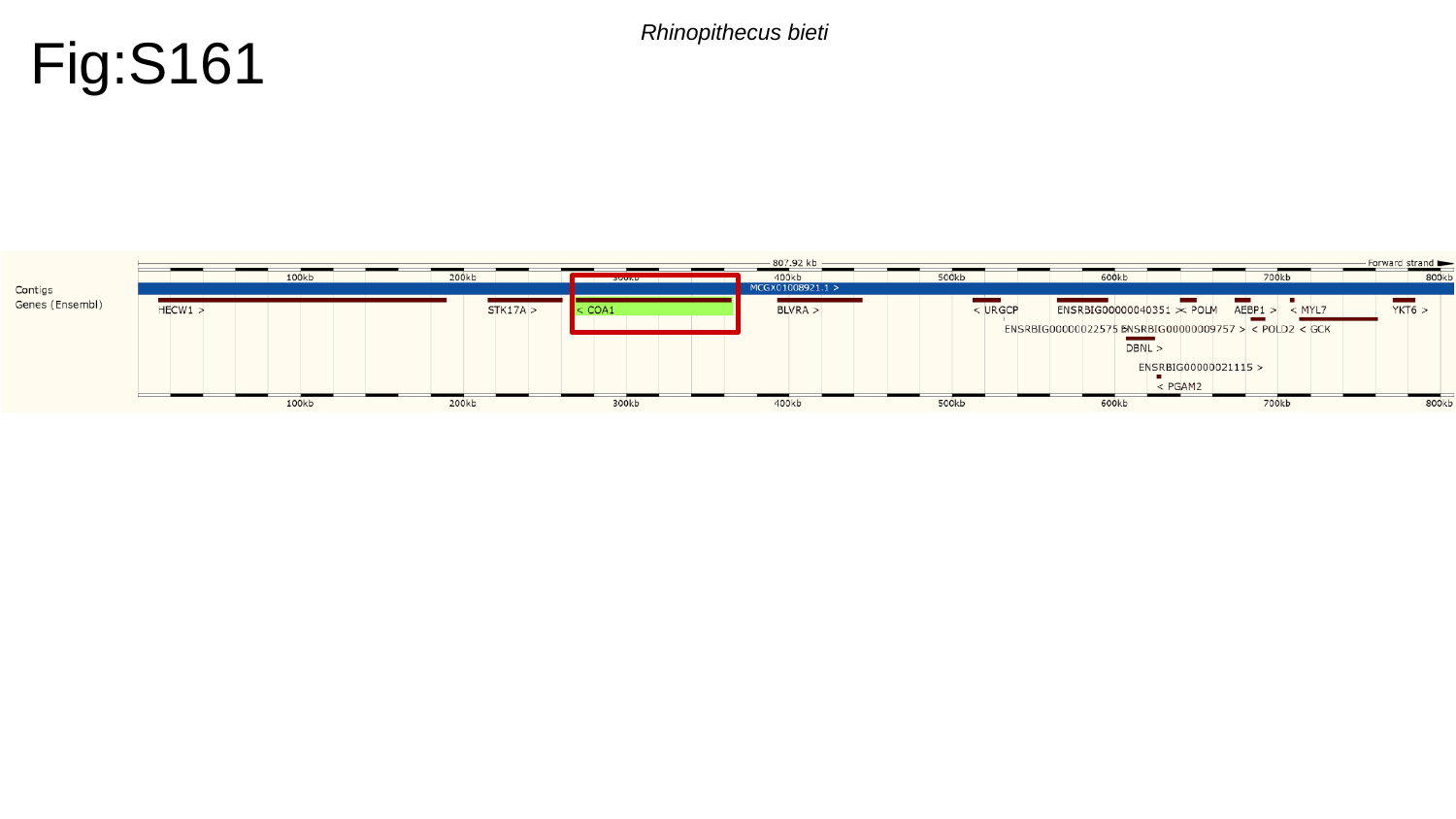

Fig:S161
Rhinopithecus bieti

### Slide 162
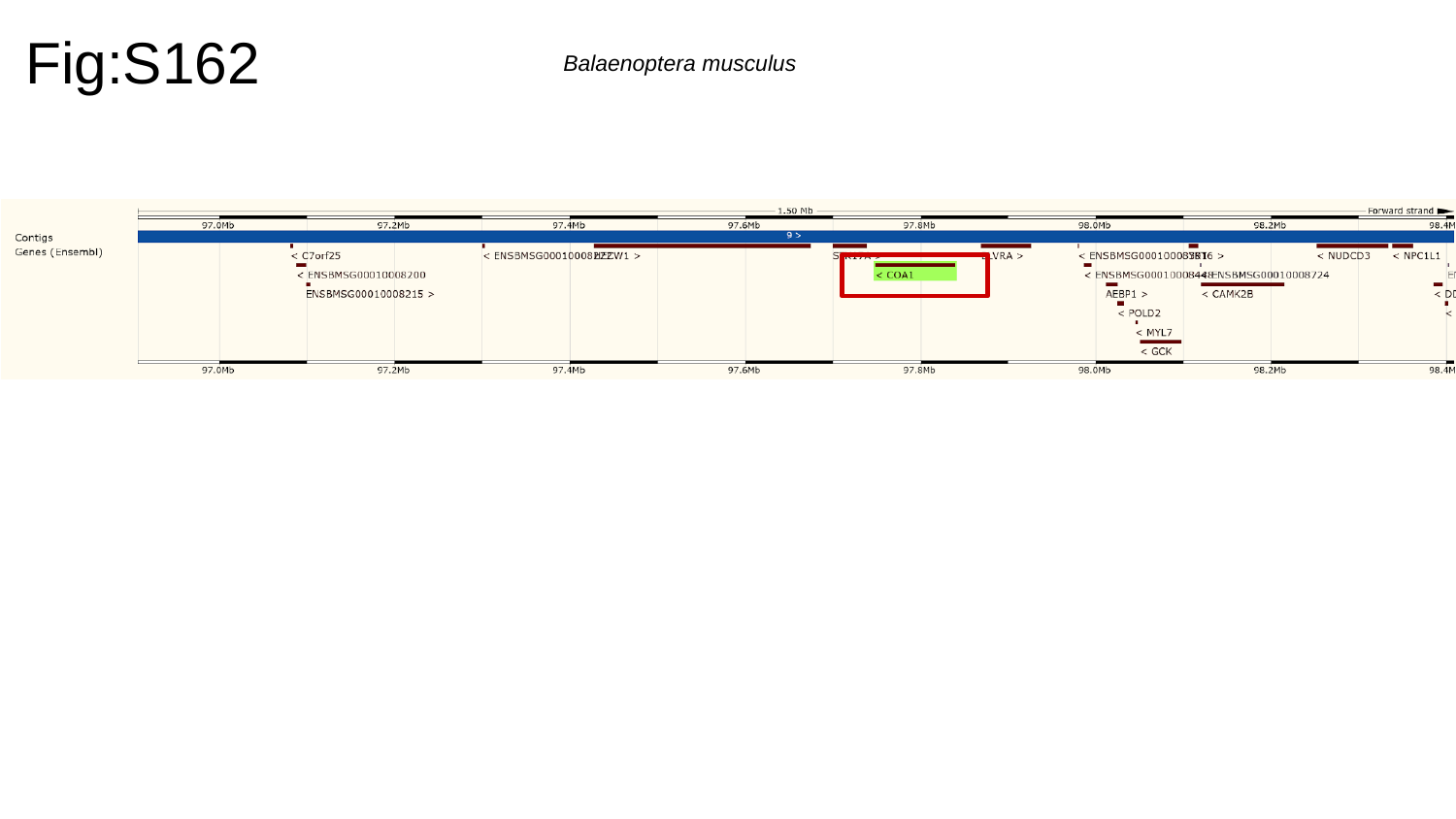

Fig:S162
Balaenoptera musculus

### Slide 163
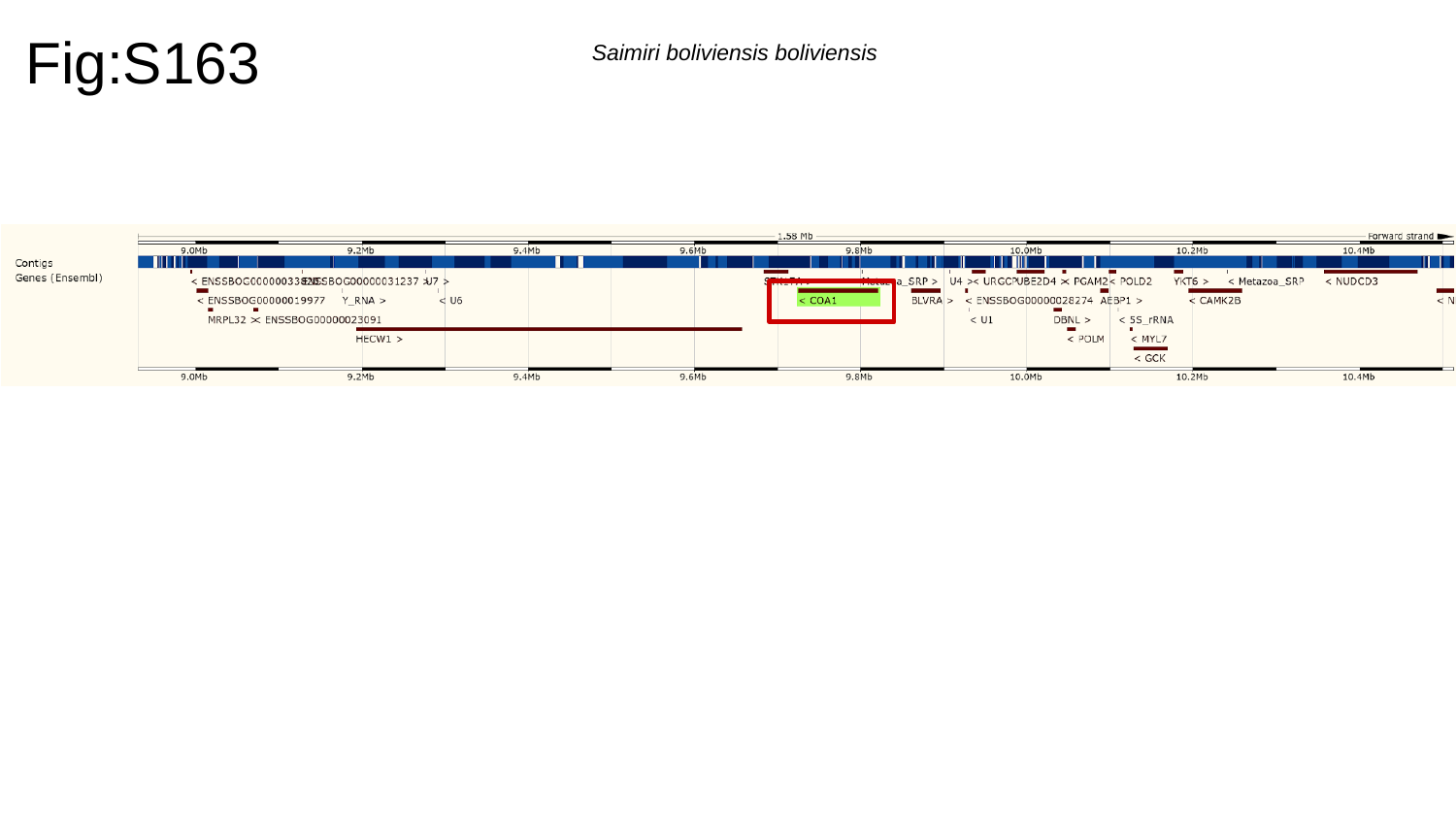

Fig:S163
Saimiri boliviensis boliviensis

### Slide 164
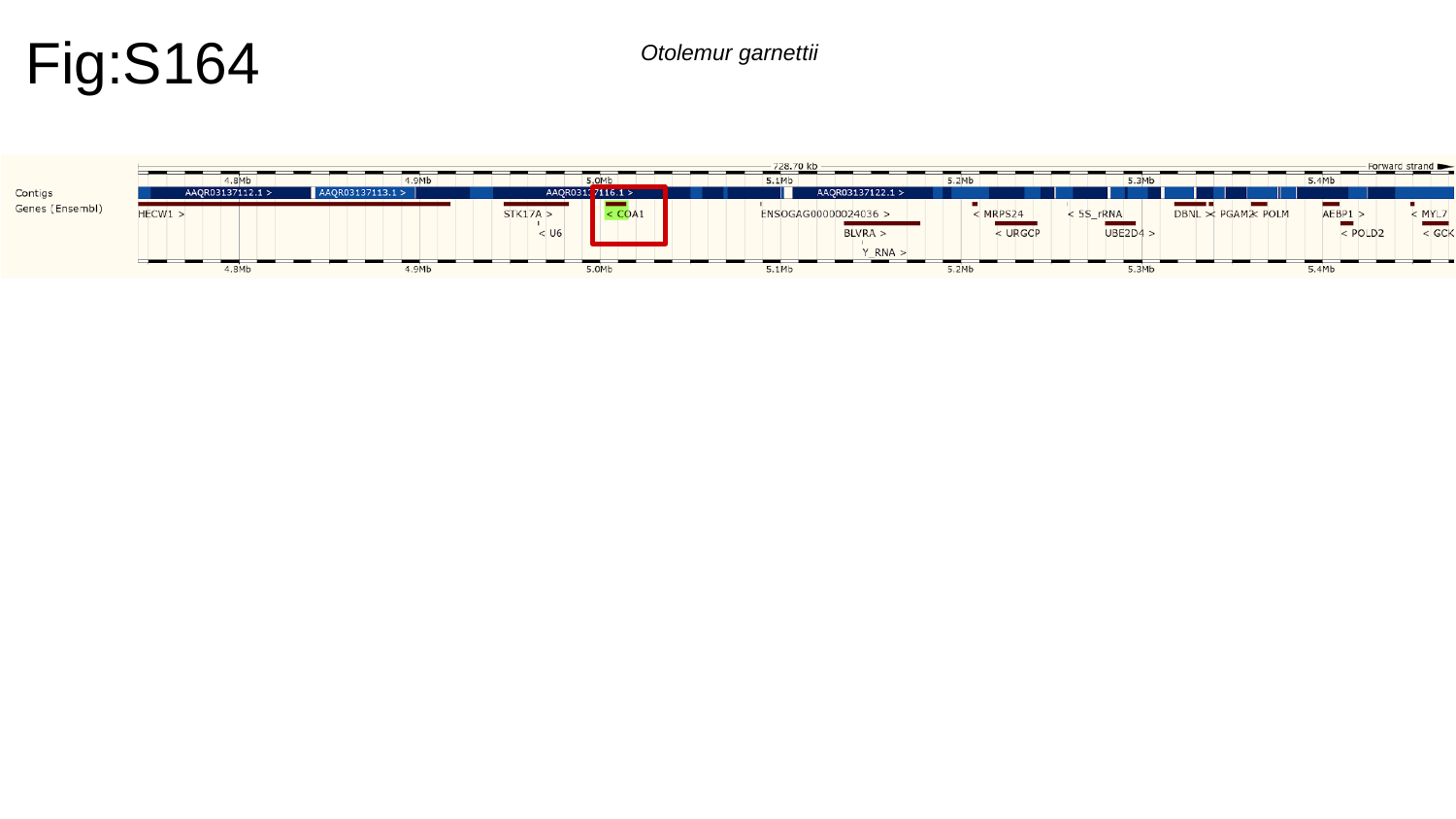

Fig:S164
Otolemur garnettii

### Slide 165
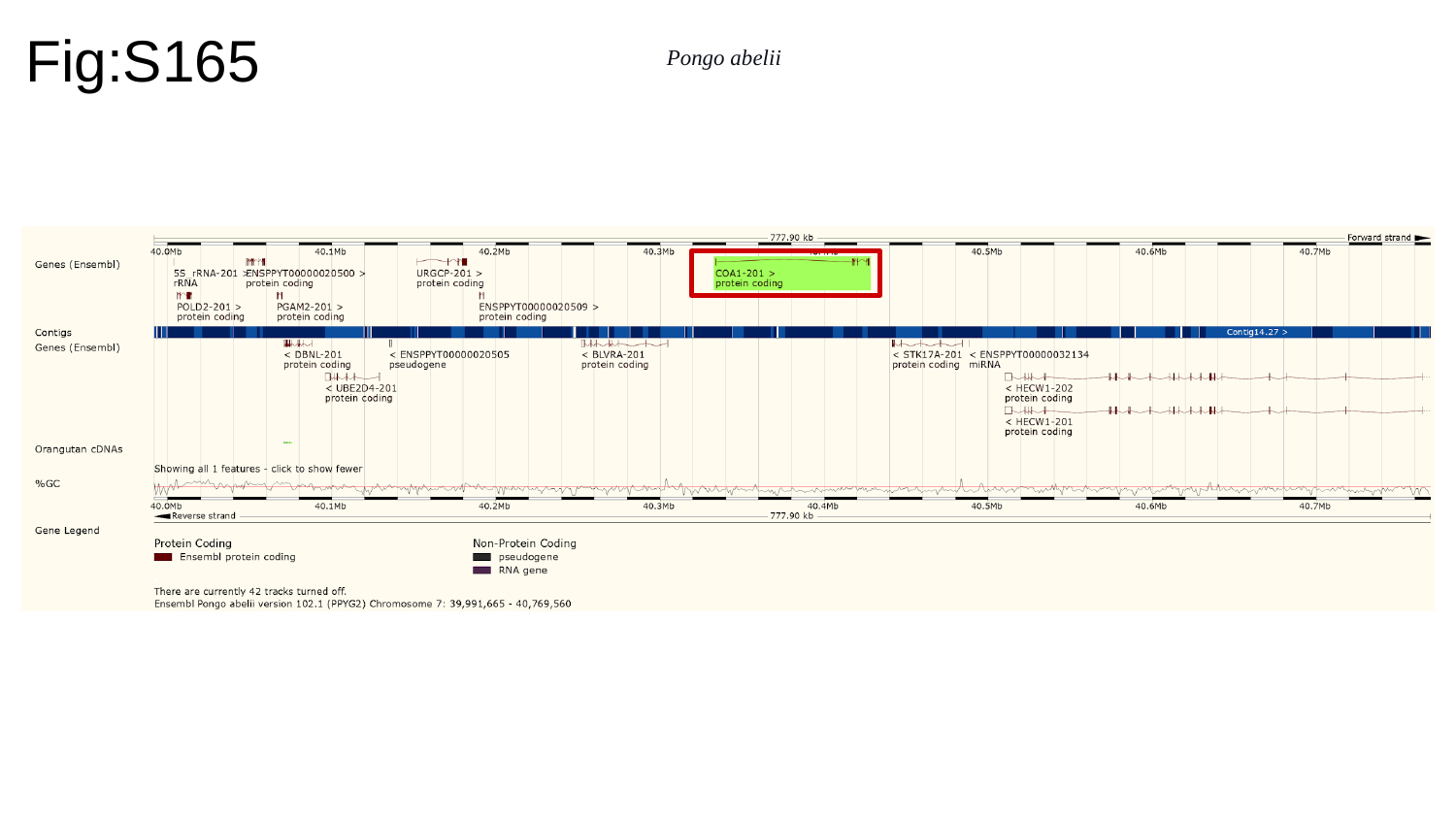

Fig:S165
Pongo abelii

### Slide 166
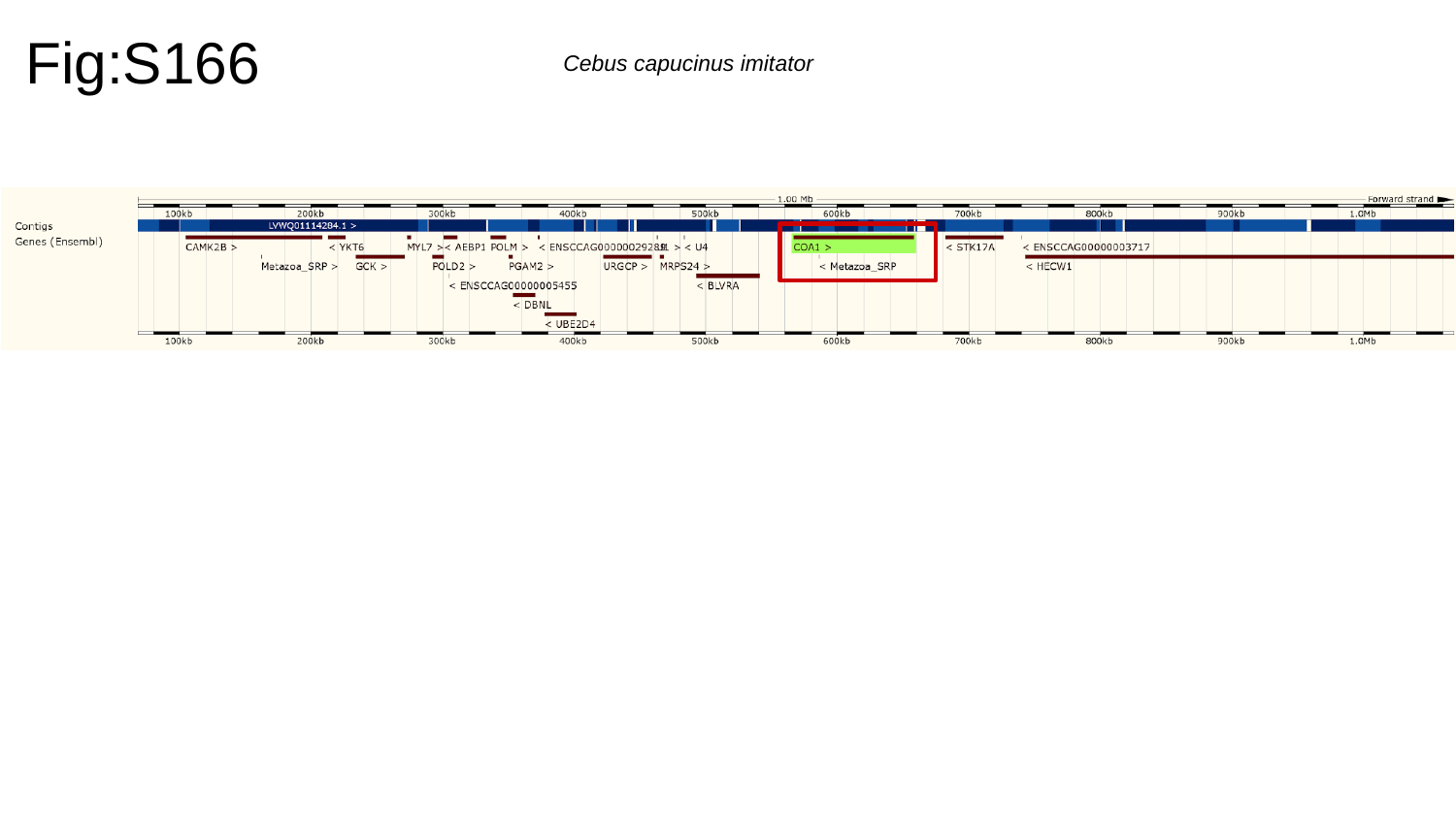

Fig:S166
Cebus capucinus imitator

### Slide 167
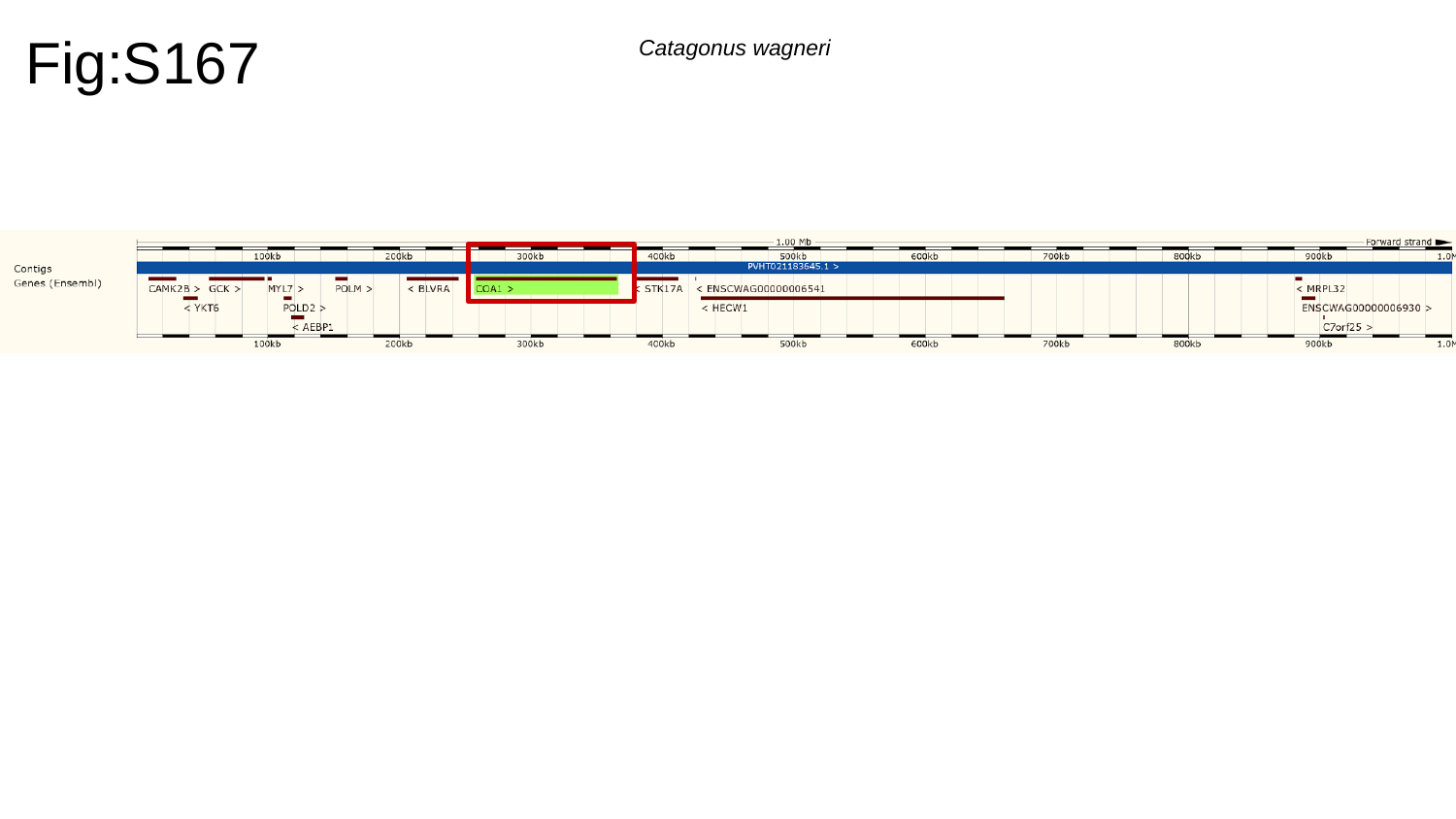

Fig:S167
Catagonus wagneri

### Slide 168
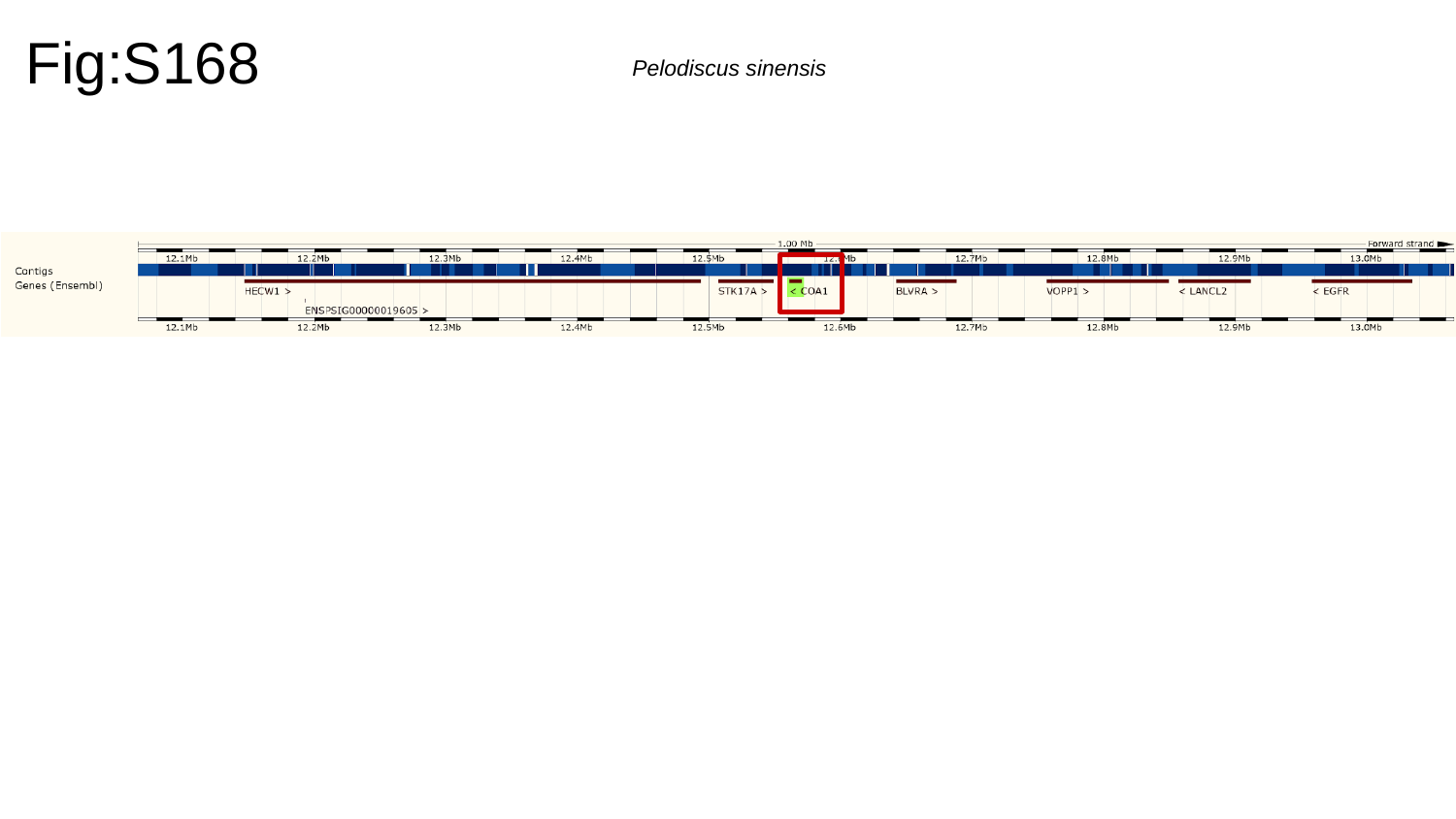

Fig:S168
Pelodiscus sinensis

### Slide 169
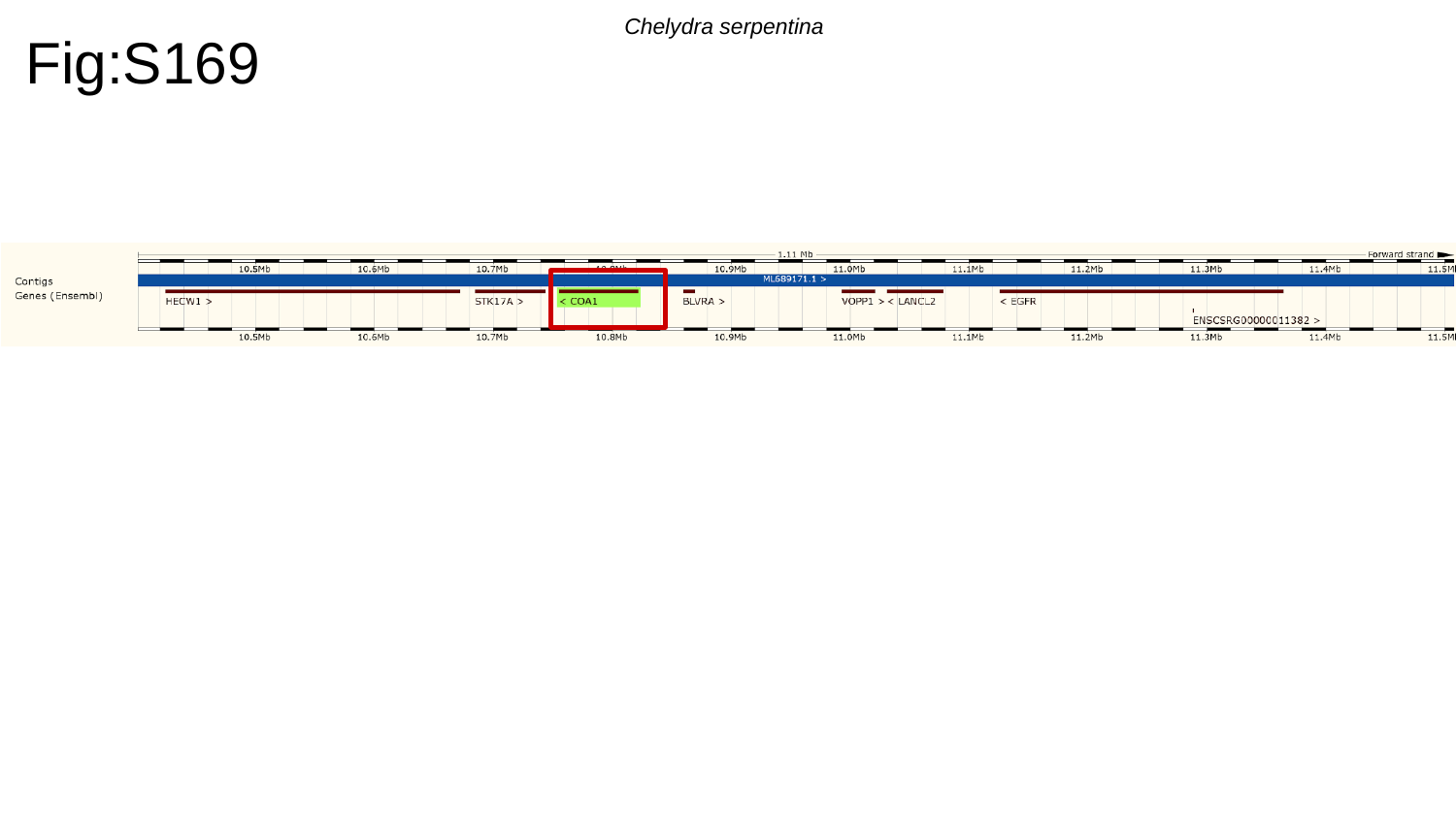

Fig:S169
Chelydra serpentina

### Slide 170
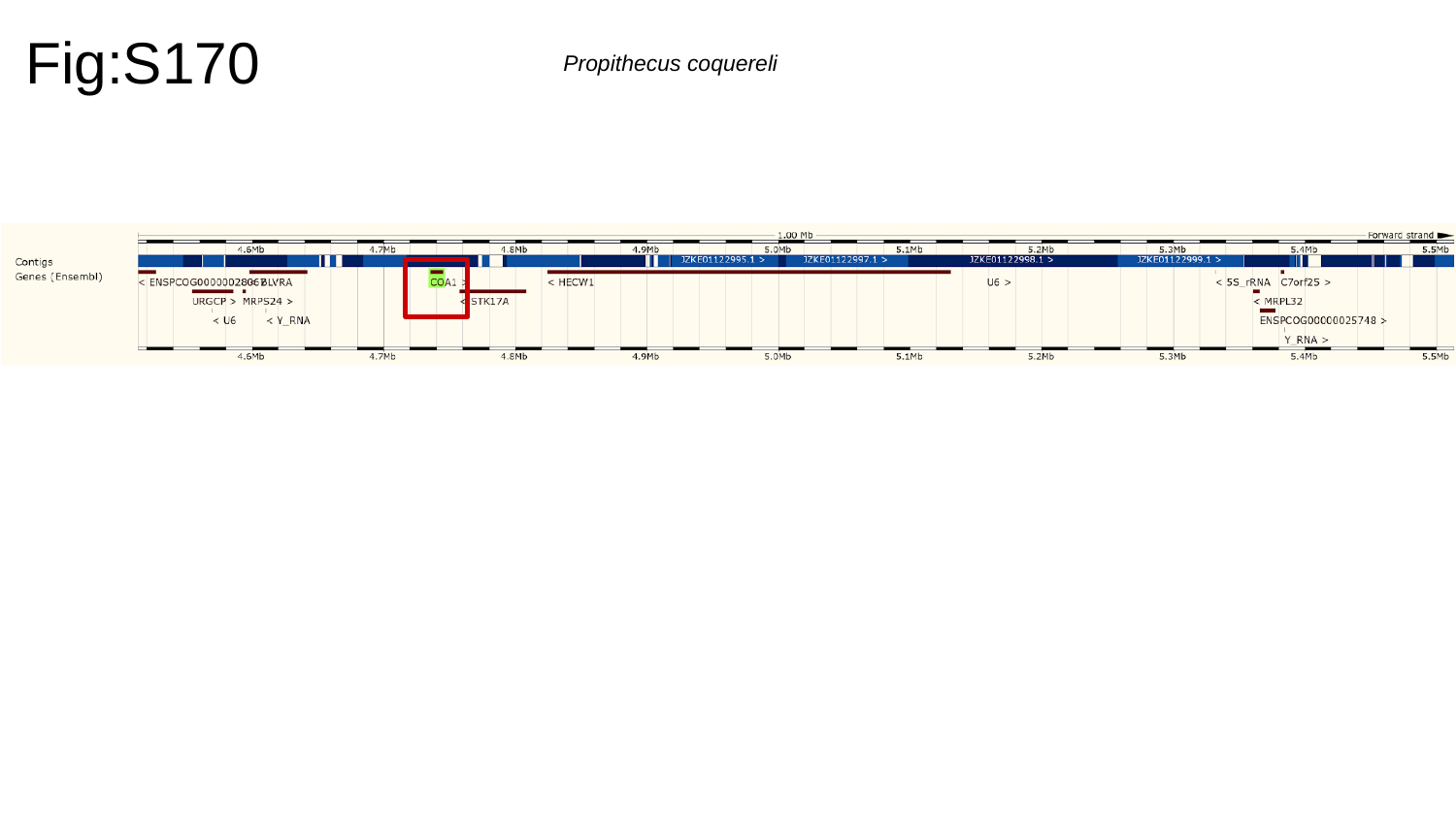

Fig:S170
Propithecus coquereli

### Slide 171
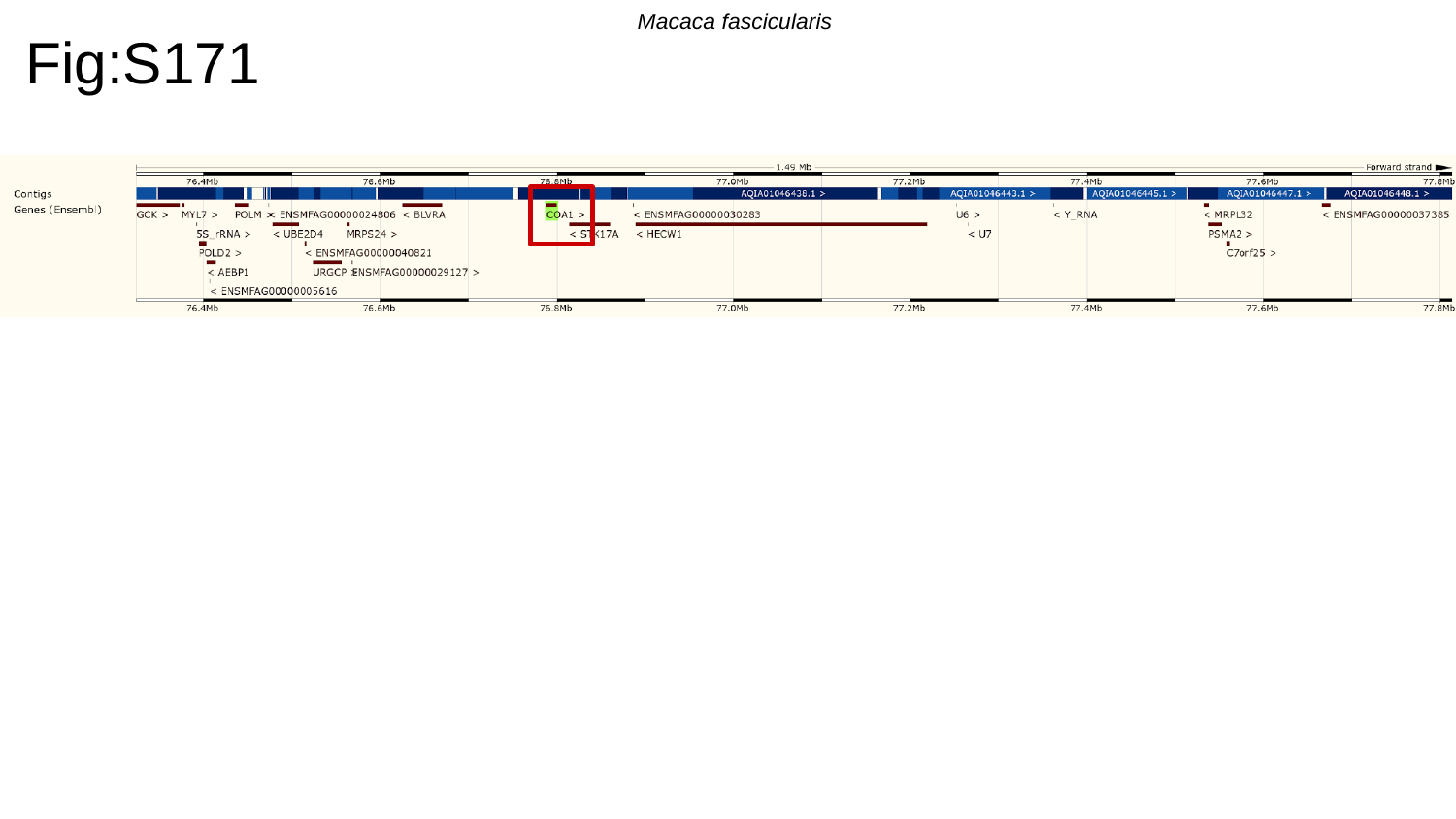

Macaca fascicularis
Fig:S171

### Slide 172
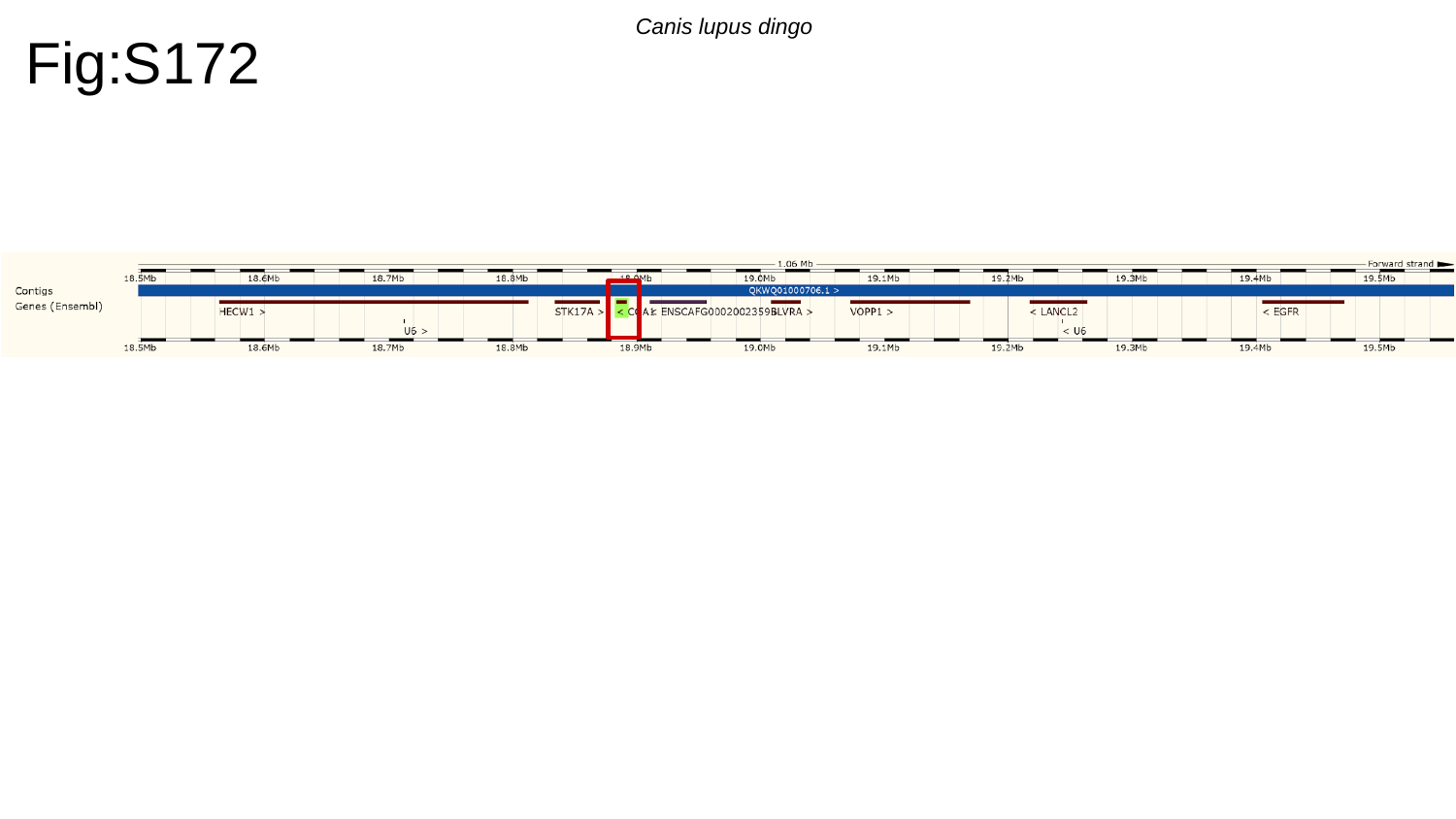

Fig:S172
Canis lupus dingo

### Slide 173
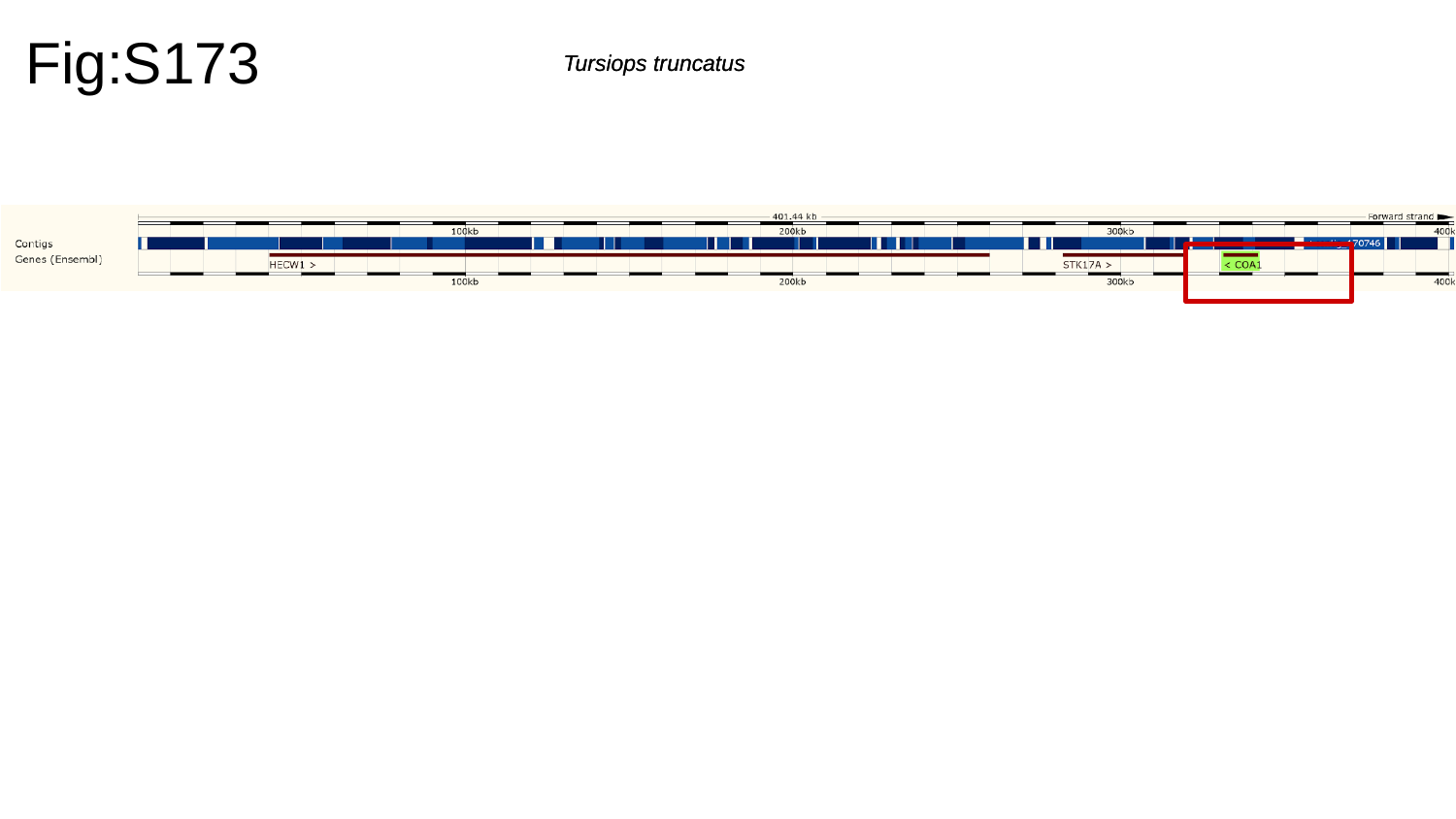

Fig:S173
Tursiops truncatus
Tursiops truncatus

### Slide 174
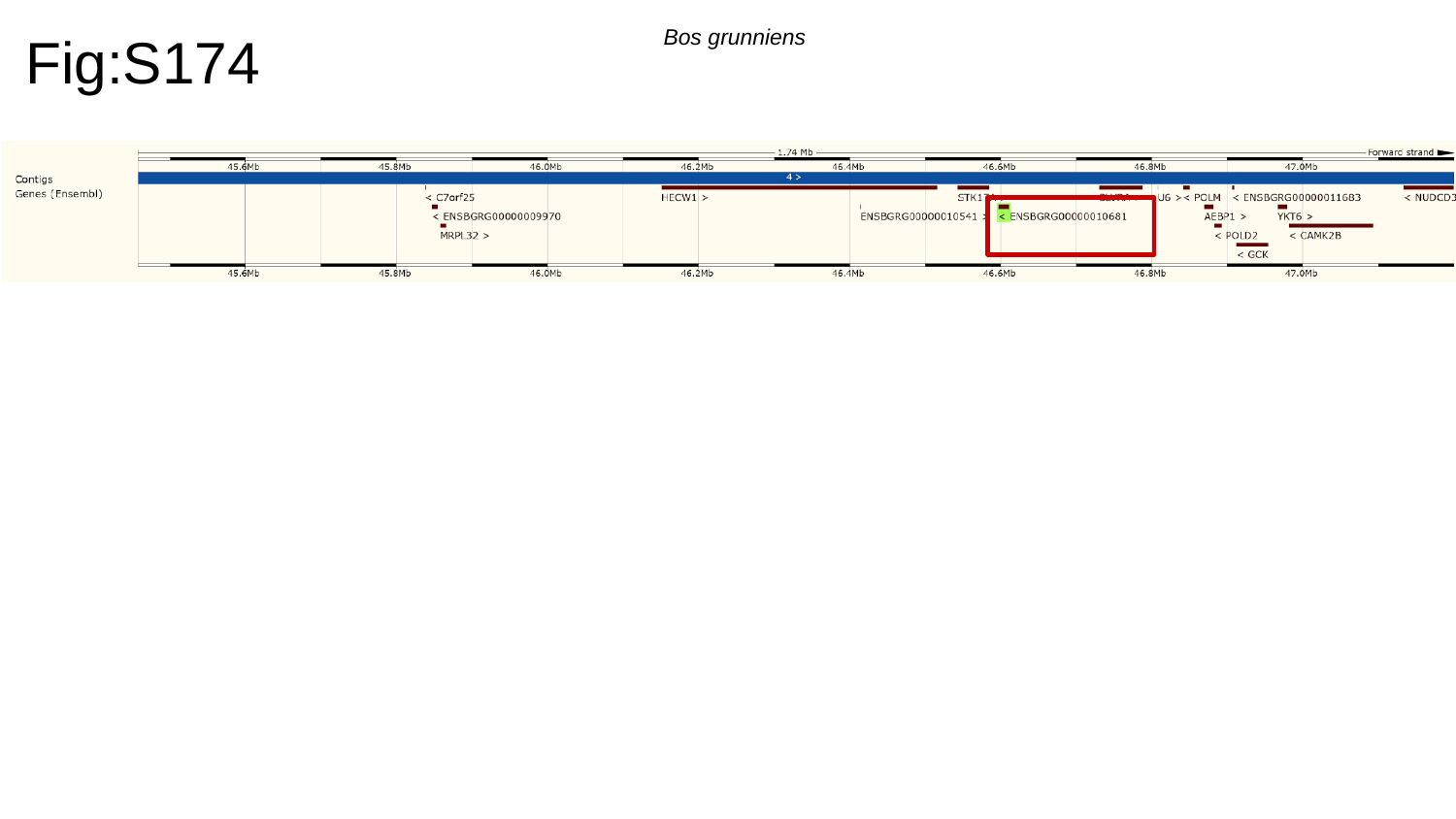

Fig:S174
Bos grunniens

### Slide 175
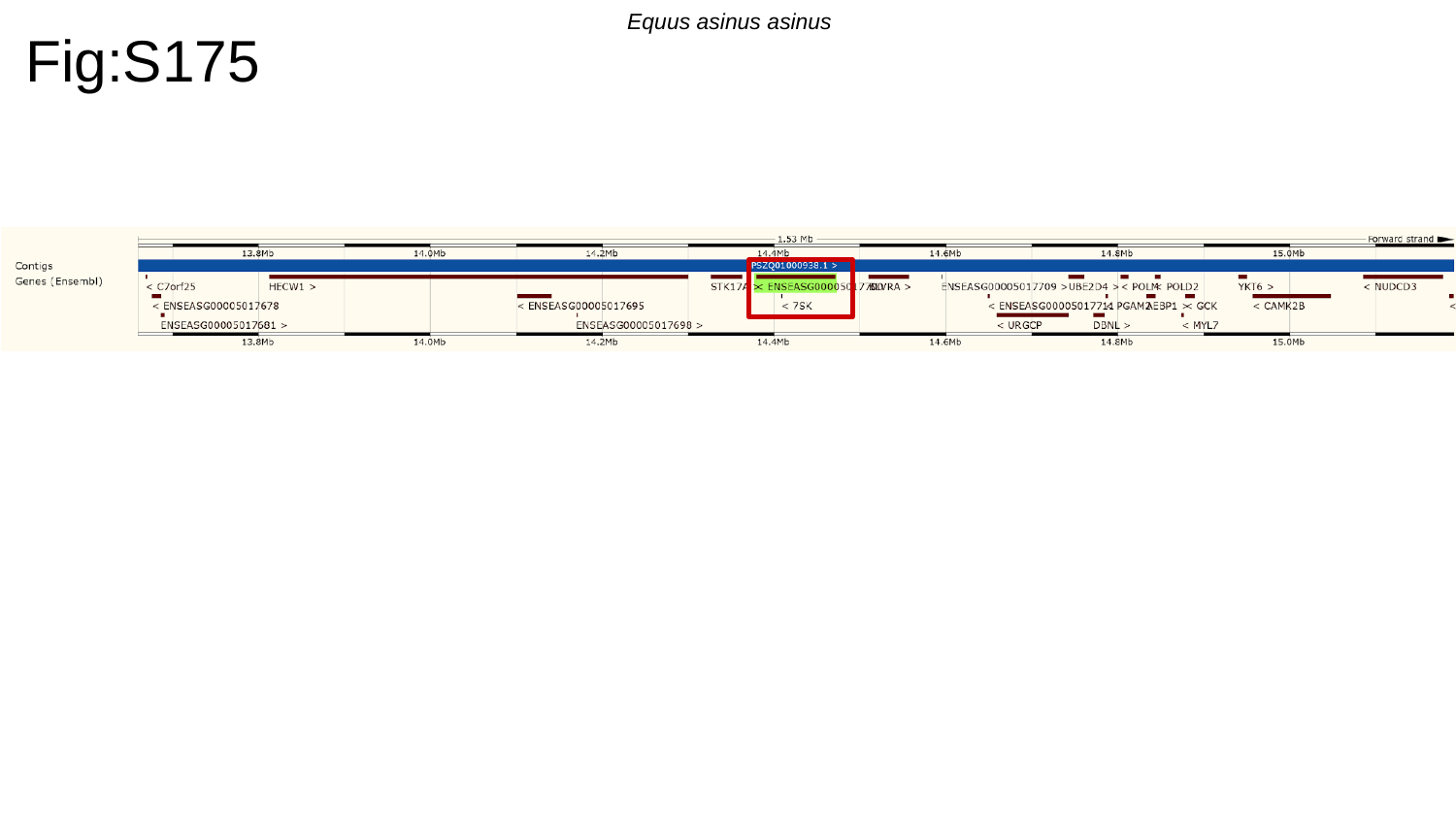

Equus asinus asinus
Fig:S175

### Slide 176
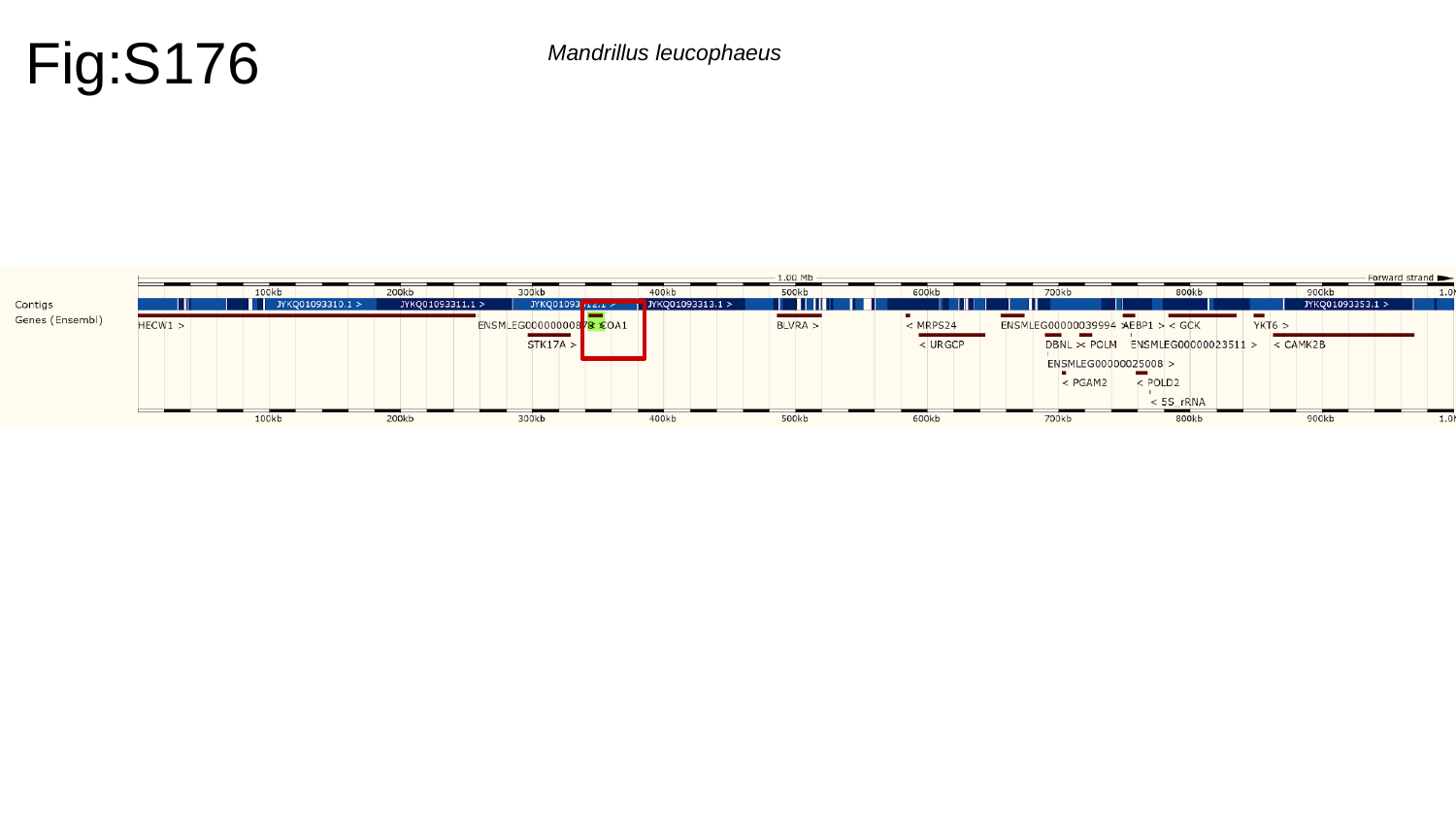

Fig:S176
Mandrillus leucophaeus

### Slide 177
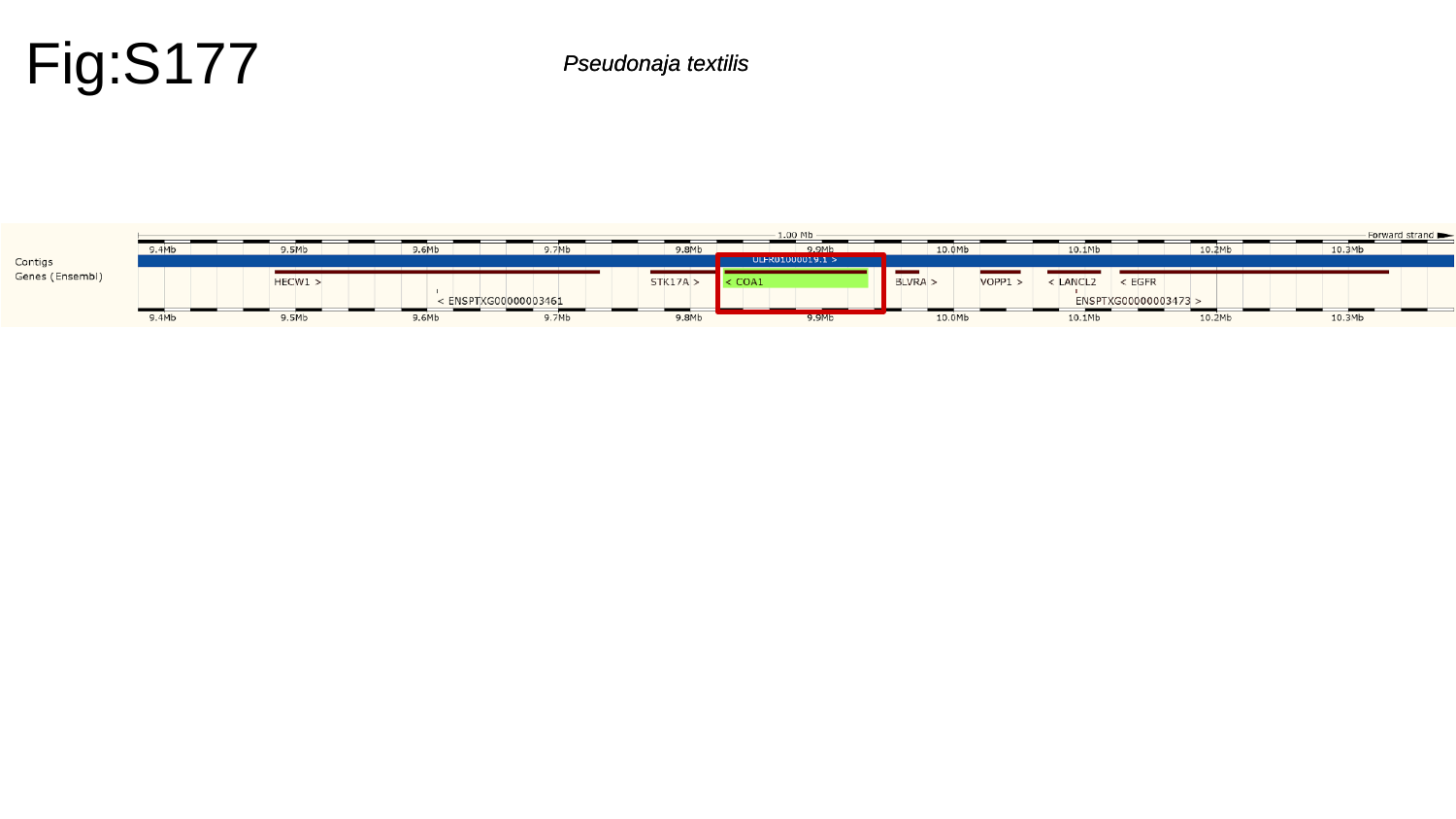

Fig:S177
Pseudonaja textilis
Pseudonaja textilis

### Slide 178
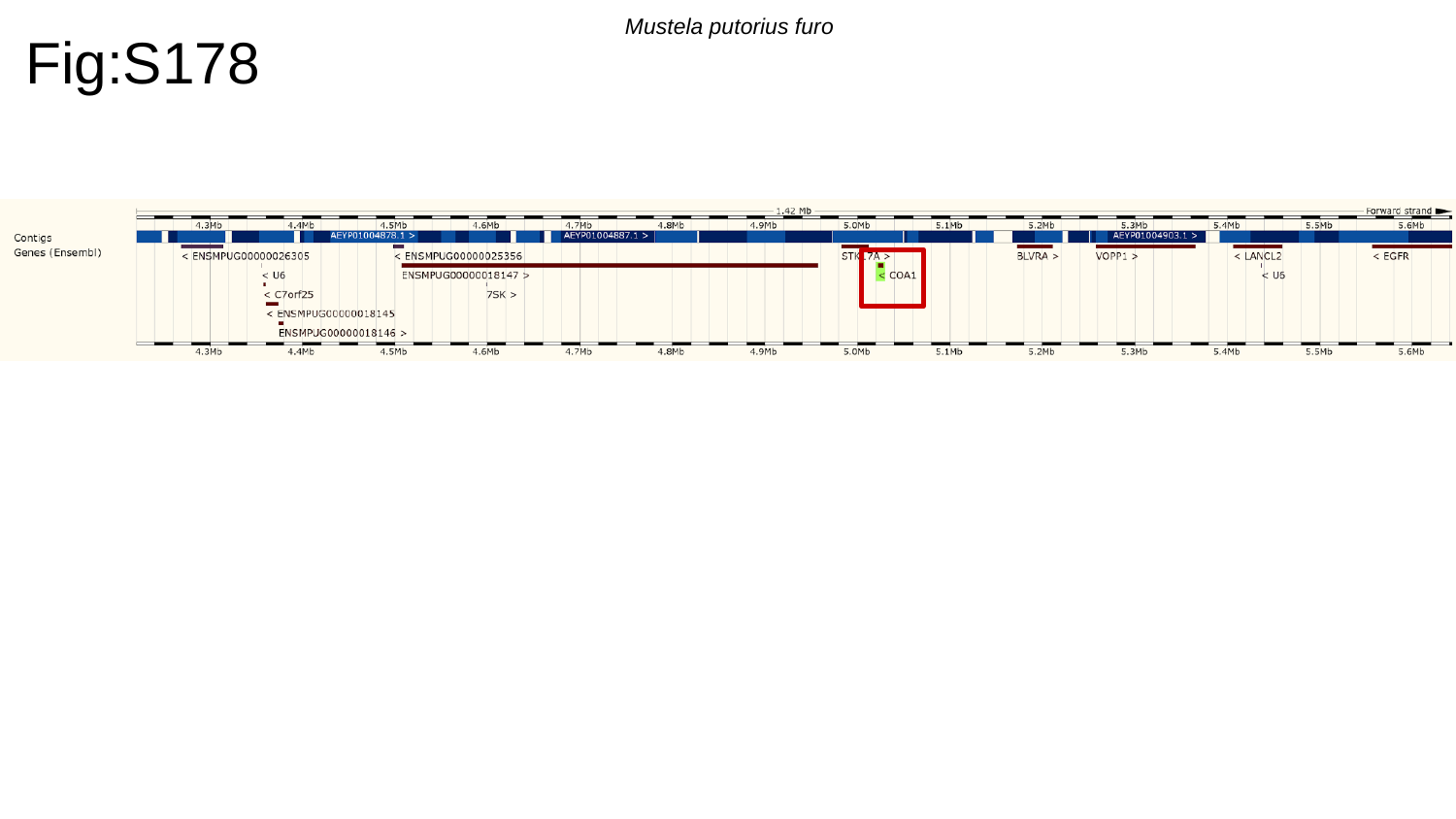

Fig:S178
Mustela putorius furo

### Slide 179
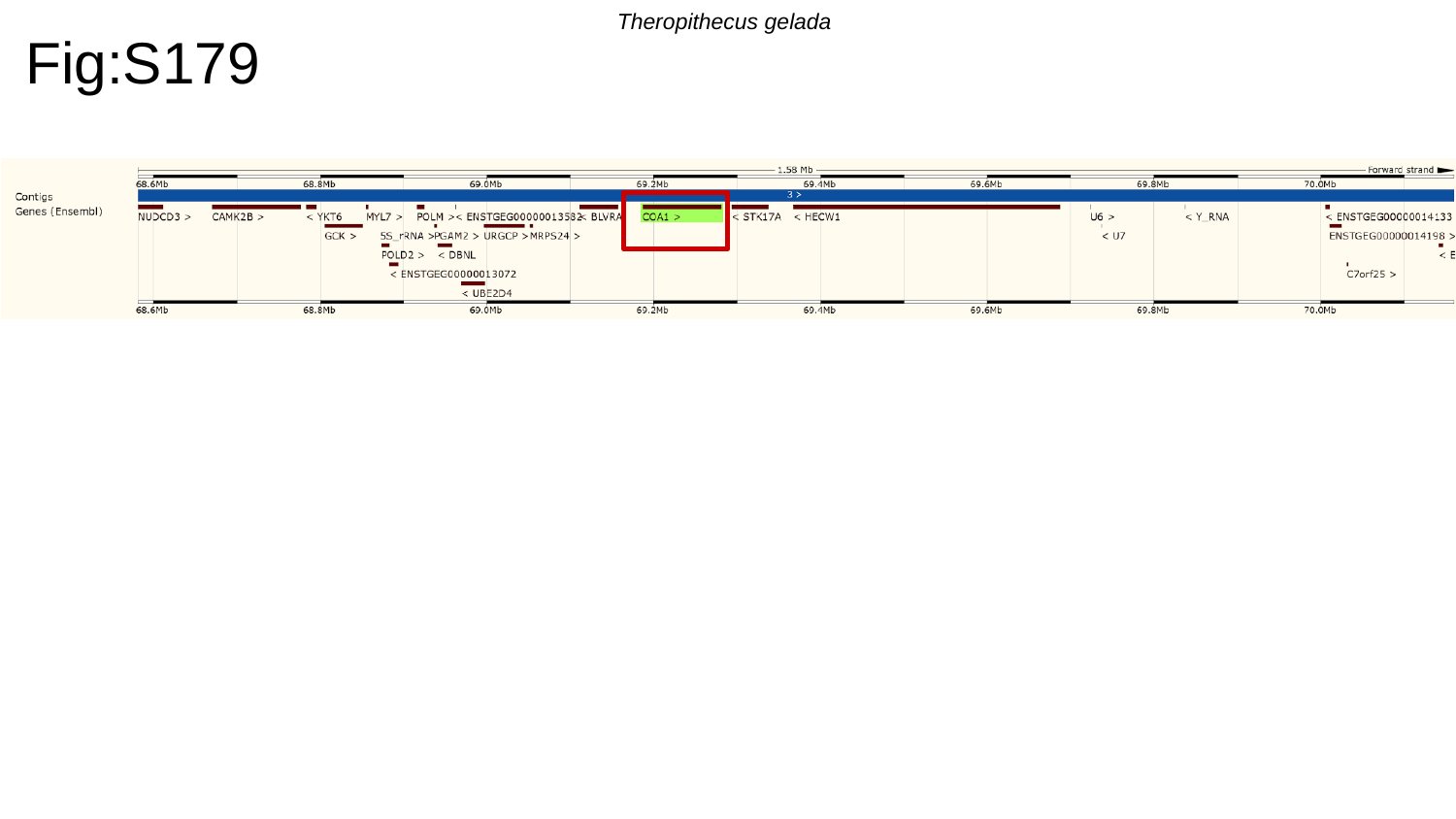

Theropithecus gelada
Fig:S179

### Slide 180
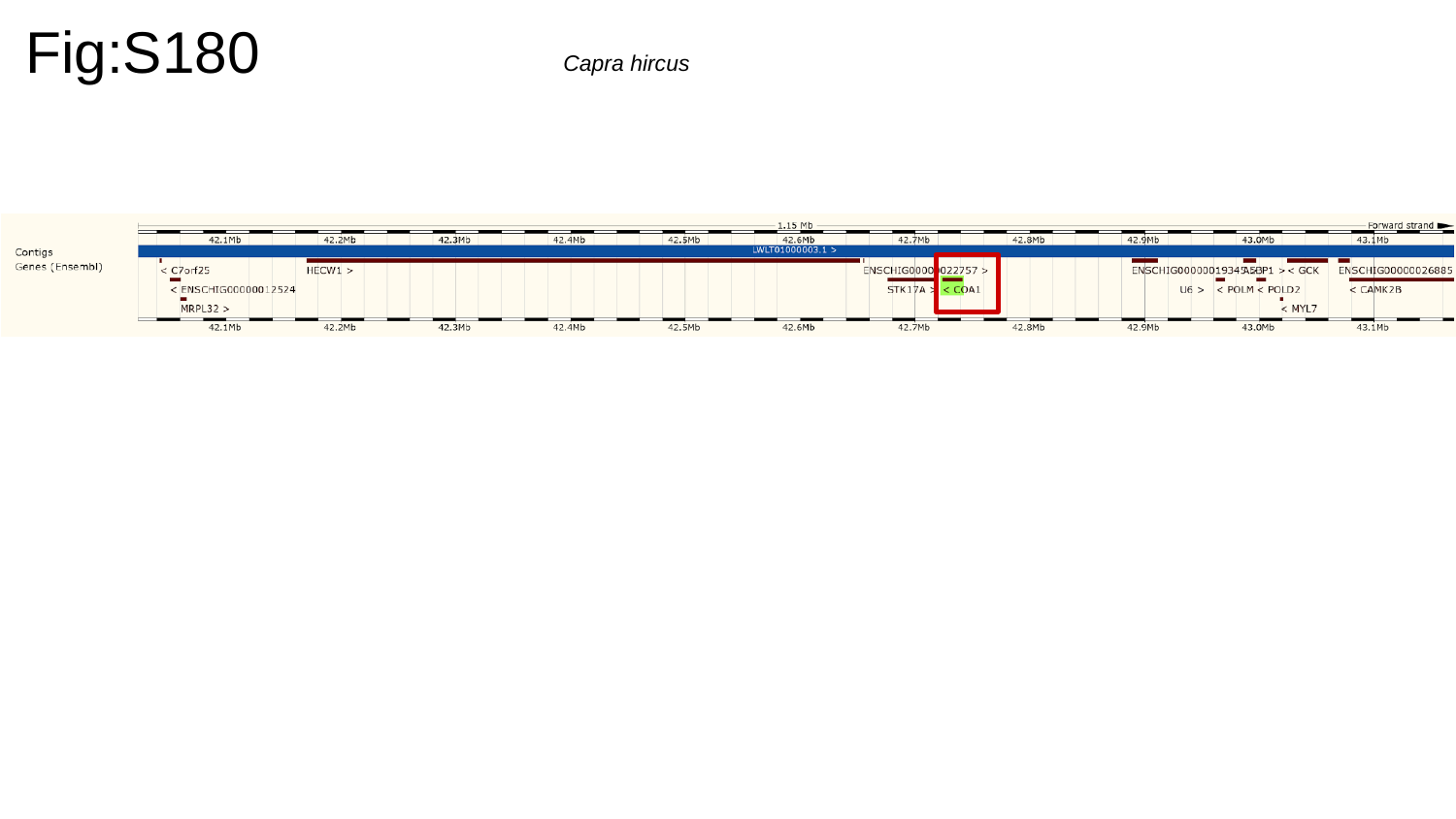

Fig:S180
Capra hircus

### Slide 181
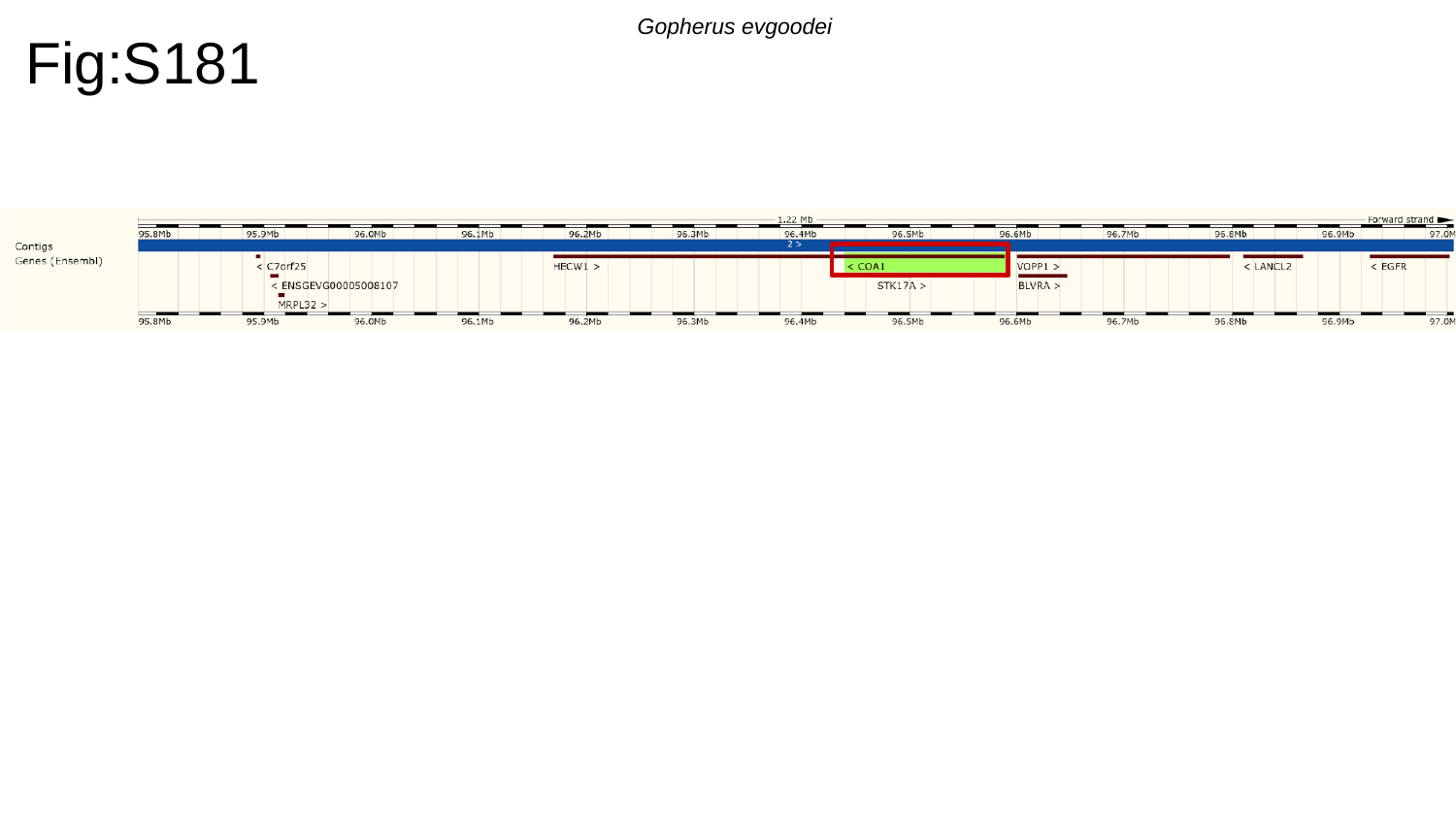

Fig:S181
Gopherus evgoodei

### Slide 182
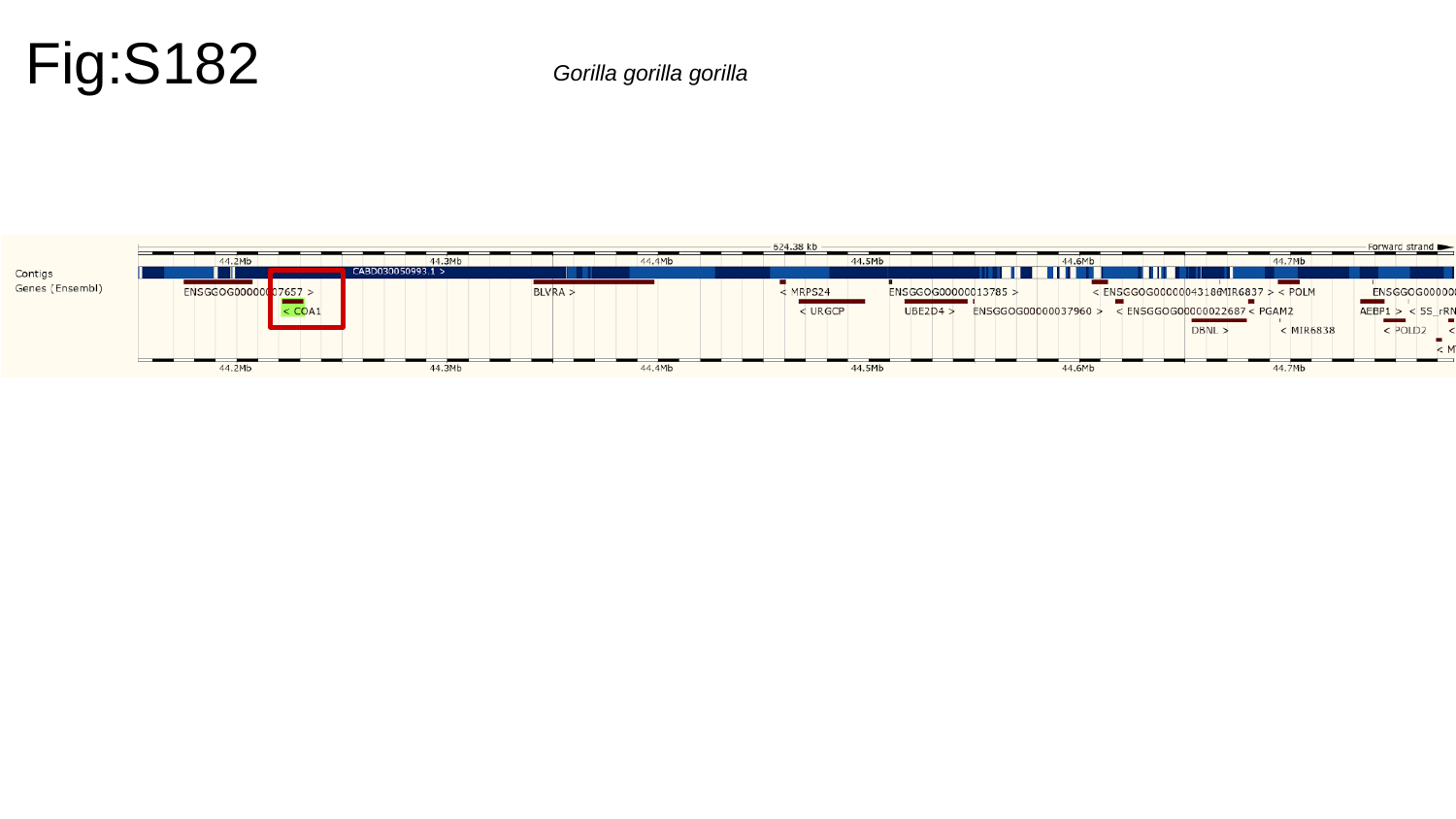

Fig:S182
Gorilla gorilla gorilla

### Slide 183
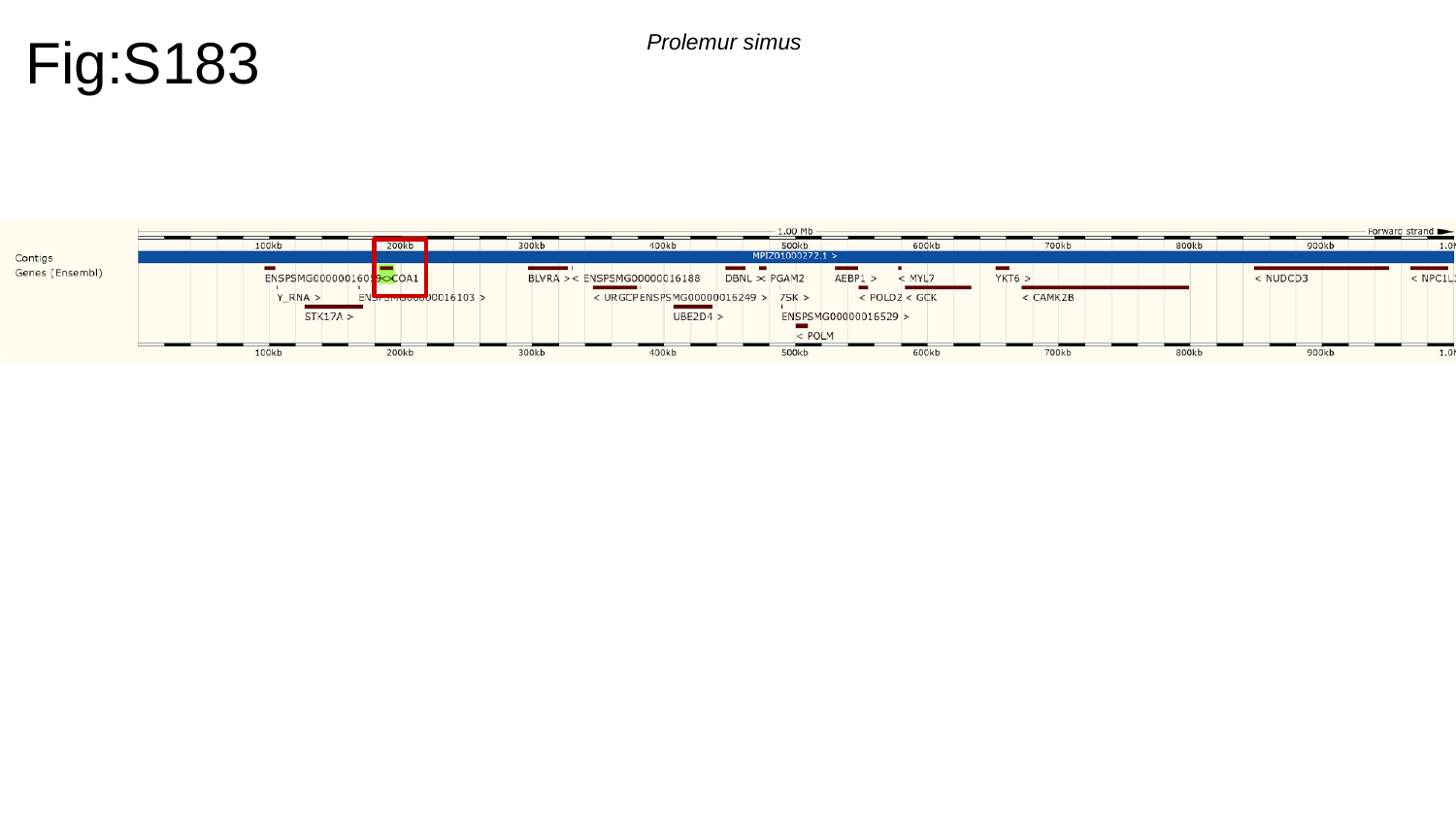

Fig:S183
Prolemur simus

### Slide 184
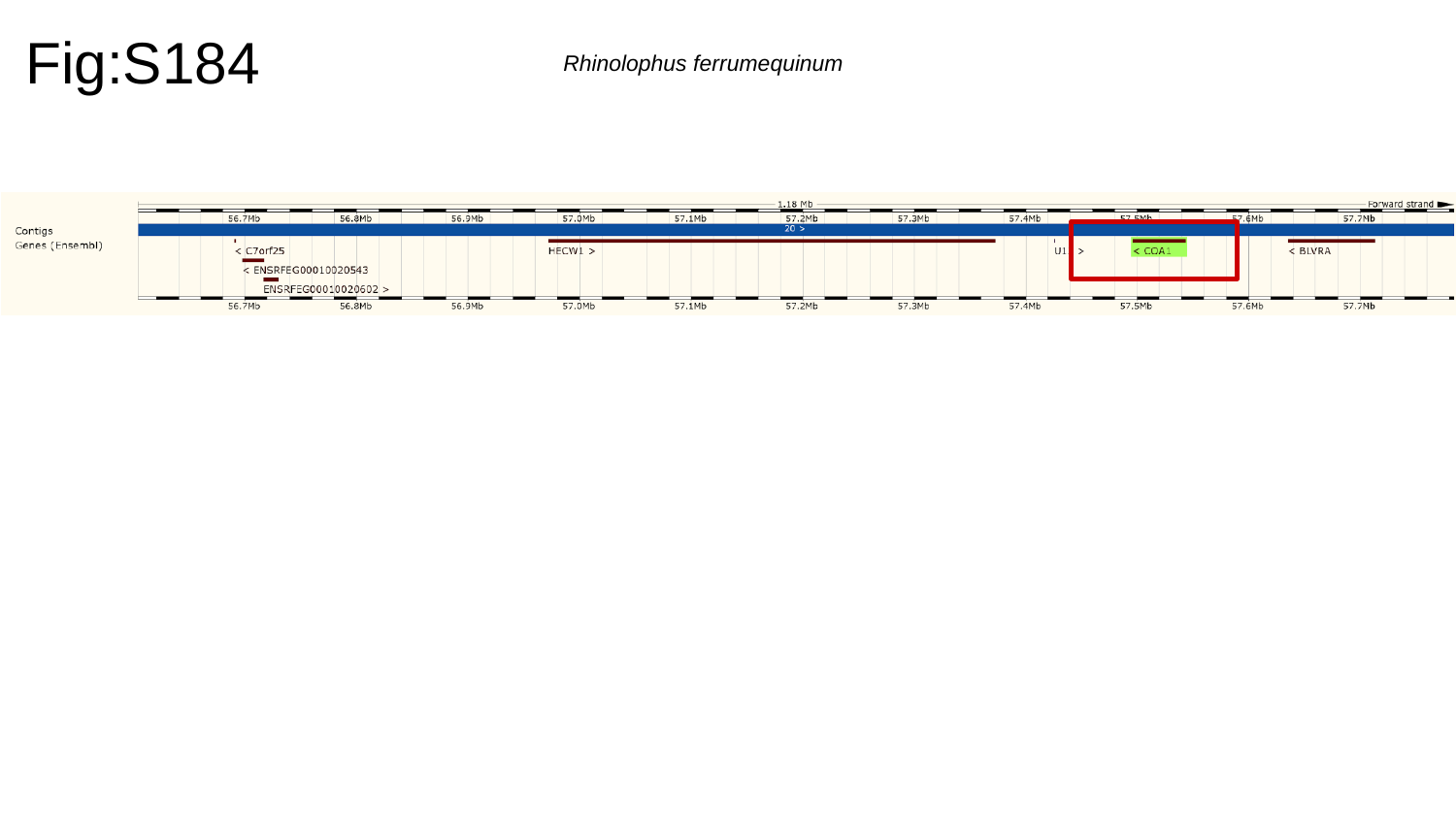

Fig:S184
Rhinolophus ferrumequinum

### Slide 185
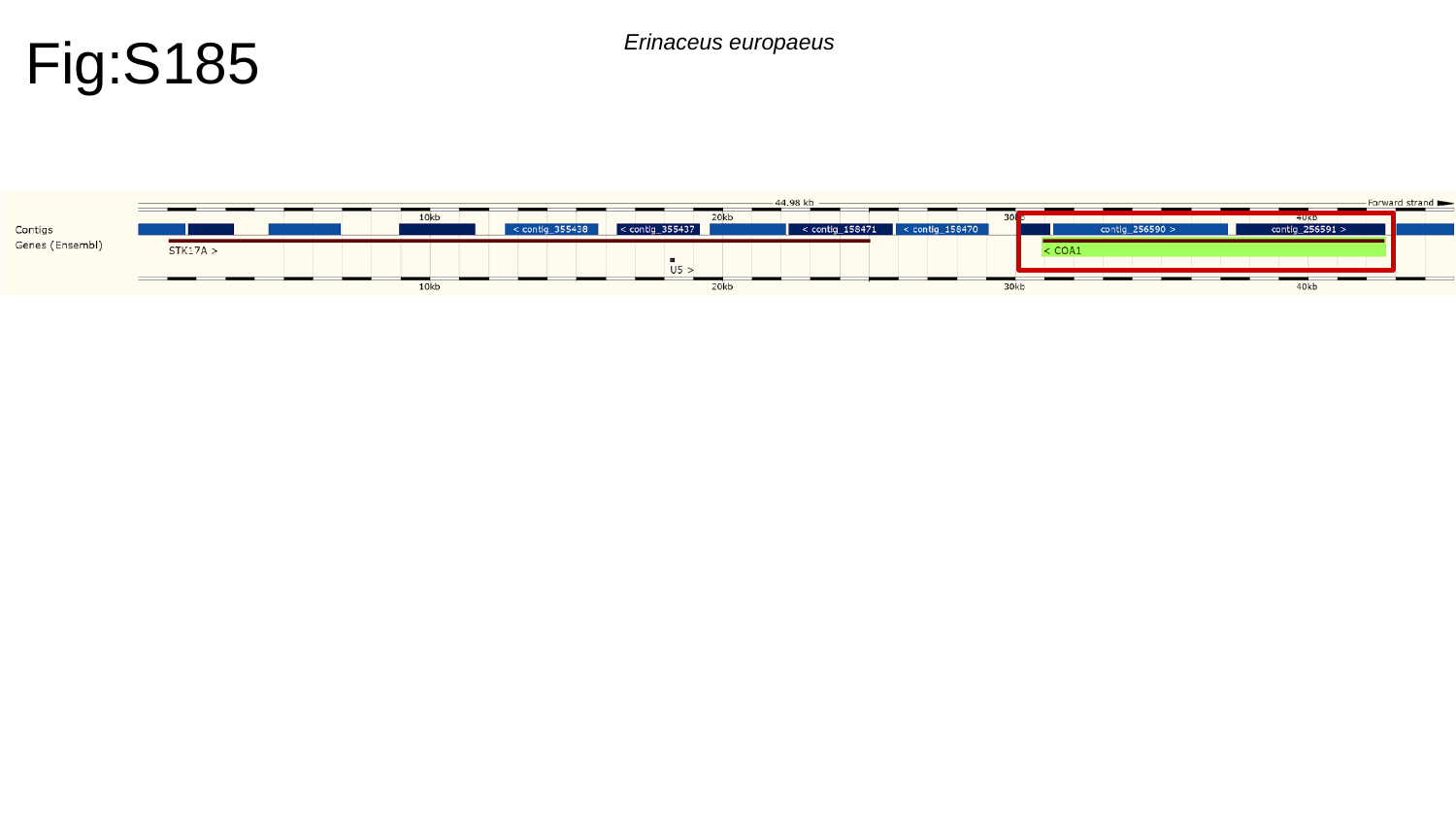

Fig:S185
Erinaceus europaeus

### Slide 186
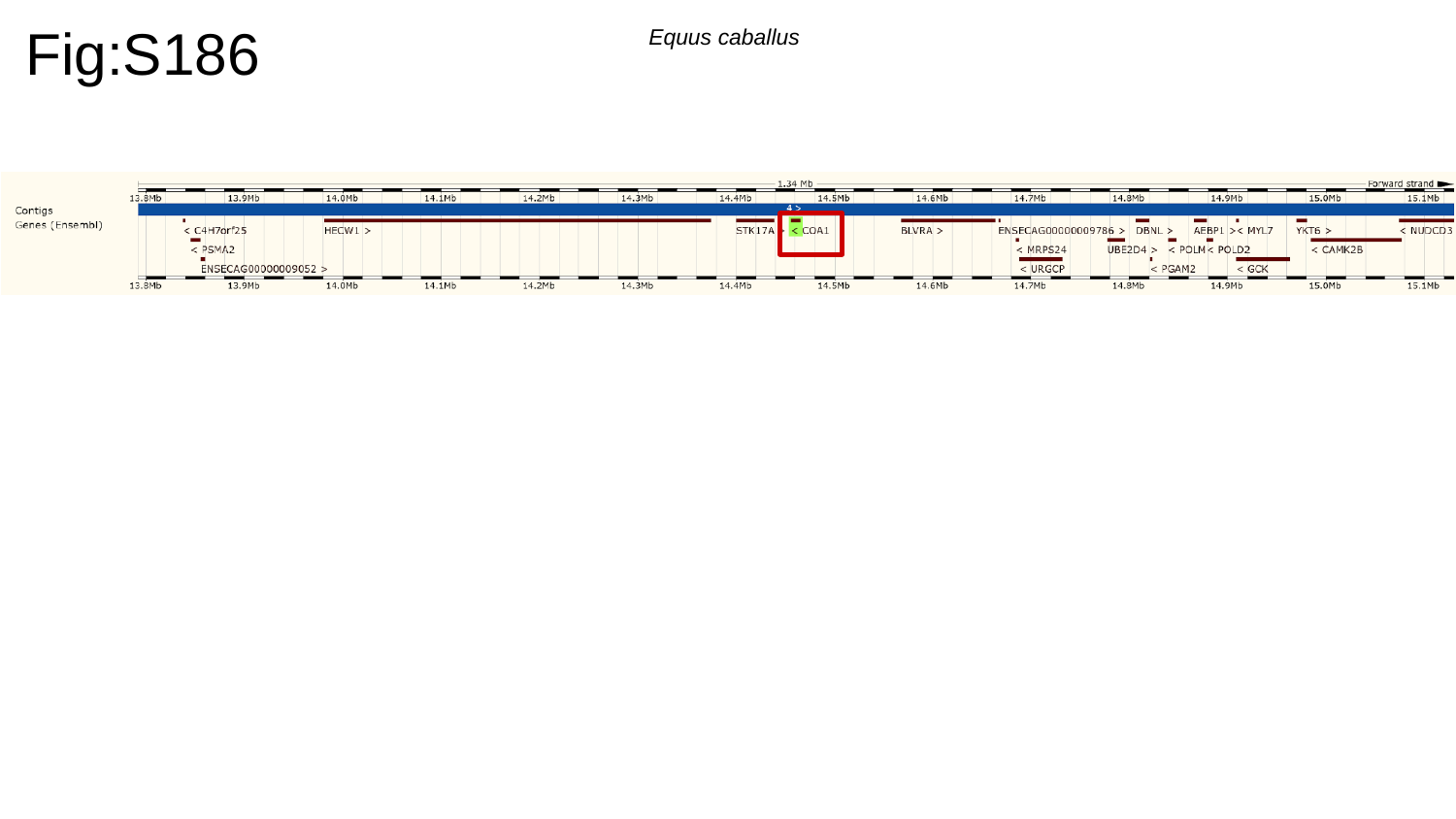

Fig:S186
Equus caballus

### Slide 187
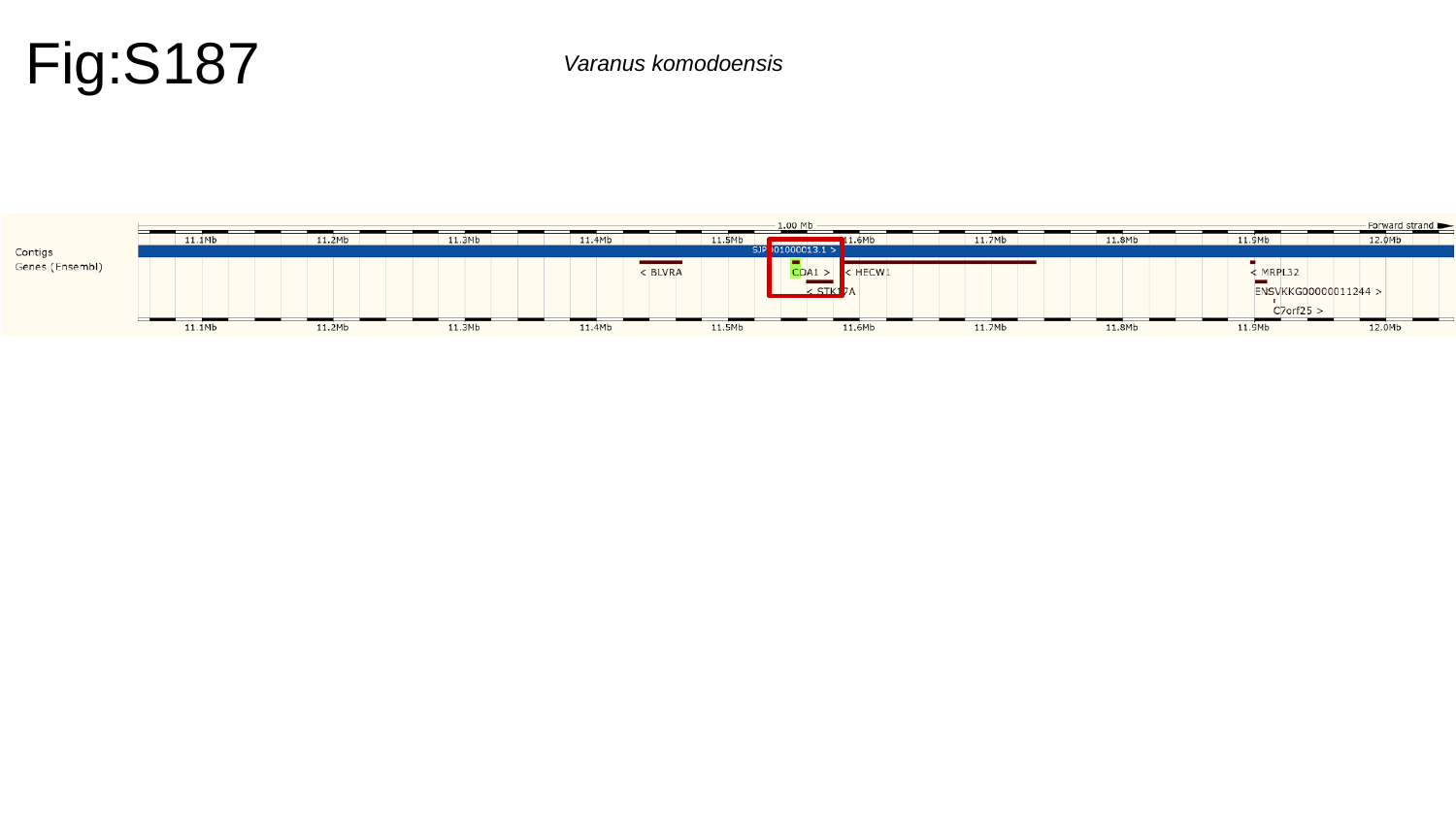

Fig:S187
Varanus komodoensis

### Slide 188
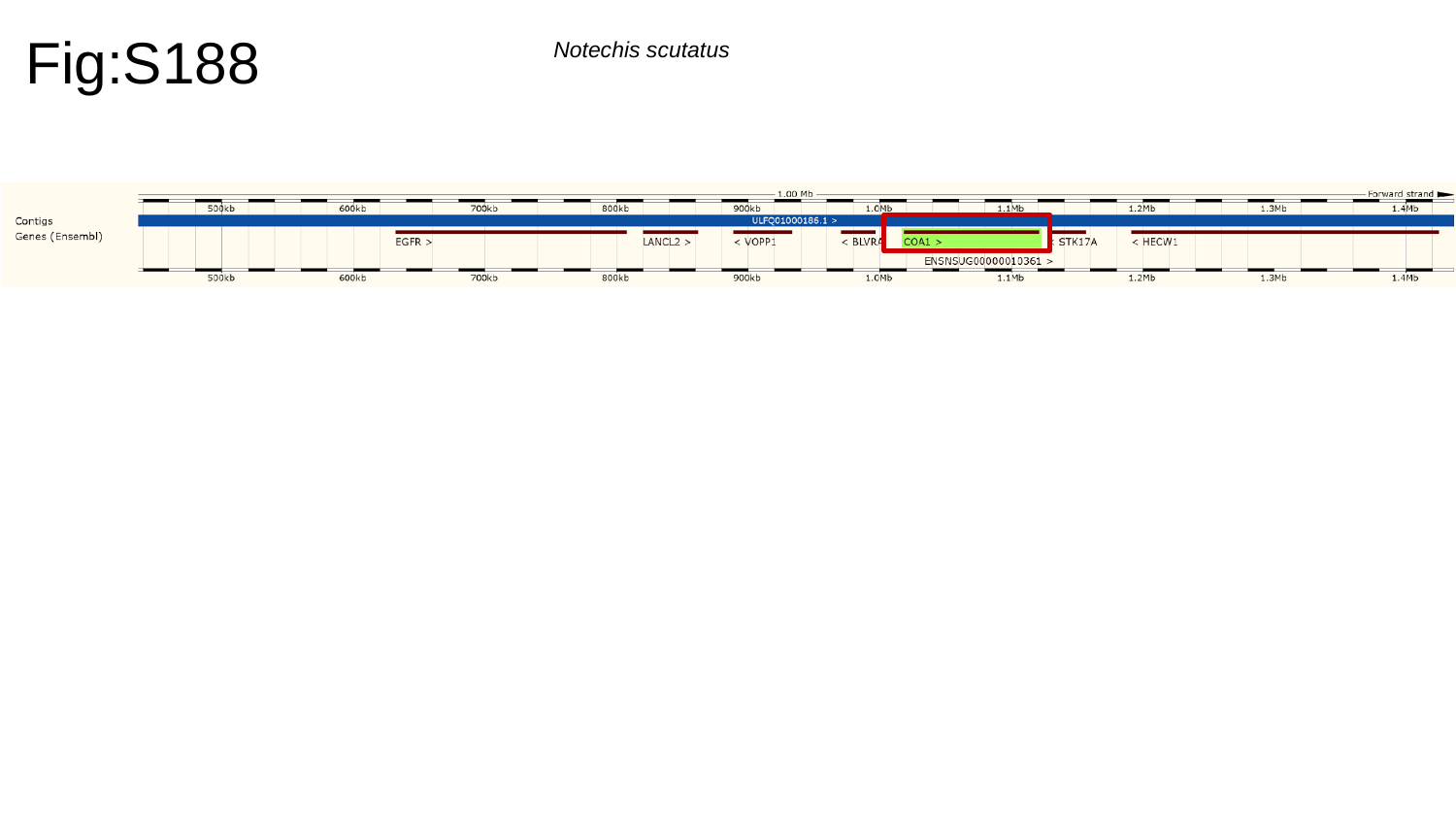

Fig:S188
Notechis scutatus

### Slide 189
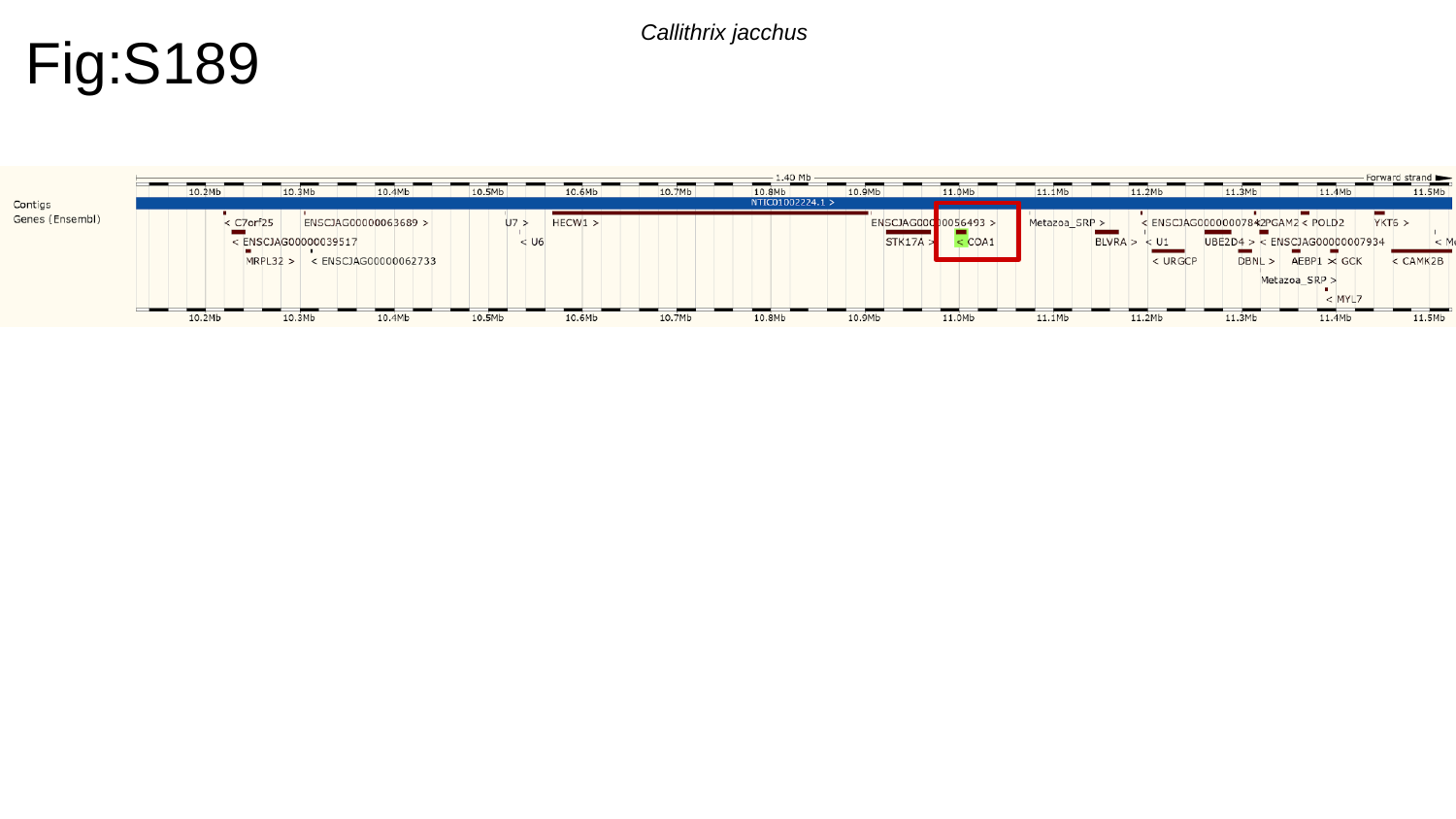

Fig:S189
Callithrix jacchus

### Slide 190
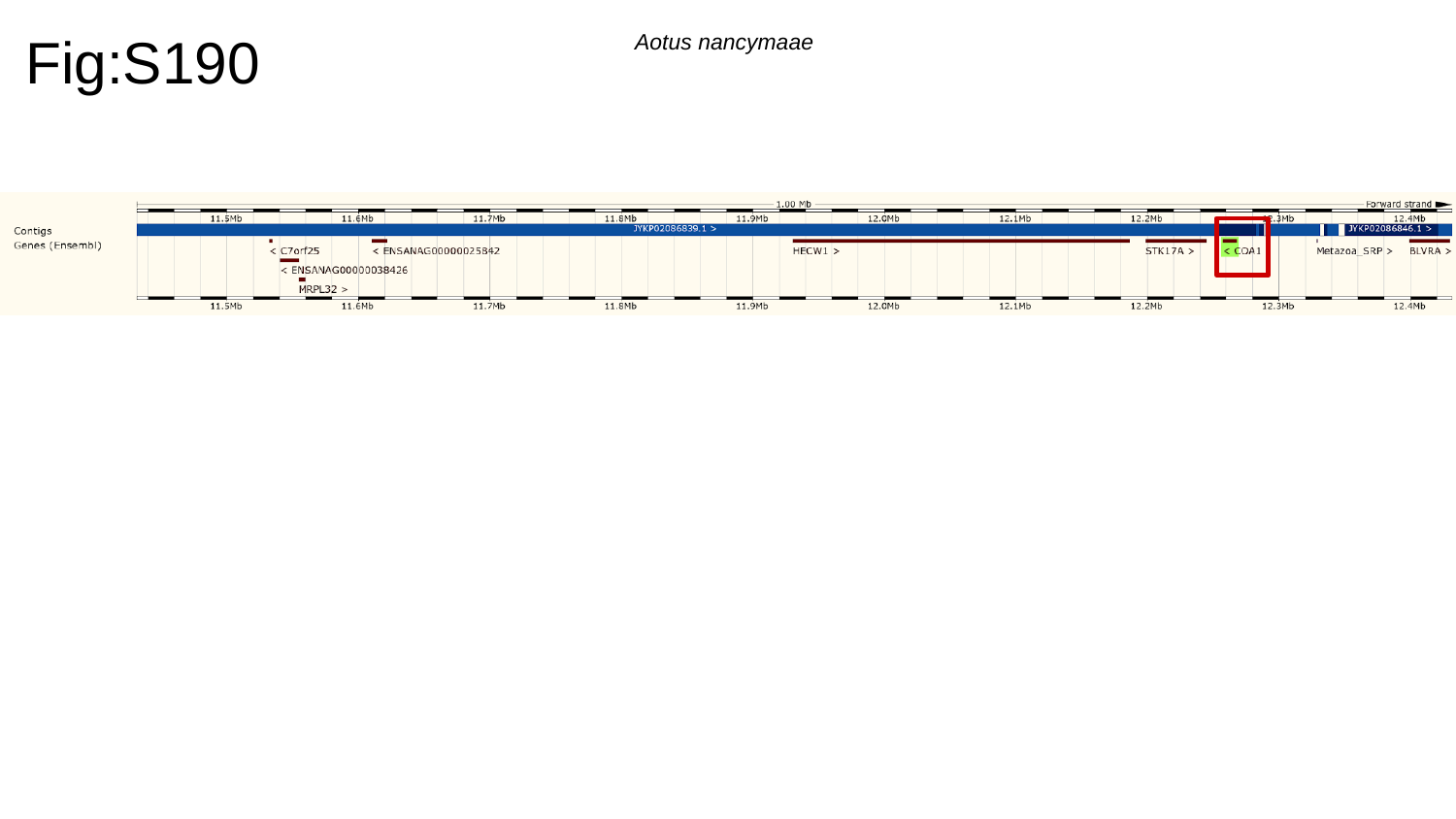

Fig:S190
Aotus nancymaae

### Slide 191
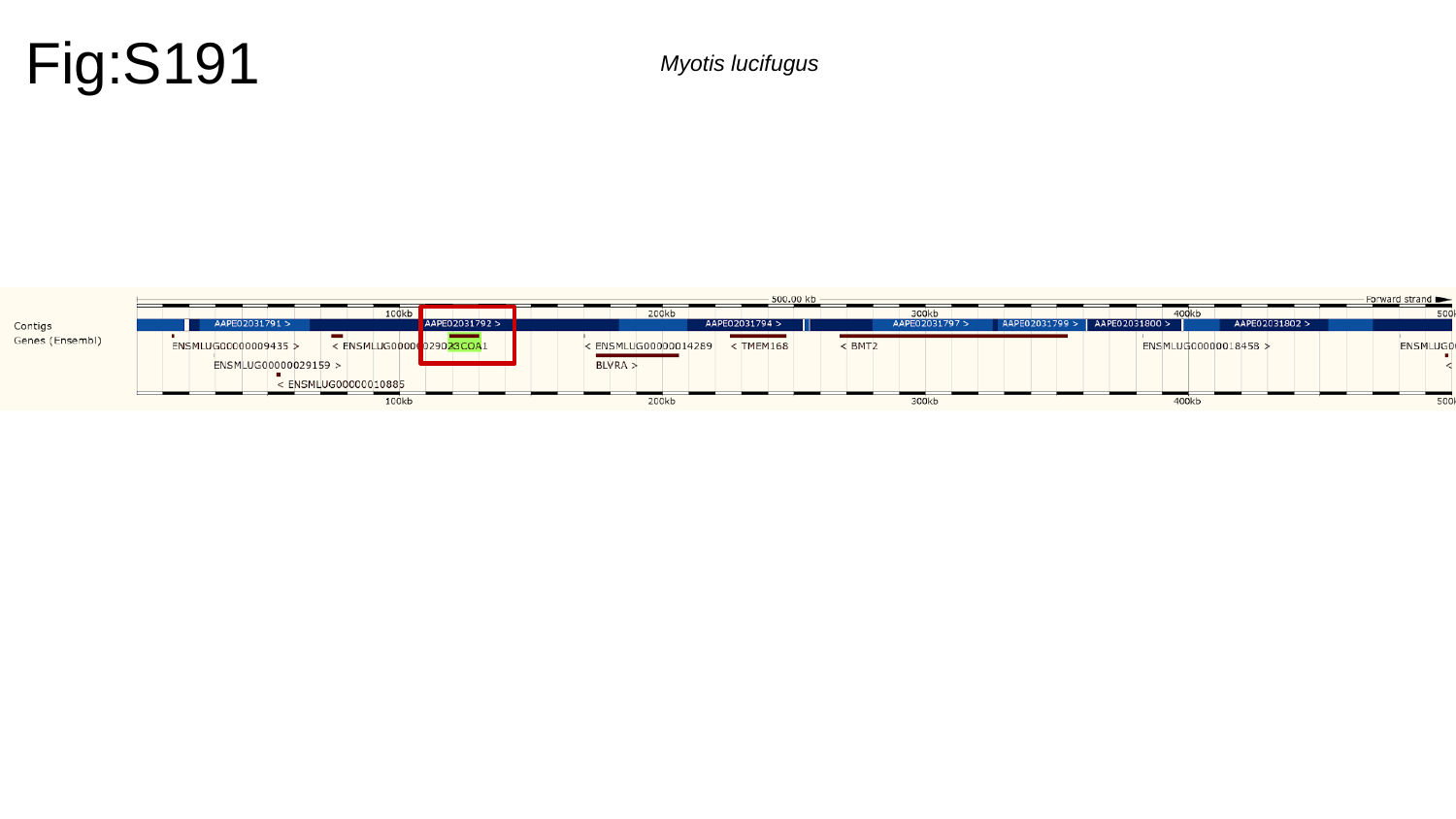

Fig:S191
Myotis lucifugus

### Slide 192
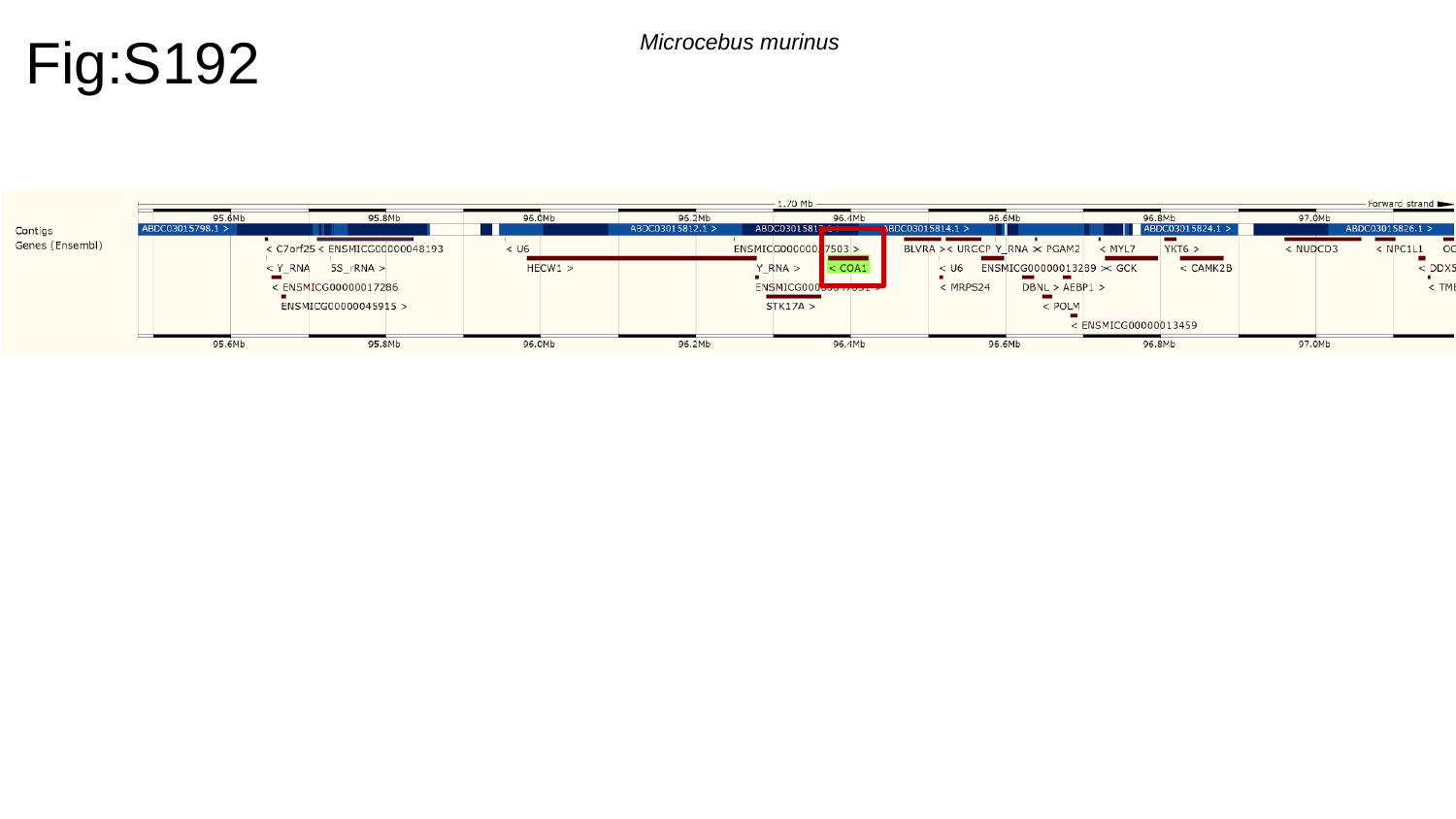

Fig:S192
Microcebus murinus

### Slide 193
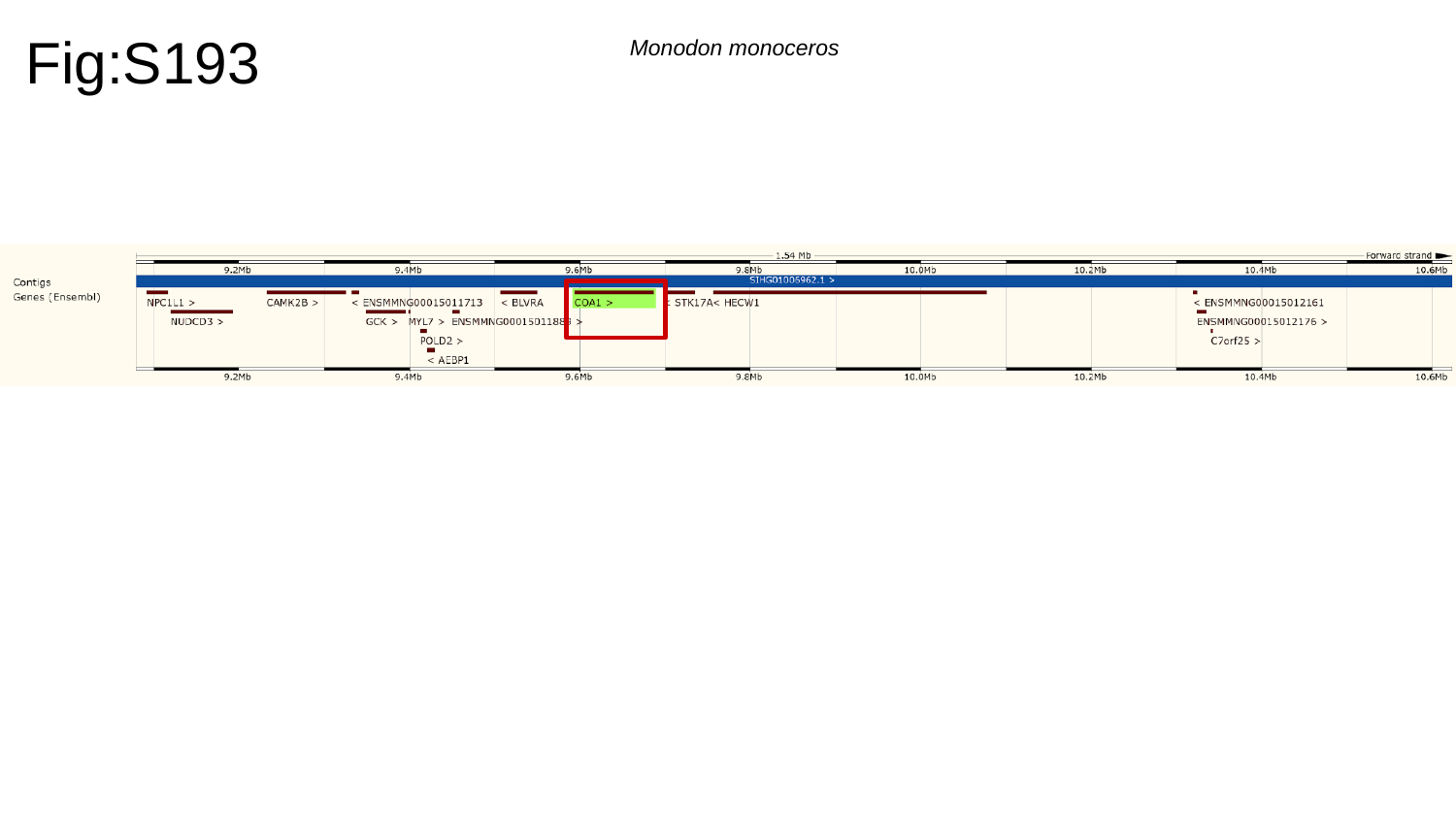

Fig:S193
Monodon monoceros

### Slide 194
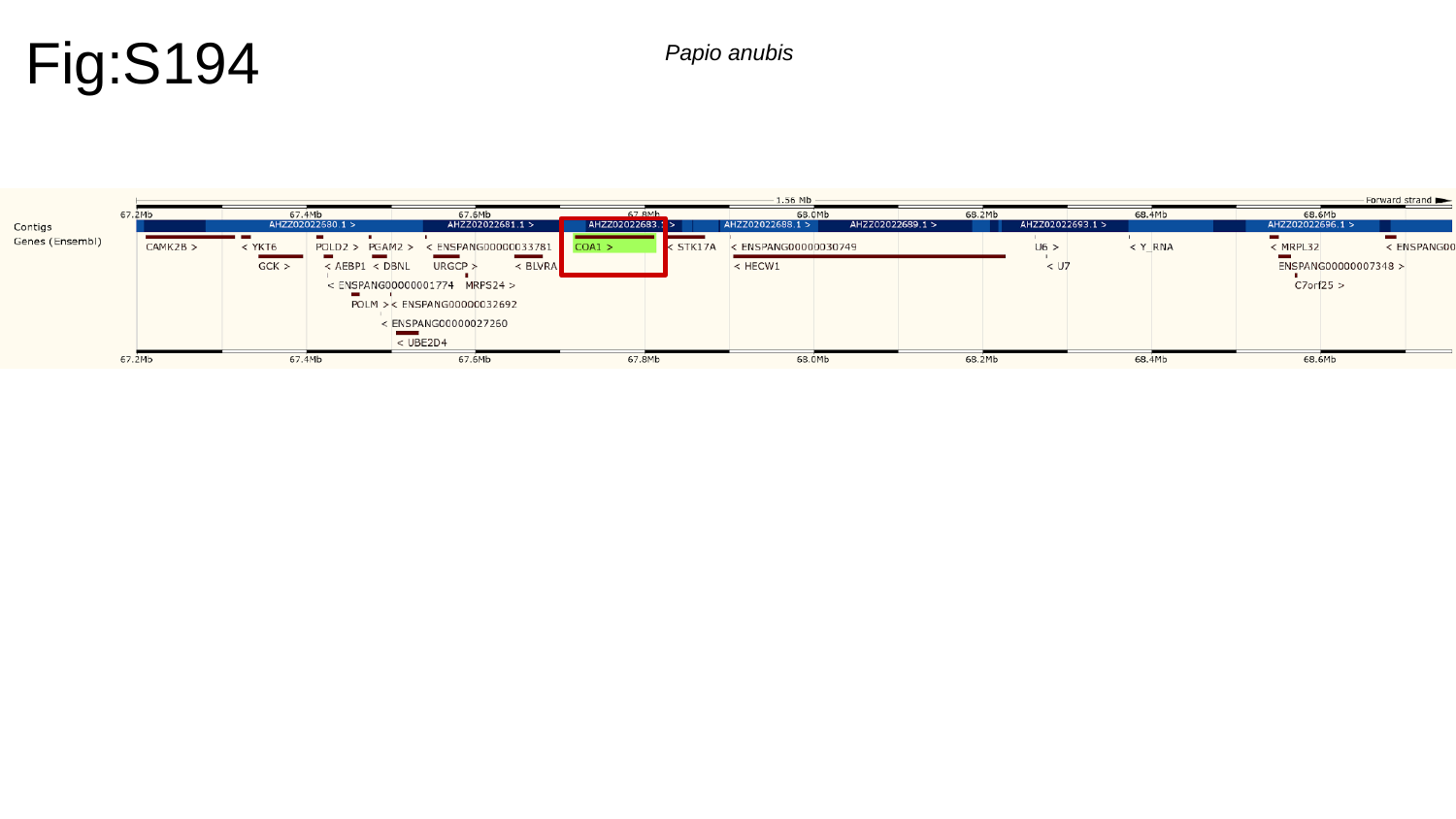

Fig:S194
Papio anubis

### Slide 195
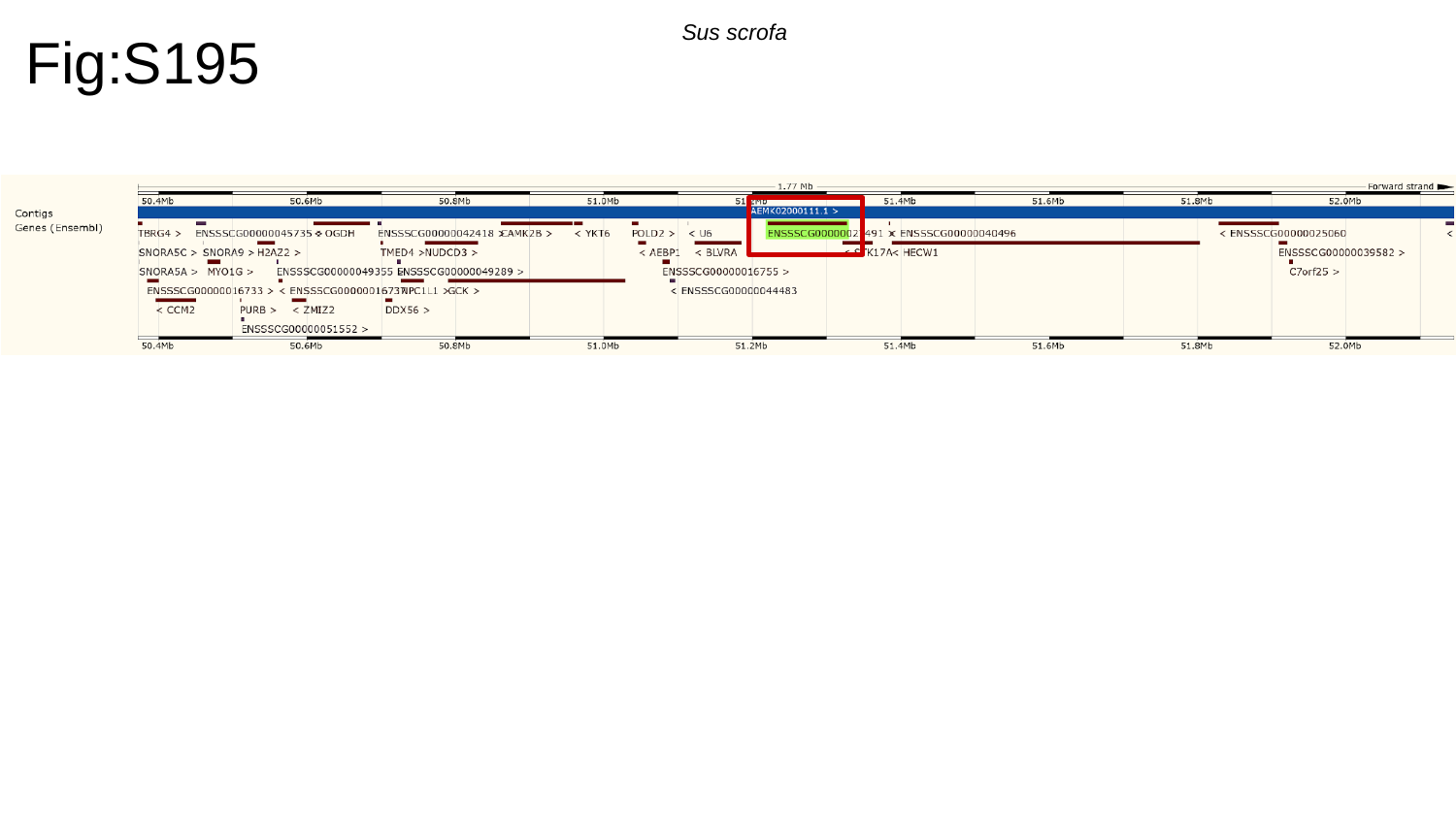

Fig:S195
Sus scrofa

### Slide 196
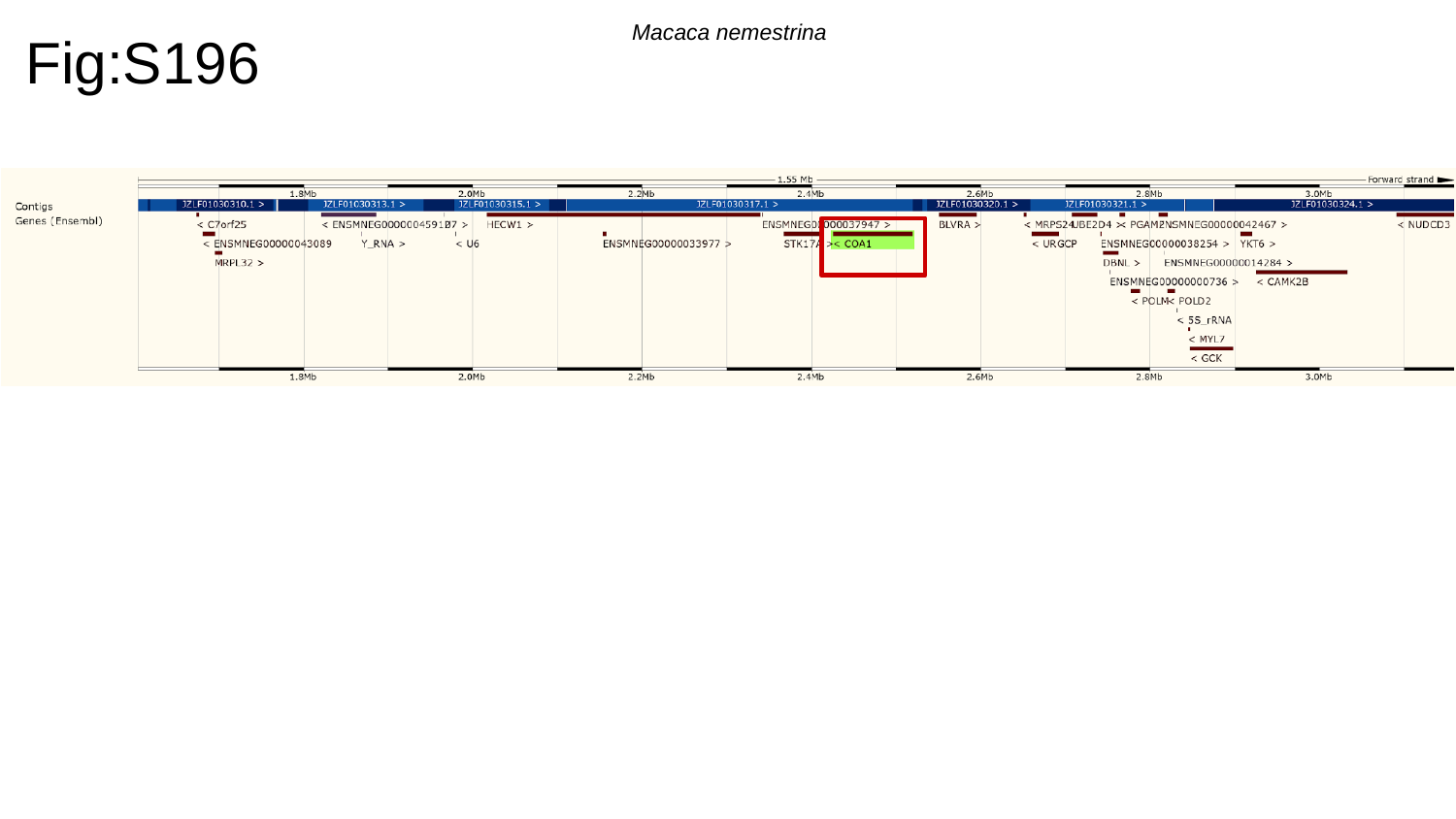

Fig:S196
Macaca nemestrina

### Slide 197
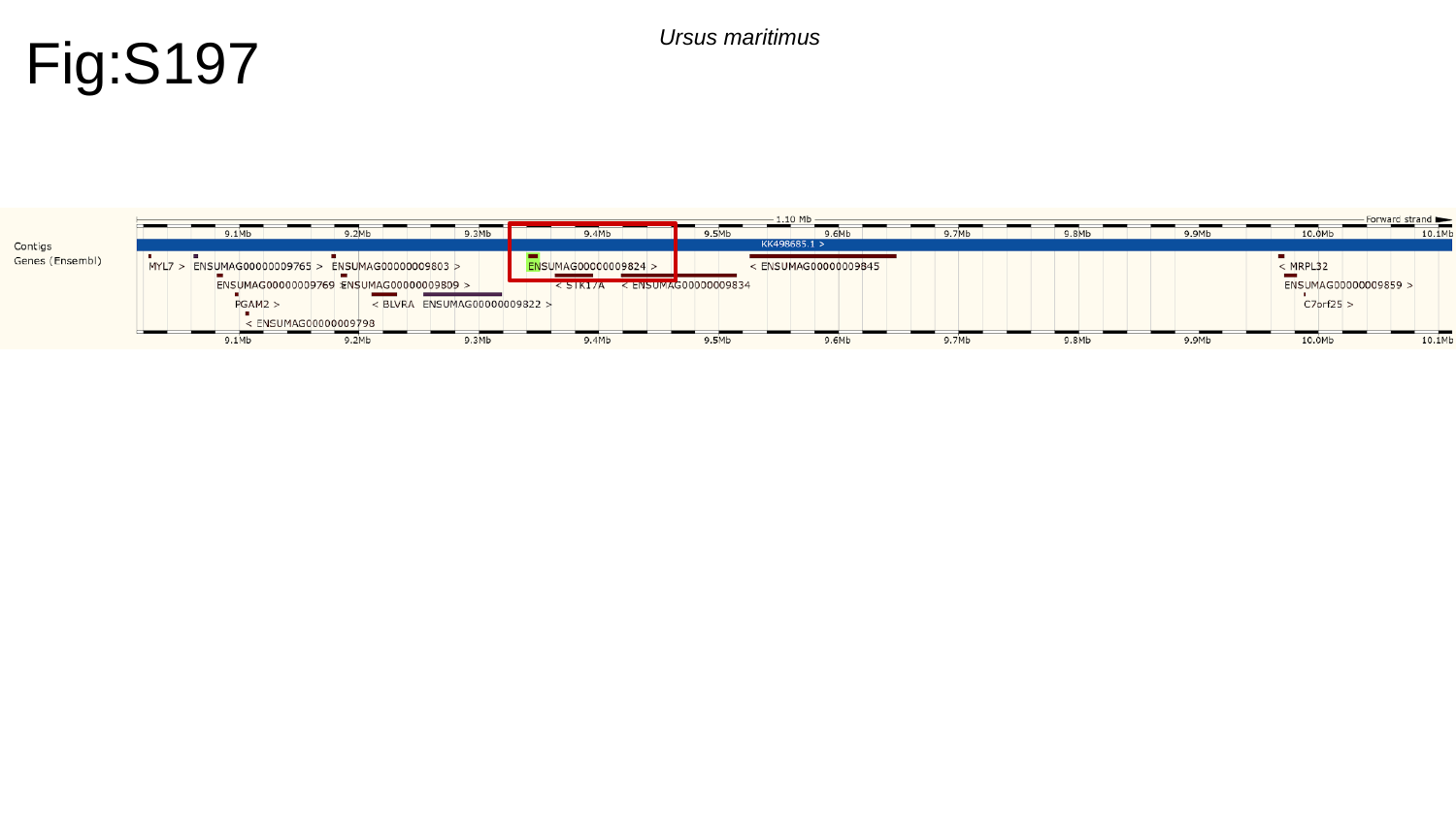

Fig:S197
Ursus maritimus

### Slide 198
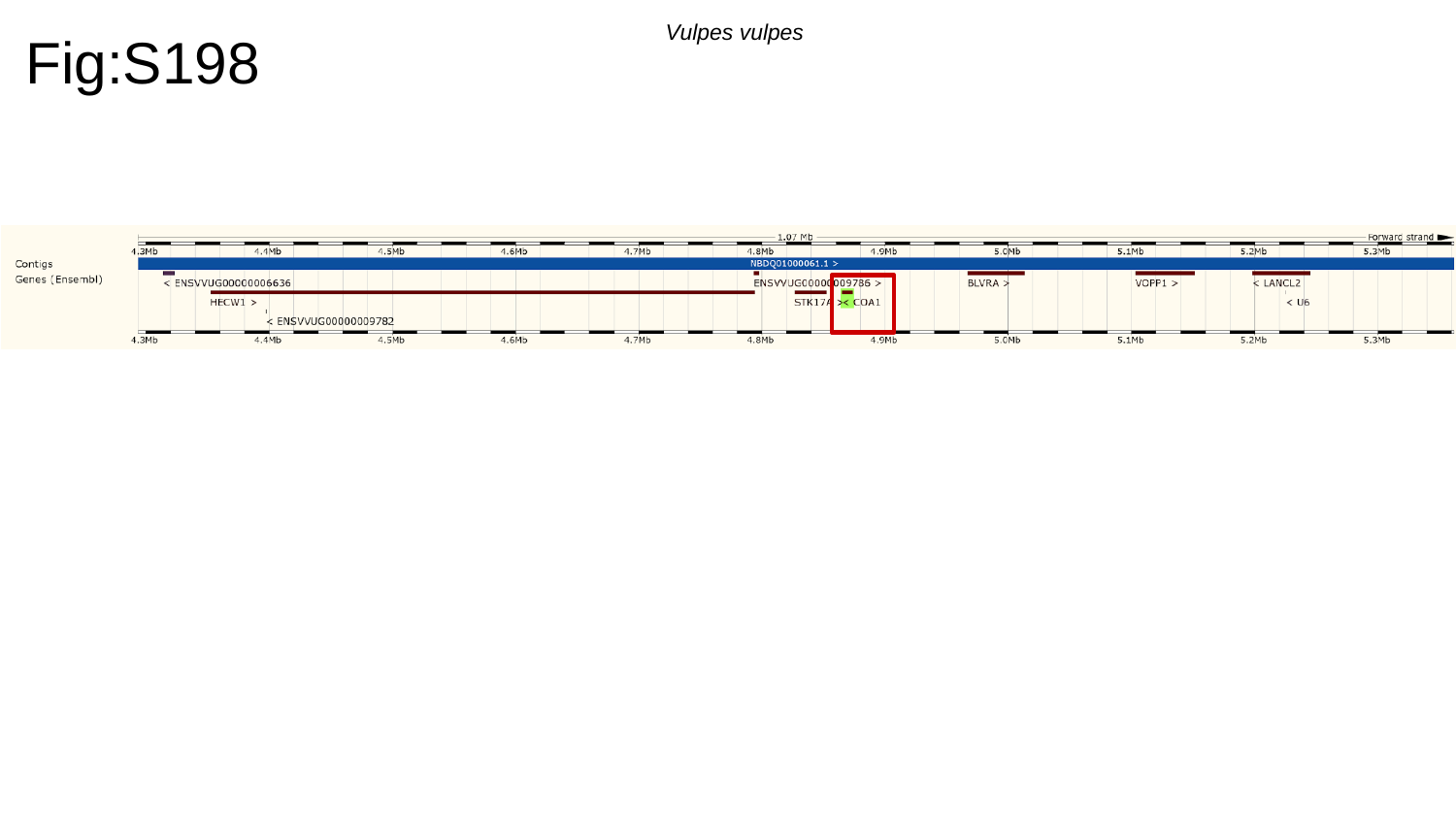

Fig:S198
Vulpes vulpes

### Slide 199
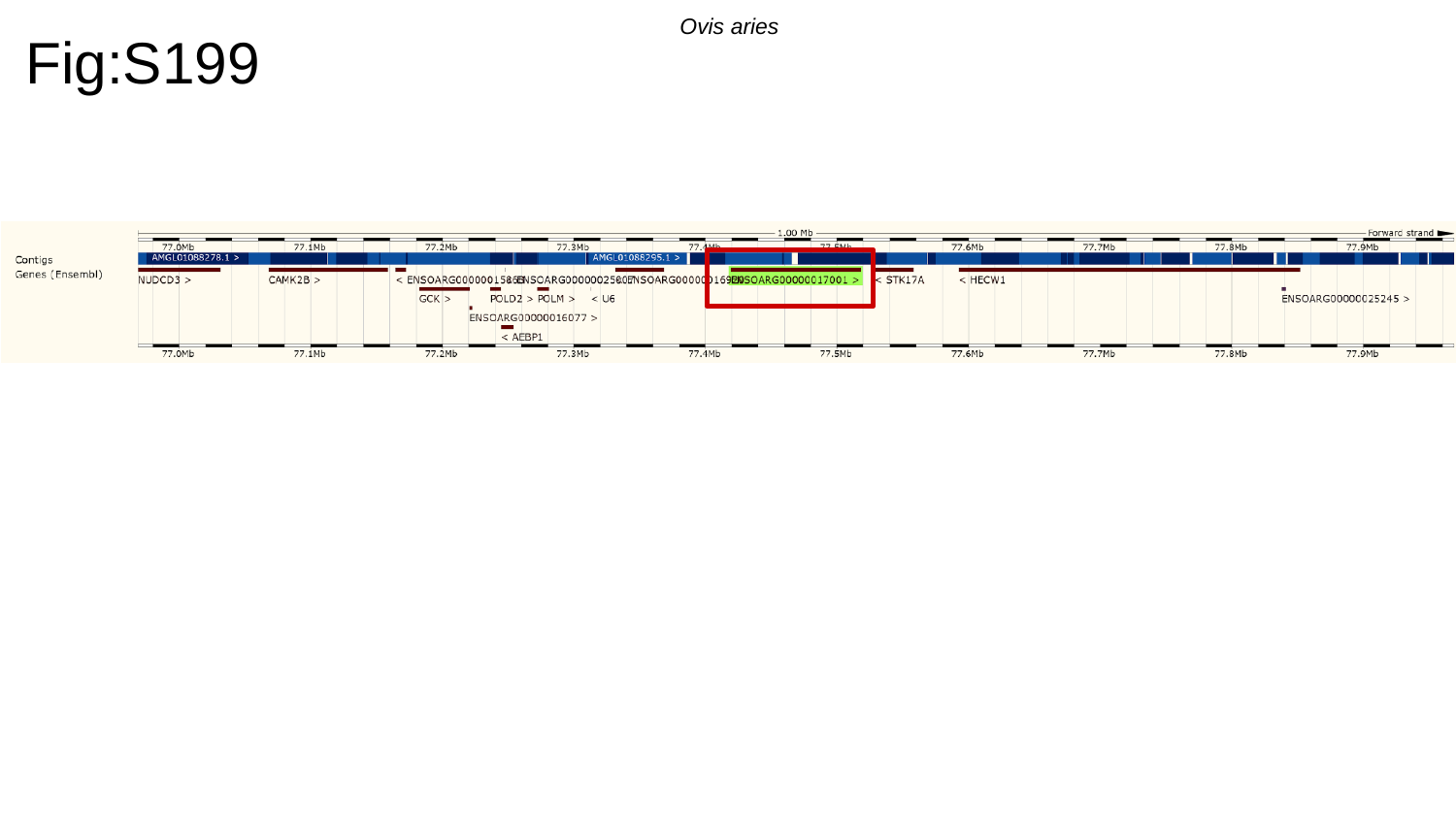

Fig:S199
Ovis aries

### Slide 200
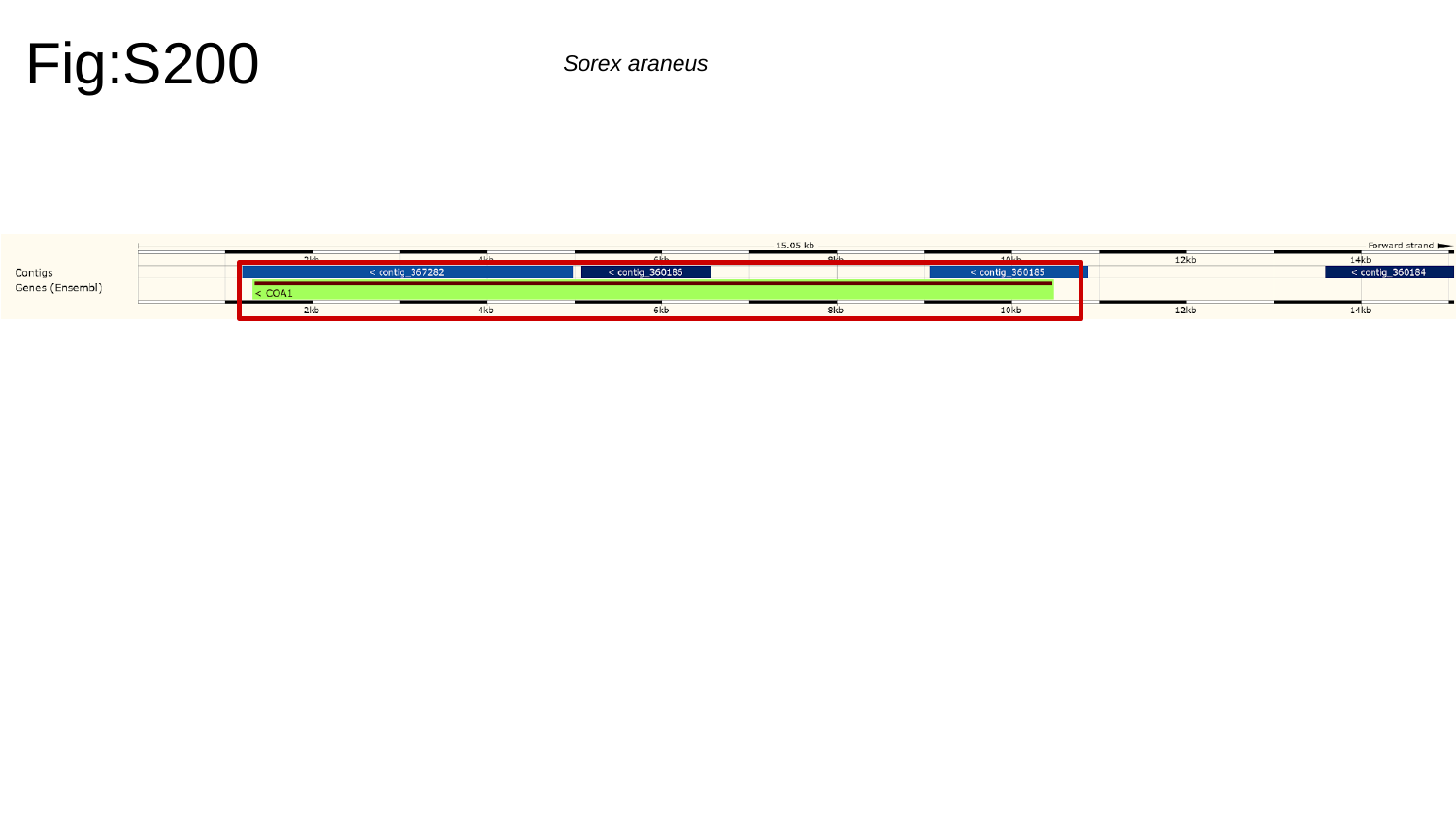

Fig:S200
Sorex araneus

### Slide 201
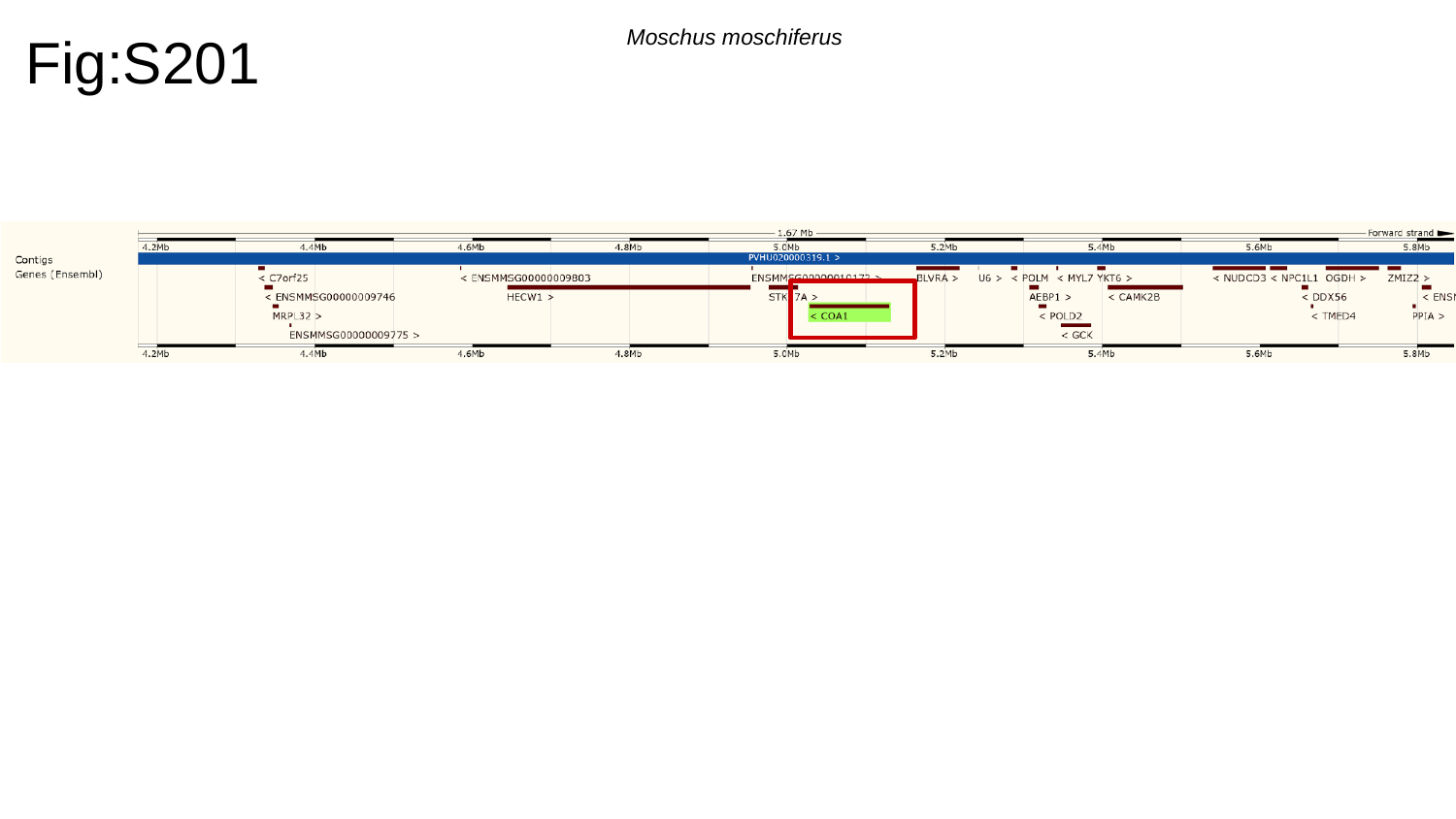

Fig:S201
Moschus moschiferus

### Slide 202
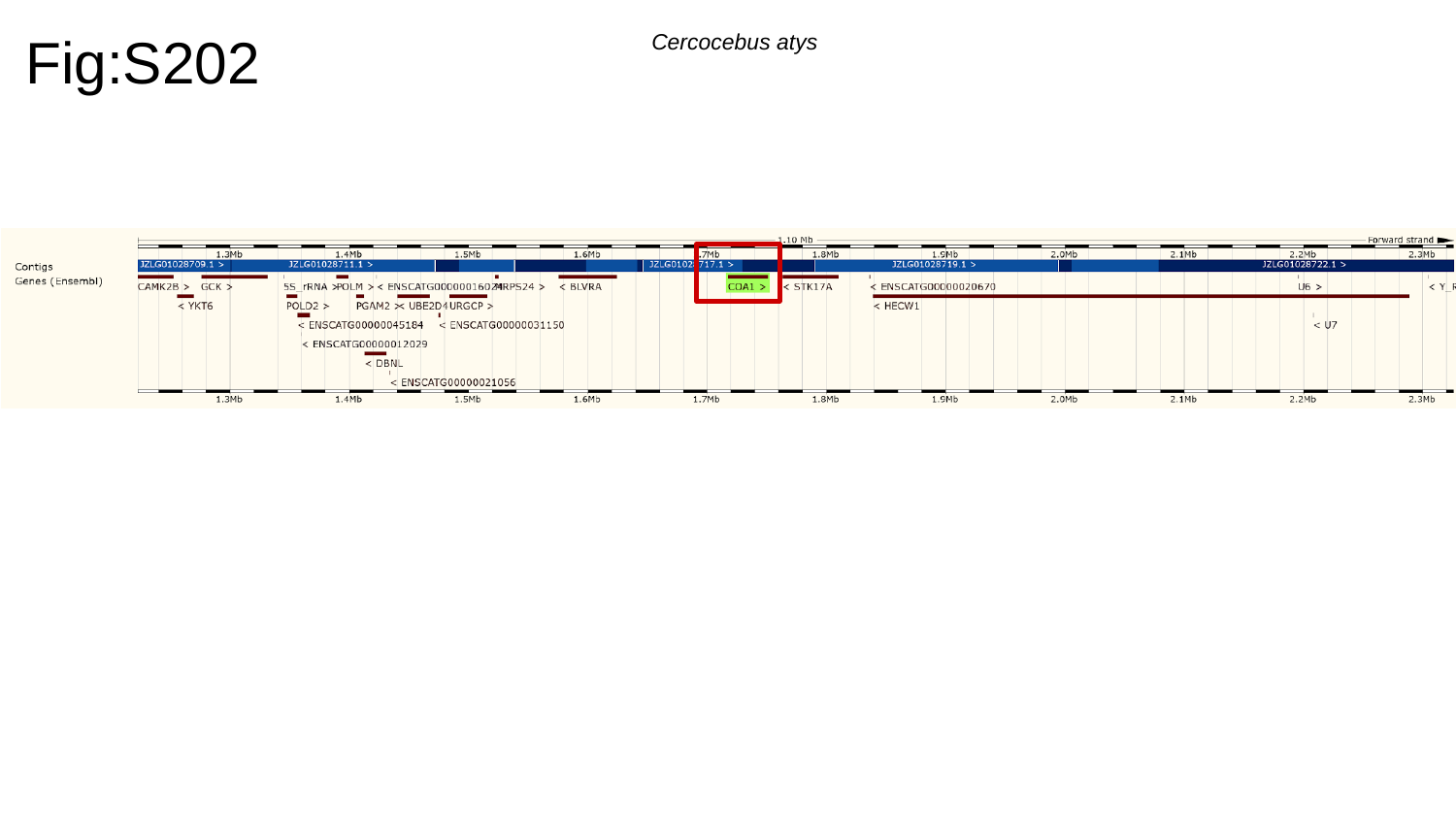

Fig:S202
Cercocebus atys

### Slide 203
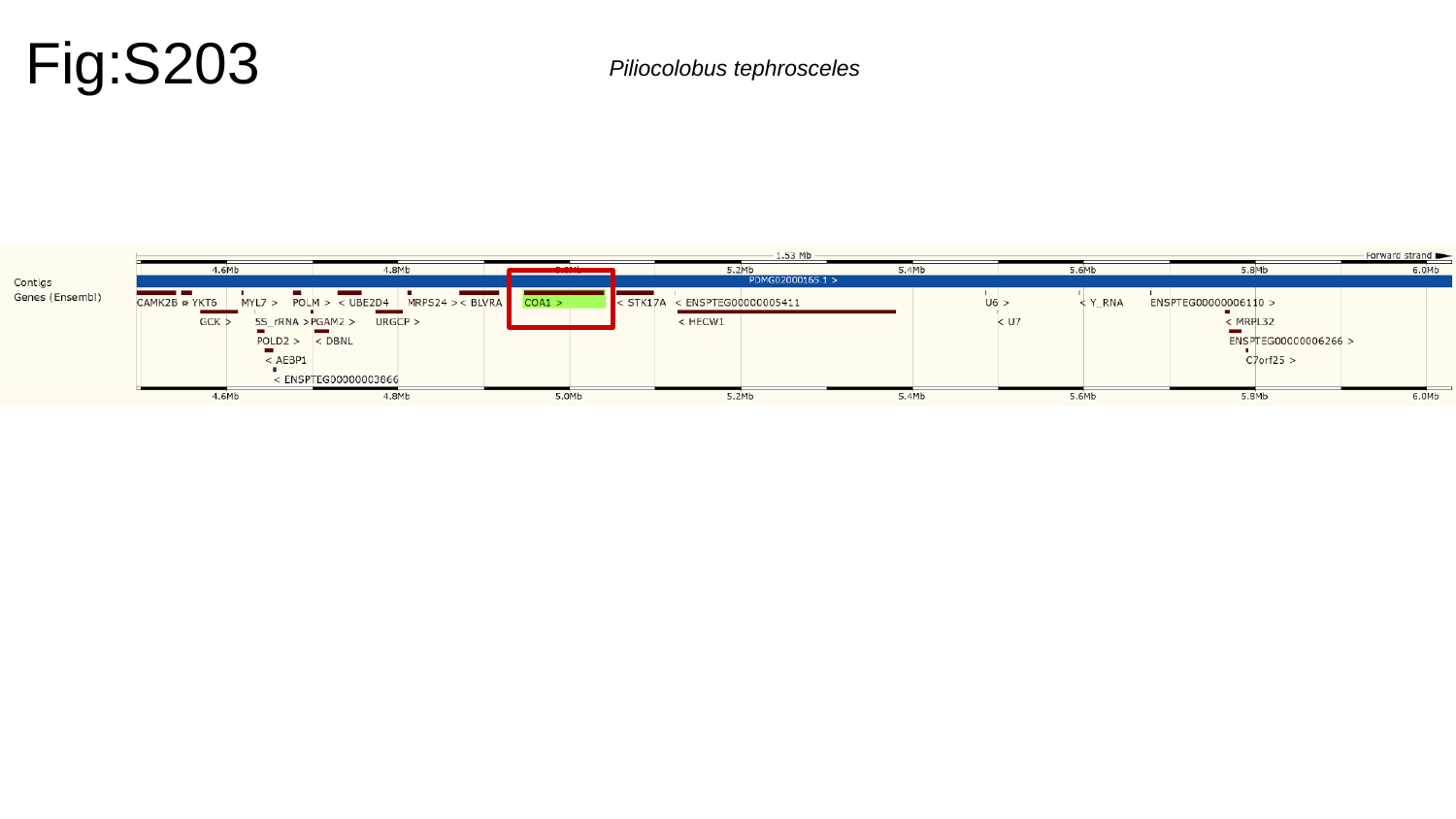

Fig:S203
Piliocolobus tephrosceles

### Slide 204
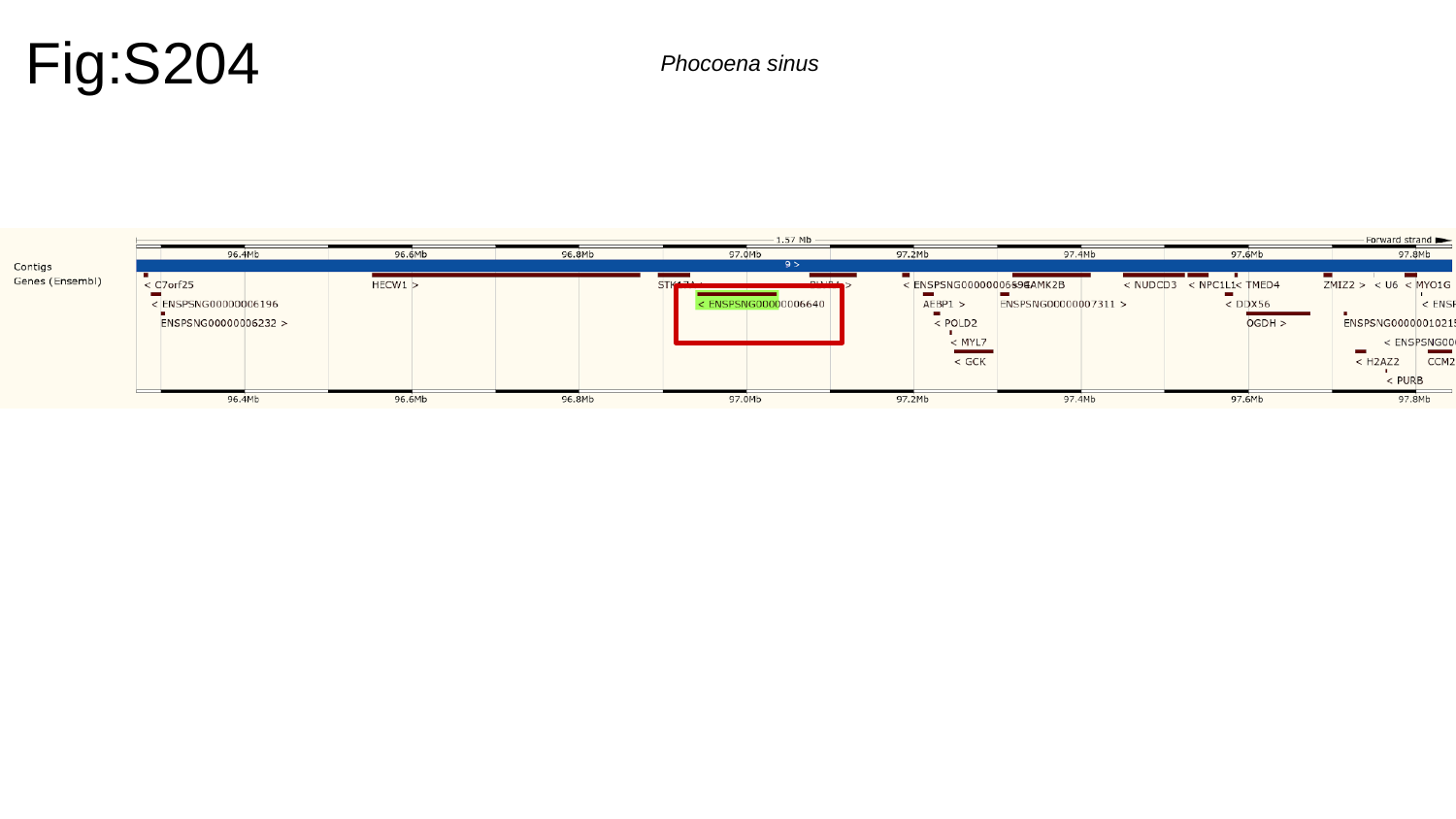

Fig:S204
Phocoena sinus

### Slide 205
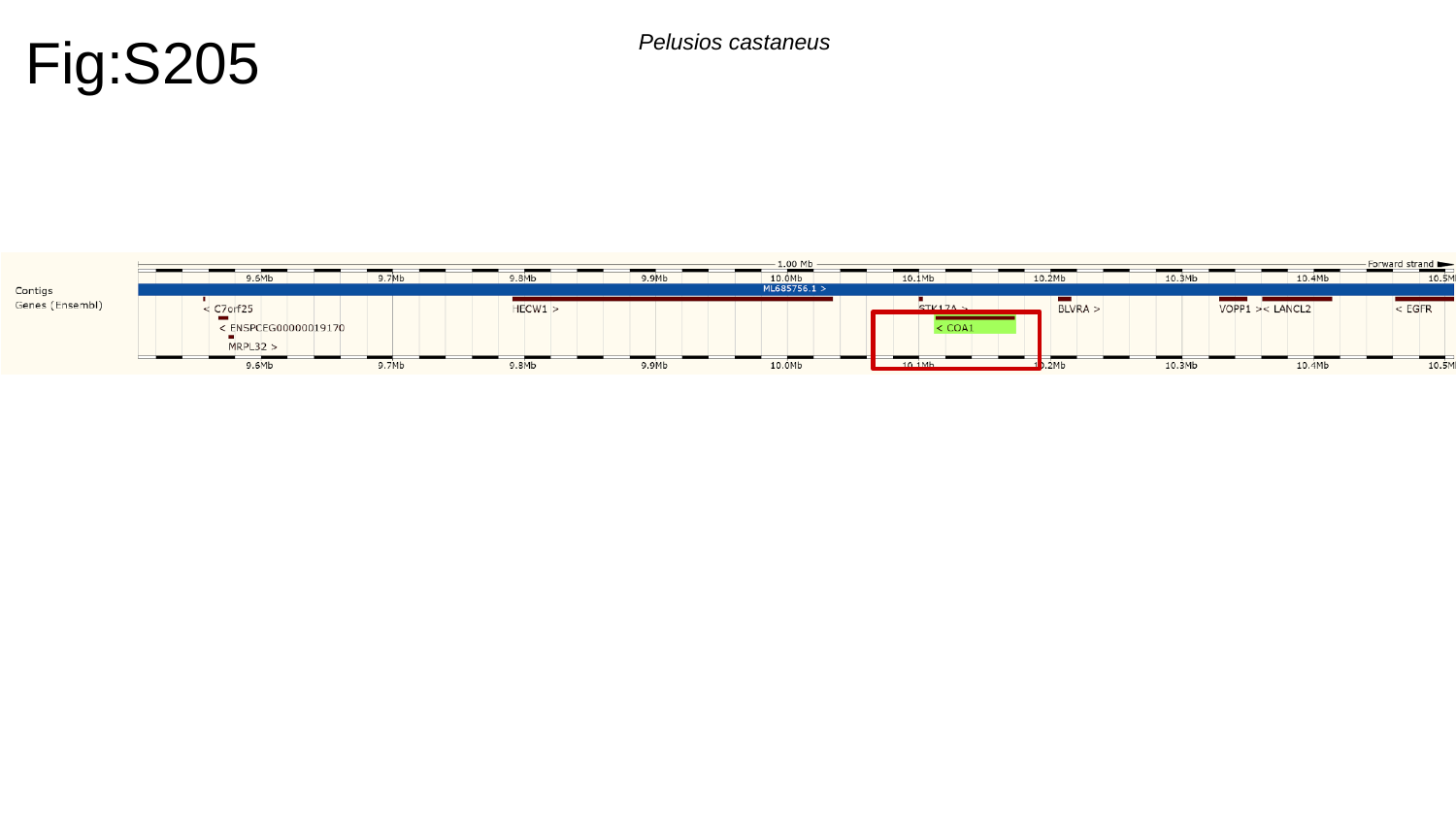

Fig:S205
Pelusios castaneus

### Slide 206
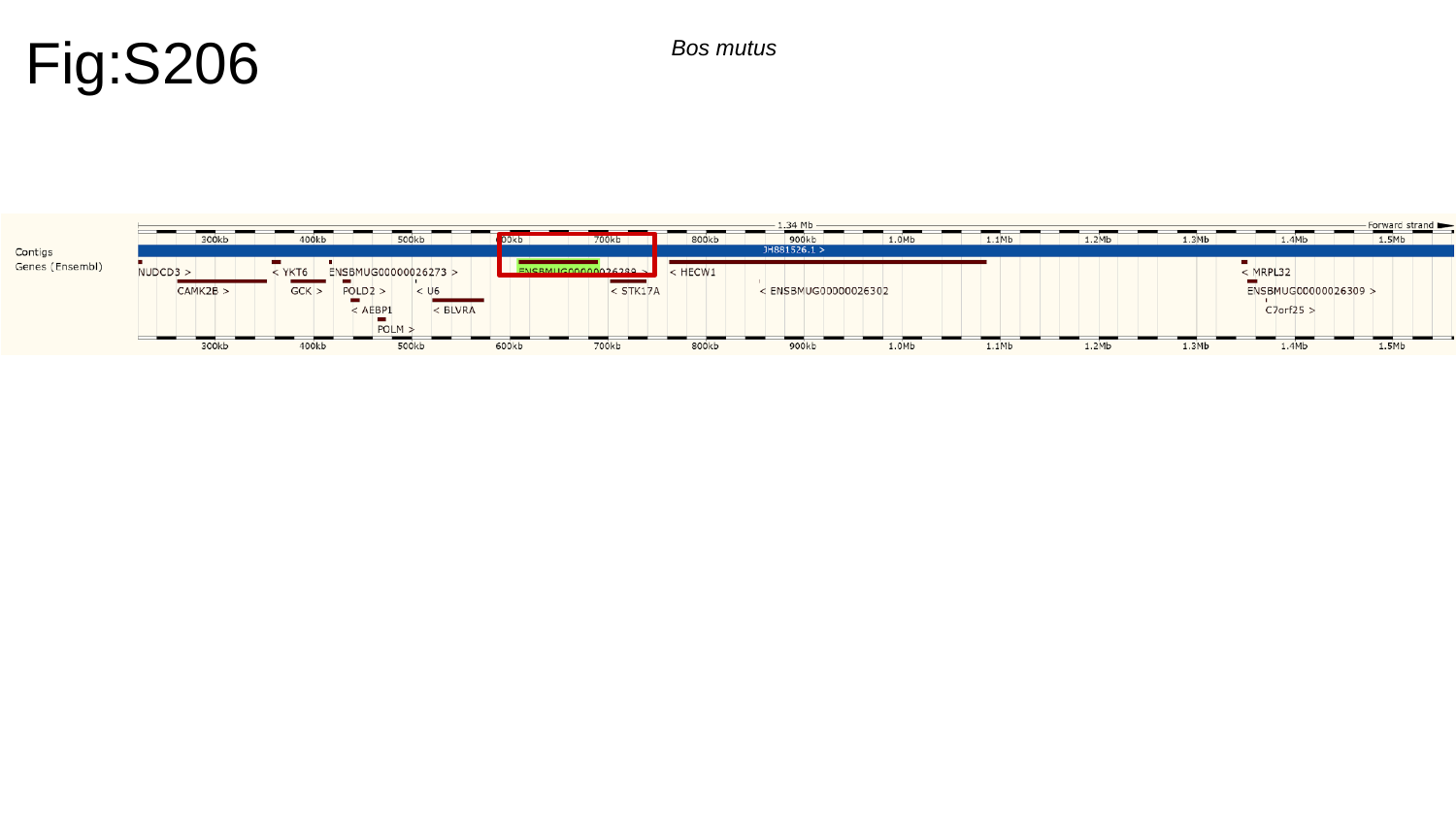

Fig:S206
Bos mutus

### Slide 207
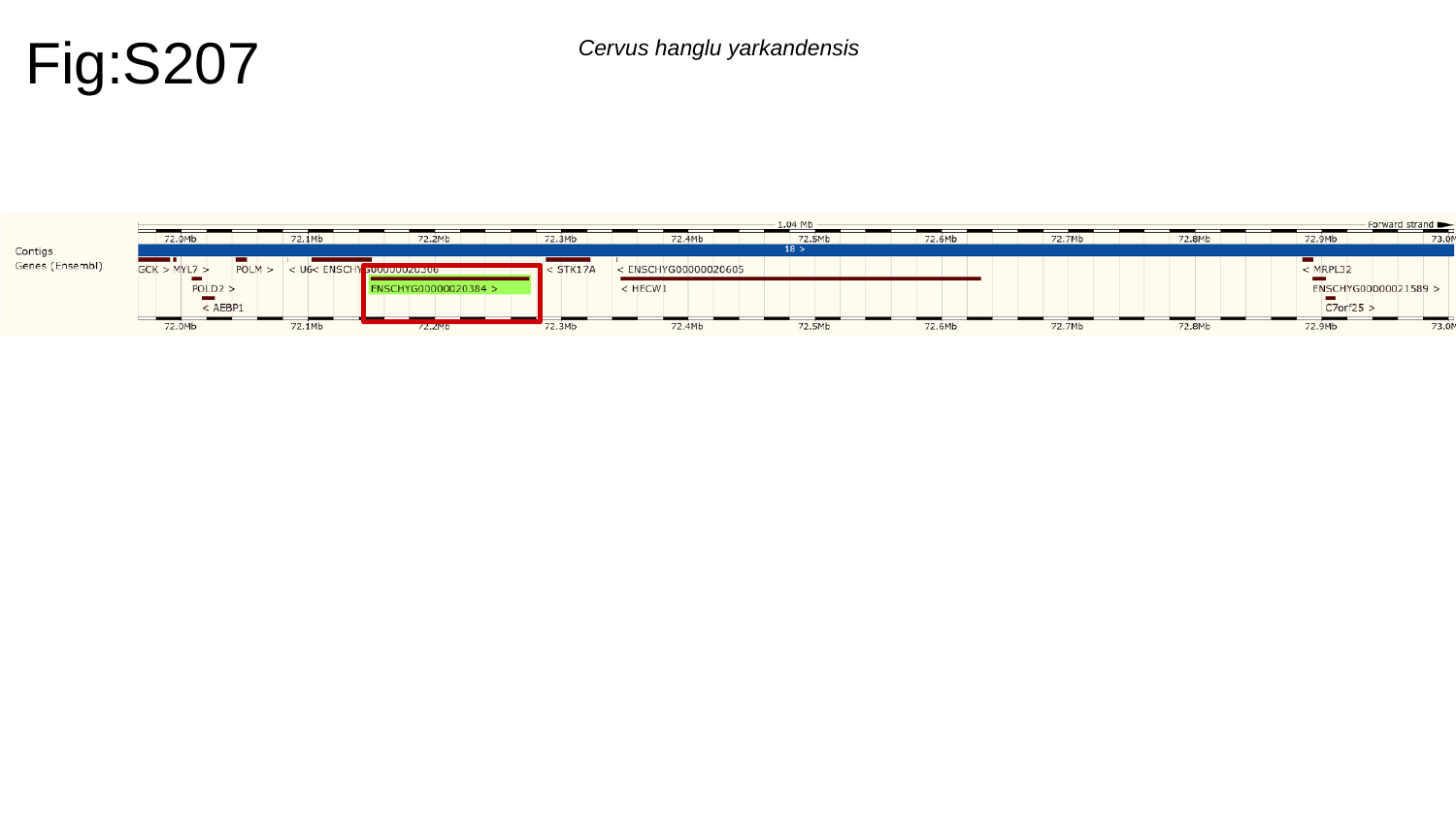

Fig:S207
Cervus hanglu yarkandensis

### Slide 208
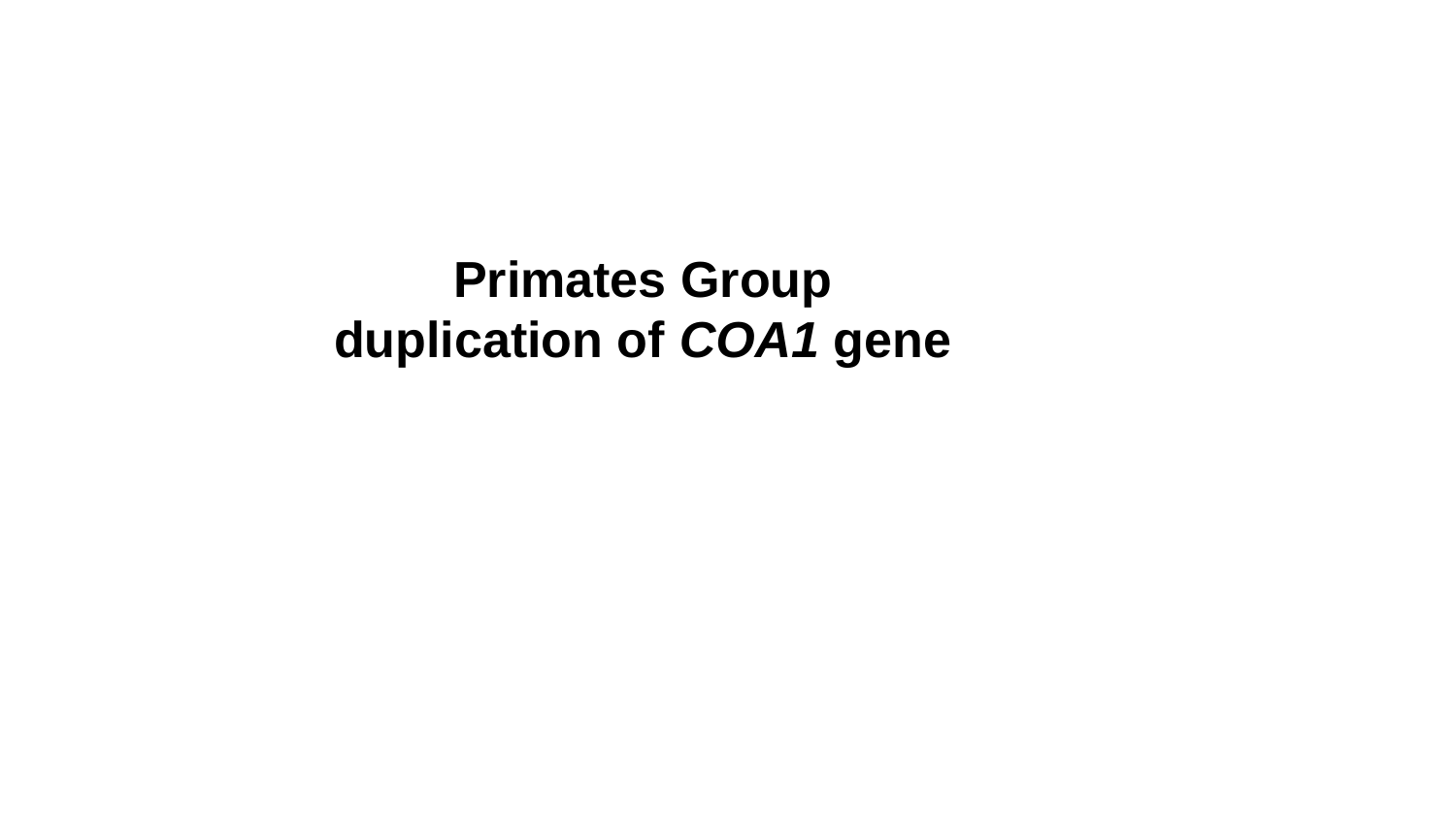

Primates Group duplication of COA1 gene

### Slide 209
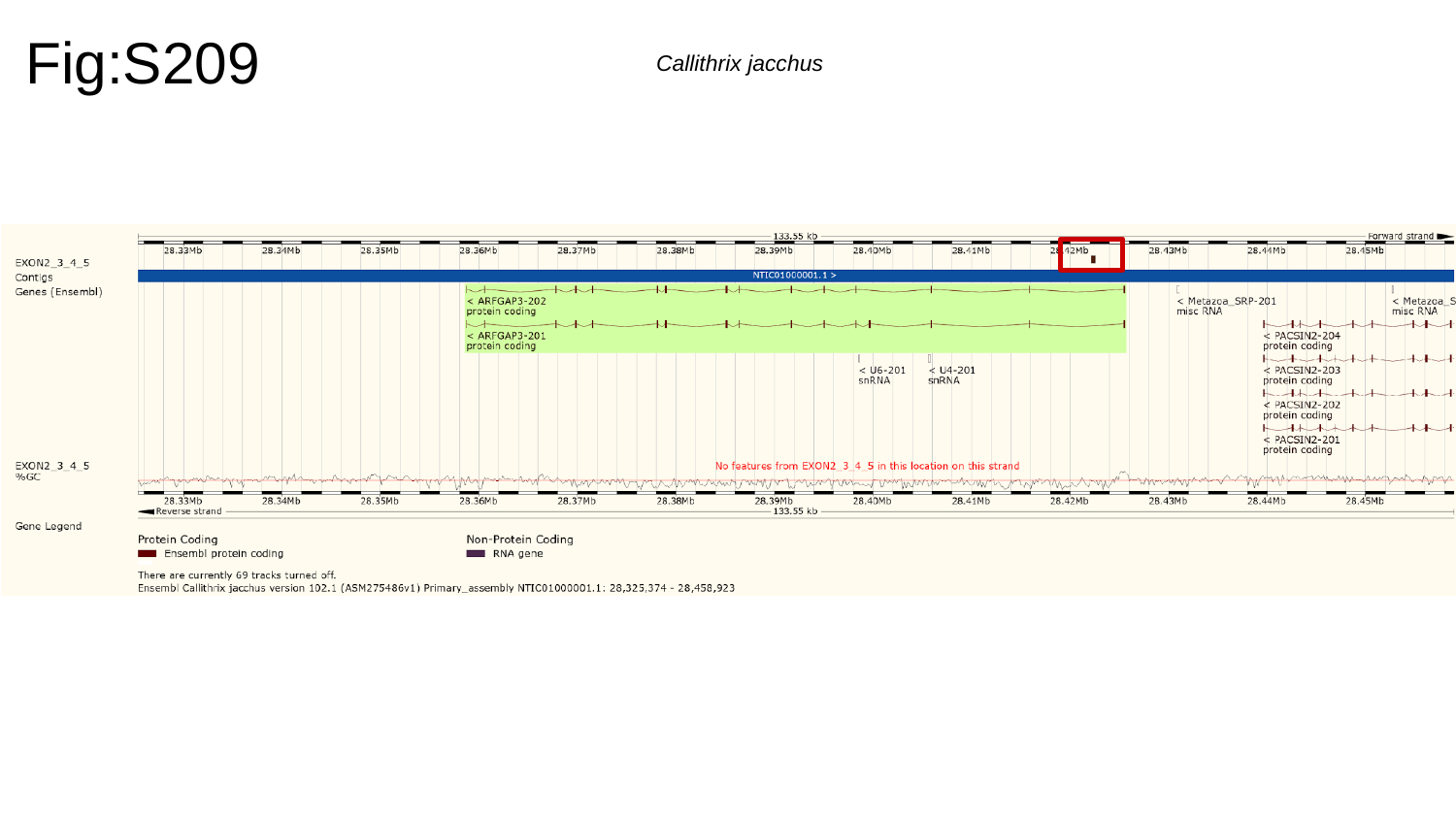

Fig:S209
Callithrix jacchus

### Slide 210
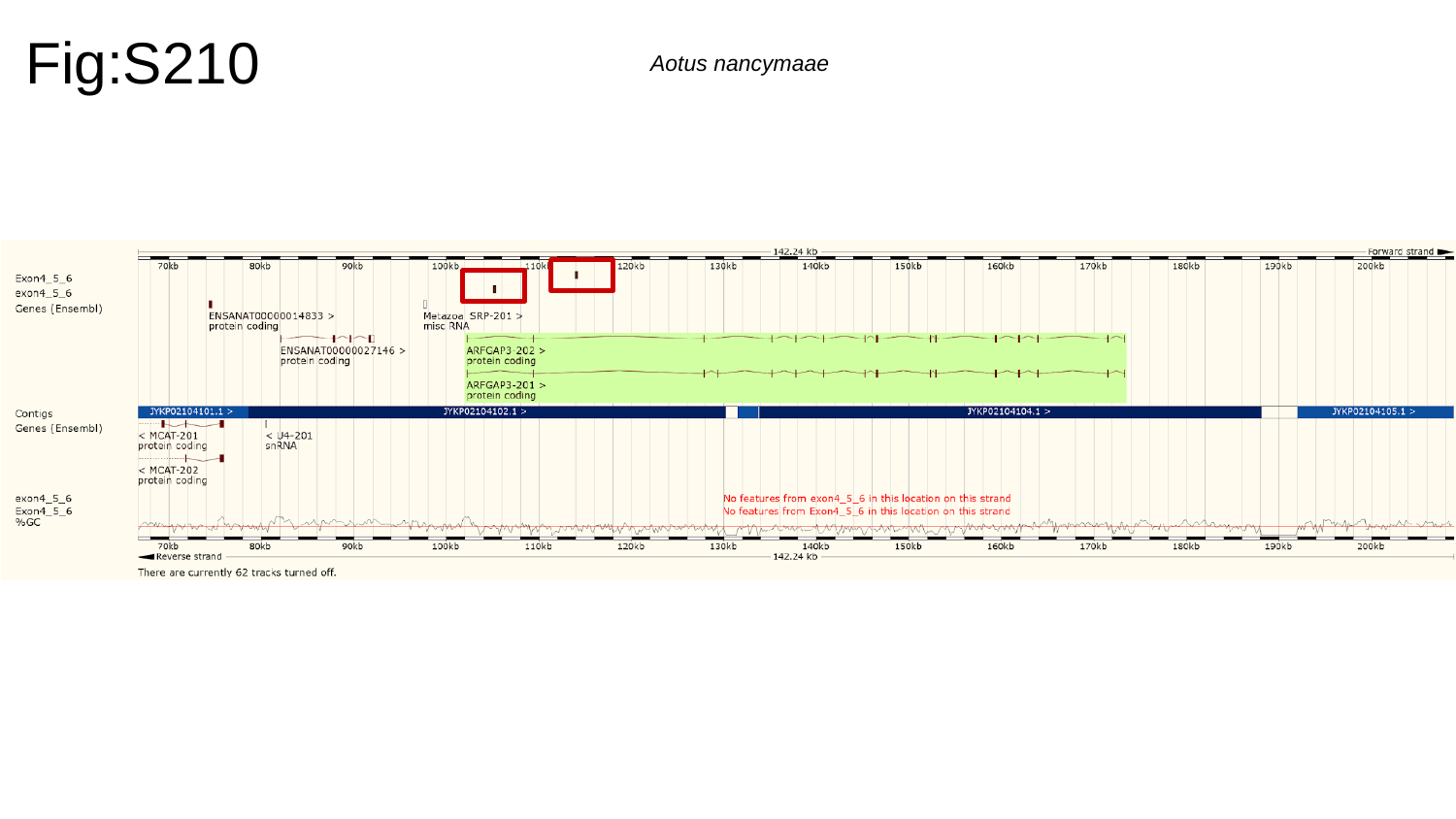

Fig:S210
Aotus nancymaae

### Slide 211
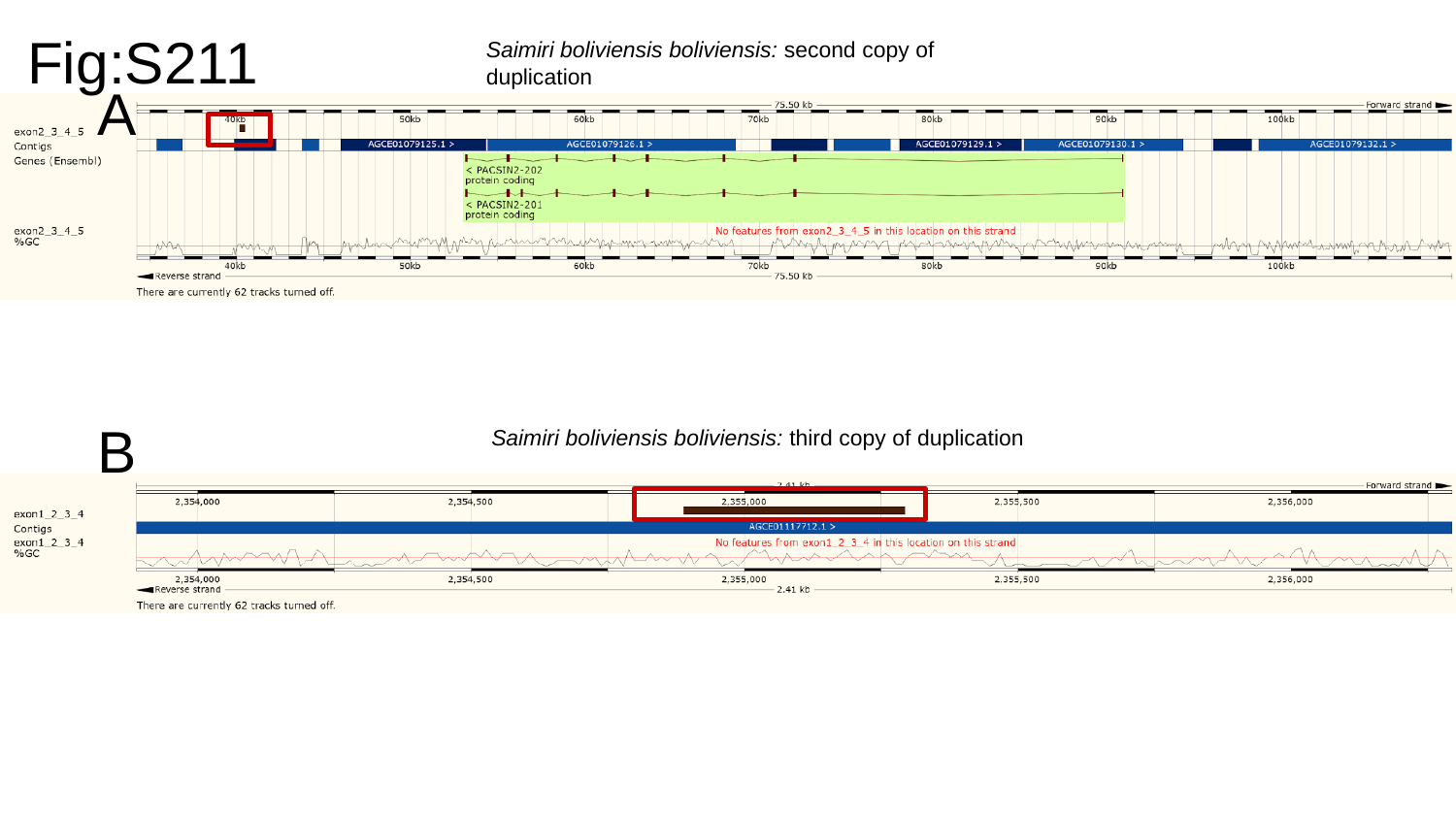

Fig:S211
Saimiri boliviensis boliviensis: second copy of duplication
A
B
Saimiri boliviensis boliviensis: third copy of duplication

### Slide 212
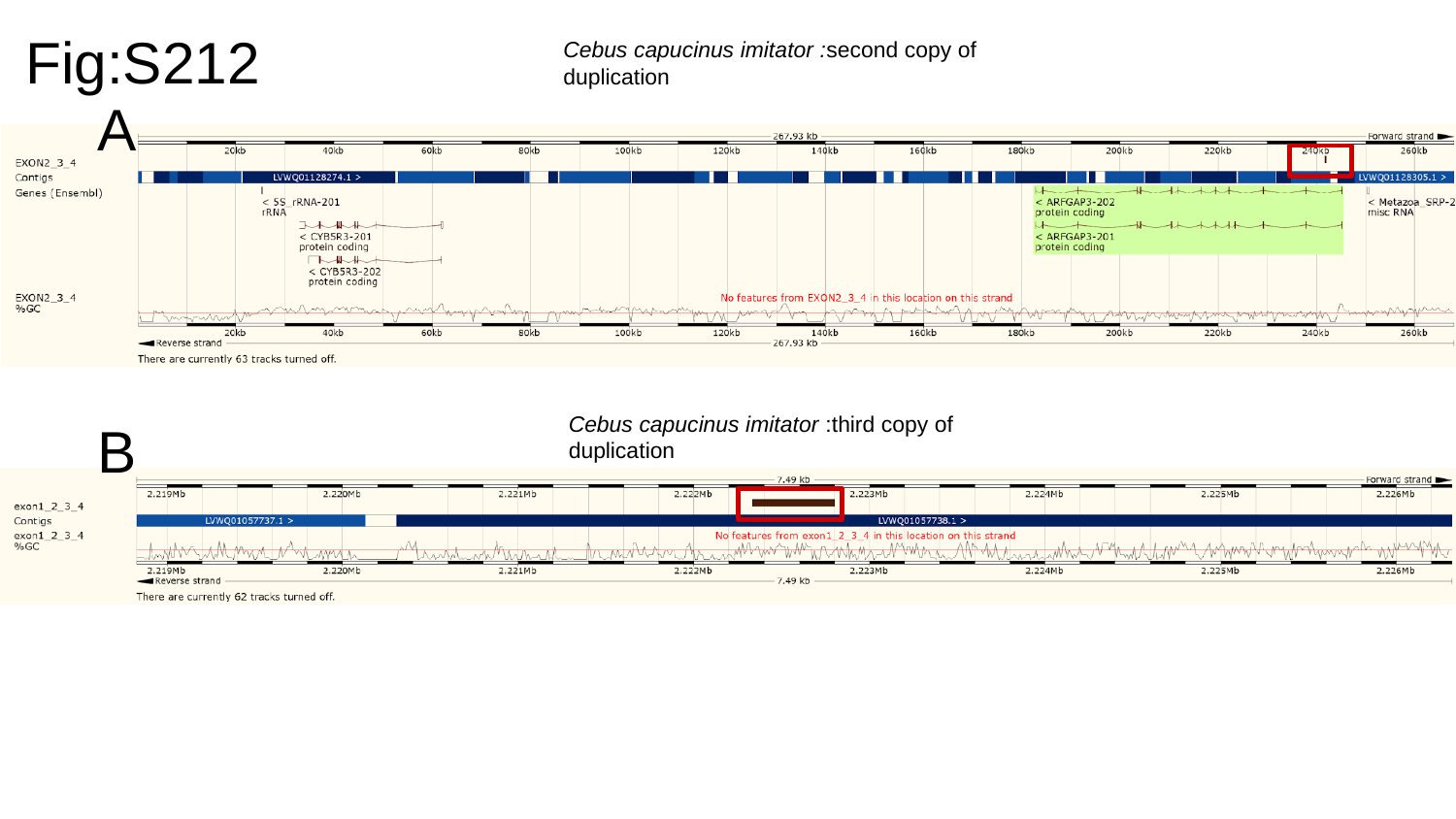

Fig:S212
Cebus capucinus imitator :second copy of duplication
A
B
Cebus capucinus imitator :third copy of duplication

### Slide 213
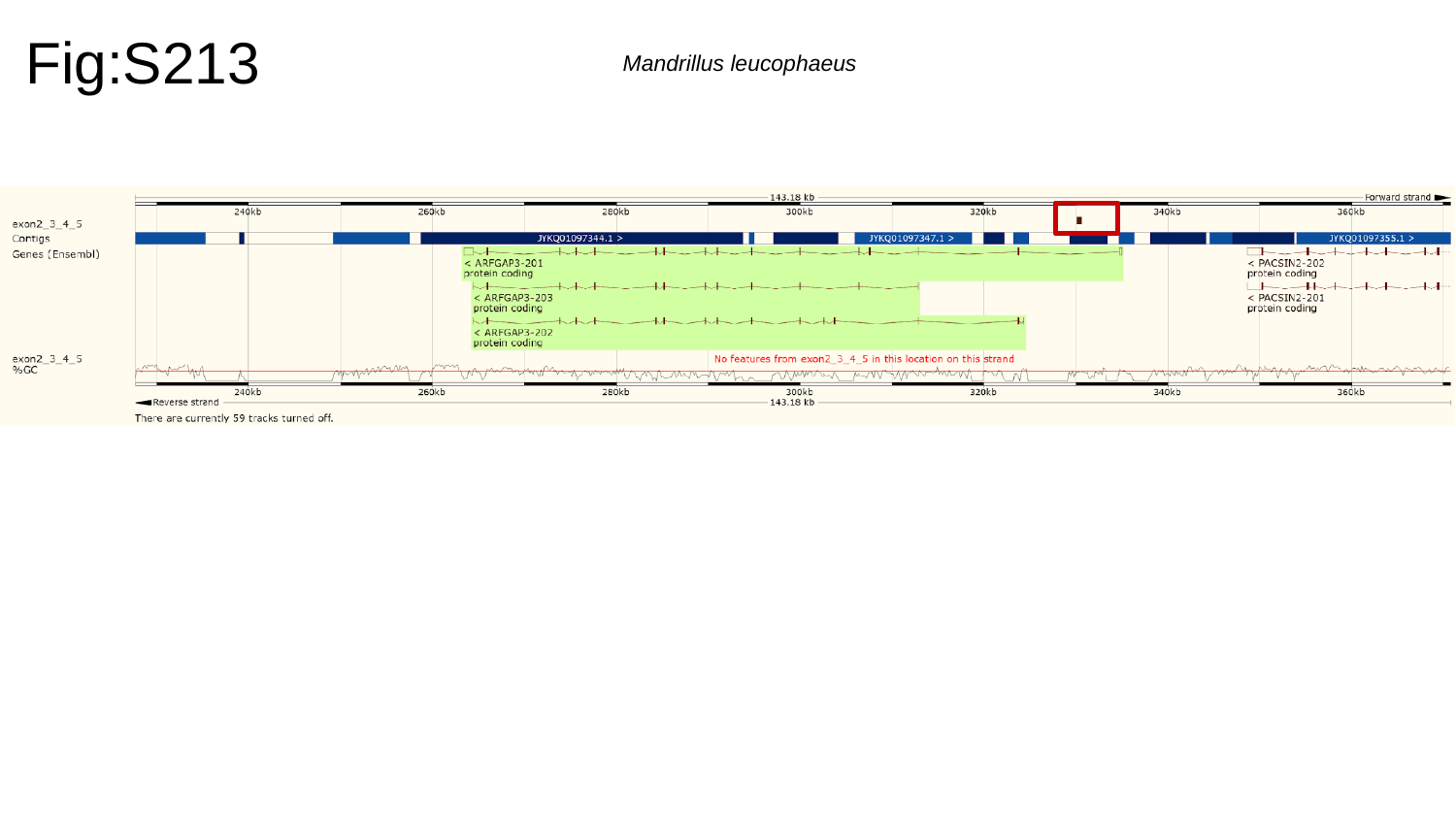

Fig:S213
Mandrillus leucophaeus

### Slide 214
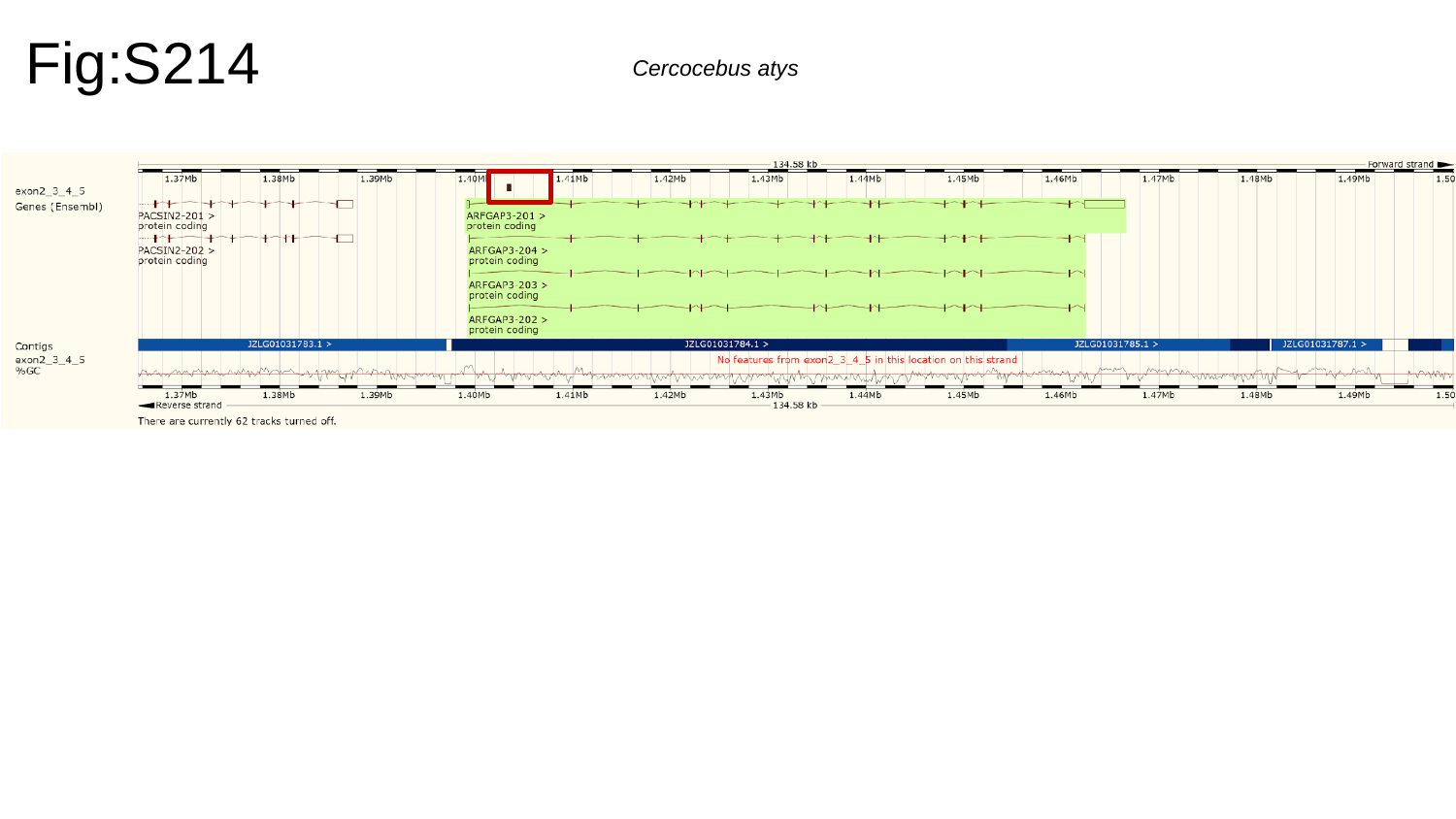

Fig:S214
Cercocebus atys

### Slide 215
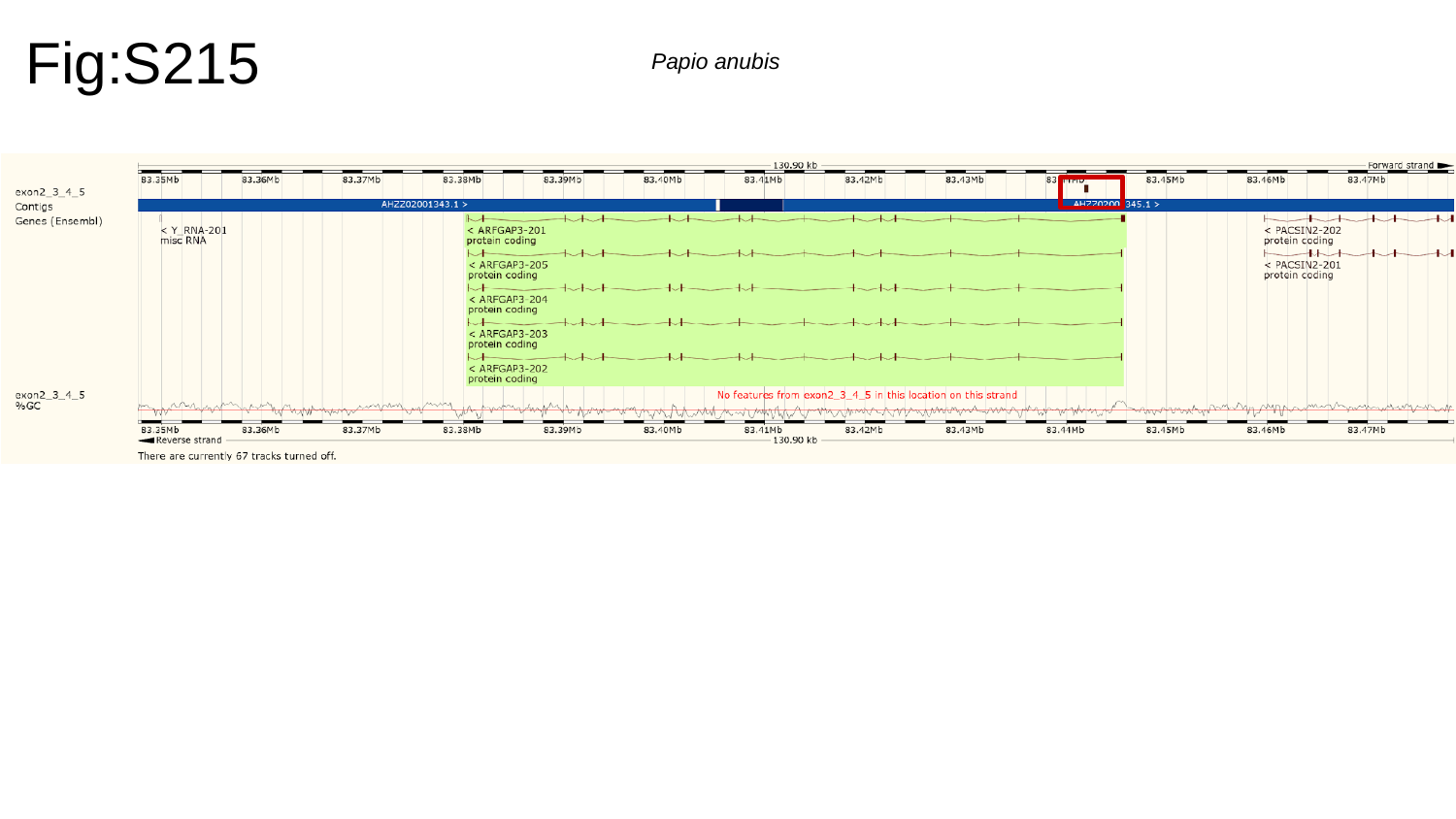

Fig:S215
Papio anubis

### Slide 216
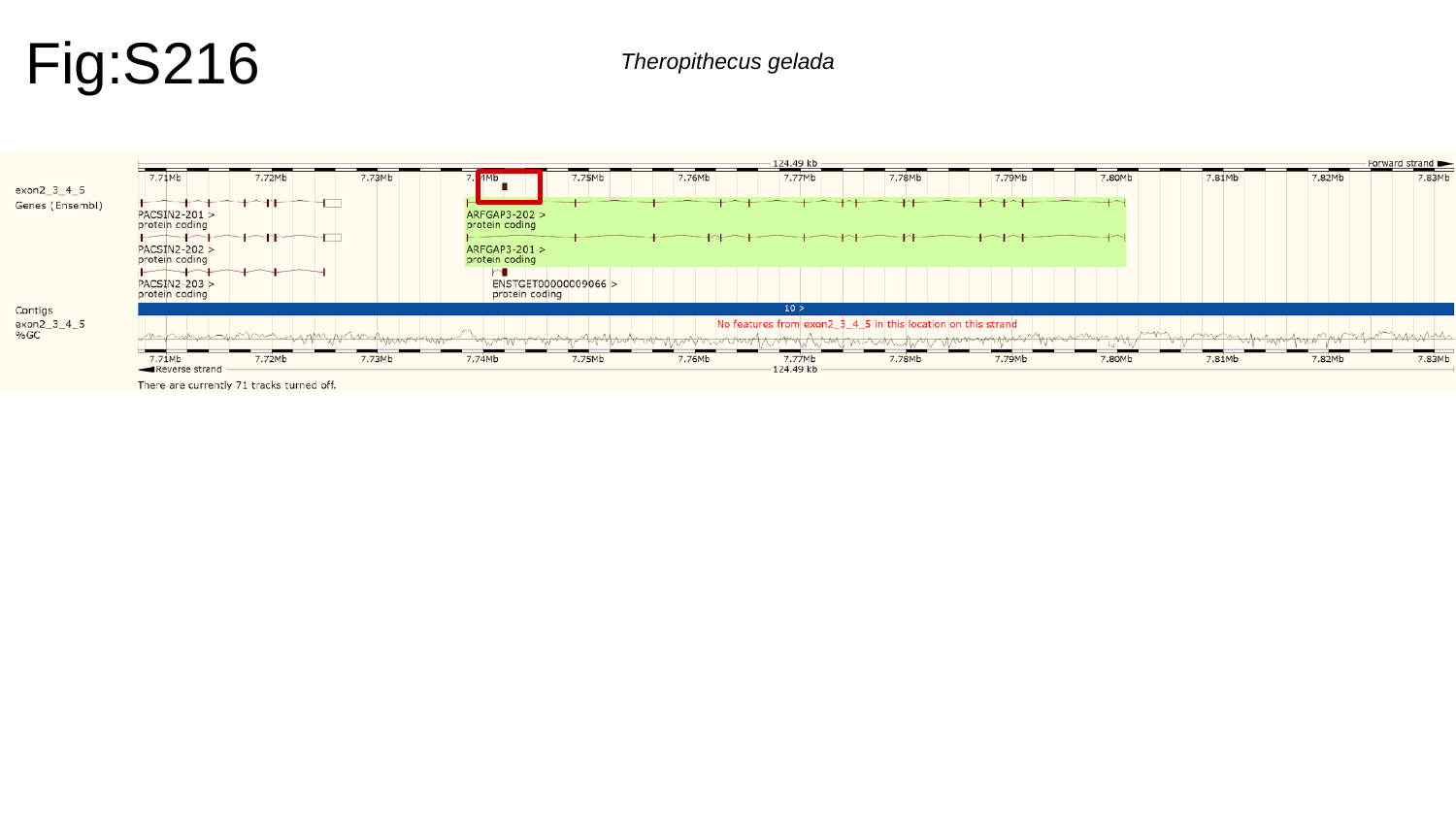

Fig:S216
Theropithecus gelada

### Slide 217
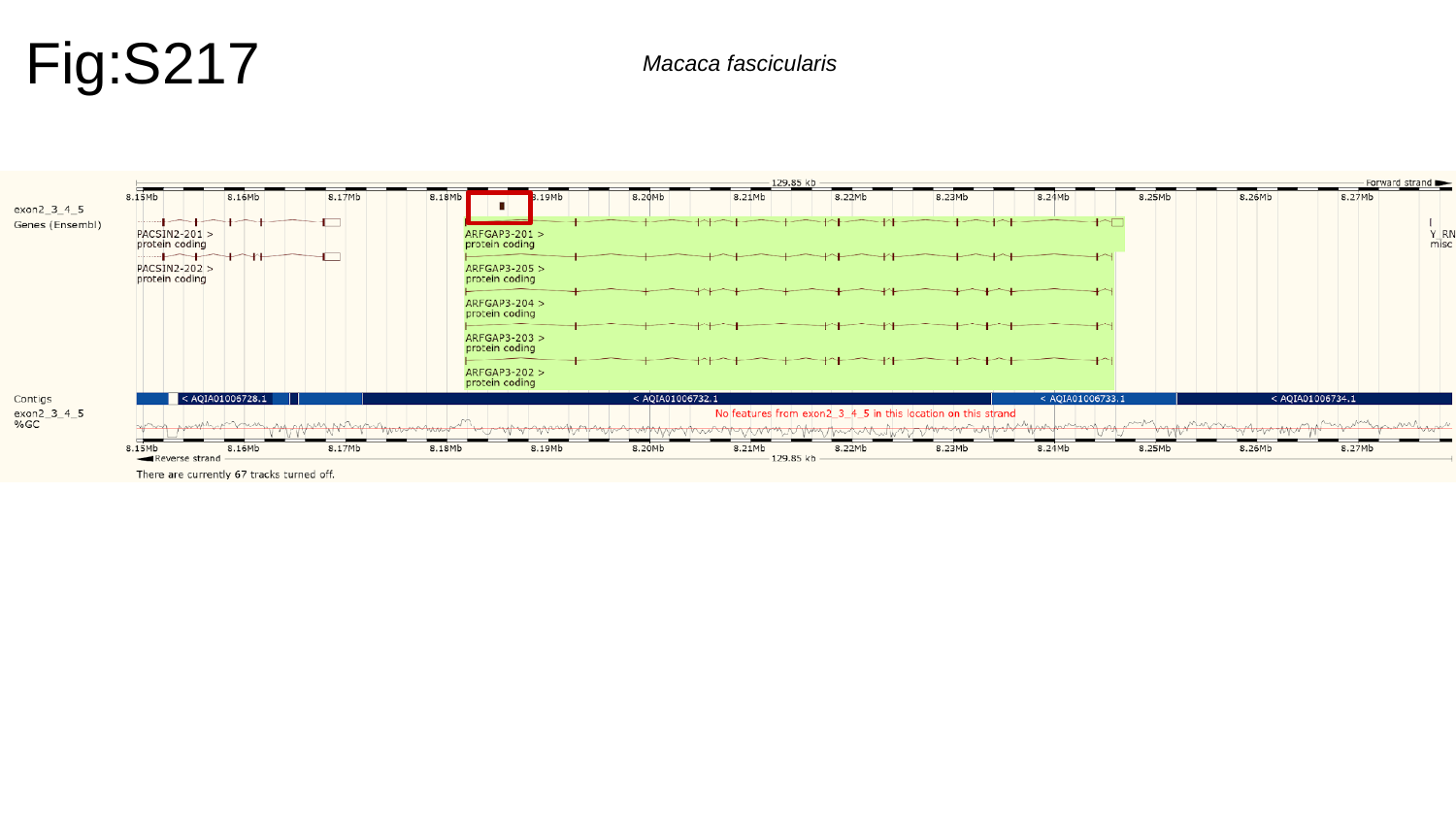

Fig:S217
Macaca fascicularis

### Slide 218
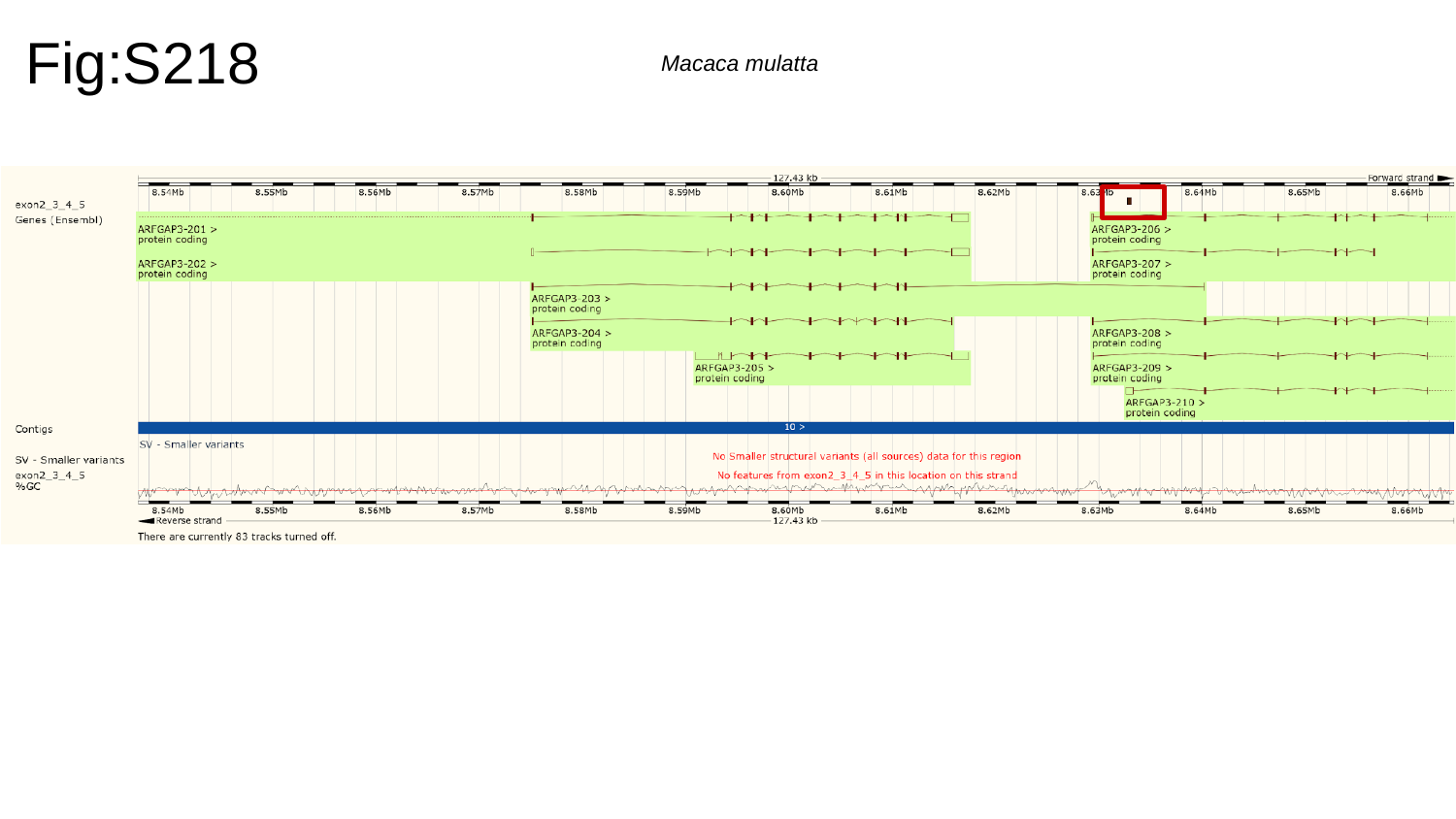

Fig:S218
Macaca mulatta

### Slide 219
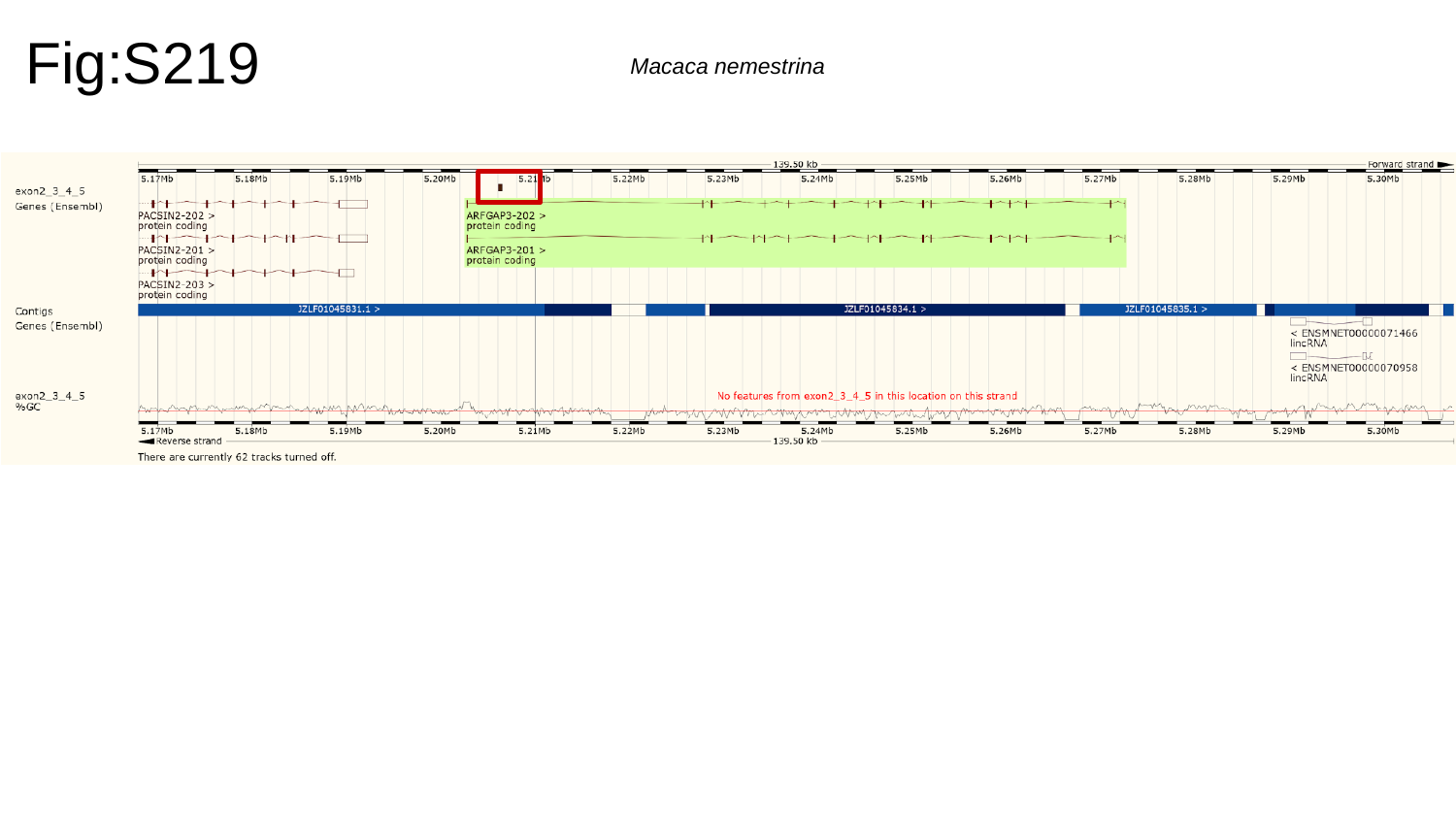

Fig:S219
Macaca nemestrina

### Slide 220
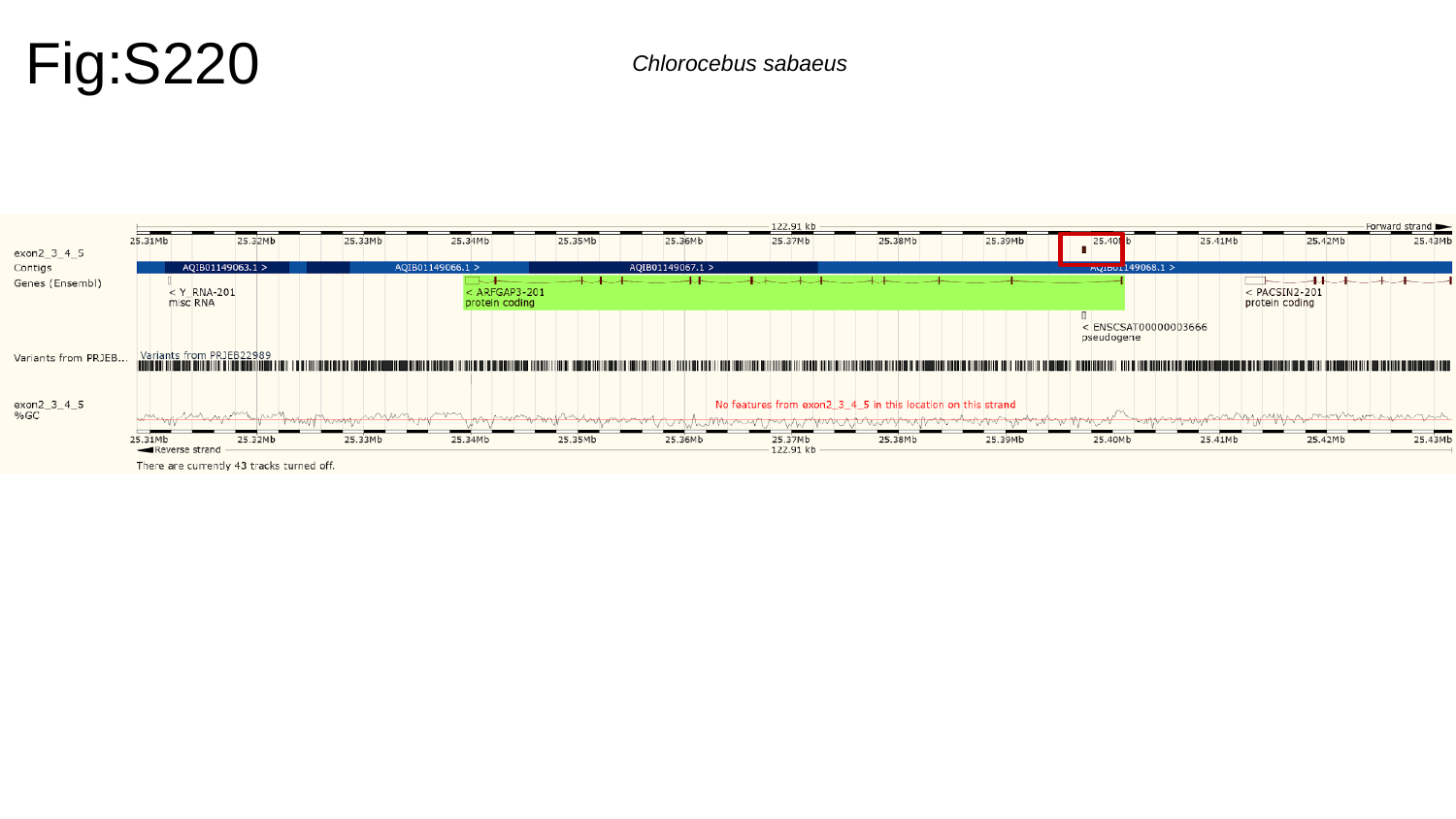

Fig:S220
Chlorocebus sabaeus

### Slide 221
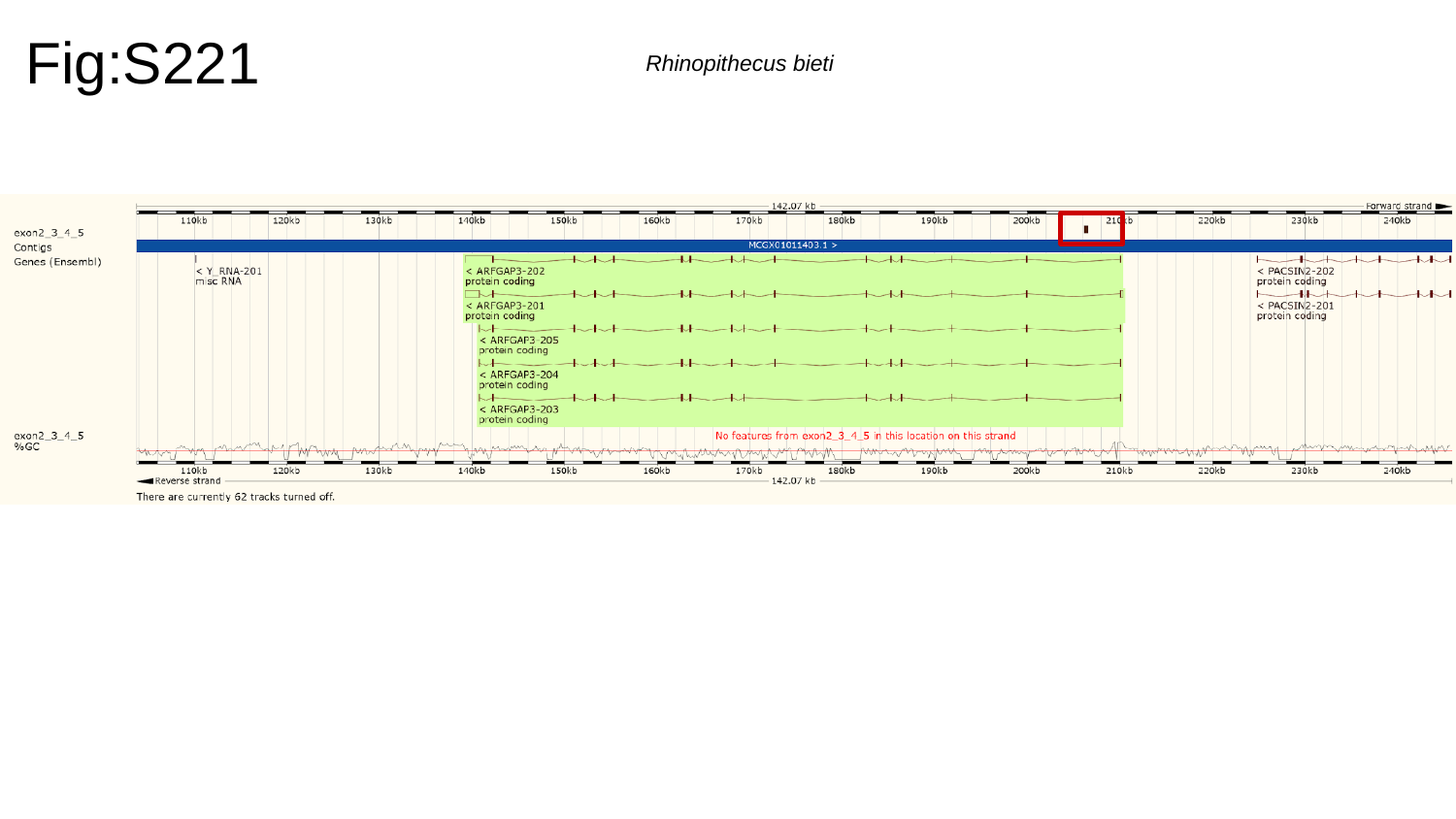

Fig:S221
Rhinopithecus bieti

### Slide 222
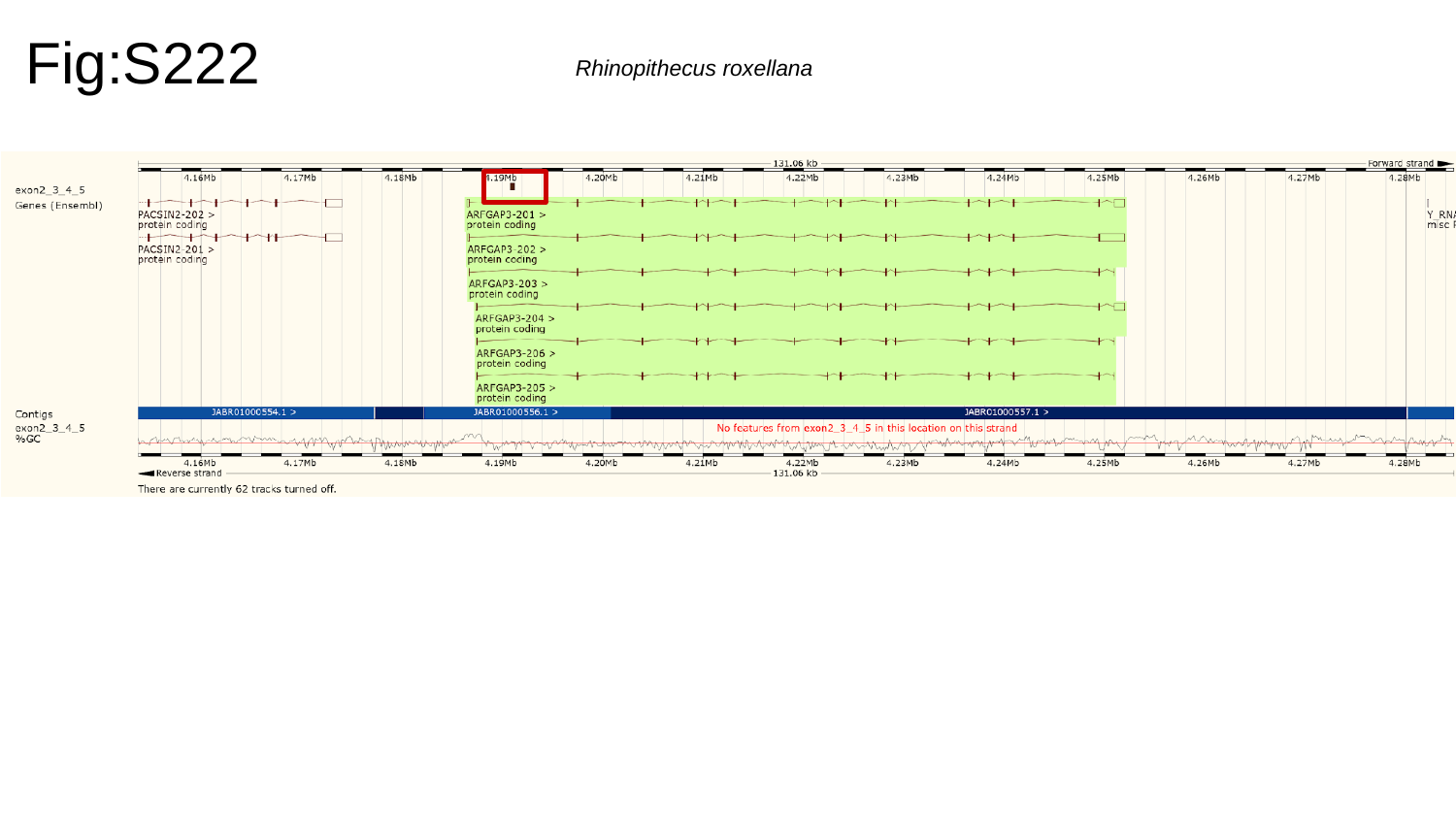

Fig:S222
Rhinopithecus roxellana

### Slide 223
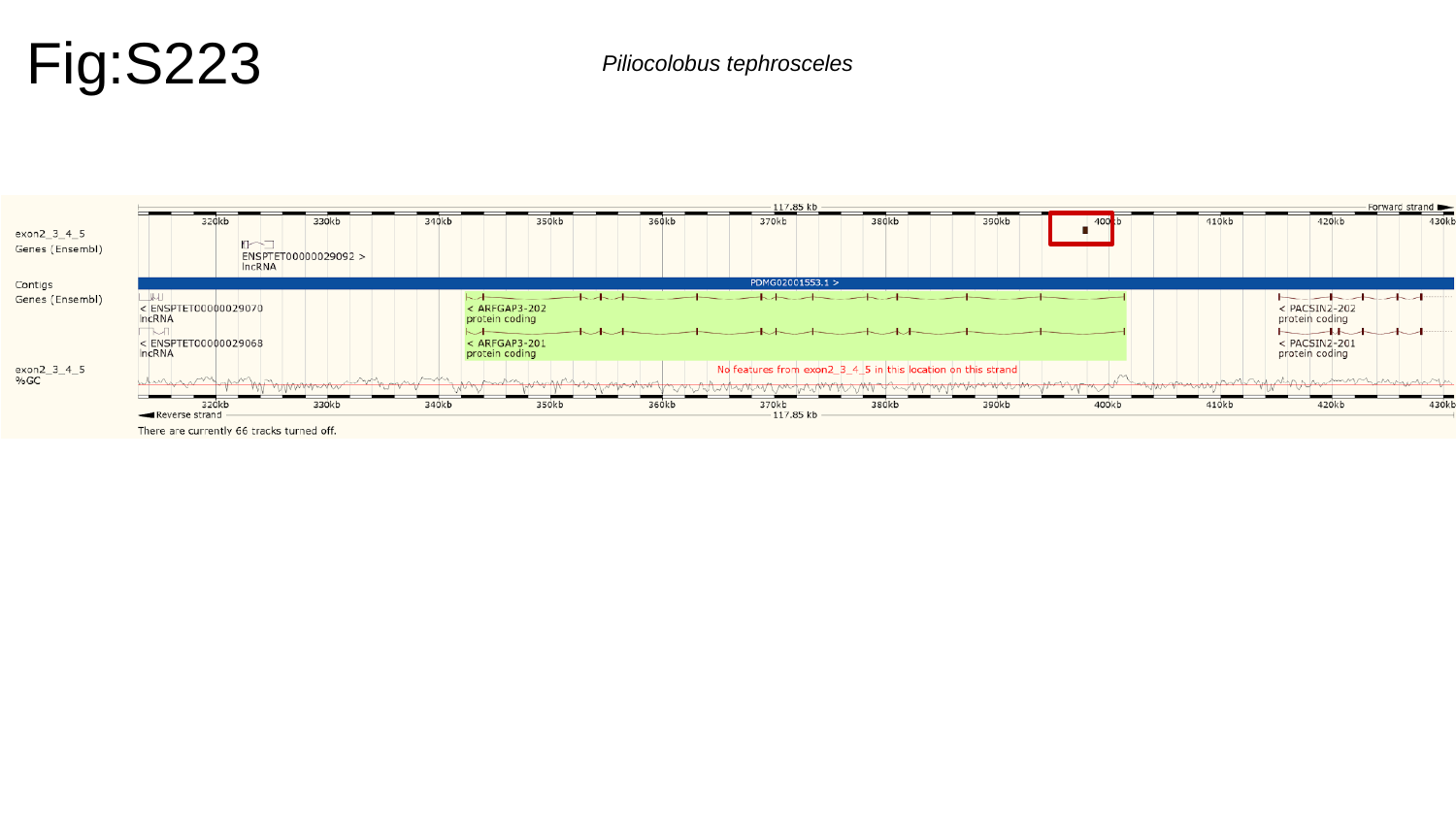

Fig:S223
Piliocolobus tephrosceles

### Slide 224
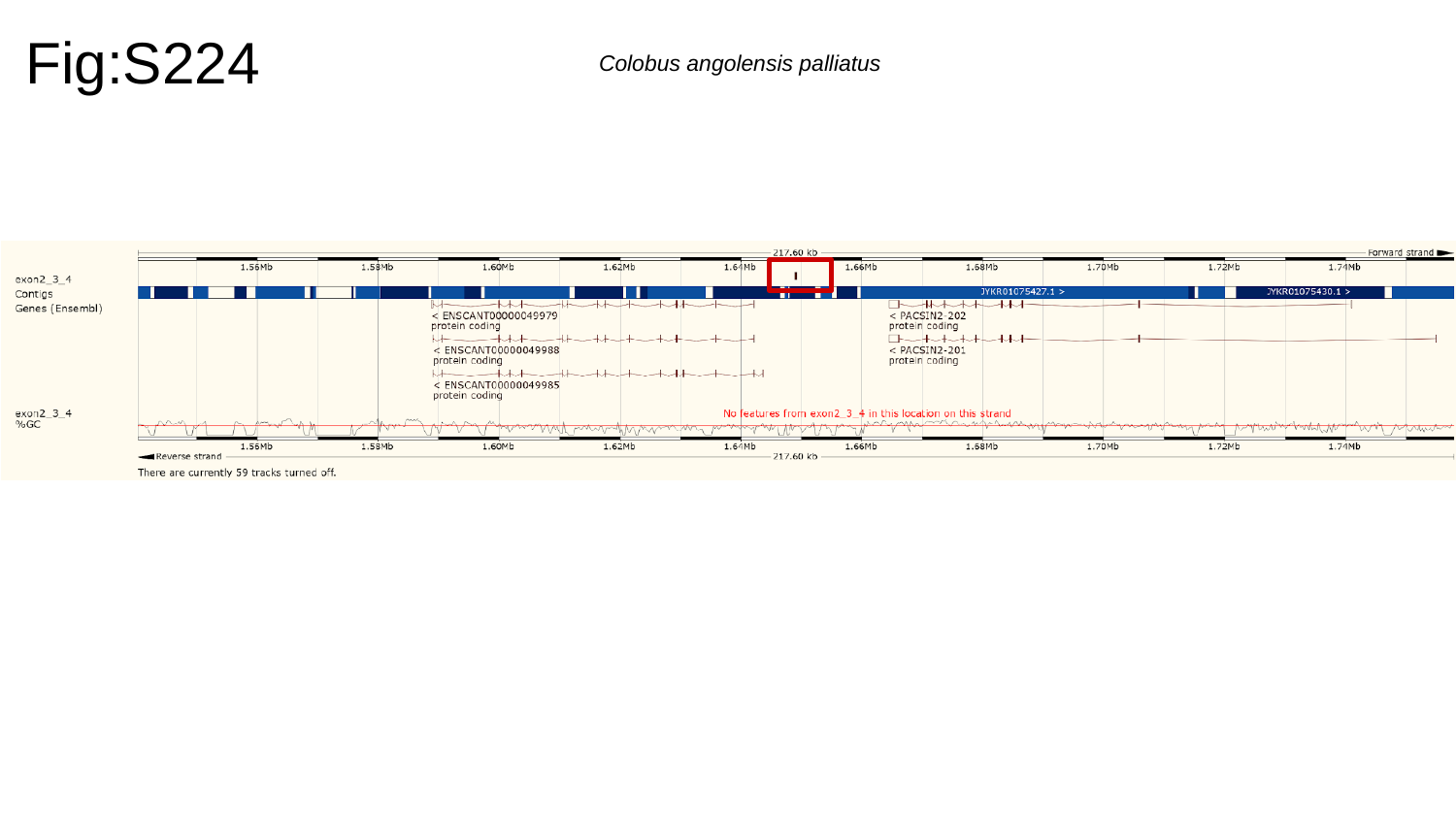

Fig:S224
Colobus angolensis palliatus

### Slide 225
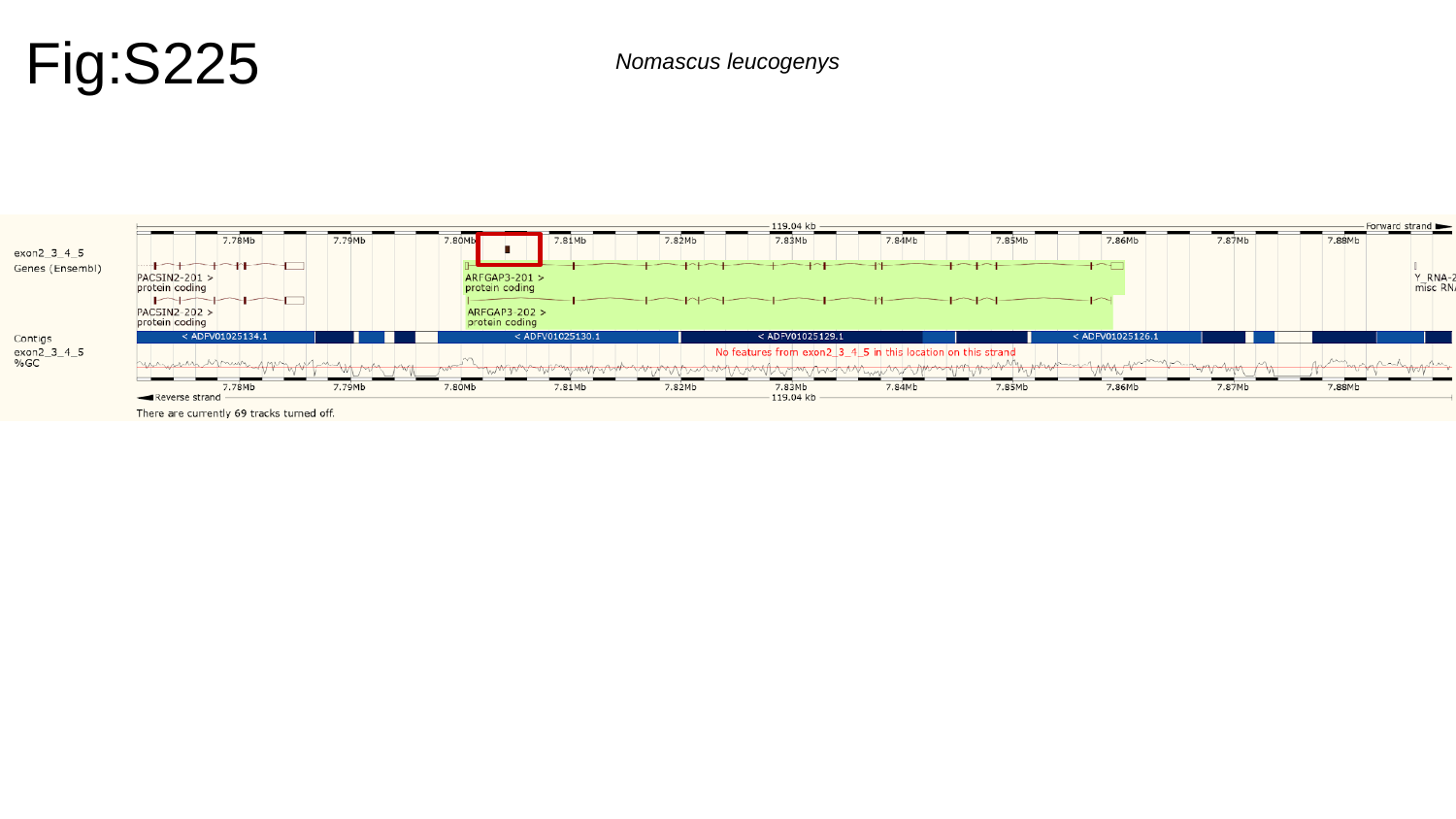

Fig:S225
Nomascus leucogenys

### Slide 226
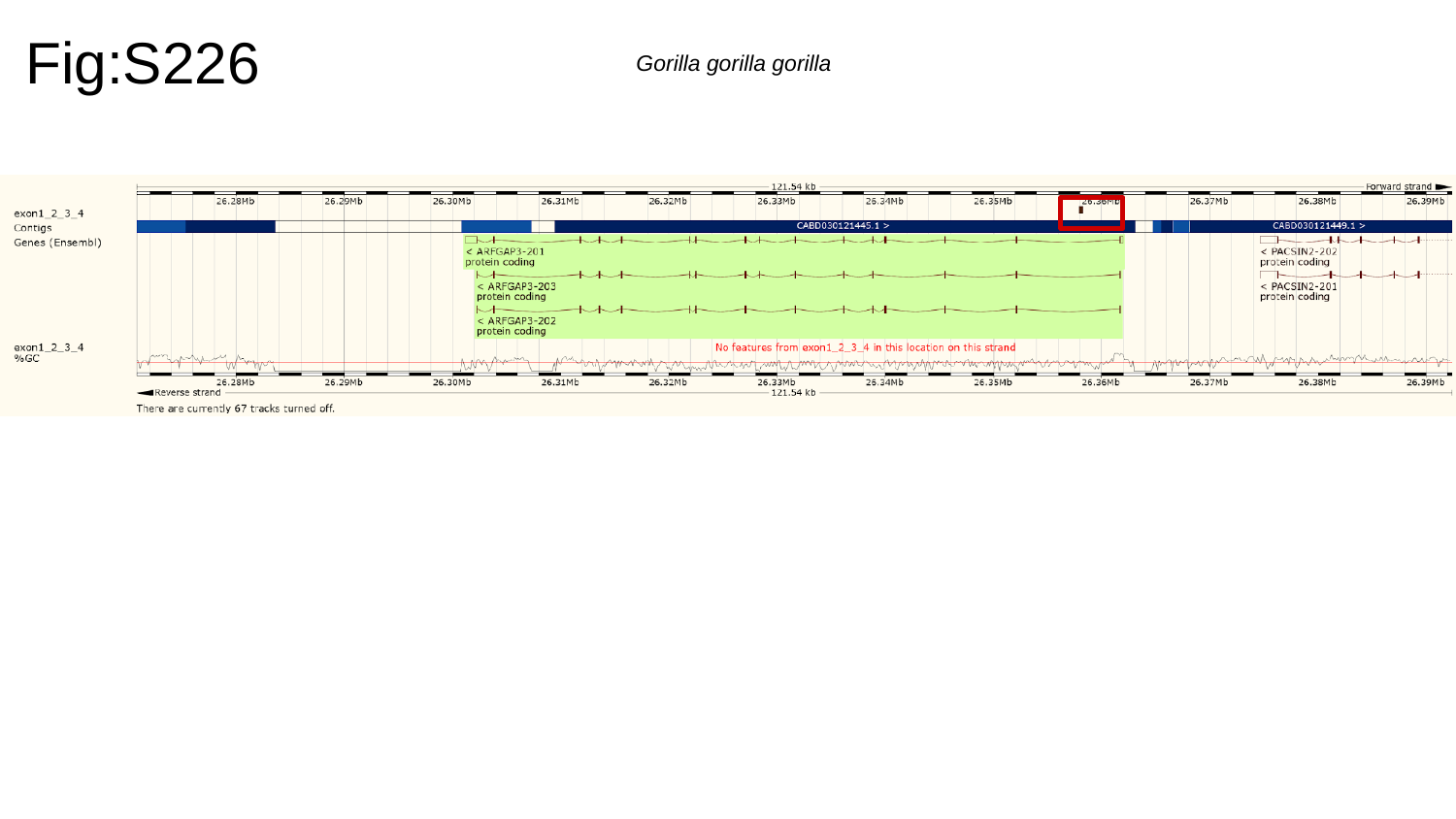

Fig:S226
Gorilla gorilla gorilla

### Slide 227
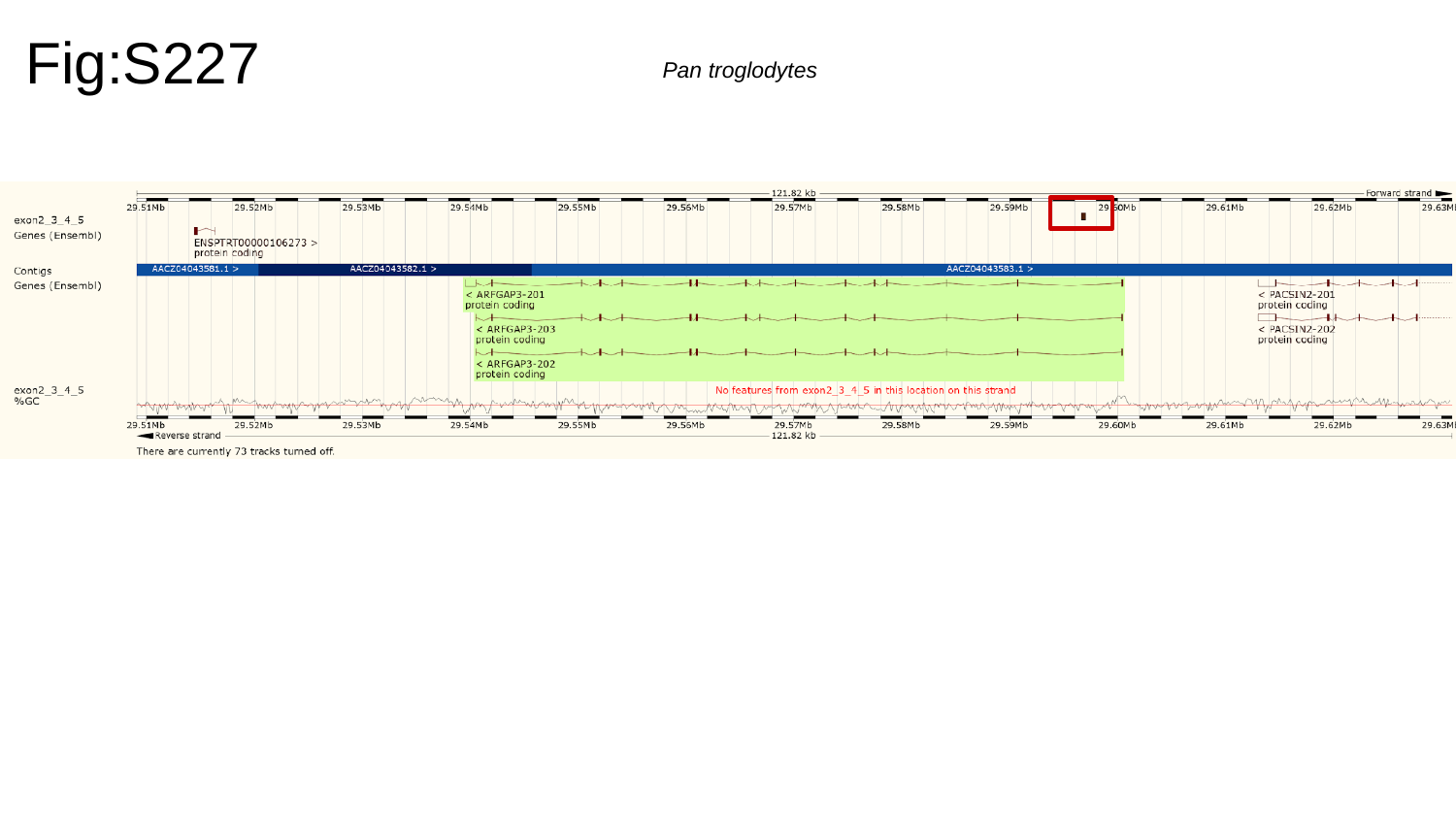

Fig:S227
Pan troglodytes

### Slide 228
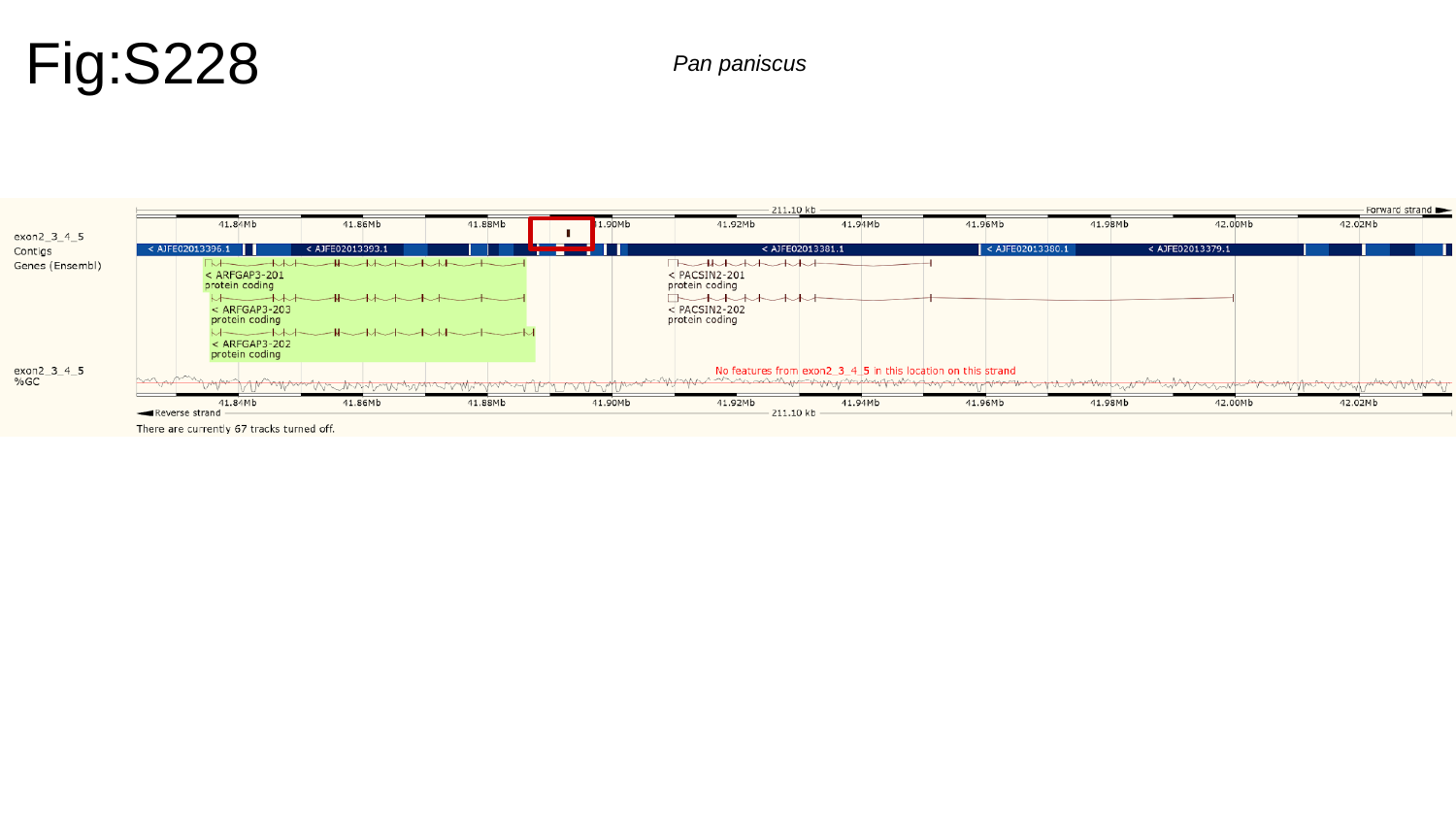

Fig:S228
Pan paniscus

### Slide 229
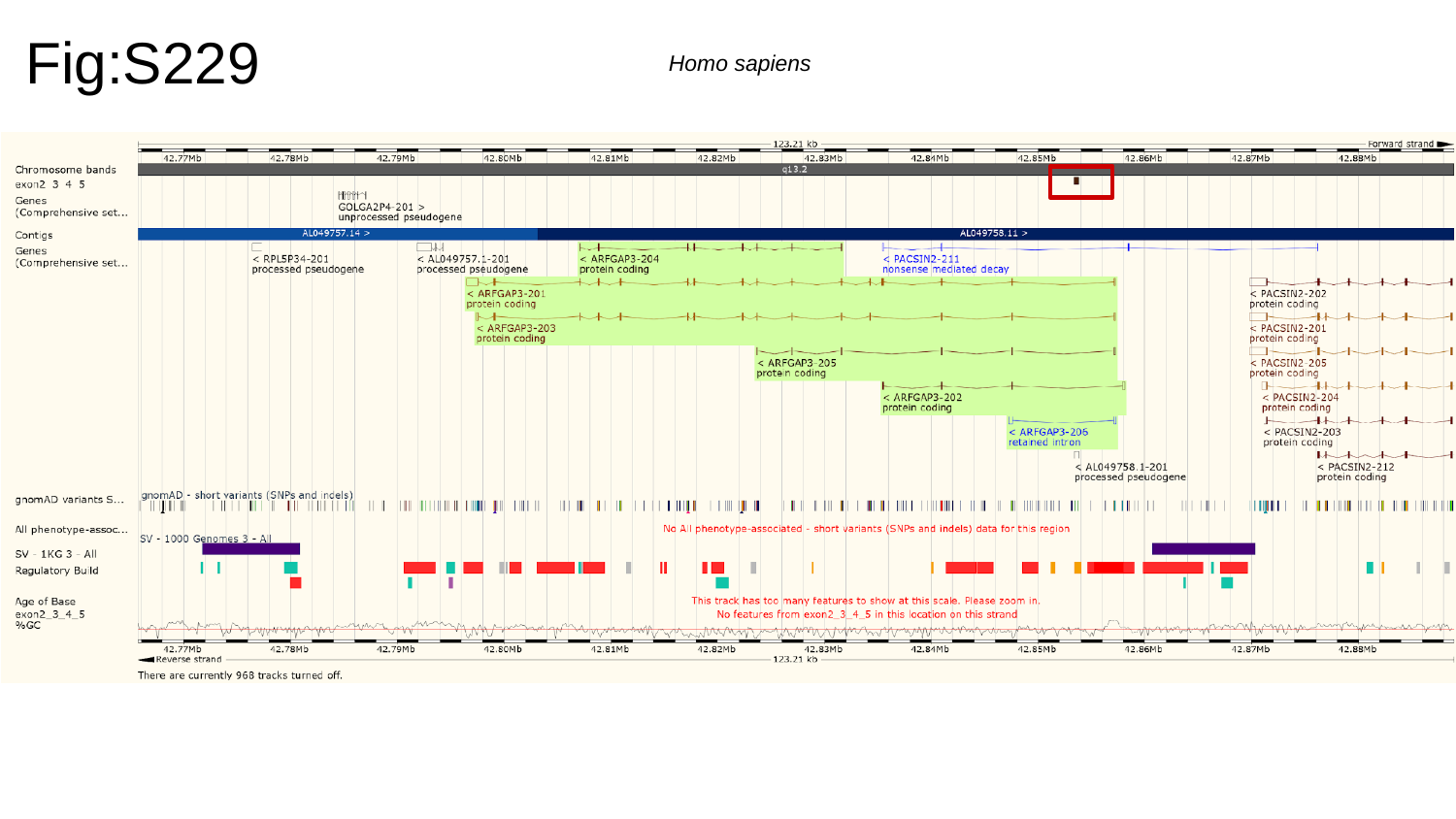

Fig:S229
Homo sapiens

### Slide 230
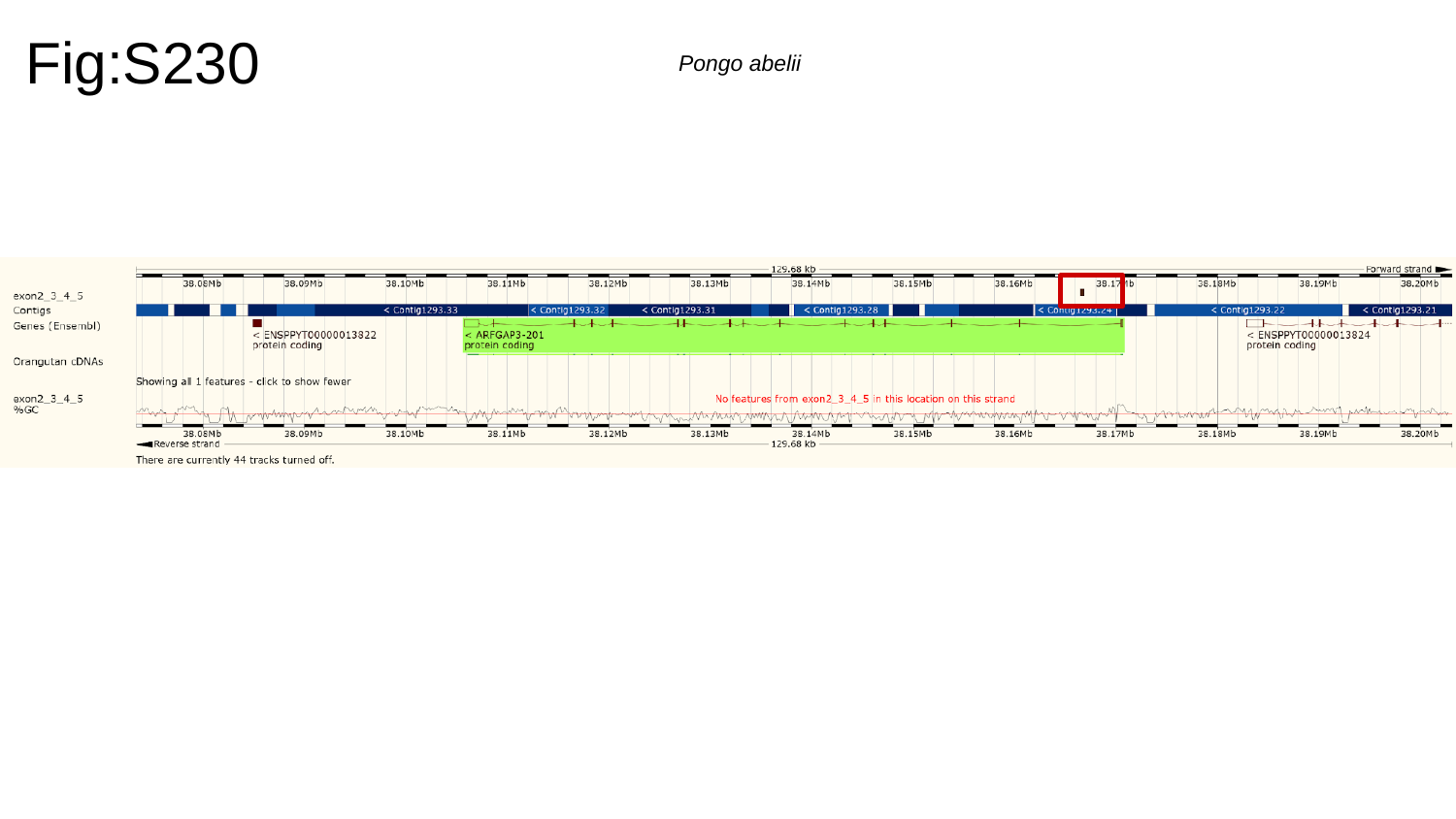

Fig:S230
Pongo abelii

### Slide 231
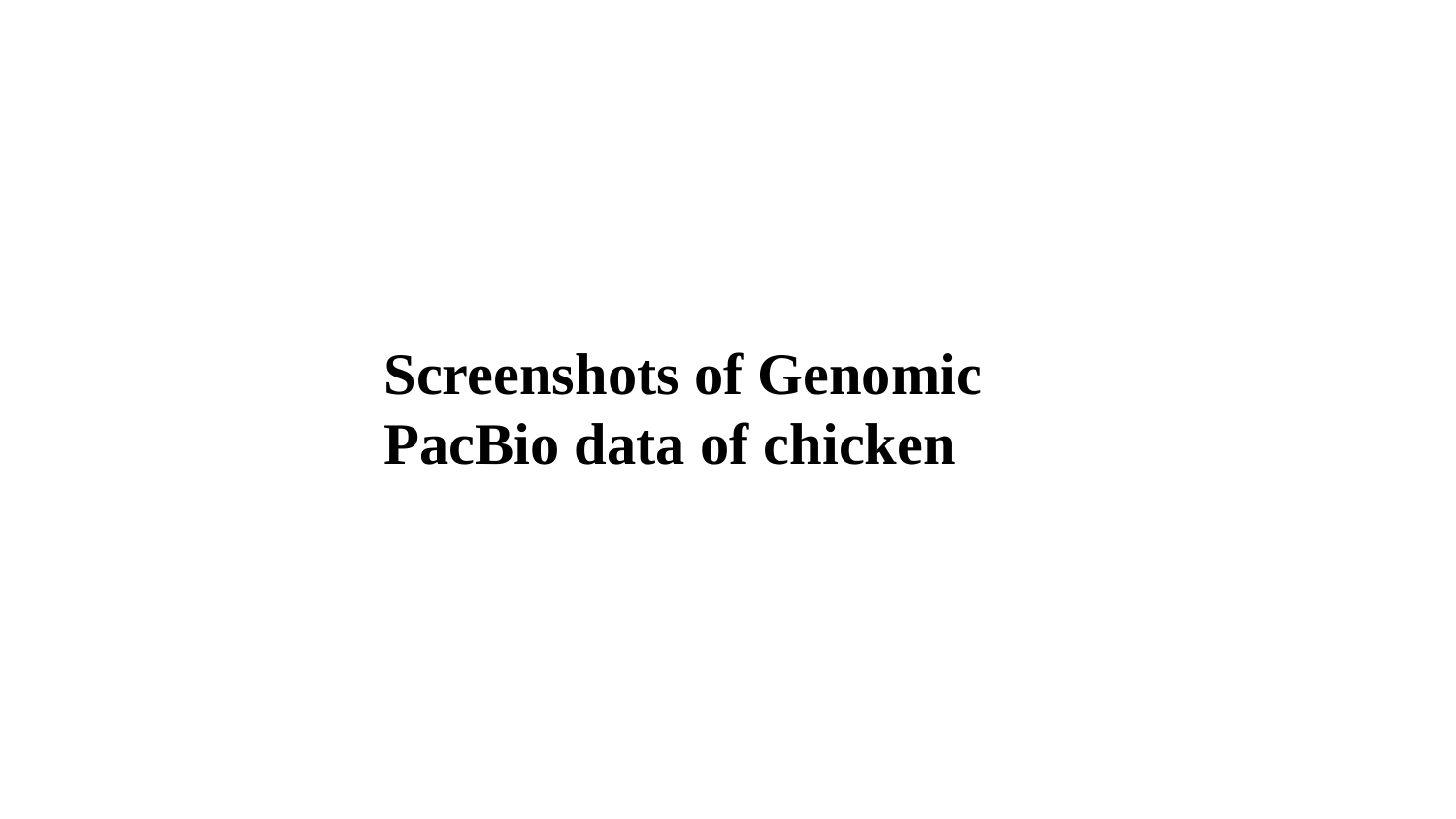

Screenshots of Genomic PacBio data of chicken

### Slide 232
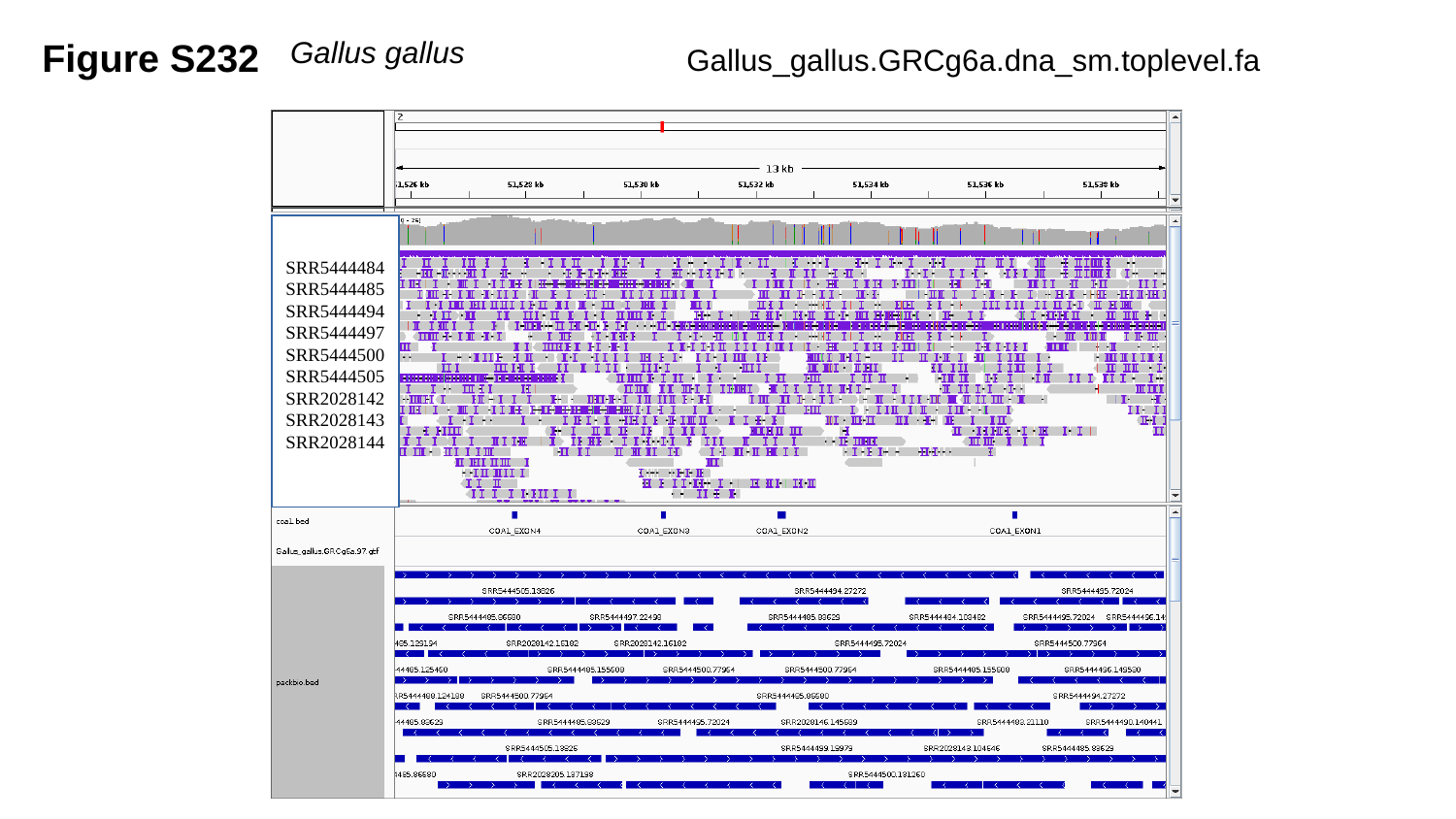

Figure S232
Gallus gallus
Gallus_gallus.GRCg6a.dna_sm.toplevel.fa
SRR5444484
SRR5444485
SRR5444494
SRR5444497
SRR5444500
SRR5444505
SRR2028142
SRR2028143
SRR2028144

### Slide 233
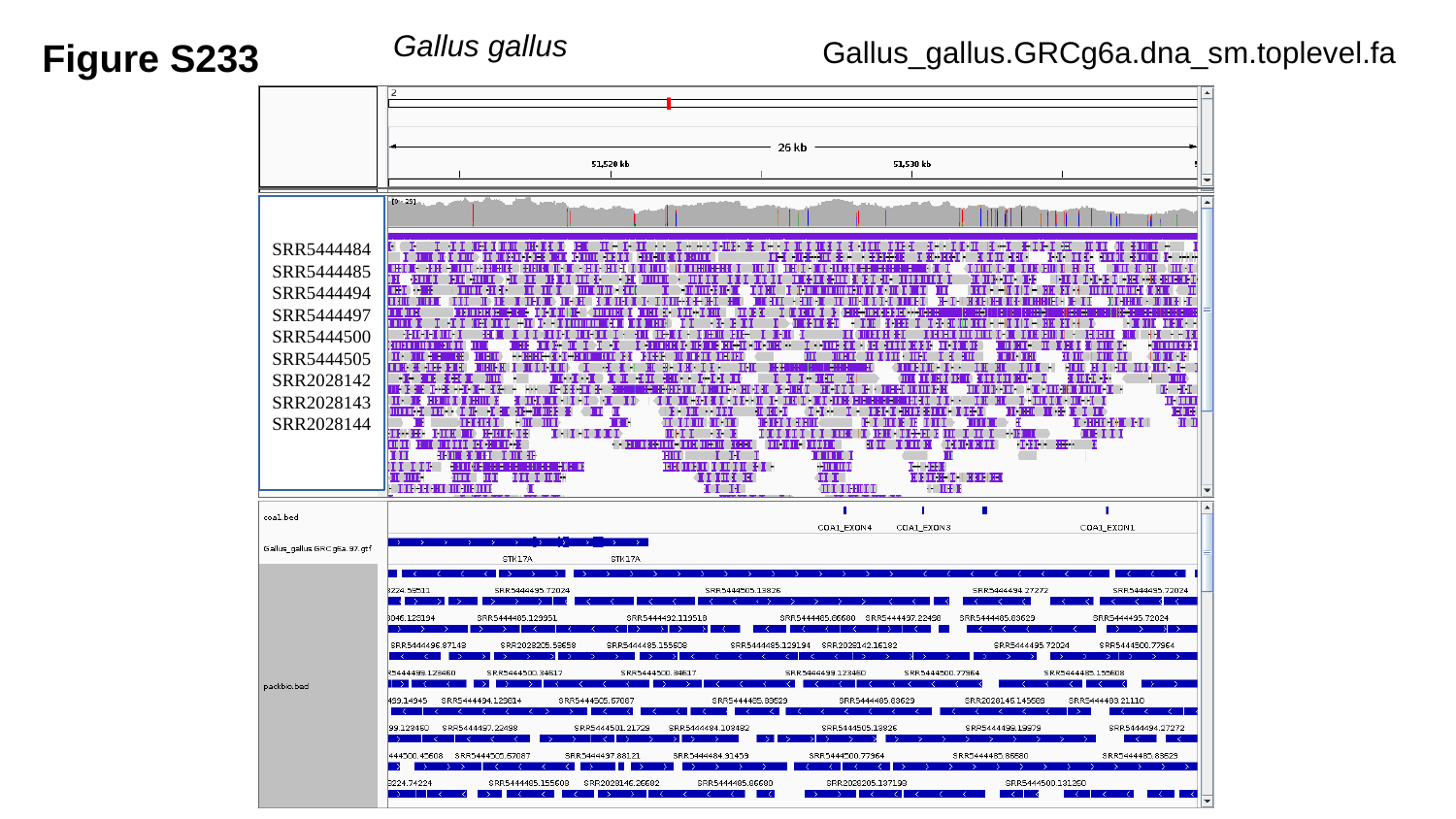

Figure S233
Gallus gallus
Gallus_gallus.GRCg6a.dna_sm.toplevel.fa
SRR5444484
SRR5444485
SRR5444494
SRR5444497
SRR5444500
SRR5444505
SRR2028142
SRR2028143
SRR2028144

### Slide 234
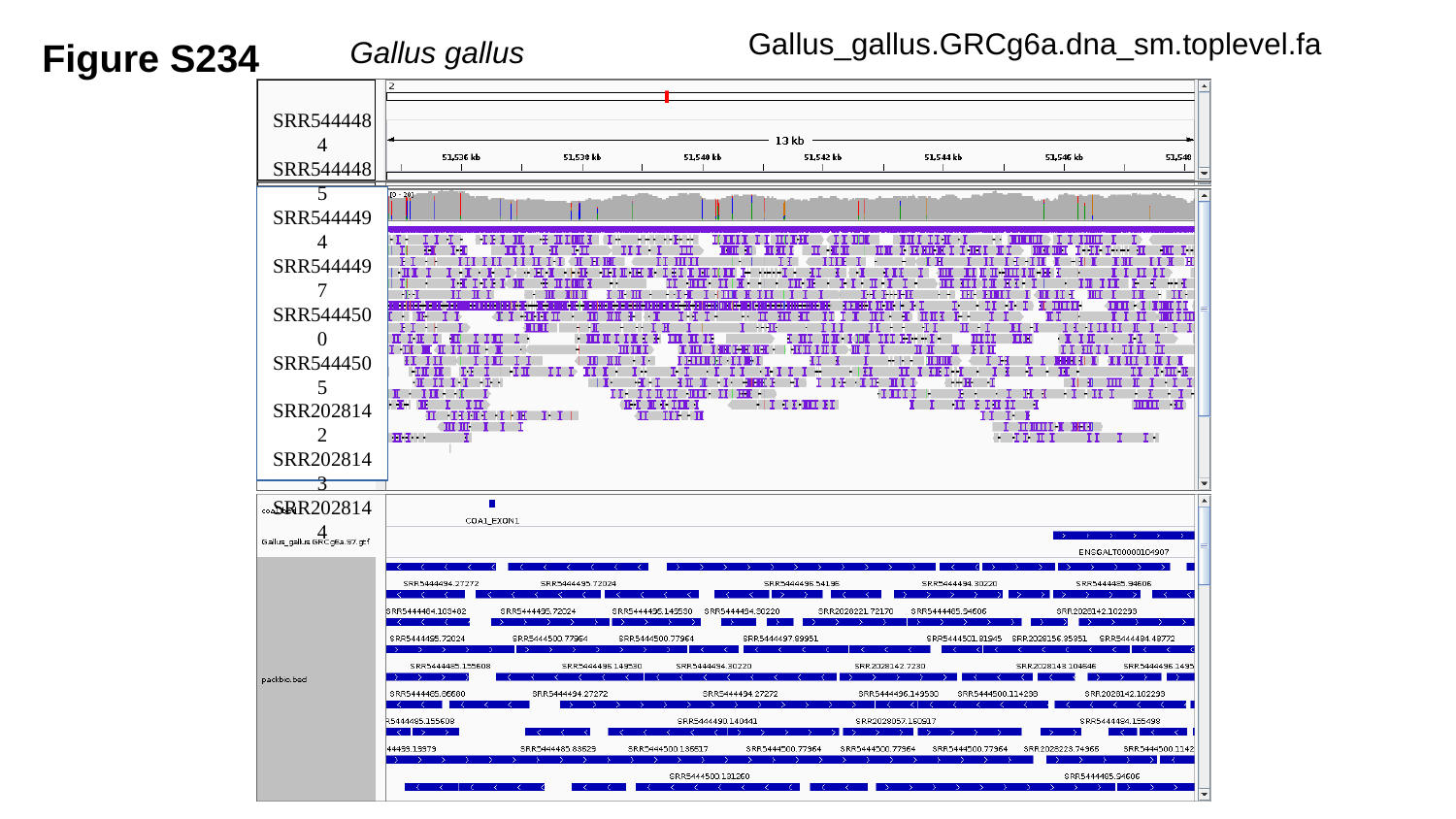

Gallus_gallus.GRCg6a.dna_sm.toplevel.fa
Figure S234
Gallus gallus
SRR5444484
SRR5444485
SRR5444494
SRR5444497
SRR5444500
SRR5444505
SRR2028142
SRR2028143
SRR2028144

### Slide 235
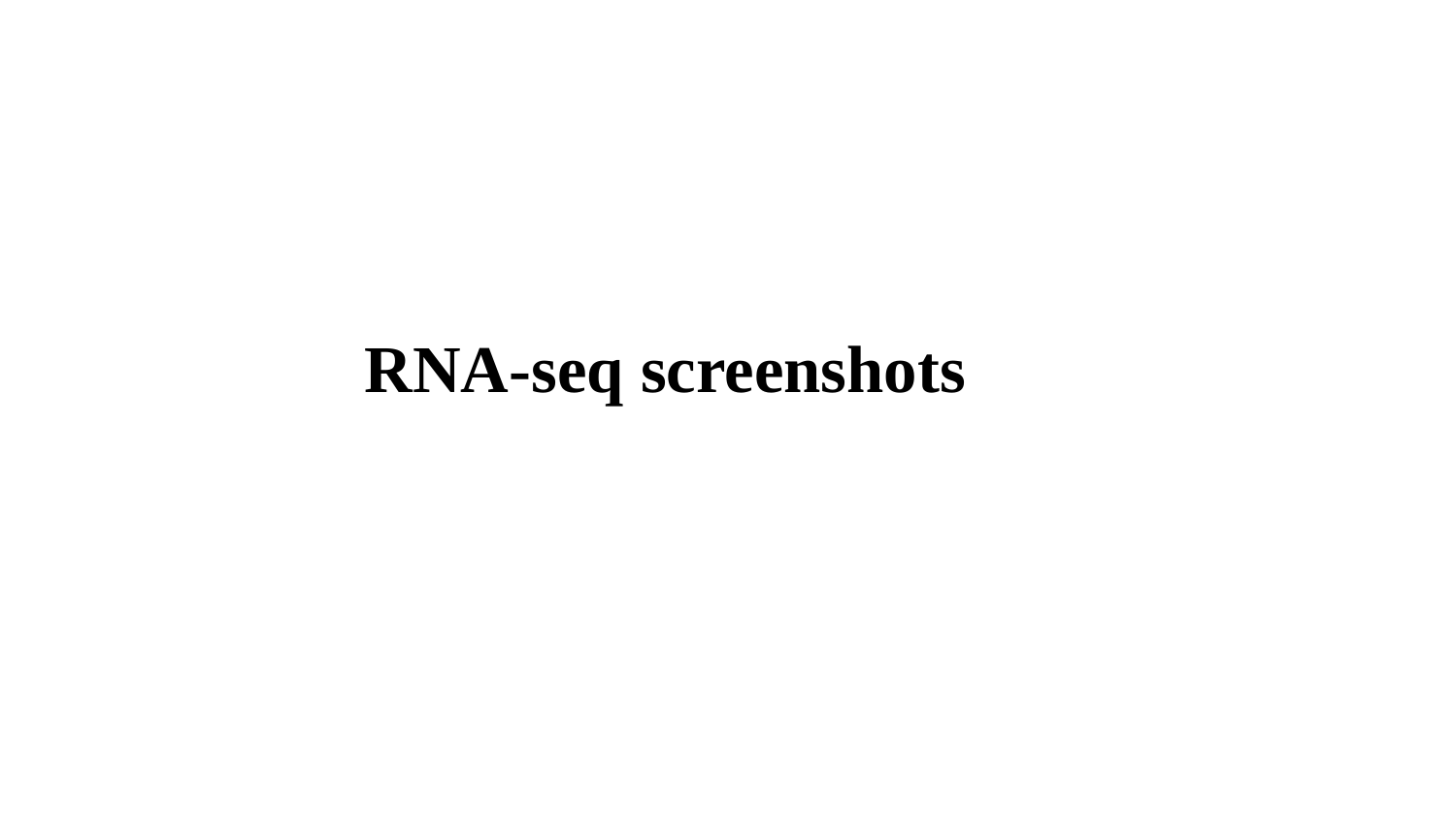

RNA-seq screenshots

### Slide 236
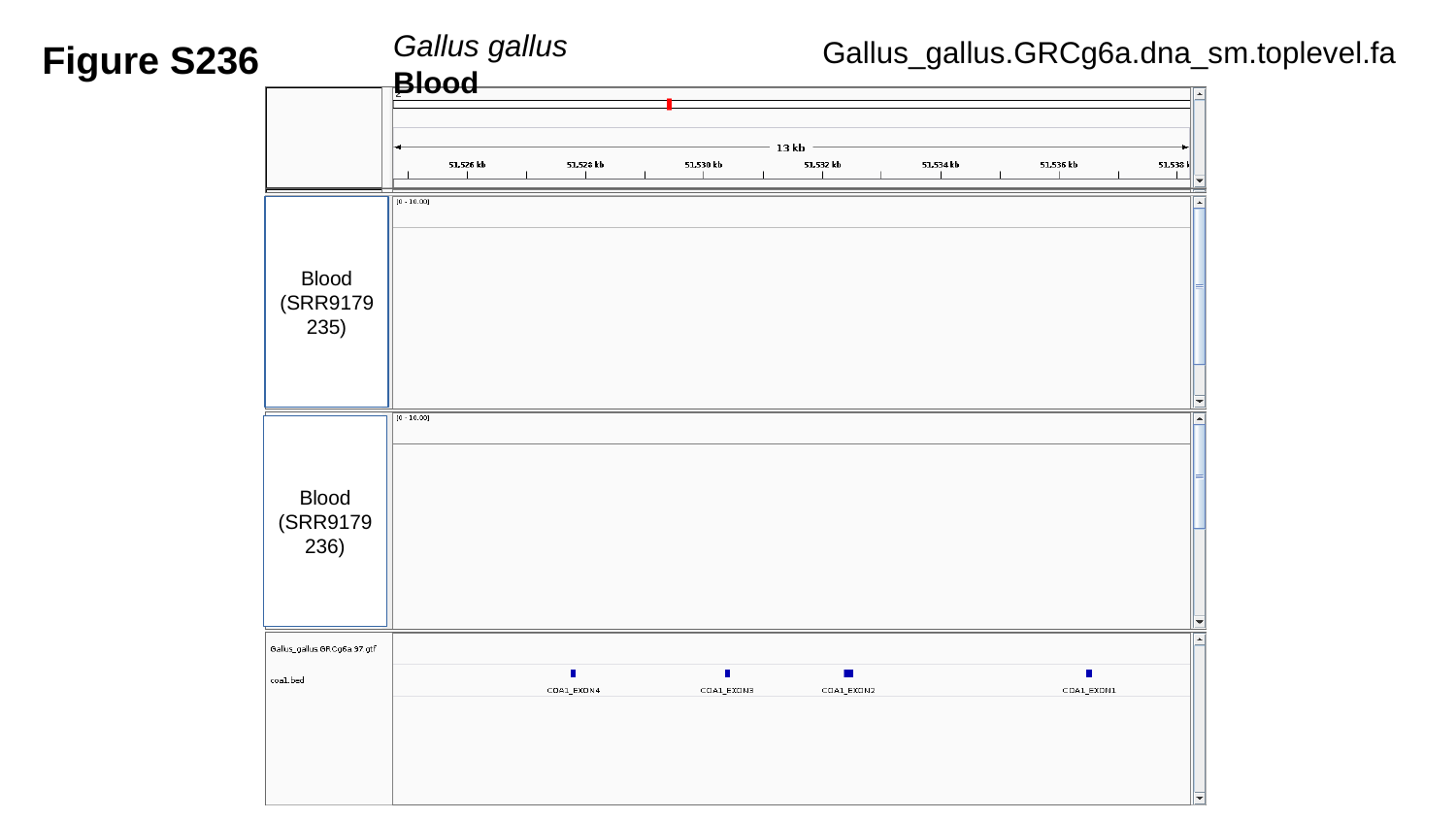

Gallus gallus	 Blood
Figure S236
Gallus_gallus.GRCg6a.dna_sm.toplevel.fa
Blood
(SRR9179235)
Blood
(SRR9179236)

### Slide 237
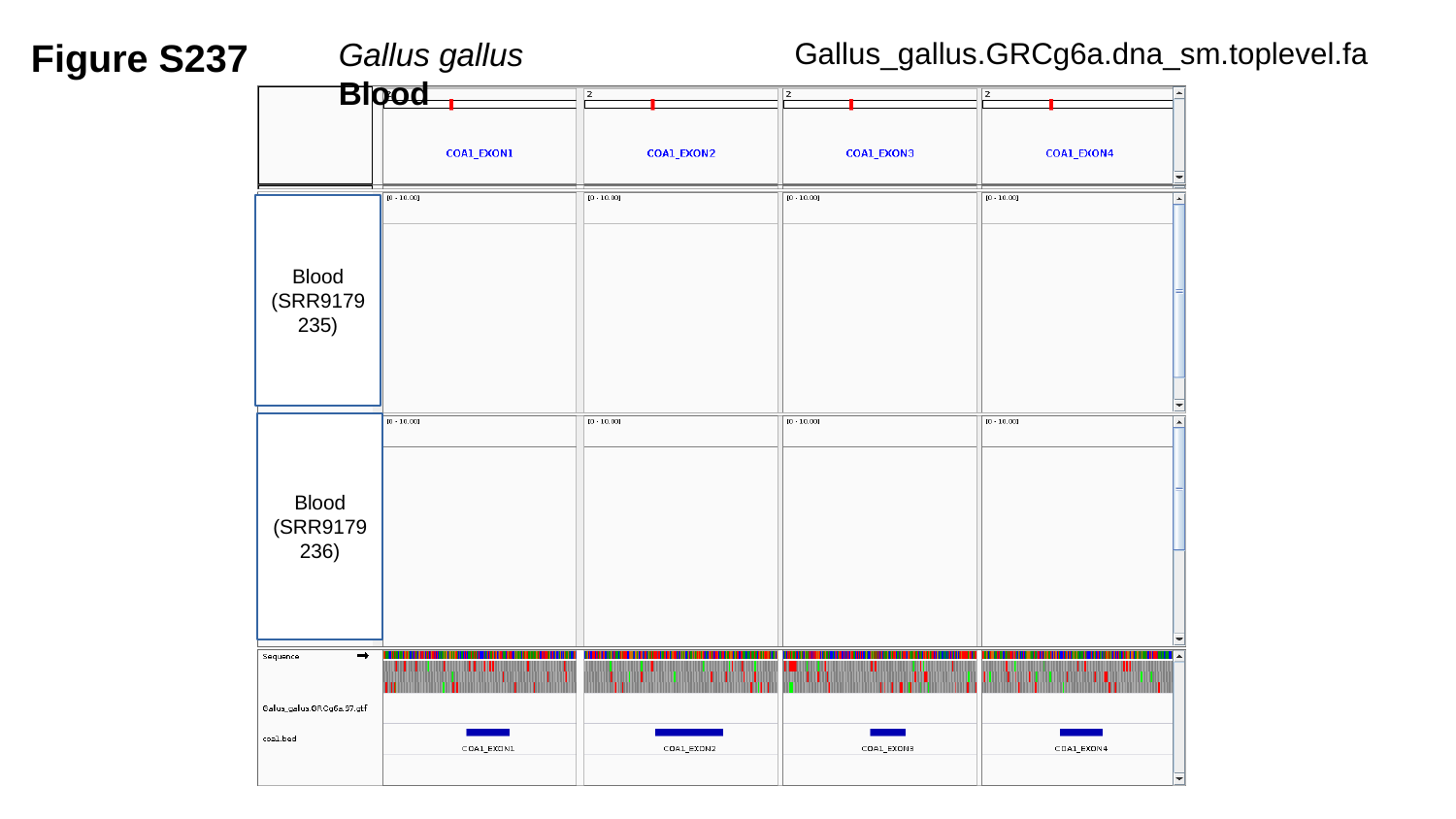

Figure S237
Gallus gallus	 Blood
Gallus_gallus.GRCg6a.dna_sm.toplevel.fa
Blood
(SRR9179235)
Blood
(SRR9179236)

### Slide 238
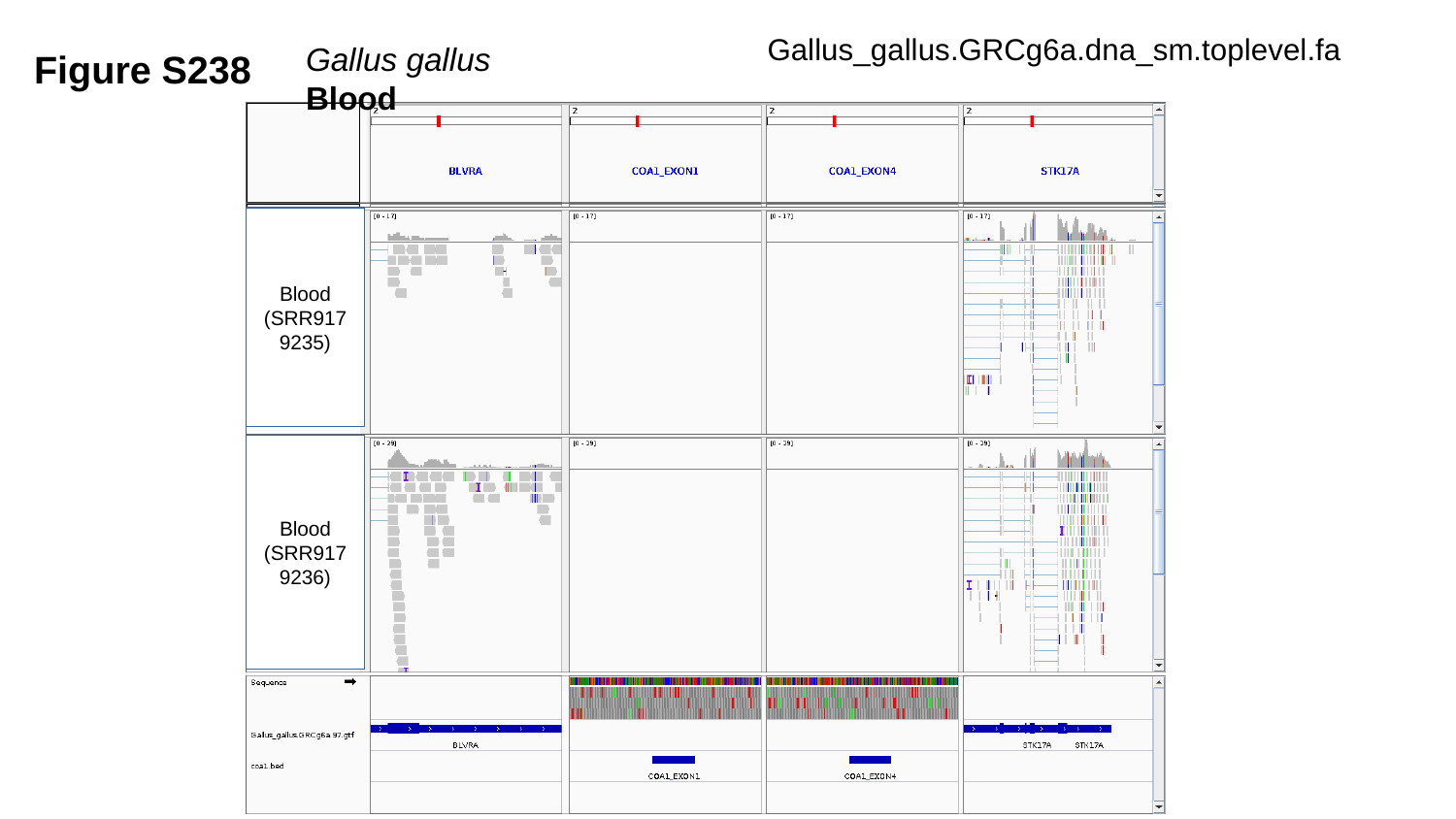

Gallus_gallus.GRCg6a.dna_sm.toplevel.fa
Gallus gallus	 Blood
Figure S238
Blood
(SRR9179235)
Blood
(SRR9179236)

### Slide 239
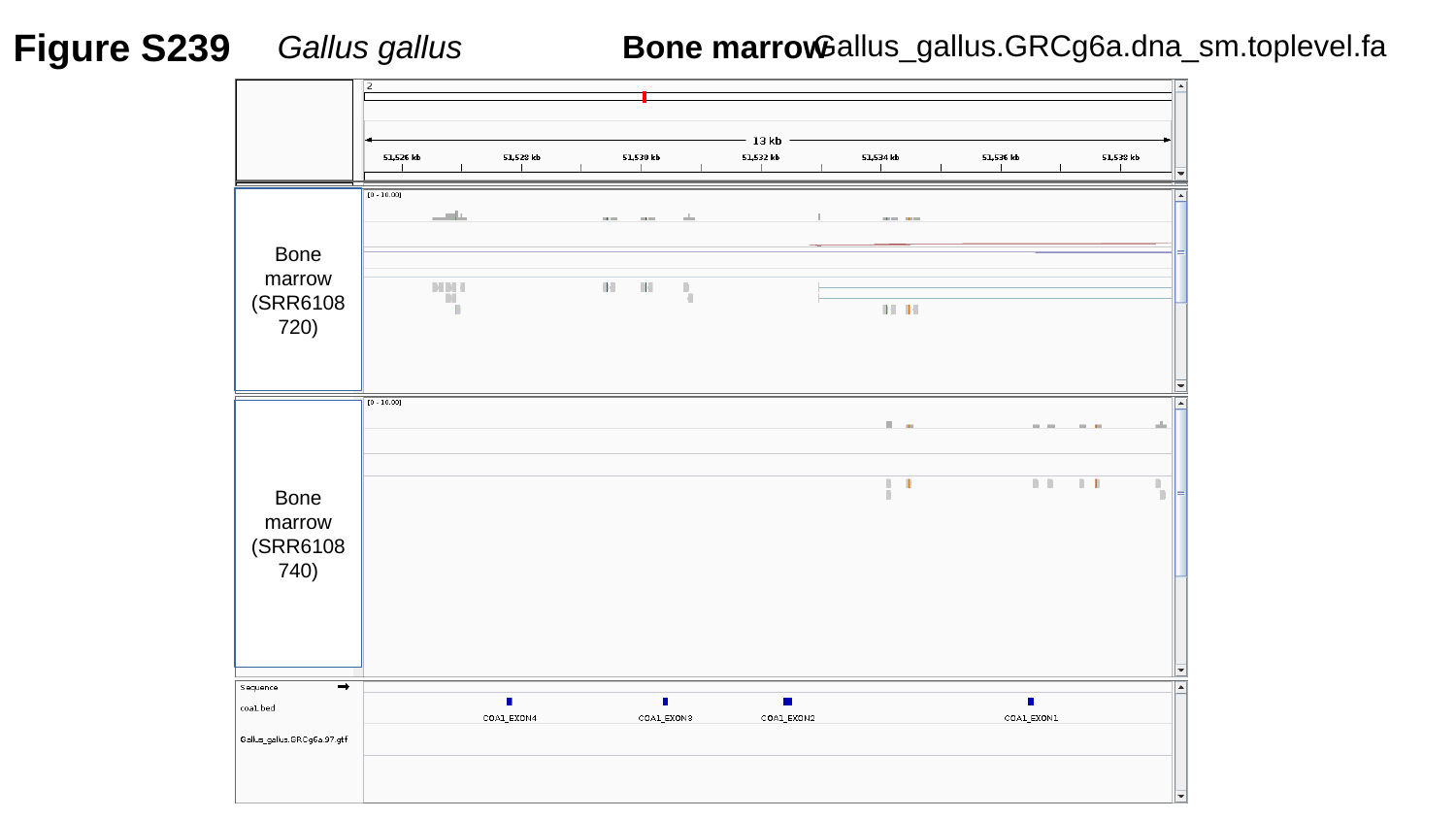

Figure S239
Gallus gallus	 Bone marrow
Gallus_gallus.GRCg6a.dna_sm.toplevel.fa
Bone marrow
(SRR6108720)
Bone marrow
(SRR6108740)

### Slide 240
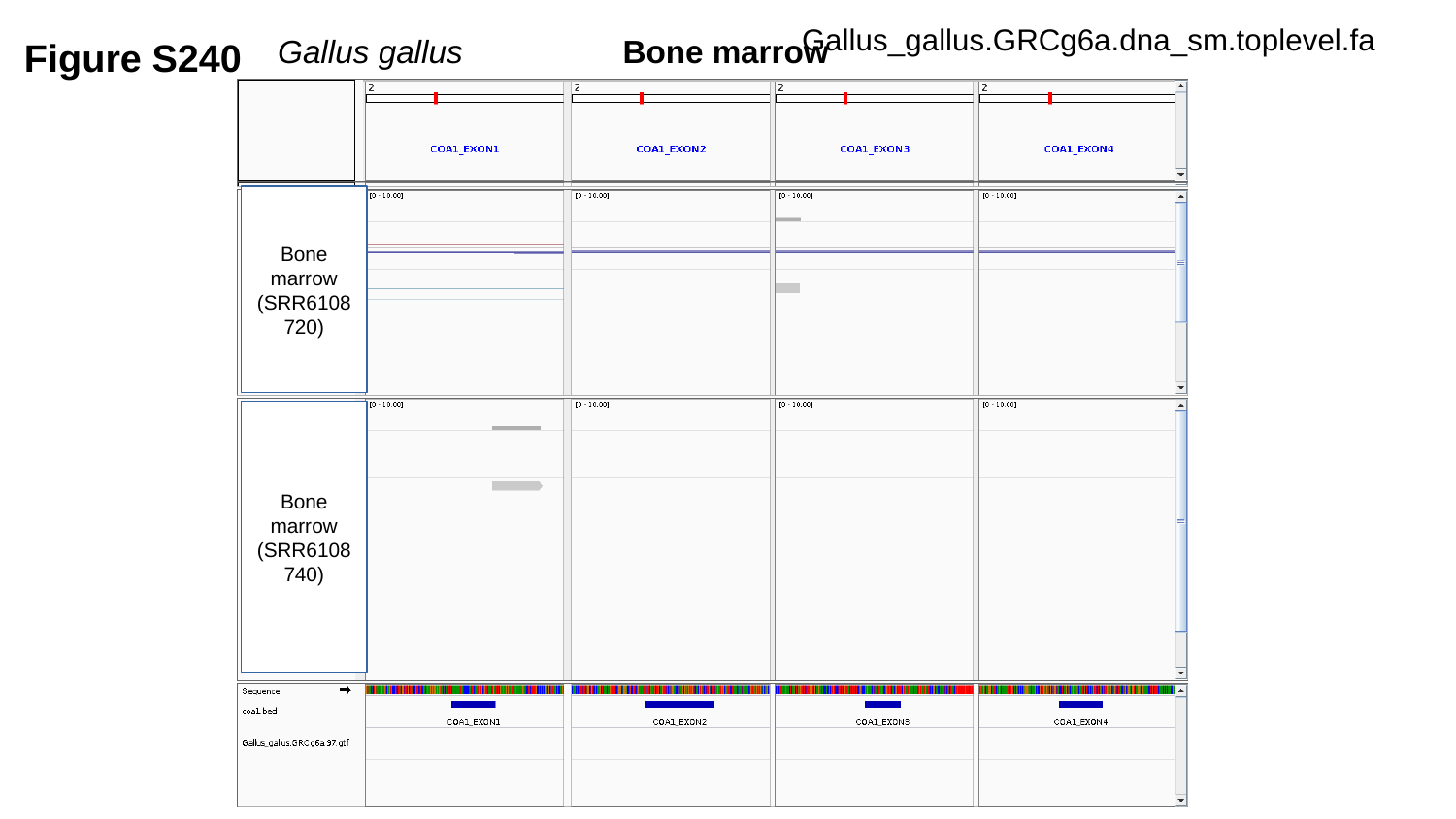

Gallus_gallus.GRCg6a.dna_sm.toplevel.fa
Figure S240
Gallus gallus	 Bone marrow
Bone marrow
(SRR6108720)
Bone marrow
(SRR6108740)

### Slide 241
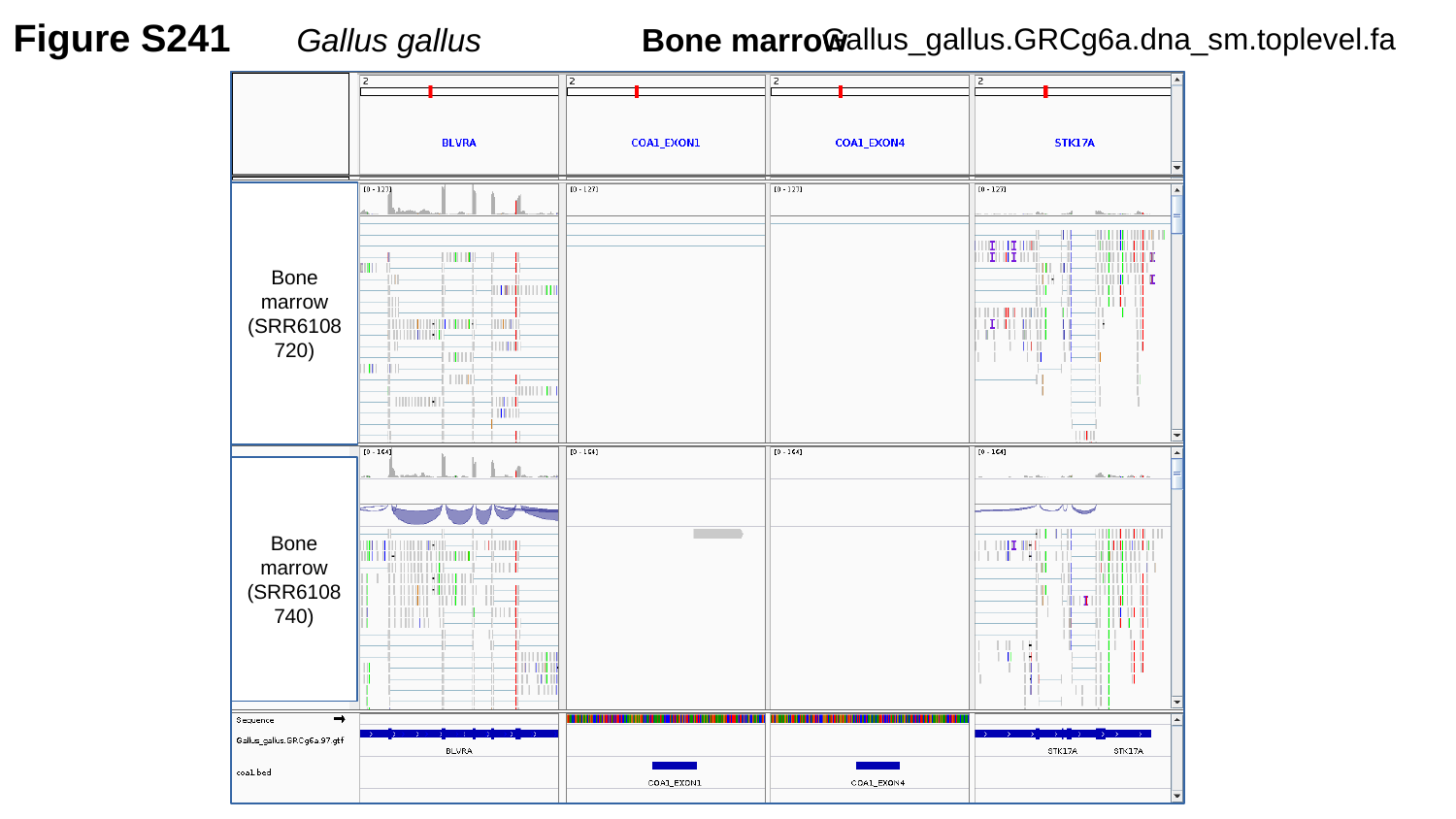

Figure S241
Gallus gallus	 Bone marrow
Gallus_gallus.GRCg6a.dna_sm.toplevel.fa
Bone marrow
(SRR6108720)
Bone marrow
(SRR6108740)

### Slide 242
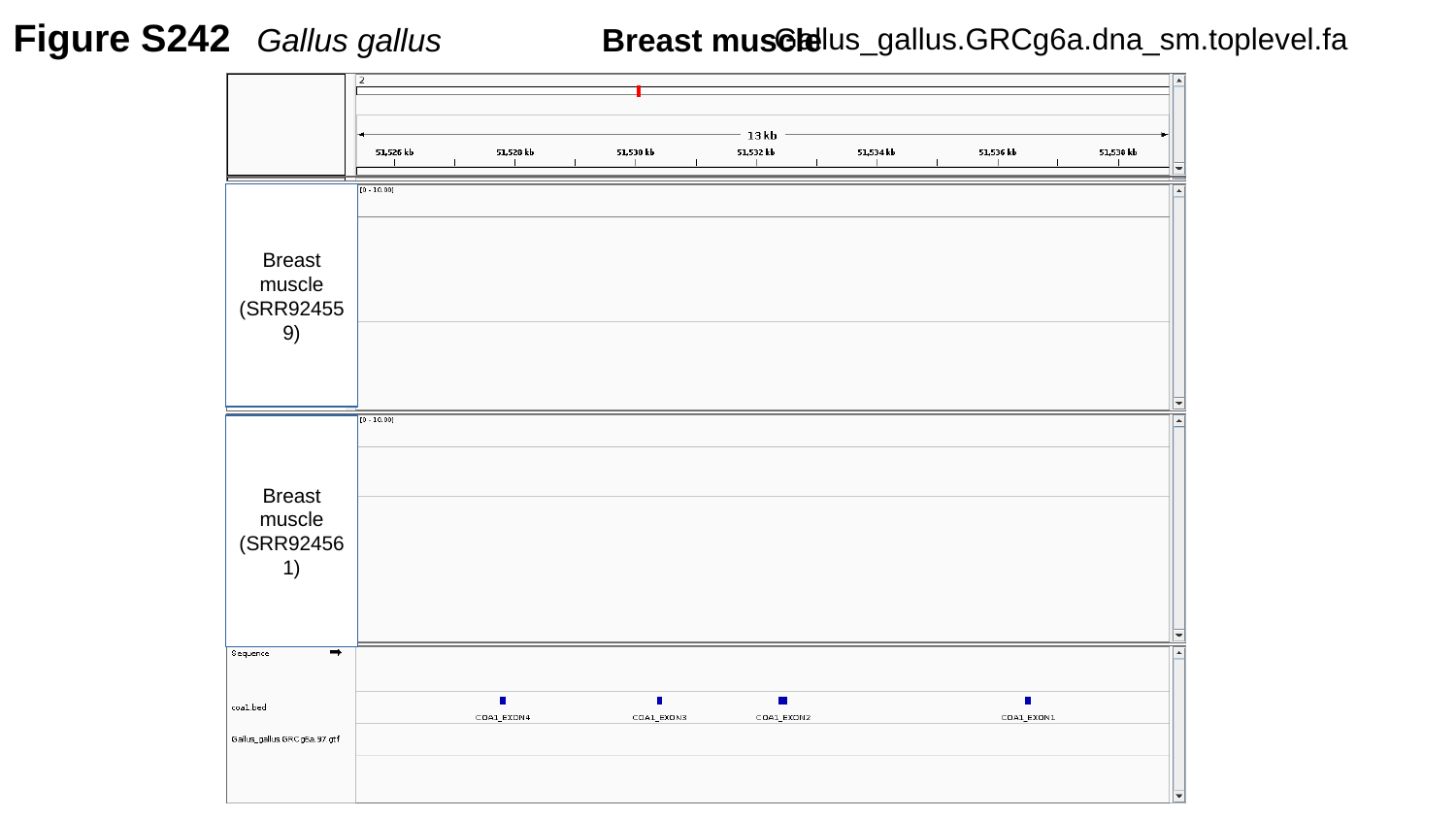

Figure S242
Gallus gallus	 Breast muscle
Gallus_gallus.GRCg6a.dna_sm.toplevel.fa
Breast muscle
(SRR924559)
Breast muscle
(SRR924561)

### Slide 243
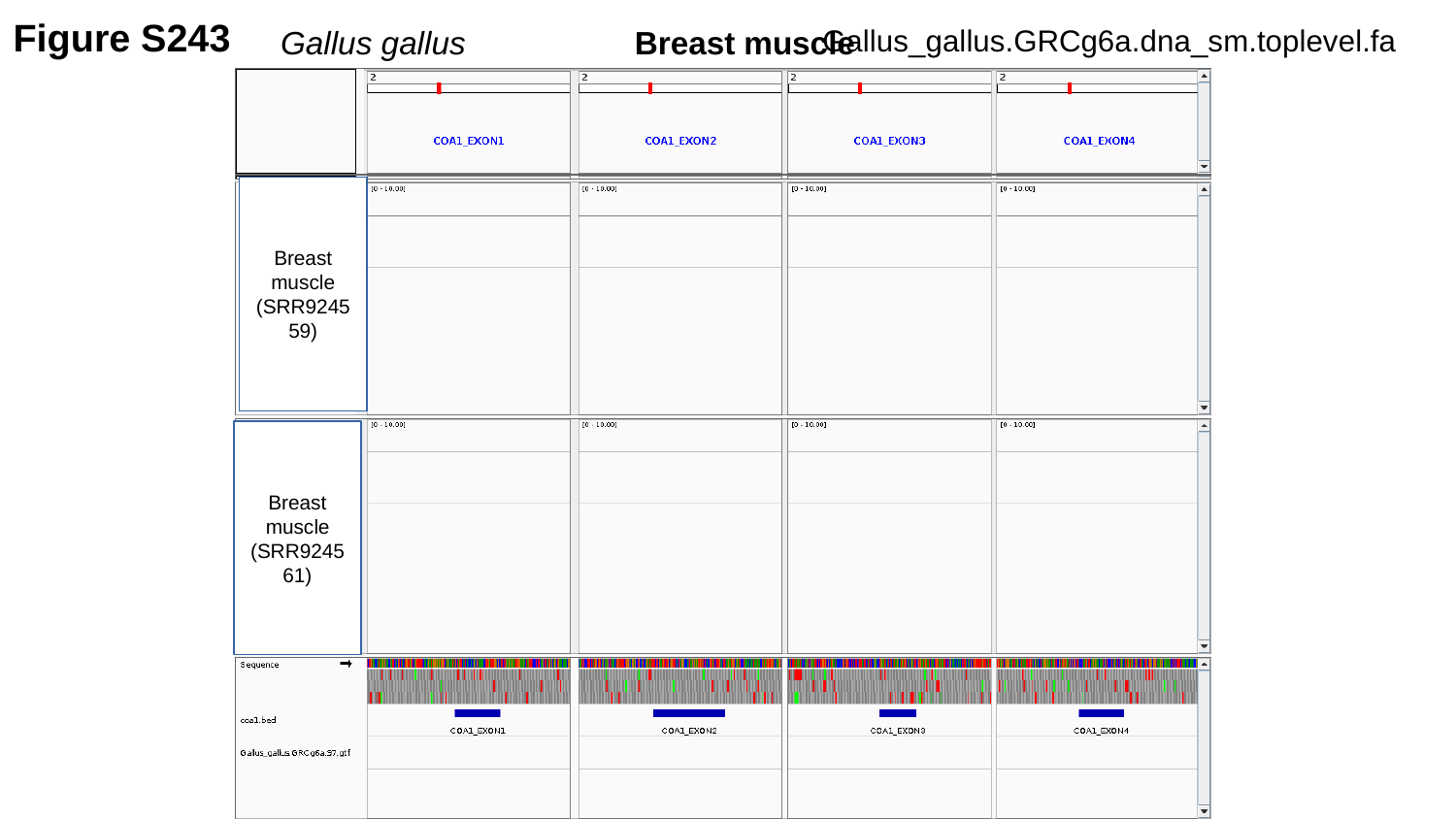

Figure S243
Gallus_gallus.GRCg6a.dna_sm.toplevel.fa
Gallus gallus	 Breast muscle
Breast muscle
(SRR924559)
Breast muscle
(SRR924561)

### Slide 244
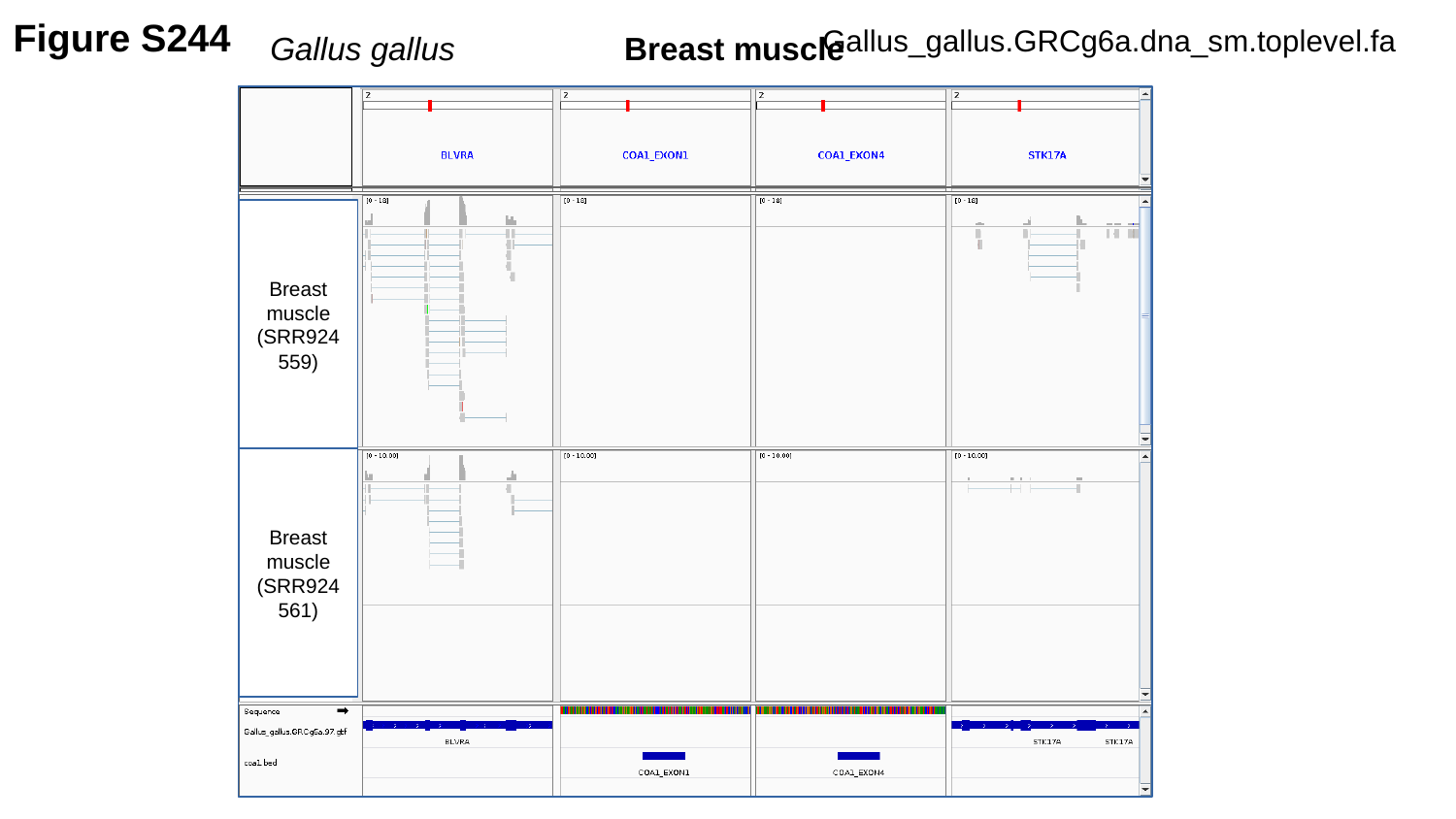

Figure S244
Gallus_gallus.GRCg6a.dna_sm.toplevel.fa
Gallus gallus	 Breast muscle
Breast muscle
(SRR924559)
Breast muscle
(SRR924561)

### Slide 245
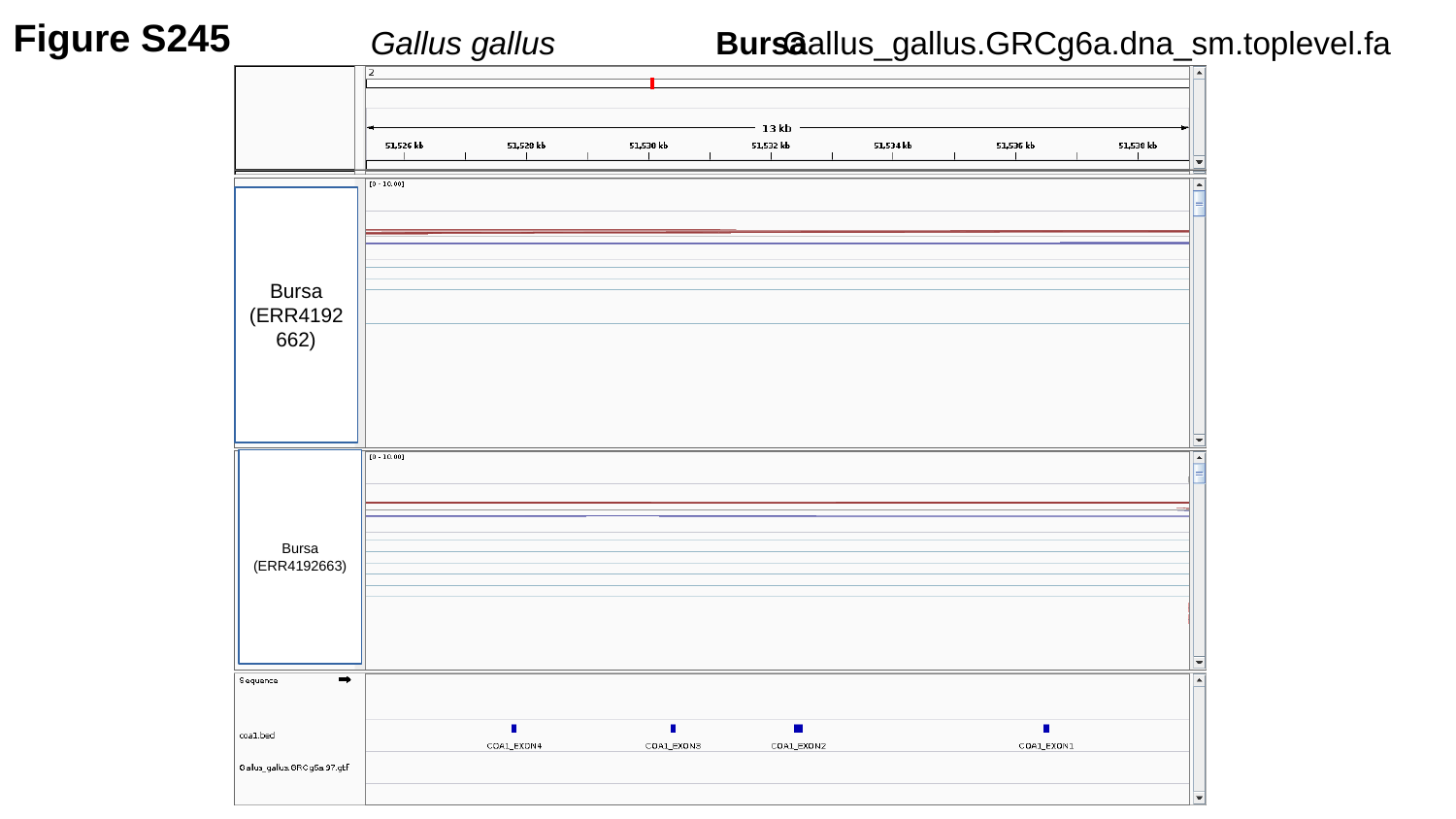

Figure S245
Gallus_gallus.GRCg6a.dna_sm.toplevel.fa
Gallus gallus	 Bursa
Bursa
(ERR4192662)
Bursa
(ERR4192663)

### Slide 246
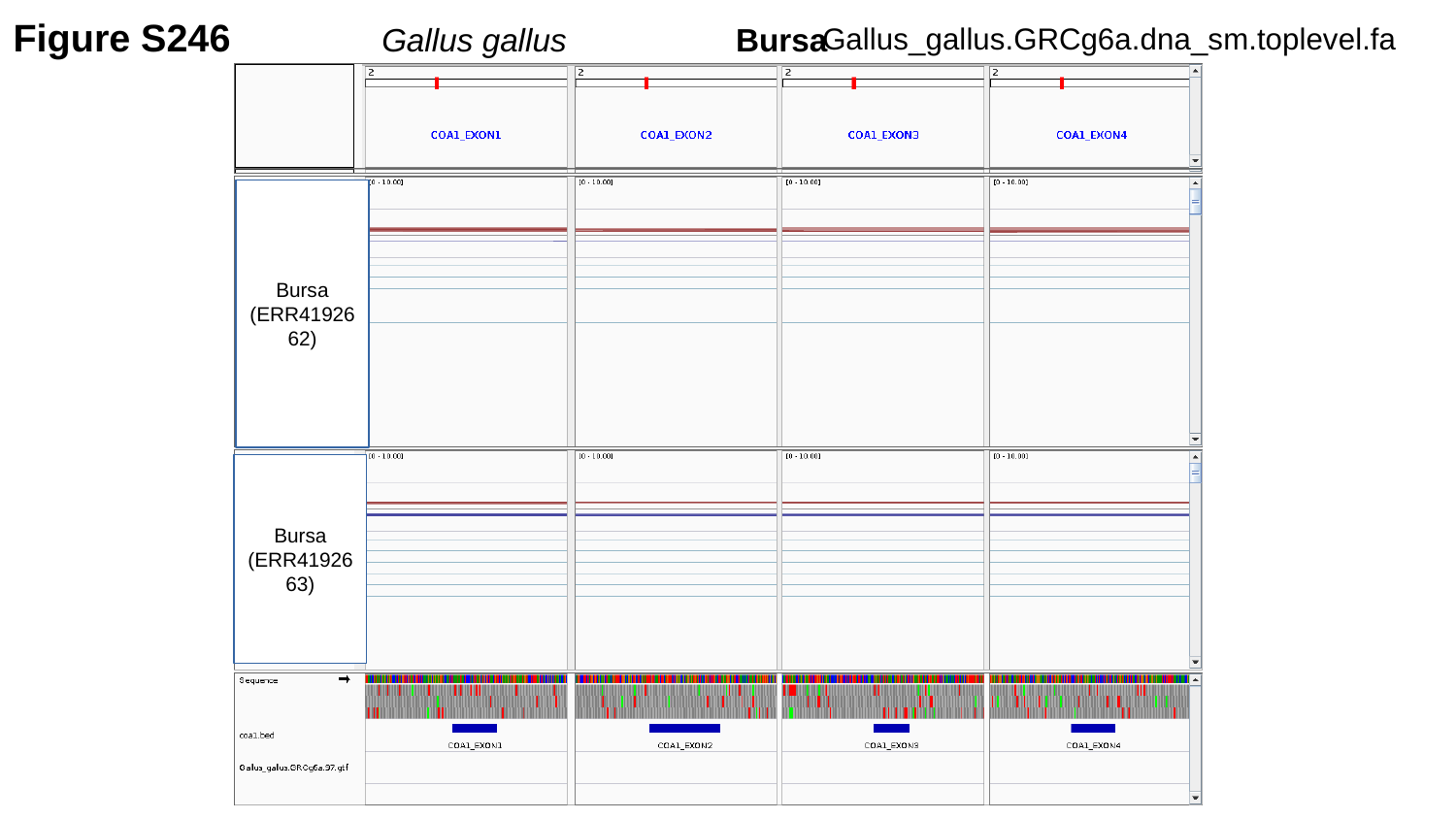

Figure S246
Gallus gallus	 Bursa
Gallus_gallus.GRCg6a.dna_sm.toplevel.fa
Bursa
(ERR4192662)
Bursa
(ERR4192663)

### Slide 247
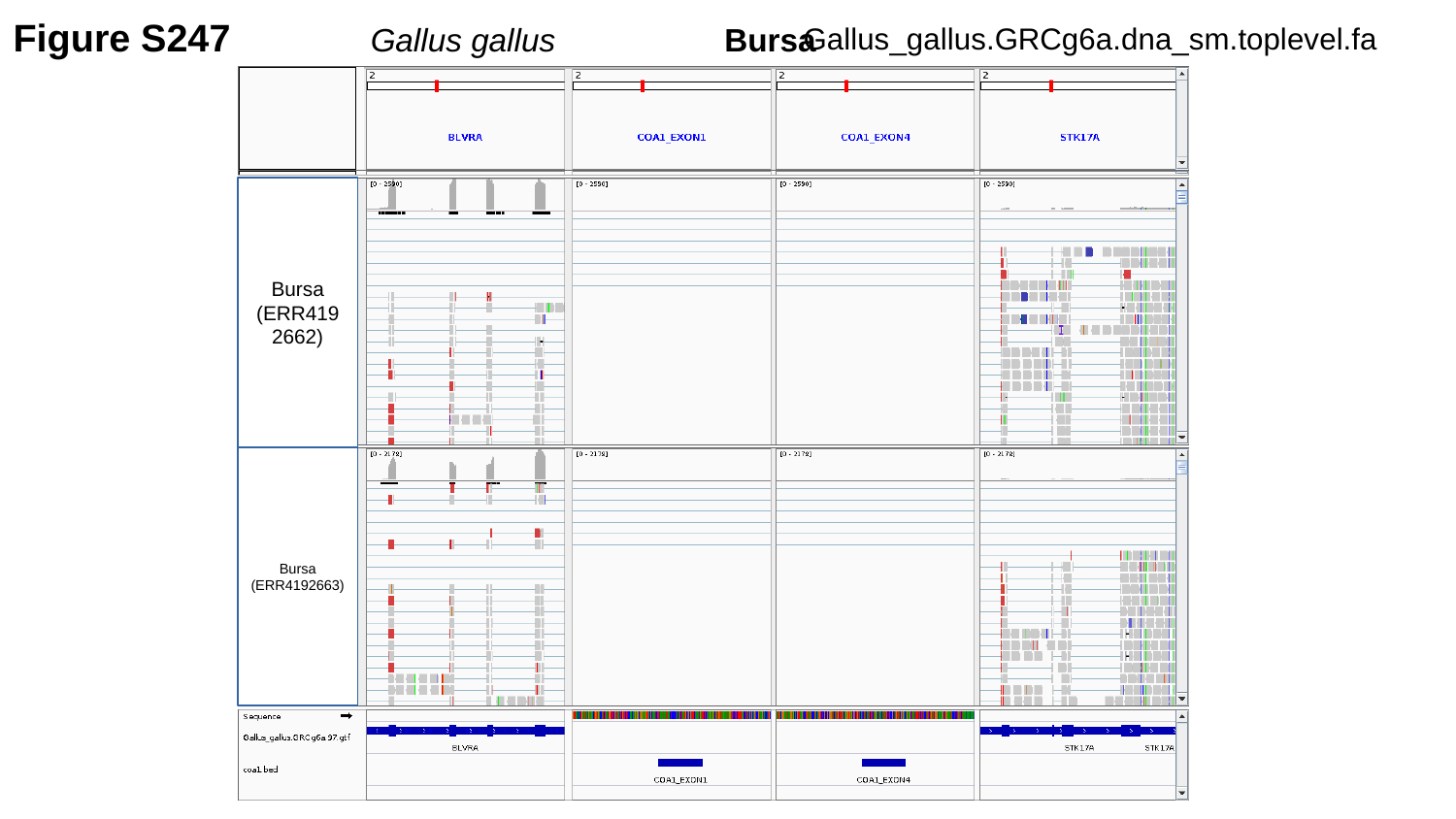

Figure S247
Gallus gallus	 Bursa
Gallus_gallus.GRCg6a.dna_sm.toplevel.fa
Bursa
(ERR4192662)
Bursa
(ERR4192663)

### Slide 248
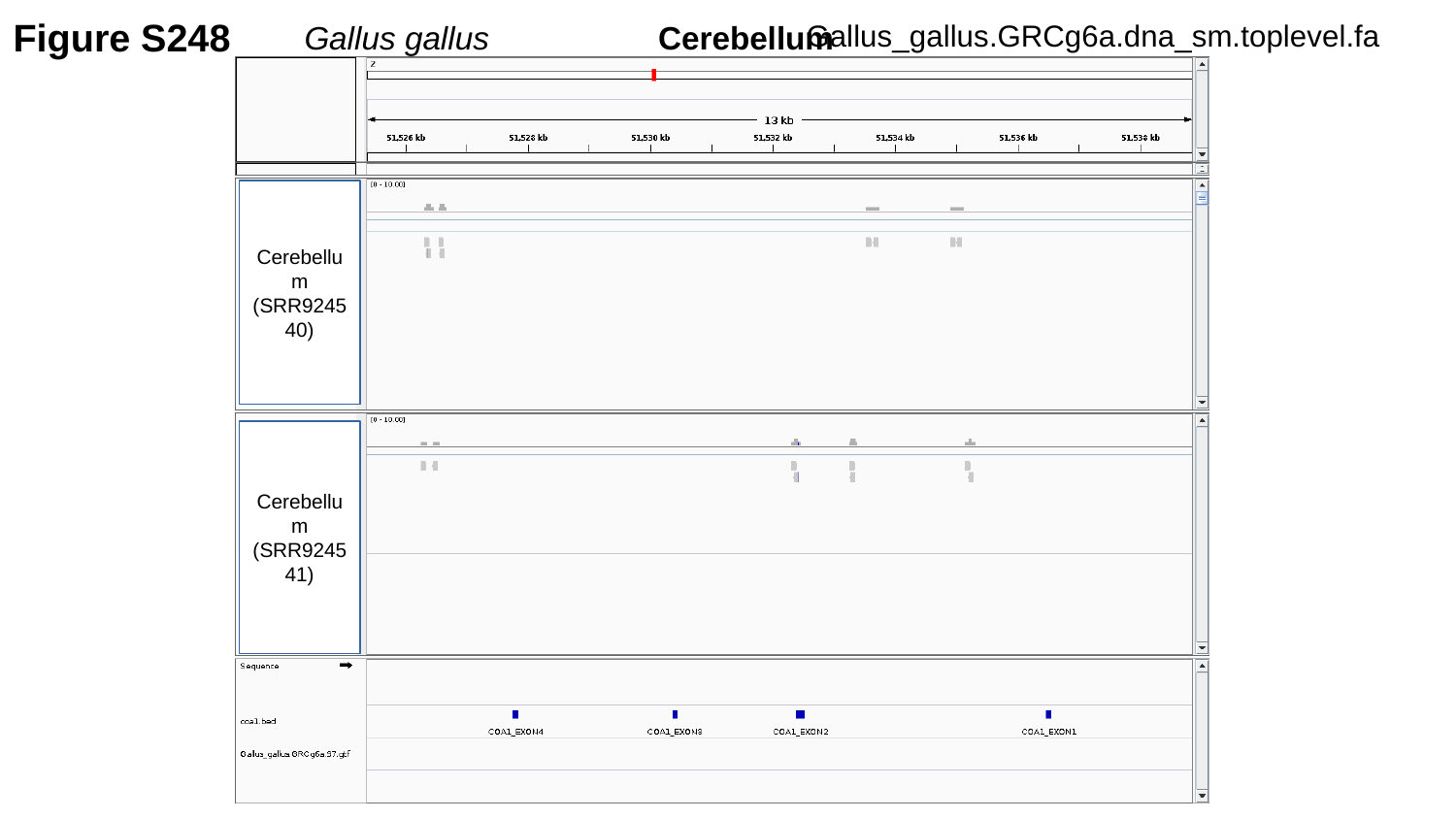

Figure S248
Gallus gallus	 Cerebellum
Gallus_gallus.GRCg6a.dna_sm.toplevel.fa
Cerebellum
(SRR924540)
Cerebellum
(SRR924541)

### Slide 249
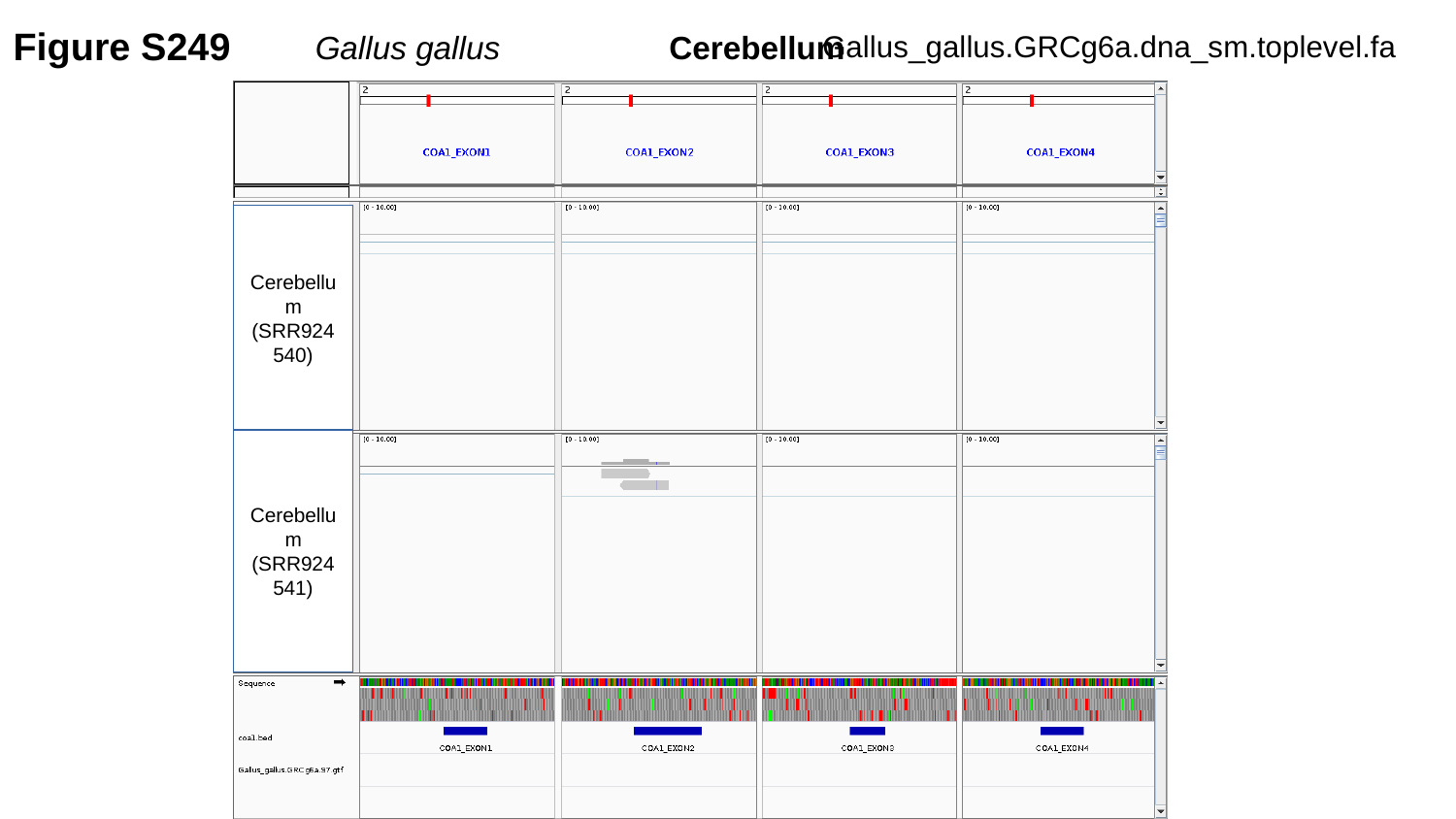

Figure S249
Gallus_gallus.GRCg6a.dna_sm.toplevel.fa
Gallus gallus	 Cerebellum
Cerebellum
(SRR924540)
Cerebellum
(SRR924541)

### Slide 250
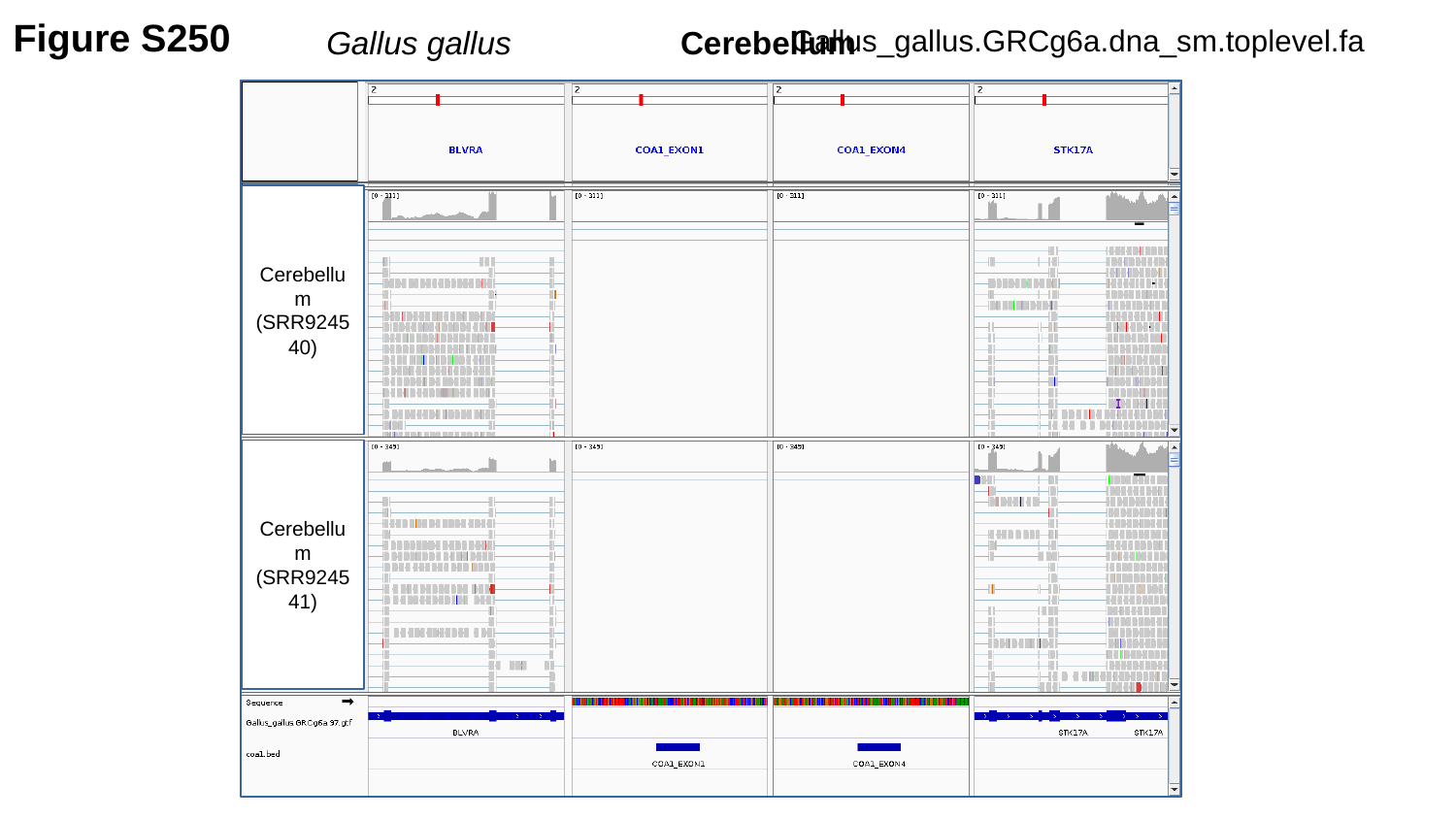

Figure S250
Gallus gallus	 Cerebellum
Gallus_gallus.GRCg6a.dna_sm.toplevel.fa
Cerebellum
(SRR924540)
Cerebellum
(SRR924541)

### Slide 251
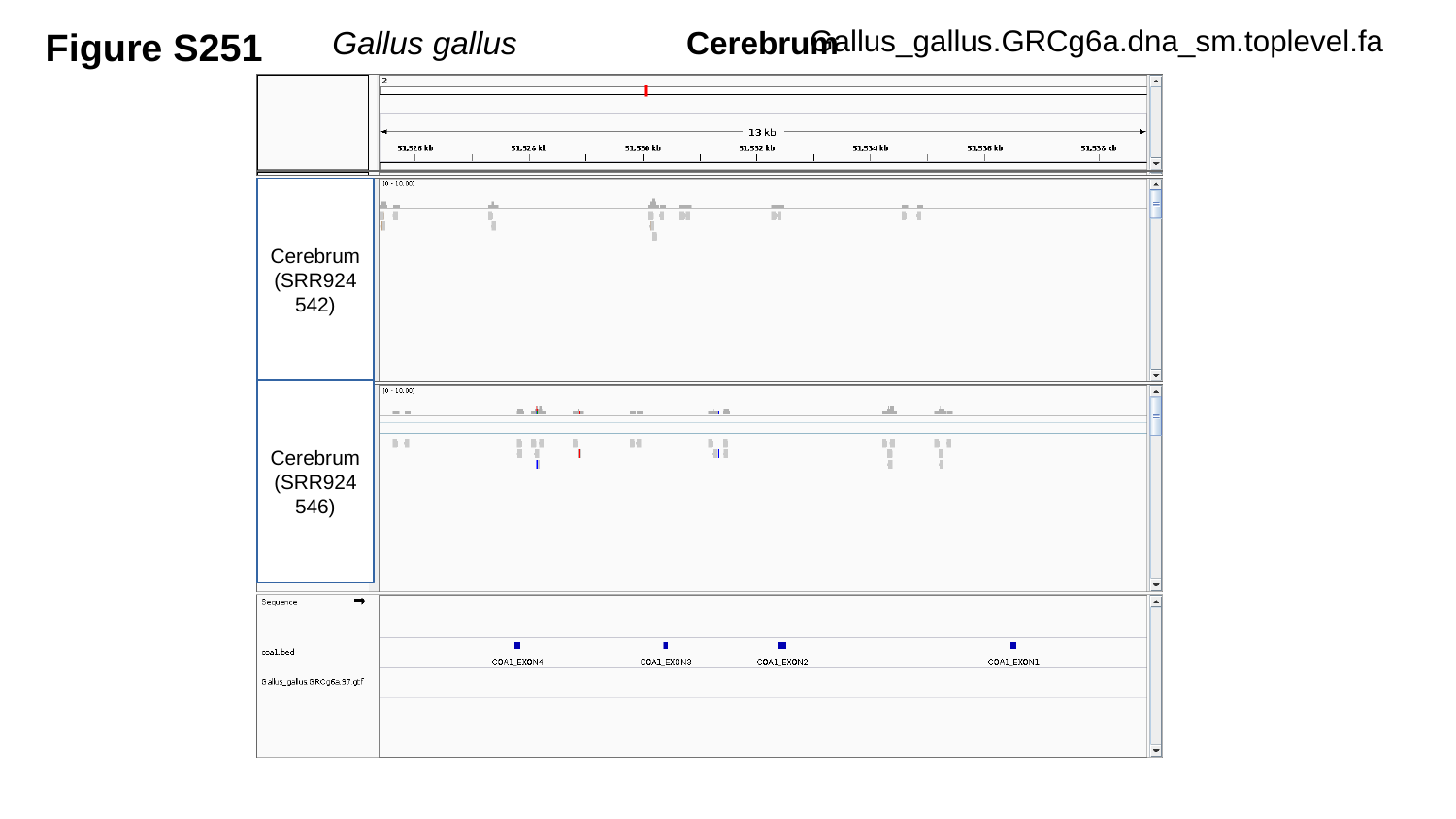

Figure S251
Gallus gallus	 Cerebrum
Gallus_gallus.GRCg6a.dna_sm.toplevel.fa
Cerebrum
(SRR924542)
Cerebrum
(SRR924546)

### Slide 252
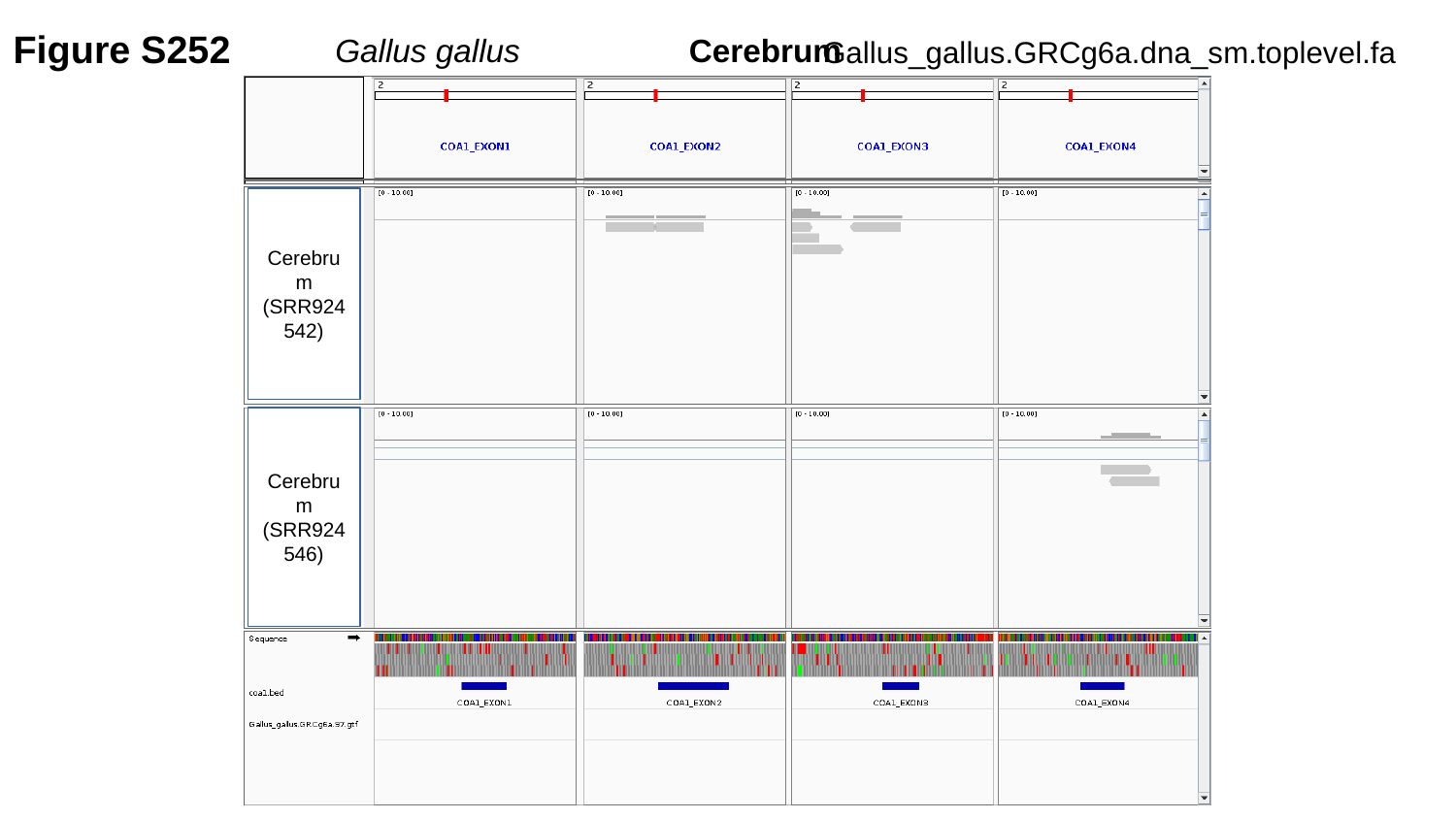

Figure S252
Gallus gallus	 Cerebrum
Gallus_gallus.GRCg6a.dna_sm.toplevel.fa
Cerebrum
(SRR924542)
Cerebrum
(SRR924546)

### Slide 253
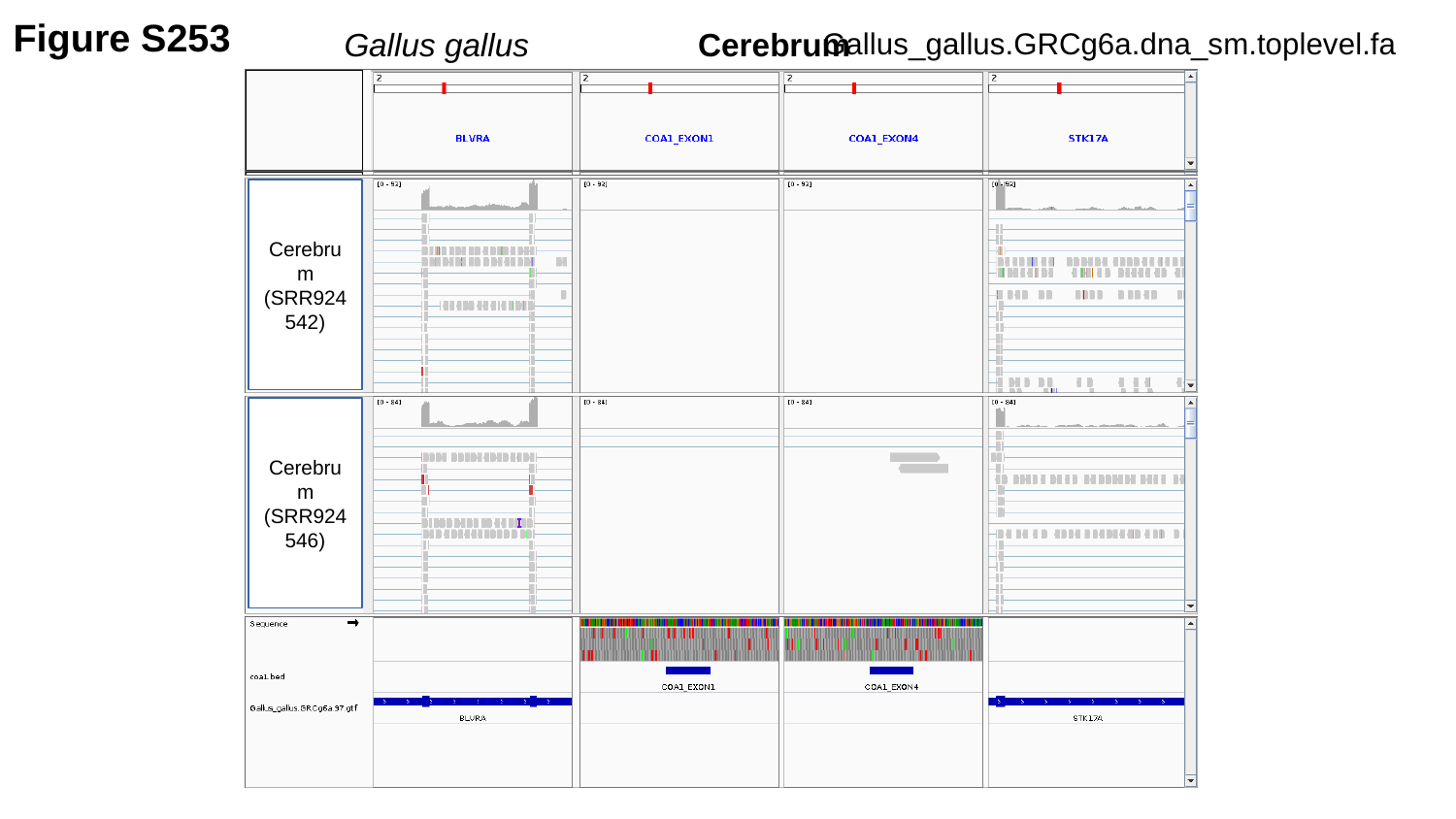

Figure S253
Gallus gallus	 Cerebrum
Gallus_gallus.GRCg6a.dna_sm.toplevel.fa
Cerebrum
(SRR924542)
Cerebrum
(SRR924546)

### Slide 254
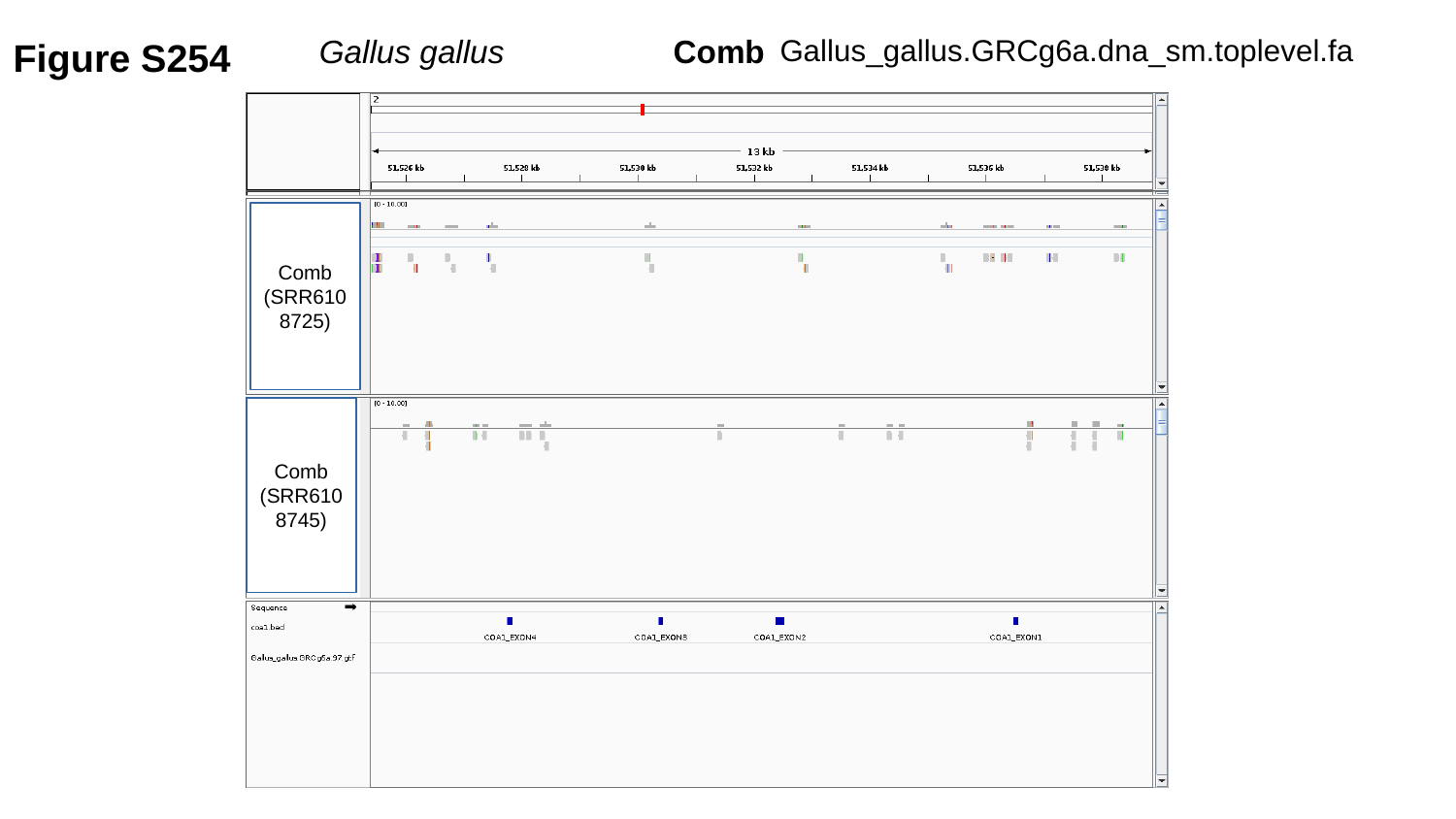

Figure S254
Gallus gallus	 Comb
Gallus_gallus.GRCg6a.dna_sm.toplevel.fa
Comb
(SRR6108725)
Comb
(SRR6108745)

### Slide 255
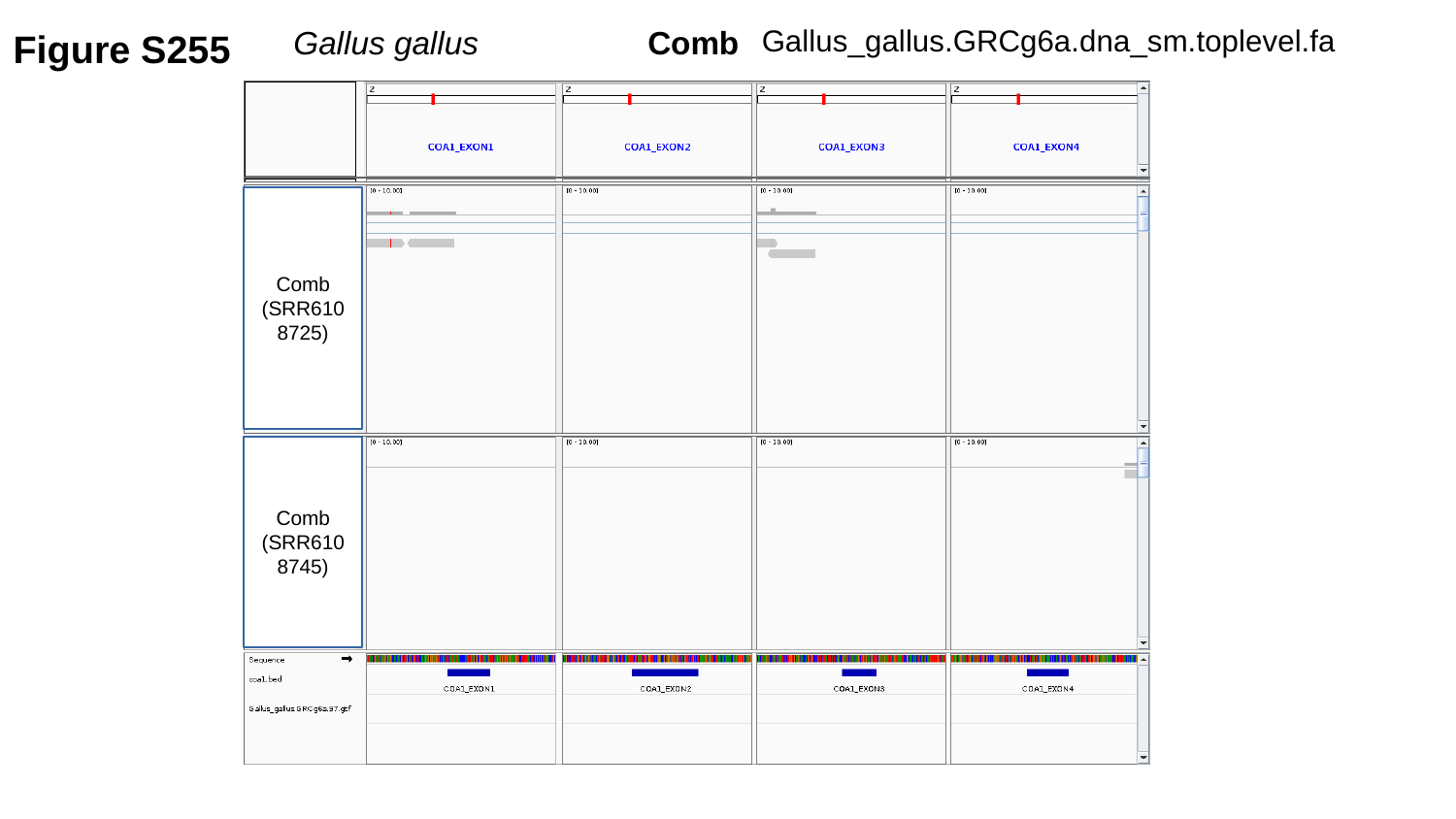

Figure S255
Gallus gallus	 Comb
Gallus_gallus.GRCg6a.dna_sm.toplevel.fa
Comb
(SRR6108725)
Comb
(SRR6108745)

### Slide 256
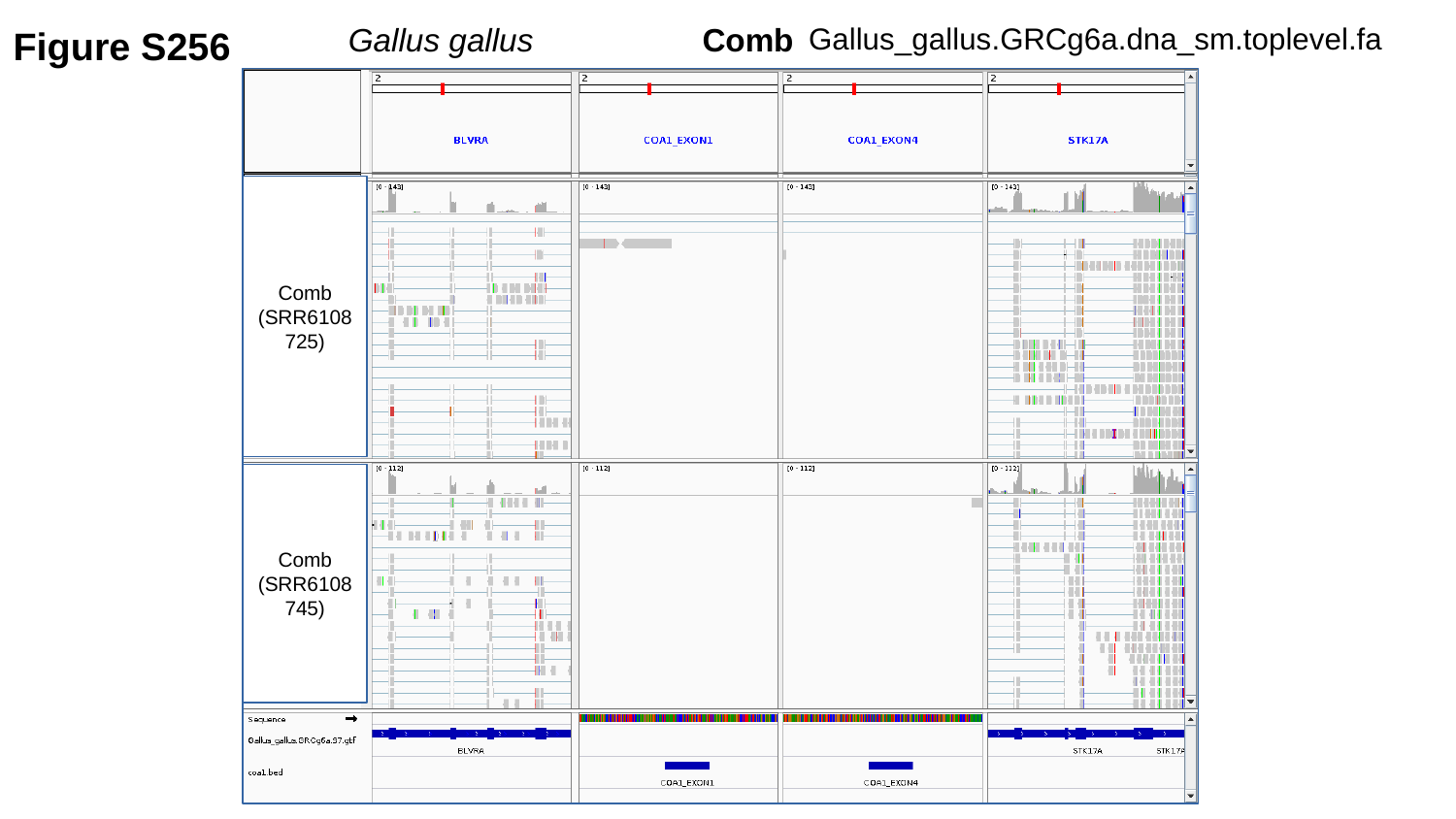

Figure S256
Gallus gallus	 Comb
Gallus_gallus.GRCg6a.dna_sm.toplevel.fa
Comb
(SRR6108725)
Comb
(SRR6108745)

### Slide 257
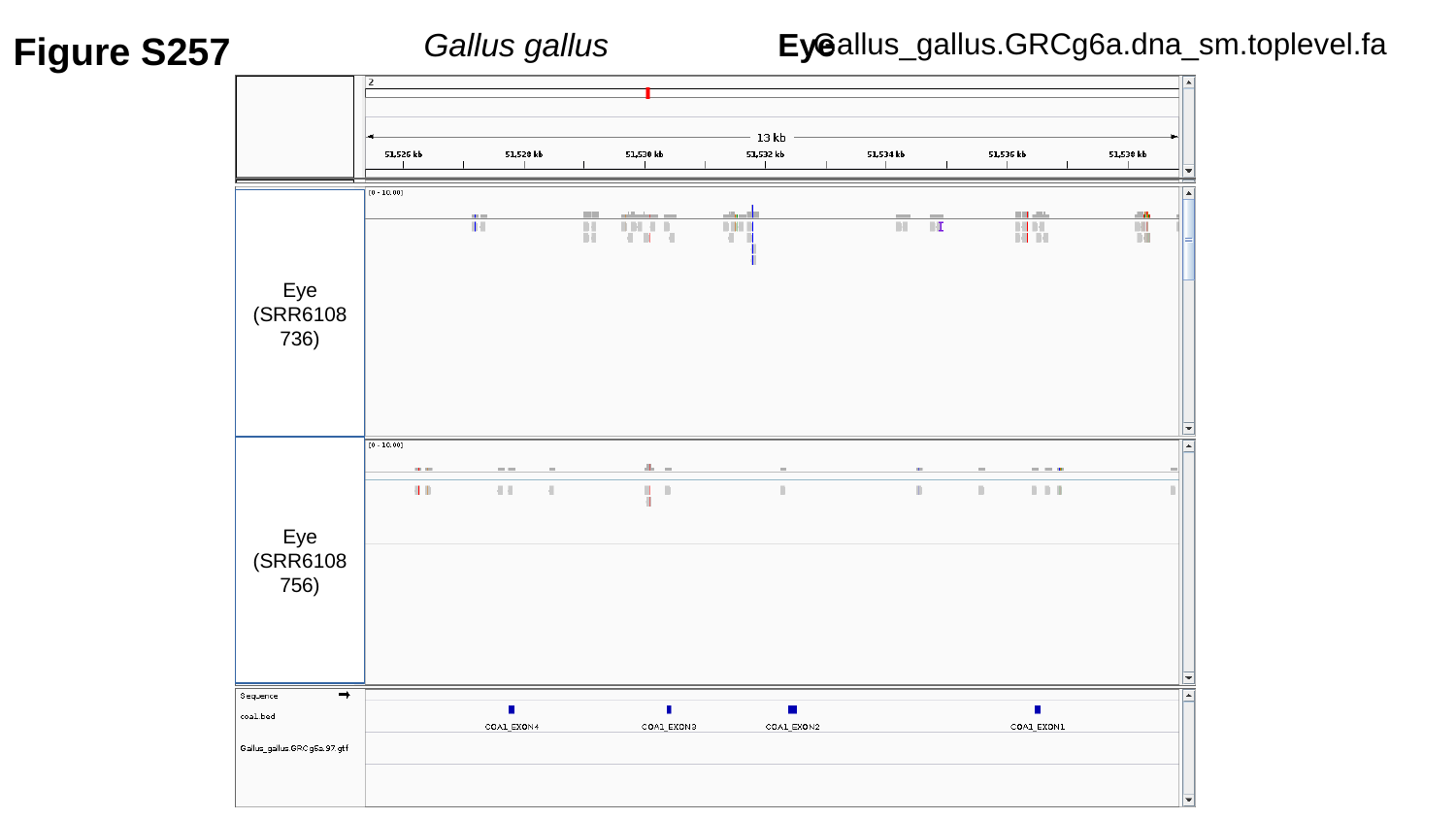

Figure S257
Gallus gallus	 Eye
Gallus_gallus.GRCg6a.dna_sm.toplevel.fa
Eye
(SRR6108736)
Eye
(SRR6108756)

### Slide 258
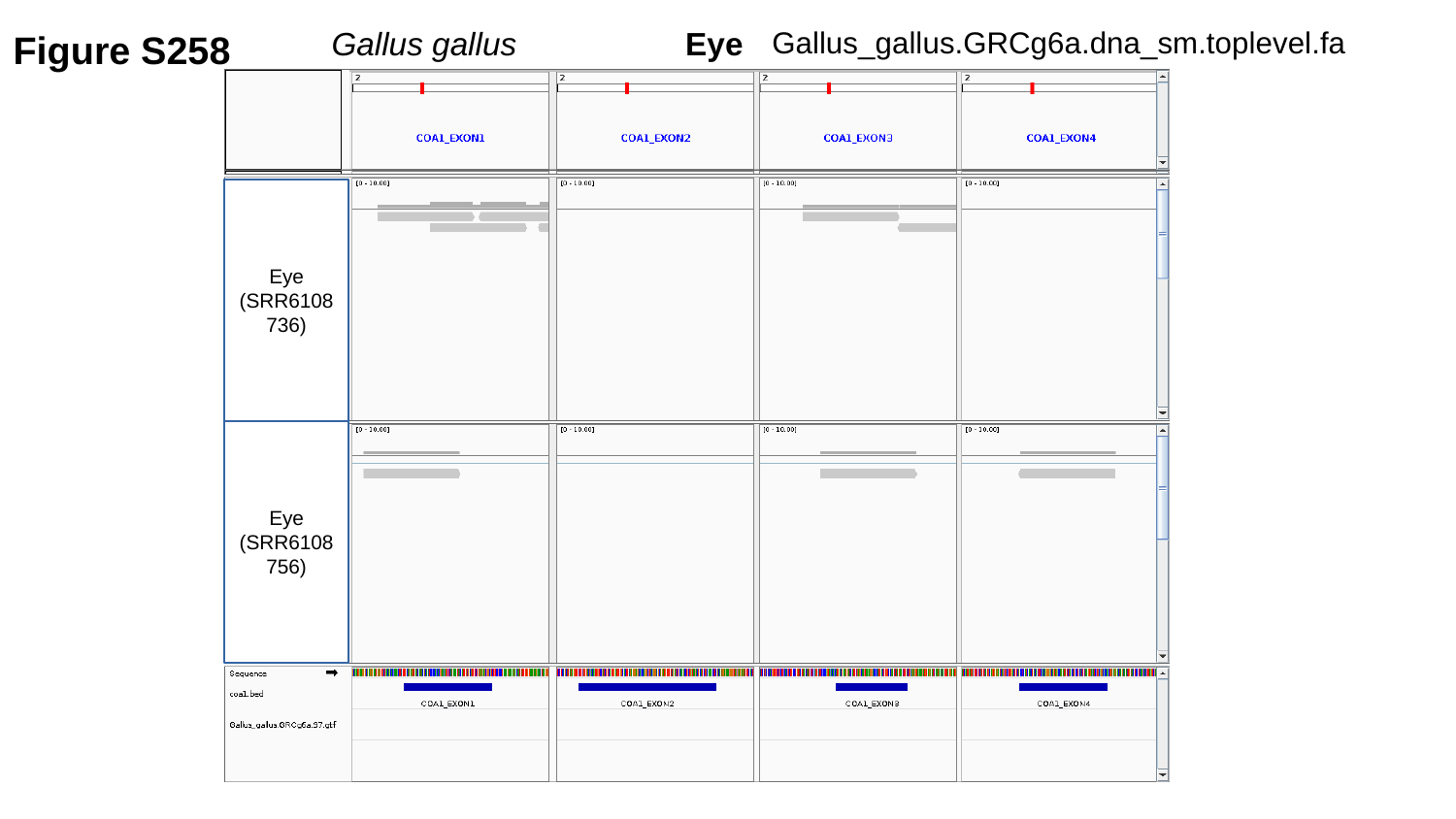

Figure S258
Gallus gallus	 Eye
Gallus_gallus.GRCg6a.dna_sm.toplevel.fa
Eye
(SRR6108736)
Eye
(SRR6108756)

### Slide 259
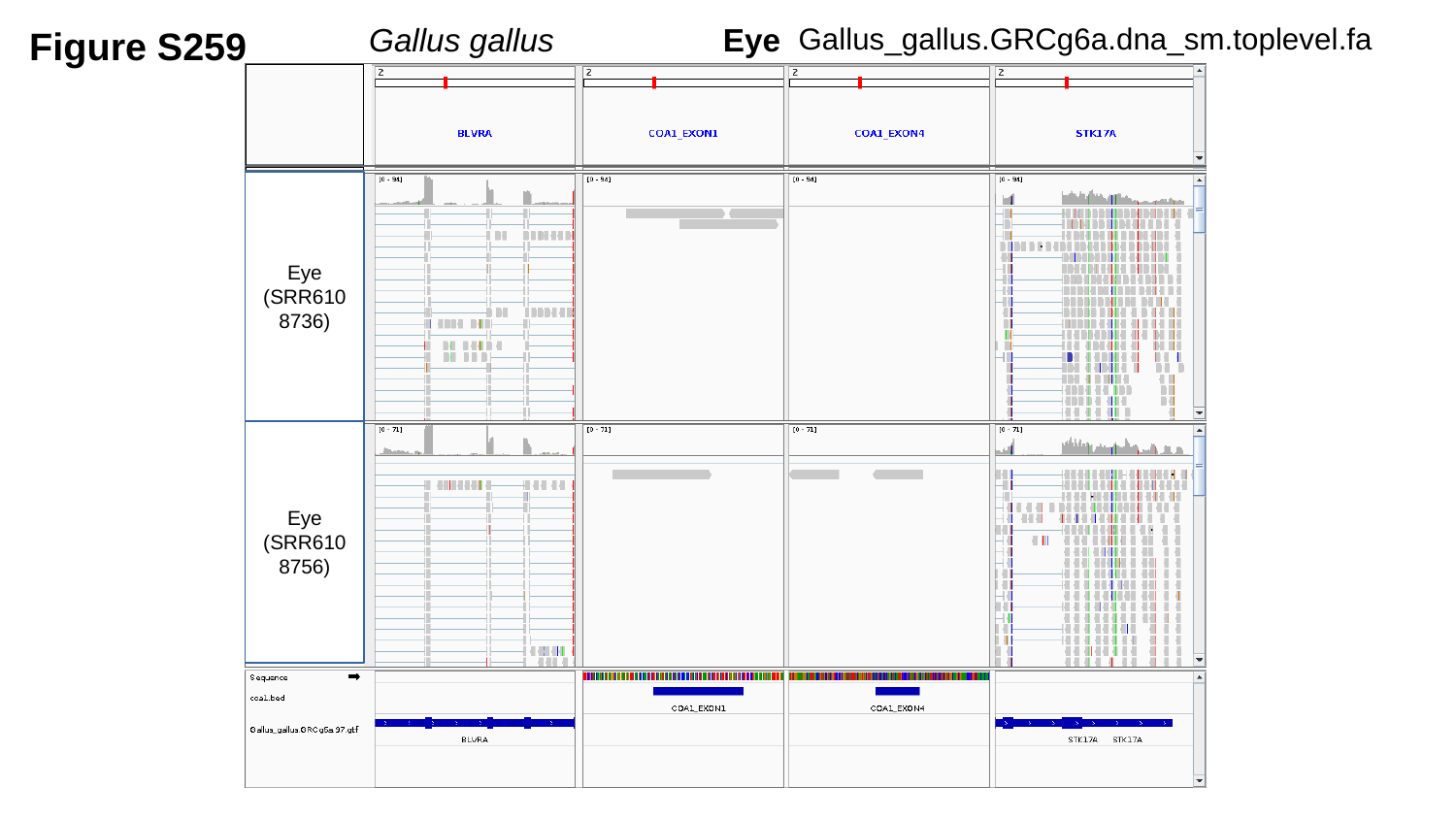

Figure S259
Gallus gallus	 Eye
Gallus_gallus.GRCg6a.dna_sm.toplevel.fa
Eye
(SRR6108736)
Eye
(SRR6108756)

### Slide 260
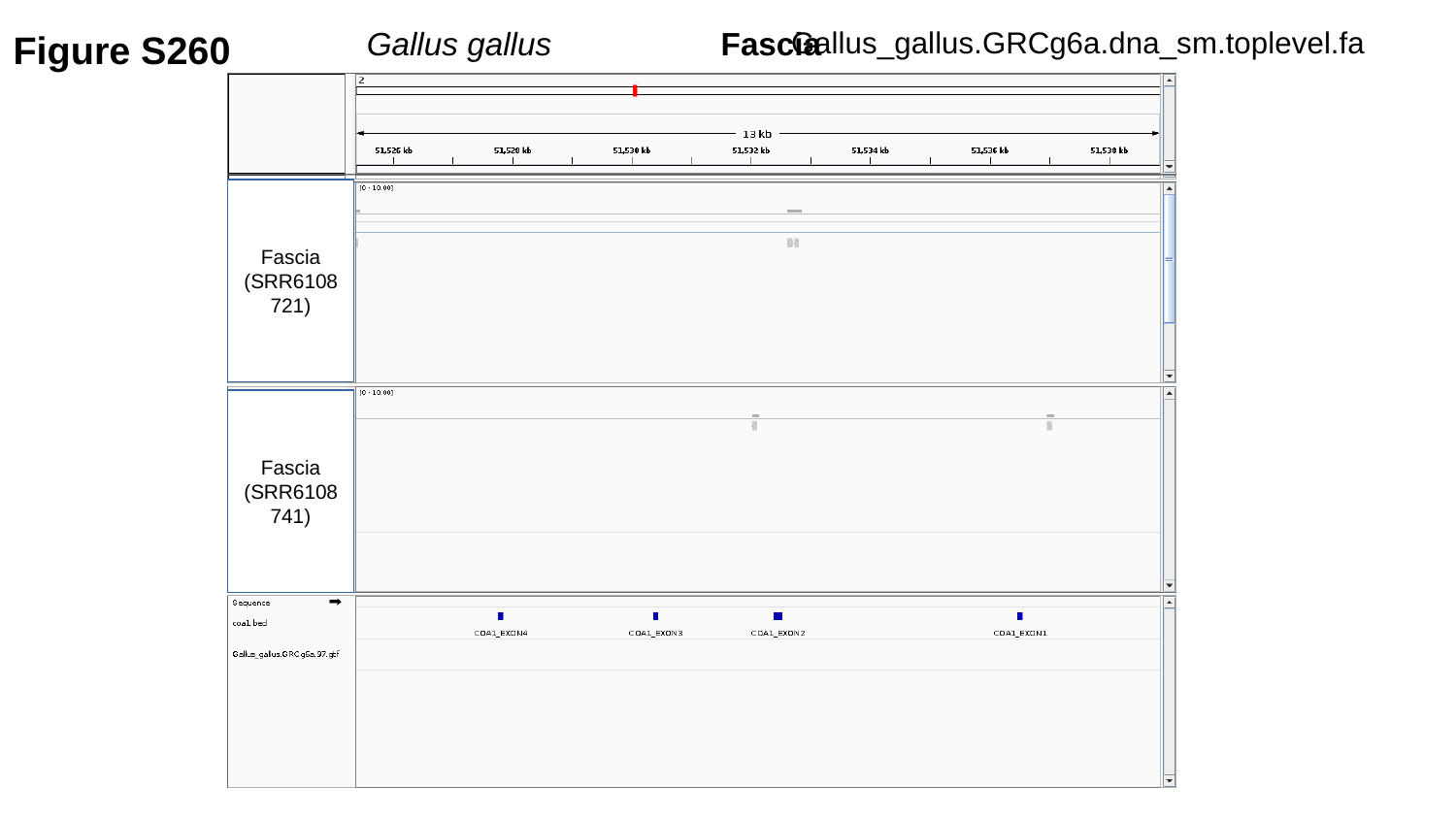

Figure S260
Gallus_gallus.GRCg6a.dna_sm.toplevel.fa
Gallus gallus	 Fascia
Fascia
(SRR6108721)
Fascia
(SRR6108741)

### Slide 261
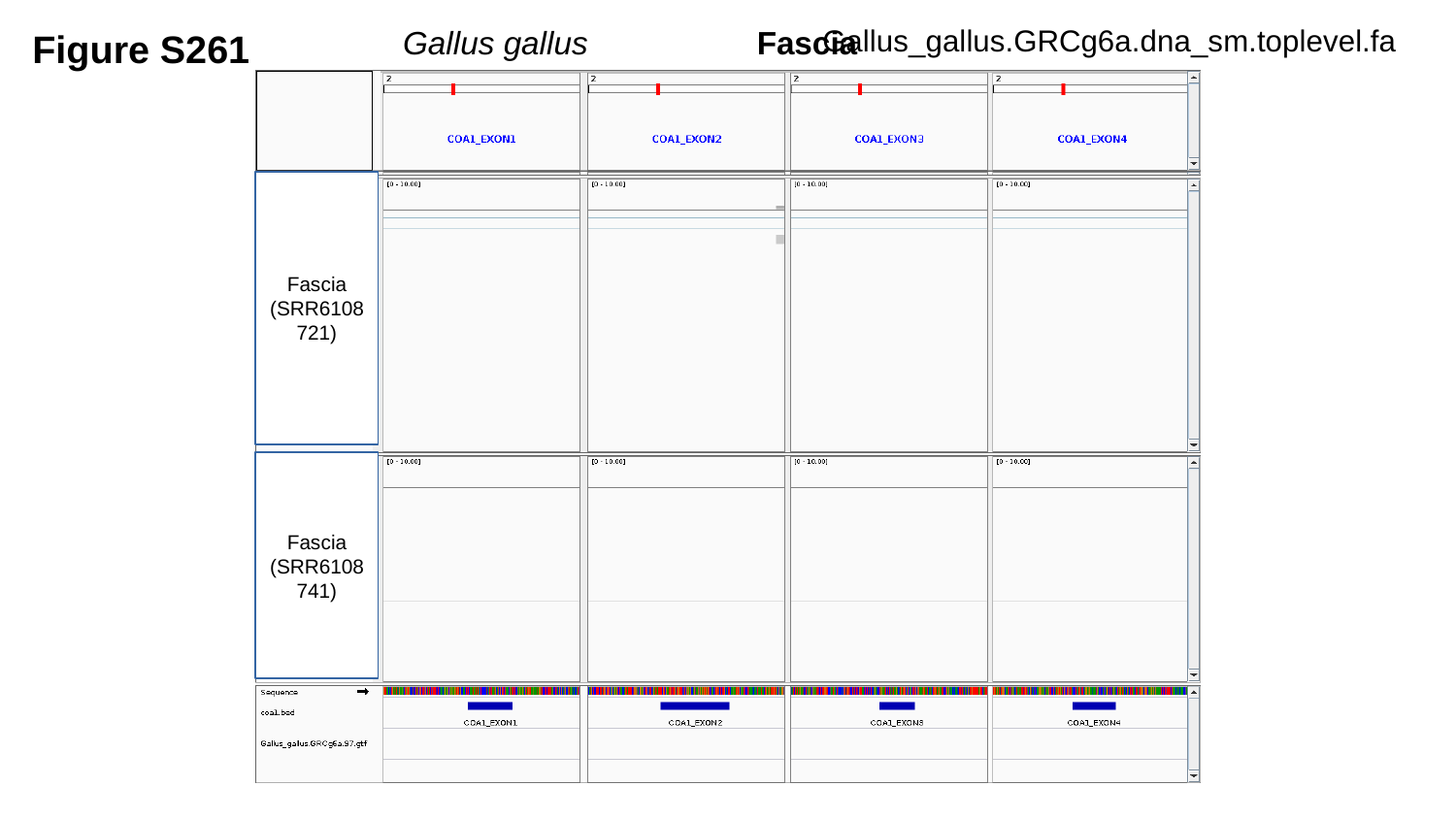

Figure S261
Gallus gallus	 Fascia
Gallus_gallus.GRCg6a.dna_sm.toplevel.fa
Fascia
(SRR6108721)
Fascia
(SRR6108741)

### Slide 262
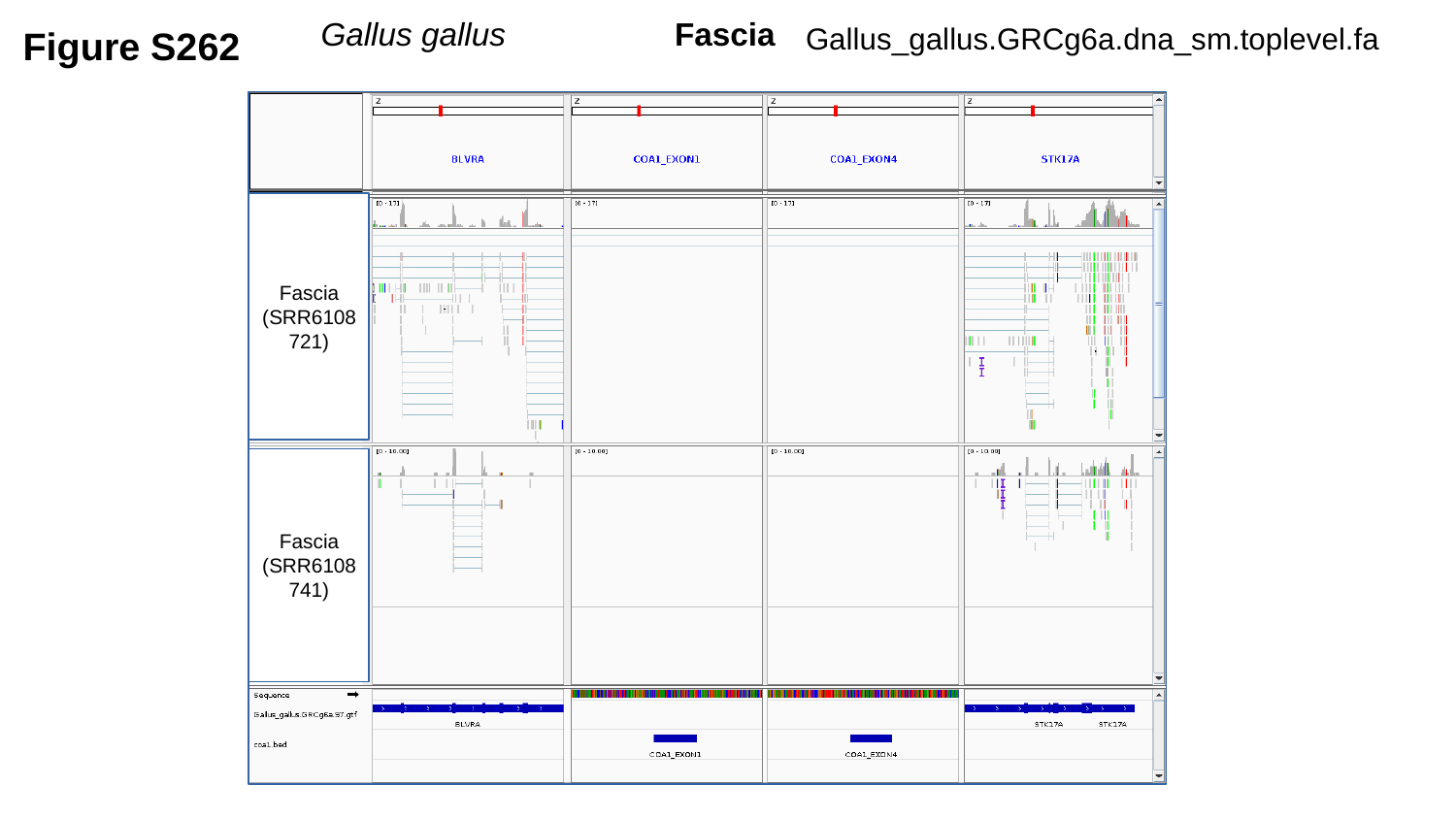

Gallus gallus	 Fascia
Figure S262
Gallus_gallus.GRCg6a.dna_sm.toplevel.fa
Fascia
(SRR6108721)
Fascia
(SRR6108741)

### Slide 263
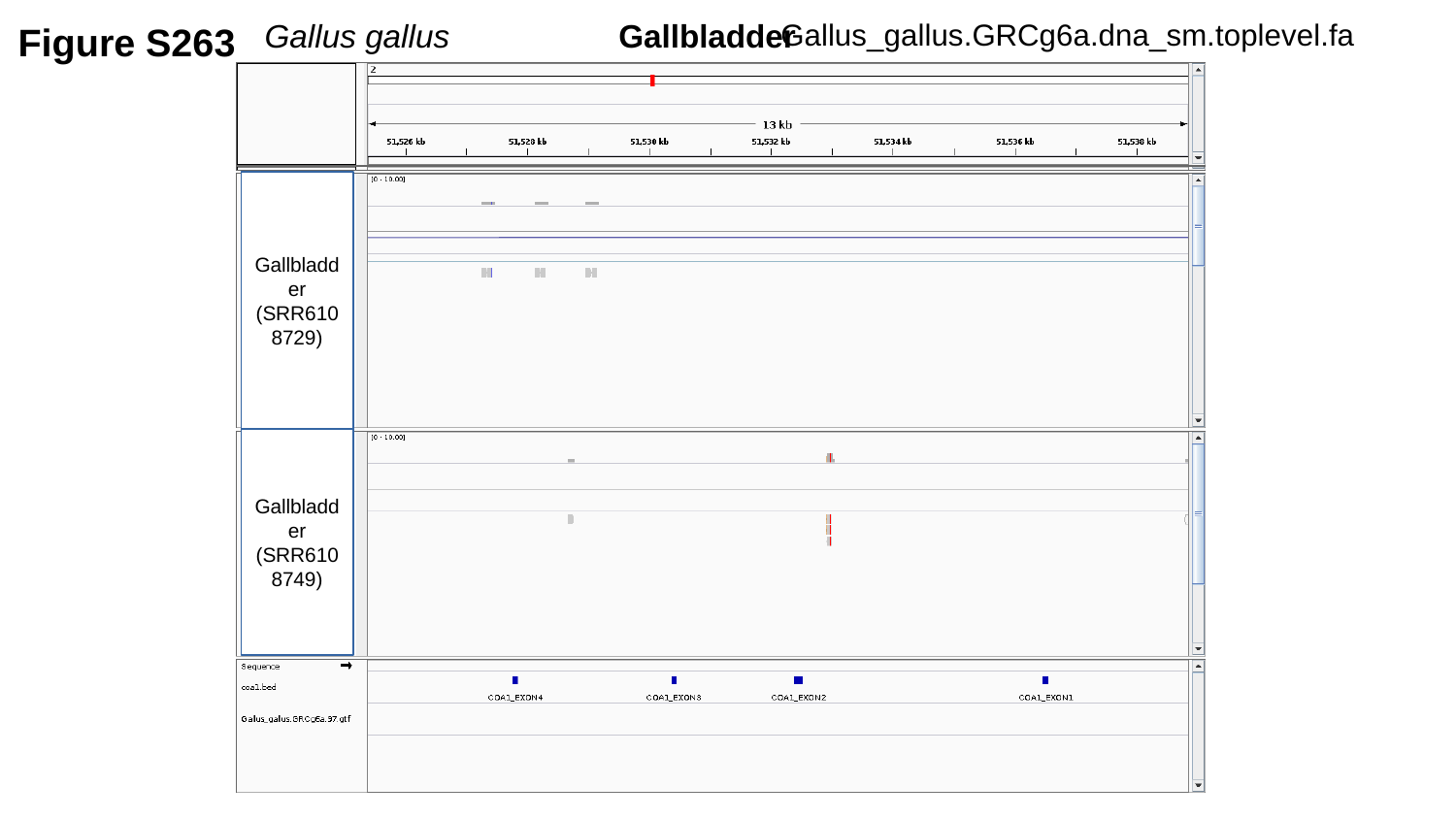

Figure S263
Gallus gallus	 Gallbladder
Gallus_gallus.GRCg6a.dna_sm.toplevel.fa
Gallbladder
(SRR6108729)
Gallbladder
(SRR6108749)

### Slide 264
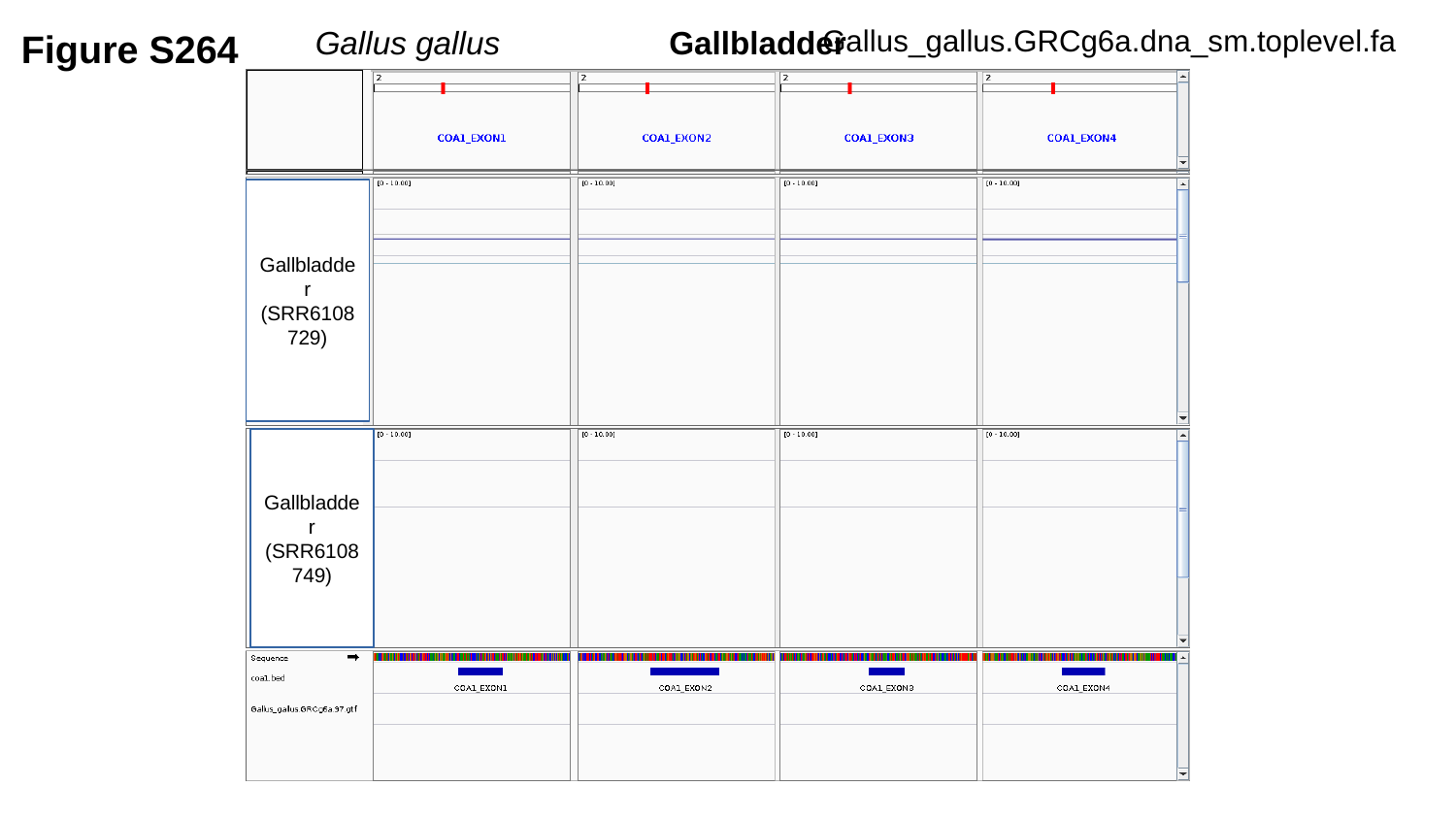

Figure S264
Gallus gallus	 Gallbladder
Gallus_gallus.GRCg6a.dna_sm.toplevel.fa
Gallbladder
(SRR6108729)
Gallbladder
(SRR6108749)

### Slide 265
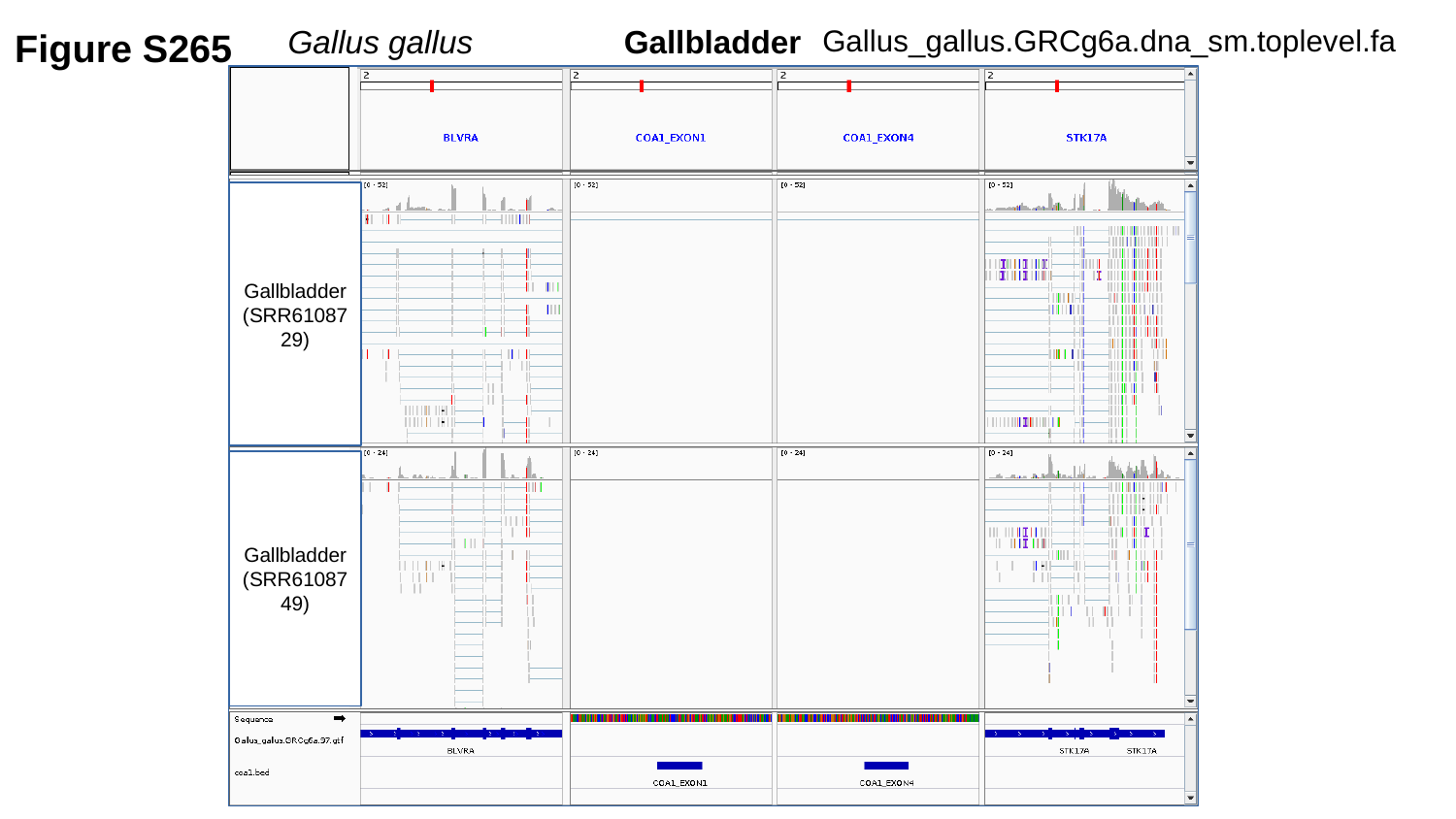

Figure S265
Gallus gallus	 Gallbladder
Gallus_gallus.GRCg6a.dna_sm.toplevel.fa
Gallbladder
(SRR6108729)
Gallbladder
(SRR6108749)

### Slide 266
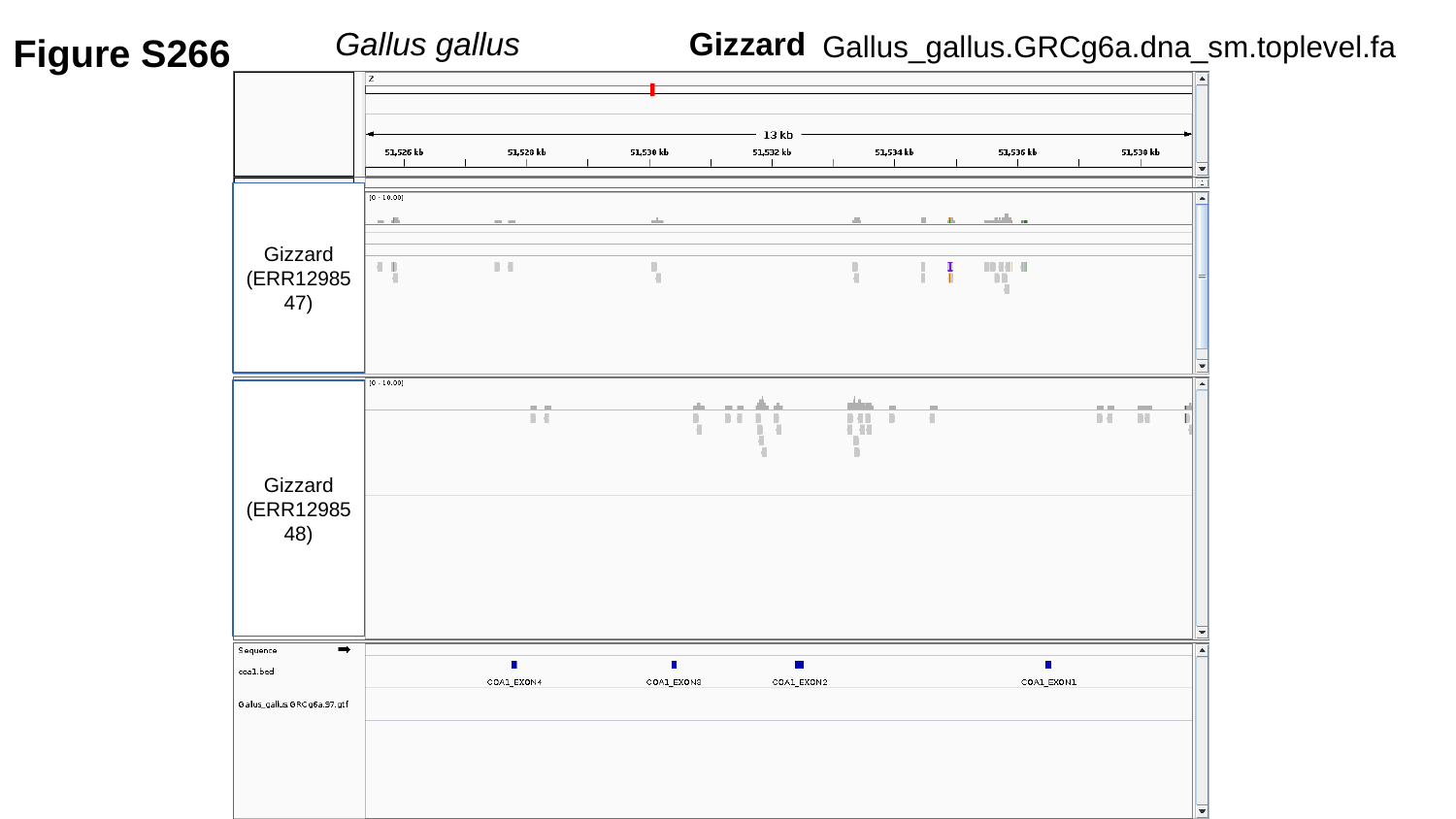

Figure S266
Gallus gallus	 Gizzard
Gallus_gallus.GRCg6a.dna_sm.toplevel.fa
Gizzard
(ERR1298547)
Gizzard
(ERR1298548)

### Slide 267
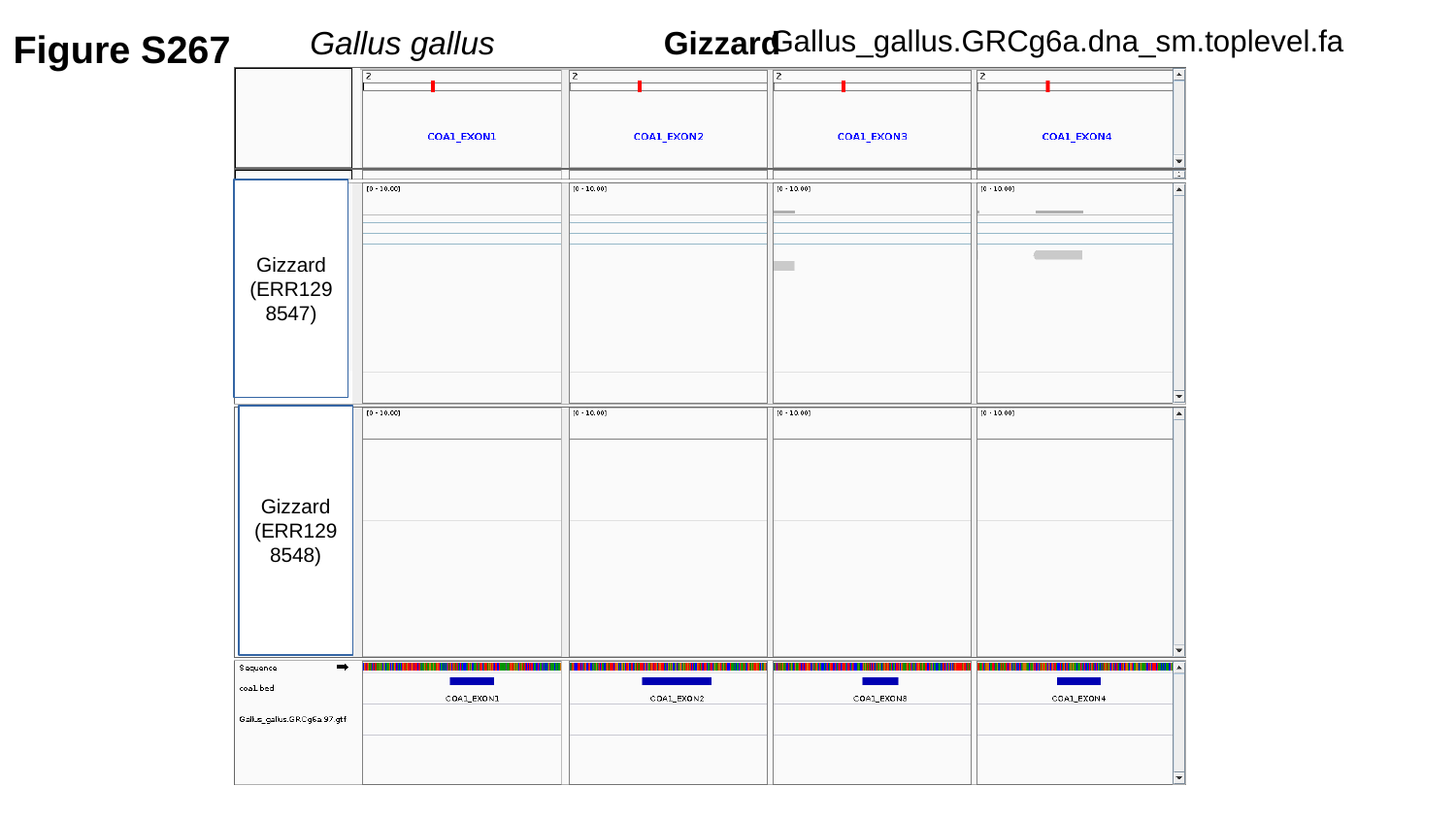

Figure S267
Gallus_gallus.GRCg6a.dna_sm.toplevel.fa
Gallus gallus	 Gizzard
Gizzard
(ERR1298547)
Gizzard
(ERR1298548)

### Slide 268
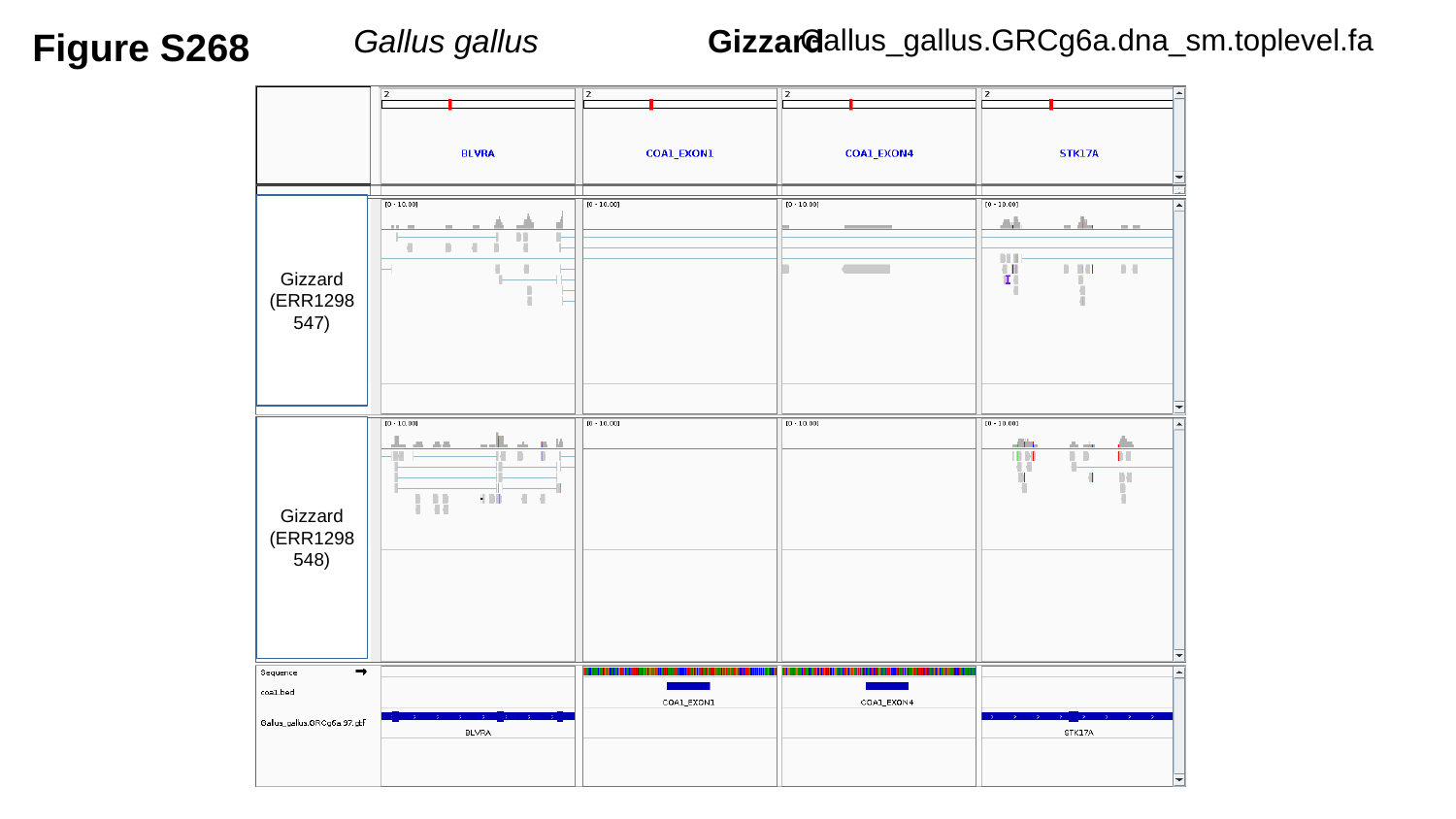

Figure S268
Gallus_gallus.GRCg6a.dna_sm.toplevel.fa
Gallus gallus	 Gizzard
Gizzard
(ERR1298547)
Gizzard
(ERR1298548)

### Slide 269
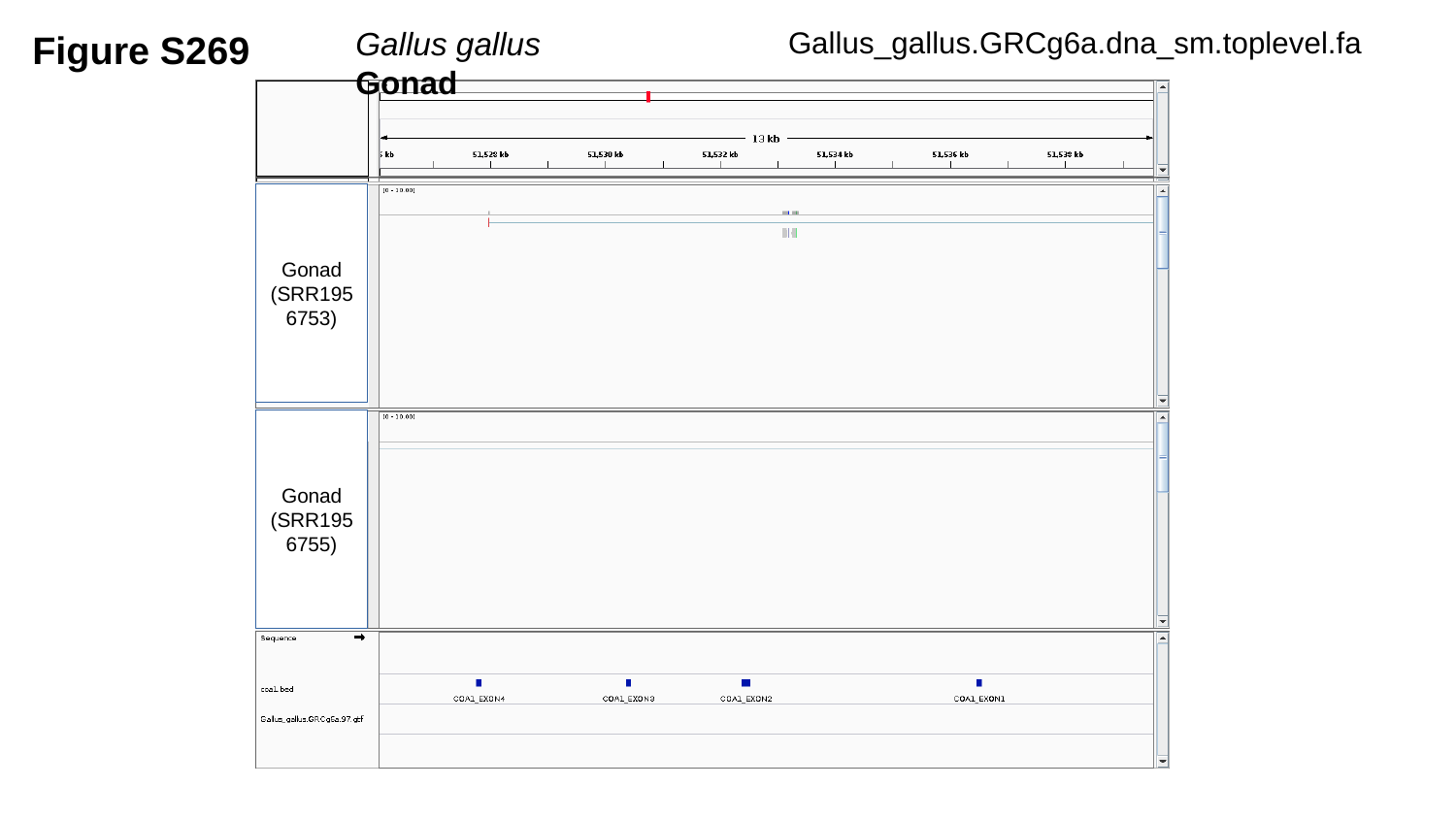

Figure S269
Gallus_gallus.GRCg6a.dna_sm.toplevel.fa
Gallus gallus	 Gonad
Gonad
(SRR1956753)
Gonad
(SRR1956755)

### Slide 270
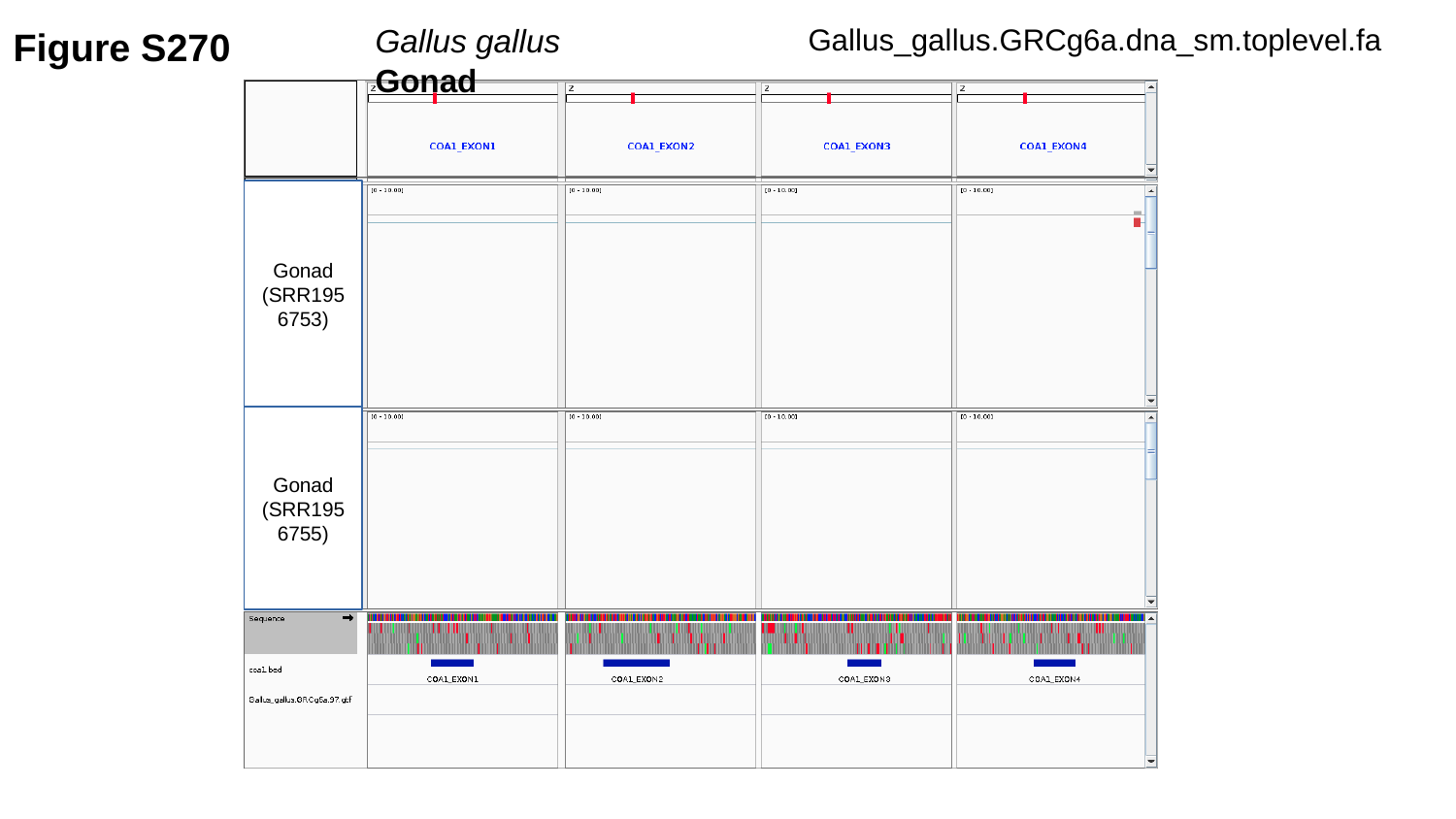

Figure S270
Gallus_gallus.GRCg6a.dna_sm.toplevel.fa
Gallus gallus	 Gonad
Gonad
(SRR1956753)
Gonad
(SRR1956755)

### Slide 271
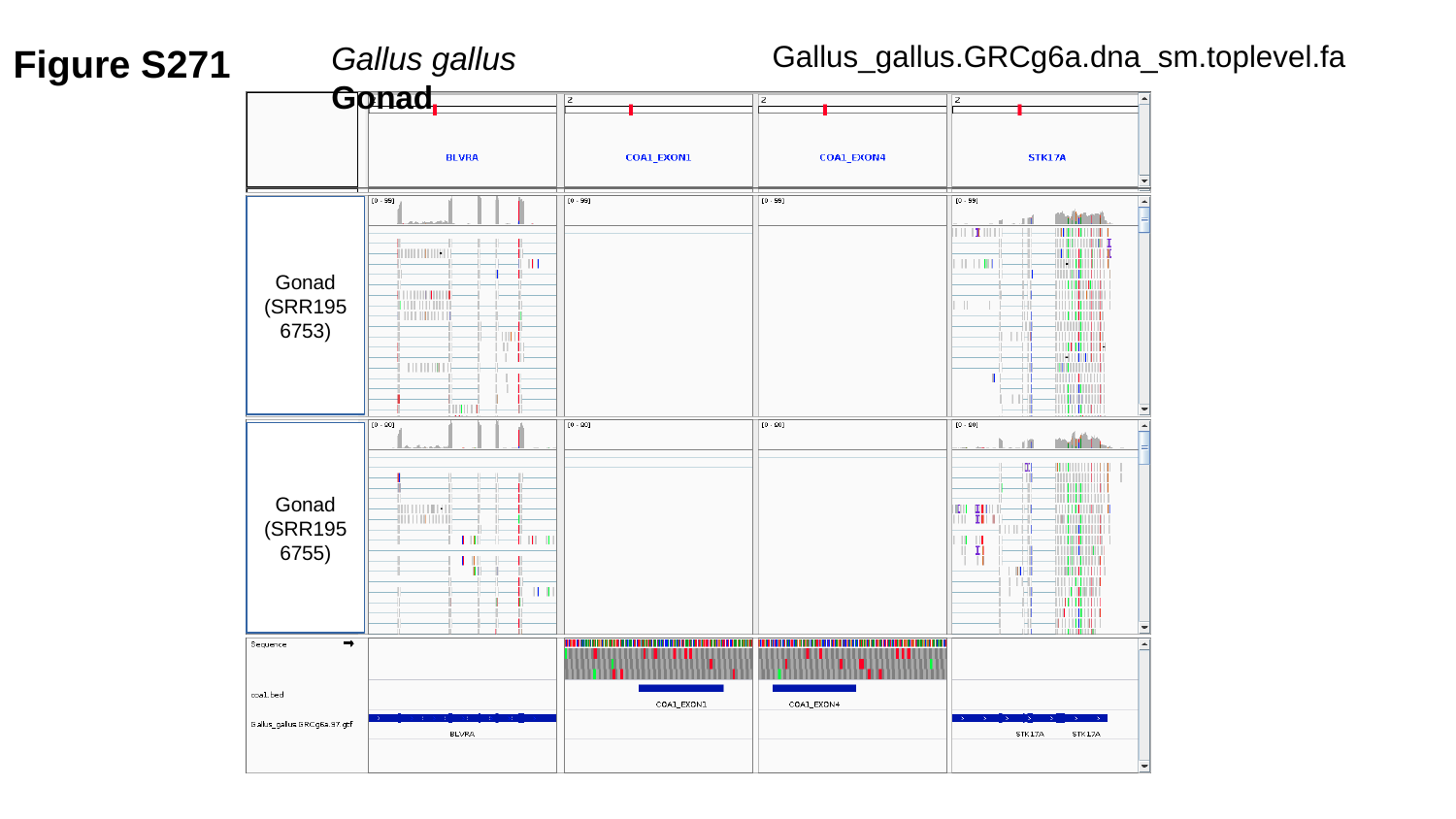

Figure S271
Gallus_gallus.GRCg6a.dna_sm.toplevel.fa
Gallus gallus	 Gonad
Gonad
(SRR1956753)
Gonad
(SRR1956755)

### Slide 272
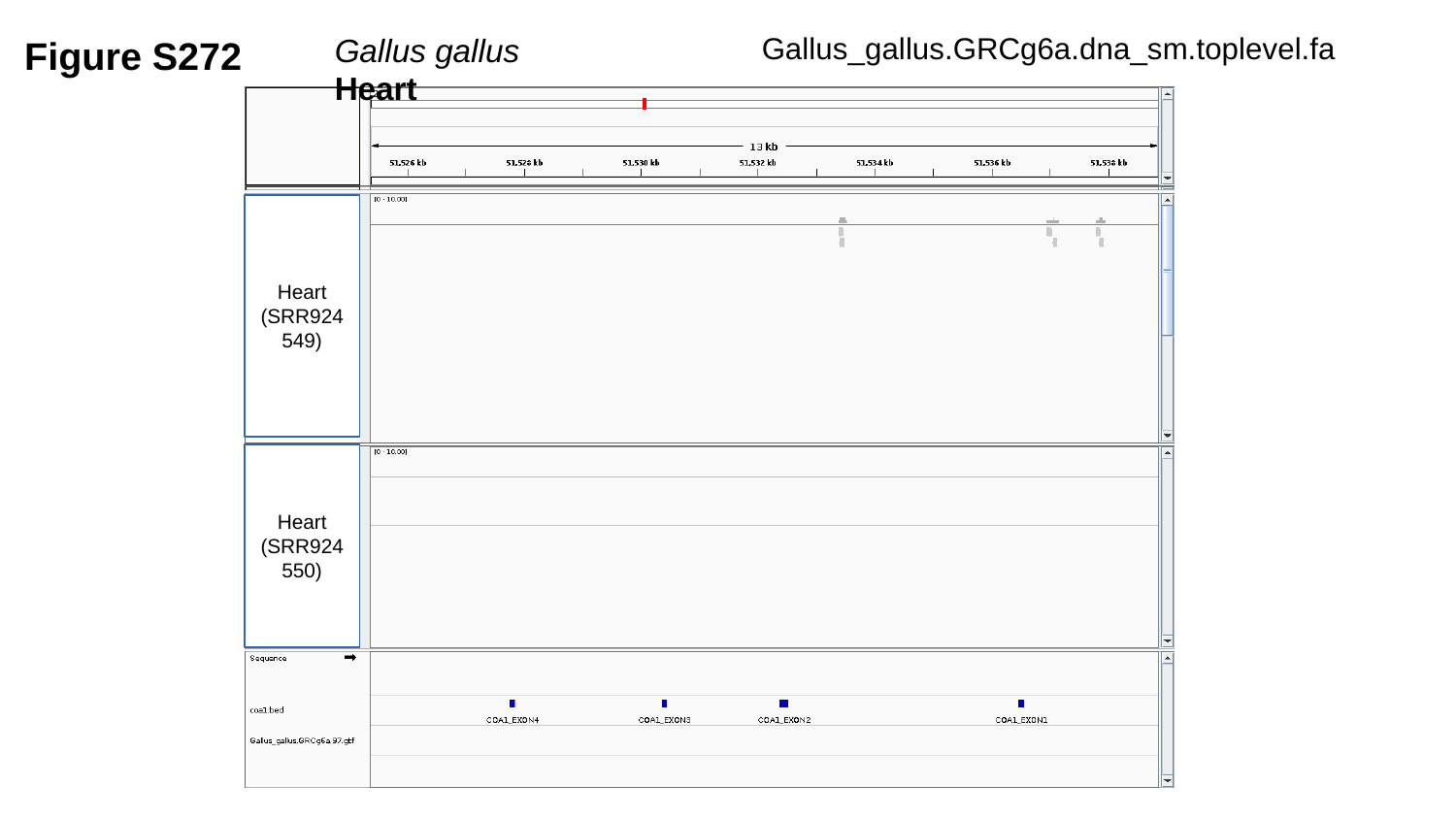

Figure S272
Gallus_gallus.GRCg6a.dna_sm.toplevel.fa
Gallus gallus	 Heart
Heart
(SRR924549)
Heart
(SRR924550)

### Slide 273
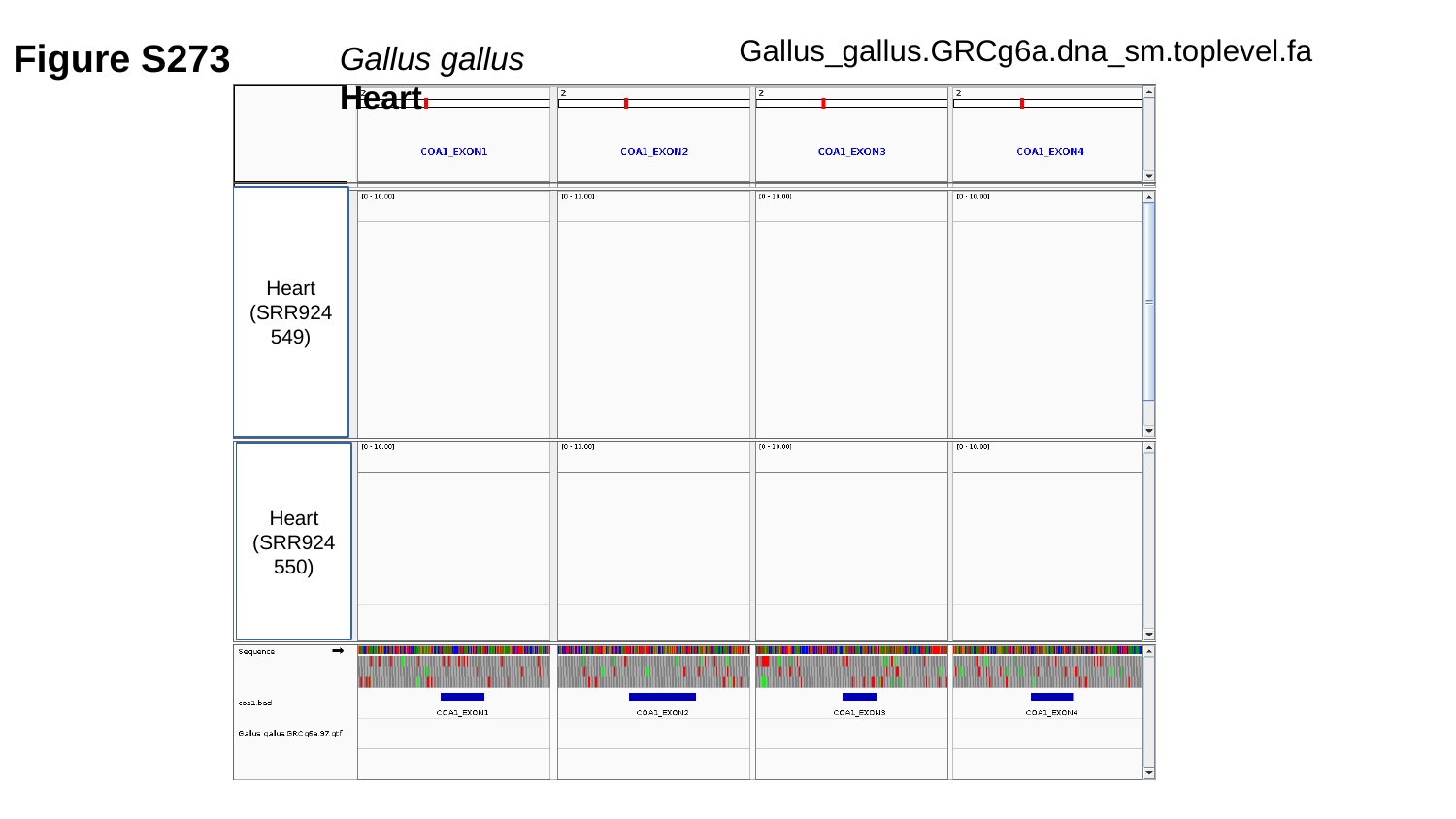

Figure S273
Gallus_gallus.GRCg6a.dna_sm.toplevel.fa
Gallus gallus	 Heart
Heart
(SRR924549)
Heart
(SRR924550)

### Slide 274
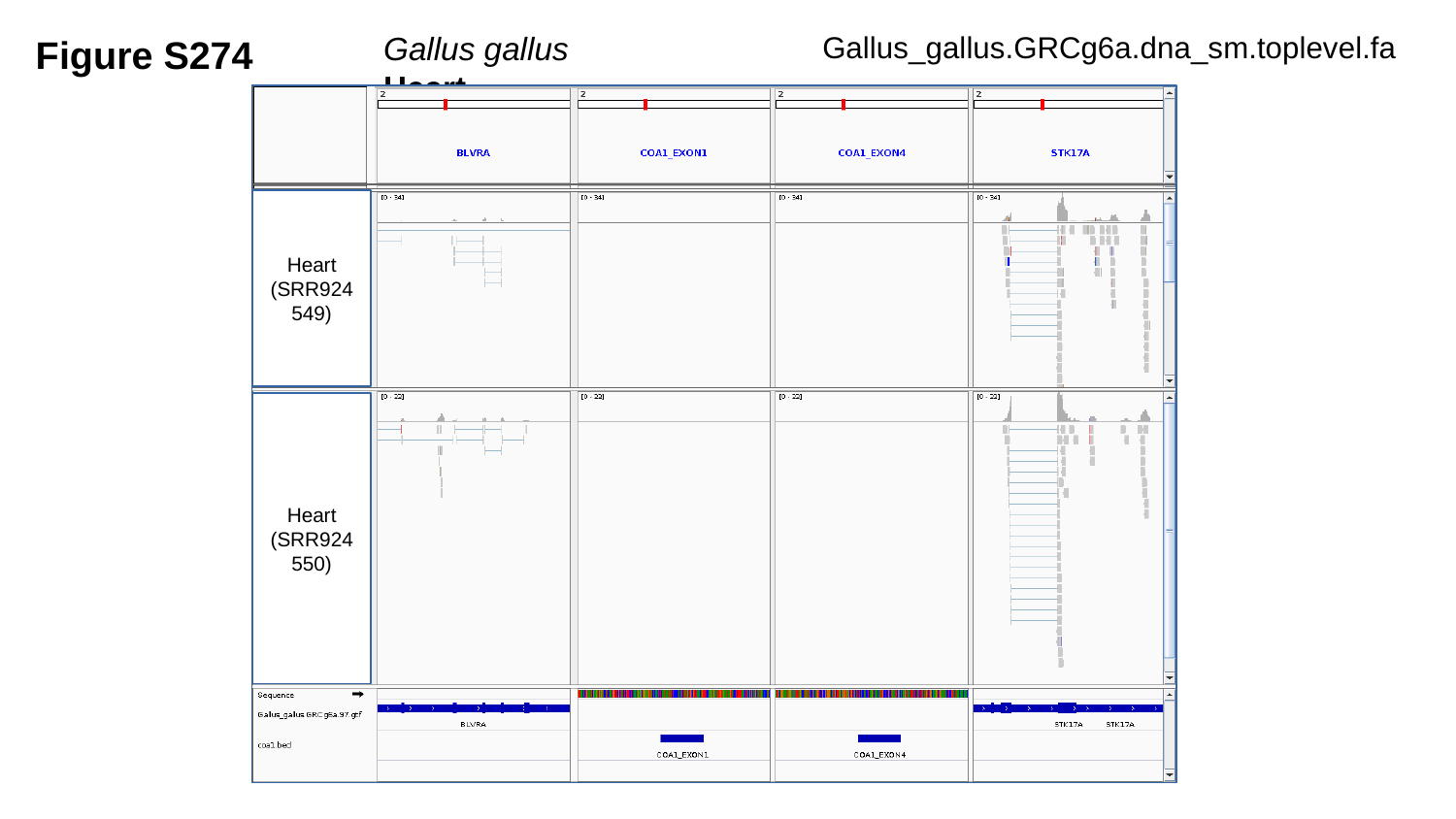

Figure S274
Gallus_gallus.GRCg6a.dna_sm.toplevel.fa
Gallus gallus	 Heart
Heart
(SRR924549)
Heart
(SRR924550)

### Slide 275
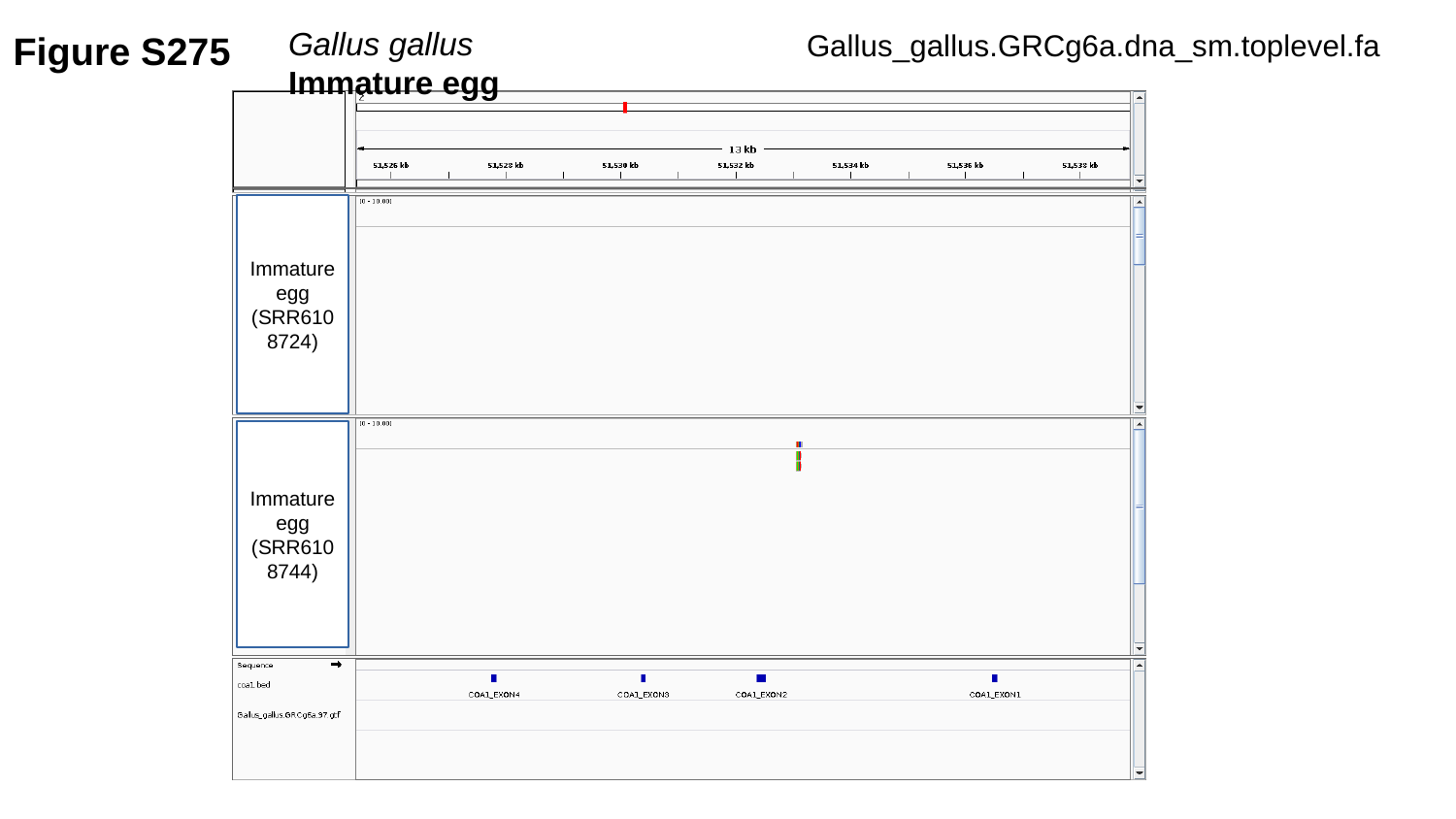

Figure S275
Gallus gallus	 Immature egg
Gallus_gallus.GRCg6a.dna_sm.toplevel.fa
Immature egg
(SRR6108724)
Immature egg
(SRR6108744)

### Slide 276
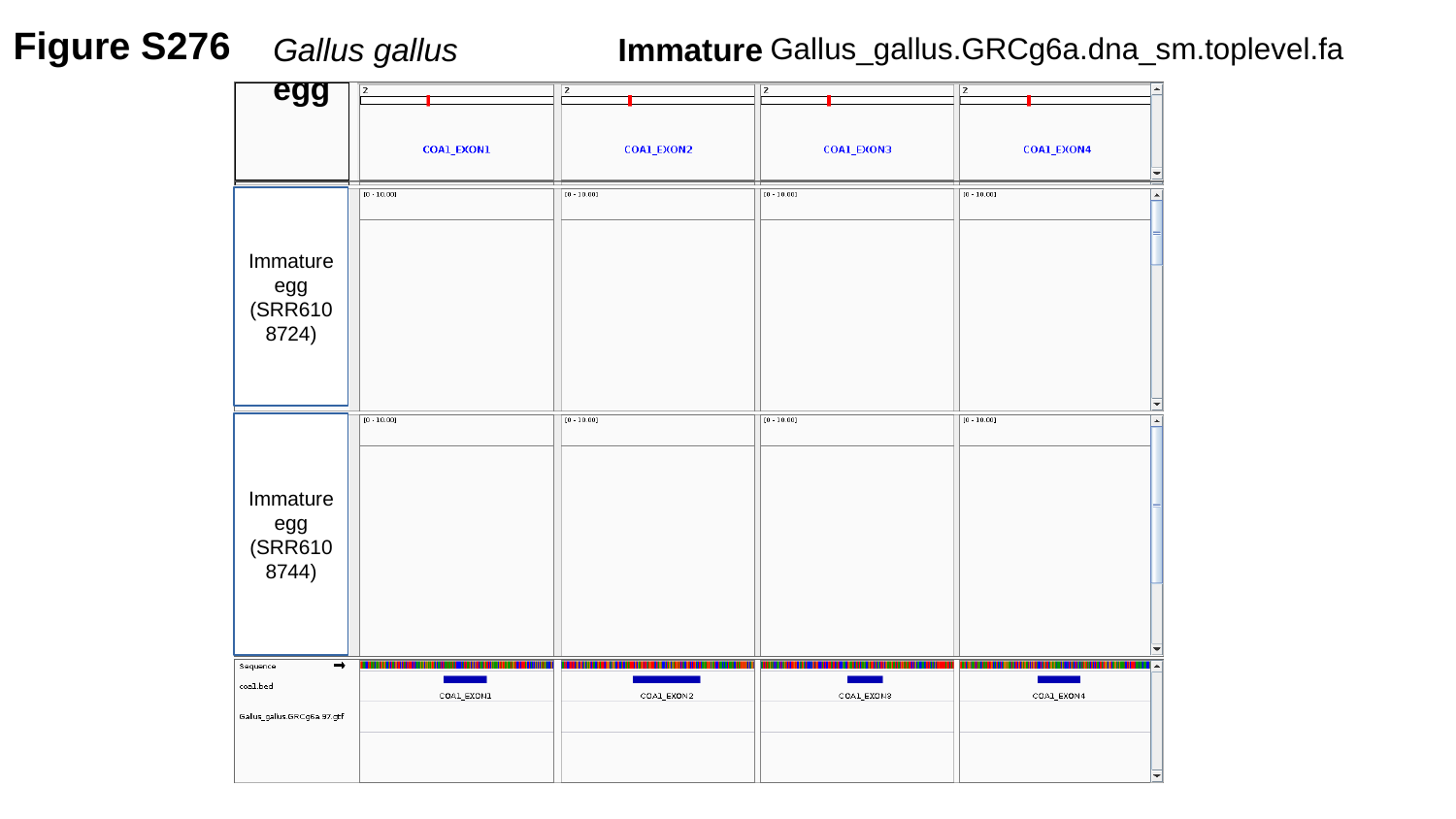

Figure S276
Gallus_gallus.GRCg6a.dna_sm.toplevel.fa
Gallus gallus	 Immature egg
Immature egg
(SRR6108724)
Immature egg
(SRR6108744)

### Slide 277
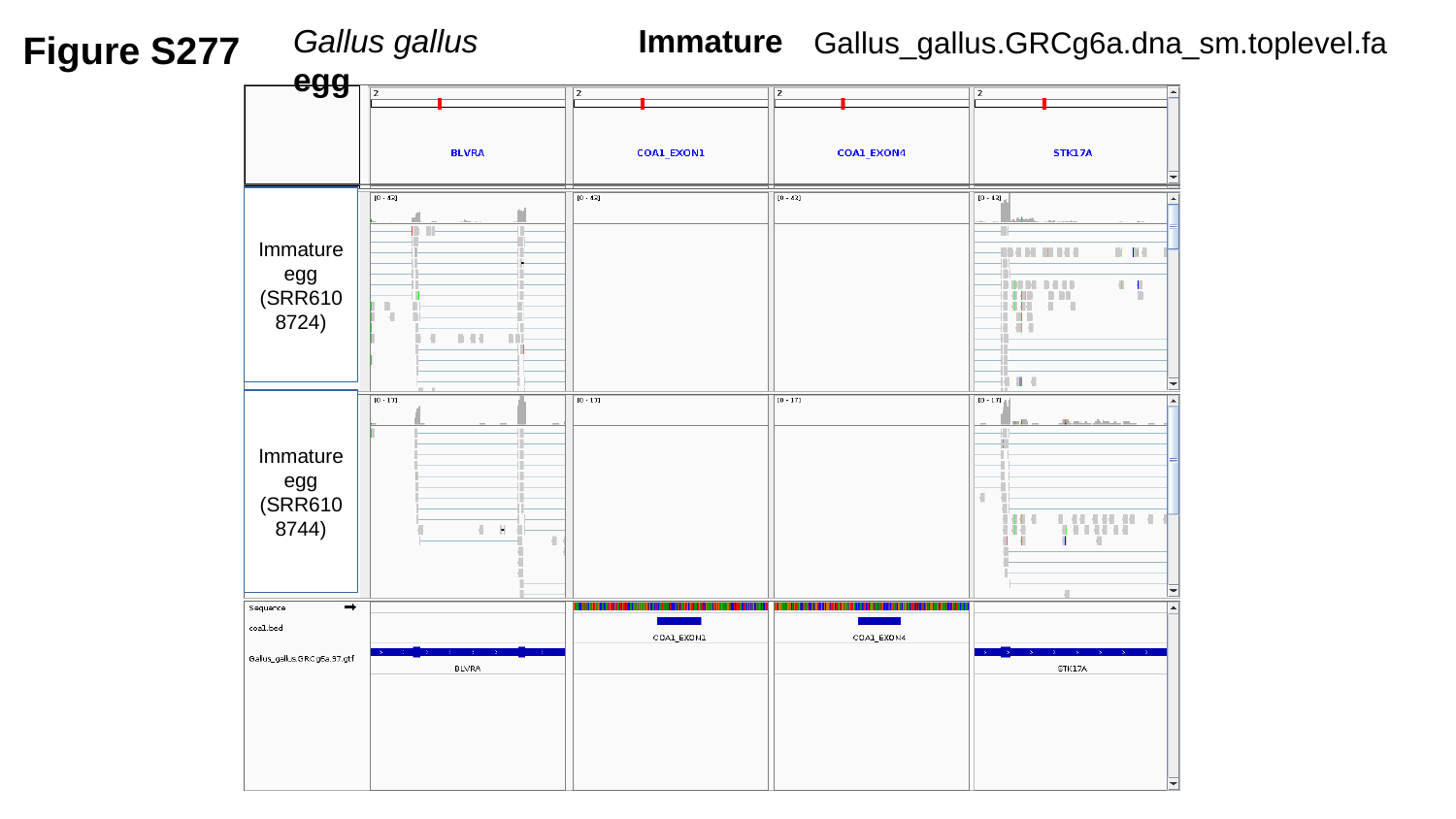

Figure S277
Gallus gallus	 Immature egg
Gallus_gallus.GRCg6a.dna_sm.toplevel.fa
Immature egg
(SRR6108724)
Immature egg
(SRR6108744)

### Slide 278
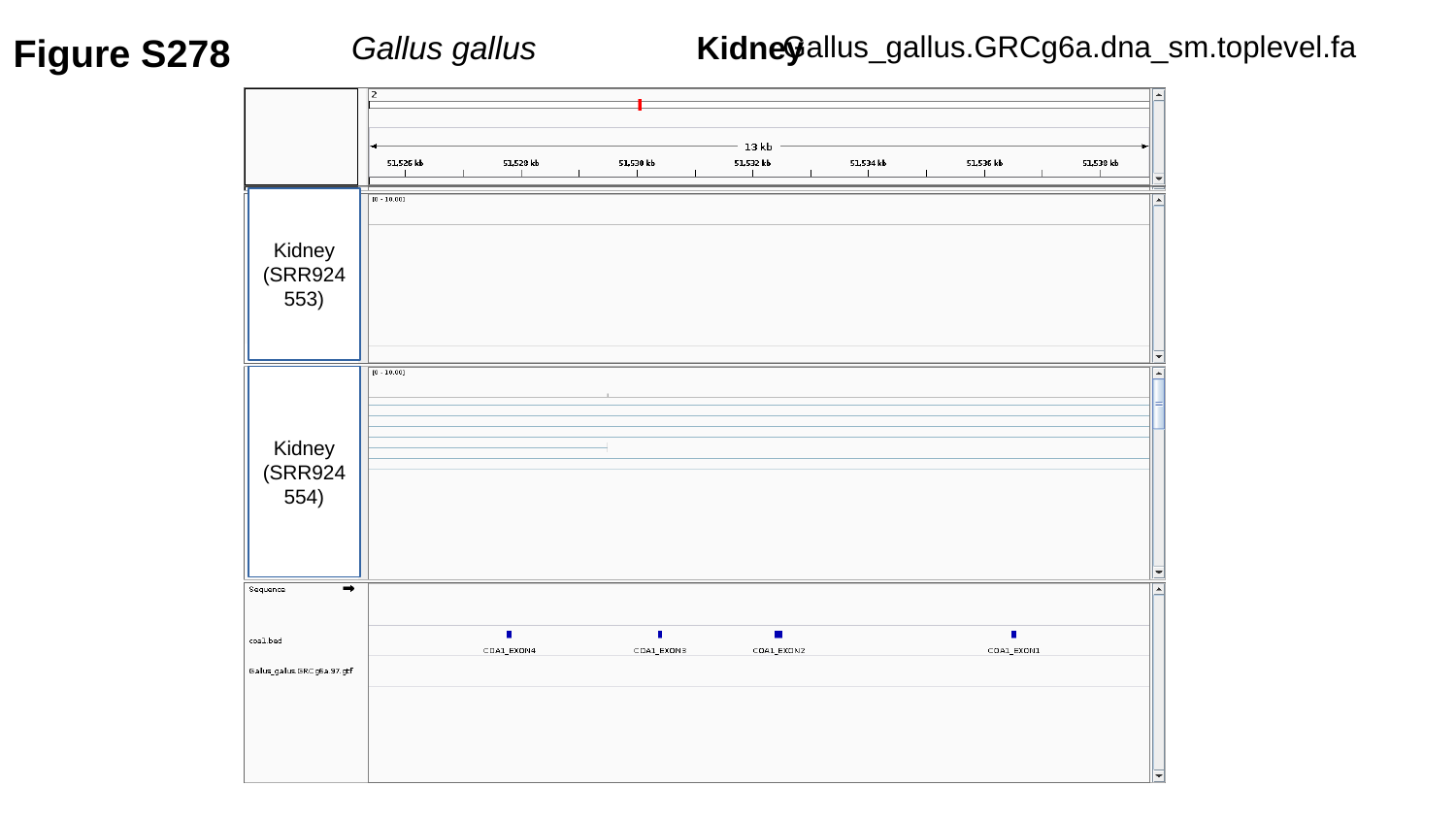

Figure S278
Gallus_gallus.GRCg6a.dna_sm.toplevel.fa
Gallus gallus	 Kidney
Kidney
(SRR924553)
Kidney
(SRR924554)

### Slide 279
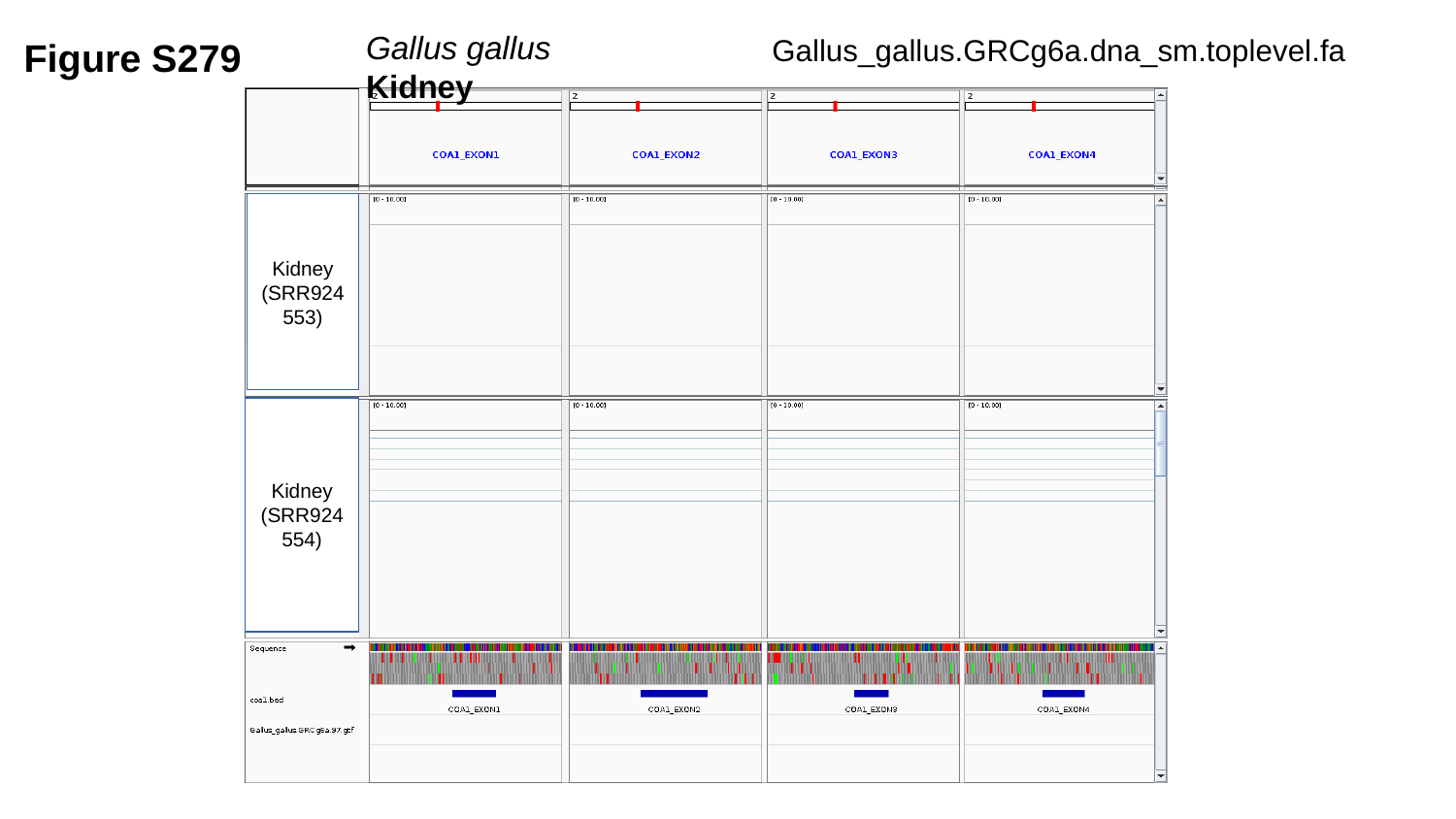

Figure S279
Gallus gallus	 Kidney
Gallus_gallus.GRCg6a.dna_sm.toplevel.fa
Kidney
(SRR924553)
Kidney
(SRR924554)

### Slide 280
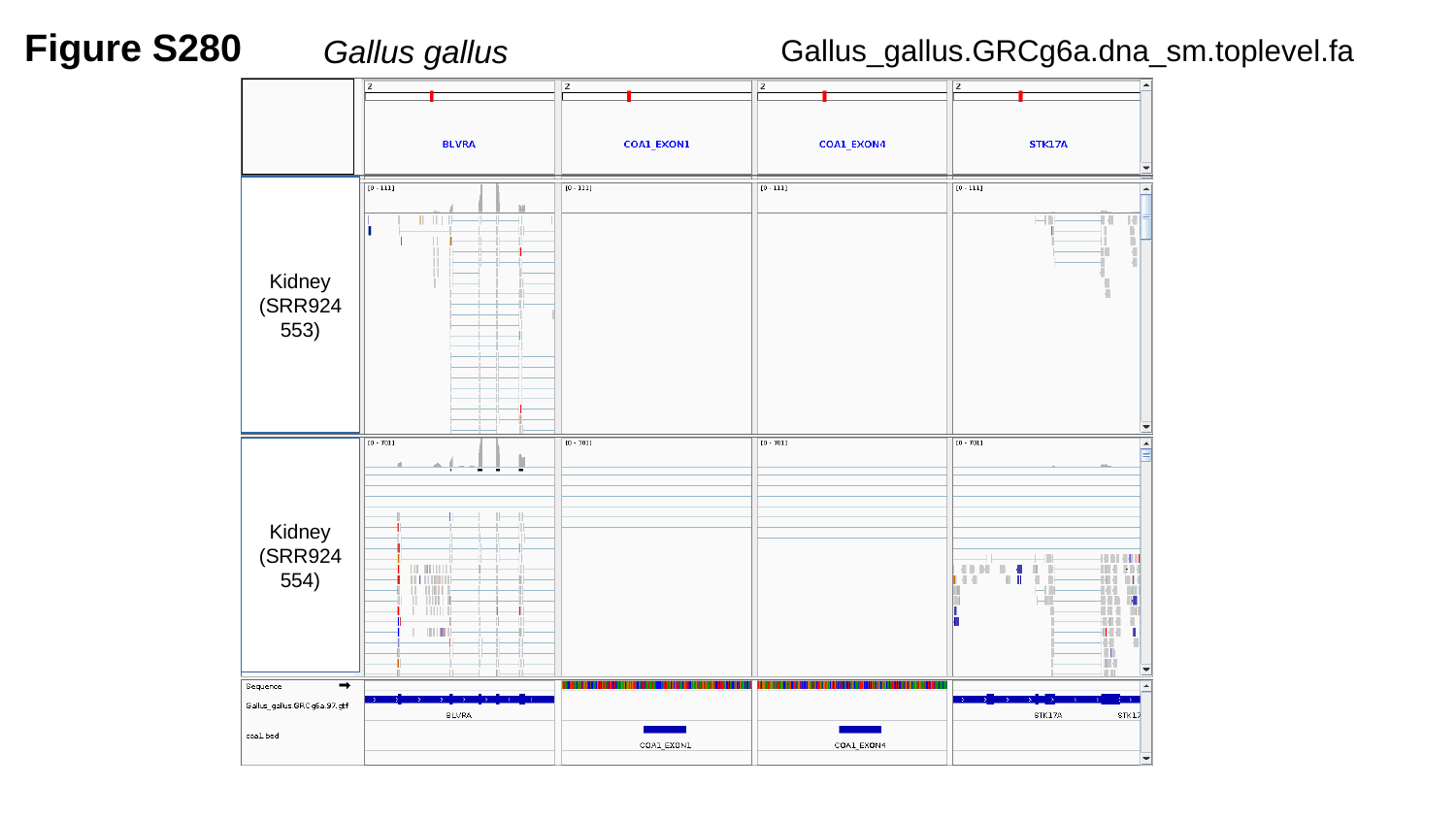

Figure S280
Gallus_gallus.GRCg6a.dna_sm.toplevel.fa
Gallus gallus	 Kidney
Kidney
(SRR924553)
Kidney
(SRR924554)

### Slide 281
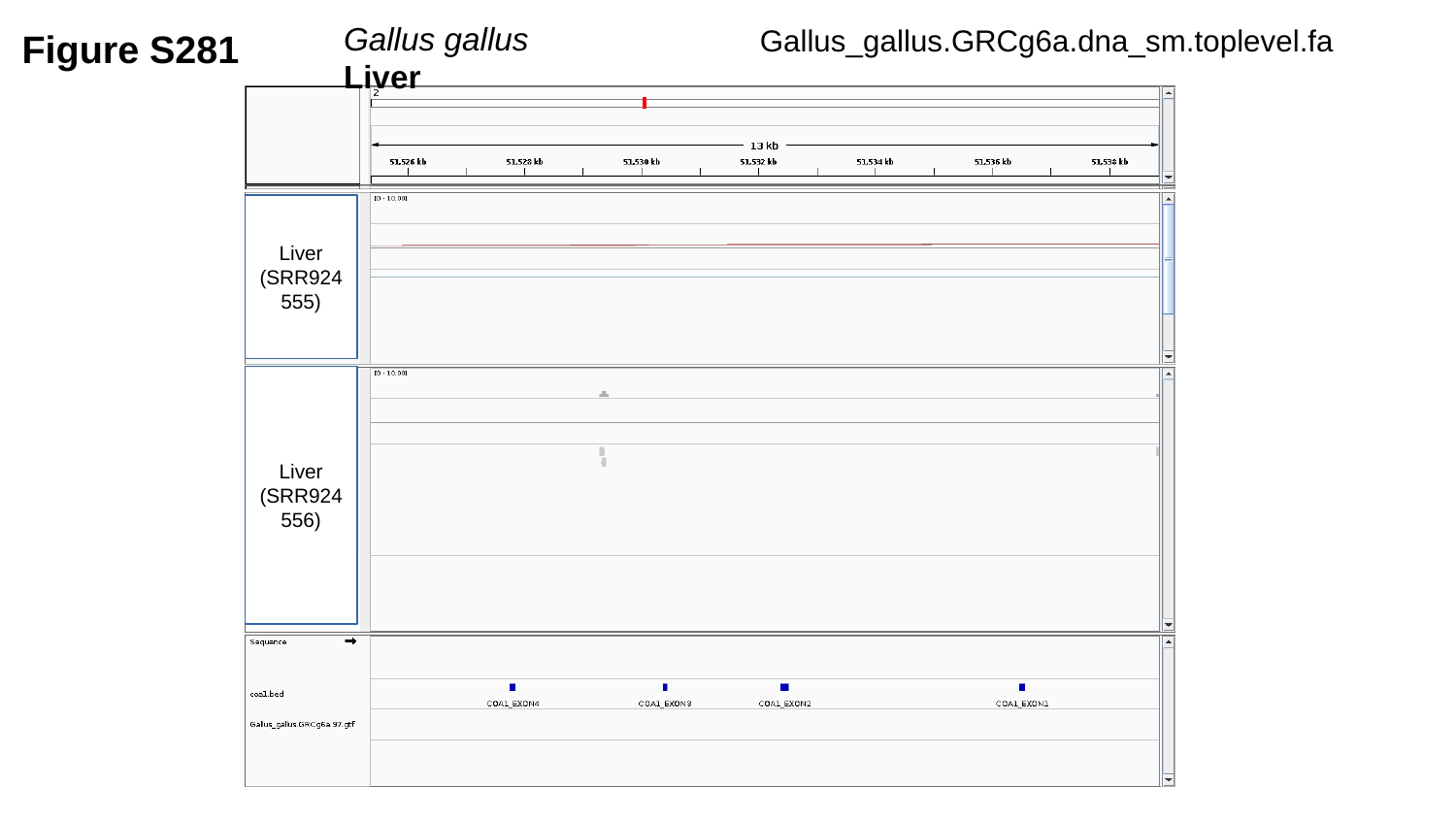

Figure S281
Gallus gallus	 Liver
Gallus_gallus.GRCg6a.dna_sm.toplevel.fa
Liver
(SRR924555)
Liver
(SRR924556)

### Slide 282
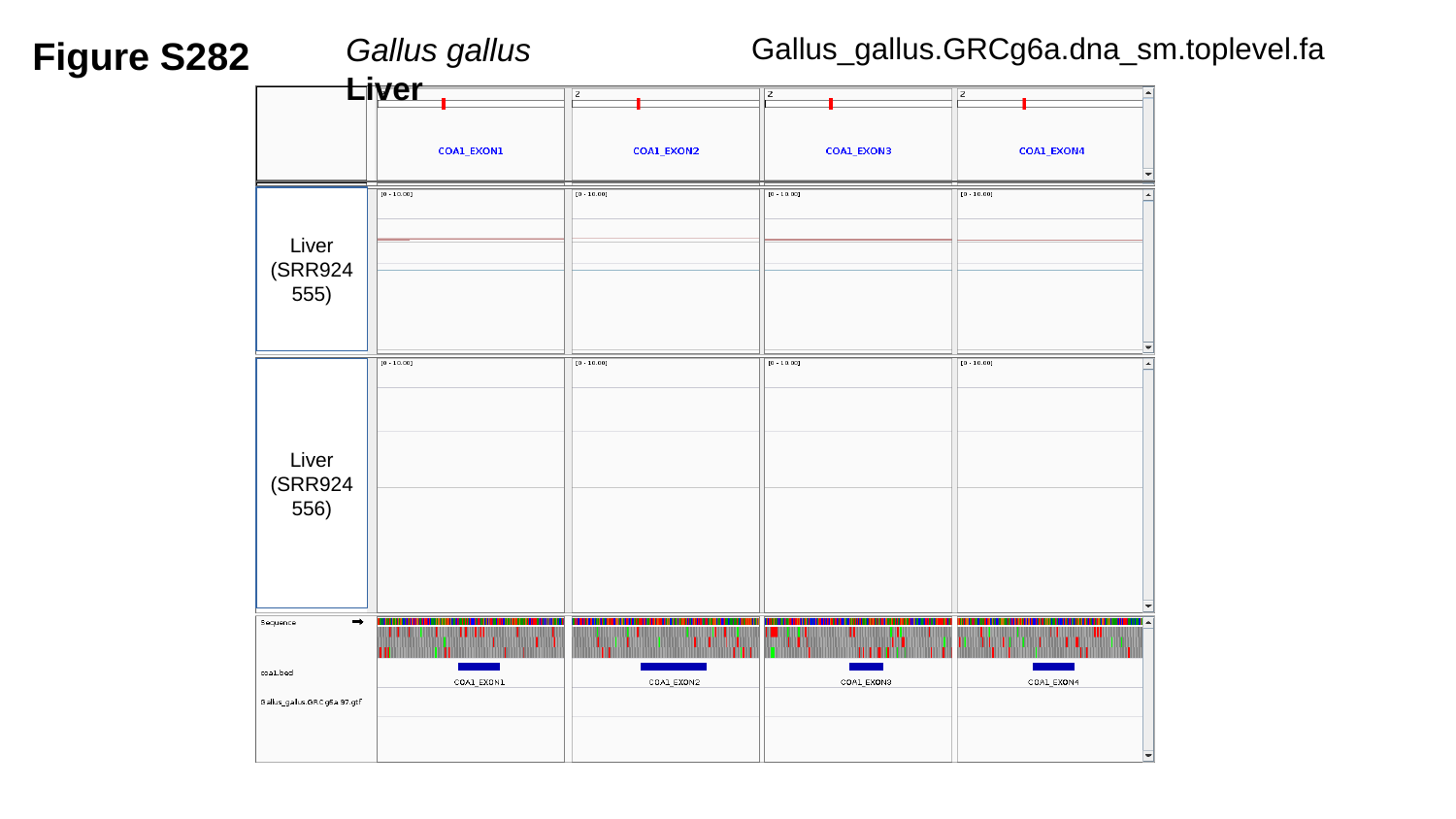

Figure S282
Gallus_gallus.GRCg6a.dna_sm.toplevel.fa
Gallus gallus	 Liver
Liver
(SRR924555)
Liver
(SRR924556)

### Slide 283
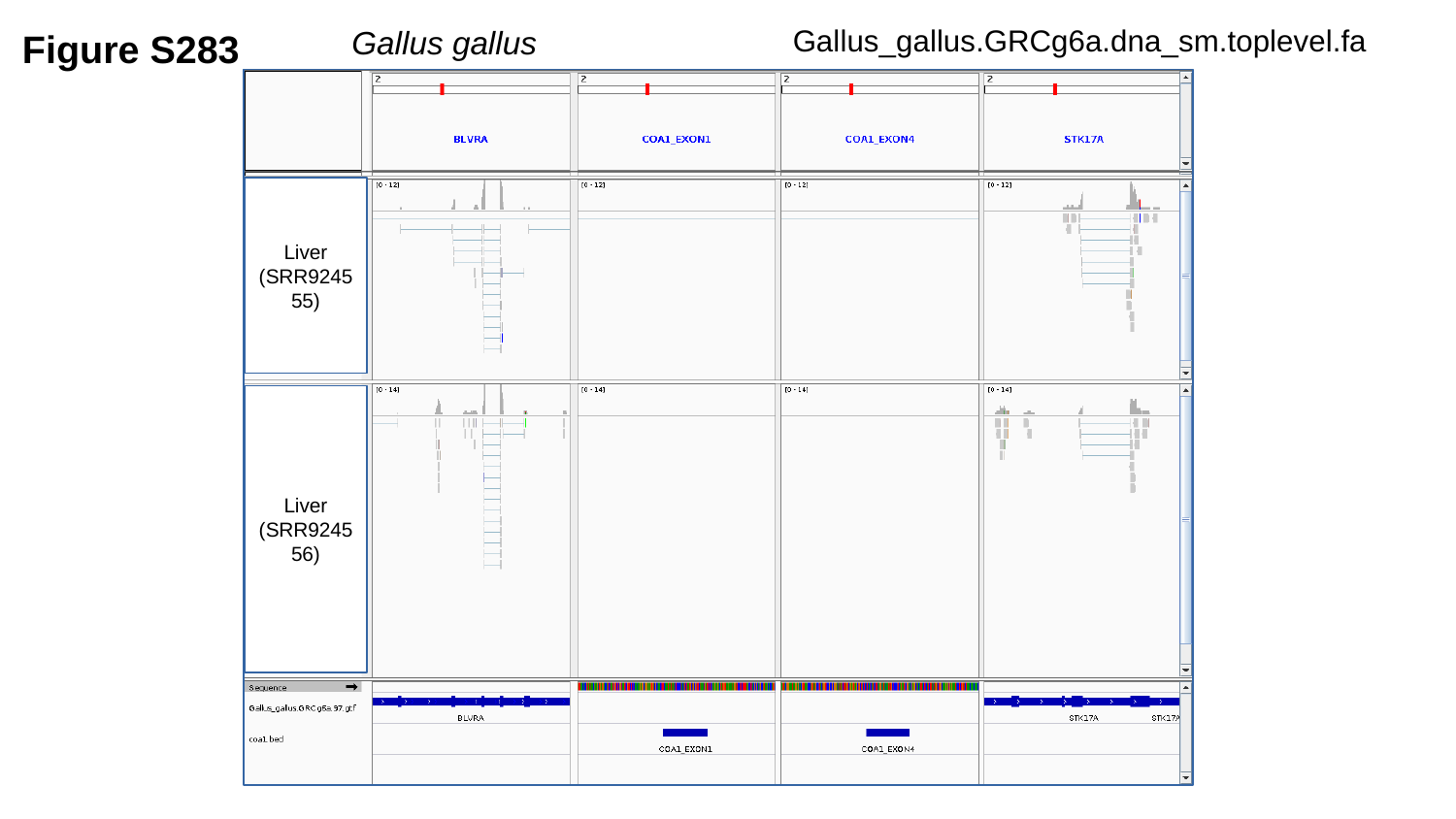

Figure S283
Gallus_gallus.GRCg6a.dna_sm.toplevel.fa
Gallus gallus	 Liver
Liver
(SRR924555)
Liver
(SRR924556)

### Slide 284
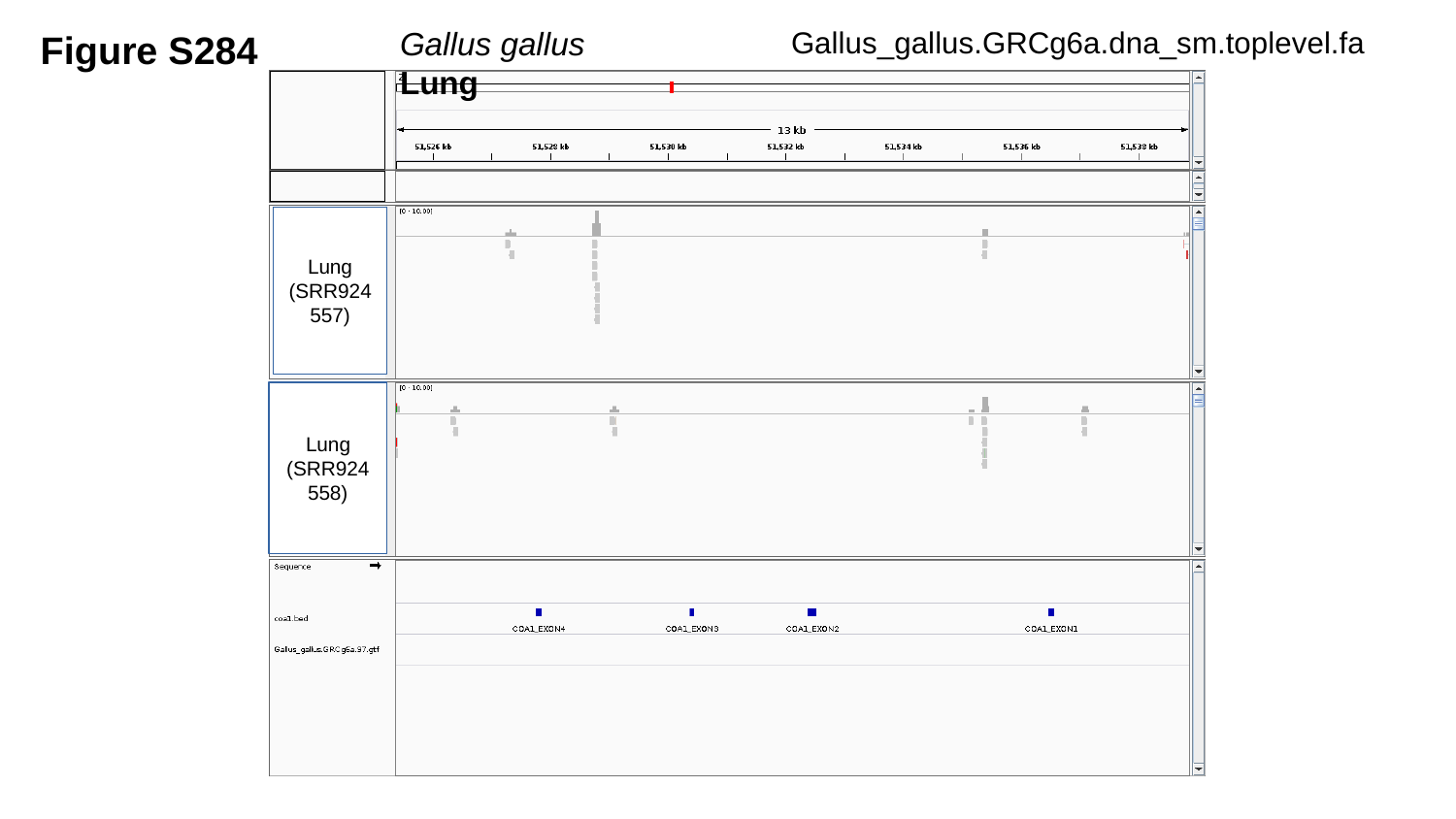

Figure S284
Gallus_gallus.GRCg6a.dna_sm.toplevel.fa
Gallus gallus	 Lung
Lung
(SRR924557)
Lung
(SRR924558)

### Slide 285
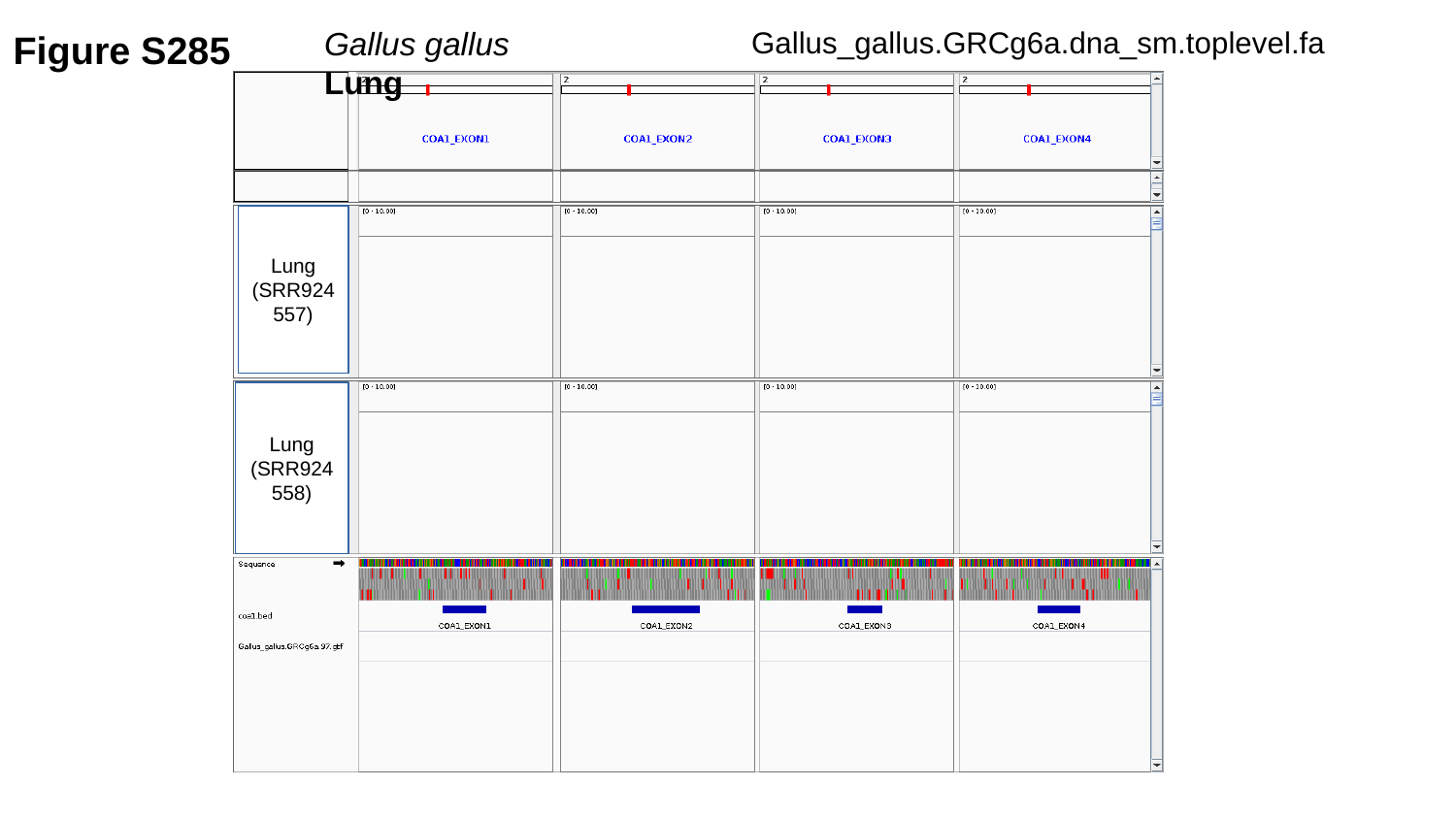

Figure S285
Gallus_gallus.GRCg6a.dna_sm.toplevel.fa
Gallus gallus	 Lung
Lung
(SRR924557)
Lung
(SRR924558)

### Slide 286
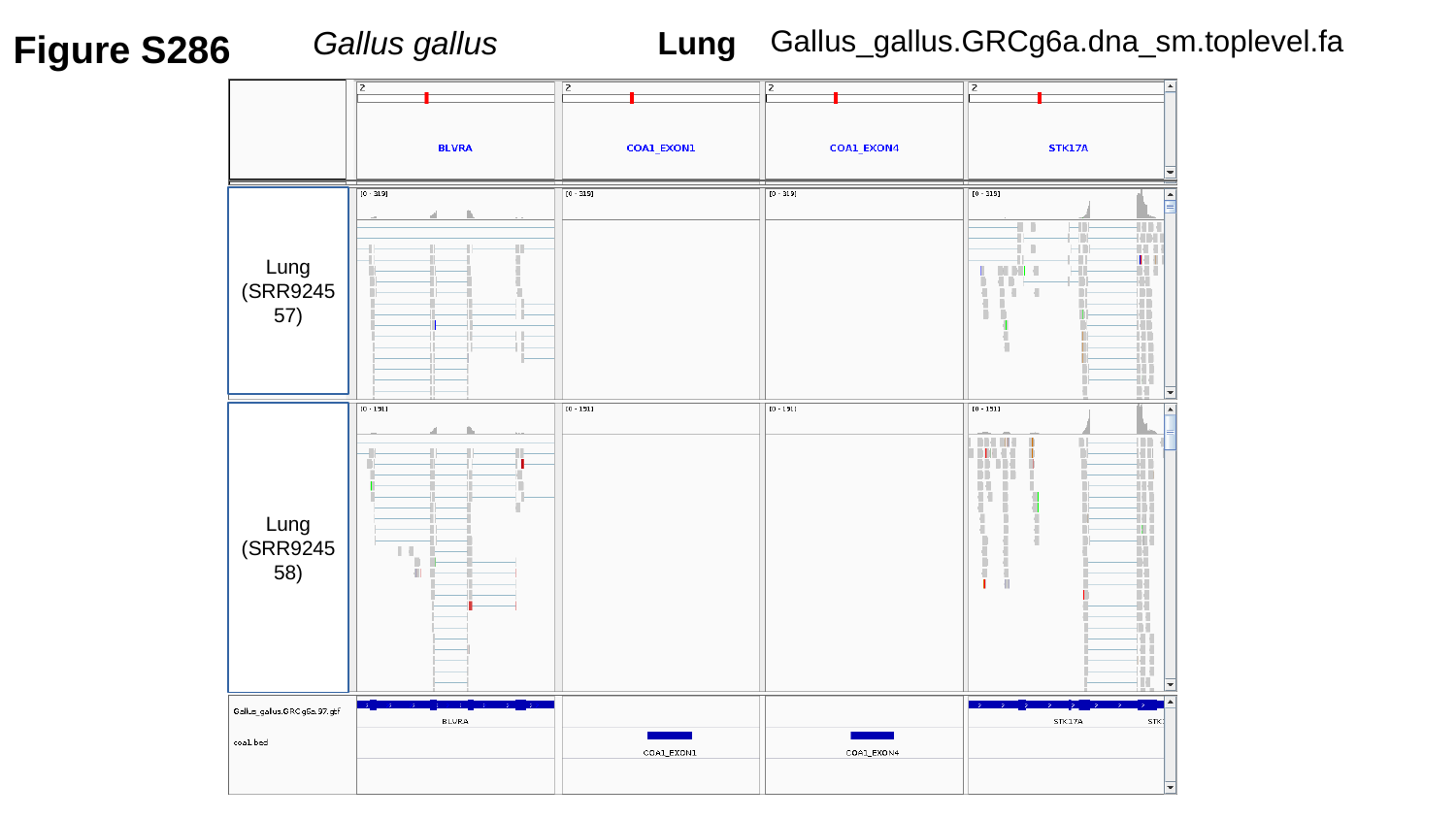

Figure S286
Gallus_gallus.GRCg6a.dna_sm.toplevel.fa
Gallus gallus	 Lung
Lung
(SRR924557)
Lung
(SRR924558)

### Slide 287
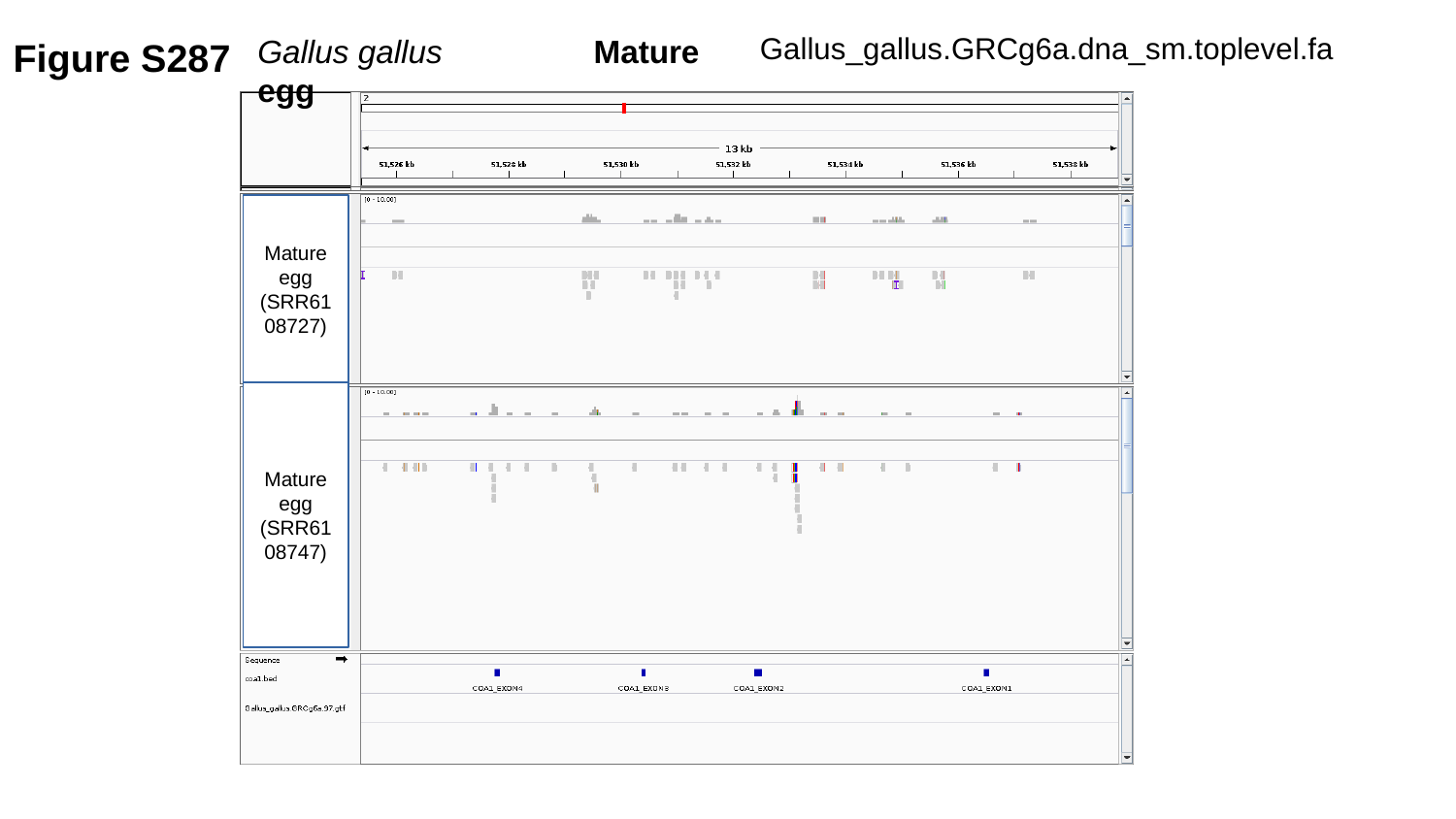

Figure S287
Gallus_gallus.GRCg6a.dna_sm.toplevel.fa
Gallus gallus	 Mature egg
Mature egg
(SRR6108727)
Mature egg
(SRR6108747)

### Slide 288
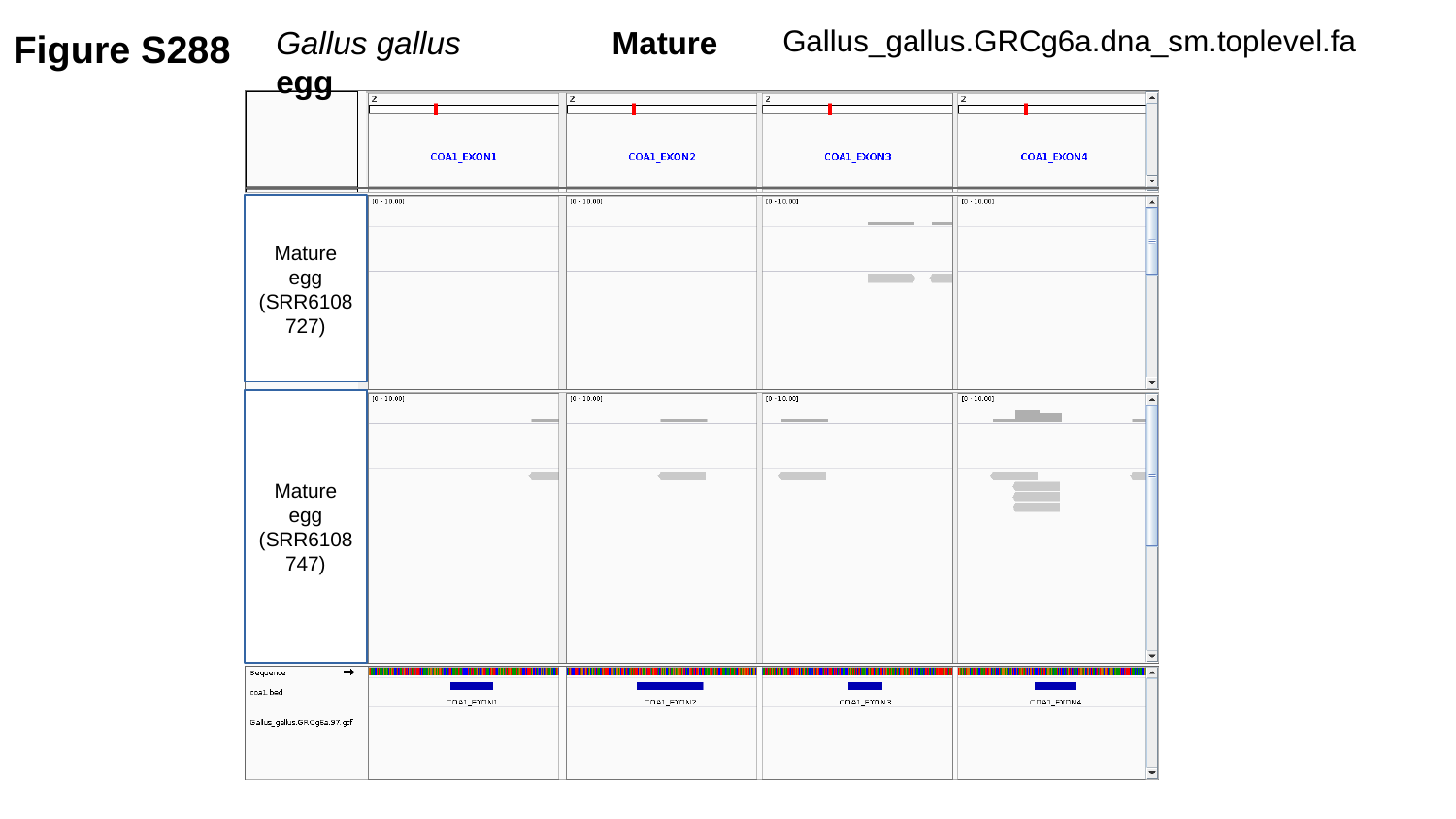

Figure S288
Gallus_gallus.GRCg6a.dna_sm.toplevel.fa
Gallus gallus	 Mature egg
Mature egg
(SRR6108727)
Mature egg
(SRR6108747)

### Slide 289
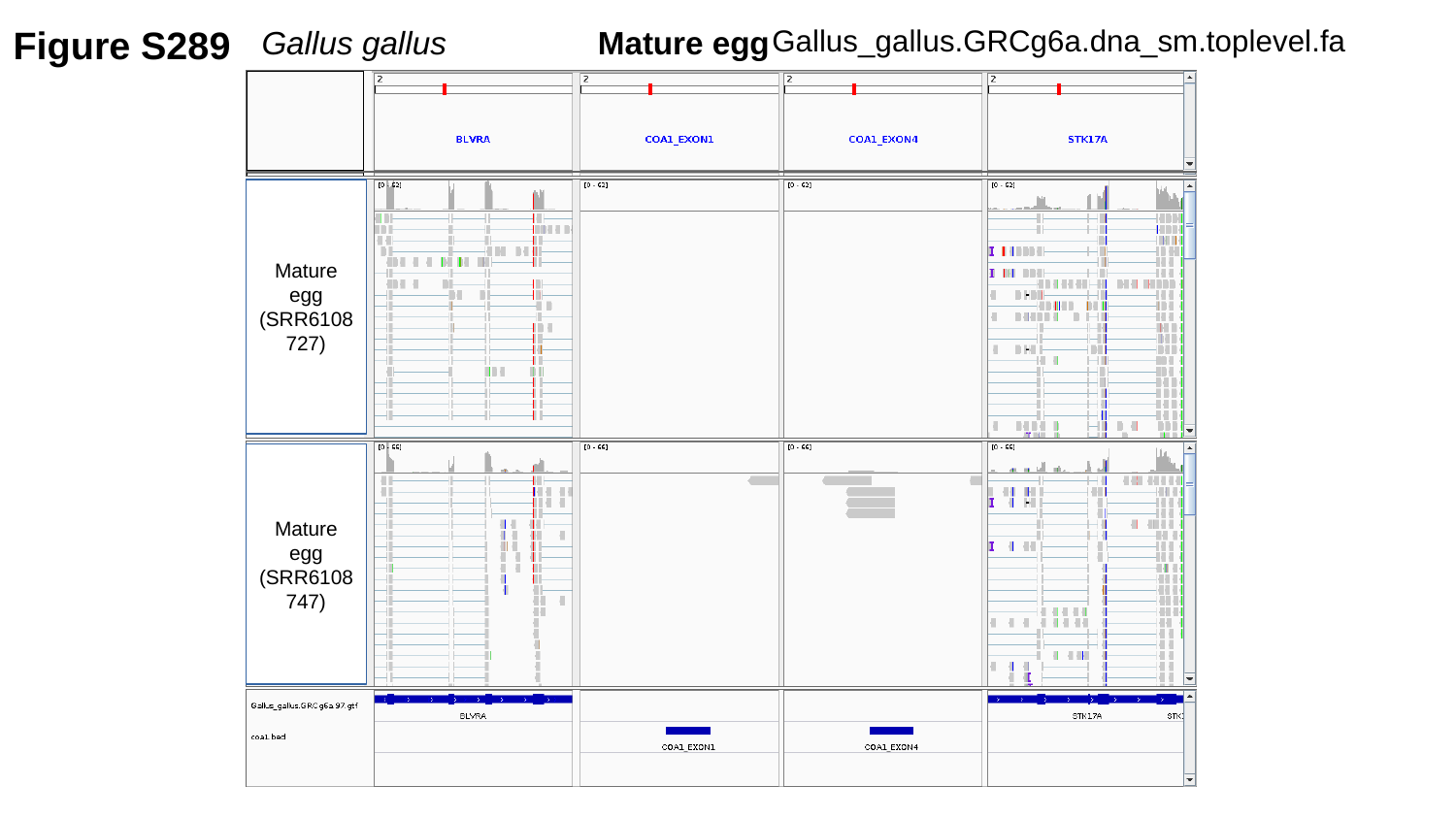

Figure S289
Gallus_gallus.GRCg6a.dna_sm.toplevel.fa
Gallus gallus	 Mature egg
Mature egg
(SRR6108727)
Mature egg
(SRR6108747)

### Slide 290
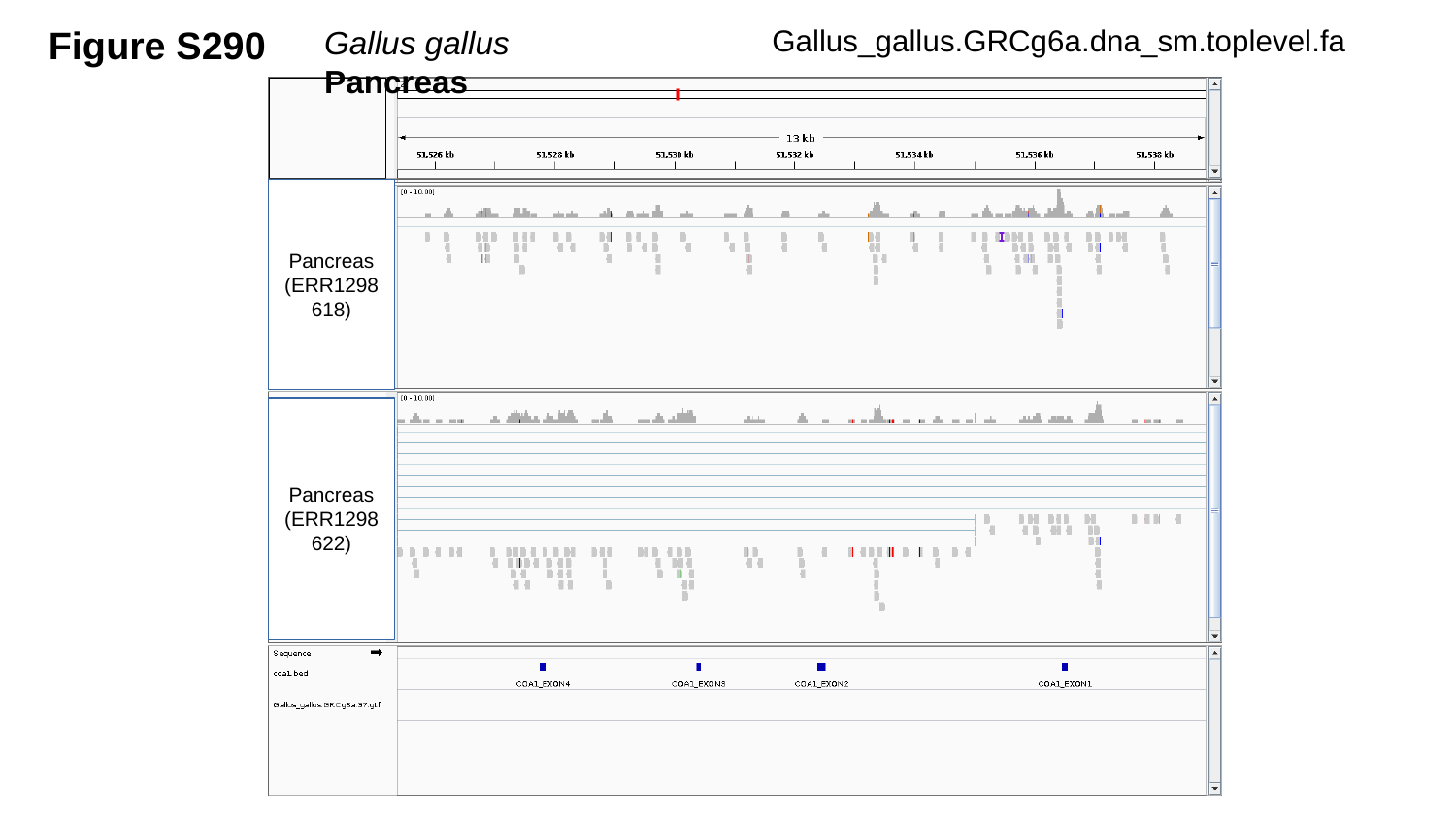

Figure S290
Gallus_gallus.GRCg6a.dna_sm.toplevel.fa
Gallus gallus	 Pancreas
Pancreas
(ERR1298618)
Pancreas
(ERR1298622)

### Slide 291
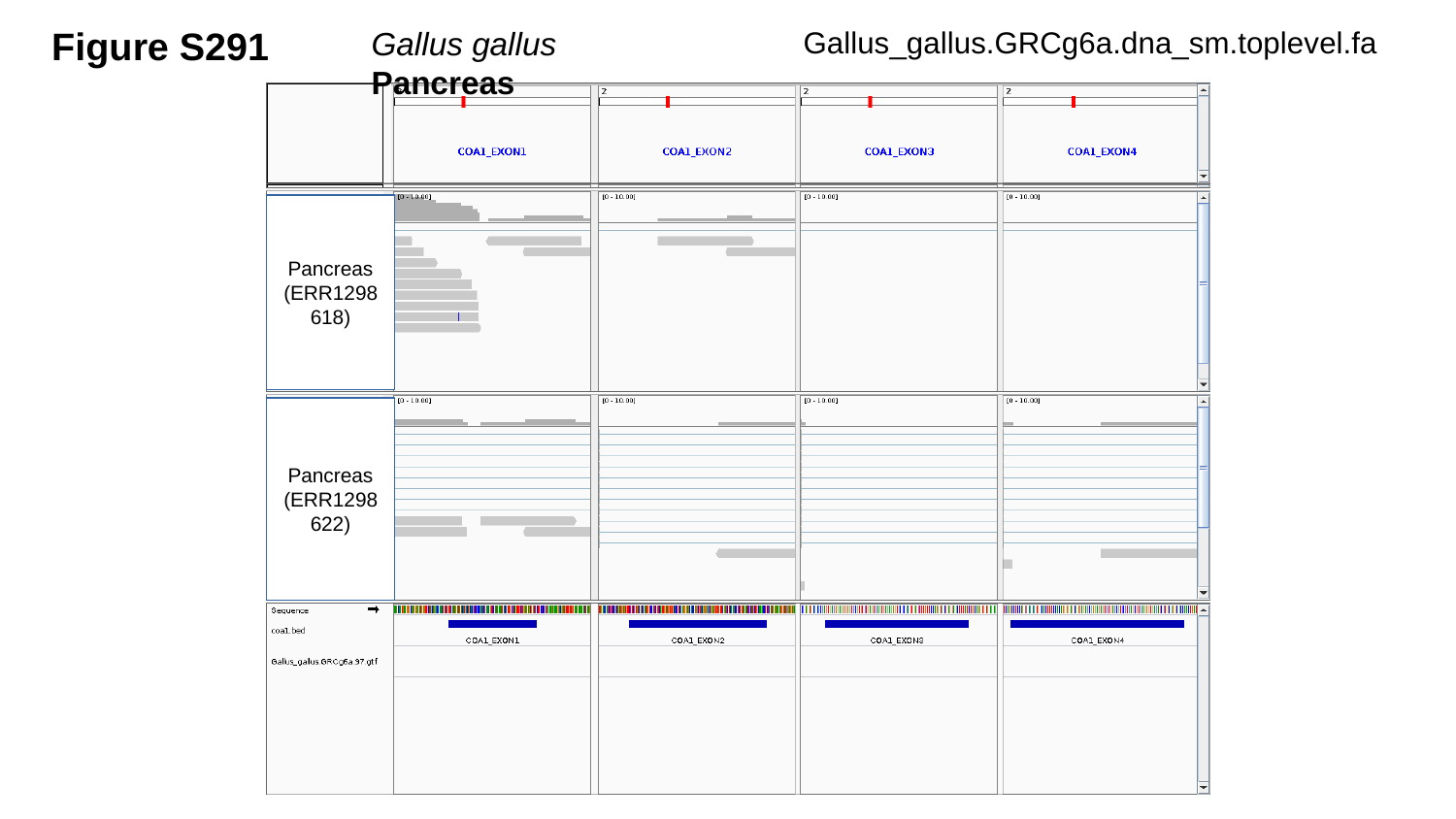

Figure S291
Gallus_gallus.GRCg6a.dna_sm.toplevel.fa
Gallus gallus	 Pancreas
Pancreas
(ERR1298618)
Pancreas
(ERR1298622)

### Slide 292
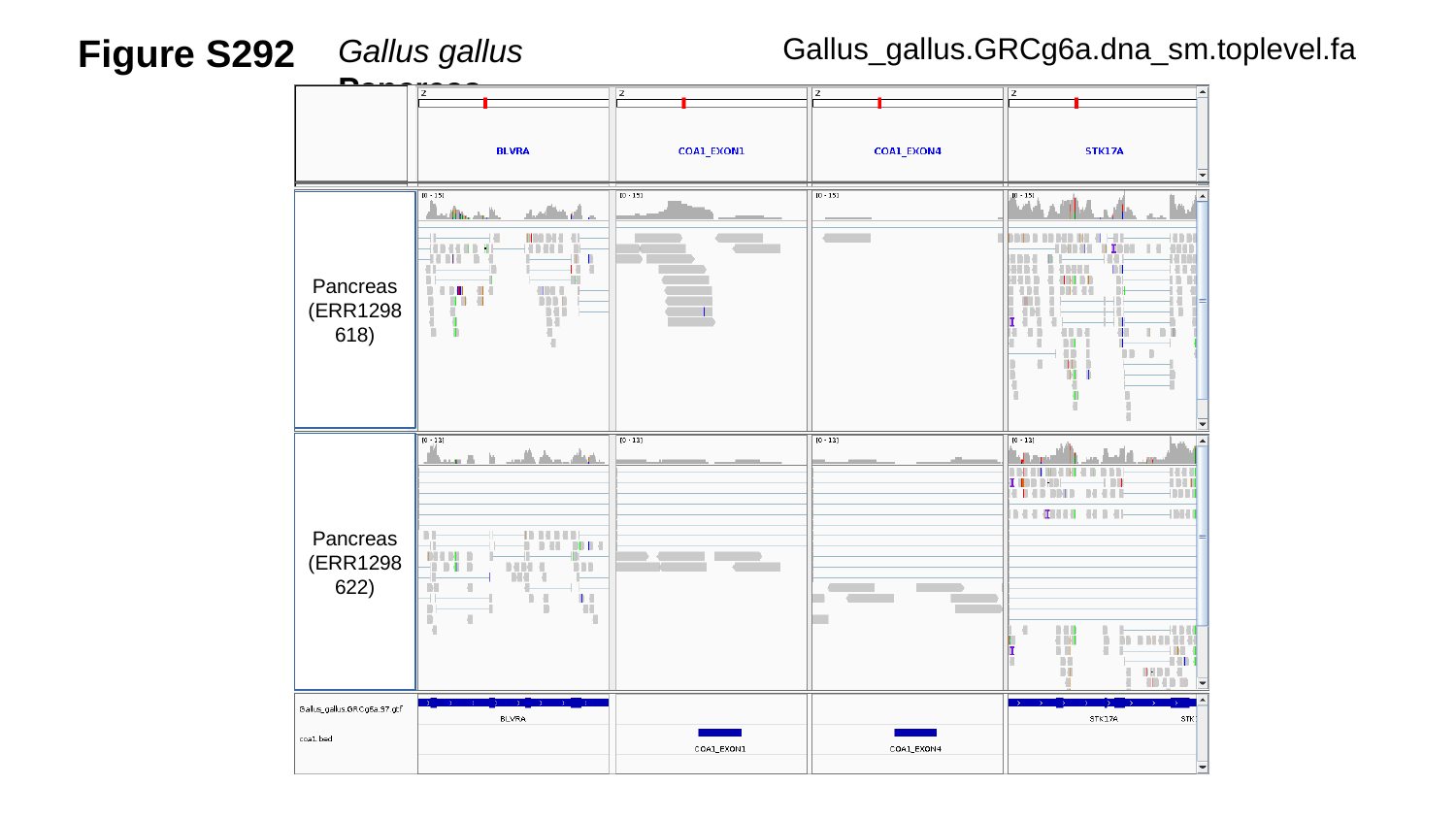

Figure S292
Gallus_gallus.GRCg6a.dna_sm.toplevel.fa
Gallus gallus	 Pancreas
Pancreas
(ERR1298618)
Pancreas
(ERR1298622)

### Slide 293
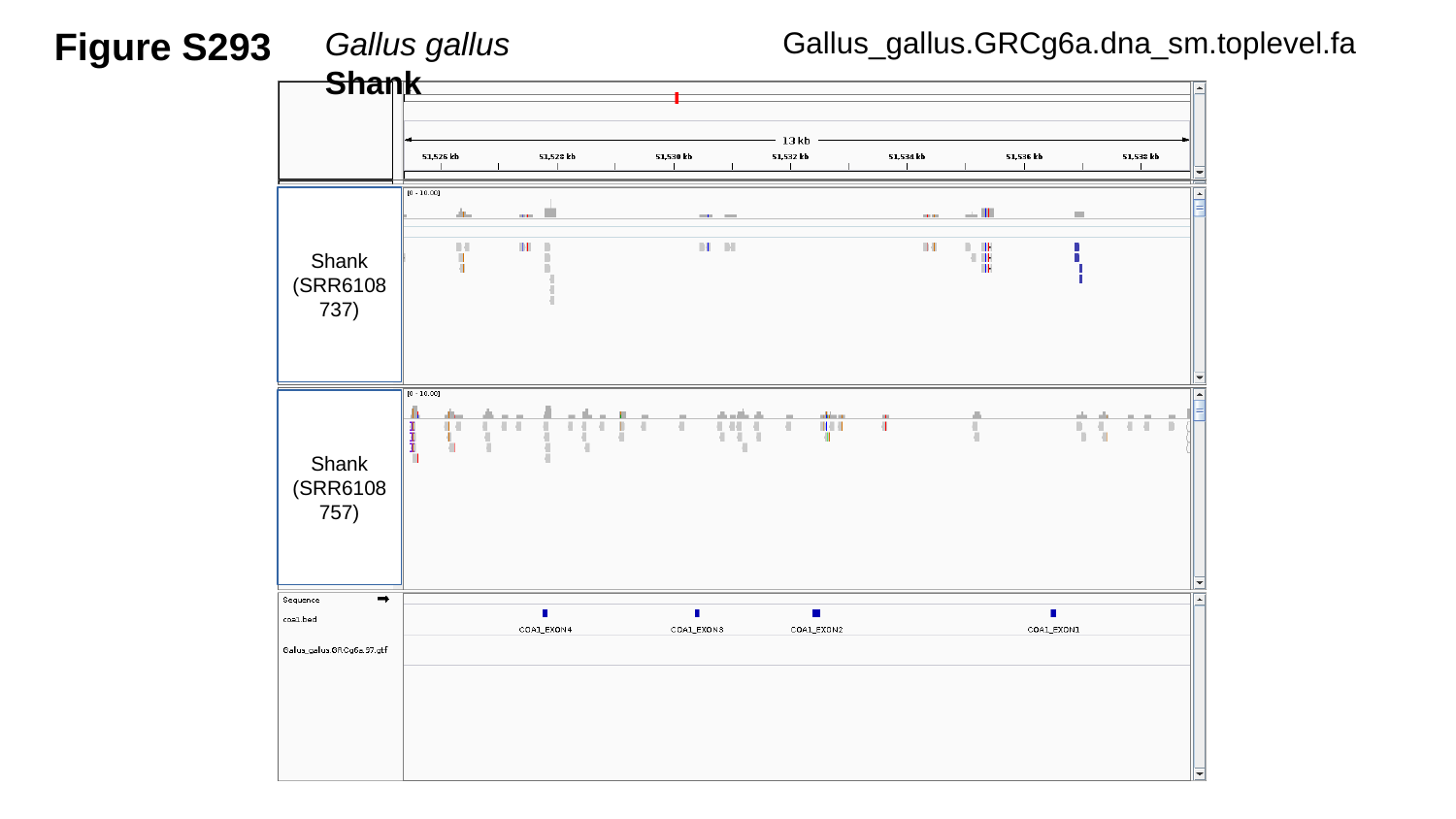

Figure S293
Gallus_gallus.GRCg6a.dna_sm.toplevel.fa
Gallus gallus	 Shank
Shank
(SRR6108737)
Shank
(SRR6108757)

### Slide 294
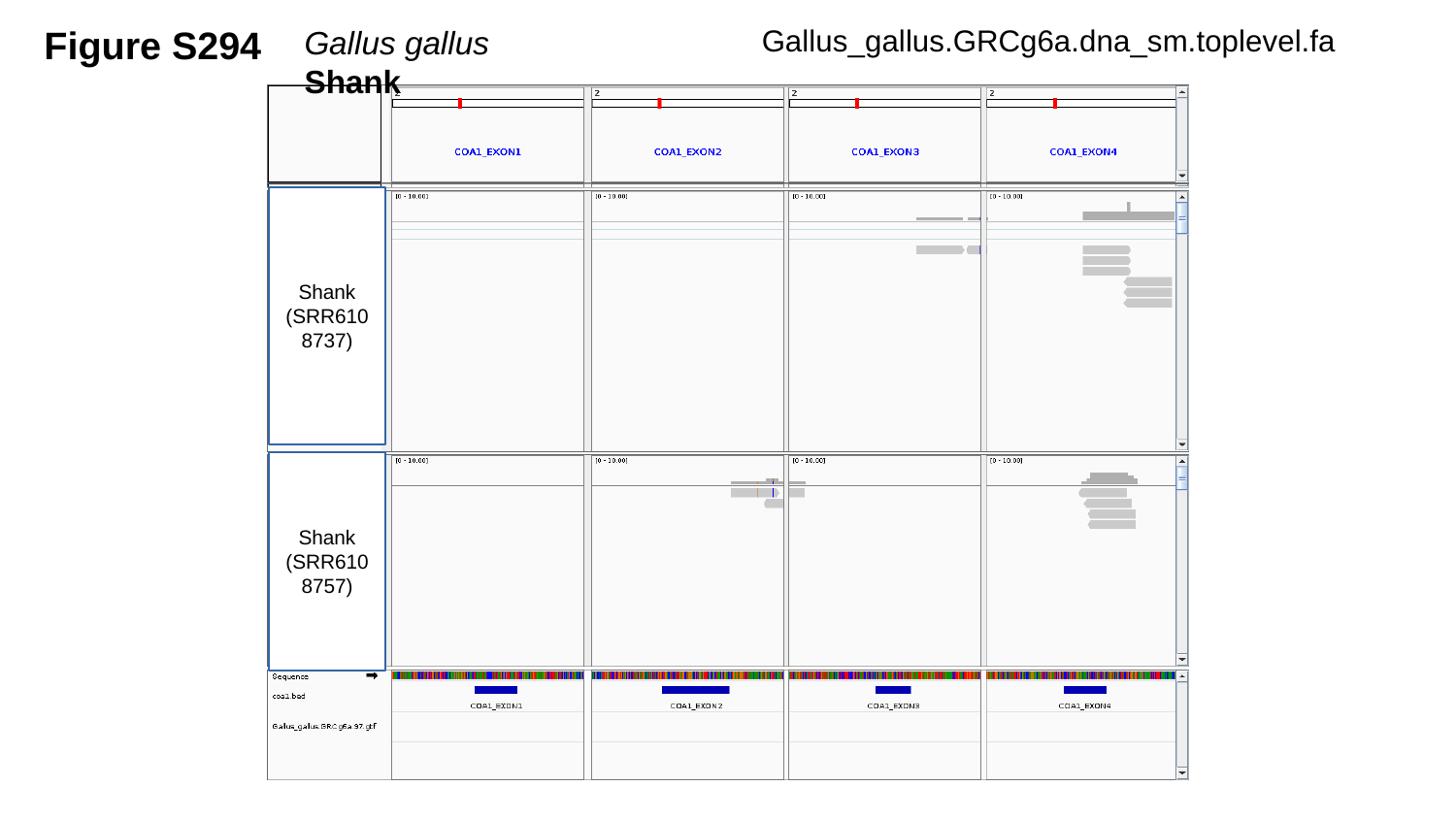

Figure S294
Gallus_gallus.GRCg6a.dna_sm.toplevel.fa
Gallus gallus	 Shank
Shank
(SRR6108737)
Shank
(SRR6108757)

### Slide 295
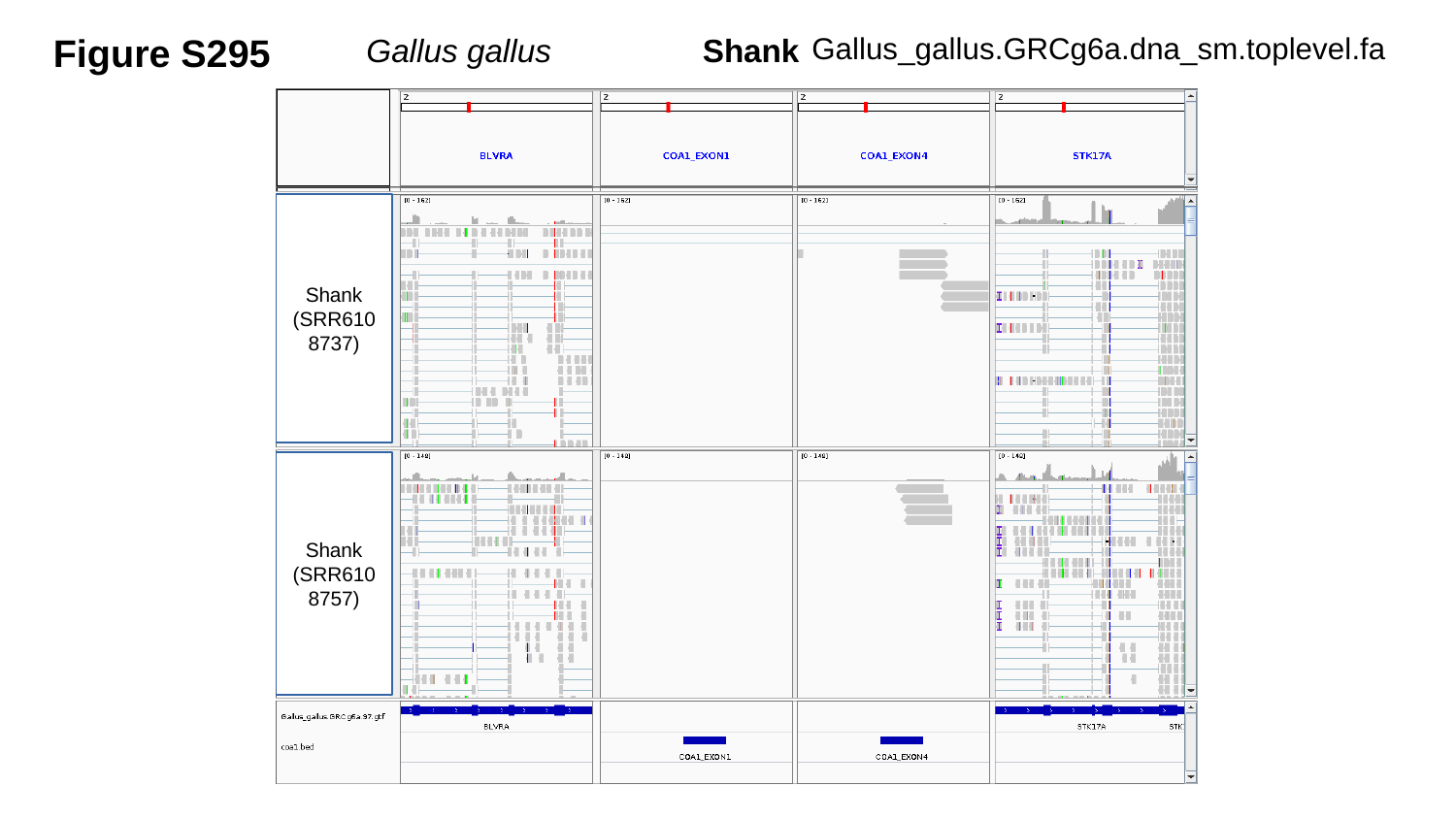

Figure S295
Gallus_gallus.GRCg6a.dna_sm.toplevel.fa
Gallus gallus	 Shank
Shank
(SRR6108737)
Shank
(SRR6108757)

### Slide 296
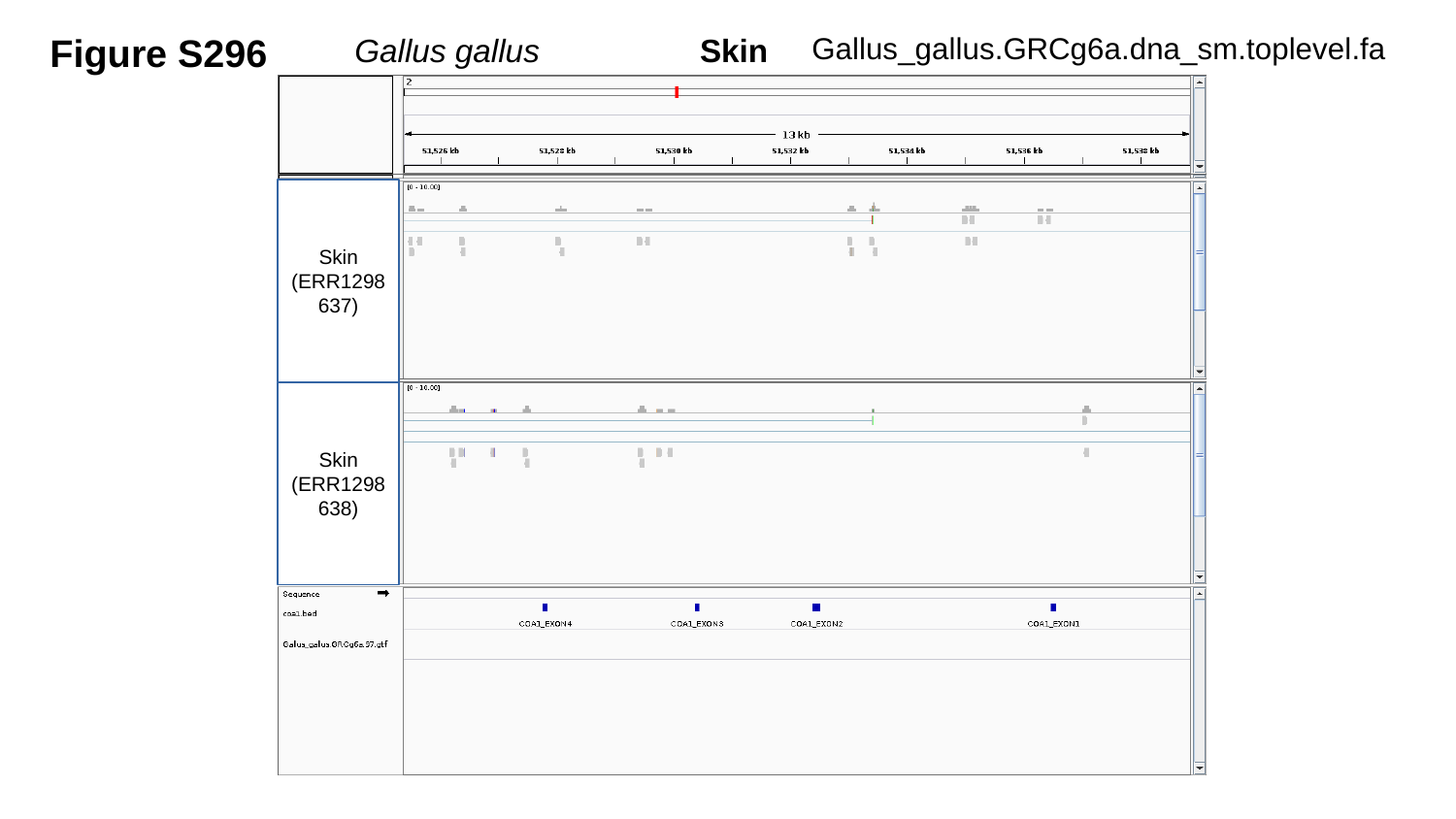

Figure S296
Gallus_gallus.GRCg6a.dna_sm.toplevel.fa
Gallus gallus	 Skin
Skin
(ERR1298637)
Skin
(ERR1298638)

### Slide 297
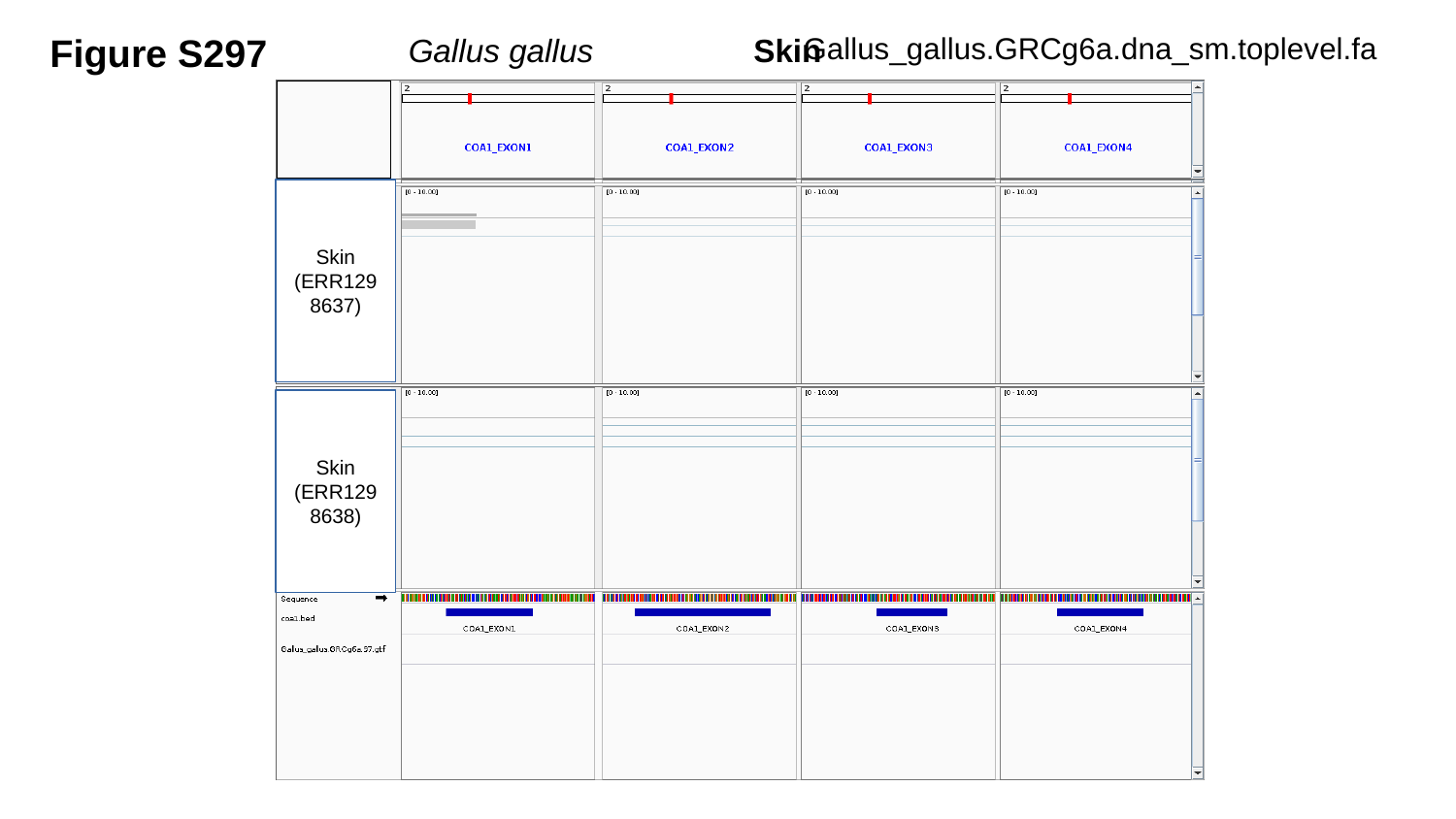

Figure S297
Gallus_gallus.GRCg6a.dna_sm.toplevel.fa
Gallus gallus	 Skin
Skin
(ERR1298637)
Skin
(ERR1298638)

### Slide 298
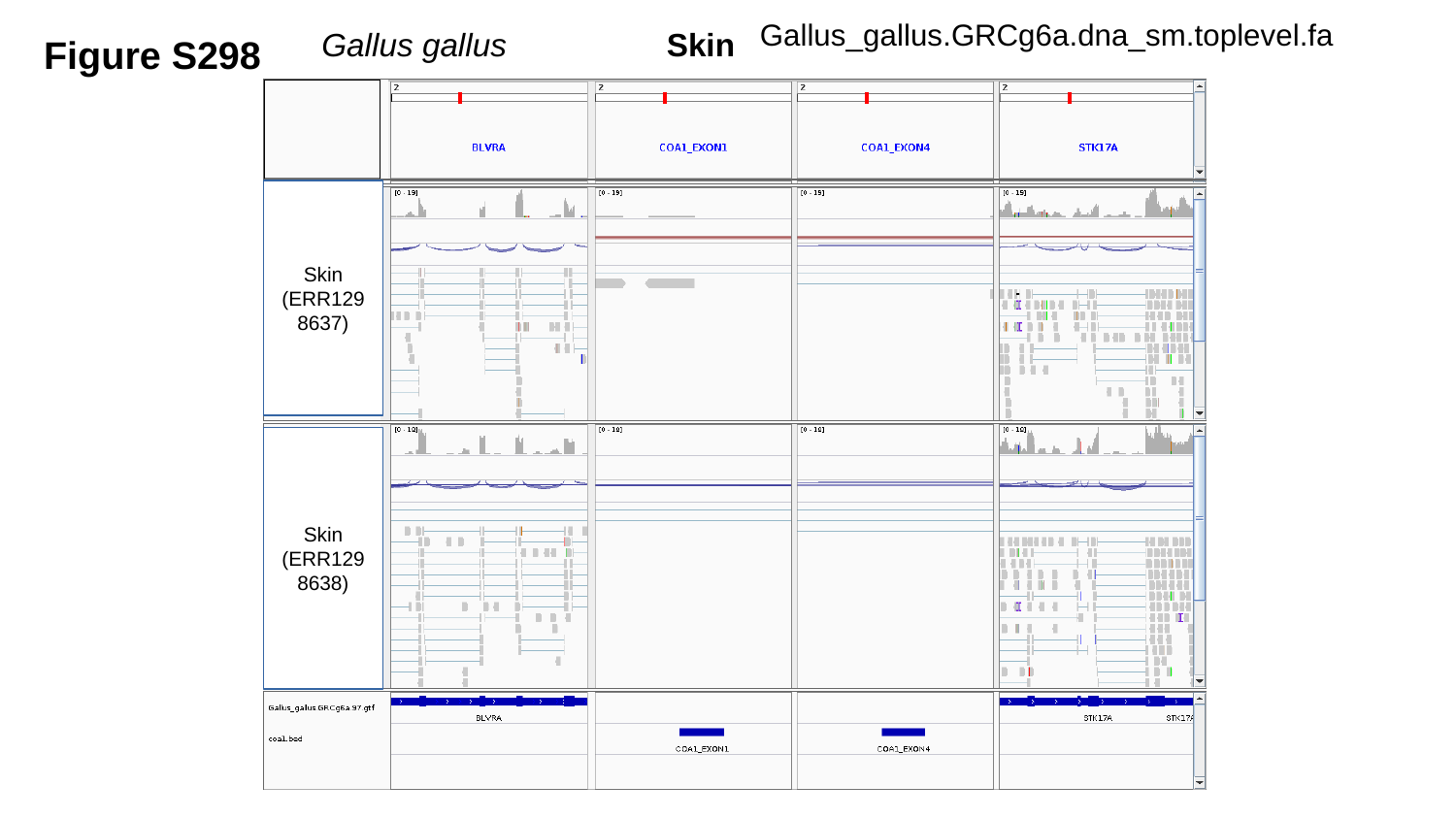

Gallus_gallus.GRCg6a.dna_sm.toplevel.fa
Figure S298
Gallus gallus	 Skin
Skin
(ERR1298637)
Skin
(ERR1298638)

### Slide 299
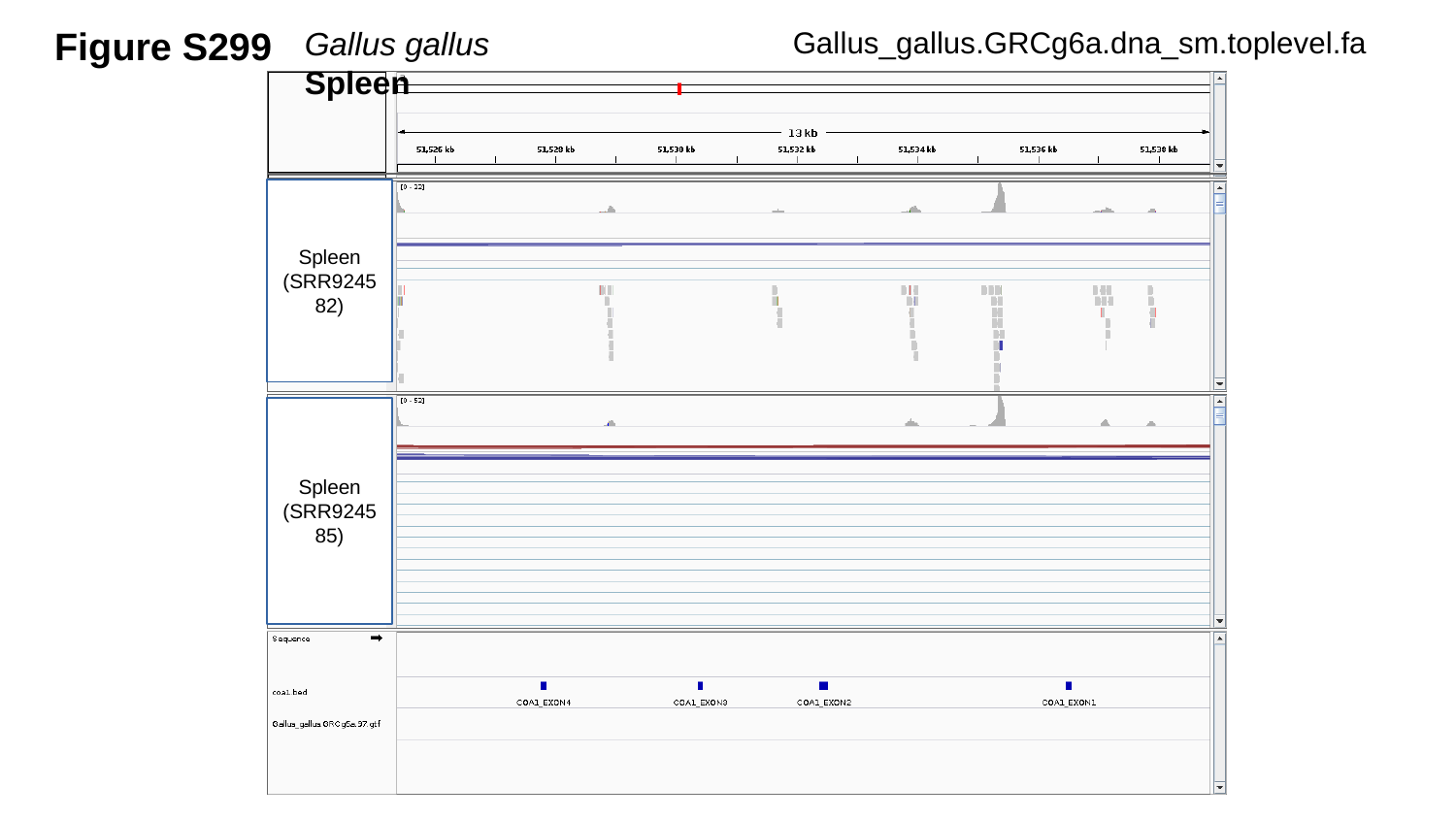

Figure S299
Gallus_gallus.GRCg6a.dna_sm.toplevel.fa
Gallus gallus	 	Spleen
Spleen
(SRR924582)
Spleen
(SRR924585)

### Slide 300
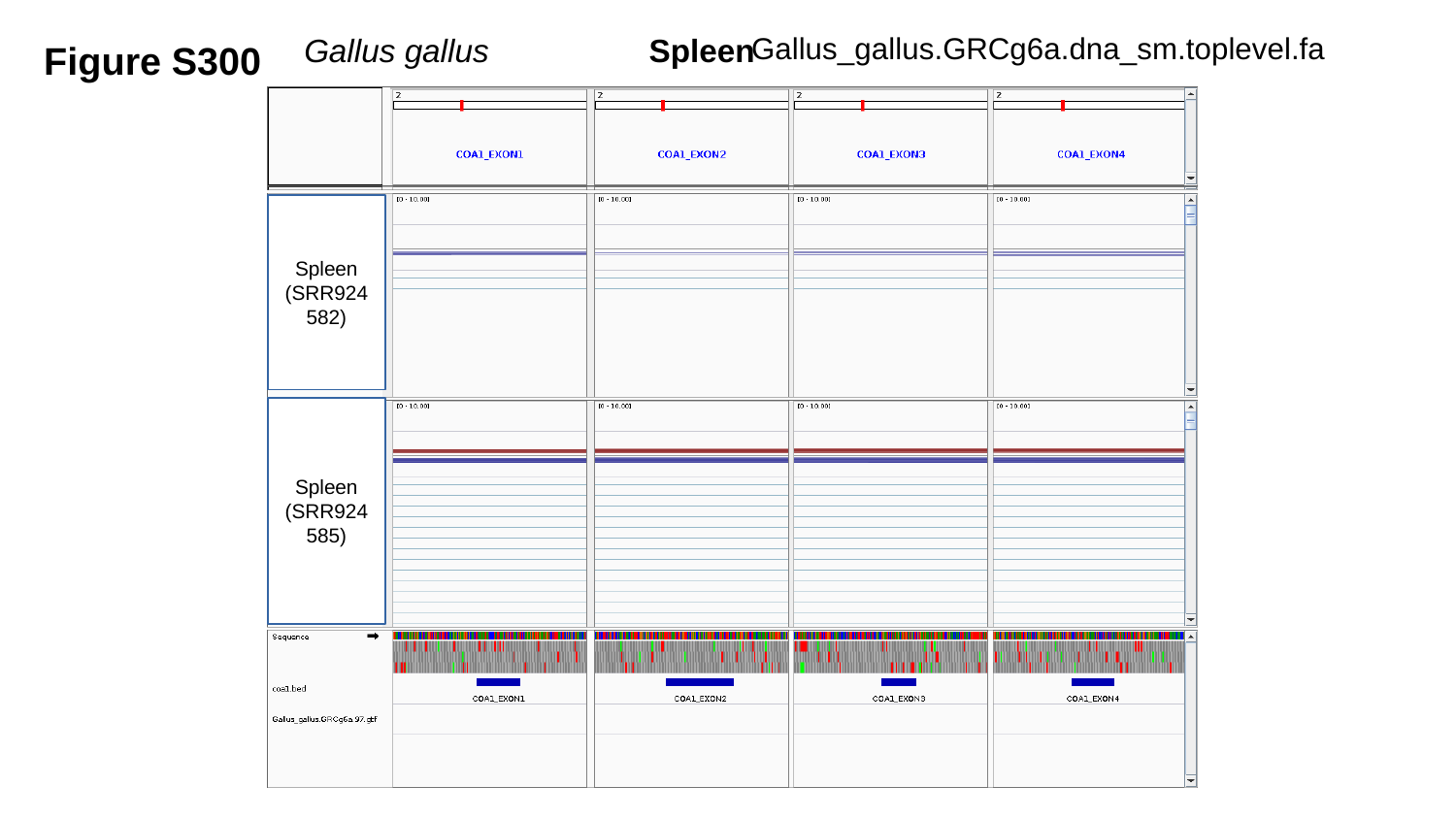

Figure S300
Gallus_gallus.GRCg6a.dna_sm.toplevel.fa
Gallus gallus	 Spleen
Spleen
(SRR924582)
Spleen
(SRR924585)

### Slide 301
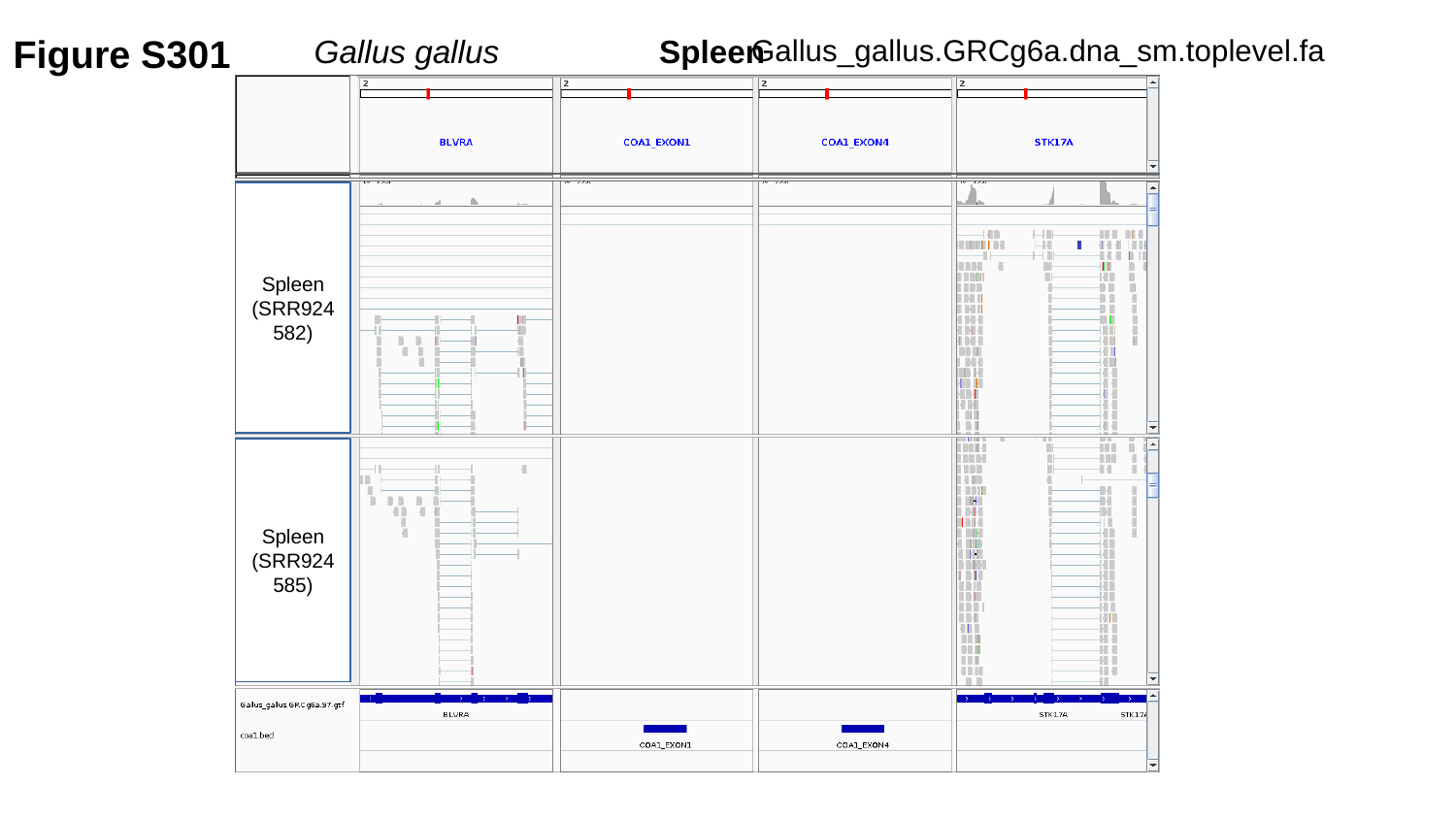

Figure S301
Gallus_gallus.GRCg6a.dna_sm.toplevel.fa
Gallus gallus	 Spleen
Spleen
(SRR924582)
Spleen
(SRR924585)

### Slide 302
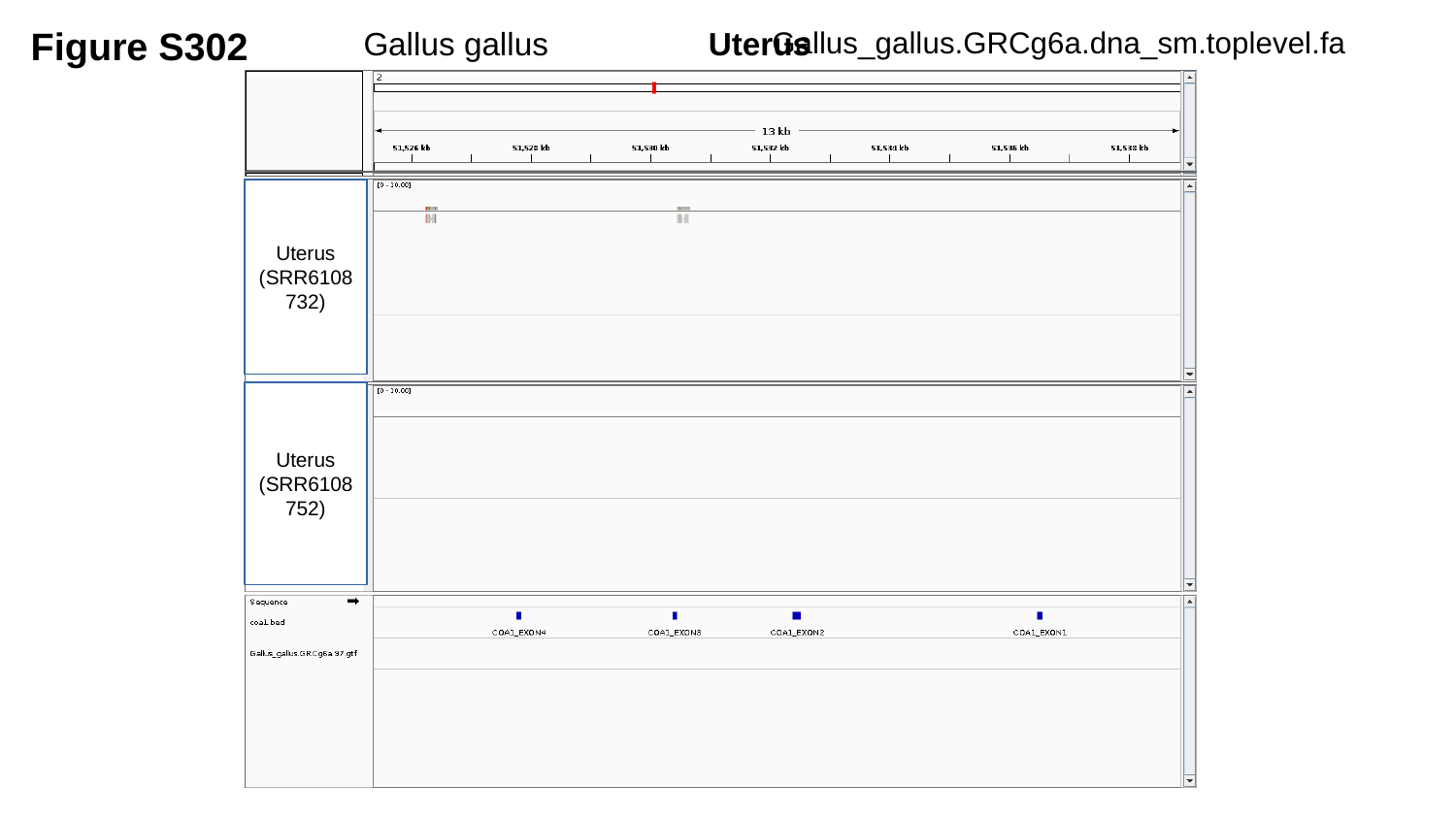

Figure S302
Gallus_gallus.GRCg6a.dna_sm.toplevel.fa
Gallus gallus	 Uterus
Uterus
(SRR6108732)
Uterus
(SRR6108752)

### Slide 303
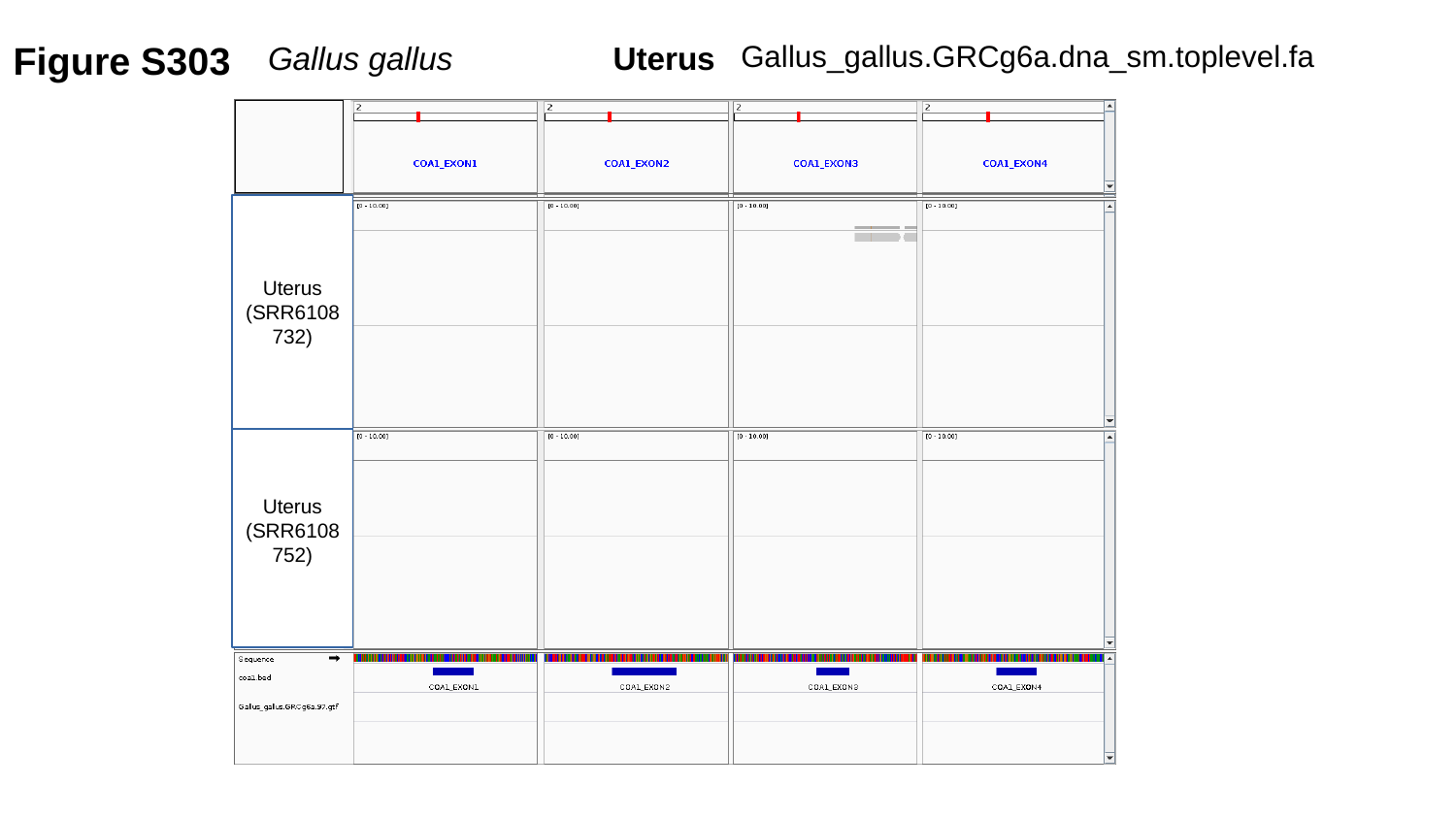

Figure S303
Gallus_gallus.GRCg6a.dna_sm.toplevel.fa
Gallus gallus	 Uterus
Uterus
(SRR6108732)
Uterus
(SRR6108752)

### Slide 304
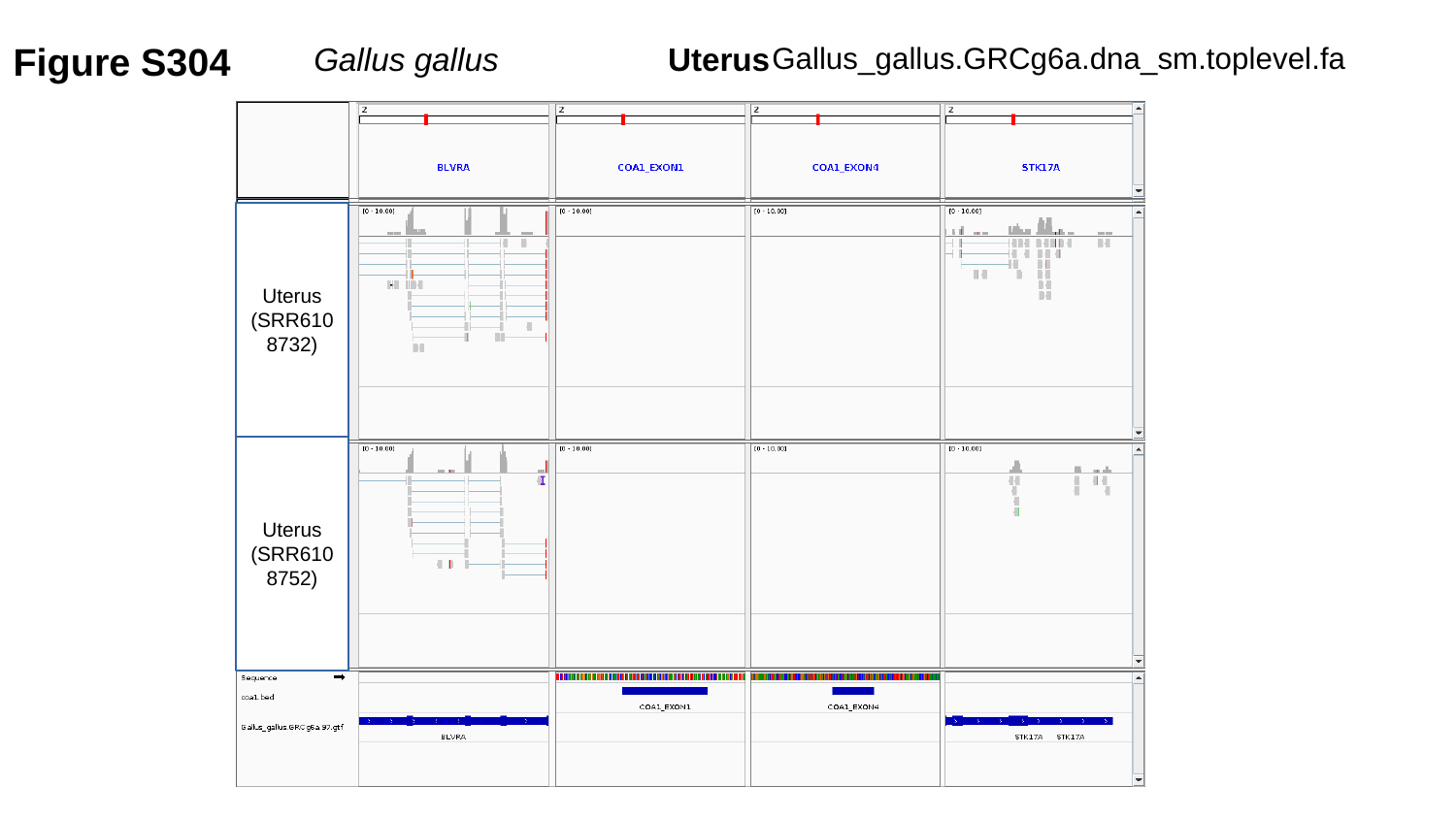

Figure S304
Gallus_gallus.GRCg6a.dna_sm.toplevel.fa
Gallus gallus	 Uterus
Uterus
(SRR6108732)
Uterus
(SRR6108752)

### Slide 305
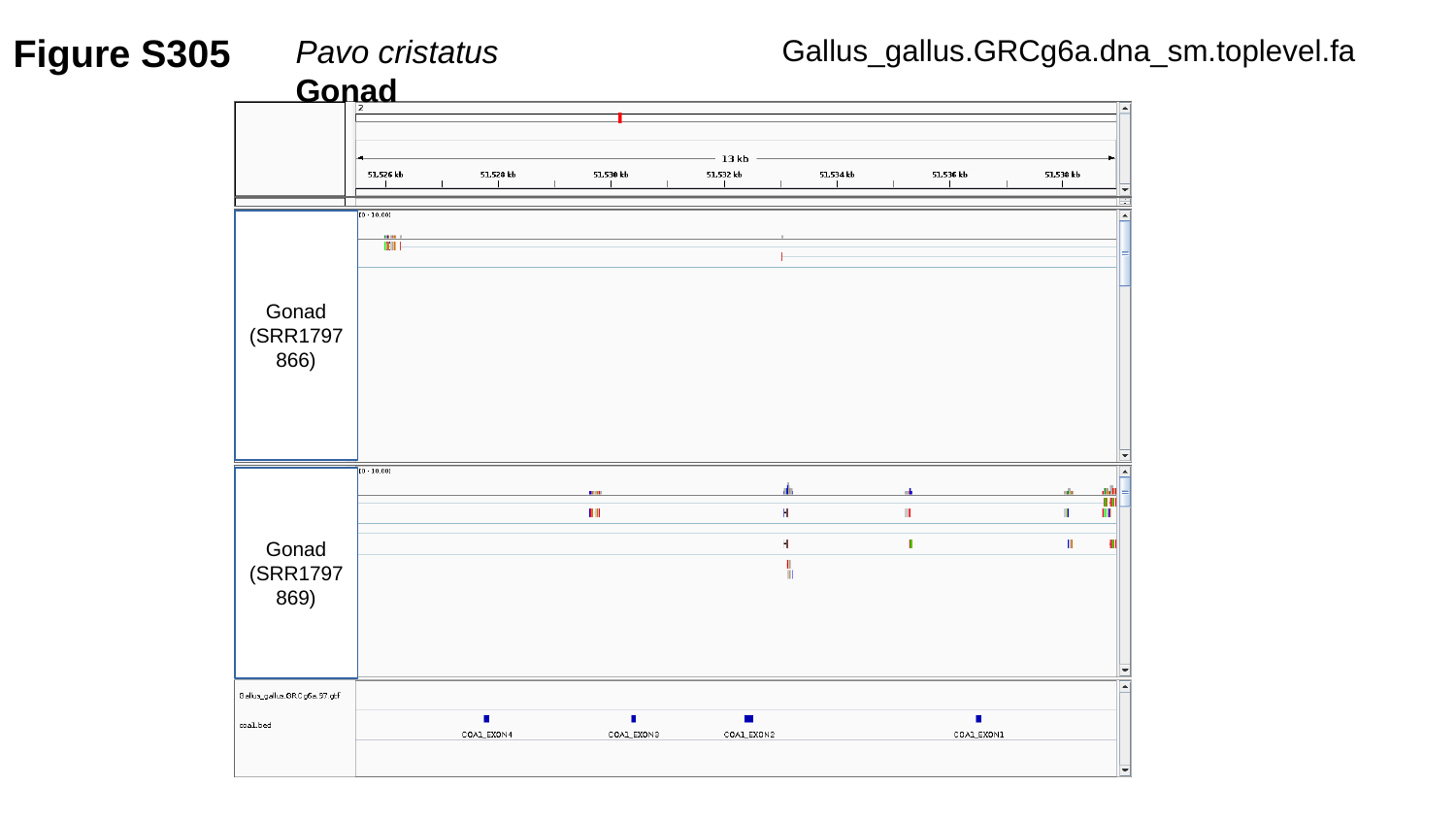

Figure S305
Gallus_gallus.GRCg6a.dna_sm.toplevel.fa
Pavo cristatus	Gonad
Gonad
(SRR1797866)
Gonad
(SRR1797869)

### Slide 306
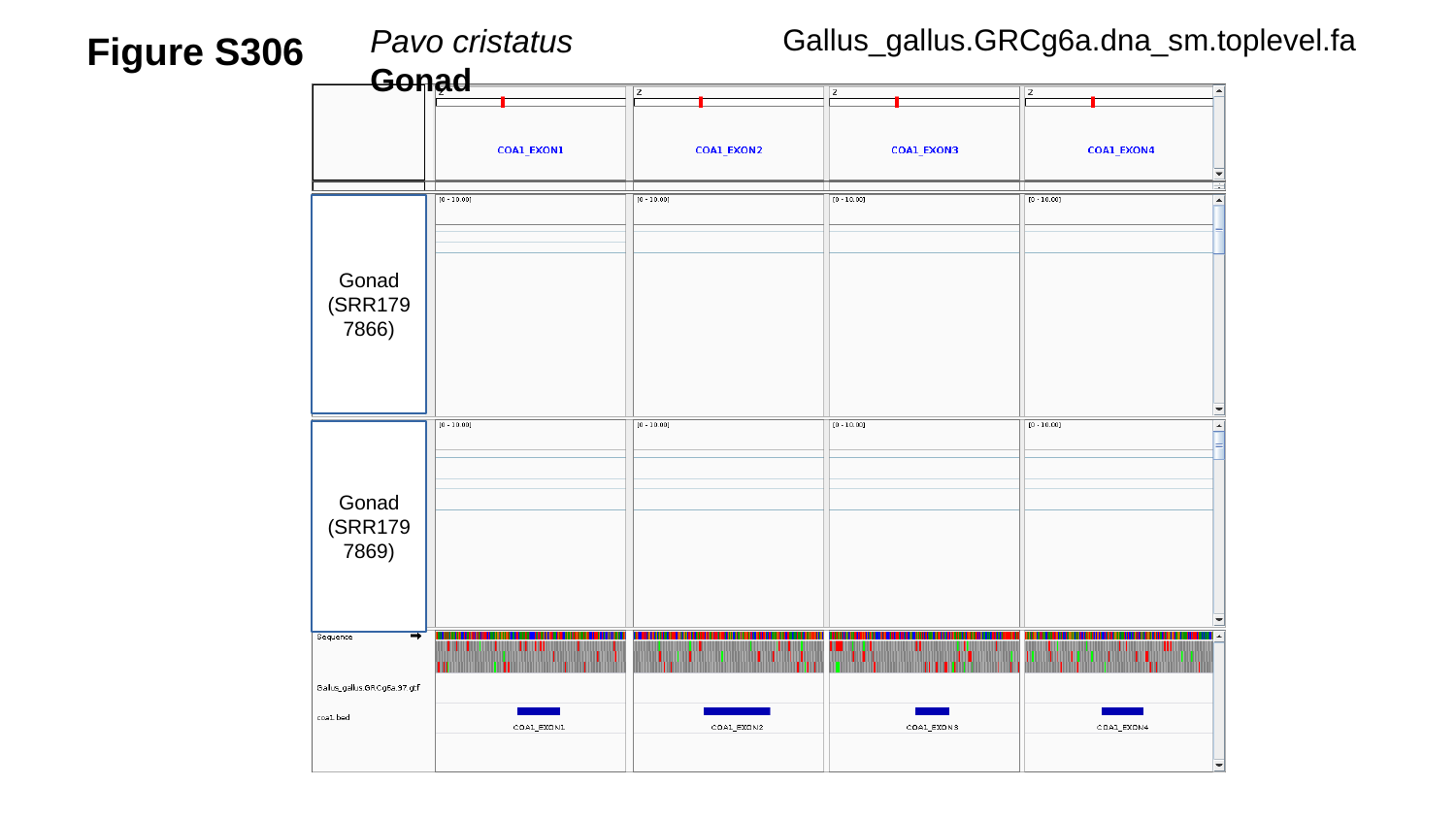

Figure S306
Pavo cristatus	Gonad
Gallus_gallus.GRCg6a.dna_sm.toplevel.fa
Gonad
(SRR1797866)
Gonad
(SRR1797869)

### Slide 307
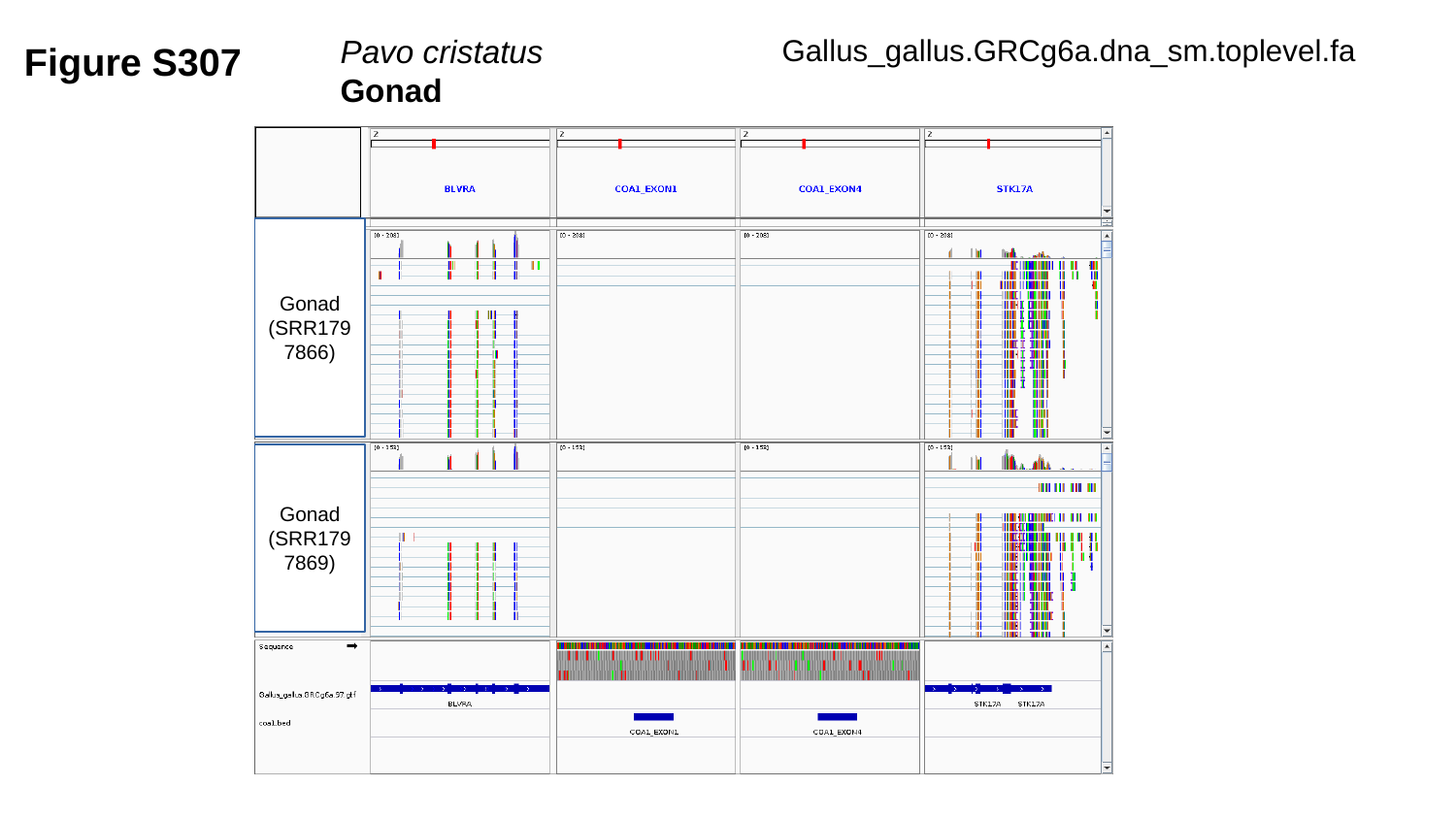

Gallus_gallus.GRCg6a.dna_sm.toplevel.fa
Figure S307
Pavo cristatus	Gonad
Gonad
(SRR1797866)
Gonad
(SRR1797869)

### Slide 308
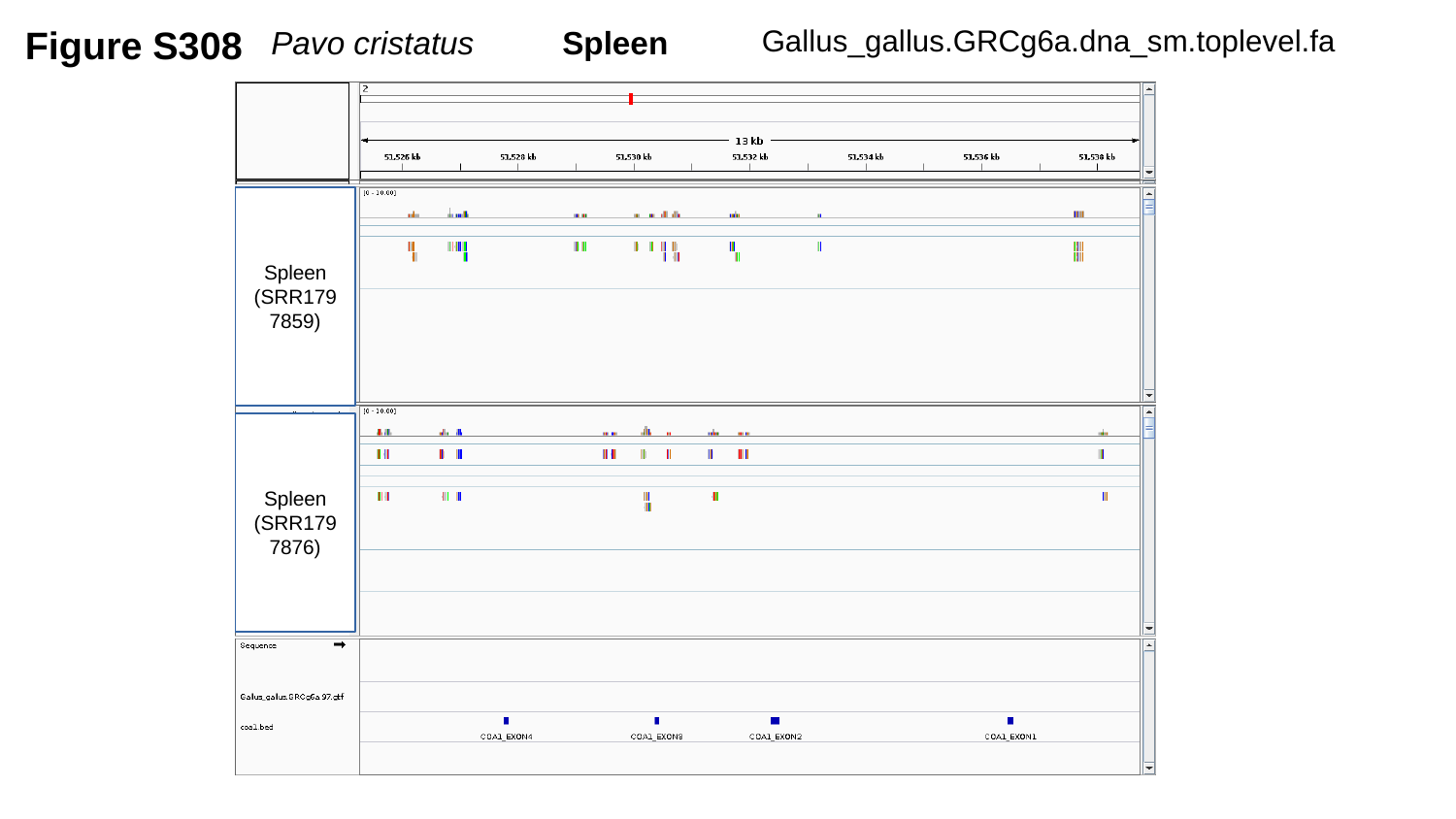

Figure S308
Gallus_gallus.GRCg6a.dna_sm.toplevel.fa
Pavo cristatus	Spleen
Spleen
(SRR1797859)
Spleen
(SRR1797876)

### Slide 309
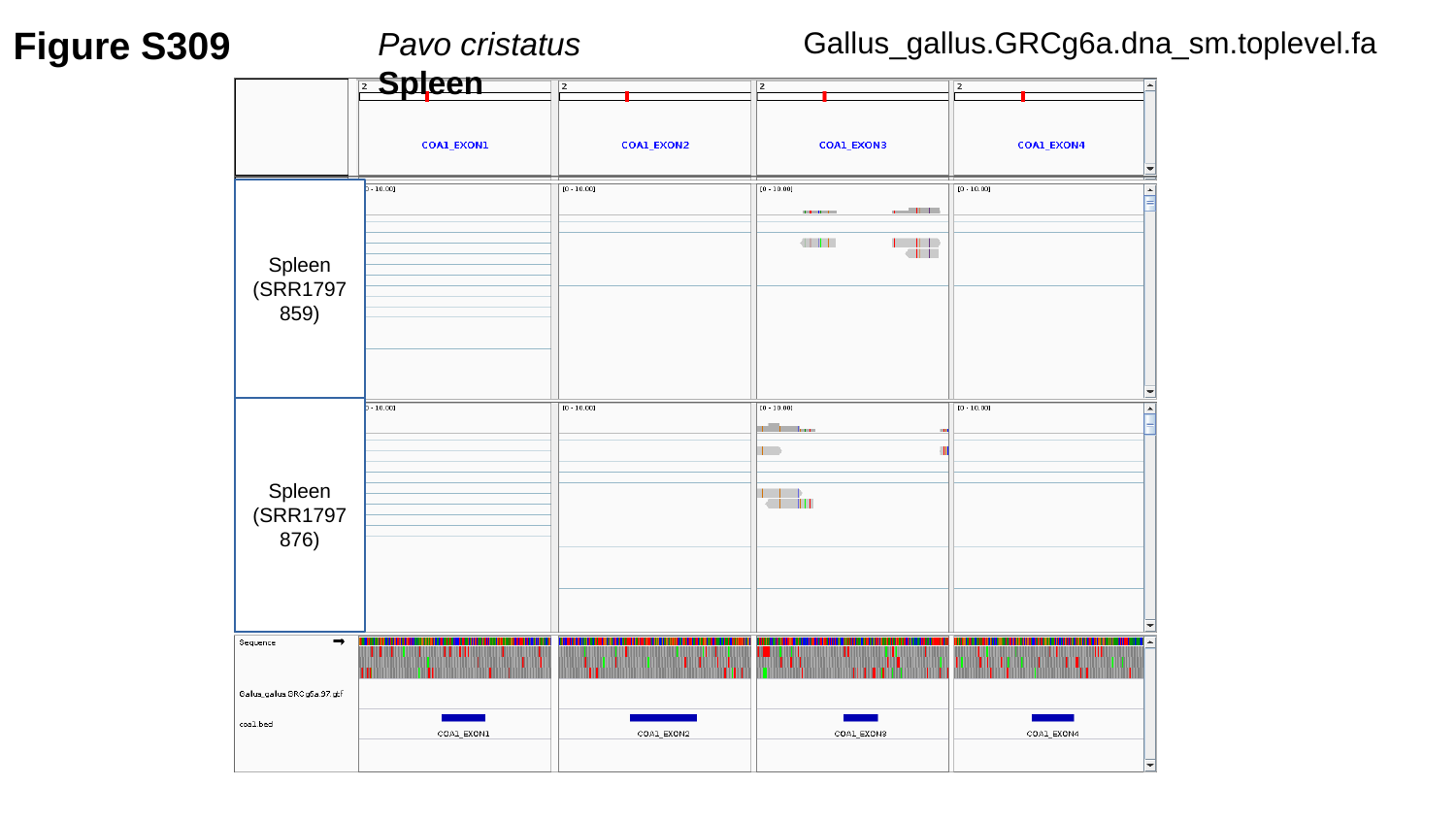

Figure S309
Gallus_gallus.GRCg6a.dna_sm.toplevel.fa
Pavo cristatus	Spleen
Spleen
(SRR1797859)
Spleen
(SRR1797876)

### Slide 310
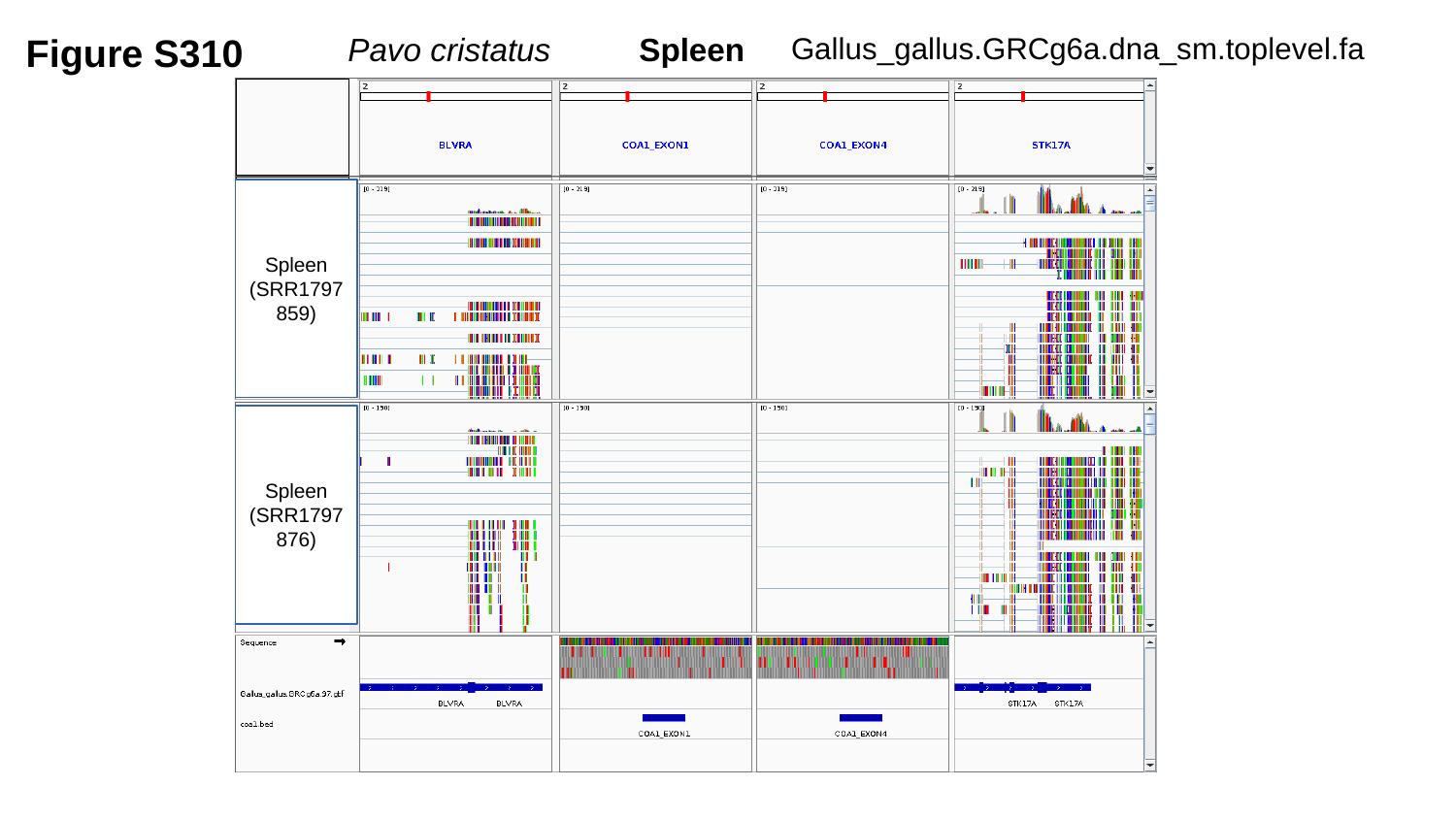

Figure S310
Gallus_gallus.GRCg6a.dna_sm.toplevel.fa
Pavo cristatus	Spleen
Spleen
(SRR1797859)
Spleen
(SRR1797876)

### Slide 311
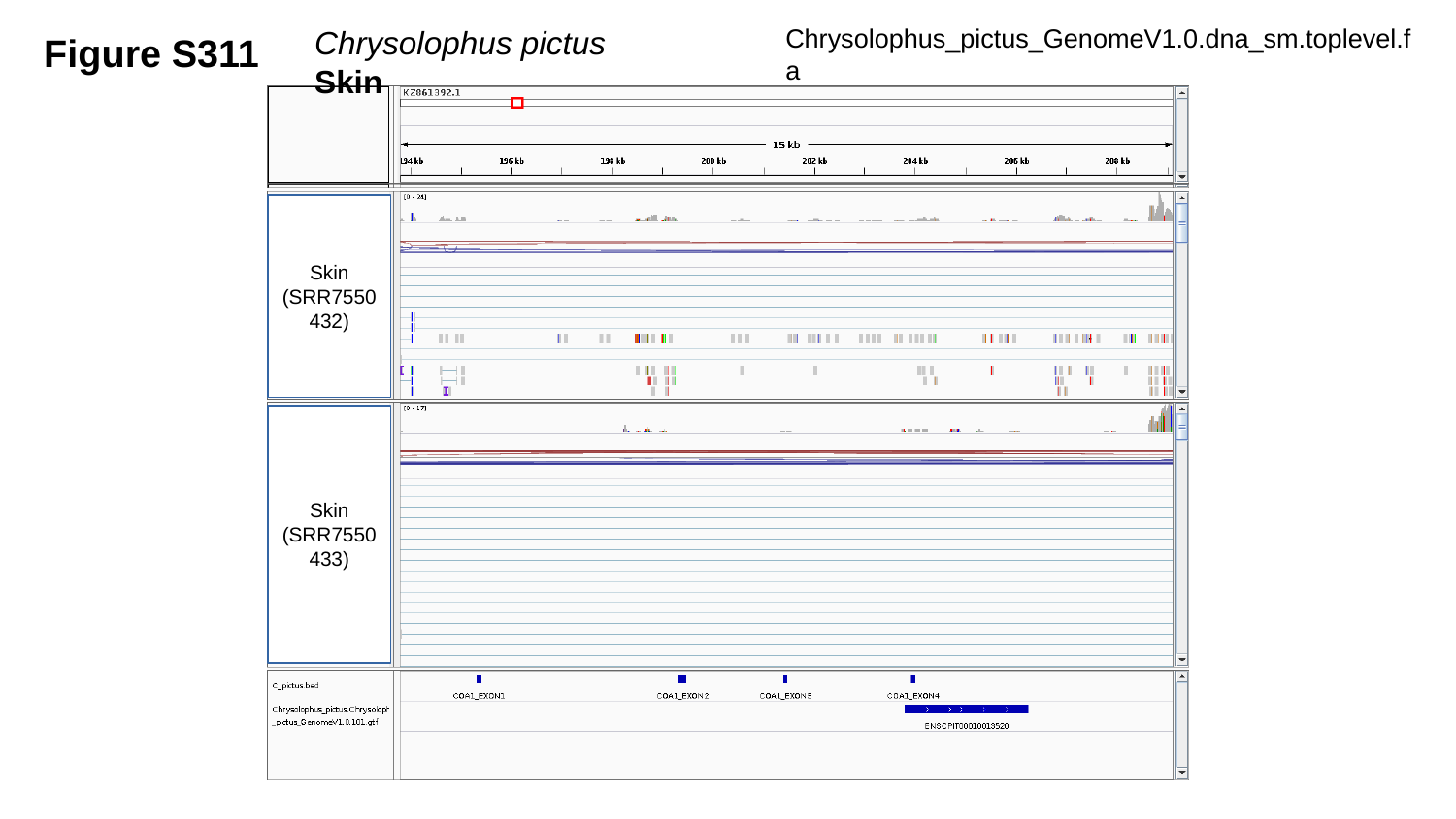

Figure S311
Chrysolophus_pictus_GenomeV1.0.dna_sm.toplevel.fa
Chrysolophus pictus	Skin
Skin
(SRR7550432)
Skin
(SRR7550433)

### Slide 312
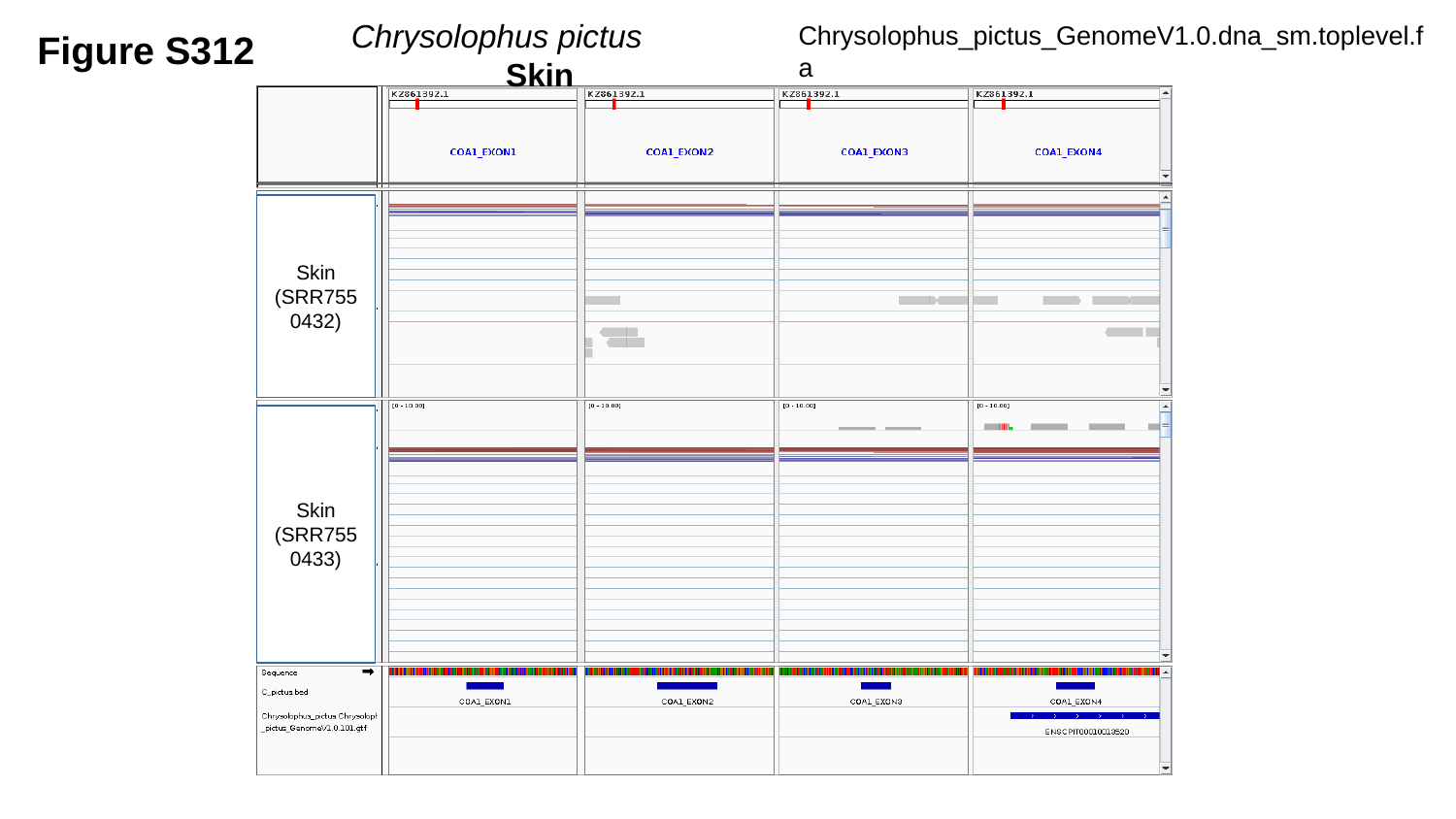

Chrysolophus pictus	 Skin
Figure S312
Chrysolophus_pictus_GenomeV1.0.dna_sm.toplevel.fa
Skin
(SRR7550432)
Skin
(SRR7550433)

### Slide 313
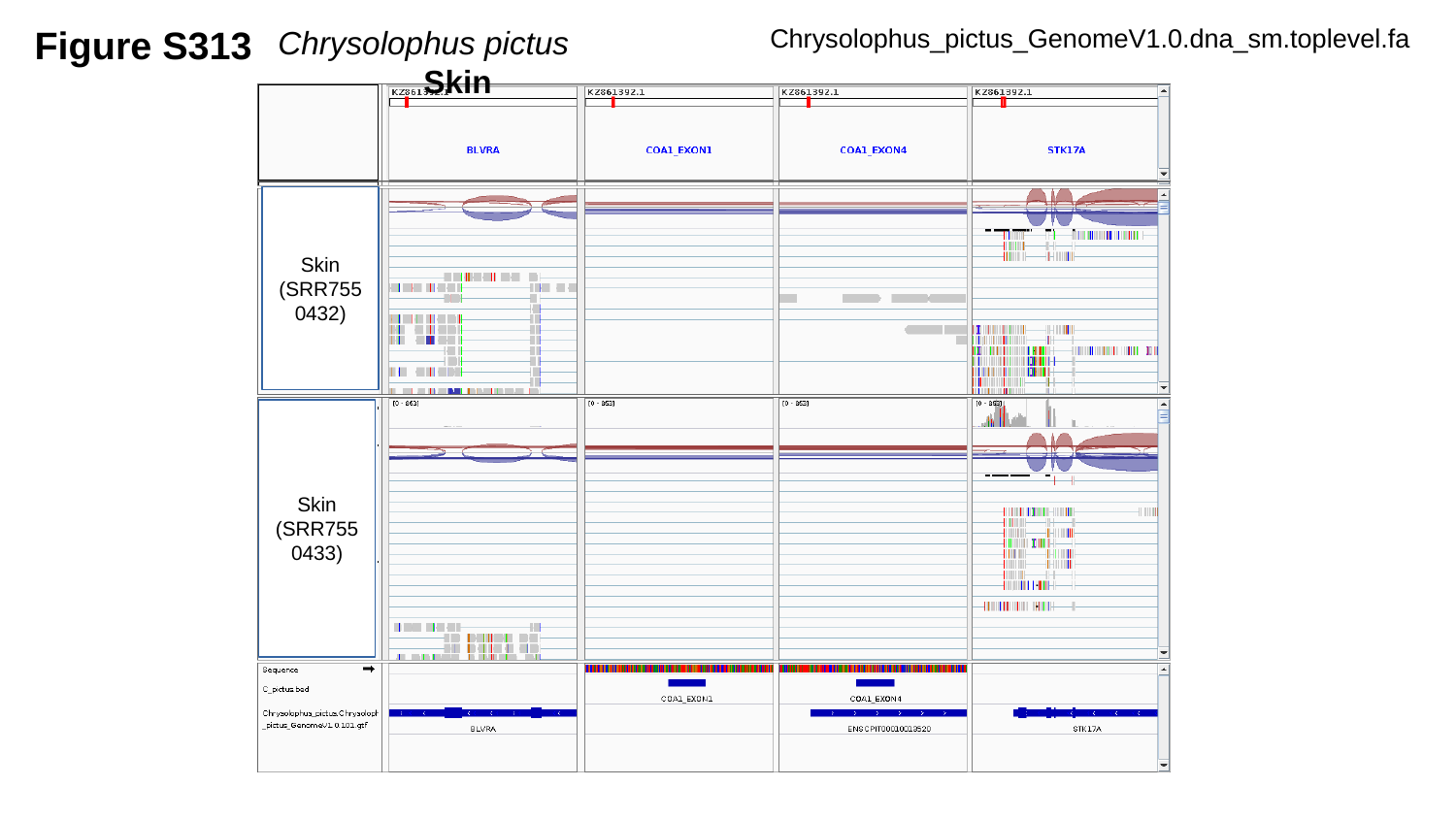

Figure S313
Chrysolophus pictus		Skin
Chrysolophus_pictus_GenomeV1.0.dna_sm.toplevel.fa
Skin
(SRR7550432)
Skin
(SRR7550433)

### Slide 314
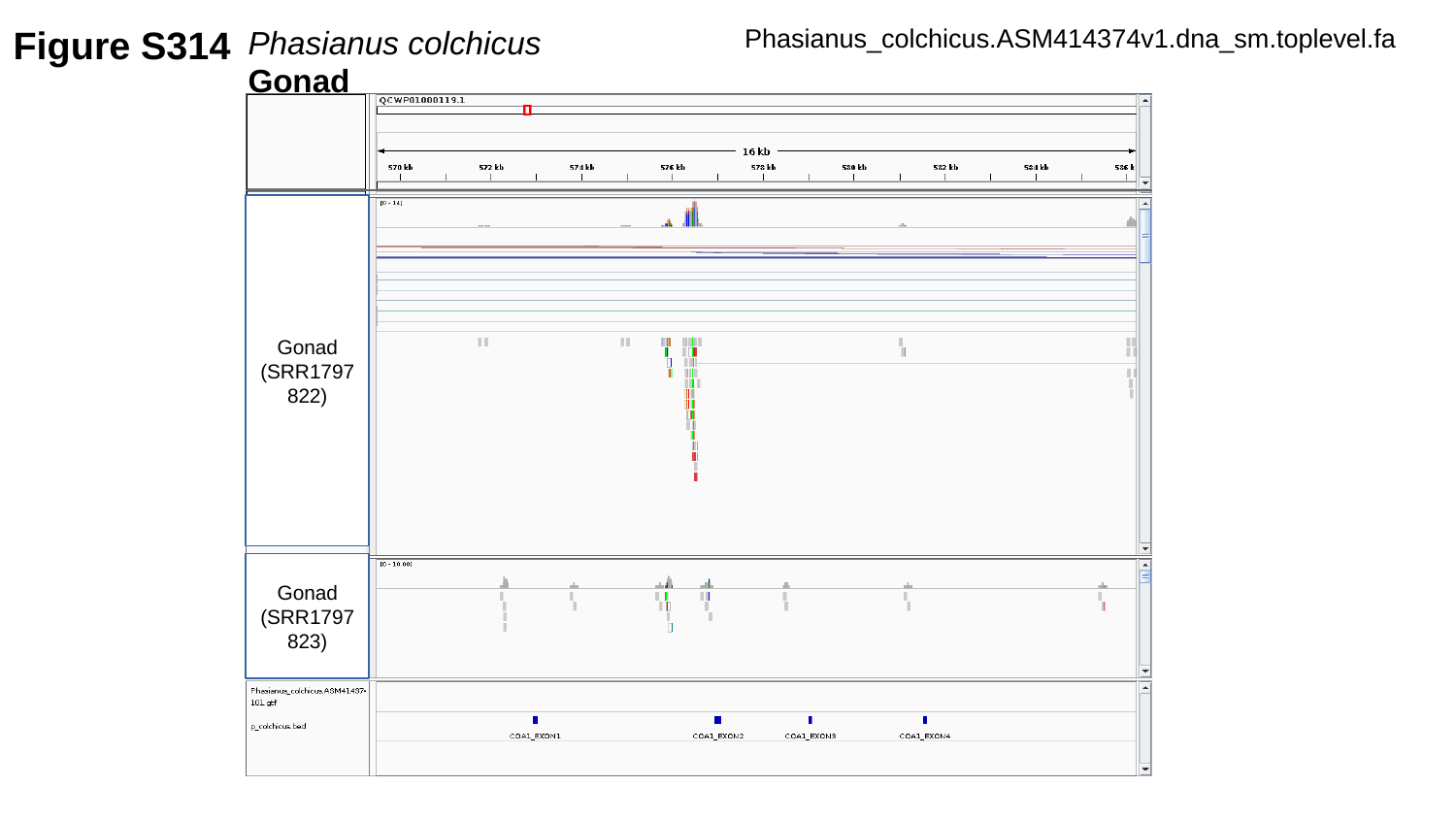

Figure S314
Phasianus colchicus	Gonad
Phasianus_colchicus.ASM414374v1.dna_sm.toplevel.fa
Gonad
(SRR1797822)
Gonad
(SRR1797823)

### Slide 315
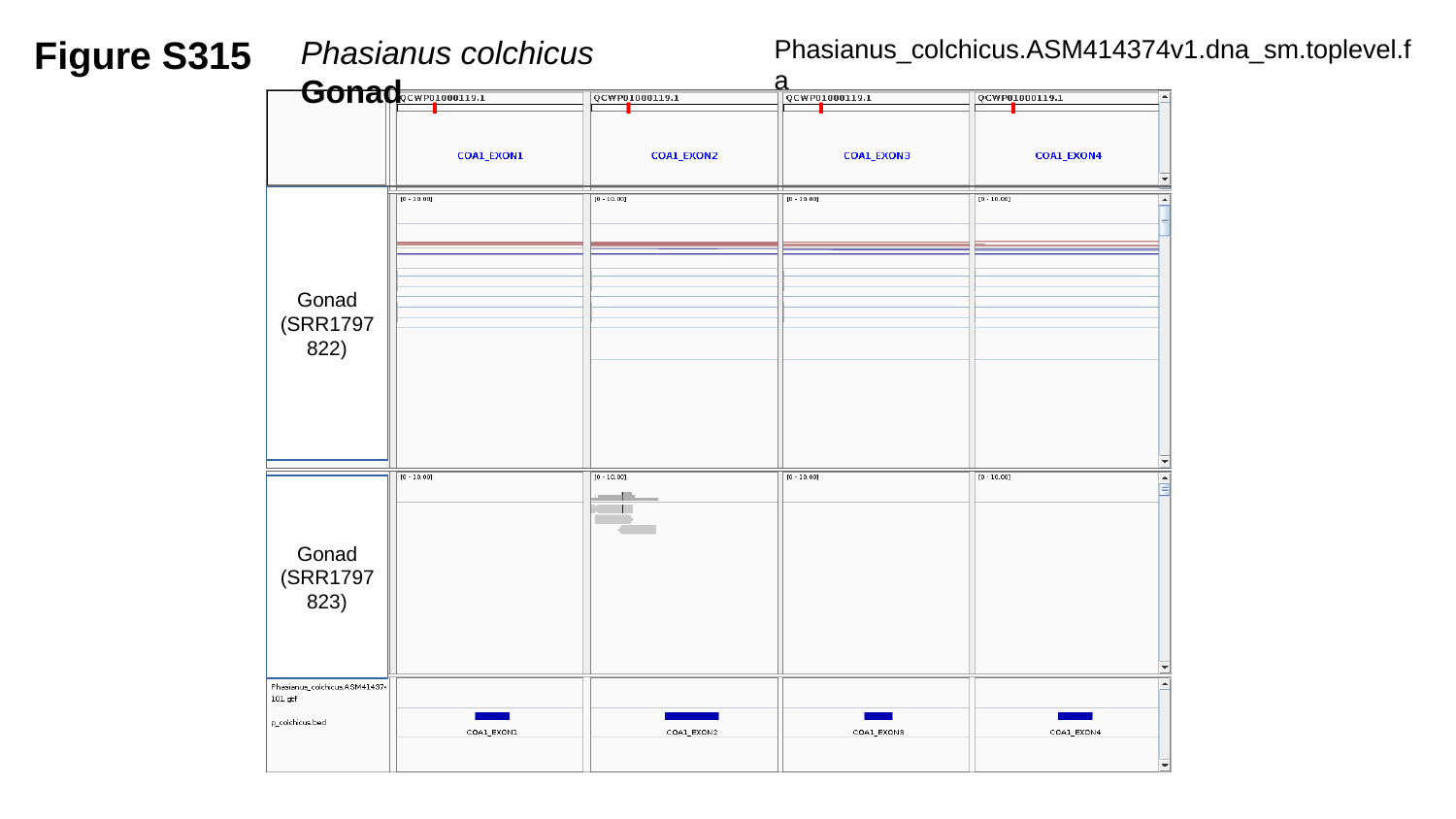

Figure S315
Phasianus colchicus	Gonad
Phasianus_colchicus.ASM414374v1.dna_sm.toplevel.fa
Gonad
(SRR1797822)
Gonad
(SRR1797823)

### Slide 316
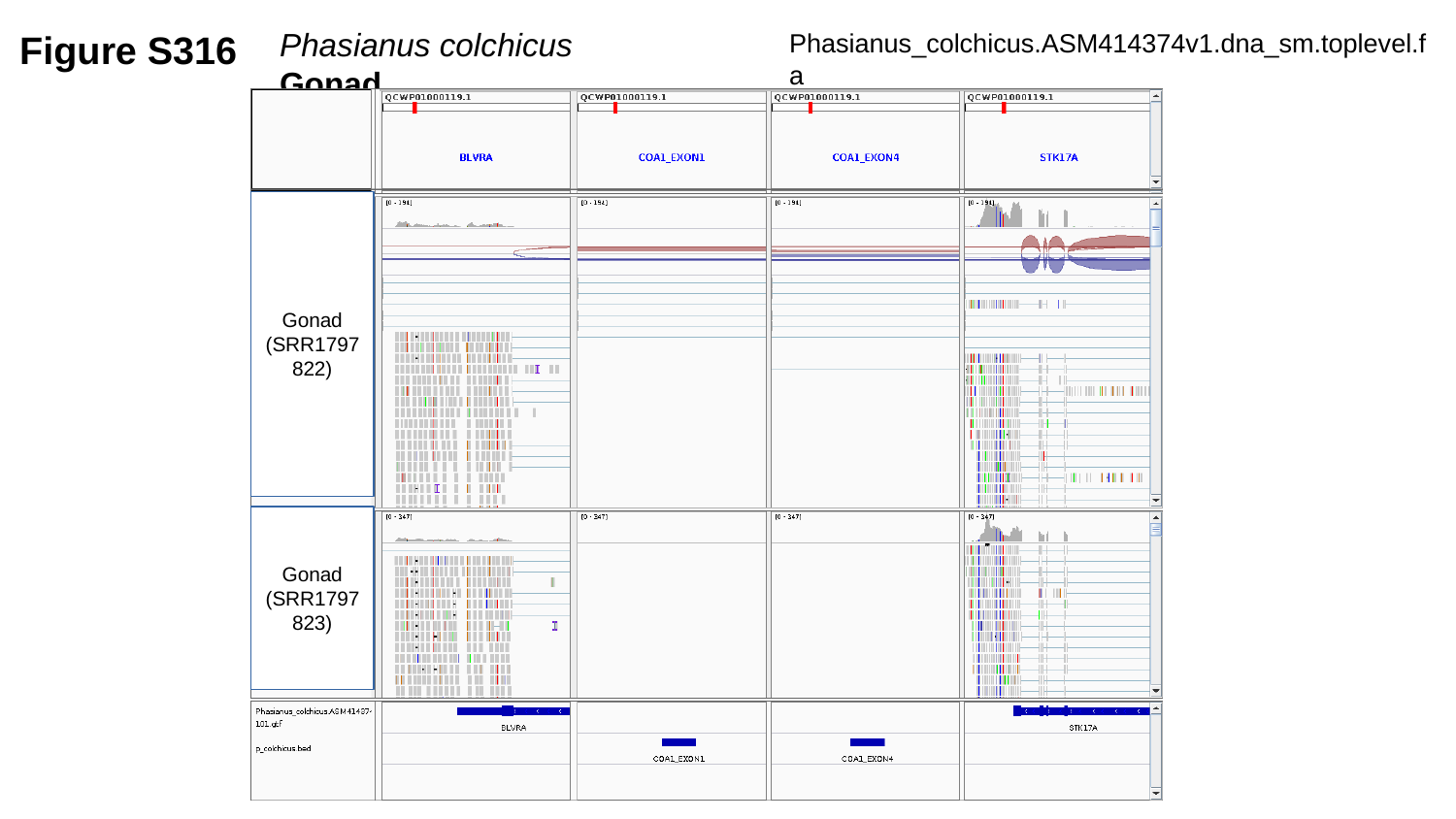

Figure S316
Phasianus colchicus	Gonad
Phasianus_colchicus.ASM414374v1.dna_sm.toplevel.fa
Gonad
(SRR1797822)
Gonad
(SRR1797823)

### Slide 317
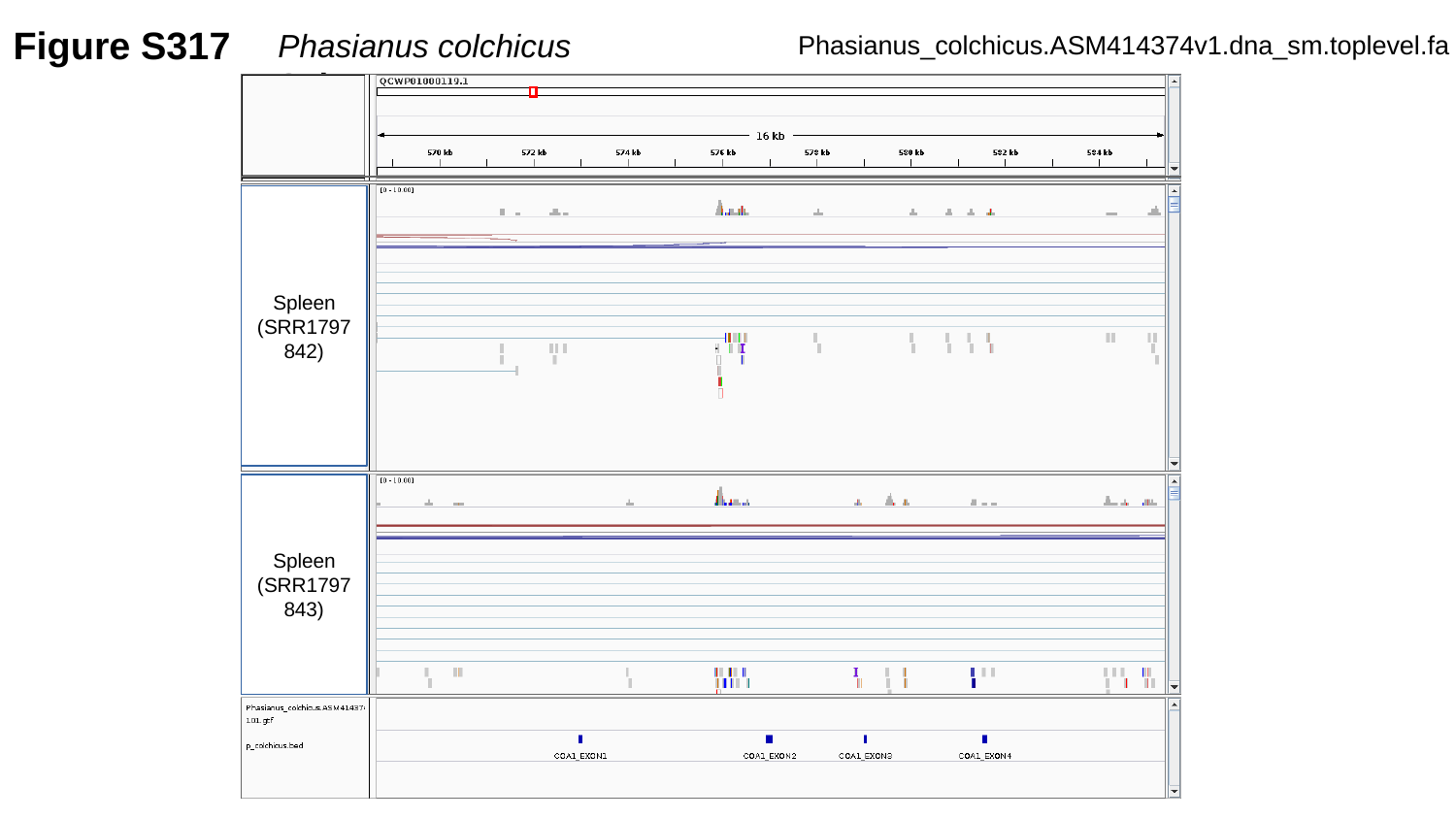

Figure S317
Phasianus colchicus	Spleen
Phasianus_colchicus.ASM414374v1.dna_sm.toplevel.fa
Spleen
(SRR1797842)
Spleen
(SRR1797843)

### Slide 318
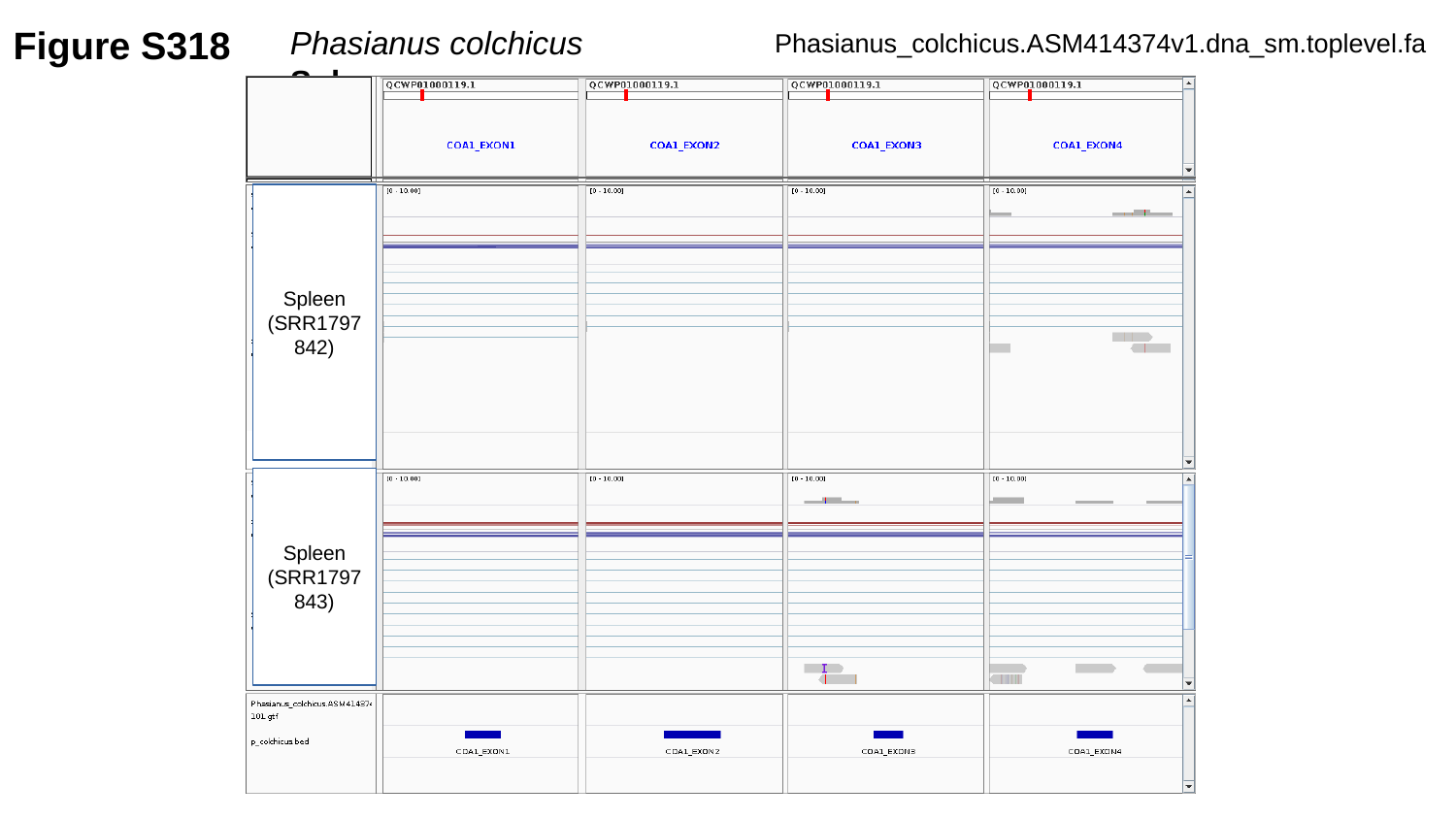

Figure S318
Phasianus colchicus	Spleen
Phasianus_colchicus.ASM414374v1.dna_sm.toplevel.fa
Spleen
(SRR1797842)
Spleen
(SRR1797843)

### Slide 319
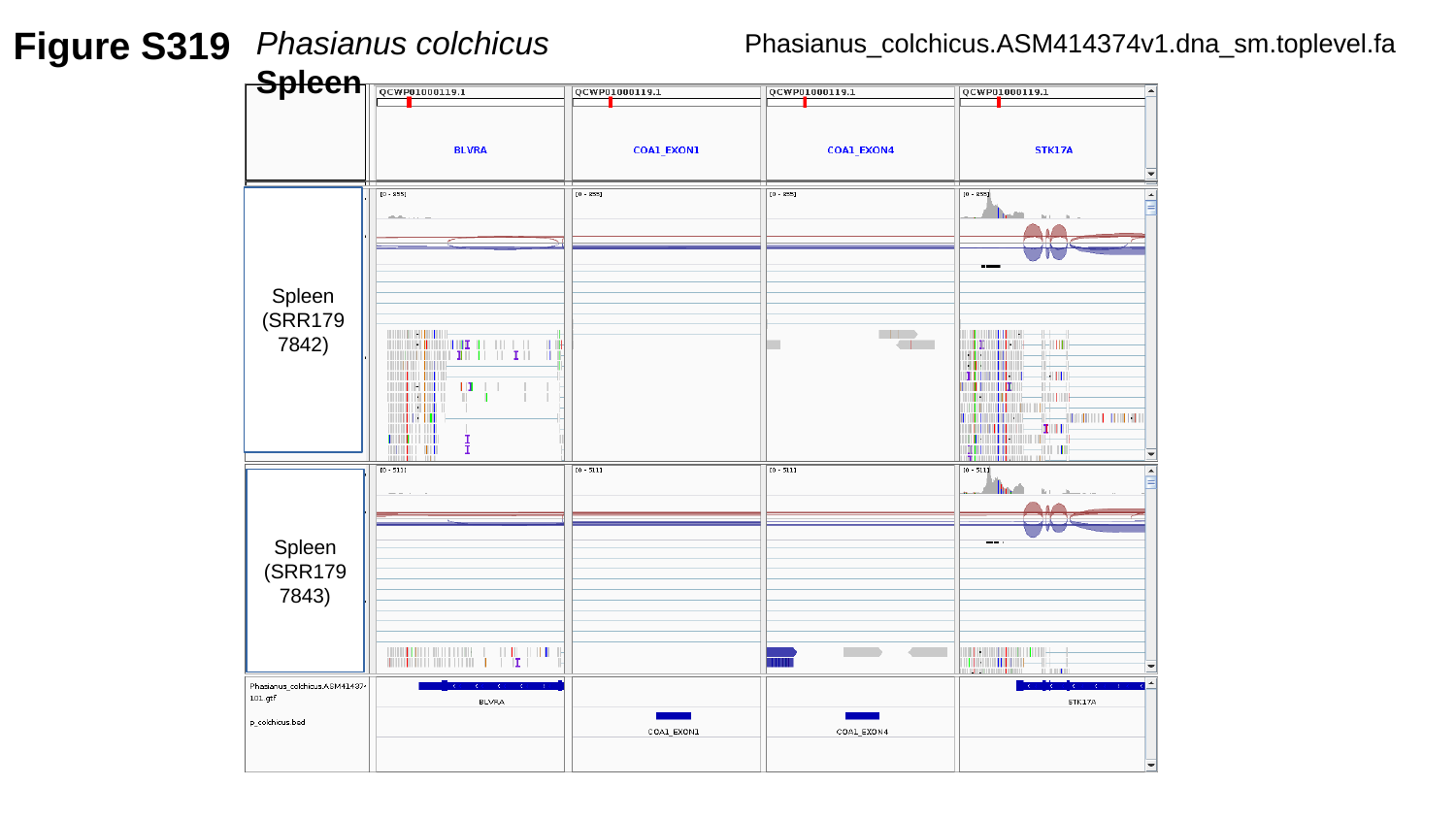

Figure S319
Phasianus colchicus	Spleen
Phasianus_colchicus.ASM414374v1.dna_sm.toplevel.fa
Spleen
(SRR1797842)
Spleen
(SRR1797843)

### Slide 320
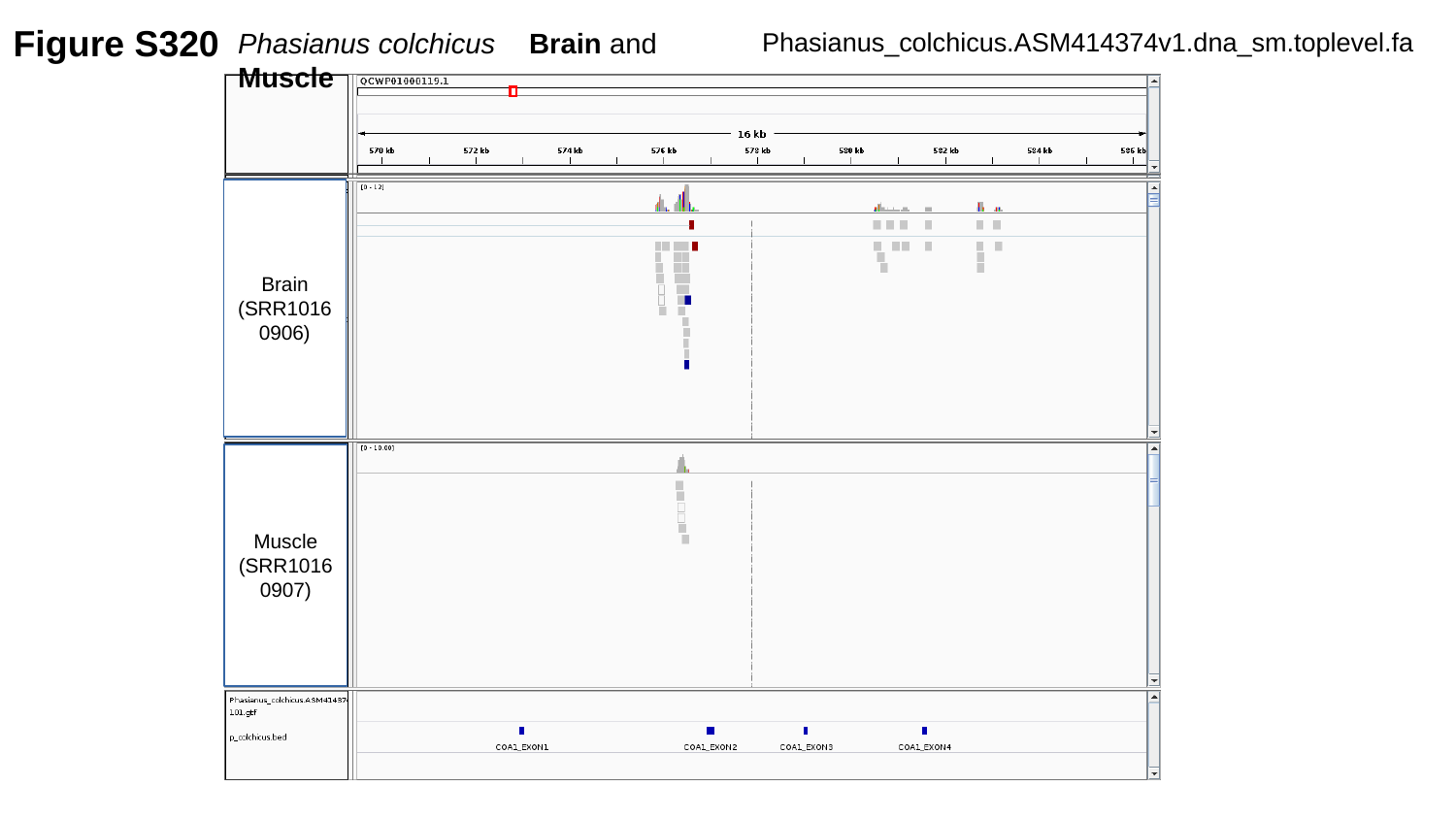

Figure S320
Phasianus colchicus	Brain and Muscle
Phasianus_colchicus.ASM414374v1.dna_sm.toplevel.fa
Brain
(SRR10160906)
Muscle
(SRR10160907)

### Slide 321
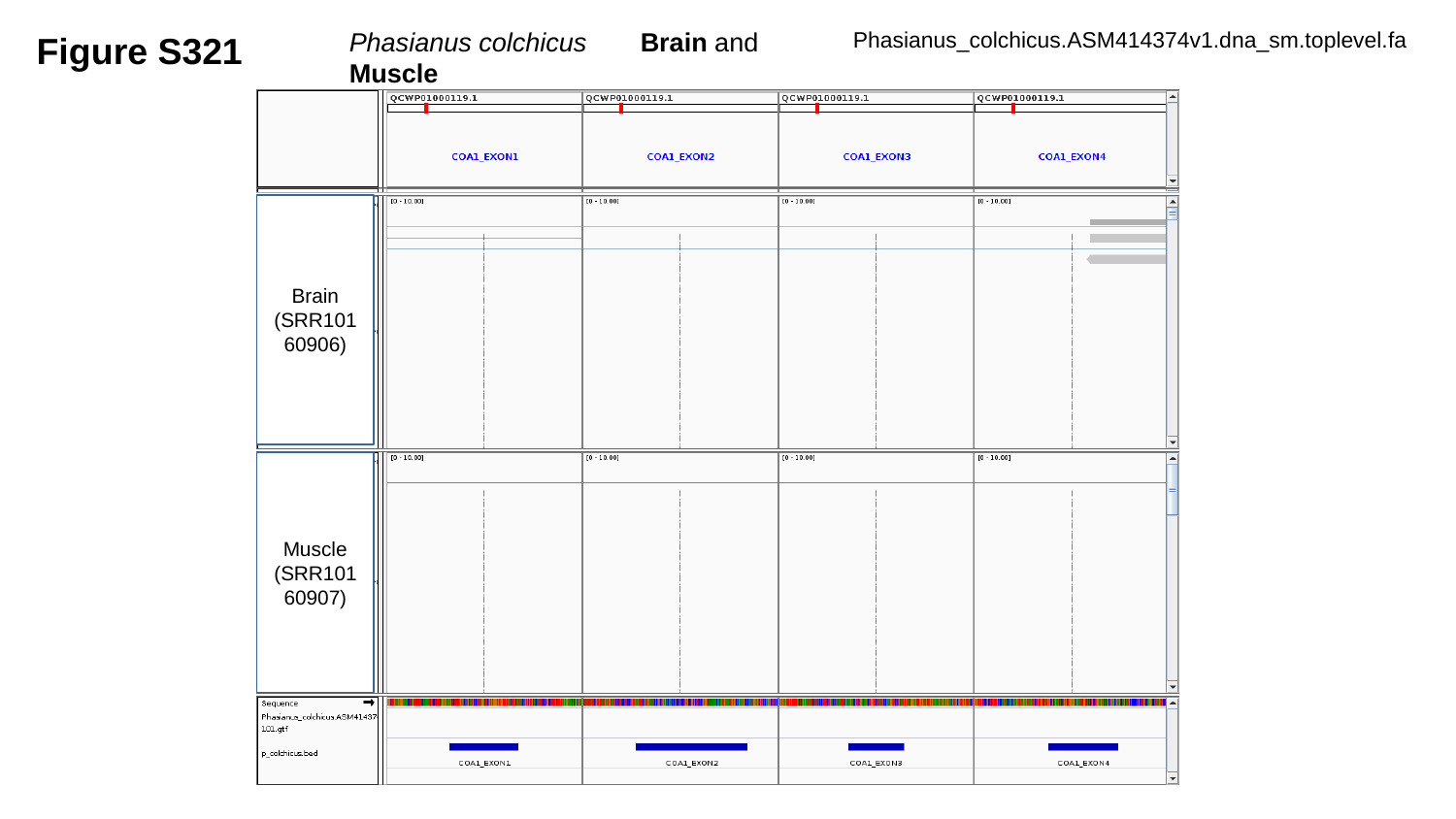

Figure S321
Phasianus colchicus	Brain and Muscle
Phasianus_colchicus.ASM414374v1.dna_sm.toplevel.fa
Brain
(SRR10160906)
Muscle
(SRR10160907)

### Slide 322
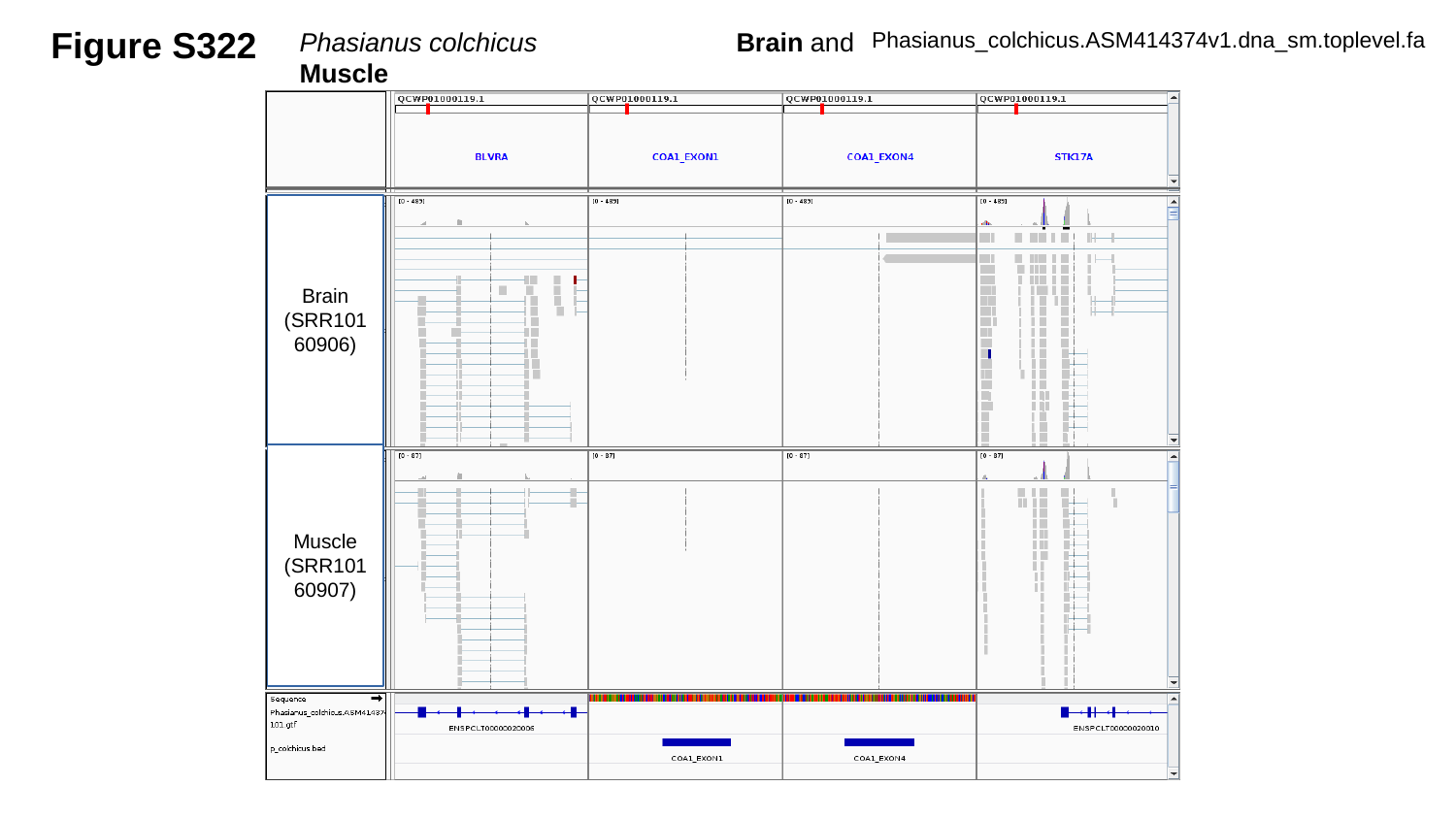

Figure S322
Phasianus colchicus		Brain and Muscle
Phasianus_colchicus.ASM414374v1.dna_sm.toplevel.fa
Brain
(SRR10160906)
Muscle
(SRR10160907)

### Slide 323
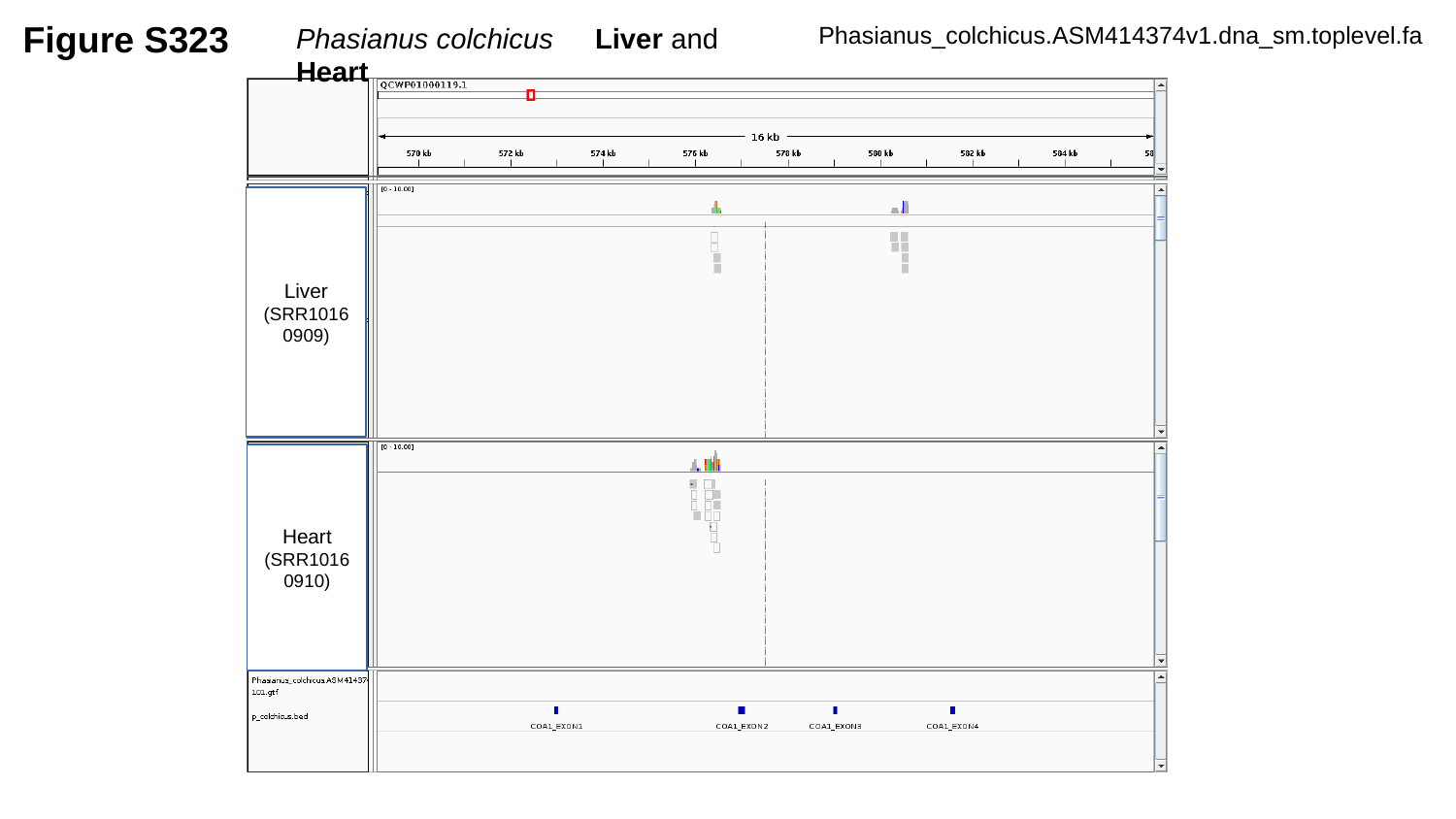

Figure S323
Phasianus colchicus	 Liver and Heart
Phasianus_colchicus.ASM414374v1.dna_sm.toplevel.fa
Liver
(SRR10160909)
Heart
(SRR10160910)

### Slide 324
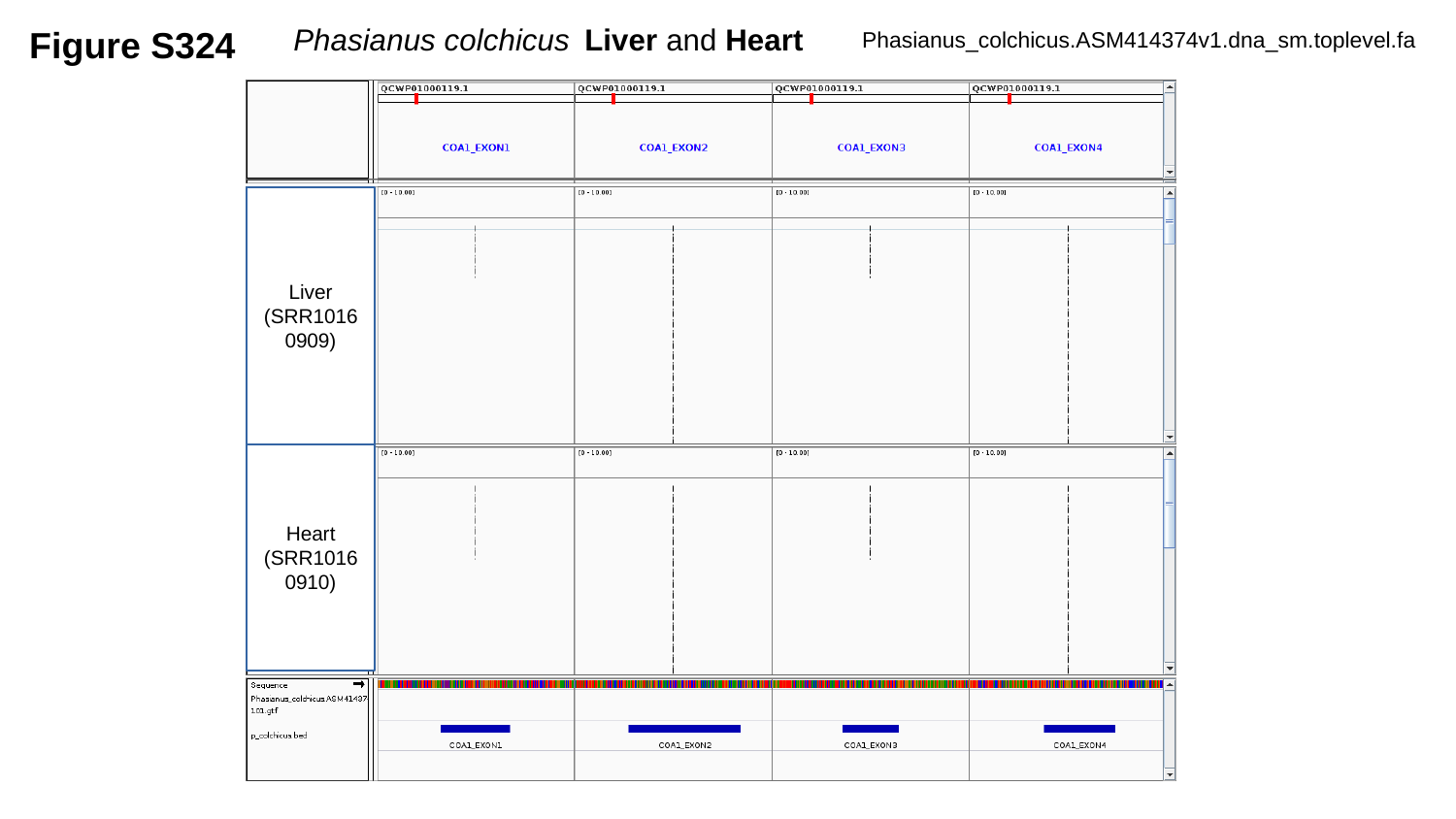

Figure S324
Phasianus colchicus	Liver and Heart
Phasianus_colchicus.ASM414374v1.dna_sm.toplevel.fa
Liver
(SRR10160909)
Heart
(SRR10160910)

### Slide 325
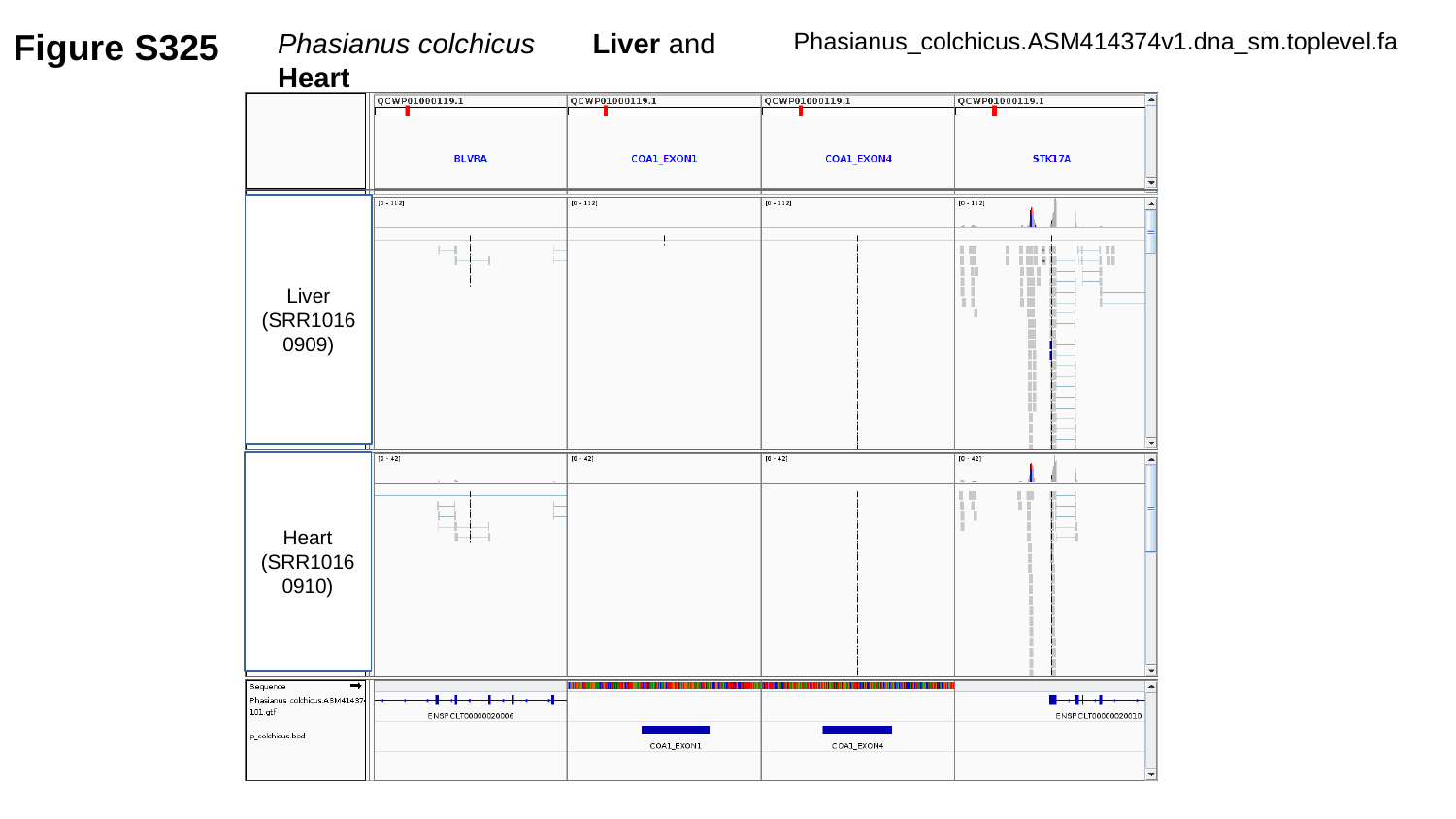

Figure S325
Phasianus colchicus	 Liver and Heart
Phasianus_colchicus.ASM414374v1.dna_sm.toplevel.fa
Liver
(SRR10160909)
Heart
(SRR10160910)

### Slide 326
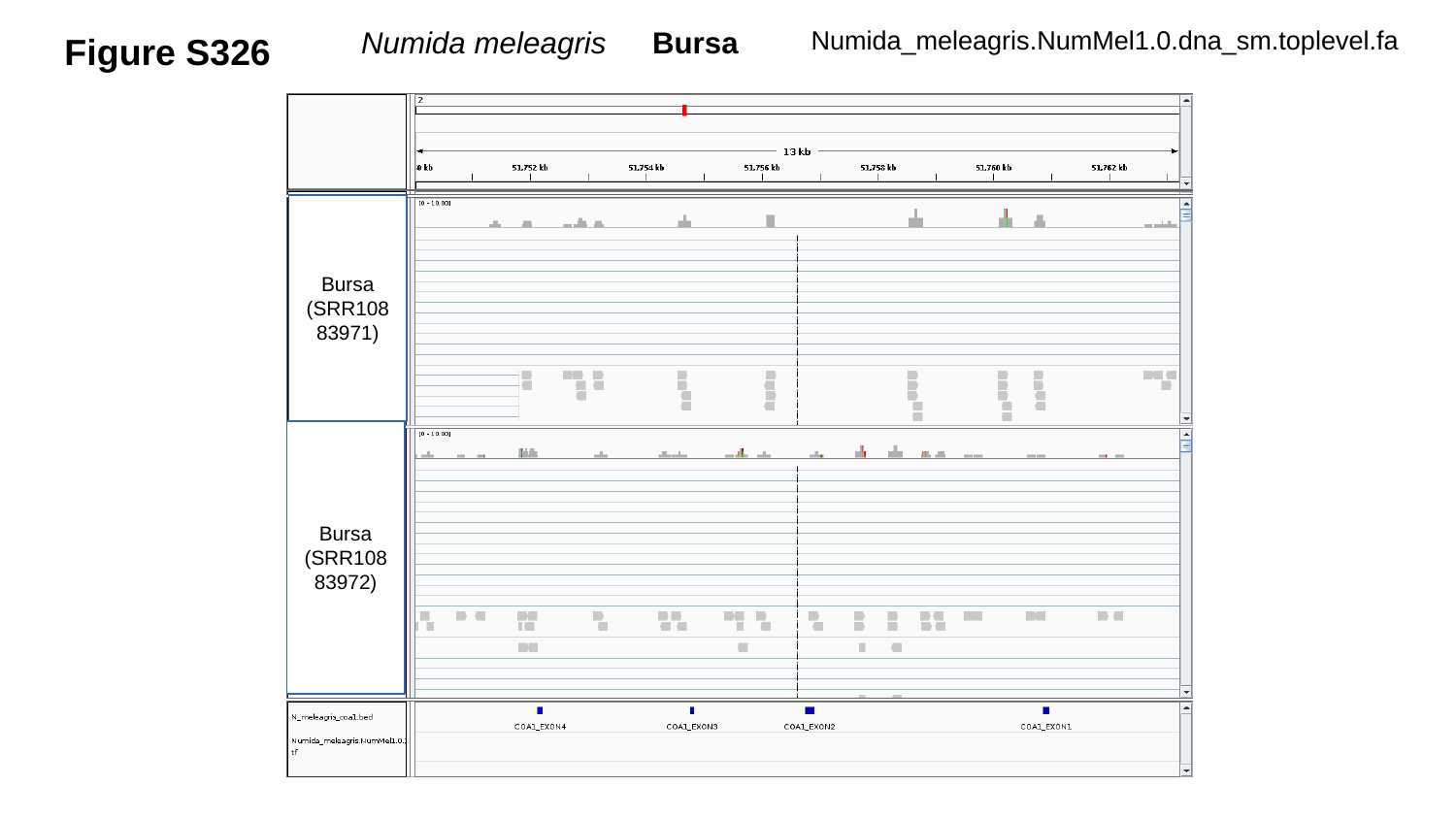

Figure S326
Numida_meleagris.NumMel1.0.dna_sm.toplevel.fa
Numida meleagris	Bursa
Bursa
(SRR10883971)
Bursa
(SRR10883972)

### Slide 327
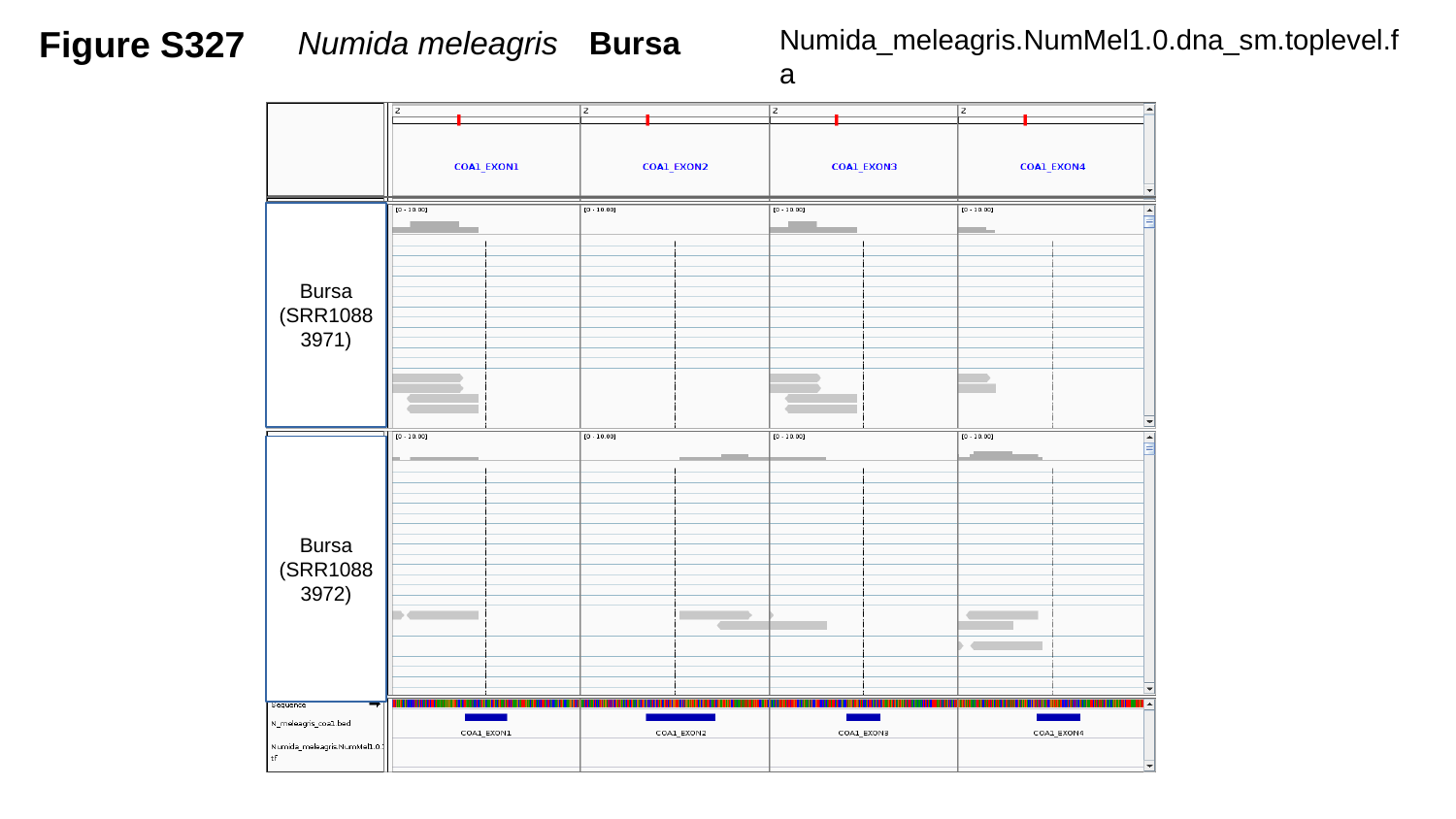

Figure S327
Numida meleagris	Bursa
Numida_meleagris.NumMel1.0.dna_sm.toplevel.fa
Bursa
(SRR10883971)
Bursa
(SRR10883972)

### Slide 328
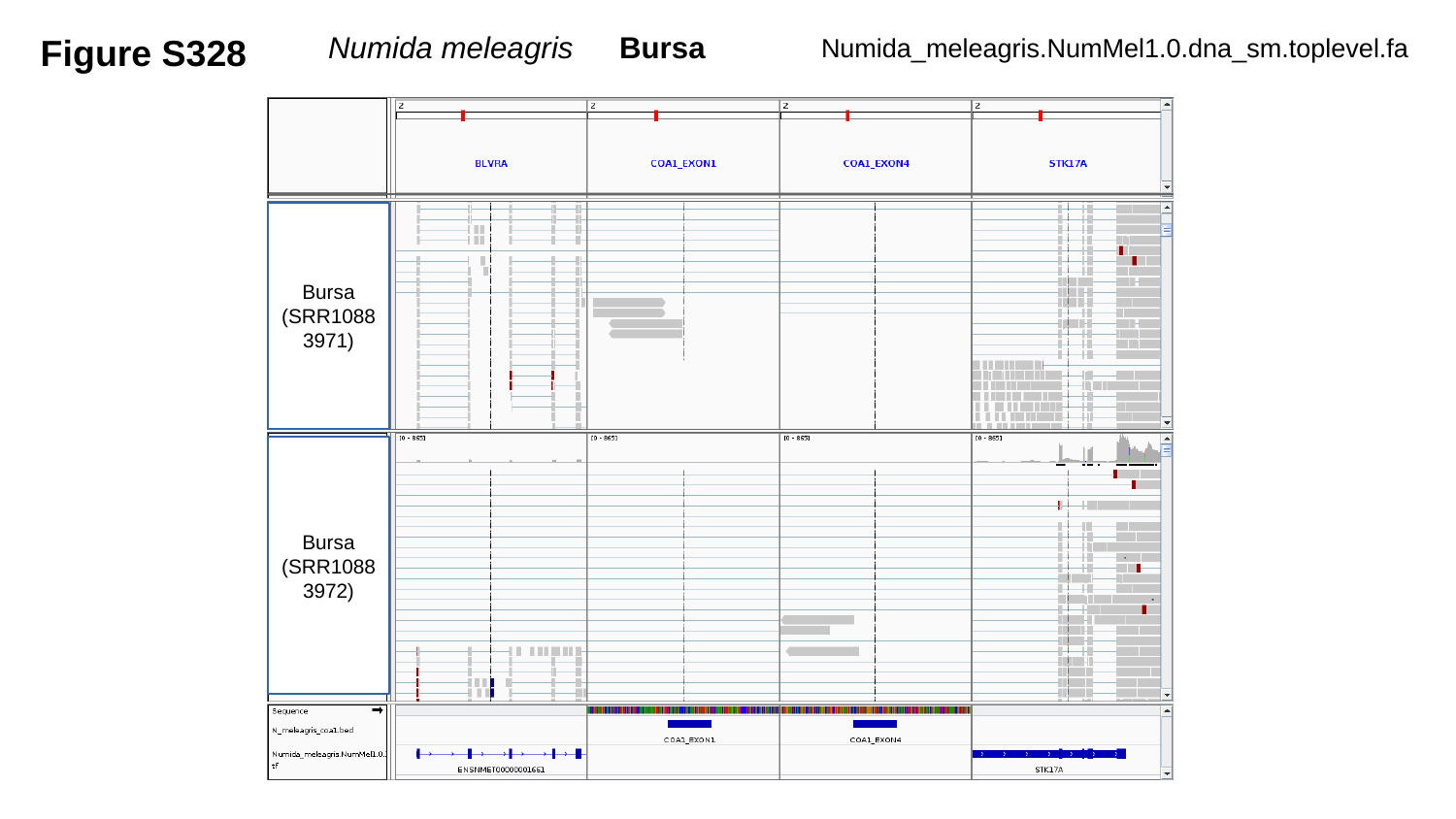

Figure S328
Numida meleagris	Bursa
Numida_meleagris.NumMel1.0.dna_sm.toplevel.fa
Bursa
(SRR10883971)
Bursa
(SRR10883972)

### Slide 329
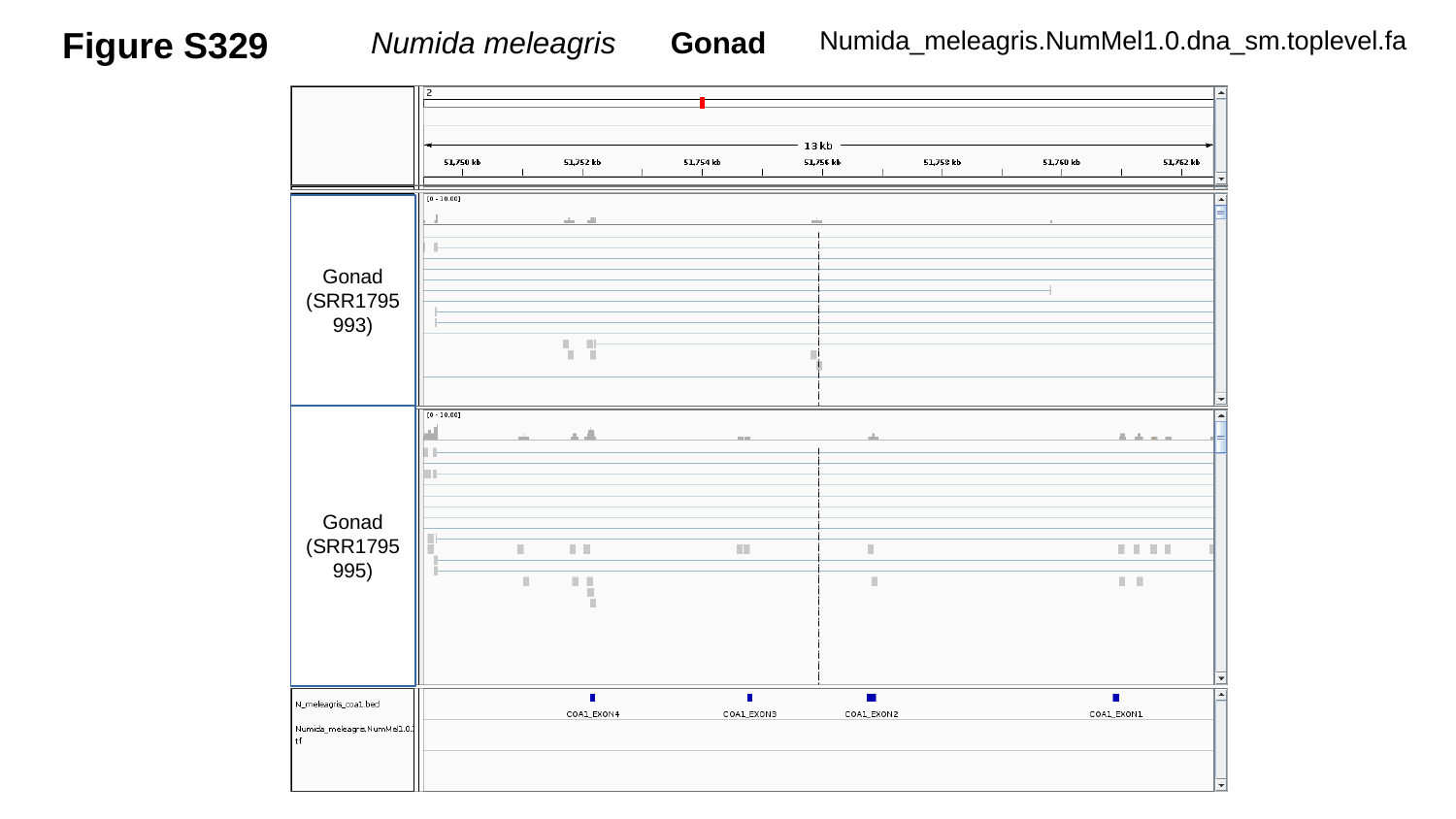

Figure S329
Numida meleagris	 Gonad
Numida_meleagris.NumMel1.0.dna_sm.toplevel.fa
Gonad
(SRR1795993)
Gonad
(SRR1795995)

### Slide 330
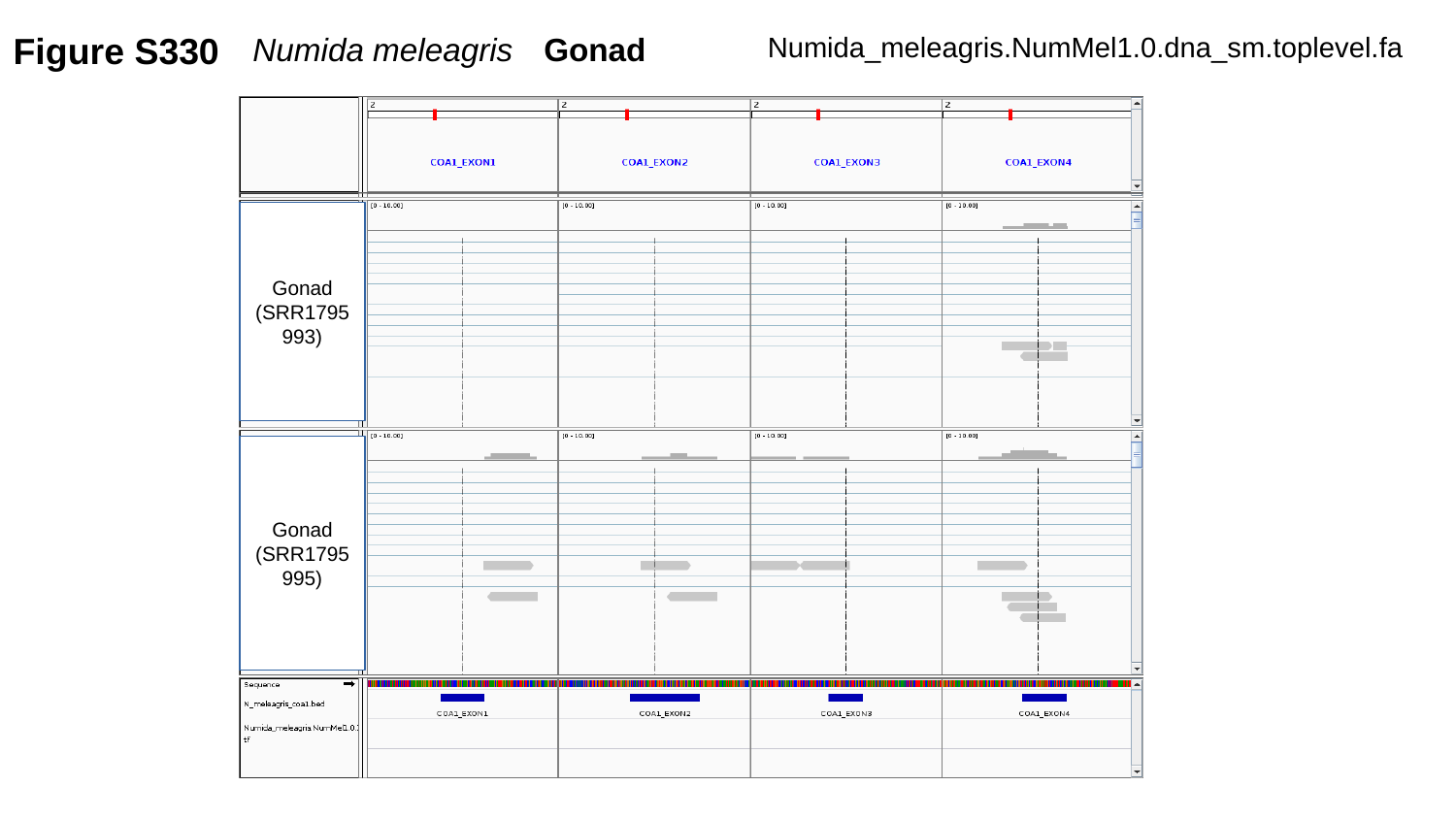

Figure S330
Numida meleagris	Gonad
Numida_meleagris.NumMel1.0.dna_sm.toplevel.fa
Gonad
(SRR1795993)
Gonad
(SRR1795995)

### Slide 331
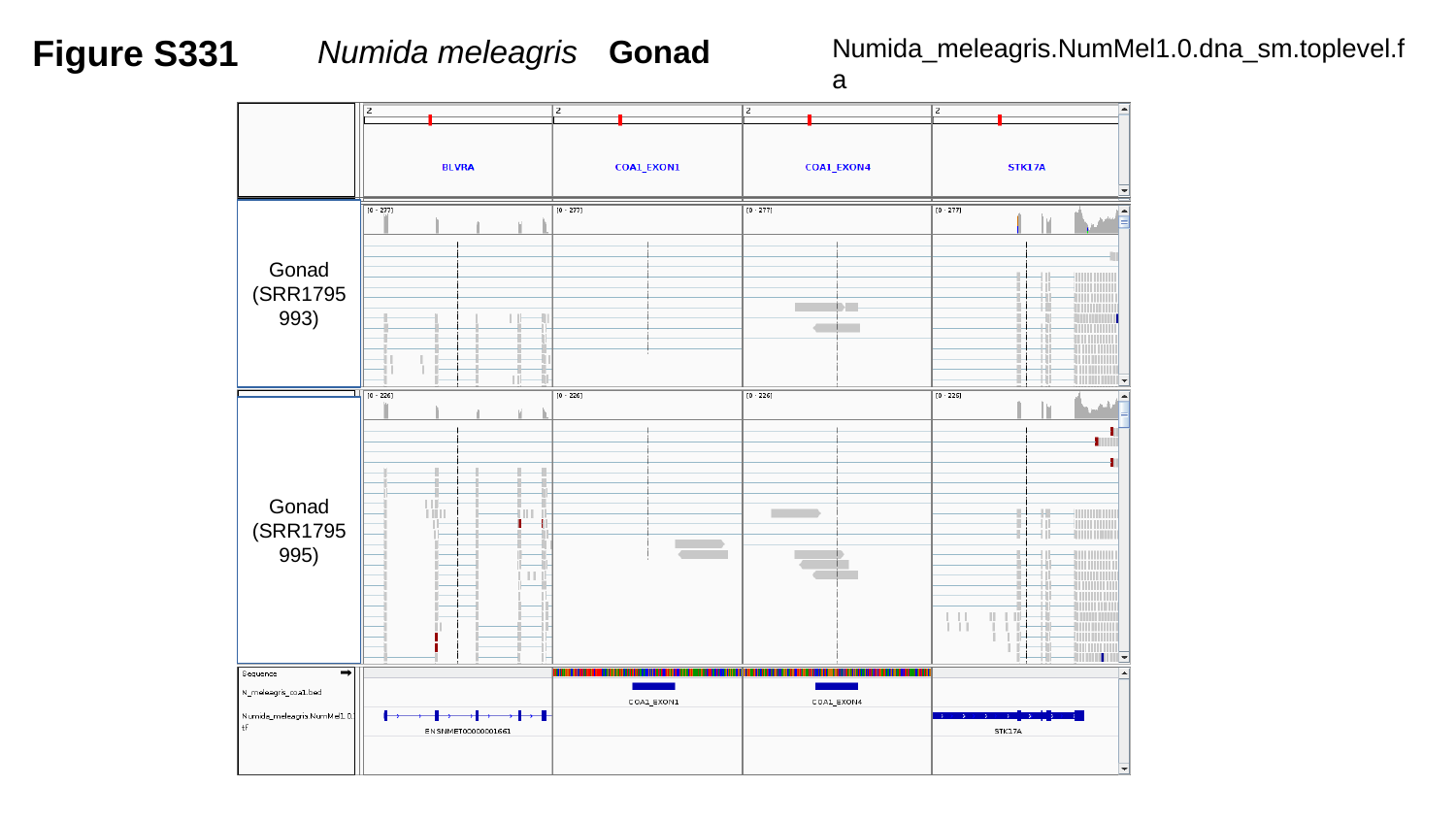

Figure S331
Numida meleagris	Gonad
Numida_meleagris.NumMel1.0.dna_sm.toplevel.fa
Gonad
(SRR1795993)
Gonad
(SRR1795995)

### Slide 332
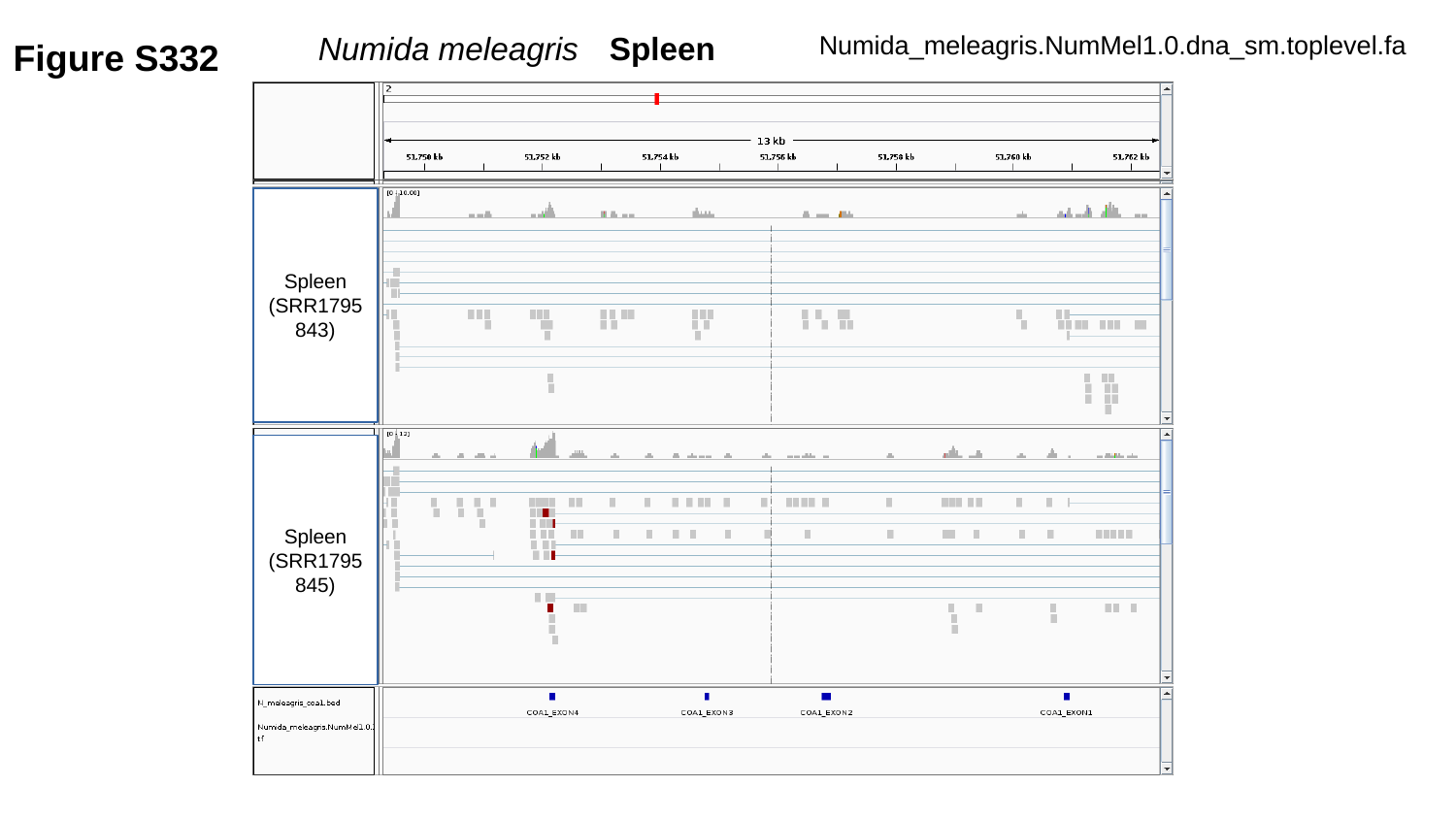

Figure S332
Numida meleagris	Spleen
Numida_meleagris.NumMel1.0.dna_sm.toplevel.fa
Spleen
(SRR1795843)
Spleen
(SRR1795845)

### Slide 333
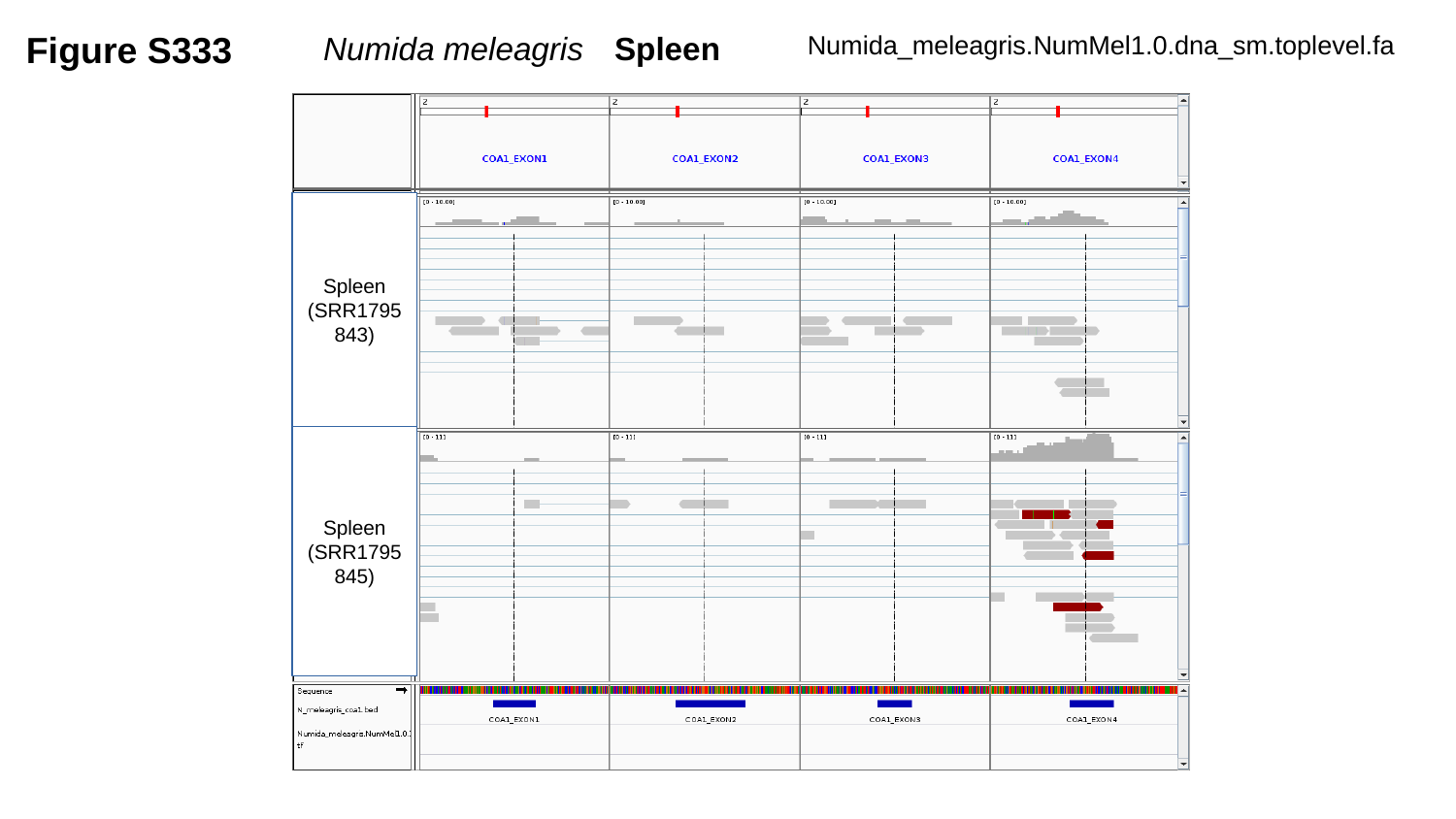

Figure S333
Numida meleagris	Spleen
Numida_meleagris.NumMel1.0.dna_sm.toplevel.fa
Spleen
(SRR1795843)
Spleen
(SRR1795845)

### Slide 334
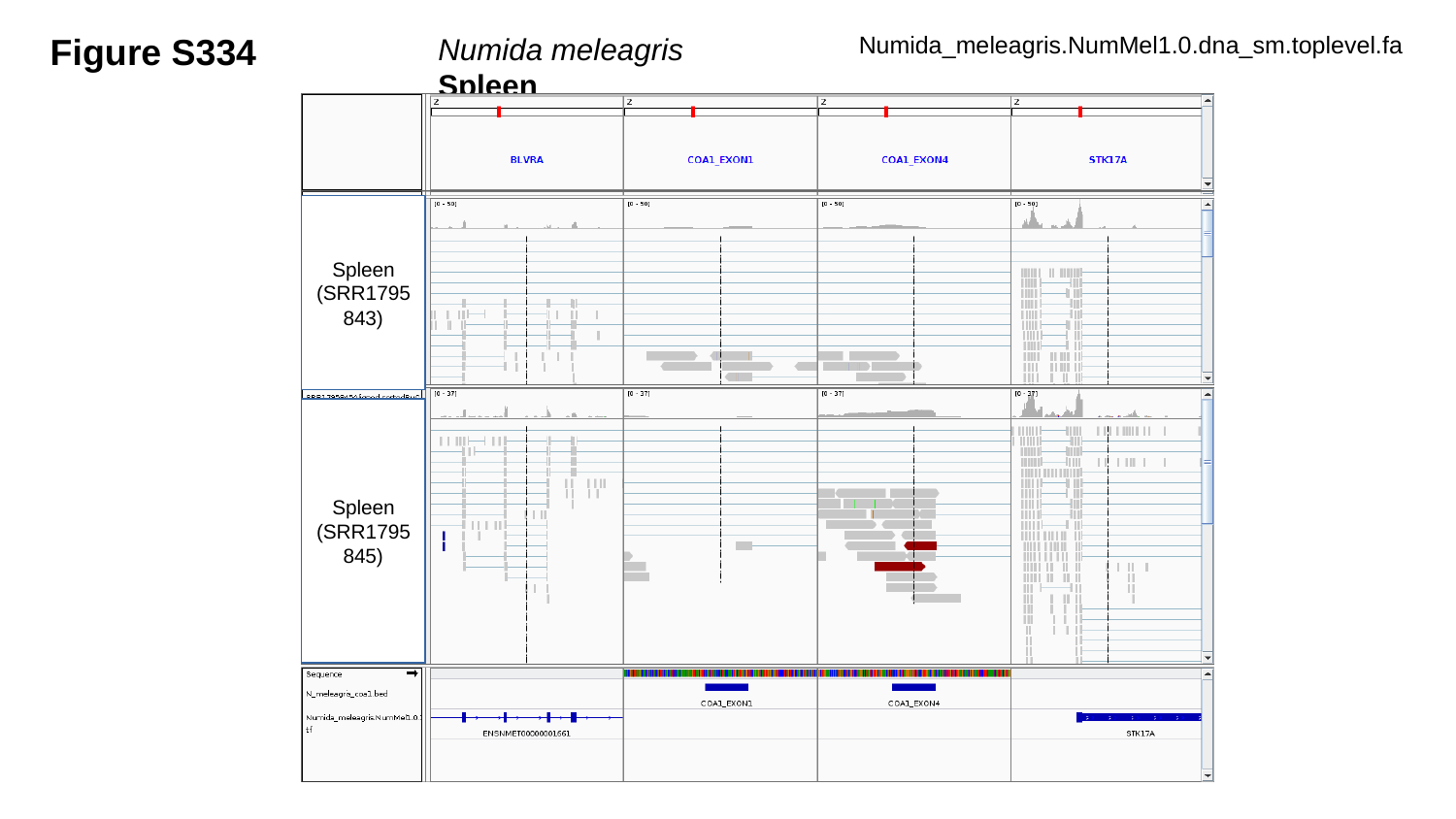

Figure S334
Numida meleagris	Spleen
Numida_meleagris.NumMel1.0.dna_sm.toplevel.fa
Spleen
(SRR1795843)
Spleen
(SRR1795845)

### Slide 335
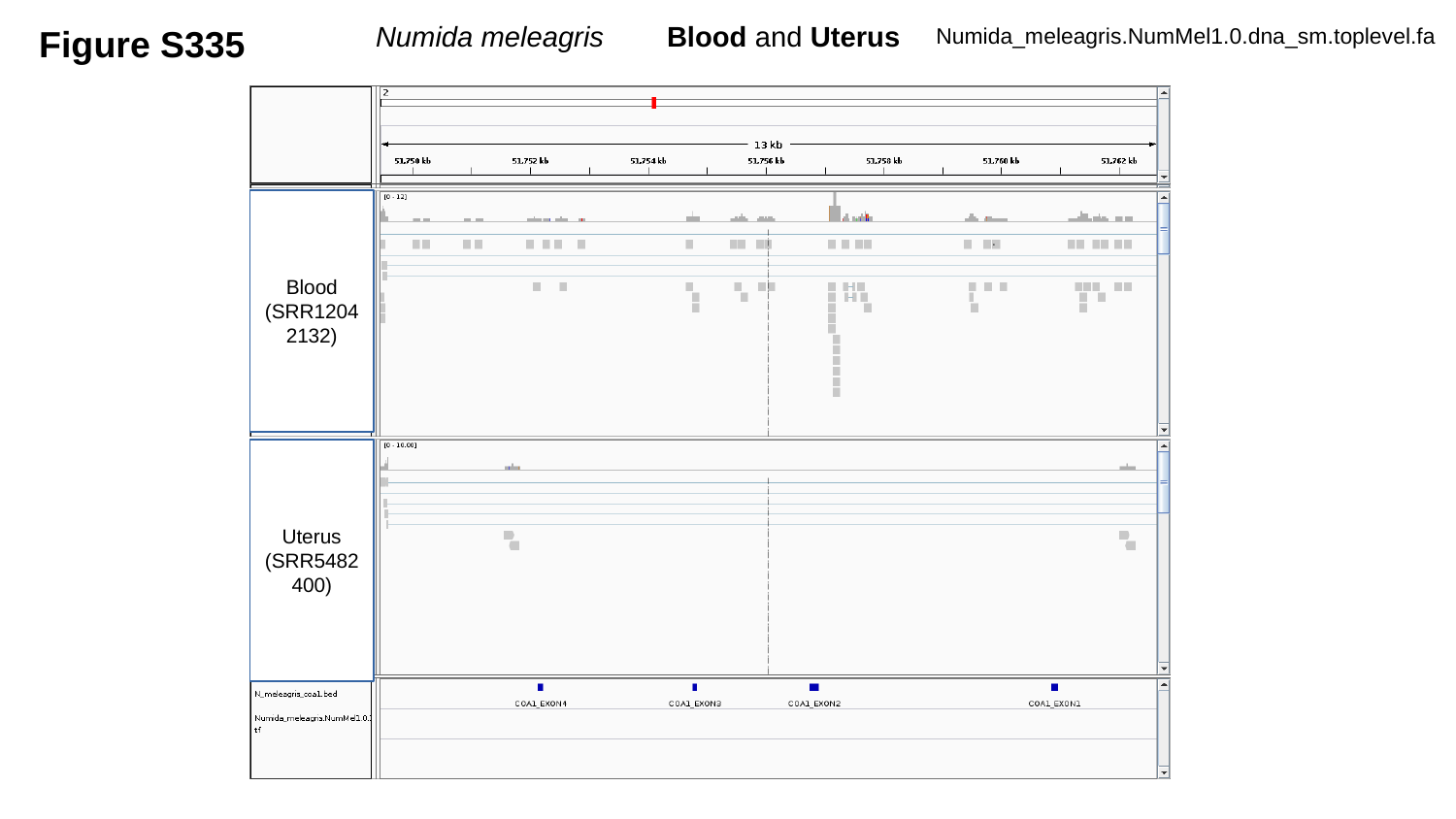

Figure S335
Numida meleagris	Blood and Uterus
Numida_meleagris.NumMel1.0.dna_sm.toplevel.fa
Blood
(SRR12042132)
Uterus
(SRR5482400)

### Slide 336
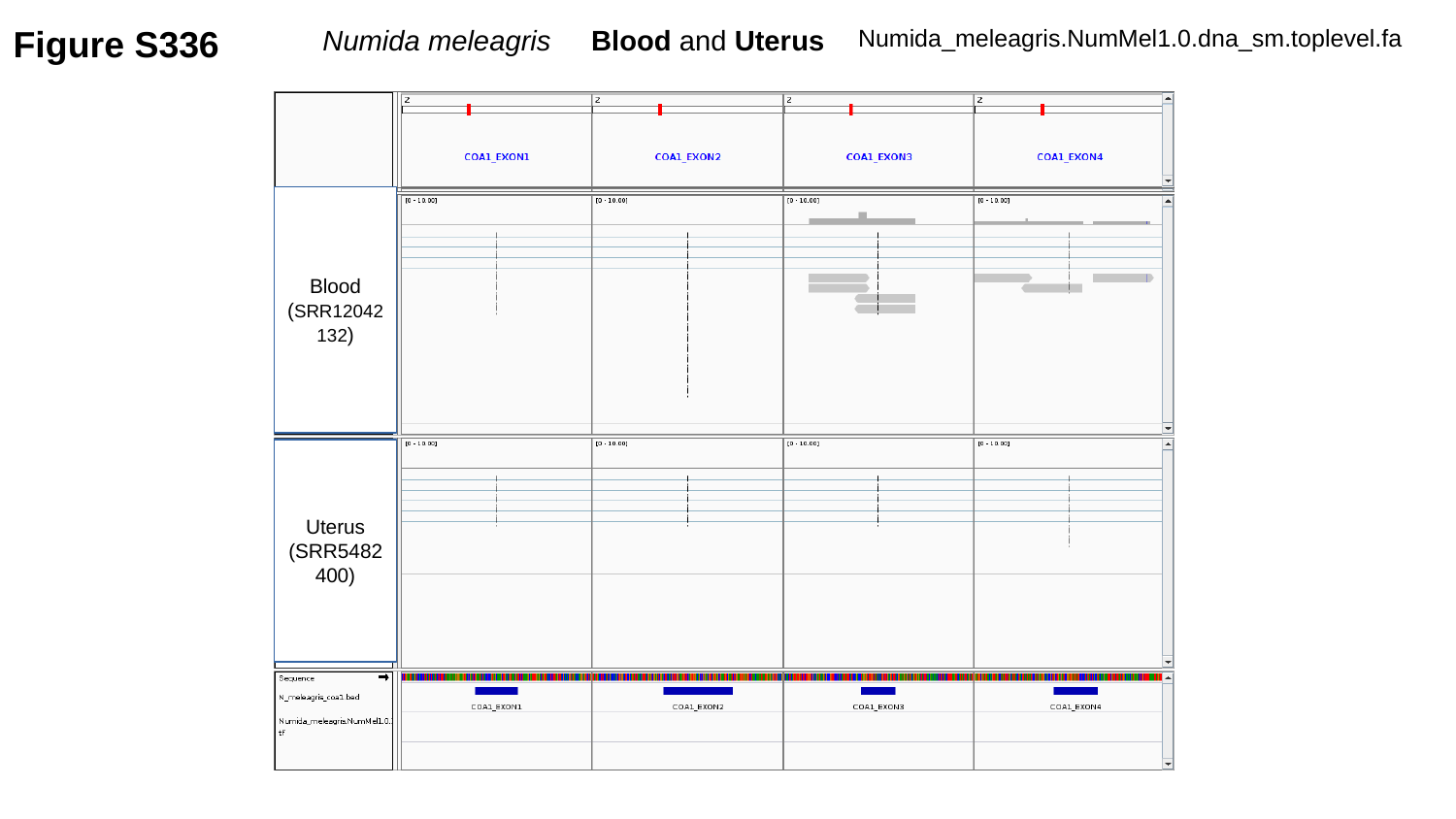

Figure S336
Numida meleagris Blood and Uterus
Numida_meleagris.NumMel1.0.dna_sm.toplevel.fa
Blood
(SRR12042132)
Uterus
(SRR5482400)

### Slide 337
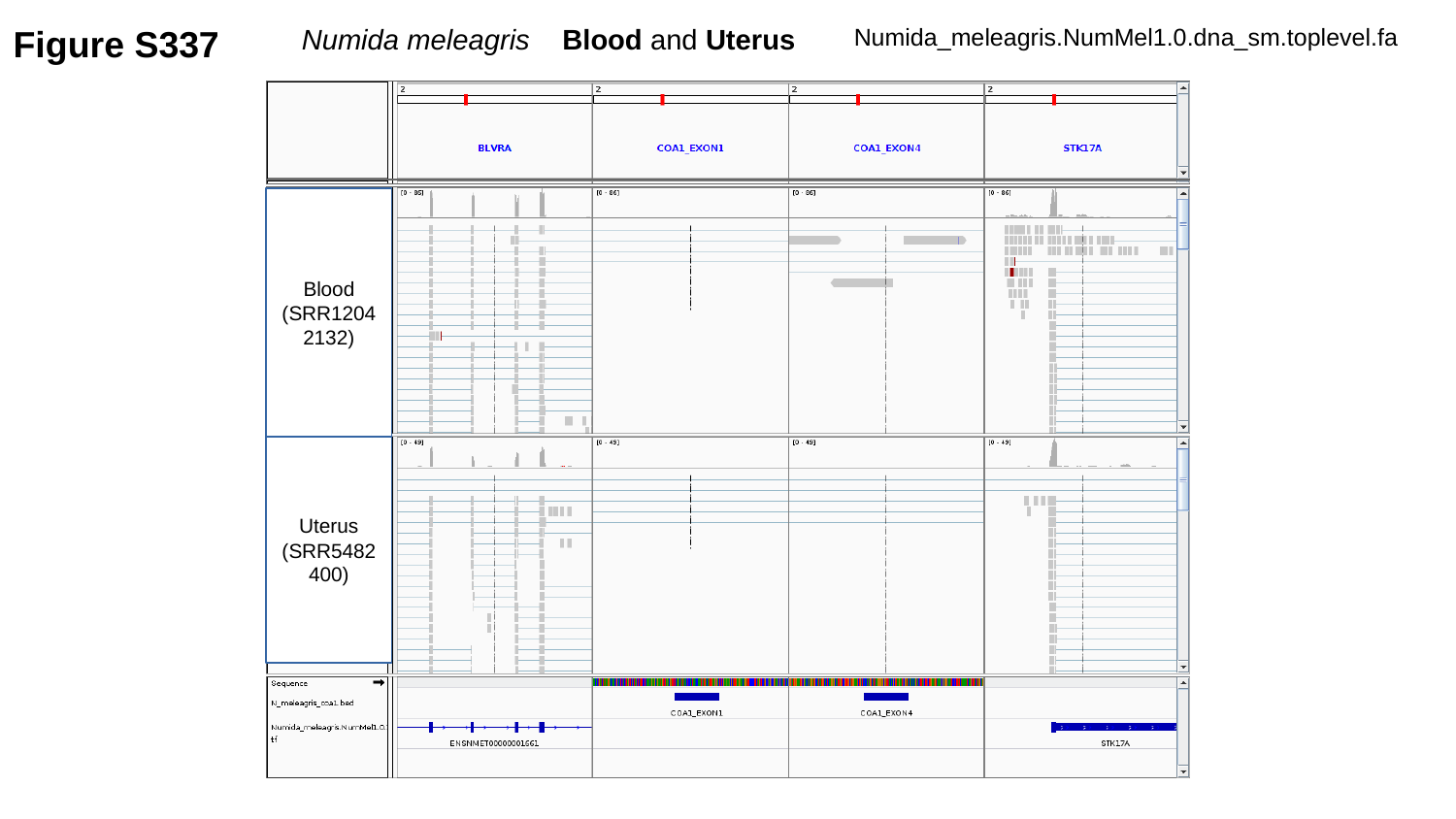

Figure S337
Numida meleagris Blood and Uterus
Numida_meleagris.NumMel1.0.dna_sm.toplevel.fa
Blood
(SRR12042132)
Uterus
(SRR5482400)

### Slide 338
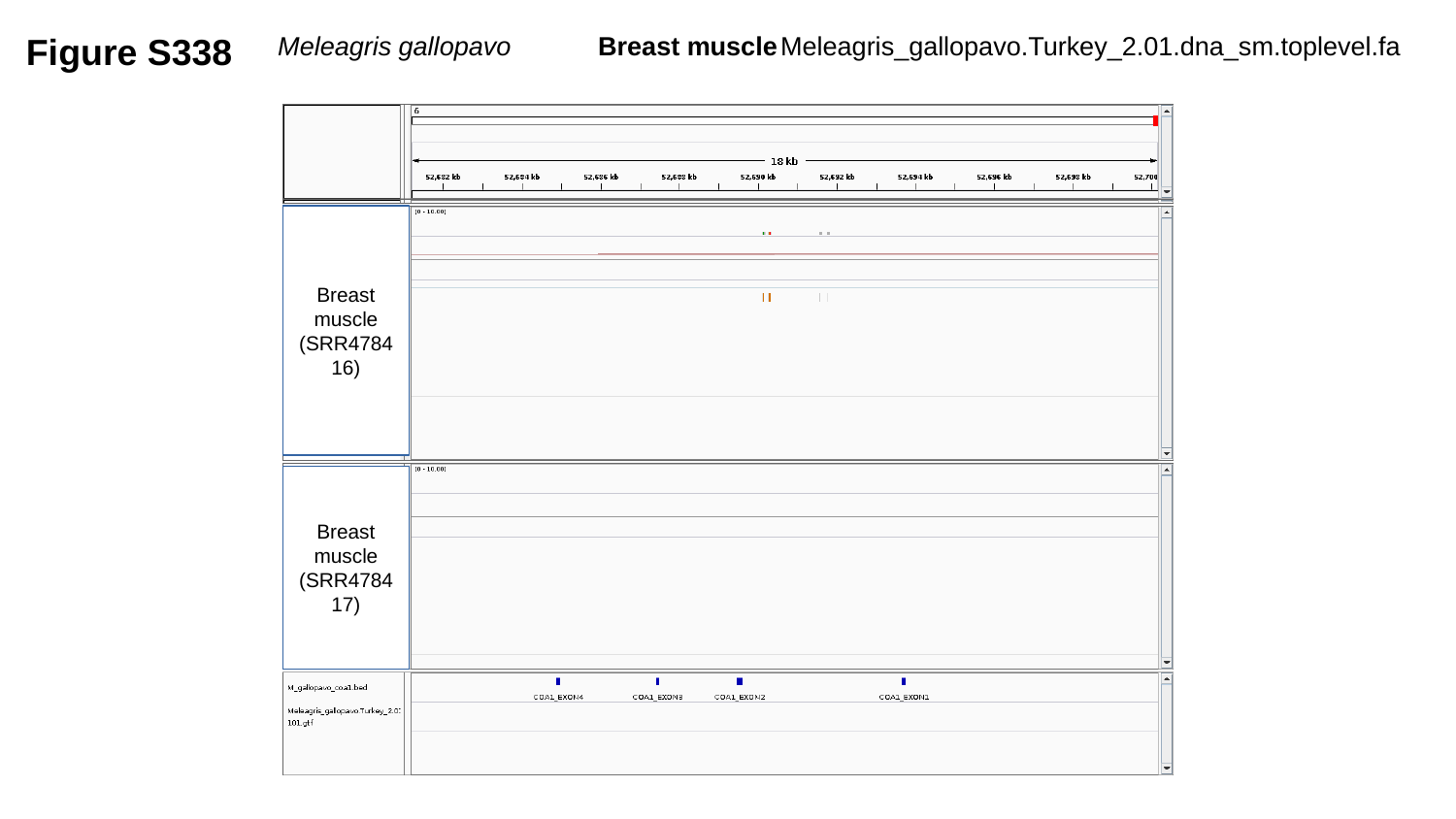

Figure S338
Meleagris gallopavo	 Breast muscle
Meleagris_gallopavo.Turkey_2.01.dna_sm.toplevel.fa
Breast muscle
(SRR478416)
Breast muscle
(SRR478417)

### Slide 339
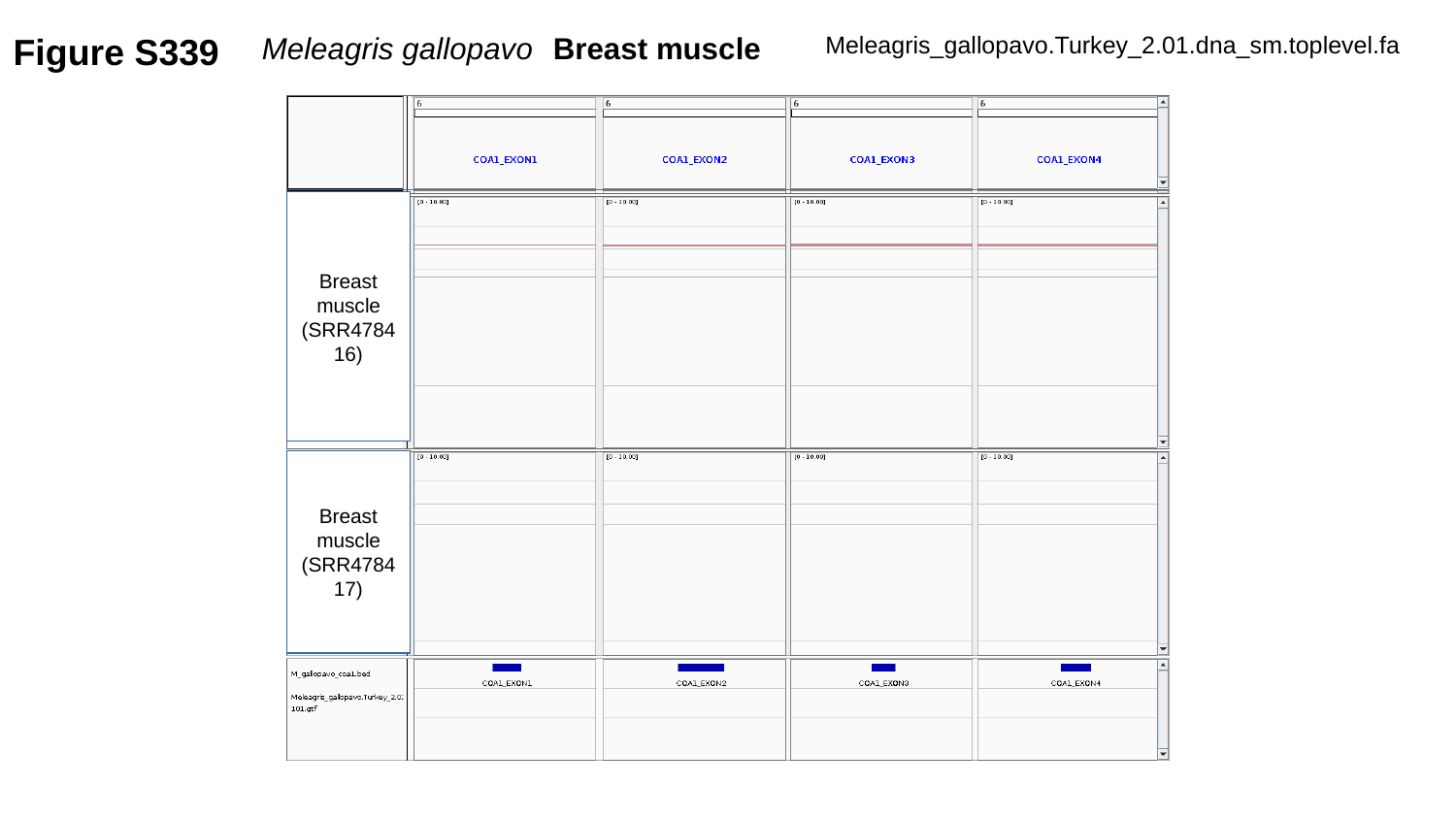

Figure S339
Meleagris gallopavo 	Breast muscle
Meleagris_gallopavo.Turkey_2.01.dna_sm.toplevel.fa
Breast muscle
(SRR478416)
Breast muscle
(SRR478417)

### Slide 340
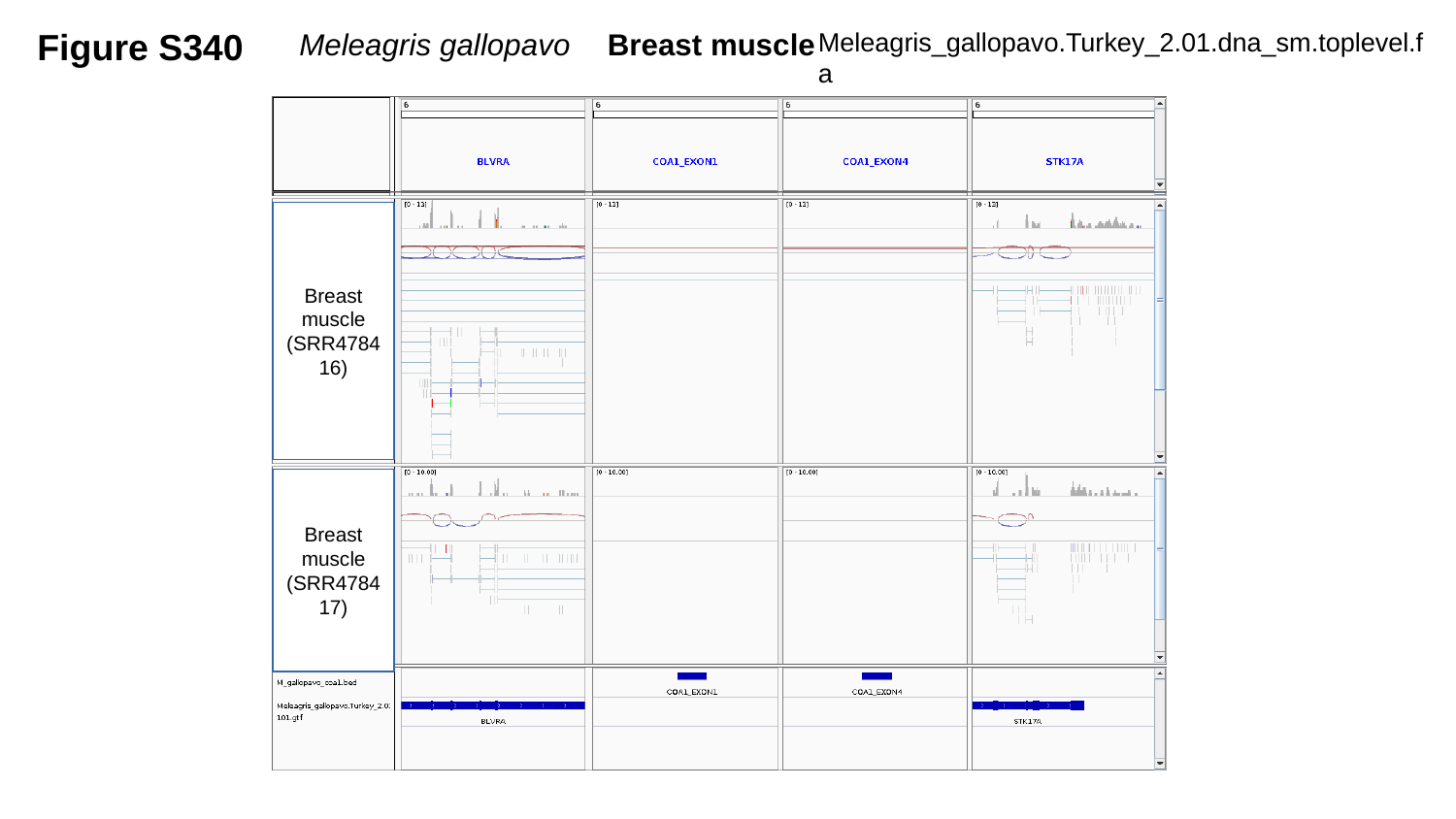

Figure S340
Meleagris gallopavo	 Breast muscle
Meleagris_gallopavo.Turkey_2.01.dna_sm.toplevel.fa
Breast muscle
(SRR478416)
Breast muscle
(SRR478417)

### Slide 341
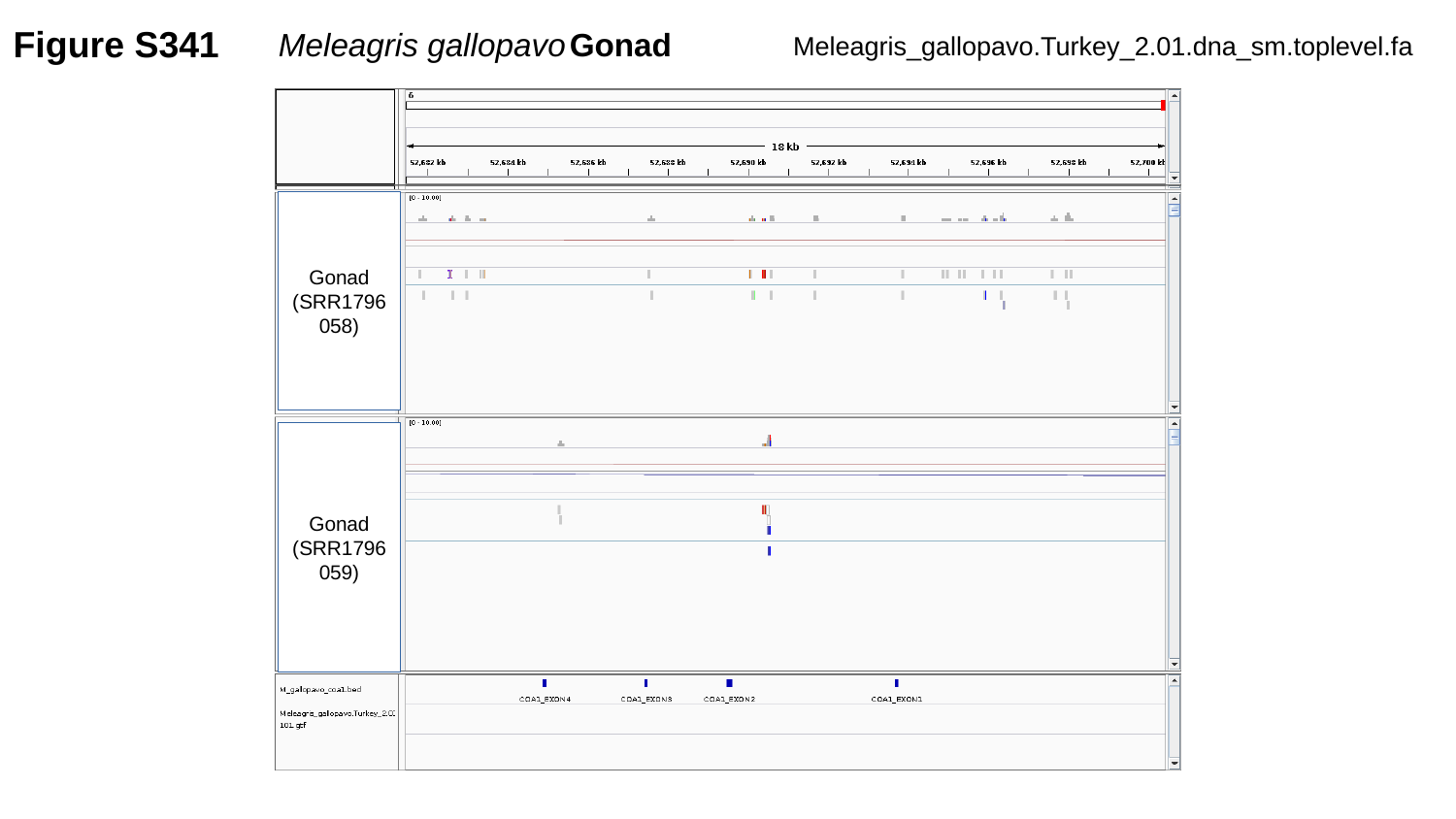

Figure S341
Meleagris gallopavo	Gonad
Meleagris_gallopavo.Turkey_2.01.dna_sm.toplevel.fa
Gonad
(SRR1796058)
Gonad
(SRR1796059)

### Slide 342
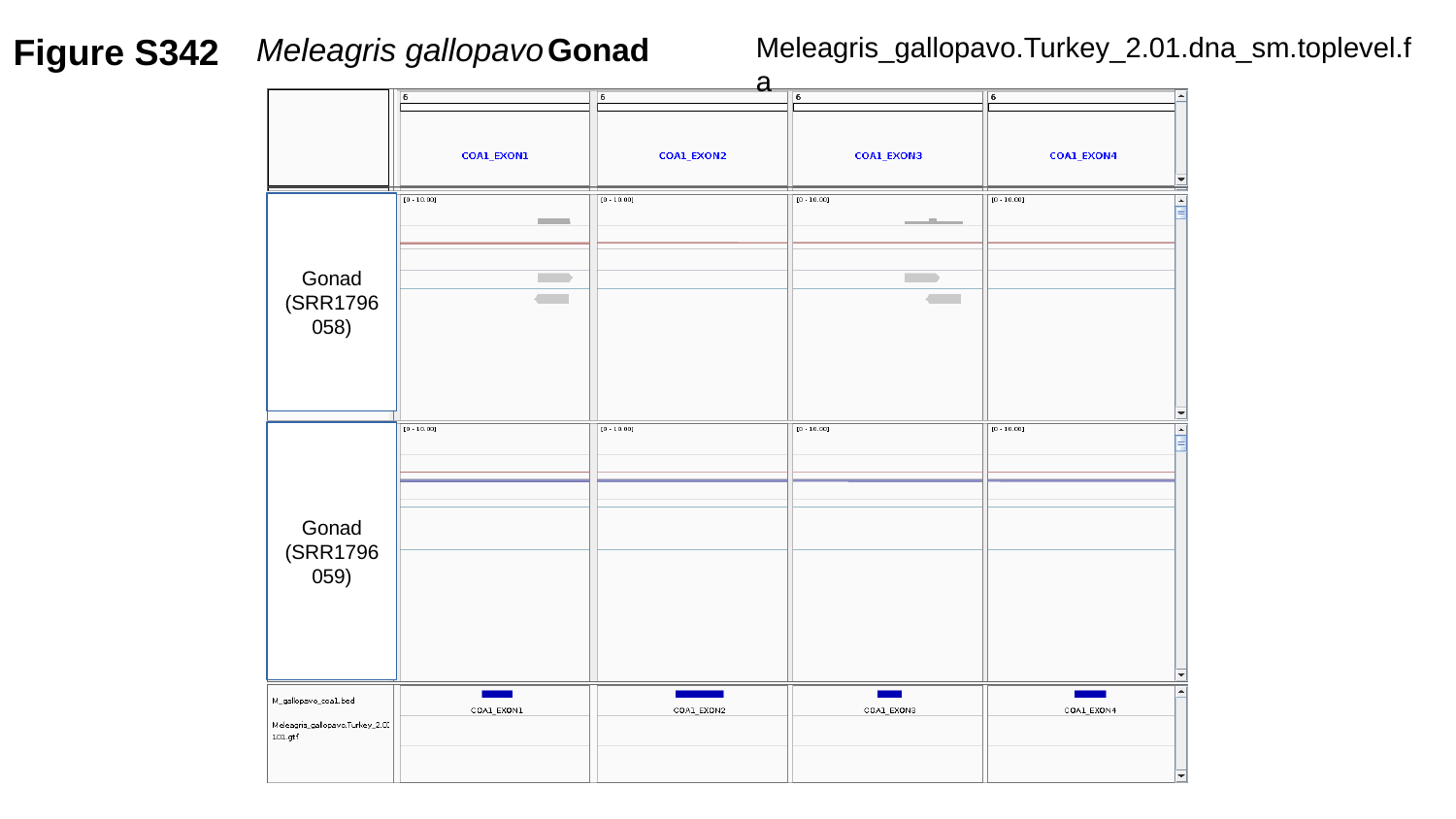

Figure S342
Meleagris gallopavo	Gonad
Meleagris_gallopavo.Turkey_2.01.dna_sm.toplevel.fa
Gonad
(SRR1796058)
Gonad
(SRR1796059)

### Slide 343
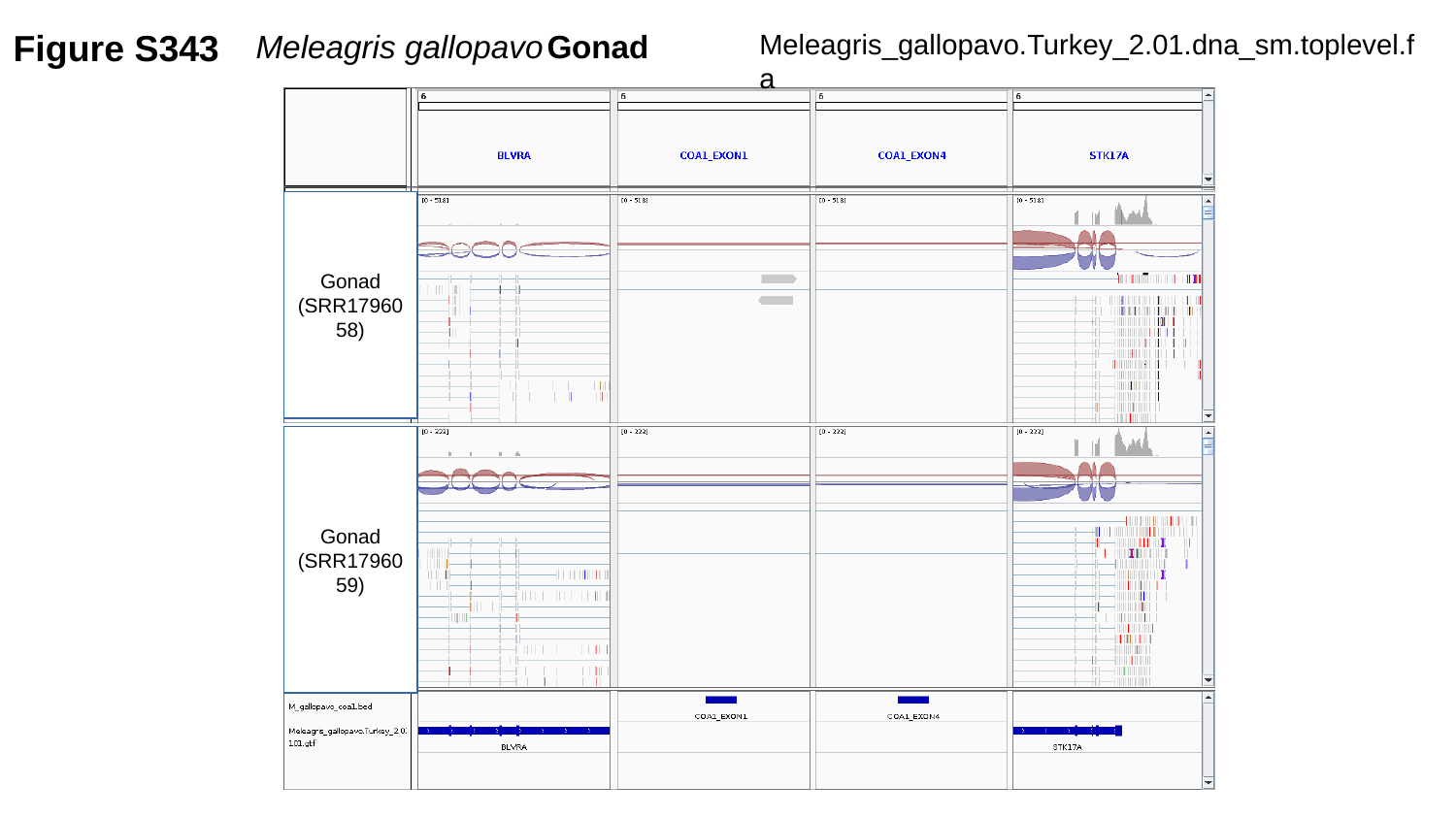

Figure S343
Meleagris gallopavo	Gonad
Meleagris_gallopavo.Turkey_2.01.dna_sm.toplevel.fa
Gonad
(SRR1796058)
Gonad
(SRR1796059)

### Slide 344
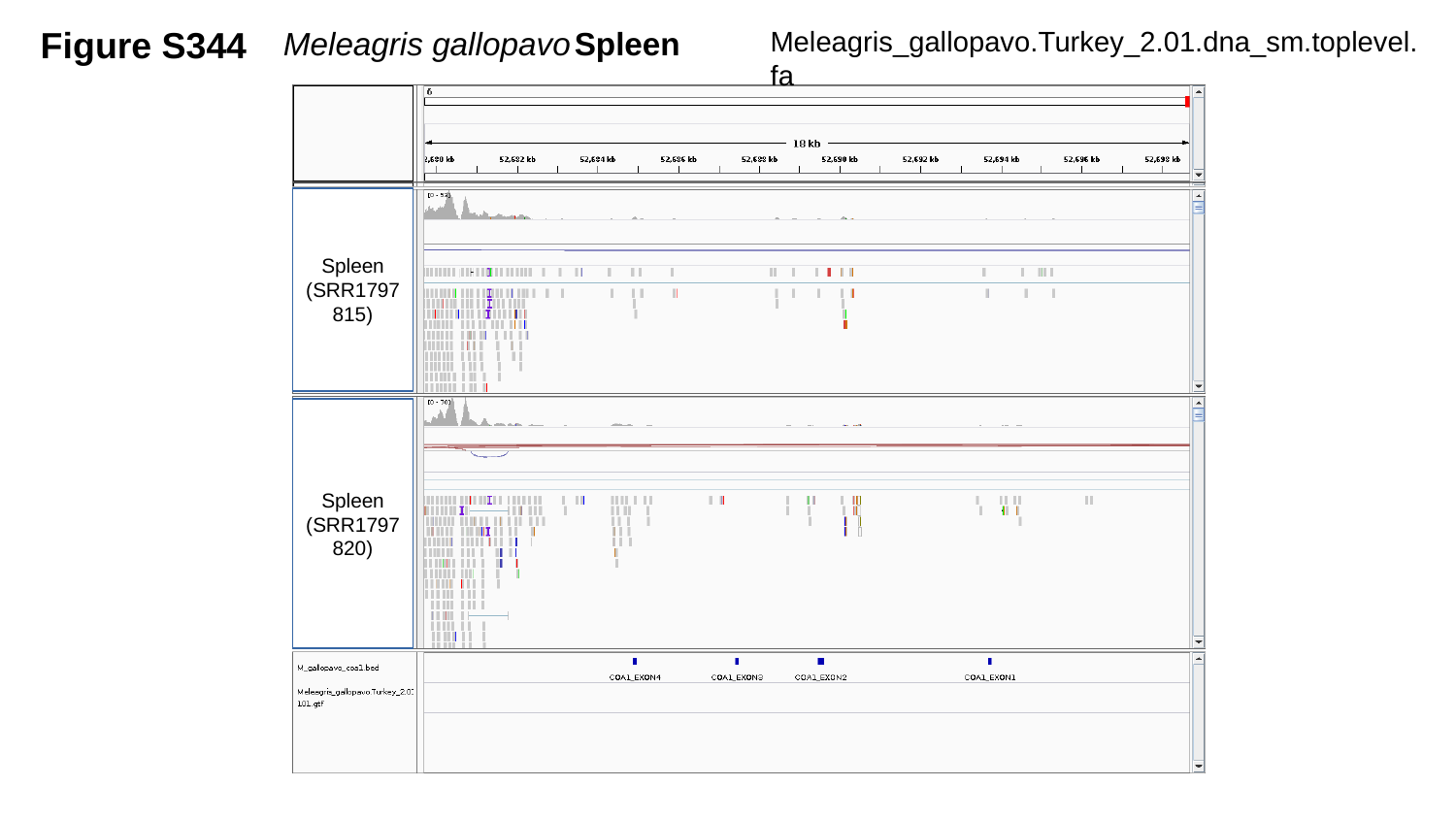

Figure S344
Meleagris gallopavo	Spleen
Meleagris_gallopavo.Turkey_2.01.dna_sm.toplevel.fa
Spleen
(SRR1797815)
Spleen
(SRR1797820)

### Slide 345
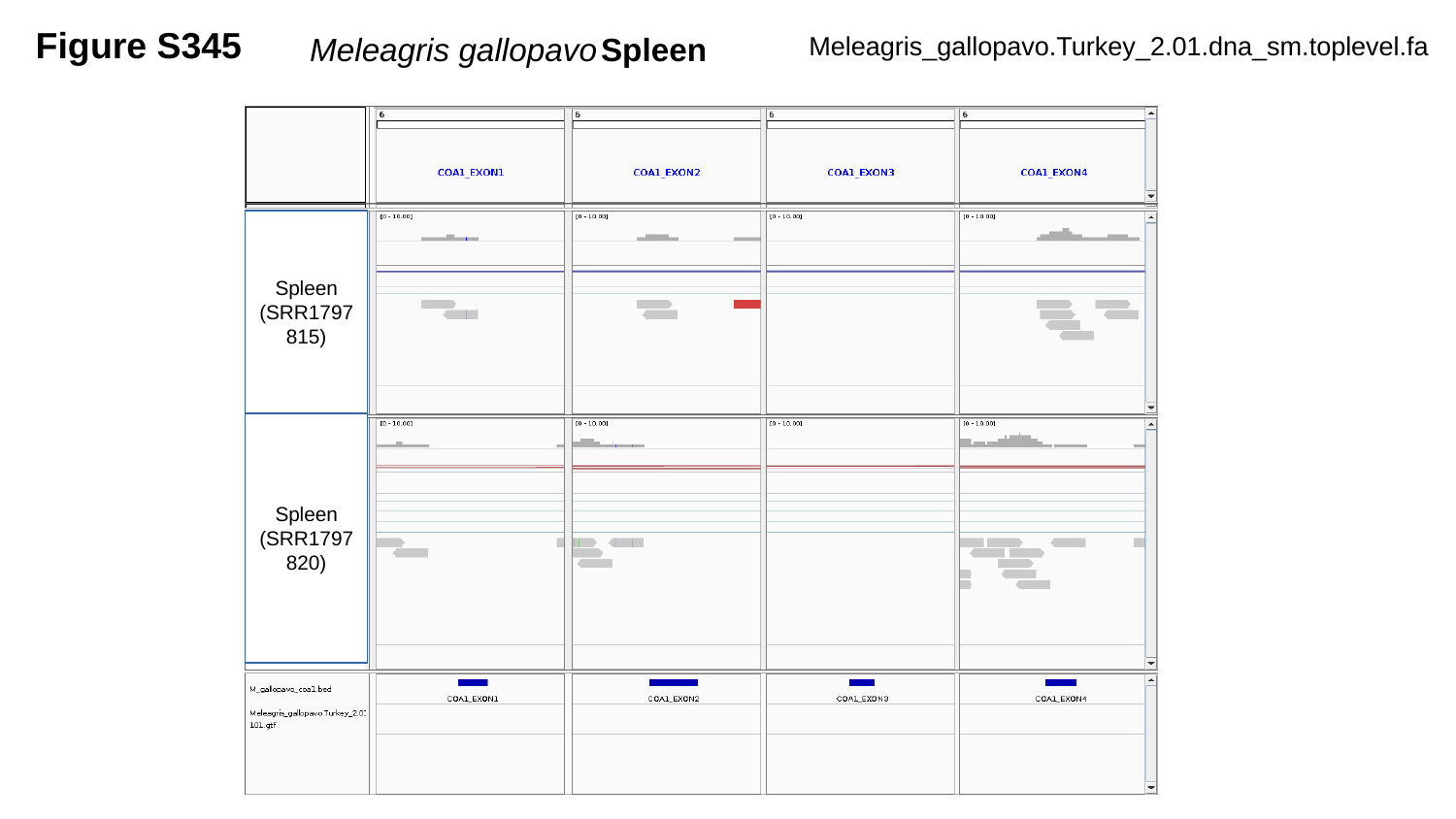

Figure S345
Meleagris gallopavo	Spleen
Meleagris_gallopavo.Turkey_2.01.dna_sm.toplevel.fa
Spleen
(SRR1797815)
Spleen
(SRR1797820)

### Slide 346
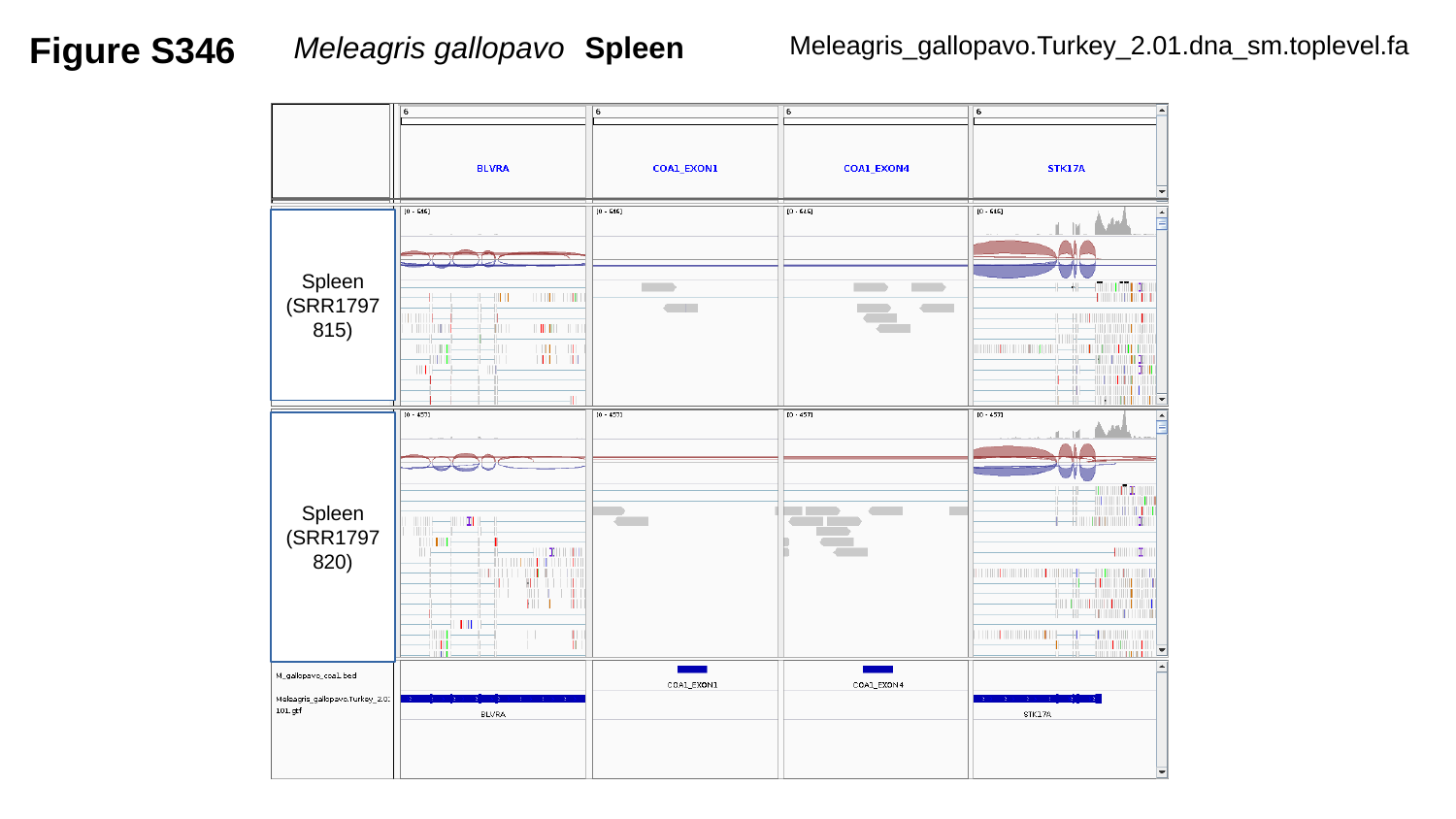

Figure S346
Meleagris gallopavo	Spleen
Meleagris_gallopavo.Turkey_2.01.dna_sm.toplevel.fa
Spleen
(SRR1797815)
Spleen
(SRR1797820)

### Slide 347
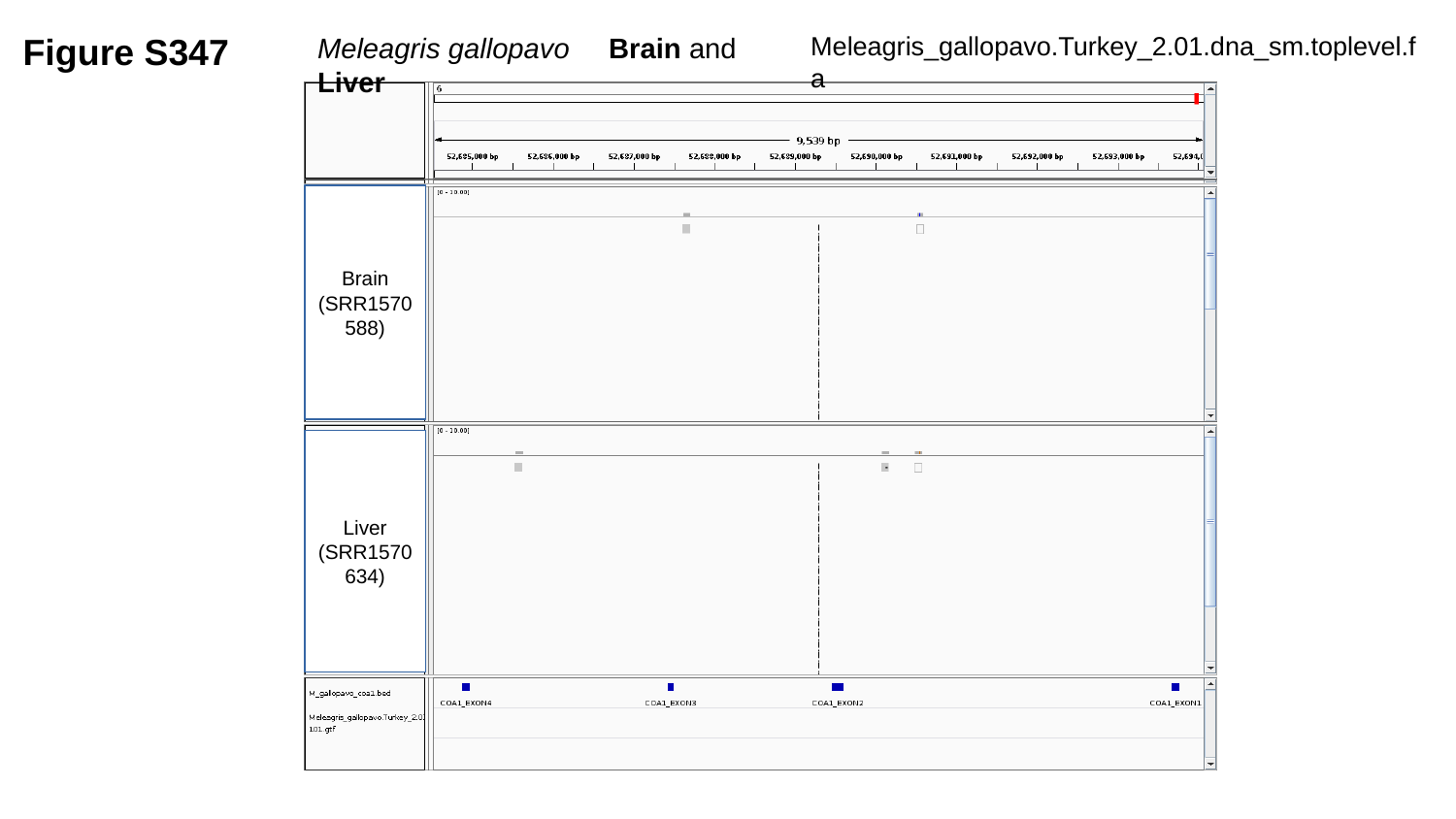

Figure S347
Meleagris gallopavo	Brain and Liver
Meleagris_gallopavo.Turkey_2.01.dna_sm.toplevel.fa
Brain
(SRR1570588)
Liver
(SRR1570634)

### Slide 348
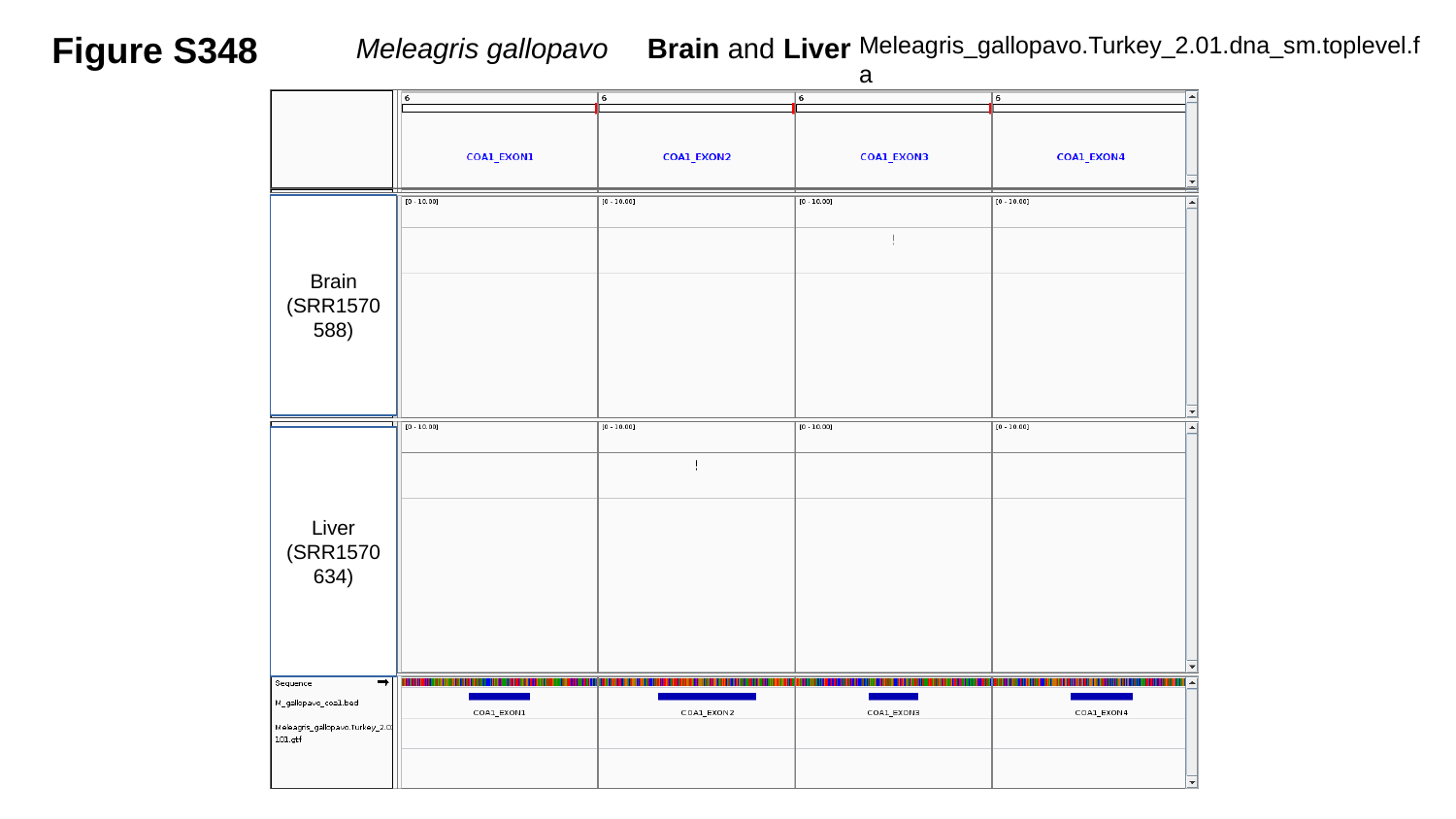

Figure S348
Meleagris gallopavo	Brain and Liver
Meleagris_gallopavo.Turkey_2.01.dna_sm.toplevel.fa
Brain
(SRR1570588)
Liver
(SRR1570634)

### Slide 349
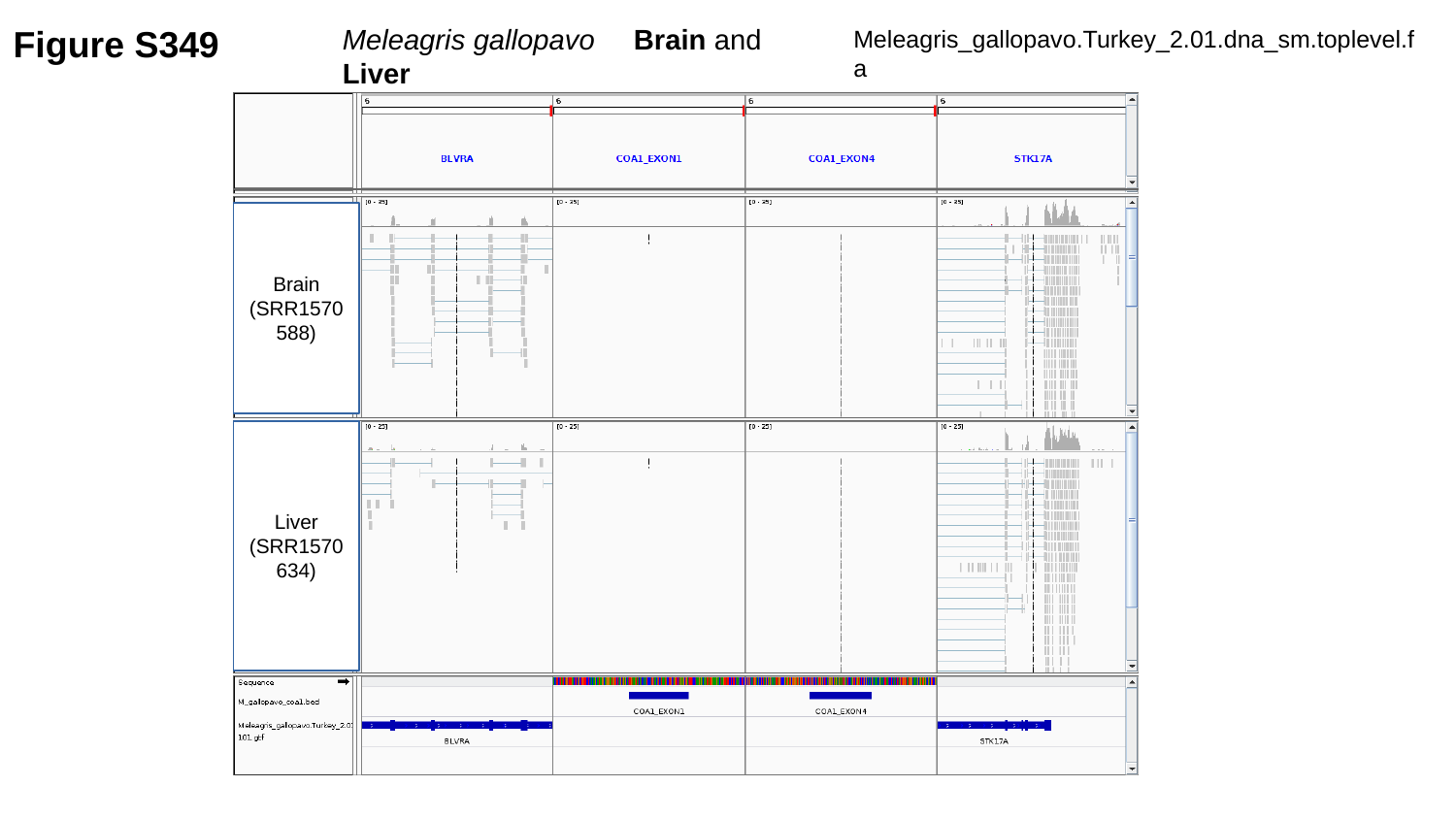

Figure S349
Meleagris gallopavo	Brain and Liver
Meleagris_gallopavo.Turkey_2.01.dna_sm.toplevel.fa
Brain
(SRR1570588)
Liver
(SRR1570634)

### Slide 350
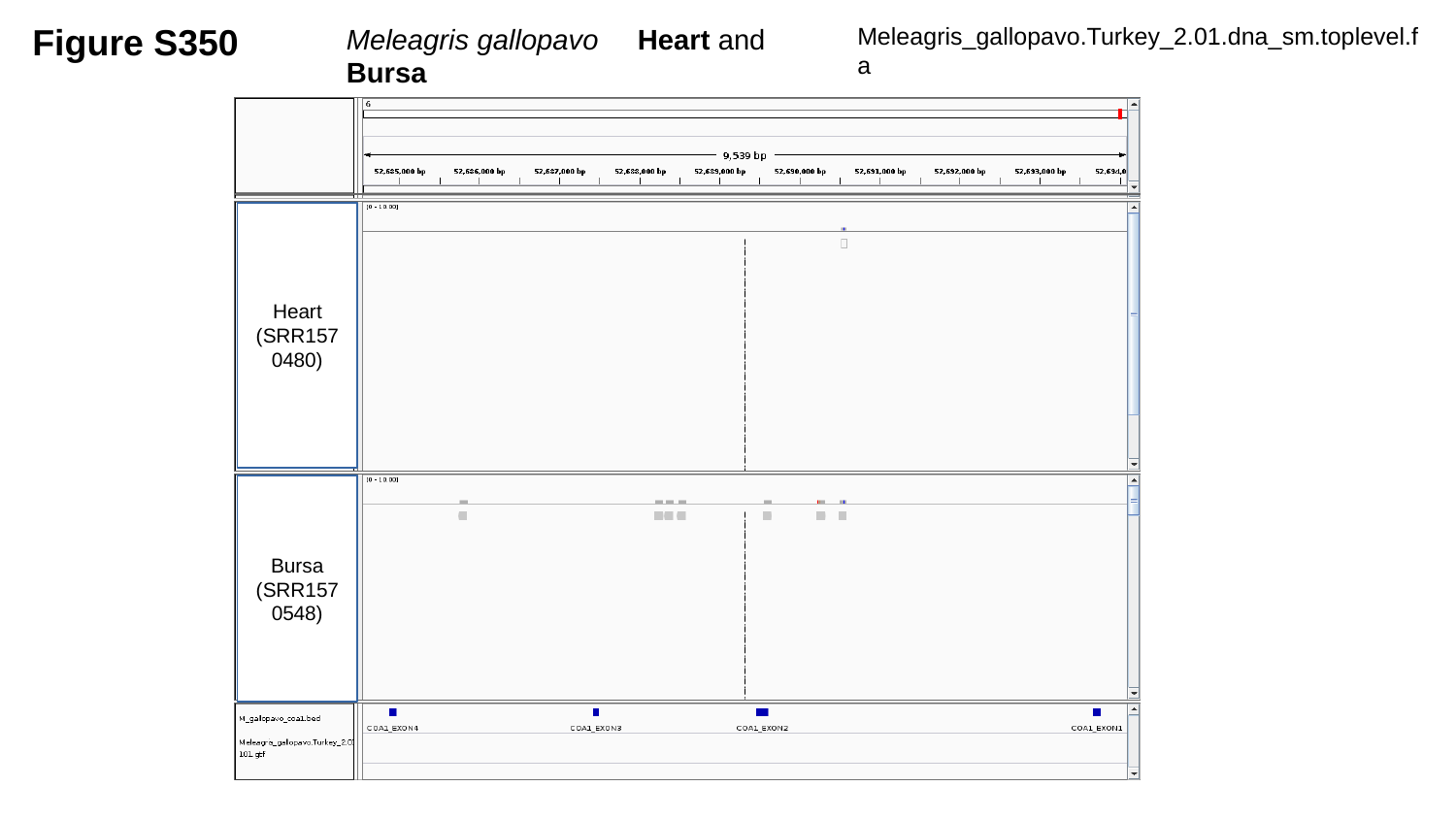

Figure S350
Meleagris gallopavo	Heart and Bursa
Meleagris_gallopavo.Turkey_2.01.dna_sm.toplevel.fa
Heart
(SRR1570480)
Bursa
(SRR1570548)

### Slide 351
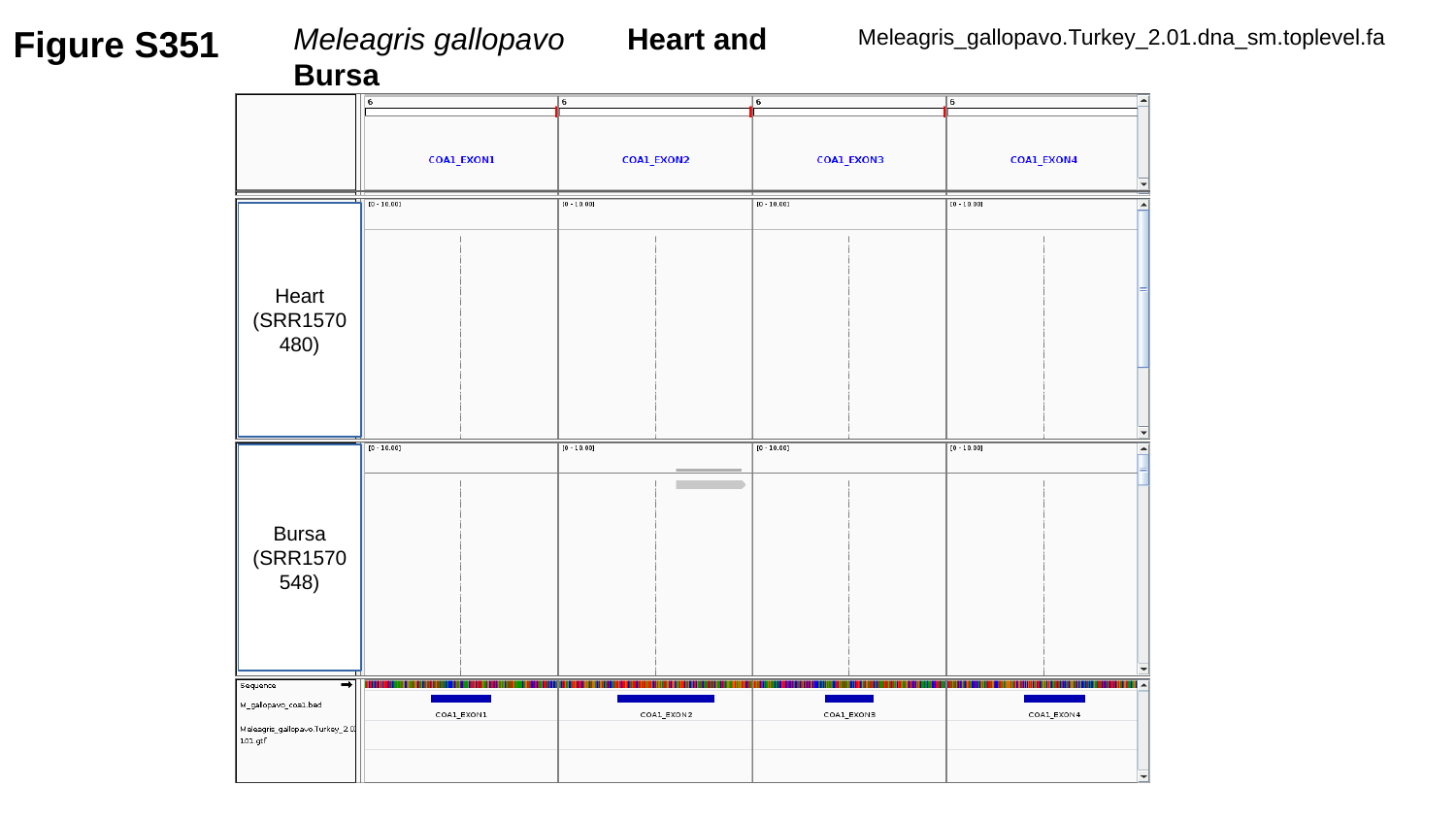

Figure S351
Meleagris gallopavo	 Heart and Bursa
Meleagris_gallopavo.Turkey_2.01.dna_sm.toplevel.fa
Heart
(SRR1570480)
Bursa
(SRR1570548)

### Slide 352
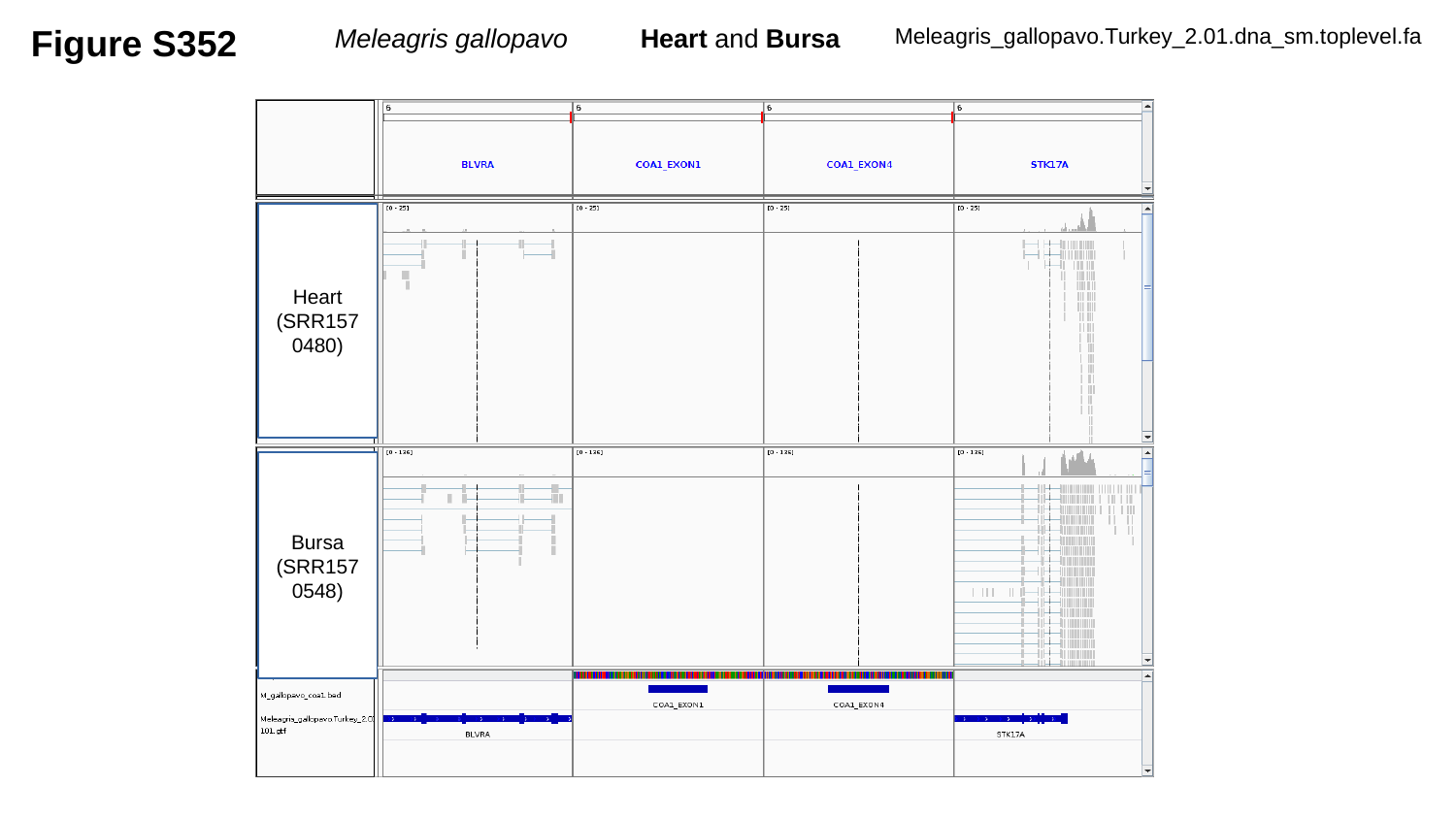

Figure S352
Meleagris gallopavo	 Heart and Bursa
Meleagris_gallopavo.Turkey_2.01.dna_sm.toplevel.fa
Heart
(SRR1570480)
Bursa
(SRR1570548)

### Slide 353
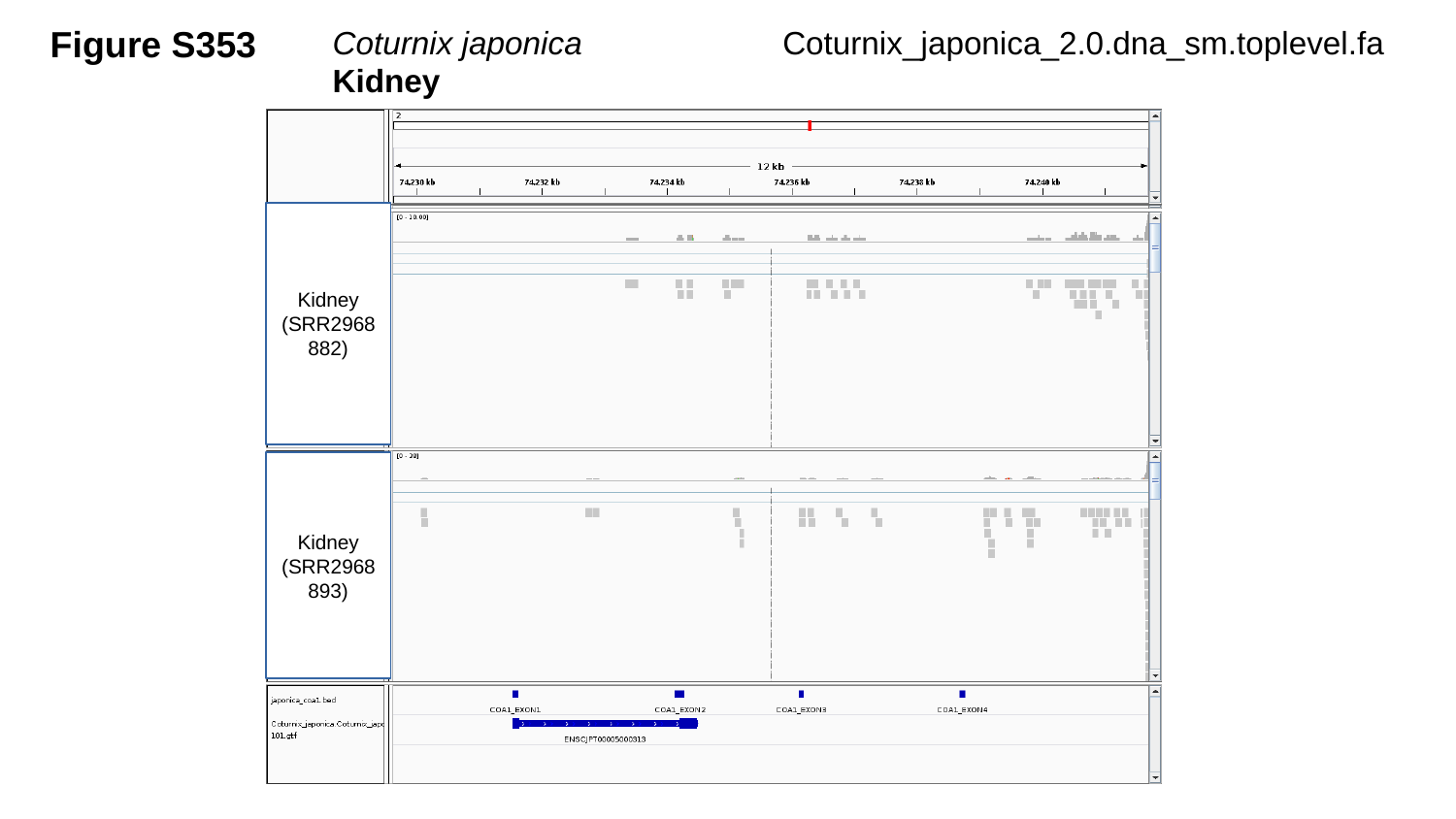

Figure S353
Coturnix japonica	 Kidney
Coturnix_japonica_2.0.dna_sm.toplevel.fa
Kidney
(SRR2968882)
Kidney
(SRR2968893)

### Slide 354
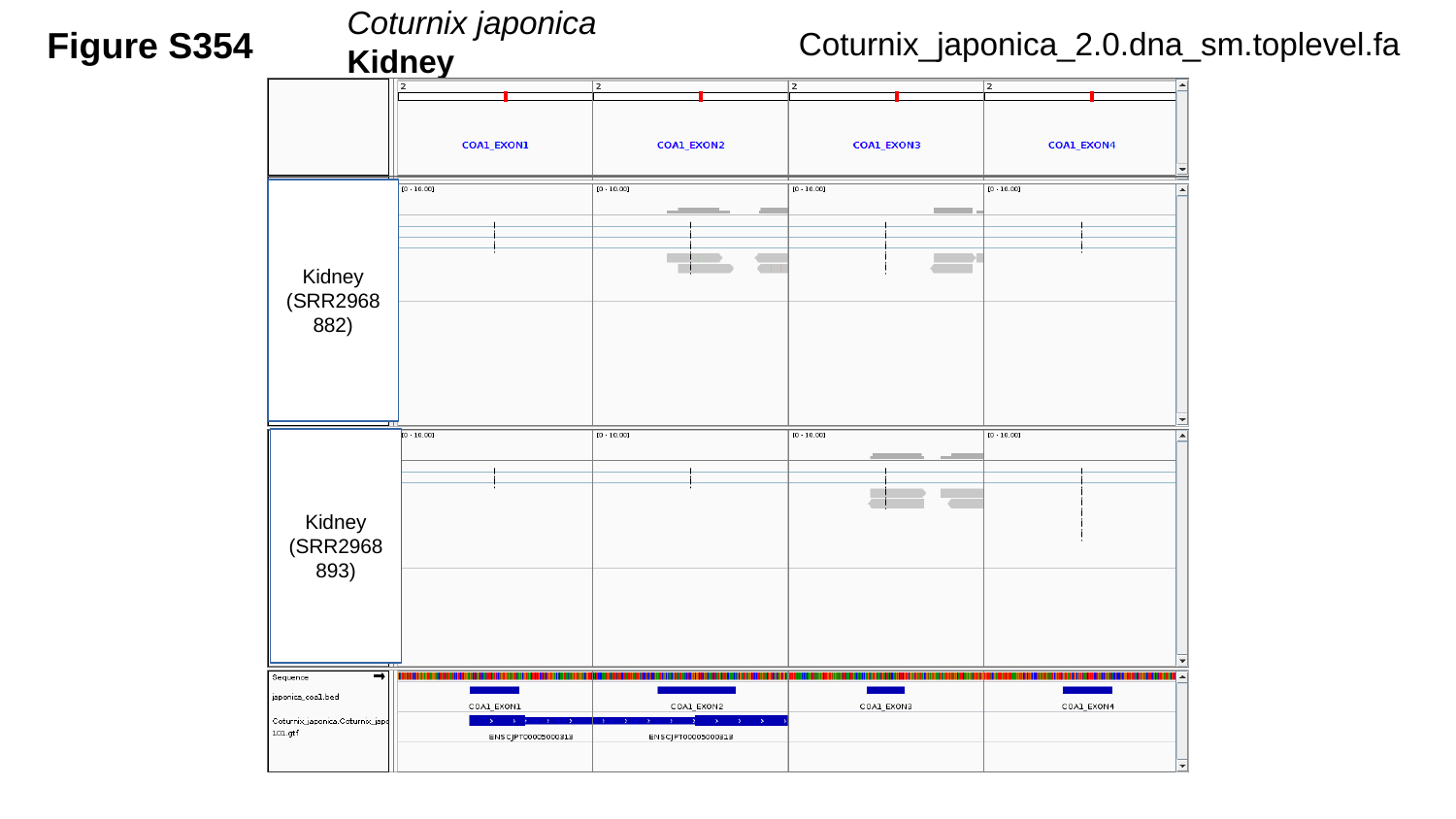

Coturnix japonica	 Kidney
Figure S354
Coturnix_japonica_2.0.dna_sm.toplevel.fa
Kidney
(SRR2968882)
Kidney
(SRR2968893)

### Slide 355
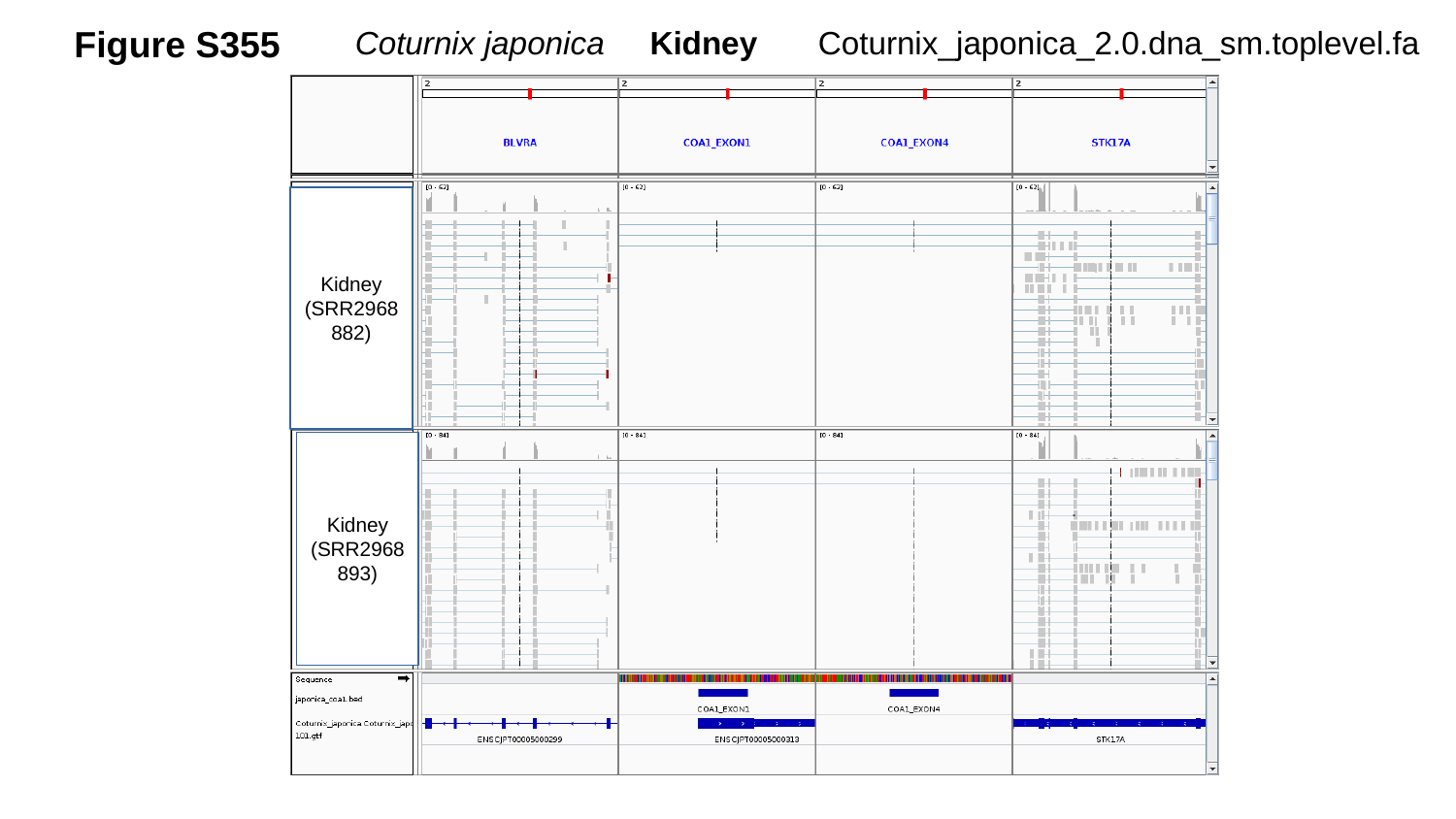

Figure S355
Coturnix_japonica_2.0.dna_sm.toplevel.fa
Coturnix japonica Kidney
Kidney
(SRR2968882)
Kidney
(SRR2968893)

### Slide 356
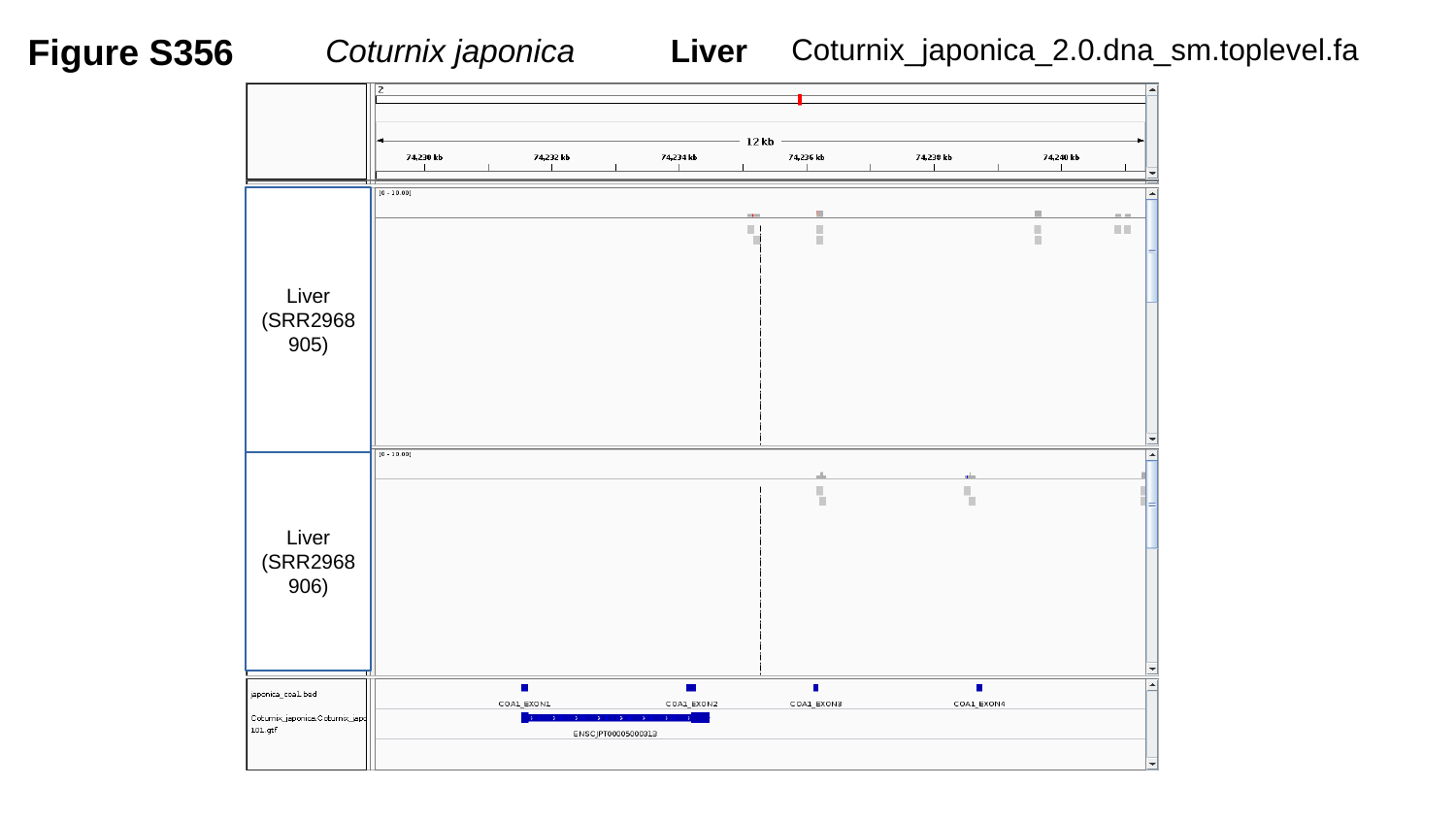

Figure S356
Coturnix japonica	 Liver
Coturnix_japonica_2.0.dna_sm.toplevel.fa
Liver
(SRR2968905)
Liver
(SRR2968906)

### Slide 357
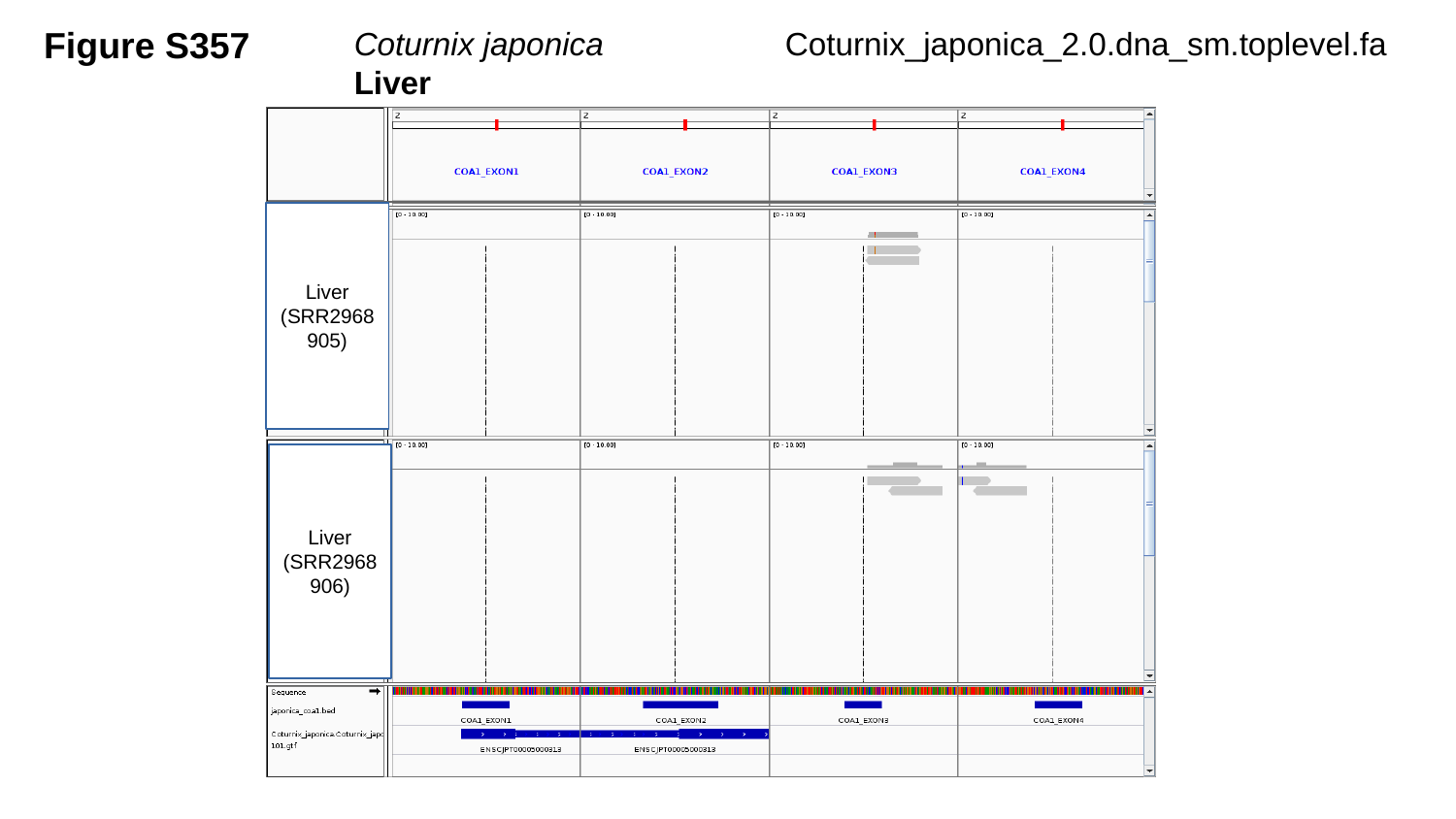

Figure S357
Coturnix japonica	 Liver
Coturnix_japonica_2.0.dna_sm.toplevel.fa
Liver
(SRR2968905)
Liver
(SRR2968906)

### Slide 358
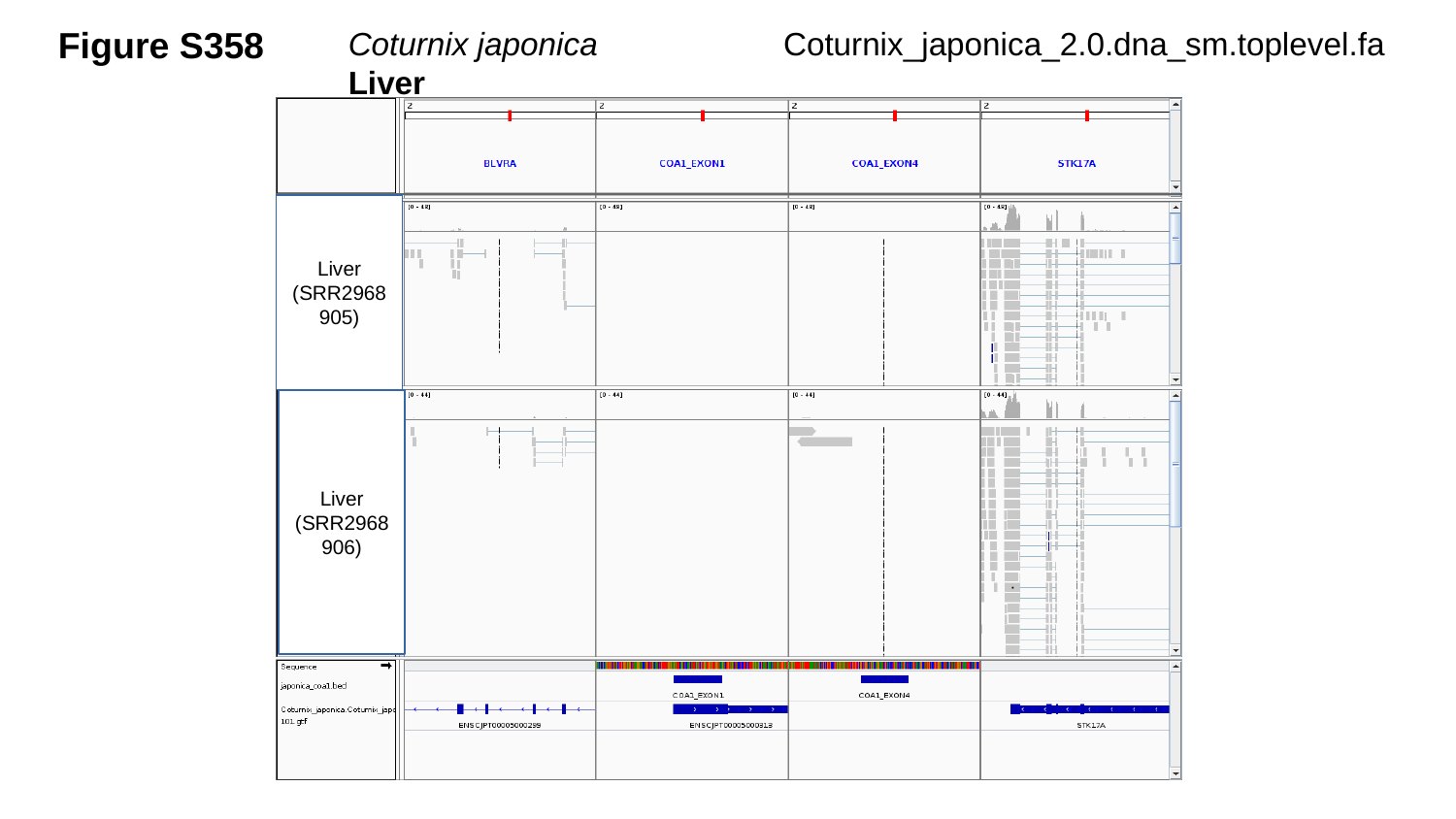

Figure S358
Coturnix japonica	 Liver
Coturnix_japonica_2.0.dna_sm.toplevel.fa
Liver
(SRR2968905)
Liver
(SRR2968906)

### Slide 359
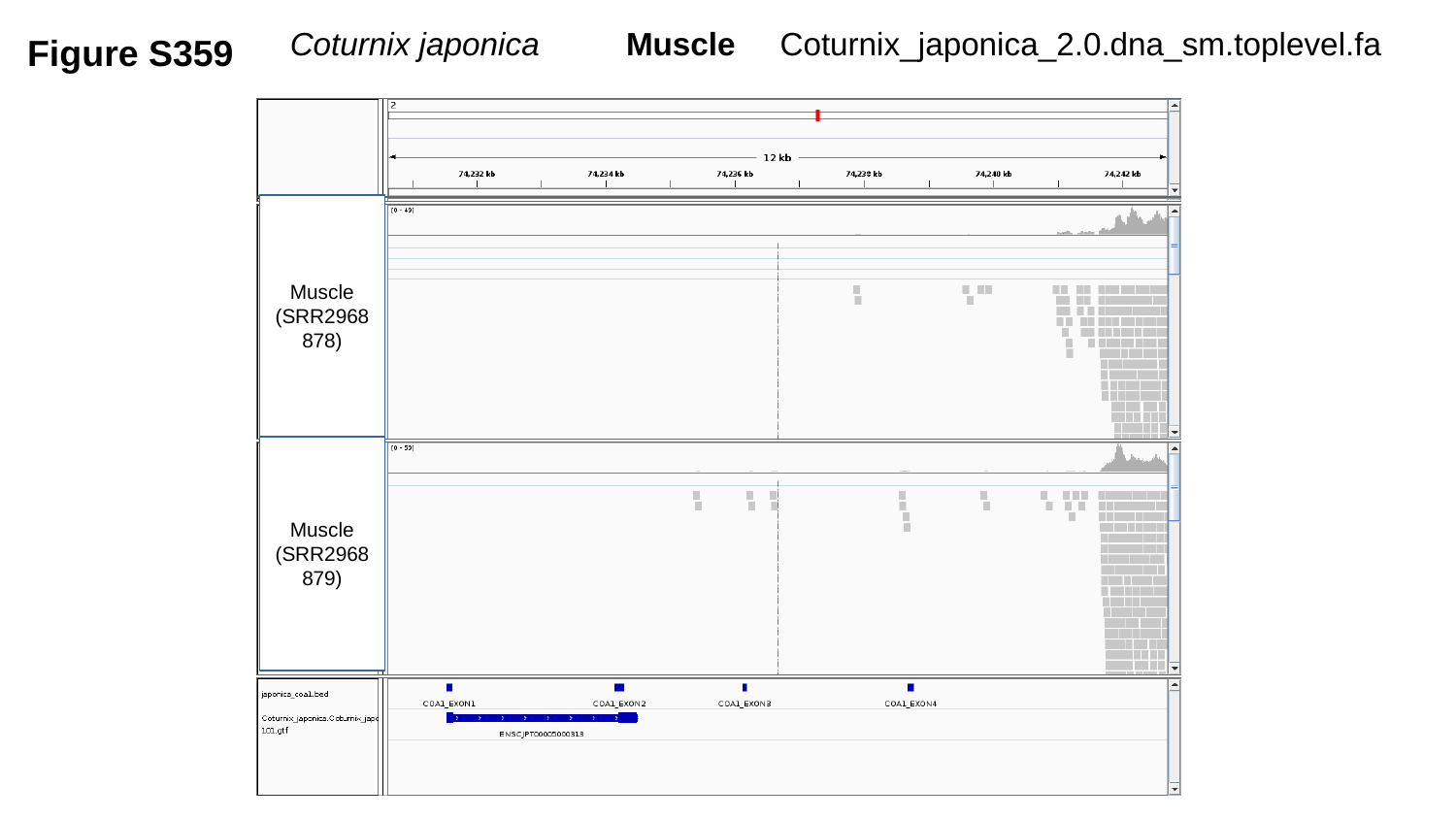

Coturnix_japonica_2.0.dna_sm.toplevel.fa
Figure S359
Coturnix japonica	 Muscle
Muscle
(SRR2968878)
Muscle
(SRR2968879)

### Slide 360
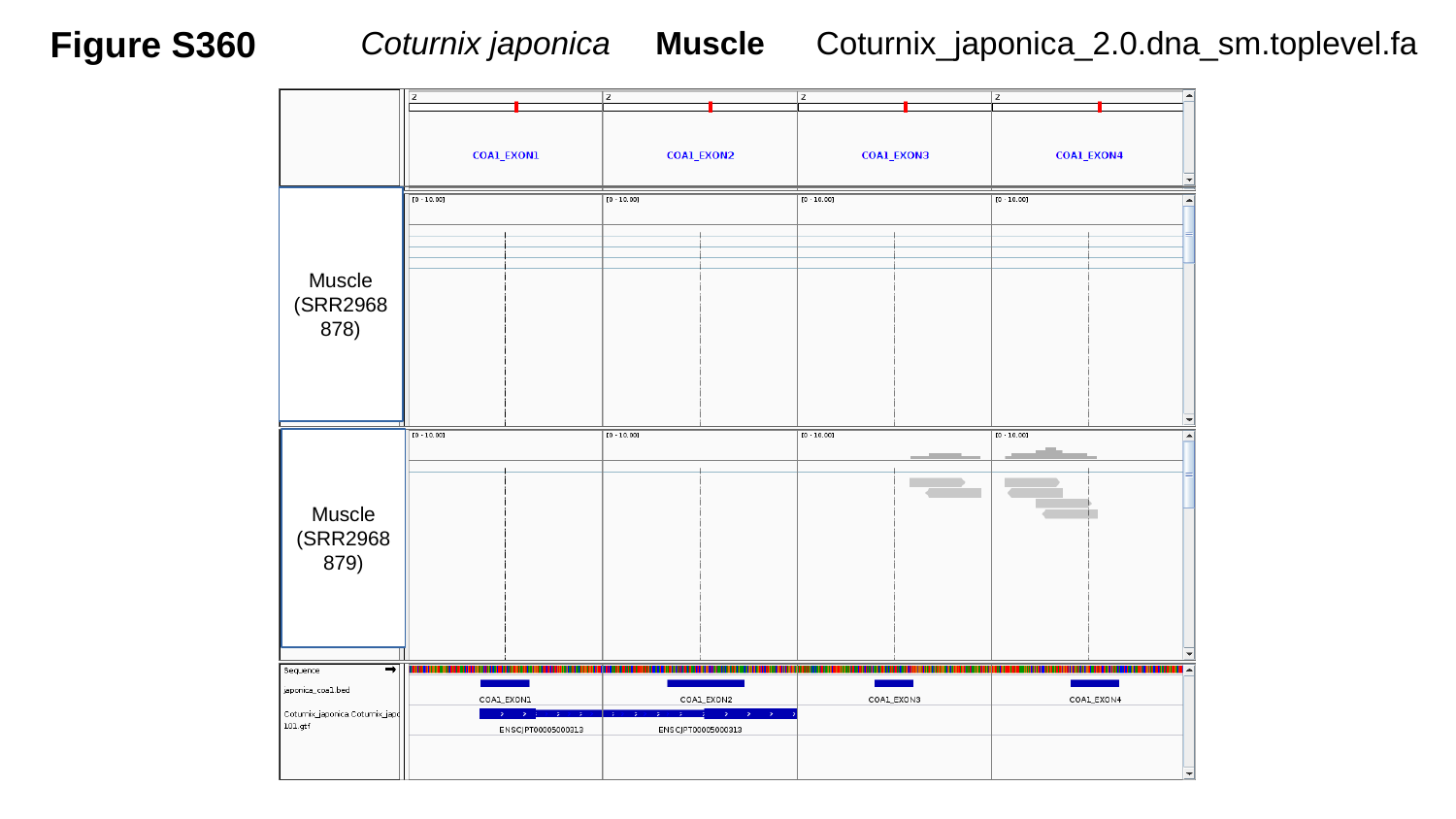

Figure S360
Coturnix japonica Muscle
Coturnix_japonica_2.0.dna_sm.toplevel.fa
Muscle
(SRR2968878)
Muscle
(SRR2968879)

### Slide 361
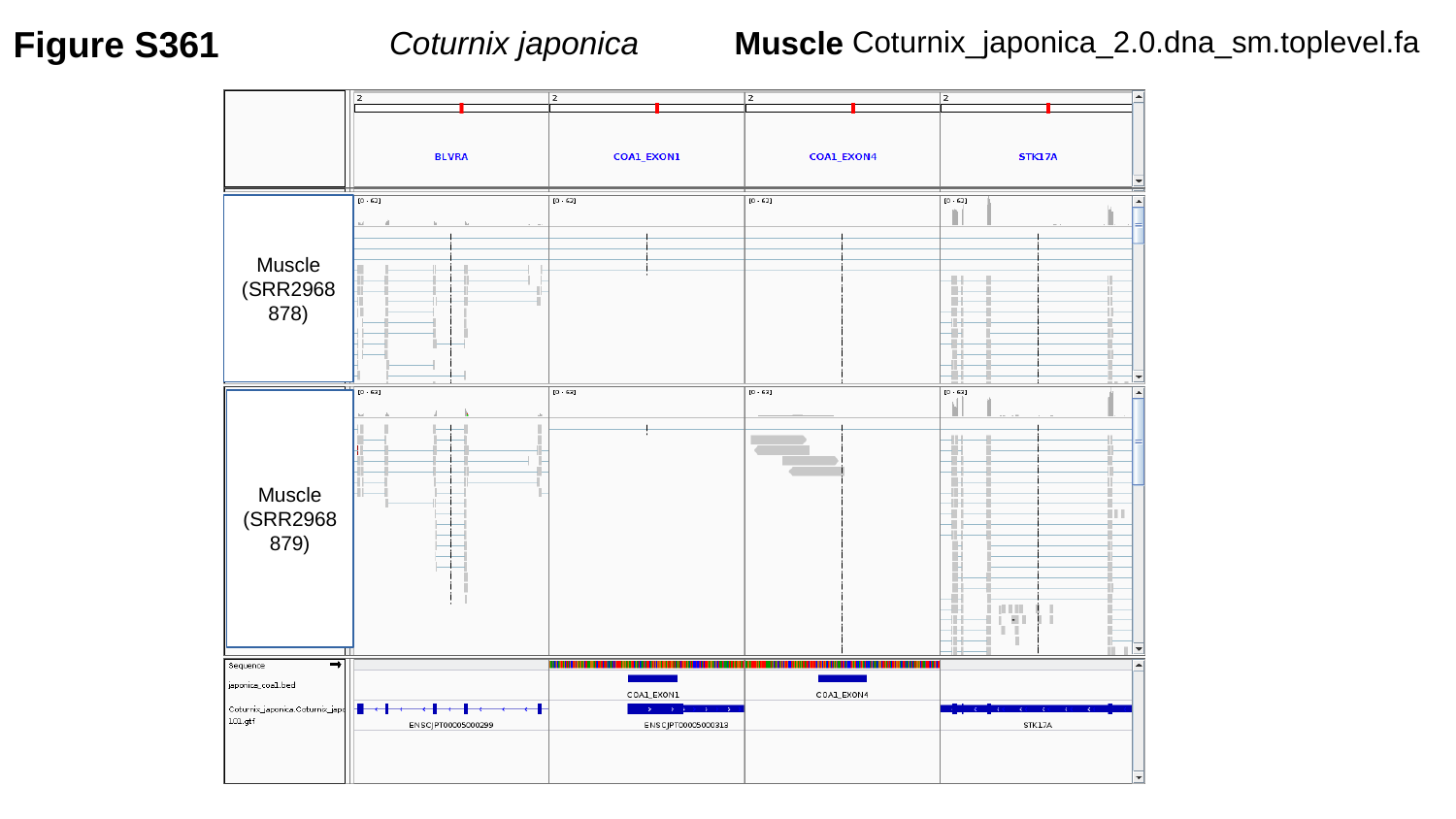

Figure S361
Coturnix japonica	 Muscle
Coturnix_japonica_2.0.dna_sm.toplevel.fa
Muscle
(SRR2968878)
Muscle
(SRR2968879)

### Slide 362
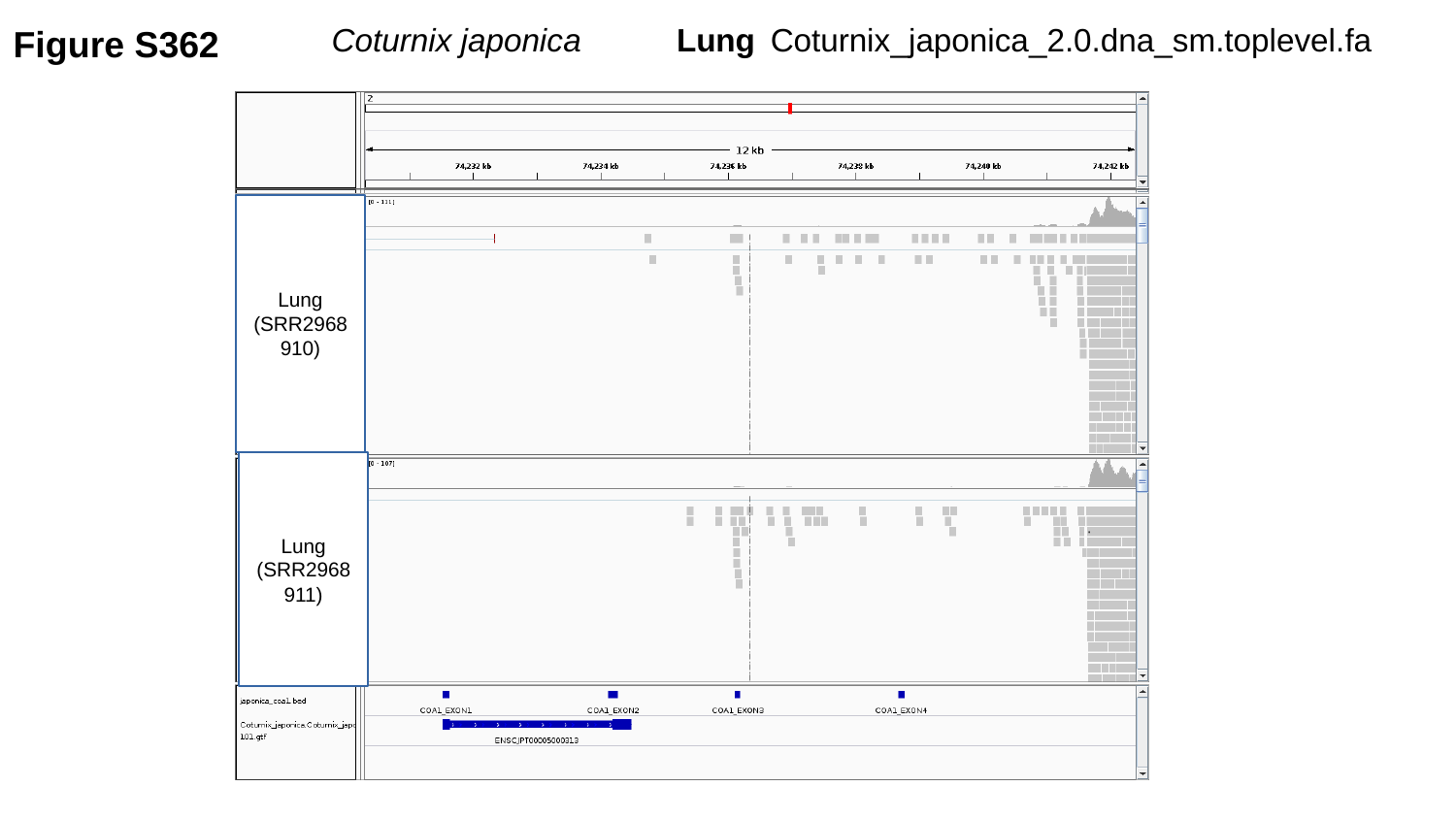

Figure S362
Coturnix_japonica_2.0.dna_sm.toplevel.fa
Coturnix japonica	 Lung
Lung
(SRR2968910)
Lung
(SRR2968911)

### Slide 363
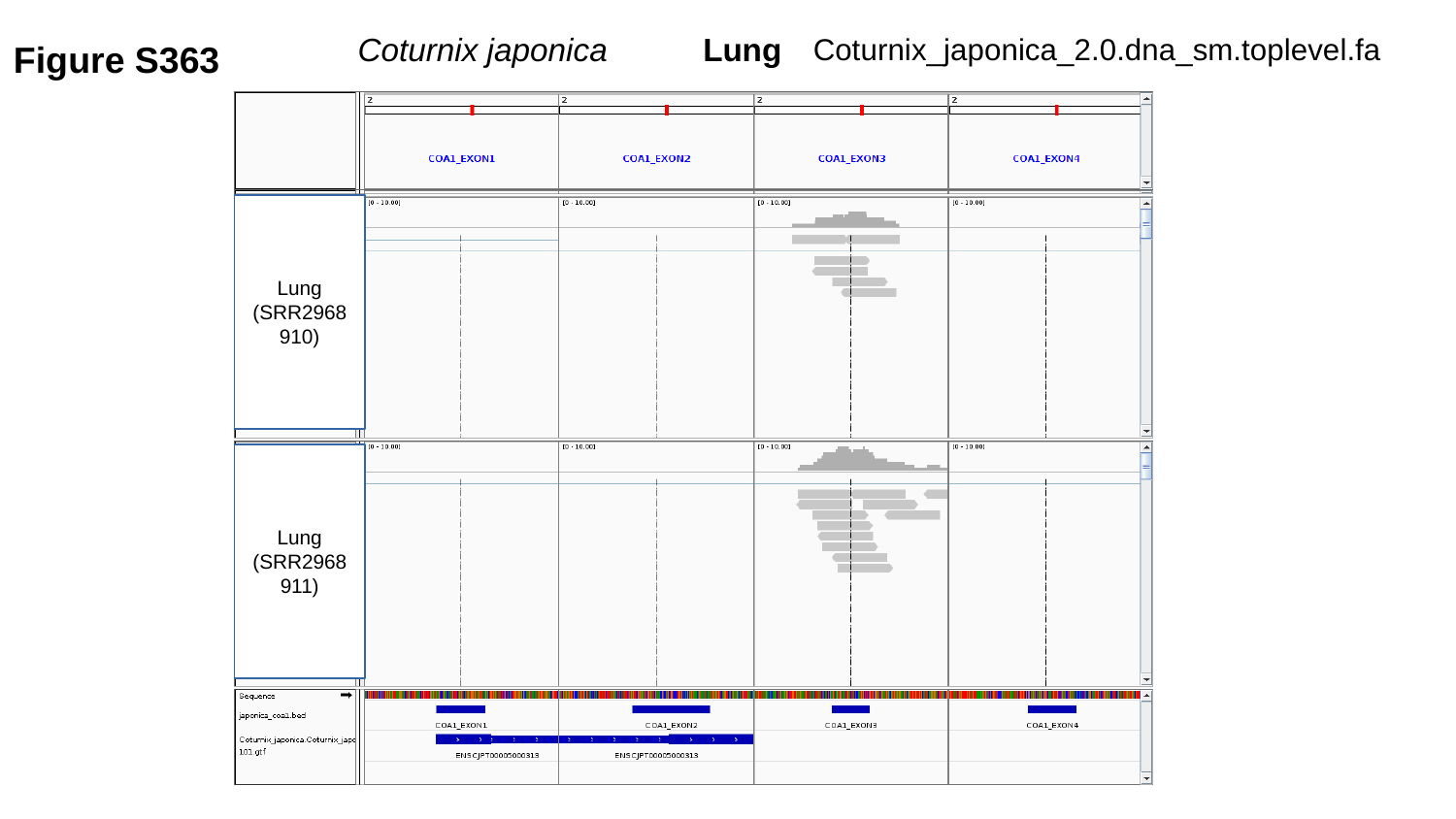

Coturnix_japonica_2.0.dna_sm.toplevel.fa
Figure S363
Coturnix japonica	 Lung
Lung
(SRR2968910)
Lung
(SRR2968911)

### Slide 364
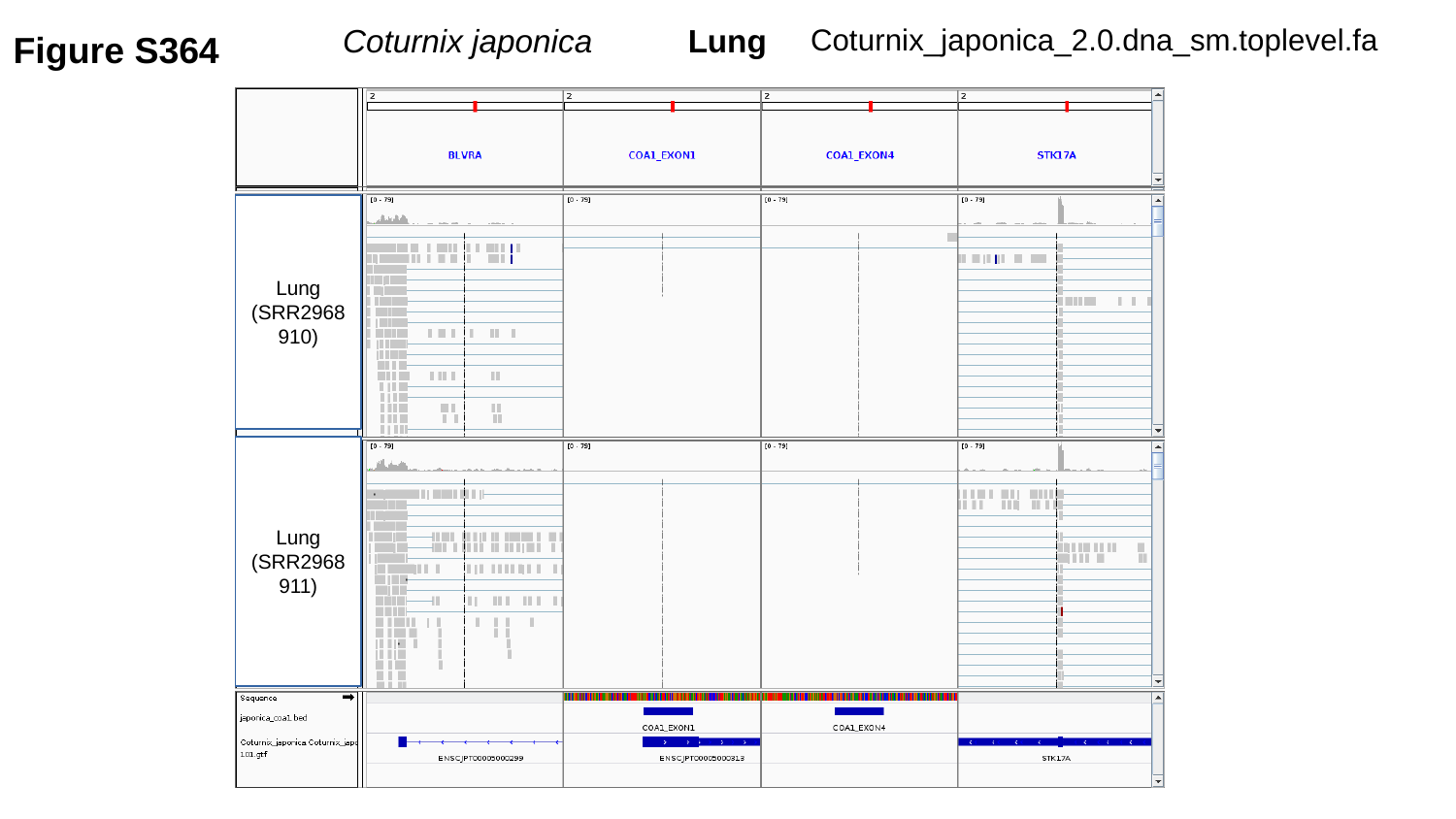

Coturnix_japonica_2.0.dna_sm.toplevel.fa
Figure S364
Coturnix japonica	 Lung
Lung
(SRR2968910)
Lung
(SRR2968911)

### Slide 365
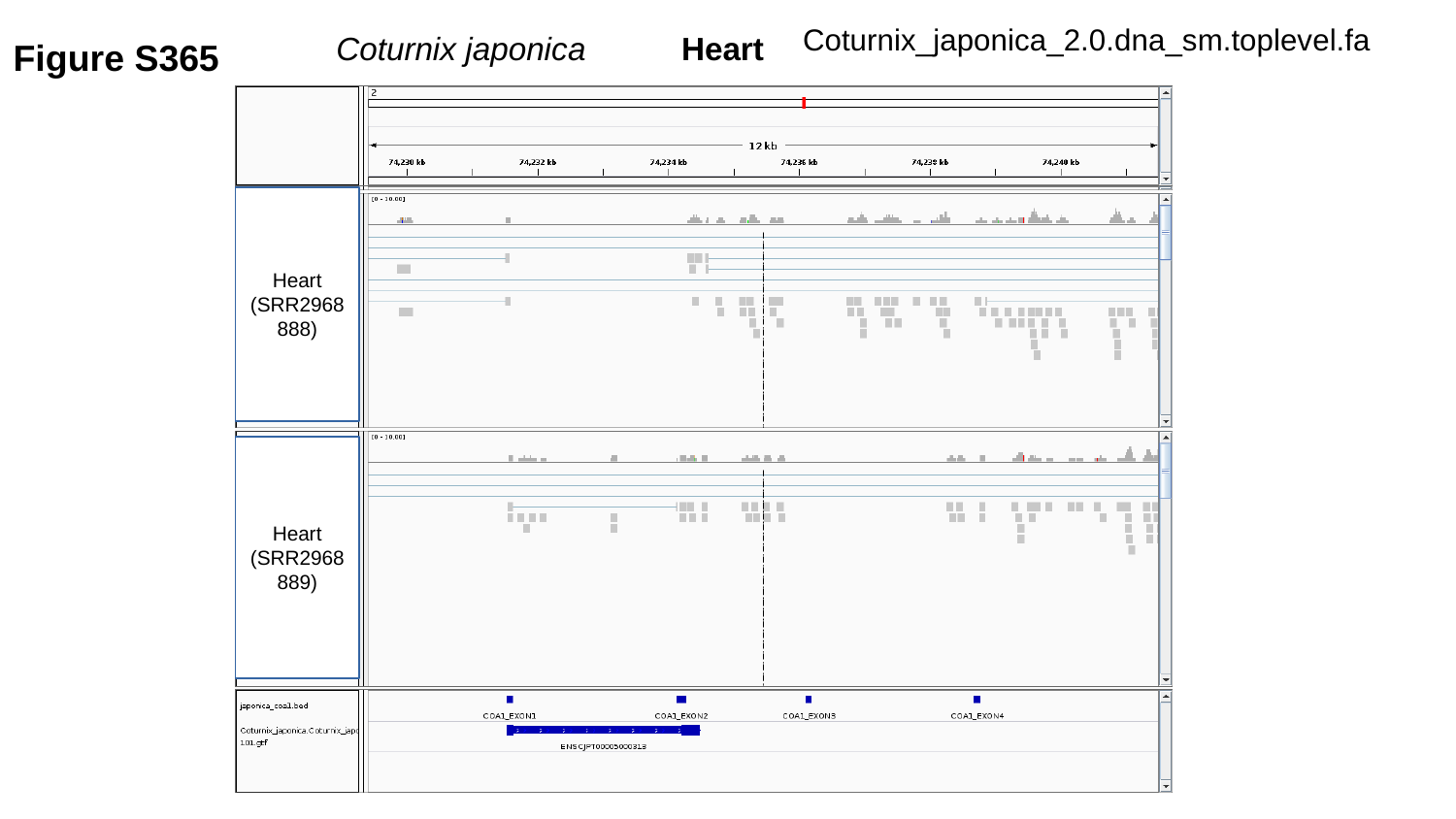

Coturnix_japonica_2.0.dna_sm.toplevel.fa
Figure S365
Coturnix japonica	 Heart
Heart
(SRR2968888)
Heart
(SRR2968889)

### Slide 366
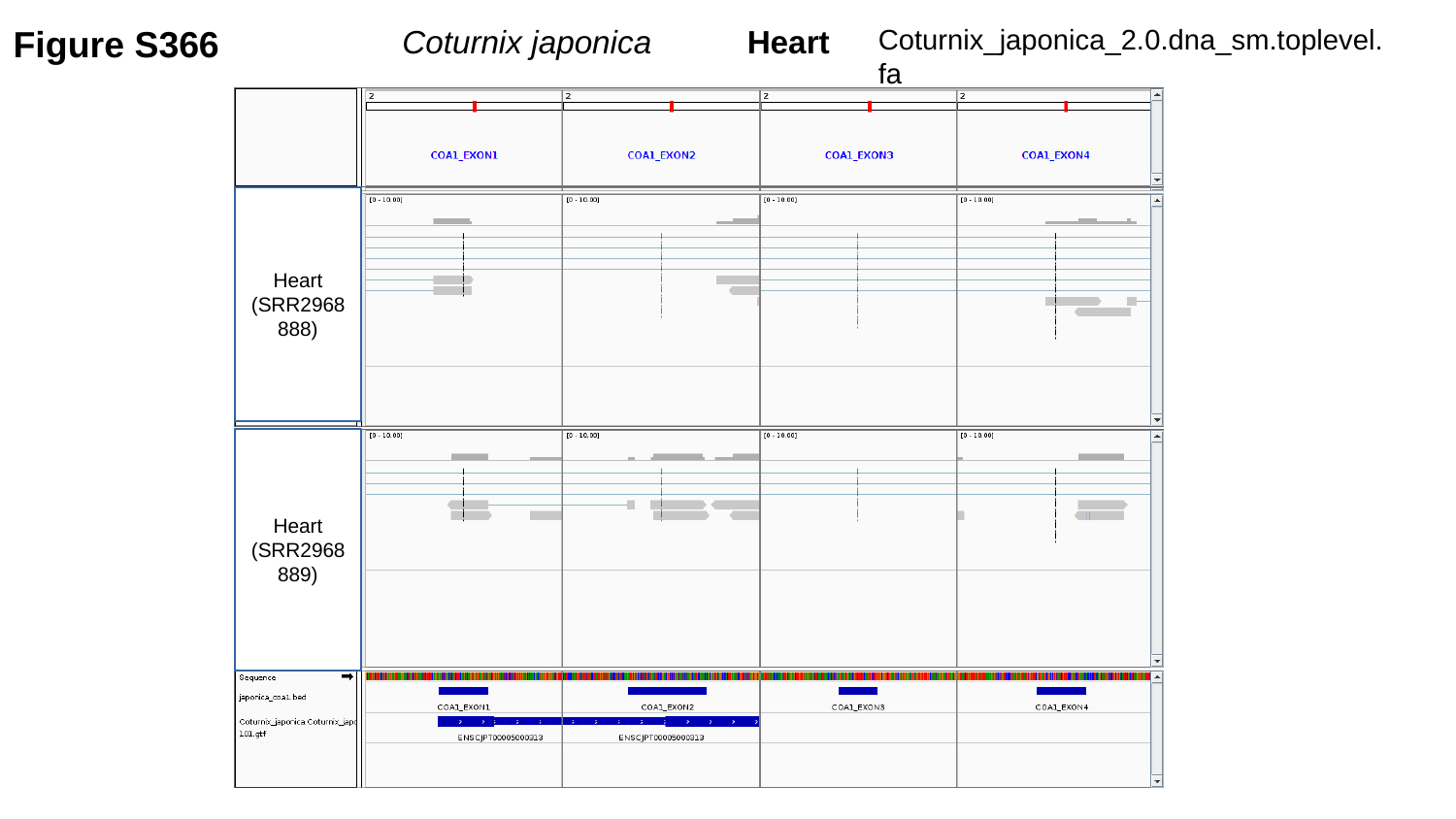

Figure S366
Coturnix_japonica_2.0.dna_sm.toplevel.fa
Coturnix japonica	 Heart
Heart
(SRR2968888)
Heart
(SRR2968889)

### Slide 367
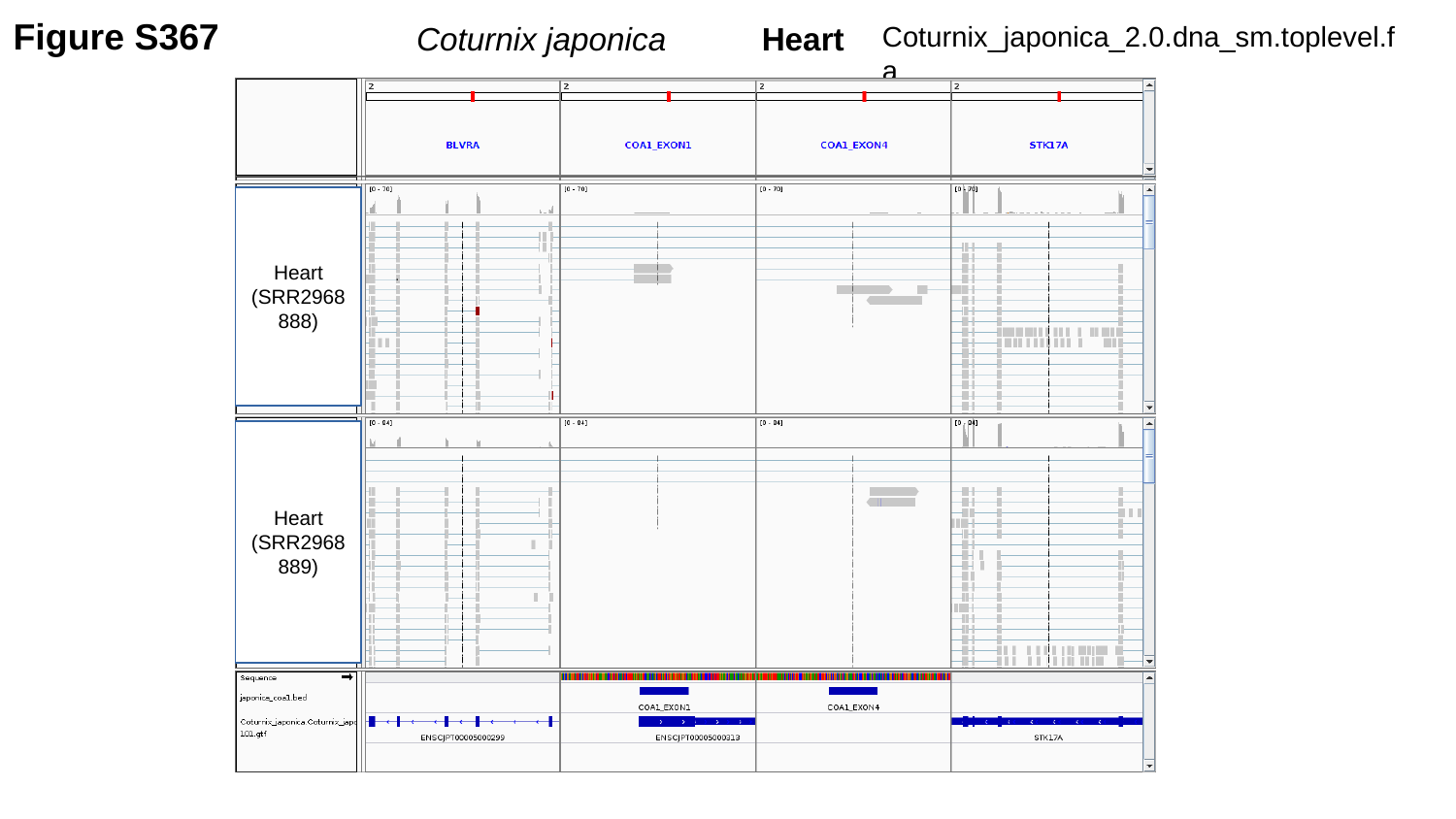

Figure S367
Coturnix_japonica_2.0.dna_sm.toplevel.fa
Coturnix japonica	 Heart
Heart
(SRR2968888)
Heart
(SRR2968889)

### Slide 368
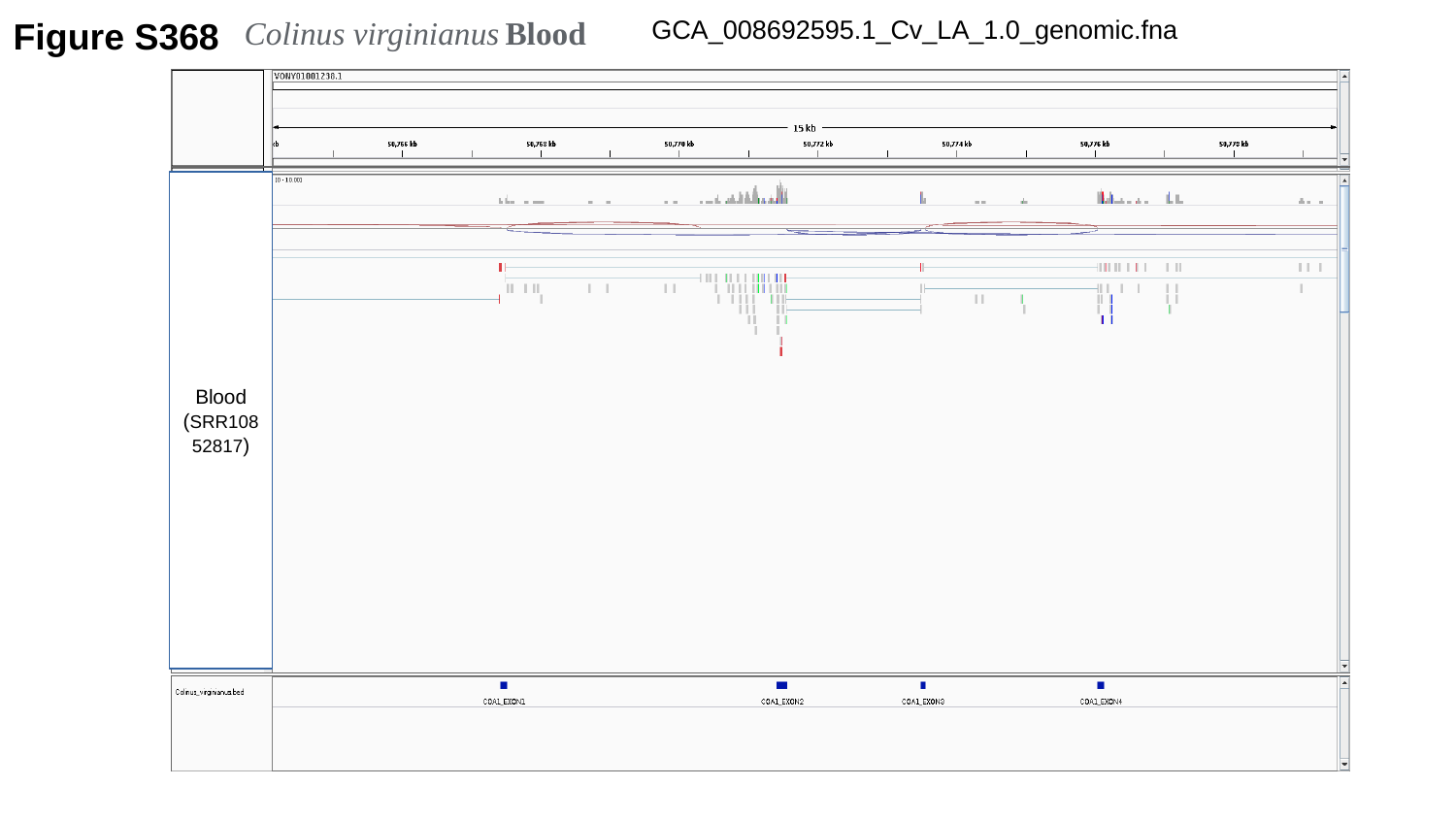

Figure S368
Colinus virginianus Blood
GCA_008692595.1_Cv_LA_1.0_genomic.fna
Blood
(SRR10852817)

### Slide 369
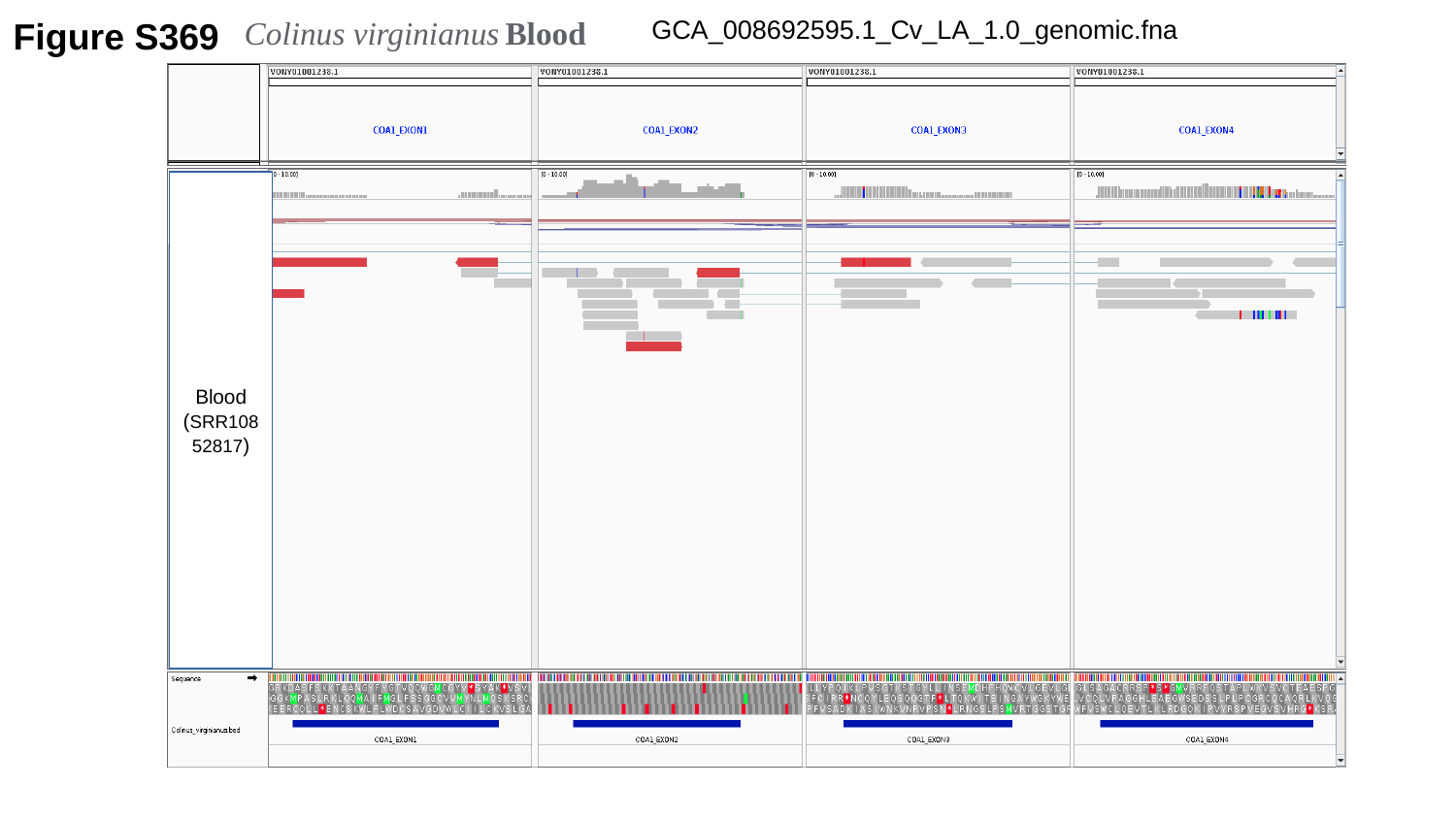

Figure S369
Colinus virginianus Blood
GCA_008692595.1_Cv_LA_1.0_genomic.fna
Blood
(SRR10852817)

### Slide 370
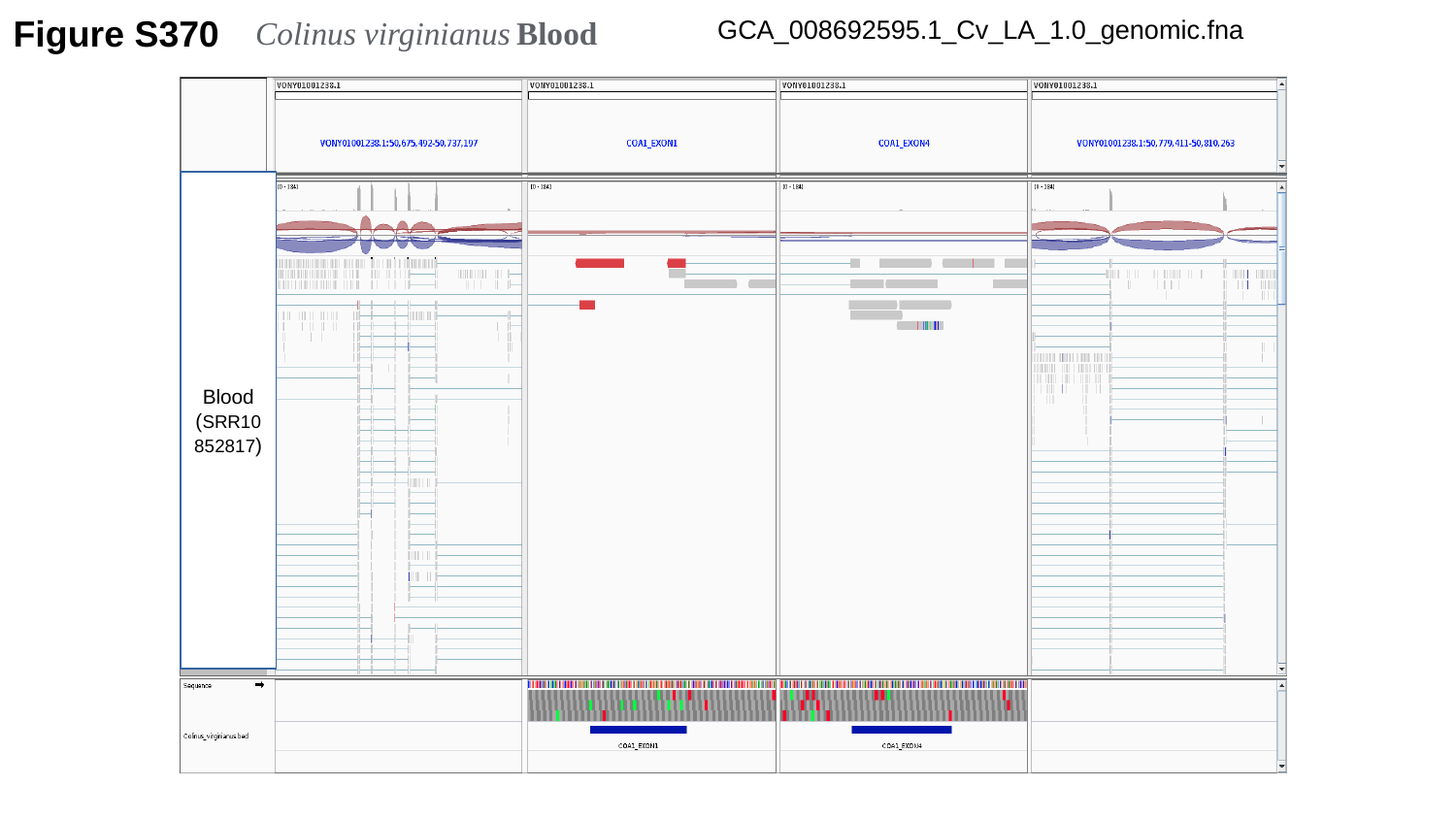

Colinus virginianus Blood
GCA_008692595.1_Cv_LA_1.0_genomic.fna
Figure S370
Blood
(SRR10852817)

### Slide 371
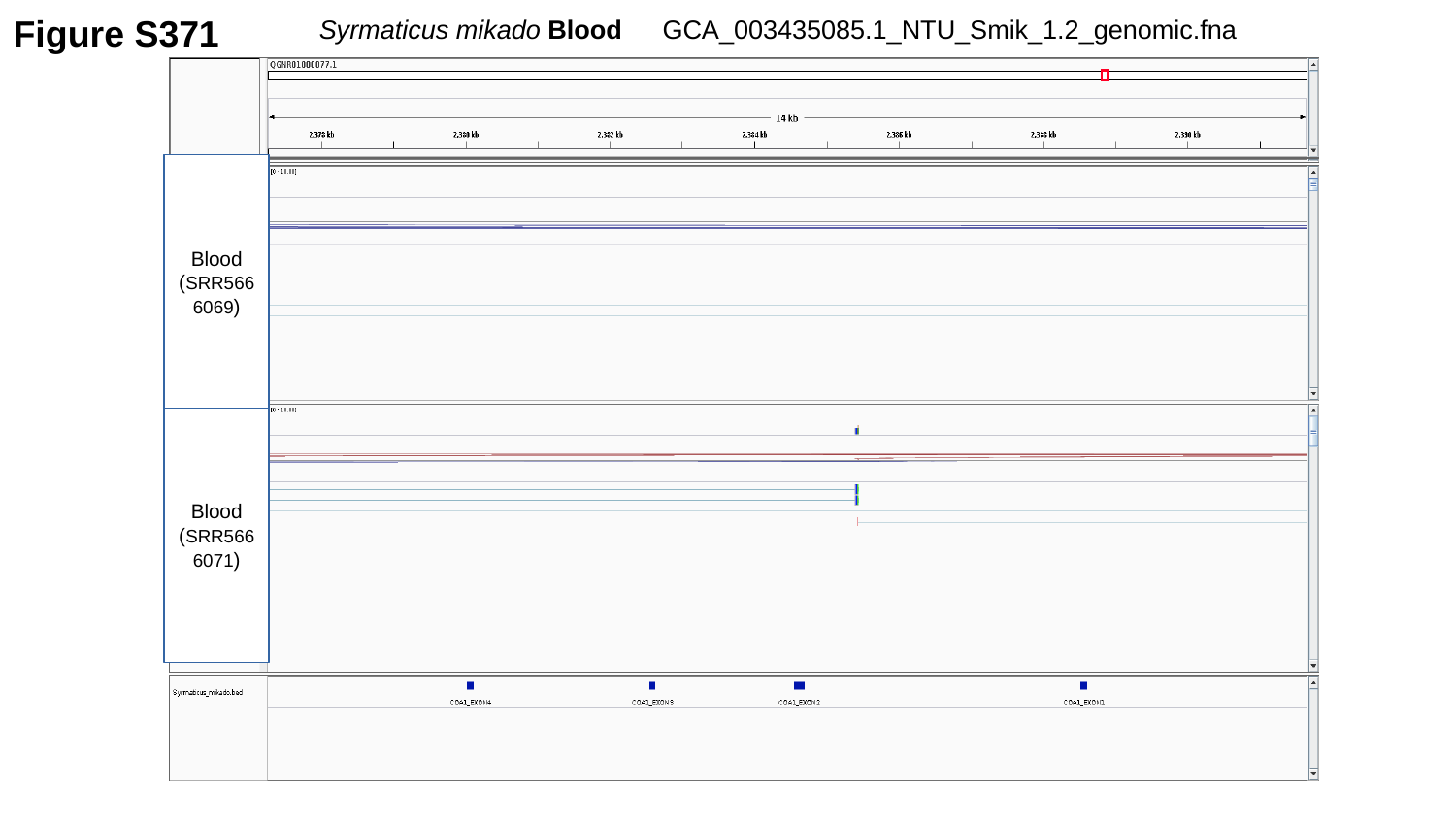

Syrmaticus mikado Blood
GCA_003435085.1_NTU_Smik_1.2_genomic.fna
Figure S371
Blood
(SRR5666069)
Blood
(SRR5666071)

### Slide 372
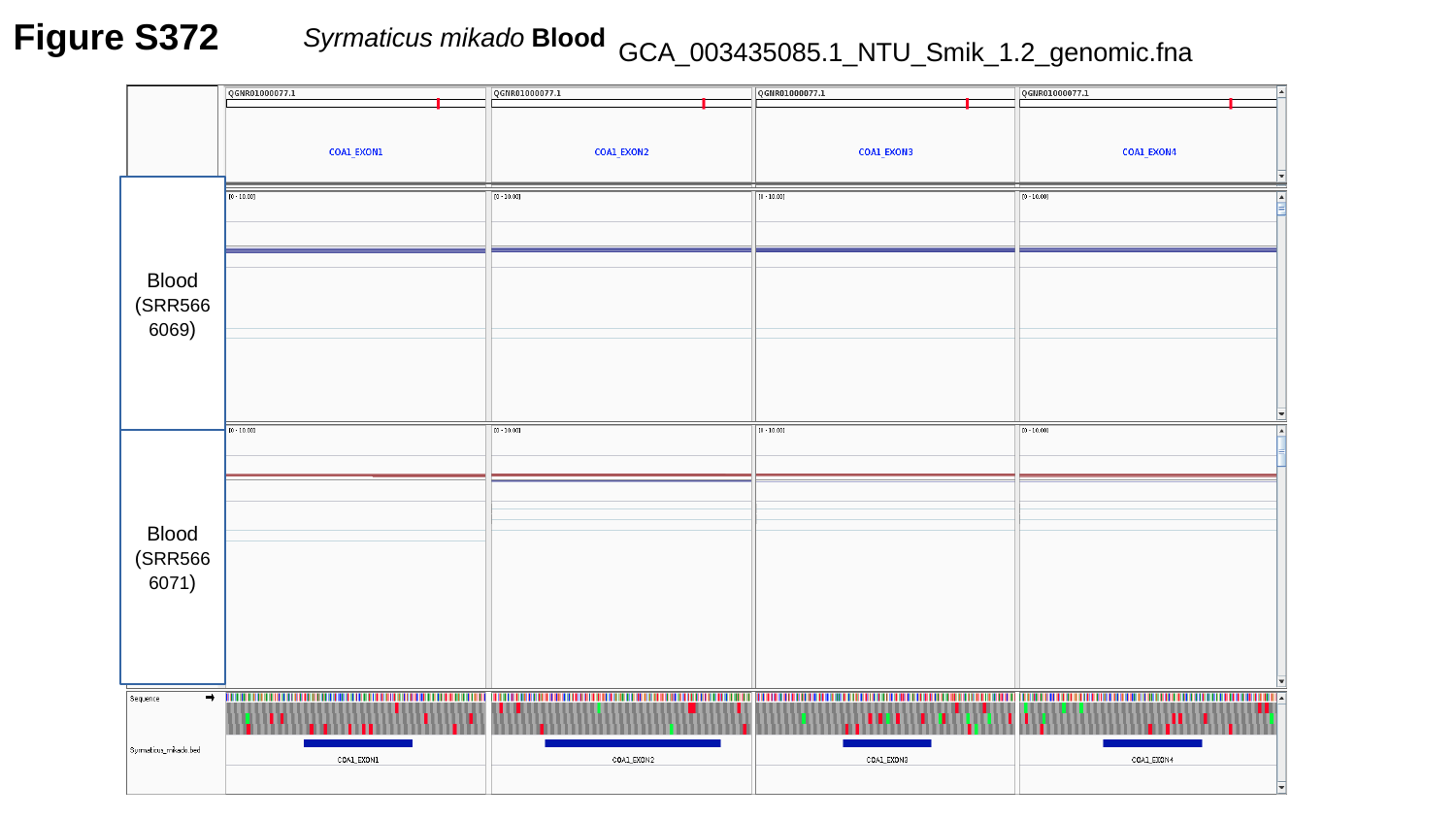

Figure S372
Syrmaticus mikado Blood
GCA_003435085.1_NTU_Smik_1.2_genomic.fna
Blood
(SRR5666069)
Blood
(SRR5666071)

### Slide 373
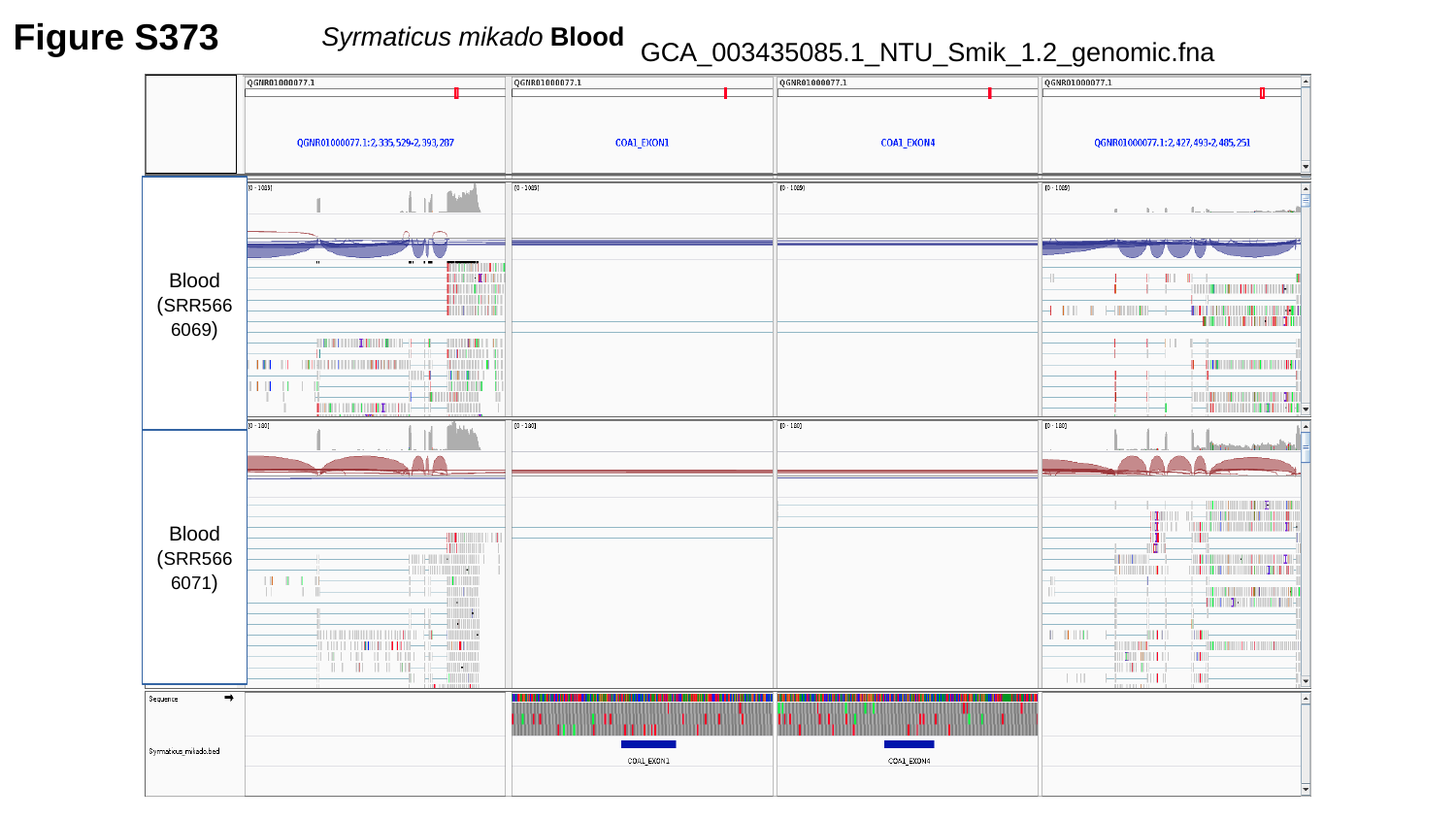

Figure S373
Syrmaticus mikado Blood
GCA_003435085.1_NTU_Smik_1.2_genomic.fna
Blood
(SRR5666069)
Blood
(SRR5666071)

### Slide 374
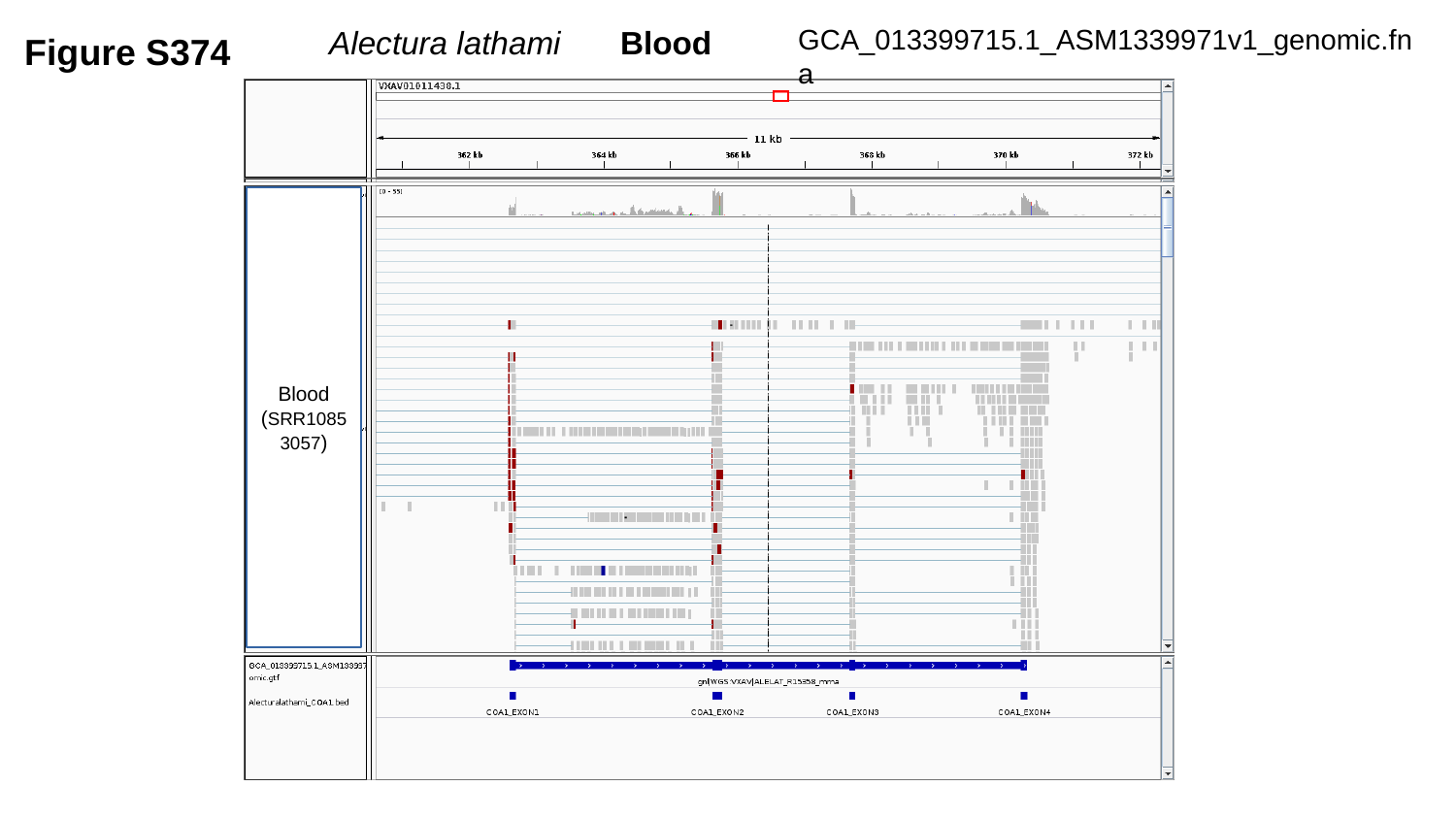

GCA_013399715.1_ASM1339971v1_genomic.fna
Figure S374
Alectura lathami	Blood
Blood
(SRR10853057)

### Slide 375
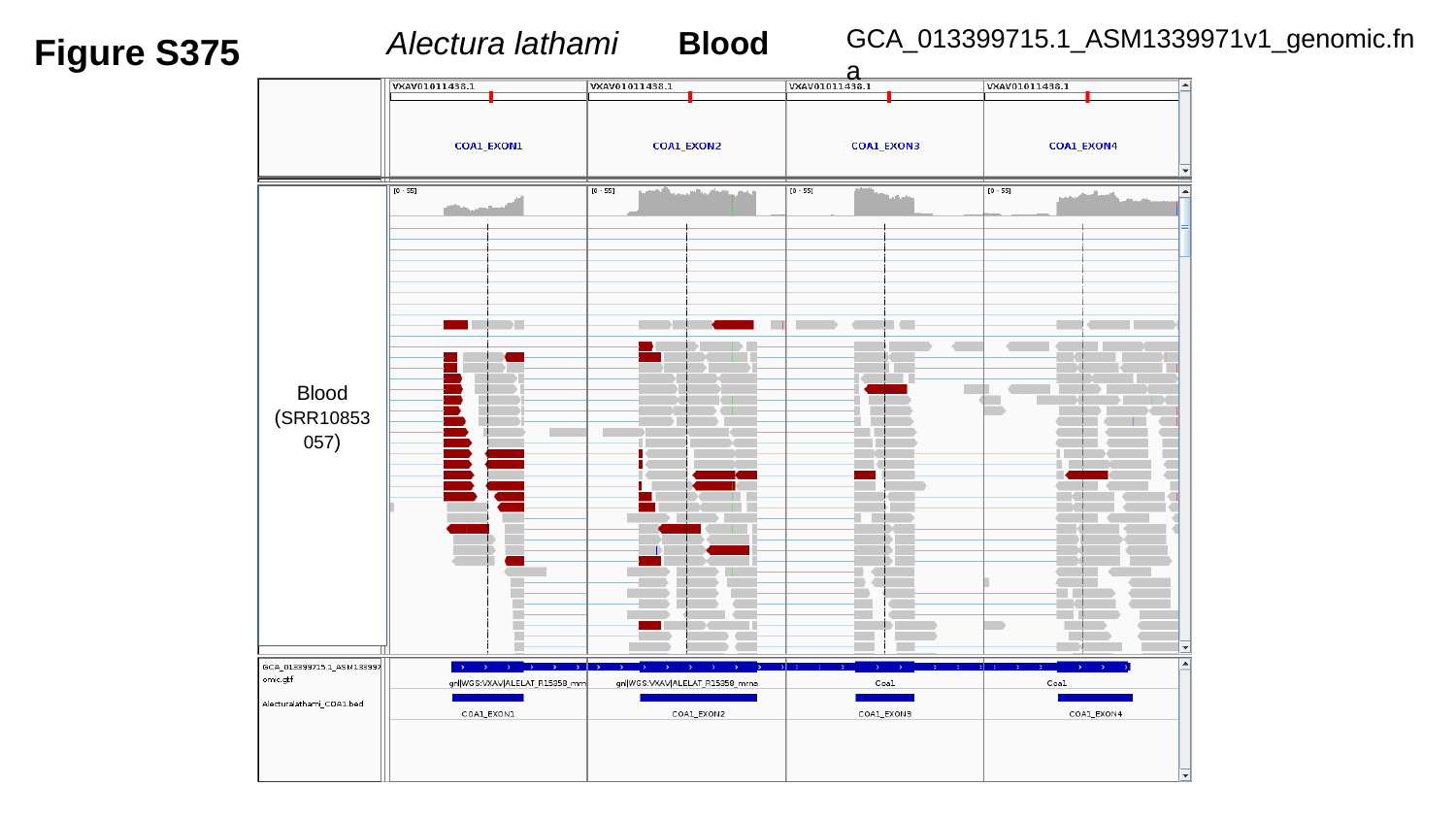

Figure S375
Alectura lathami	Blood
GCA_013399715.1_ASM1339971v1_genomic.fna
Blood
(SRR10853057)

### Slide 376
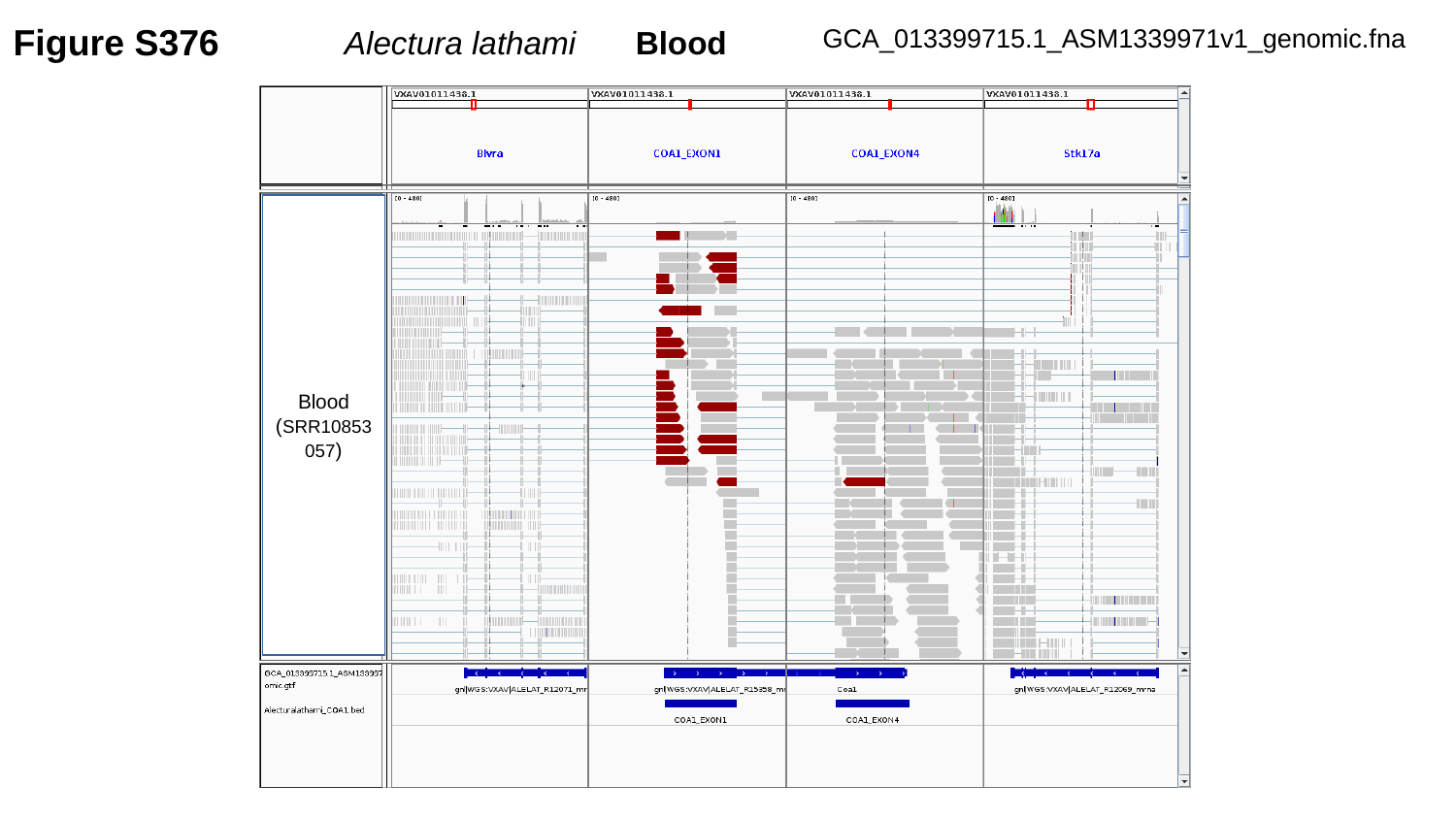

Figure S376
GCA_013399715.1_ASM1339971v1_genomic.fna
Alectura lathami	Blood
Blood
(SRR10853057)

### Slide 377
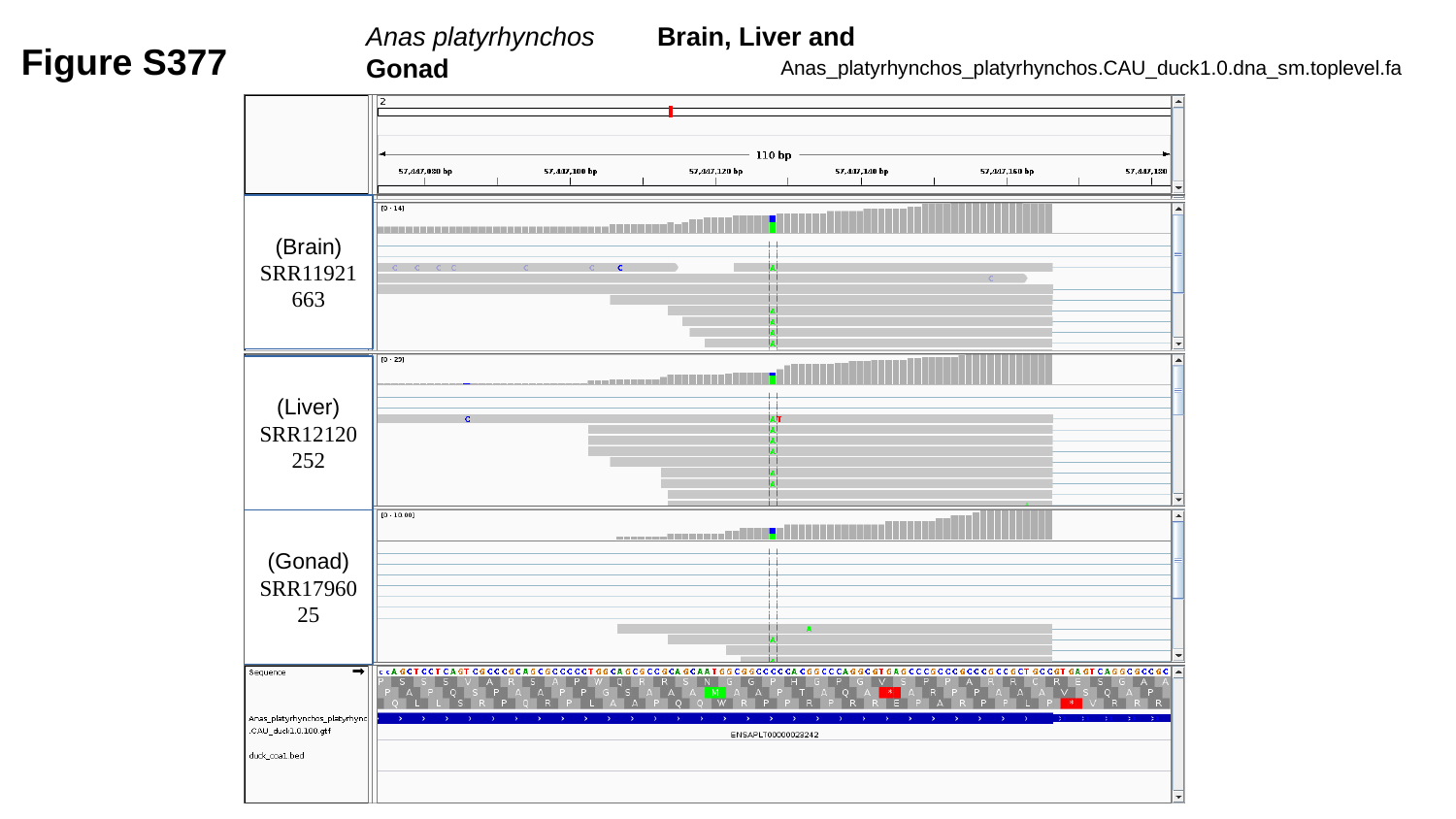

Anas platyrhynchos	Brain, Liver and Gonad
Figure S377
Anas_platyrhynchos_platyrhynchos.CAU_duck1.0.dna_sm.toplevel.fa
(Brain)
SRR11921663
(Liver)
SRR12120252
(Gonad)
SRR1796025

### Slide 378
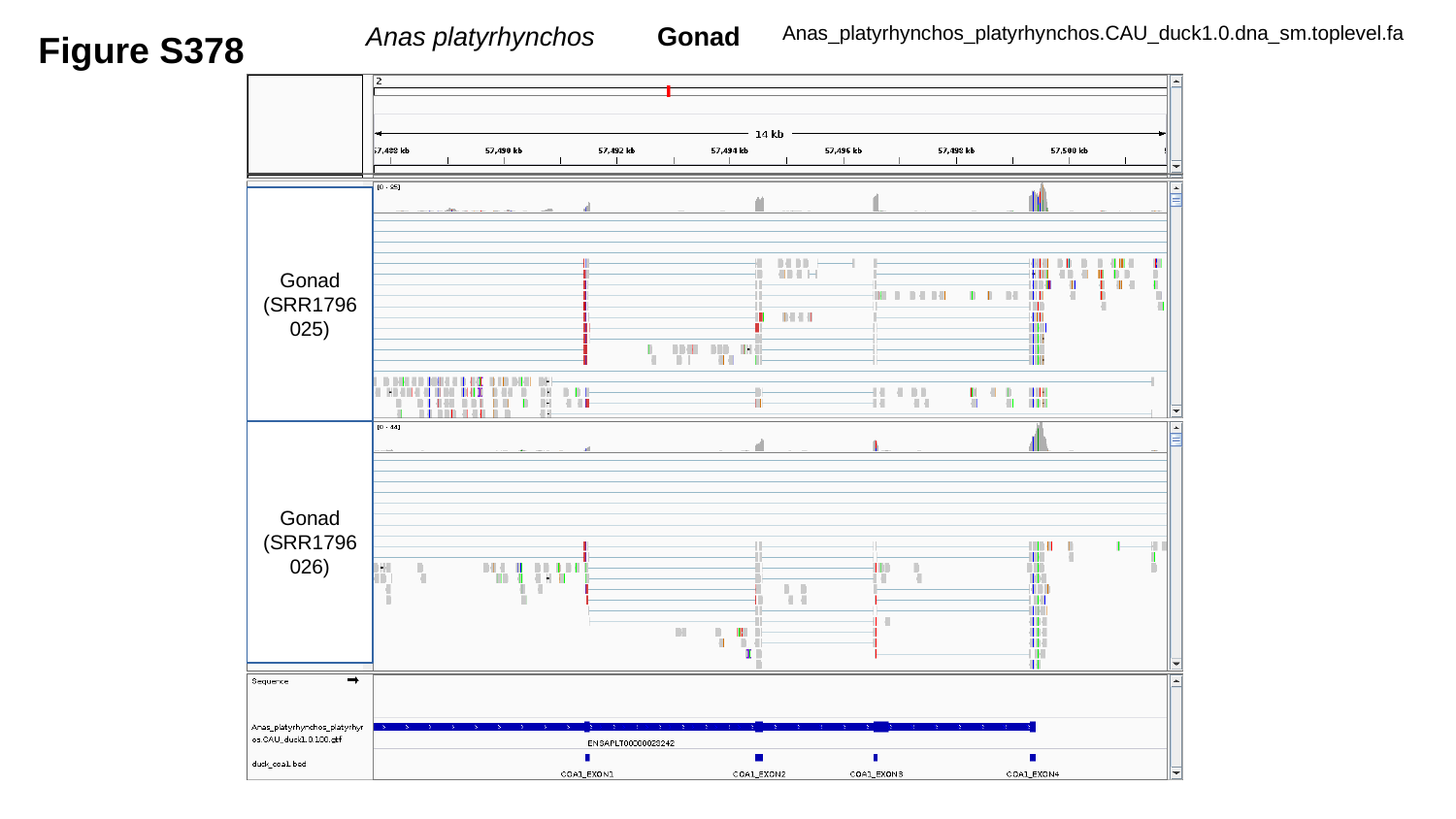

Figure S378
Anas platyrhynchos	Gonad
Anas_platyrhynchos_platyrhynchos.CAU_duck1.0.dna_sm.toplevel.fa
Gonad
(SRR1796025)
Gonad
(SRR1796026)

### Slide 379
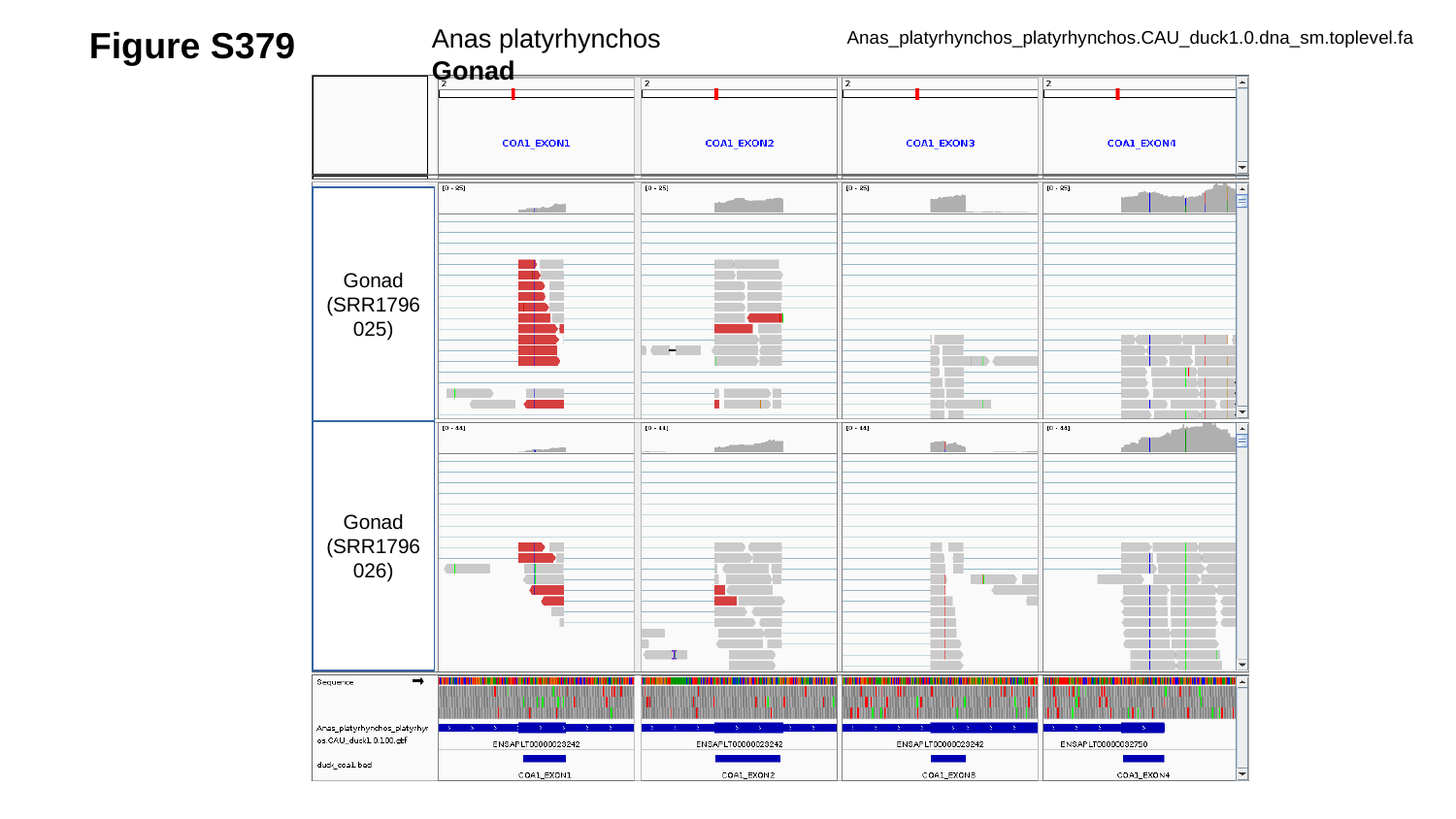

Figure S379
Anas platyrhynchos	Gonad
Anas_platyrhynchos_platyrhynchos.CAU_duck1.0.dna_sm.toplevel.fa
Gonad
(SRR1796025)
Gonad
(SRR1796026)

### Slide 380
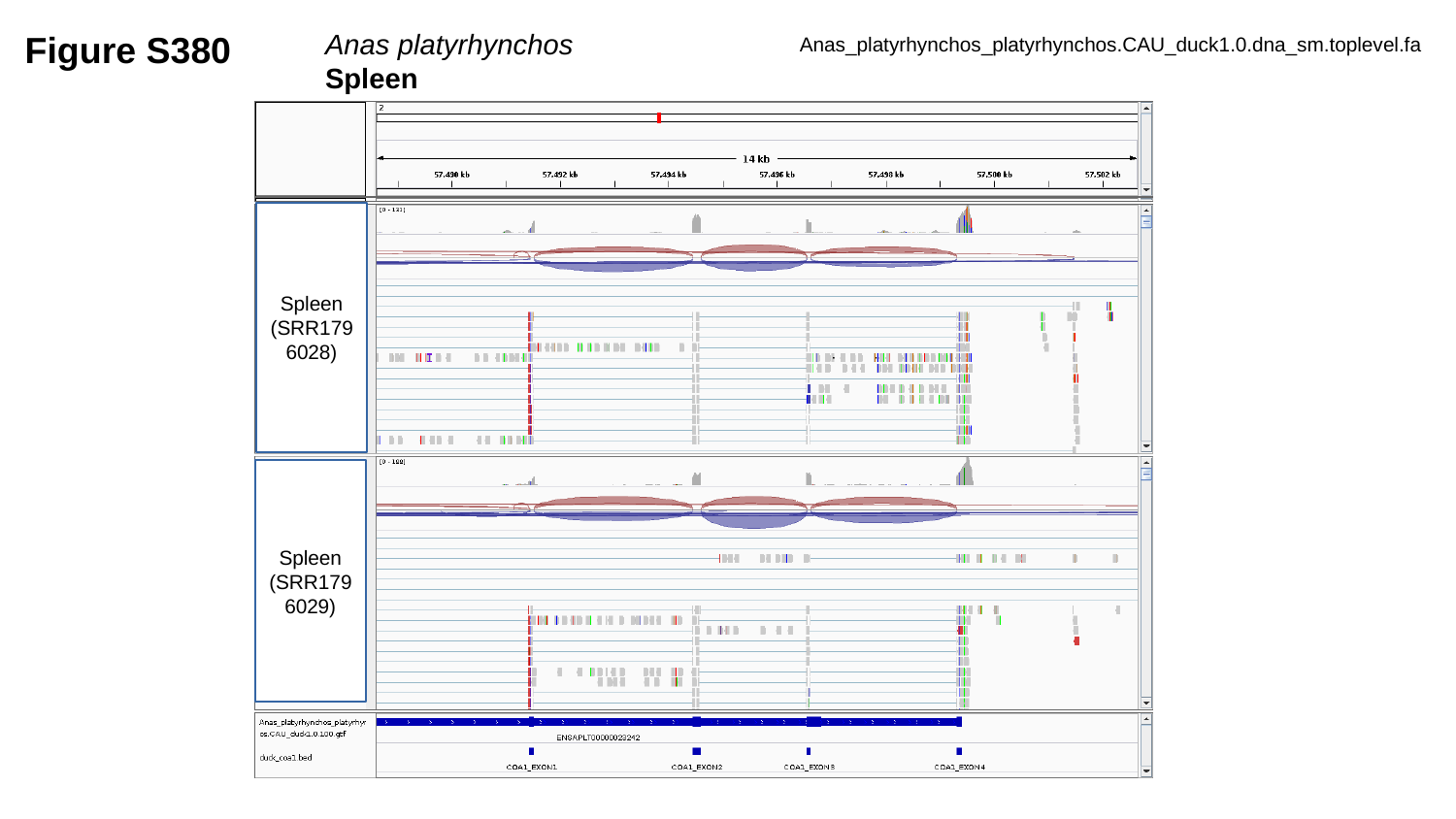

Figure S380
Anas platyrhynchos		Spleen
Anas_platyrhynchos_platyrhynchos.CAU_duck1.0.dna_sm.toplevel.fa
Spleen
(SRR1796028)
Spleen
(SRR1796029)

### Slide 381
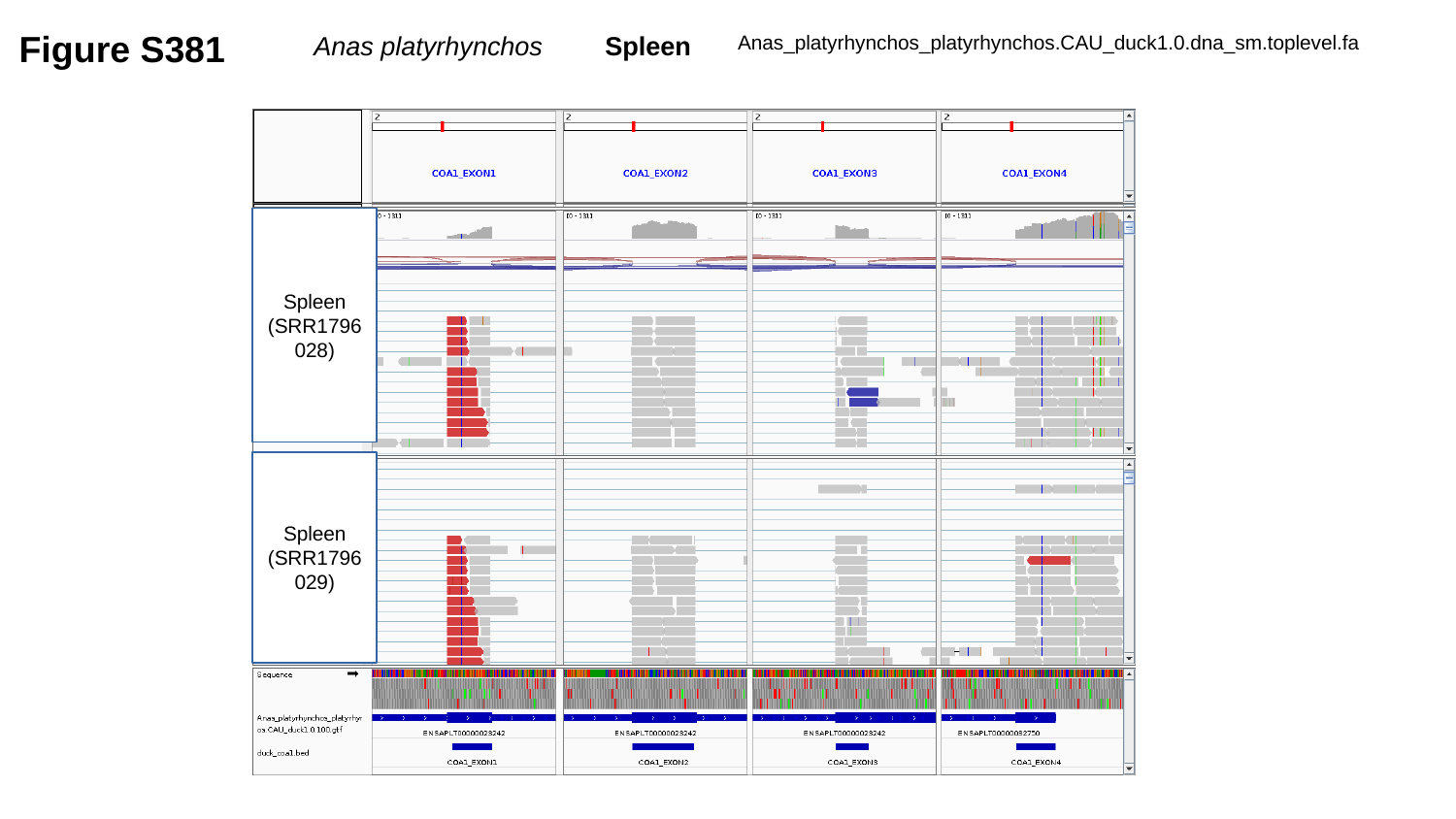

Figure S381
Anas platyrhynchos	Spleen
Anas_platyrhynchos_platyrhynchos.CAU_duck1.0.dna_sm.toplevel.fa
Spleen
(SRR1796028)
Spleen
(SRR1796029)

### Slide 382
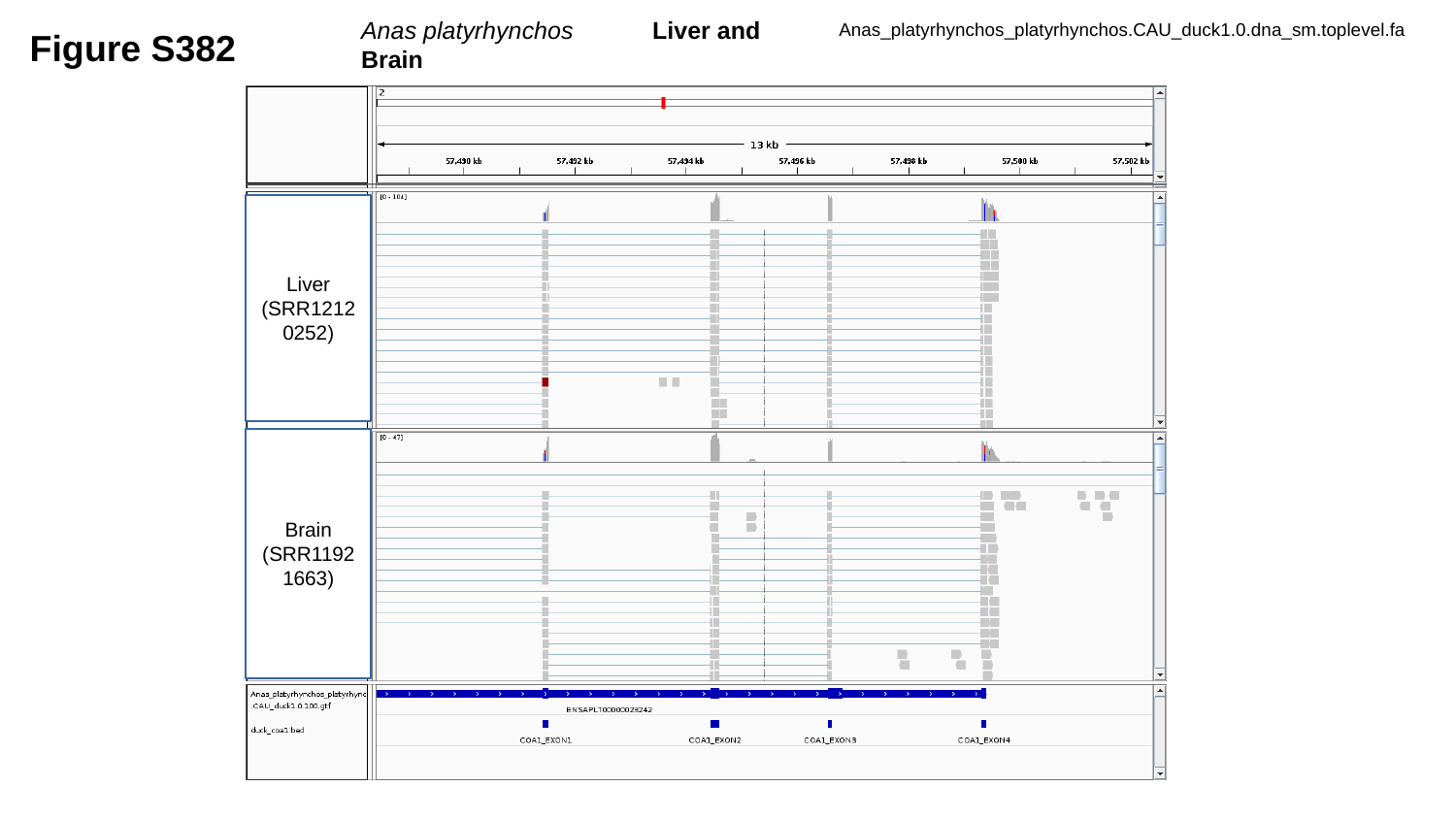

Anas platyrhynchos	Liver and Brain
Figure S382
Anas_platyrhynchos_platyrhynchos.CAU_duck1.0.dna_sm.toplevel.fa
Liver
(SRR12120252)
Brain
(SRR11921663)

### Slide 383
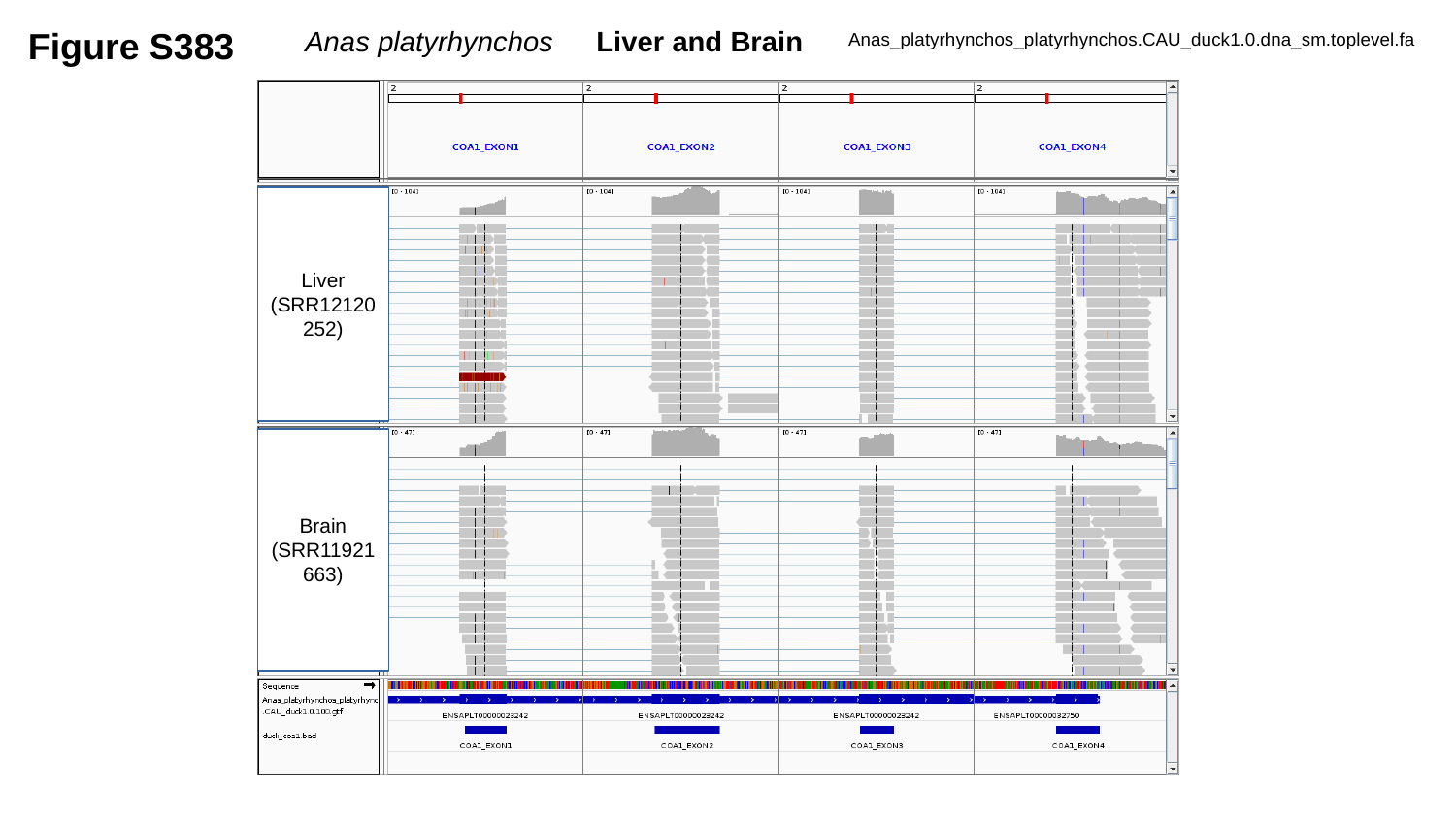

Figure S383
Anas platyrhynchos	Liver and Brain
Anas_platyrhynchos_platyrhynchos.CAU_duck1.0.dna_sm.toplevel.fa
Liver
(SRR12120252)
Brain
(SRR11921663)

### Slide 384
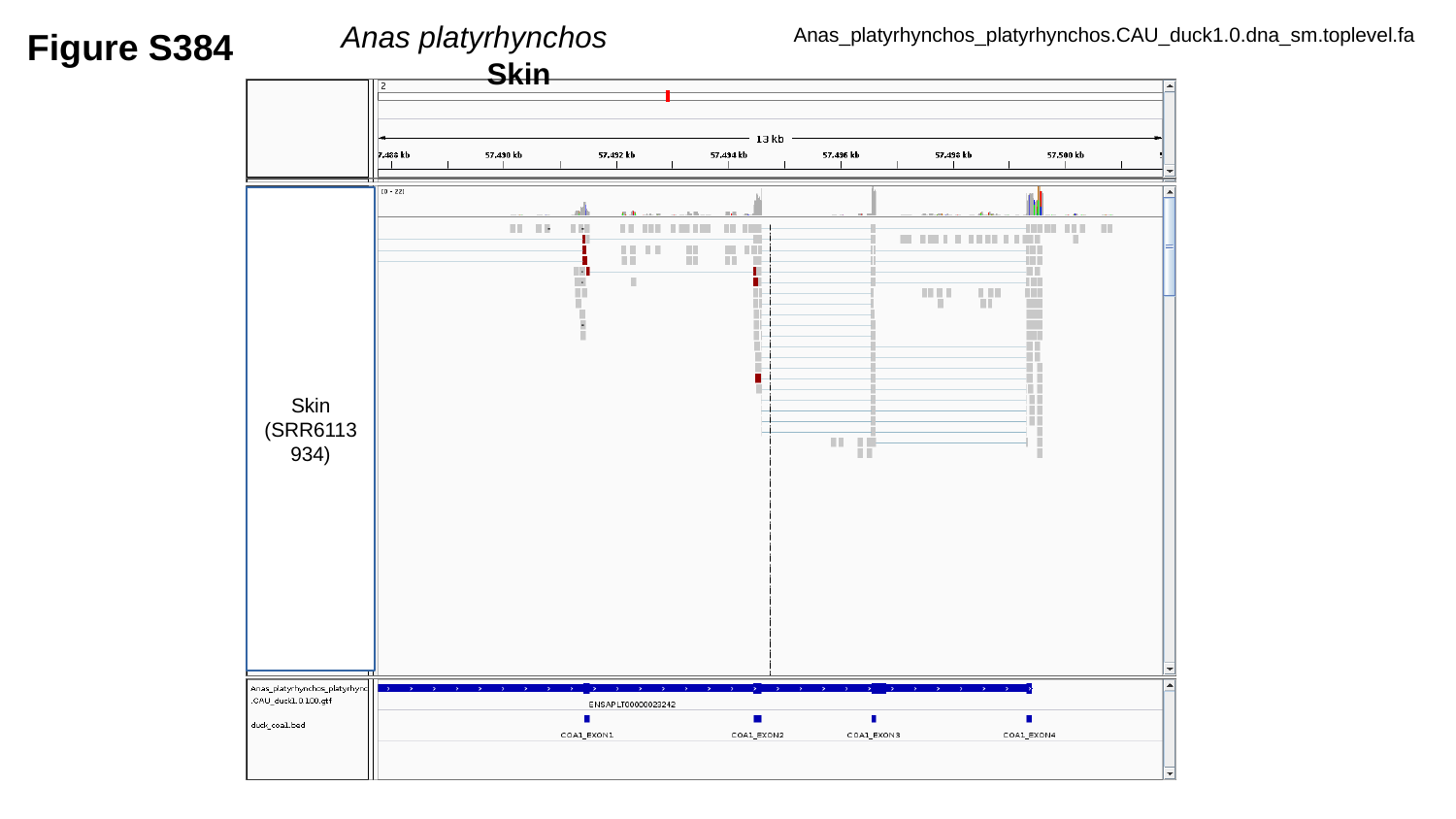

Figure S384
Anas platyrhynchos		Skin
Anas_platyrhynchos_platyrhynchos.CAU_duck1.0.dna_sm.toplevel.fa
Skin
(SRR6113934)

### Slide 385
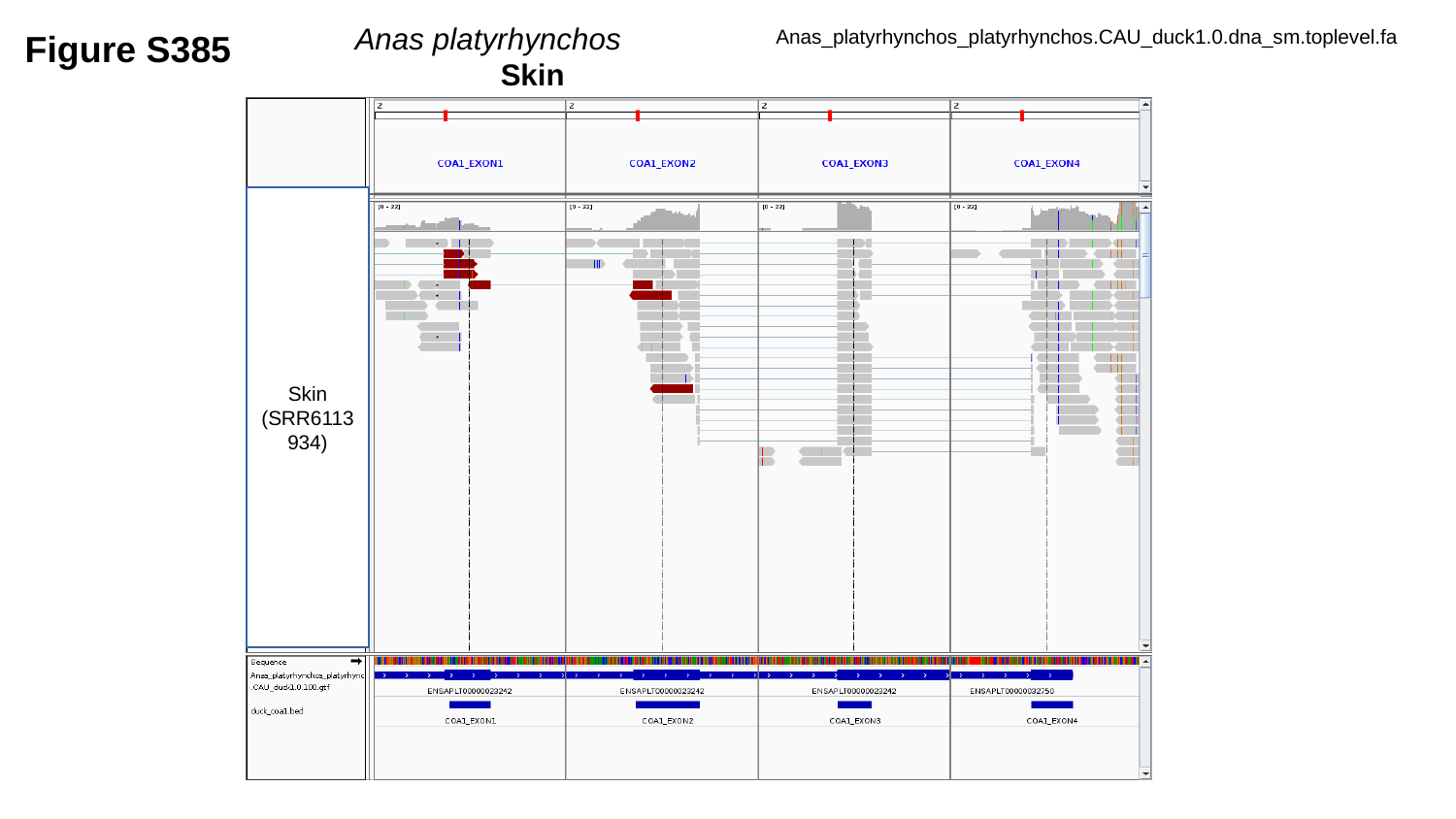

Figure S385
Anas platyrhynchos		Skin
Anas_platyrhynchos_platyrhynchos.CAU_duck1.0.dna_sm.toplevel.fa
Skin
(SRR6113934)

### Slide 386
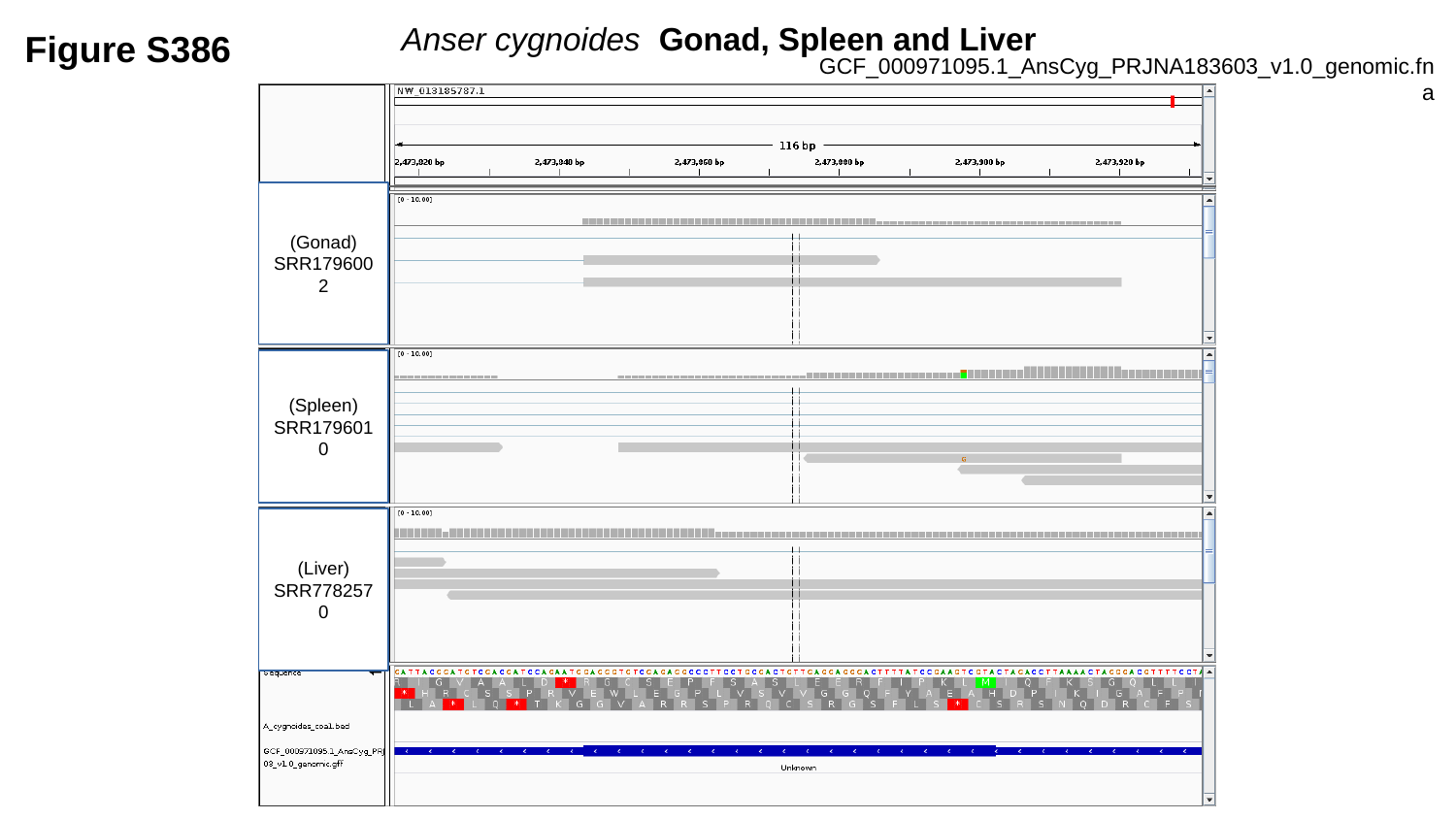

Anser cygnoides Gonad, Spleen and Liver
Figure S386
GCF_000971095.1_AnsCyg_PRJNA183603_v1.0_genomic.fna
(Gonad)
SRR1796002
(Spleen)
SRR1796010
(Liver)
SRR7782570

### Slide 387
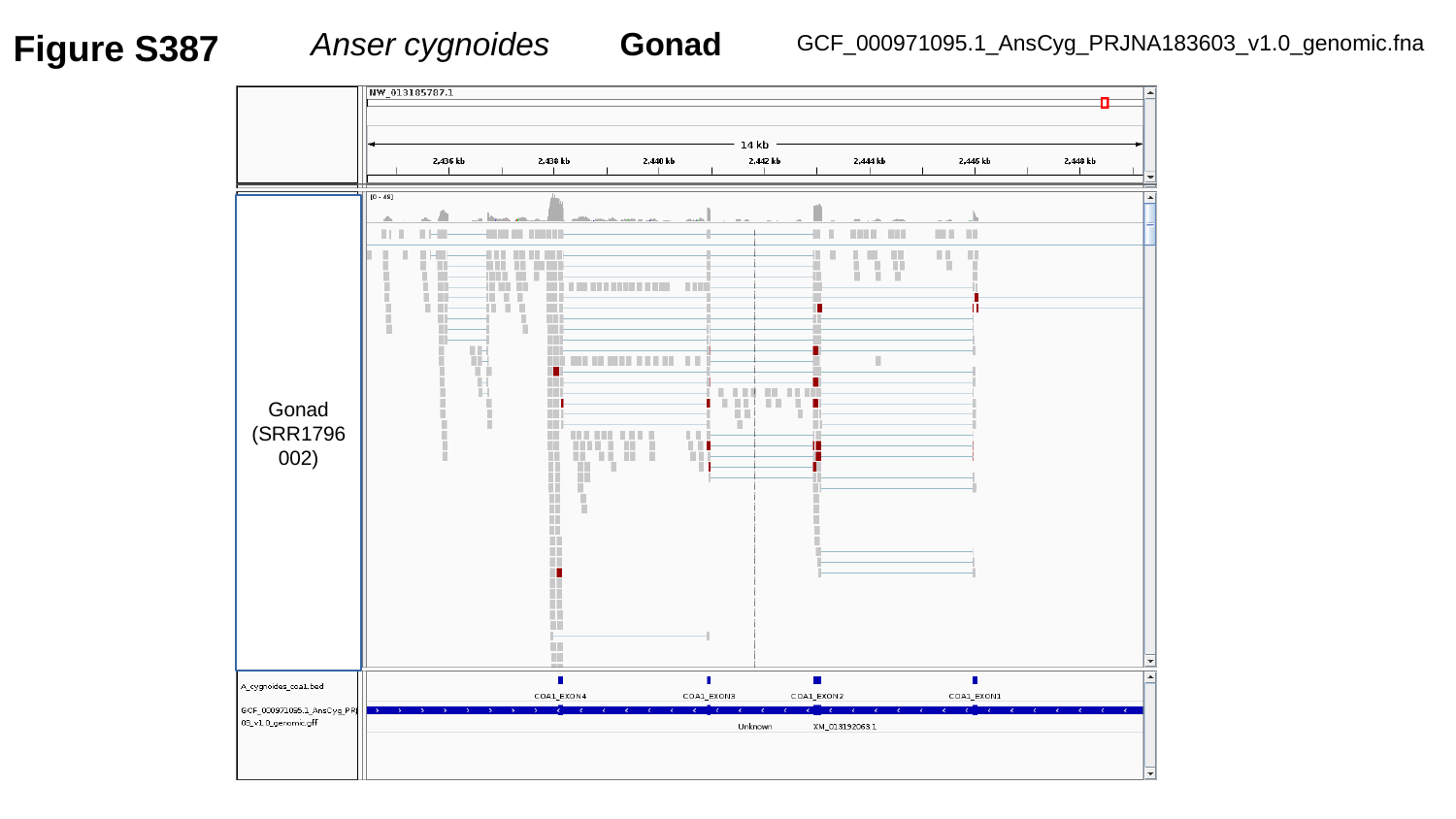

Figure S387
Anser cygnoides	 Gonad
GCF_000971095.1_AnsCyg_PRJNA183603_v1.0_genomic.fna
Gonad
(SRR1796002)

### Slide 388
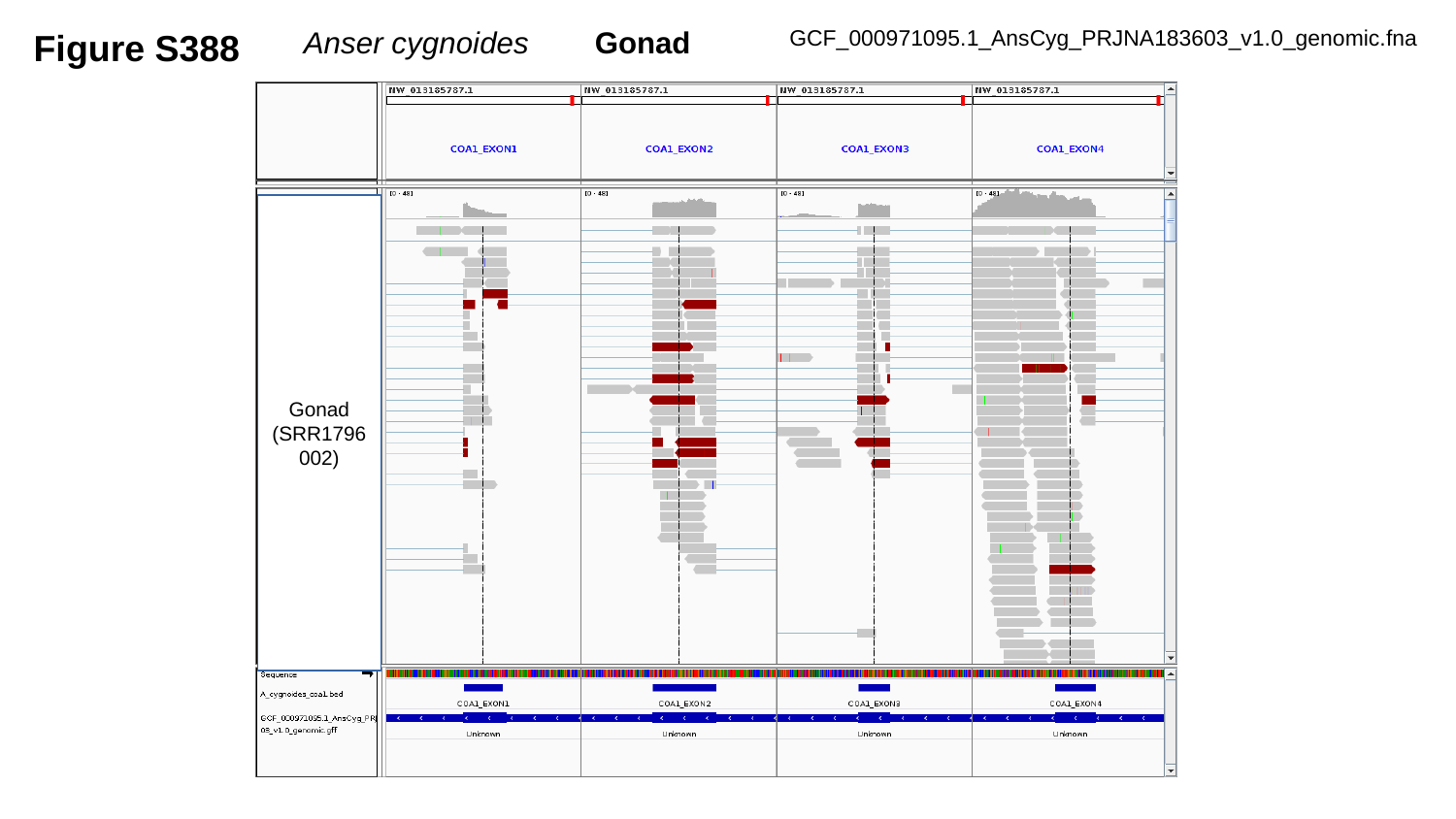

Figure S388
Anser cygnoides	Gonad
GCF_000971095.1_AnsCyg_PRJNA183603_v1.0_genomic.fna
Gonad
(SRR1796002)

### Slide 389
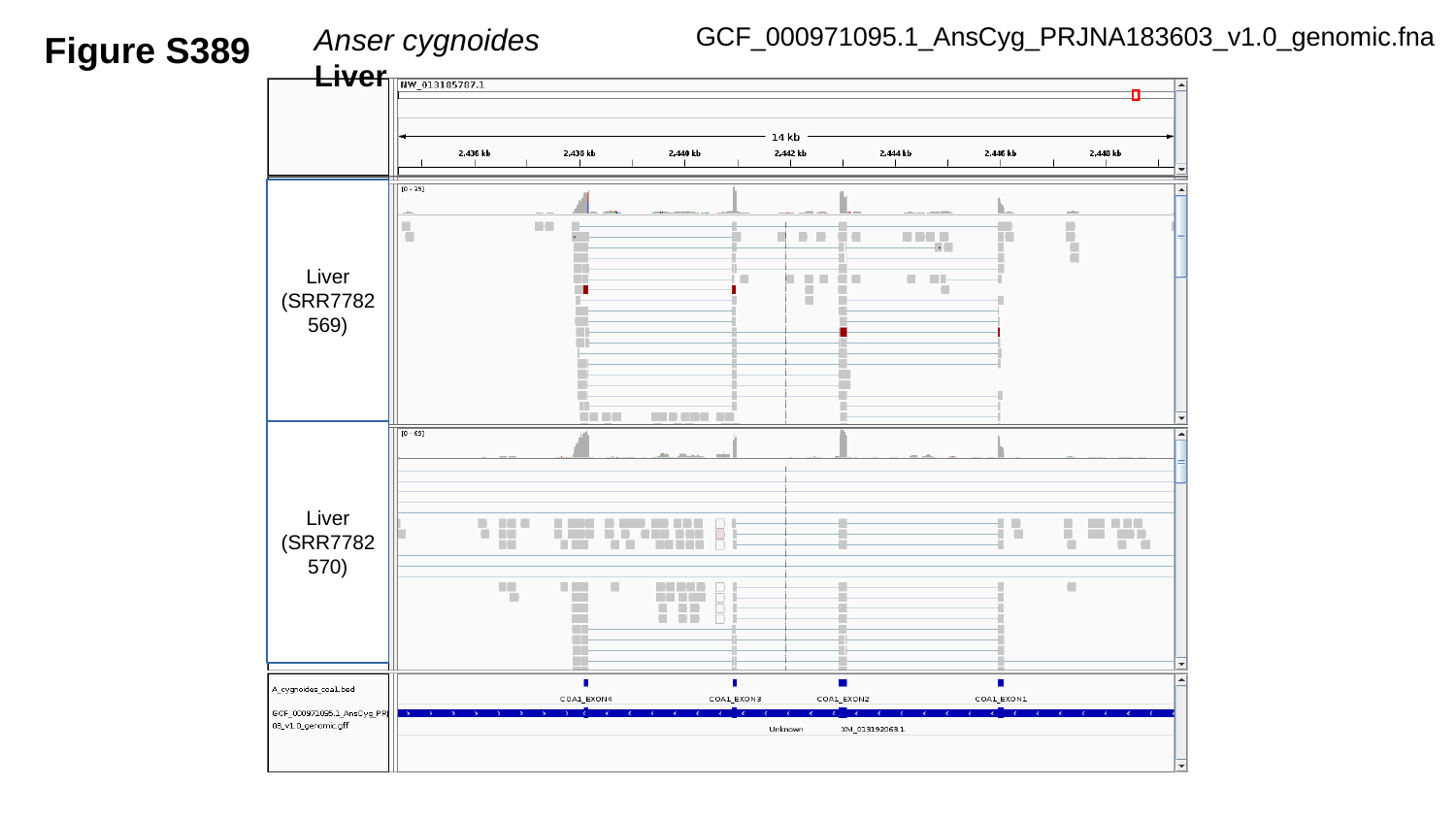

Figure S389
Anser cygnoides	Liver
GCF_000971095.1_AnsCyg_PRJNA183603_v1.0_genomic.fna
Liver
(SRR7782569)
Liver
(SRR7782570)

### Slide 390
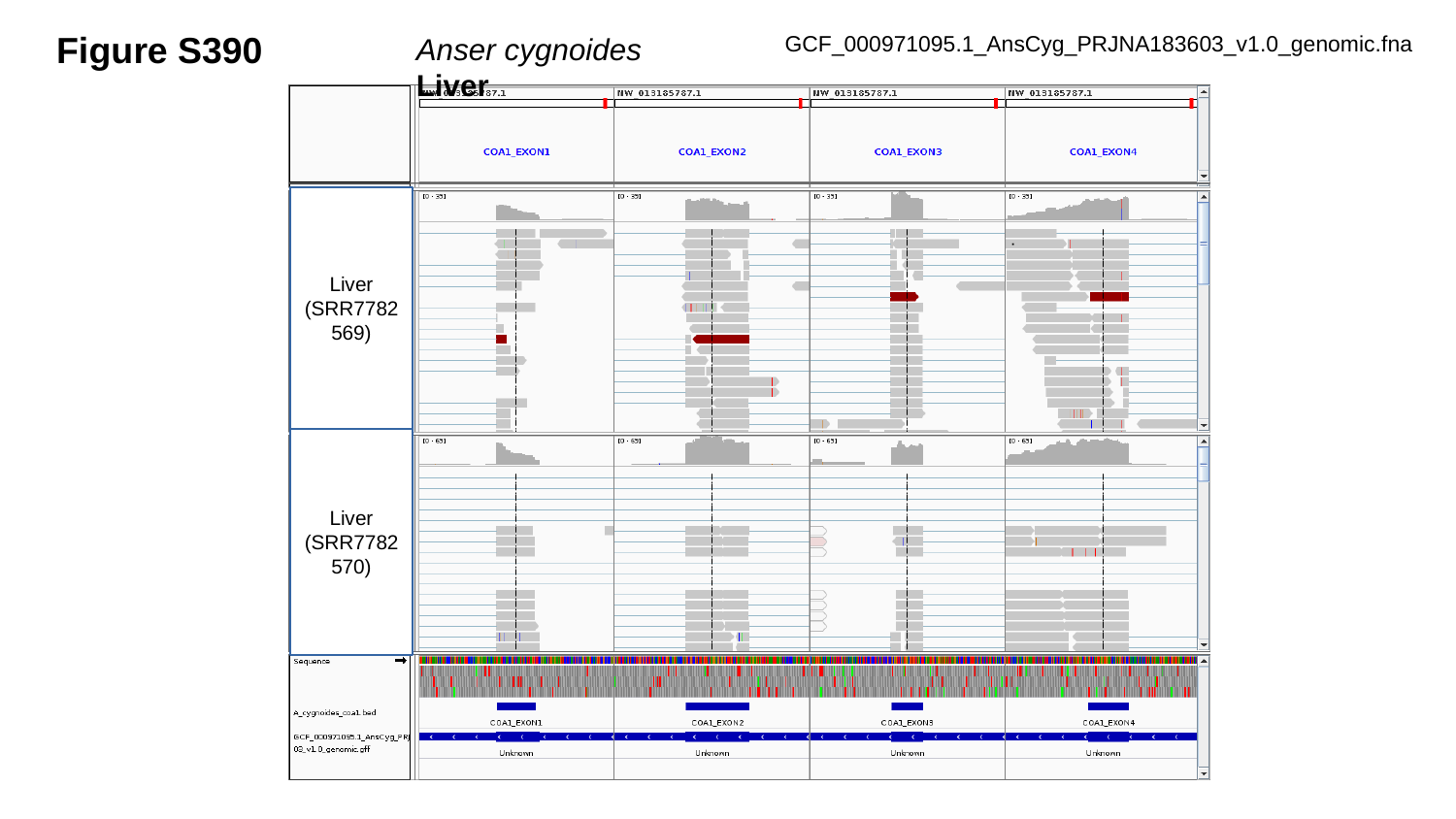

Figure S390
GCF_000971095.1_AnsCyg_PRJNA183603_v1.0_genomic.fna
Anser cygnoides	Liver
Liver
(SRR7782569)
Liver
(SRR7782570)

### Slide 391
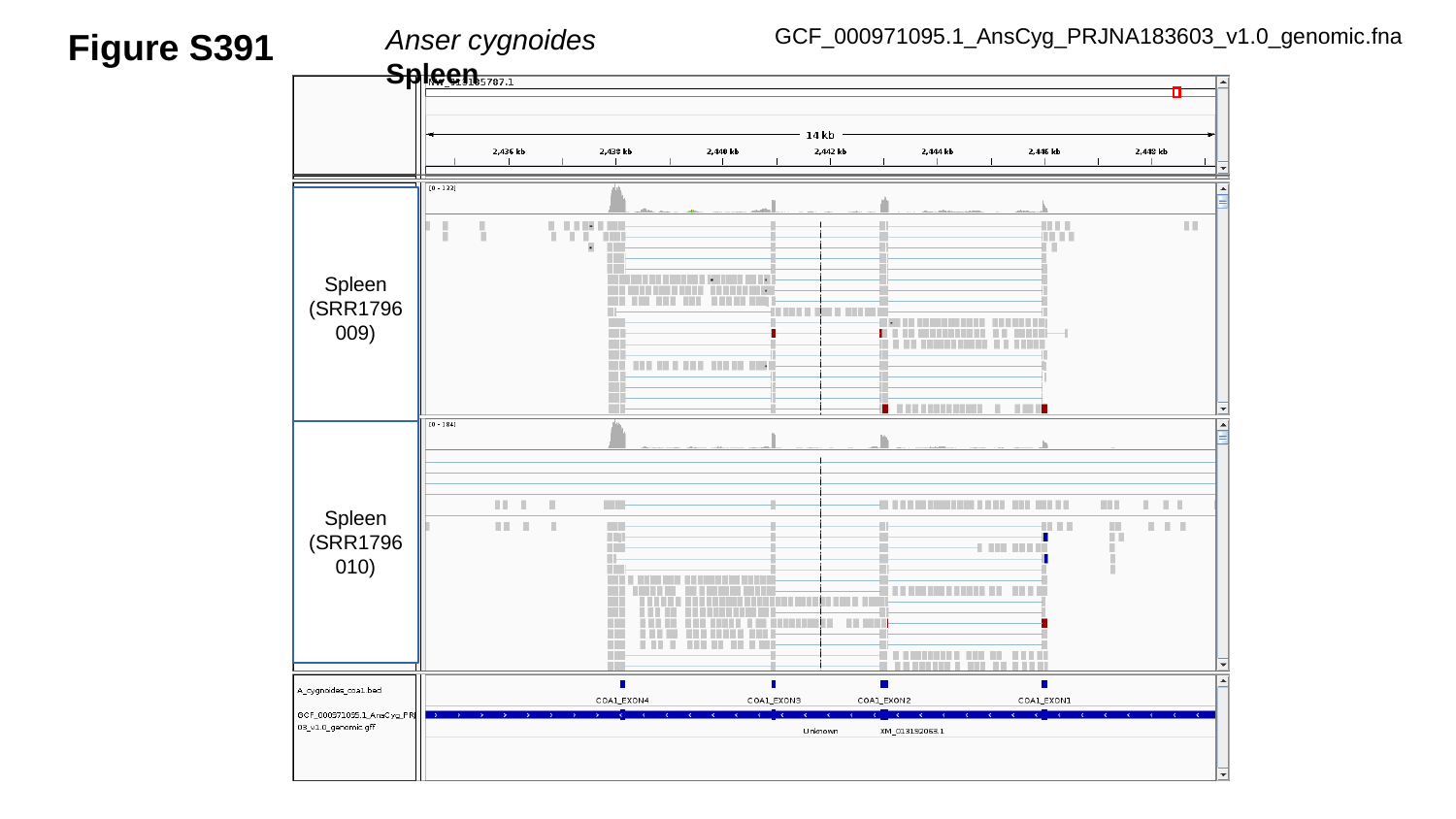

Figure S391
Anser cygnoides	Spleen
GCF_000971095.1_AnsCyg_PRJNA183603_v1.0_genomic.fna
Spleen
(SRR1796009)
Spleen
(SRR1796010)

### Slide 392
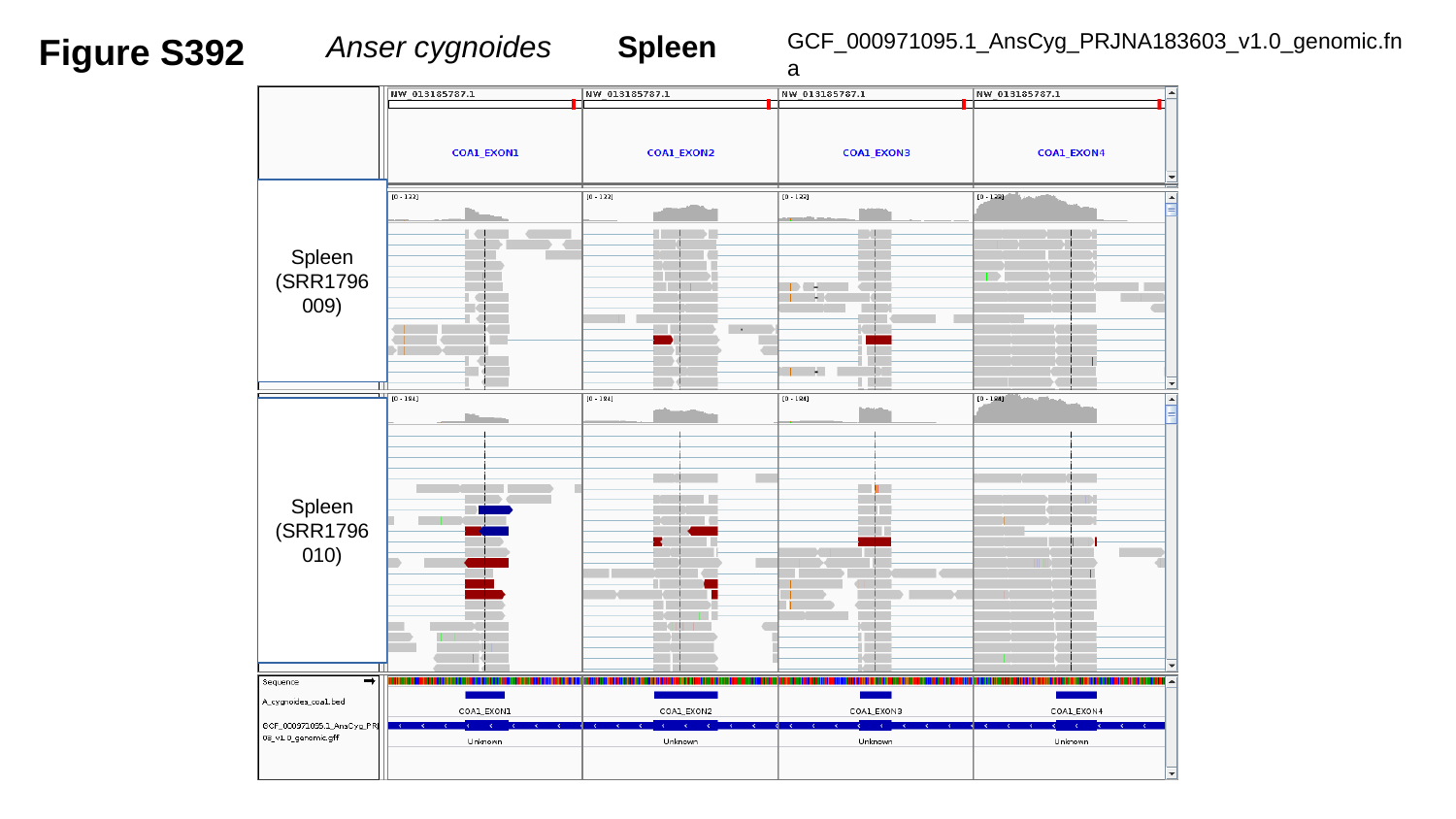

Figure S392
Anser cygnoides	Spleen
GCF_000971095.1_AnsCyg_PRJNA183603_v1.0_genomic.fna
Spleen
(SRR1796009)
Spleen
(SRR1796010)

### Slide 393
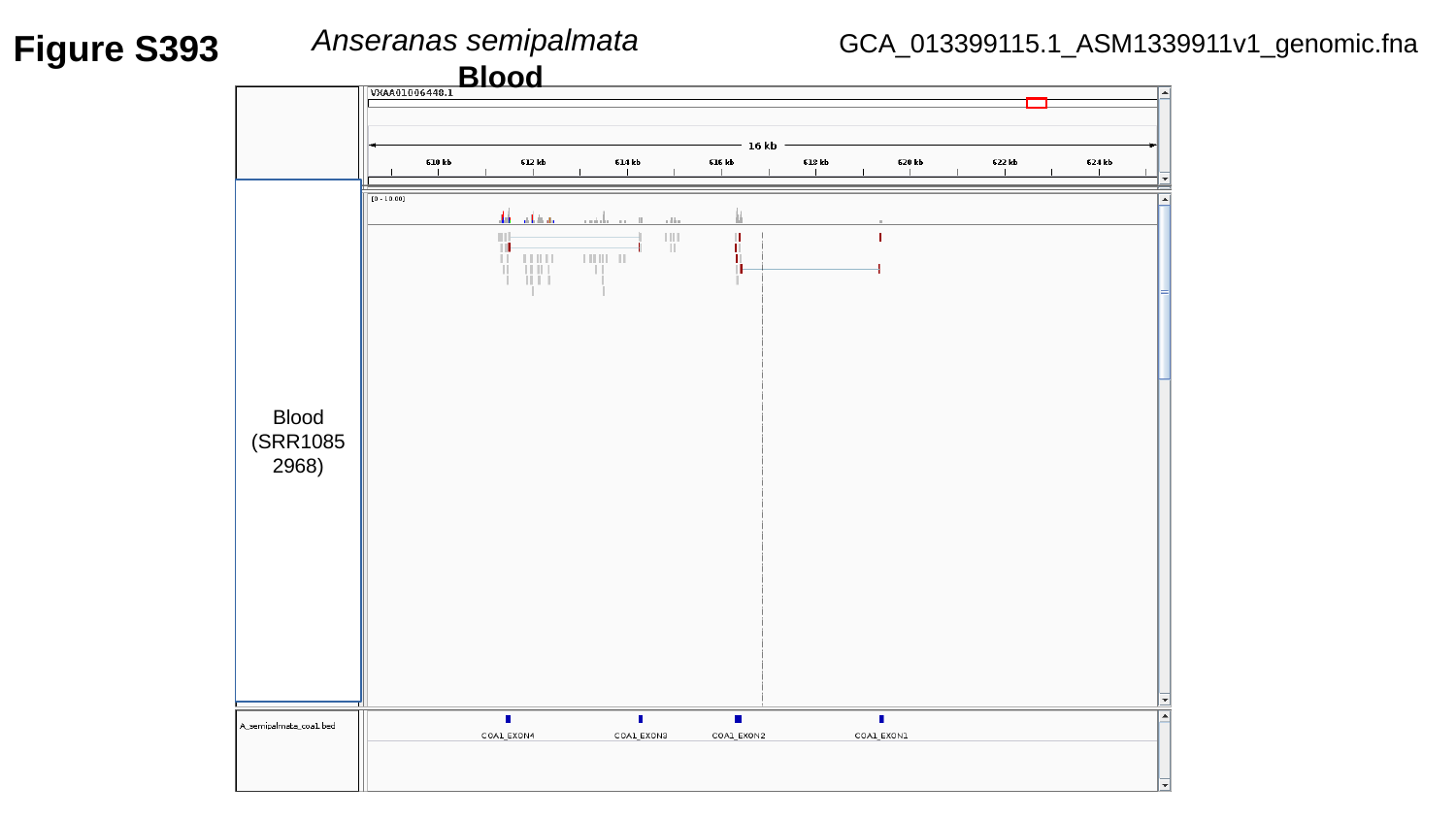

Figure S393
Anseranas semipalmata		Blood
GCA_013399115.1_ASM1339911v1_genomic.fna
Blood
(SRR10852968)

### Slide 394
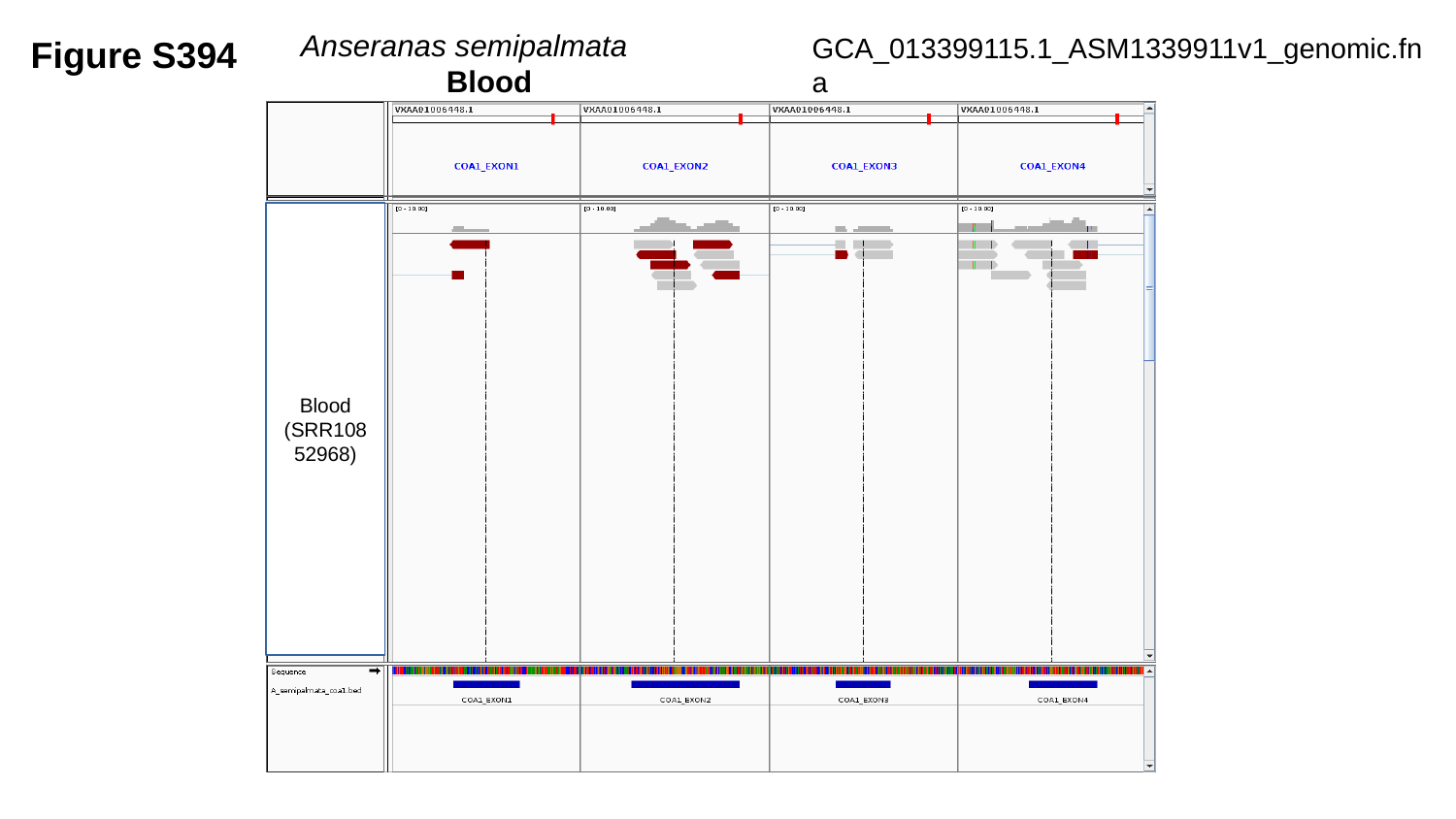

Figure S394
Anseranas semipalmata		Blood
GCA_013399115.1_ASM1339911v1_genomic.fna
Blood
(SRR10852968)

### Slide 395
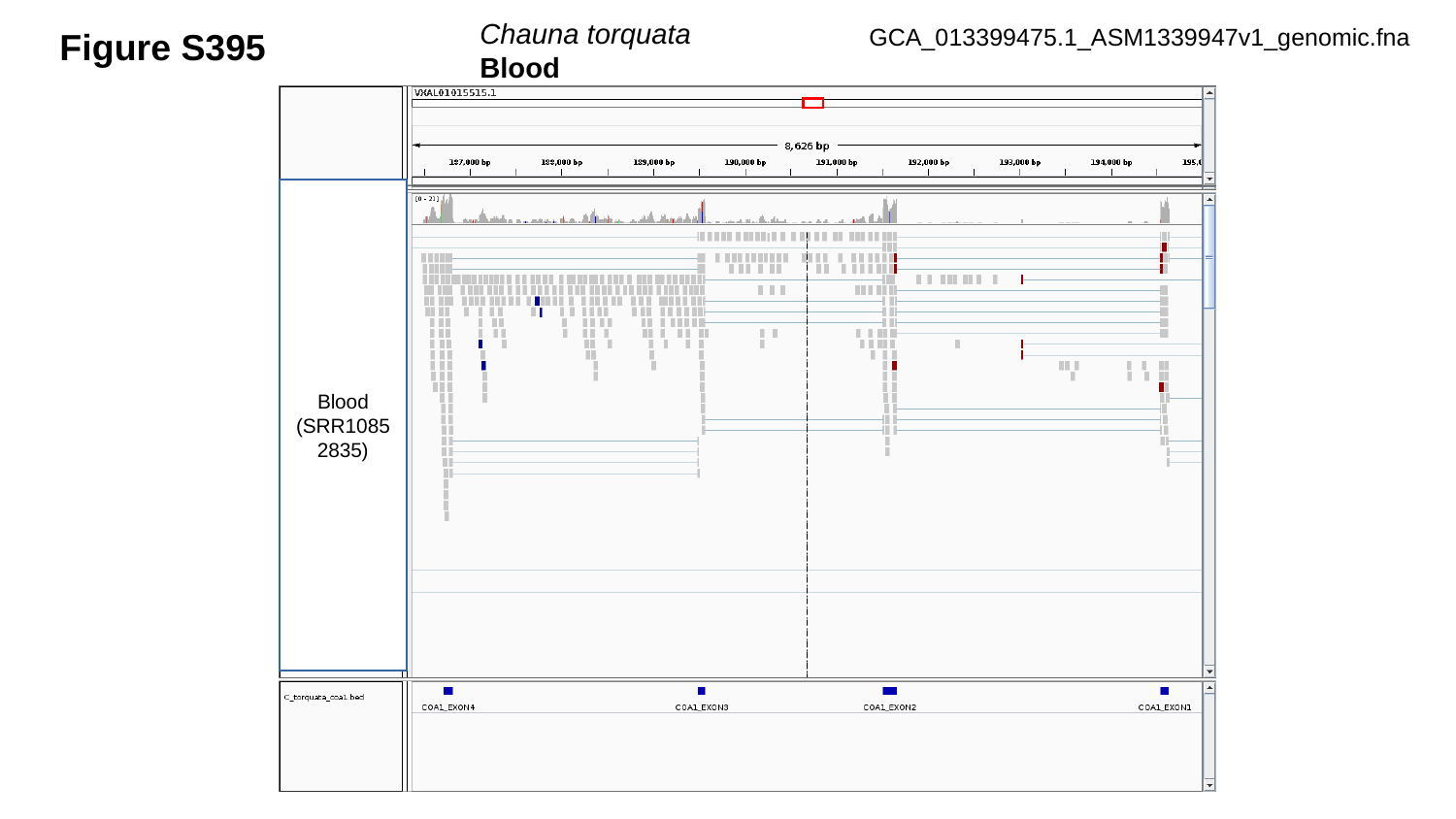

Chauna torquata	Blood
Figure S395
GCA_013399475.1_ASM1339947v1_genomic.fna
Blood
(SRR10852835)

### Slide 396
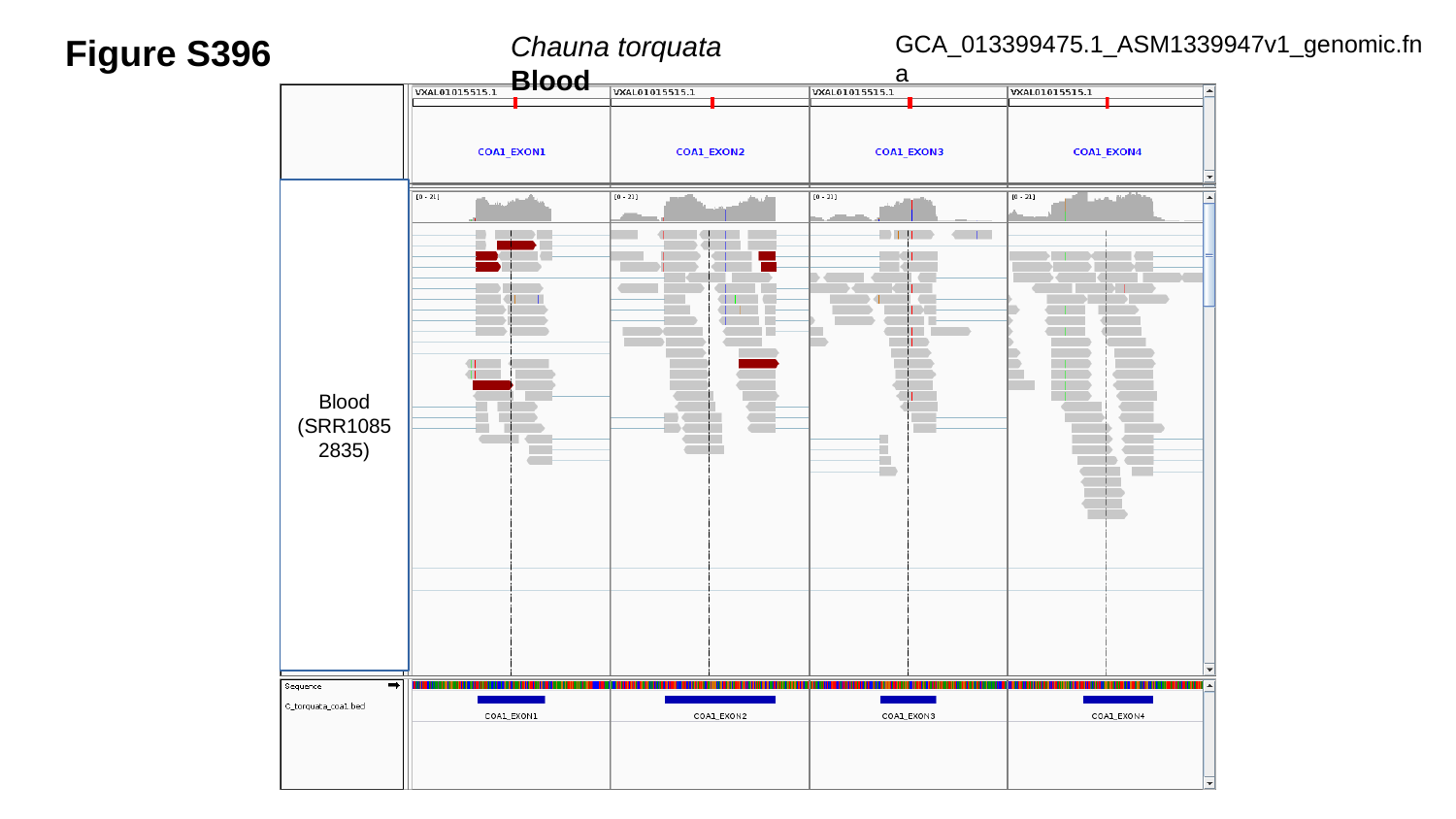

Figure S396
Chauna torquata	Blood
GCA_013399475.1_ASM1339947v1_genomic.fna
Blood
(SRR10852835)

### Slide 397
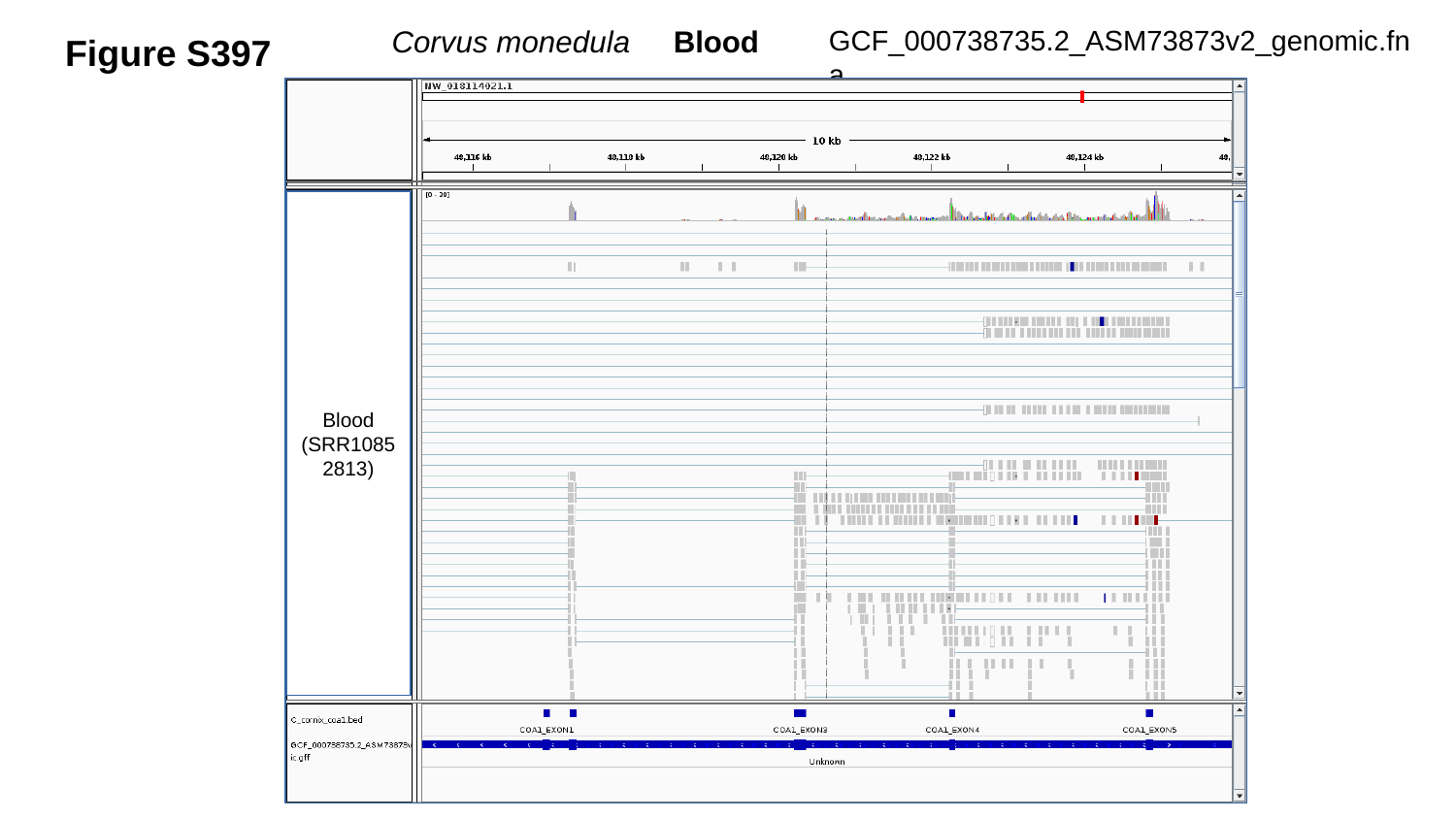

GCF_000738735.2_ASM73873v2_genomic.fna
Corvus monedula Blood
Figure S397
Blood
(SRR10852813)

### Slide 398
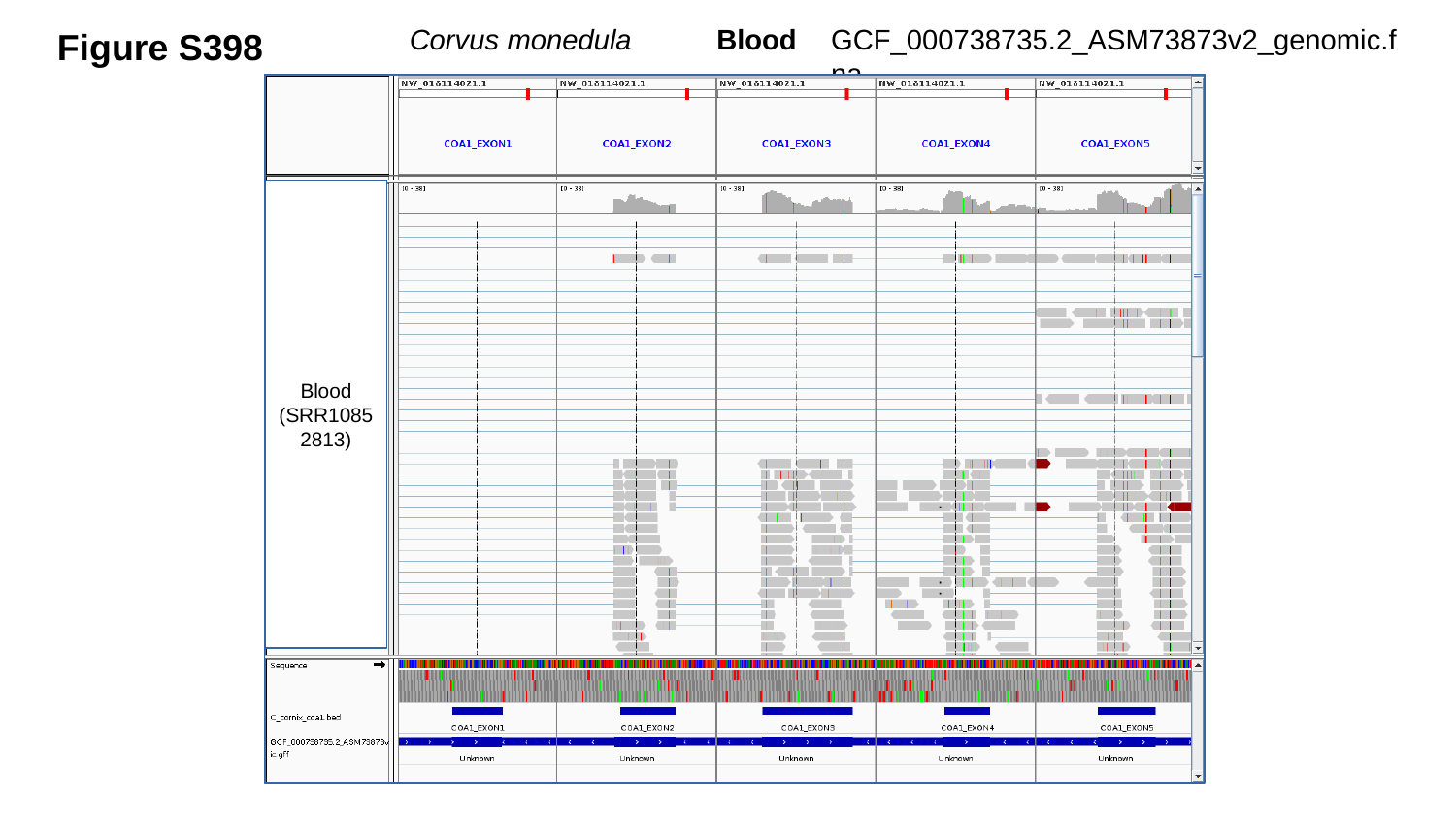

Figure S398
GCF_000738735.2_ASM73873v2_genomic.fna
Corvus monedula	 Blood
Blood
(SRR10852813)

### Slide 399
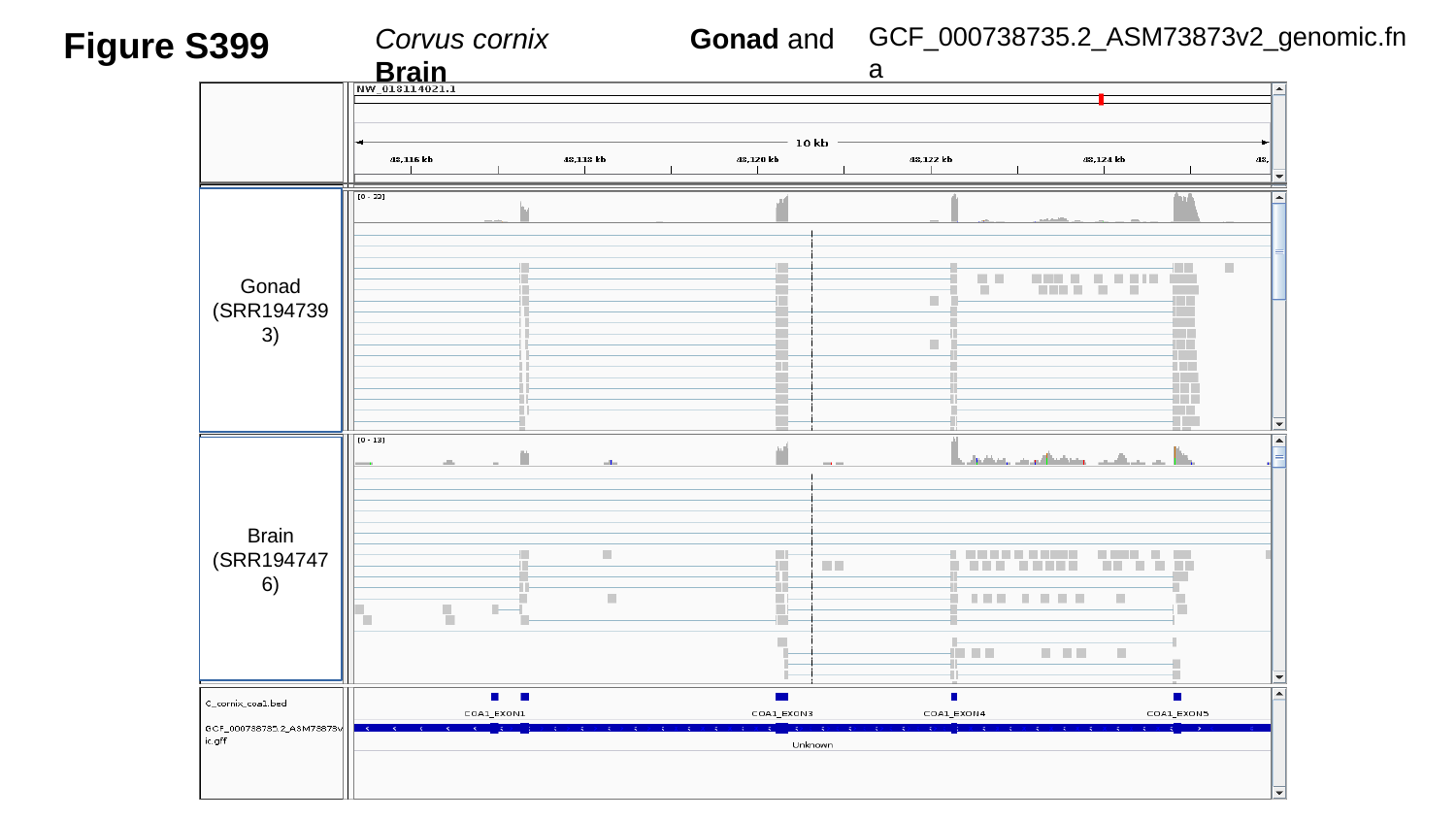

Figure S399
Corvus cornix	 Gonad and Brain
GCF_000738735.2_ASM73873v2_genomic.fna
Gonad
(SRR1947393)
Brain
(SRR1947476)

### Slide 400
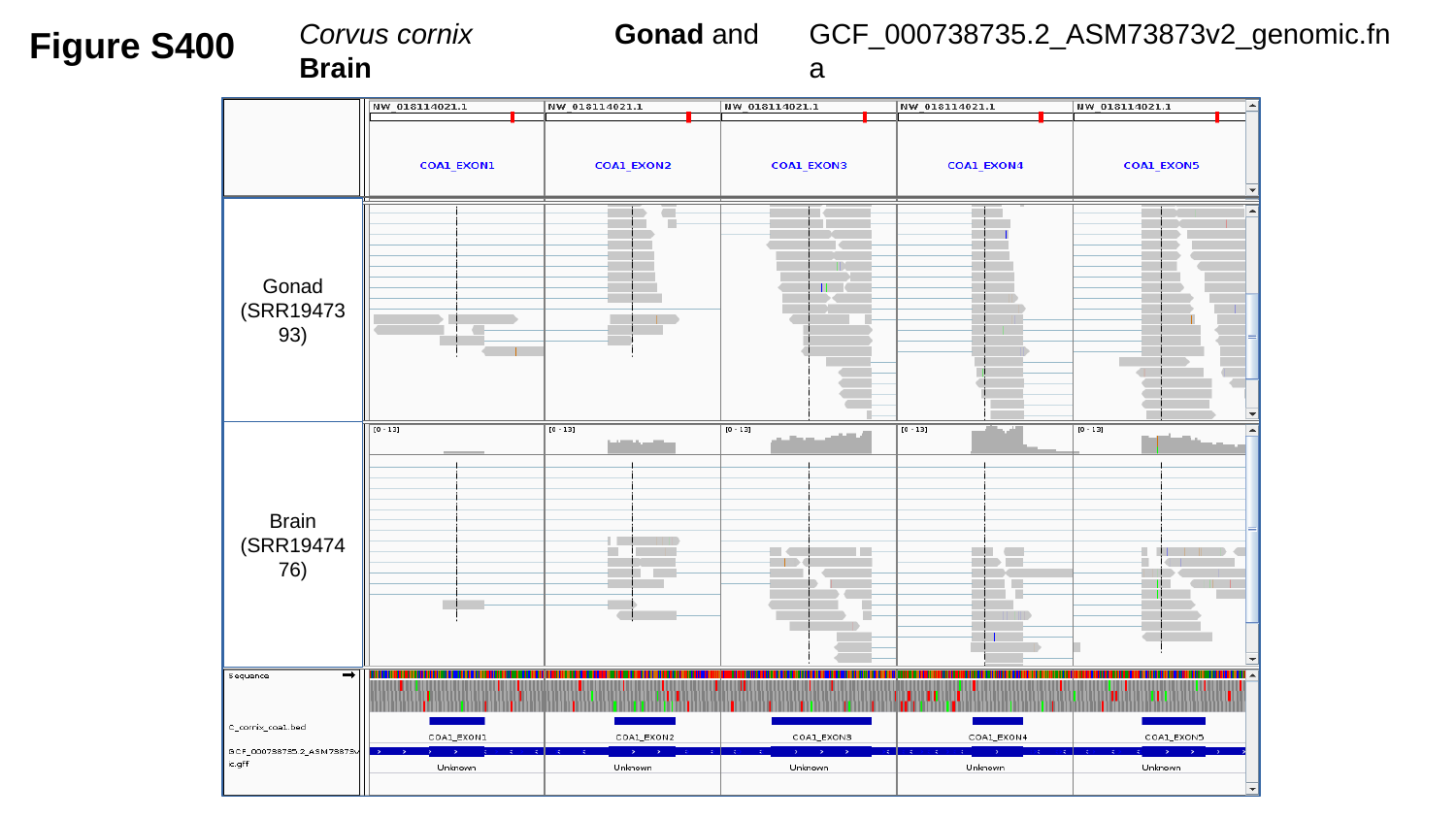

Figure S400
Corvus cornix	 Gonad and Brain
GCF_000738735.2_ASM73873v2_genomic.fna
Gonad
(SRR1947393)
Brain
(SRR1947476)

### Slide 401
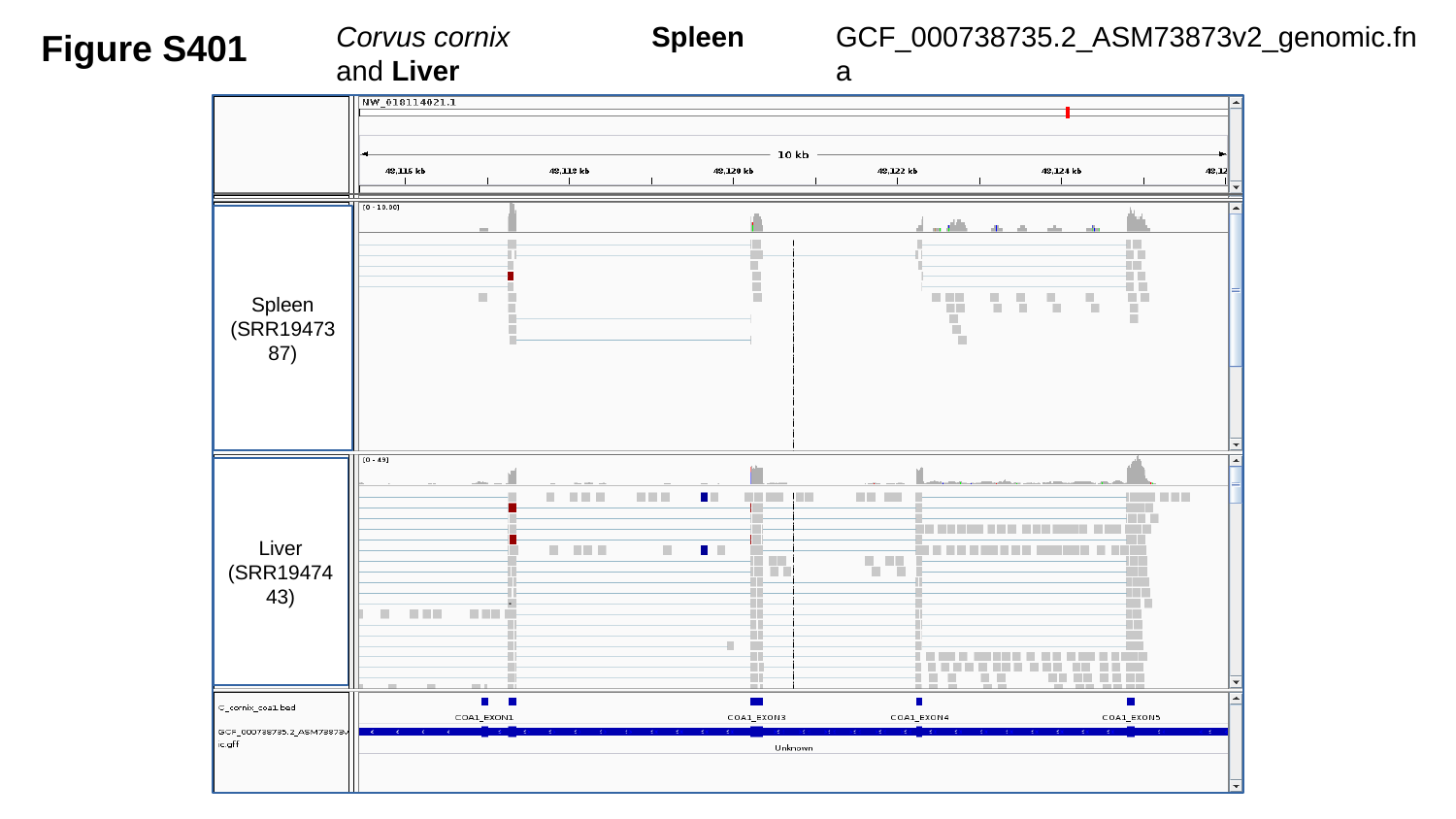

GCF_000738735.2_ASM73873v2_genomic.fna
Figure S401
Corvus cornix	 Spleen and Liver
Spleen
(SRR1947387)
Liver
(SRR1947443)

### Slide 402
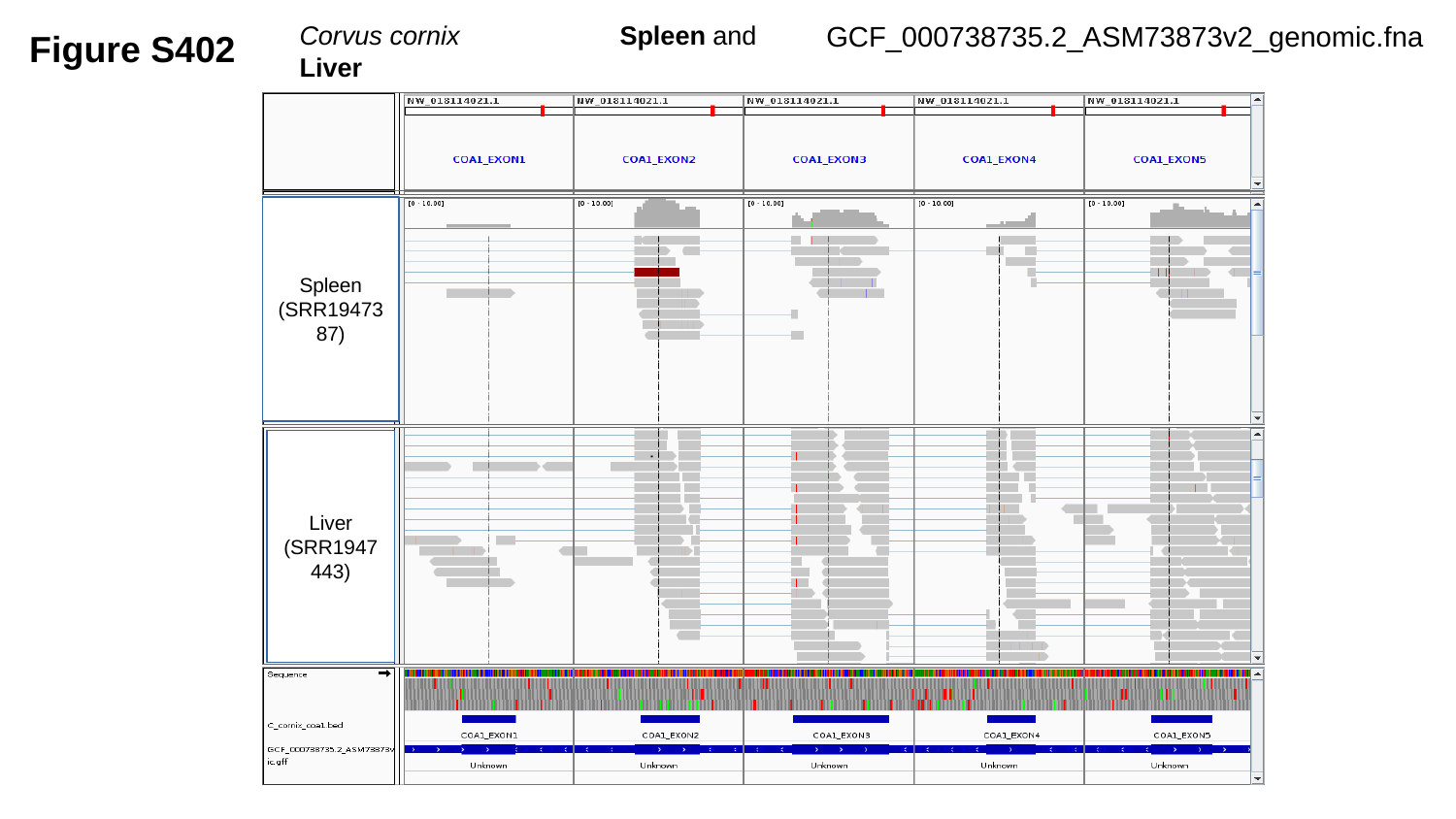

GCF_000738735.2_ASM73873v2_genomic.fna
Figure S402
Corvus cornix	 Spleen and Liver
Spleen
(SRR1947387)
Liver
(SRR1947443)

### Slide 403
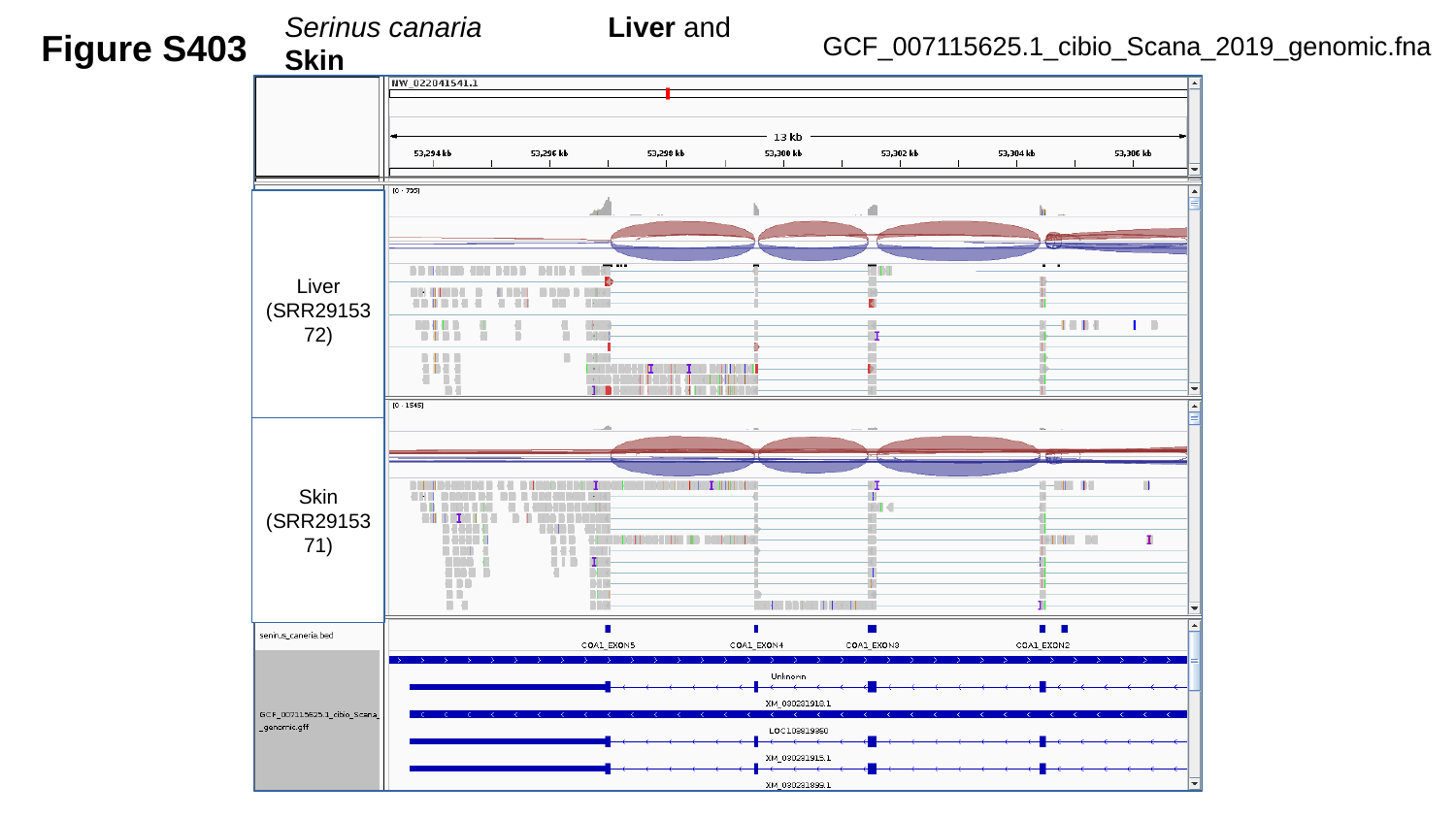

Serinus canaria	 Liver and Skin
Figure S403
GCF_007115625.1_cibio_Scana_2019_genomic.fna
Liver
(SRR2915372)
Skin
(SRR2915371)

### Slide 404
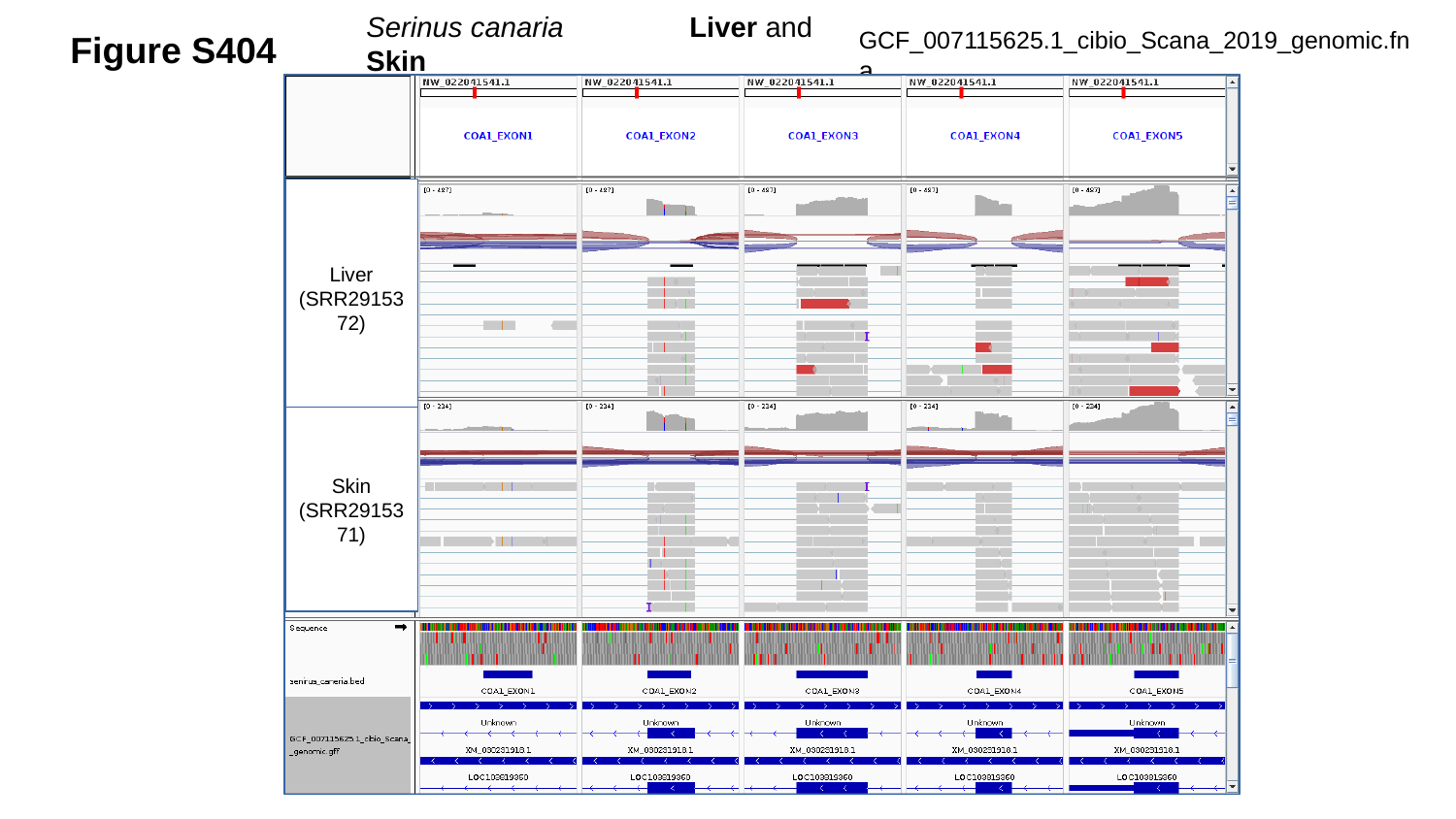

Serinus canaria	 Liver and Skin
Figure S404
GCF_007115625.1_cibio_Scana_2019_genomic.fna
Liver
(SRR2915372)
Skin
(SRR2915371)

### Slide 405
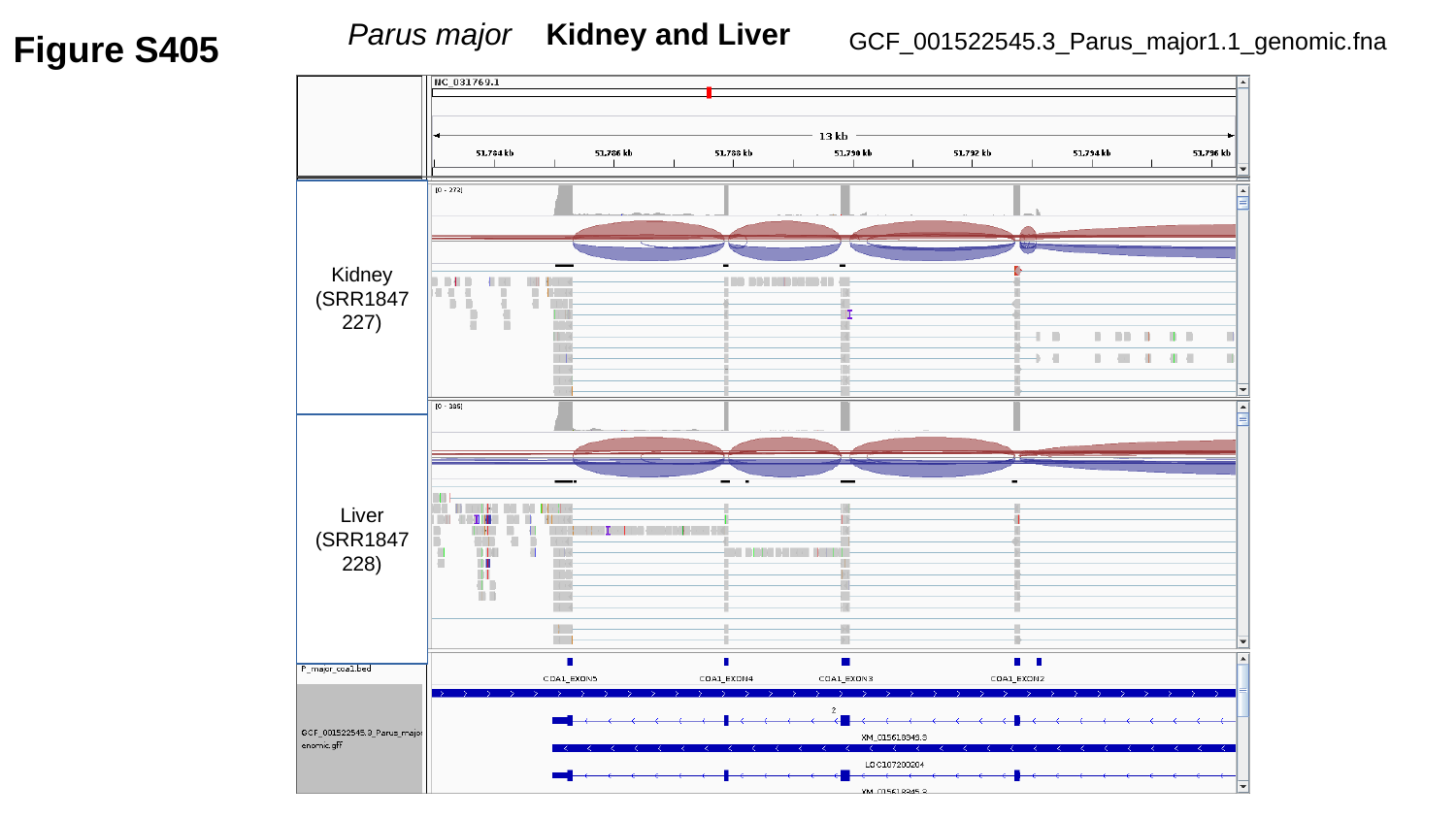

Parus major Kidney and Liver
Figure S405
GCF_001522545.3_Parus_major1.1_genomic.fna
Kidney
(SRR1847227)
Liver
(SRR1847228)

### Slide 406
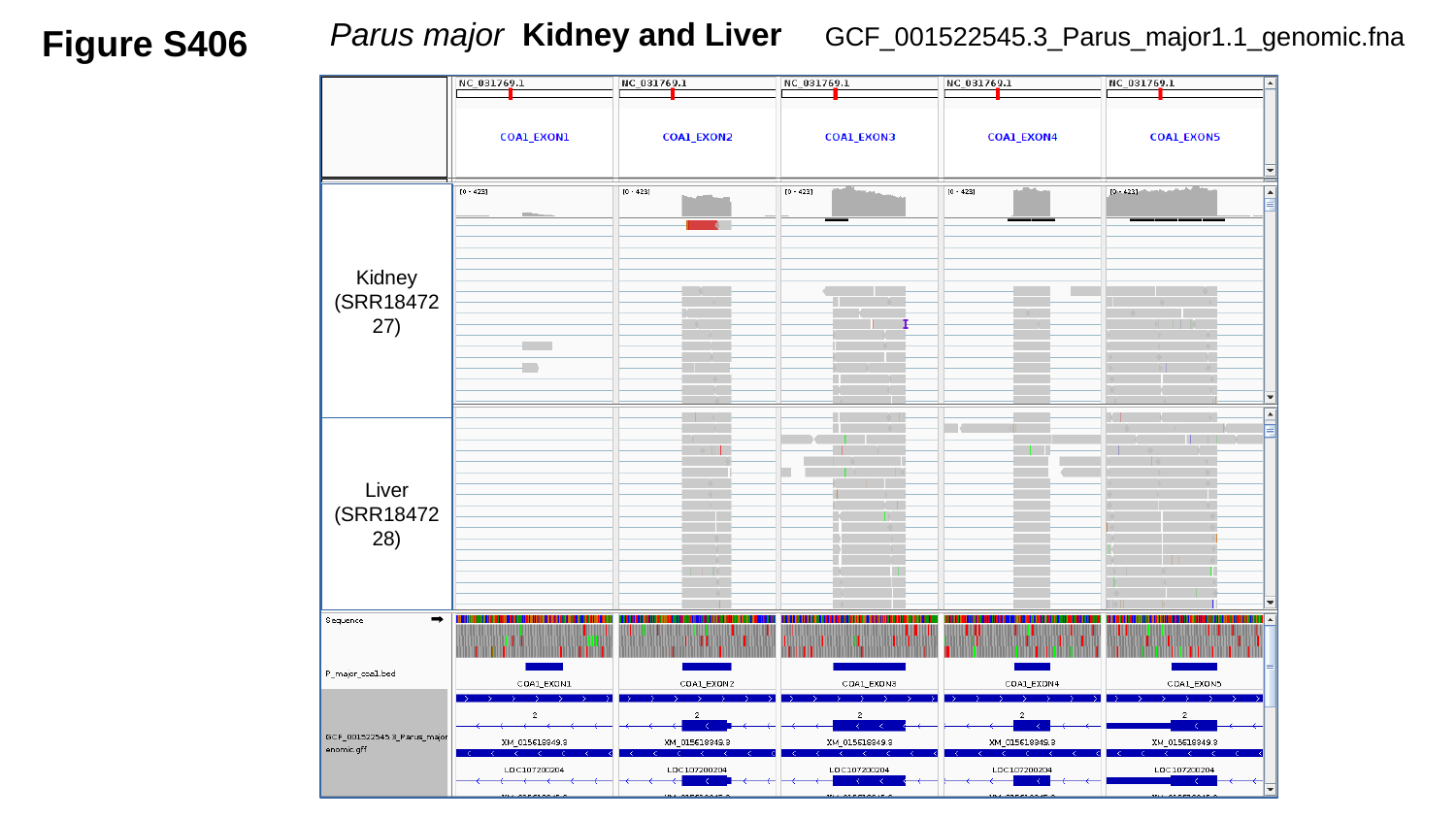

Figure S406
GCF_001522545.3_Parus_major1.1_genomic.fna
 Parus major Kidney and Liver
Kidney
(SRR1847227)
Liver
(SRR1847228)

### Slide 407
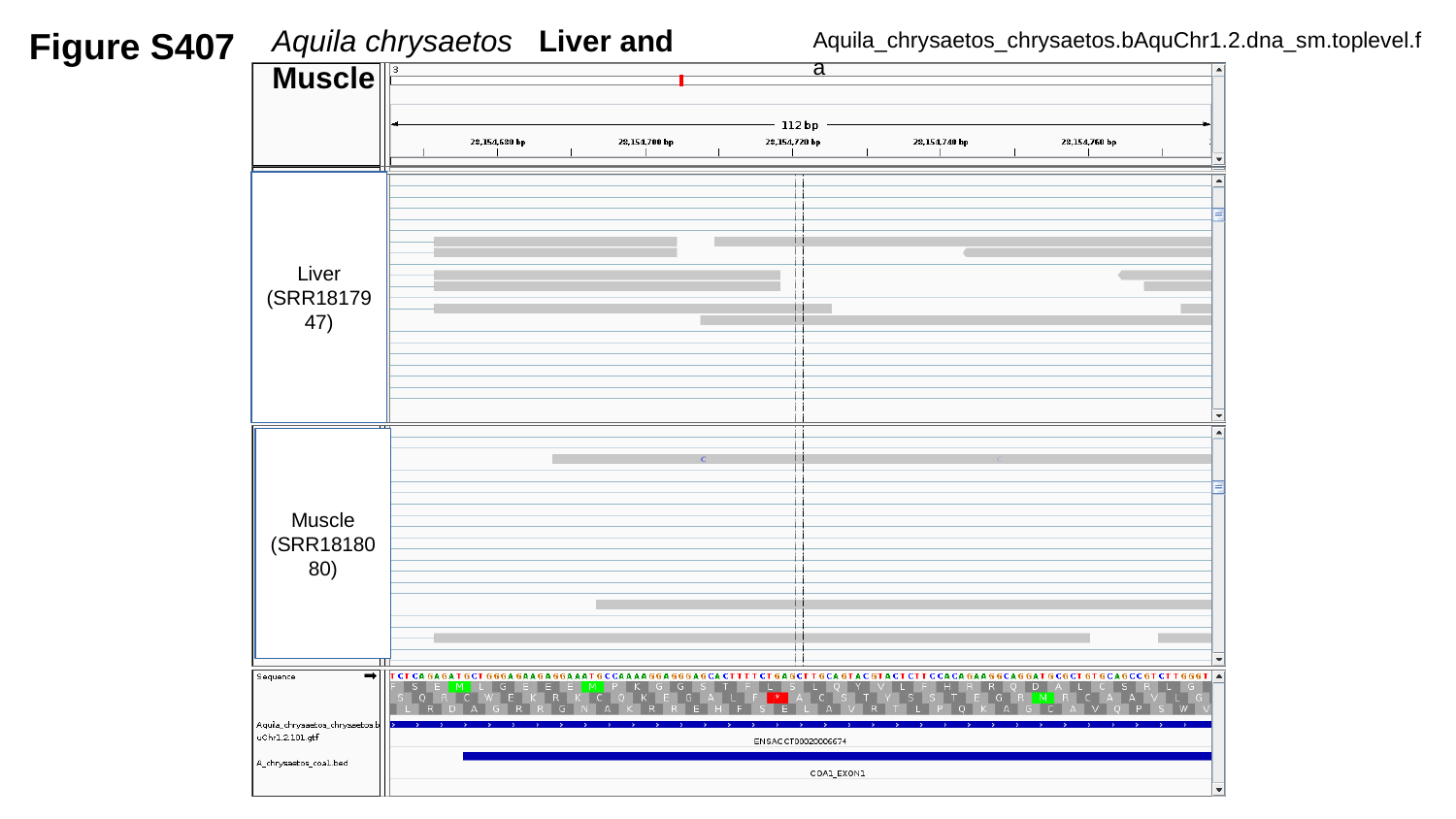

Figure S407
Aquila chrysaetos Liver and Muscle
Aquila_chrysaetos_chrysaetos.bAquChr1.2.dna_sm.toplevel.fa
Liver
(SRR1817947)
Muscle
(SRR1818080)

### Slide 408
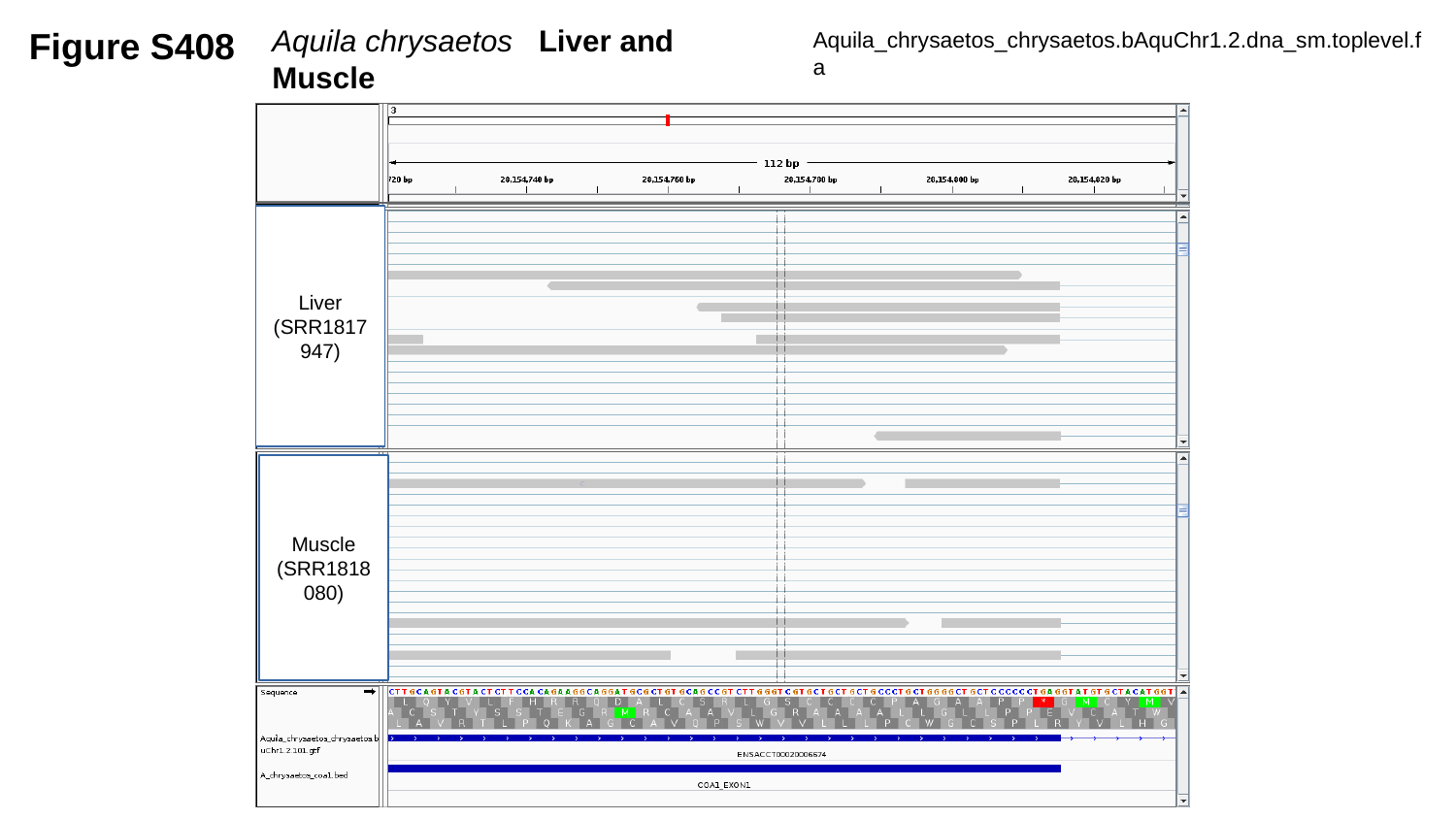

Figure S408
Aquila chrysaetos Liver and Muscle
Aquila_chrysaetos_chrysaetos.bAquChr1.2.dna_sm.toplevel.fa
Liver
(SRR1817947)
Muscle
(SRR1818080)

### Slide 409
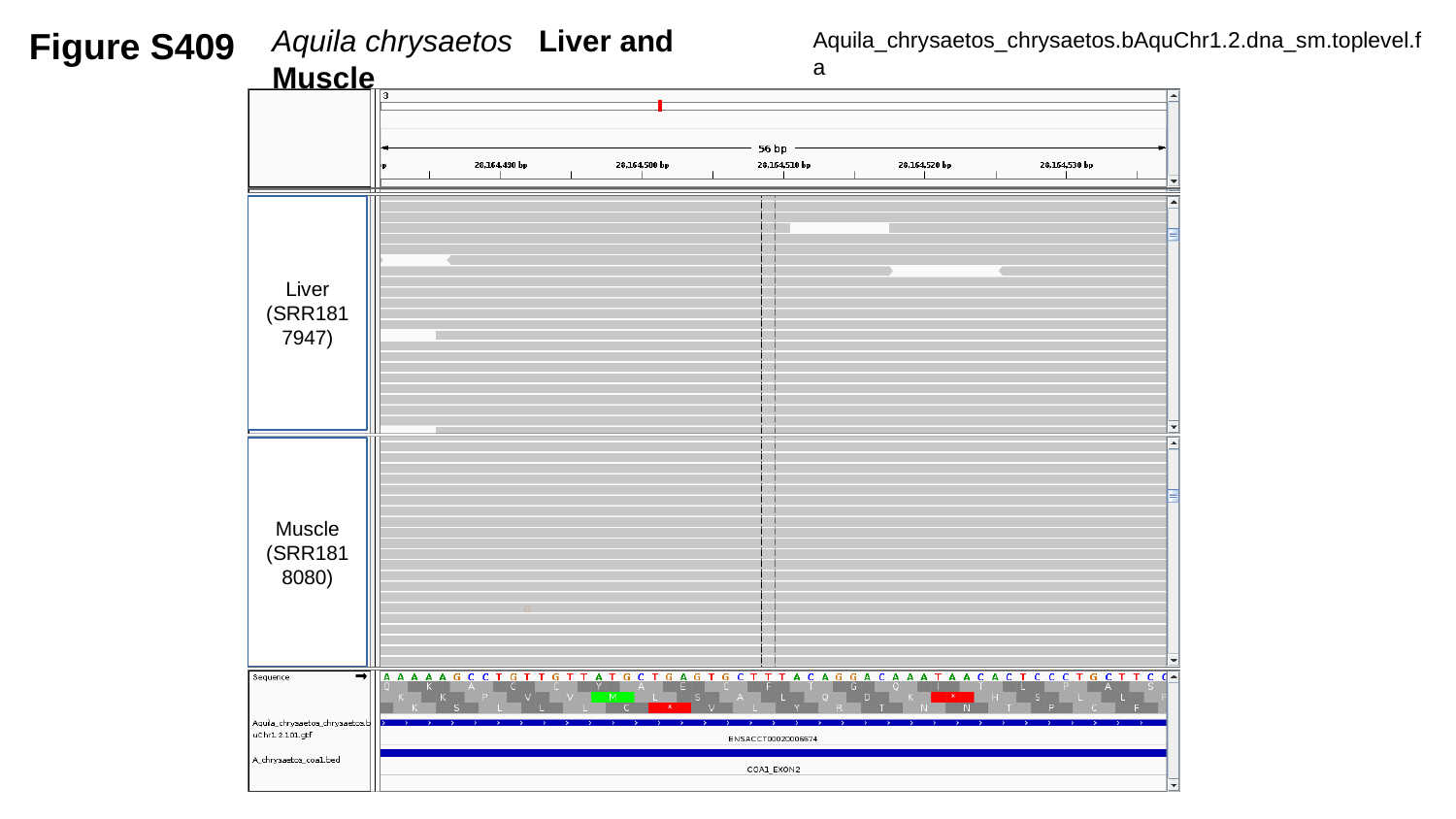

Figure S409
Aquila chrysaetos Liver and Muscle
Aquila_chrysaetos_chrysaetos.bAquChr1.2.dna_sm.toplevel.fa
Liver
(SRR1817947)
Muscle
(SRR1818080)

### Slide 410
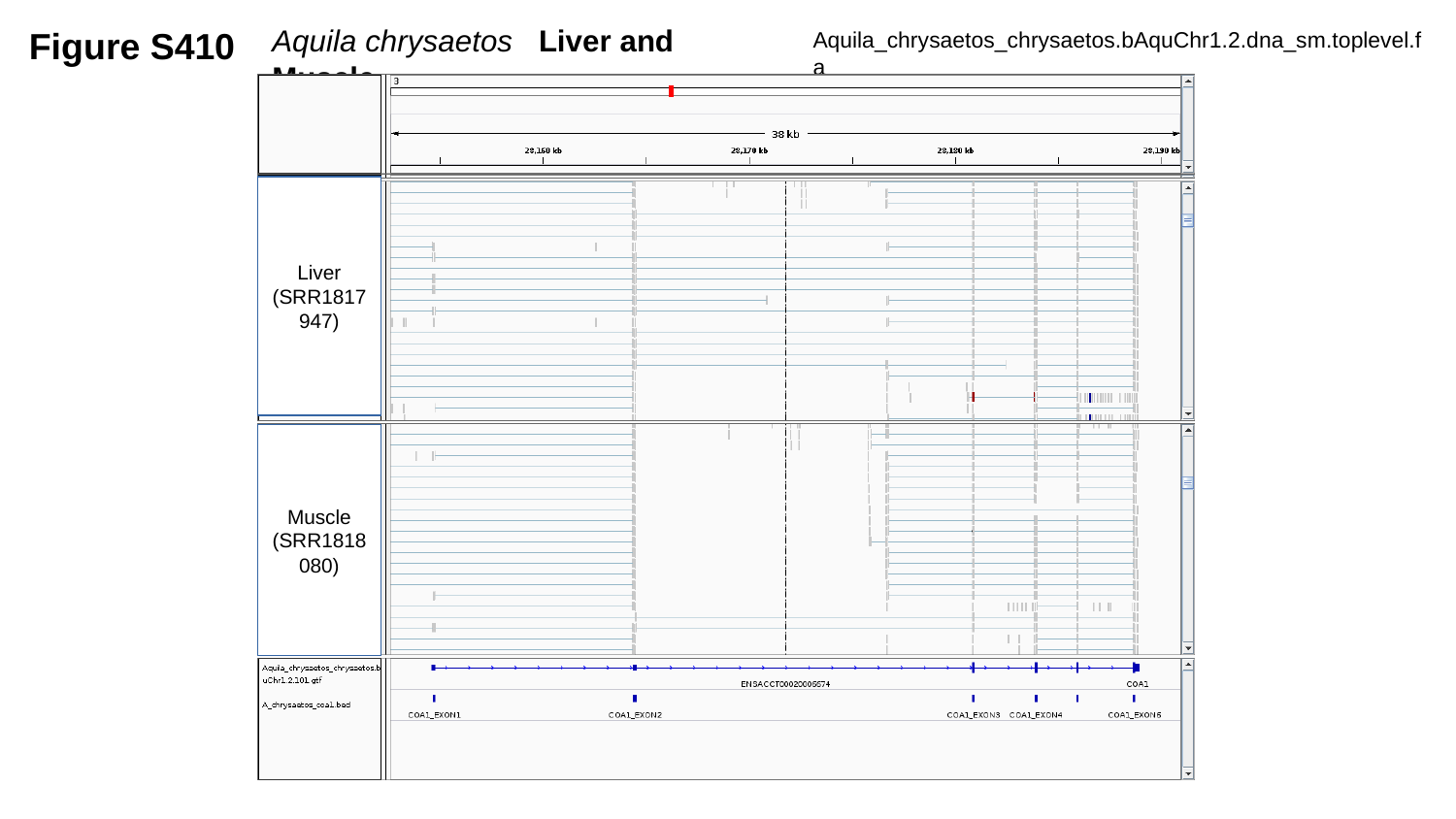

Figure S410
Aquila chrysaetos Liver and Muscle
Aquila_chrysaetos_chrysaetos.bAquChr1.2.dna_sm.toplevel.fa
Liver
(SRR1817947)
Muscle
(SRR1818080)

### Slide 411
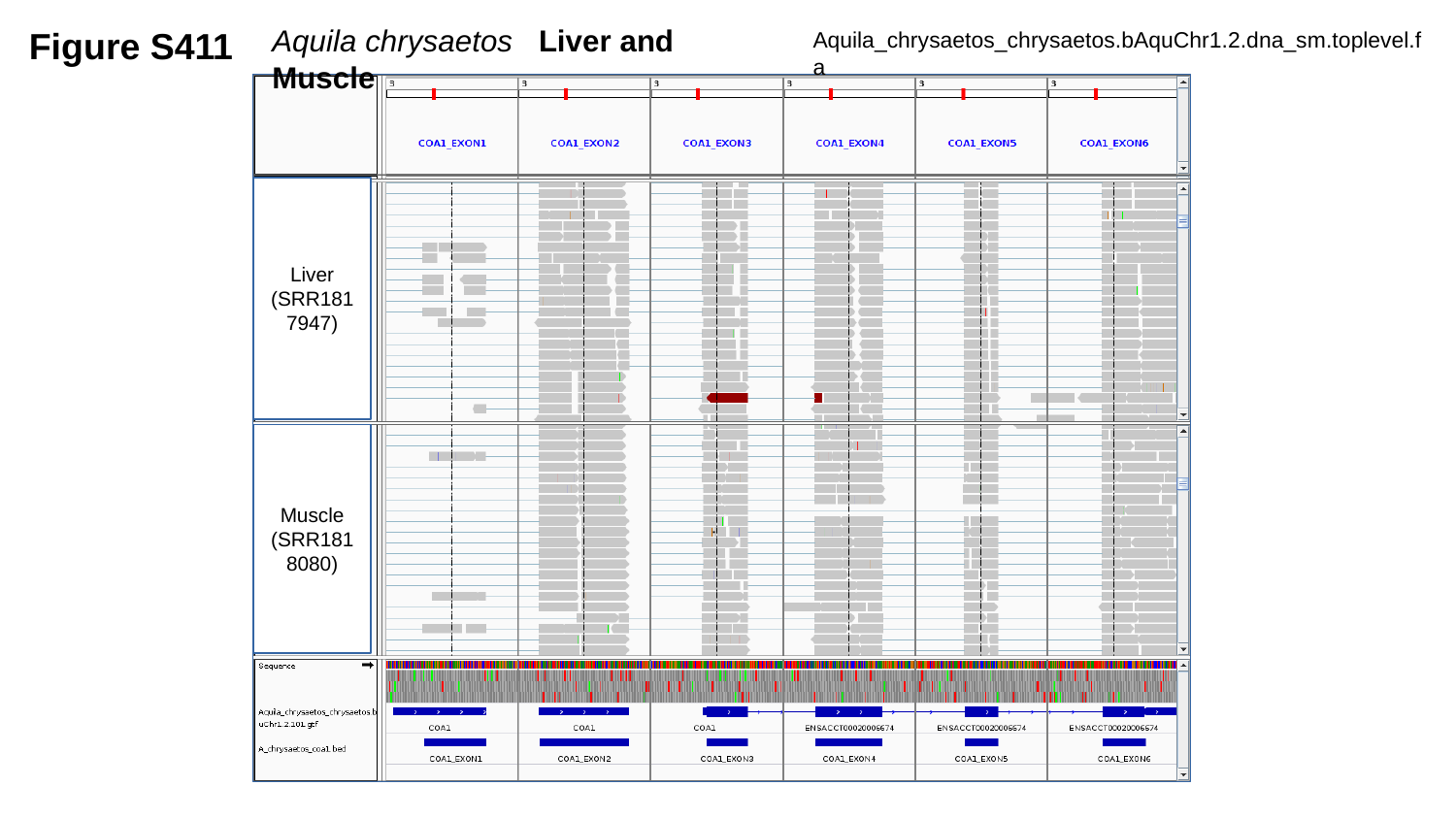

Figure S411
Aquila chrysaetos Liver and Muscle
Aquila_chrysaetos_chrysaetos.bAquChr1.2.dna_sm.toplevel.fa
Liver
(SRR1817947)
Muscle
(SRR1818080)

### Slide 412
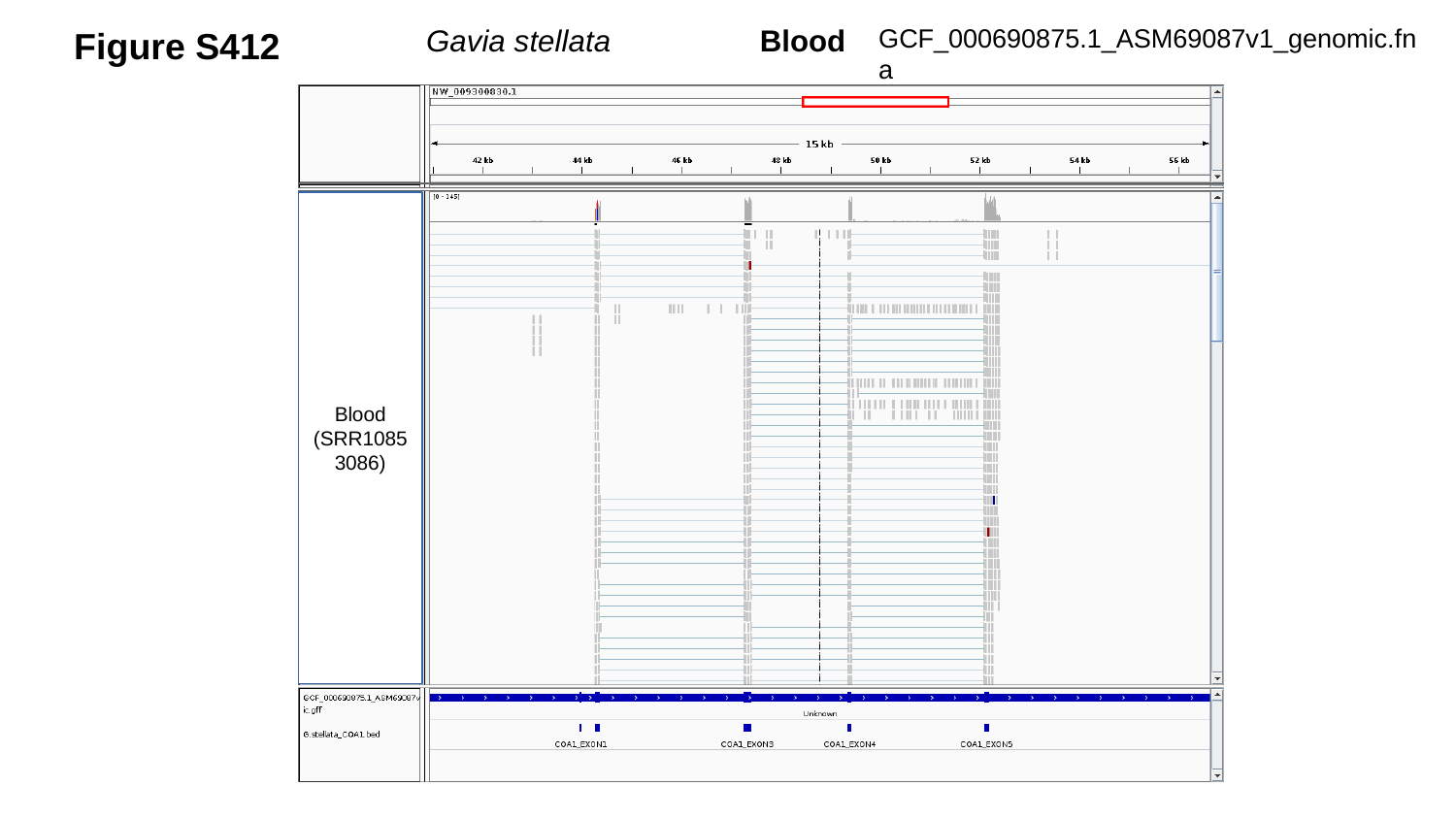

Figure S412
GCF_000690875.1_ASM69087v1_genomic.fna
Gavia stellata	 Blood
Blood
(SRR10853086)

### Slide 413
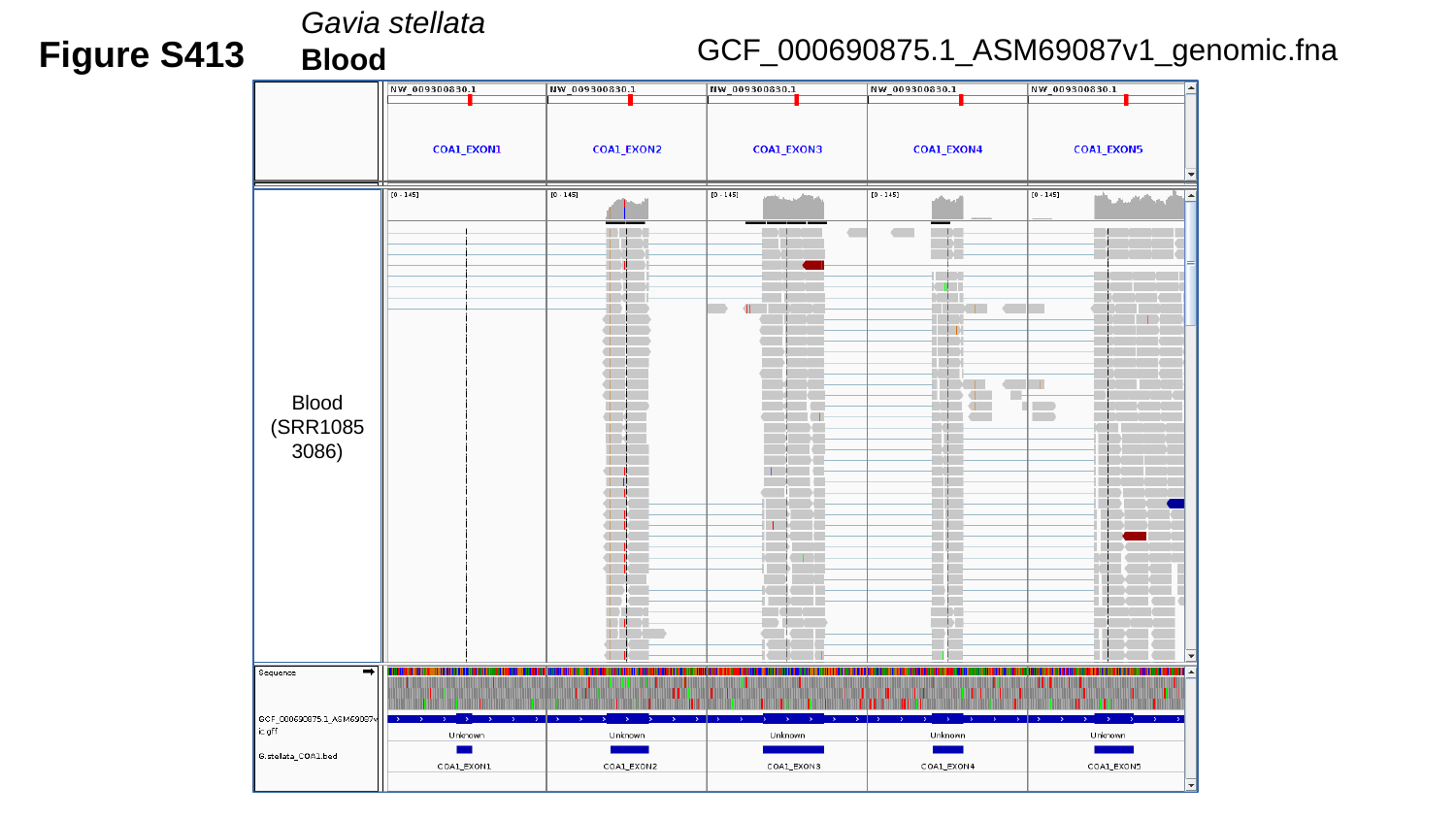

Gavia stellata	Blood
Figure S413
GCF_000690875.1_ASM69087v1_genomic.fna
Blood
(SRR10853086)

### Slide 414
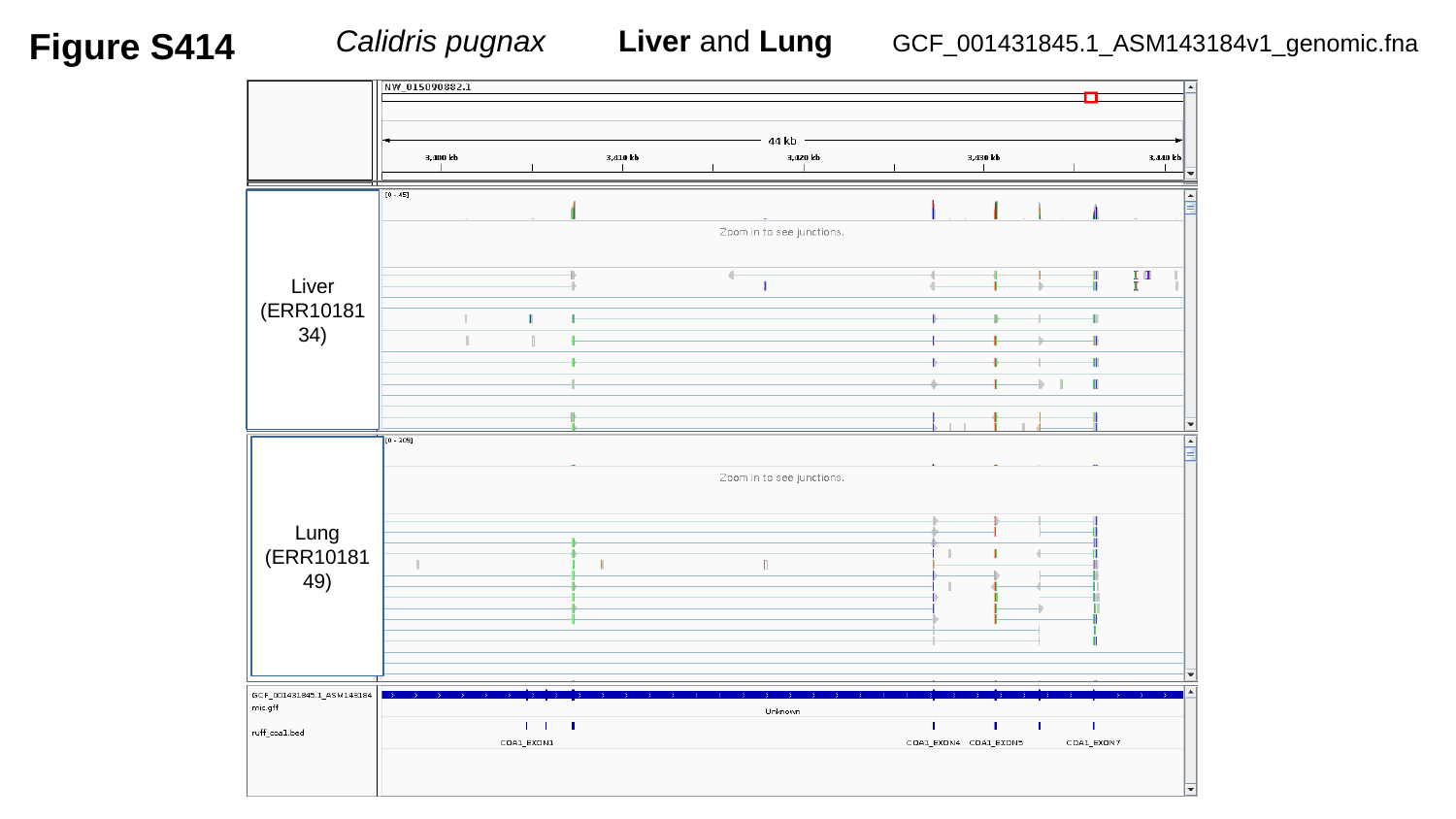

Figure S414
 Calidris pugnax	Liver and Lung
GCF_001431845.1_ASM143184v1_genomic.fna
Liver
(ERR1018134)
Lung
(ERR1018149)

### Slide 415
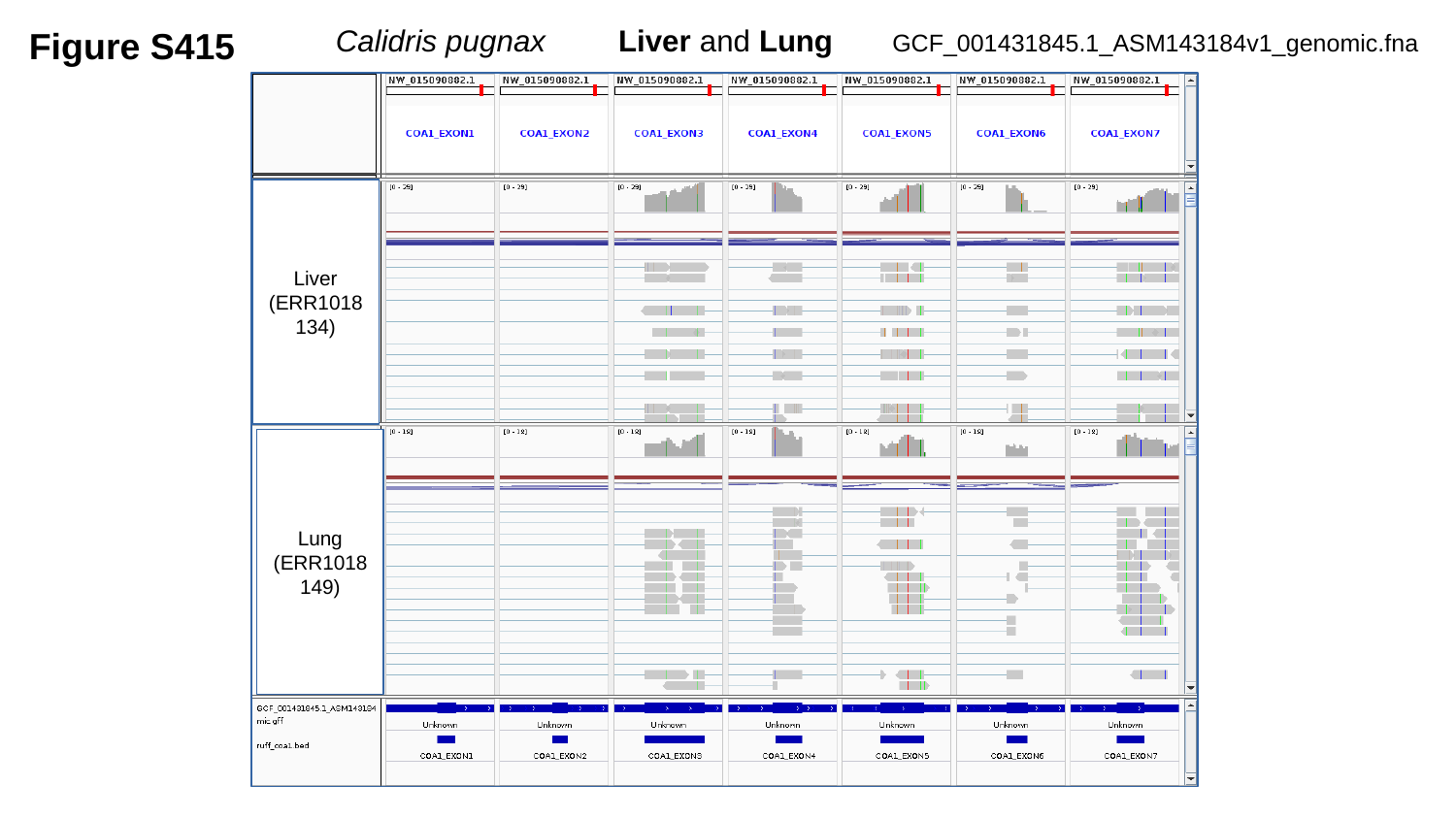

Figure S415
 Calidris pugnax	Liver and Lung
GCF_001431845.1_ASM143184v1_genomic.fna
Liver
(ERR1018134)
Lung
(ERR1018149)

### Slide 416
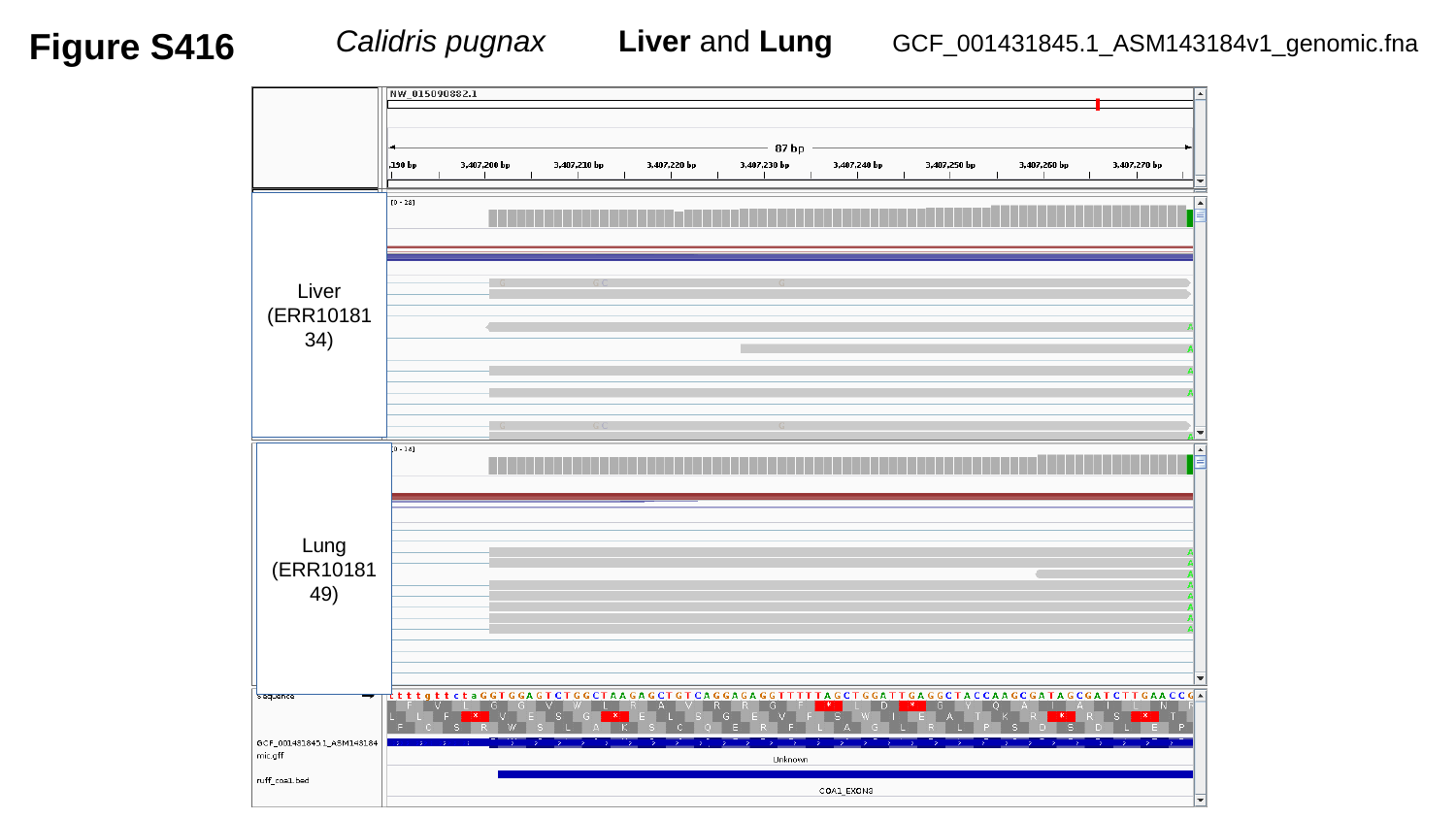

Figure S416
 Calidris pugnax	Liver and Lung
GCF_001431845.1_ASM143184v1_genomic.fna
Liver
(ERR1018134)
Lung
(ERR1018149)

### Slide 417
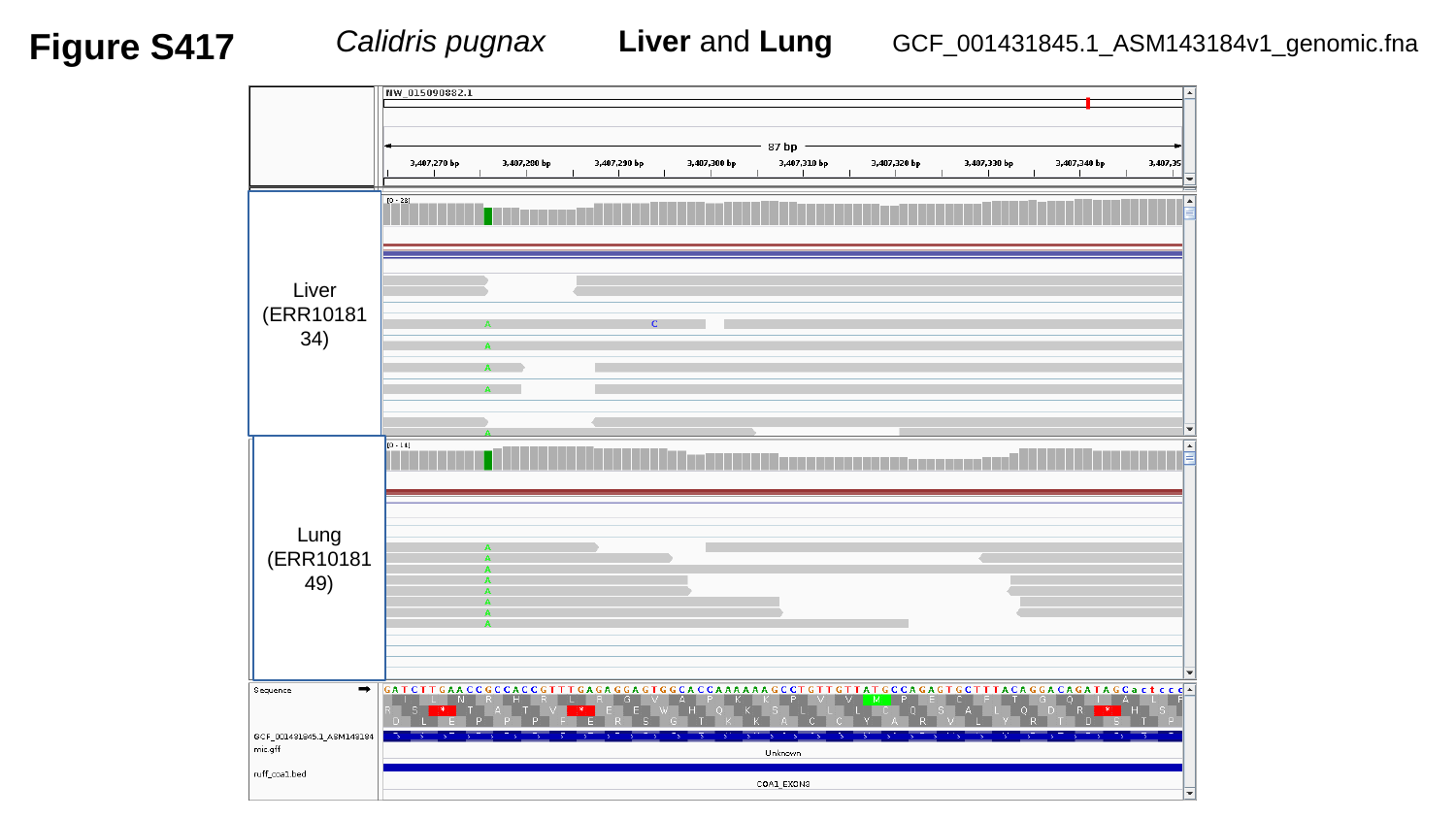

Figure S417
 Calidris pugnax	Liver and Lung
GCF_001431845.1_ASM143184v1_genomic.fna
Liver
(ERR1018134)
Lung
(ERR1018149)

### Slide 418
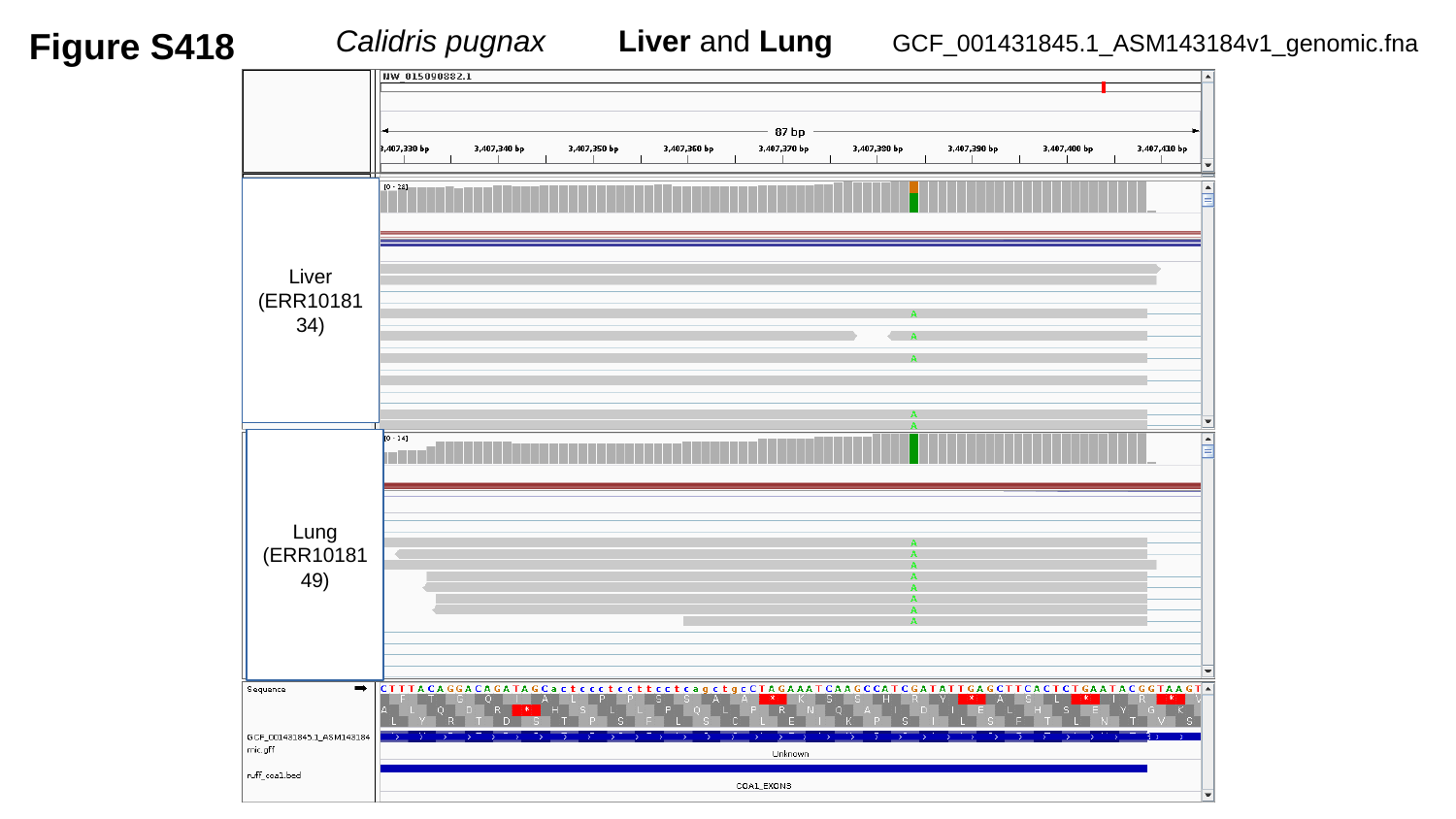

Figure S418
 Calidris pugnax	Liver and Lung
GCF_001431845.1_ASM143184v1_genomic.fna
Liver
(ERR1018134)
Lung
(ERR1018149)

### Slide 419
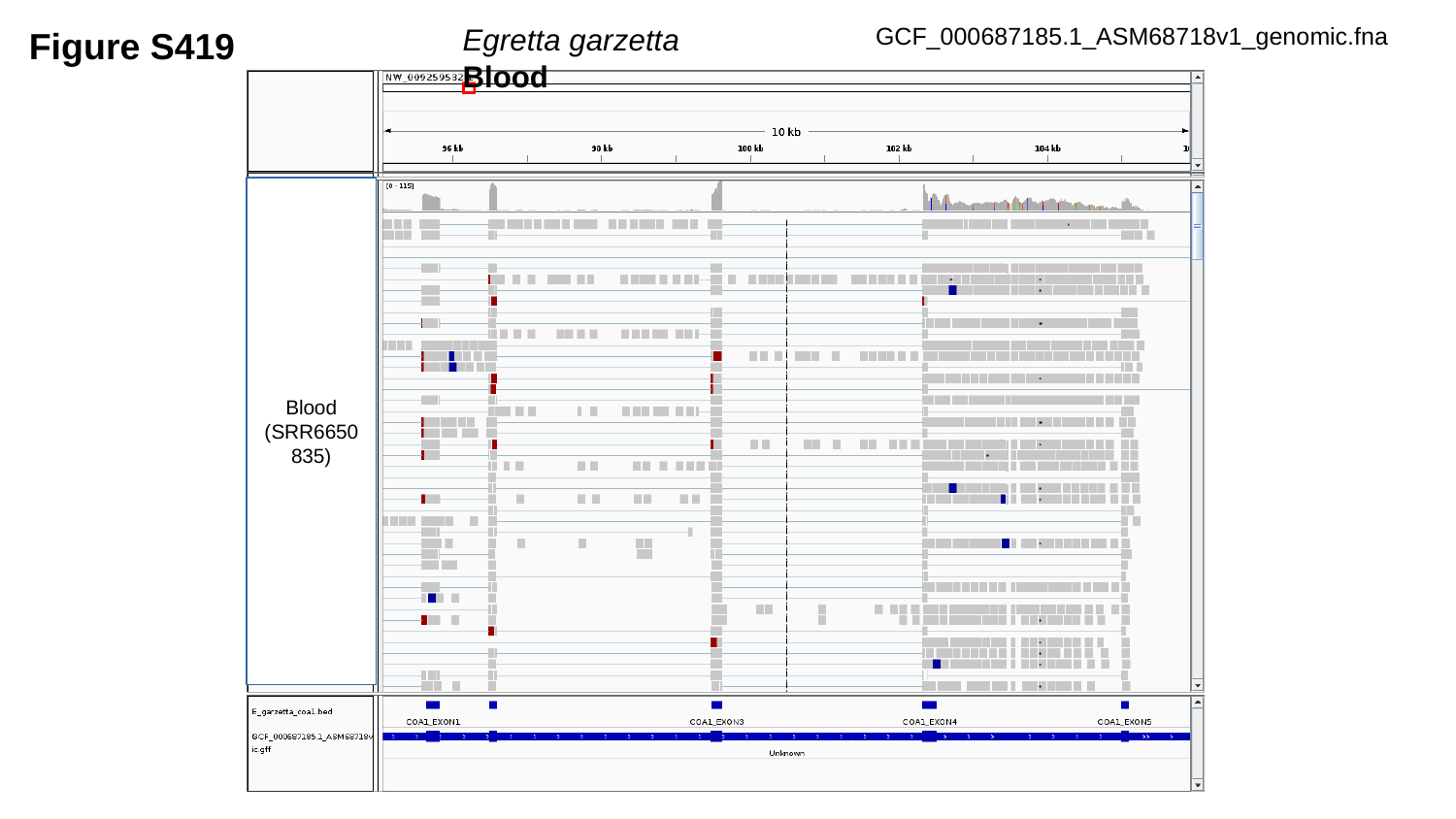

Egretta garzetta	Blood
GCF_000687185.1_ASM68718v1_genomic.fna
Figure S419
Blood
(SRR6650835)

### Slide 420
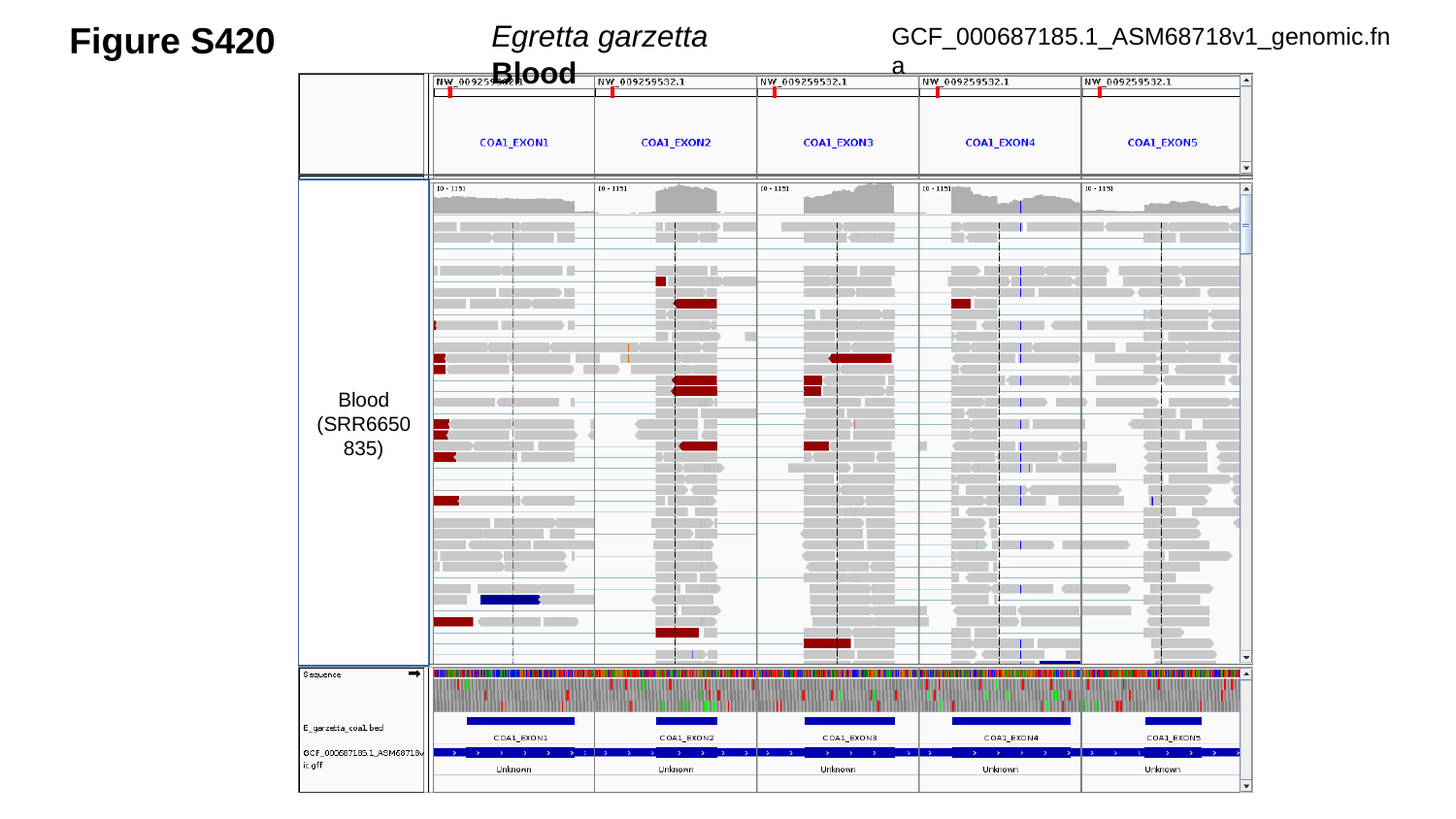

Egretta garzetta	 Blood
Figure S420
GCF_000687185.1_ASM68718v1_genomic.fna
Blood
(SRR6650835)

### Slide 421
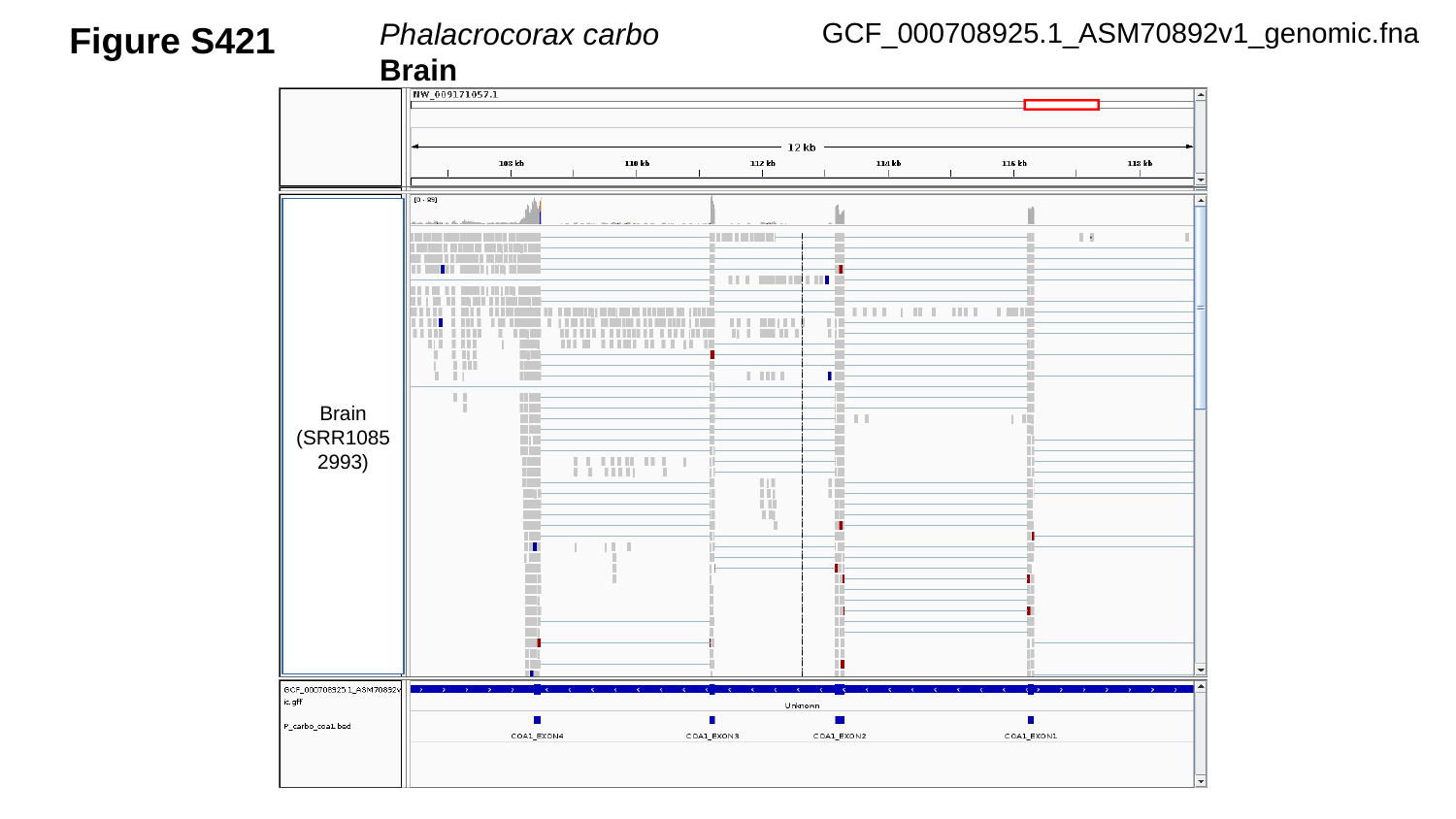

Figure S421
GCF_000708925.1_ASM70892v1_genomic.fna
Phalacrocorax carbo Brain
Brain
(SRR10852993)

### Slide 422
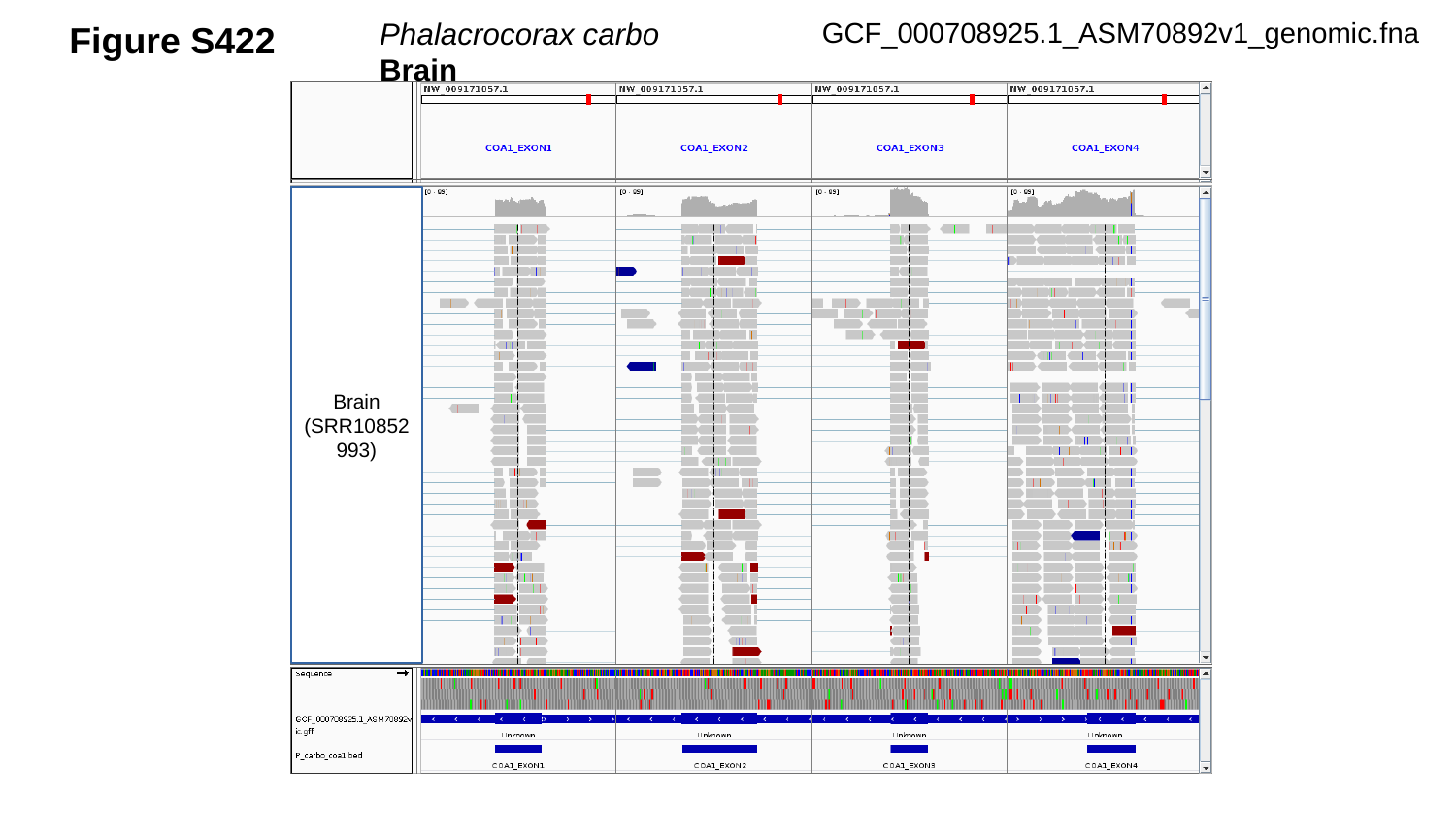

Figure S422
GCF_000708925.1_ASM70892v1_genomic.fna
Phalacrocorax carbo Brain
Brain
(SRR10852993)

### Slide 423
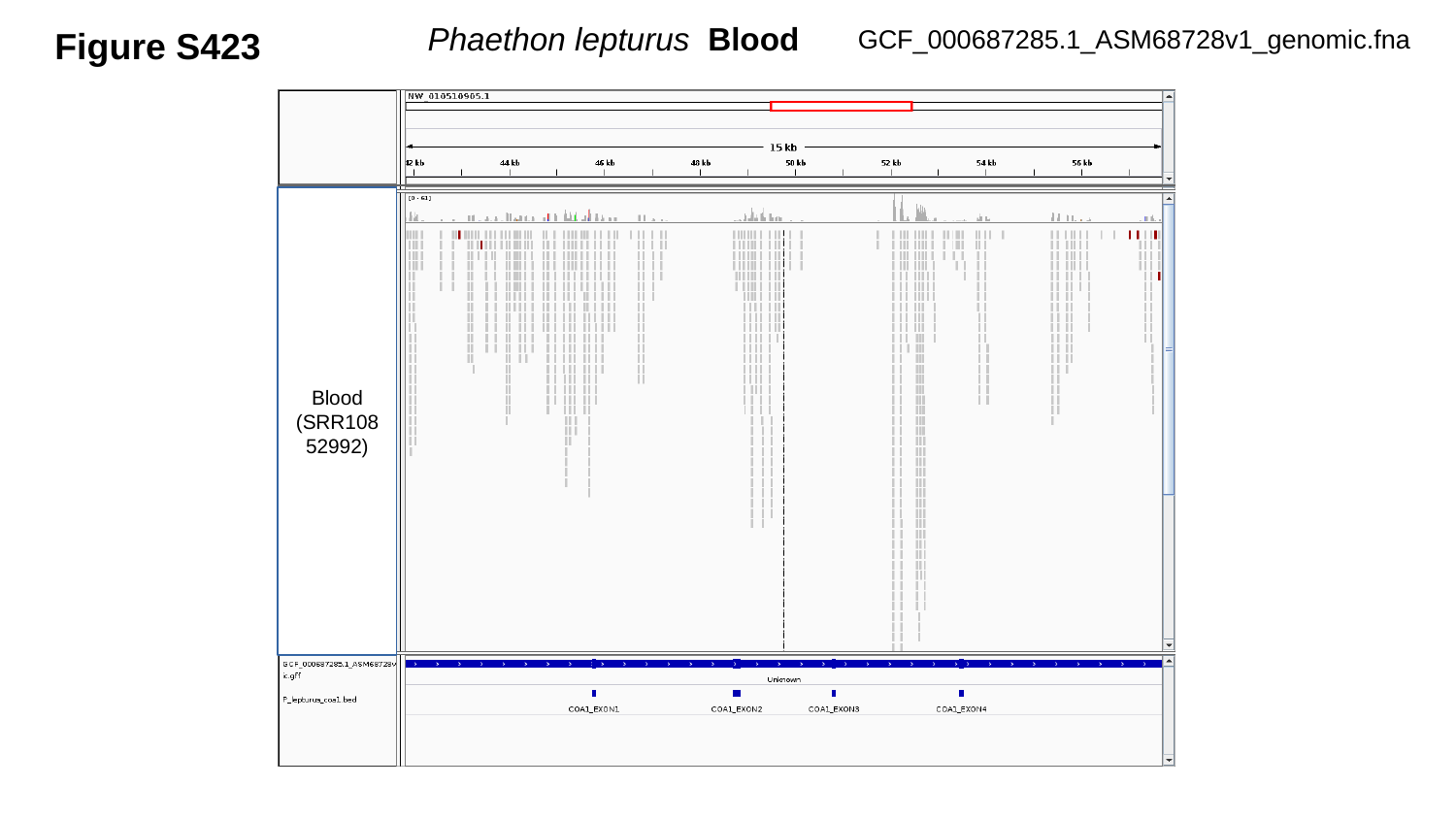

GCF_000687285.1_ASM68728v1_genomic.fna
Figure S423
 Phaethon lepturus Blood
Blood
(SRR10852992)

### Slide 424
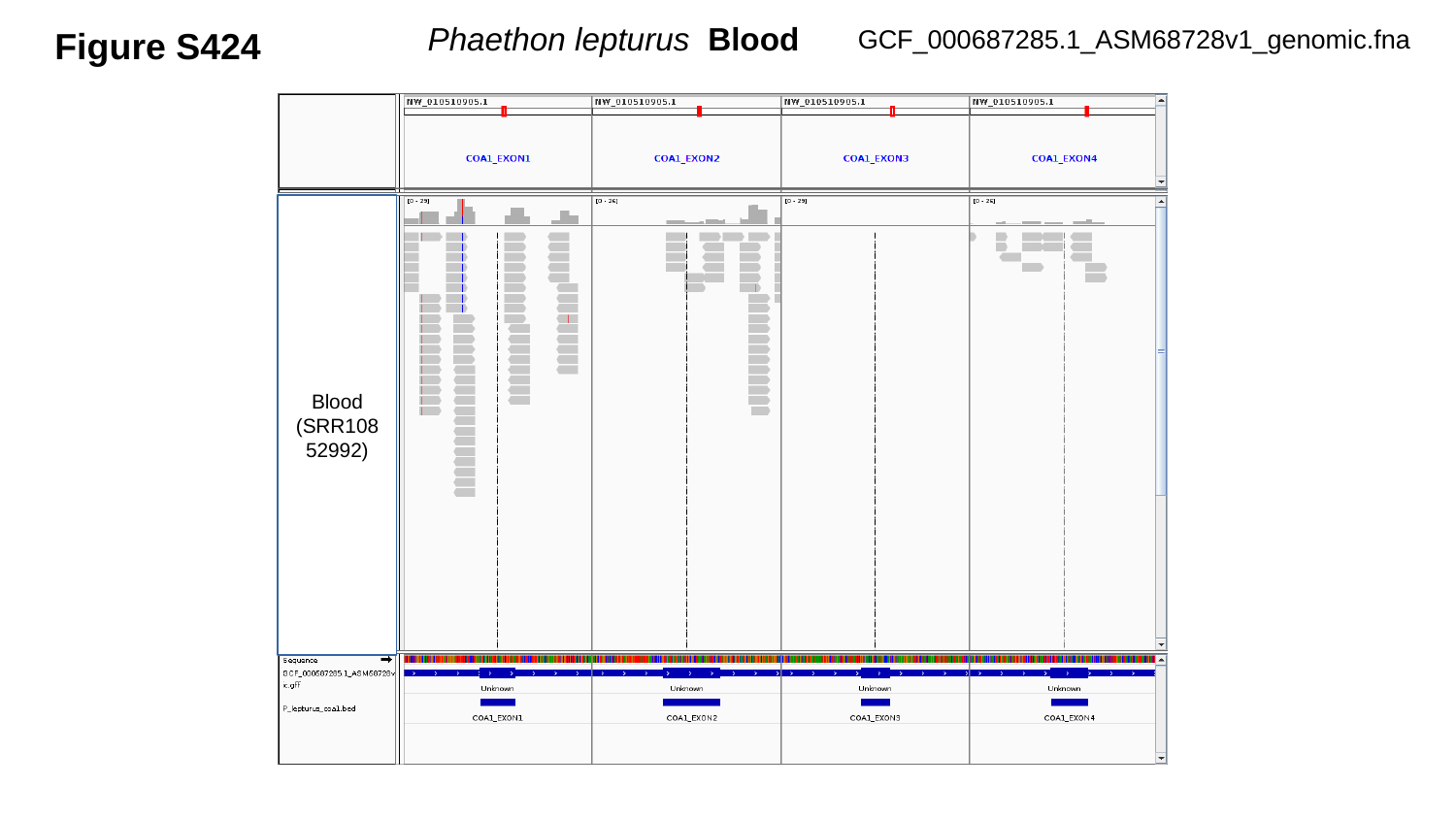

GCF_000687285.1_ASM68728v1_genomic.fna
Figure S424
 Phaethon lepturus Blood
Blood
(SRR10852992)

### Slide 425
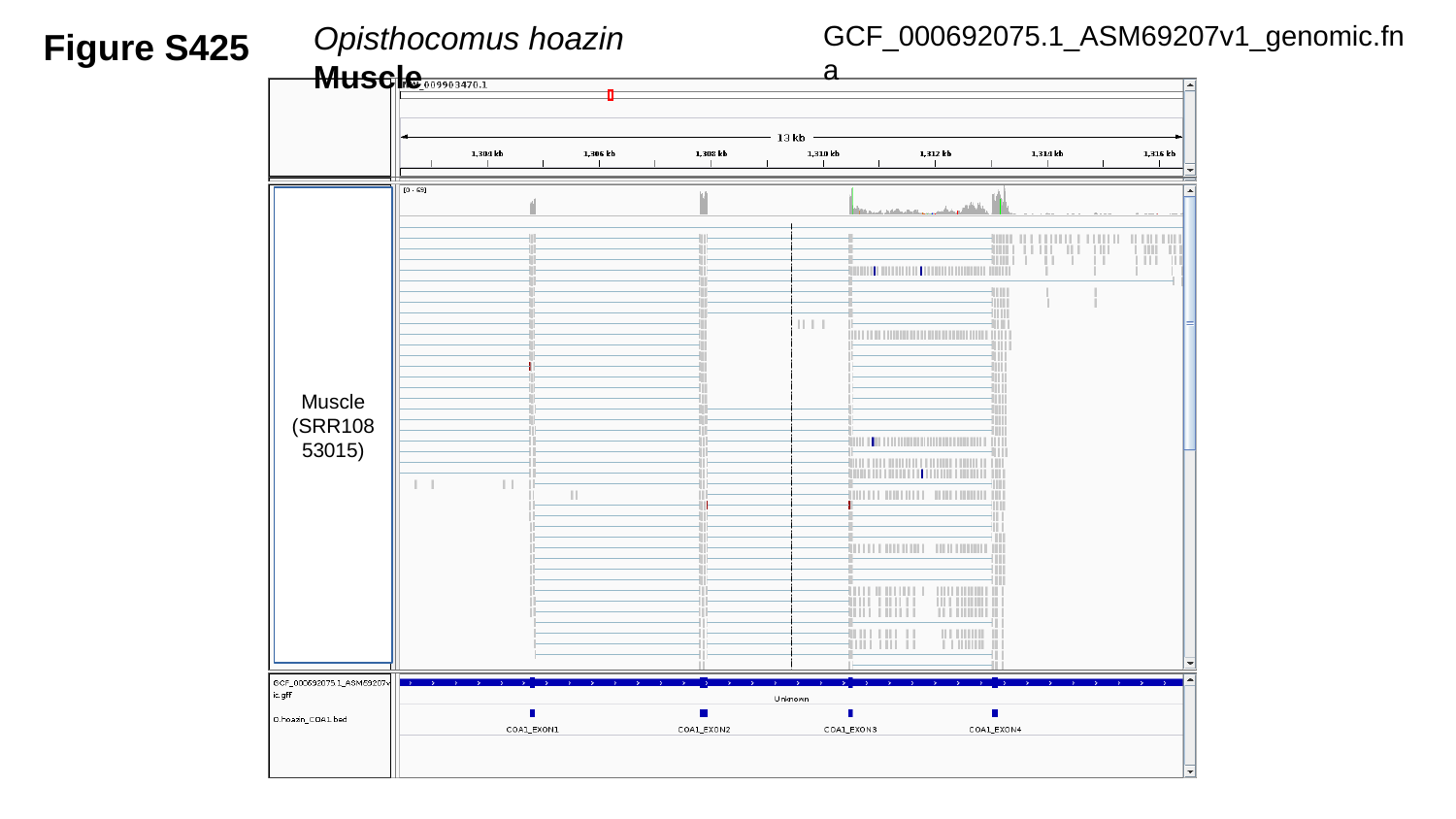

GCF_000692075.1_ASM69207v1_genomic.fna
Figure S425
Opisthocomus hoazin	 Muscle
Muscle
(SRR10853015)

### Slide 426
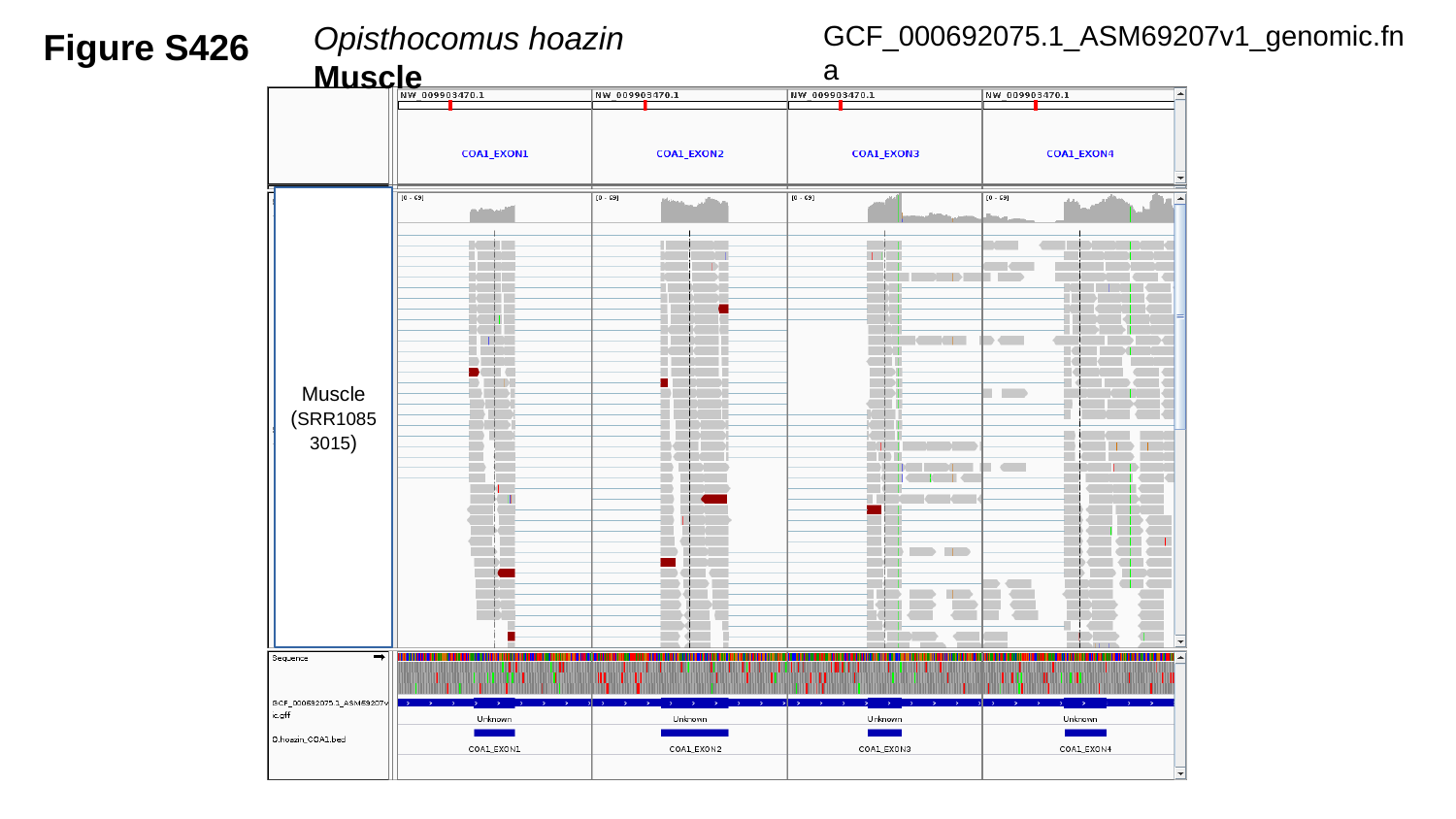

GCF_000692075.1_ASM69207v1_genomic.fna
Figure S426
Opisthocomus hoazin	 Muscle
Muscle
(SRR10853015)

### Slide 427
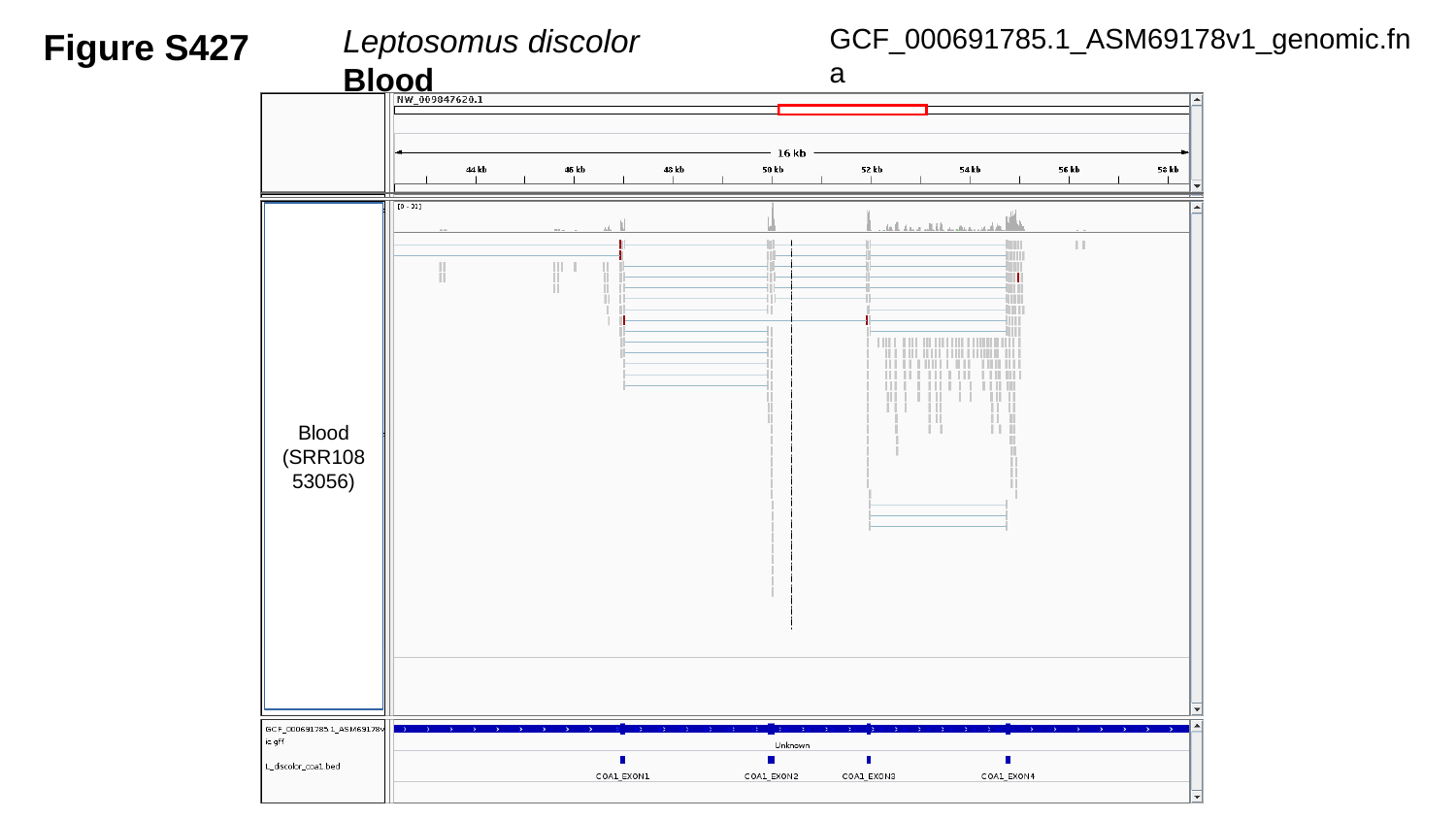

Figure S427
GCF_000691785.1_ASM69178v1_genomic.fna
Leptosomus discolor	 Blood
Blood
(SRR10853056)

### Slide 428
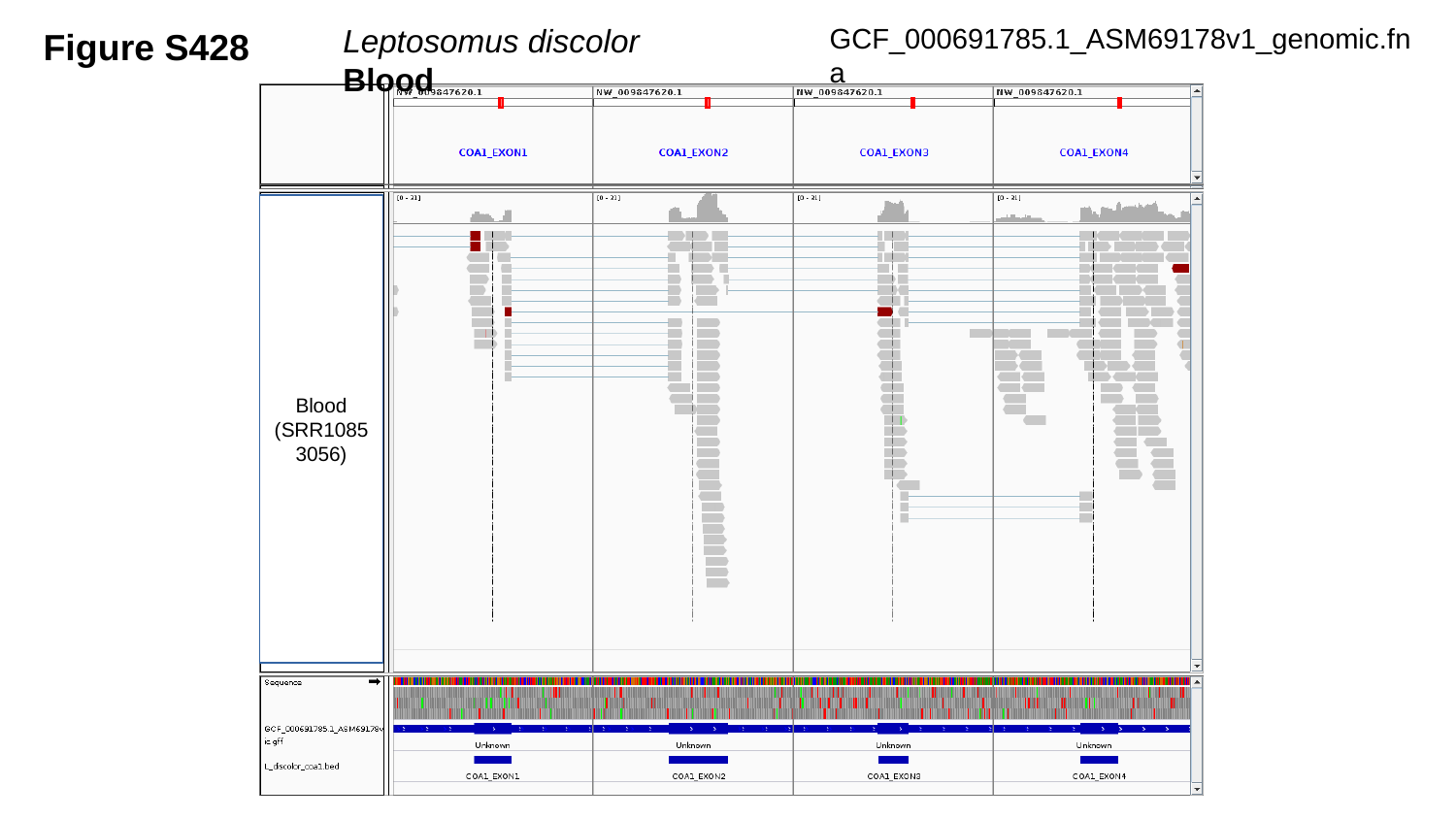

Figure S428
GCF_000691785.1_ASM69178v1_genomic.fna
Leptosomus discolor	 Blood
Blood
(SRR10853056)

### Slide 429
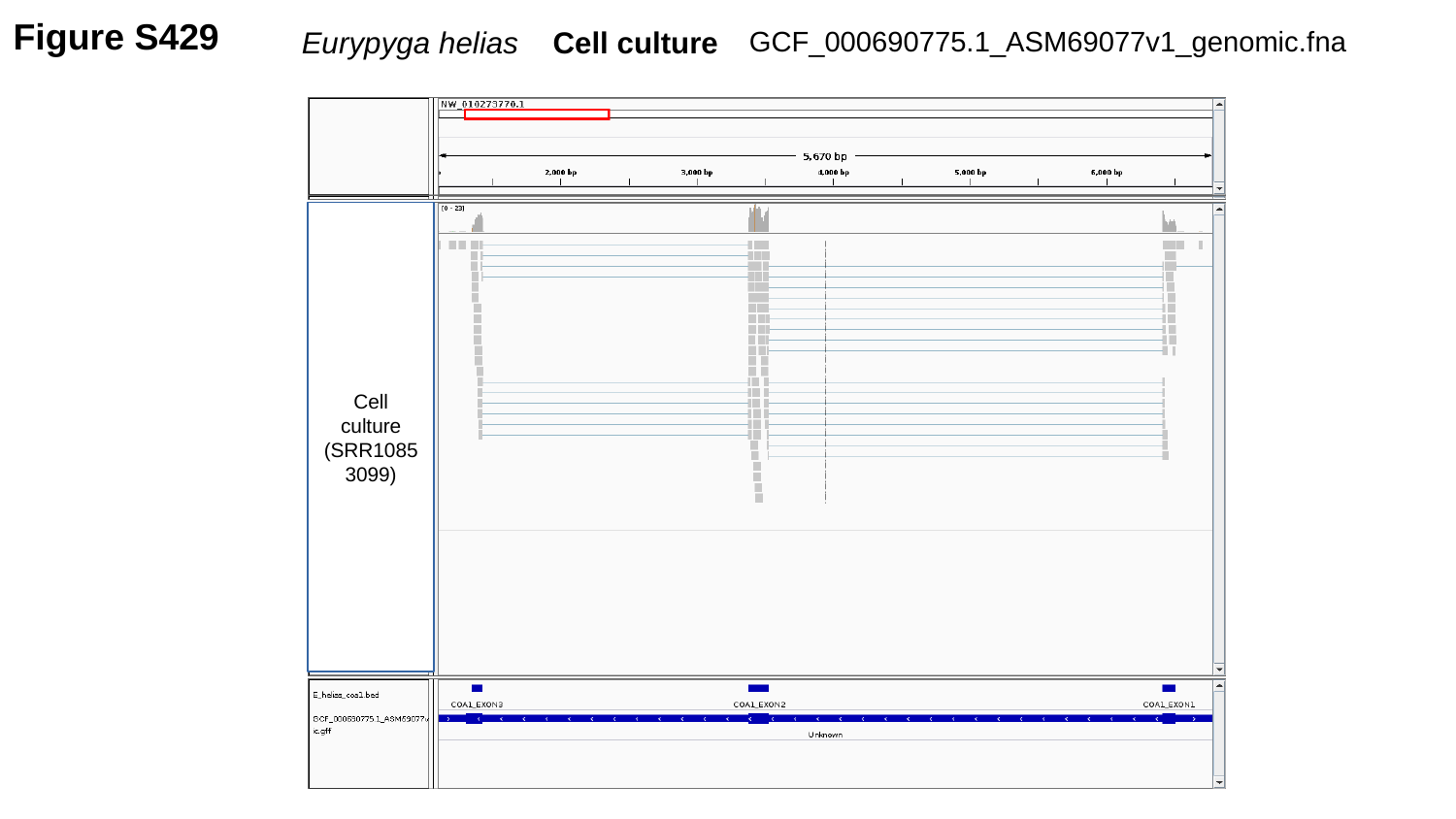

Figure S429
GCF_000690775.1_ASM69077v1_genomic.fna
Eurypyga helias Cell culture
Cell culture
(SRR10853099)

### Slide 430
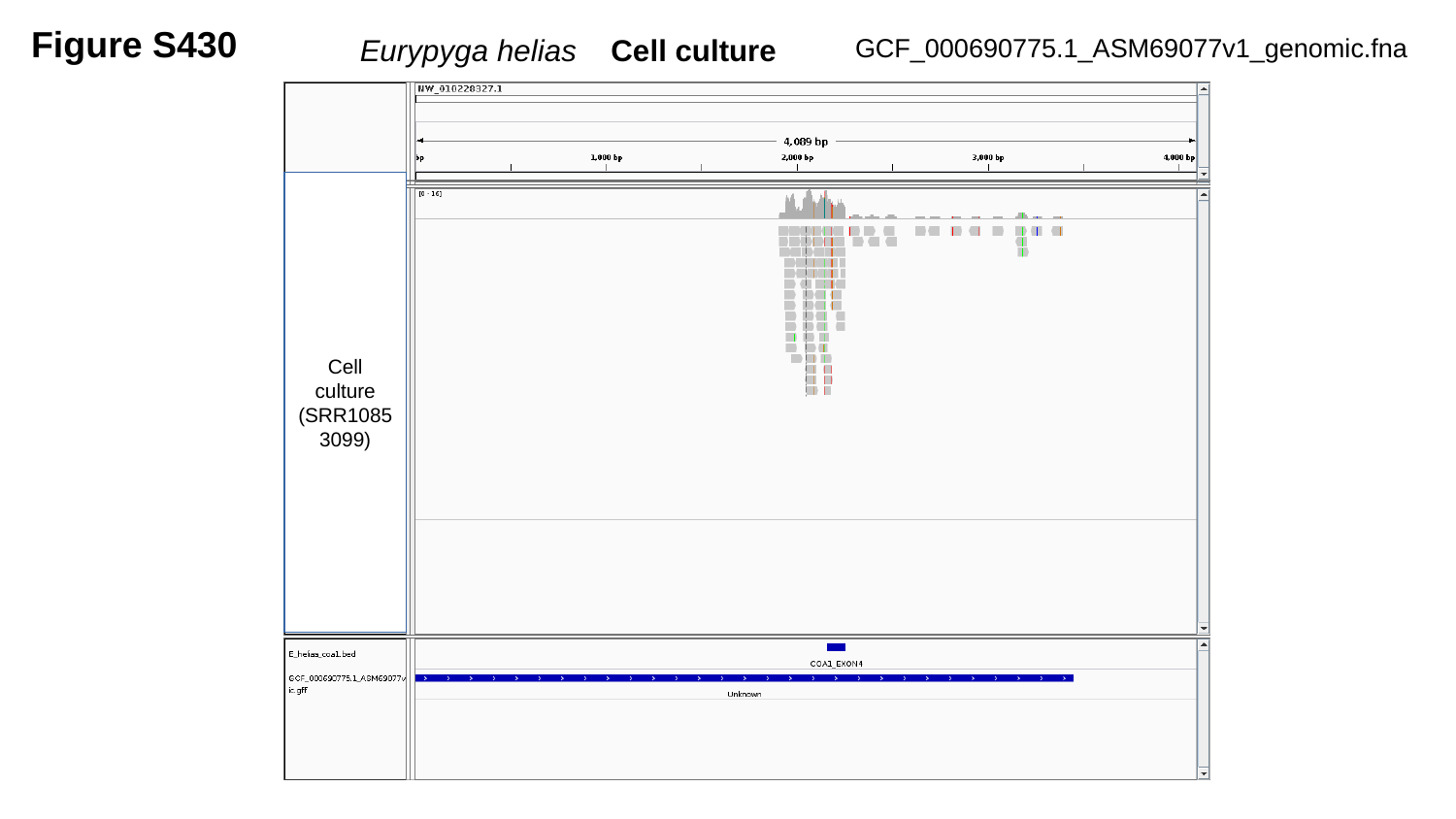

Figure S430
Eurypyga helias Cell culture
GCF_000690775.1_ASM69077v1_genomic.fna
Cell culture
(SRR10853099)

### Slide 431
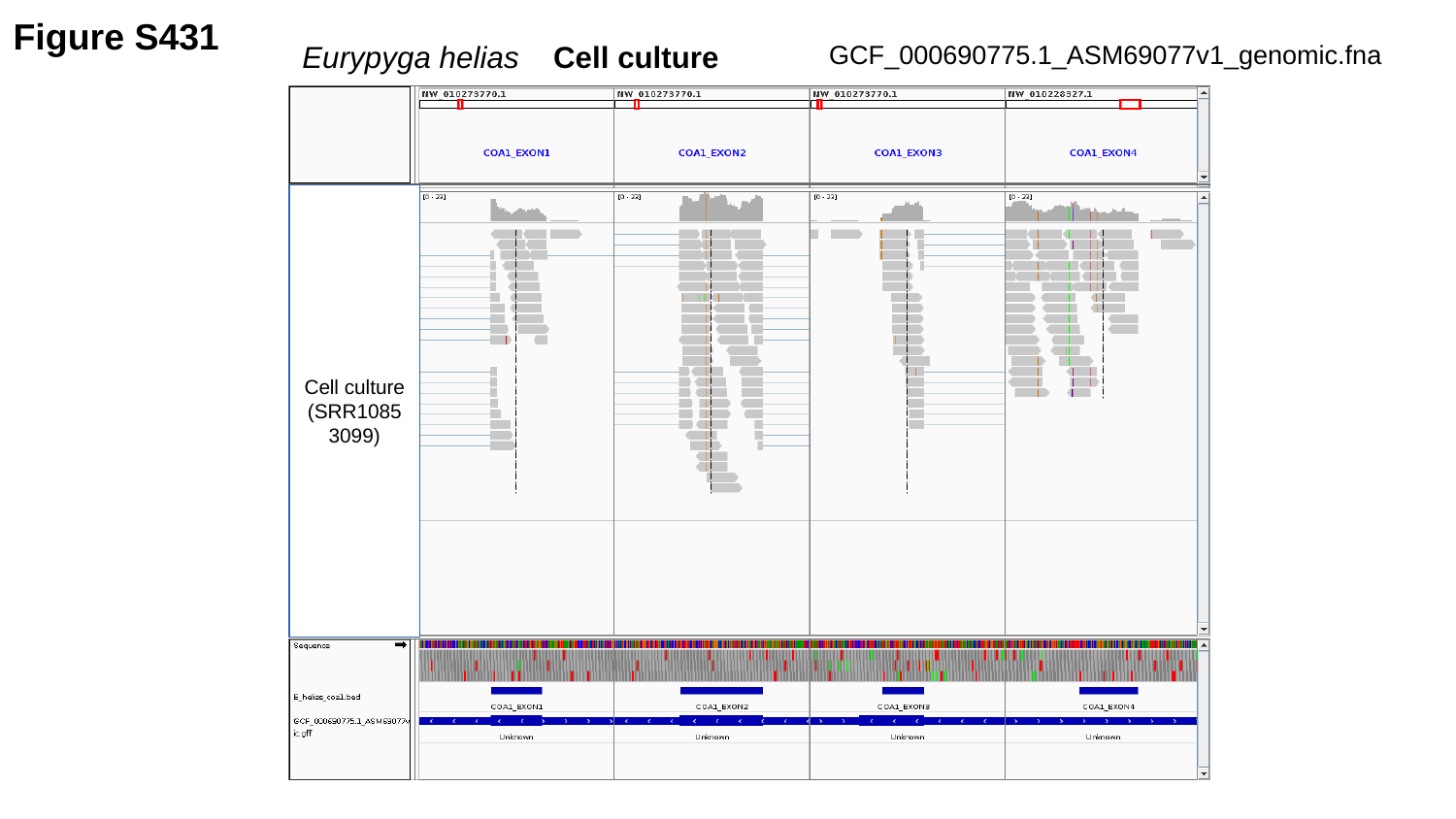

Figure S431
Eurypyga helias Cell culture
GCF_000690775.1_ASM69077v1_genomic.fna
Cell culture
(SRR10853099)

### Slide 432
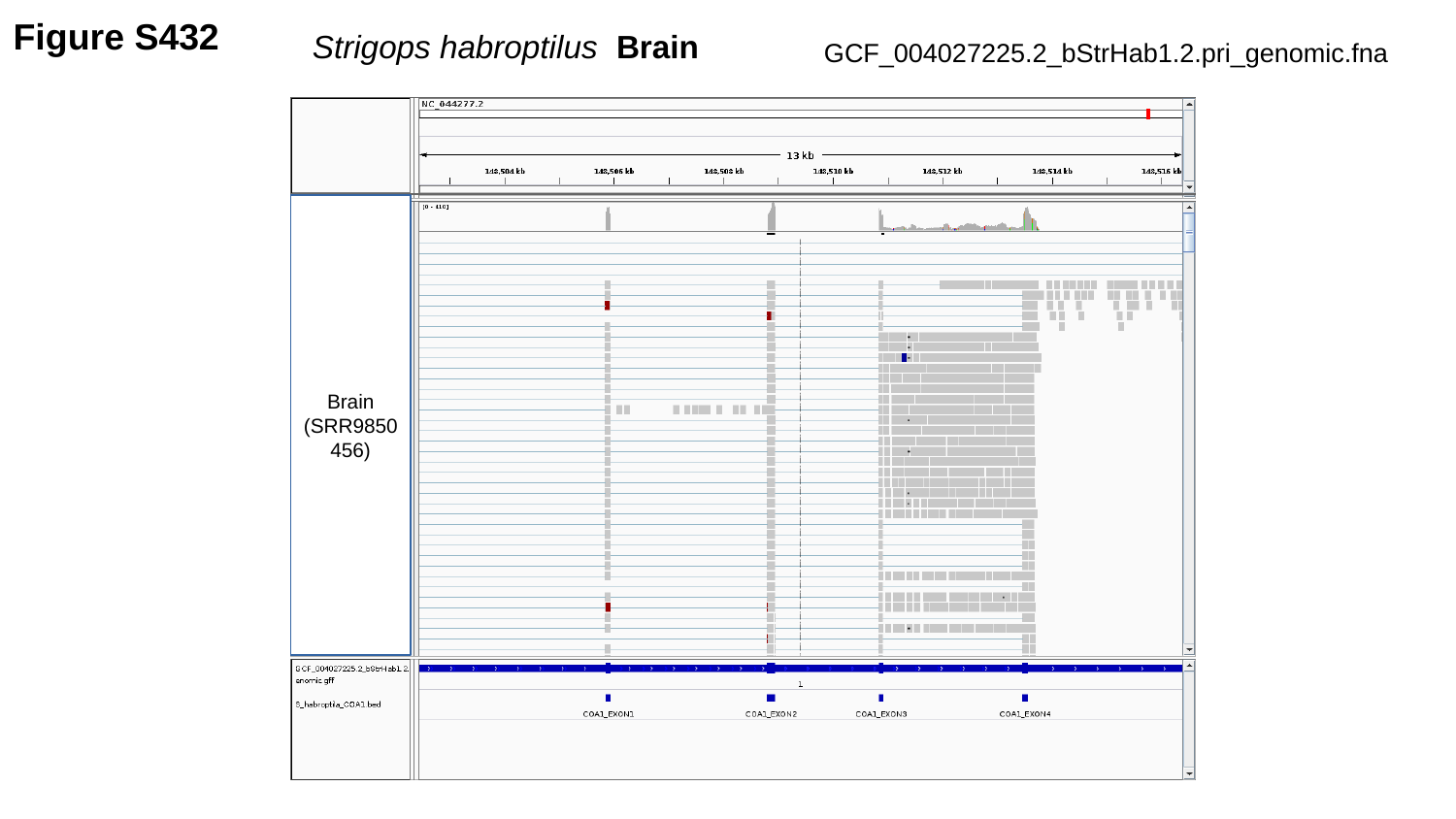

Figure S432
 Strigops habroptilus Brain
GCF_004027225.2_bStrHab1.2.pri_genomic.fna
Brain
(SRR9850456)

### Slide 433
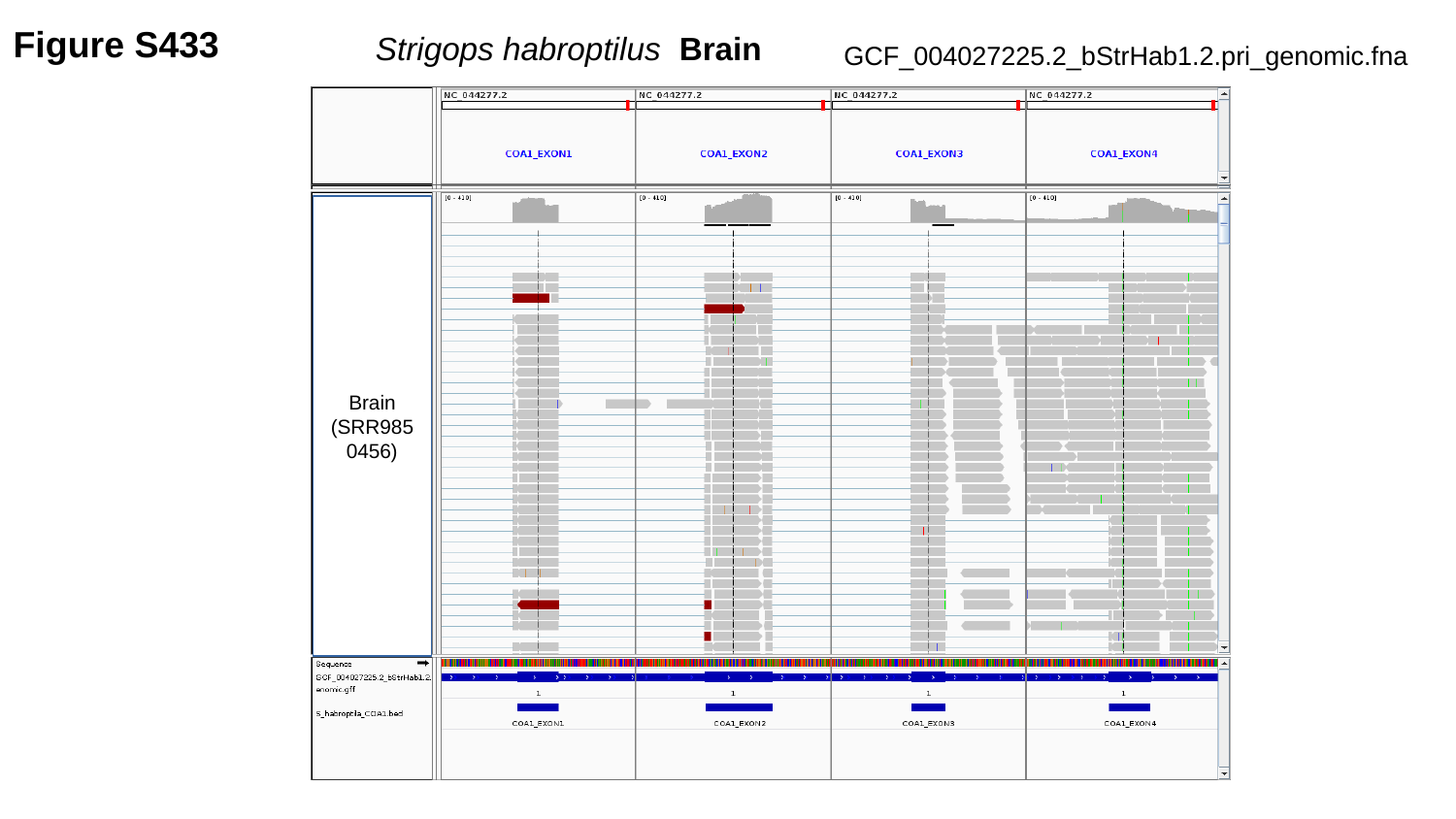

Figure S433
 Strigops habroptilus Brain
GCF_004027225.2_bStrHab1.2.pri_genomic.fna
Brain
(SRR9850456)

### Slide 434
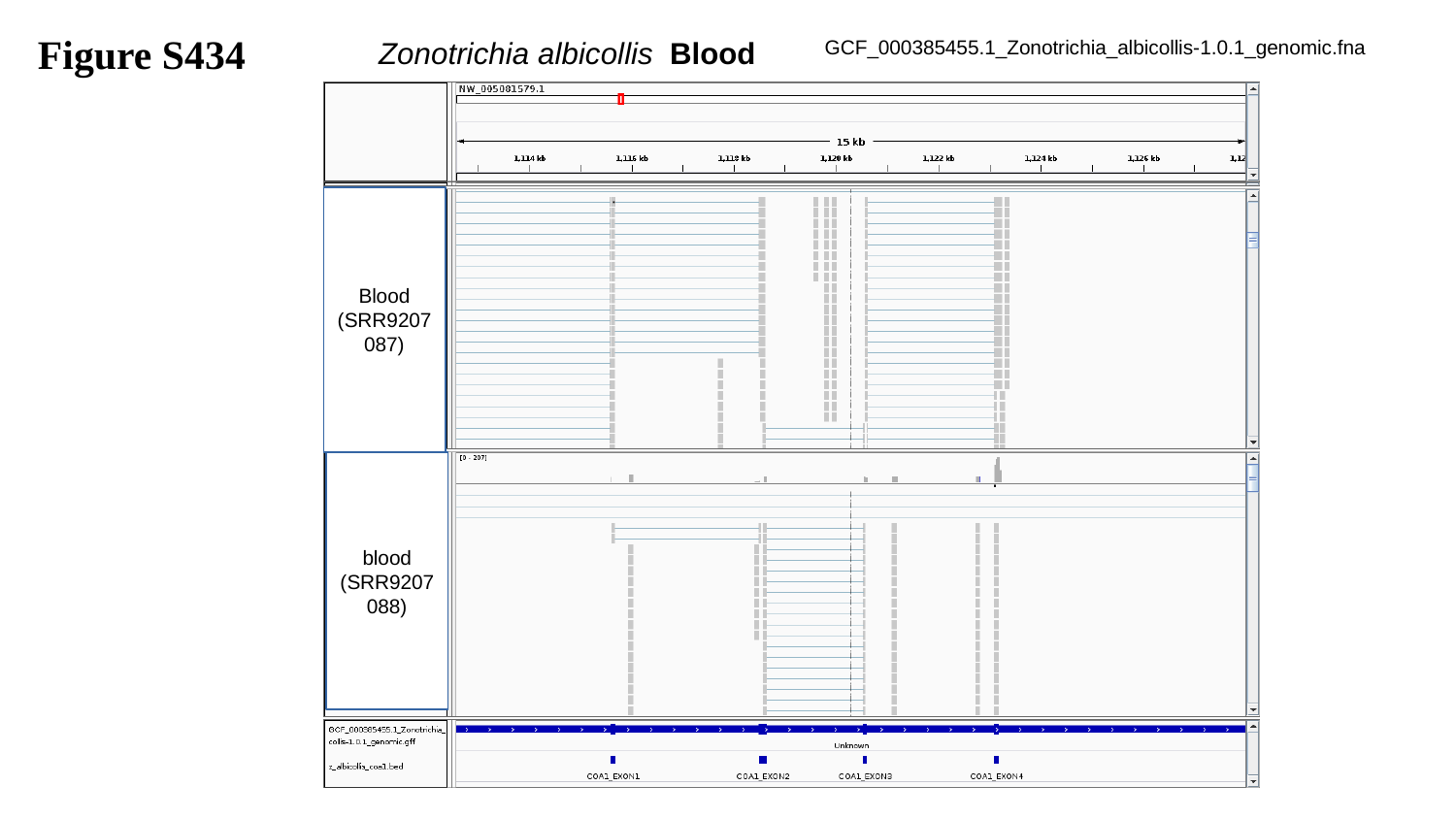

Figure S434
GCF_000385455.1_Zonotrichia_albicollis-1.0.1_genomic.fna
Zonotrichia albicollis	Blood
Blood
(SRR9207087)
blood
(SRR9207088)

### Slide 435
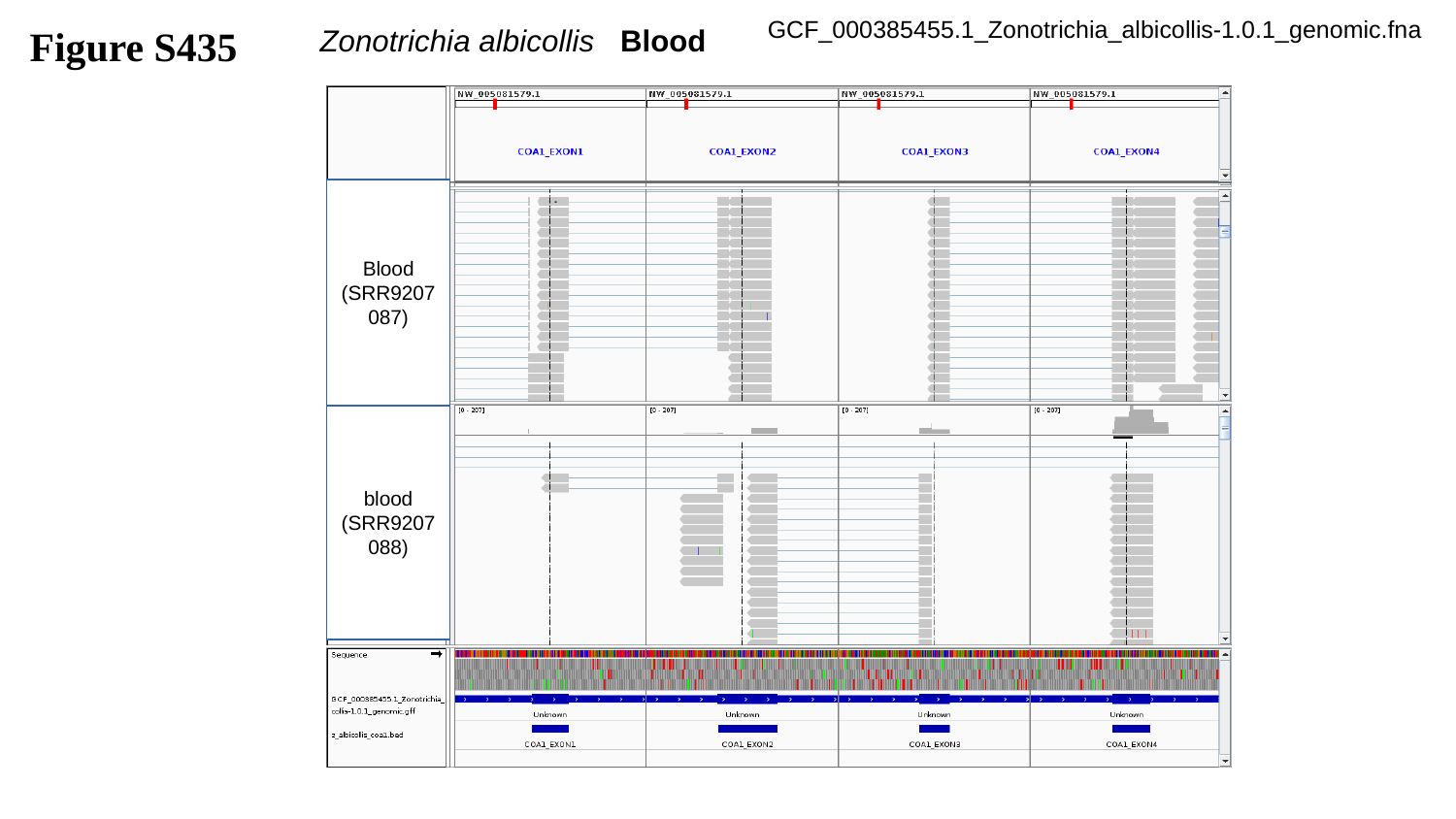

Figure S435
GCF_000385455.1_Zonotrichia_albicollis-1.0.1_genomic.fna
Zonotrichia albicollis Blood
Blood
(SRR9207087)
blood
(SRR9207088)

### Slide 436
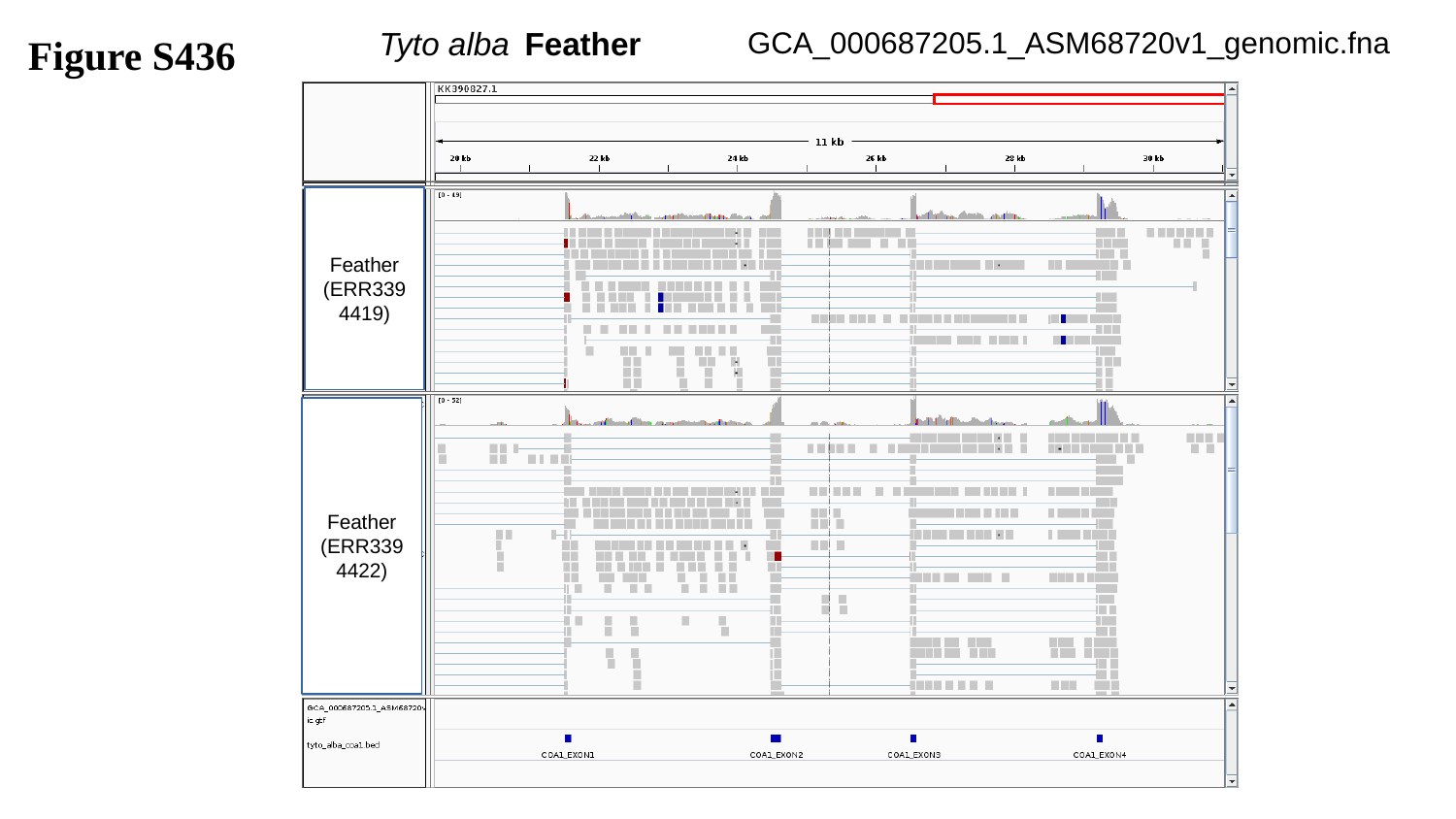

GCA_000687205.1_ASM68720v1_genomic.fna
Figure S436
Tyto alba	Feather
Feather
(ERR3394419)
Feather
(ERR3394422)

### Slide 437
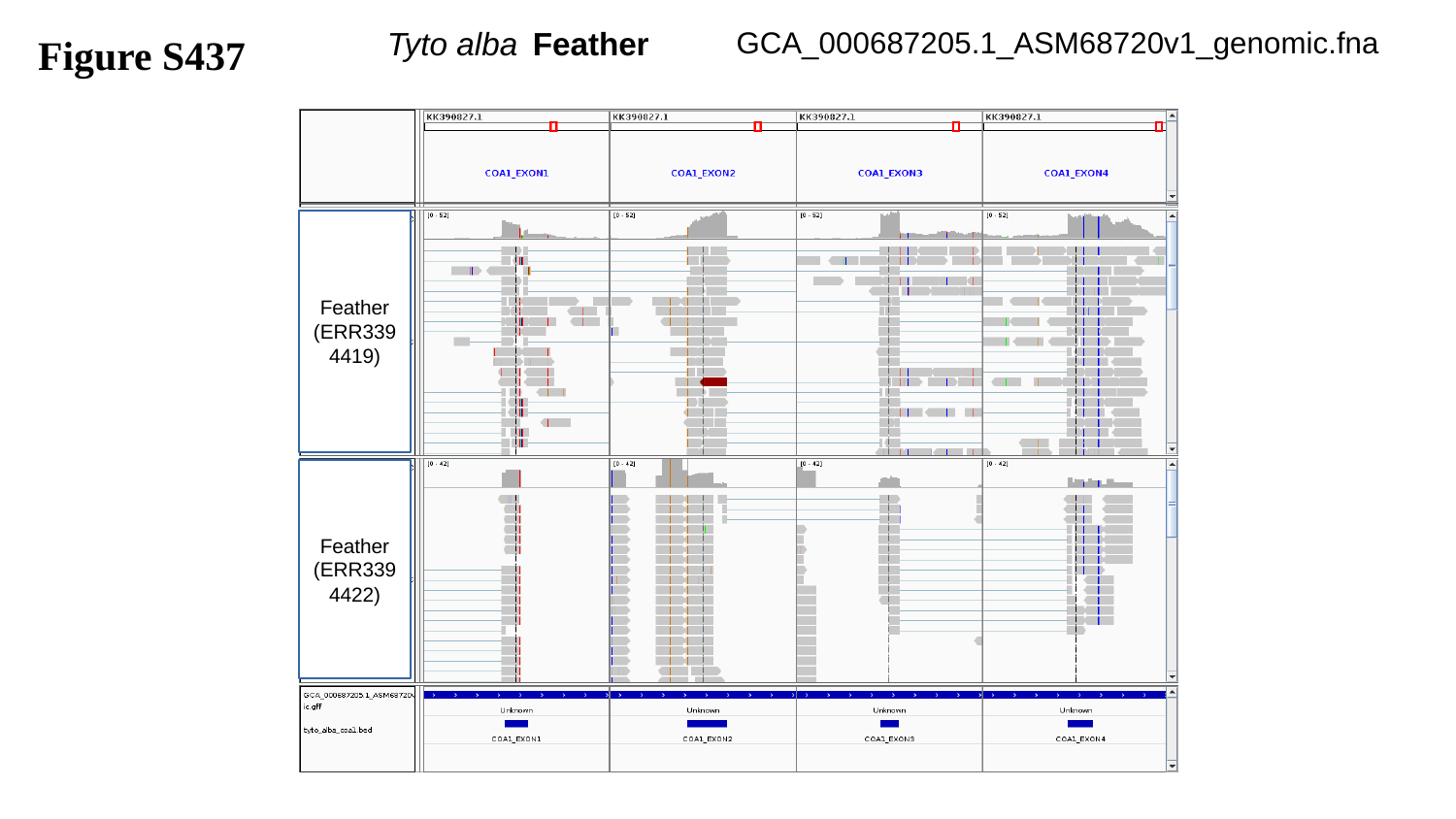

Figure S437
GCA_000687205.1_ASM68720v1_genomic.fna
Tyto alba	Feather
Feather
(ERR3394419)
Feather
(ERR3394422)

### Slide 438
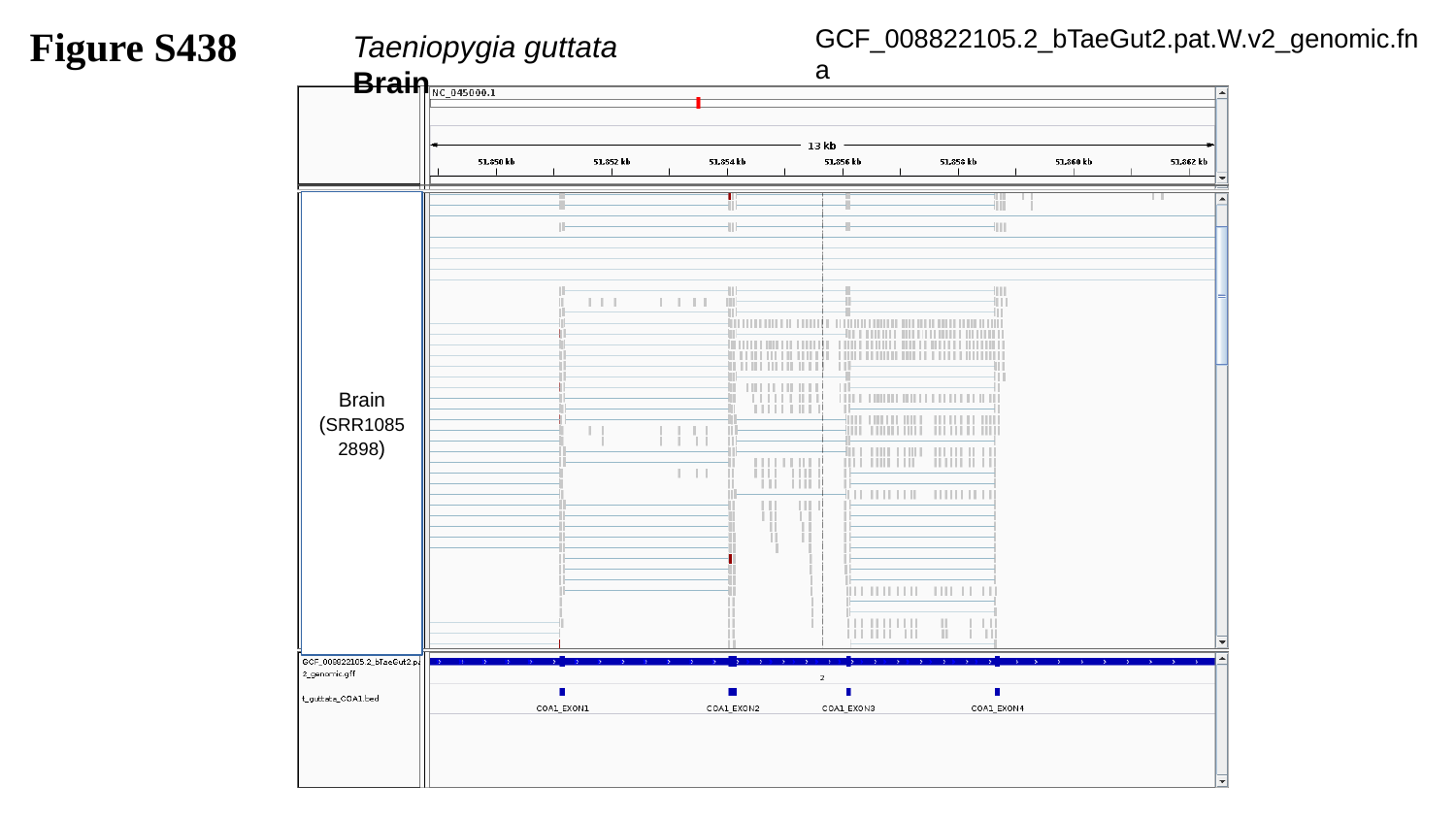

Figure S438
GCF_008822105.2_bTaeGut2.pat.W.v2_genomic.fna
Taeniopygia guttata		Brain
Brain
(SRR10852898)

### Slide 439
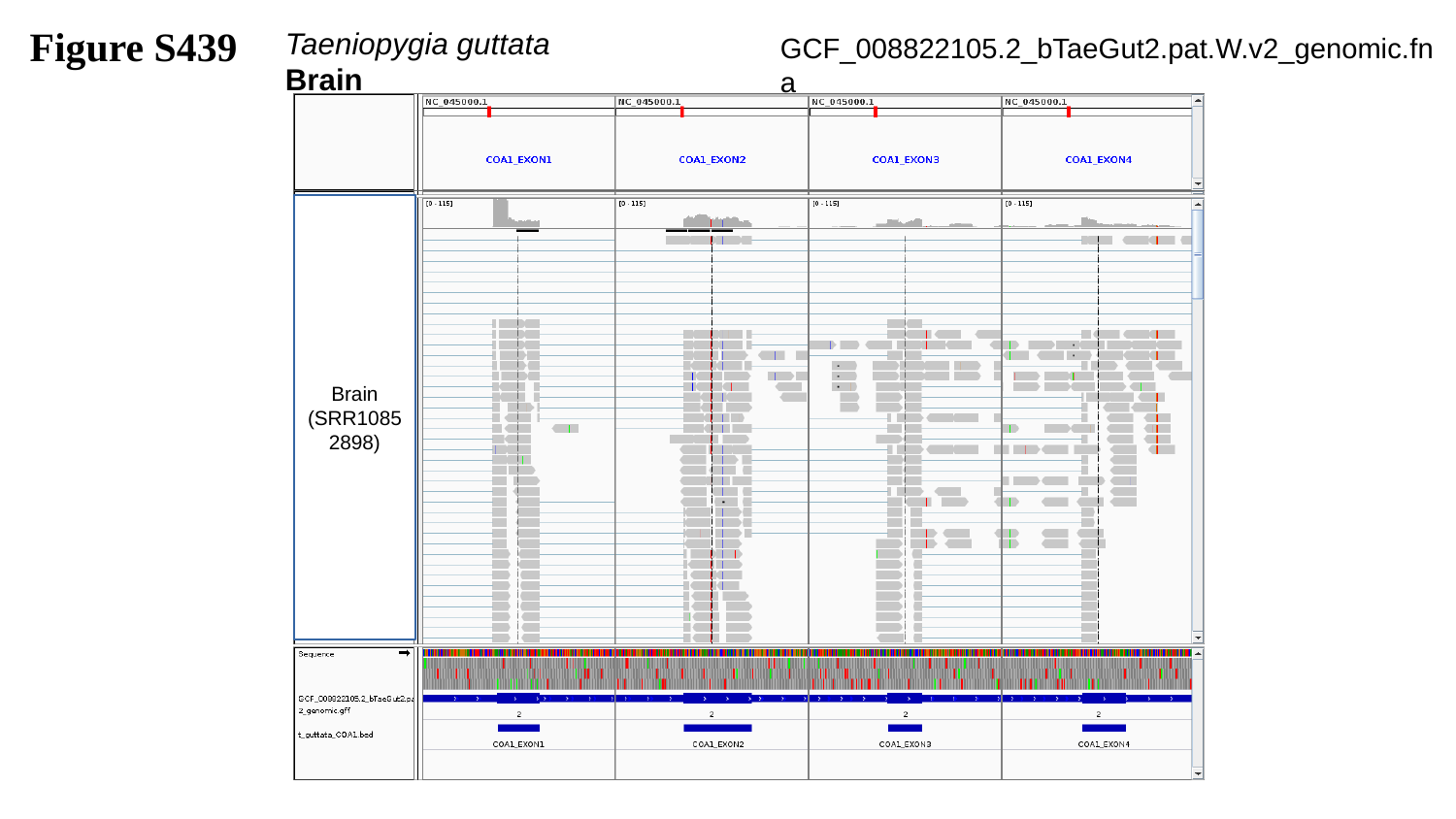

Figure S439
Taeniopygia guttata		Brain
GCF_008822105.2_bTaeGut2.pat.W.v2_genomic.fna
Brain
(SRR10852898)

### Slide 440
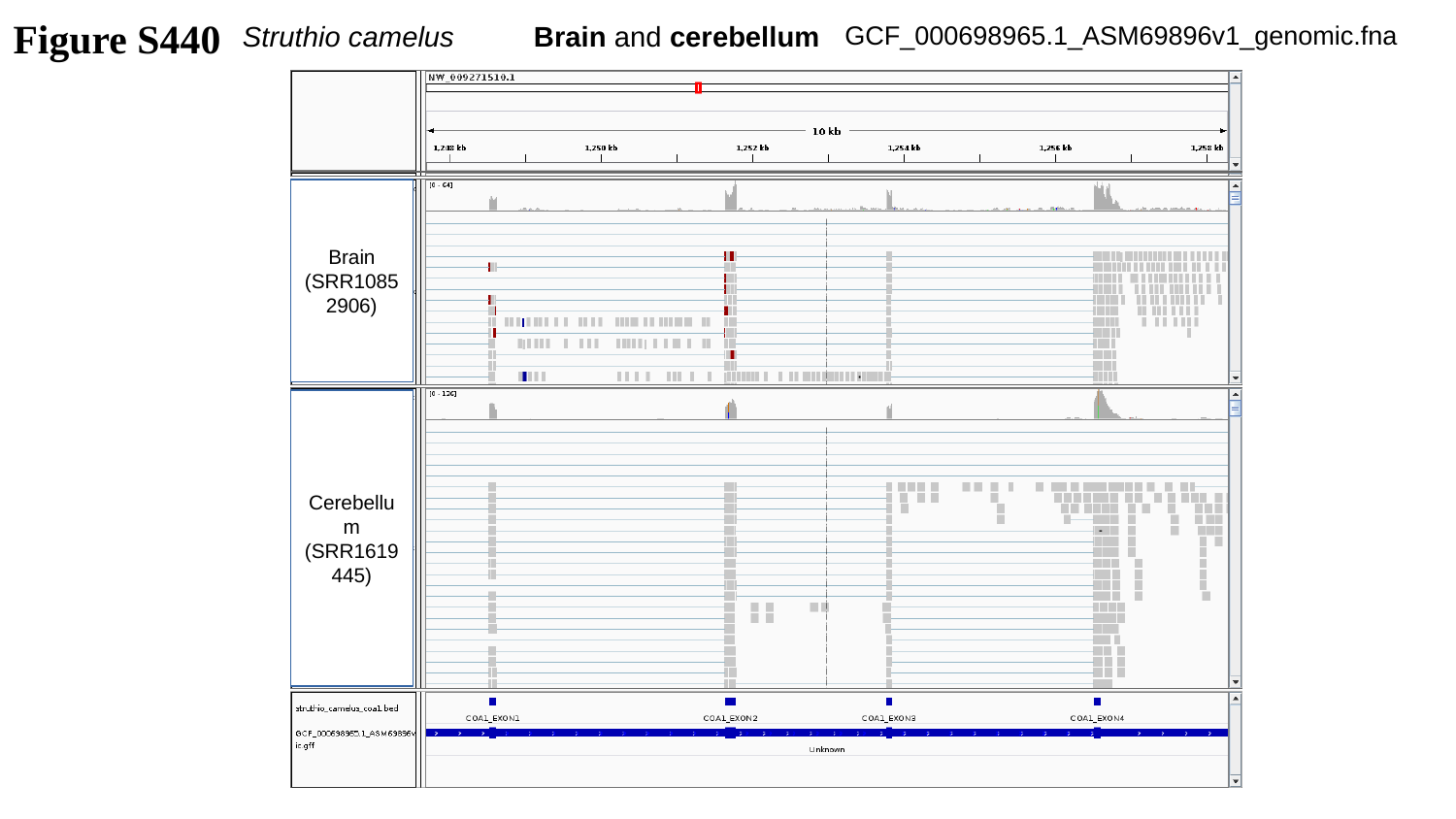

Figure S440
Struthio camelus	Brain and cerebellum
GCF_000698965.1_ASM69896v1_genomic.fna
Brain
(SRR10852906)
Cerebellum
(SRR1619445)

### Slide 441
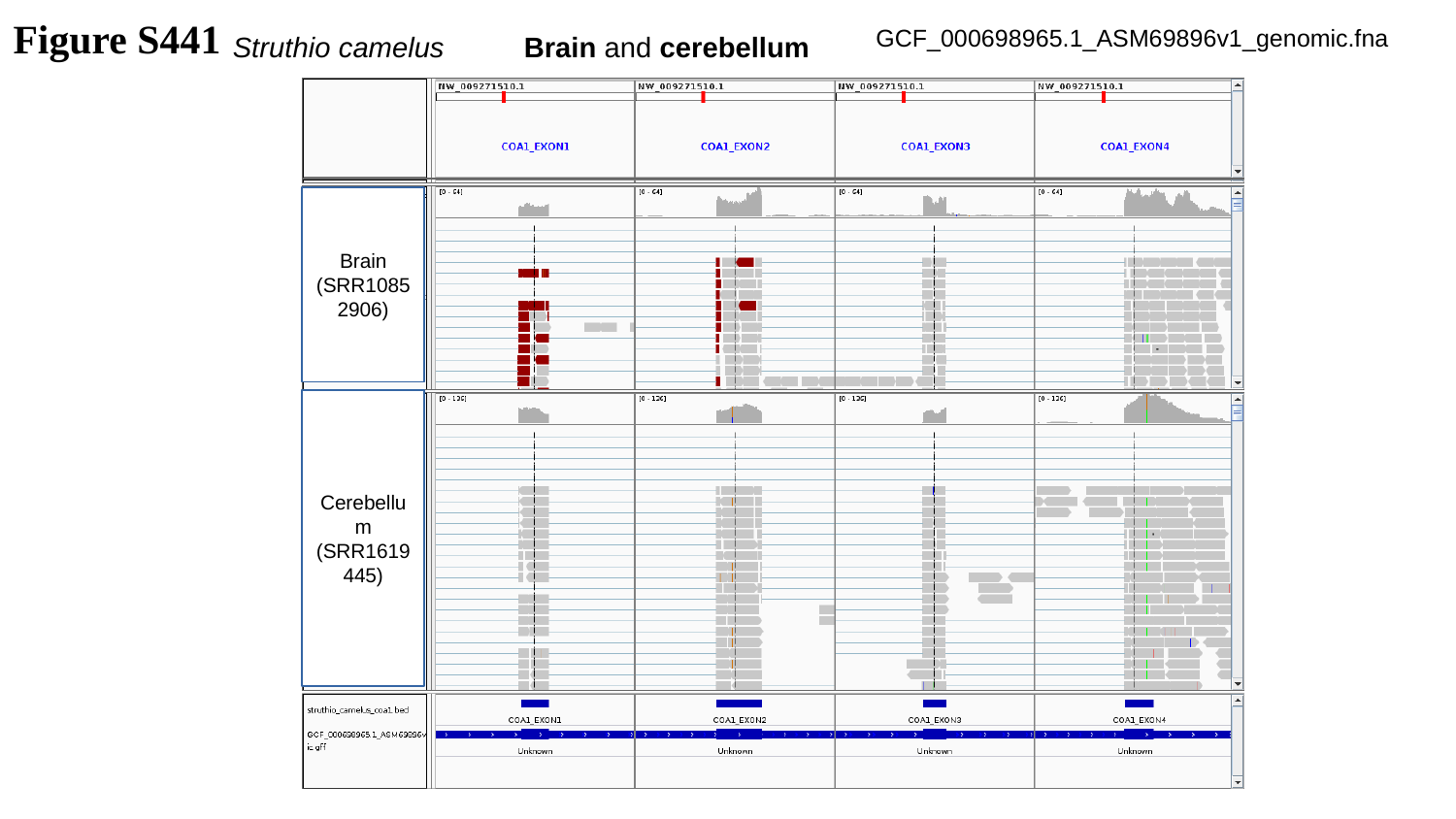

Figure S441
GCF_000698965.1_ASM69896v1_genomic.fna
Struthio camelus	Brain and cerebellum
Brain
(SRR10852906)
Cerebellum
(SRR1619445)

### Slide 442
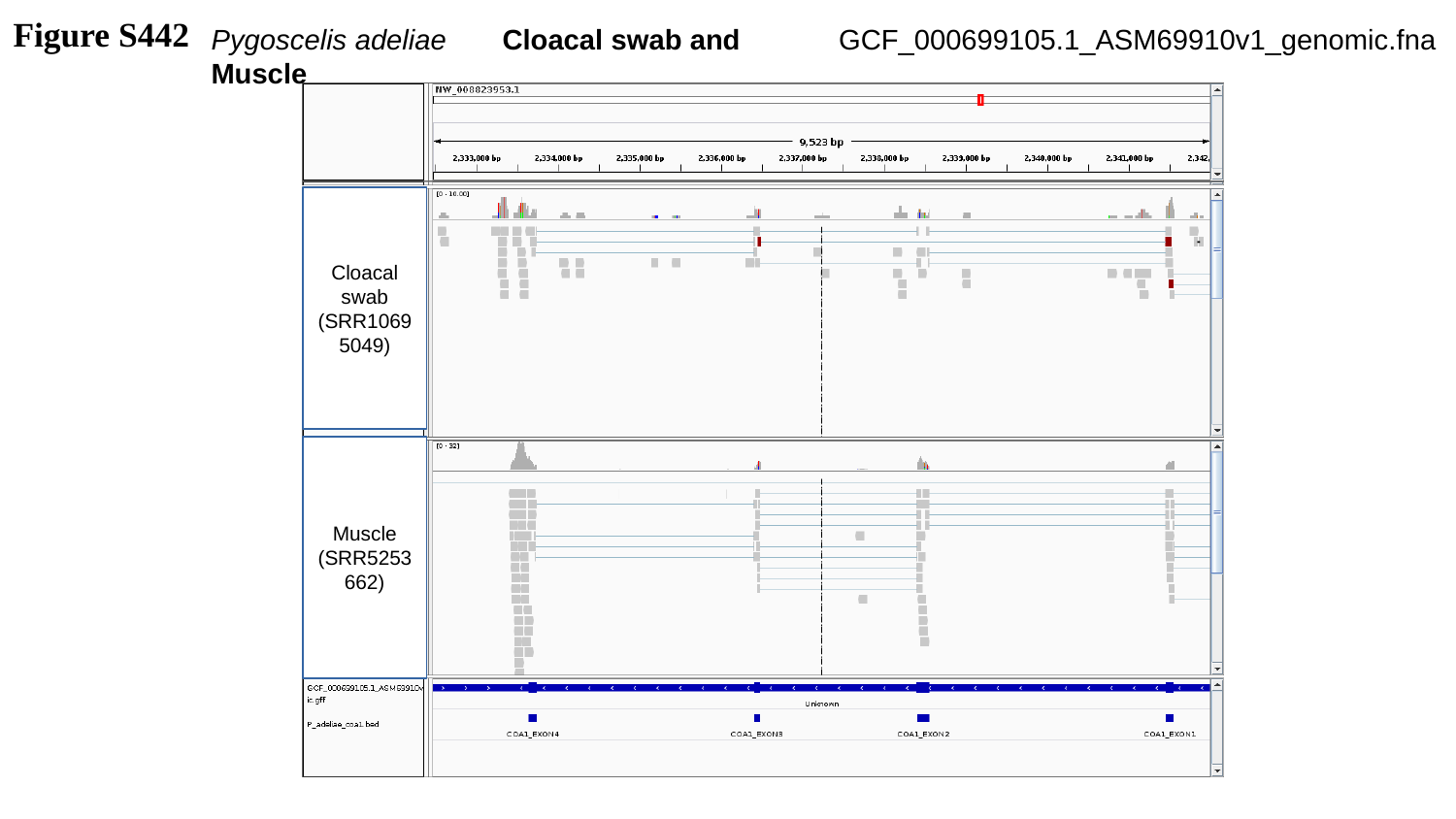

Figure S442
Pygoscelis adeliae	Cloacal swab and Muscle
GCF_000699105.1_ASM69910v1_genomic.fna
Cloacal swab
(SRR10695049)
Muscle
(SRR5253662)

### Slide 443
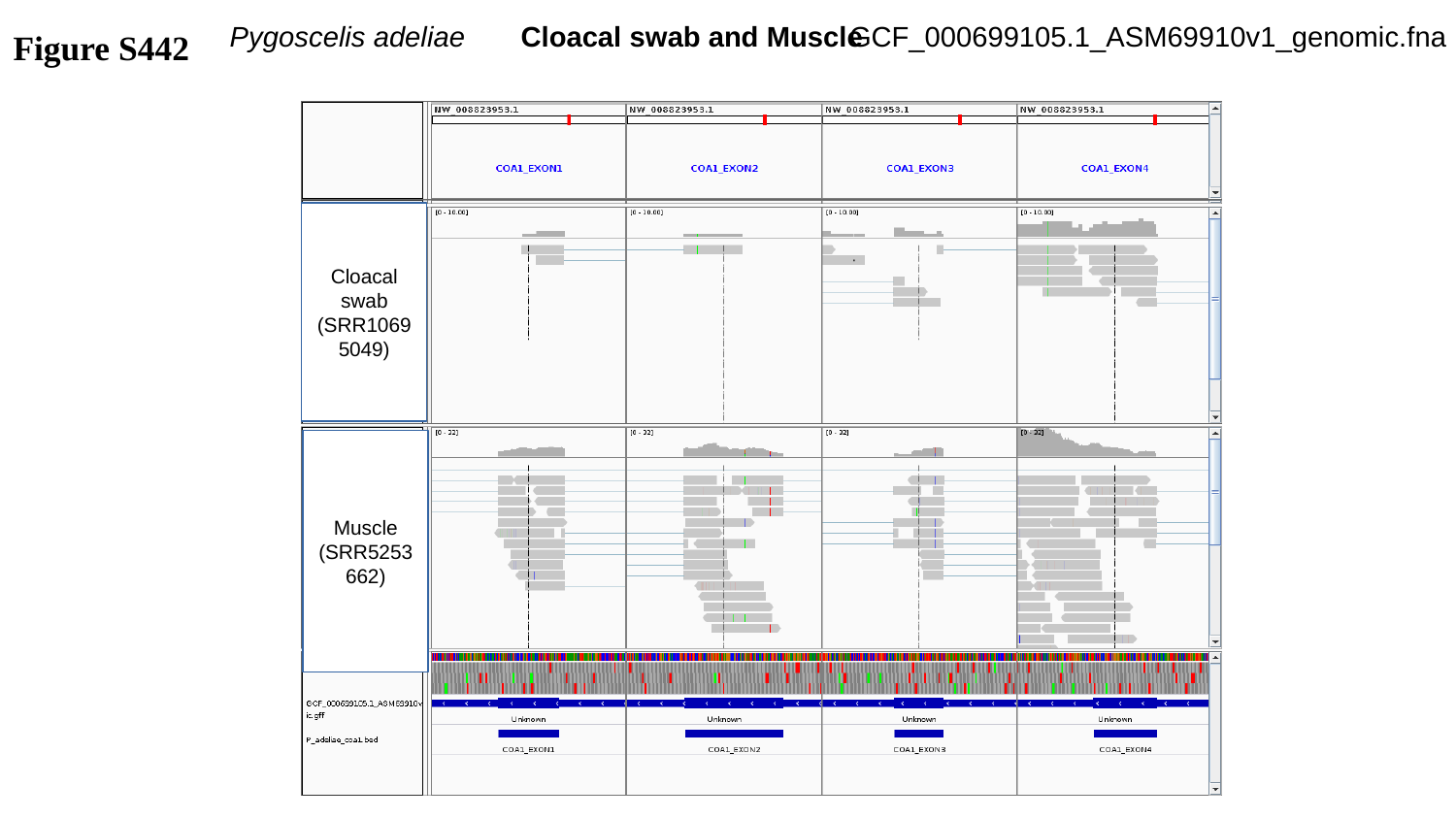

GCF_000699105.1_ASM69910v1_genomic.fna
Figure S442
Pygoscelis adeliae	Cloacal swab and Muscle
Cloacal swab
(SRR10695049)
Muscle
(SRR5253662)

### Slide 444
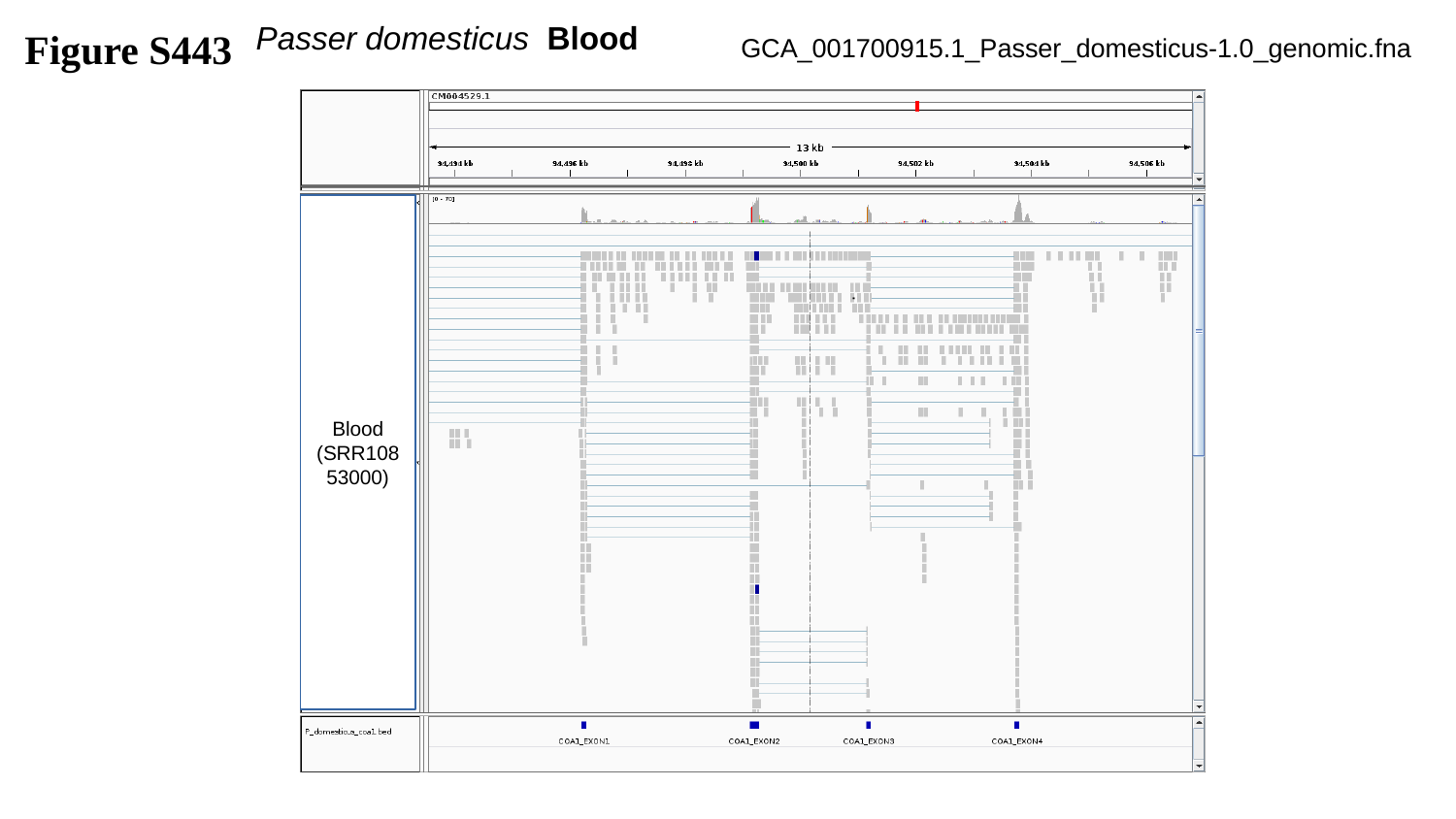

Figure S443
Passer domesticus	Blood
GCA_001700915.1_Passer_domesticus-1.0_genomic.fna
Blood
(SRR10853000)

### Slide 445
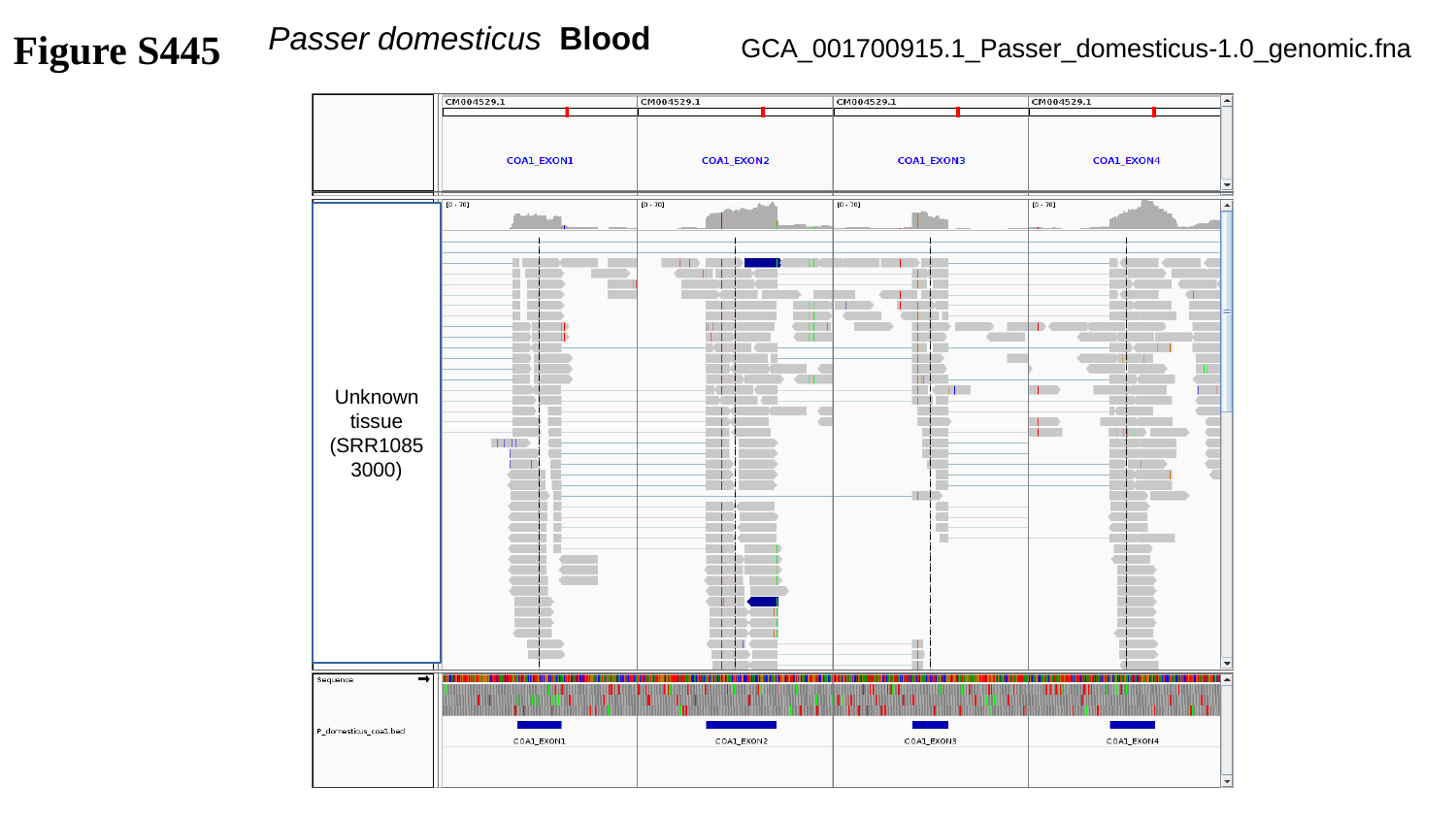

Figure S445
Passer domesticus	Blood
GCA_001700915.1_Passer_domesticus-1.0_genomic.fna
Unknown tissue
(SRR10853000)

### Slide 446
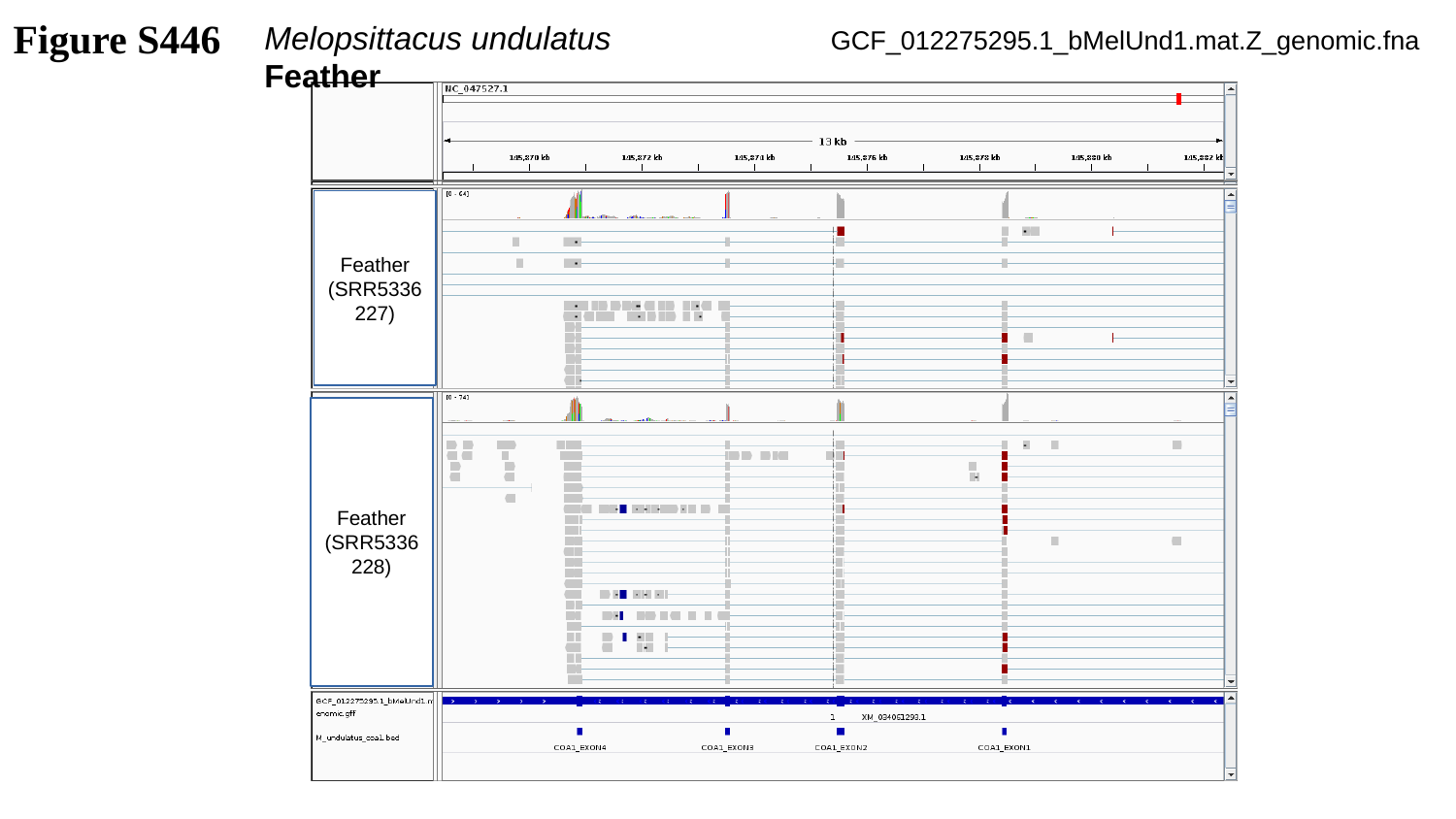

Figure S446
Melopsittacus undulatus	 Feather
GCF_012275295.1_bMelUnd1.mat.Z_genomic.fna
Feather
(SRR5336227)
Feather
(SRR5336228)

### Slide 447
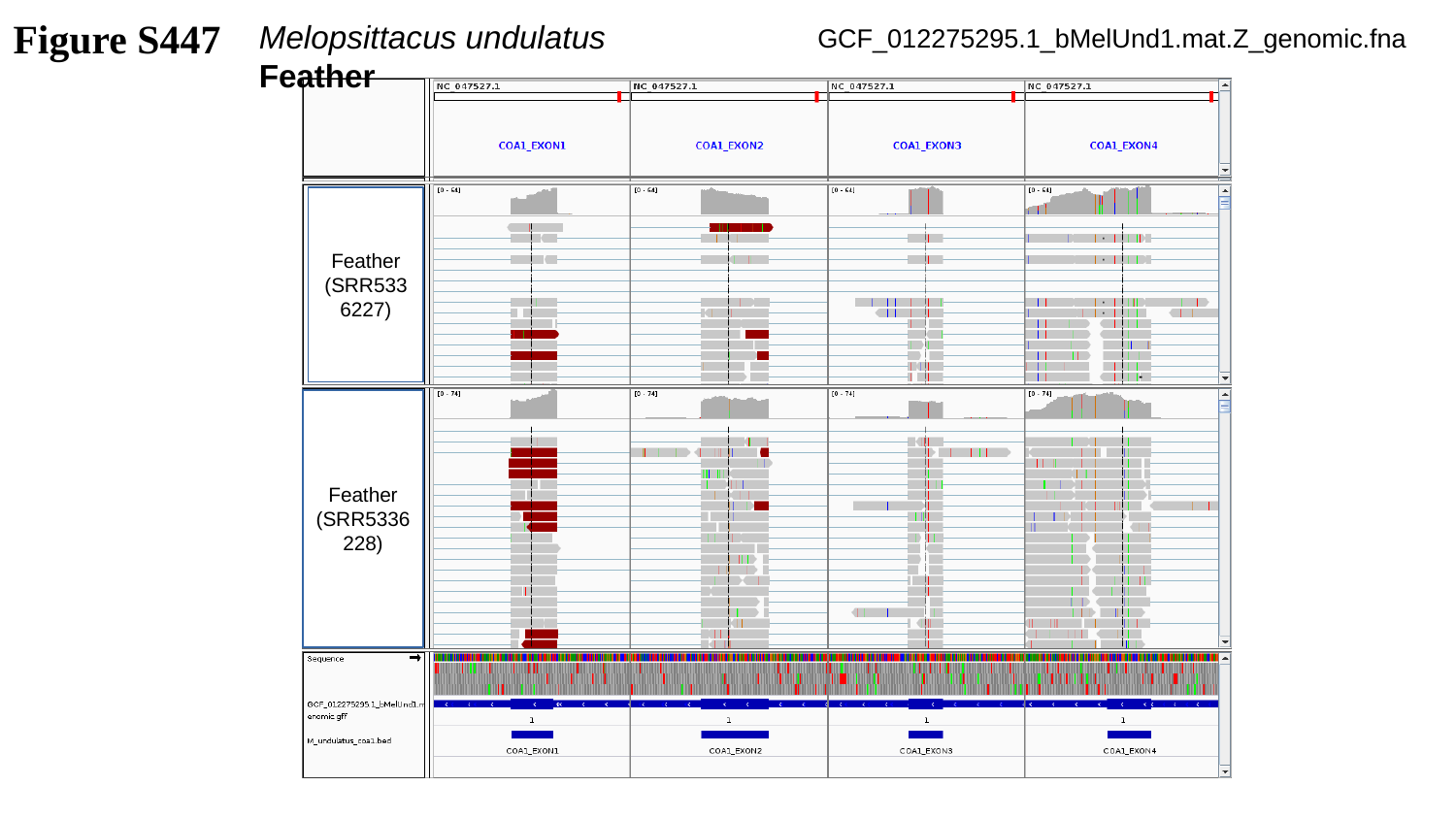

Figure S447
Melopsittacus undulatus	 Feather
GCF_012275295.1_bMelUnd1.mat.Z_genomic.fna
Feather
(SRR5336227)
Feather
(SRR5336228)

### Slide 448
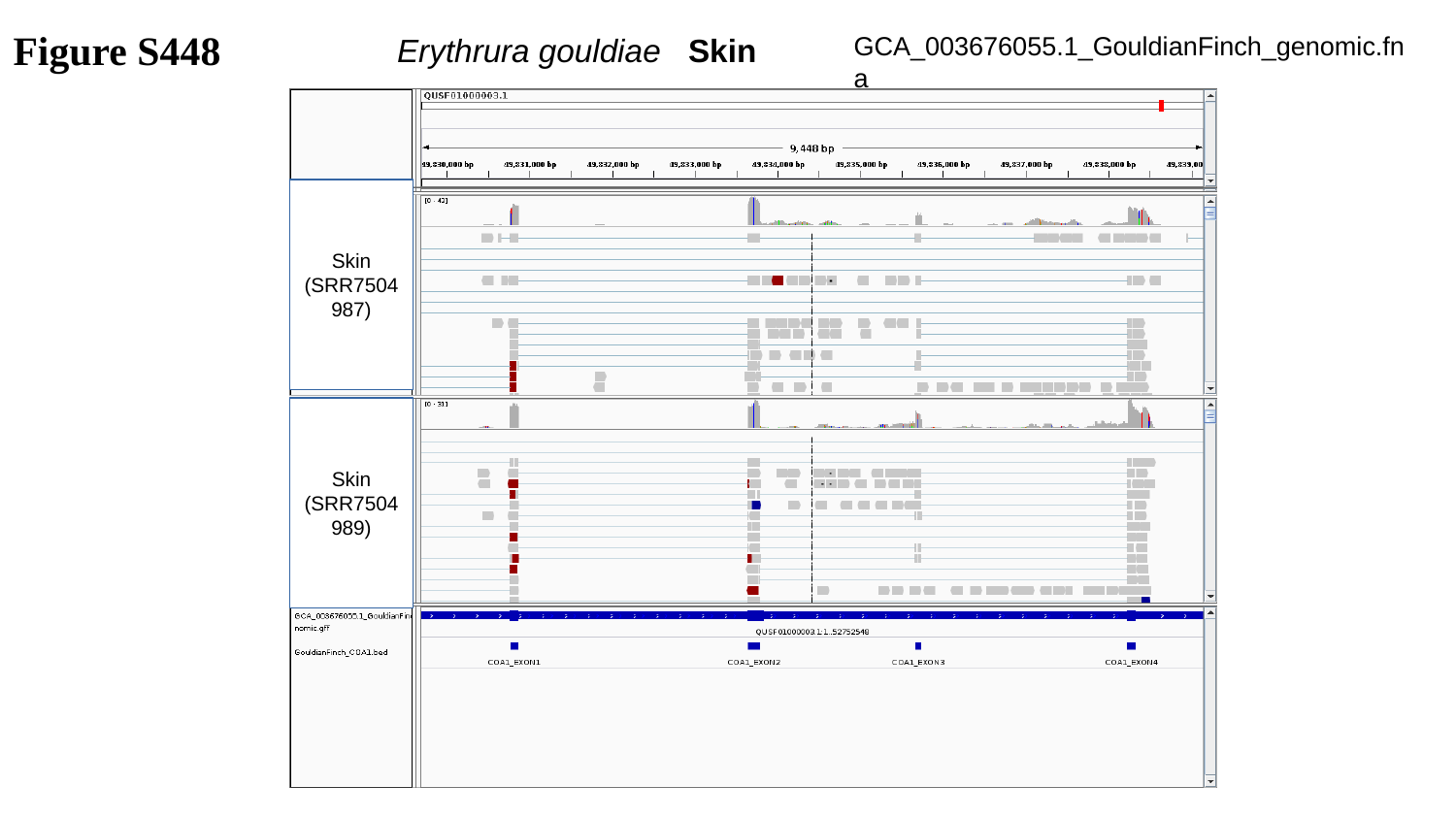

Figure S448
GCA_003676055.1_GouldianFinch_genomic.fna
Erythrura gouldiae	Skin
Skin
(SRR7504987)
Skin
(SRR7504989)

### Slide 449
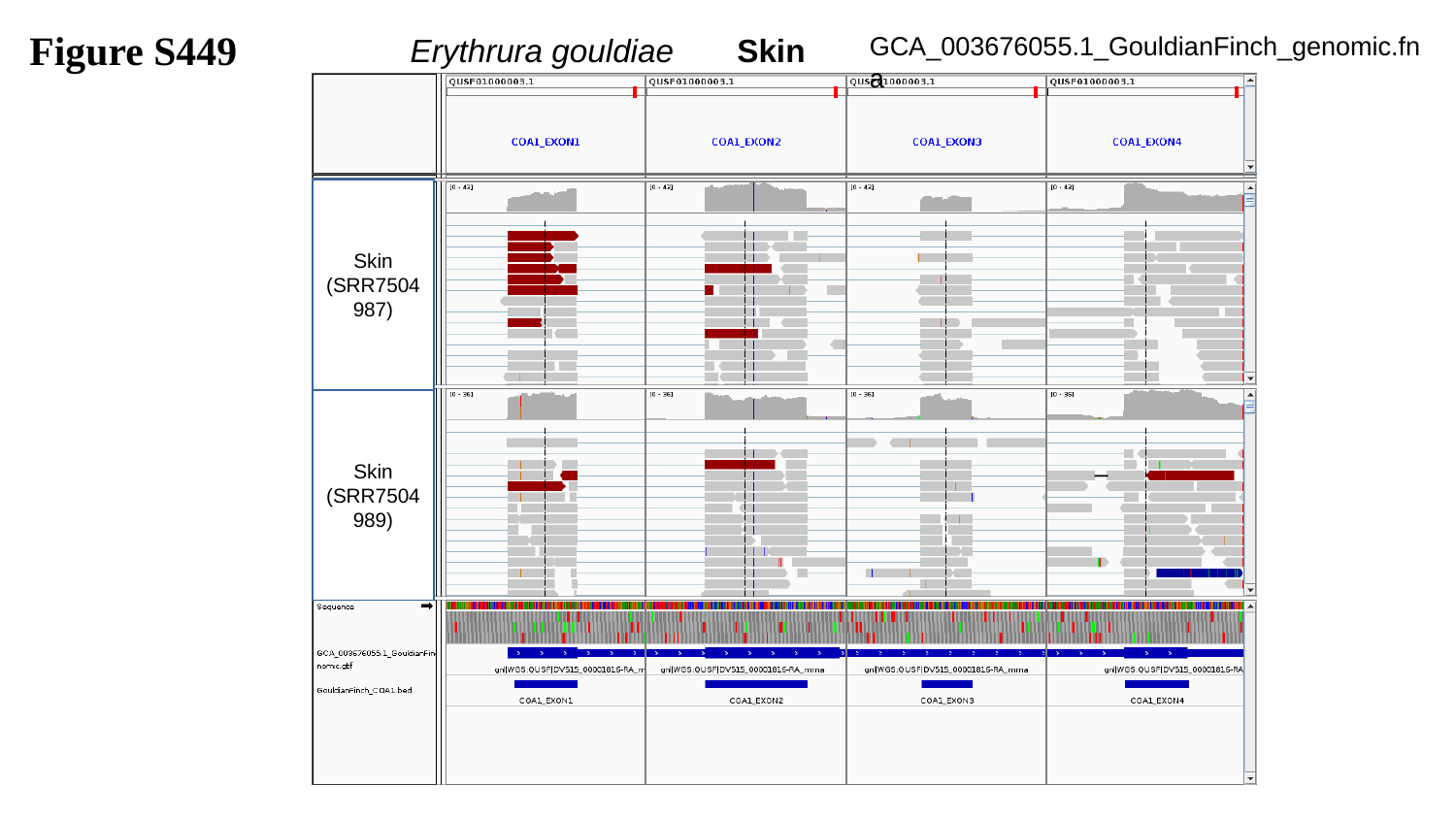

Figure S449
GCA_003676055.1_GouldianFinch_genomic.fna
Erythrura gouldiae Skin
Skin
(SRR7504987)
Skin
(SRR7504989)

### Slide 450
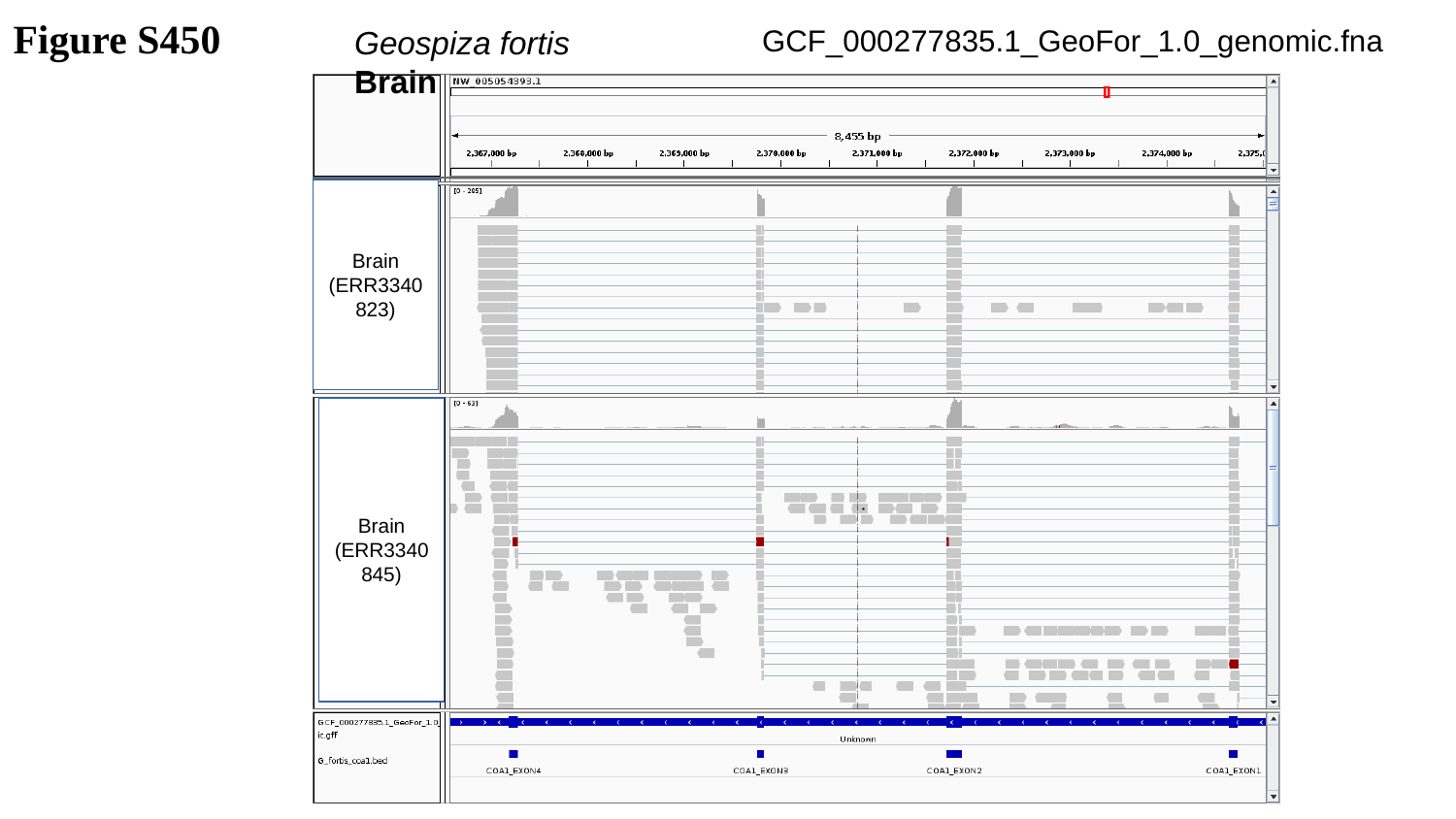

Figure S450
GCF_000277835.1_GeoFor_1.0_genomic.fna
Geospiza fortis	 Brain
Brain
(ERR3340823)
Brain
(ERR3340845)

### Slide 451
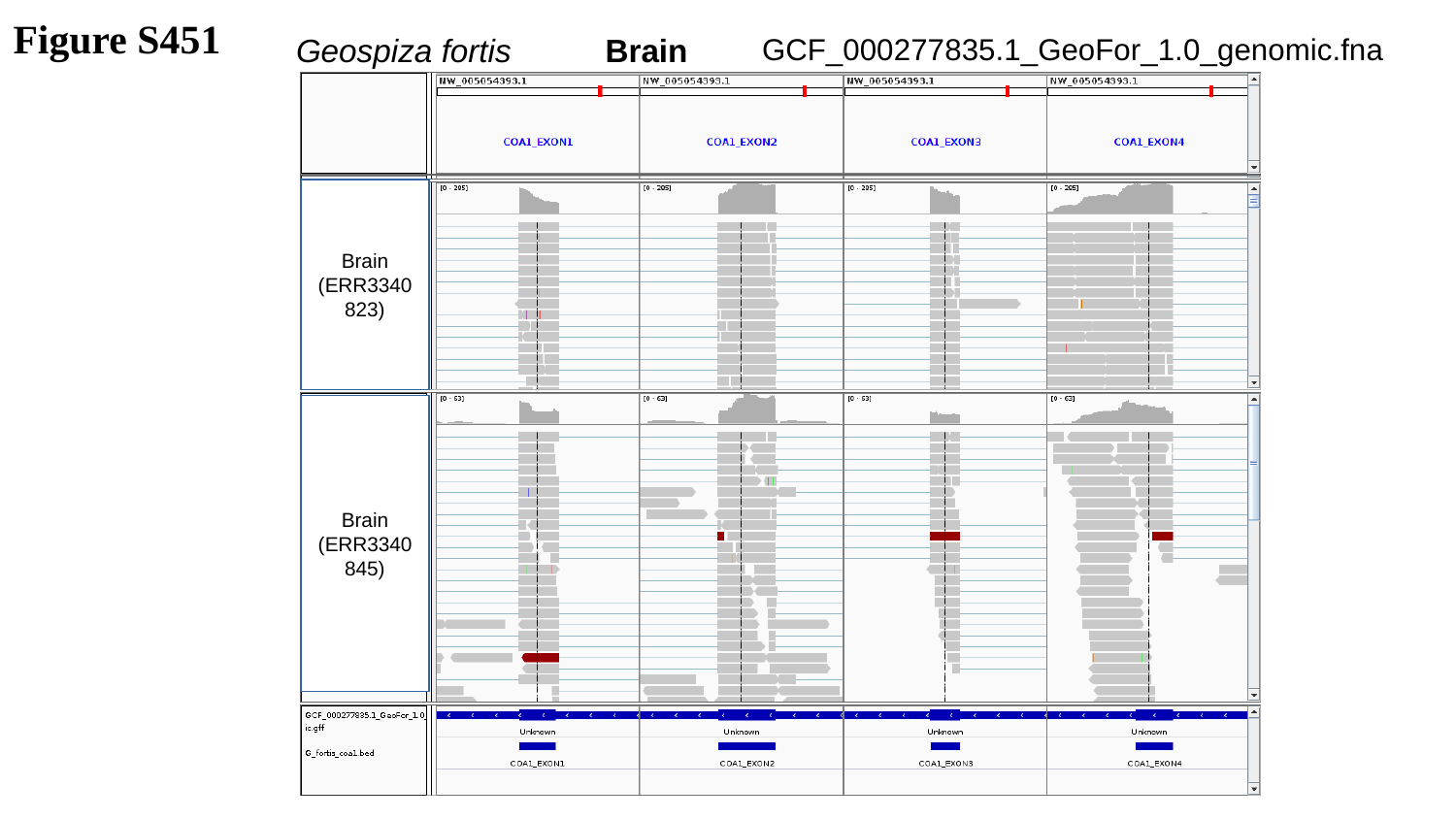

Figure S451
GCF_000277835.1_GeoFor_1.0_genomic.fna
Geospiza fortis	 Brain
Brain
(ERR3340823)
Brain
(ERR3340845)

### Slide 452
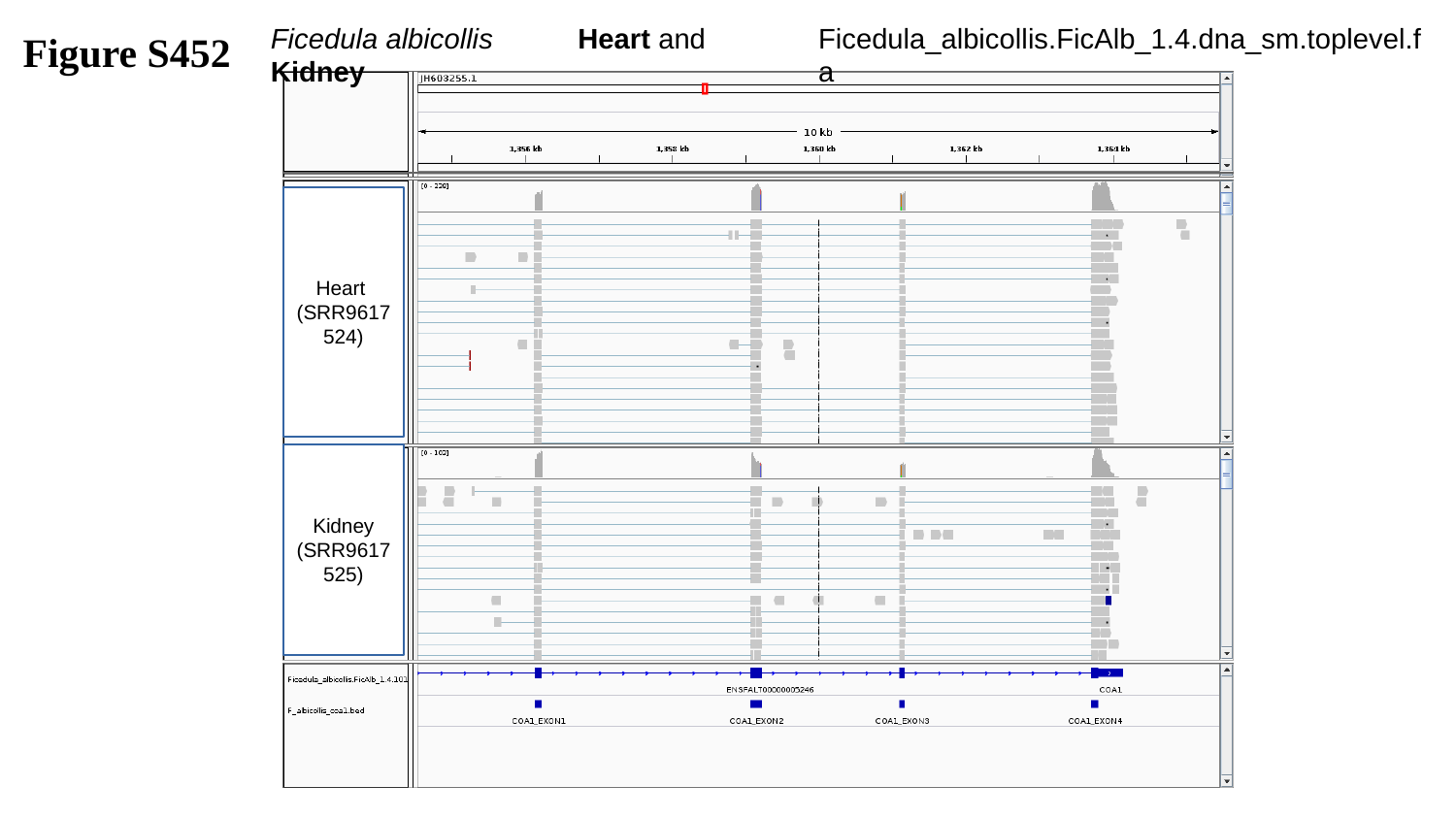

Figure S452
Ficedula_albicollis.FicAlb_1.4.dna_sm.toplevel.fa
Ficedula albicollis	 Heart and Kidney
Heart
(SRR9617524)
Kidney
(SRR9617525)

### Slide 453
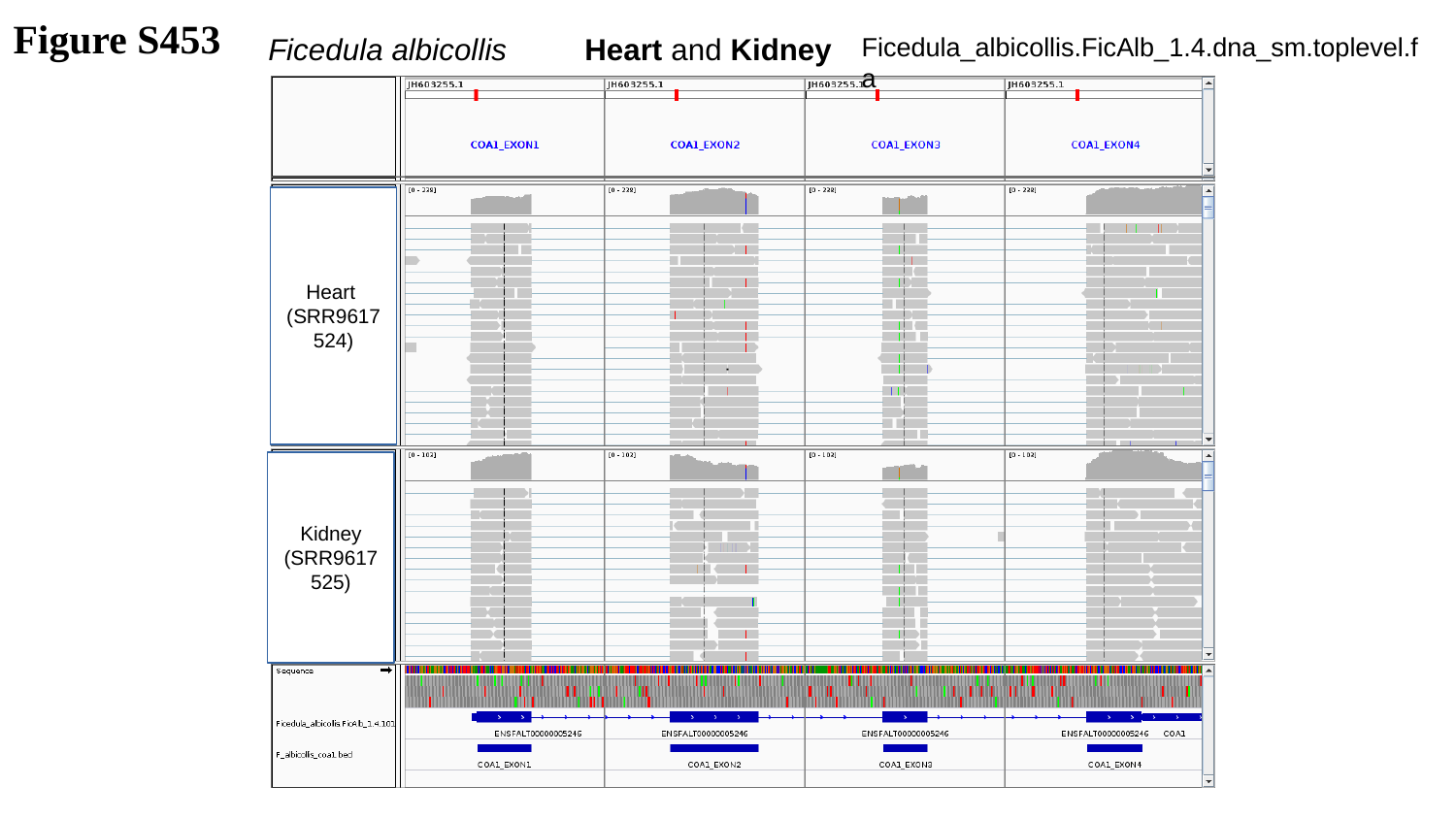

Figure S453
Ficedula_albicollis.FicAlb_1.4.dna_sm.toplevel.fa
Ficedula albicollis	 Heart and Kidney
Heart
(SRR9617524)
Kidney
(SRR9617525)

### Slide 454
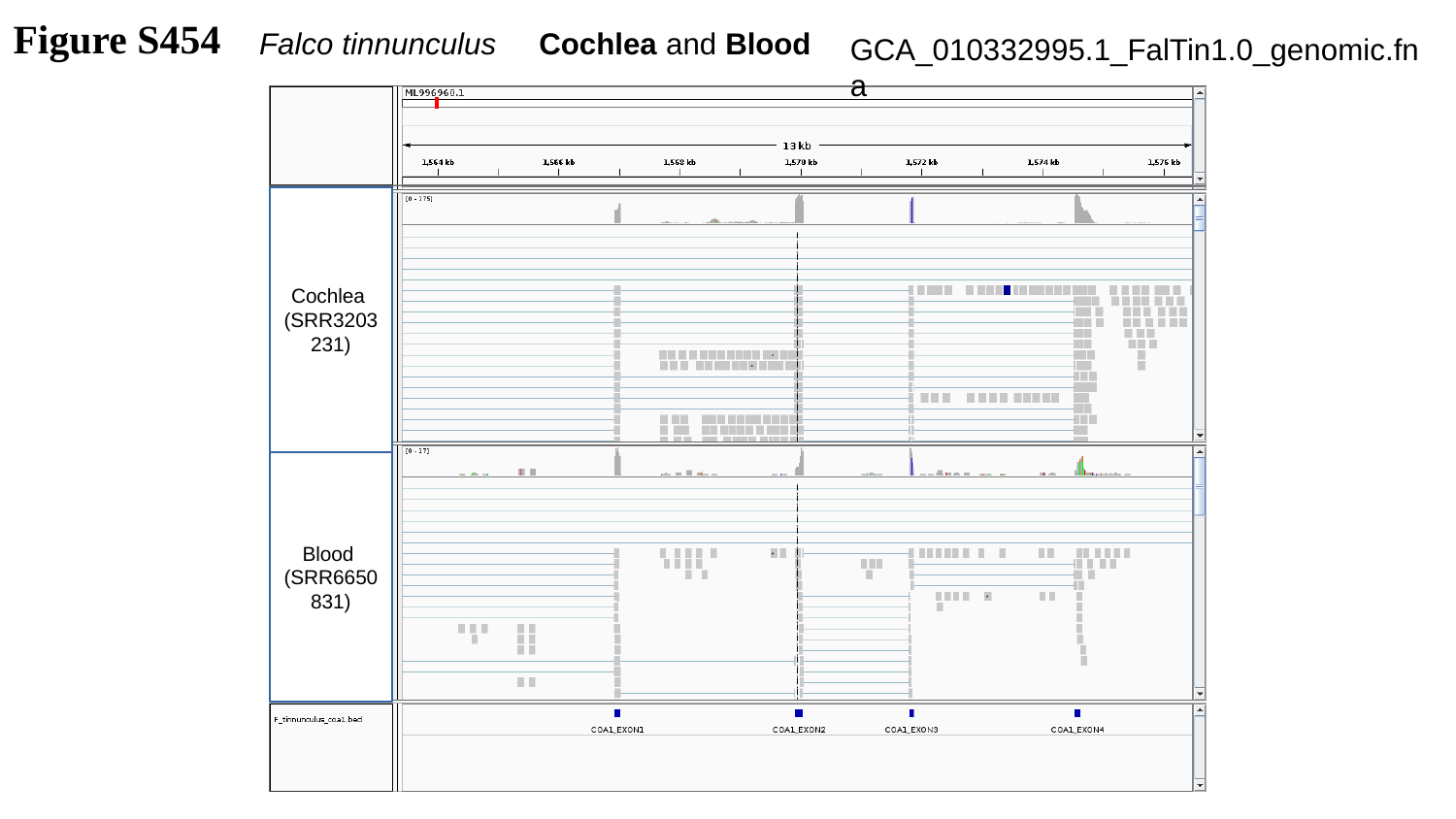

Figure S454
Falco tinnunculus Cochlea and Blood
GCA_010332995.1_FalTin1.0_genomic.fna
Cochlea
(SRR3203231)
Blood
(SRR6650831)

### Slide 455
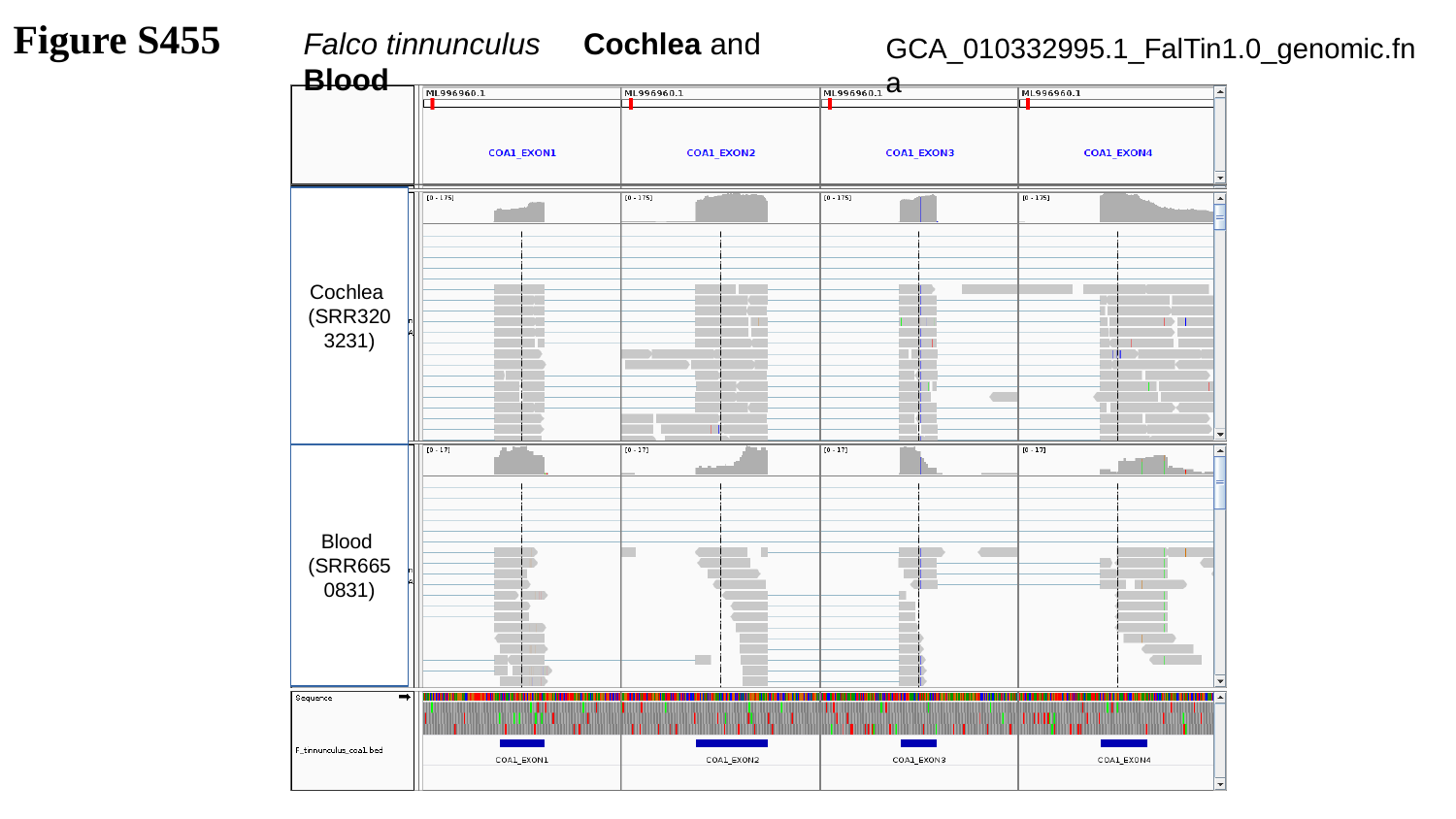

Figure S455
Falco tinnunculus Cochlea and Blood
GCA_010332995.1_FalTin1.0_genomic.fna
Cochlea
(SRR3203231)
Blood
(SRR6650831)

### Slide 456
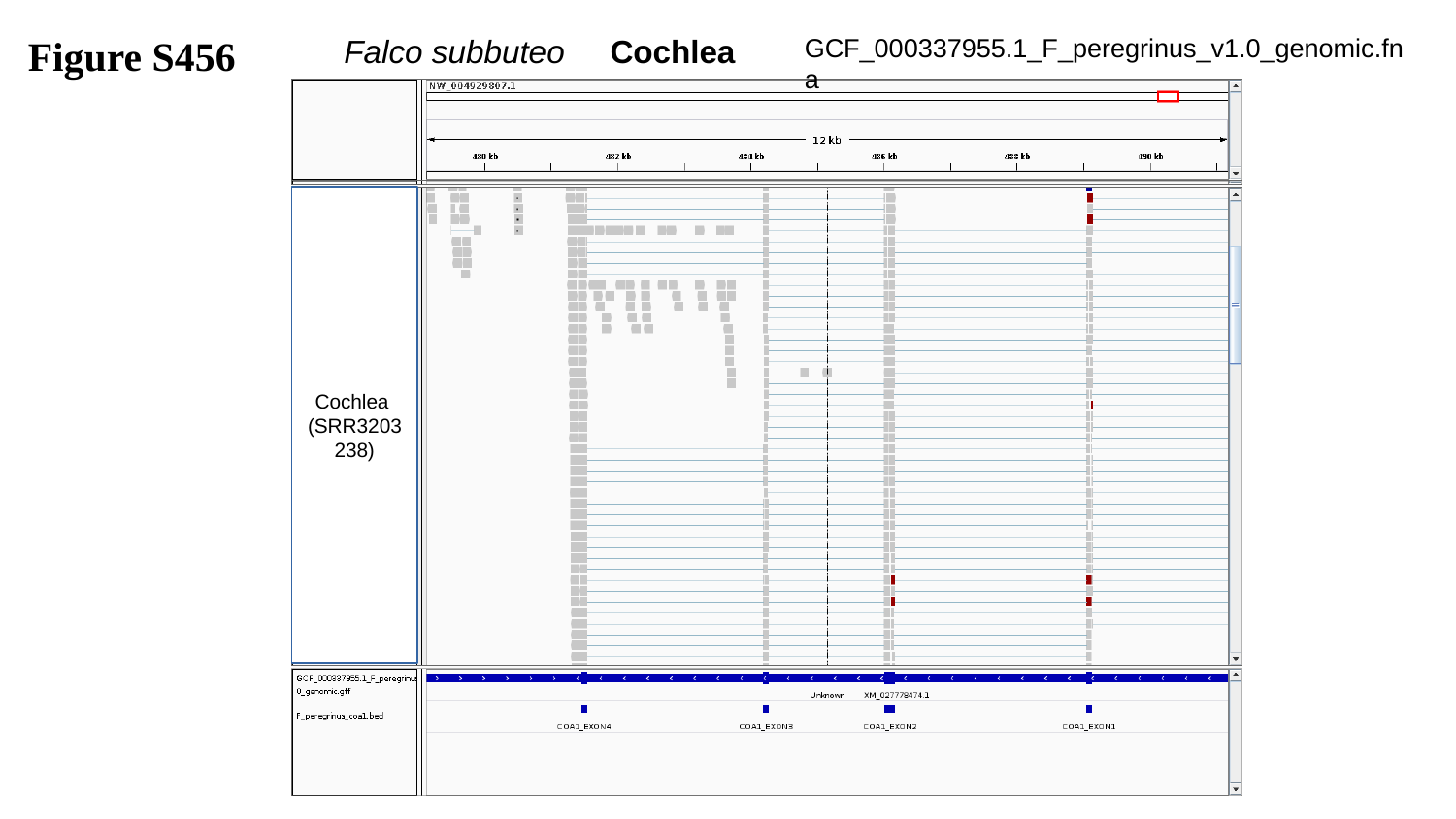

Figure S456
GCF_000337955.1_F_peregrinus_v1.0_genomic.fna
Falco subbuteo Cochlea
Cochlea
(SRR3203238)

### Slide 457
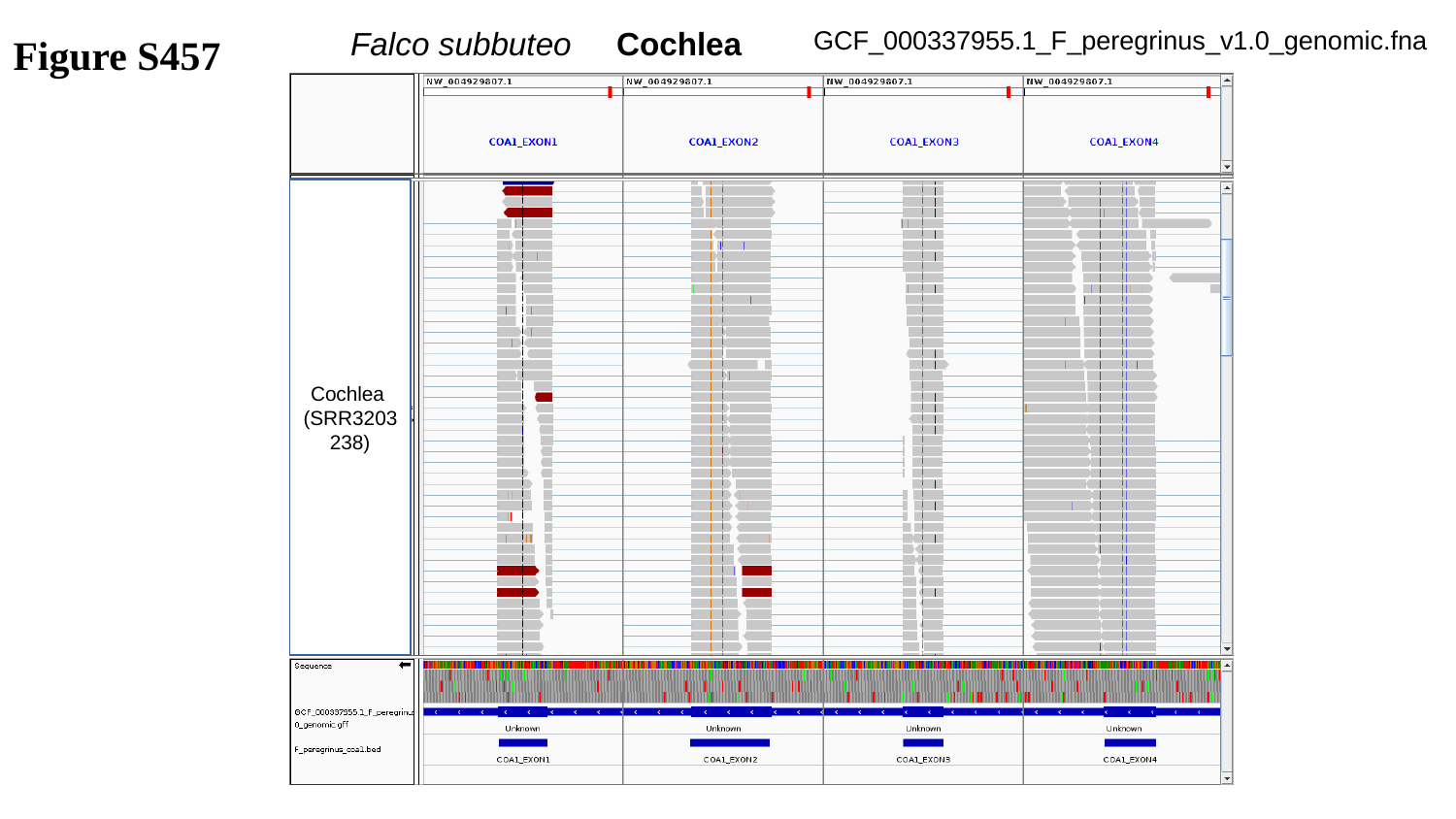

Figure S457
GCF_000337955.1_F_peregrinus_v1.0_genomic.fna
Falco subbuteo Cochlea
Cochlea
(SRR3203238)

### Slide 458
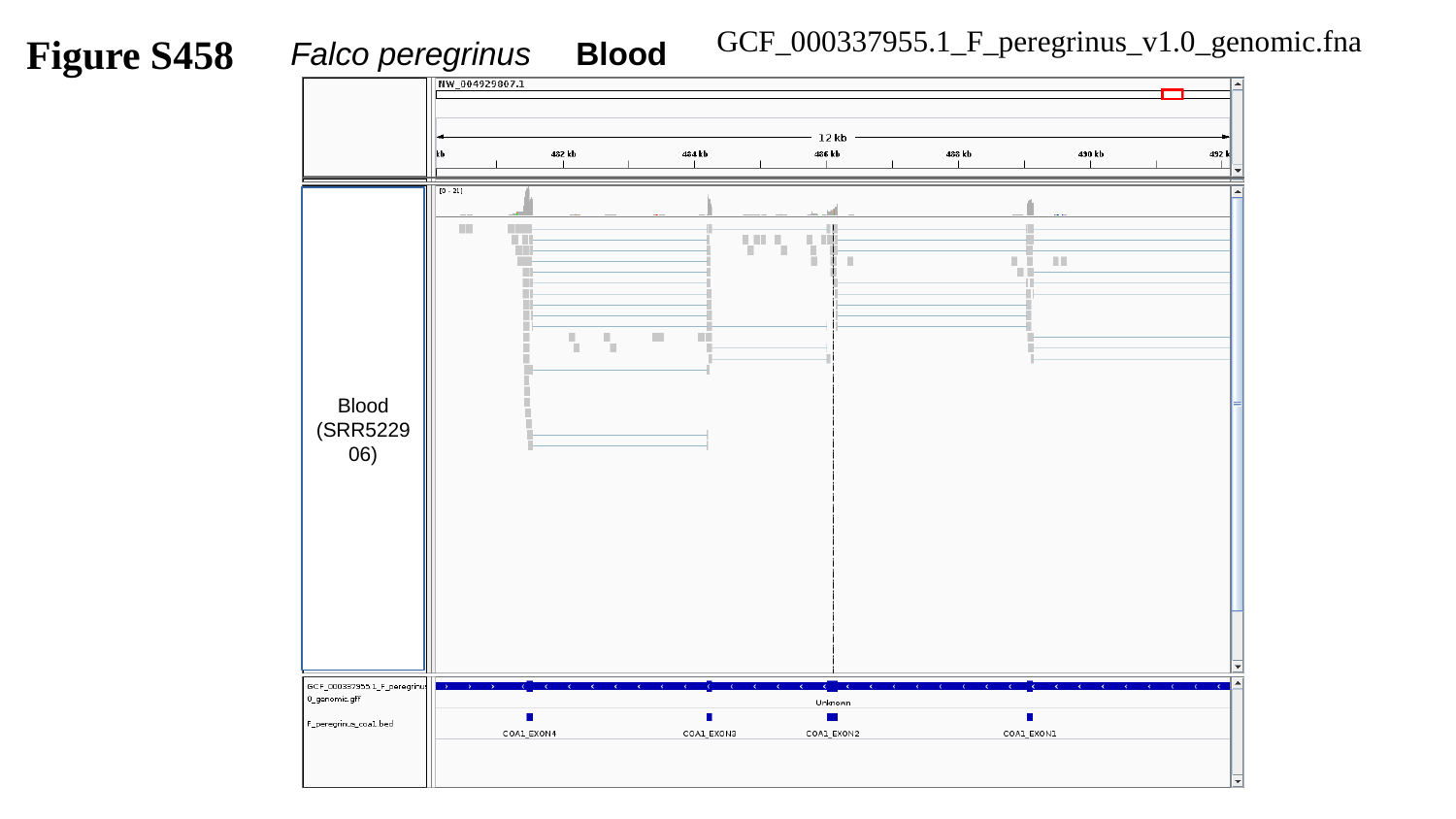

Figure S458
GCF_000337955.1_F_peregrinus_v1.0_genomic.fna
Falco peregrinus Blood
Blood
(SRR522906)

### Slide 459
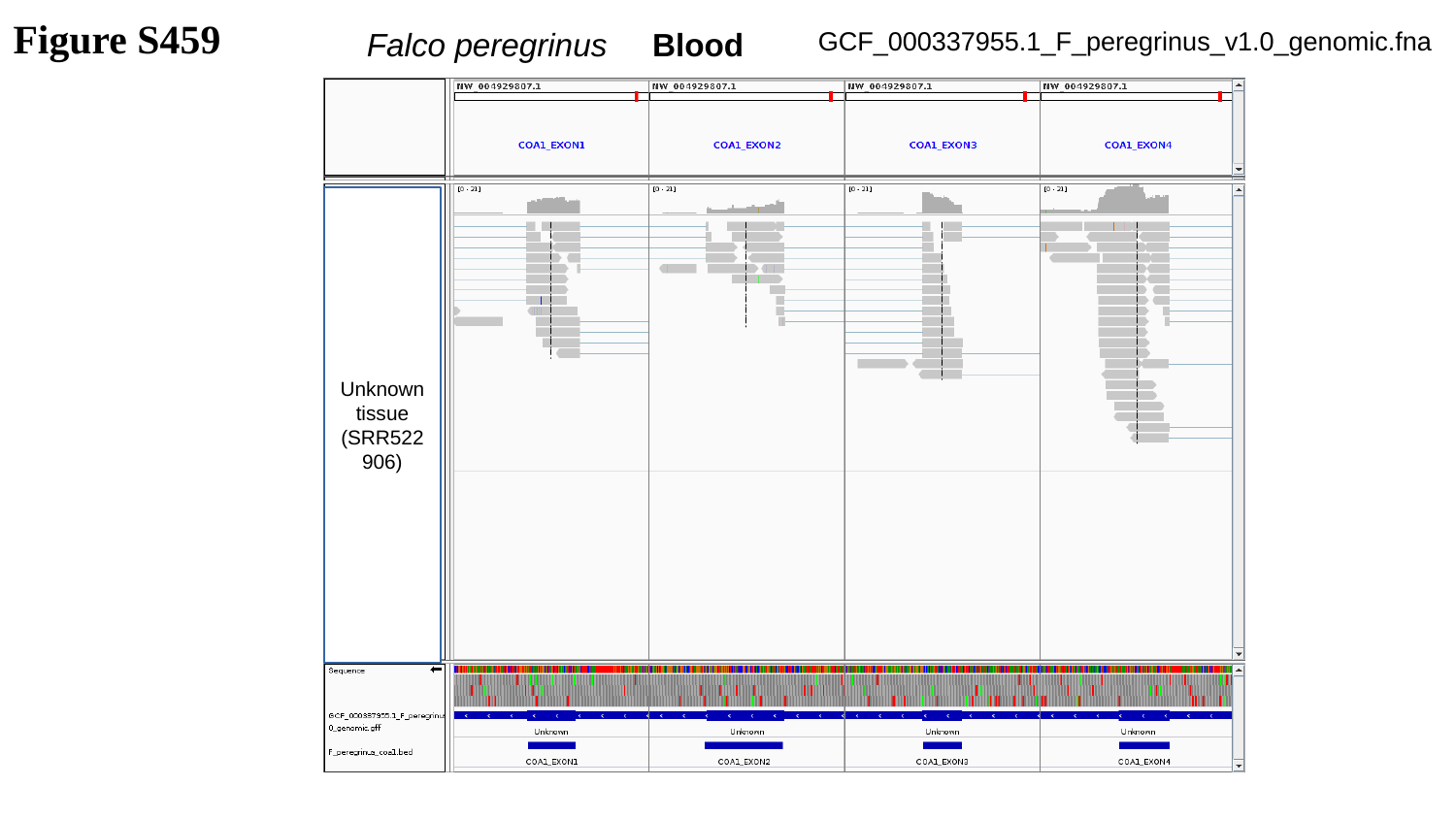

Figure S459
GCF_000337955.1_F_peregrinus_v1.0_genomic.fna
Falco peregrinus Blood
Unknown tissue
(SRR522906)

### Slide 460
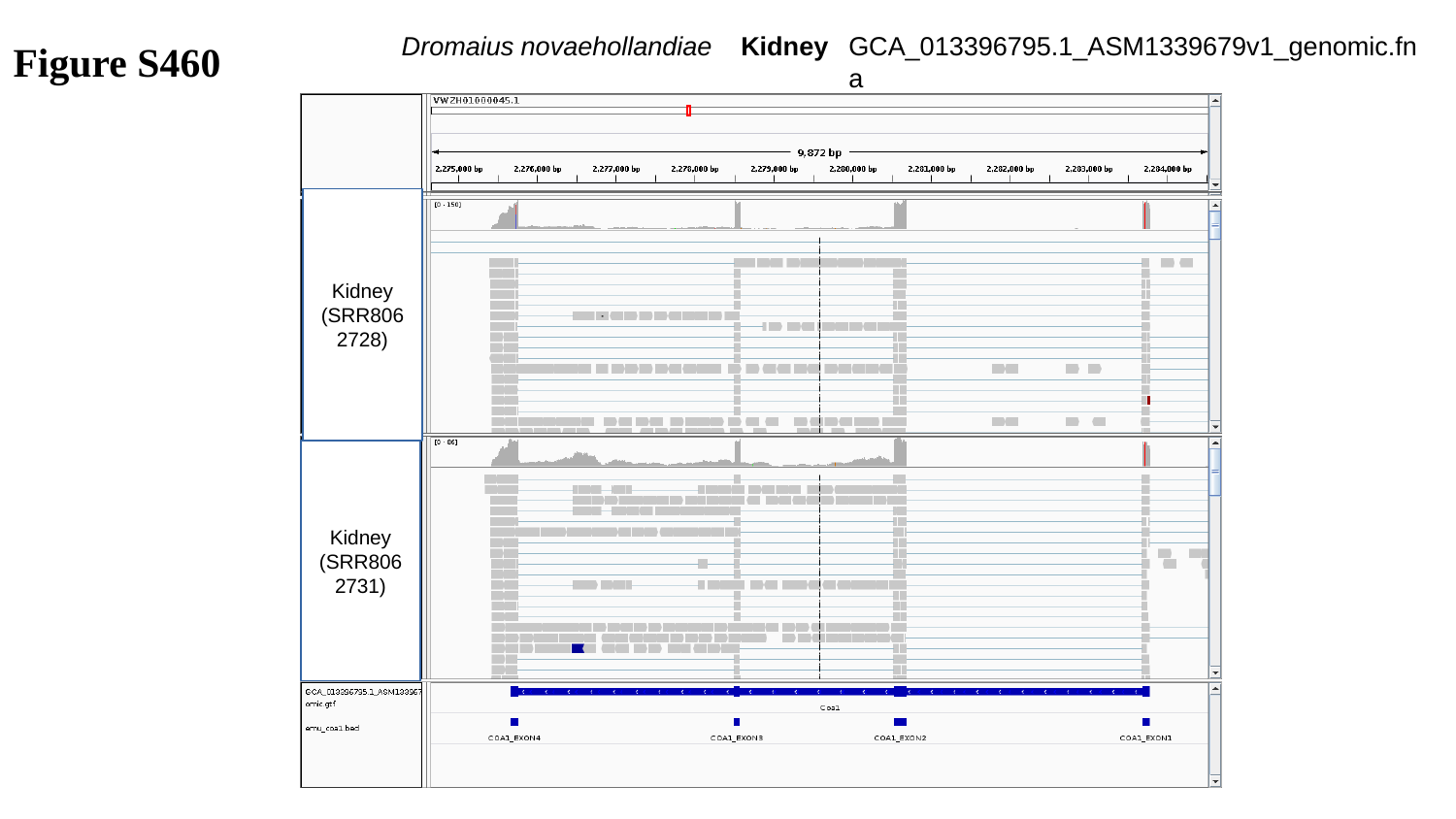

Figure S460
Dromaius novaehollandiae Kidney
GCA_013396795.1_ASM1339679v1_genomic.fna
Kidney
(SRR8062728)
Kidney
(SRR8062731)

### Slide 461
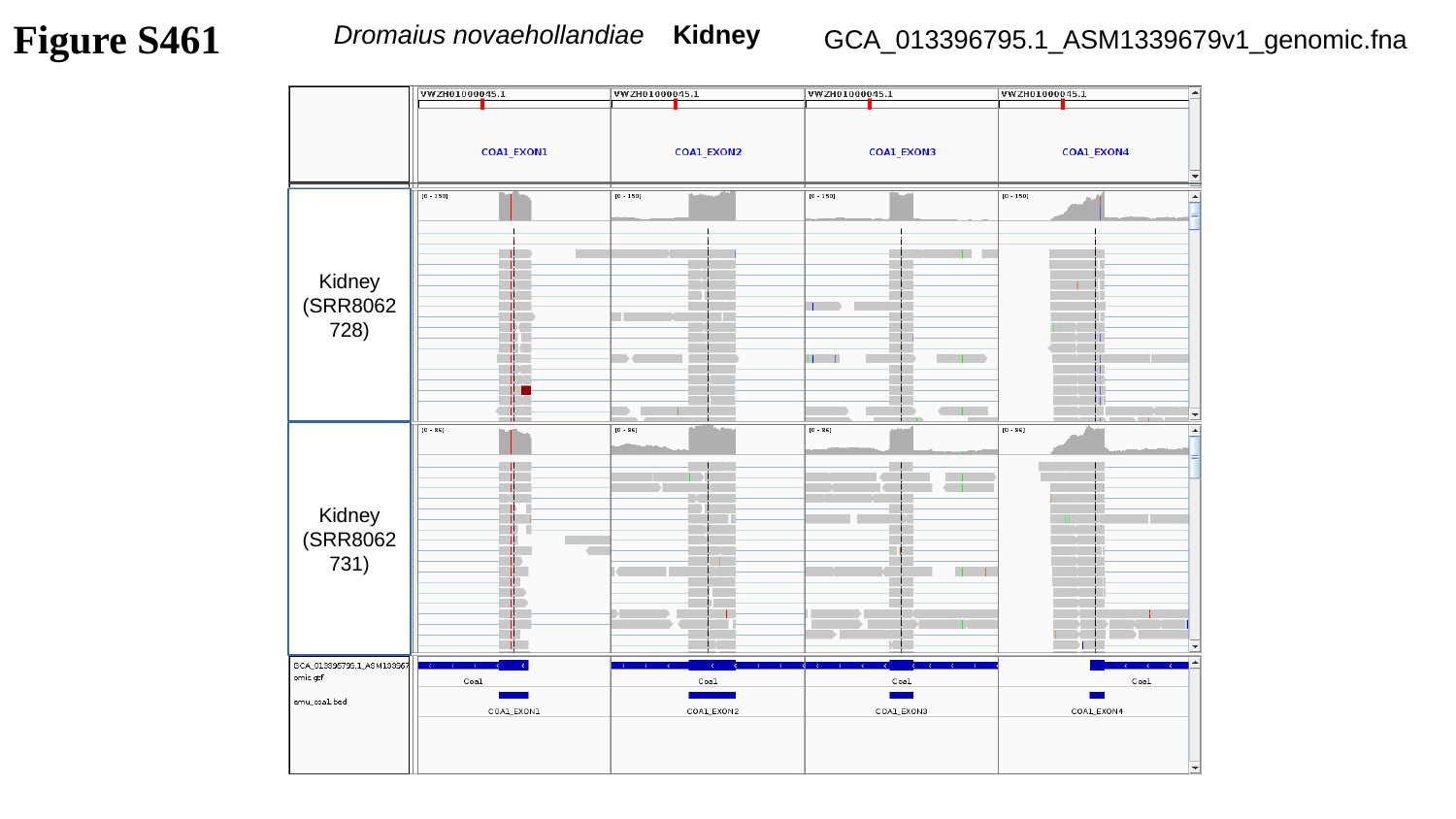

Figure S461
Dromaius novaehollandiae Kidney
GCA_013396795.1_ASM1339679v1_genomic.fna
Kidney
(SRR8062728)
Kidney
(SRR8062731)

### Slide 462
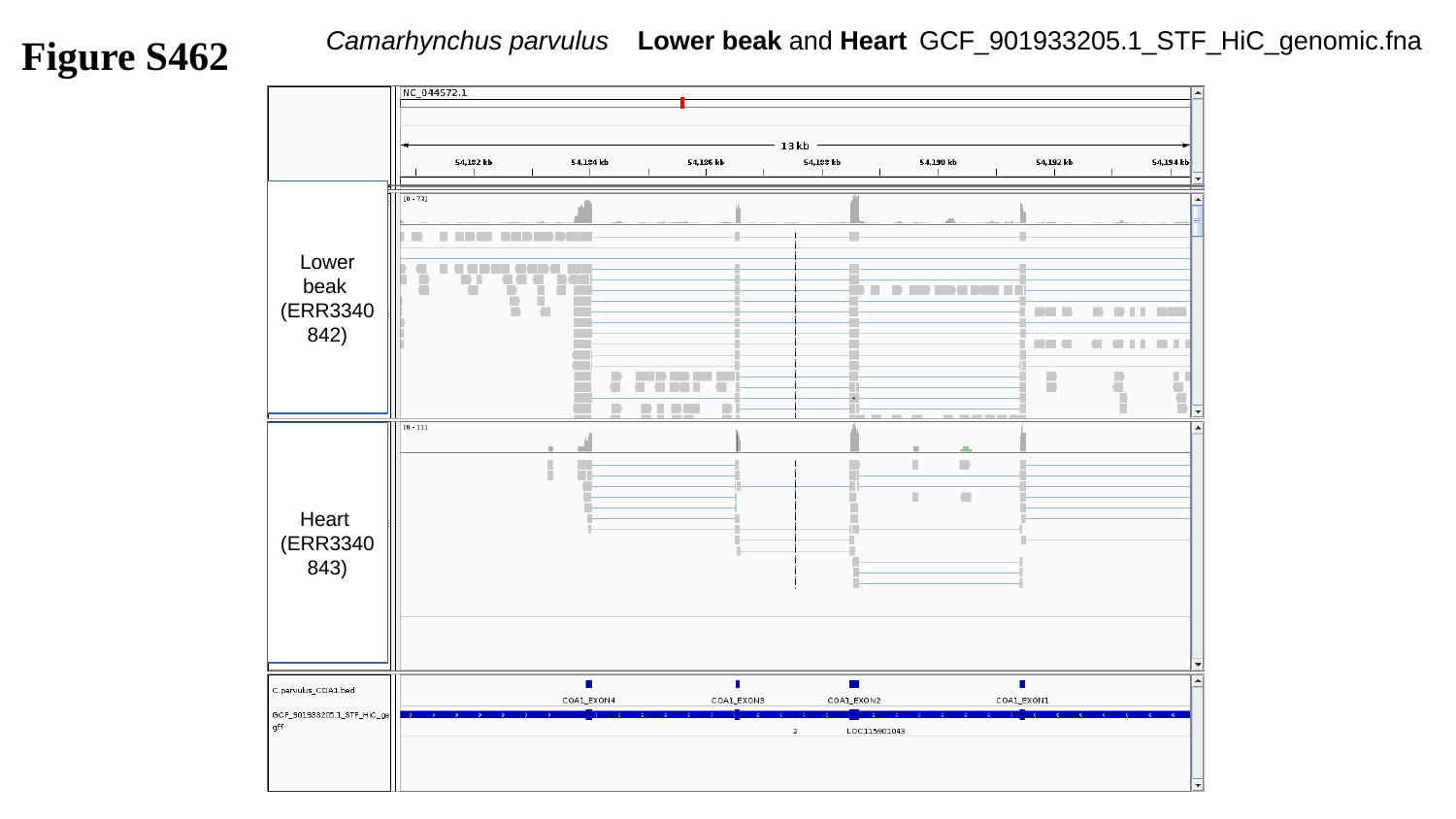

Figure S462
Camarhynchus parvulus Lower beak and Heart
GCF_901933205.1_STF_HiC_genomic.fna
Lower beak
(ERR3340842)
Heart
(ERR3340843)

### Slide 463
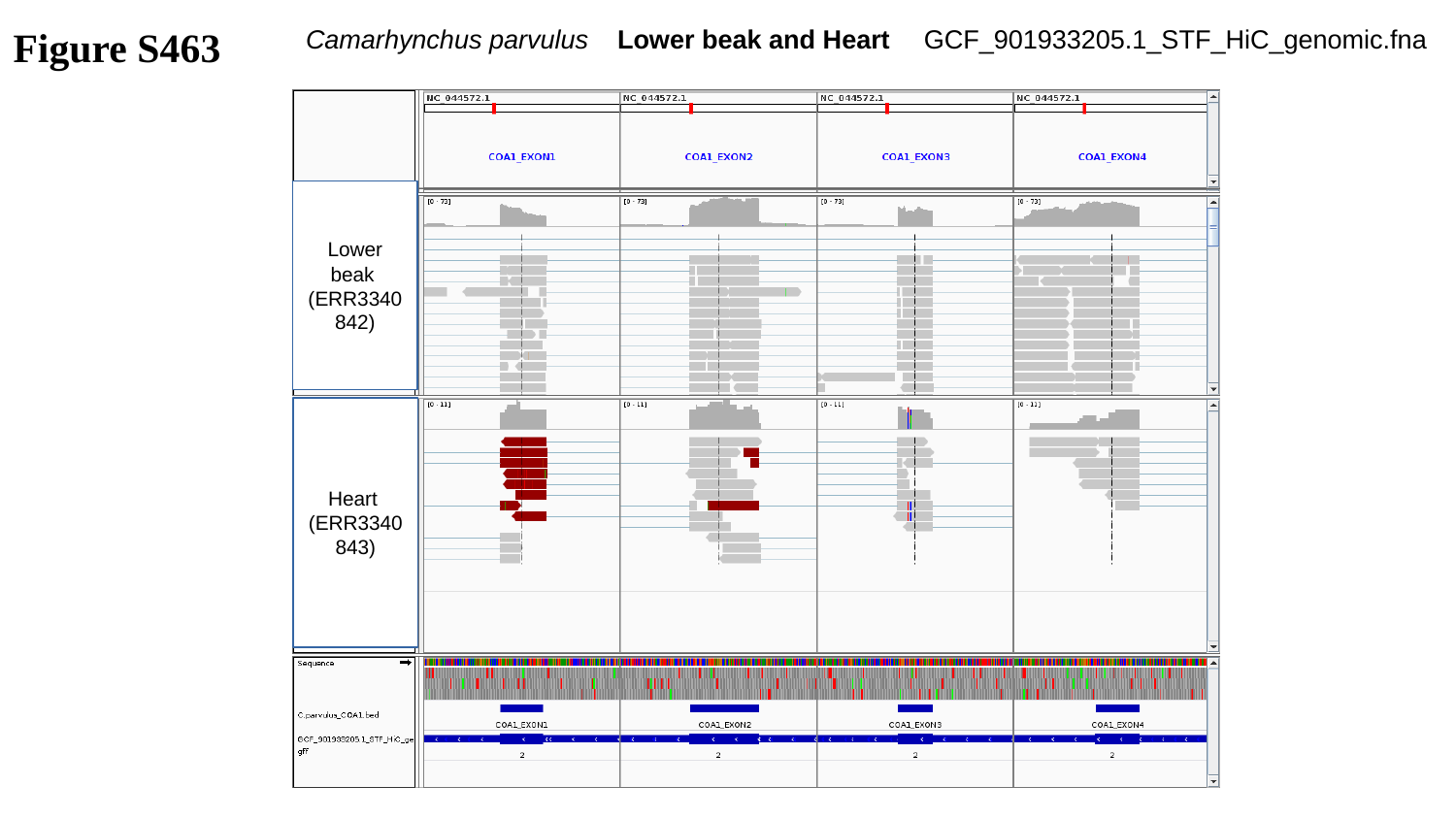

Figure S463
Camarhynchus parvulus Lower beak and Heart
GCF_901933205.1_STF_HiC_genomic.fna
Lower beak
(ERR3340842)
Heart
(ERR3340843)

### Slide 464
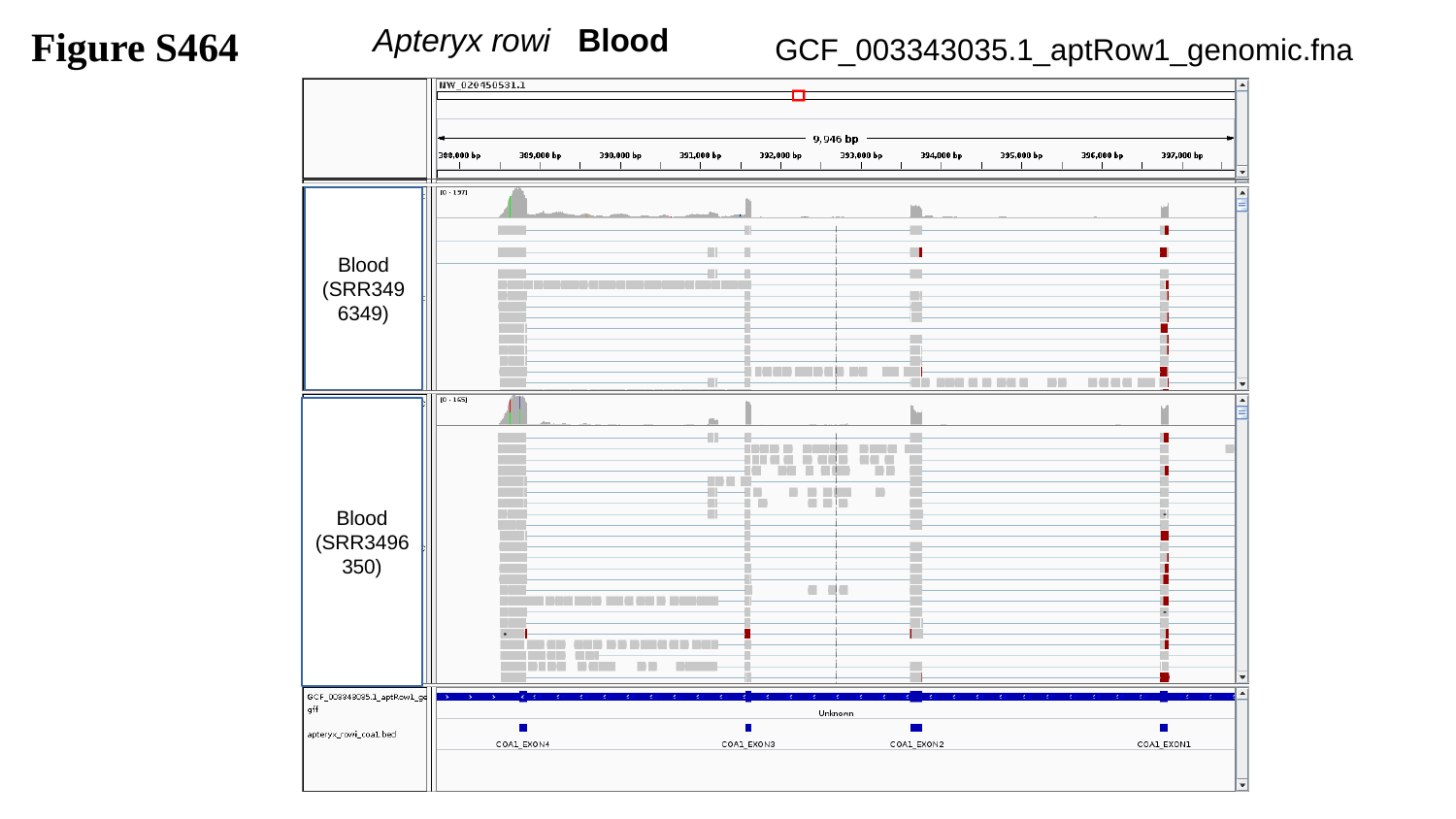

Figure S464
Apteryx rowi Blood
GCF_003343035.1_aptRow1_genomic.fna
Blood
(SRR3496349)
Blood
(SRR3496350)

### Slide 465
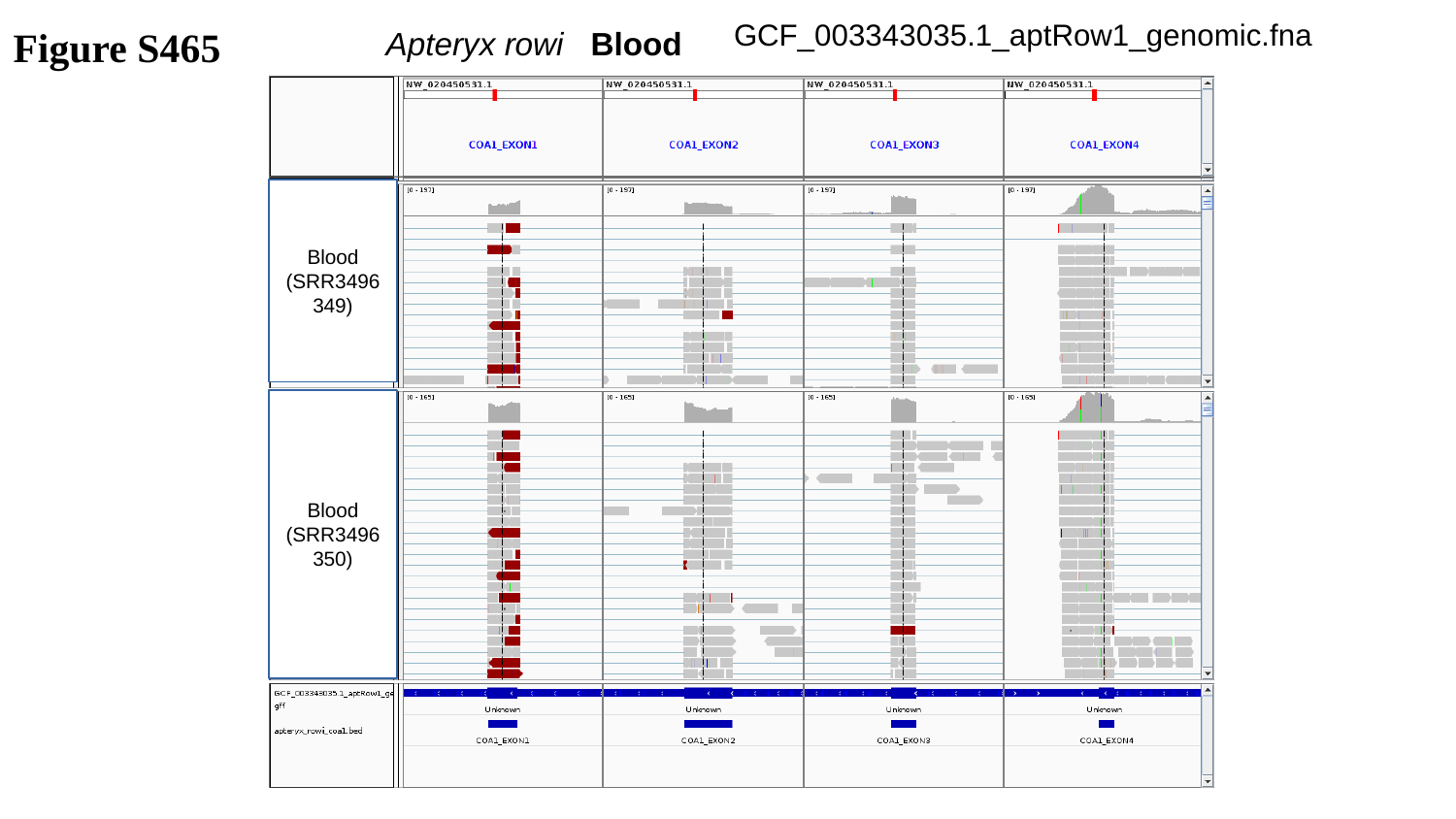

GCF_003343035.1_aptRow1_genomic.fna
Figure S465
Apteryx rowi Blood
Blood
(SRR3496349)
Blood
(SRR3496350)

### Slide 466
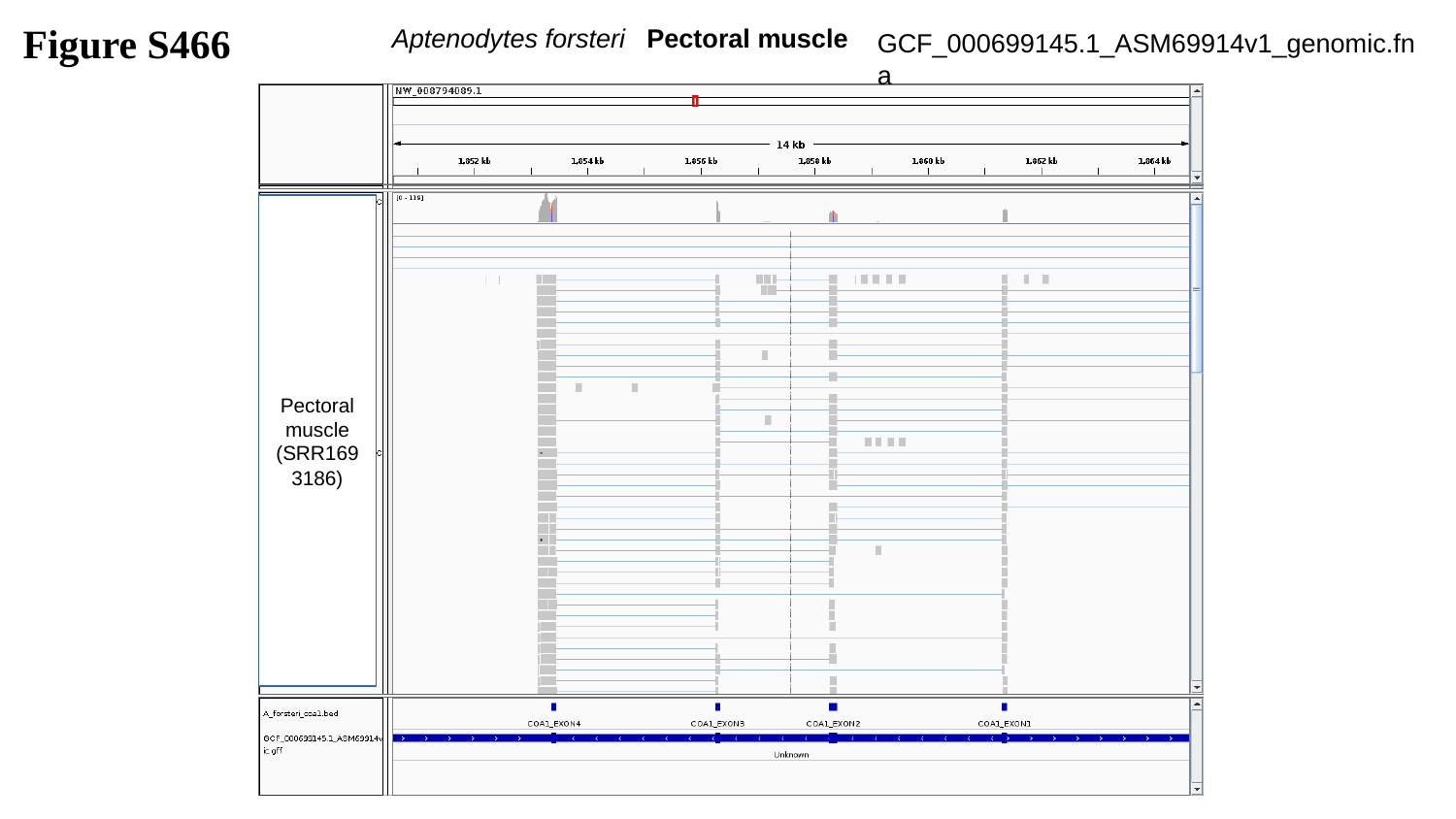

Figure S466
 Aptenodytes forsteri Pectoral muscle
GCF_000699145.1_ASM69914v1_genomic.fna
Pectoral muscle
(SRR1693186)

### Slide 467
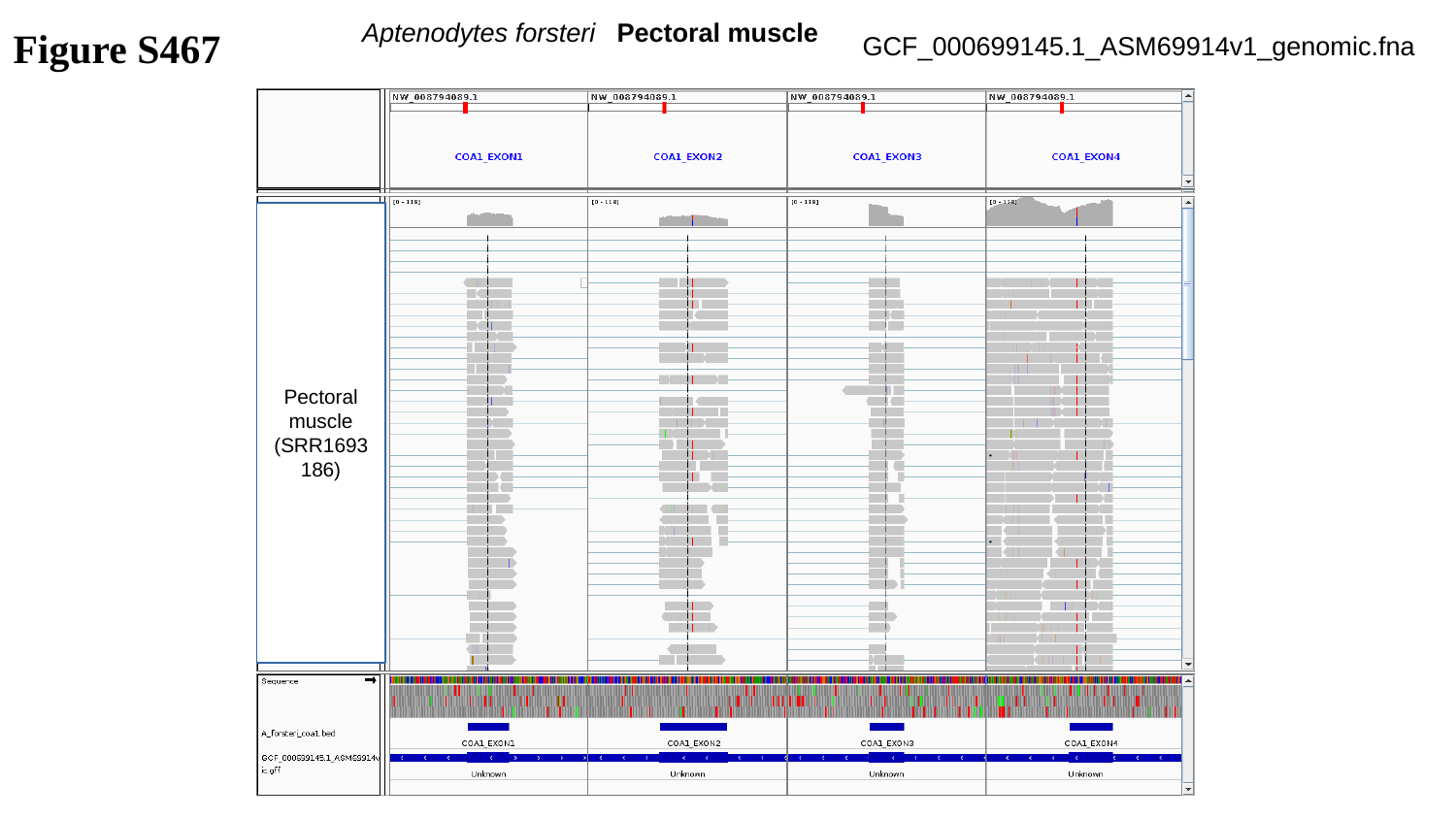

Aptenodytes forsteri Pectoral muscle
Figure S467
GCF_000699145.1_ASM69914v1_genomic.fna
Pectoral muscle
(SRR1693186)

### Slide 468
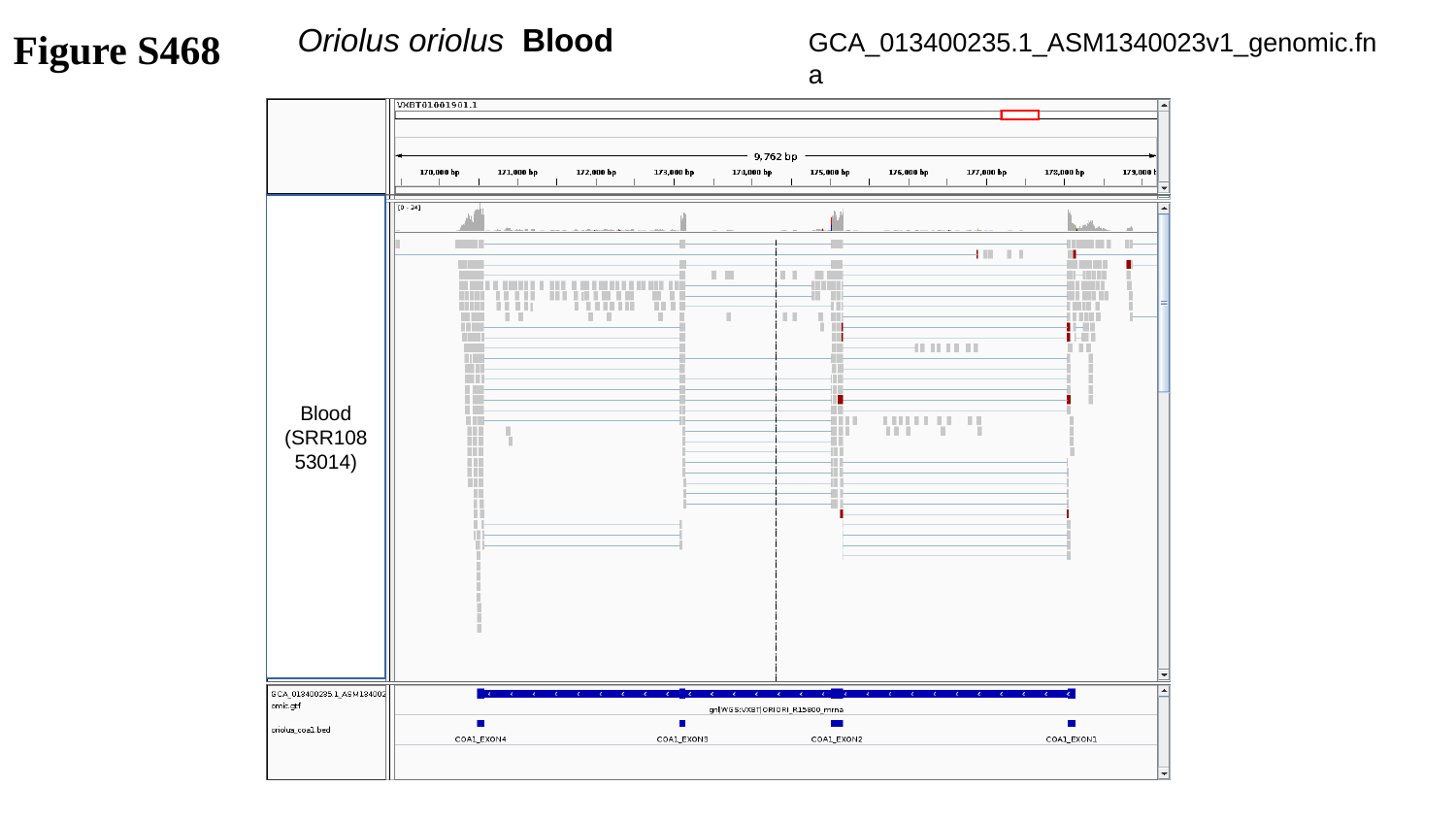

Figure S468
GCA_013400235.1_ASM1340023v1_genomic.fna
 Oriolus oriolus Blood
Blood
(SRR10853014)

### Slide 469
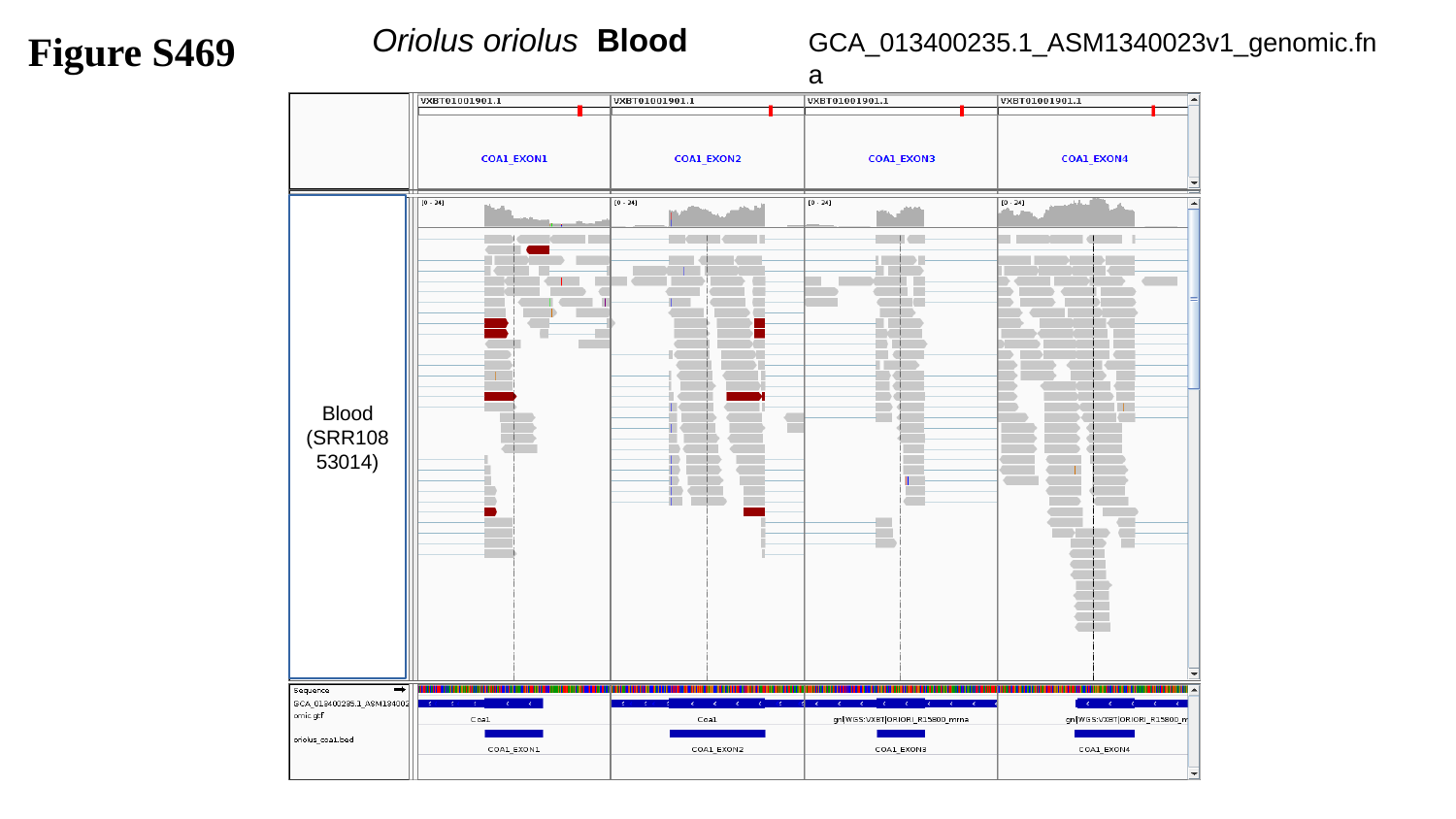

GCA_013400235.1_ASM1340023v1_genomic.fna
 Oriolus oriolus Blood
Figure S469
Blood
(SRR10853014)

### Slide 470
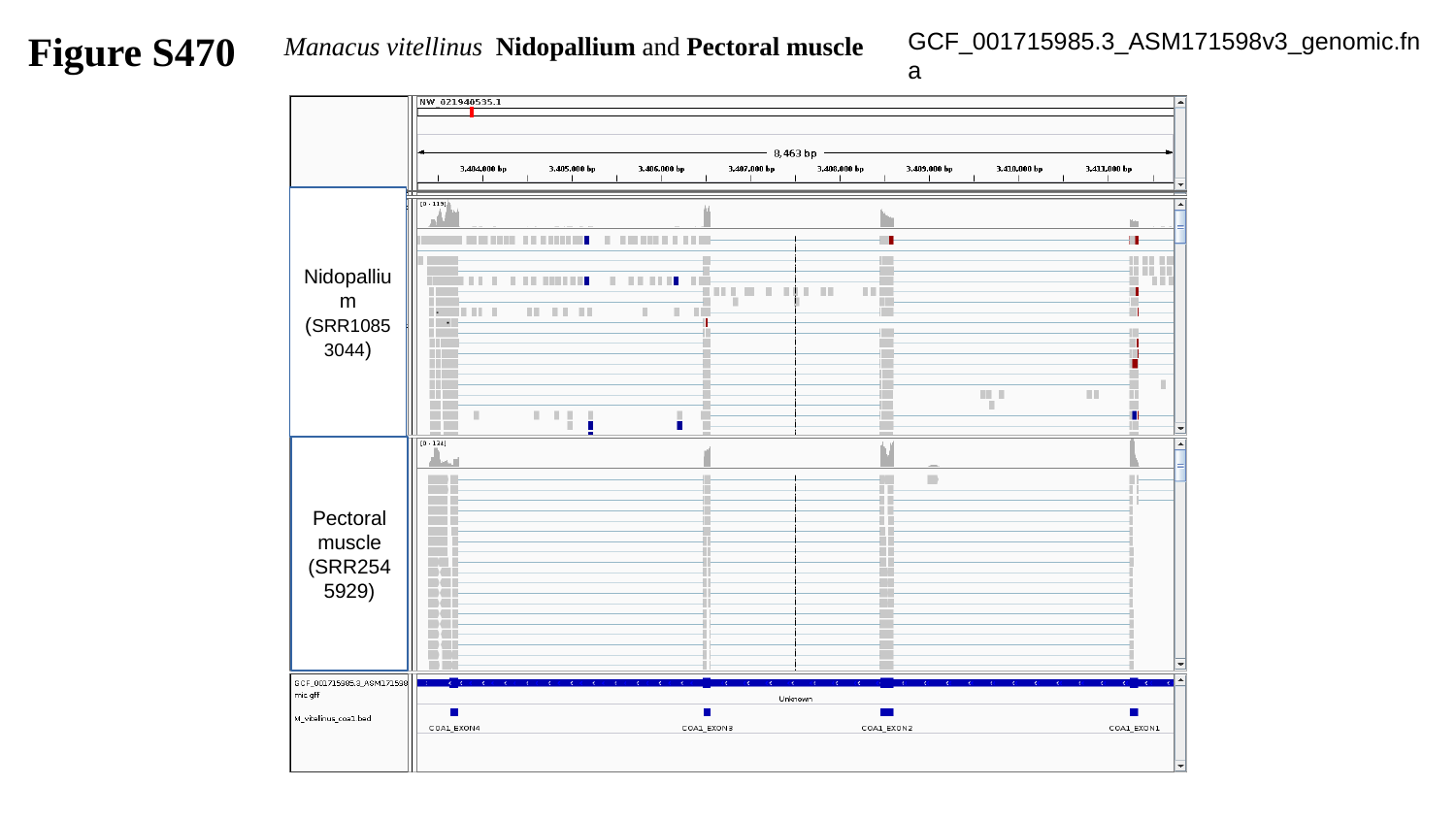

Figure S470
GCF_001715985.3_ASM171598v3_genomic.fna
 Manacus vitellinus Nidopallium and Pectoral muscle
Nidopallium
(SRR10853044)
Pectoral muscle
(SRR2545929)

### Slide 471
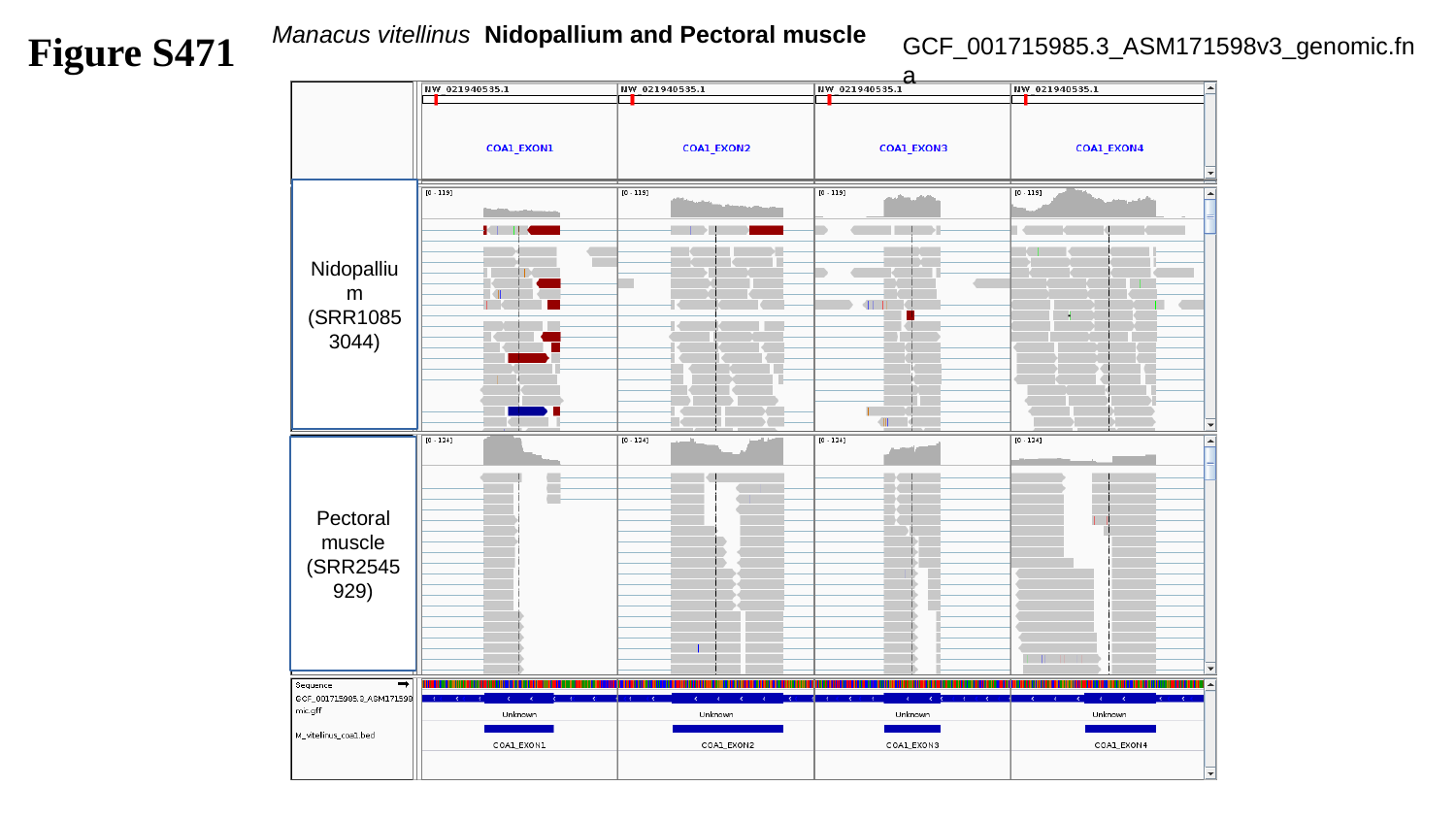

Figure S471
 Manacus vitellinus Nidopallium and Pectoral muscle
GCF_001715985.3_ASM171598v3_genomic.fna
Nidopallium
(SRR10853044)
Pectoral muscle
(SRR2545929)

### Slide 472
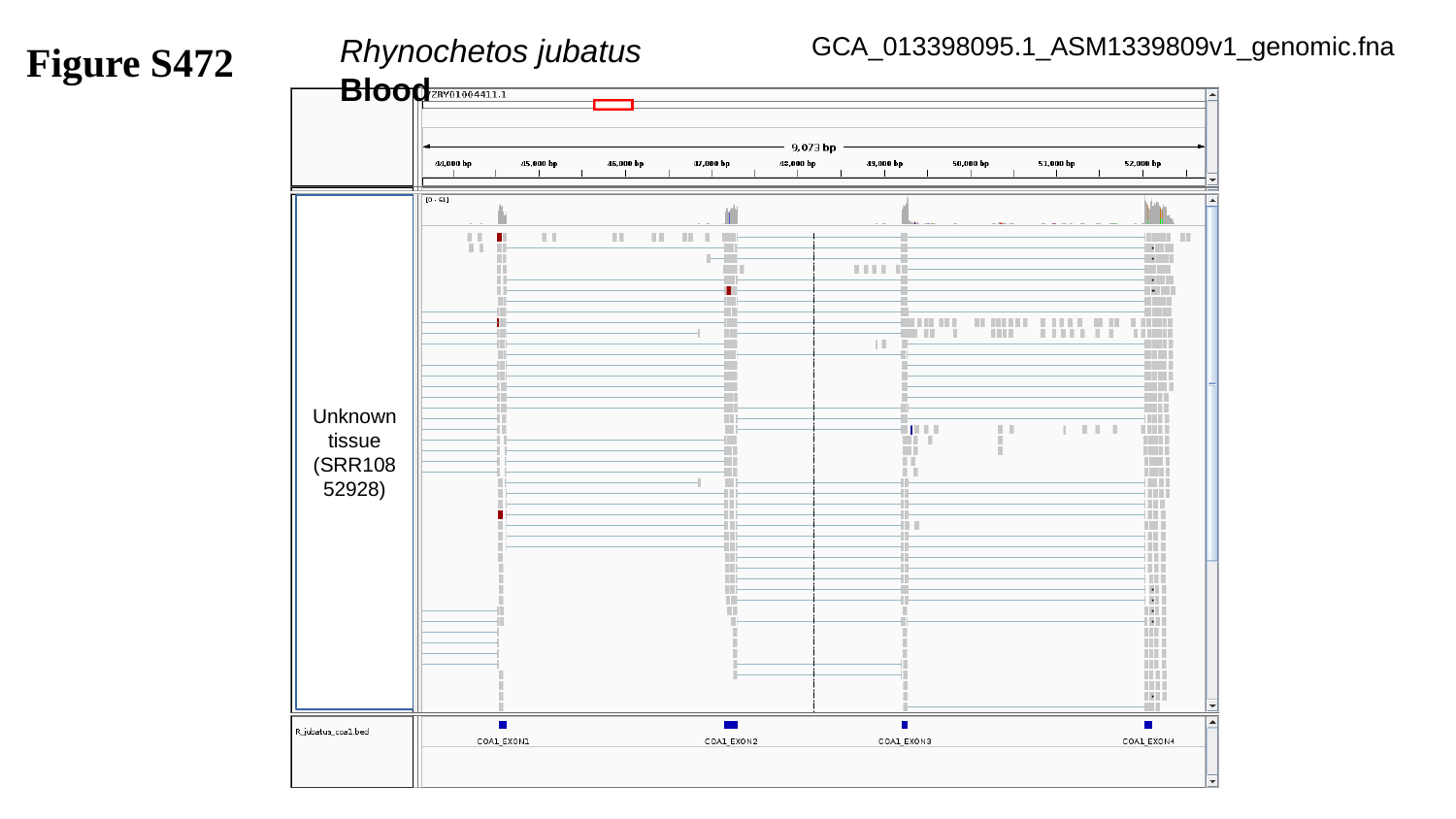

GCA_013398095.1_ASM1339809v1_genomic.fna
Figure S472
Rhynochetos jubatus	Blood
Unknown tissue
(SRR10852928)

### Slide 473
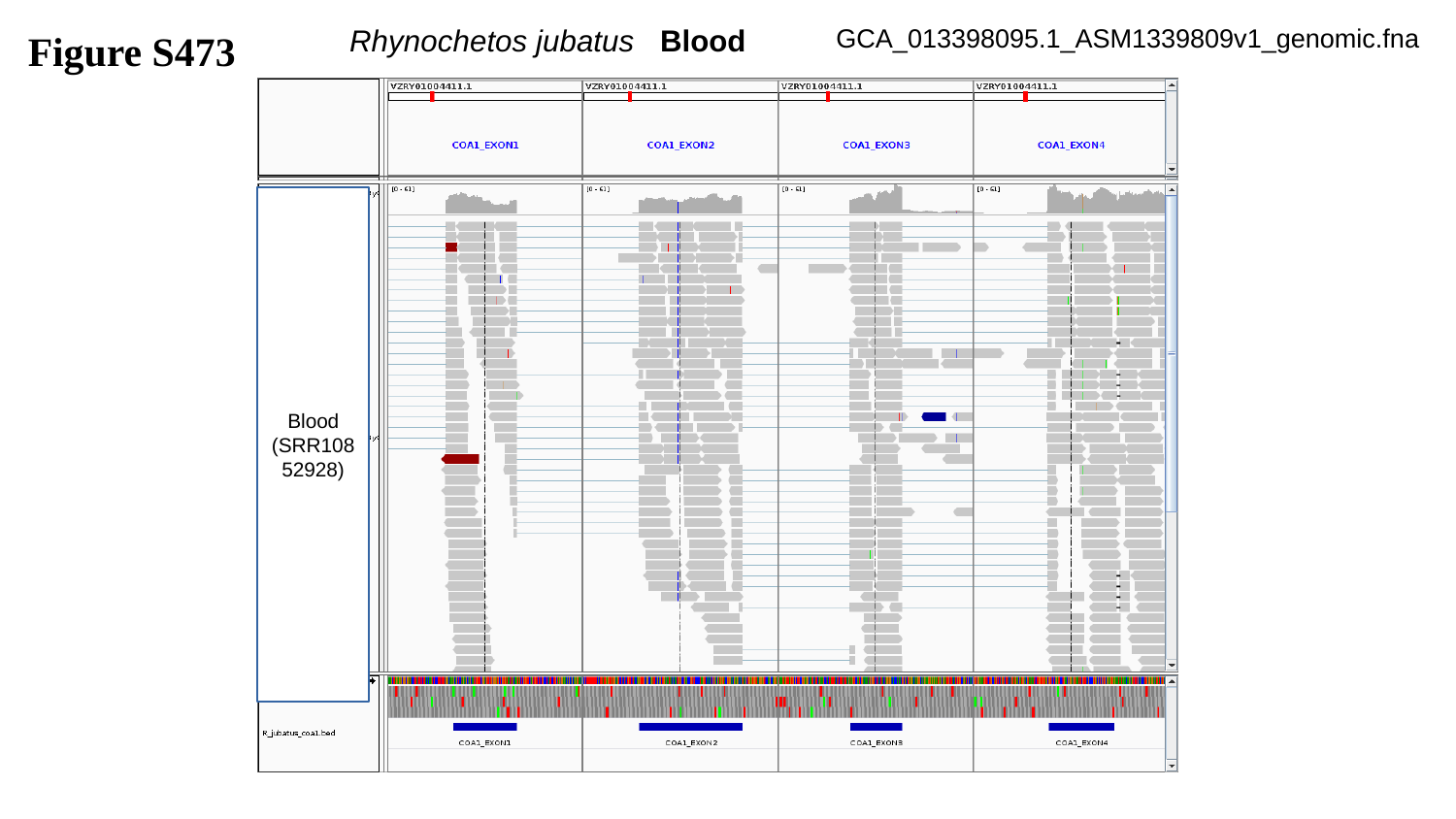

Figure S473
Rhynochetos jubatus Blood
GCA_013398095.1_ASM1339809v1_genomic.fna
Blood
(SRR10852928)

### Slide 474
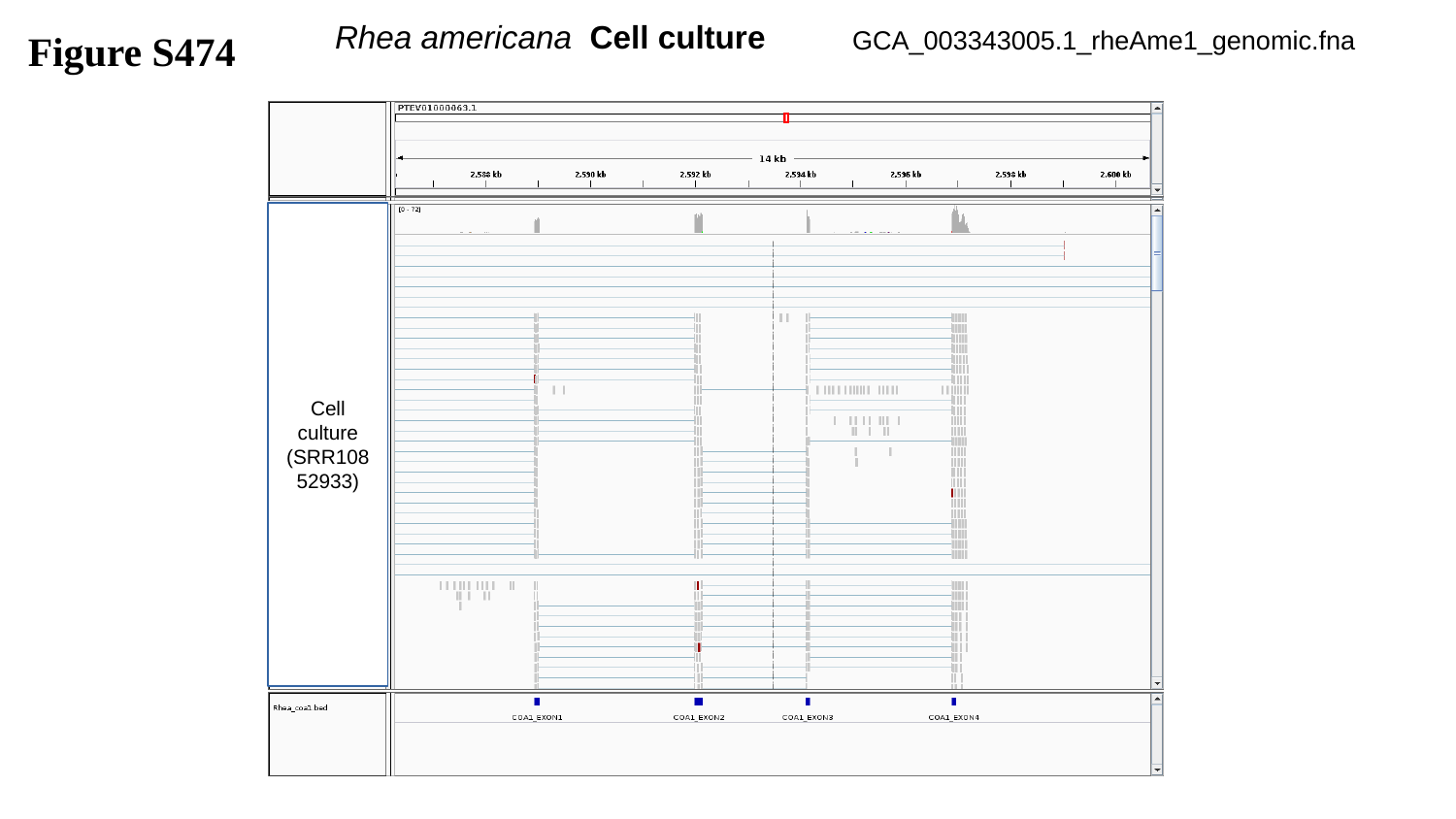

Rhea americana Cell culture
GCA_003343005.1_rheAme1_genomic.fna
Figure S474
Cell culture
(SRR10852933)

### Slide 475
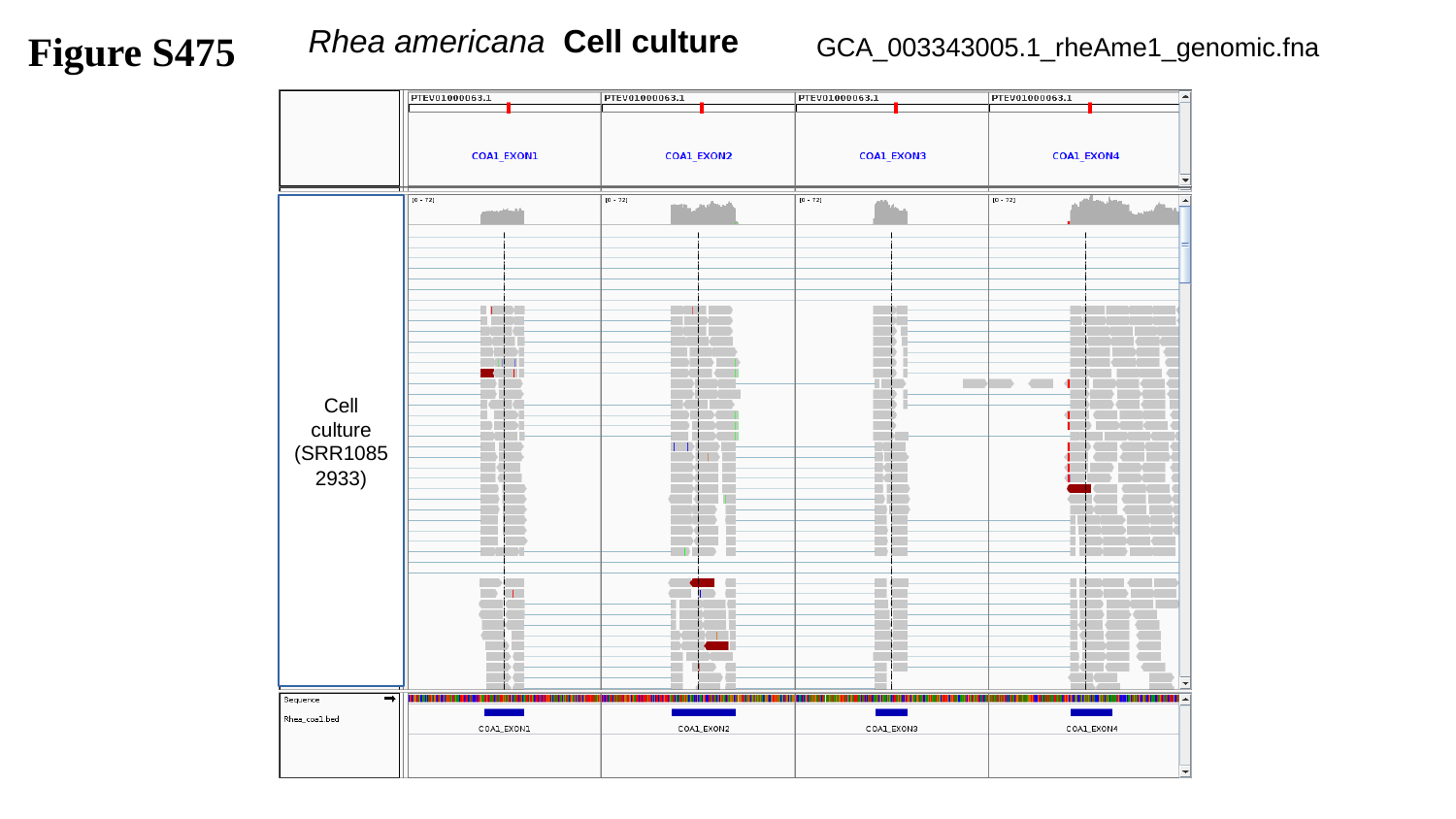

Figure S475
 Rhea americana Cell culture
GCA_003343005.1_rheAme1_genomic.fna
Cell culture
(SRR10852933)

### Slide 476
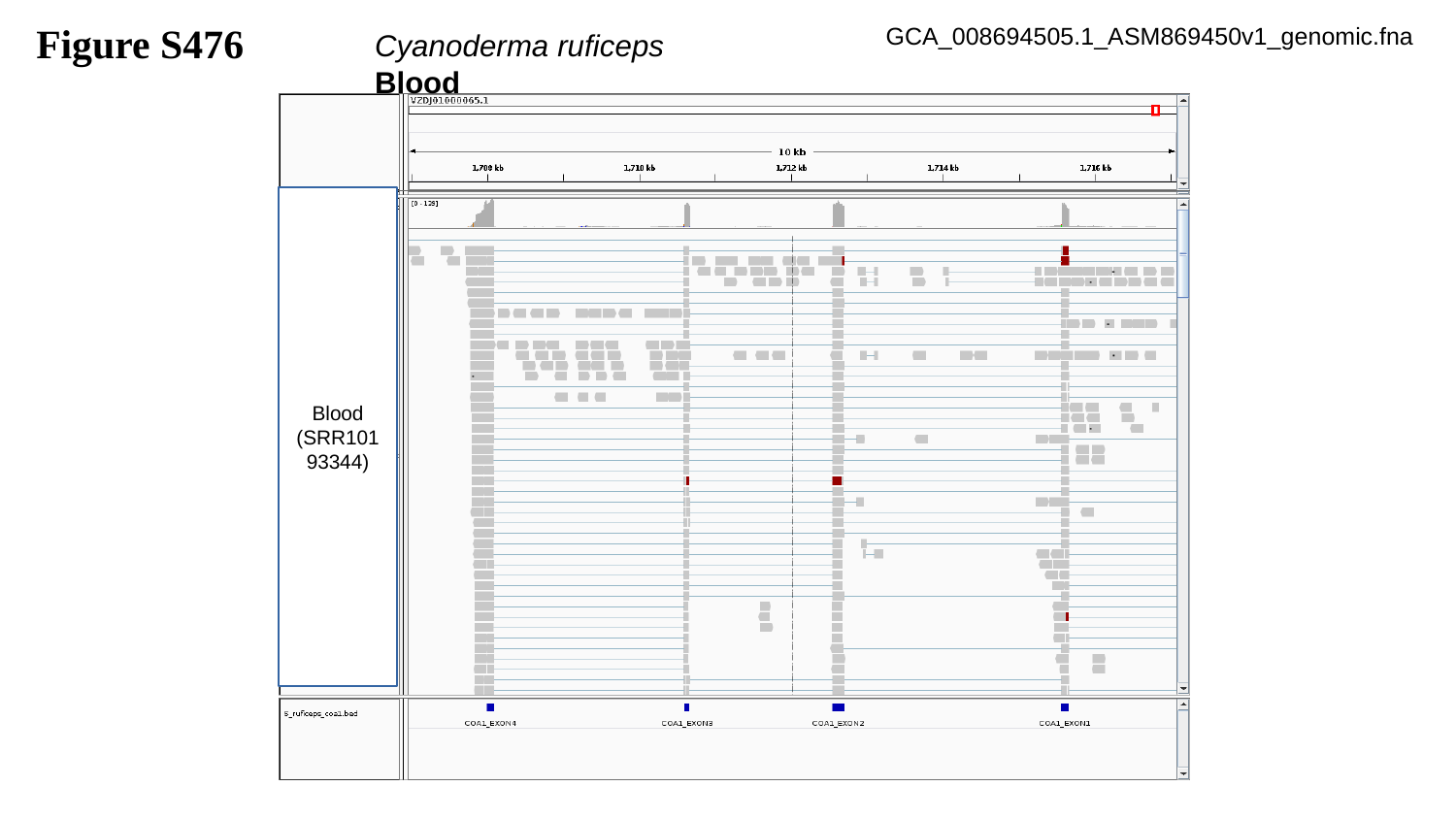

Figure S476
GCA_008694505.1_ASM869450v1_genomic.fna
Cyanoderma ruficeps		Blood
Blood
(SRR10193344)

### Slide 477
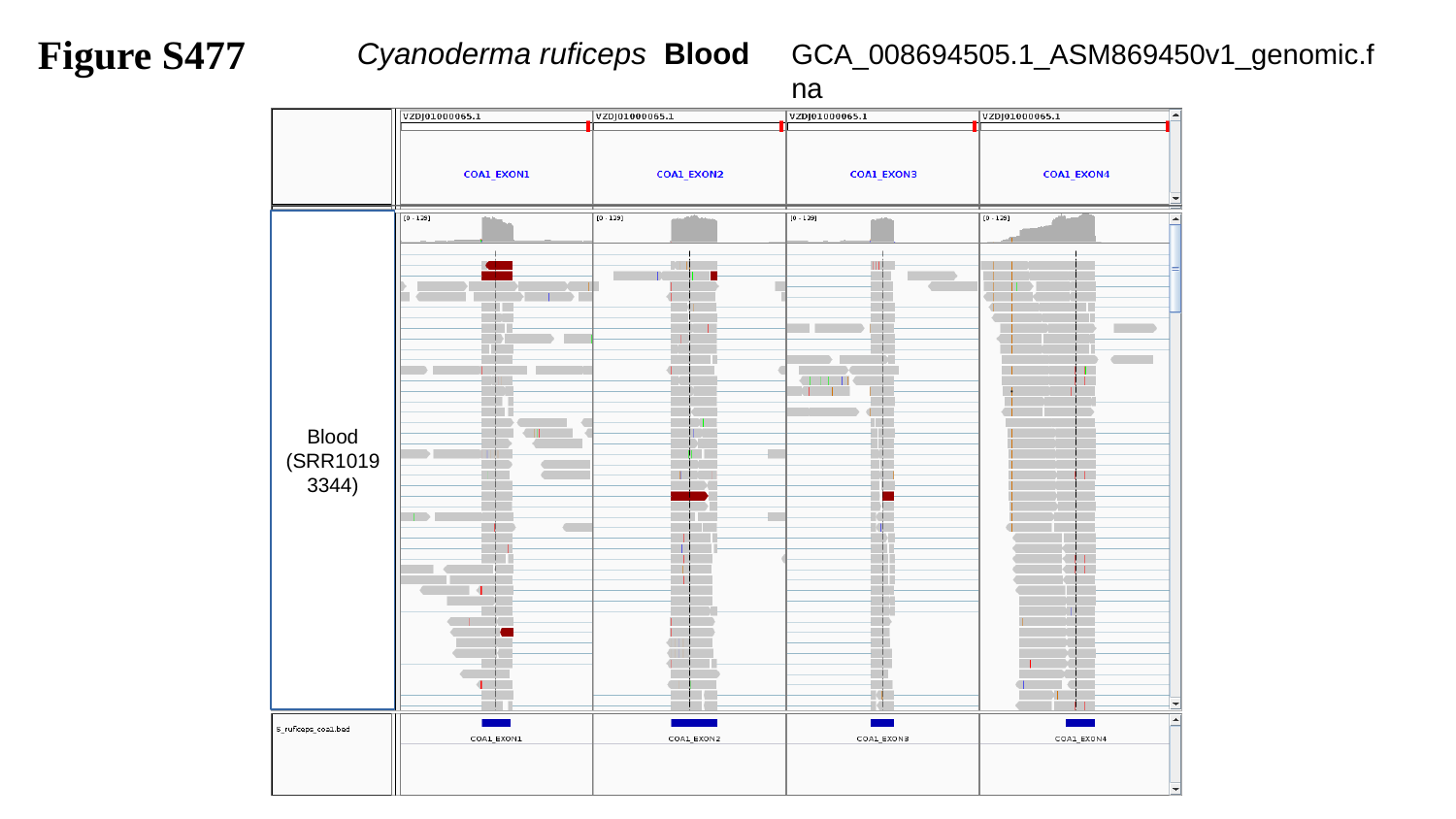

Figure S477
Cyanoderma ruficeps Blood
GCA_008694505.1_ASM869450v1_genomic.fna
Blood
(SRR10193344)

### Slide 478
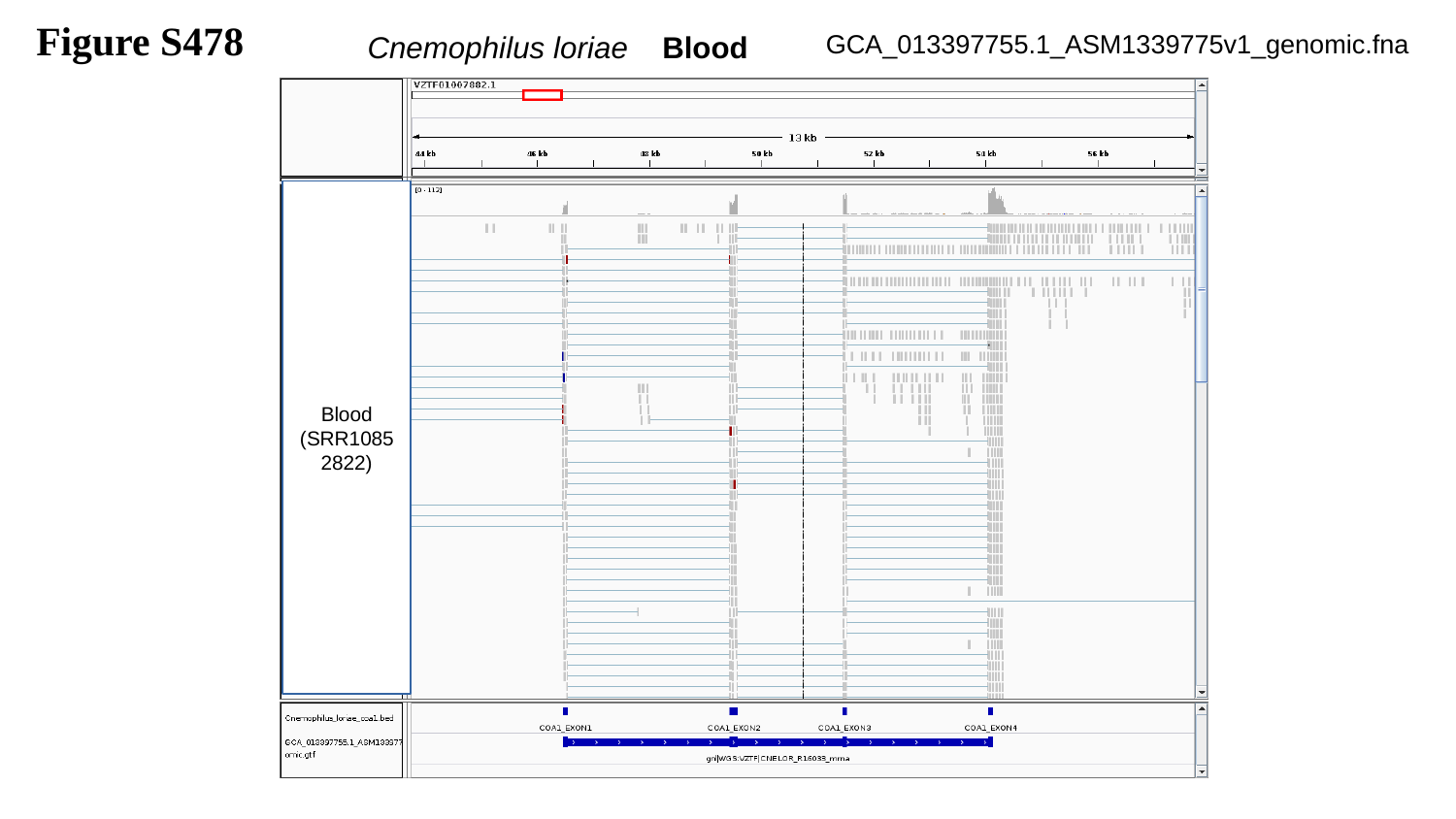

Figure S478
Cnemophilus loriae Blood
GCA_013397755.1_ASM1339775v1_genomic.fna
Blood
(SRR10852822)

### Slide 479
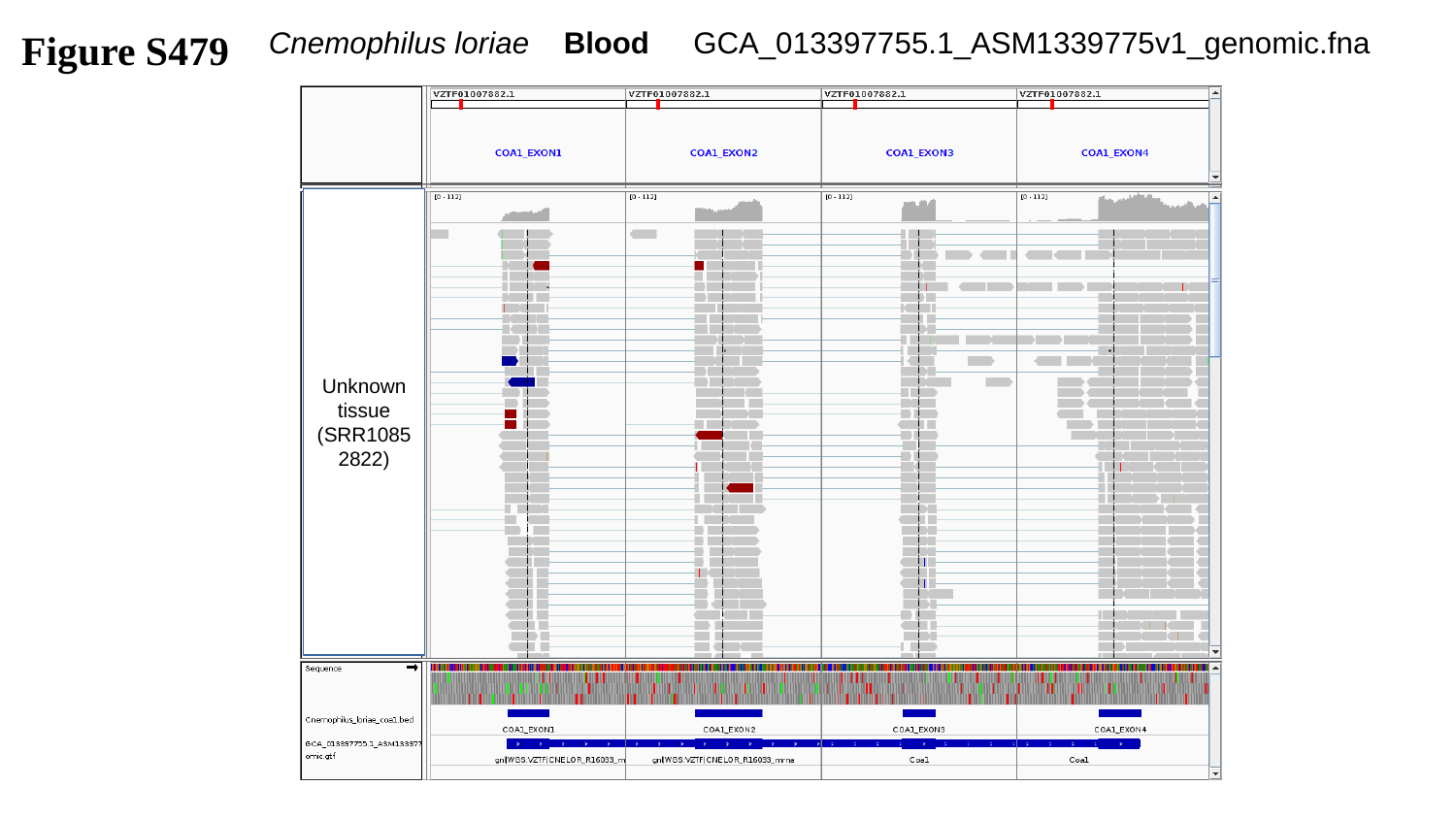

Figure S479
Cnemophilus loriae Blood
GCA_013397755.1_ASM1339775v1_genomic.fna
Unknown tissue
(SRR10852822)

### Slide 480
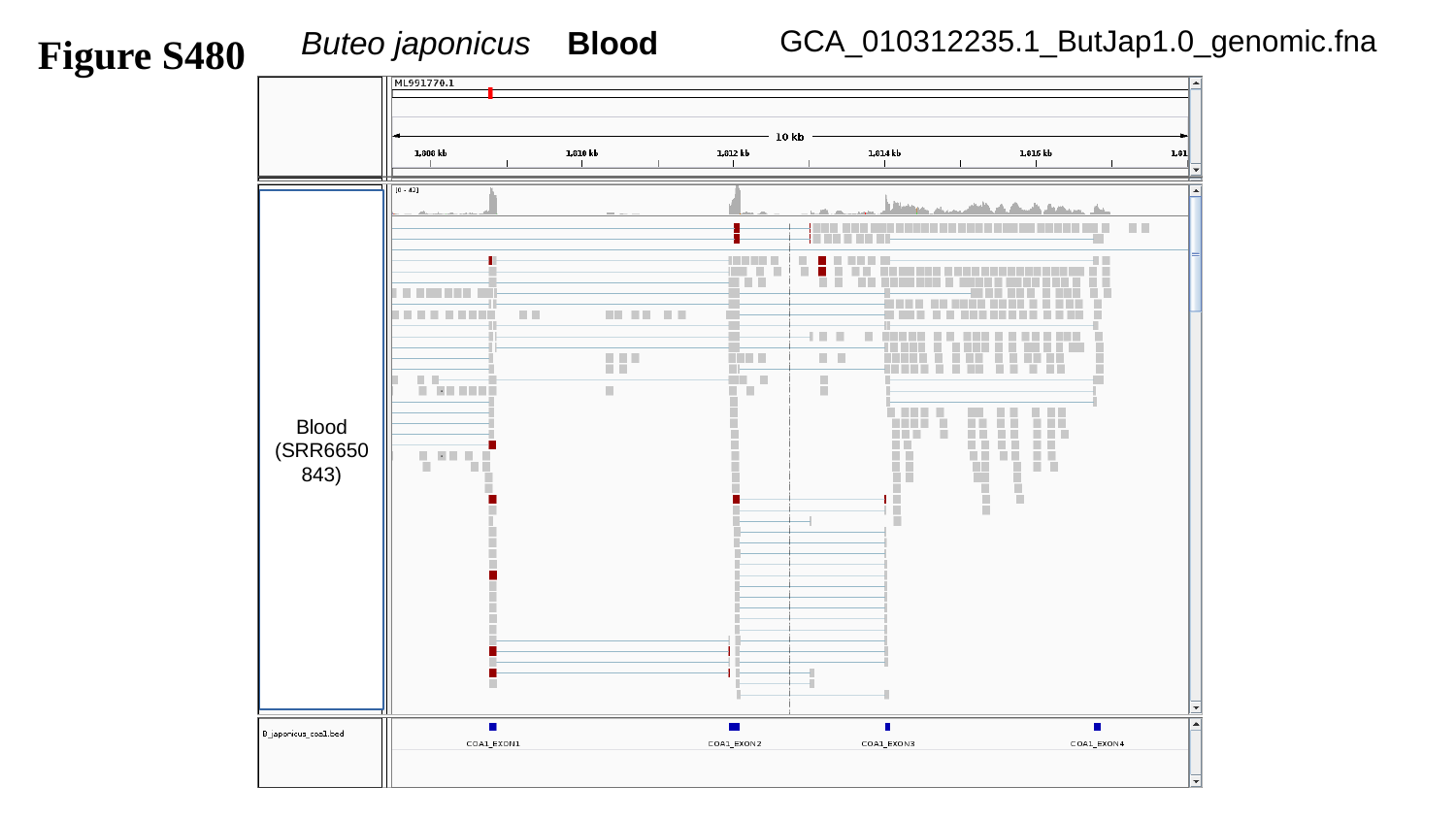

Buteo japonicus Blood
Figure S480
GCA_010312235.1_ButJap1.0_genomic.fna
Blood
(SRR6650843)

### Slide 481
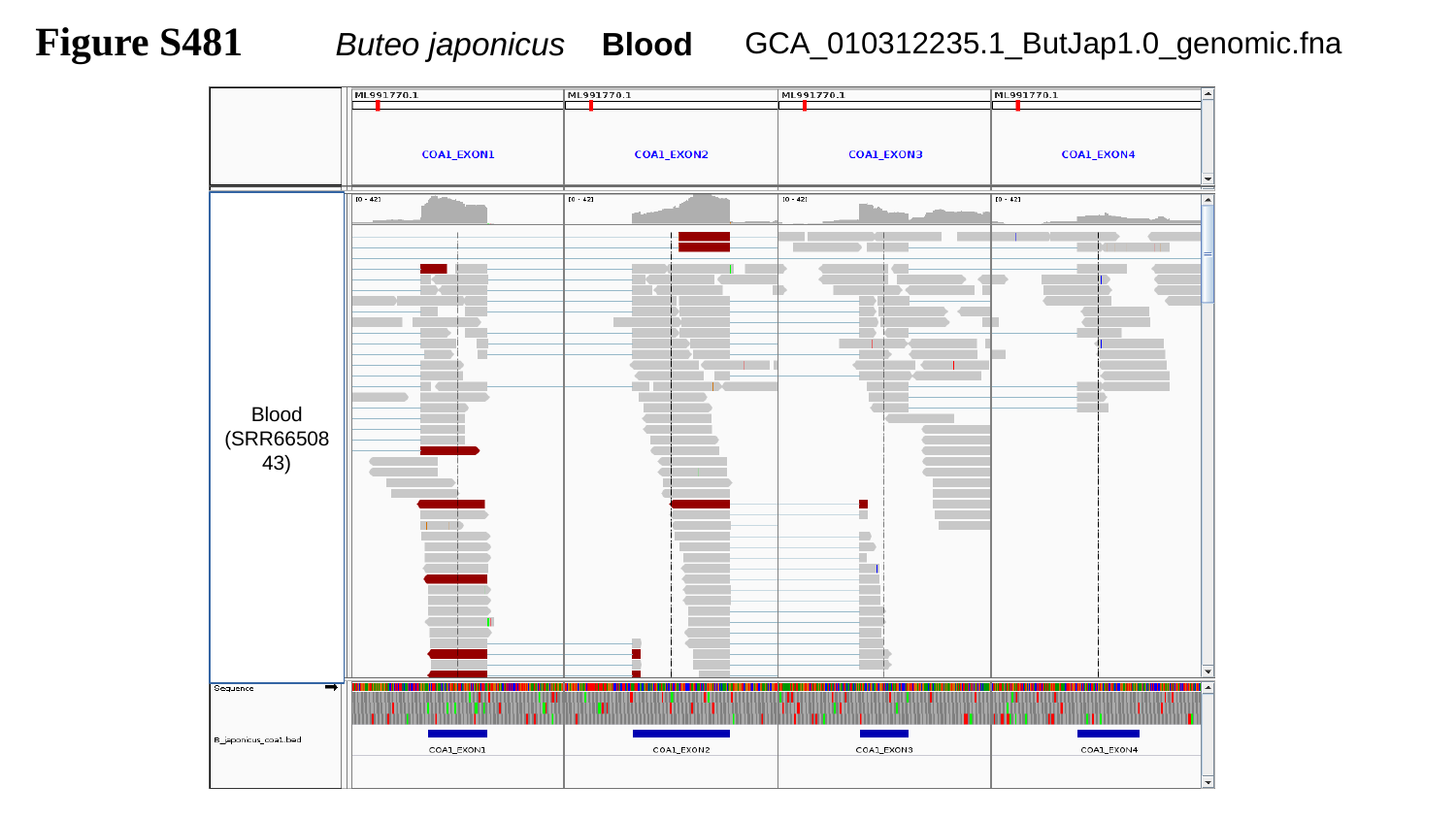

Figure S481
Buteo japonicus Blood
GCA_010312235.1_ButJap1.0_genomic.fna
Blood
(SRR6650843)

### Slide 482
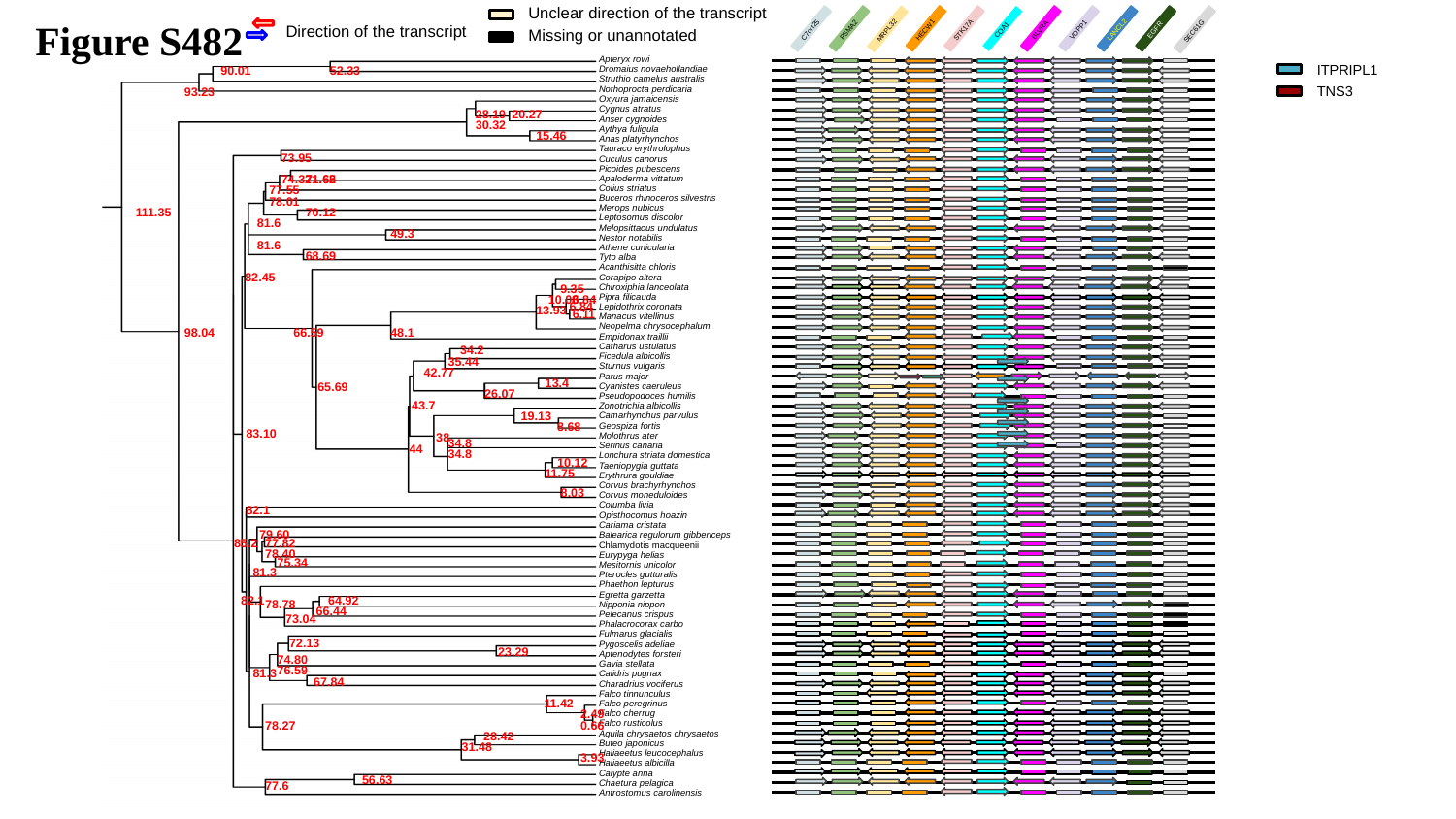

Unclear direction of the transcript
Figure S482
Direction of the transcript
Missing or unannotated
LANCL2
EGFR
PSMA2
HECW1
STK17A
BLVRA
VOPP1
C7orf25
COA1
MRPL32
SEC61G
ITPRIPL1
Apteryx rowi
90.01
52.33
Dromaius novaehollandiae
Struthio camelus australis
TNS3
93.23
Nothoprocta perdicaria
Oxyura jamaicensis
Cygnus atratus
28.19
20.27
Anser cygnoides
30.32
Aythya fuligula
15.46
Anas platyrhynchos
Tauraco erythrolophus
73.95
Cuculus canorus
Picoides pubescens
74.32
71.62
71.65
Apaloderma vittatum
77.55
Colius striatus
78.01
Buceros rhinoceros silvestris
111.35
70.12
Merops nubicus
Leptosomus discolor
81.6
Melopsittacus undulatus
49.3
Nestor notabilis
81.6
Athene cunicularia
68.69
Tyto alba
Acanthisitta chloris
82.45
Corapipo altera
9.35
Chiroxiphia lanceolata
10.08
6.84
Pipra filicauda
6.84
13.93
Lepidothrix coronata
6.11
Manacus vitellinus
Neopelma chrysocephalum
98.04
66.59
48.1
Empidonax traillii
34.2
Catharus ustulatus
Ficedula albicollis
35.44
Sturnus vulgaris
42.77
Parus major
13.4
65.69
Cyanistes caeruleus
26.07
Pseudopodoces humilis
43.7
Zonotrichia albicollis
19.13
Camarhynchus parvulus
8.68
Geospiza fortis
83.10
38
Molothrus ater
34.8
44
Serinus canaria
34.8
Lonchura striata domestica
10.12
Taeniopygia guttata
11.75
Erythrura gouldiae
Corvus brachyrhynchos
8.03
Corvus moneduloides
Columba livia
82.1
Opisthocomus hoazin
Cariama cristata
79.60
Balearica regulorum gibbericeps
85.2
77.82
Chlamydotis macqueenii
78.40
Eurypyga helias
75.34
Mesitornis unicolor
81.3
Pterocles gutturalis
Phaethon lepturus
Egretta garzetta
82.1
64.92
78.78
Nipponia nippon
66.44
73.04
Pelecanus crispus
Phalacrocorax carbo
Fulmarus glacialis
72.13
Pygoscelis adeliae
23.29
Aptenodytes forsteri
74.80
Gavia stellata
76.59
81.3
Calidris pugnax
67.84
Charadrius vociferus
Falco tinnunculus
11.42
Falco peregrinus
2.49
Falco cherrug
78.27
0.66
Falco rusticolus
28.42
Aquila chrysaetos chrysaetos
31.48
Buteo japonicus
Haliaeetus leucocephalus
3.93
Haliaeetus albicilla
Calypte anna
56.63
77.6
Chaetura pelagica
Antrostomus carolinensis

### Slide 483
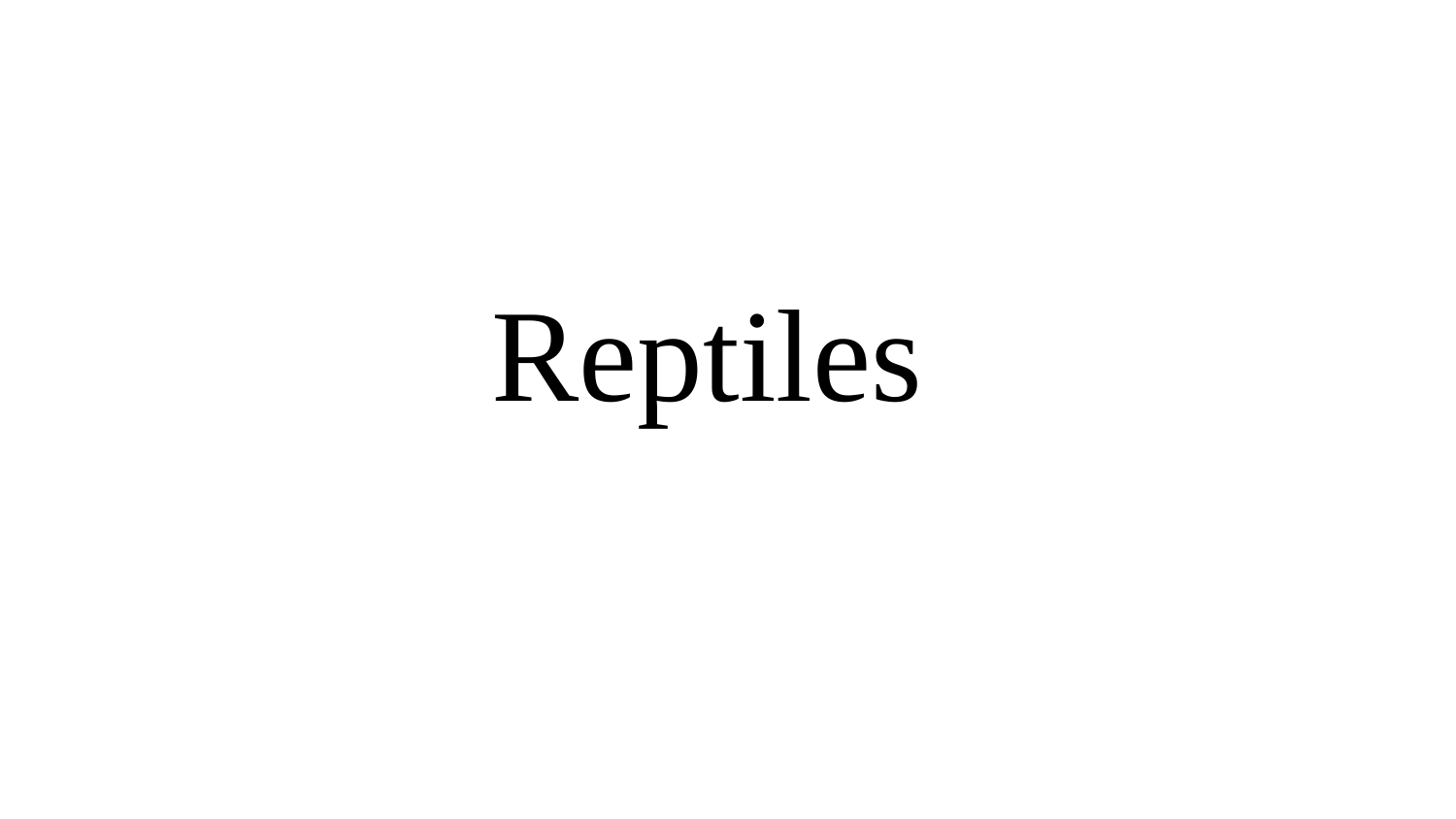

Reptiles

### Slide 484
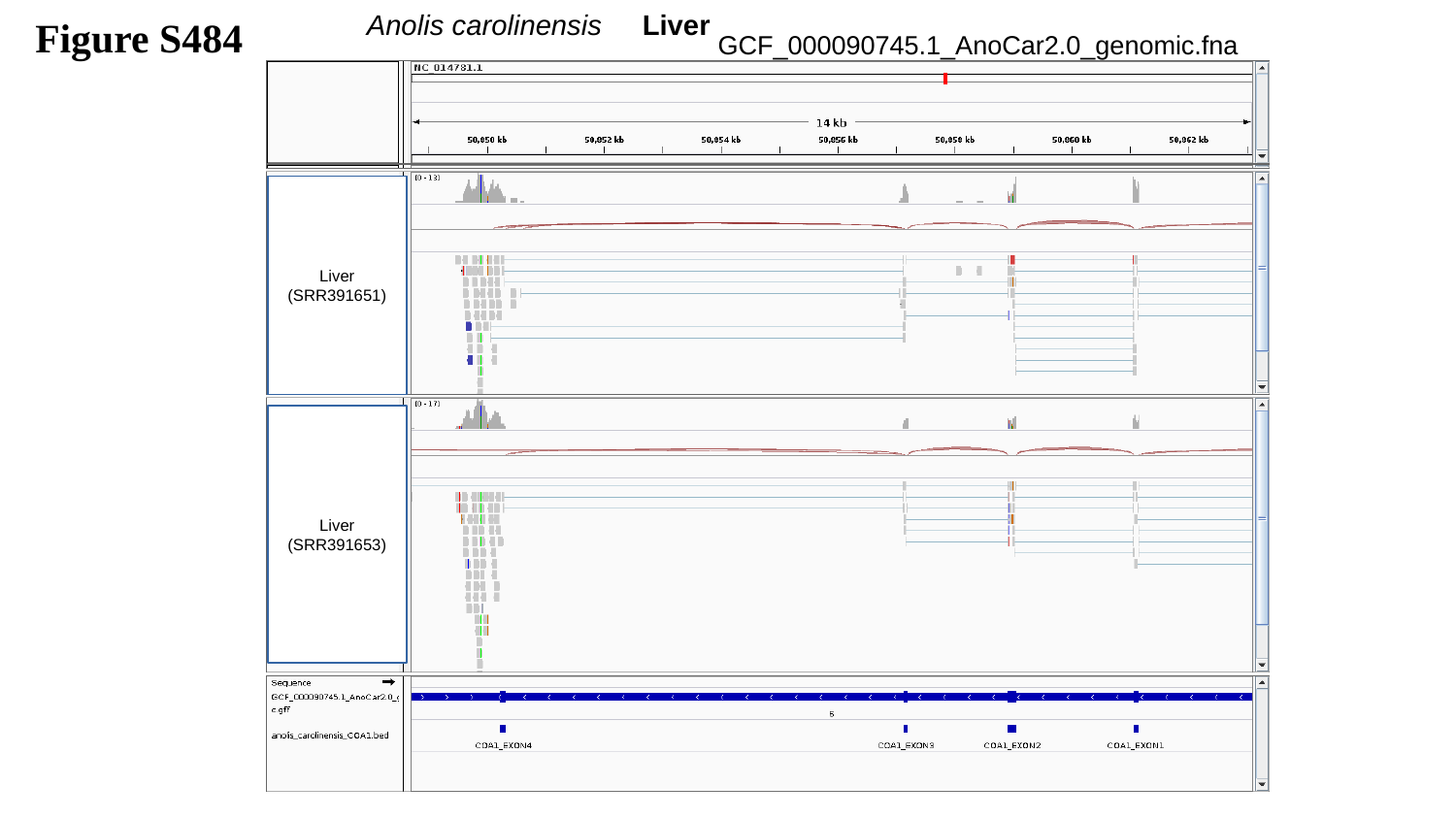

Figure S484
Anolis carolinensis Liver
GCF_000090745.1_AnoCar2.0_genomic.fna
Liver
(SRR391651)
Liver
(SRR391653)

### Slide 485
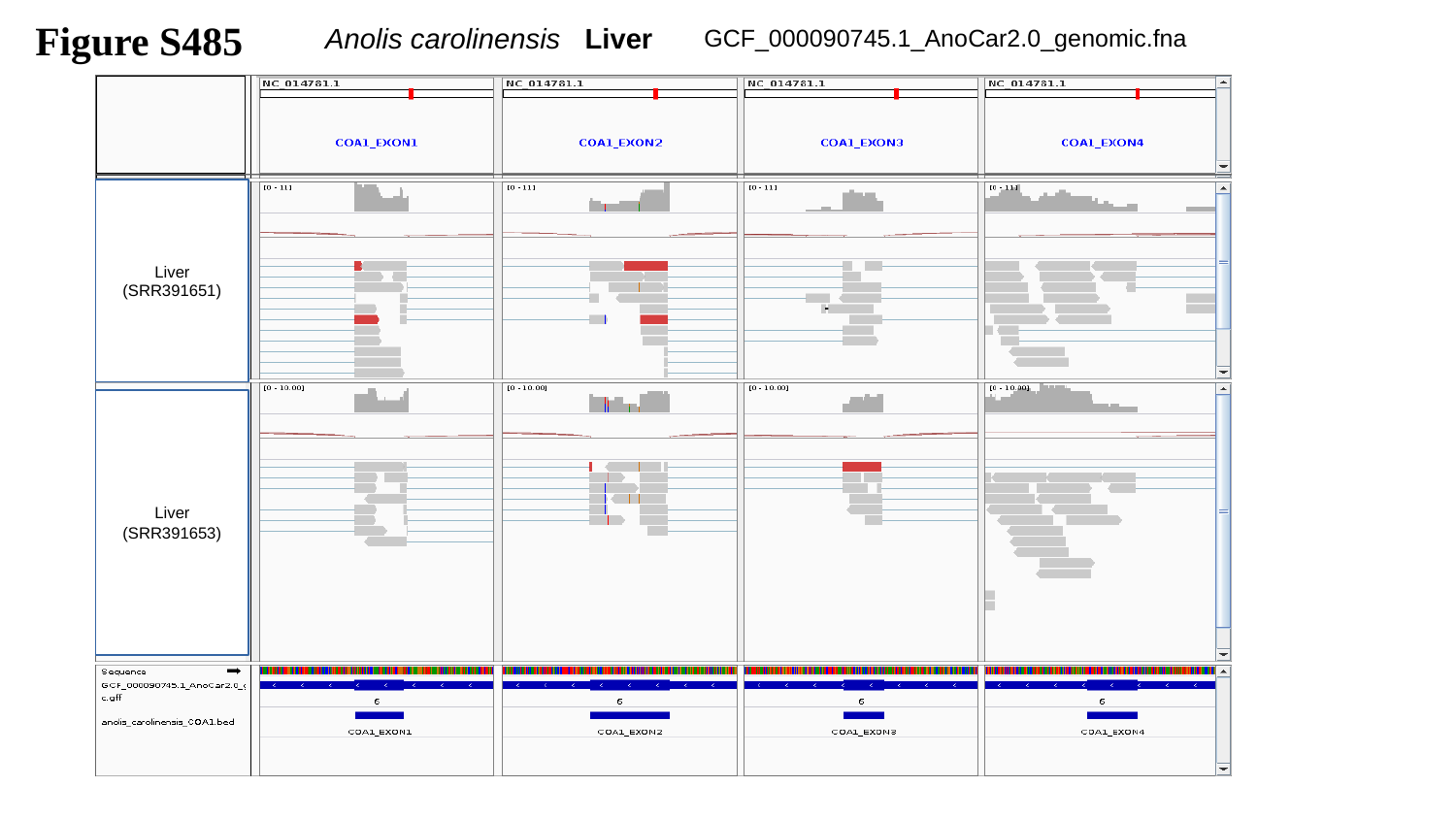

Figure S485
 Anolis carolinensis Liver
GCF_000090745.1_AnoCar2.0_genomic.fna
Liver
(SRR391651)
Liver
(SRR391653)

### Slide 486
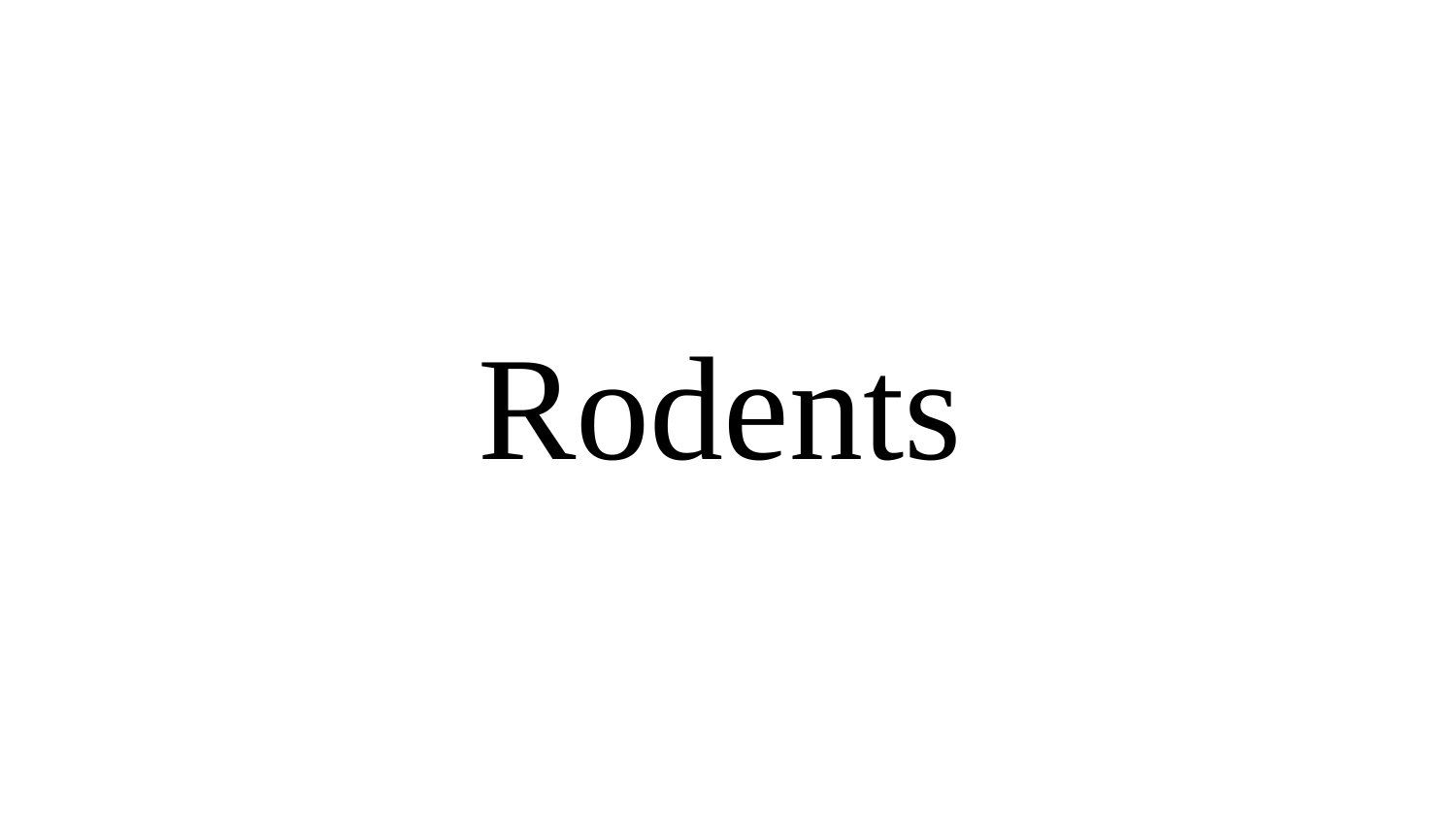

Rodents

### Slide 487
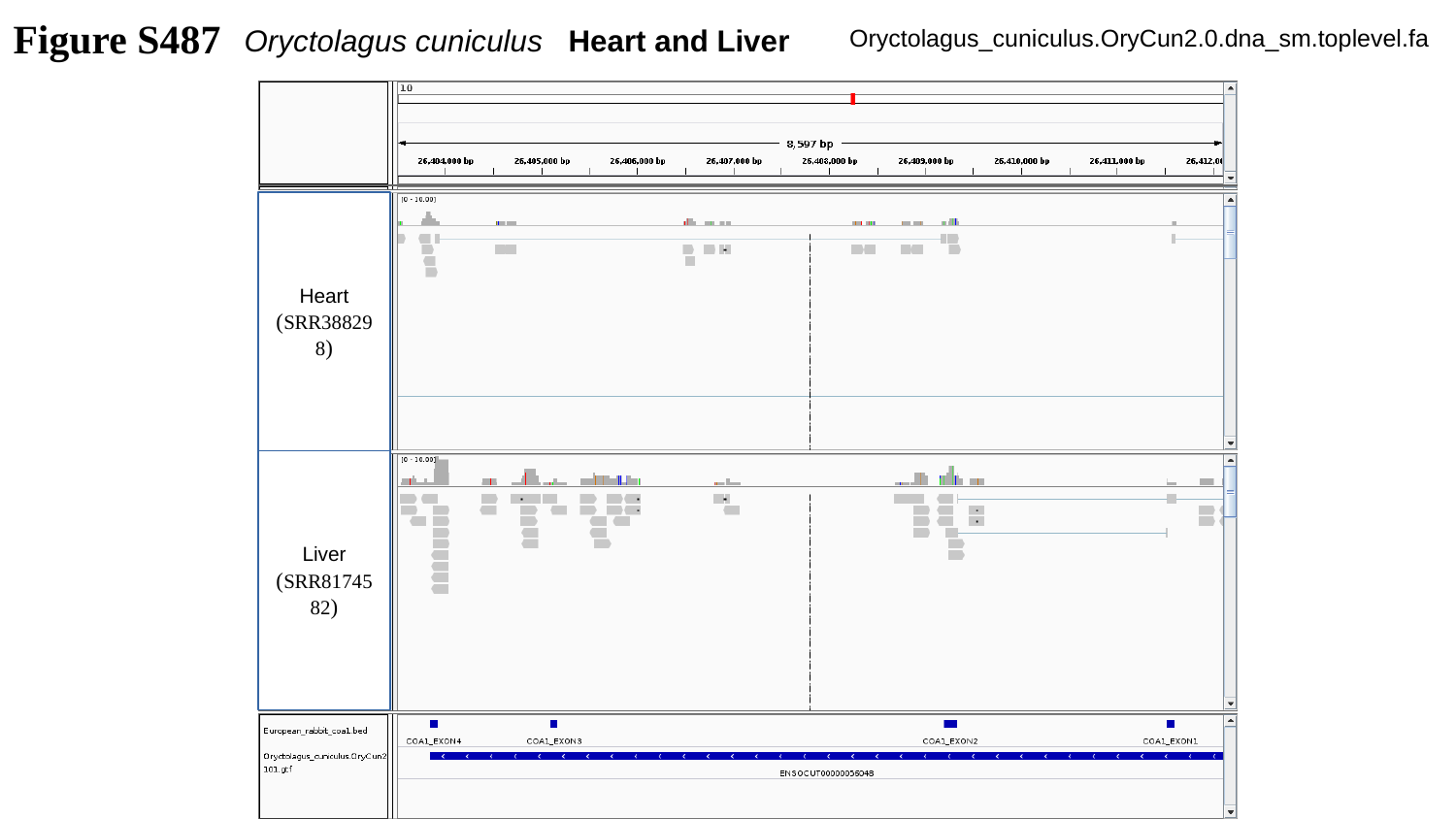

Figure S487
Oryctolagus cuniculus Heart and Liver
Oryctolagus_cuniculus.OryCun2.0.dna_sm.toplevel.fa
Heart
(SRR388298)
Liver
(SRR8174582)

### Slide 488
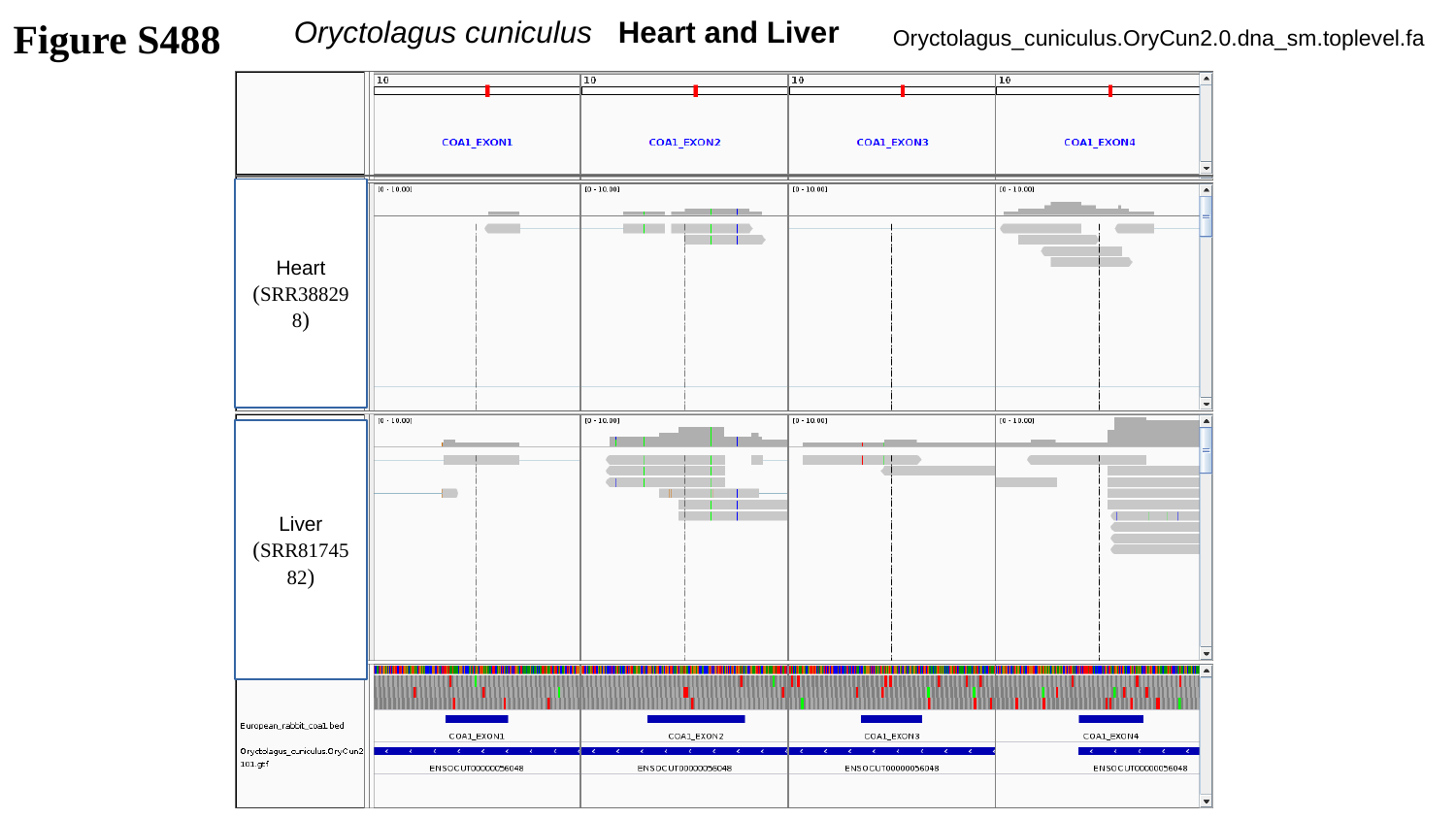

Figure S488
Oryctolagus cuniculus Heart and Liver
Oryctolagus_cuniculus.OryCun2.0.dna_sm.toplevel.fa
Heart
(SRR388298)
Liver
(SRR8174582)

### Slide 489
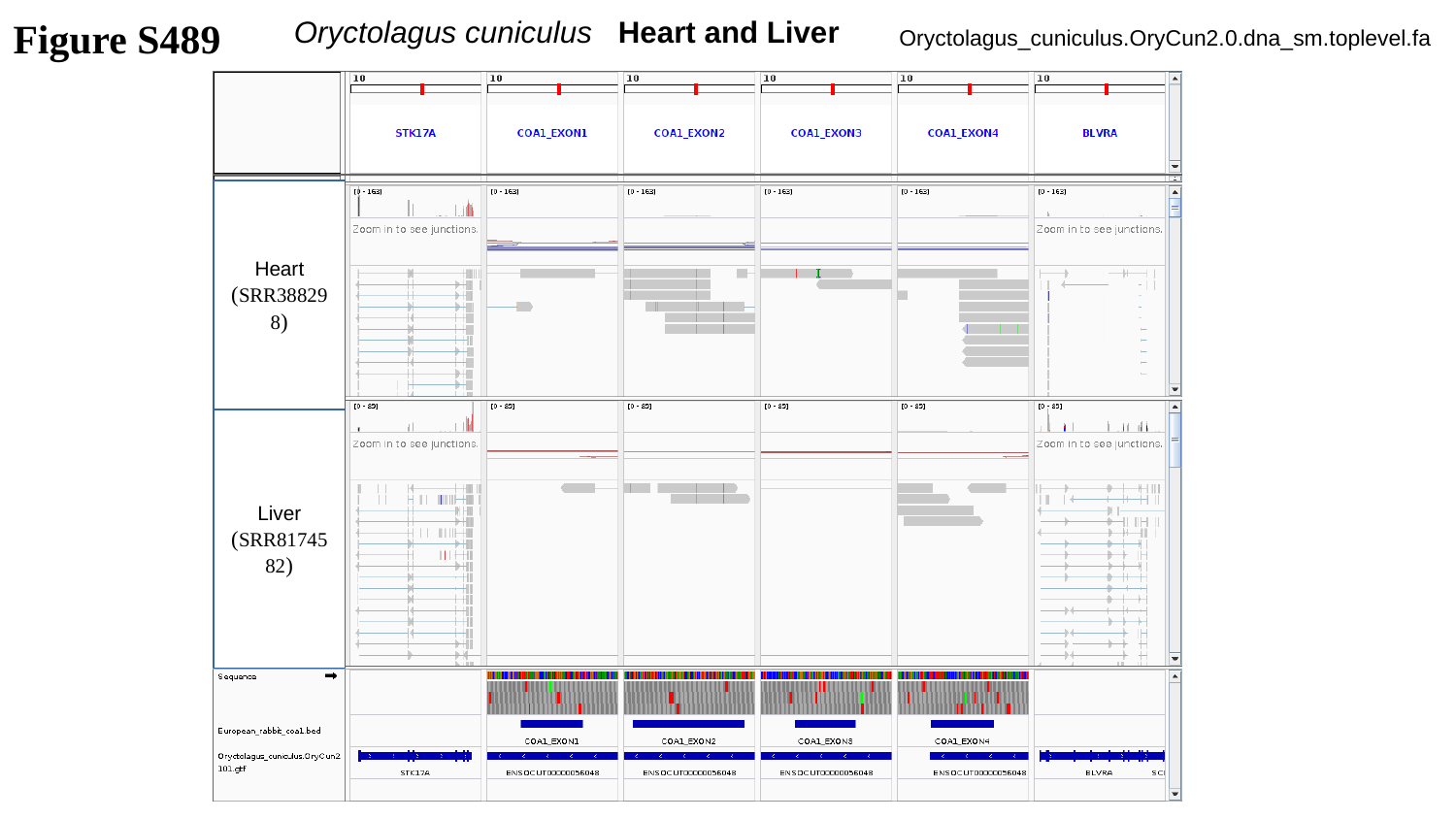

Figure S489
Oryctolagus cuniculus Heart and Liver
Oryctolagus_cuniculus.OryCun2.0.dna_sm.toplevel.fa
Heart
(SRR388298)
Liver
(SRR8174582)

### Slide 490
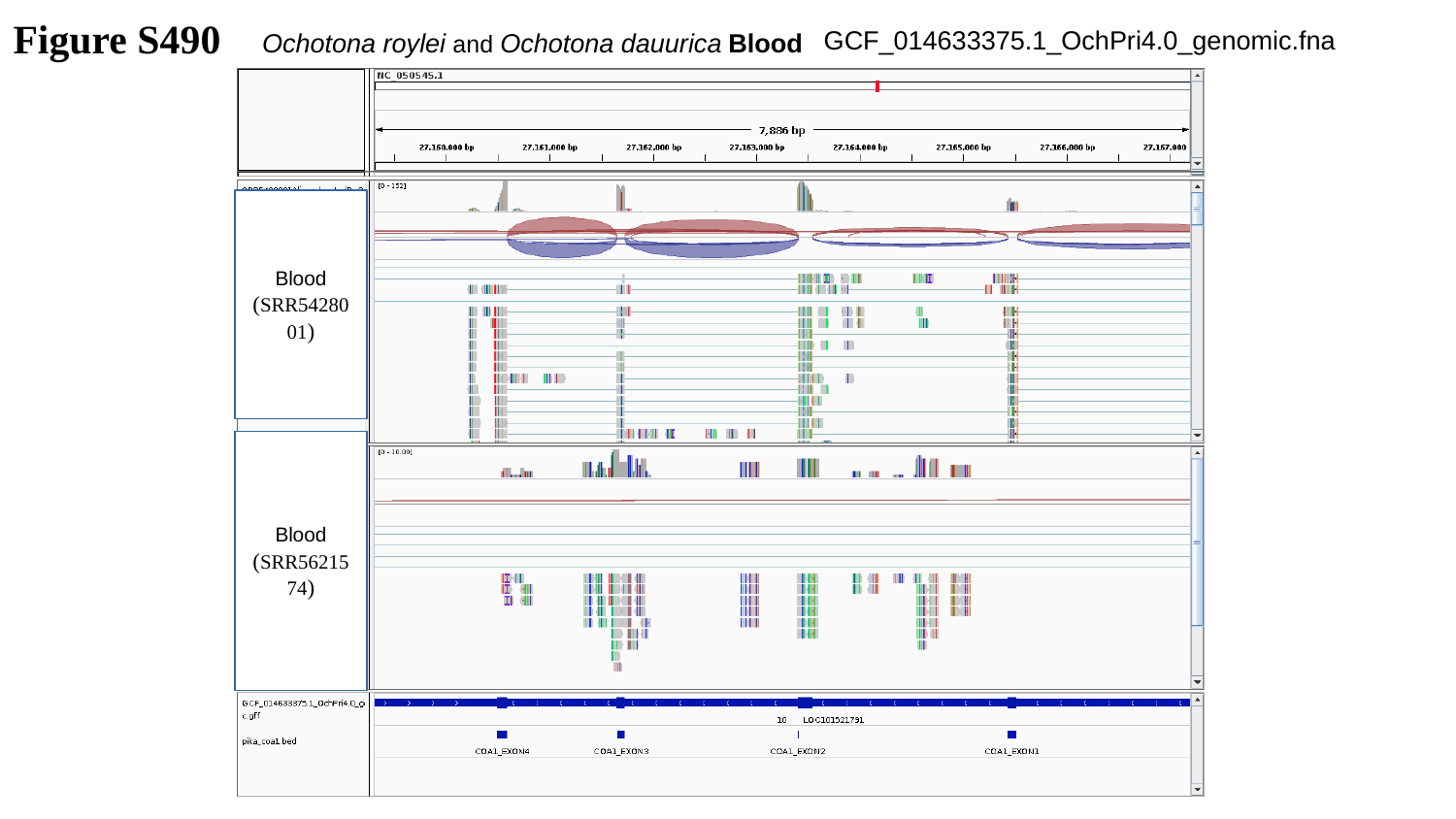

Figure S490
GCF_014633375.1_OchPri4.0_genomic.fna
Ochotona roylei and Ochotona dauurica Blood
Blood
(SRR5428001)
Blood
(SRR5621574)

### Slide 491
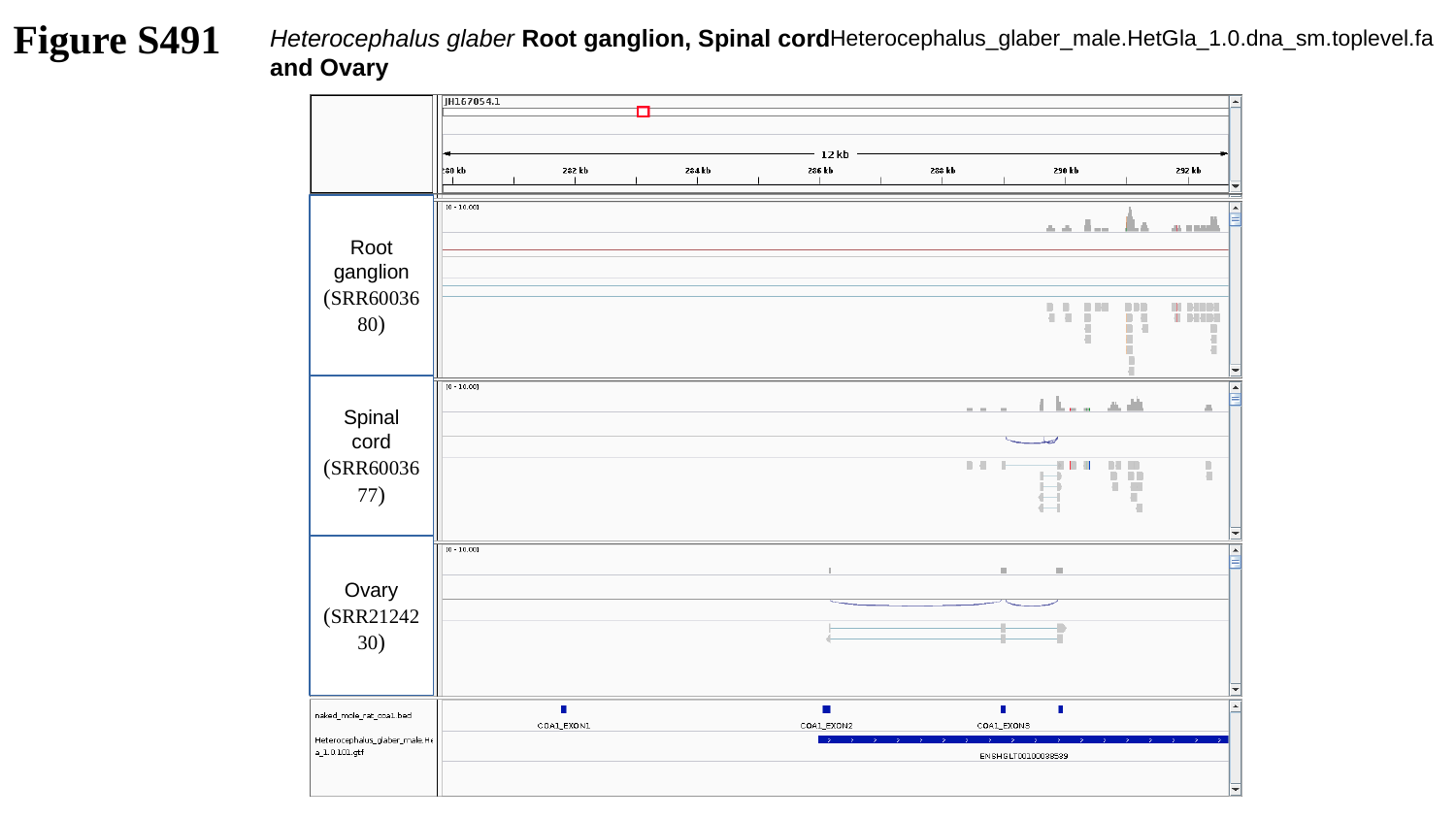

Figure S491
Heterocephalus glaber Root ganglion, Spinal cord and Ovary
Heterocephalus_glaber_male.HetGla_1.0.dna_sm.toplevel.fa
Root ganglion
(SRR6003680)
Spinal cord
(SRR6003677)
Ovary
(SRR2124230)

### Slide 492
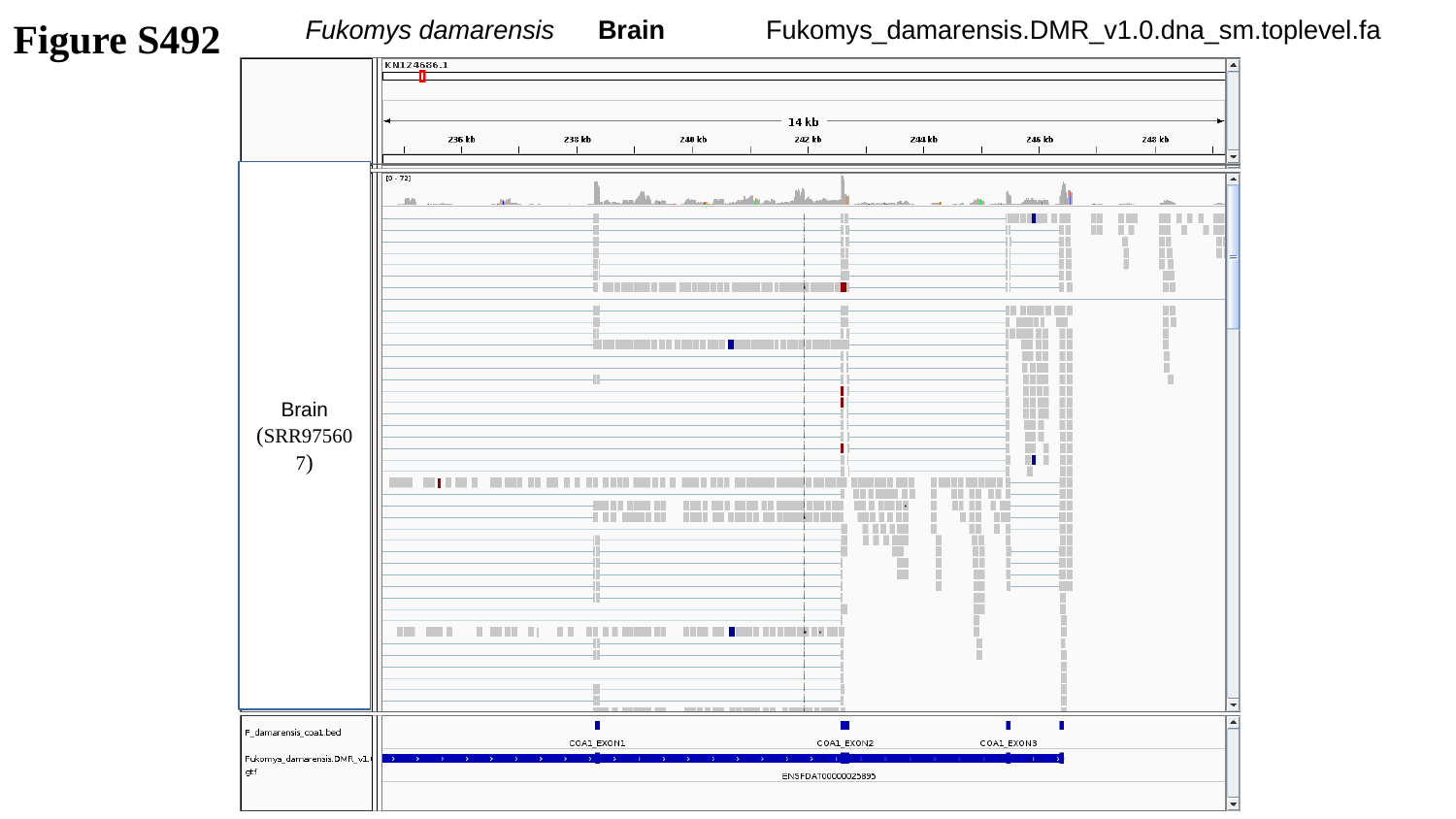

Figure S492
Fukomys damarensis Brain
Fukomys_damarensis.DMR_v1.0.dna_sm.toplevel.fa
Brain
(SRR975607)

### Slide 493
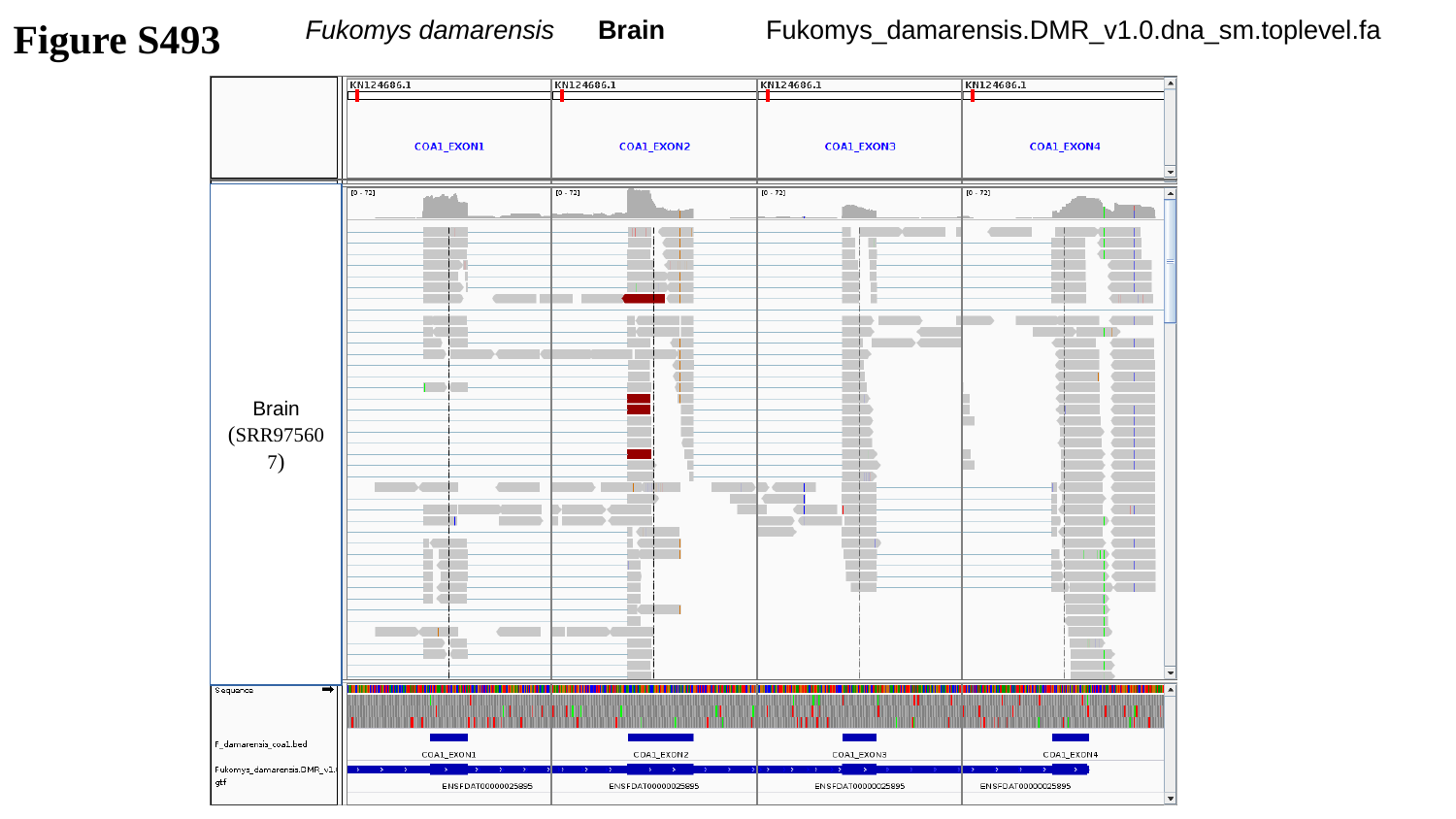

Figure S493
Fukomys damarensis Brain
Fukomys_damarensis.DMR_v1.0.dna_sm.toplevel.fa
Brain
(SRR975607)

### Slide 494
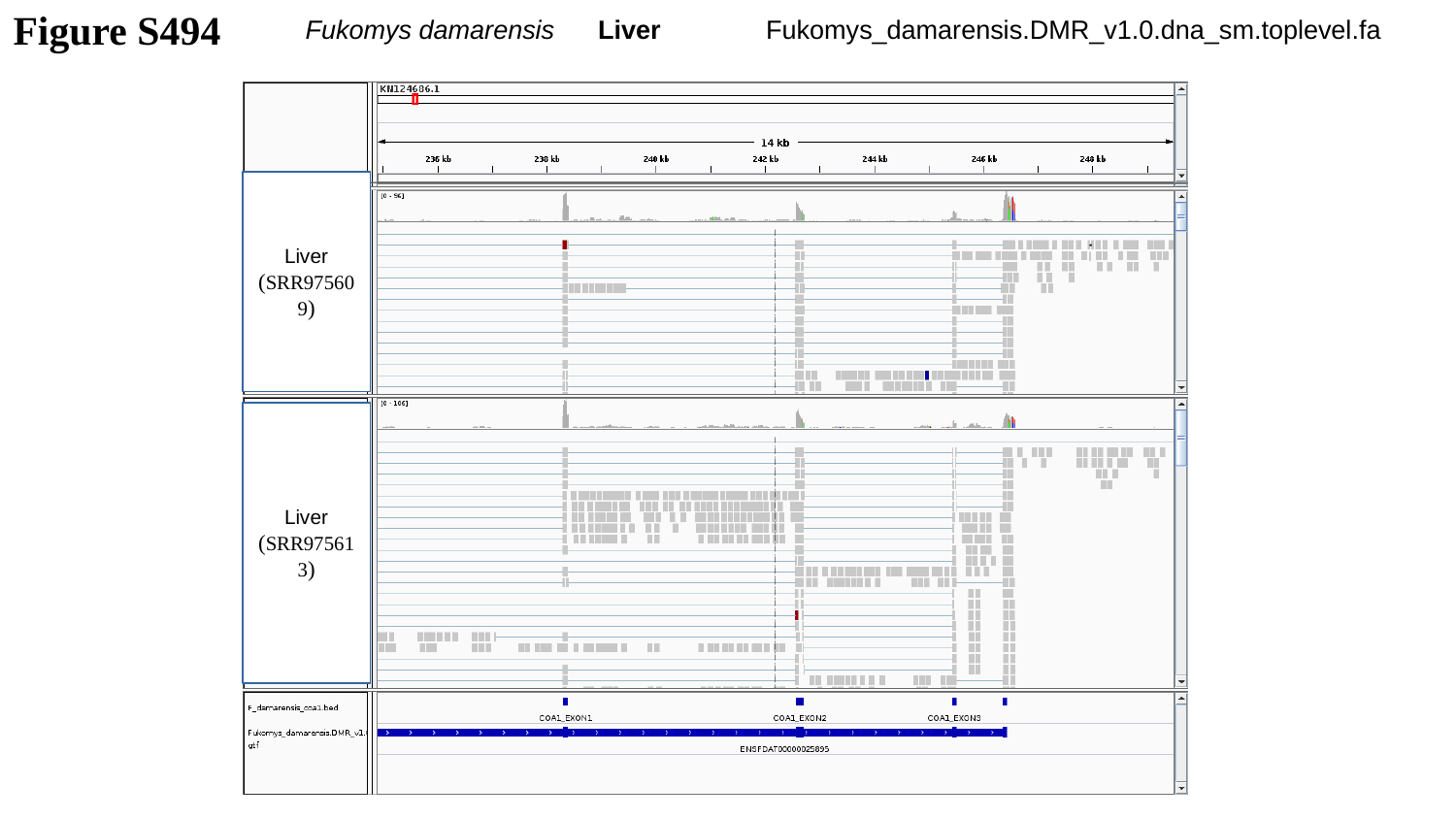

Fukomys damarensis Liver
Fukomys_damarensis.DMR_v1.0.dna_sm.toplevel.fa
Figure S494
Liver
(SRR975609)
Liver
(SRR975613)

### Slide 495
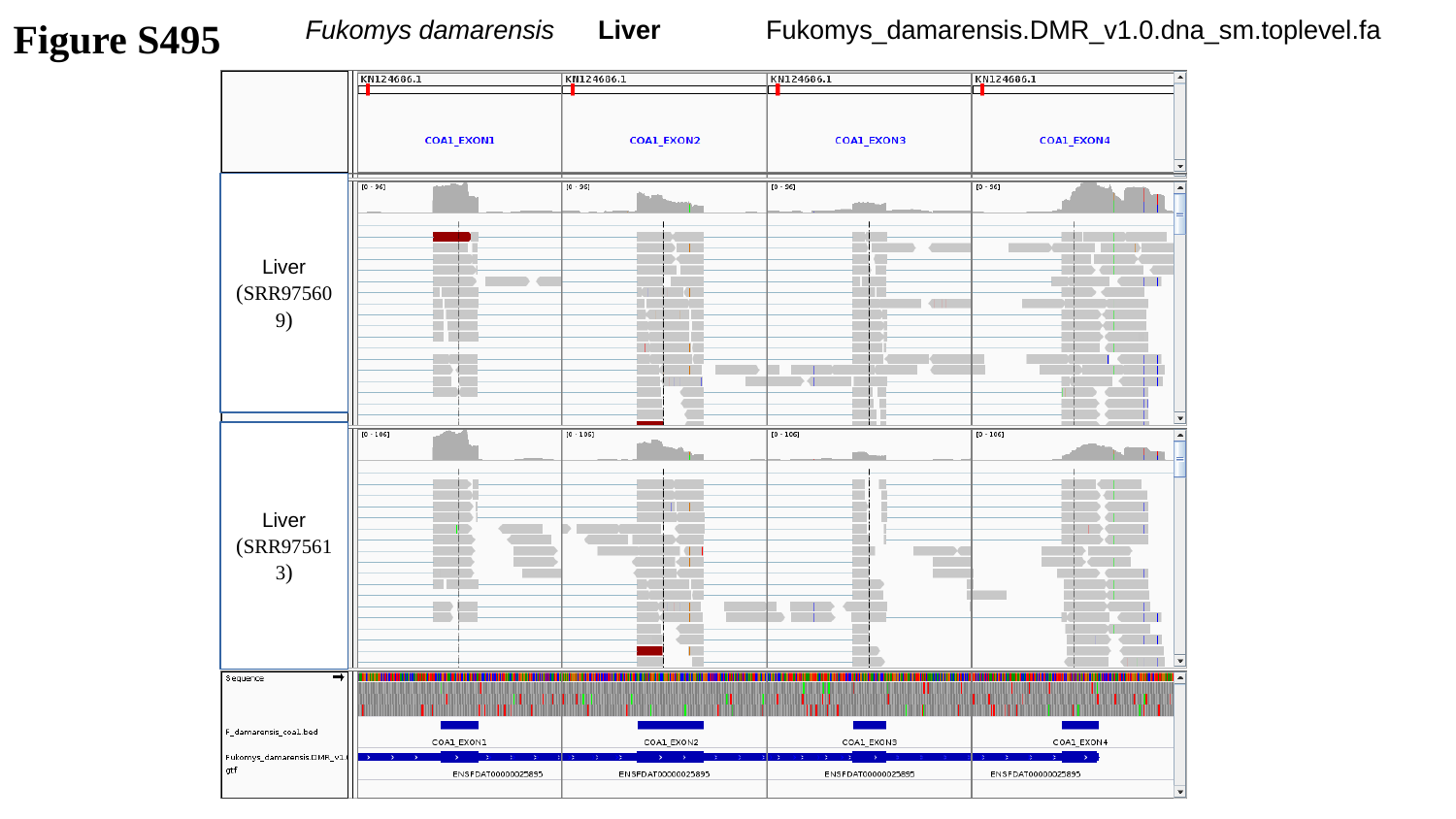

Figure S495
Fukomys damarensis Liver
Fukomys_damarensis.DMR_v1.0.dna_sm.toplevel.fa
Liver
(SRR975609)
Liver
(SRR975613)

### Slide 496
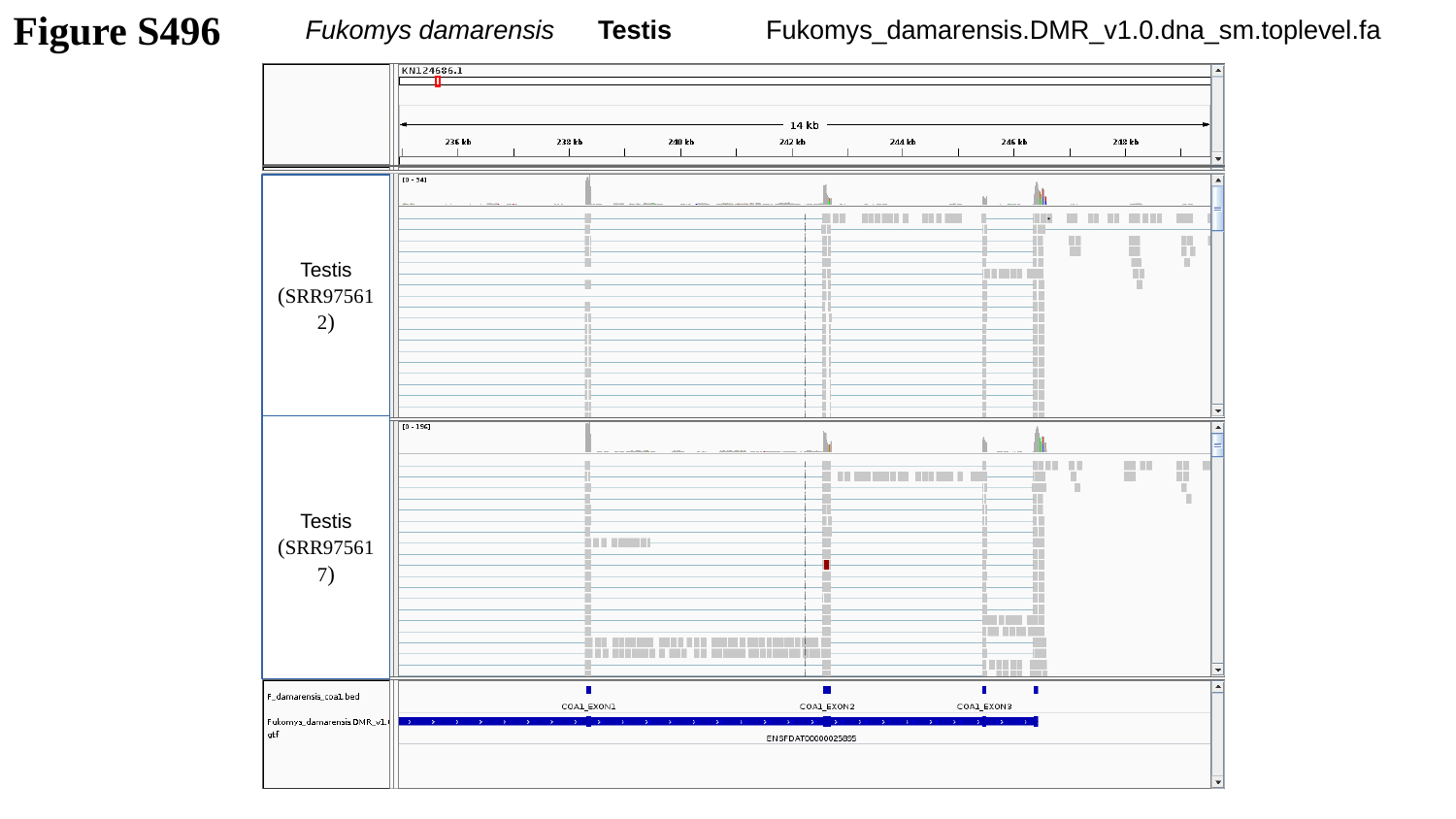

Fukomys damarensis Testis
Fukomys_damarensis.DMR_v1.0.dna_sm.toplevel.fa
Figure S496
Testis
(SRR975612)
Testis
(SRR975617)

### Slide 497
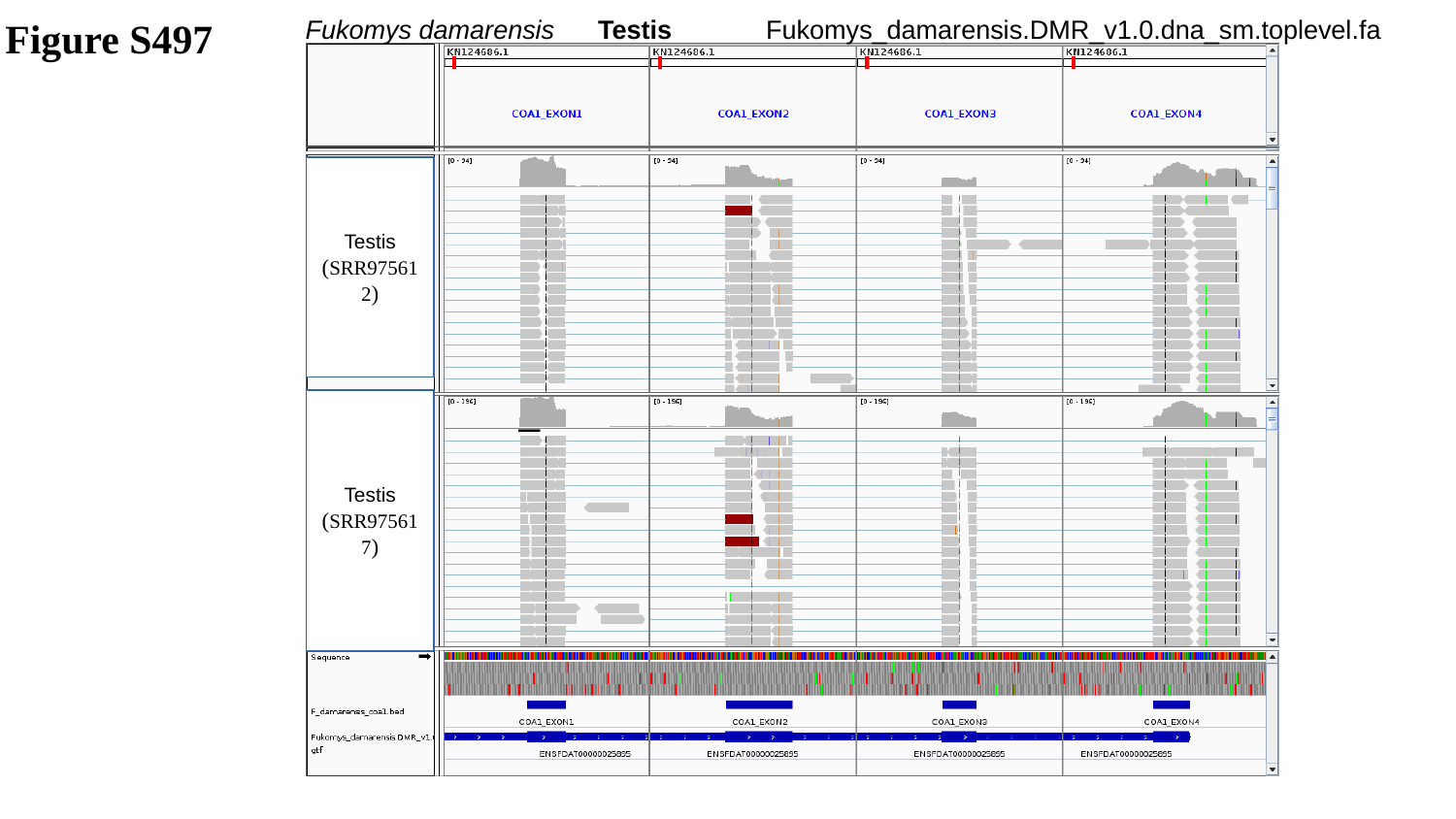

Figure S497
Fukomys damarensis Testis
Fukomys_damarensis.DMR_v1.0.dna_sm.toplevel.fa
Testis
(SRR975612)
Testis
(SRR975617)

### Slide 498
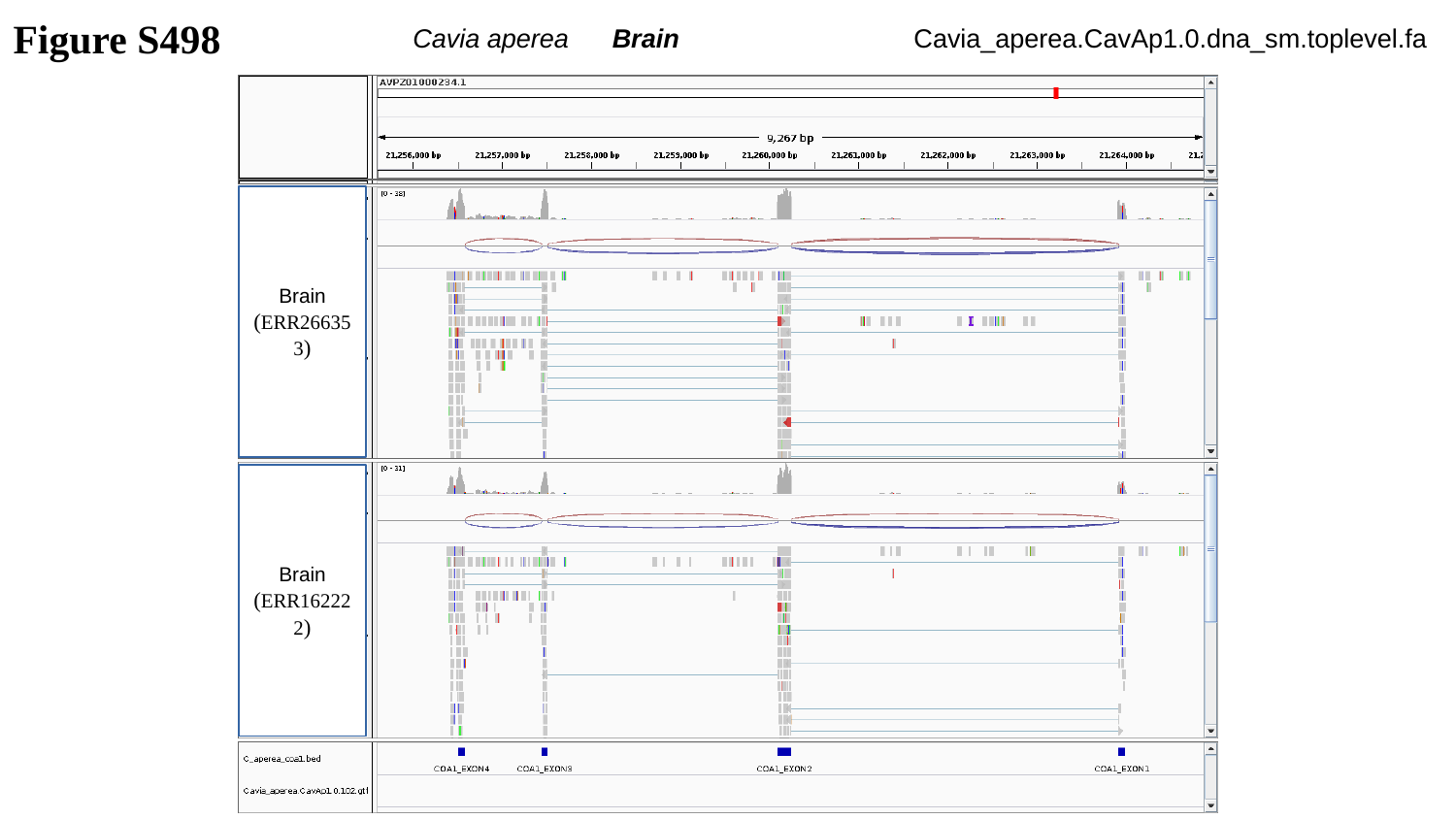

Figure S498
Cavia aperea Brain
Cavia_aperea.CavAp1.0.dna_sm.toplevel.fa
Brain
(ERR266353)
Brain
(ERR162222)

### Slide 499
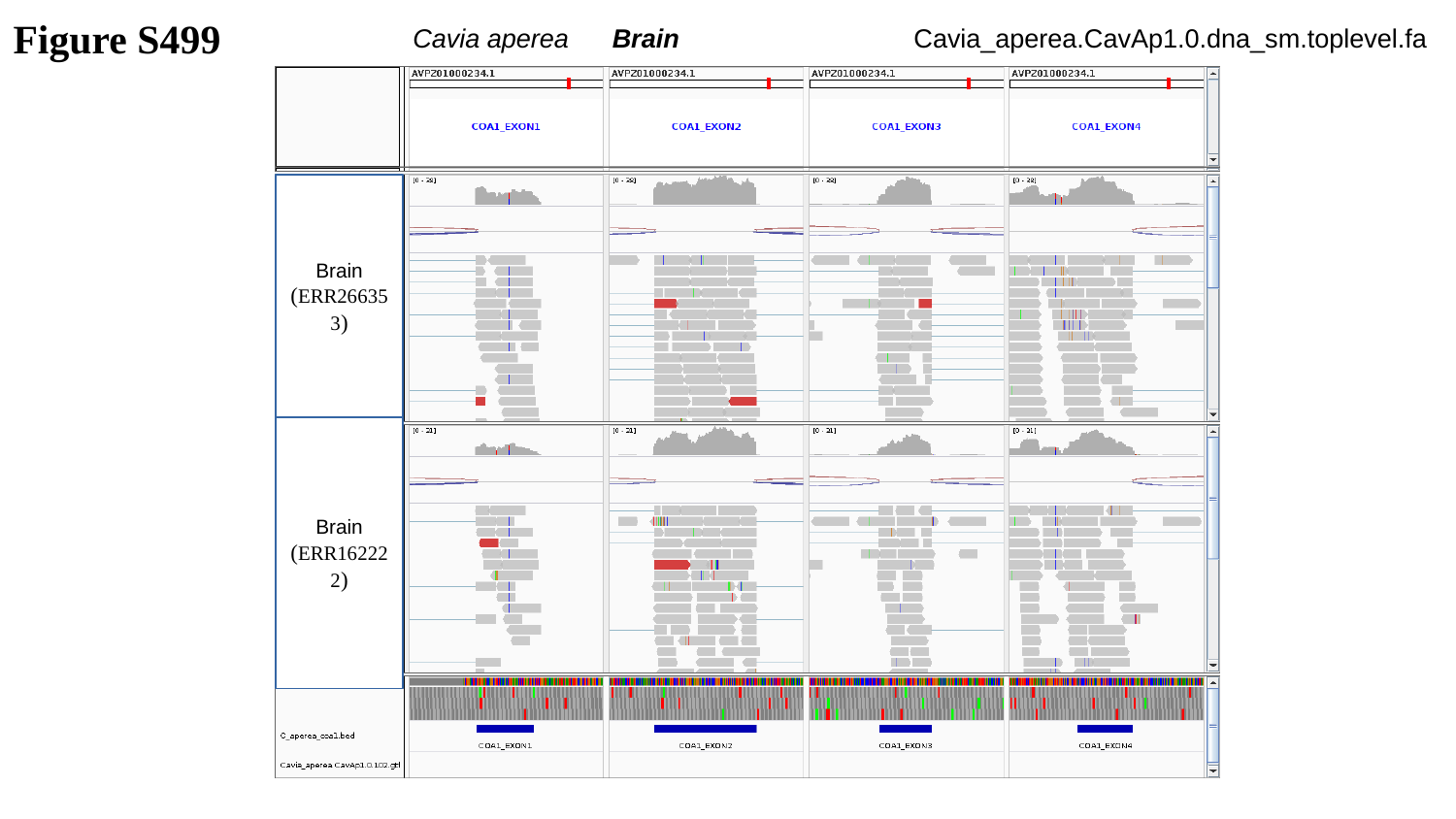

Figure S499
Cavia aperea Brain
Cavia_aperea.CavAp1.0.dna_sm.toplevel.fa
Brain
(ERR266353)
Brain
(ERR162222)

### Slide 500
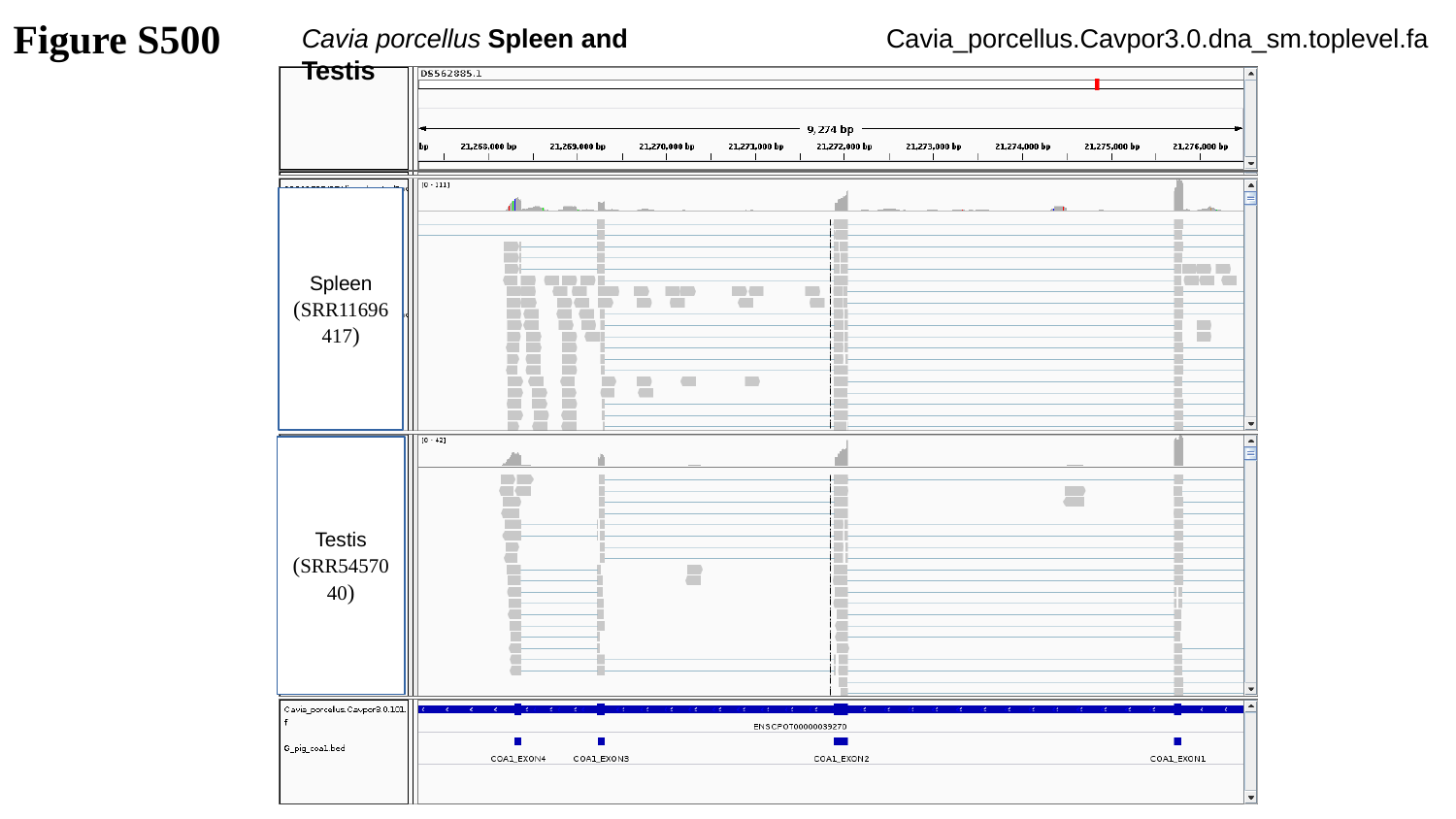

Figure S500
Cavia porcellus Spleen and Testis
Cavia_porcellus.Cavpor3.0.dna_sm.toplevel.fa
Spleen
(SRR11696417)
Testis
(SRR5457040)

### Slide 501
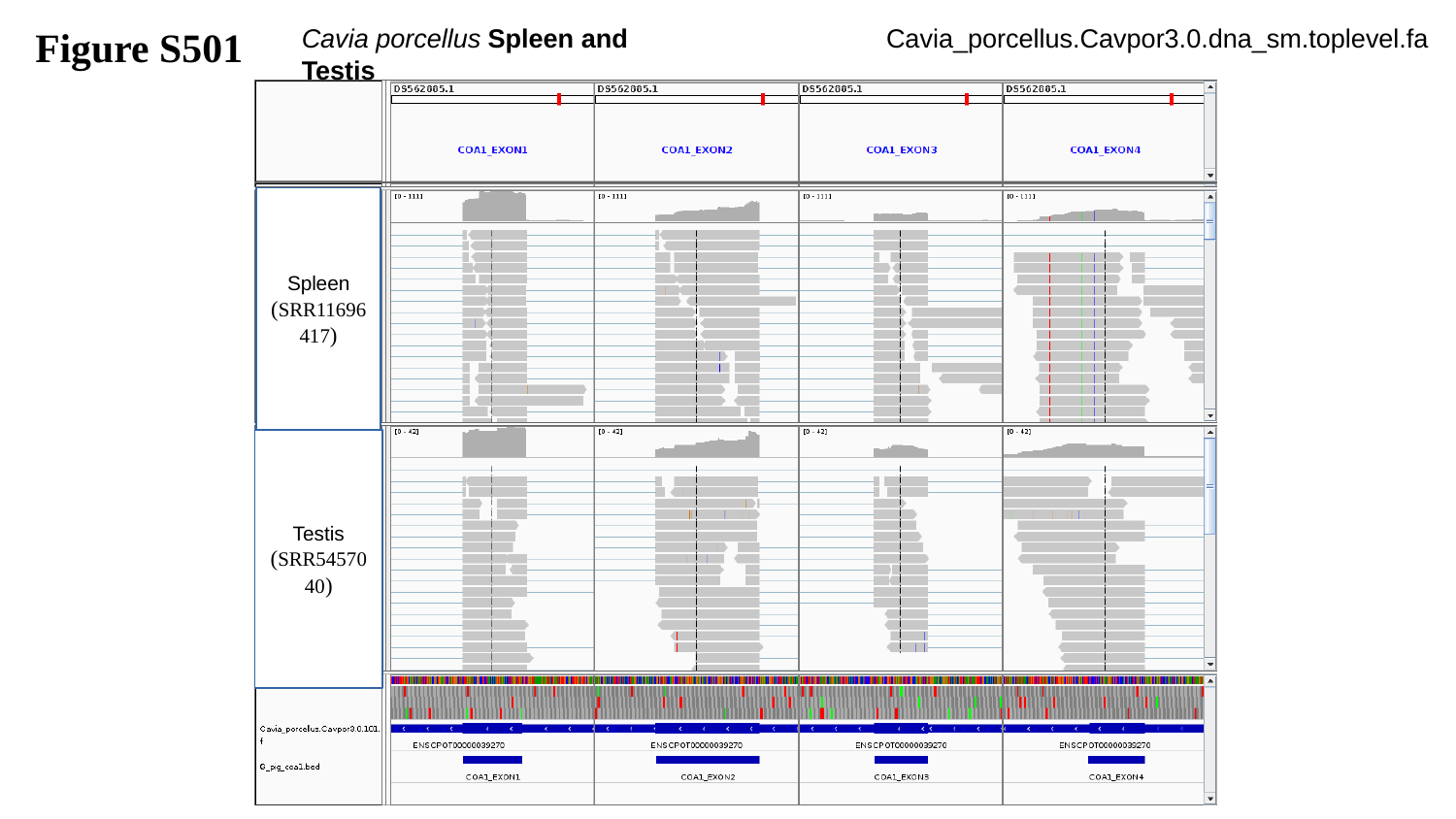

Figure S501
Cavia porcellus Spleen and Testis
Cavia_porcellus.Cavpor3.0.dna_sm.toplevel.fa
Spleen
(SRR11696417)
Testis
(SRR5457040)

### Slide 502
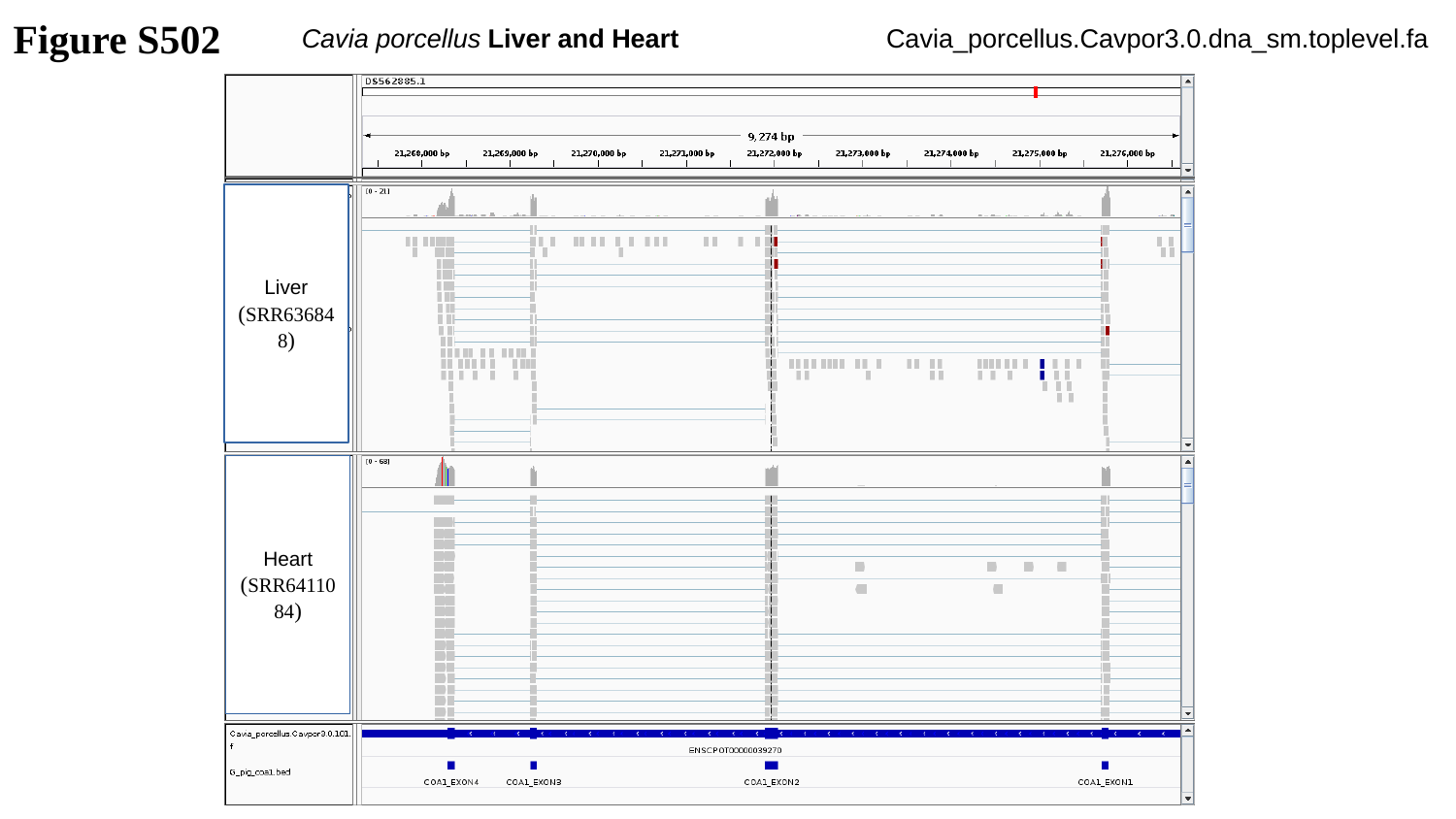

Figure S502
Cavia porcellus Liver and Heart
Cavia_porcellus.Cavpor3.0.dna_sm.toplevel.fa
Liver
(SRR636848)
Heart
(SRR6411084)

### Slide 503
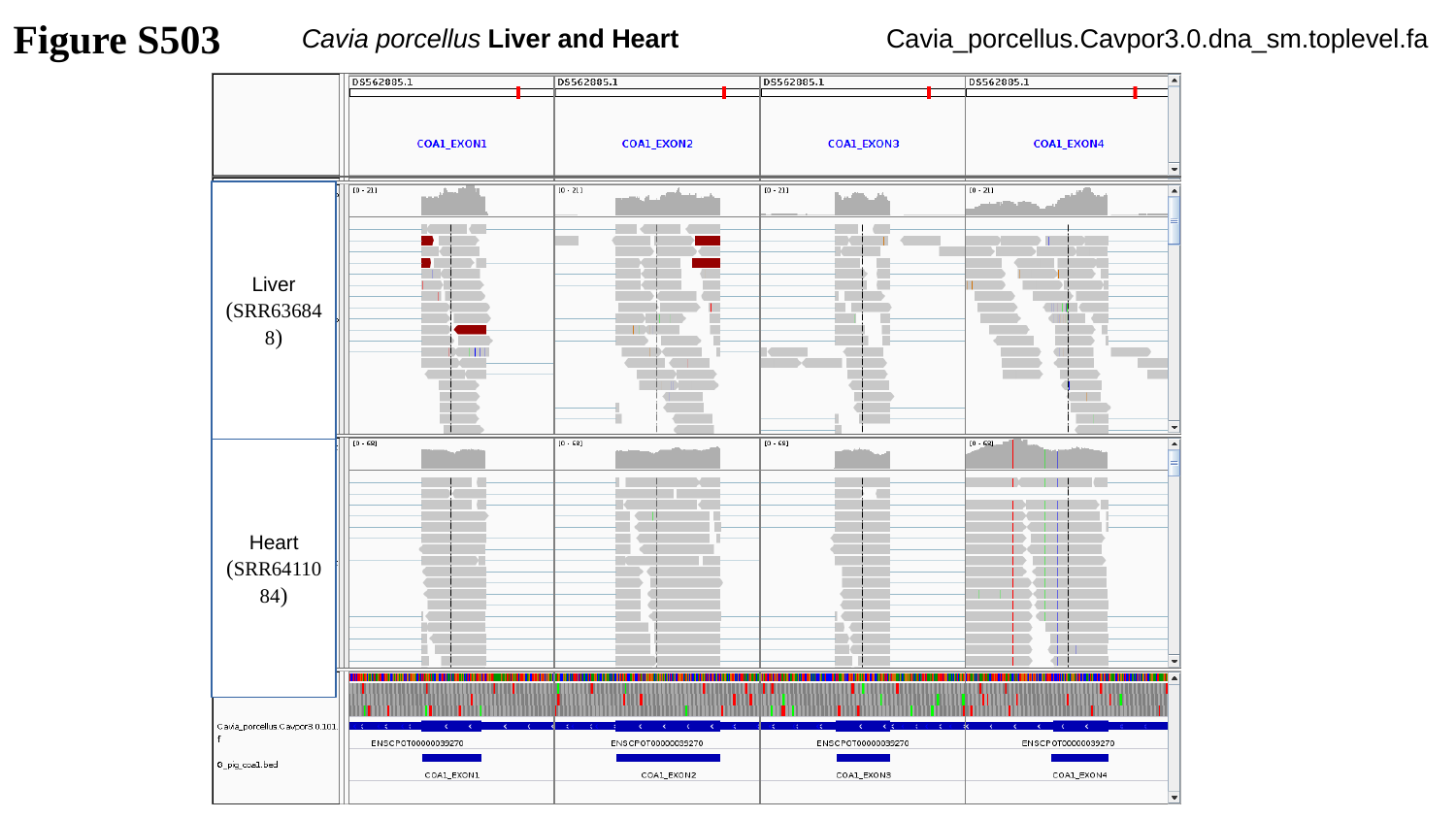

Figure S503
Cavia porcellus Liver and Heart
Cavia_porcellus.Cavpor3.0.dna_sm.toplevel.fa
Liver
(SRR636848)
Heart
(SRR6411084)

### Slide 504
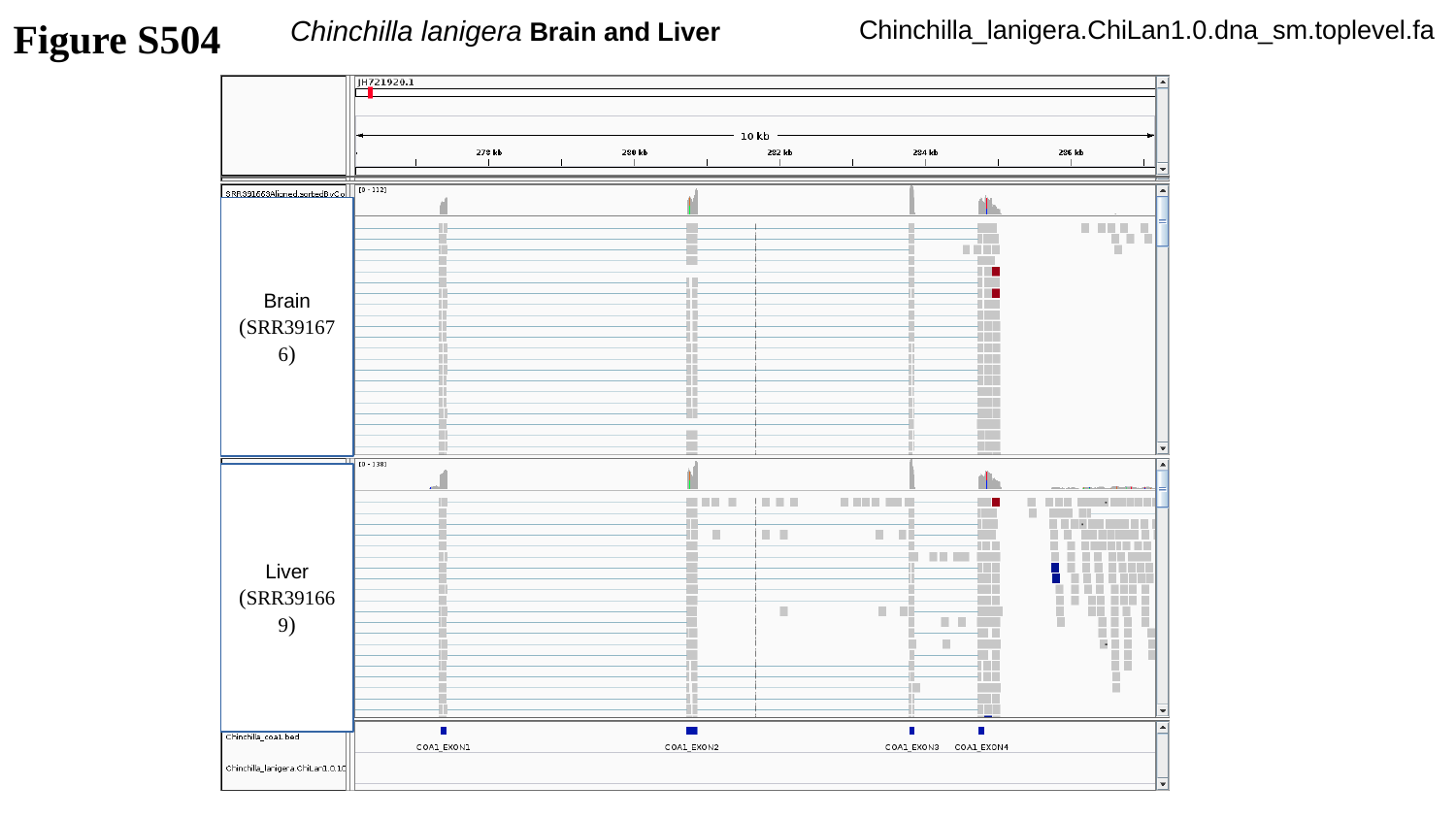

Figure S504
Chinchilla lanigera Brain and Liver
Chinchilla_lanigera.ChiLan1.0.dna_sm.toplevel.fa
Brain
(SRR391676)
Liver
(SRR391669)

### Slide 505
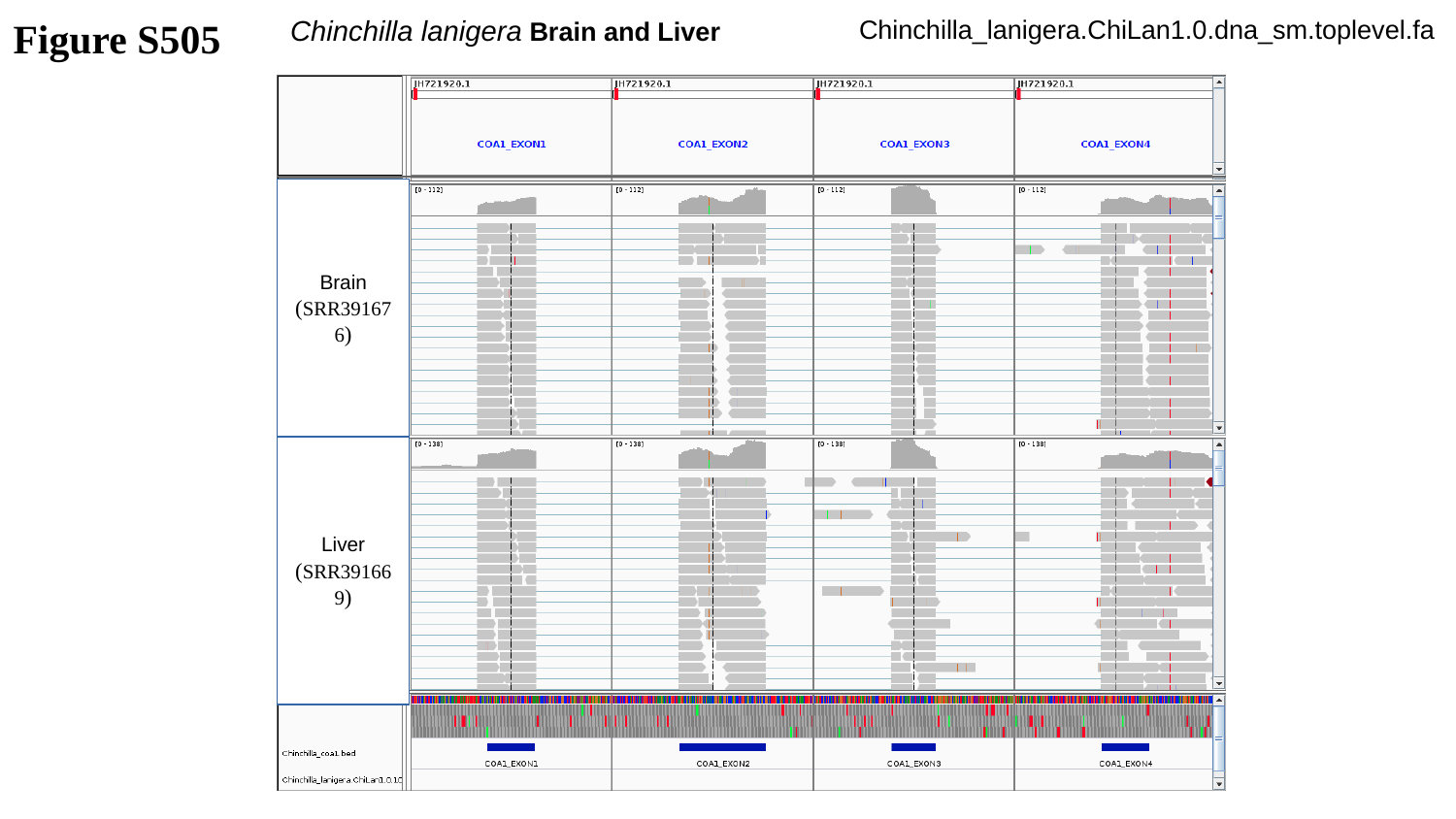

Figure S505
Chinchilla lanigera Brain and Liver
Chinchilla_lanigera.ChiLan1.0.dna_sm.toplevel.fa
Brain
(SRR391676)
Liver
(SRR391669)

### Slide 506
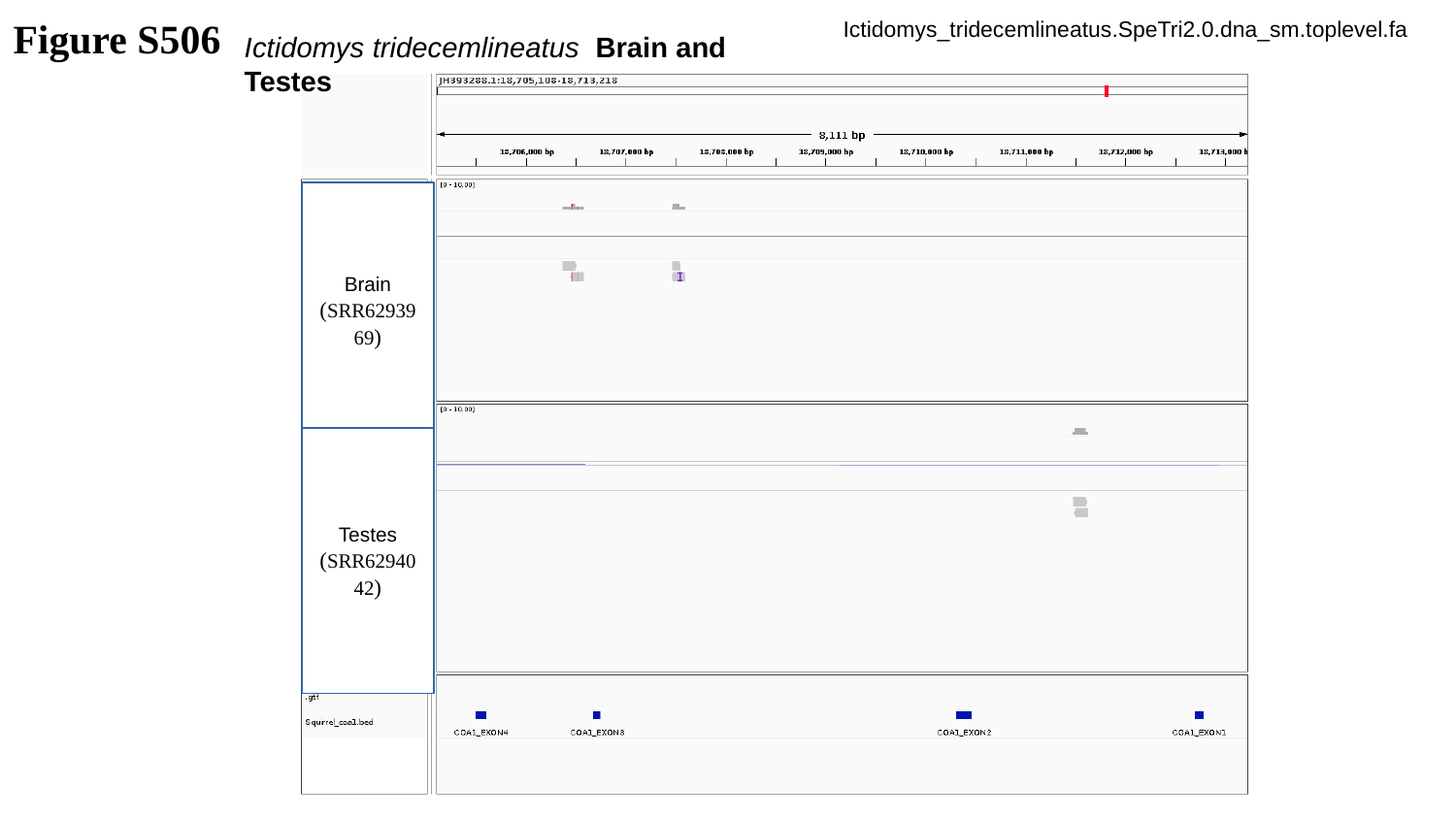

Figure S506
Ictidomys_tridecemlineatus.SpeTri2.0.dna_sm.toplevel.fa
Ictidomys tridecemlineatus Brain and Testes
Brain
(SRR6293969)
Testes
(SRR6294042)

### Slide 507
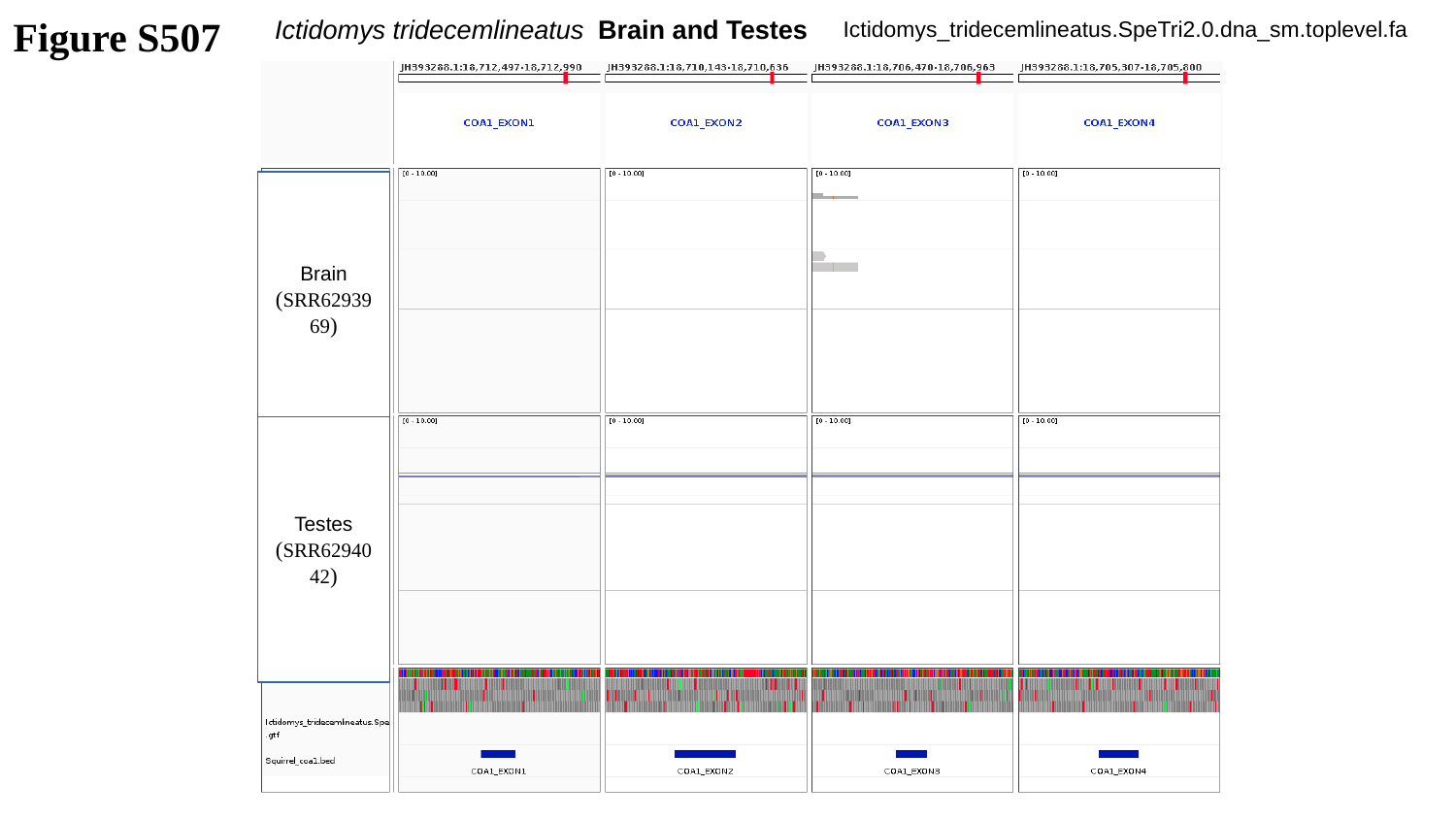

Ictidomys tridecemlineatus Brain and Testes
Figure S507
Ictidomys_tridecemlineatus.SpeTri2.0.dna_sm.toplevel.fa
Brain
(SRR6293969)
Testes
(SRR6294042)

### Slide 508
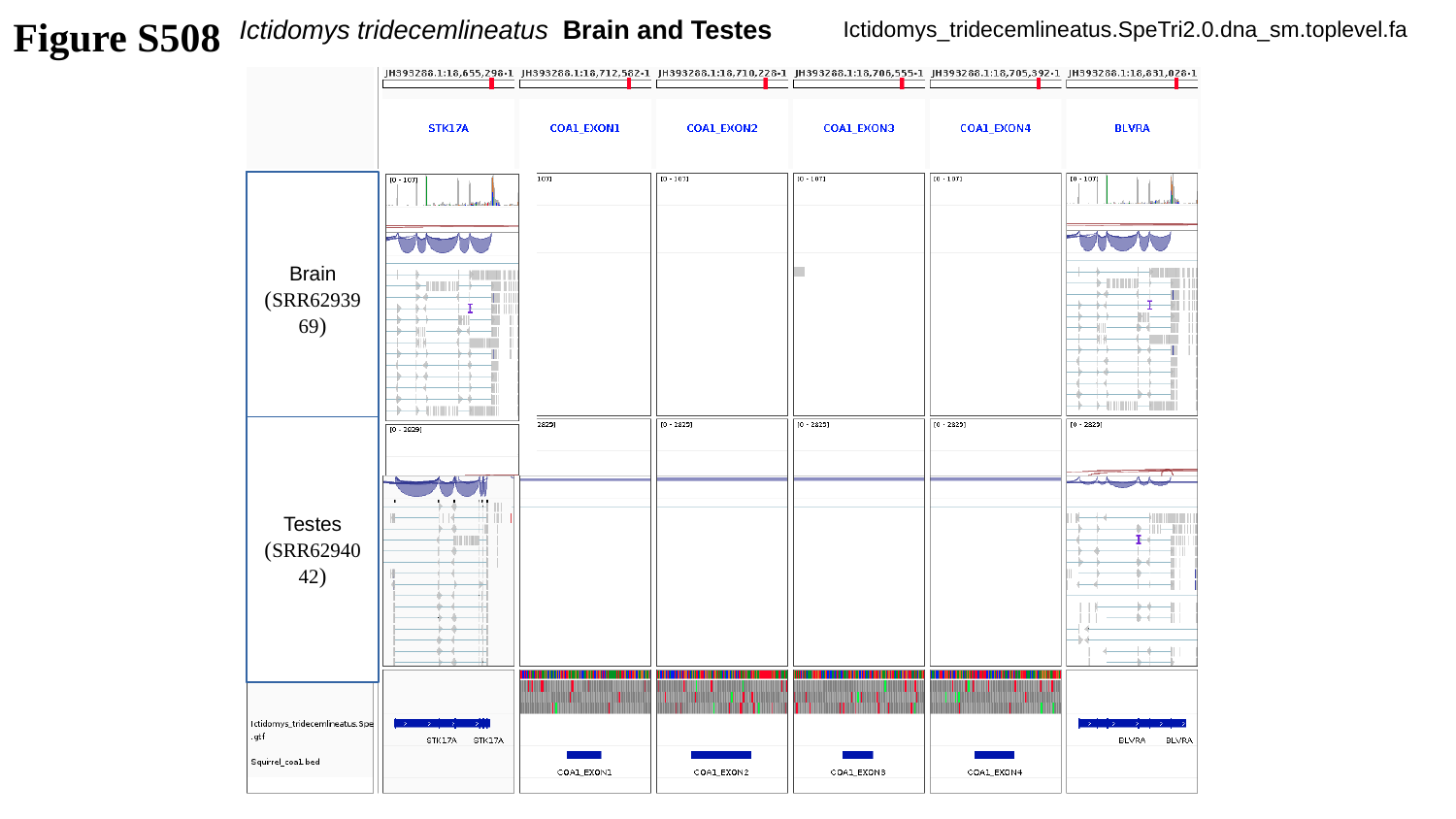

Ictidomys tridecemlineatus Brain and Testes
Figure S508
Ictidomys_tridecemlineatus.SpeTri2.0.dna_sm.toplevel.fa
Brain
(SRR6293969)
Brain
(SRR6293969)
Testes
(SRR6294042)
Testes
(SRR6294042)

### Slide 509
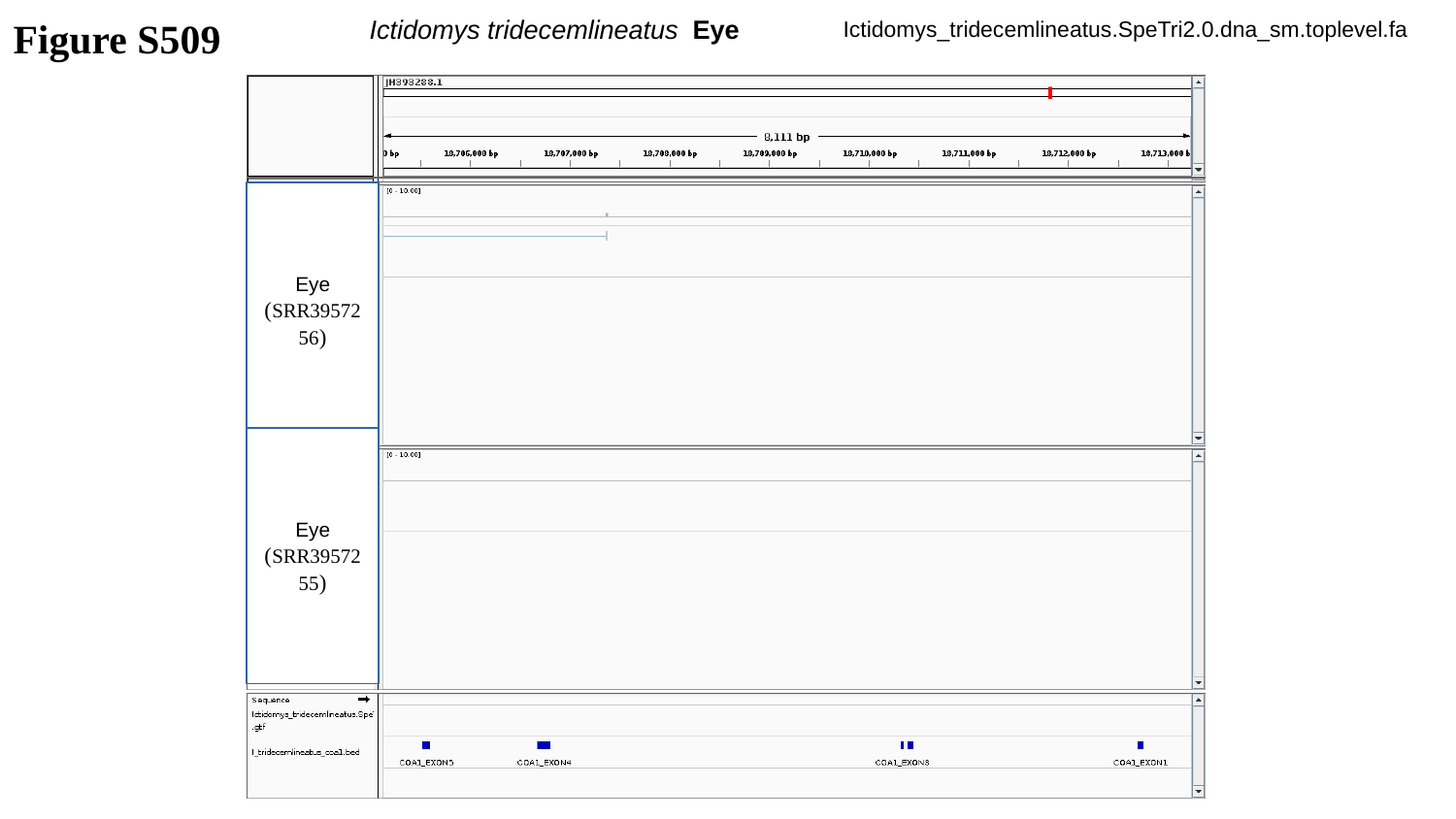

Figure S509
Ictidomys tridecemlineatus Eye
Ictidomys_tridecemlineatus.SpeTri2.0.dna_sm.toplevel.fa
Eye
(SRR3957256)
Eye
(SRR3957255)

### Slide 510
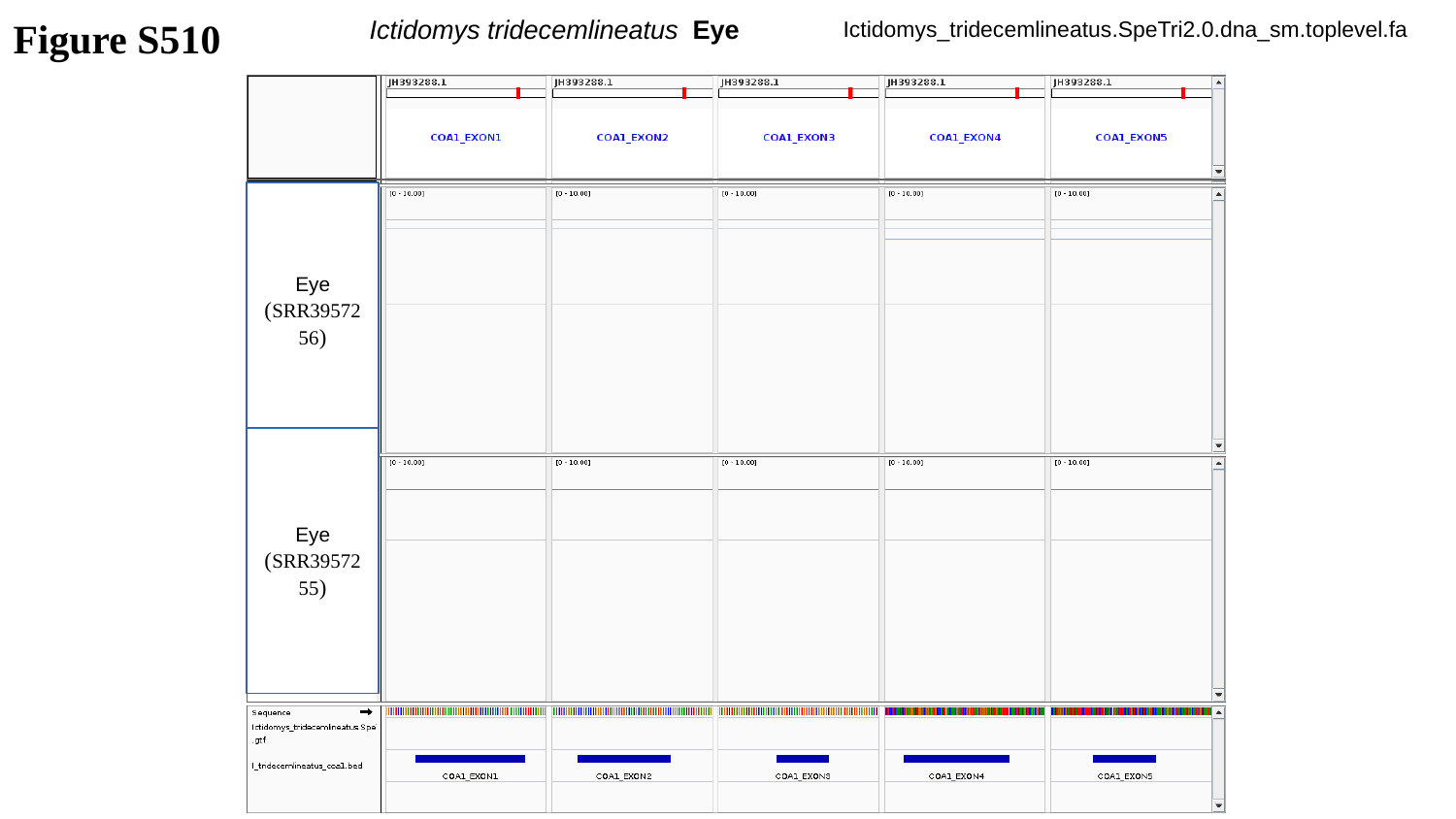

Figure S510
Ictidomys tridecemlineatus Eye
Ictidomys_tridecemlineatus.SpeTri2.0.dna_sm.toplevel.fa
Eye
(SRR3957256)
Eye
(SRR3957255)

### Slide 511
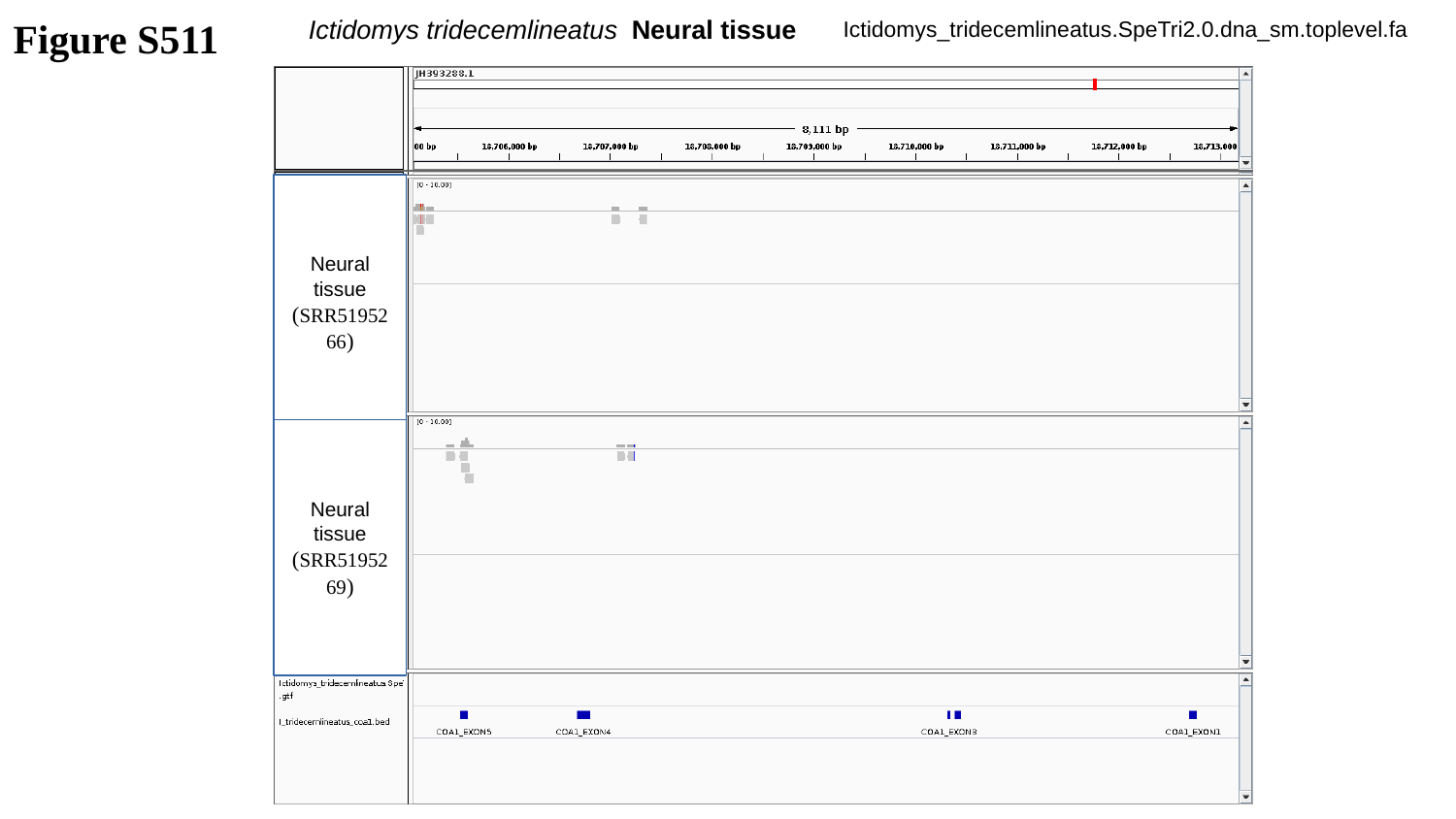

Figure S511
Ictidomys tridecemlineatus Neural tissue
Ictidomys_tridecemlineatus.SpeTri2.0.dna_sm.toplevel.fa
Neural tissue
(SRR5195266)
Neural tissue
(SRR5195269)

### Slide 512
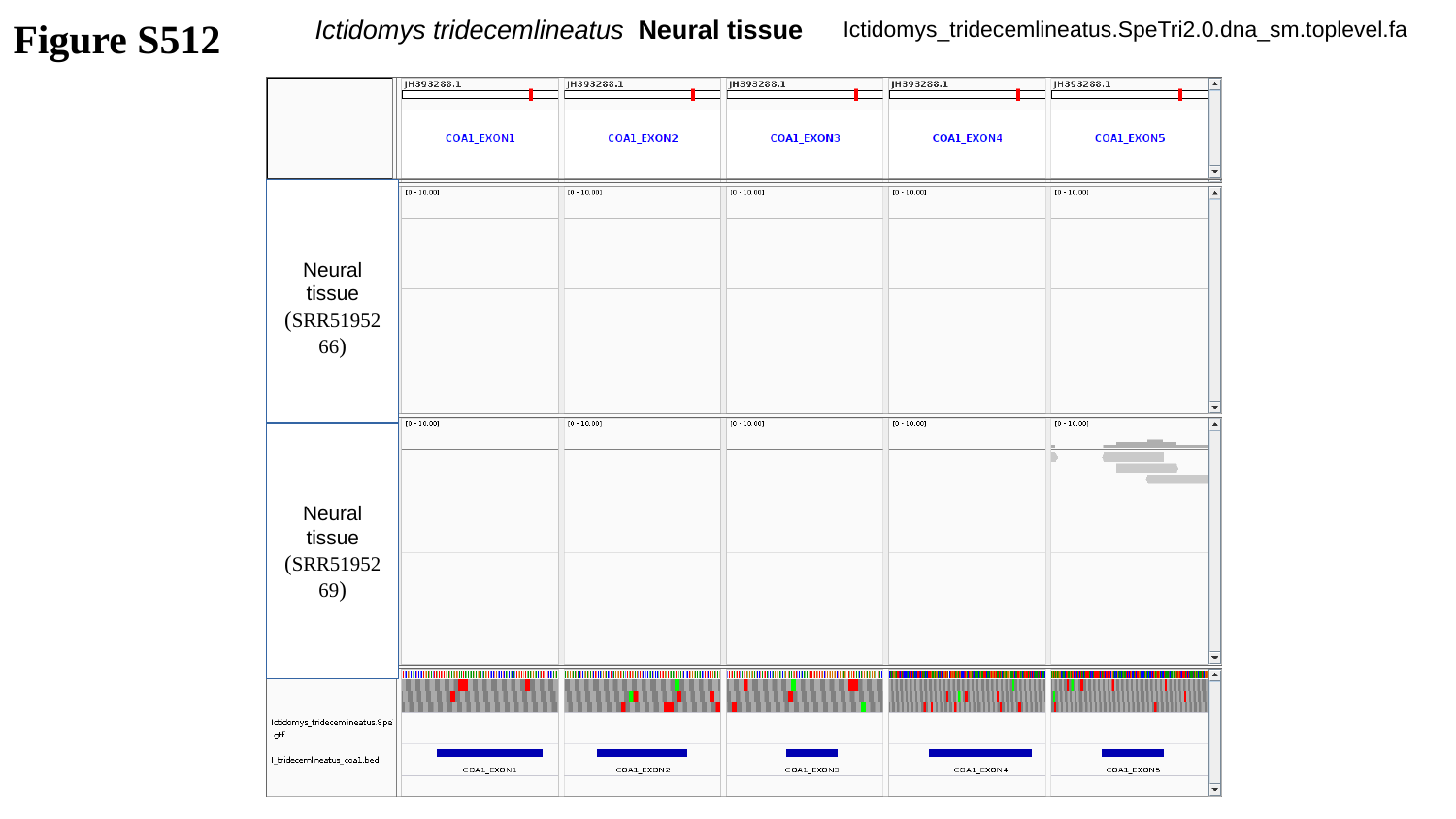

Figure S512
Ictidomys tridecemlineatus Neural tissue
Ictidomys_tridecemlineatus.SpeTri2.0.dna_sm.toplevel.fa
Neural tissue
(SRR5195266)
Neural tissue
(SRR5195269)

### Slide 513
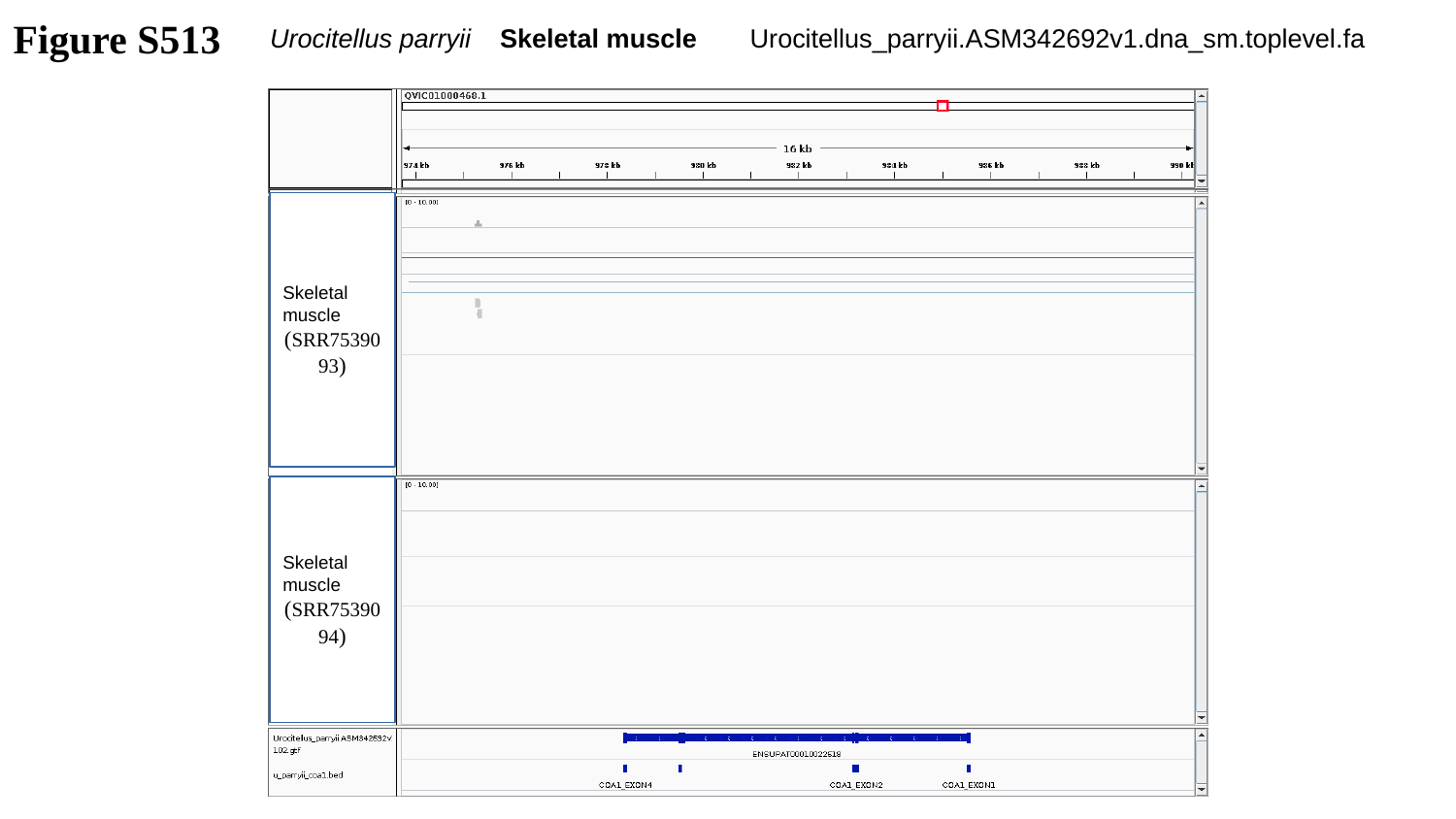

Figure S513
Urocitellus parryii Skeletal muscle
Urocitellus_parryii.ASM342692v1.dna_sm.toplevel.fa
Skeletal muscle
(SRR7539093)
Skeletal muscle
(SRR7539094)

### Slide 514
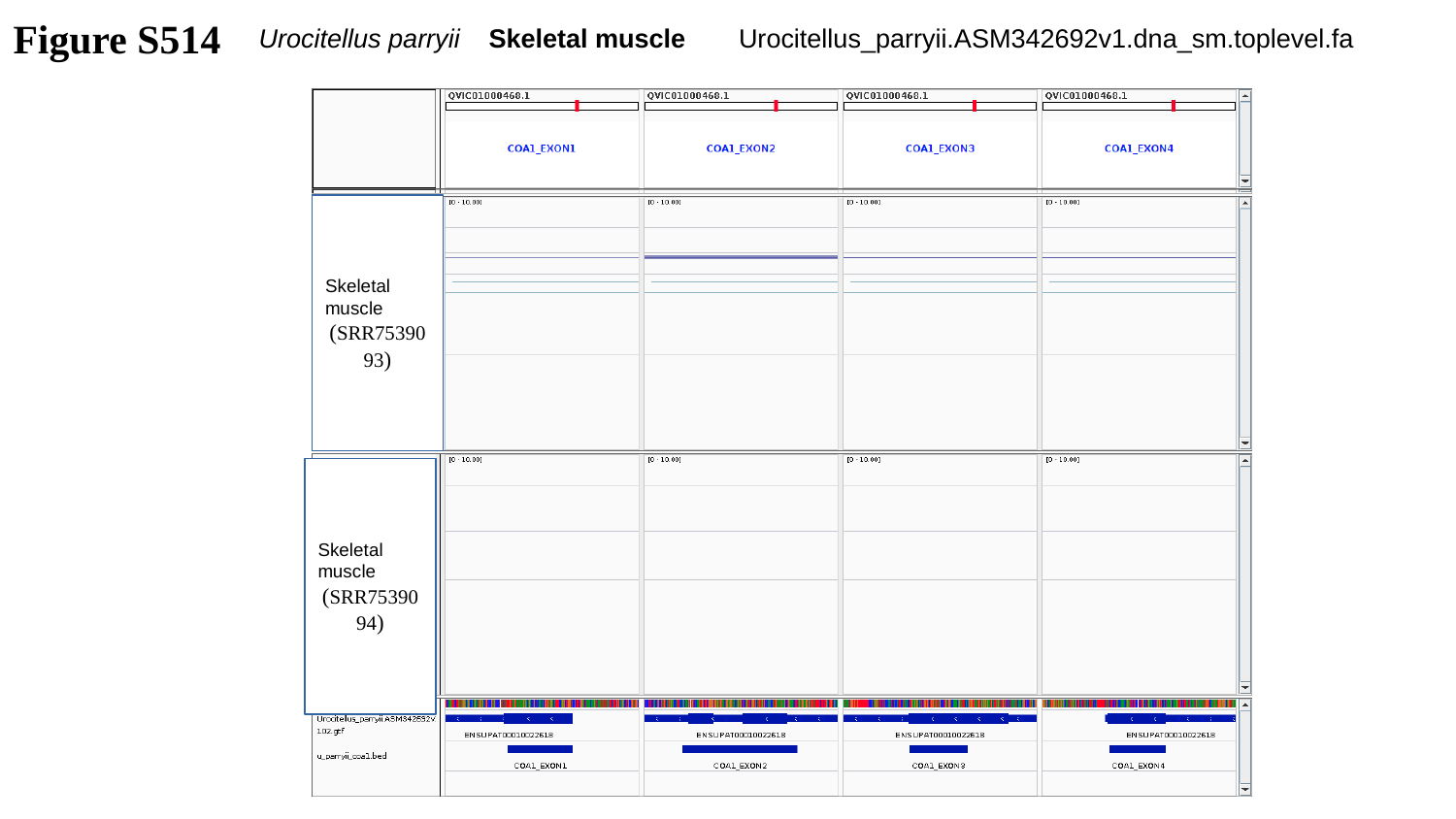

Figure S514
Urocitellus parryii Skeletal muscle
Urocitellus_parryii.ASM342692v1.dna_sm.toplevel.fa
Skeletal muscle
(SRR7539093)
Skeletal muscle
(SRR7539094)

### Slide 515
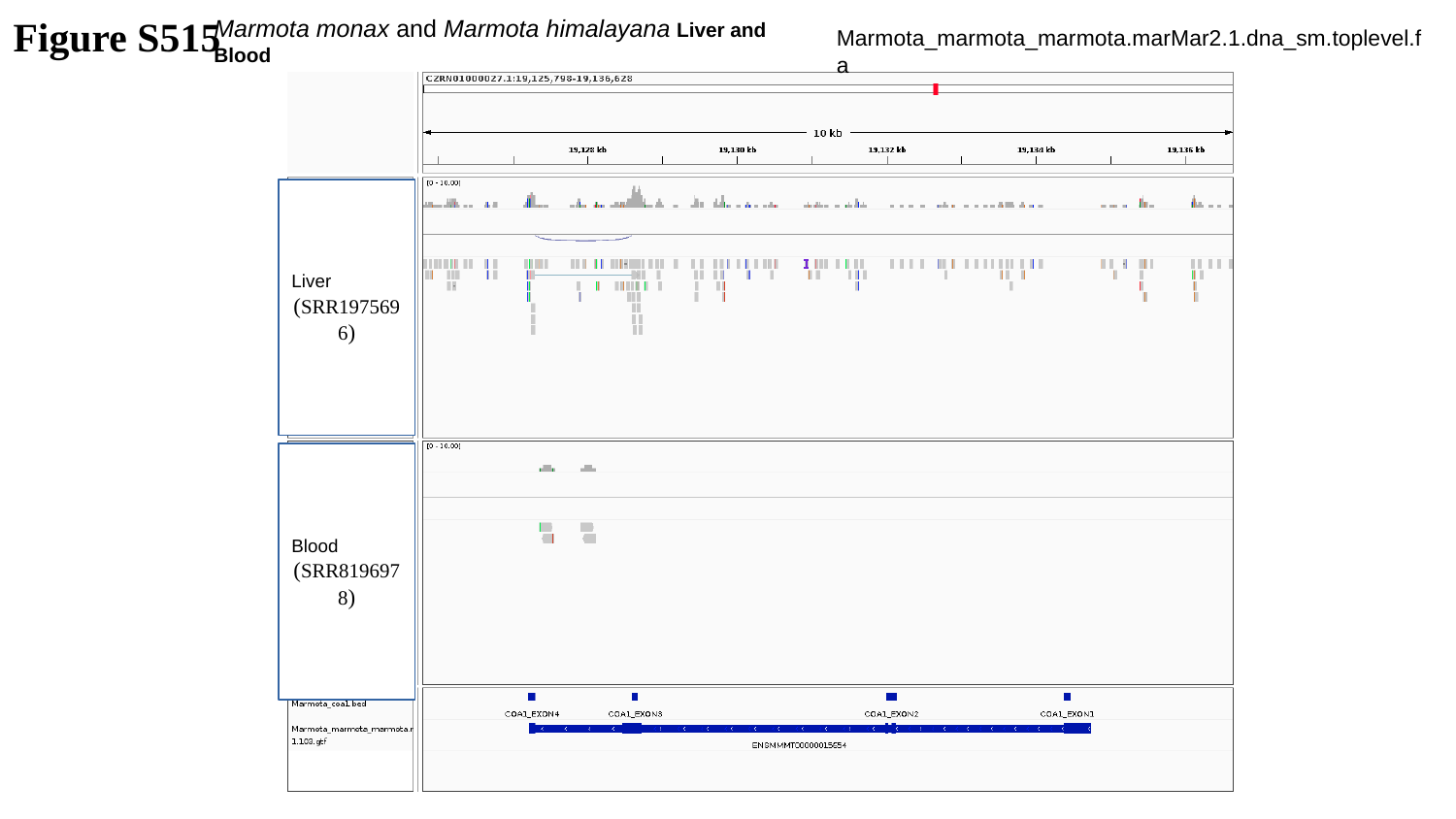

Marmota monax and Marmota himalayana Liver and Blood
Figure S515
Marmota_marmota_marmota.marMar2.1.dna_sm.toplevel.fa
Liver
(SRR1975696)
Blood
(SRR8196978)

### Slide 516
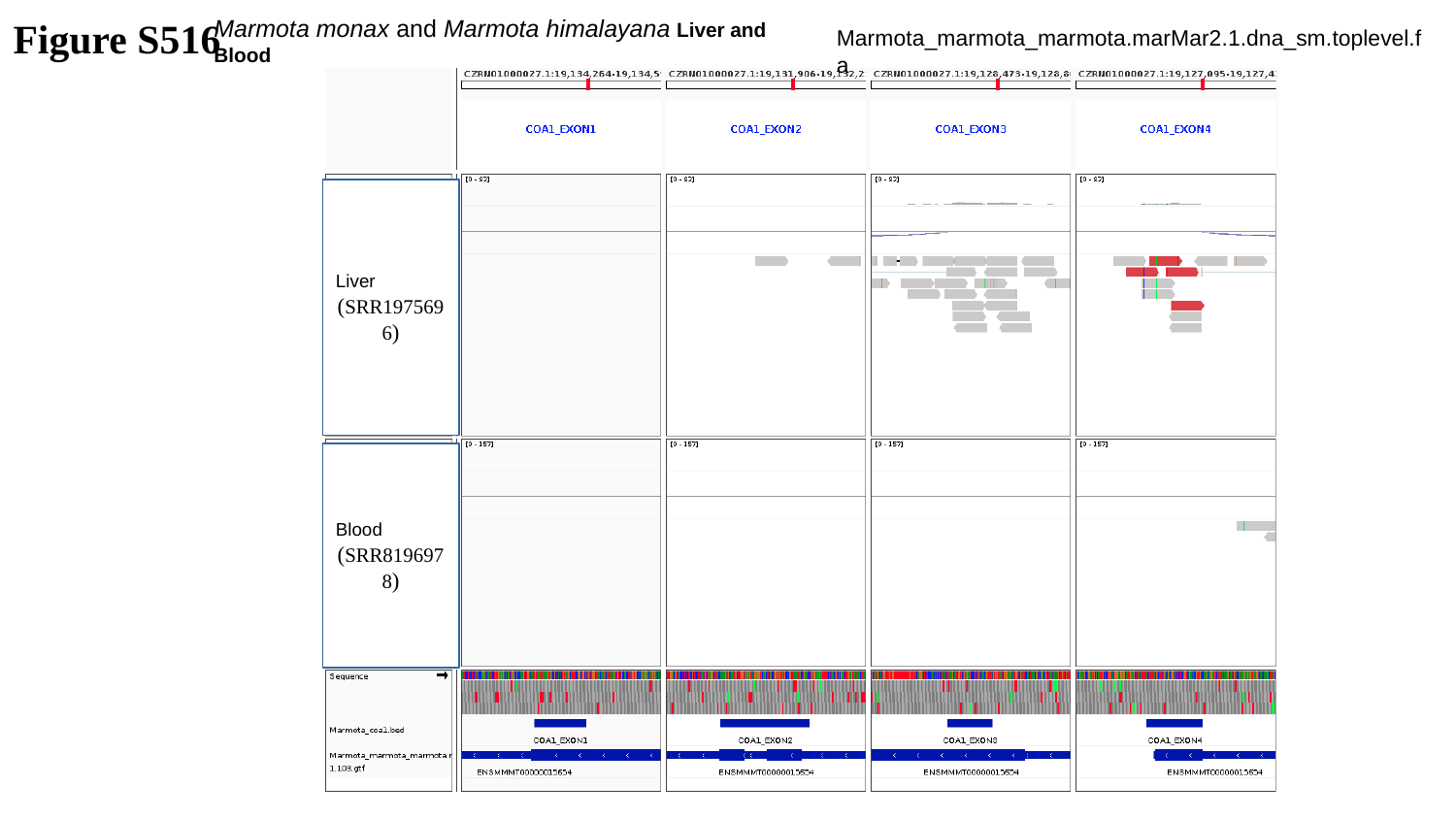

Figure S516
Marmota monax and Marmota himalayana Liver and Blood
Marmota_marmota_marmota.marMar2.1.dna_sm.toplevel.fa
Liver
(SRR1975696)
Blood
(SRR8196978)

### Slide 517
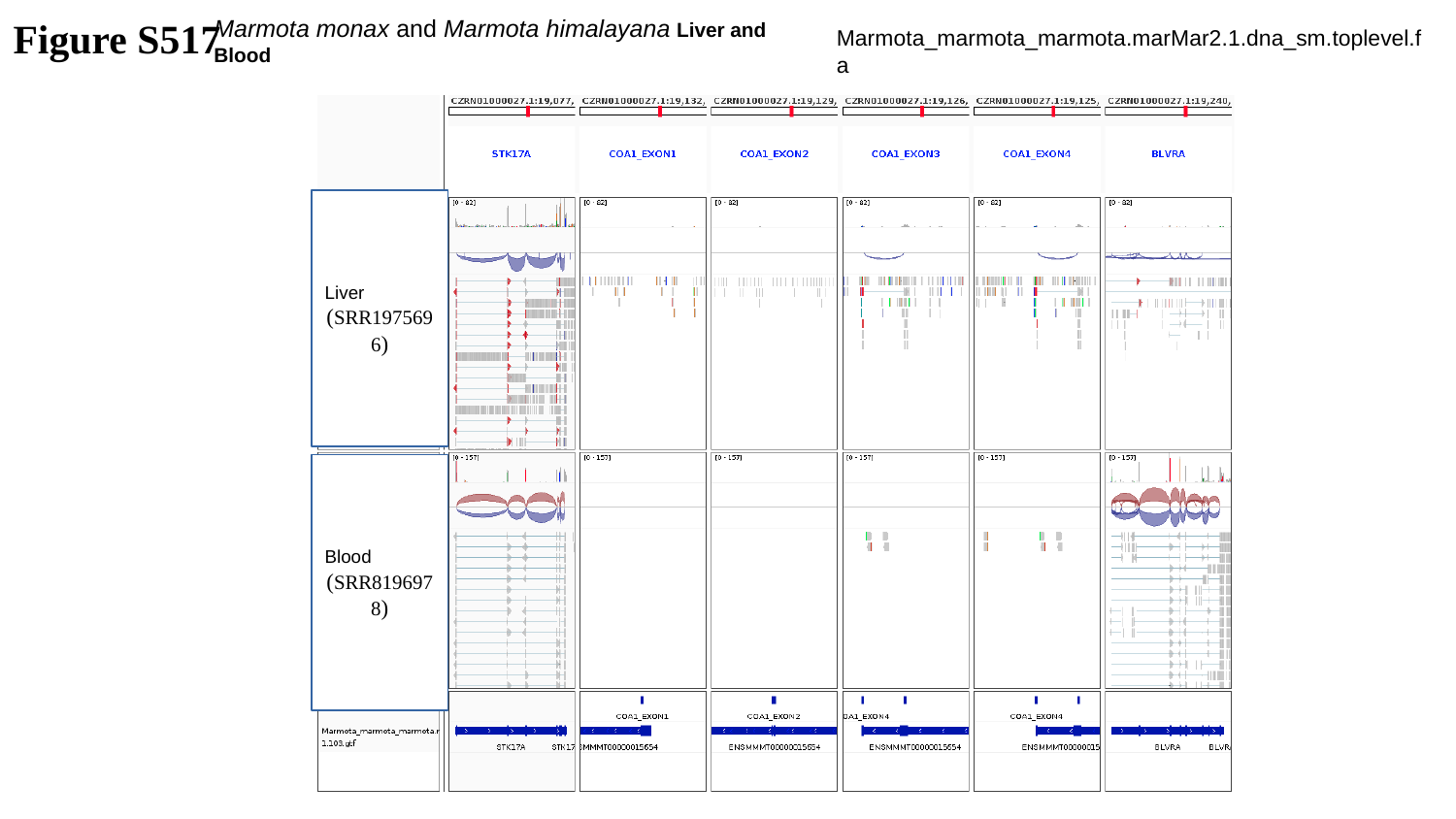

Figure S517
Marmota monax and Marmota himalayana Liver and Blood
Marmota_marmota_marmota.marMar2.1.dna_sm.toplevel.fa
Liver
(SRR1975696)
Blood
(SRR8196978)

### Slide 518
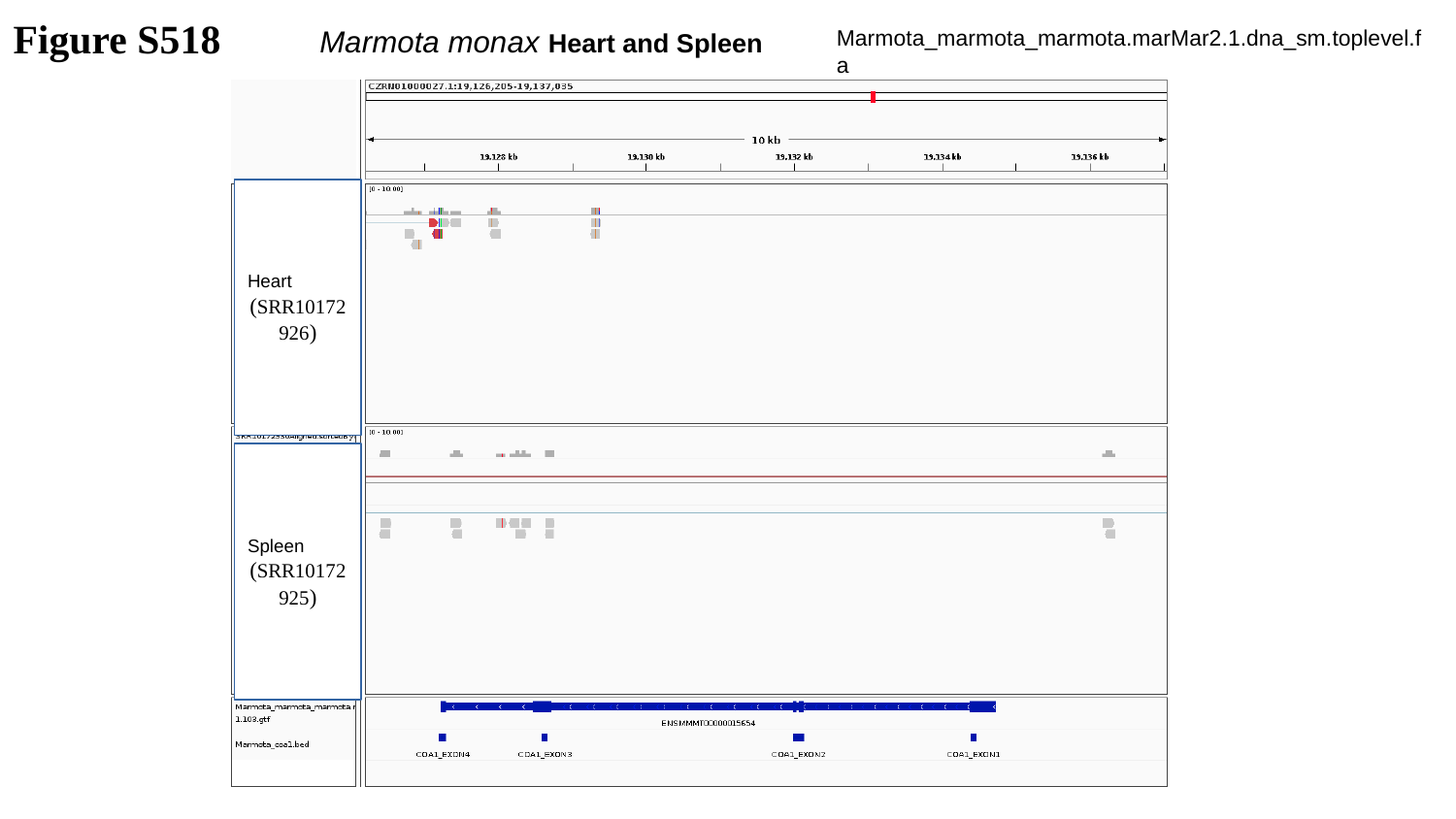

Figure S518
Marmota monax Heart and Spleen
Marmota_marmota_marmota.marMar2.1.dna_sm.toplevel.fa
Heart
(SRR10172926)
Spleen
(SRR10172925)

### Slide 519
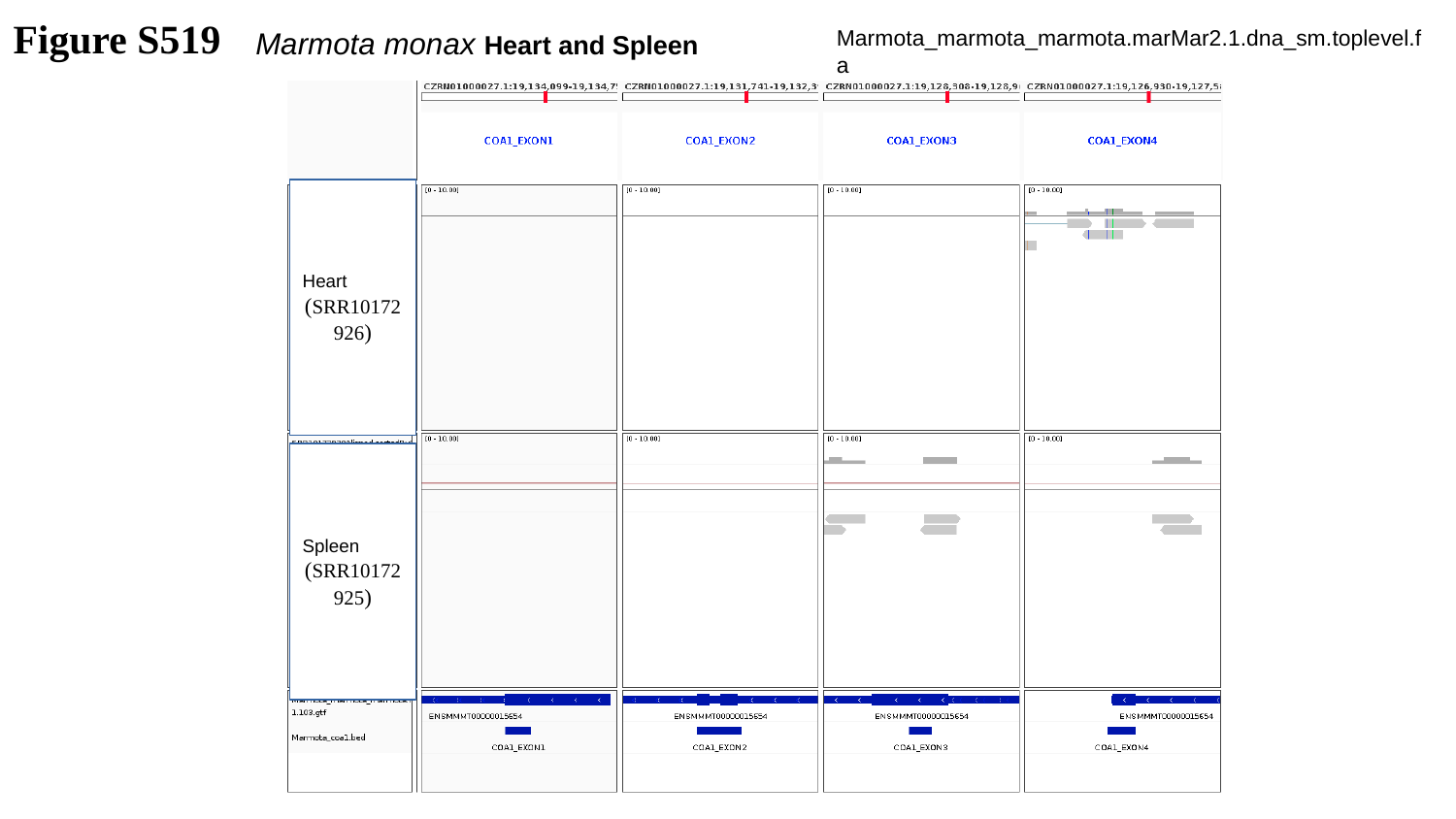

Figure S519
Marmota monax Heart and Spleen
Marmota_marmota_marmota.marMar2.1.dna_sm.toplevel.fa
Heart
(SRR10172926)
Spleen
(SRR10172925)

### Slide 520
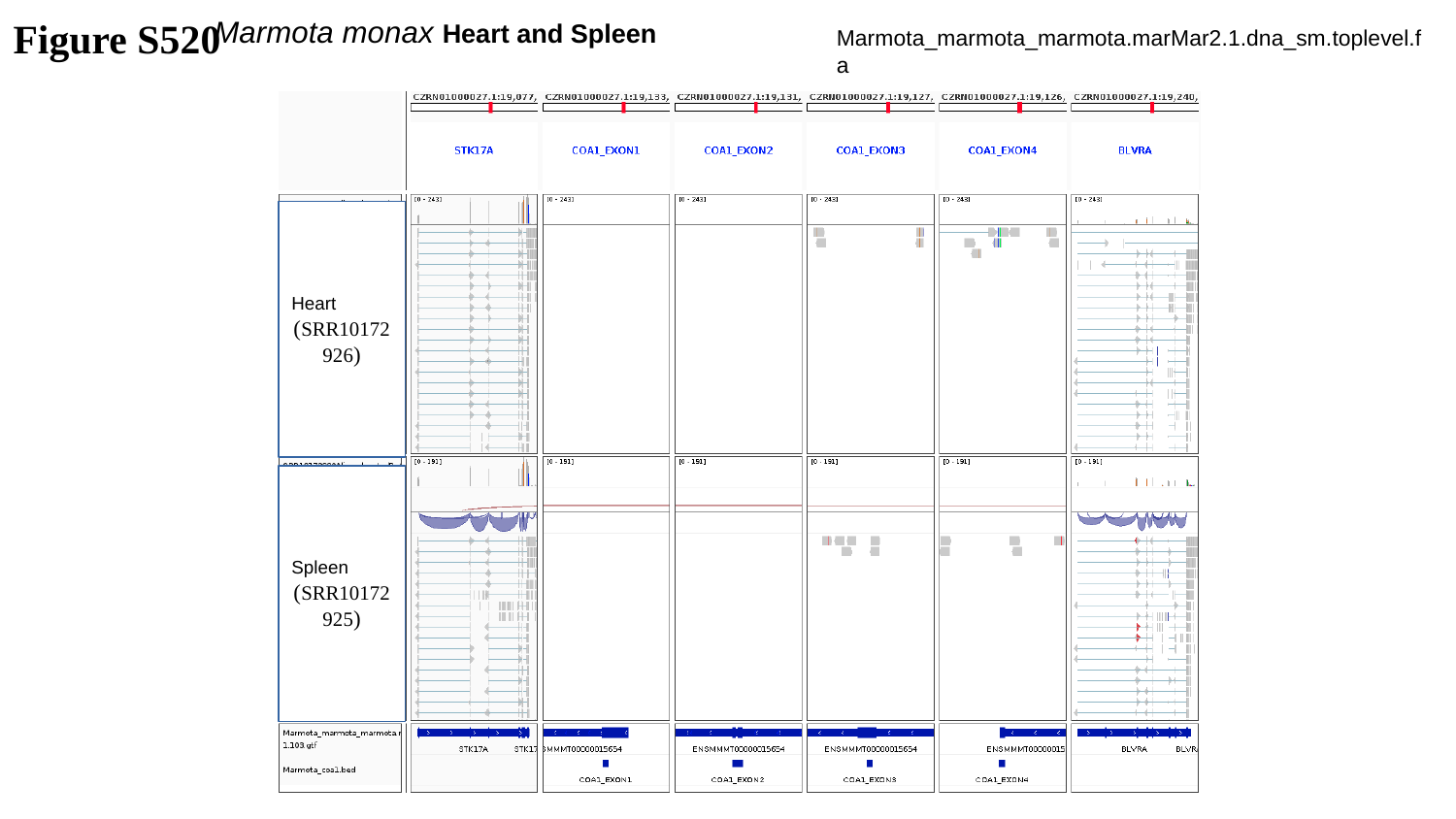

Figure S520
Marmota monax Heart and Spleen
Marmota_marmota_marmota.marMar2.1.dna_sm.toplevel.fa
Heart
(SRR10172926)
Spleen
(SRR10172925)

### Slide 521
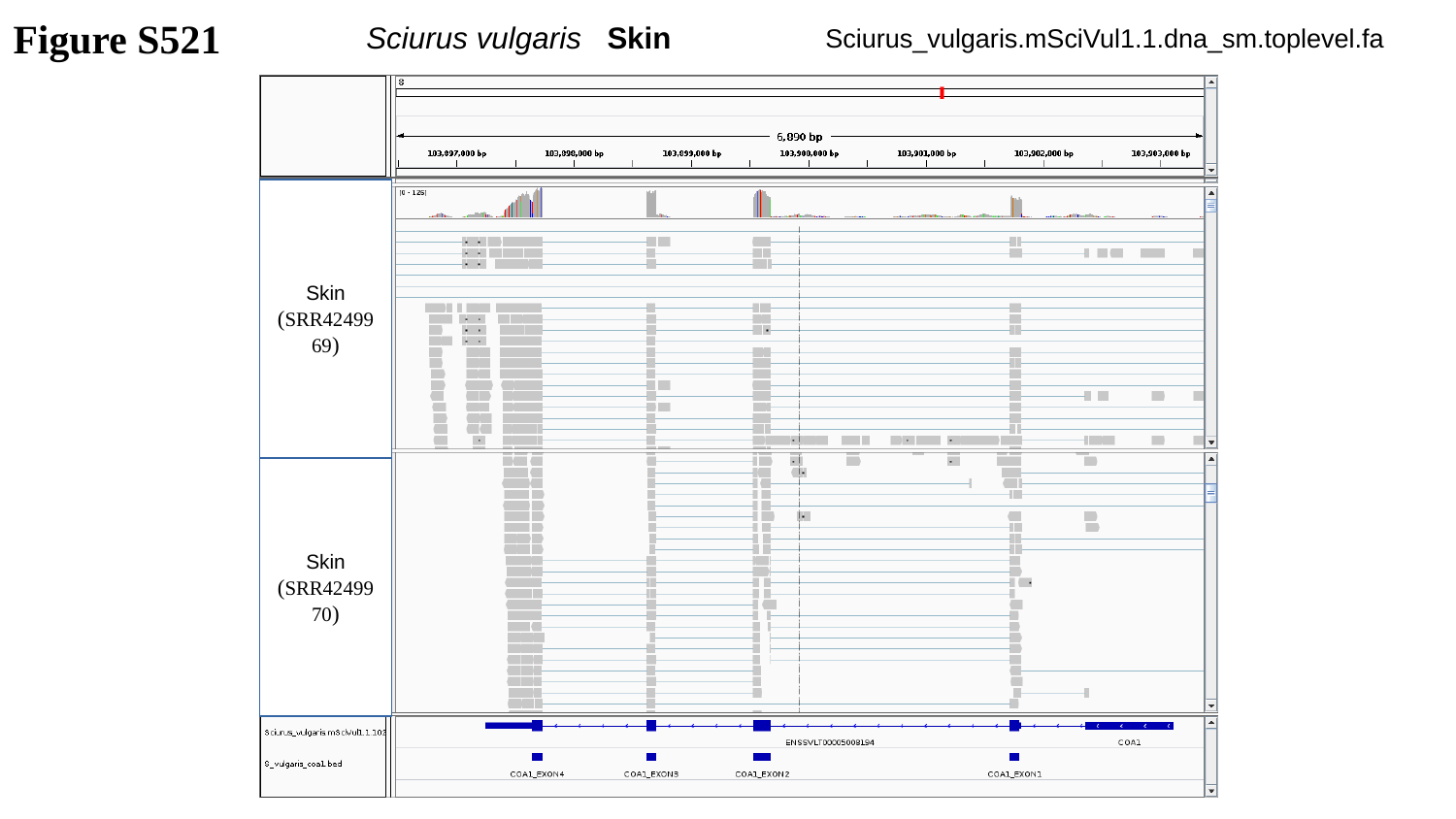

Figure S521
Sciurus vulgaris Skin
Sciurus_vulgaris.mSciVul1.1.dna_sm.toplevel.fa
Skin
(SRR4249969)
Skin
(SRR4249970)

### Slide 522
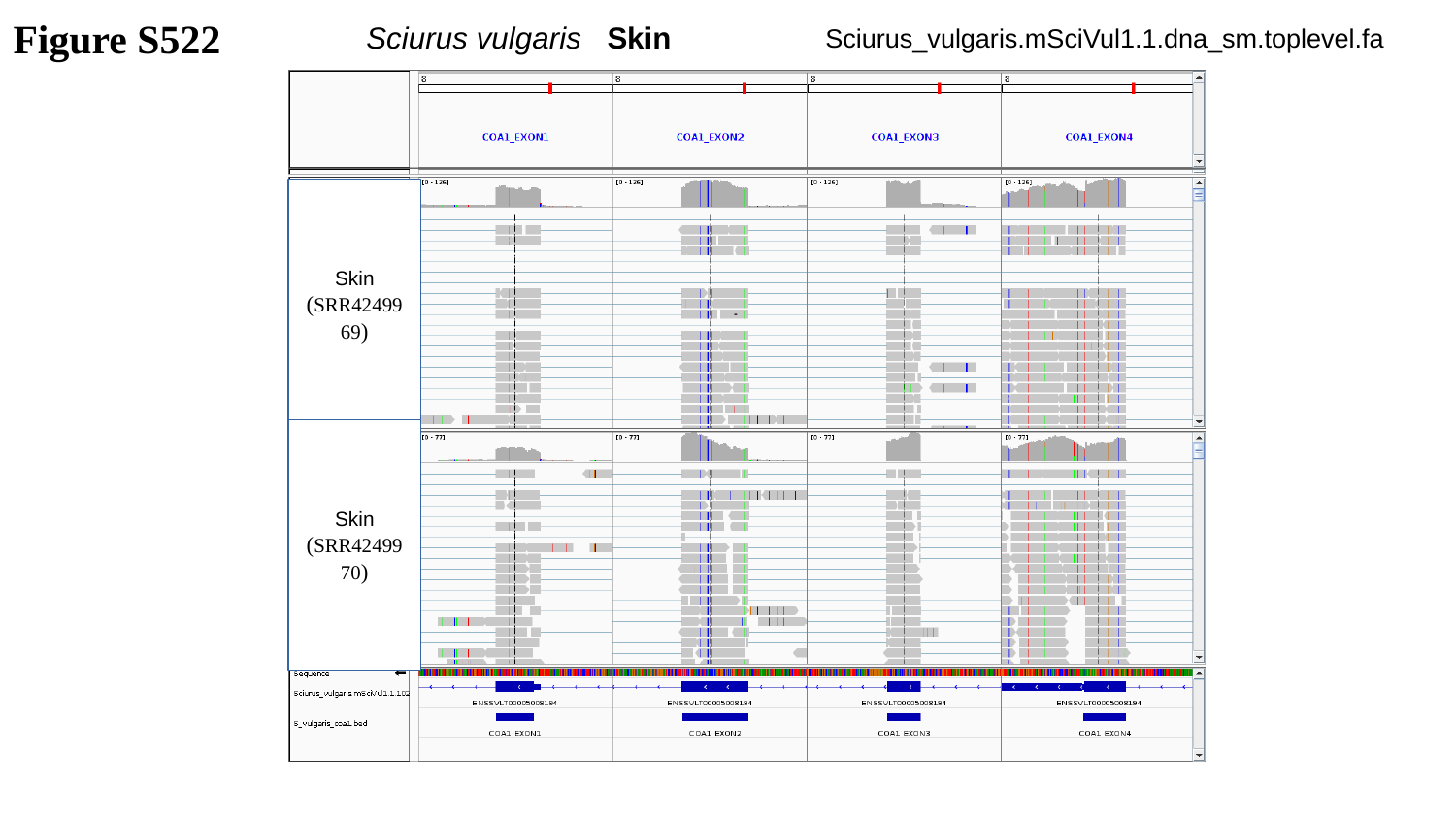

Figure S522
Sciurus vulgaris Skin
Sciurus_vulgaris.mSciVul1.1.dna_sm.toplevel.fa
Skin
(SRR4249969)
Skin
(SRR4249970)

### Slide 523
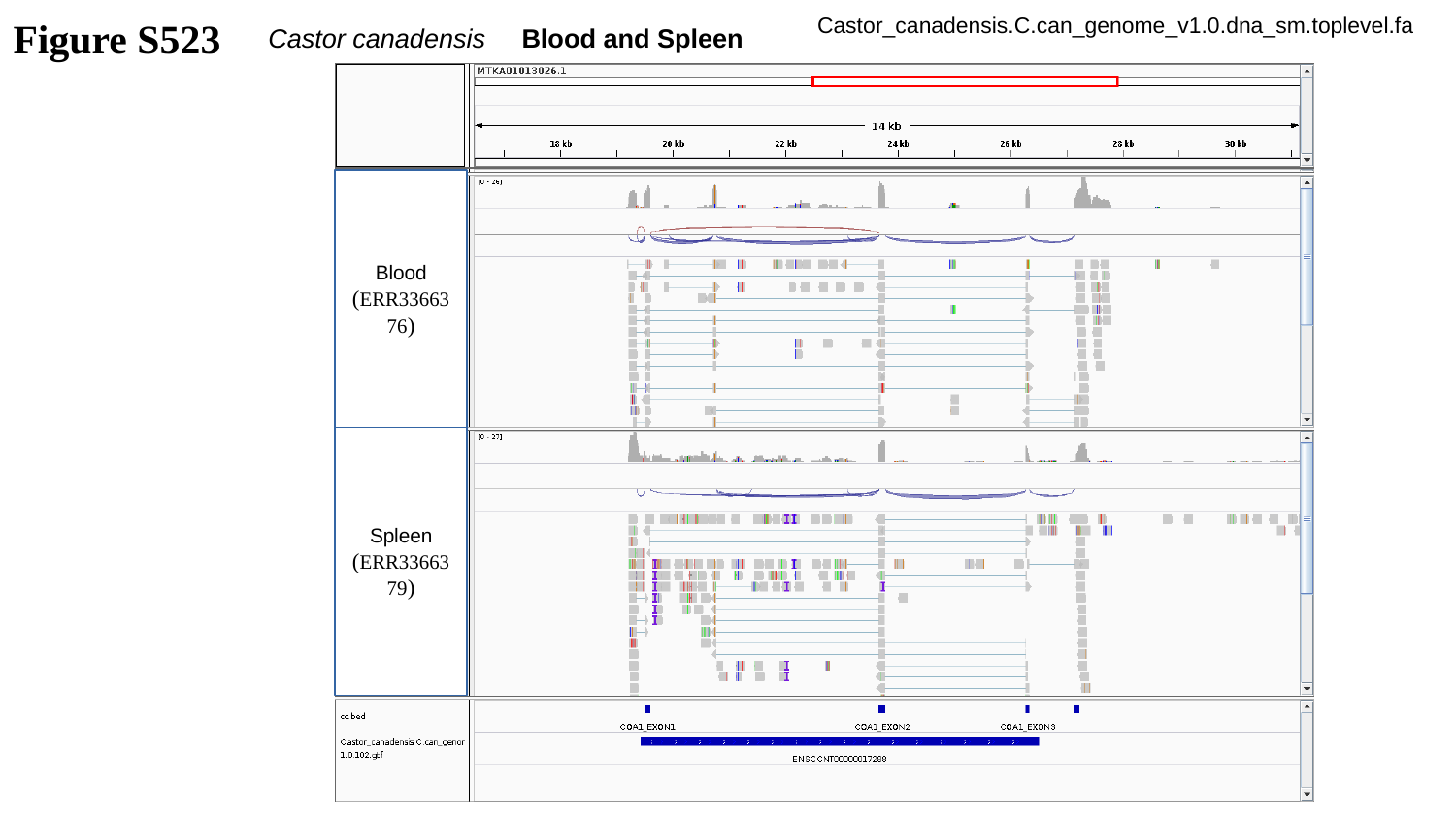

Figure S523
Castor_canadensis.C.can_genome_v1.0.dna_sm.toplevel.fa
Castor canadensis Blood and Spleen
Blood
(ERR3366376)
Spleen
(ERR3366379)

### Slide 524
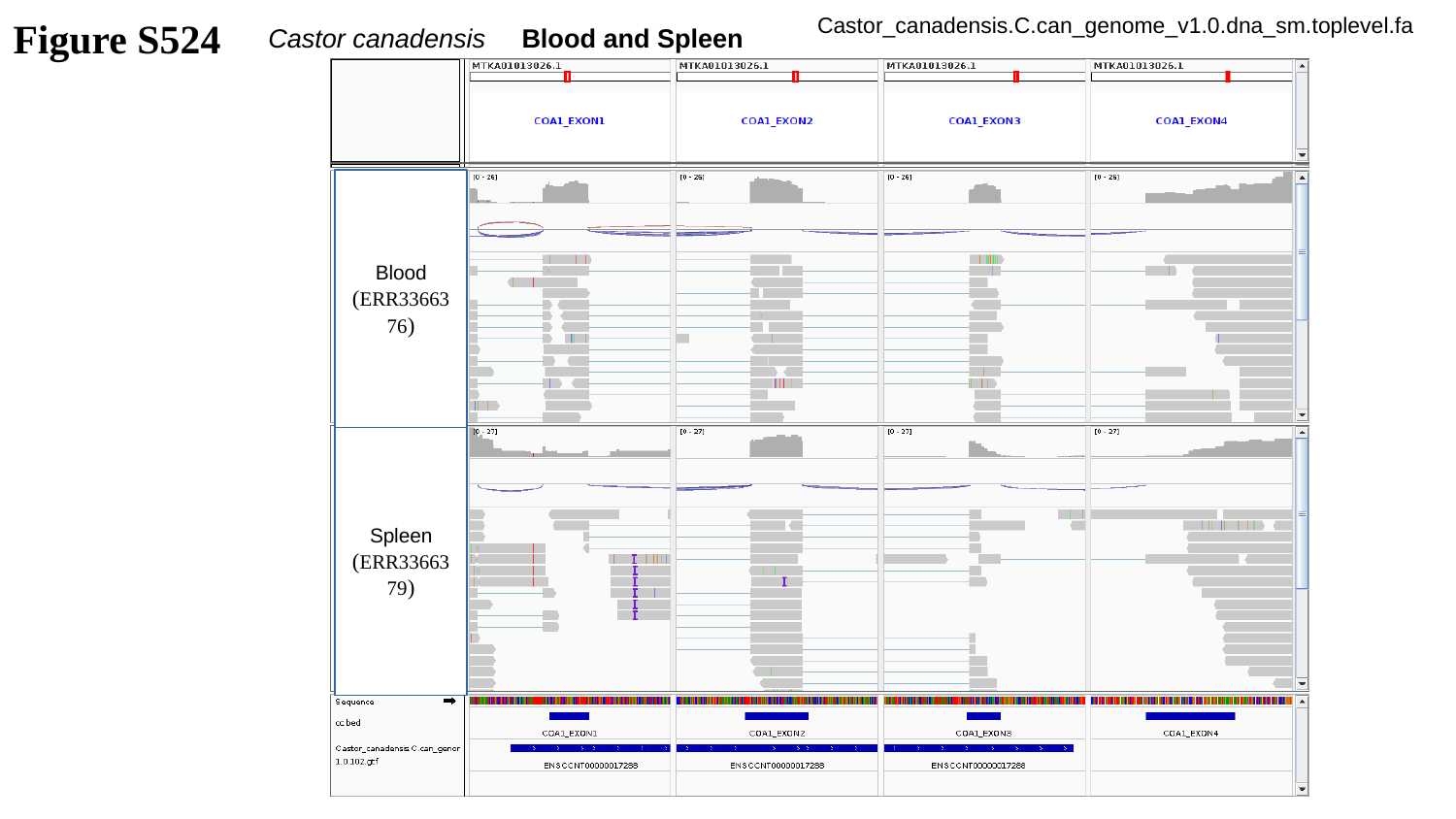

Figure S524
Castor_canadensis.C.can_genome_v1.0.dna_sm.toplevel.fa
Castor canadensis Blood and Spleen
Blood
(ERR3366376)
Spleen
(ERR3366379)

### Slide 525
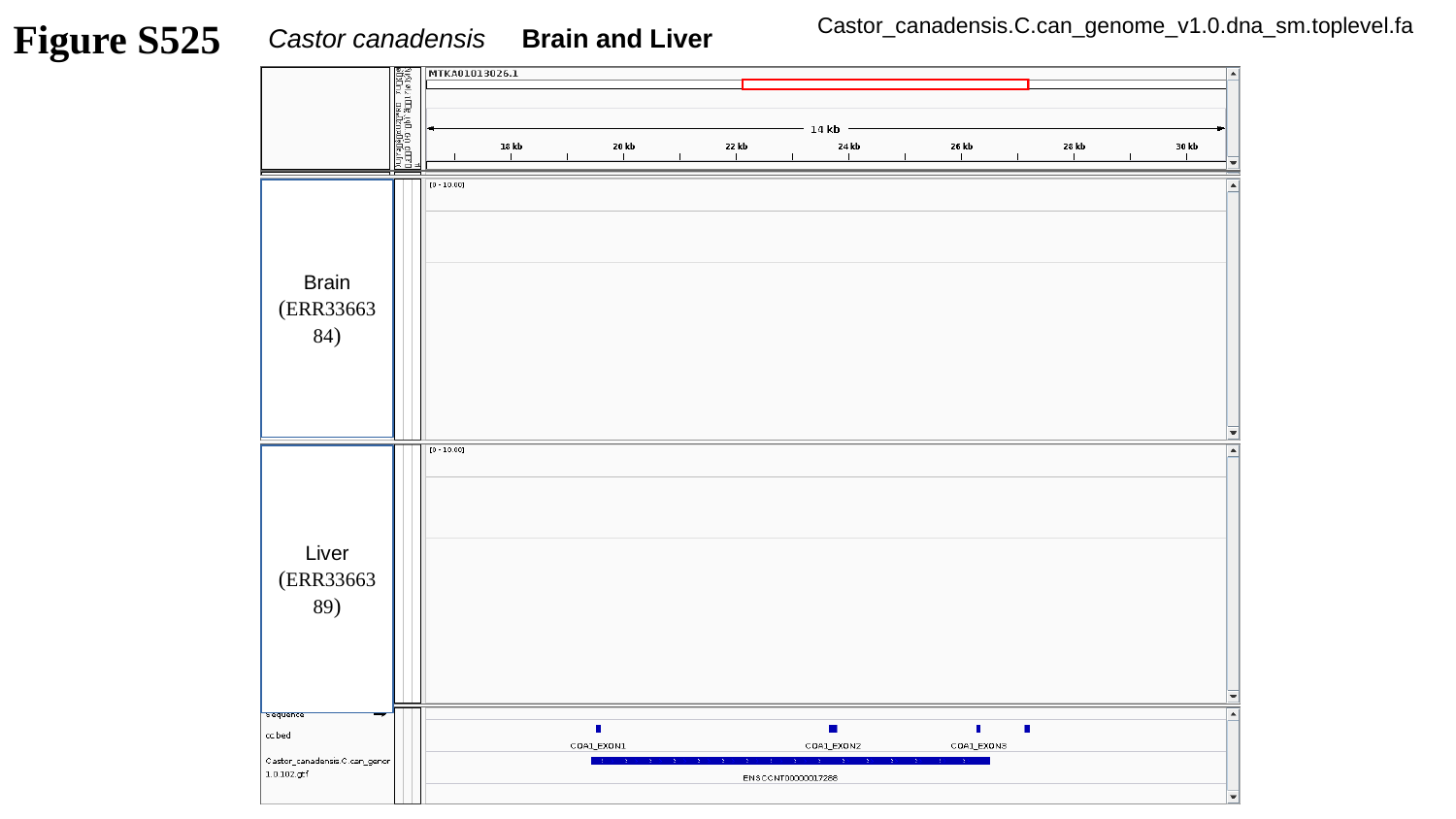

Figure S525
Castor_canadensis.C.can_genome_v1.0.dna_sm.toplevel.fa
Castor canadensis Brain and Liver
Brain
(ERR3366384)
Liver
(ERR3366389)

### Slide 526
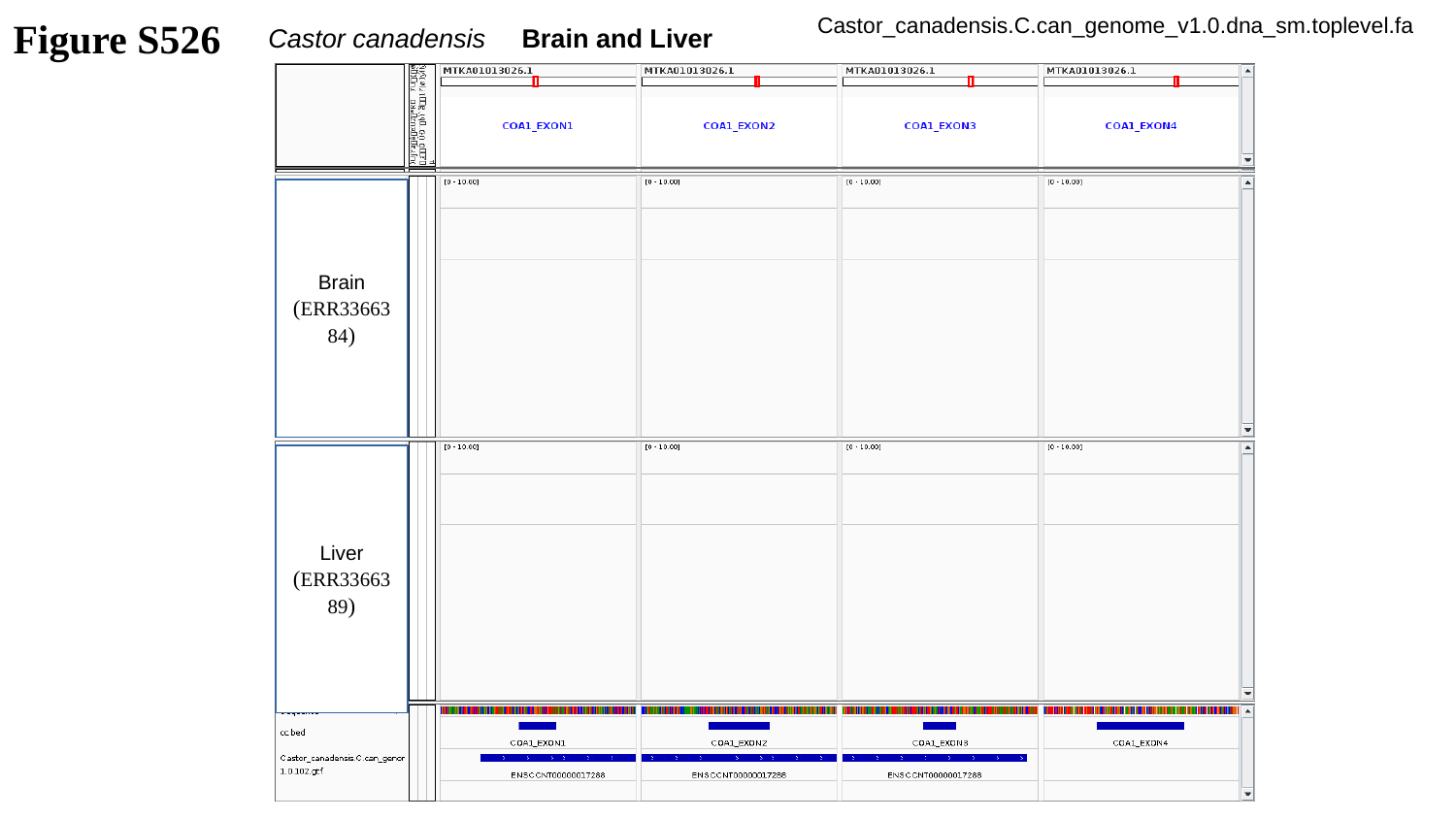

Figure S526
Castor_canadensis.C.can_genome_v1.0.dna_sm.toplevel.fa
Castor canadensis Brain and Liver
Brain
(ERR3366384)
Liver
(ERR3366389)

### Slide 527
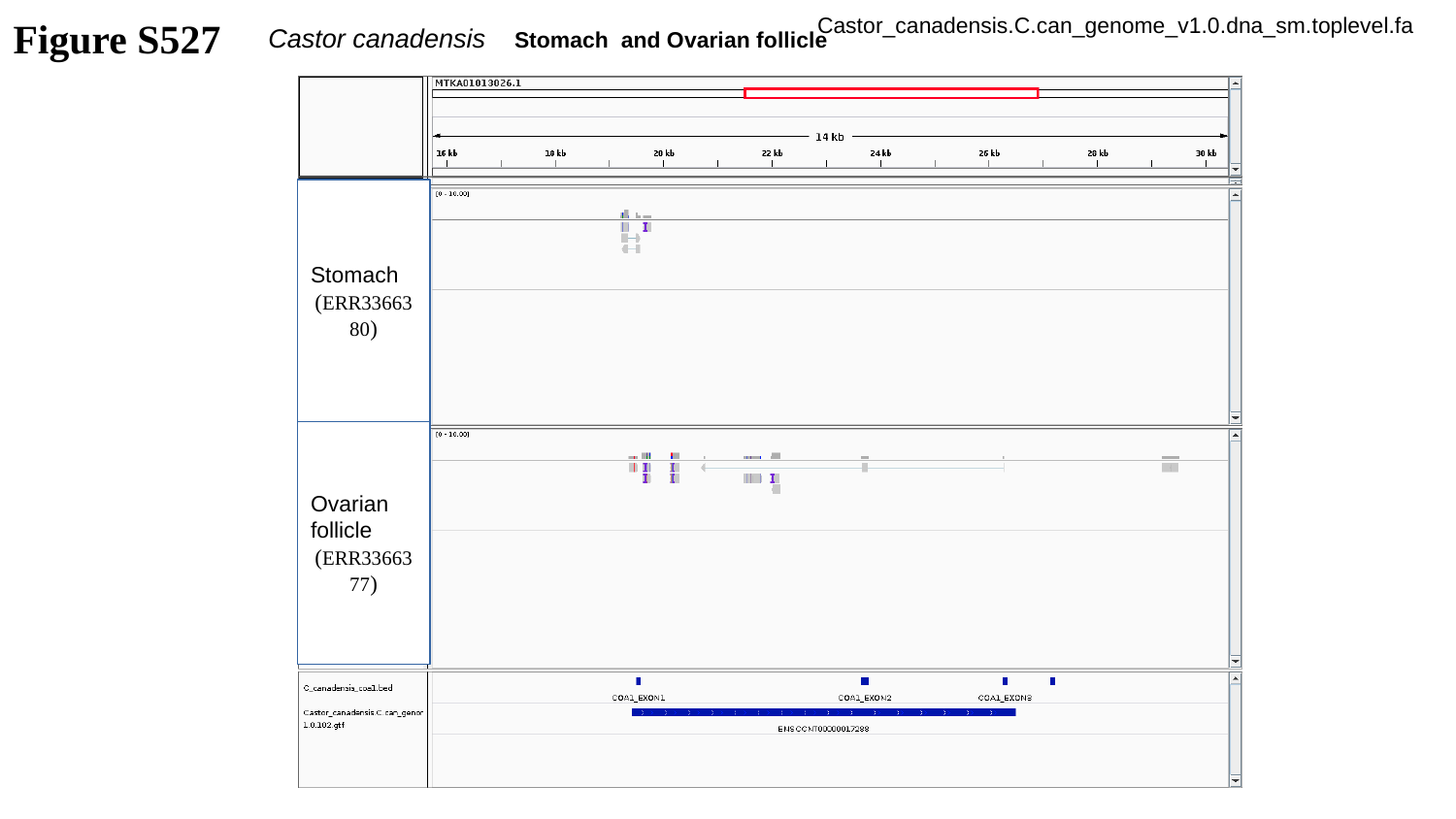

Figure S527
Castor_canadensis.C.can_genome_v1.0.dna_sm.toplevel.fa
Castor canadensis Stomach and Ovarian follicle
Stomach
(ERR3366380)
Ovarian follicle
(ERR3366377)

### Slide 528
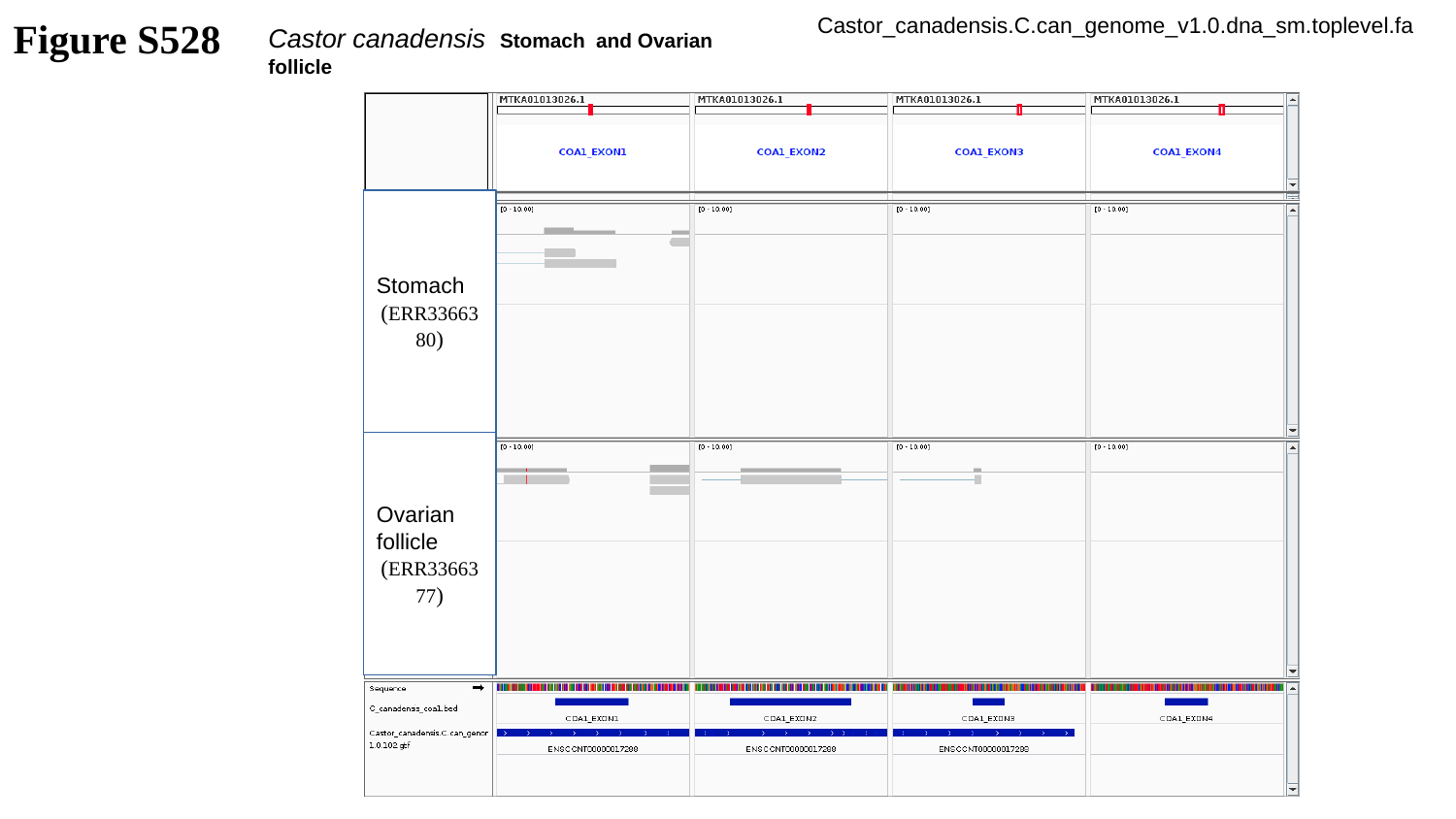

Figure S528
Castor_canadensis.C.can_genome_v1.0.dna_sm.toplevel.fa
Castor canadensis Stomach and Ovarian follicle
Stomach
(ERR3366380)
Ovarian follicle
(ERR3366377)

### Slide 529
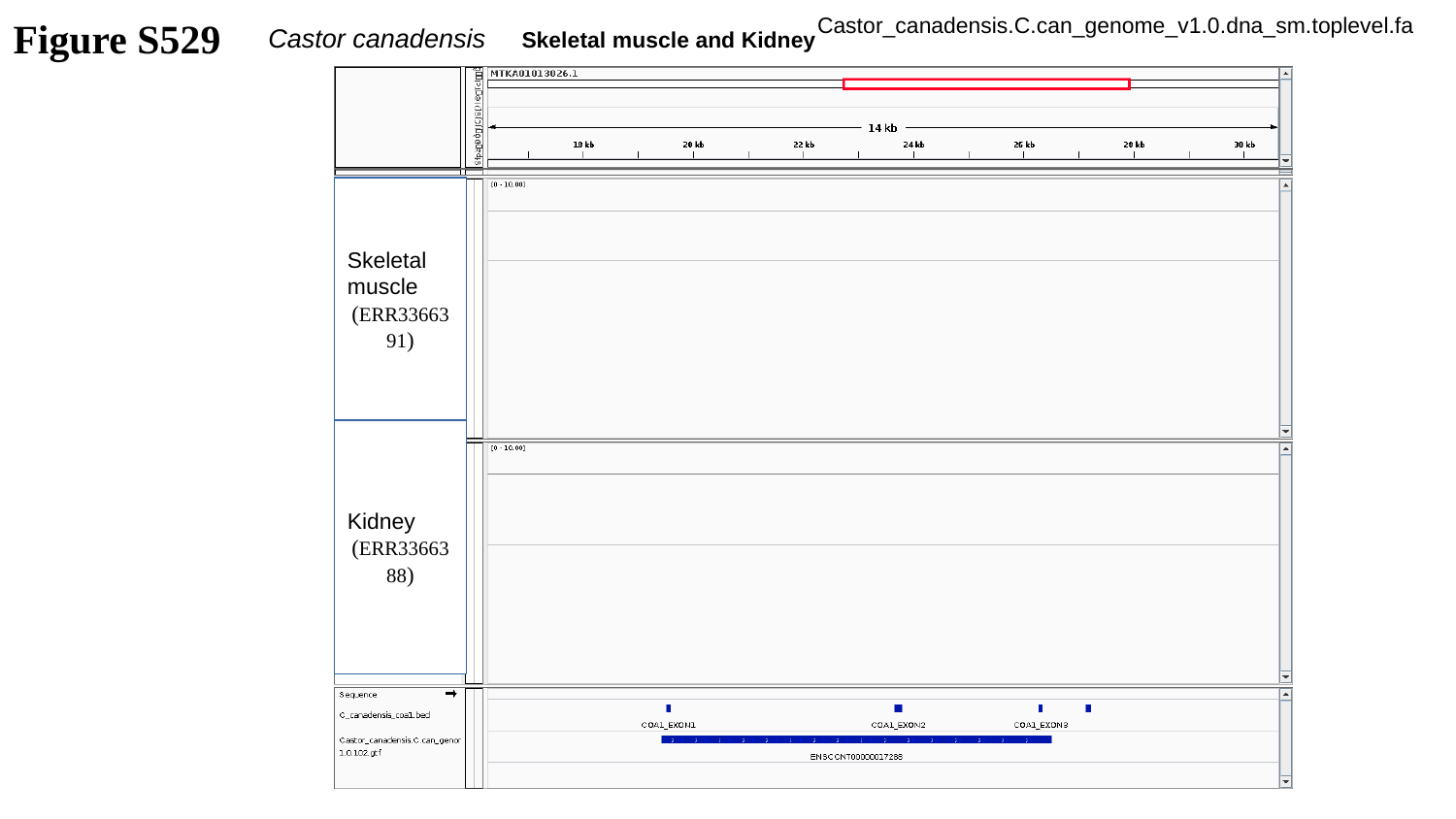

Figure S529
Castor_canadensis.C.can_genome_v1.0.dna_sm.toplevel.fa
Castor canadensis Skeletal muscle and Kidney
Skeletal muscle
(ERR3366391)
Kidney
(ERR3366388)

### Slide 530
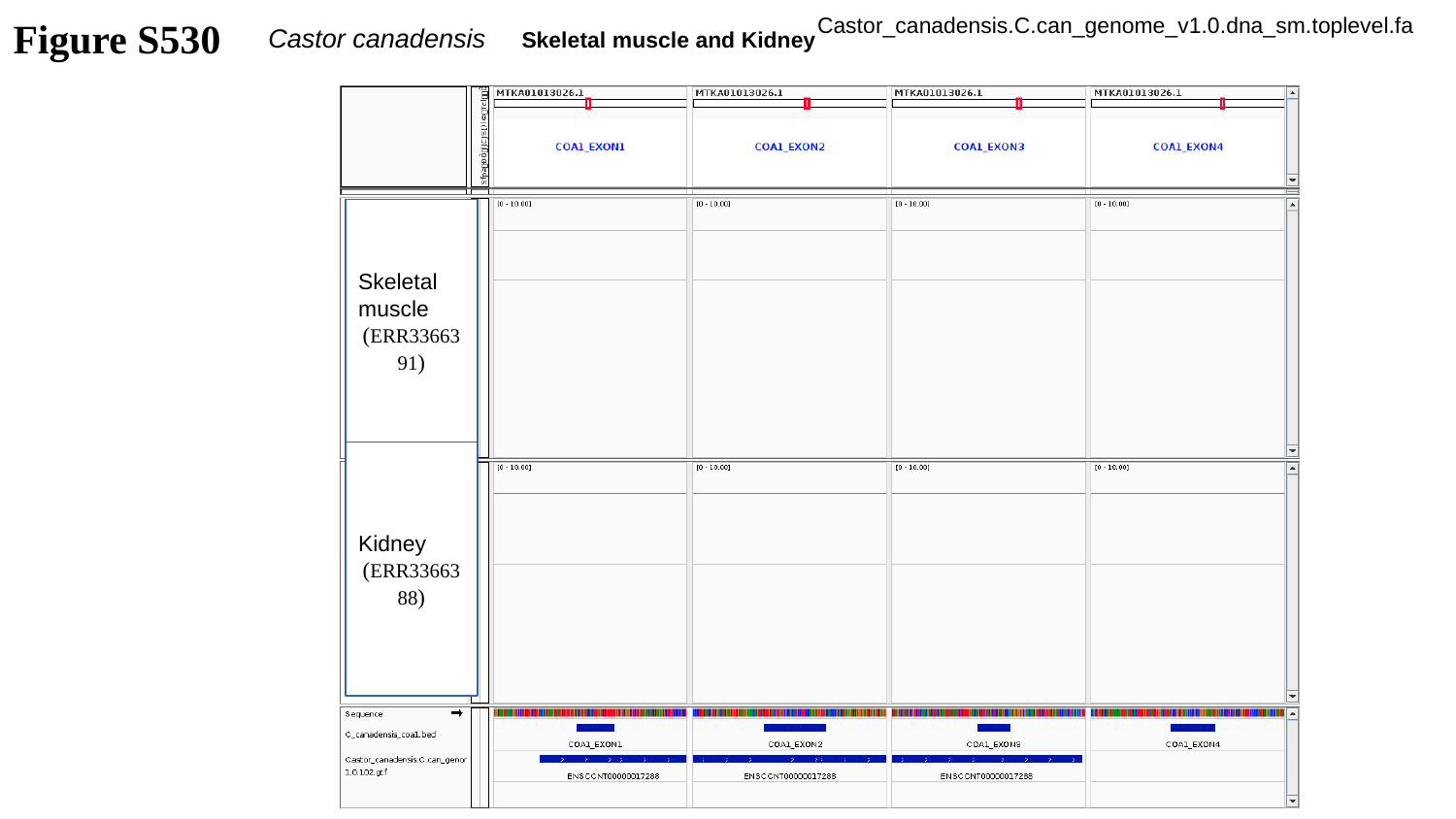

Figure S530
Castor_canadensis.C.can_genome_v1.0.dna_sm.toplevel.fa
Castor canadensis Skeletal muscle and Kidney
Skeletal muscle
(ERR3366391)
Kidney
(ERR3366388)

### Slide 531
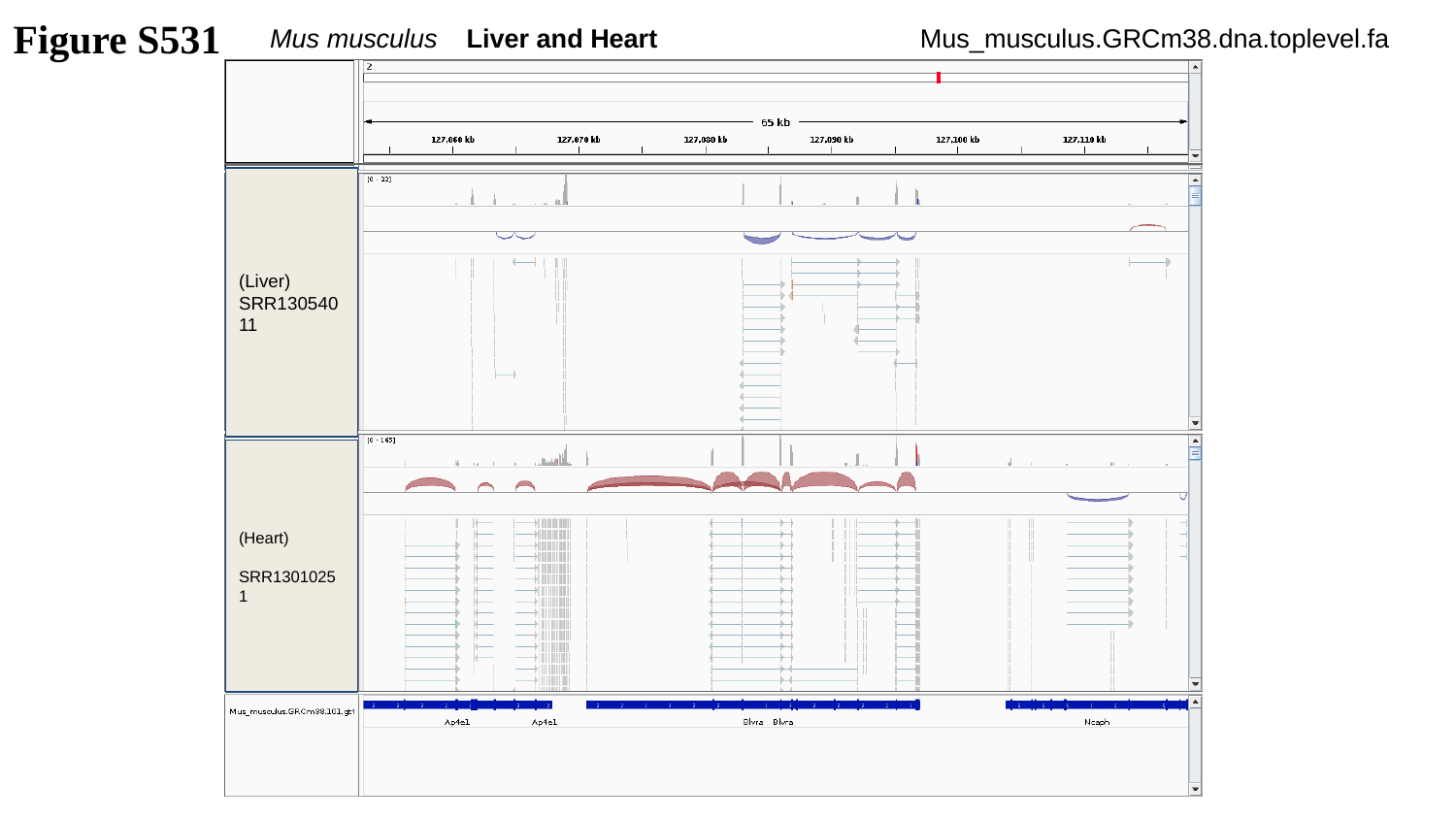

Figure S531
Mus musculus Liver and Heart
Mus_musculus.GRCm38.dna.toplevel.fa
(Liver)
SRR13054011
(Heart)
SRR13010251

### Slide 532
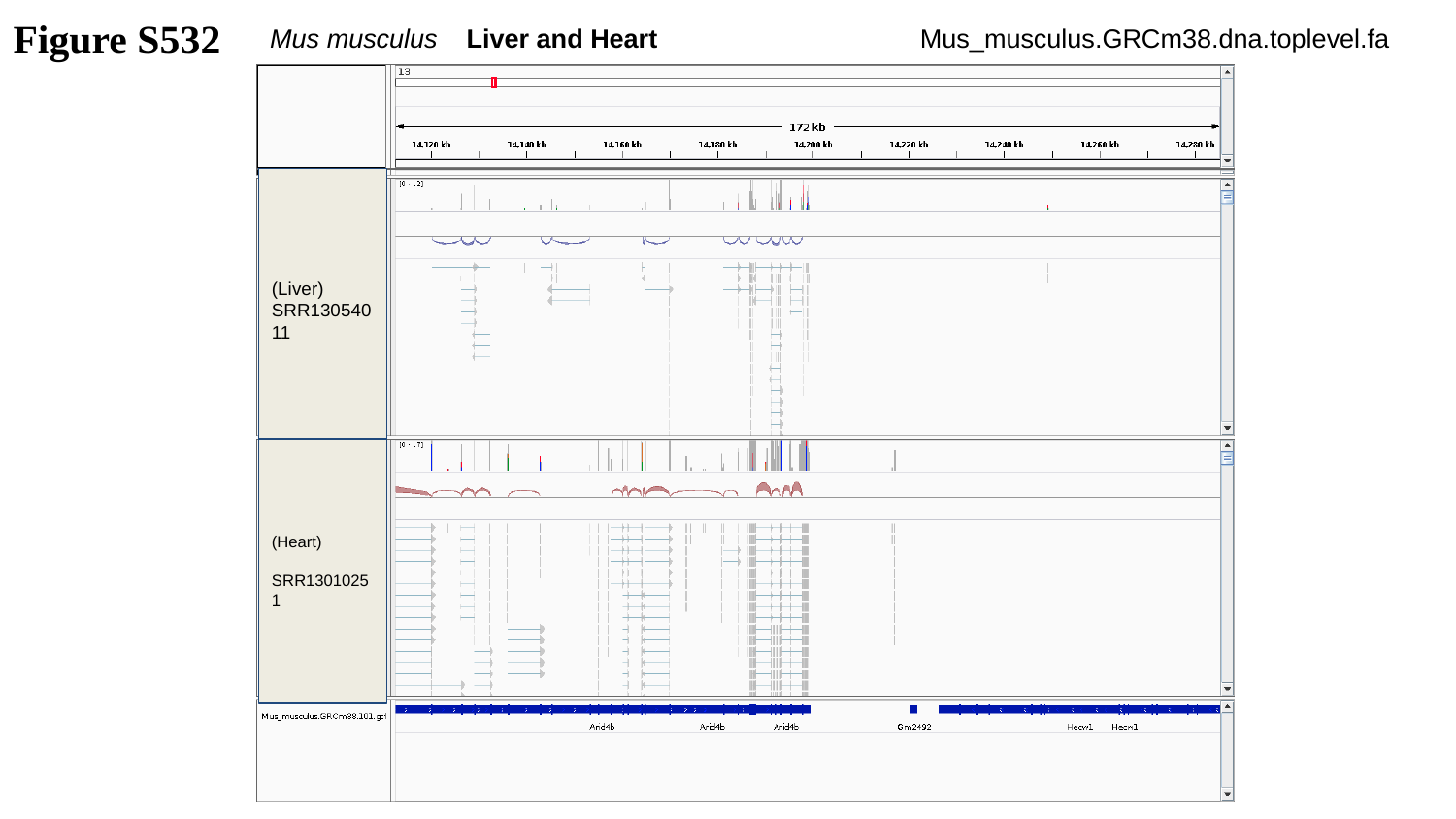

Figure S532
Mus musculus Liver and Heart
Mus_musculus.GRCm38.dna.toplevel.fa
(Liver)
SRR13054011
(Heart)
SRR13010251

### Slide 533
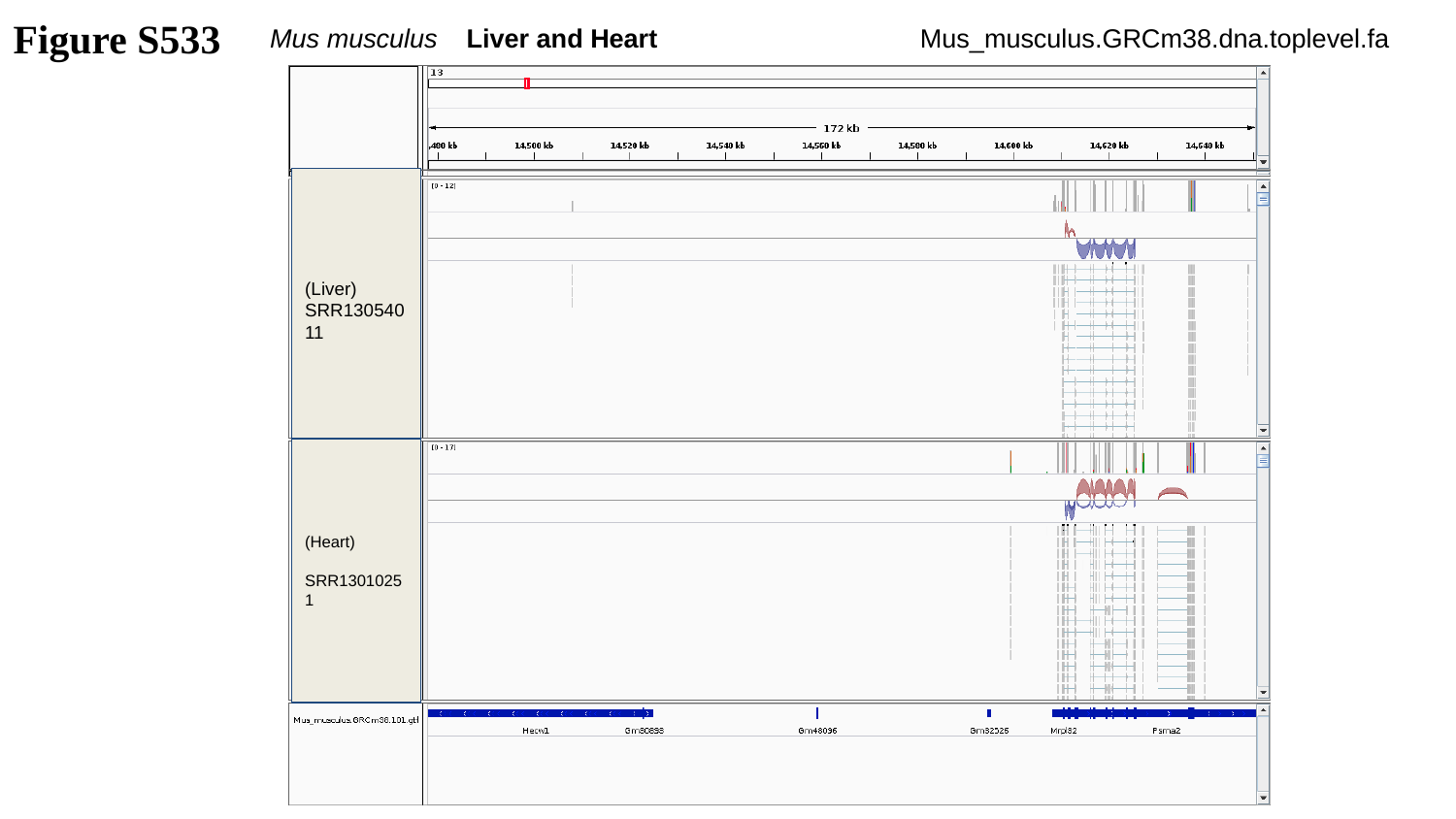

Figure S533
Mus musculus Liver and Heart
Mus_musculus.GRCm38.dna.toplevel.fa
(Liver)
SRR13054011
(Heart)
SRR13010251

### Slide 534
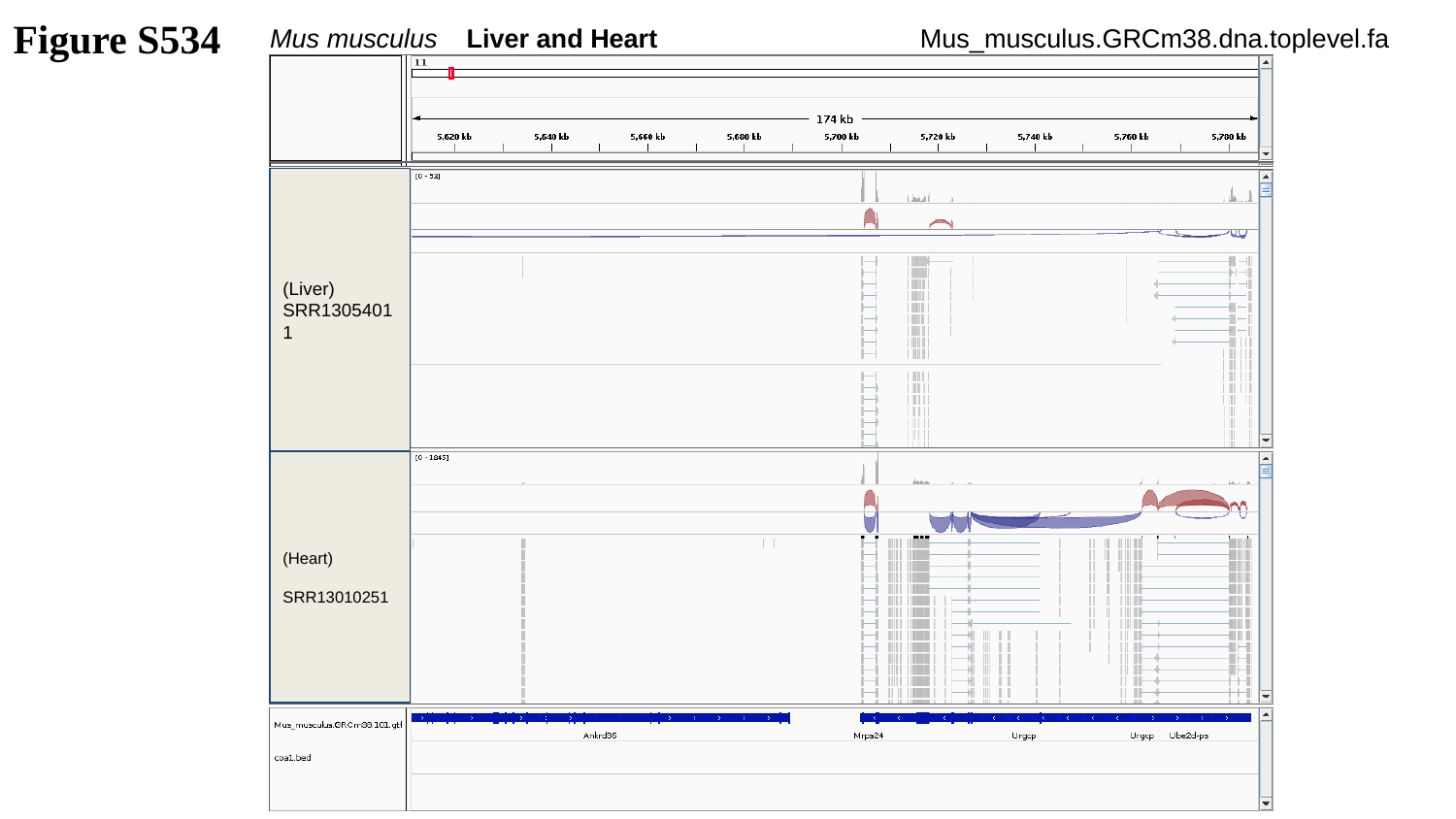

Figure S534
Mus musculus Liver and Heart
Mus_musculus.GRCm38.dna.toplevel.fa
(Liver)
SRR13054011
(Heart)
SRR13010251

### Slide 535
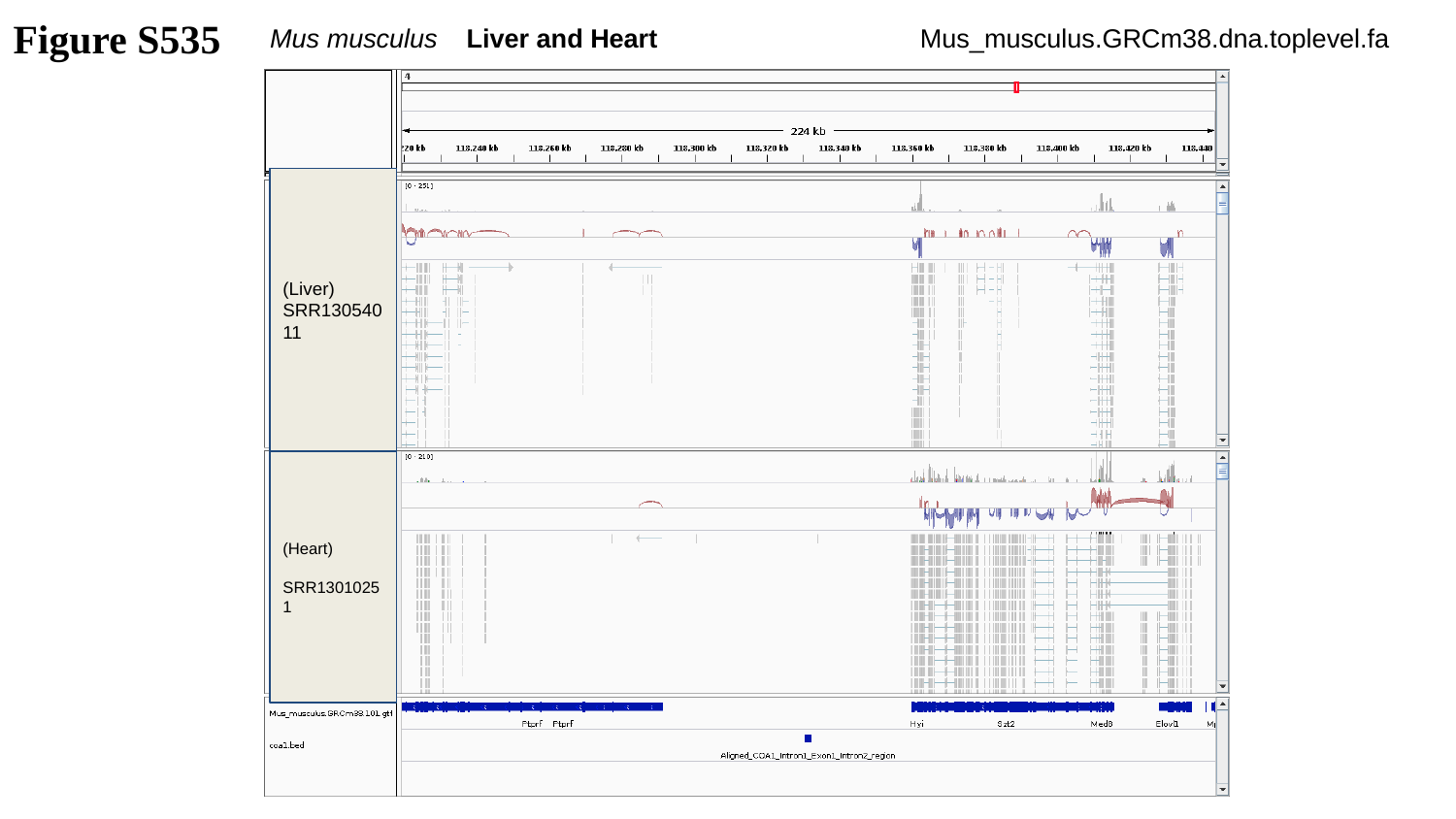

Figure S535
Mus musculus Liver and Heart
Mus_musculus.GRCm38.dna.toplevel.fa
(Liver)
SRR13054011
(Heart)
SRR13010251

### Slide 536
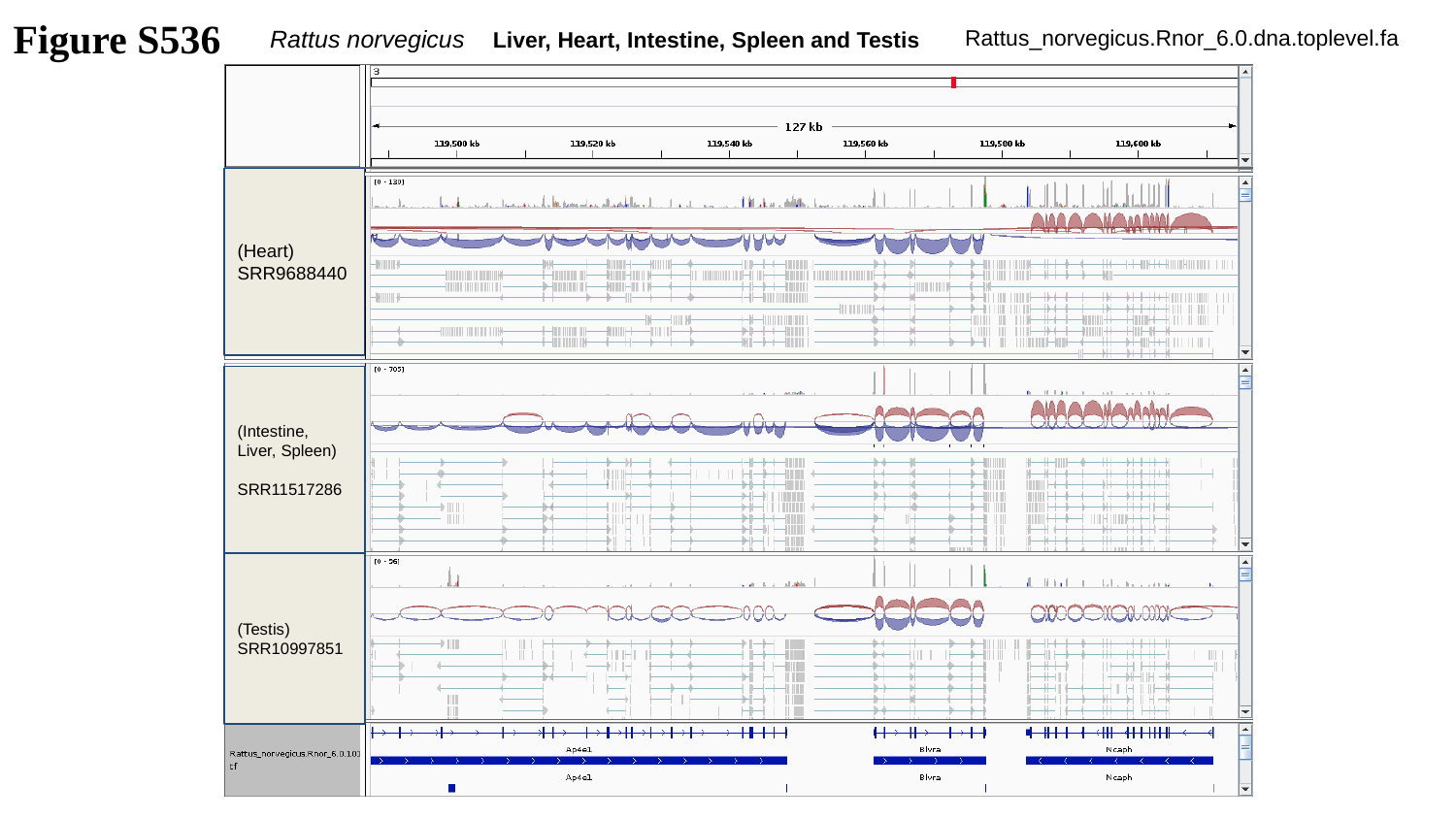

Figure S536
Rattus norvegicus Liver, Heart, Intestine, Spleen and Testis
Rattus_norvegicus.Rnor_6.0.dna.toplevel.fa
(Heart)
SRR9688440
(Intestine, Liver, Spleen)
SRR11517286
(Testis)
SRR10997851

### Slide 537
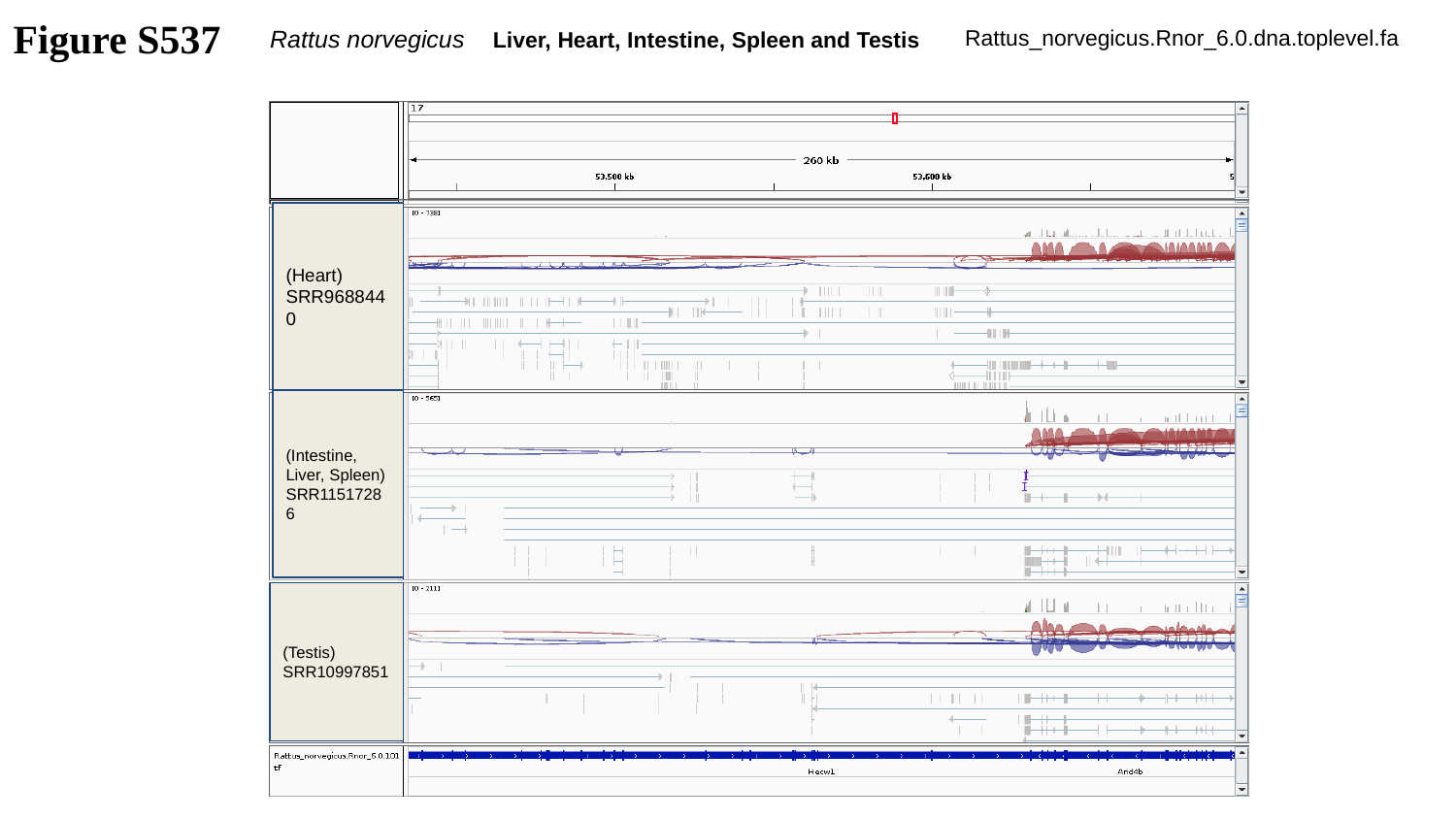

Figure S537
Rattus norvegicus Liver, Heart, Intestine, Spleen and Testis
Rattus_norvegicus.Rnor_6.0.dna.toplevel.fa
(Heart)
SRR9688440
(Intestine, Liver, Spleen)
SRR11517286
(Testis)
SRR10997851

### Slide 538
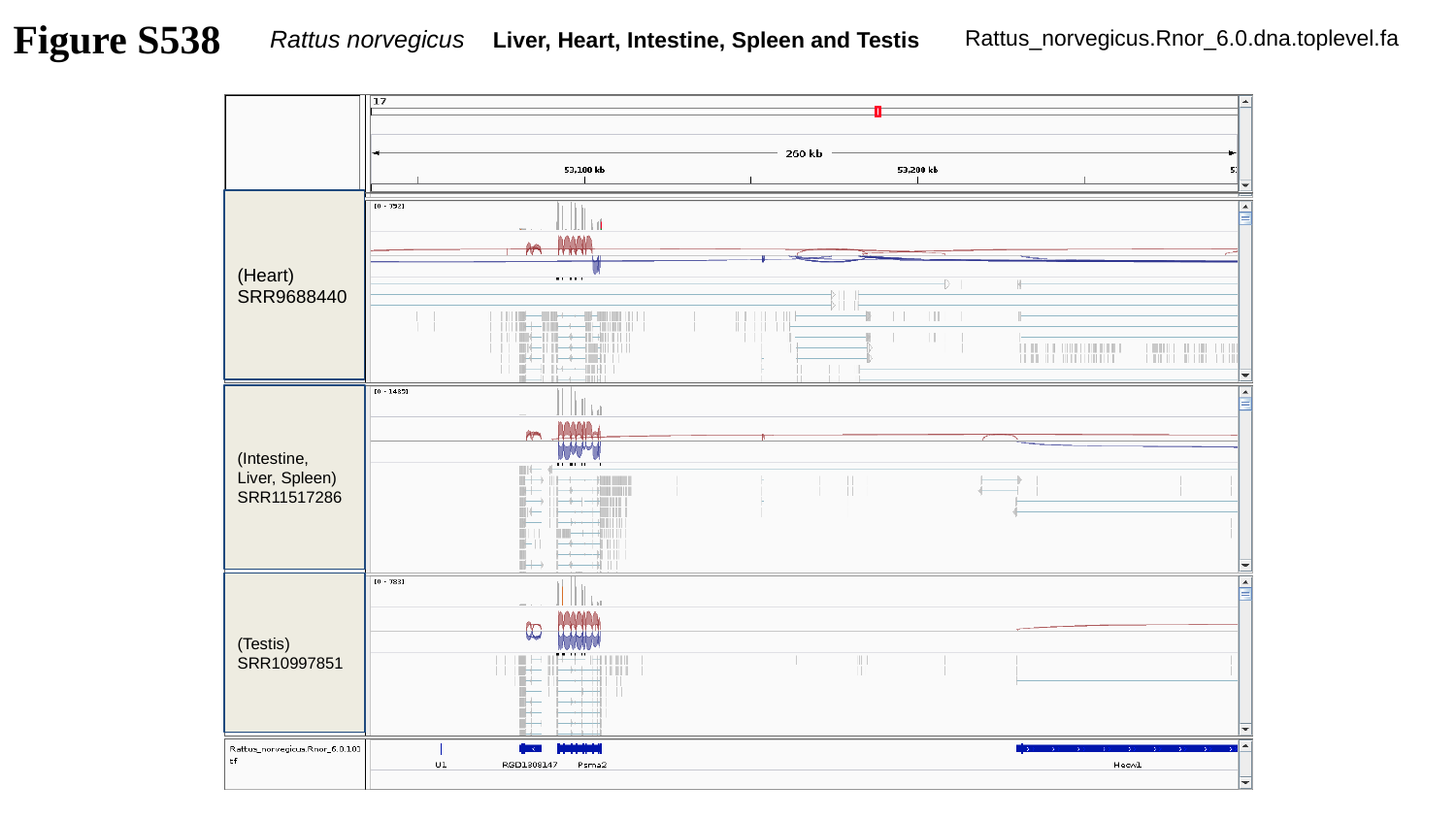

Figure S538
Rattus norvegicus Liver, Heart, Intestine, Spleen and Testis
Rattus_norvegicus.Rnor_6.0.dna.toplevel.fa
(Heart)
SRR9688440
(Intestine, Liver, Spleen)
SRR11517286
(Testis)
SRR10997851

### Slide 539
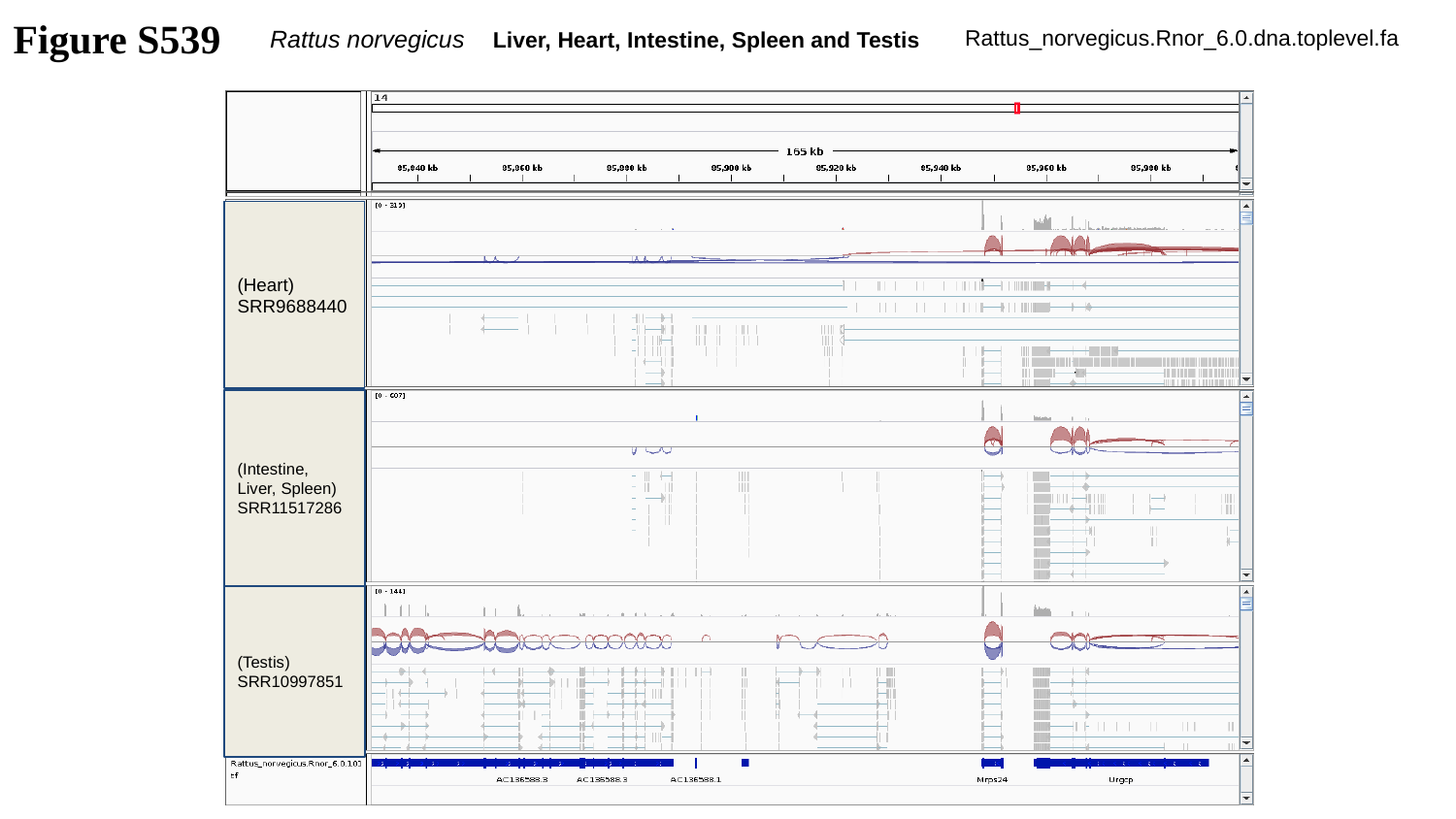

Figure S539
Rattus norvegicus Liver, Heart, Intestine, Spleen and Testis
Rattus_norvegicus.Rnor_6.0.dna.toplevel.fa
(Heart)
SRR9688440
(Intestine, Liver, Spleen)
SRR11517286
(Testis)
SRR10997851

### Slide 540
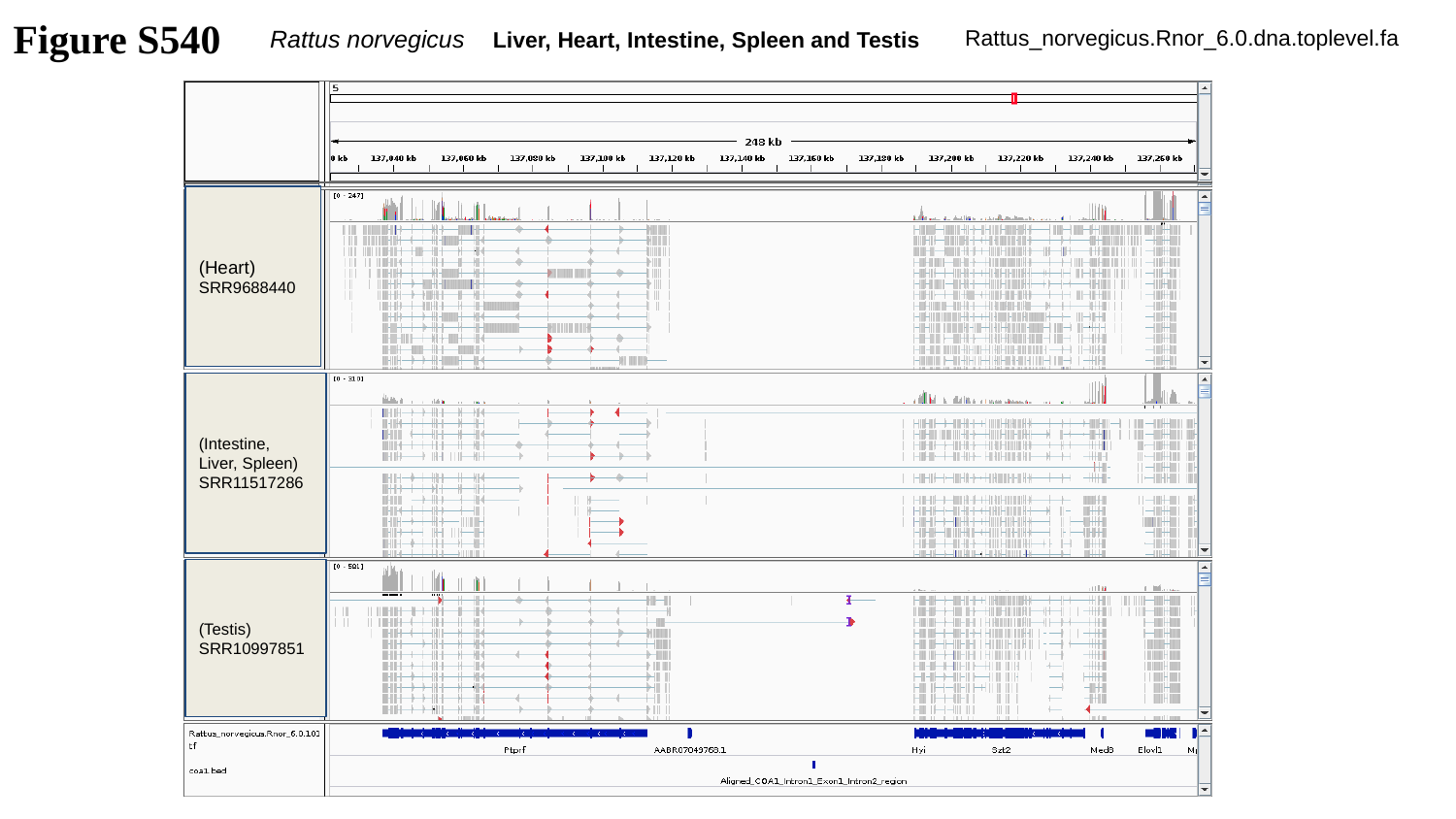

Figure S540
Rattus norvegicus Liver, Heart, Intestine, Spleen and Testis
Rattus_norvegicus.Rnor_6.0.dna.toplevel.fa
(Heart)
SRR9688440
(Intestine, Liver, Spleen)
SRR11517286
(Testis)
SRR10997851

### Slide 541
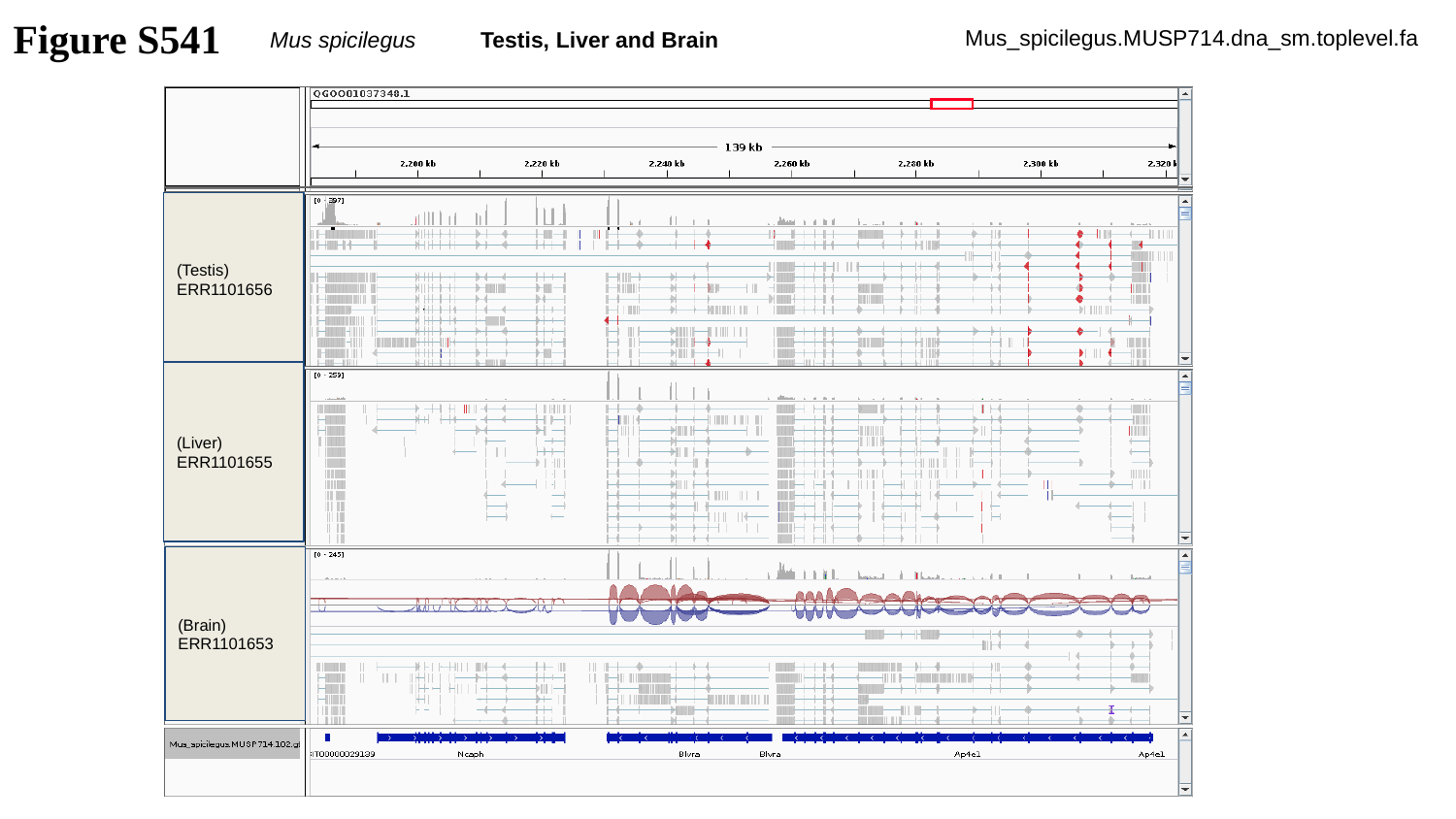

Figure S541
Mus spicilegus Testis, Liver and Brain
Mus_spicilegus.MUSP714.dna_sm.toplevel.fa
(Testis)
ERR1101656
(Liver)
ERR1101655
(Brain)
ERR1101653

### Slide 542
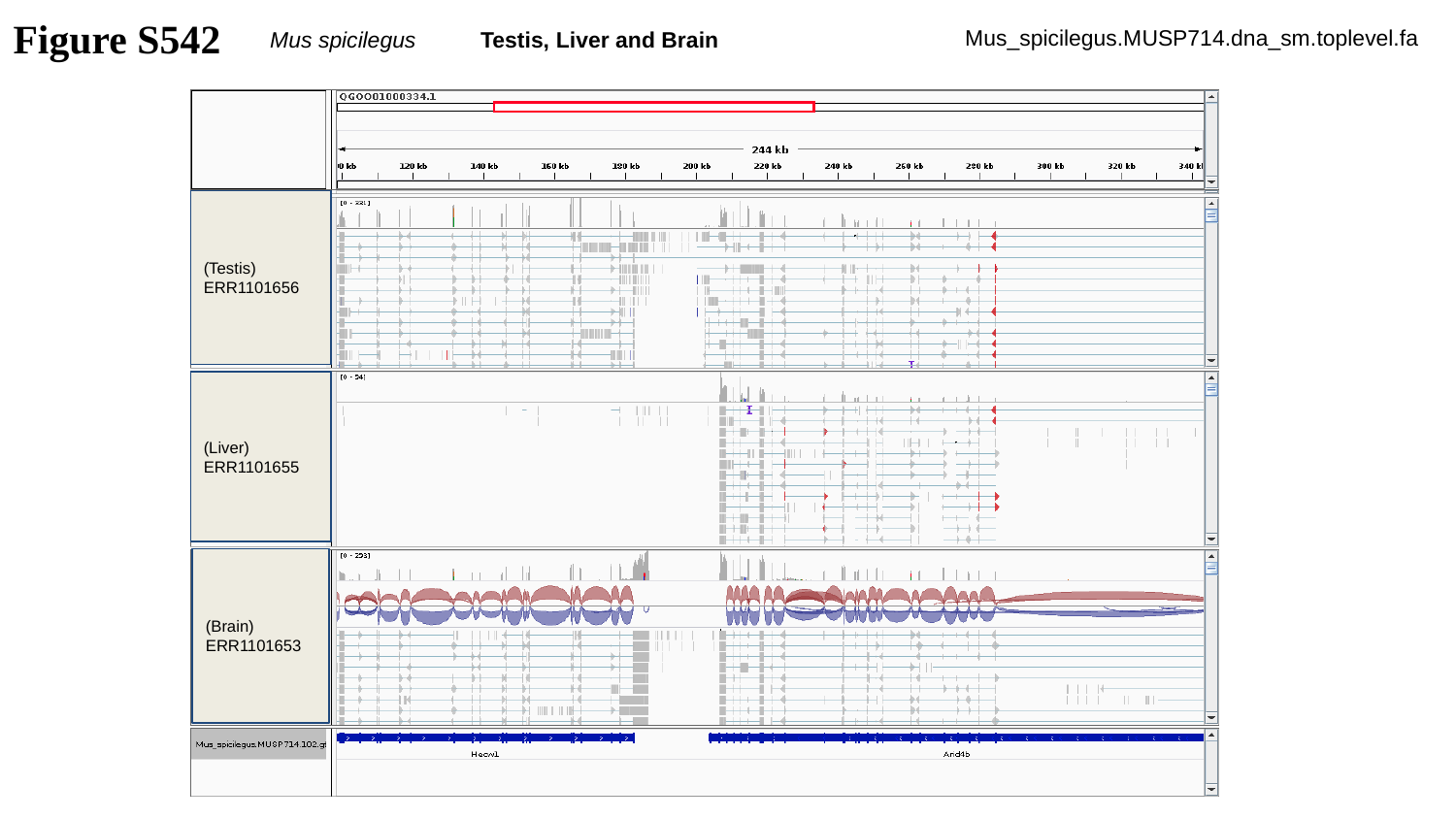

Figure S542
Mus spicilegus Testis, Liver and Brain
Mus_spicilegus.MUSP714.dna_sm.toplevel.fa
(Testis)
ERR1101656
(Liver)
ERR1101655
(Brain)
ERR1101653

### Slide 543
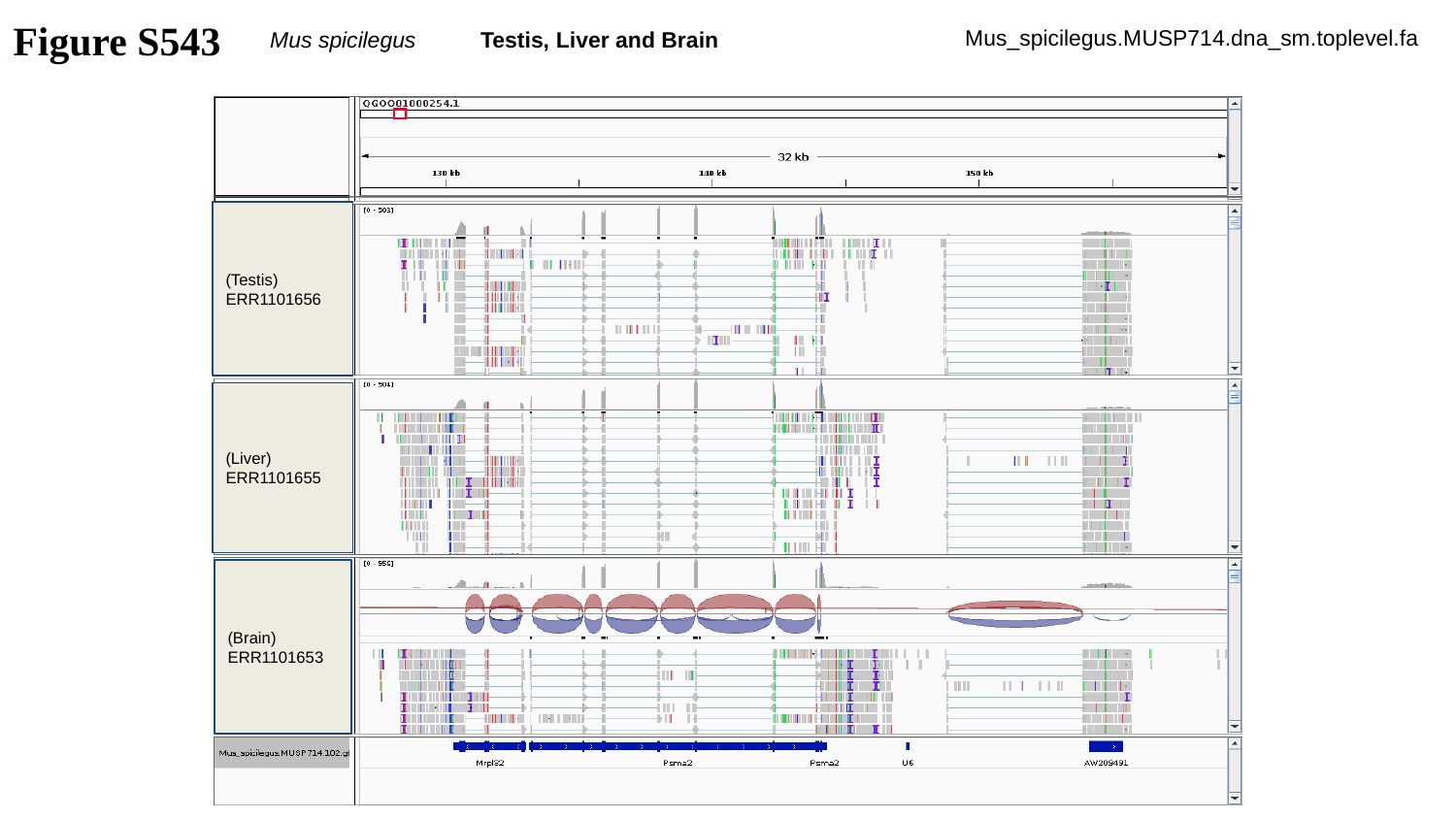

Figure S543
Mus spicilegus Testis, Liver and Brain
Mus_spicilegus.MUSP714.dna_sm.toplevel.fa
(Testis)
ERR1101656
(Liver)
ERR1101655
(Brain)
ERR1101653

### Slide 544
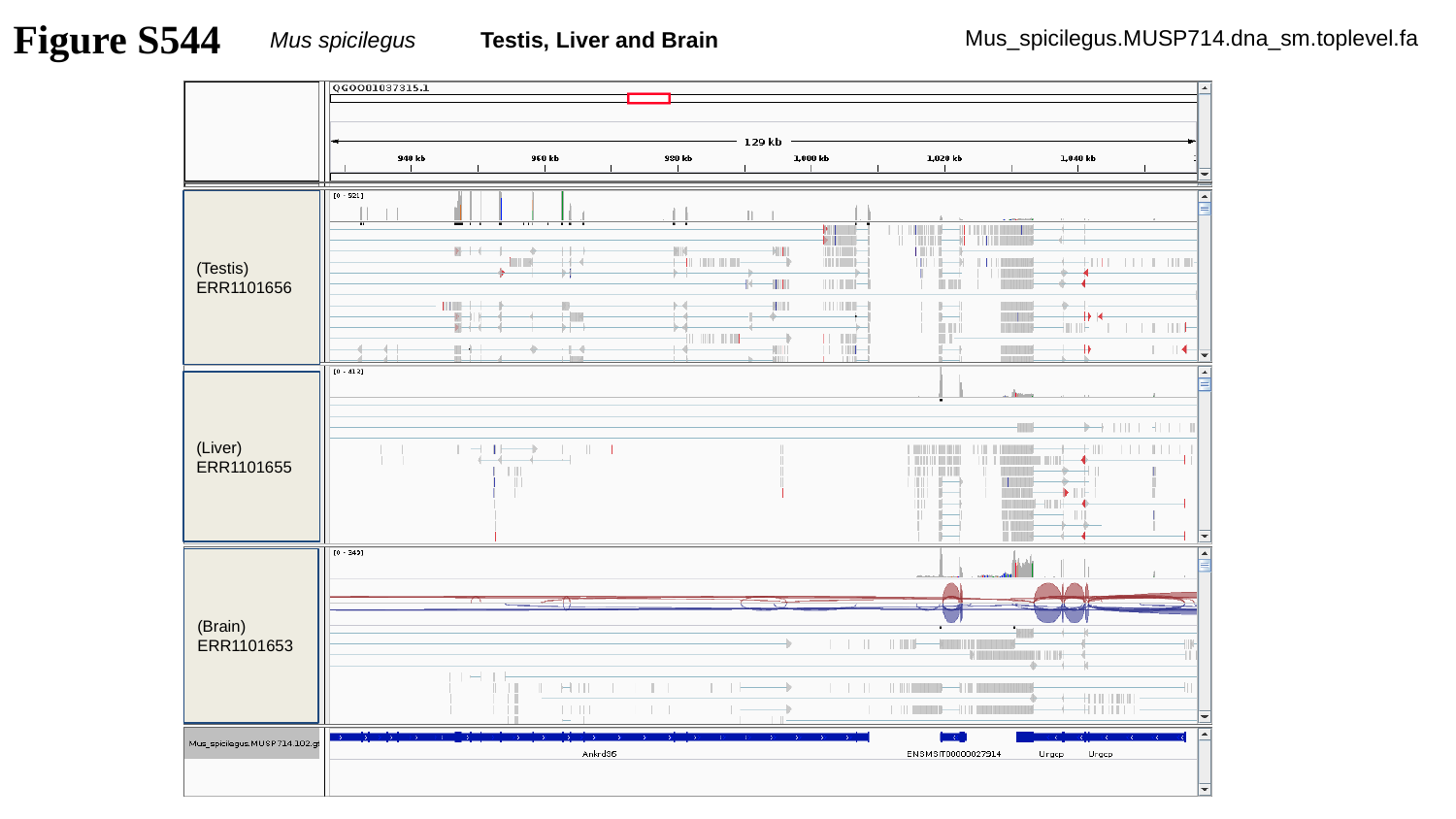

Figure S544
Mus spicilegus Testis, Liver and Brain
Mus_spicilegus.MUSP714.dna_sm.toplevel.fa
(Testis)
ERR1101656
(Liver)
ERR1101655
(Brain)
ERR1101653

### Slide 545
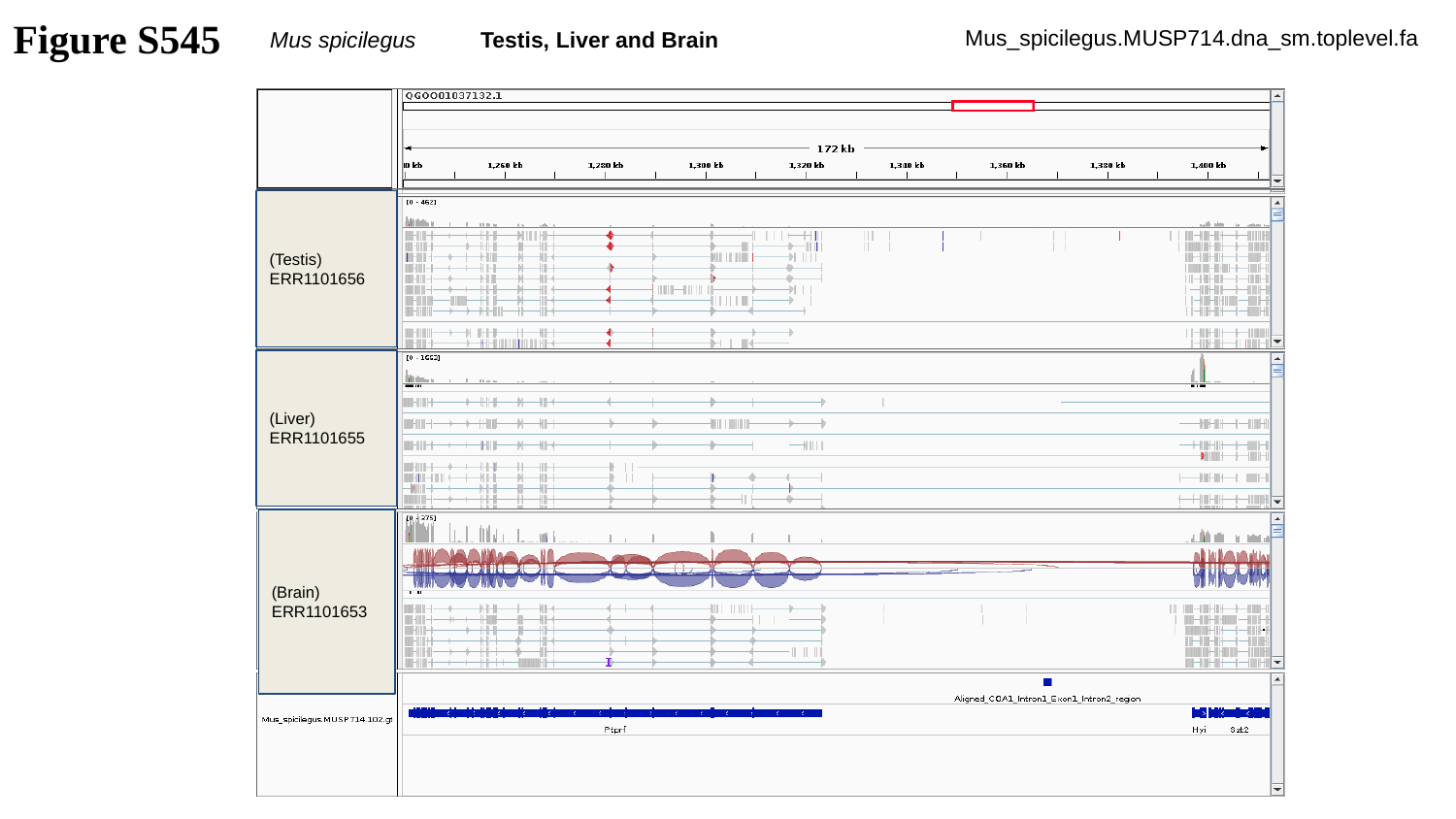

Figure S545
Mus spicilegus Testis, Liver and Brain
Mus_spicilegus.MUSP714.dna_sm.toplevel.fa
(Testis)
ERR1101656
(Liver)
ERR1101655
(Brain)
ERR1101653

### Slide 546
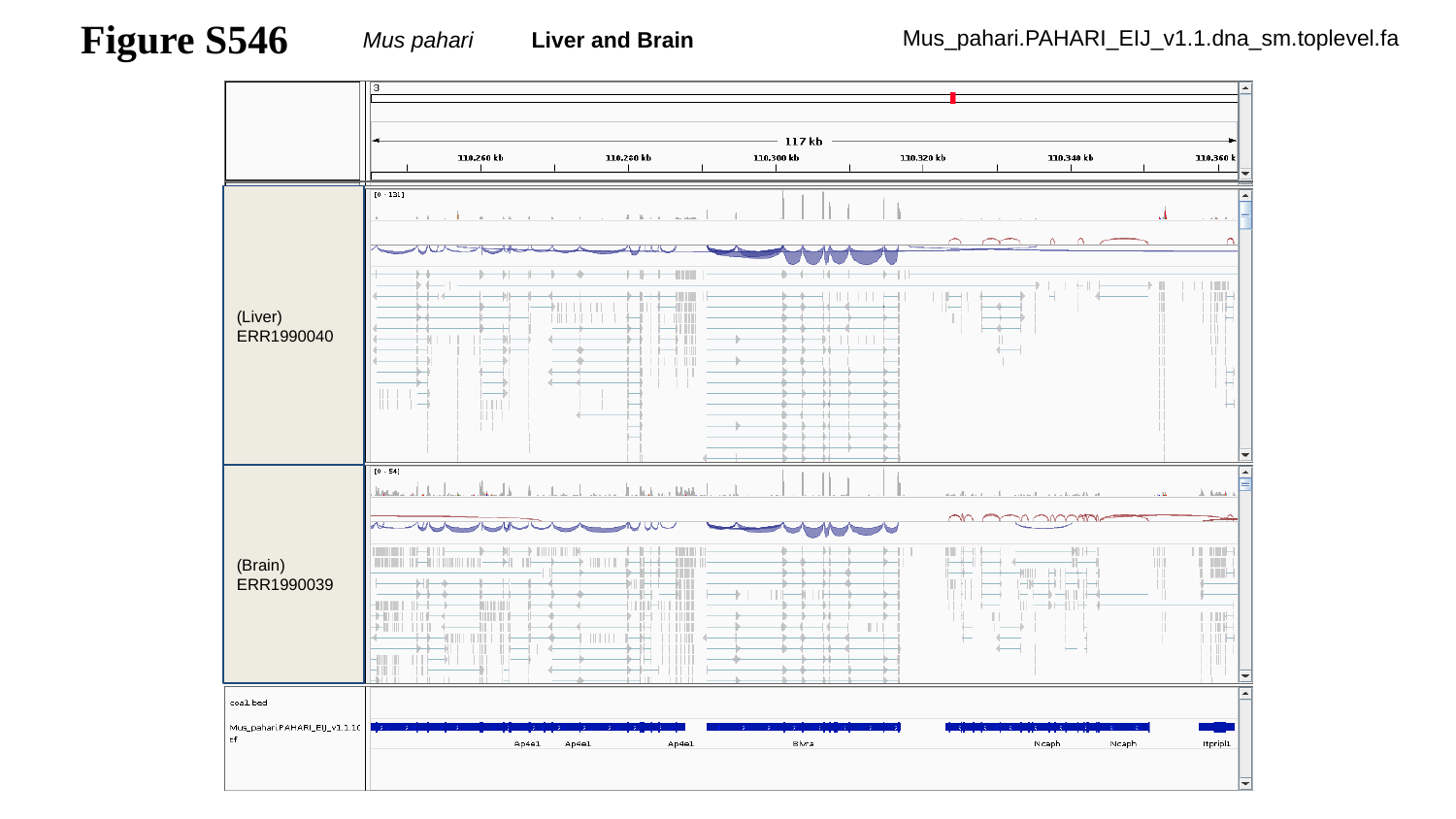

Figure S546
Mus pahari Liver and Brain
Mus_pahari.PAHARI_EIJ_v1.1.dna_sm.toplevel.fa
(Liver)
ERR1990040
(Brain)
ERR1990039

### Slide 547
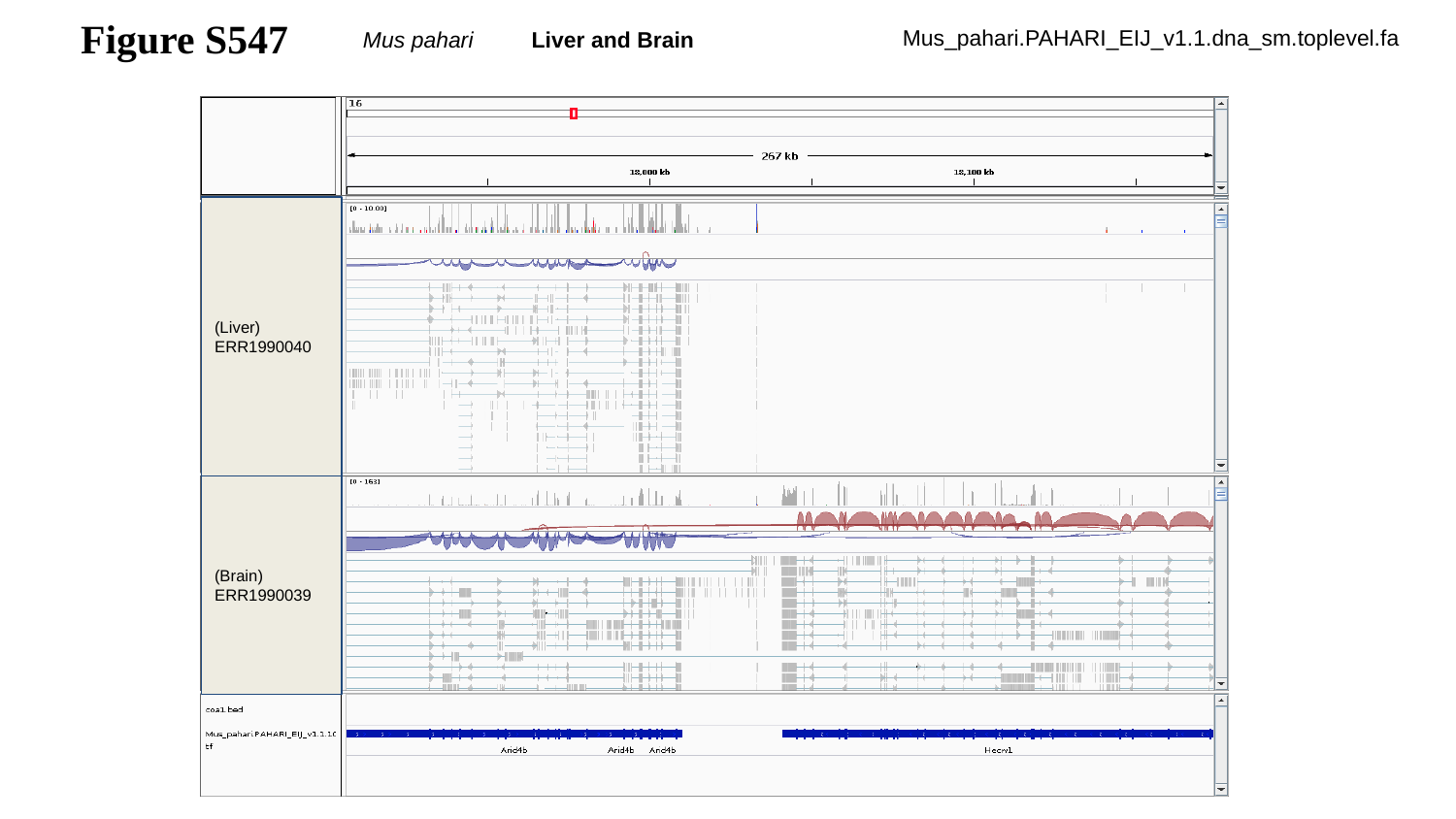

Figure S547
Mus pahari Liver and Brain
Mus_pahari.PAHARI_EIJ_v1.1.dna_sm.toplevel.fa
(Liver)
ERR1990040
(Brain)
ERR1990039

### Slide 548
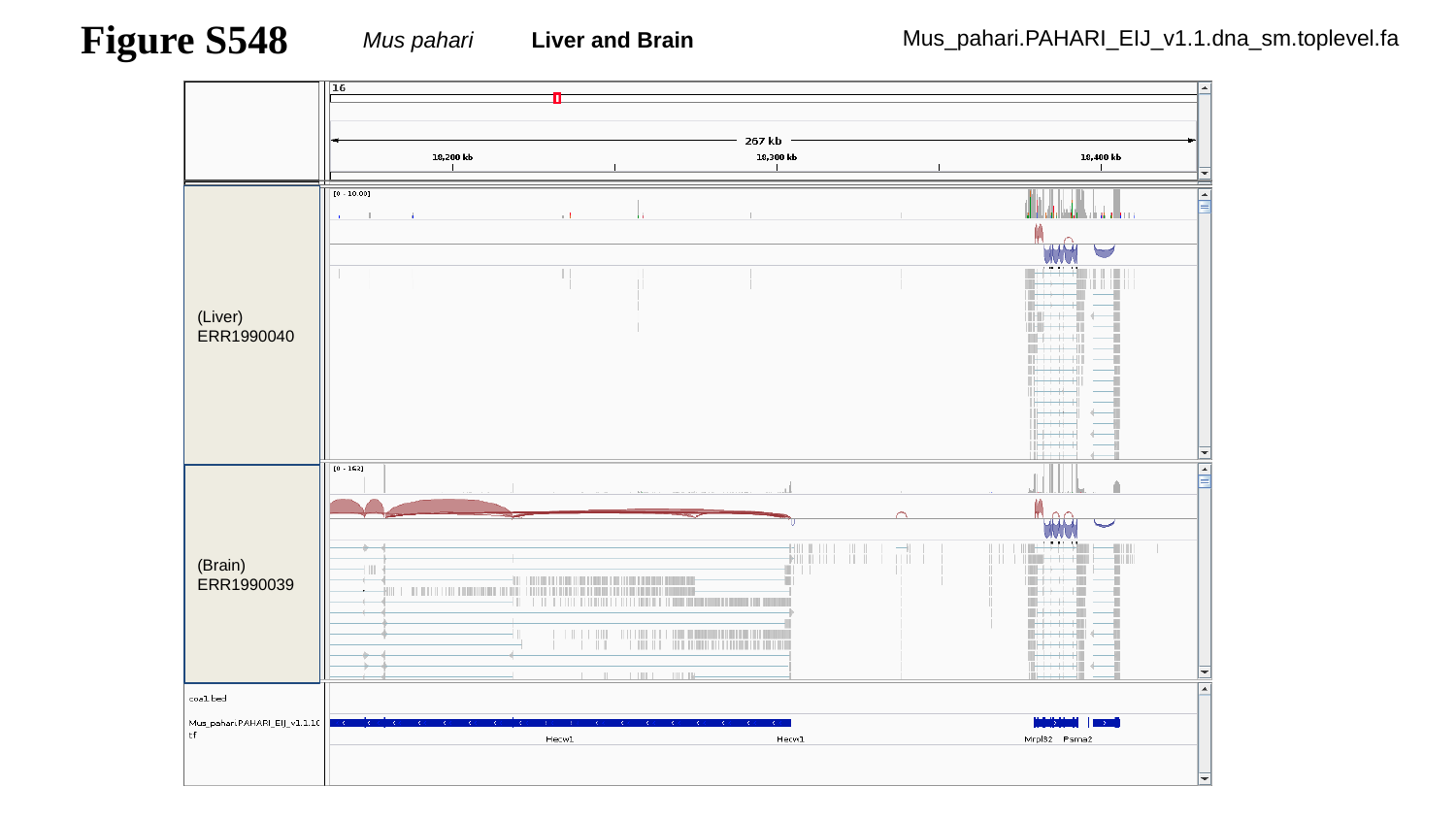

Figure S548
Mus pahari Liver and Brain
Mus_pahari.PAHARI_EIJ_v1.1.dna_sm.toplevel.fa
(Liver)
ERR1990040
(Brain)
ERR1990039

### Slide 549
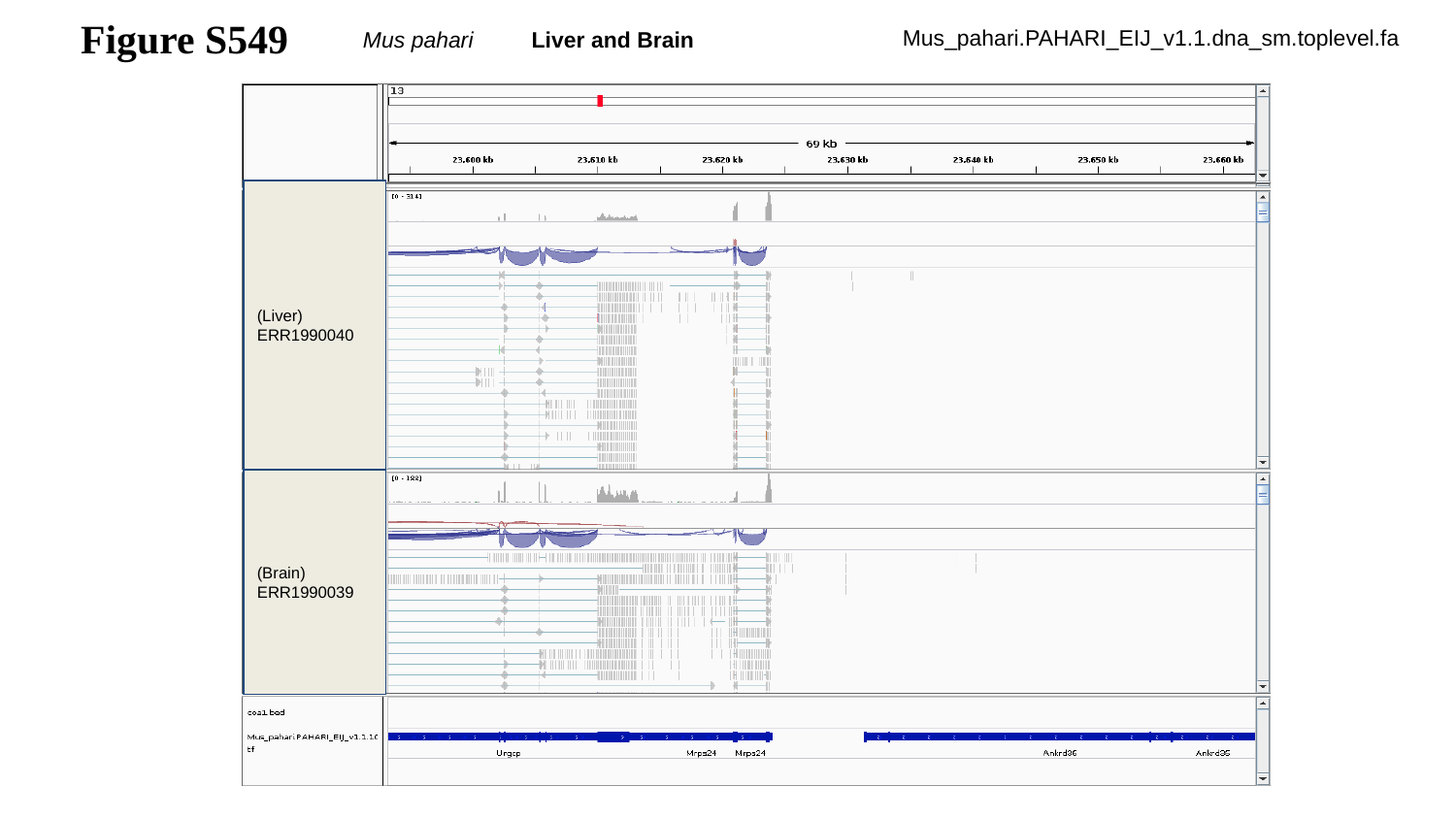

Figure S549
Mus pahari Liver and Brain
Mus_pahari.PAHARI_EIJ_v1.1.dna_sm.toplevel.fa
(Liver)
ERR1990040
(Brain)
ERR1990039

### Slide 550
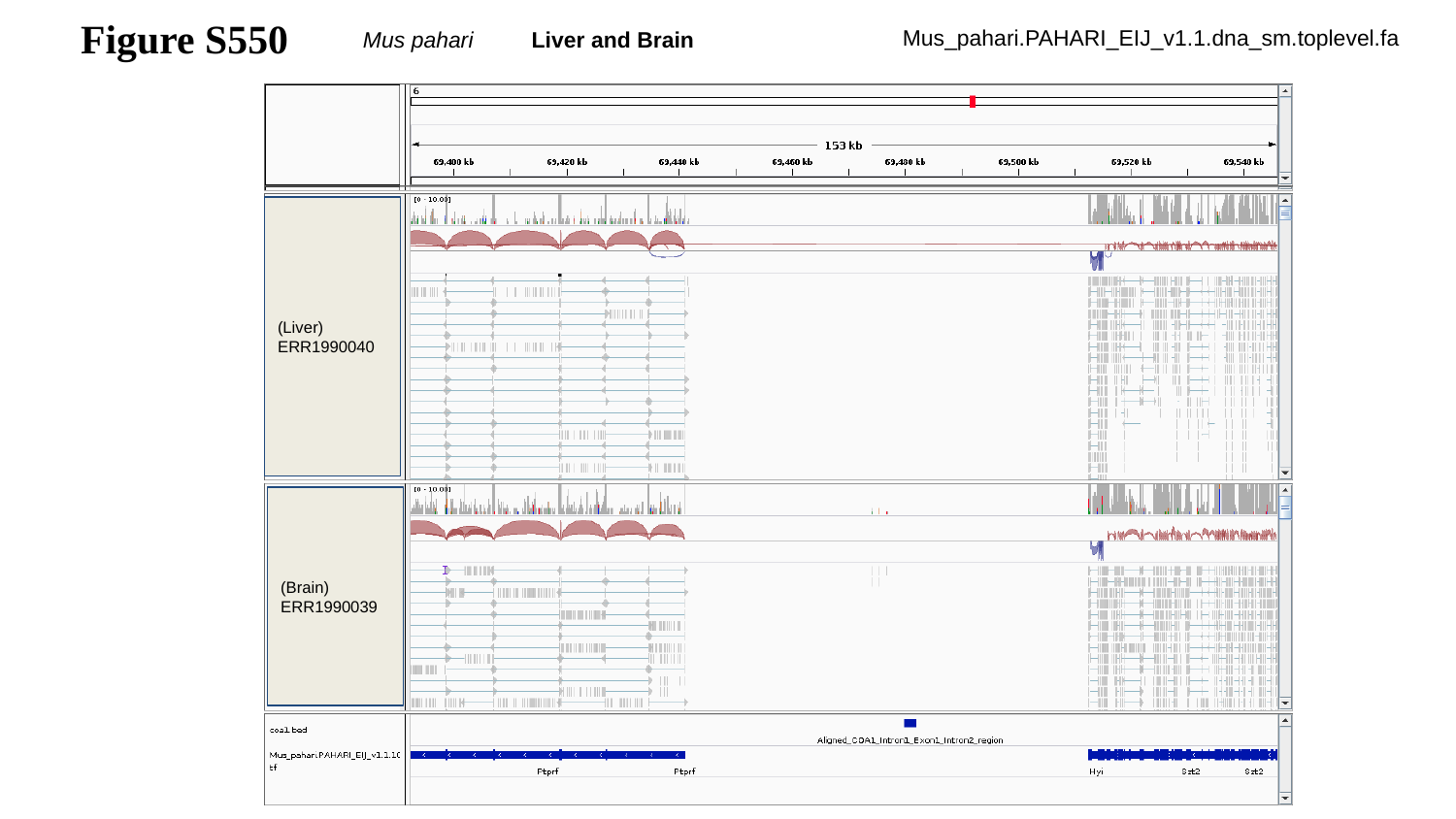

Figure S550
Mus pahari Liver and Brain
Mus_pahari.PAHARI_EIJ_v1.1.dna_sm.toplevel.fa
(Liver)
ERR1990040
(Brain)
ERR1990039

### Slide 551
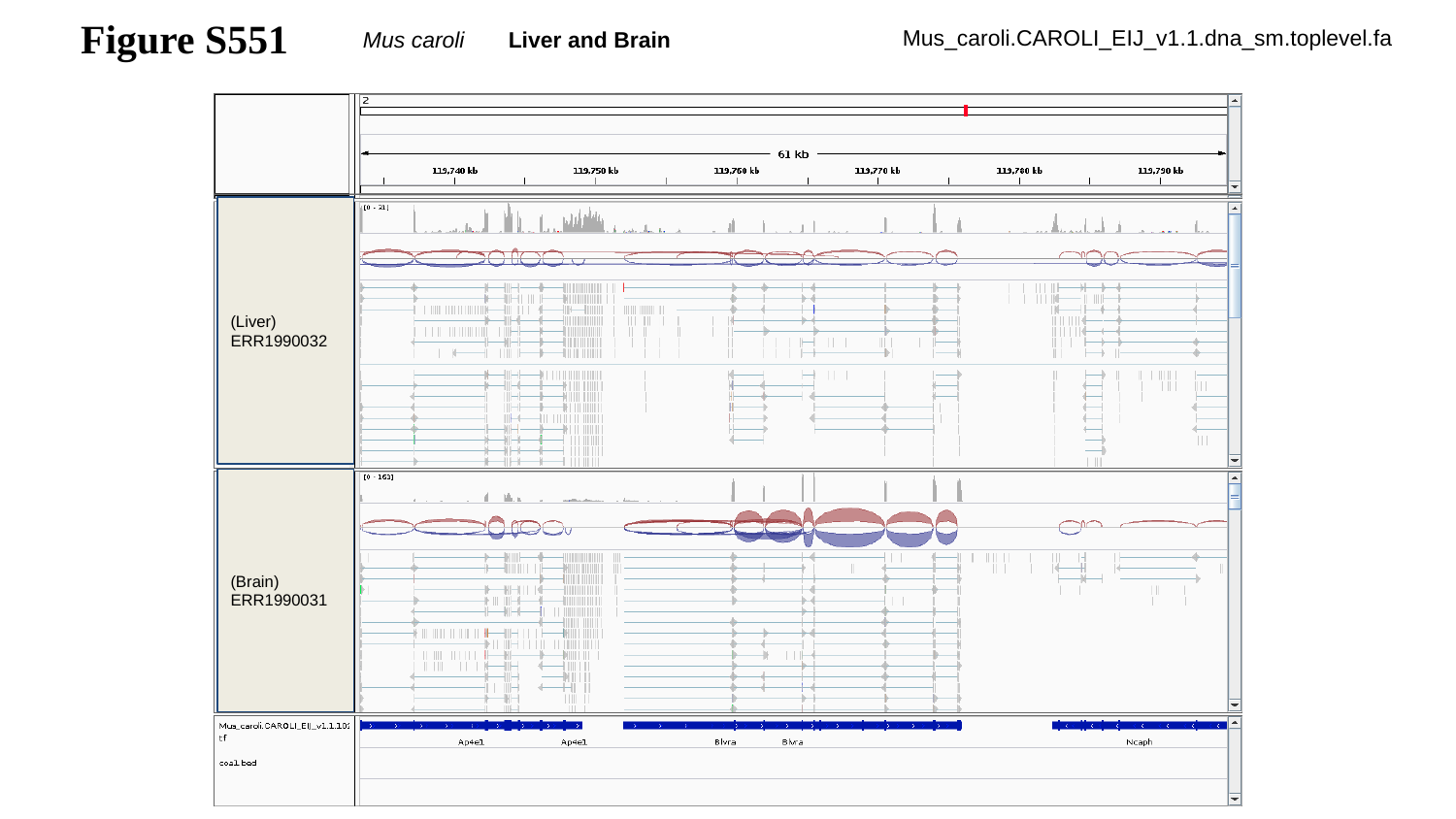

Figure S551
Mus caroli Liver and Brain
Mus_caroli.CAROLI_EIJ_v1.1.dna_sm.toplevel.fa
(Liver)
ERR1990032
(Brain)
ERR1990031

### Slide 552
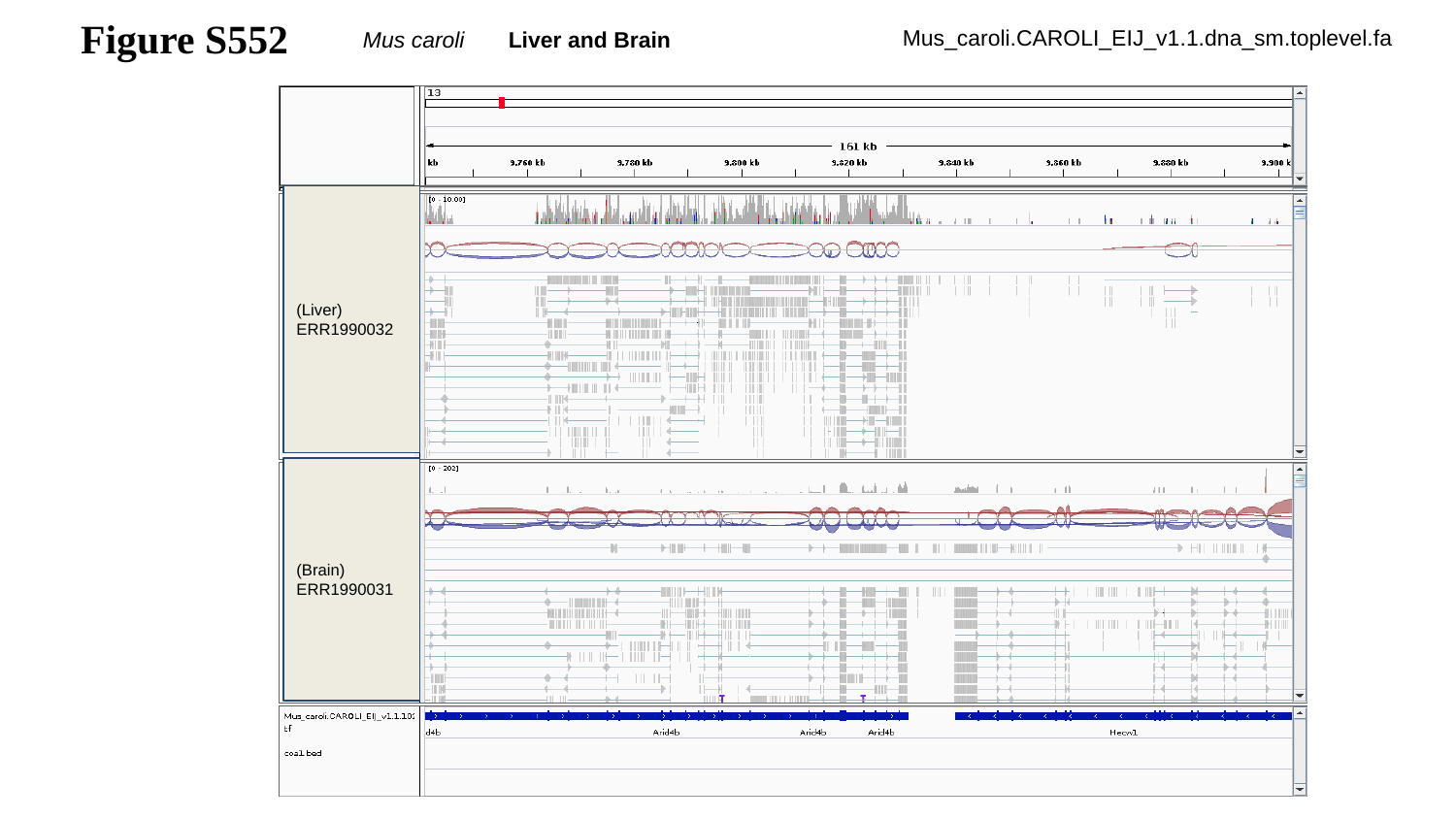

Figure S552
Mus caroli Liver and Brain
Mus_caroli.CAROLI_EIJ_v1.1.dna_sm.toplevel.fa
(Liver)
ERR1990032
(Brain)
ERR1990031

### Slide 553
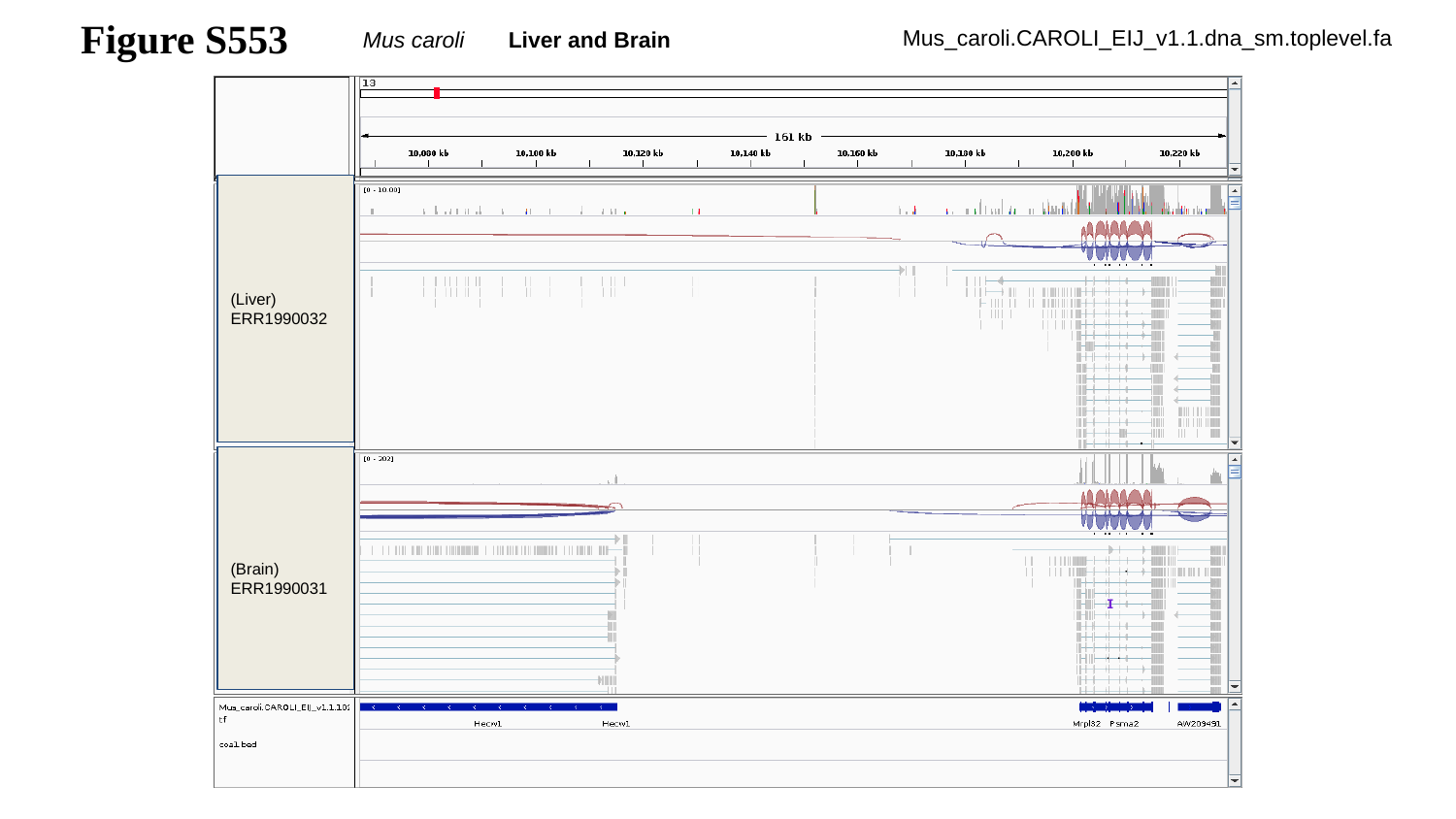

Figure S553
Mus caroli Liver and Brain
Mus_caroli.CAROLI_EIJ_v1.1.dna_sm.toplevel.fa
(Liver)
ERR1990032
(Brain)
ERR1990031

### Slide 554
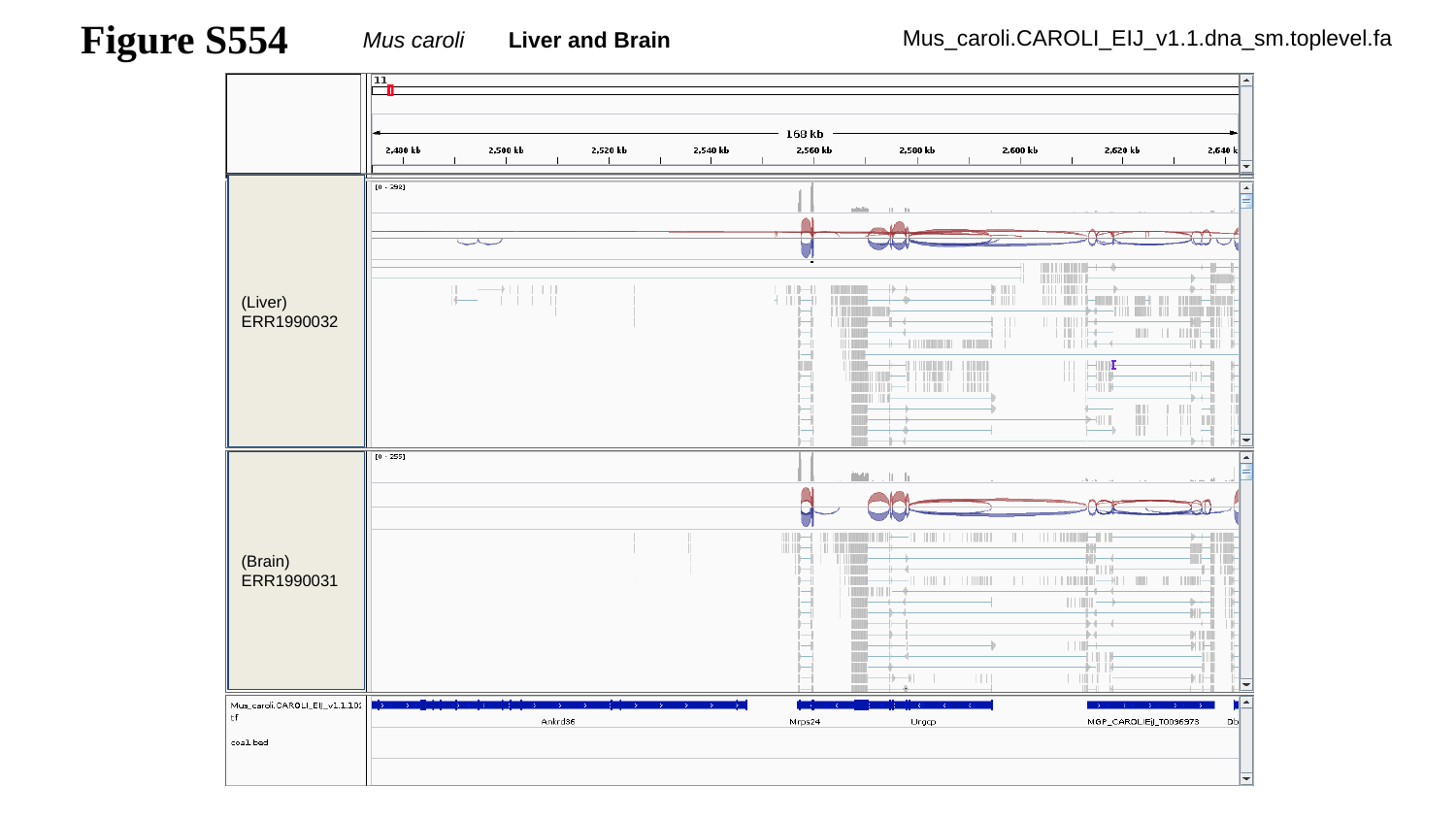

Figure S554
Mus caroli Liver and Brain
Mus_caroli.CAROLI_EIJ_v1.1.dna_sm.toplevel.fa
(Liver)
ERR1990032
(Brain)
ERR1990031

### Slide 555
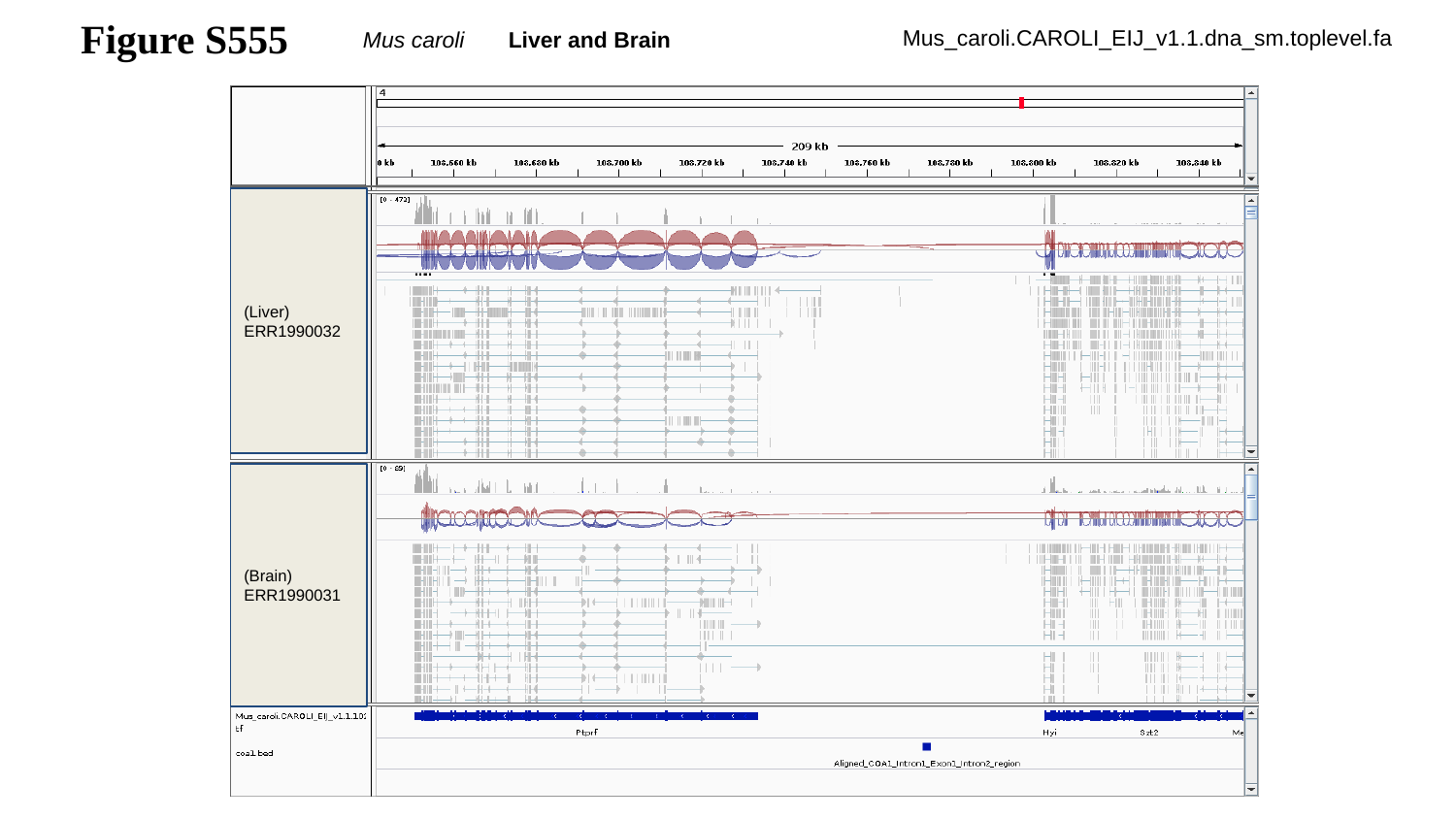

Figure S555
Mus caroli Liver and Brain
Mus_caroli.CAROLI_EIJ_v1.1.dna_sm.toplevel.fa
(Liver)
ERR1990032
(Brain)
ERR1990031

### Slide 556
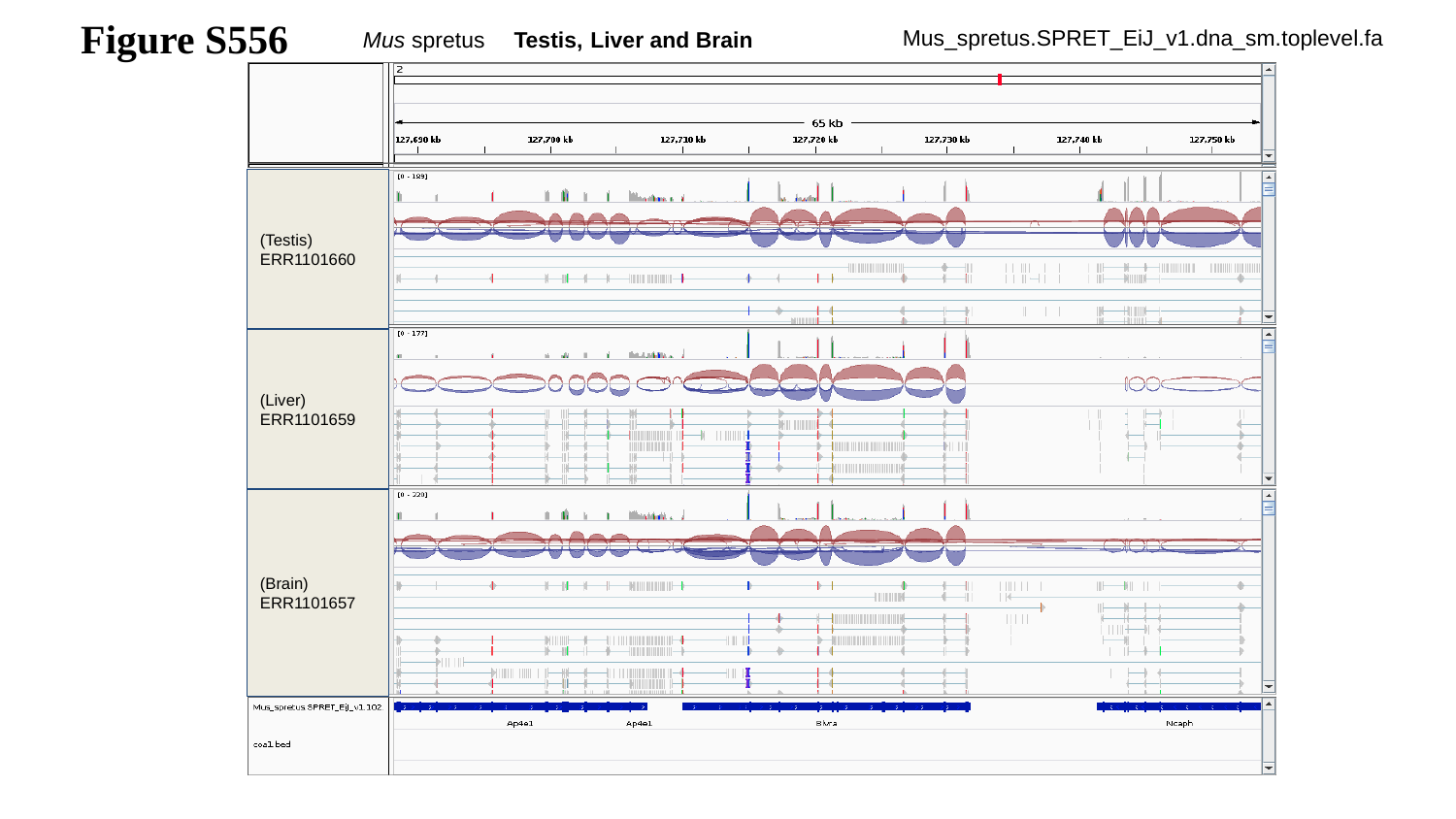

Figure S556
Mus spretus Testis, Liver and Brain
Mus_spretus.SPRET_EiJ_v1.dna_sm.toplevel.fa
(Testis)
ERR1101660
(Liver)
ERR1101659
(Brain)
ERR1101657

### Slide 557
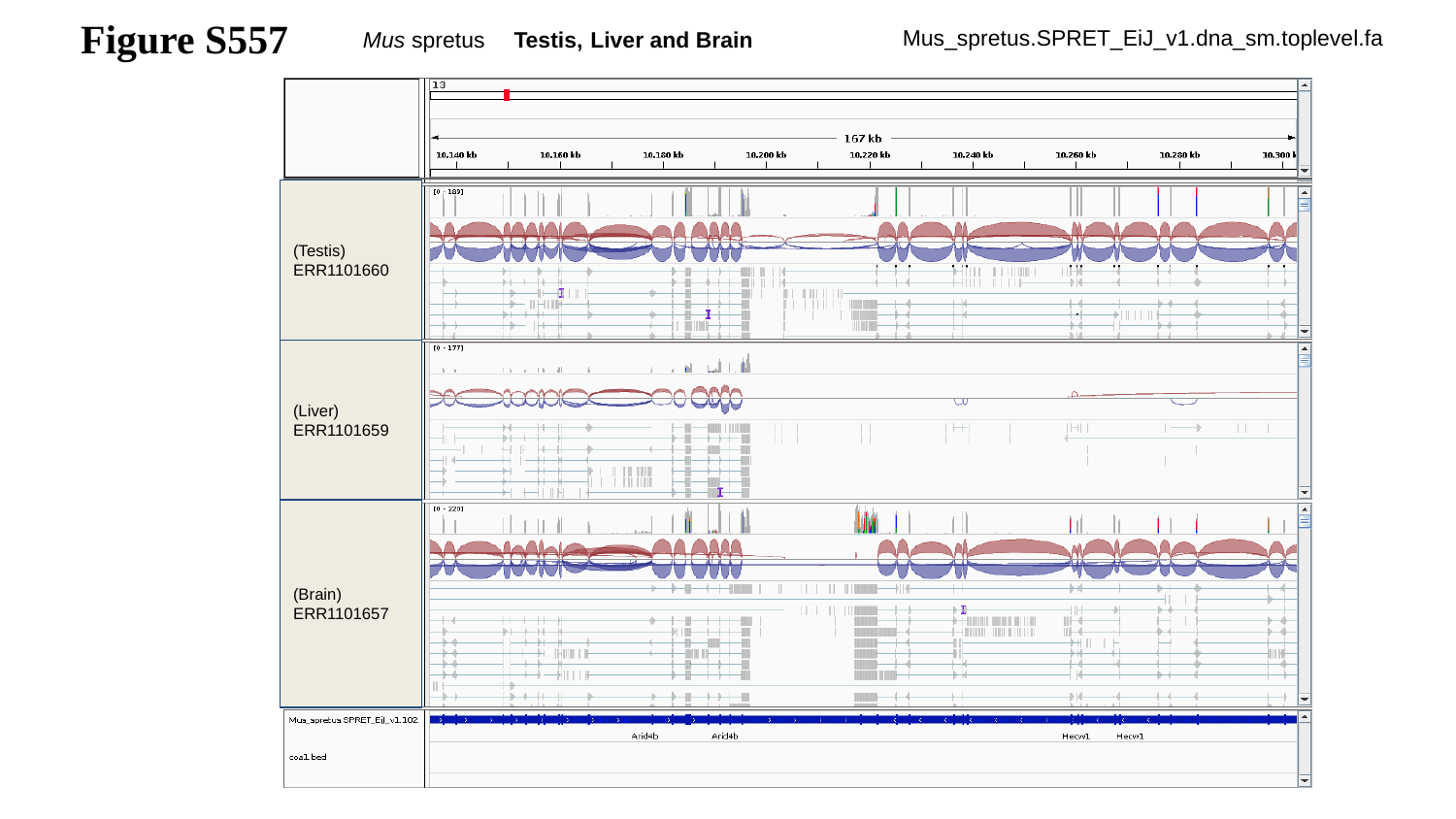

Figure S557
Mus spretus Testis, Liver and Brain
Mus_spretus.SPRET_EiJ_v1.dna_sm.toplevel.fa
(Testis)
ERR1101660
(Liver)
ERR1101659
(Brain)
ERR1101657

### Slide 558
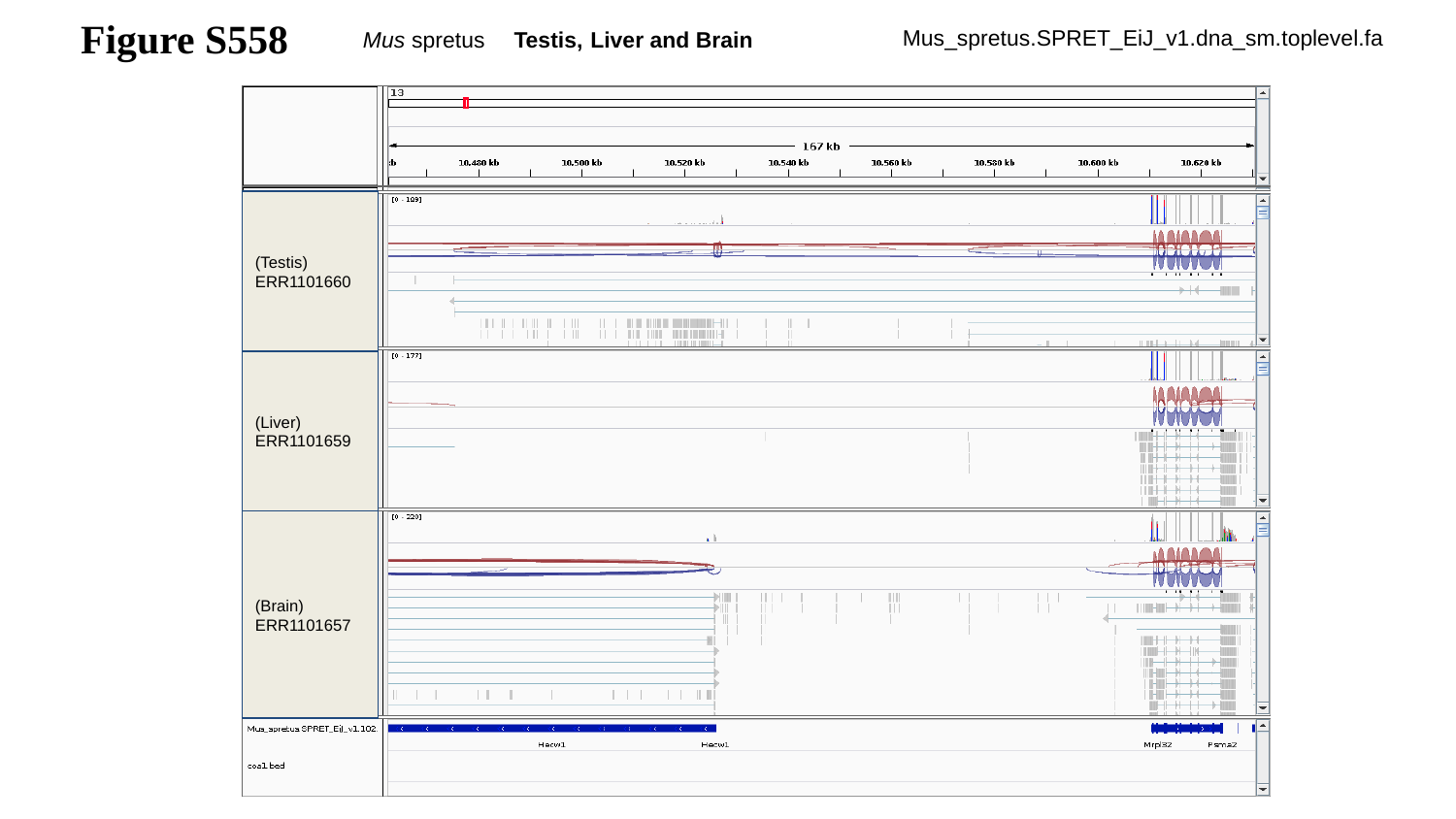

Figure S558
Mus spretus Testis, Liver and Brain
Mus_spretus.SPRET_EiJ_v1.dna_sm.toplevel.fa
(Testis)
ERR1101660
(Liver)
ERR1101659
(Brain)
ERR1101657

### Slide 559
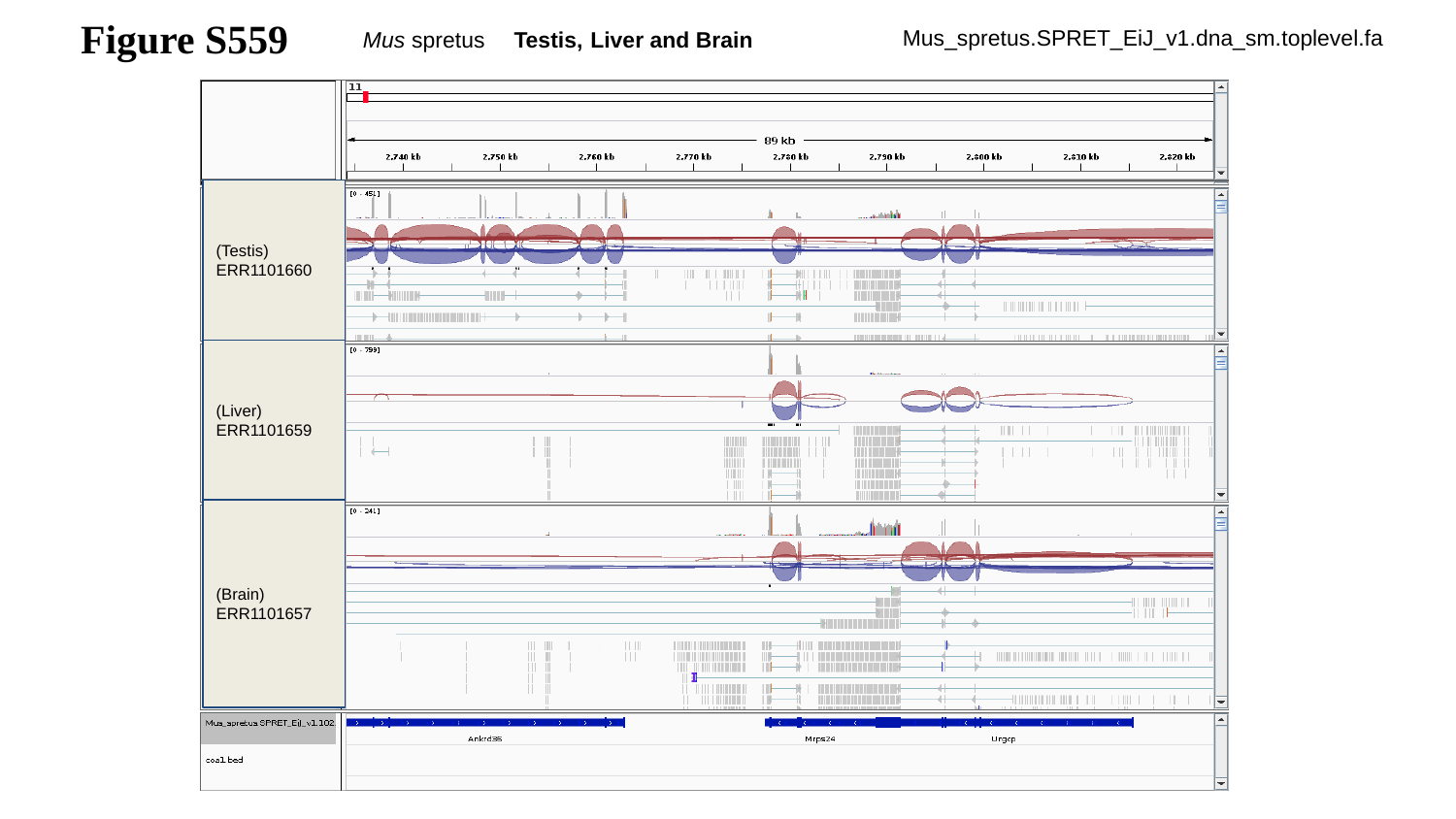

Figure S559
Mus spretus Testis, Liver and Brain
Mus_spretus.SPRET_EiJ_v1.dna_sm.toplevel.fa
(Testis)
ERR1101660
(Liver)
ERR1101659
(Brain)
ERR1101657

### Slide 560
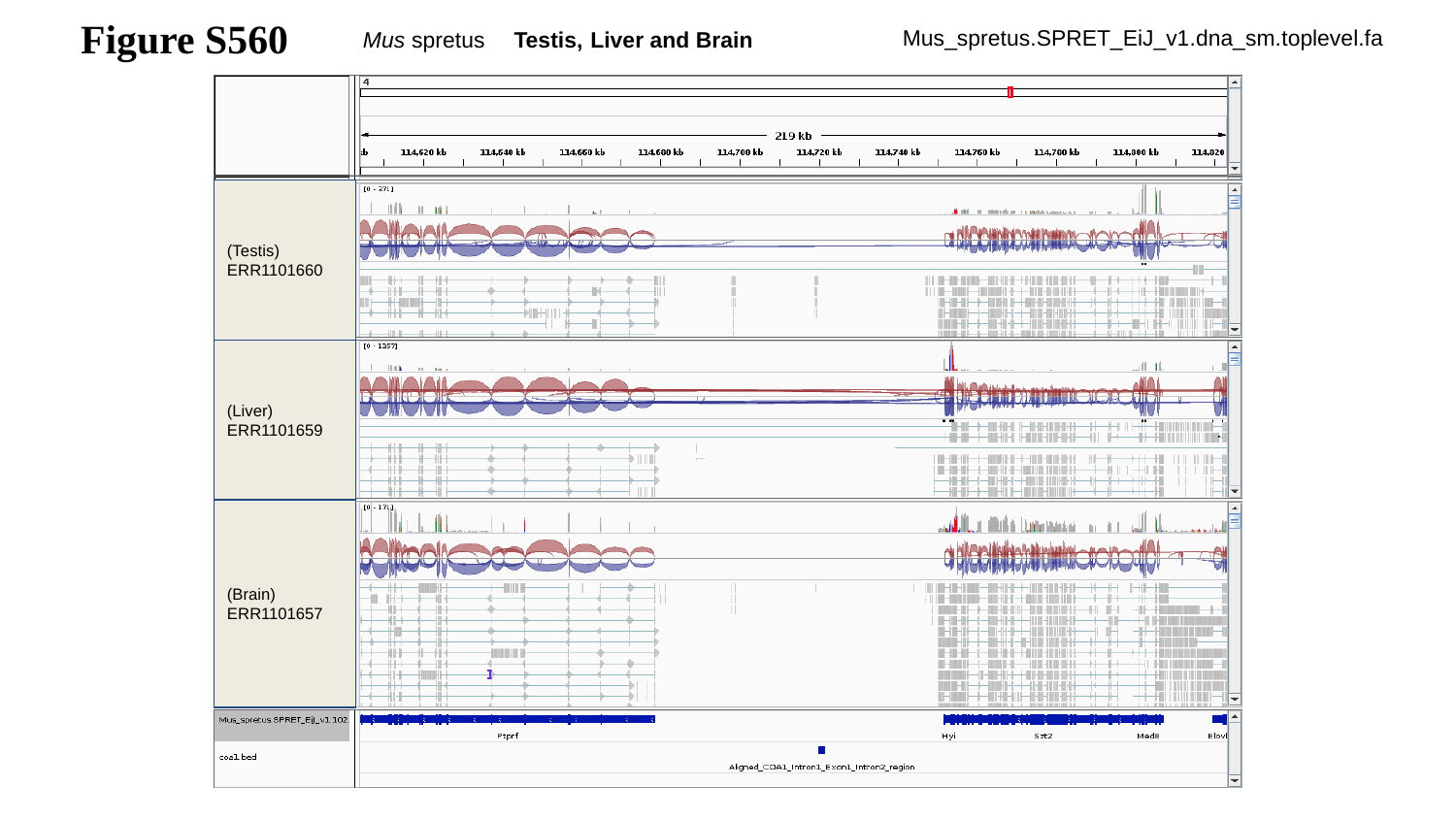

Figure S560
Mus spretus Testis, Liver and Brain
Mus_spretus.SPRET_EiJ_v1.dna_sm.toplevel.fa
(Testis)
ERR1101660
(Liver)
ERR1101659
(Brain)
ERR1101657

### Slide 561
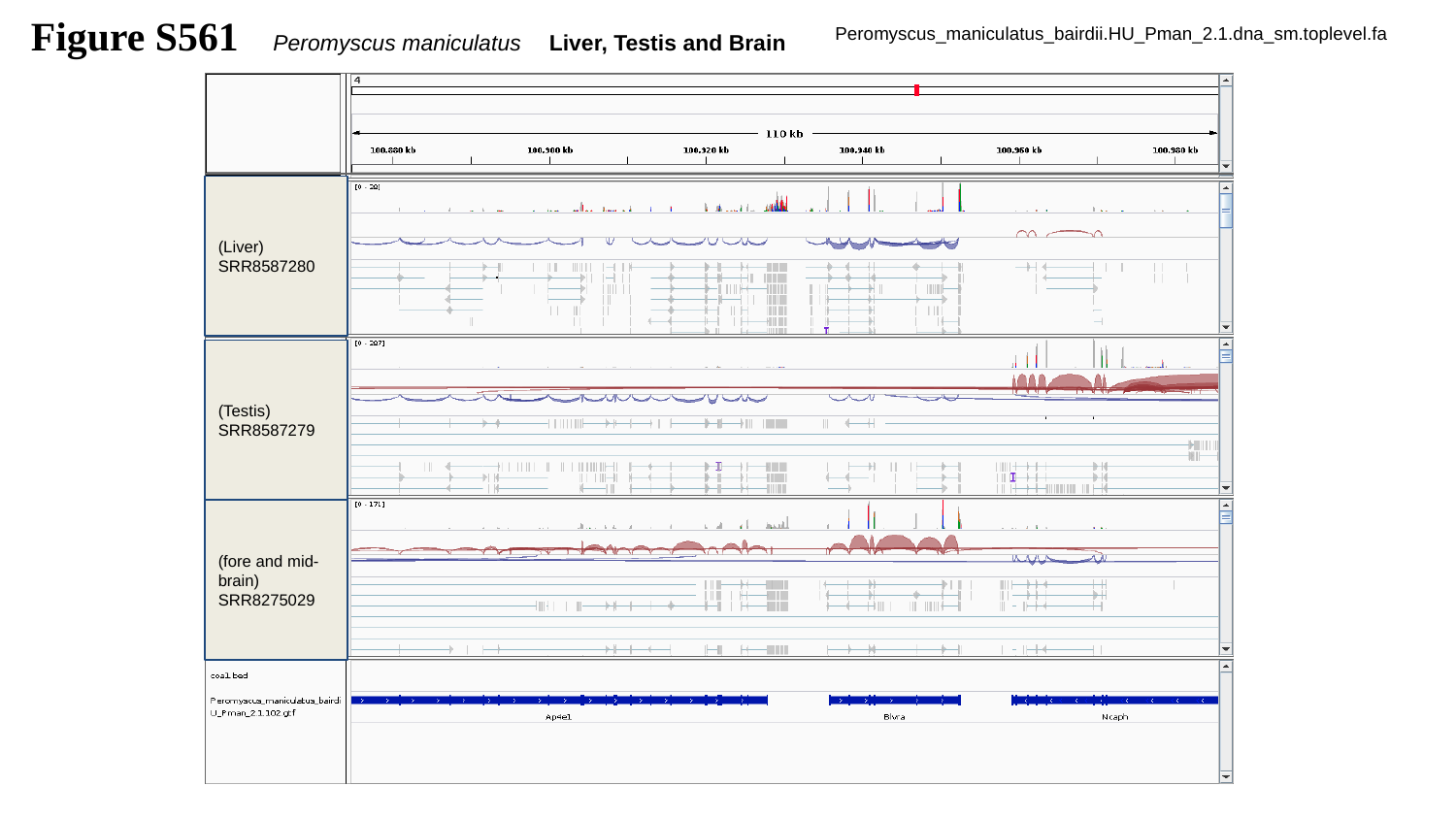

Figure S561
Peromyscus maniculatus Liver, Testis and Brain
Peromyscus_maniculatus_bairdii.HU_Pman_2.1.dna_sm.toplevel.fa
(Liver)
SRR8587280
(Testis)
SRR8587279
(fore and mid-brain)
SRR8275029

### Slide 562
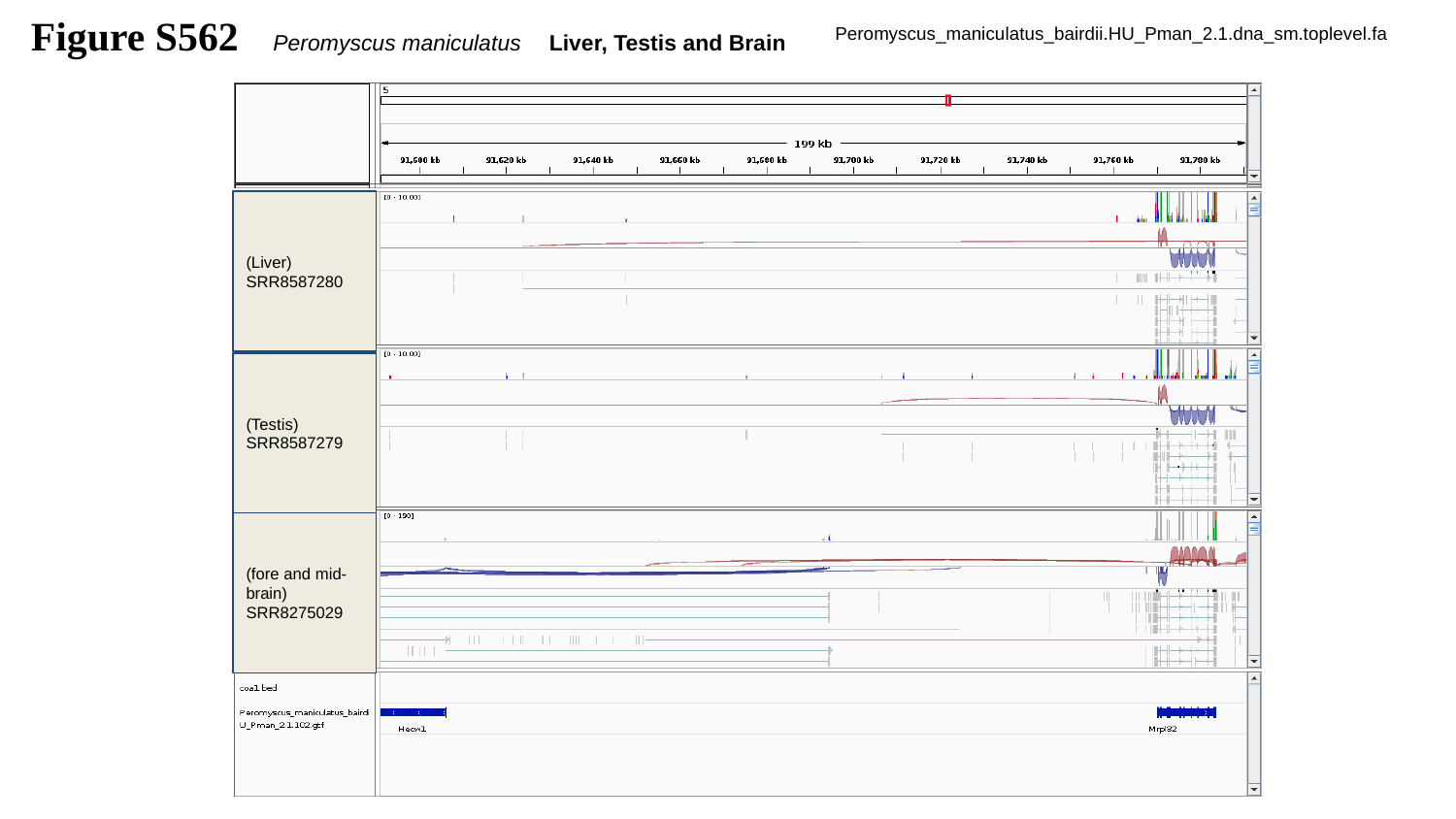

Figure S562
Peromyscus maniculatus Liver, Testis and Brain
Peromyscus_maniculatus_bairdii.HU_Pman_2.1.dna_sm.toplevel.fa
(Liver)
SRR8587280
(Testis)
SRR8587279
(fore and mid-brain)
SRR8275029

### Slide 563
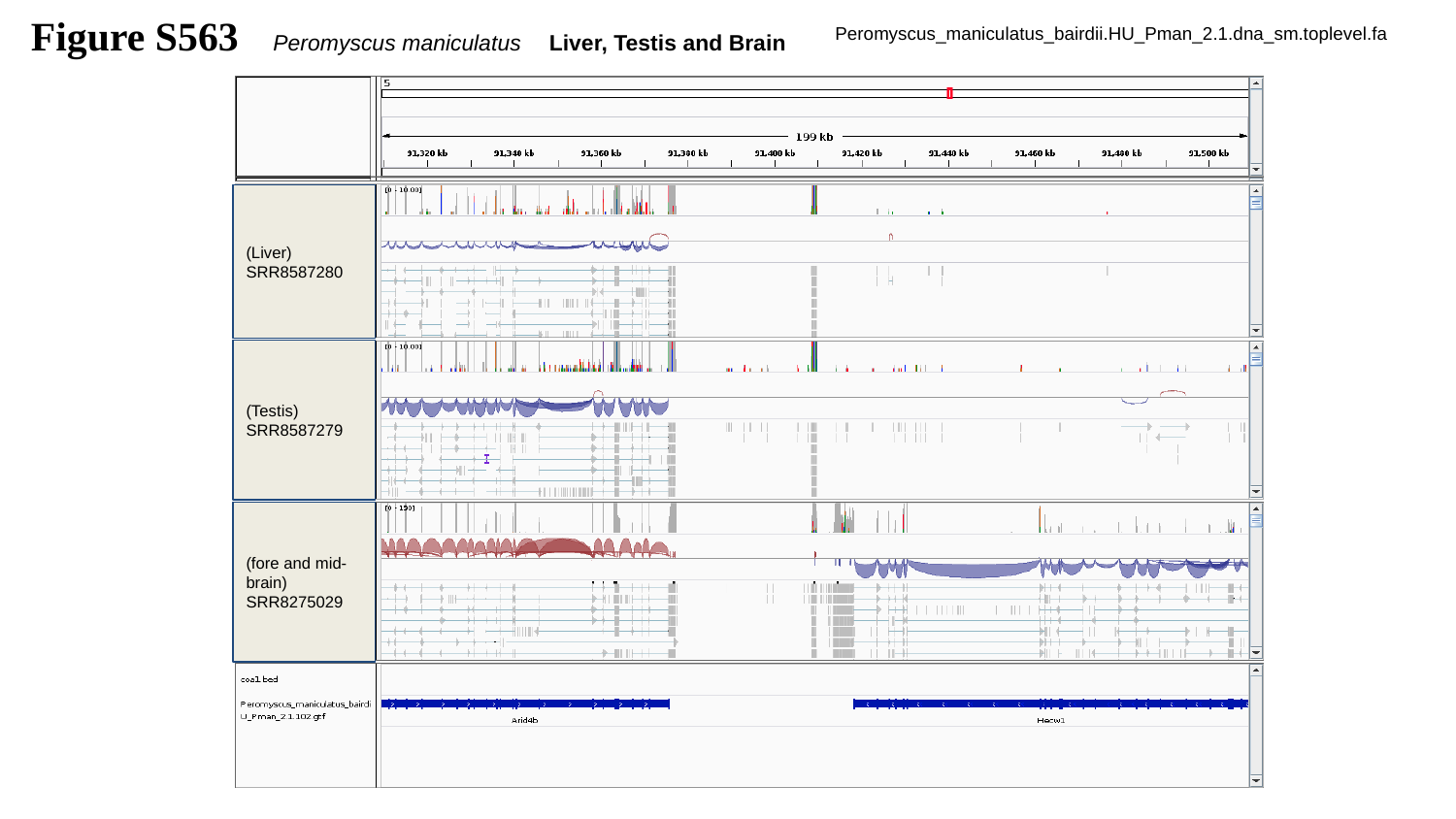

Figure S563
Peromyscus maniculatus Liver, Testis and Brain
Peromyscus_maniculatus_bairdii.HU_Pman_2.1.dna_sm.toplevel.fa
(Liver)
SRR8587280
(Testis)
SRR8587279
(fore and mid-brain)
SRR8275029

### Slide 564
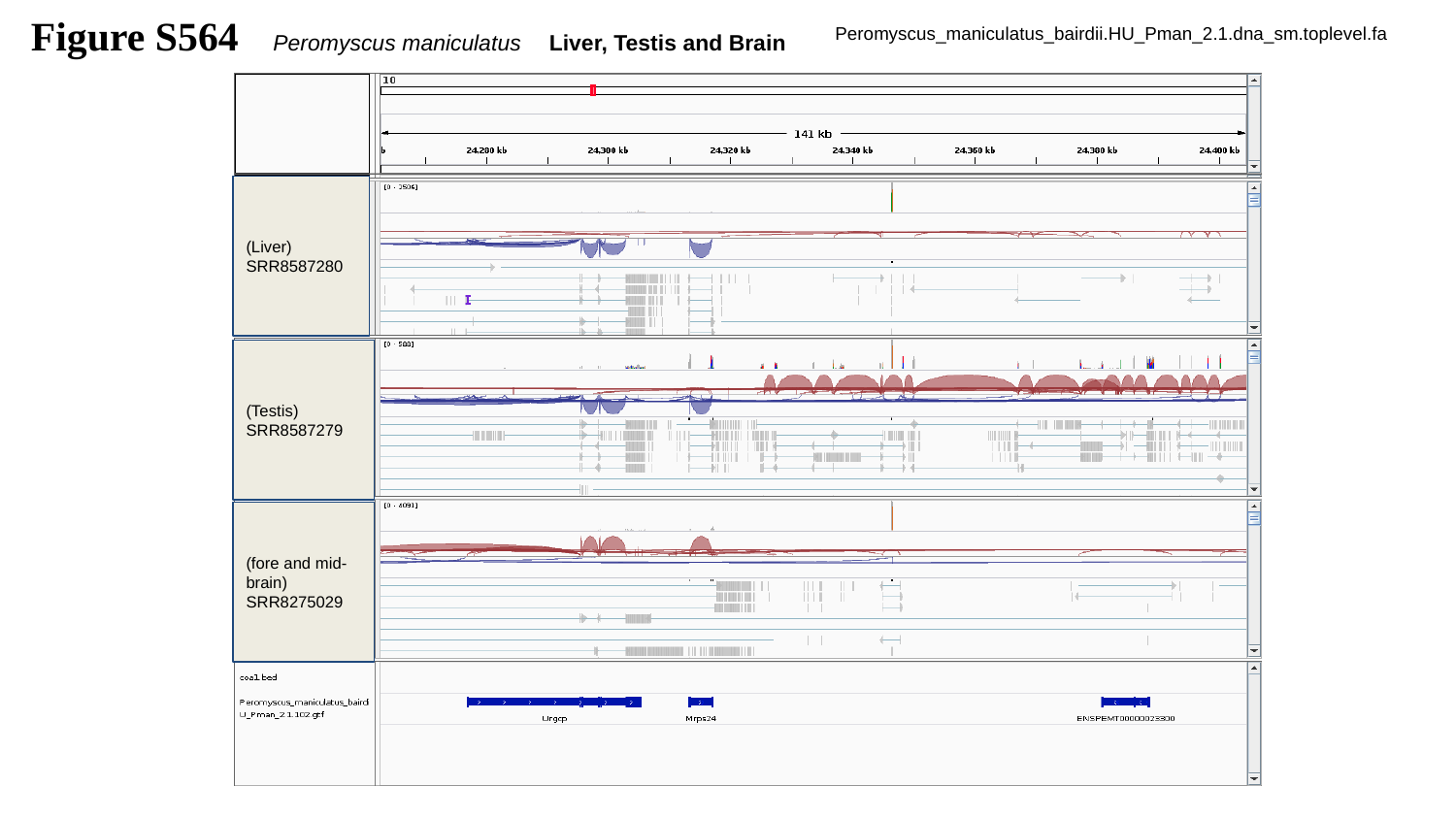

Figure S564
Peromyscus maniculatus Liver, Testis and Brain
Peromyscus_maniculatus_bairdii.HU_Pman_2.1.dna_sm.toplevel.fa
(Liver)
SRR8587280
(Testis)
SRR8587279
(fore and mid-brain)
SRR8275029

### Slide 565
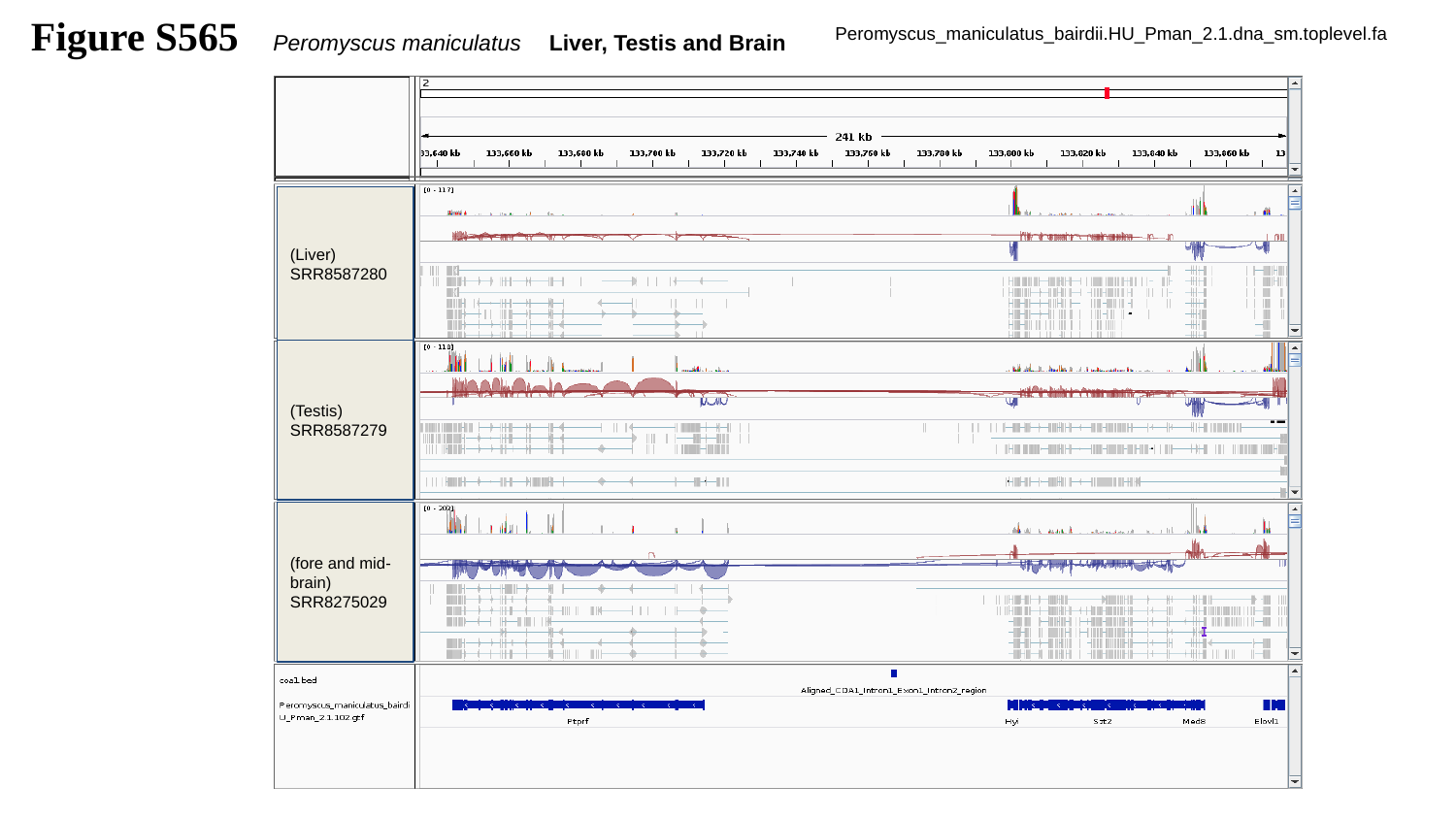

Figure S565
Peromyscus maniculatus Liver, Testis and Brain
Peromyscus_maniculatus_bairdii.HU_Pman_2.1.dna_sm.toplevel.fa
(Liver)
SRR8587280
(Testis)
SRR8587279
(fore and mid-brain)
SRR8275029

### Slide 566
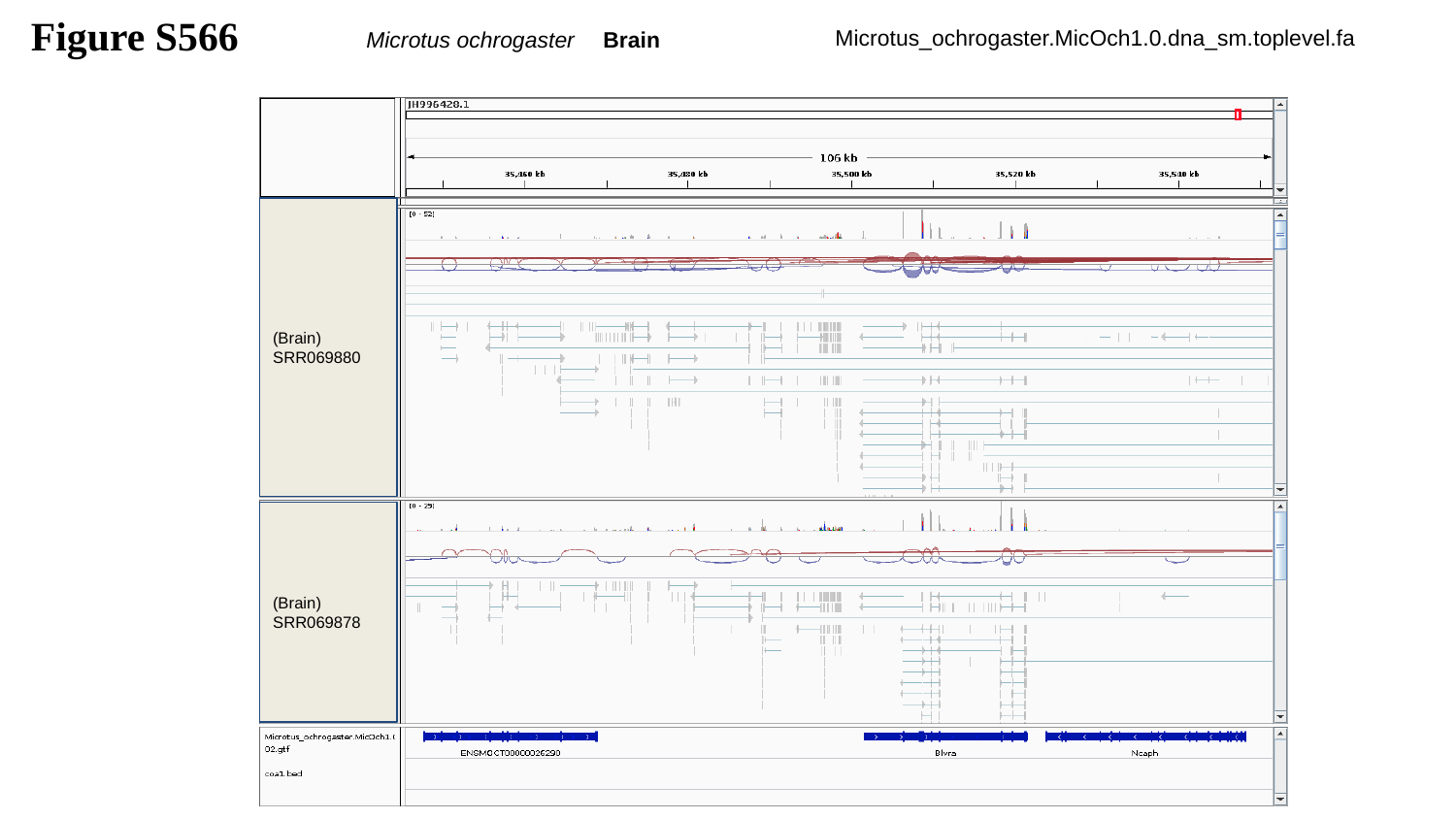

Figure S566
Microtus ochrogaster Brain
Microtus_ochrogaster.MicOch1.0.dna_sm.toplevel.fa
(Brain)
SRR069880
(Brain)
SRR069878

### Slide 567
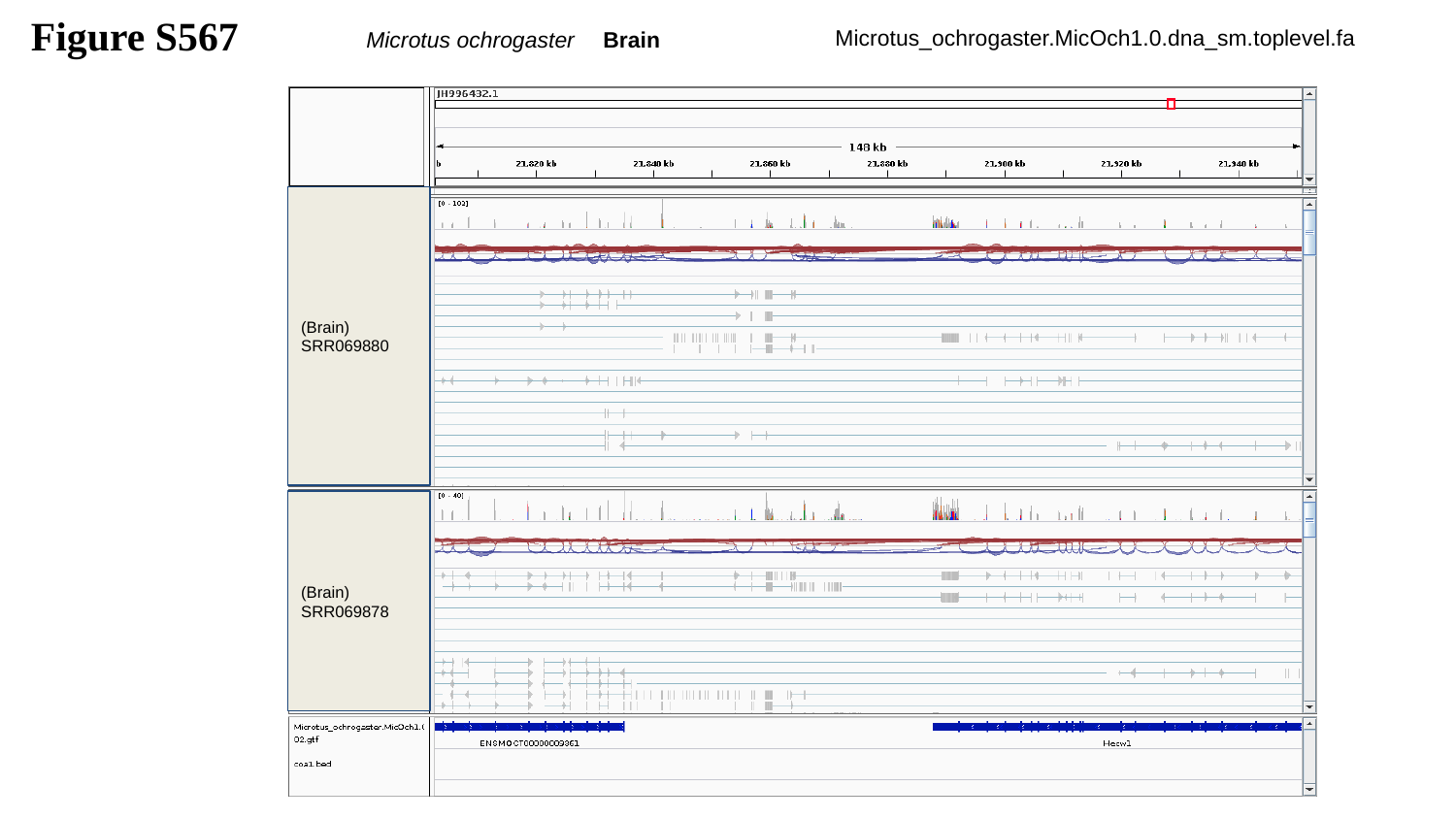

Figure S567
Microtus ochrogaster Brain
Microtus_ochrogaster.MicOch1.0.dna_sm.toplevel.fa
(Brain)
SRR069880
(Brain)
SRR069878

### Slide 568
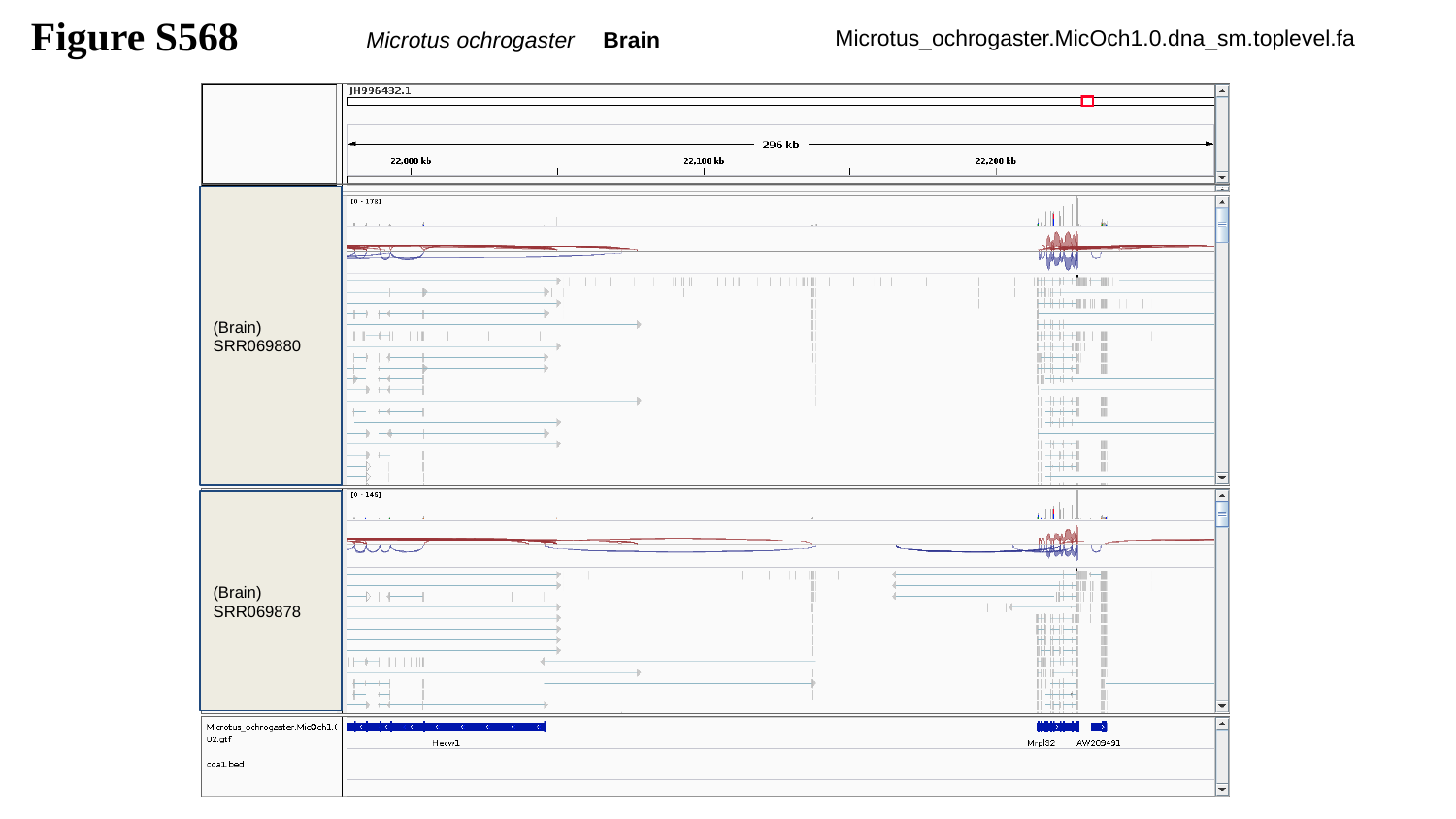

Figure S568
Microtus ochrogaster Brain
Microtus_ochrogaster.MicOch1.0.dna_sm.toplevel.fa
(Brain)
SRR069880
(Brain)
SRR069878

### Slide 569
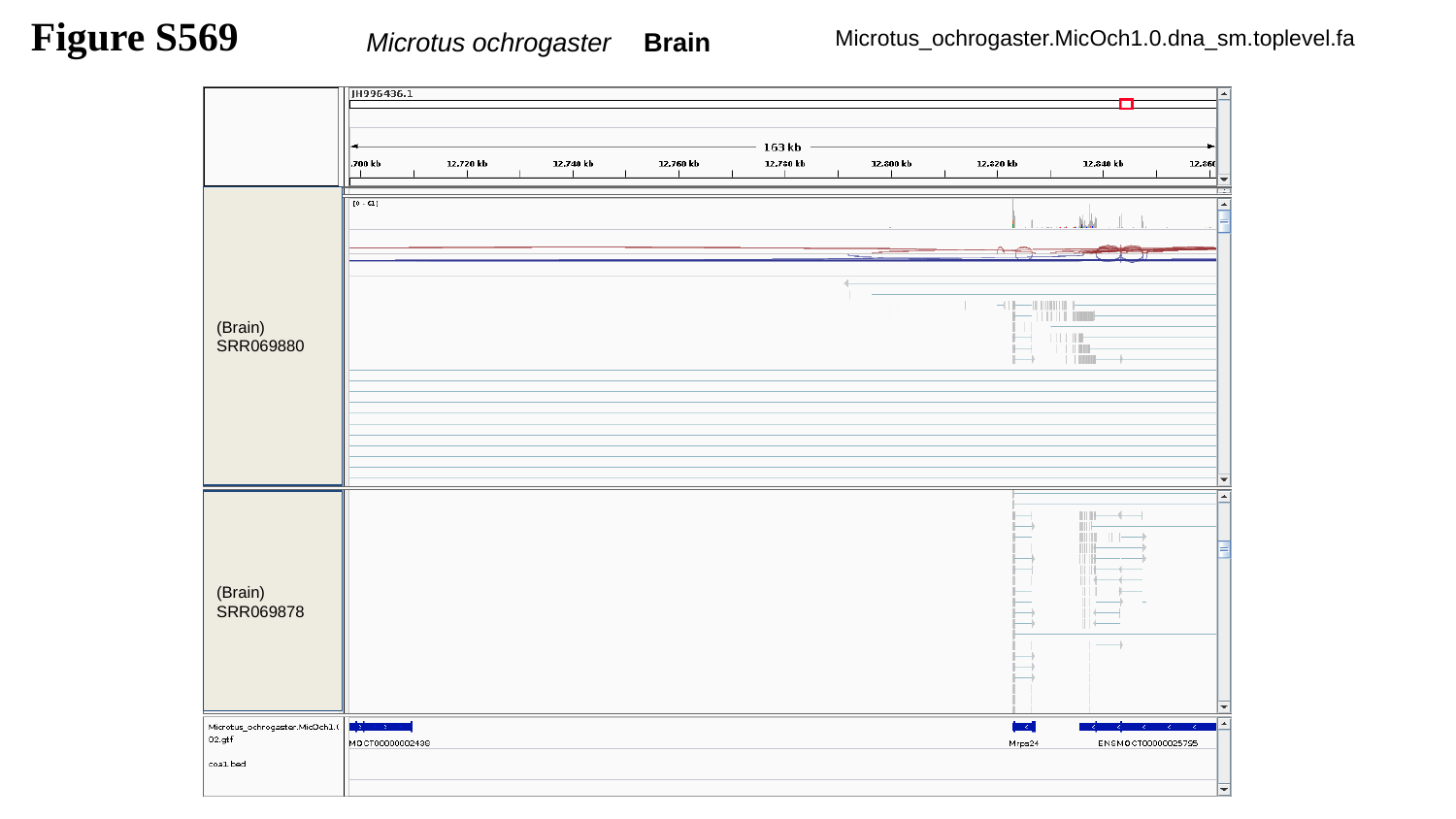

Figure S569
Microtus ochrogaster Brain
Microtus_ochrogaster.MicOch1.0.dna_sm.toplevel.fa
(Brain)
SRR069880
(Brain)
SRR069878

### Slide 570
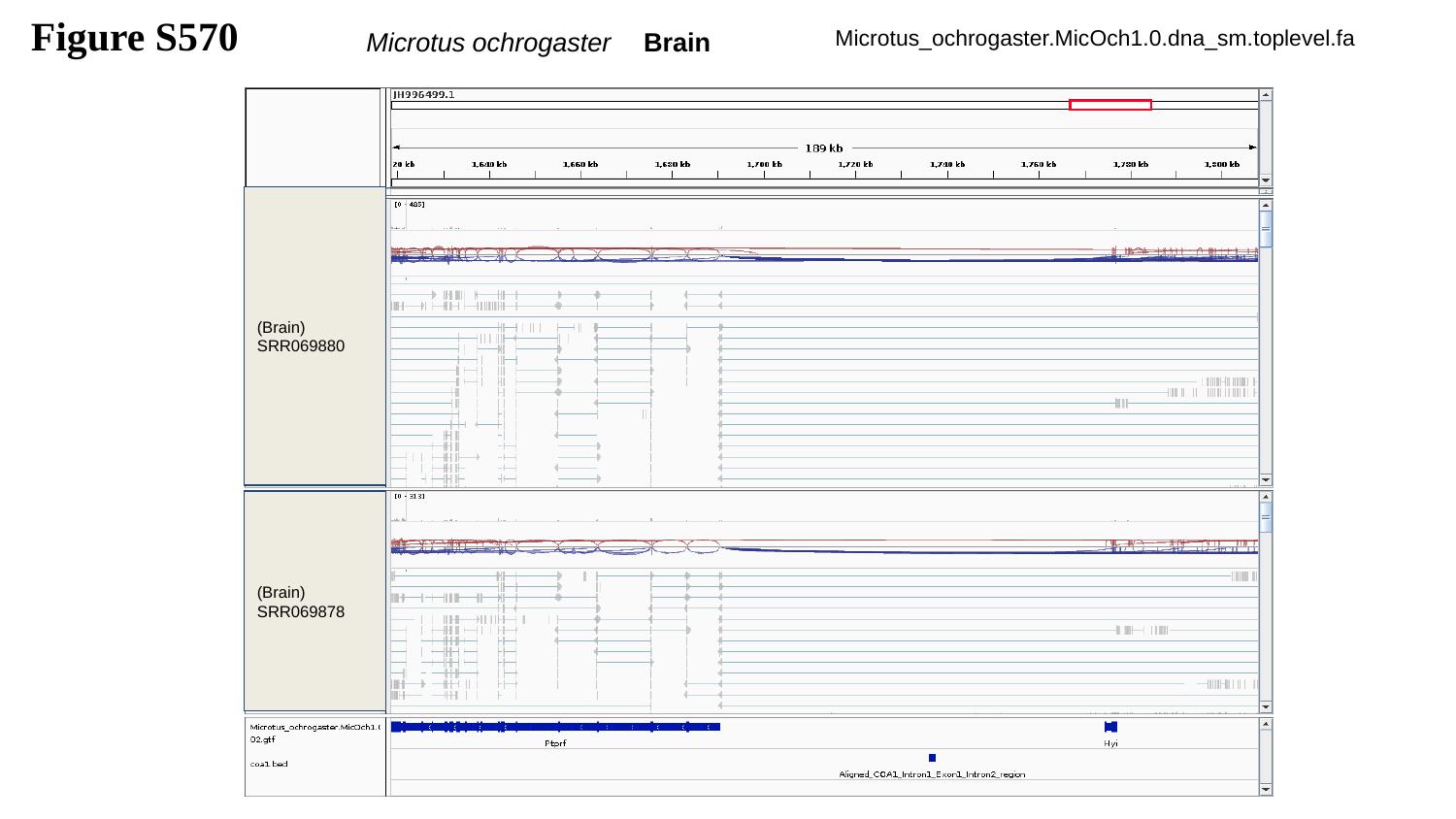

Figure S570
Microtus ochrogaster Brain
Microtus_ochrogaster.MicOch1.0.dna_sm.toplevel.fa
(Brain)
SRR069880
(Brain)
SRR069878

### Slide 571
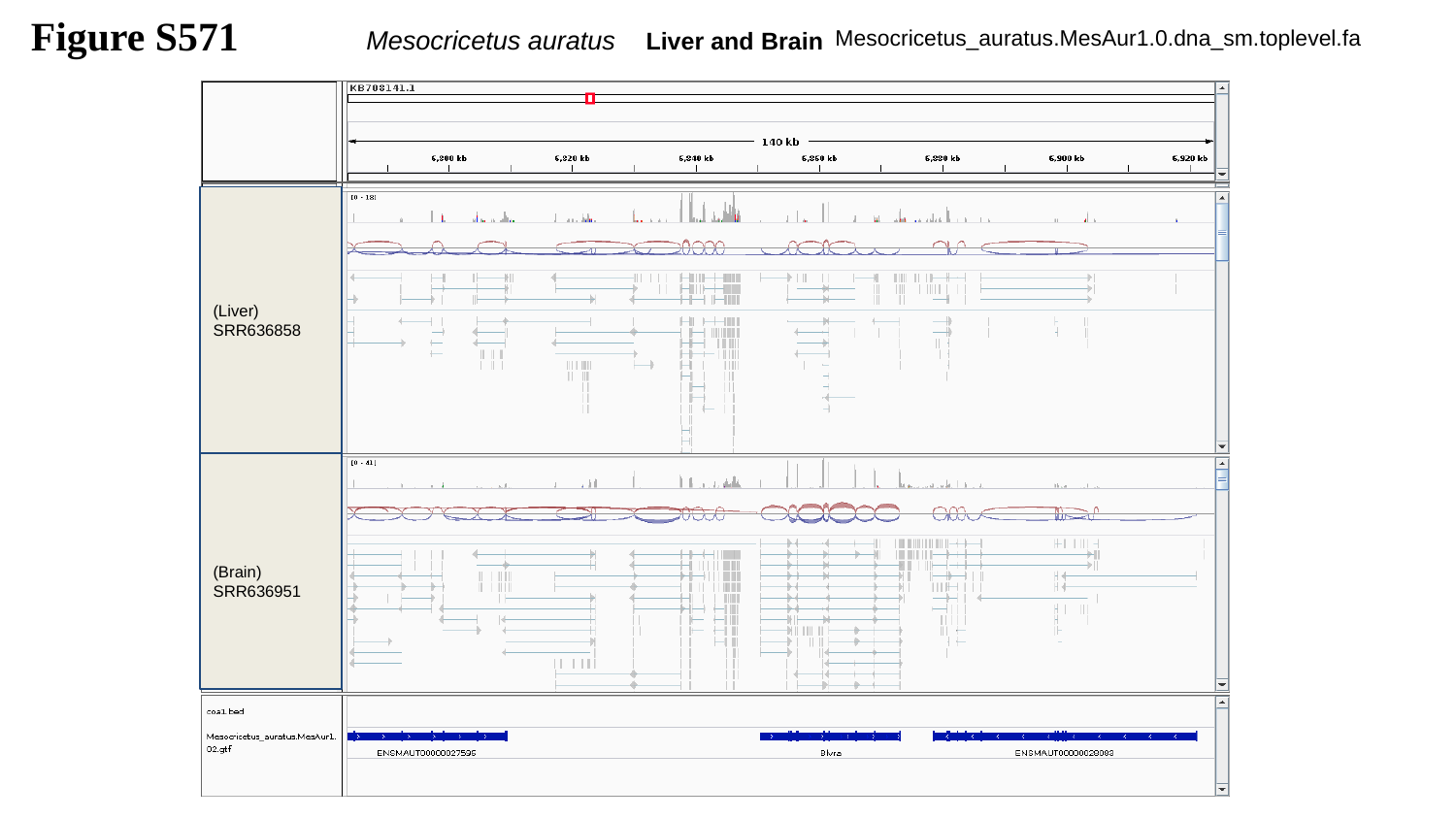

Figure S571
Mesocricetus auratus Liver and Brain
Mesocricetus_auratus.MesAur1.0.dna_sm.toplevel.fa
(Liver)
SRR636858
(Brain)
SRR636951

### Slide 572
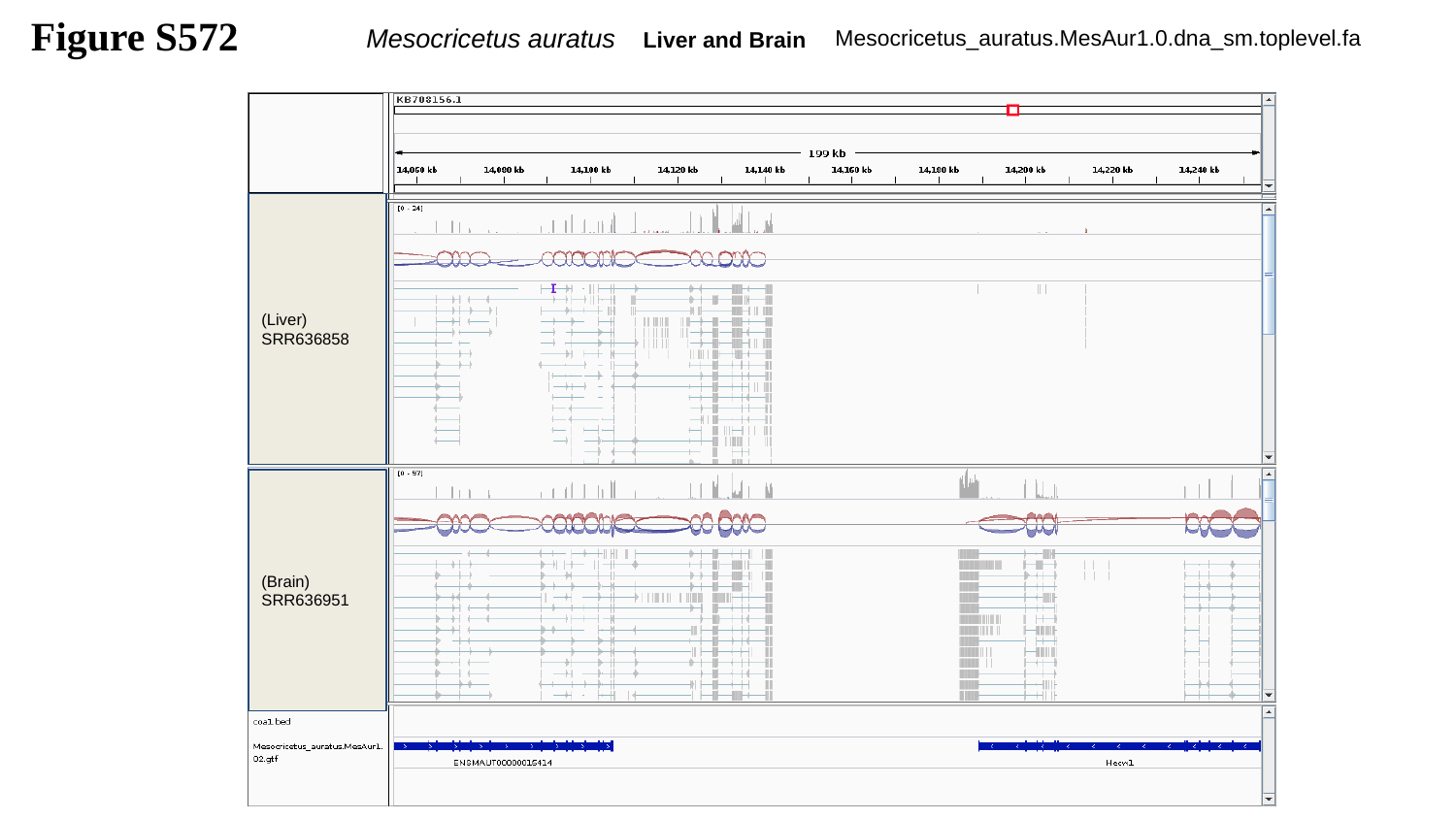

Figure S572
Mesocricetus auratus Liver and Brain
Mesocricetus_auratus.MesAur1.0.dna_sm.toplevel.fa
(Liver)
SRR636858
(Brain)
SRR636951

### Slide 573
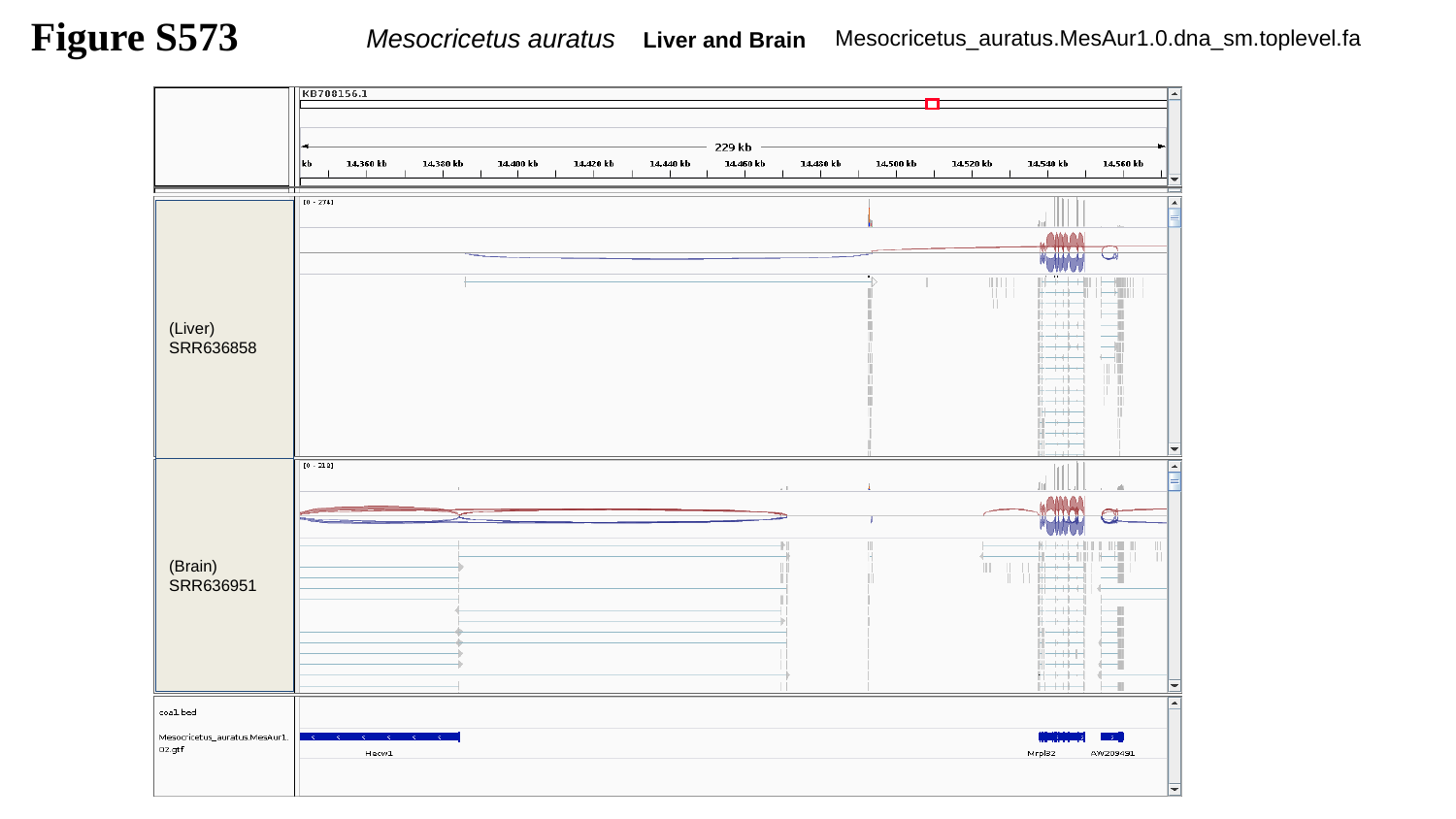

Figure S573
Mesocricetus auratus Liver and Brain
Mesocricetus_auratus.MesAur1.0.dna_sm.toplevel.fa
(Liver)
SRR636858
(Brain)
SRR636951

### Slide 574
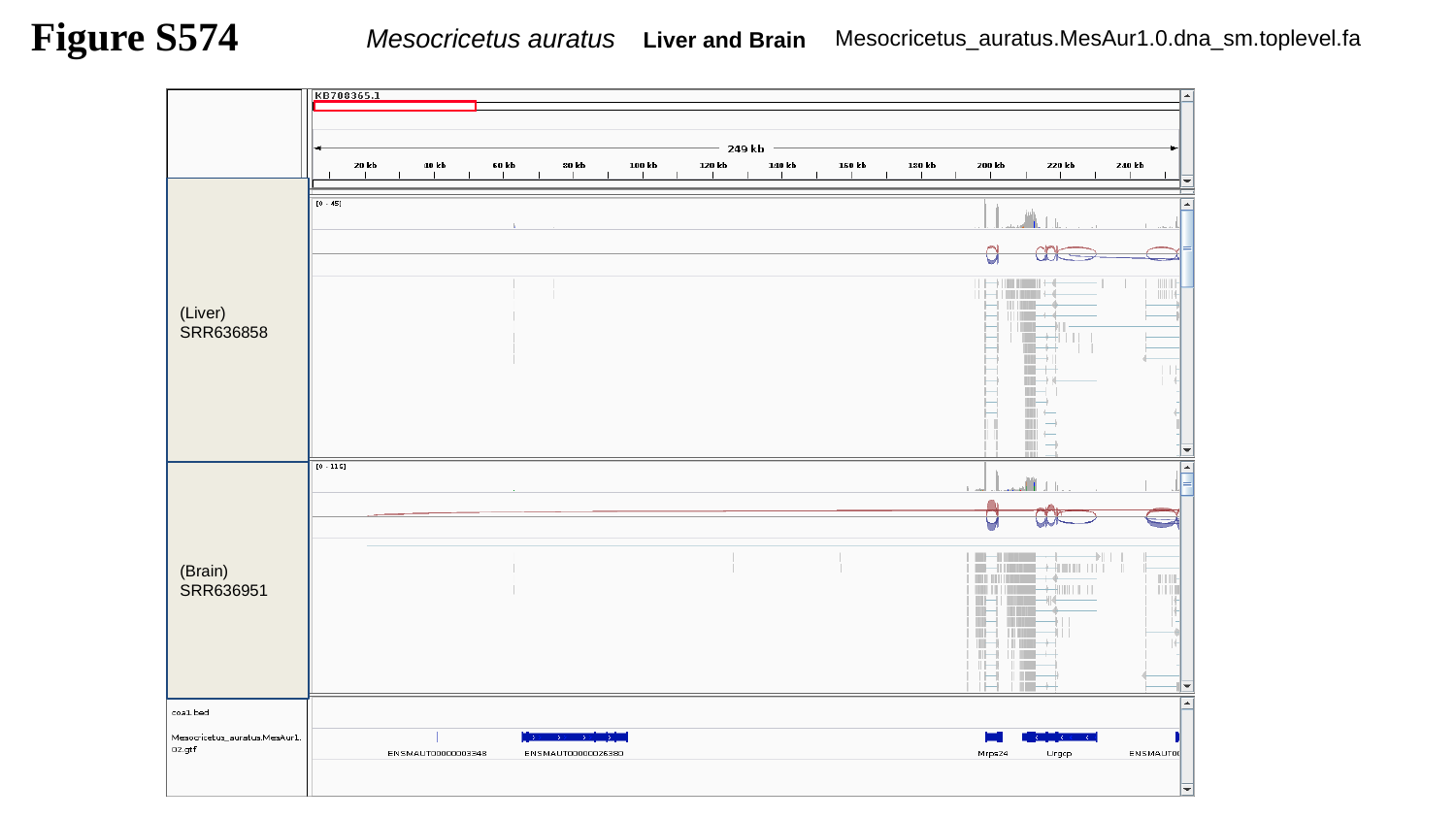

Figure S574
Mesocricetus auratus Liver and Brain
Mesocricetus_auratus.MesAur1.0.dna_sm.toplevel.fa
(Liver)
SRR636858
(Brain)
SRR636951

### Slide 575
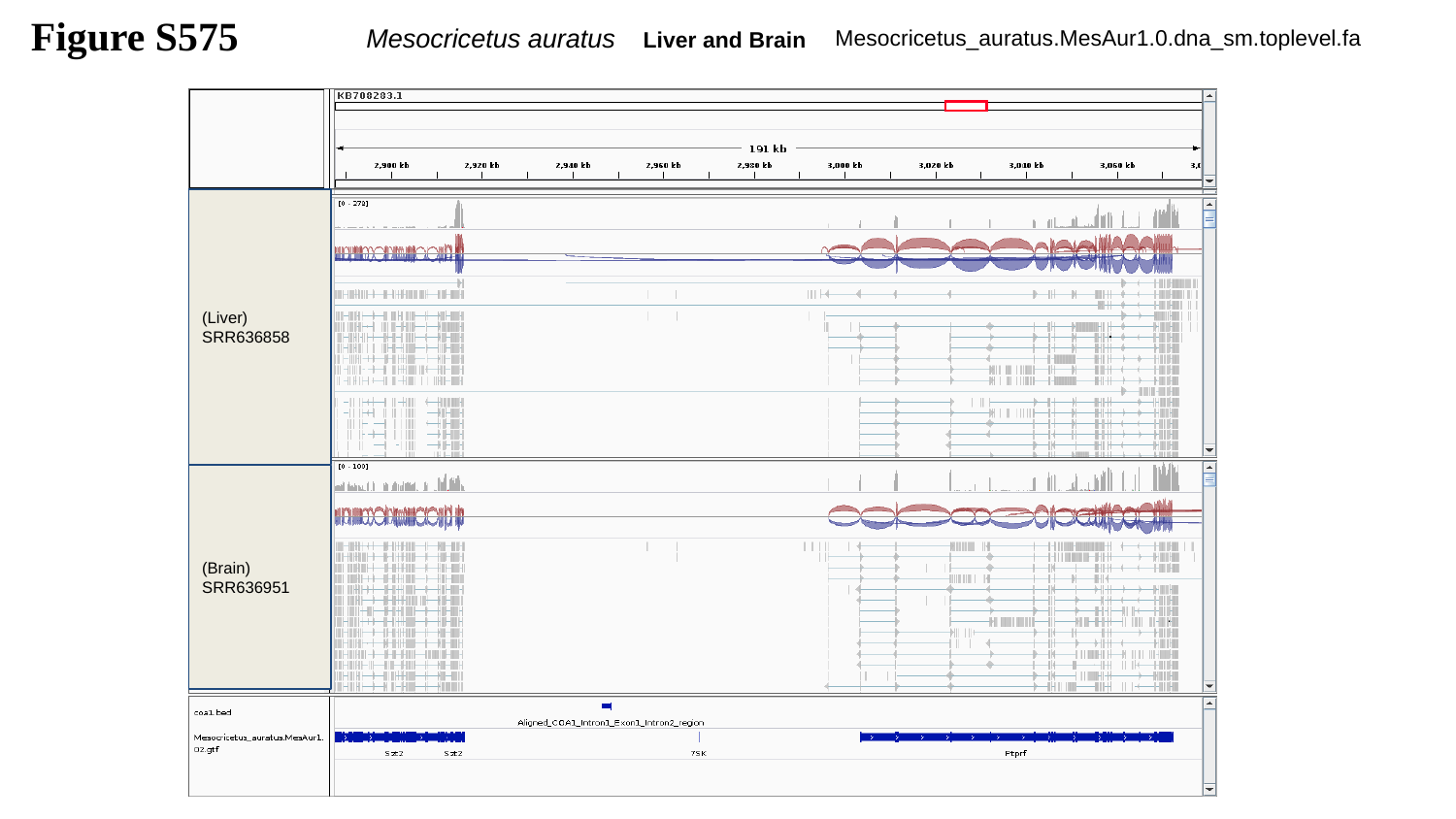

Figure S575
Mesocricetus auratus Liver and Brain
Mesocricetus_auratus.MesAur1.0.dna_sm.toplevel.fa
(Liver)
SRR636858
(Brain)
SRR636951

### Slide 576
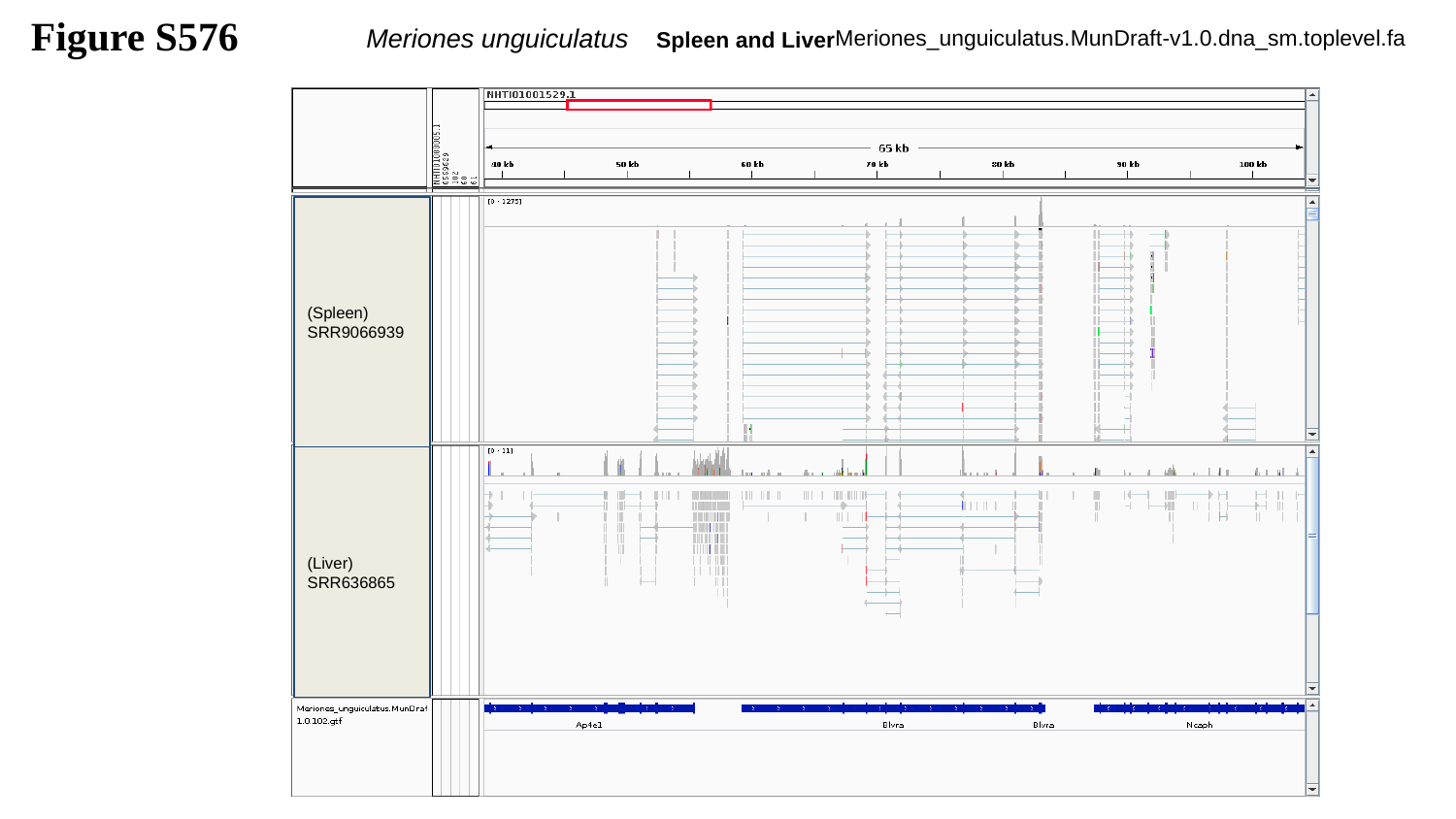

Figure S576
Meriones unguiculatus Spleen and Liver
Meriones_unguiculatus.MunDraft-v1.0.dna_sm.toplevel.fa
(Spleen)
SRR9066939
(Liver)
SRR636865

### Slide 577
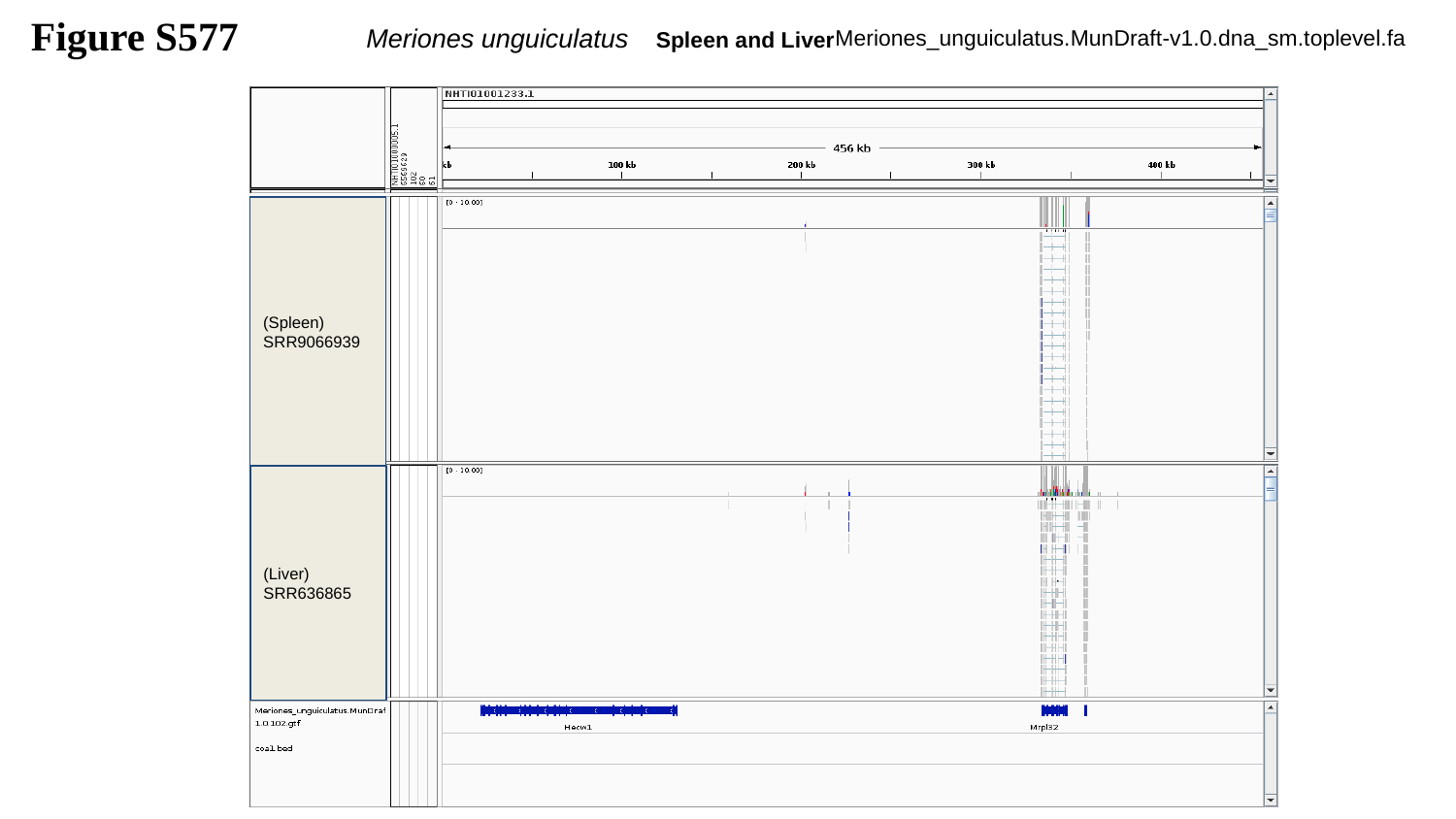

Figure S577
Meriones unguiculatus Spleen and Liver
Meriones_unguiculatus.MunDraft-v1.0.dna_sm.toplevel.fa
(Spleen)
SRR9066939
(Liver)
SRR636865

### Slide 578
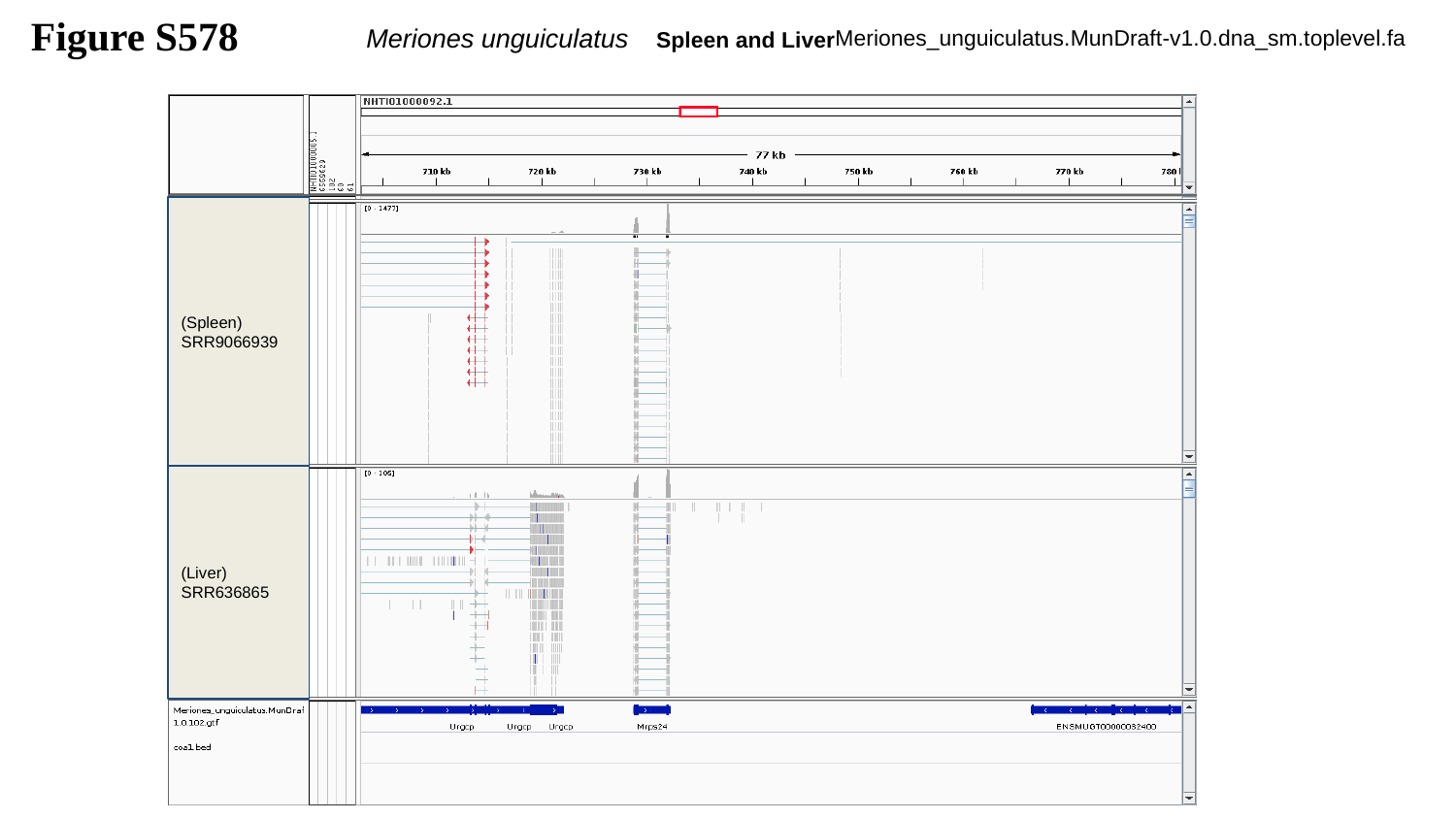

Figure S578
Meriones unguiculatus Spleen and Liver
Meriones_unguiculatus.MunDraft-v1.0.dna_sm.toplevel.fa
(Spleen)
SRR9066939
(Liver)
SRR636865

### Slide 579
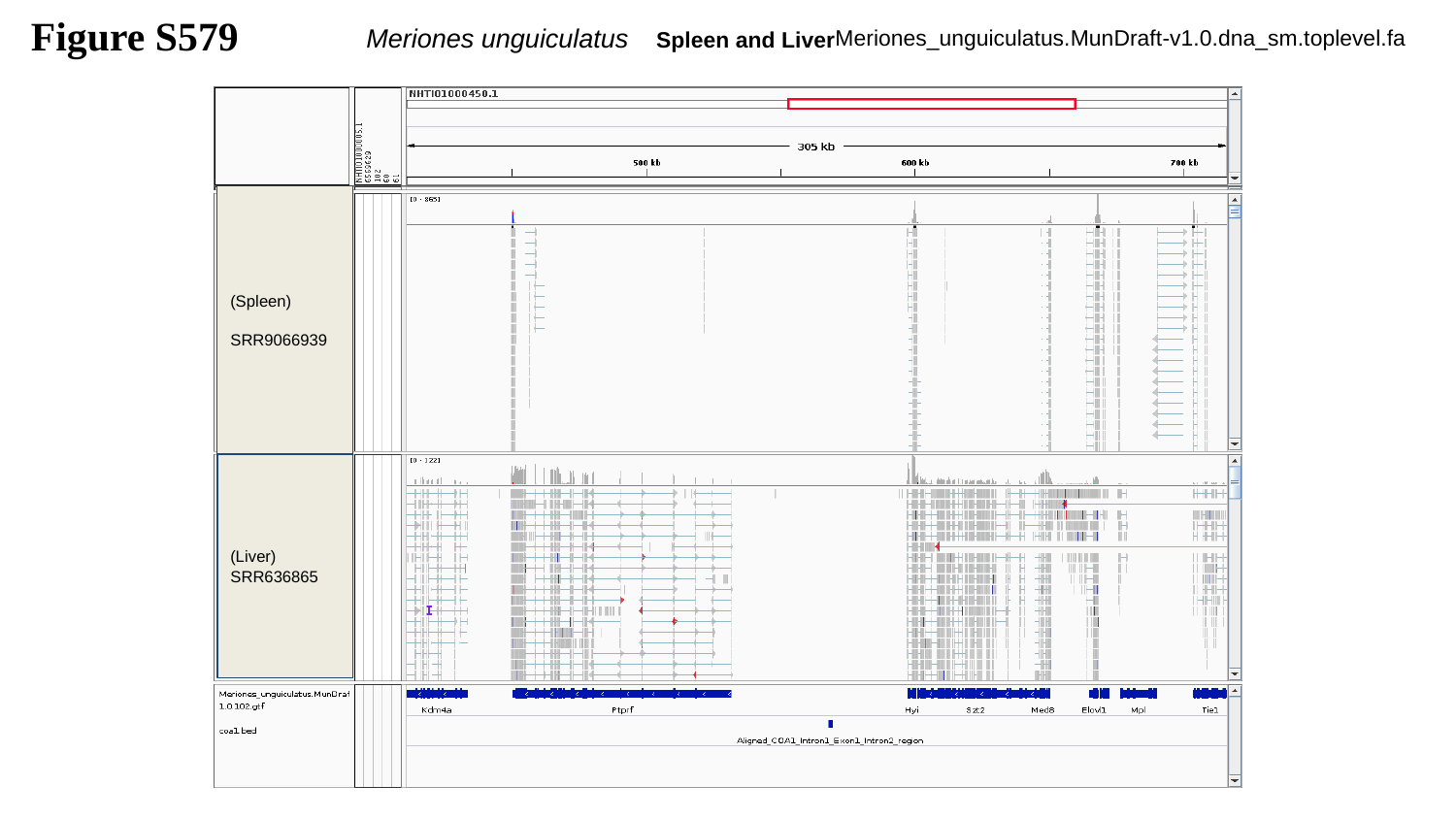

Figure S579
Meriones unguiculatus Spleen and Liver
Meriones_unguiculatus.MunDraft-v1.0.dna_sm.toplevel.fa
(Spleen)
SRR9066939
(Liver)
SRR636865

### Slide 580
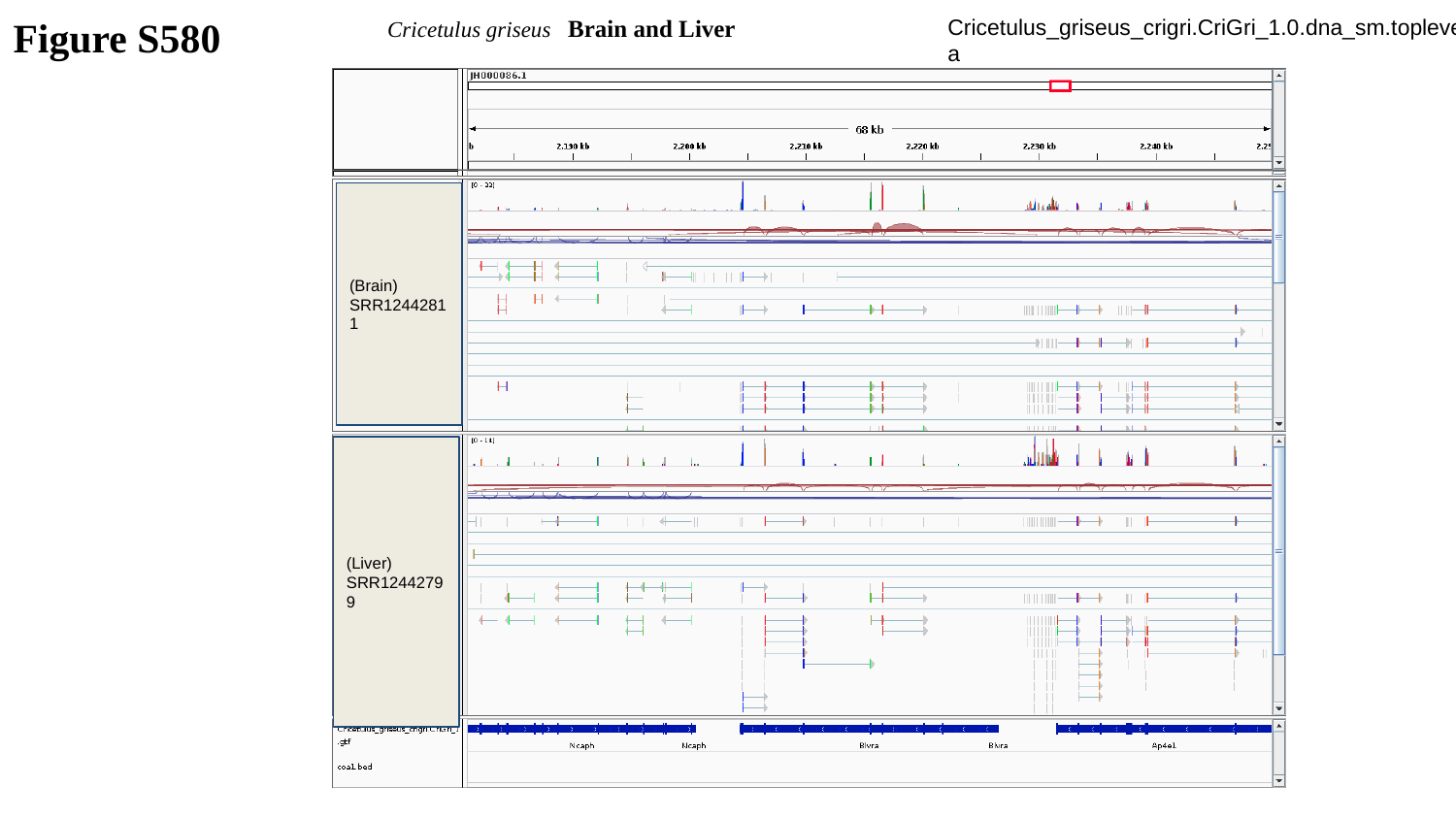

Cricetulus griseus Brain and Liver
Cricetulus_griseus_crigri.CriGri_1.0.dna_sm.toplevel.fa
Figure S580
(Brain)
SRR12442811
(Liver)
SRR12442799

### Slide 581
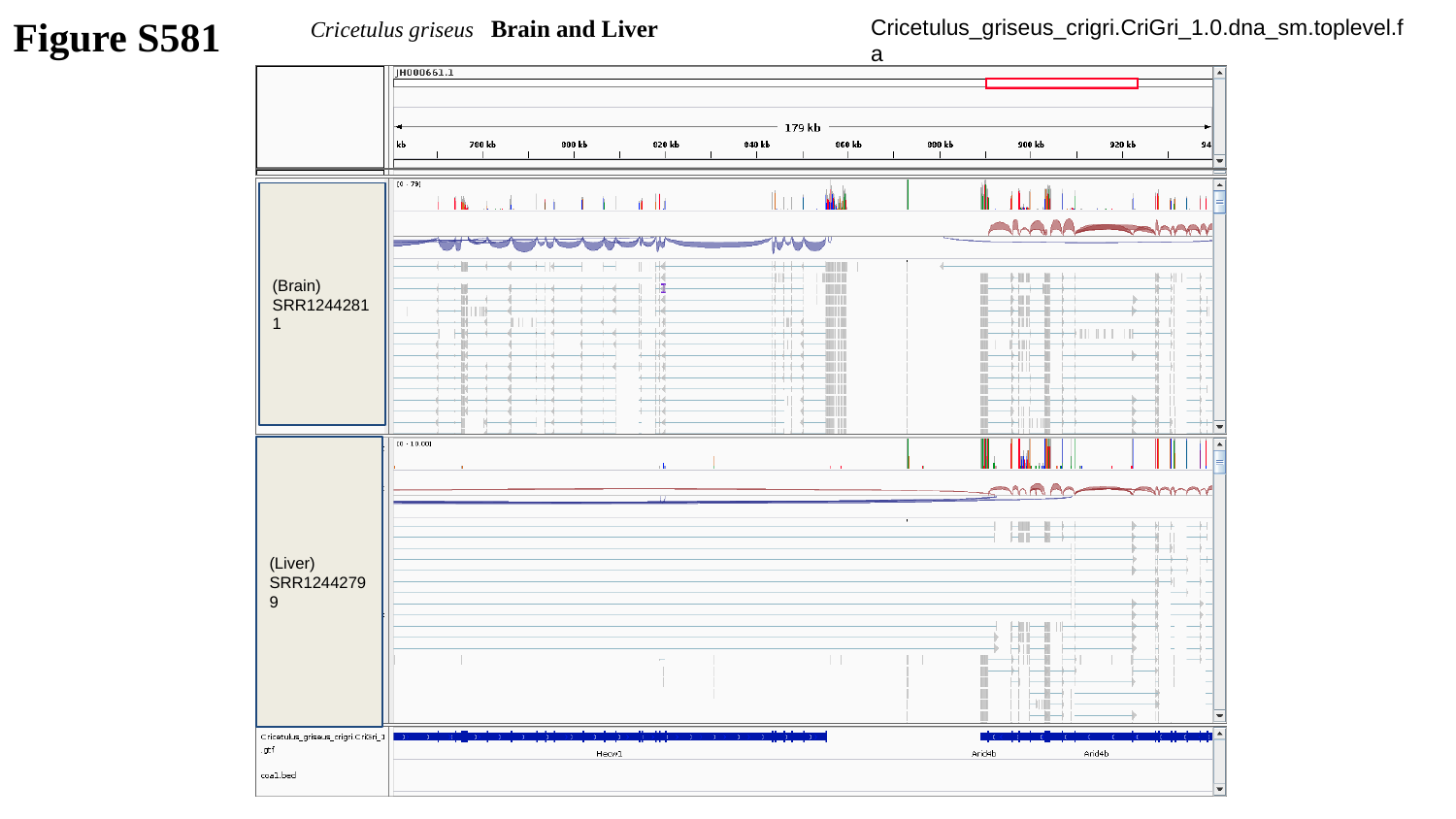

Cricetulus griseus Brain and Liver
Cricetulus_griseus_crigri.CriGri_1.0.dna_sm.toplevel.fa
Figure S581
(Brain)
SRR12442811
(Liver)
SRR12442799

### Slide 582
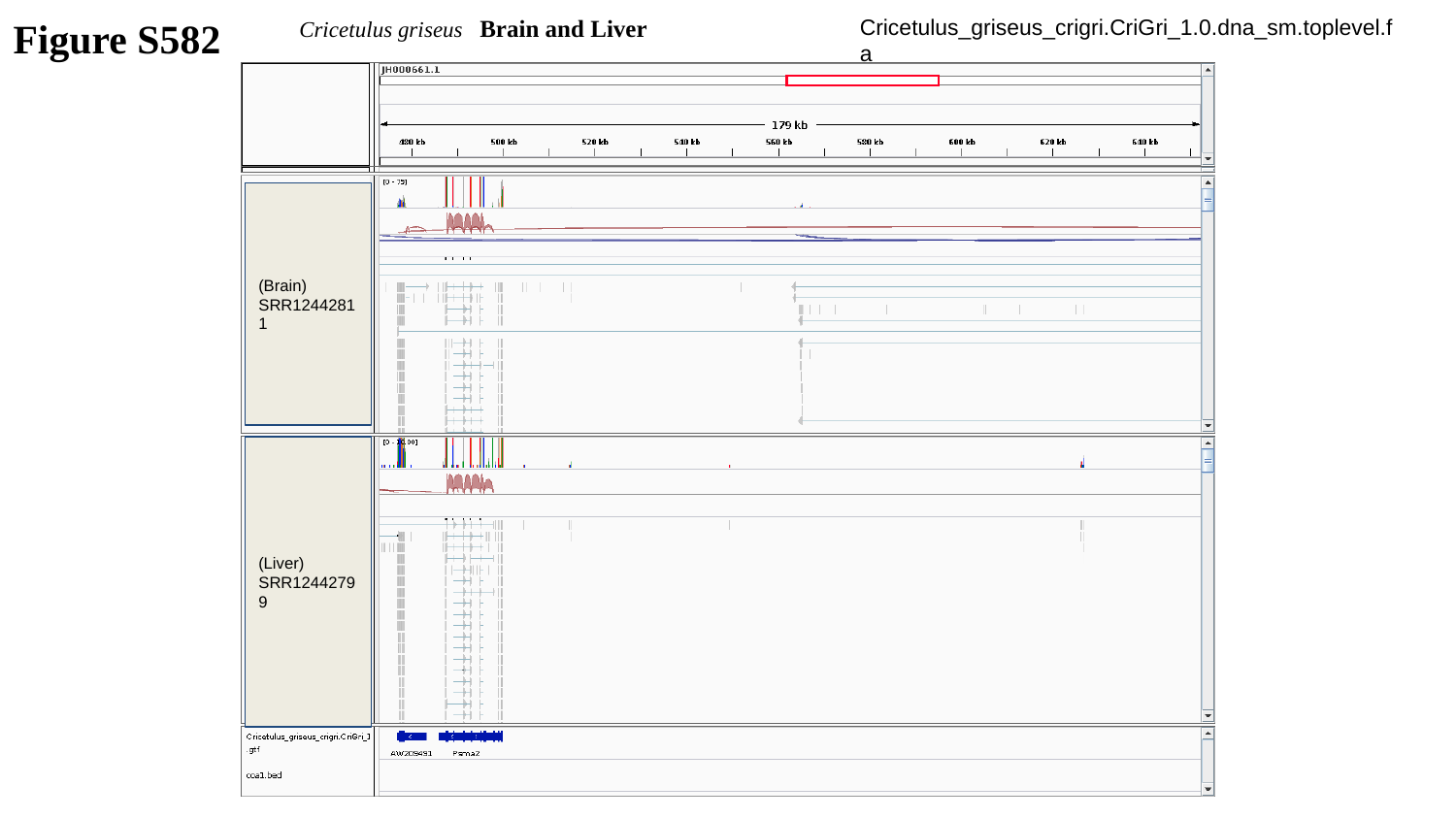

Figure S582
Cricetulus griseus Brain and Liver
Cricetulus_griseus_crigri.CriGri_1.0.dna_sm.toplevel.fa
(Brain)
SRR12442811
(Liver)
SRR12442799

### Slide 583
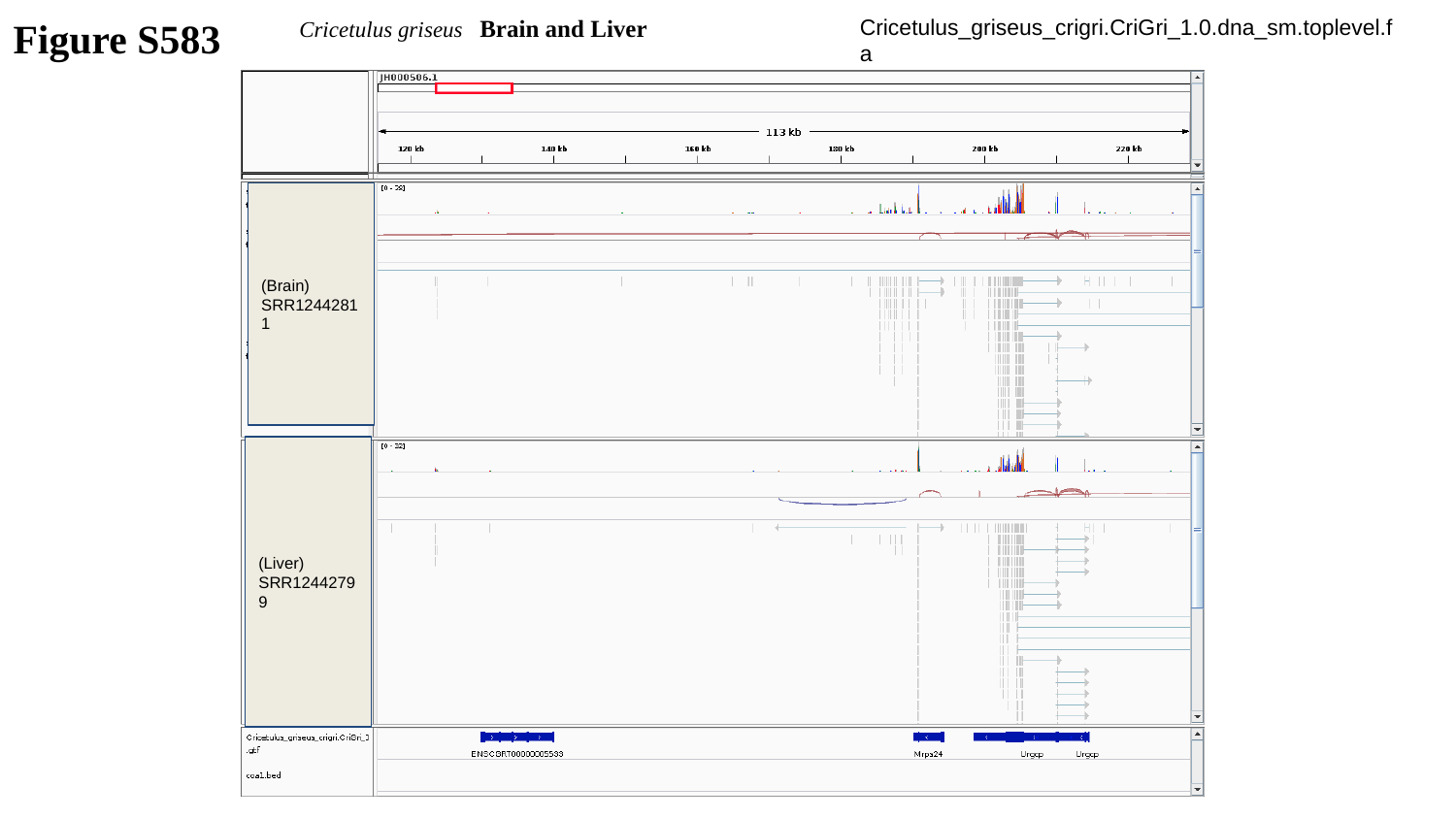

Figure S583
Cricetulus griseus Brain and Liver
Cricetulus_griseus_crigri.CriGri_1.0.dna_sm.toplevel.fa
(Brain)
SRR12442811
(Liver)
SRR12442799

### Slide 584
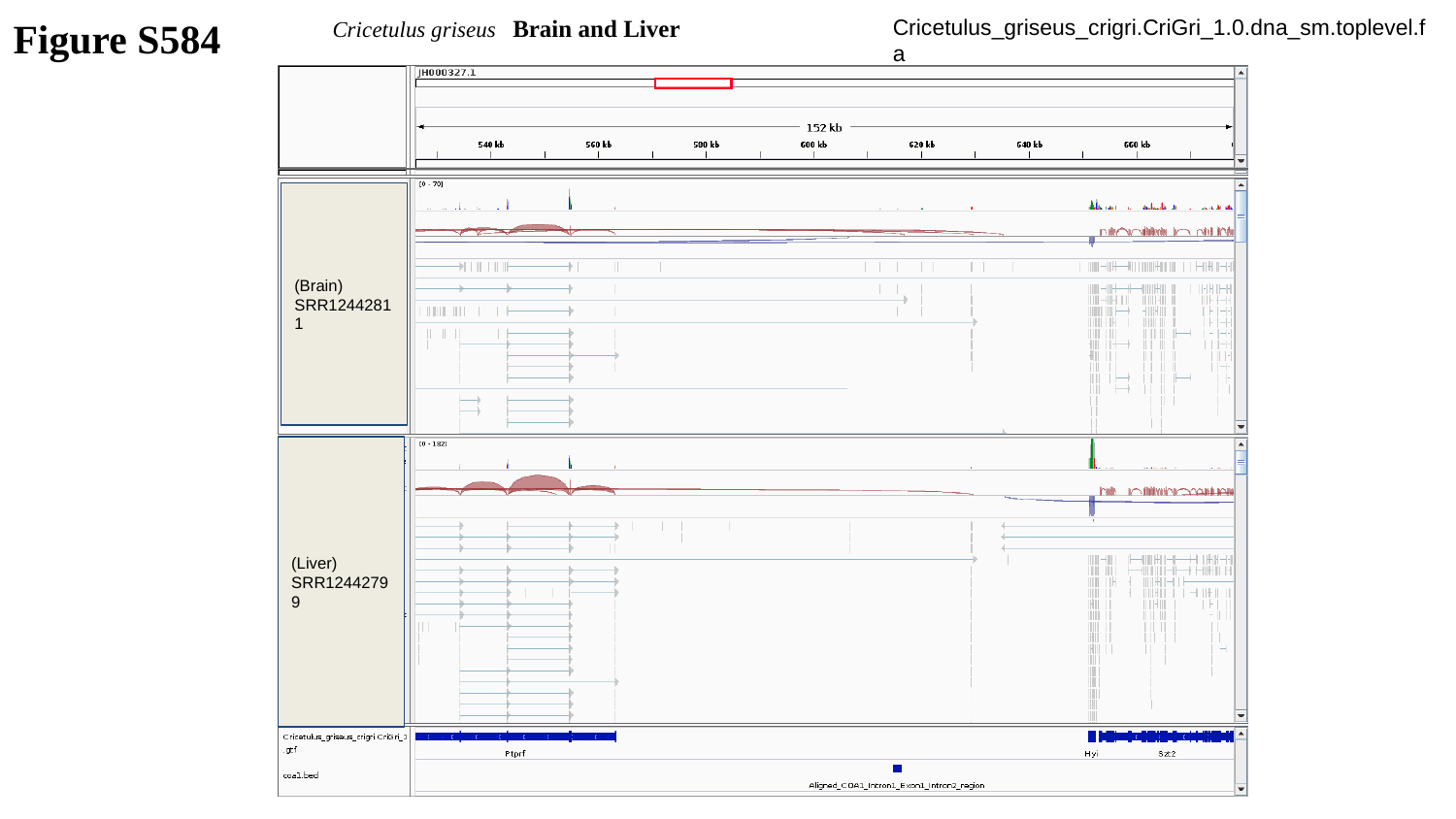

Figure S584
Cricetulus griseus Brain and Liver
Cricetulus_griseus_crigri.CriGri_1.0.dna_sm.toplevel.fa
(Brain)
SRR12442811
(Liver)
SRR12442799

### Slide 585
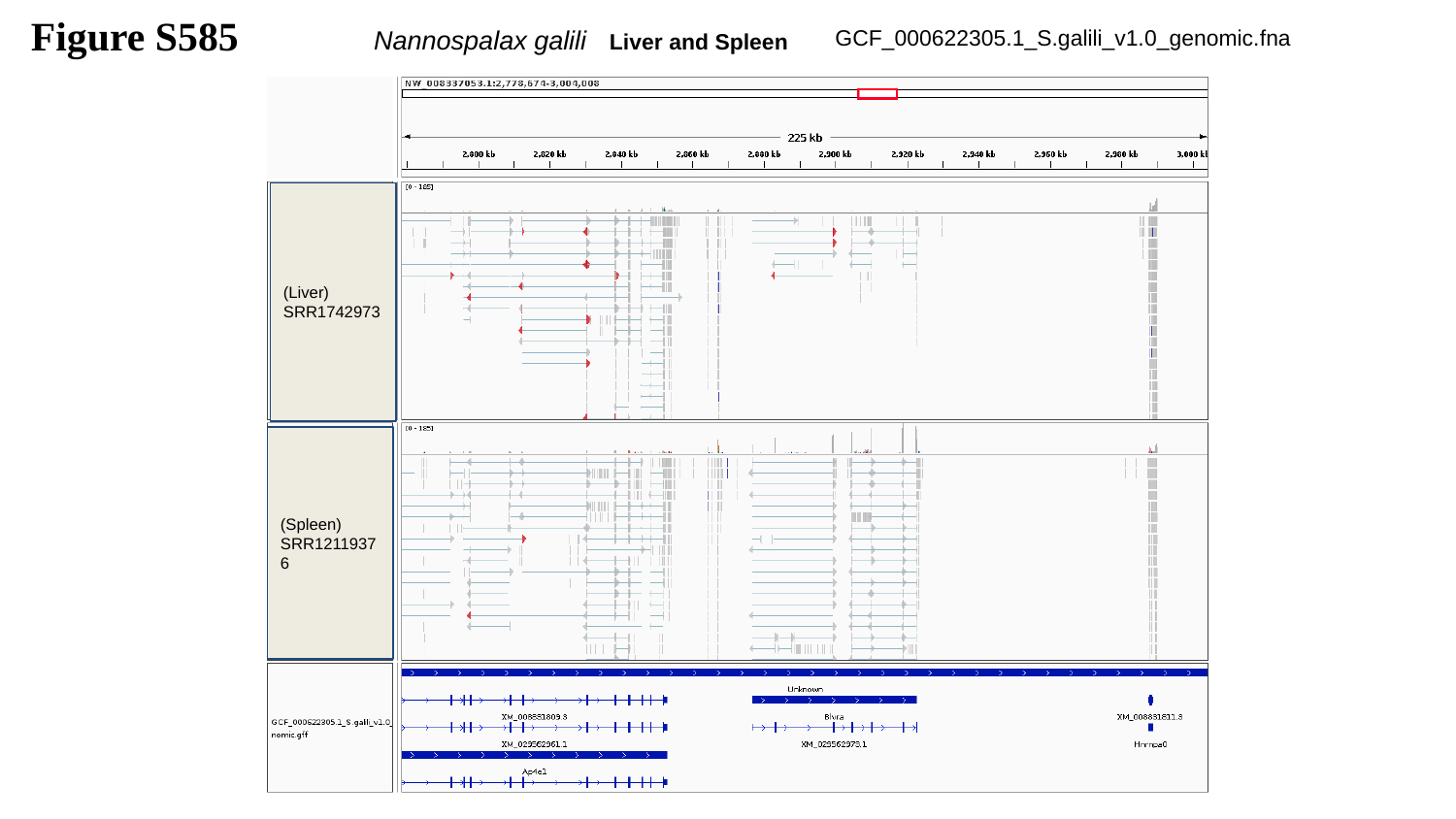

Figure S585
Nannospalax galili Liver and Spleen
GCF_000622305.1_S.galili_v1.0_genomic.fna
(Liver)
SRR1742973
(Spleen)
SRR12119376

### Slide 586
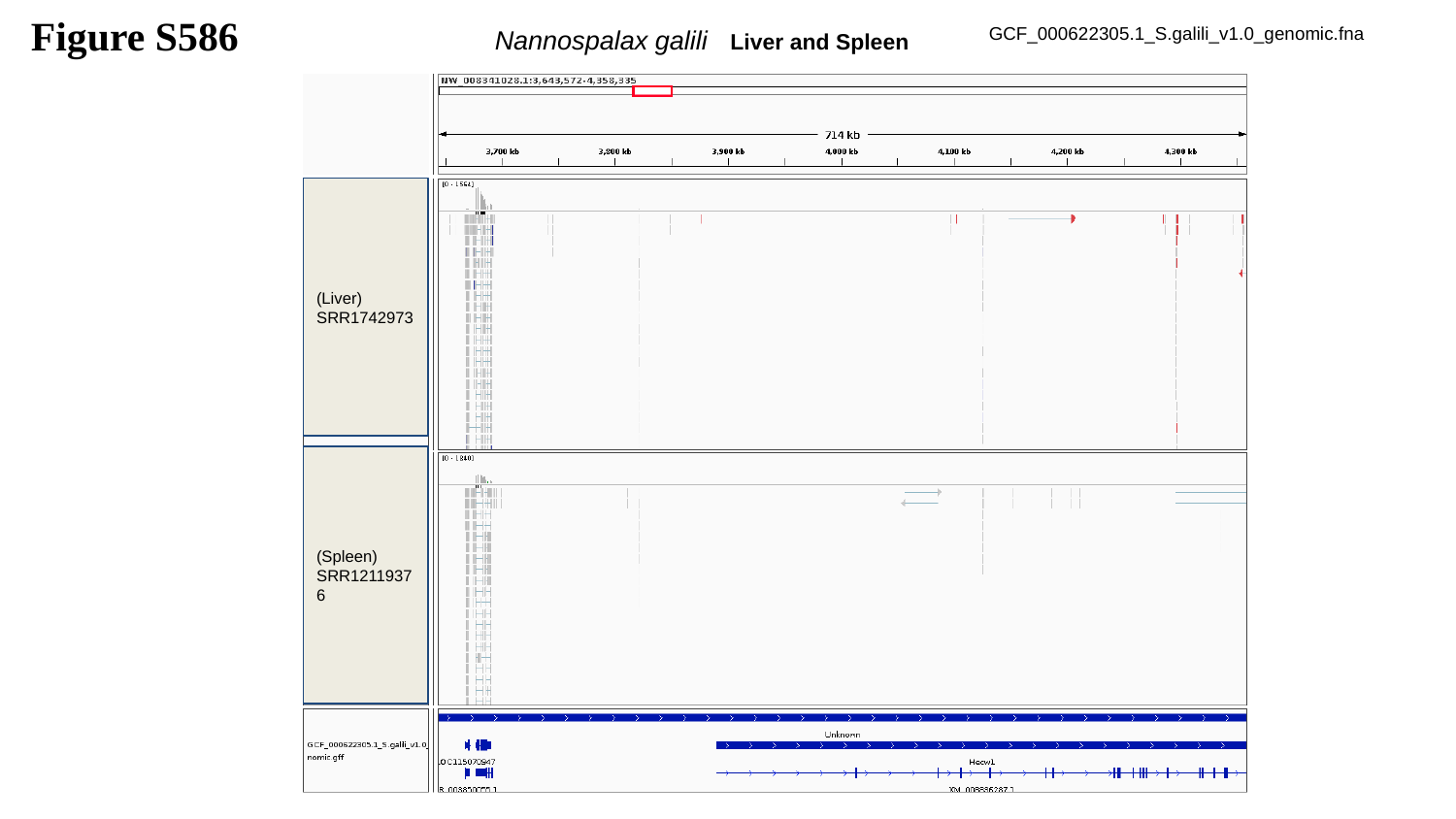

Figure S586
Nannospalax galili Liver and Spleen
GCF_000622305.1_S.galili_v1.0_genomic.fna
(Liver)
SRR1742973
(Spleen)
SRR12119376

### Slide 587
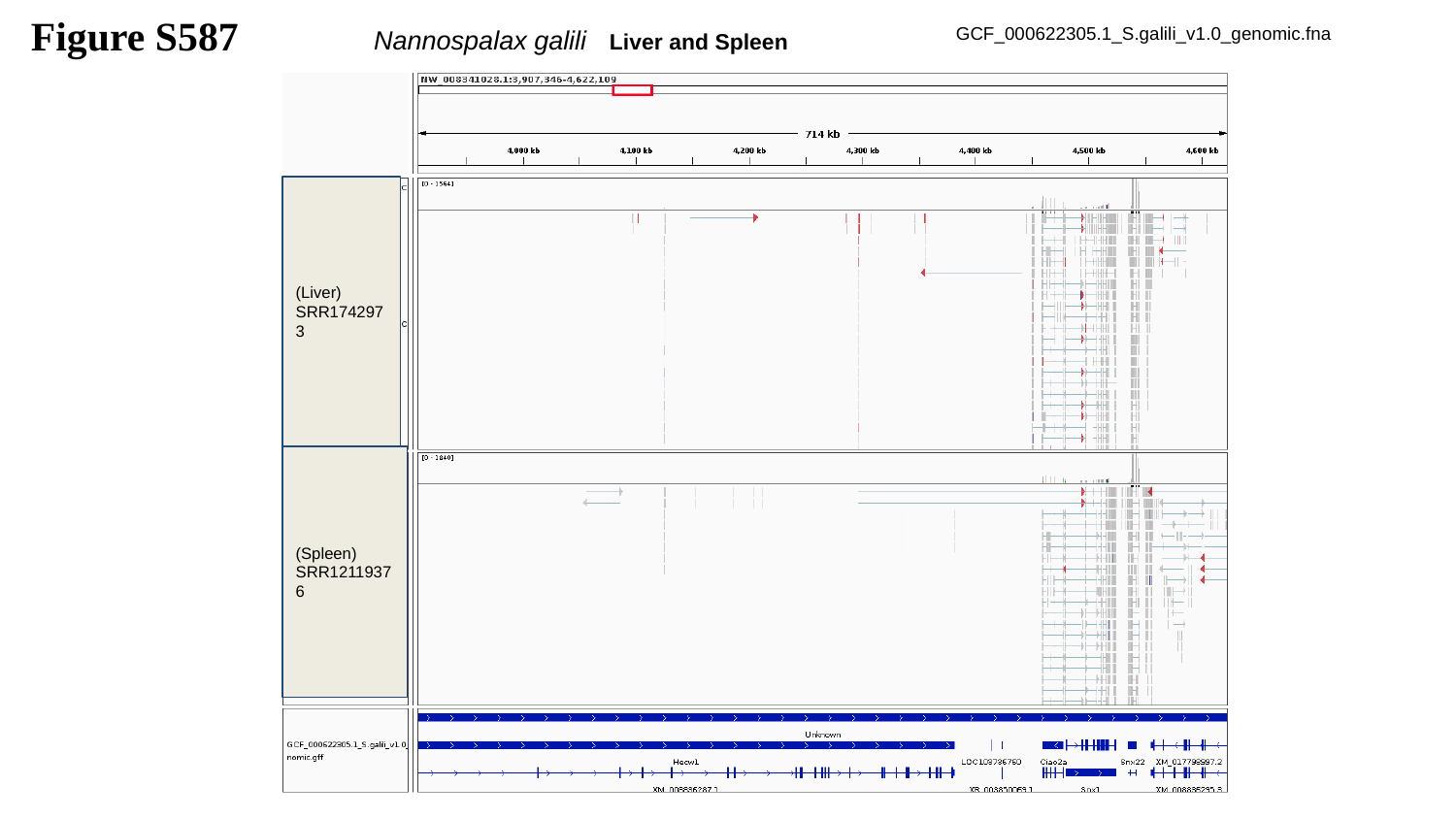

Figure S587
Nannospalax galili Liver and Spleen
GCF_000622305.1_S.galili_v1.0_genomic.fna
(Liver)
SRR1742973
(Spleen)
SRR12119376

### Slide 588
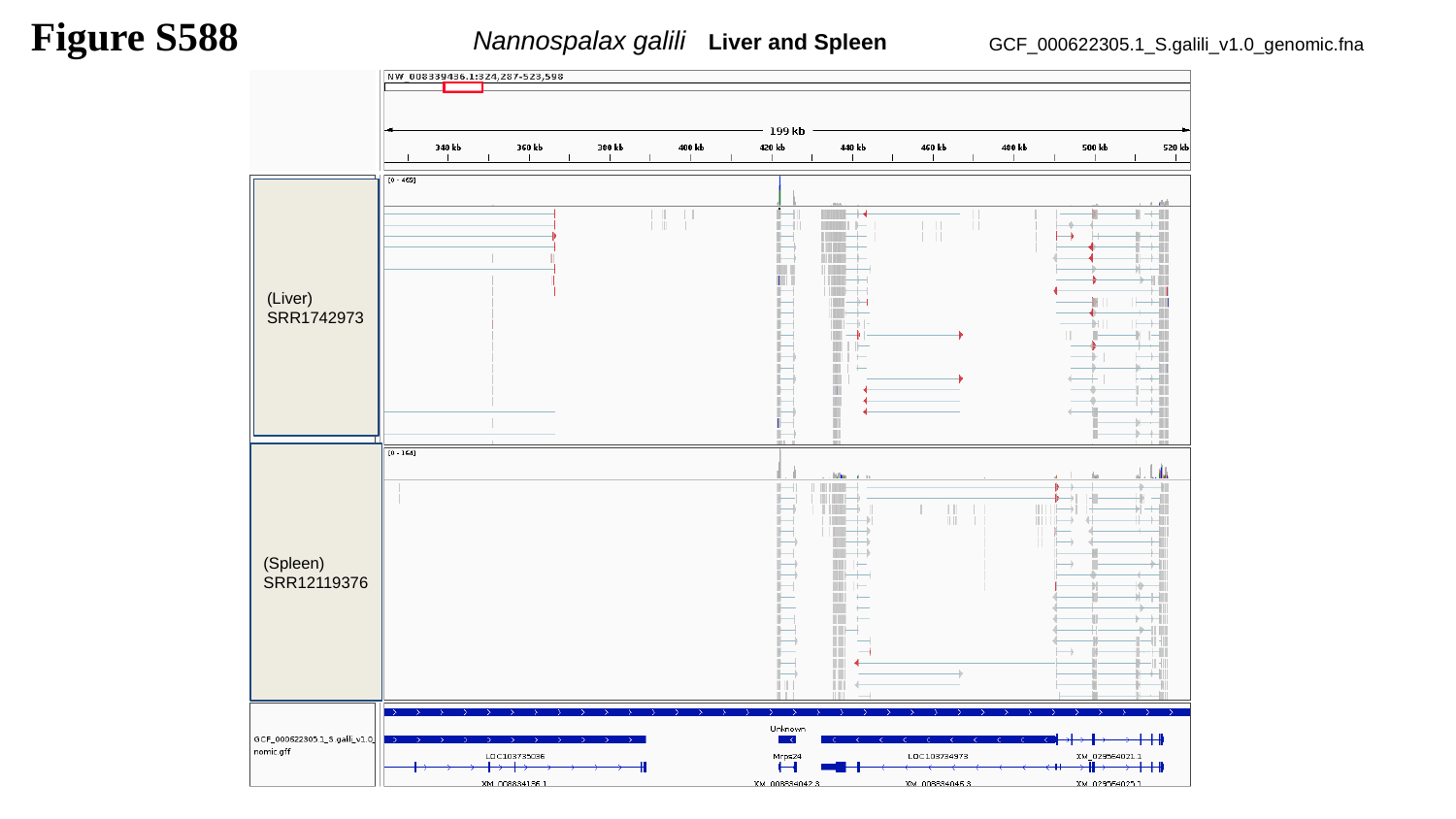

Figure S588
Nannospalax galili Liver and Spleen
GCF_000622305.1_S.galili_v1.0_genomic.fna
(Liver)
SRR1742973
(Spleen)
SRR12119376

### Slide 589
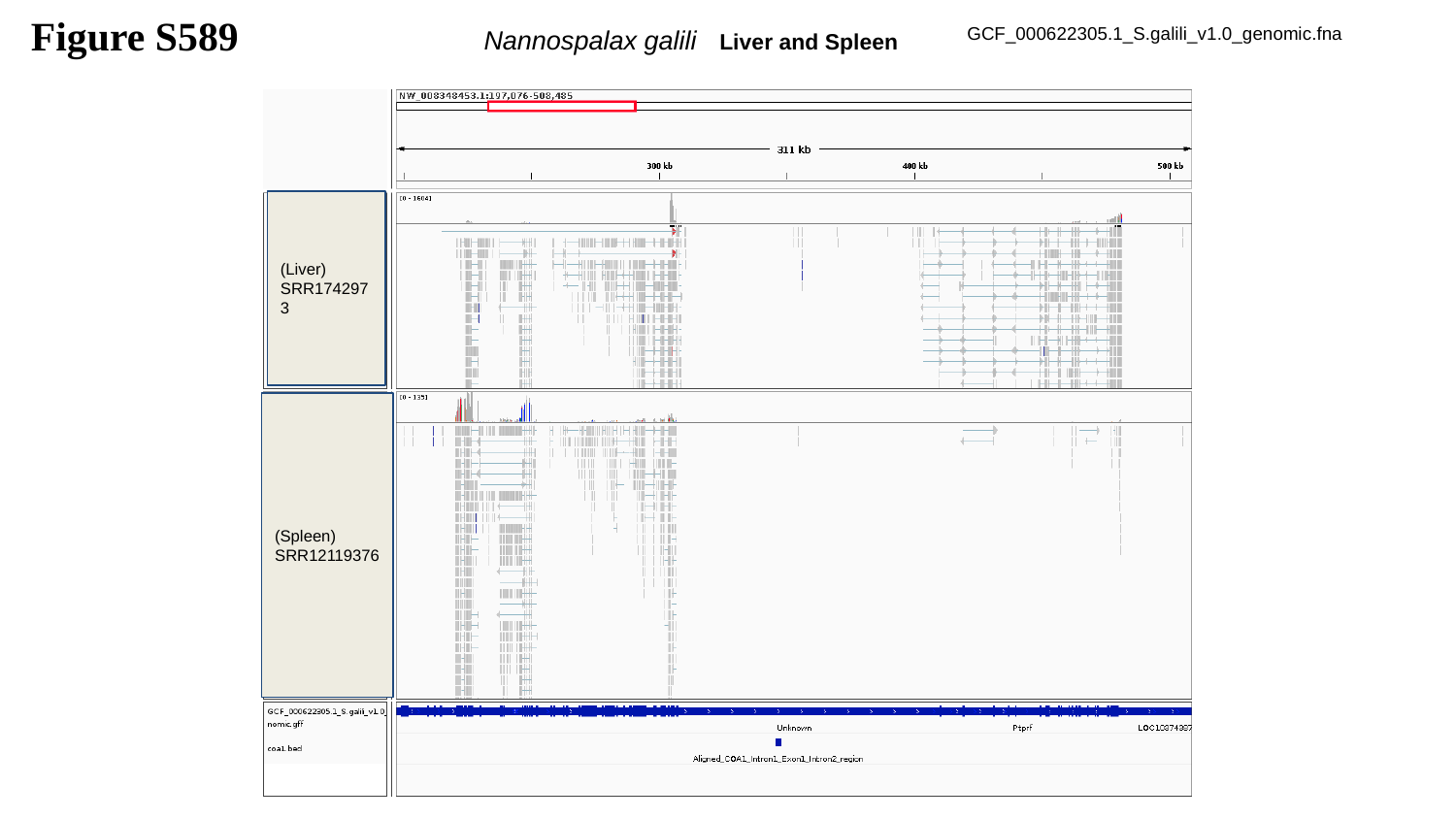

Figure S589
Nannospalax galili Liver and Spleen
GCF_000622305.1_S.galili_v1.0_genomic.fna
(Liver)
SRR1742973
(Spleen)
SRR12119376

### Slide 590
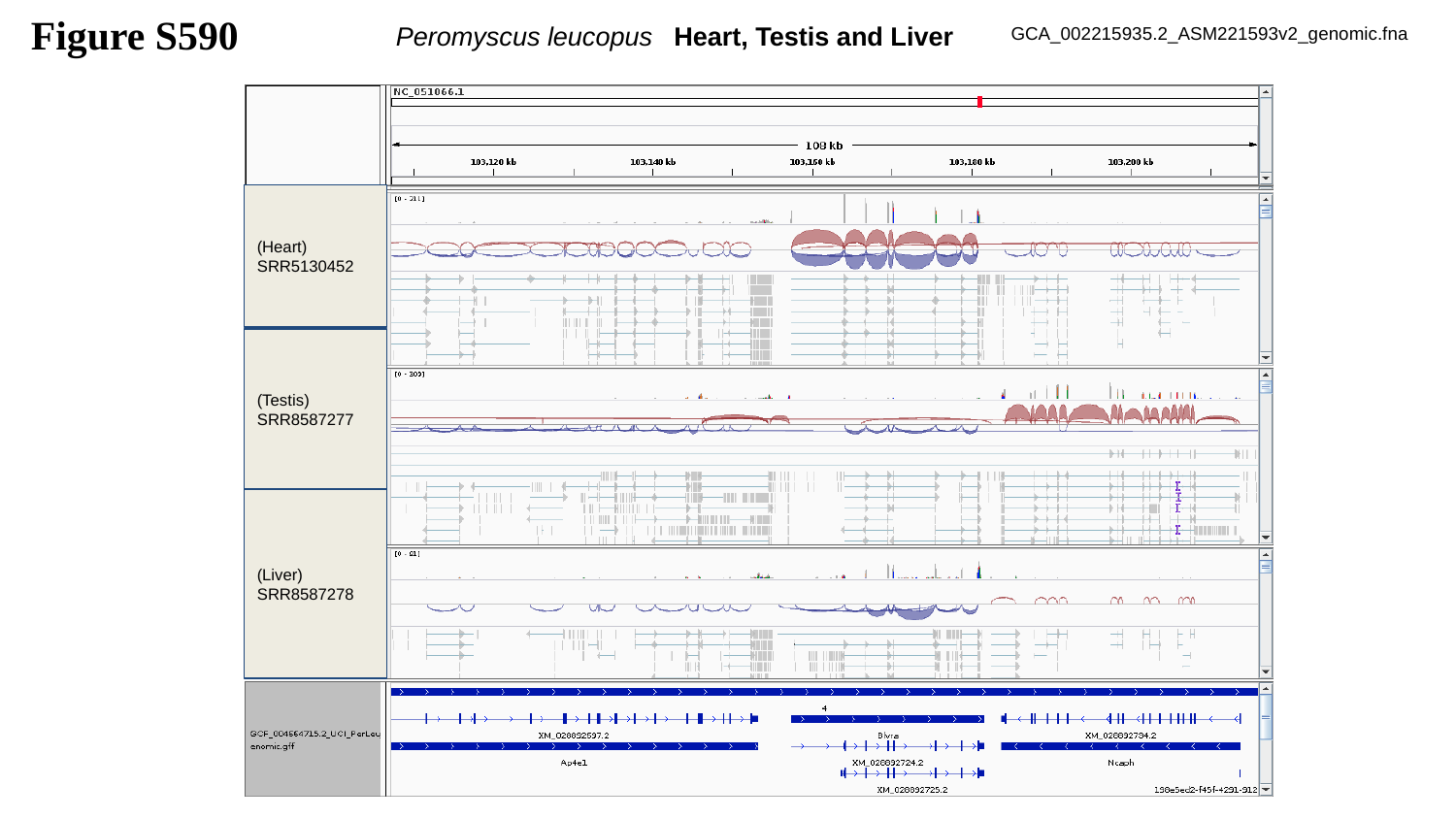

Figure S590
Peromyscus leucopus Heart, Testis and Liver
GCA_002215935.2_ASM221593v2_genomic.fna
(Heart)
SRR5130452
(Testis)
SRR8587277
(Liver)
SRR8587278

### Slide 591
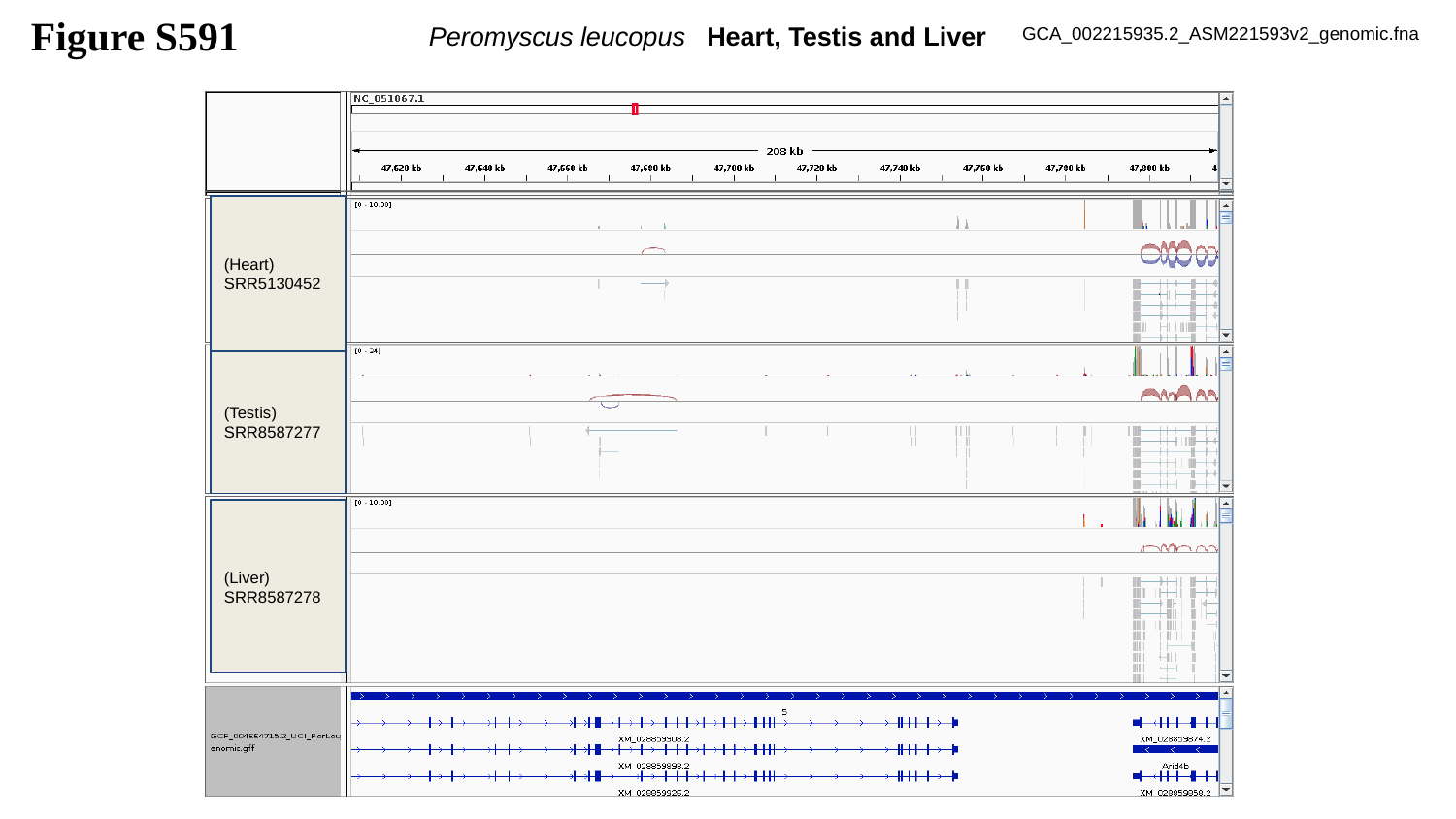

Figure S591
Peromyscus leucopus Heart, Testis and Liver
GCA_002215935.2_ASM221593v2_genomic.fna
(Heart)
SRR5130452
(Testis)
SRR8587277
(Liver)
SRR8587278

### Slide 592
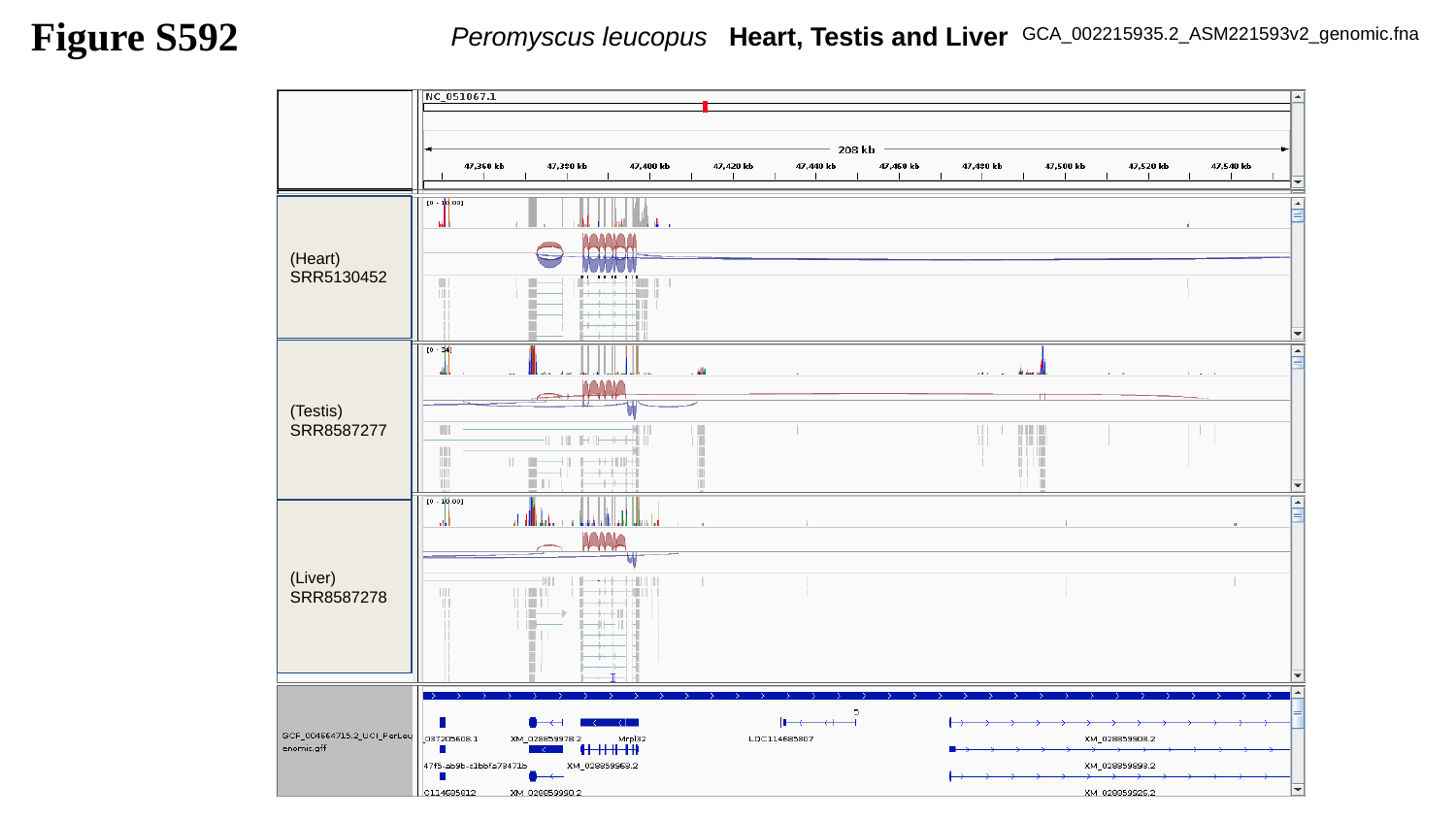

Figure S592
Peromyscus leucopus Heart, Testis and Liver
GCA_002215935.2_ASM221593v2_genomic.fna
(Heart)
SRR5130452
(Testis)
SRR8587277
(Liver)
SRR8587278

### Slide 593
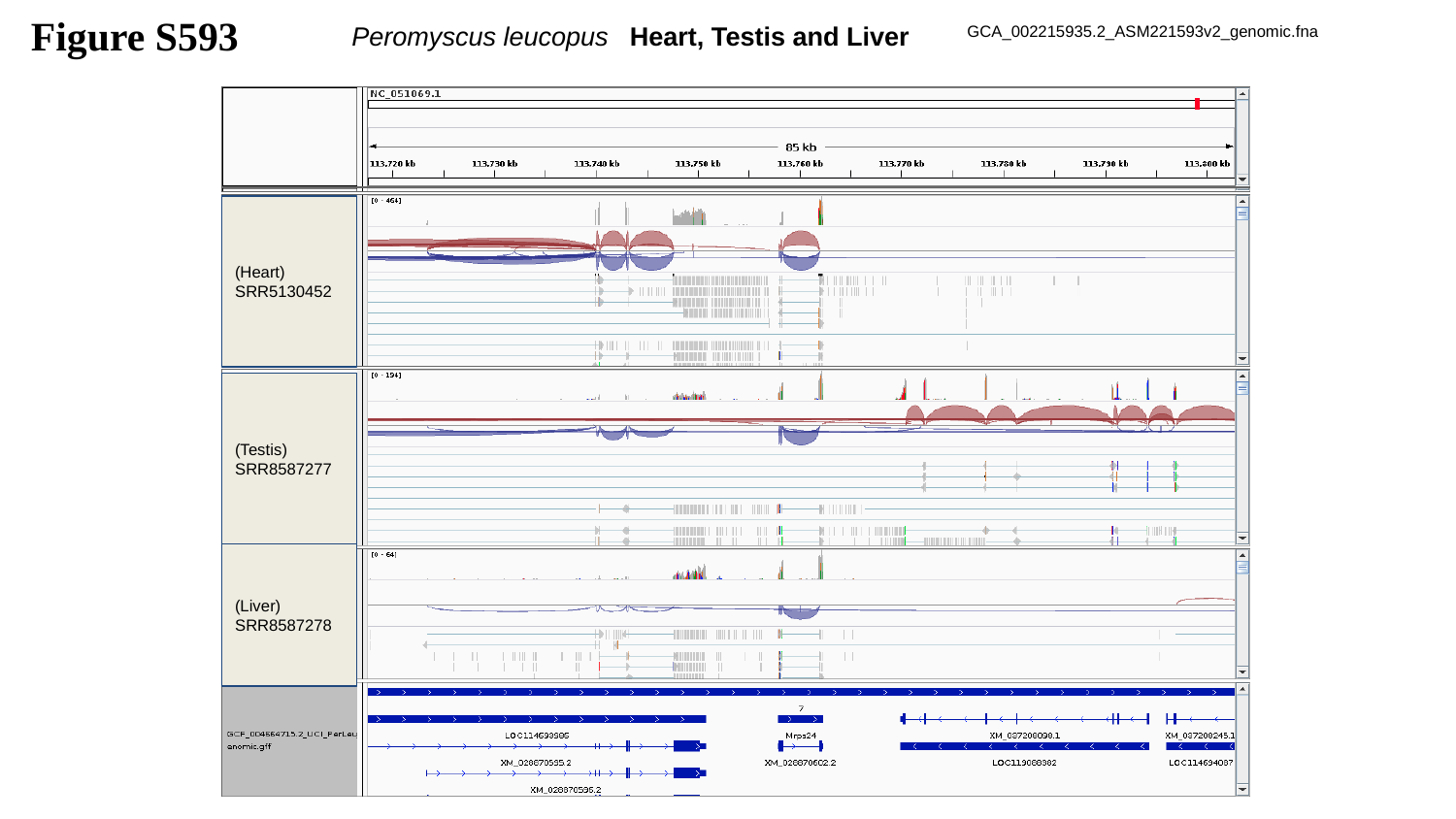

Figure S593
Peromyscus leucopus Heart, Testis and Liver
GCA_002215935.2_ASM221593v2_genomic.fna
(Heart)
SRR5130452
(Testis)
SRR8587277
(Liver)
SRR8587278

### Slide 594
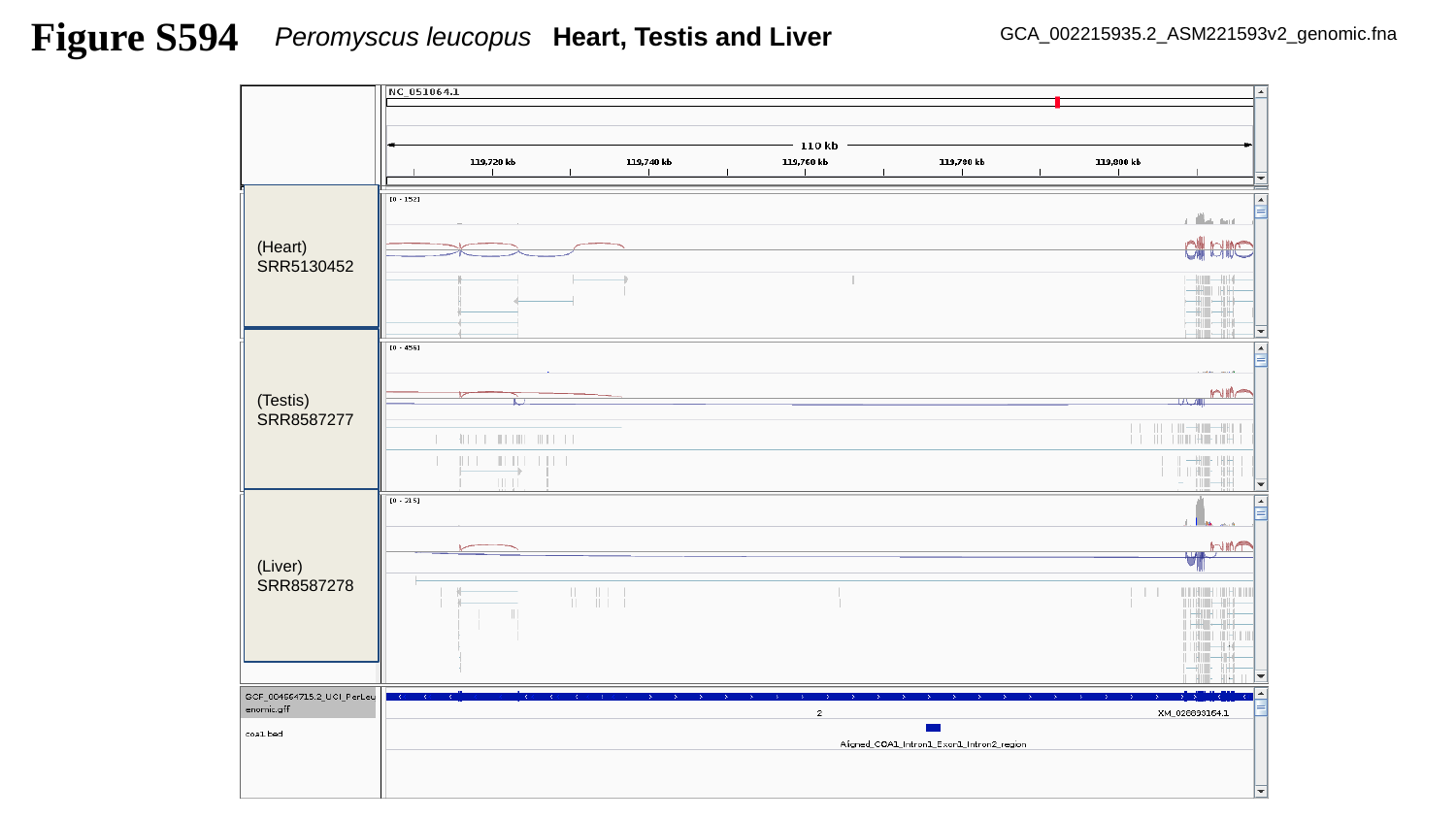

Figure S594
Peromyscus leucopus Heart, Testis and Liver
GCA_002215935.2_ASM221593v2_genomic.fna
(Heart)
SRR5130452
(Testis)
SRR8587277
(Liver)
SRR8587278

### Slide 595
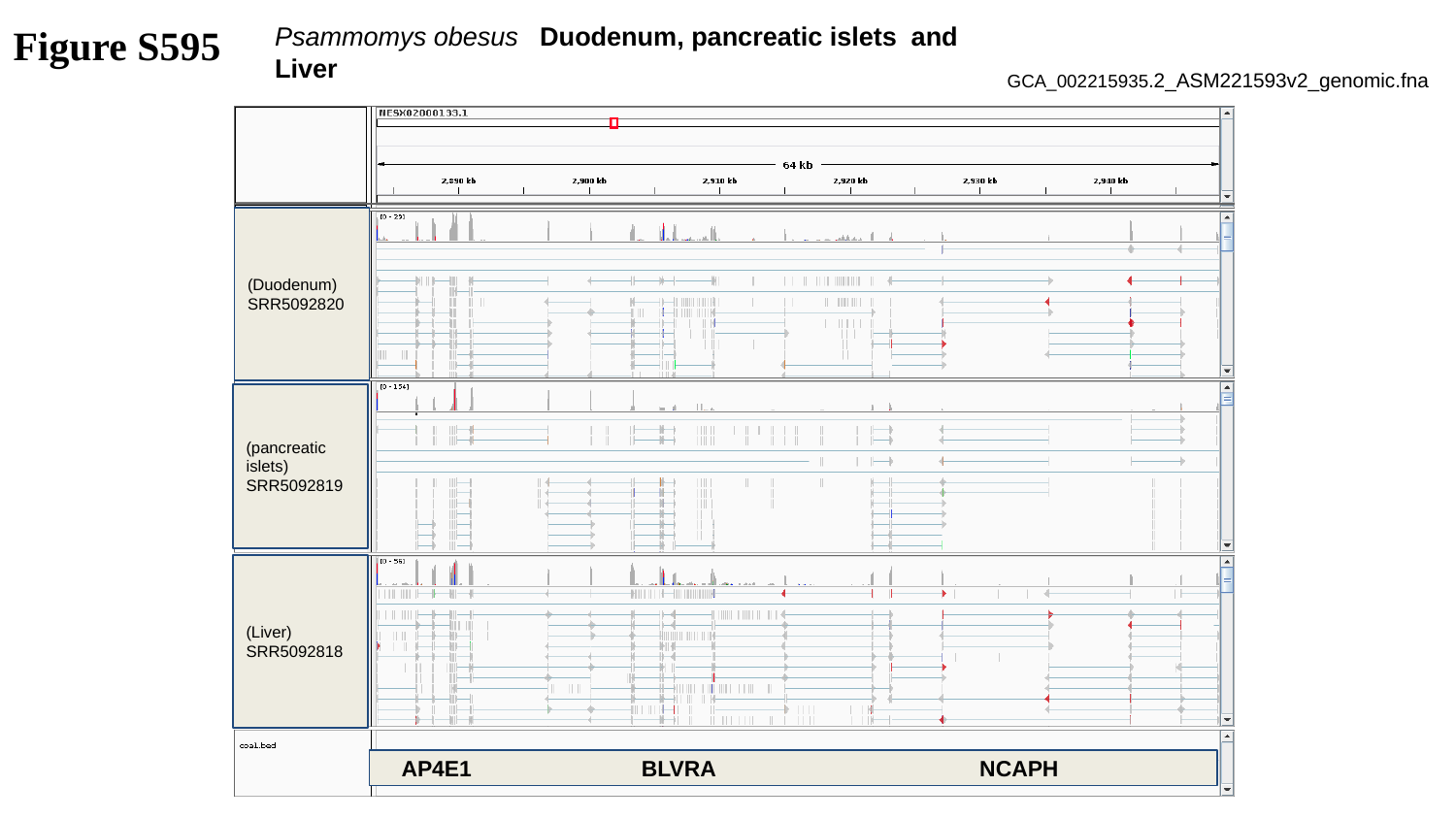

Figure S595
Psammomys obesus Duodenum, pancreatic islets and Liver
GCA_002215935.2_ASM221593v2_genomic.fna
(Duodenum)
SRR5092820
(pancreatic islets)
SRR5092819
(Liver)
SRR5092818
 AP4E1 BLVRA NCAPH

### Slide 596
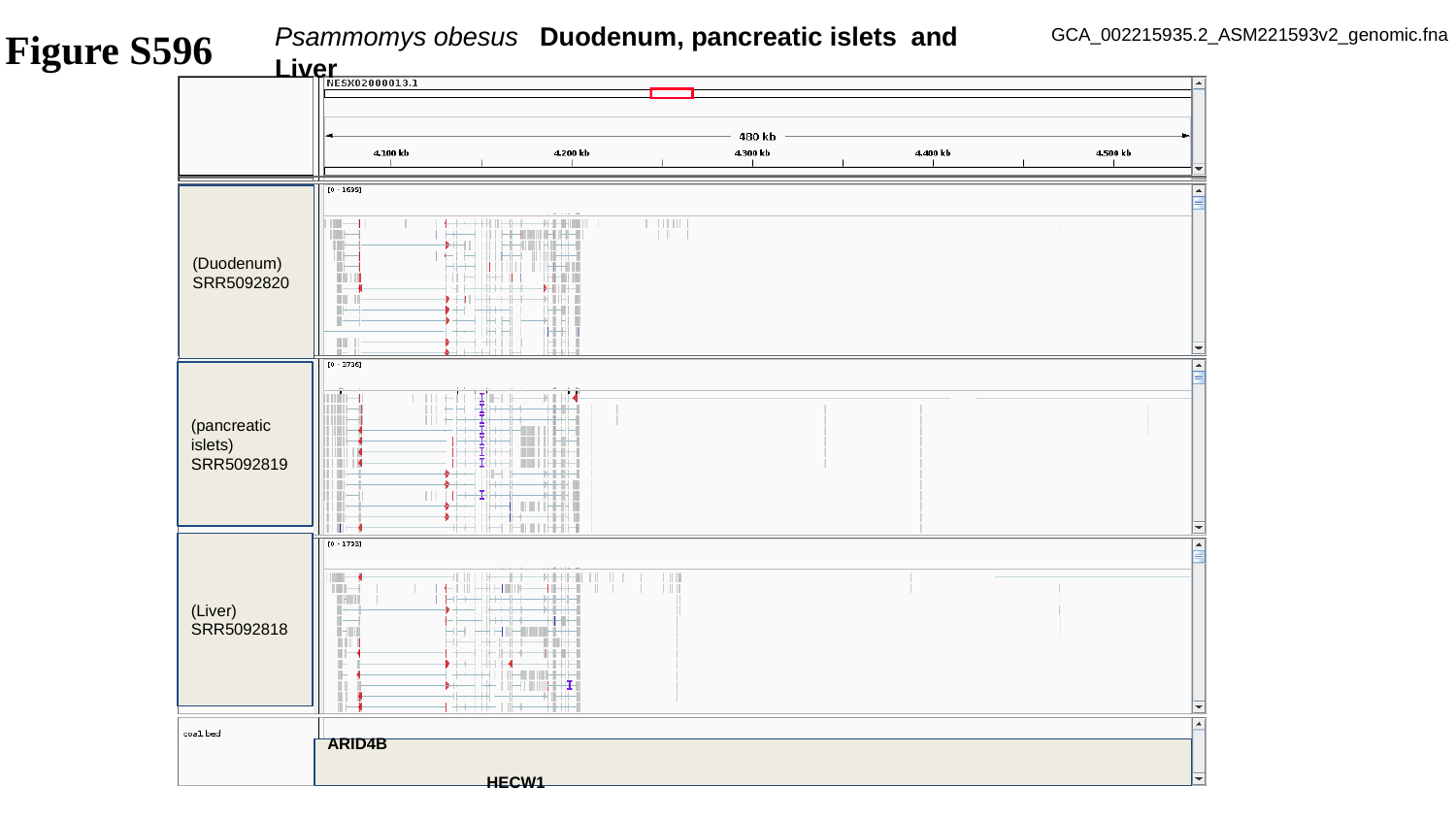

Psammomys obesus Duodenum, pancreatic islets and Liver
GCA_002215935.2_ASM221593v2_genomic.fna
Figure S596
(Duodenum)
SRR5092820
(pancreatic islets)
SRR5092819
(Liver)
SRR5092818
ARID4B											 HECW1

### Slide 597
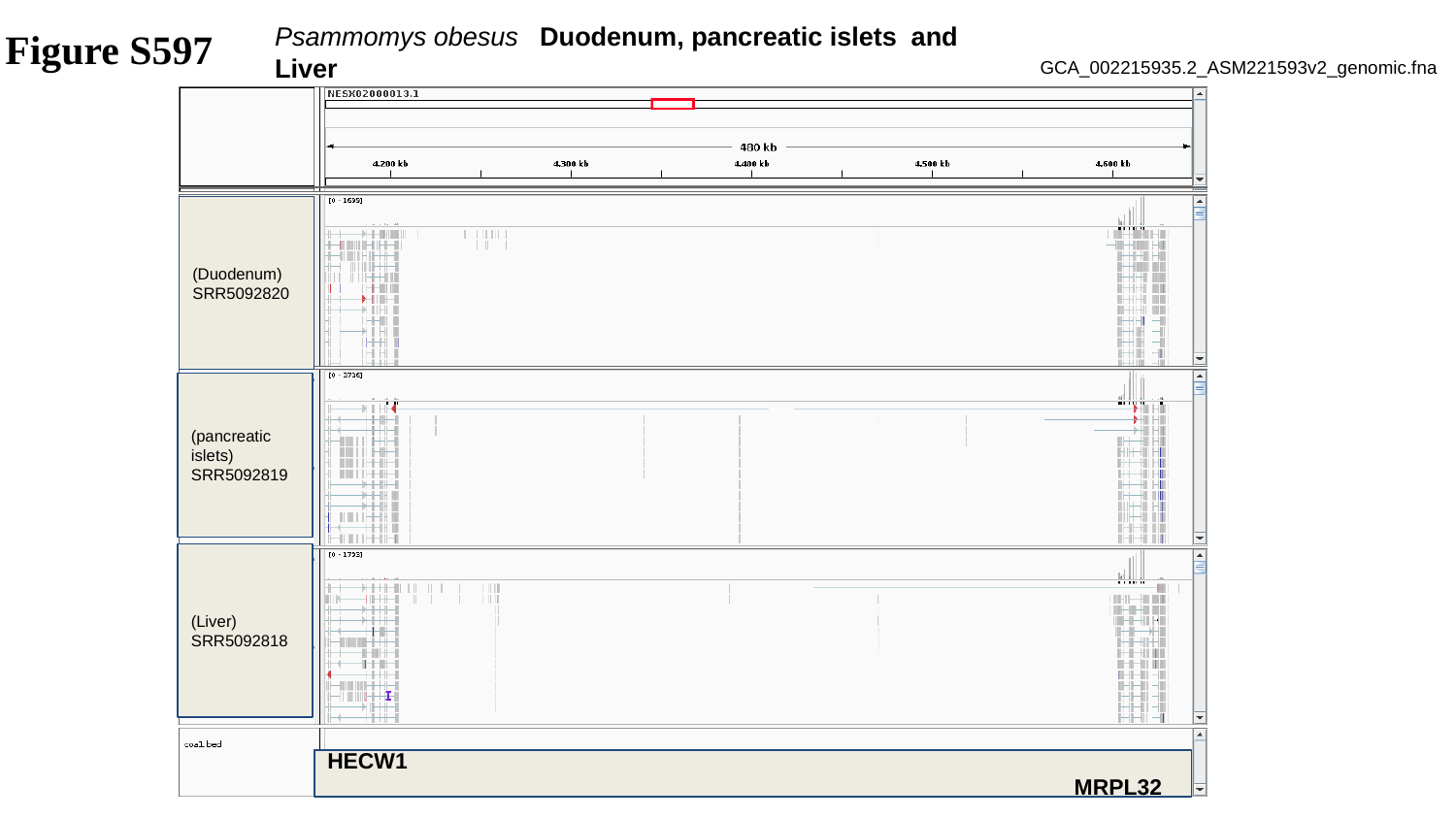

Psammomys obesus Duodenum, pancreatic islets and Liver
Figure S597
GCA_002215935.2_ASM221593v2_genomic.fna
(Duodenum)
SRR5092820
(pancreatic islets)
SRR5092819
(Liver)
SRR5092818
HECW1										 MRPL32

### Slide 598
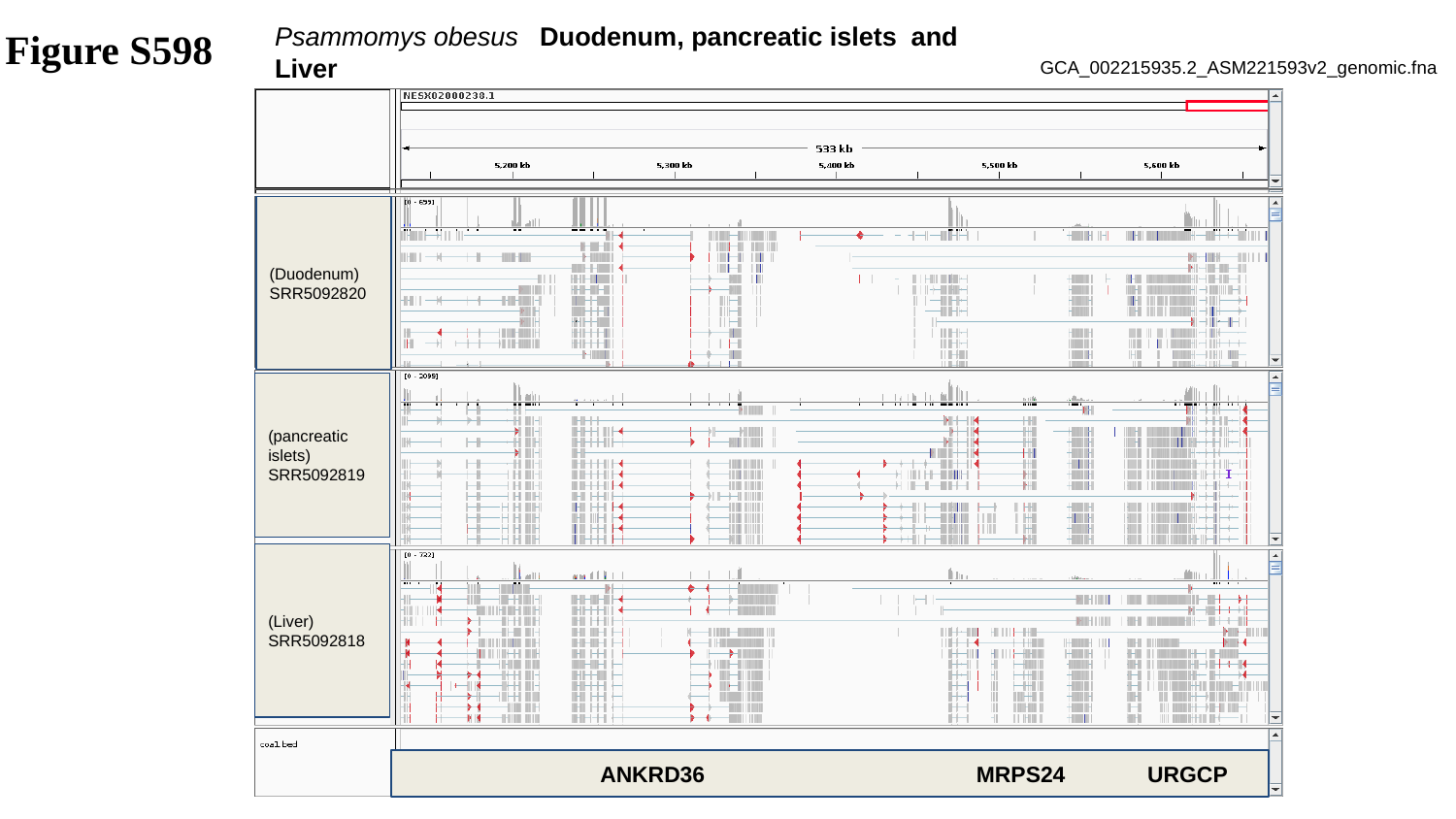

Psammomys obesus Duodenum, pancreatic islets and Liver
Figure S598
GCA_002215935.2_ASM221593v2_genomic.fna
(Duodenum)
SRR5092820
(pancreatic islets)
SRR5092819
(Liver)
SRR5092818
 ANKRD36 MRPS24 URGCP

### Slide 599
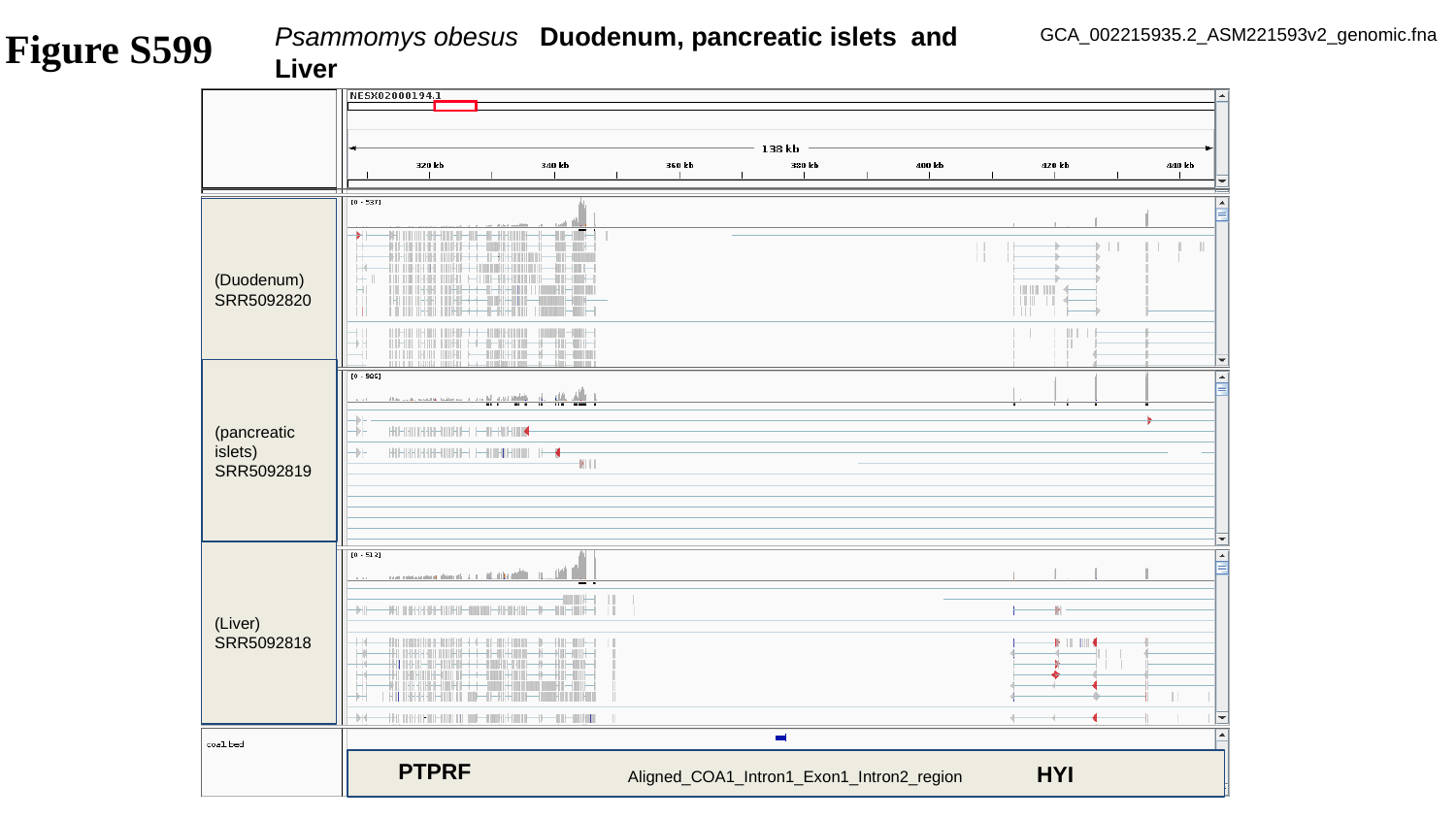

Psammomys obesus Duodenum, pancreatic islets and Liver
Figure S599
GCA_002215935.2_ASM221593v2_genomic.fna
(Duodenum)
SRR5092820
(pancreatic islets)
SRR5092819
(Liver)
SRR5092818
PTPRF
 Aligned_COA1_Intron1_Exon1_Intron2_region HYI

### Slide 600
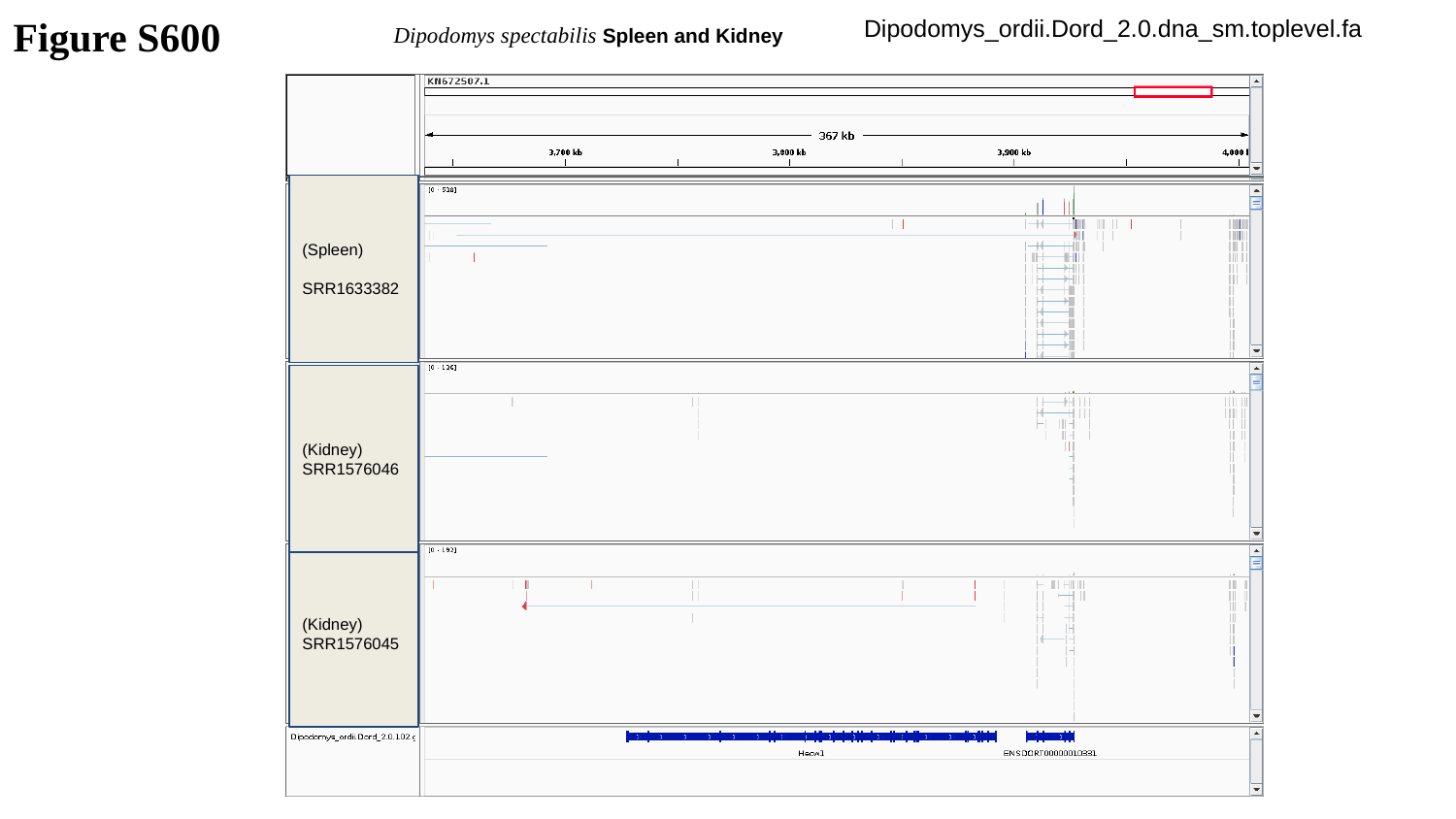

Dipodomys_ordii.Dord_2.0.dna_sm.toplevel.fa
Figure S600
Dipodomys spectabilis Spleen and Kidney
(Spleen)
SRR1633382
(Kidney)
SRR1576046
(Kidney)
SRR1576045

### Slide 601
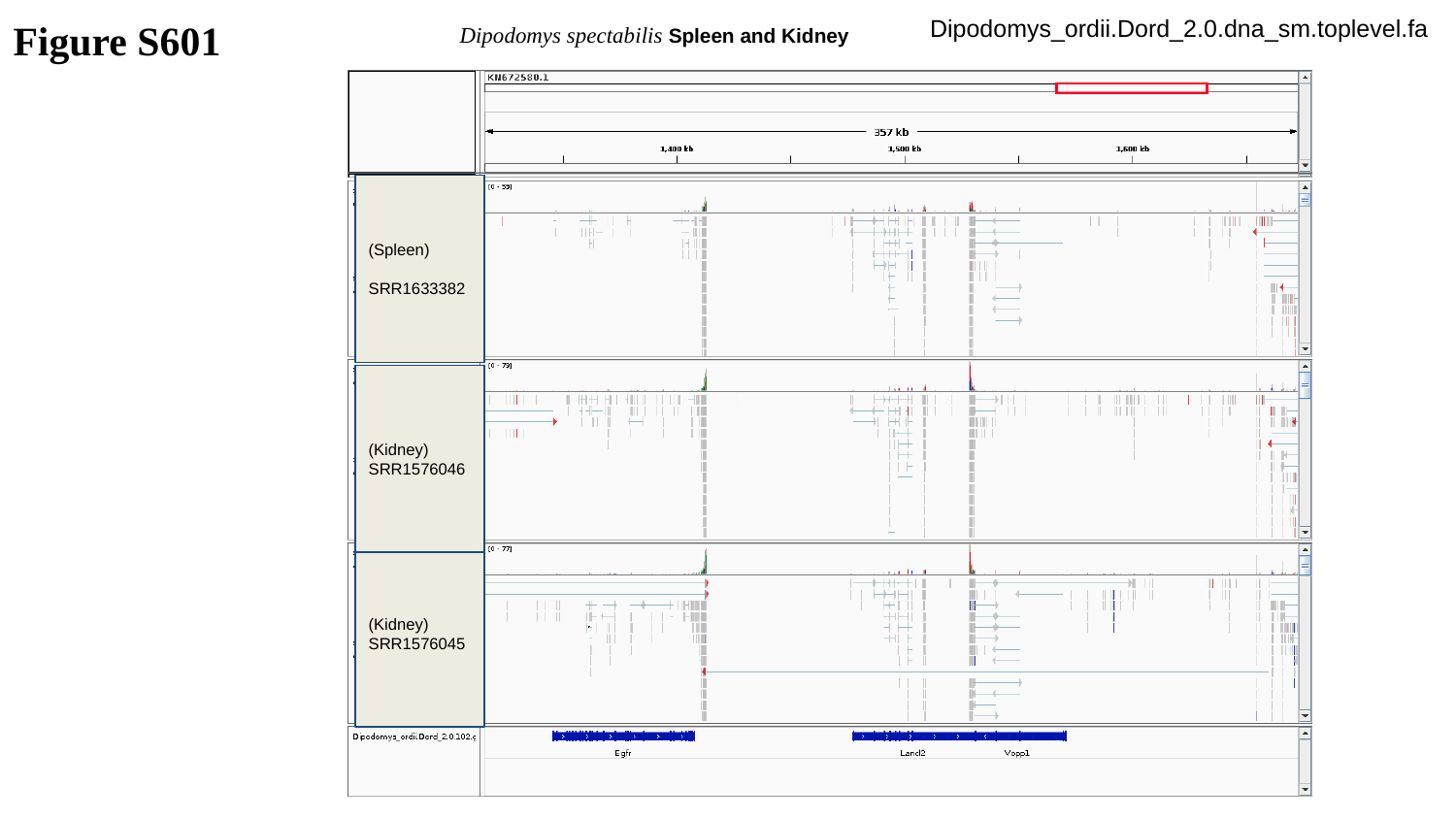

Dipodomys_ordii.Dord_2.0.dna_sm.toplevel.fa
Figure S601
Dipodomys spectabilis Spleen and Kidney
(Spleen)
SRR1633382
(Kidney)
SRR1576046
(Kidney)
SRR1576045

### Slide 602
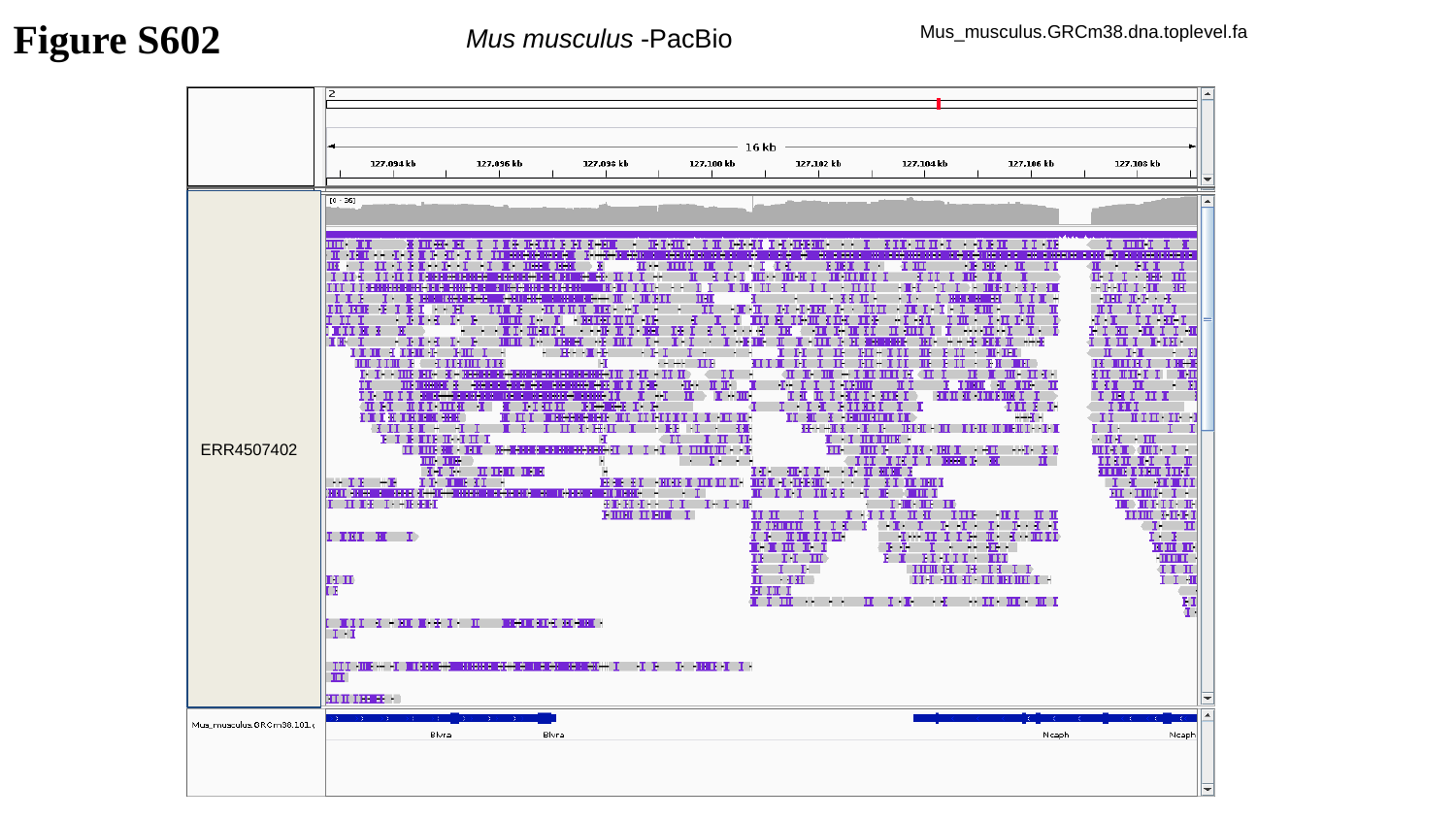

Figure S602
Mus musculus -PacBio
Mus_musculus.GRCm38.dna.toplevel.fa
ERR4507402

### Slide 603
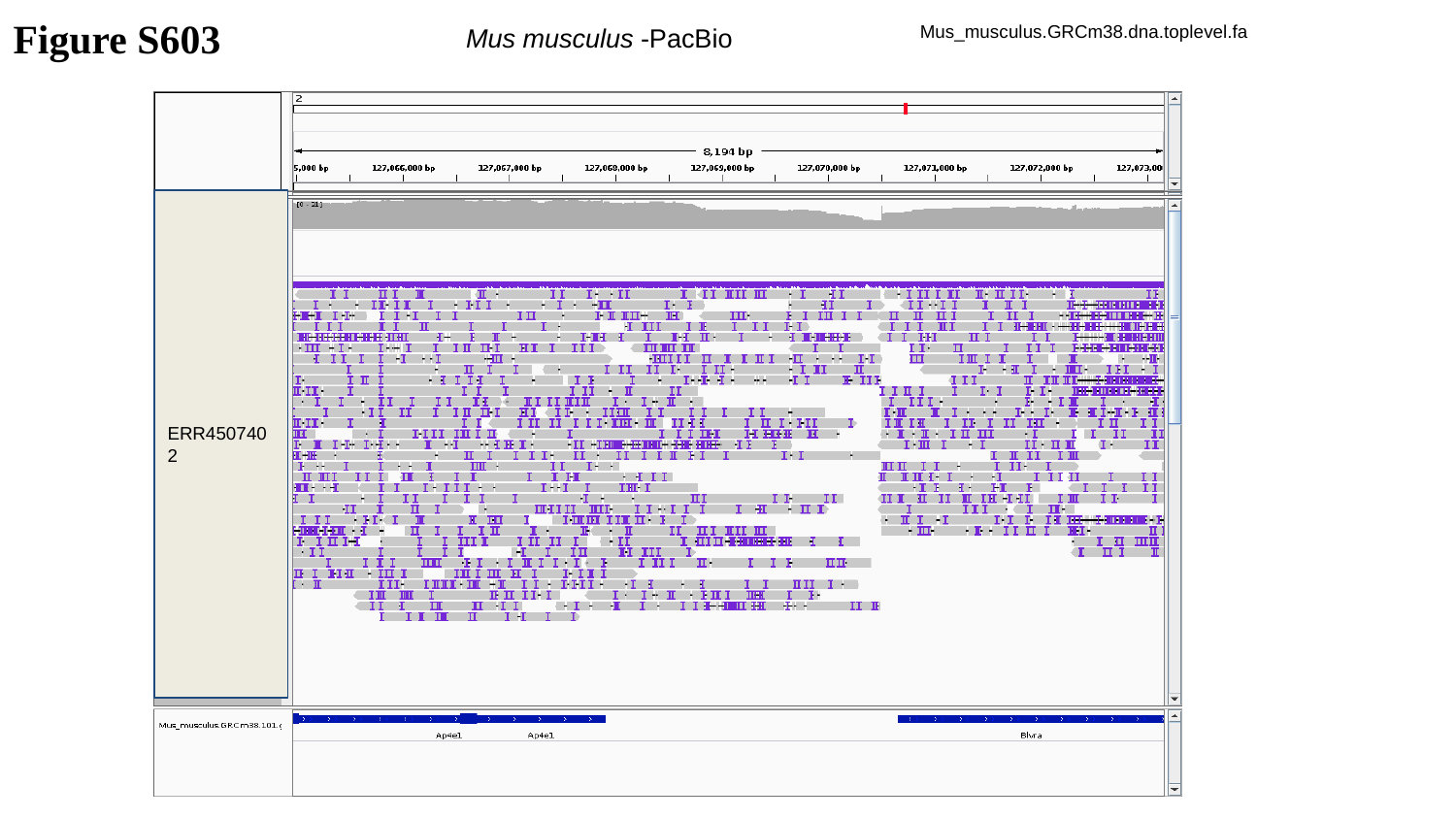

Figure S603
Mus musculus -PacBio
Mus_musculus.GRCm38.dna.toplevel.fa
ERR4507402

### Slide 604
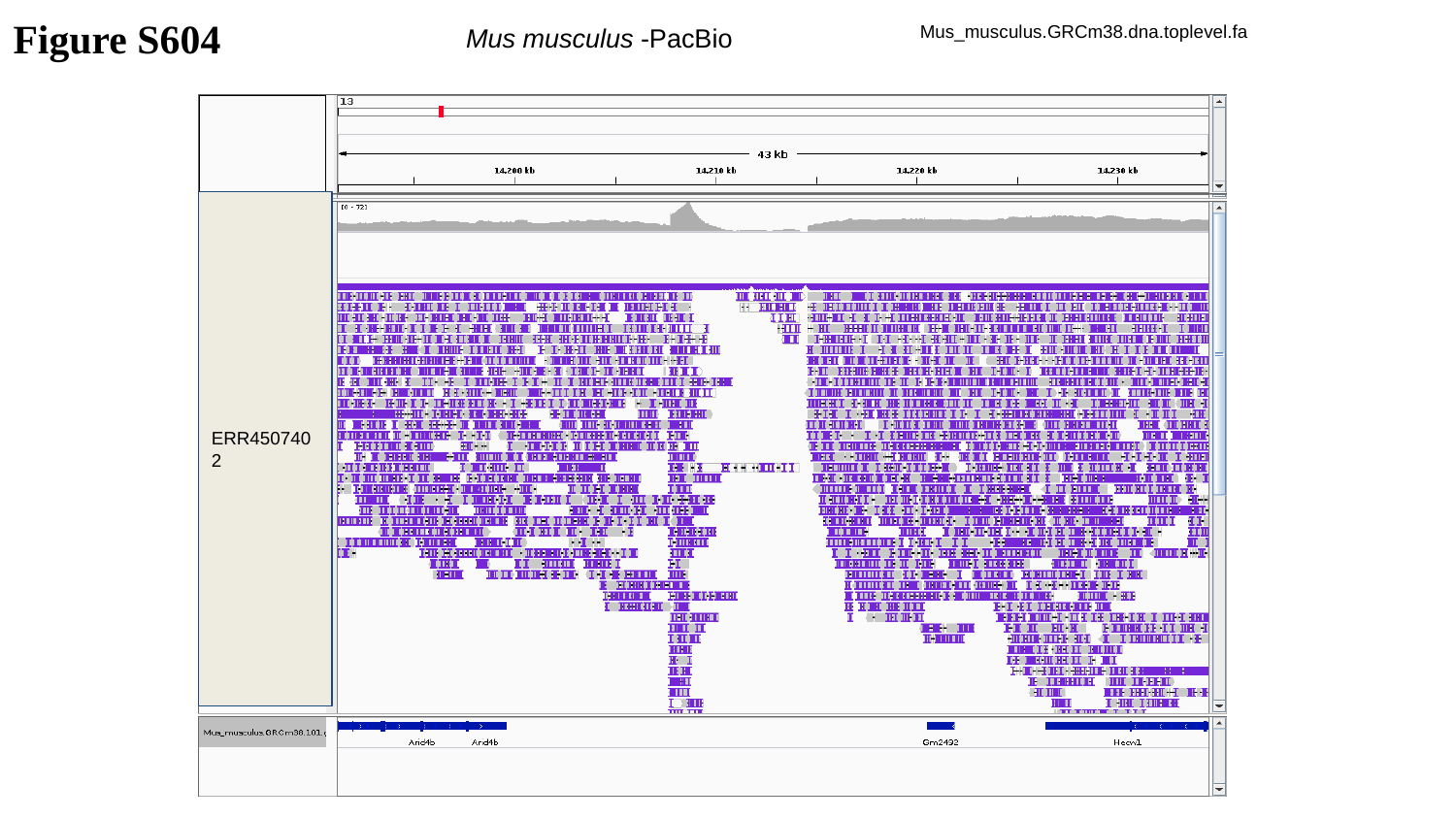

Figure S604
Mus musculus -PacBio
Mus_musculus.GRCm38.dna.toplevel.fa
ERR4507402

### Slide 605
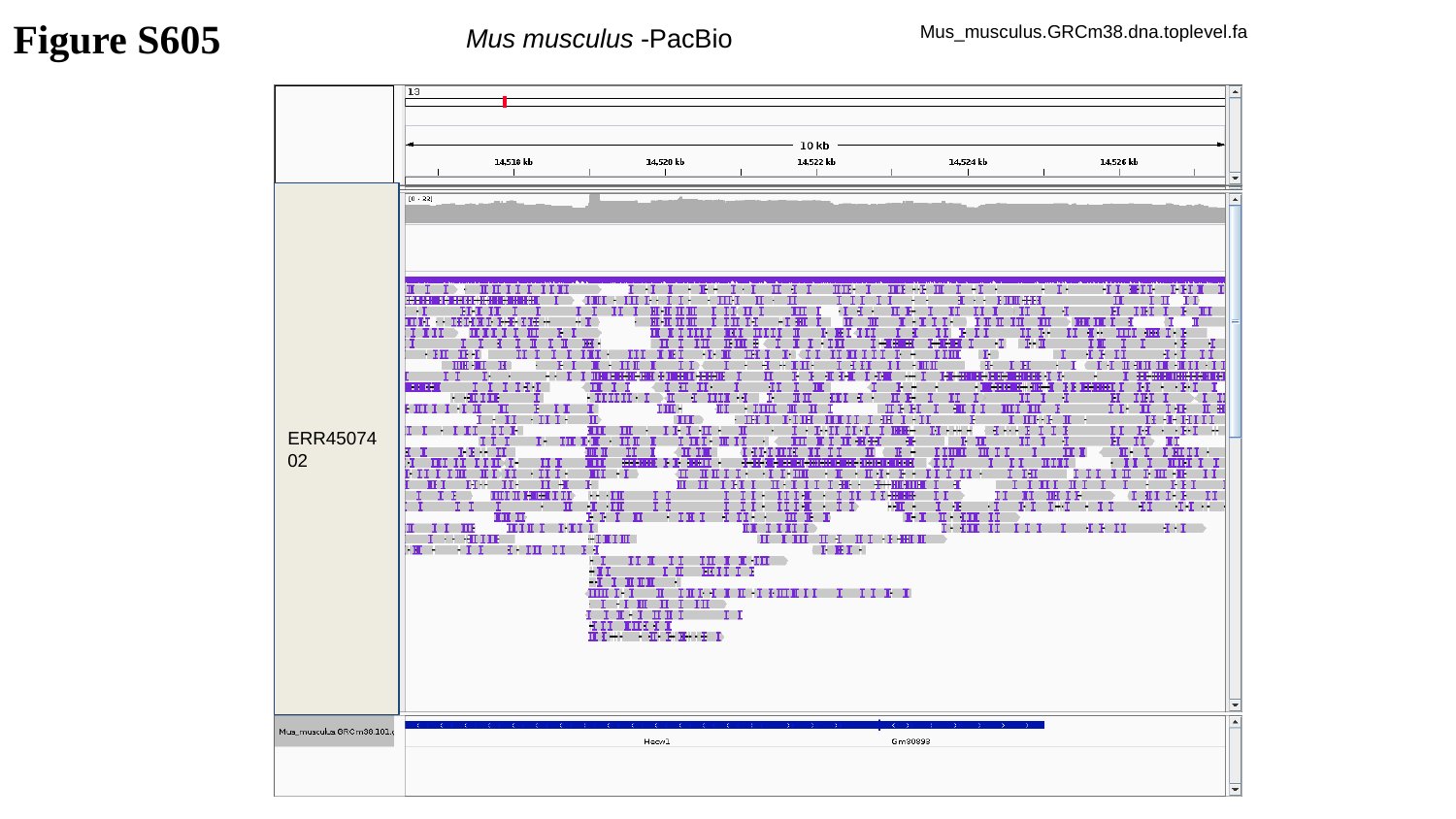

Figure S605
Mus musculus -PacBio
Mus_musculus.GRCm38.dna.toplevel.fa
ERR4507402

### Slide 606
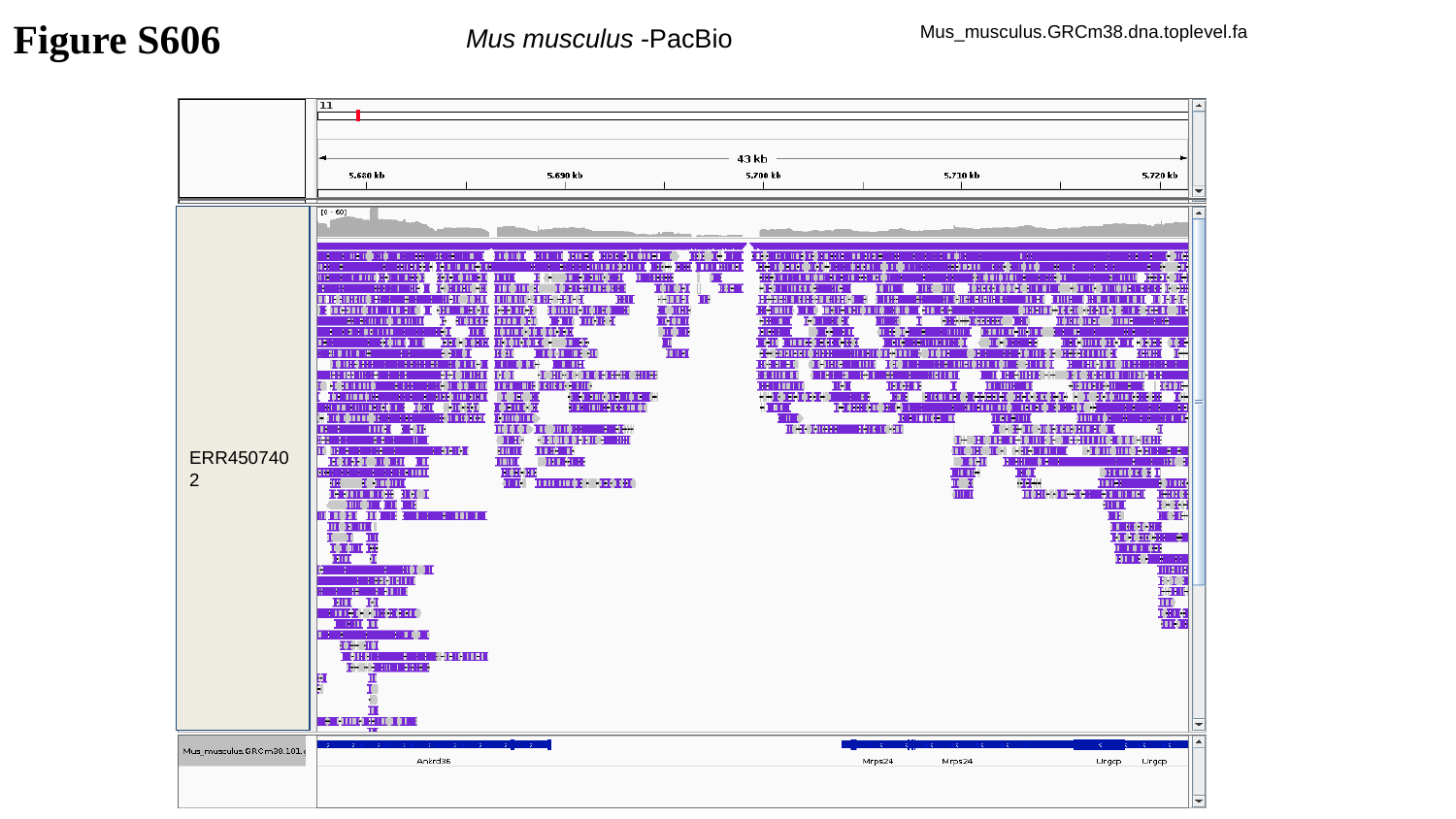

Figure S606
Mus musculus -PacBio
Mus_musculus.GRCm38.dna.toplevel.fa
ERR4507402

### Slide 607
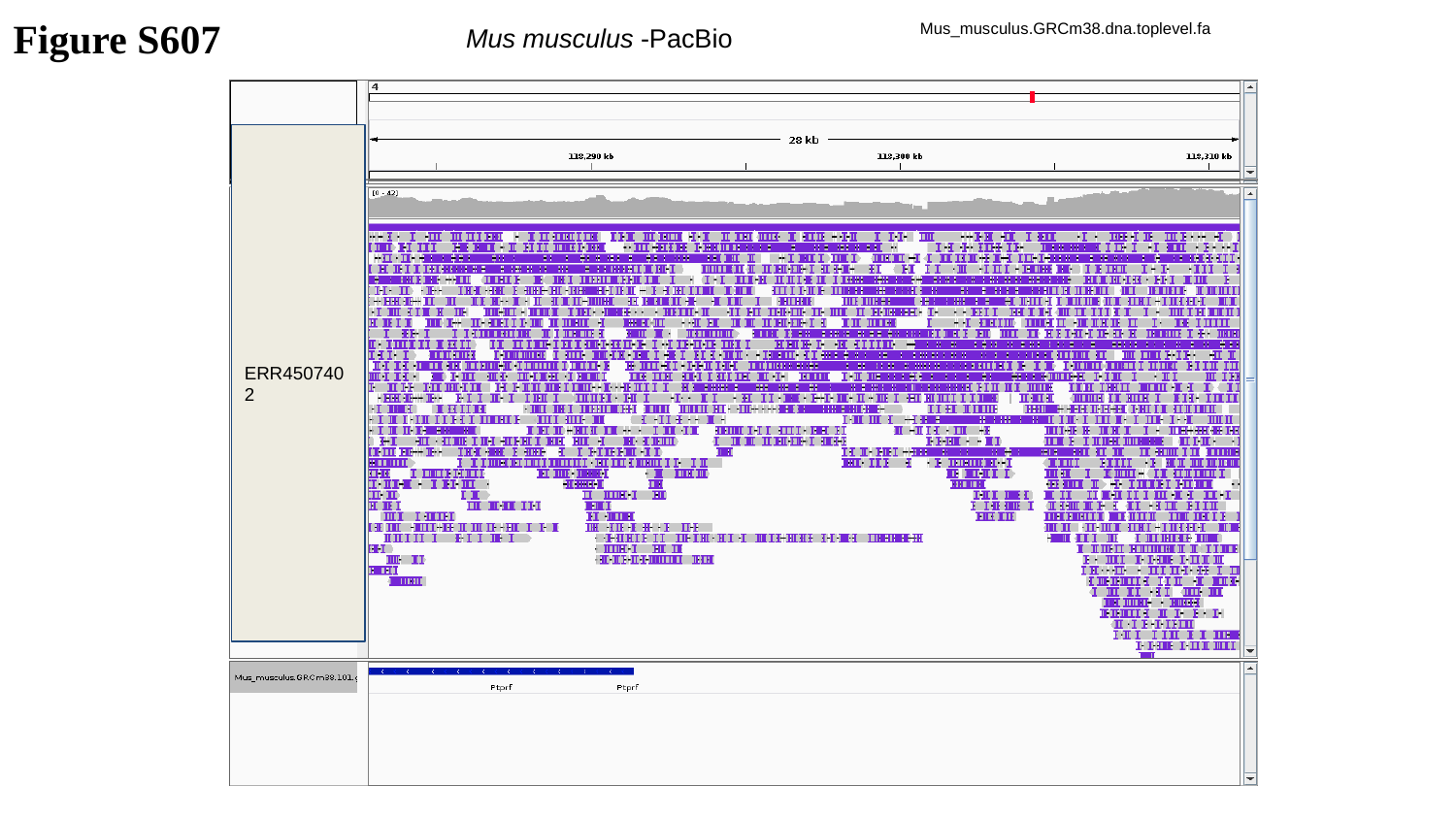

Figure S607
Mus musculus -PacBio
Mus_musculus.GRCm38.dna.toplevel.fa
ERR4507402

### Slide 608
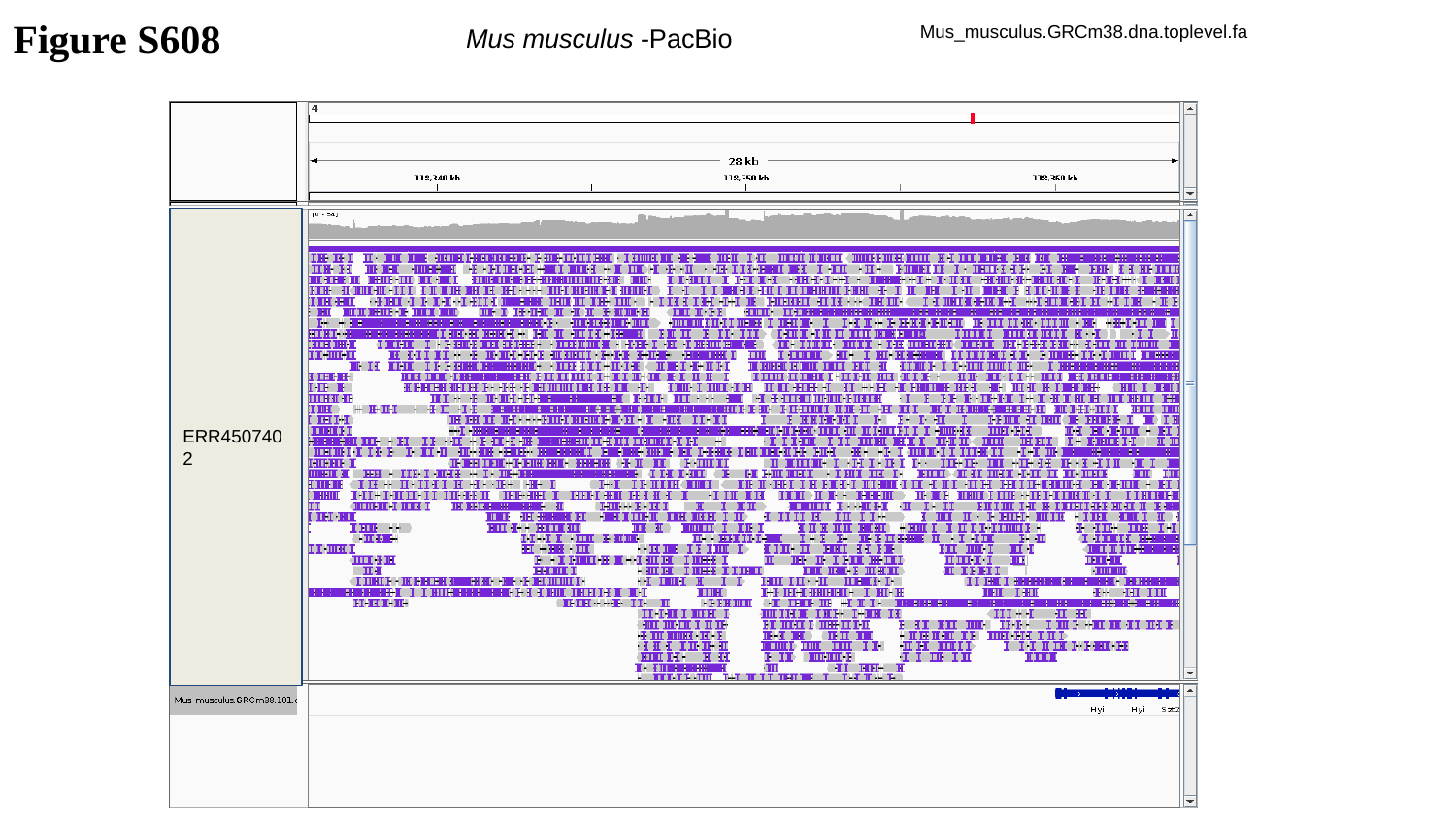

Figure S608
Mus musculus -PacBio
Mus_musculus.GRCm38.dna.toplevel.fa
ERR4507402

### Slide 609
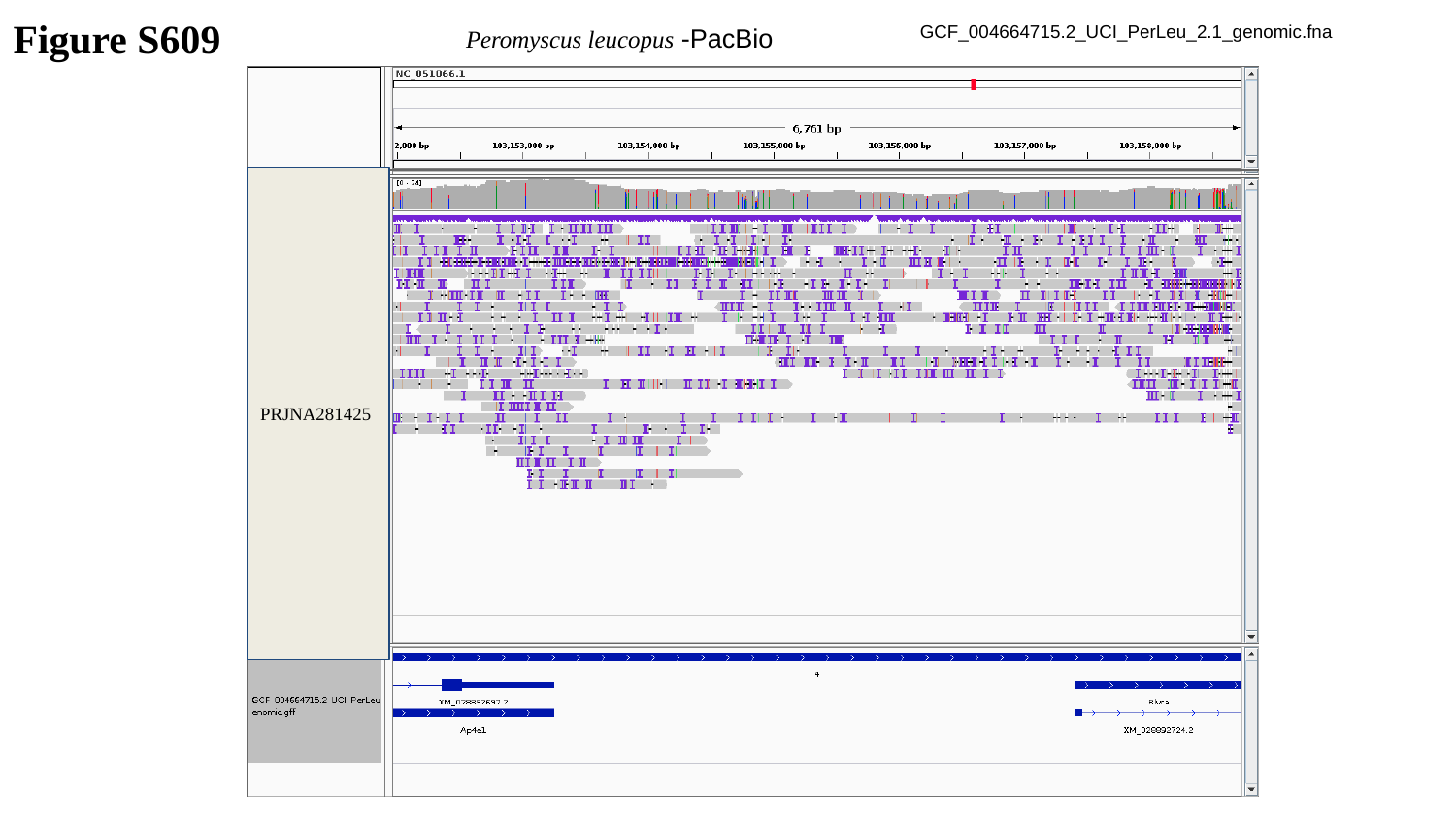

Figure S609
Peromyscus leucopus -PacBio
GCF_004664715.2_UCI_PerLeu_2.1_genomic.fna
PRJNA281425

### Slide 610
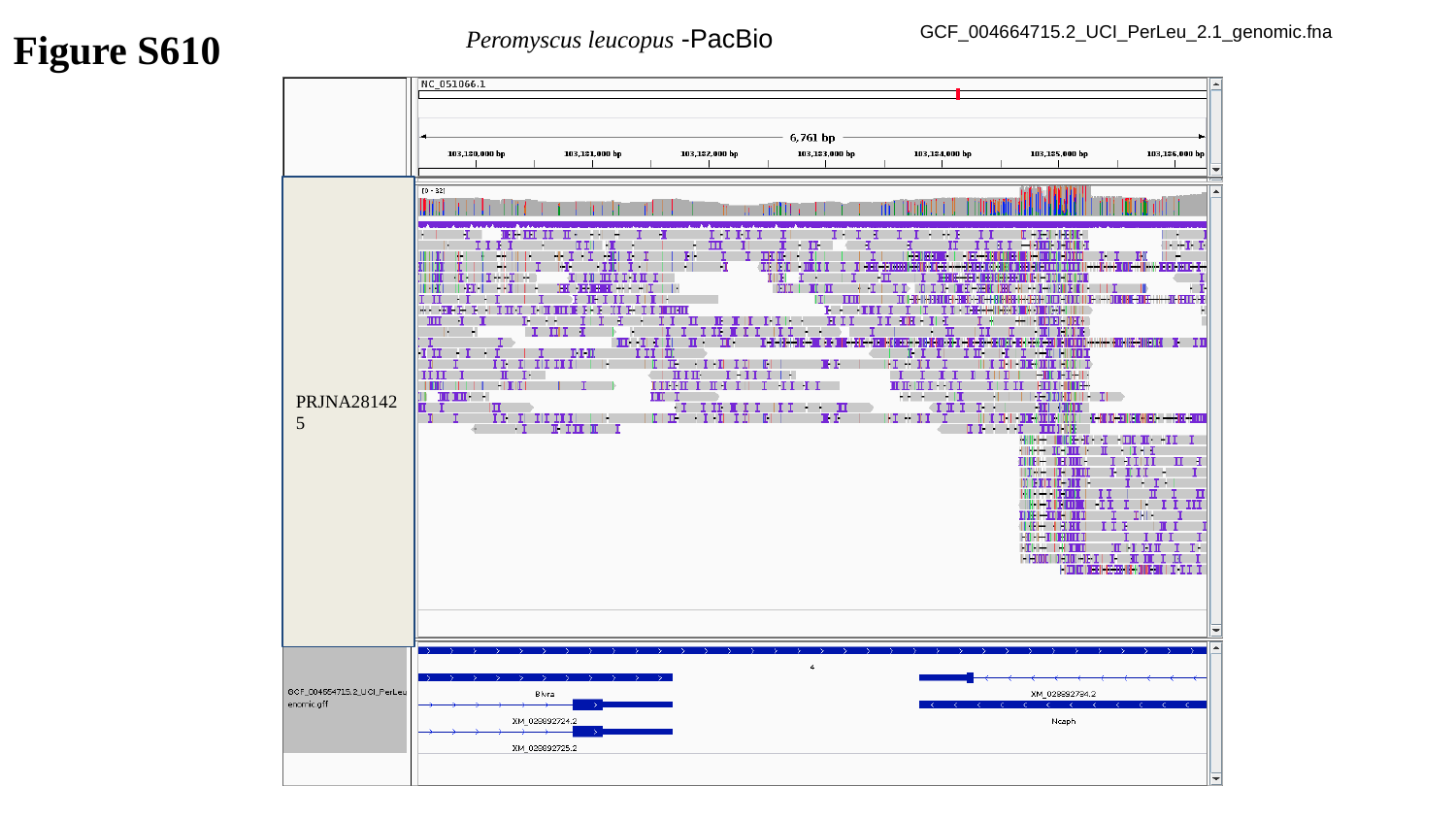

Peromyscus leucopus -PacBio
GCF_004664715.2_UCI_PerLeu_2.1_genomic.fna
Figure S610
PRJNA281425

### Slide 611
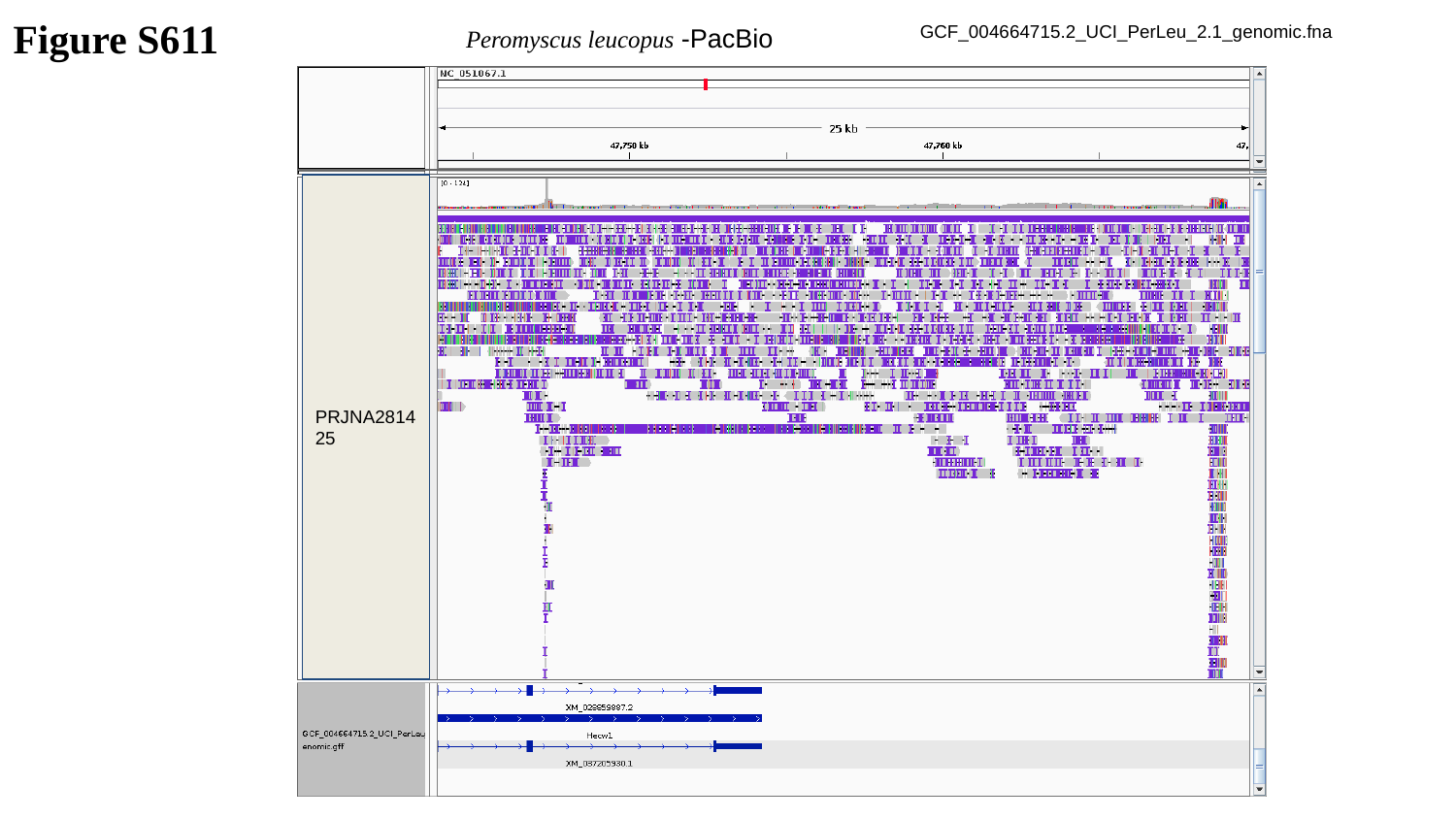

Figure S611
Peromyscus leucopus -PacBio
GCF_004664715.2_UCI_PerLeu_2.1_genomic.fna
PRJNA281425

### Slide 612
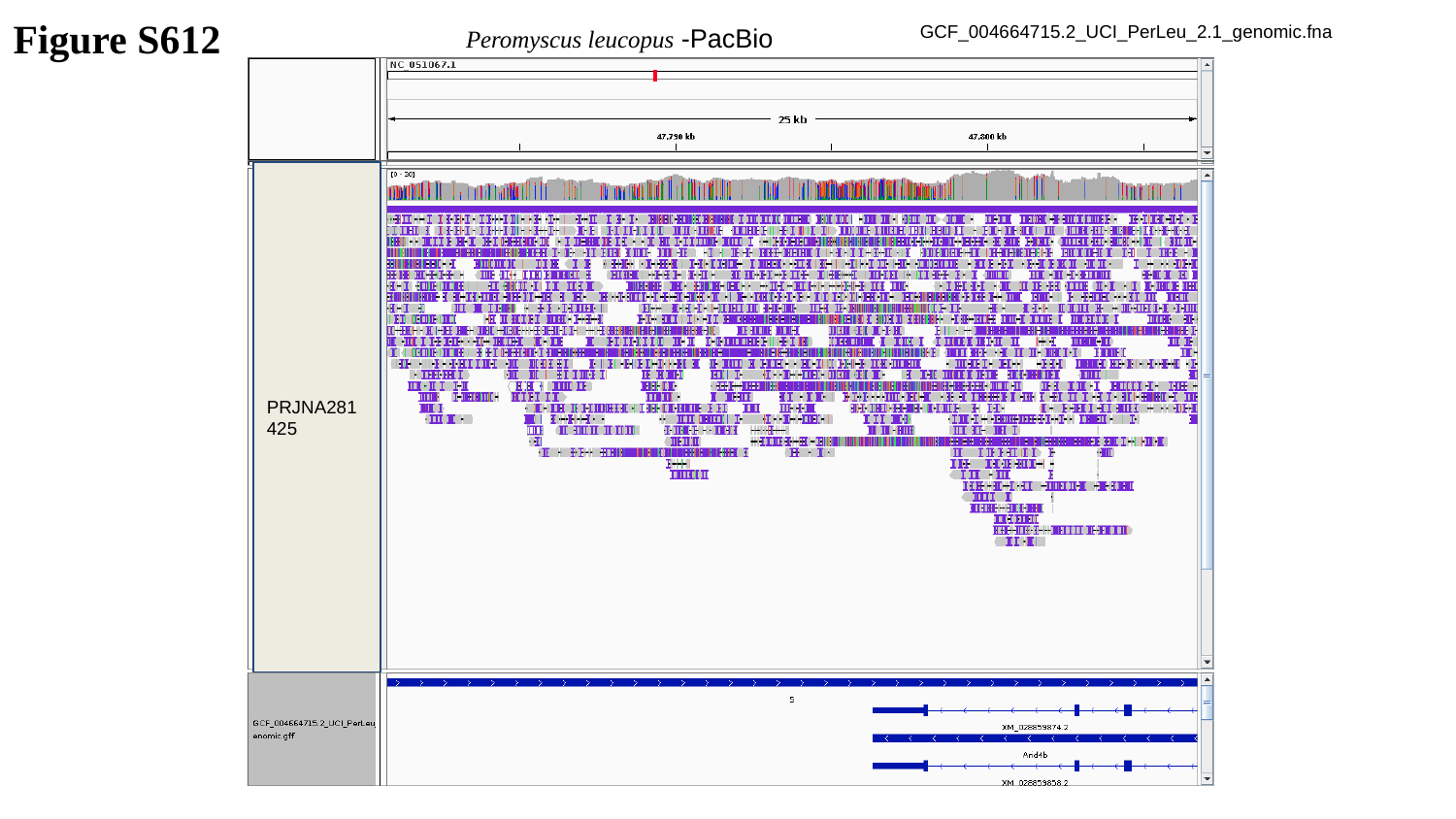

Figure S612
Peromyscus leucopus -PacBio
GCF_004664715.2_UCI_PerLeu_2.1_genomic.fna
PRJNA281425

### Slide 613
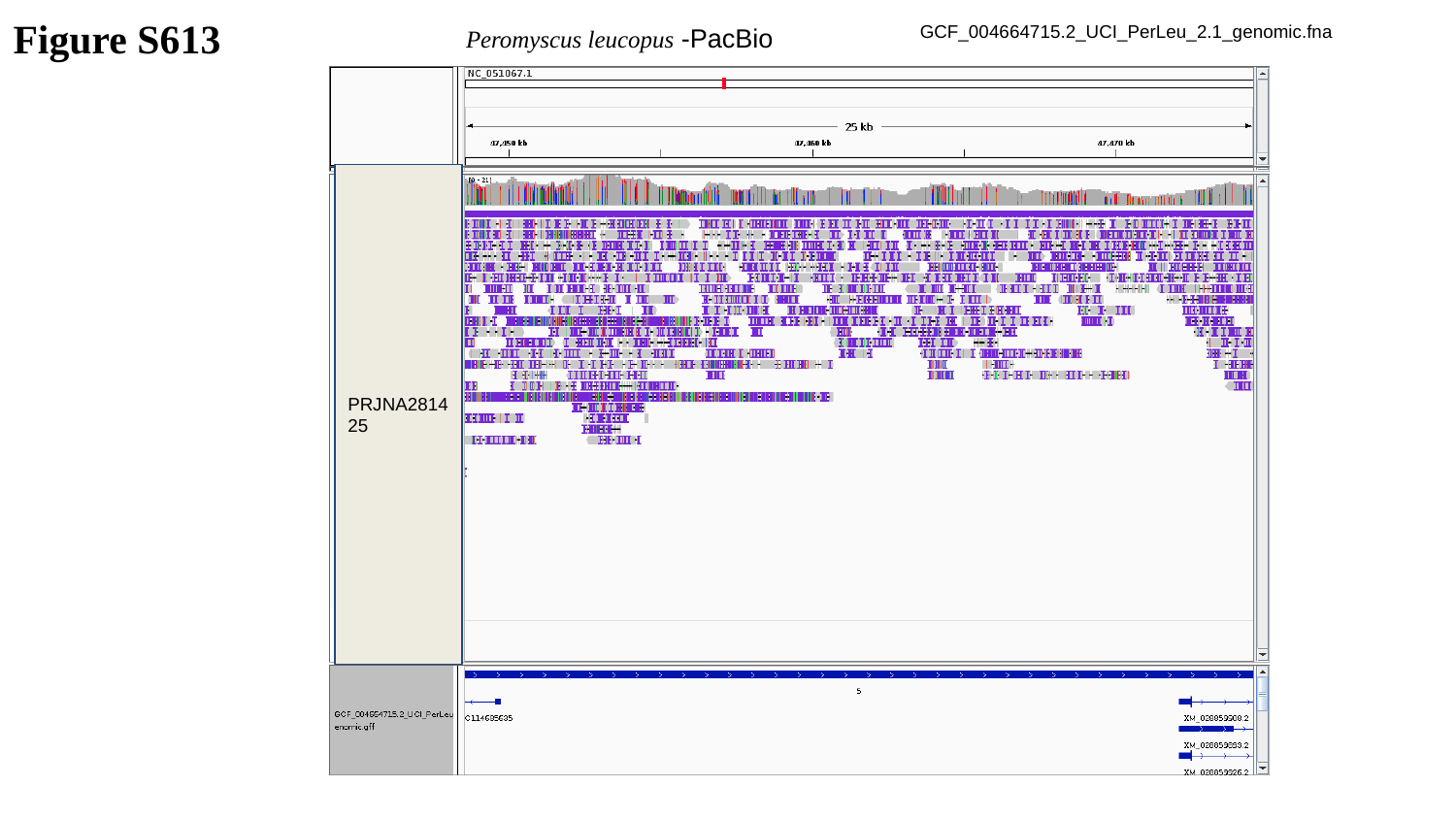

Figure S613
Peromyscus leucopus -PacBio
GCF_004664715.2_UCI_PerLeu_2.1_genomic.fna
PRJNA281425

### Slide 614
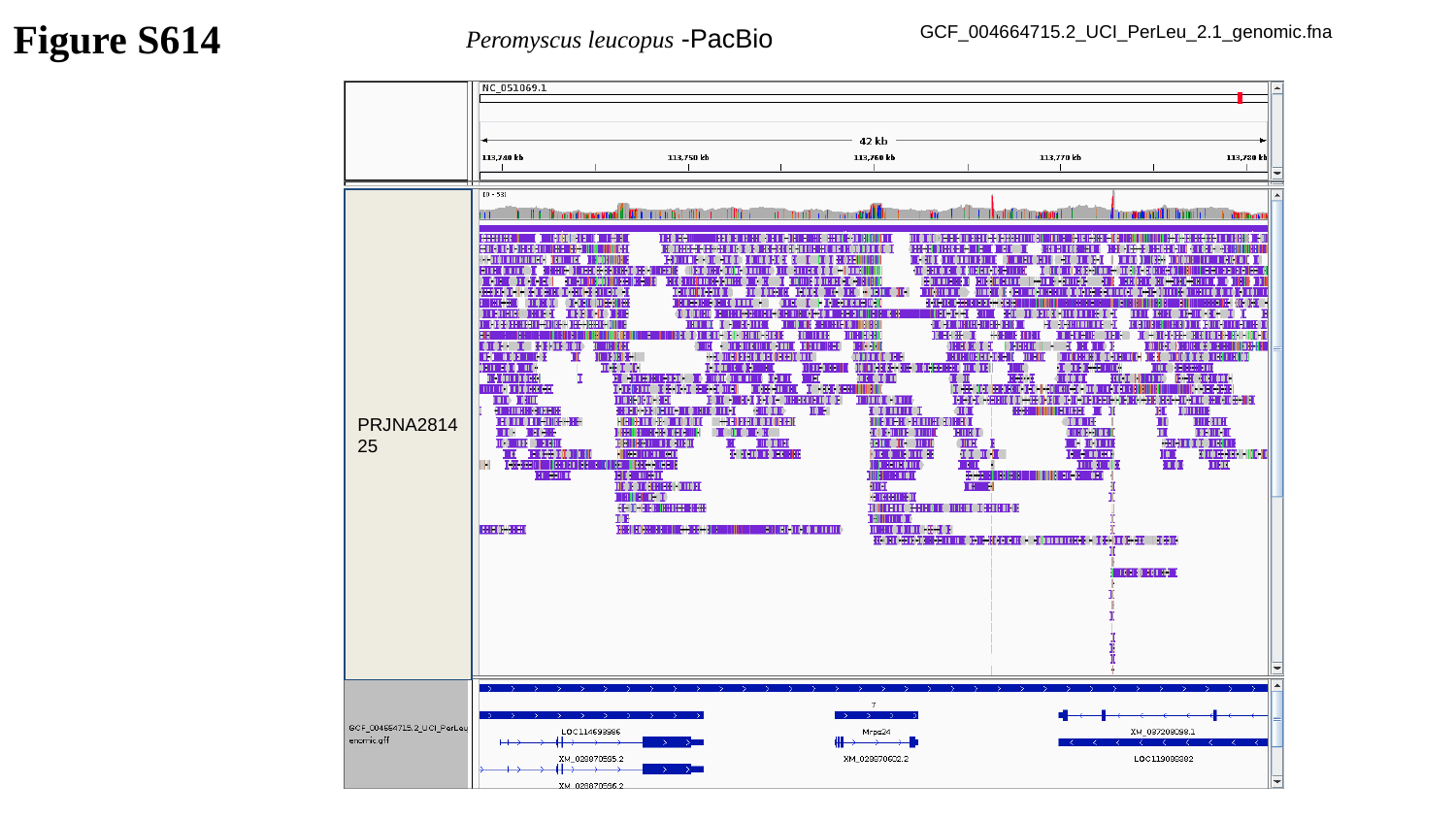

Figure S614
Peromyscus leucopus -PacBio
GCF_004664715.2_UCI_PerLeu_2.1_genomic.fna
PRJNA281425

### Slide 615
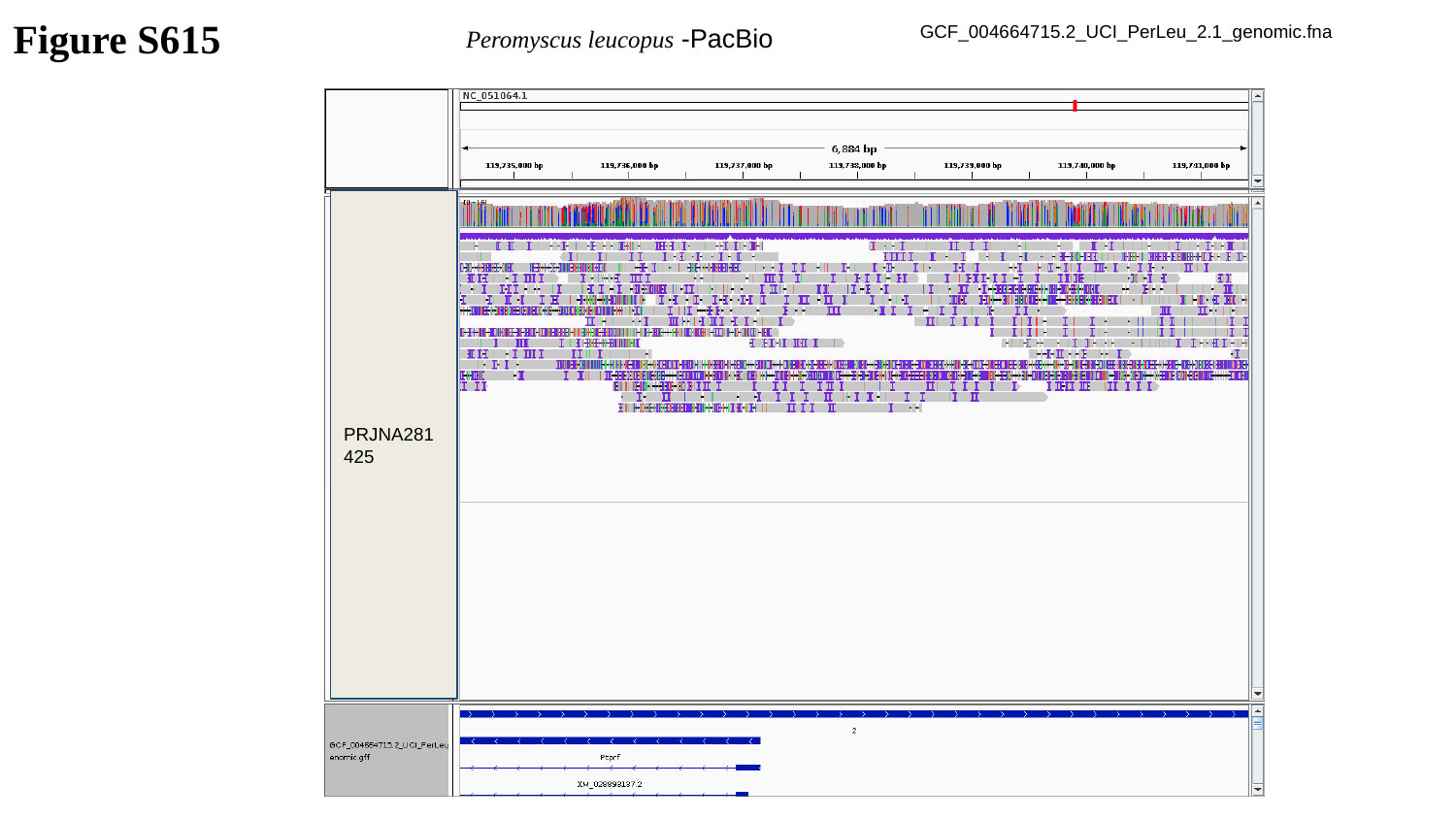

Figure S615
Peromyscus leucopus -PacBio
GCF_004664715.2_UCI_PerLeu_2.1_genomic.fna
PRJNA281425

### Slide 616
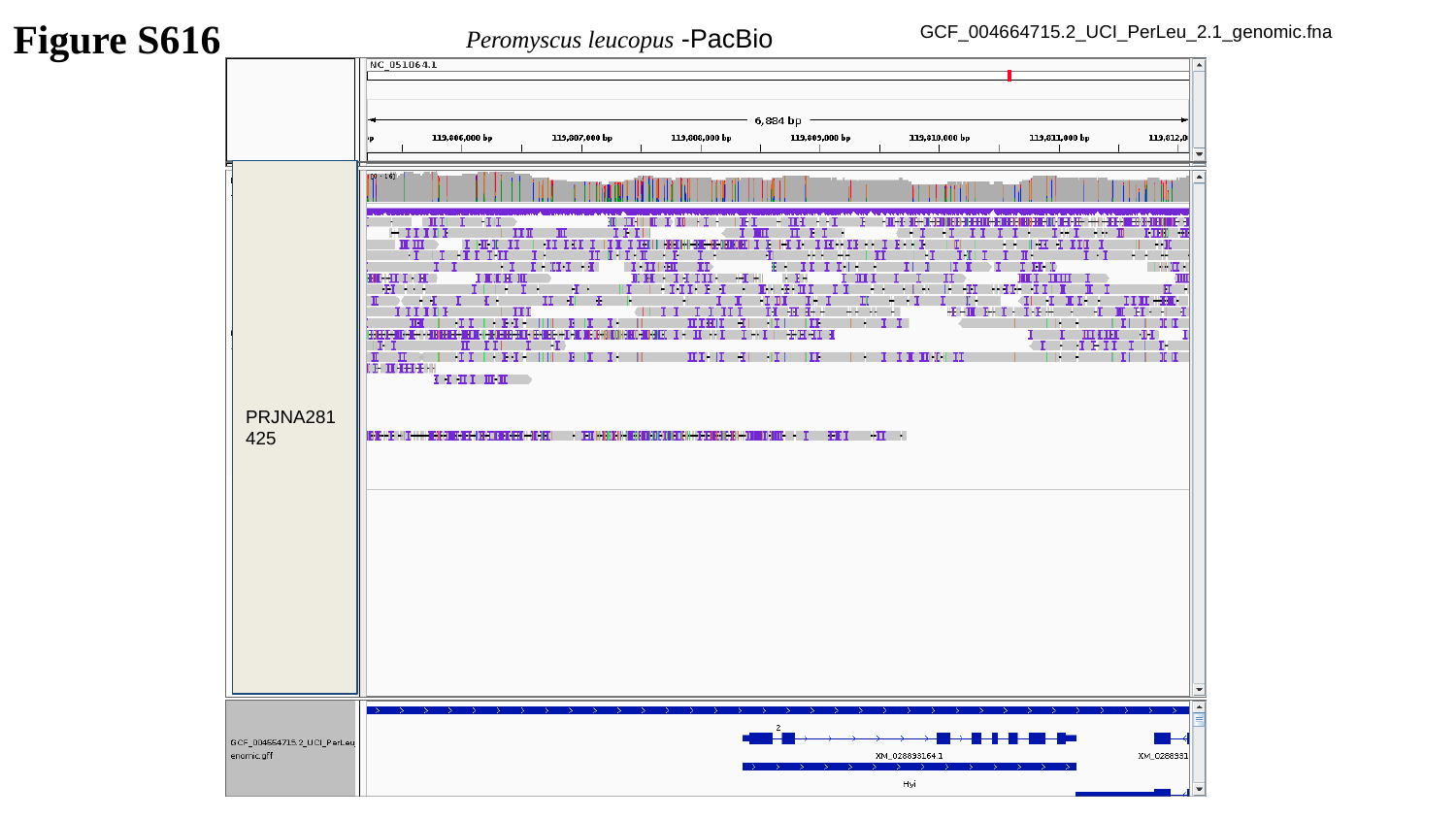

Figure S616
Peromyscus leucopus -PacBio
GCF_004664715.2_UCI_PerLeu_2.1_genomic.fna
PRJNA281425

### Slide 617
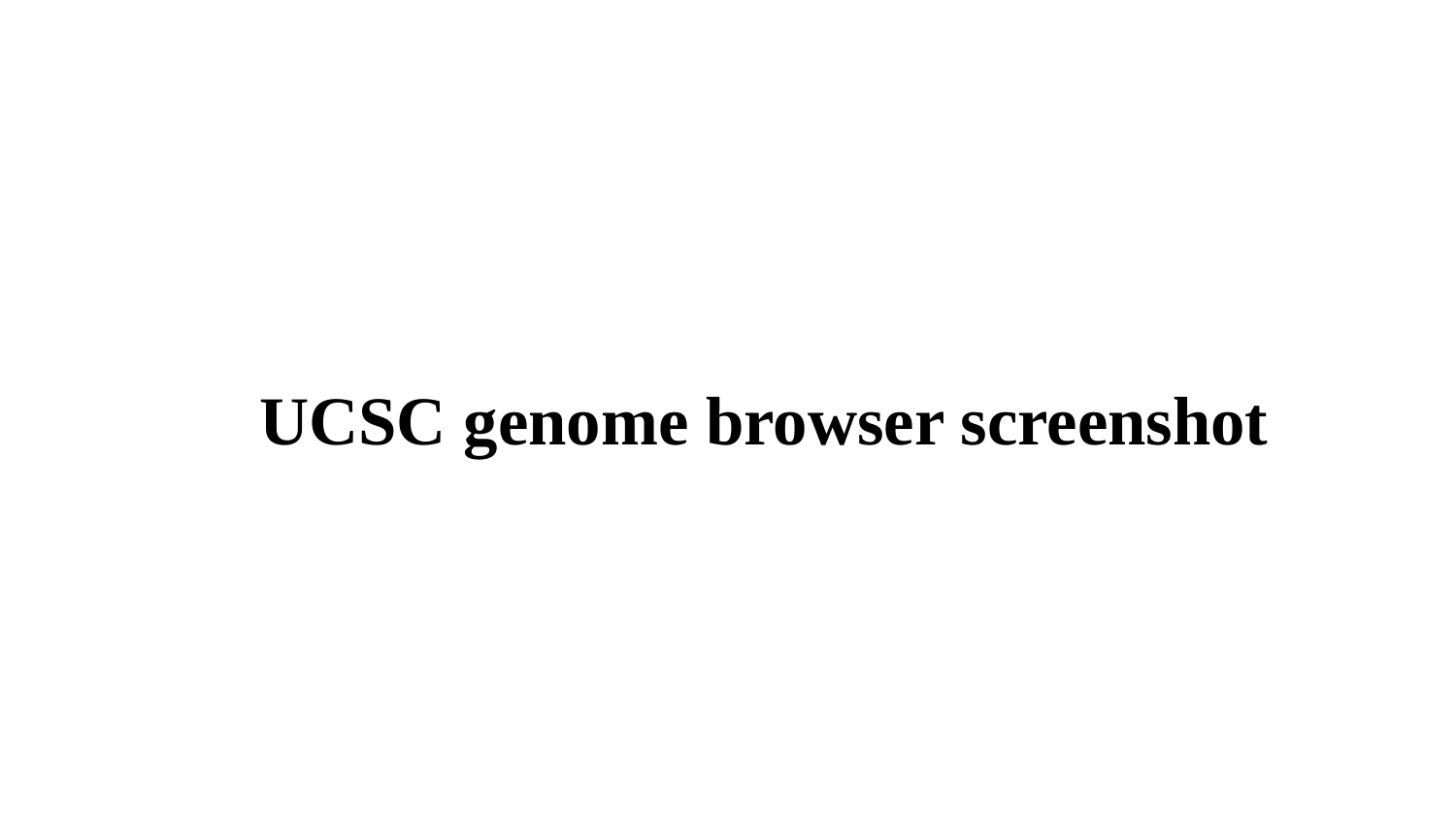

UCSC genome browser screenshot

### Slide 618
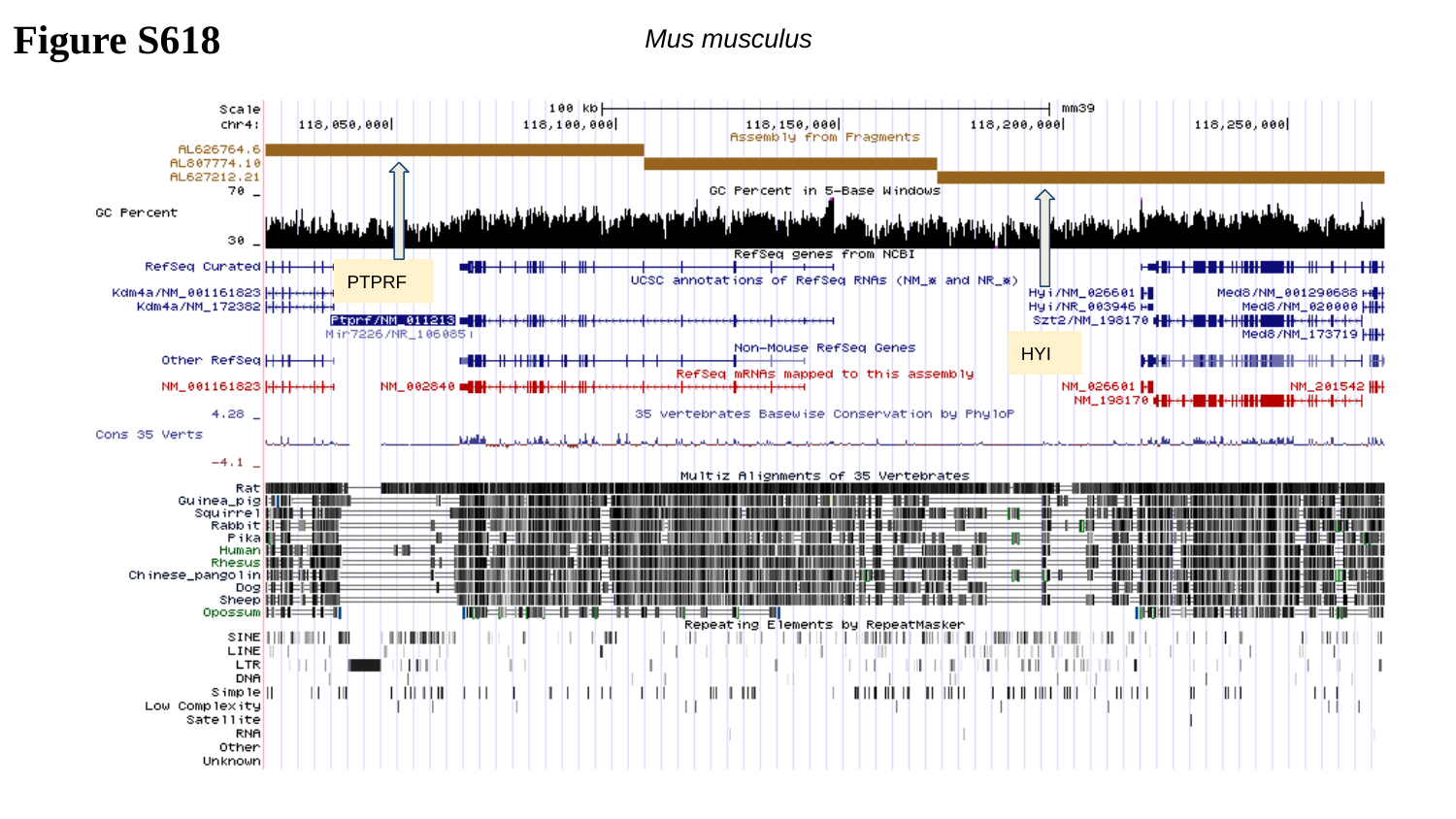

Figure S618
Mus musculus
PTPRF
HYI

### Slide 619
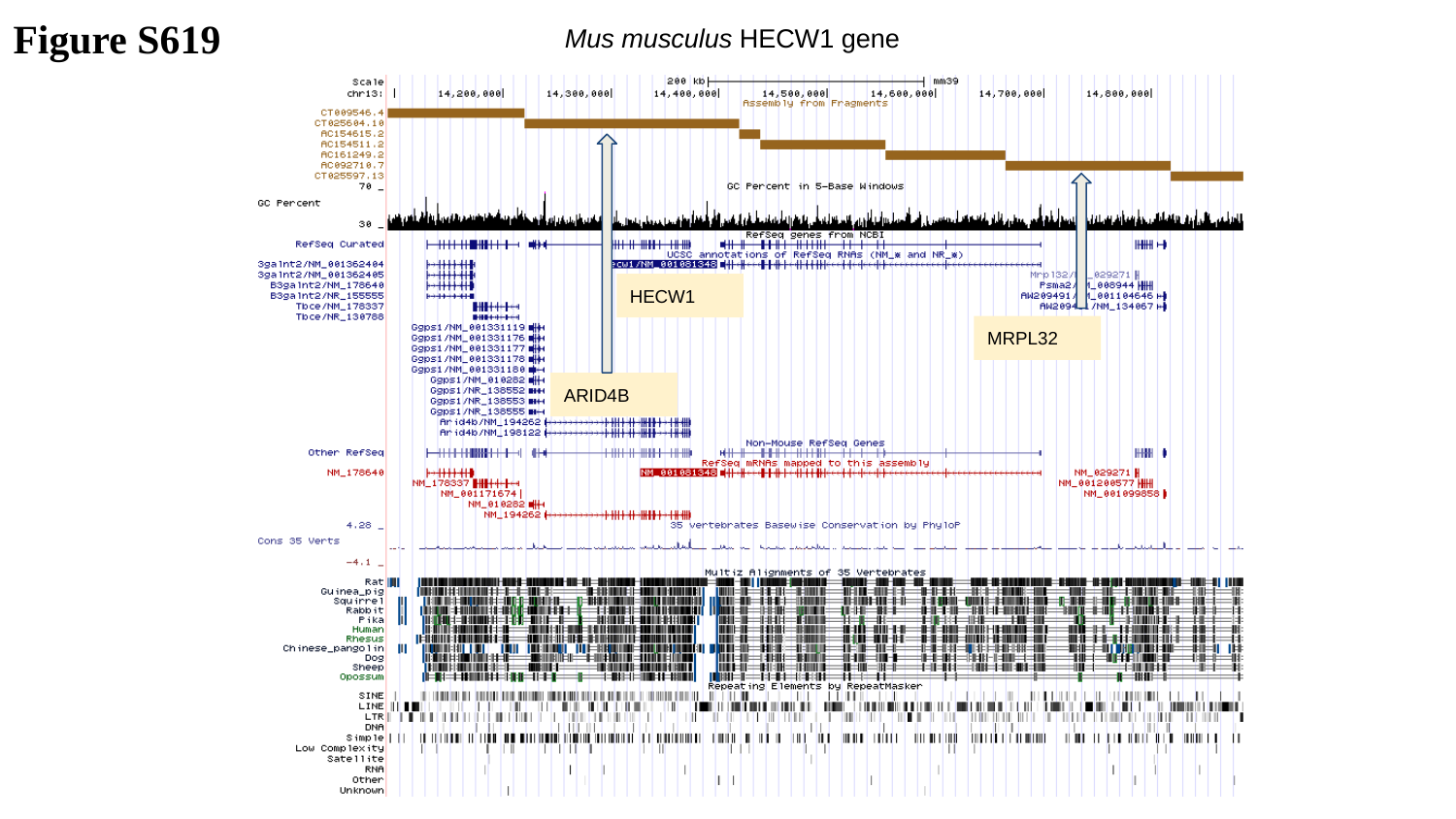

Figure S619
Mus musculus HECW1 gene
HECW1
MRPL32
ARID4B

### Slide 620
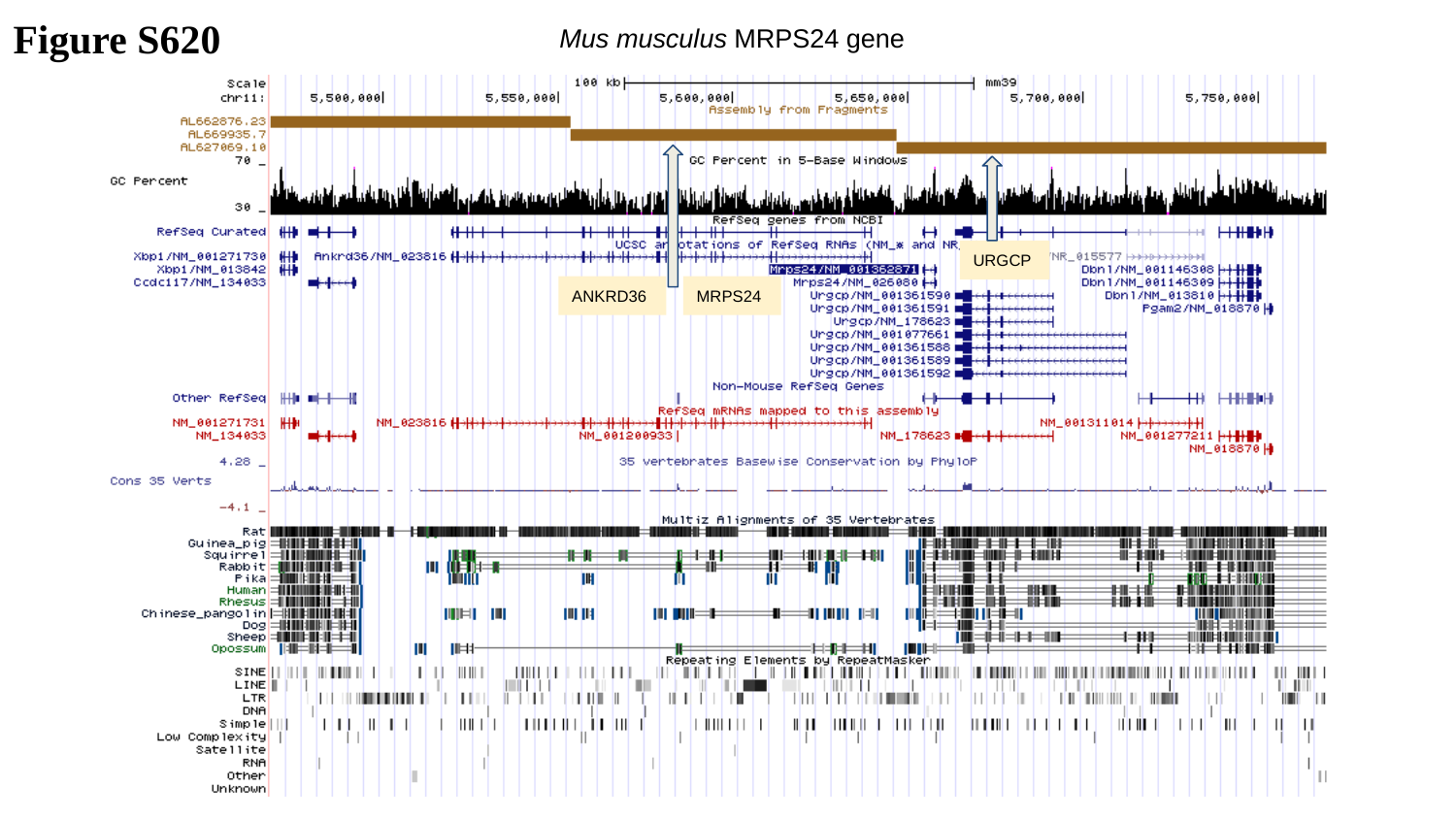

Figure S620
Mus musculus MRPS24 gene
URGCP
ANKRD36
MRPS24

### Slide 621
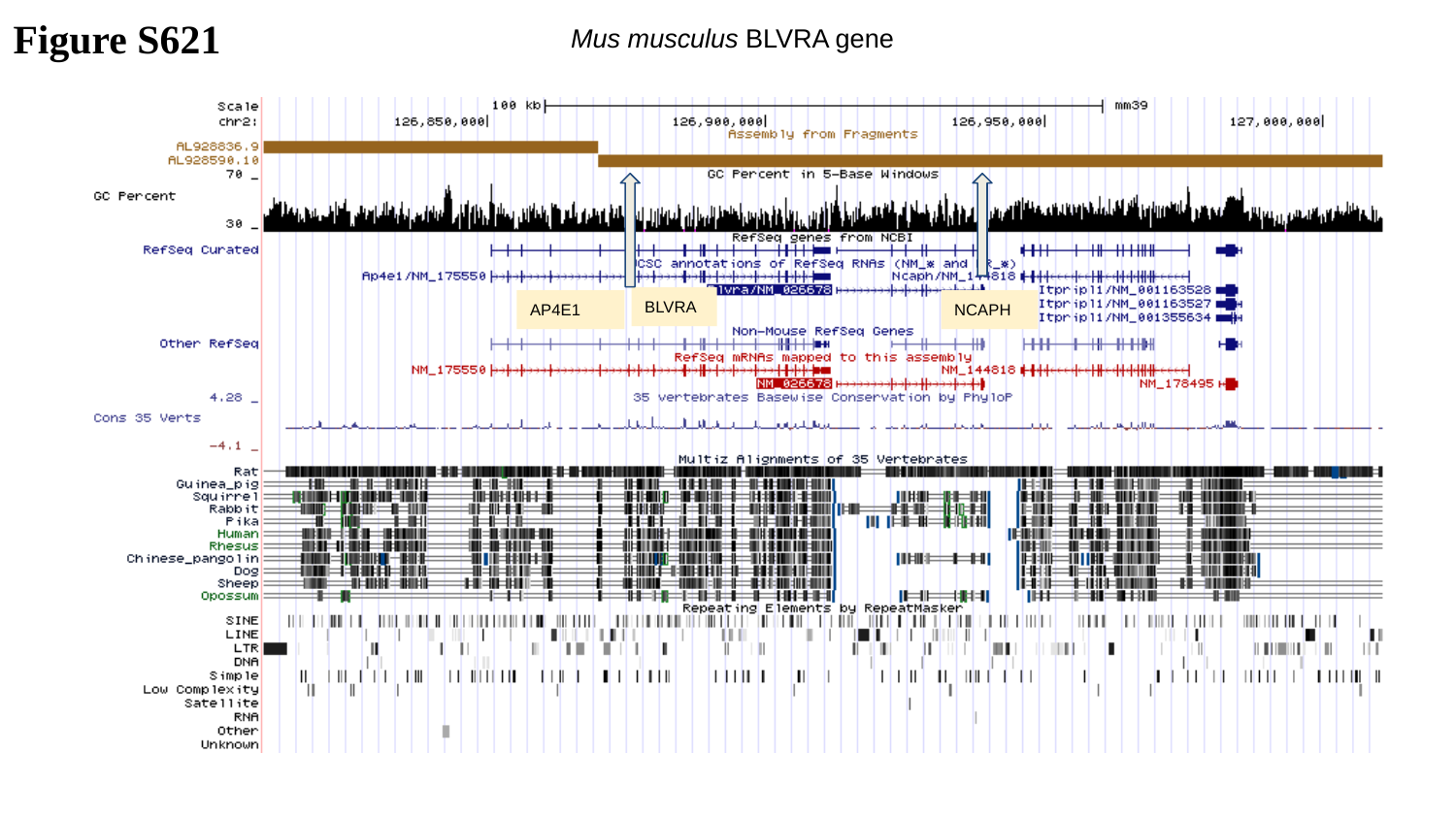

Figure S621
Mus musculus BLVRA gene
BLVRA
AP4E1
NCAPH

### Slide 622
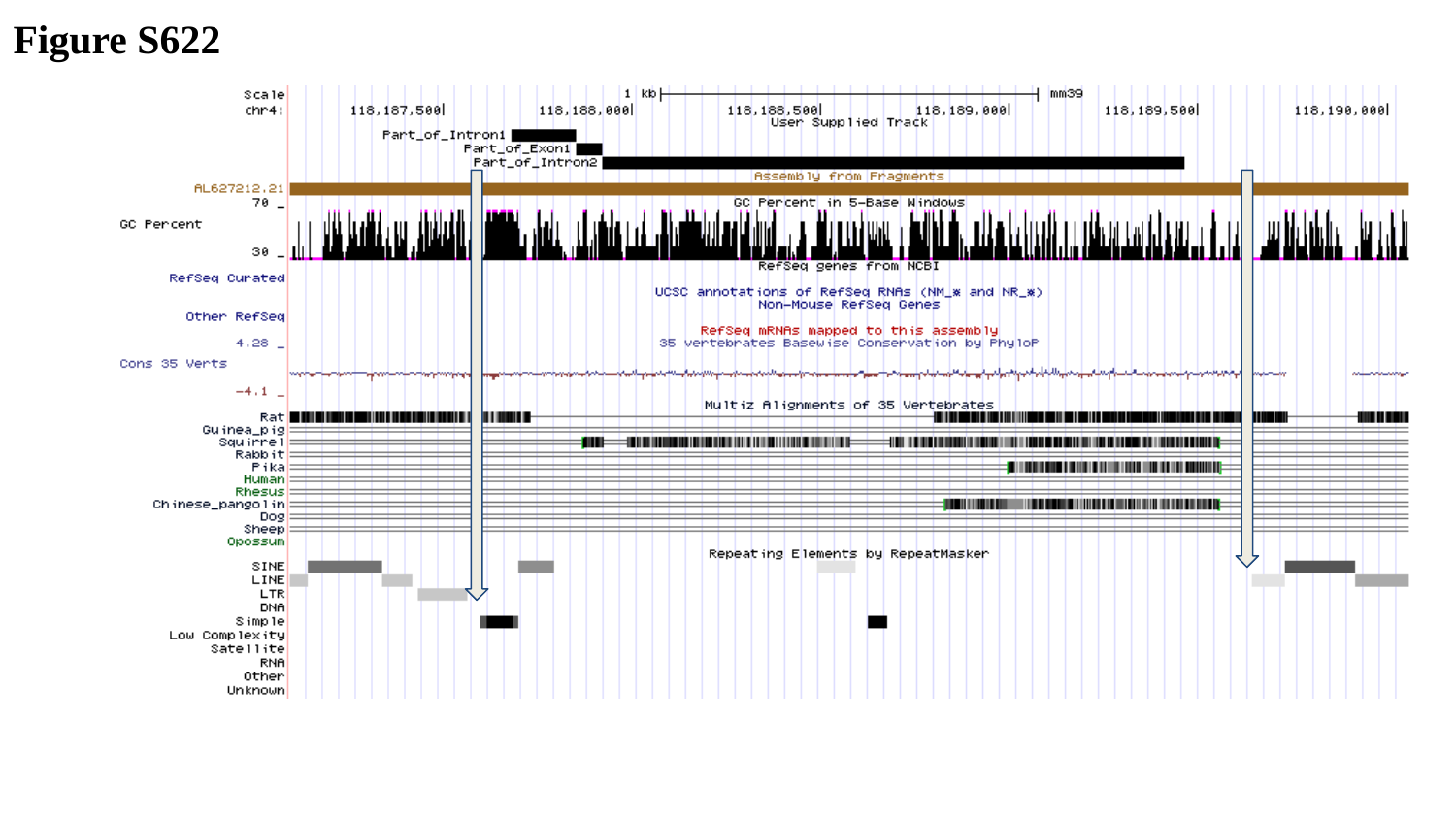

Figure S622

### Slide 623
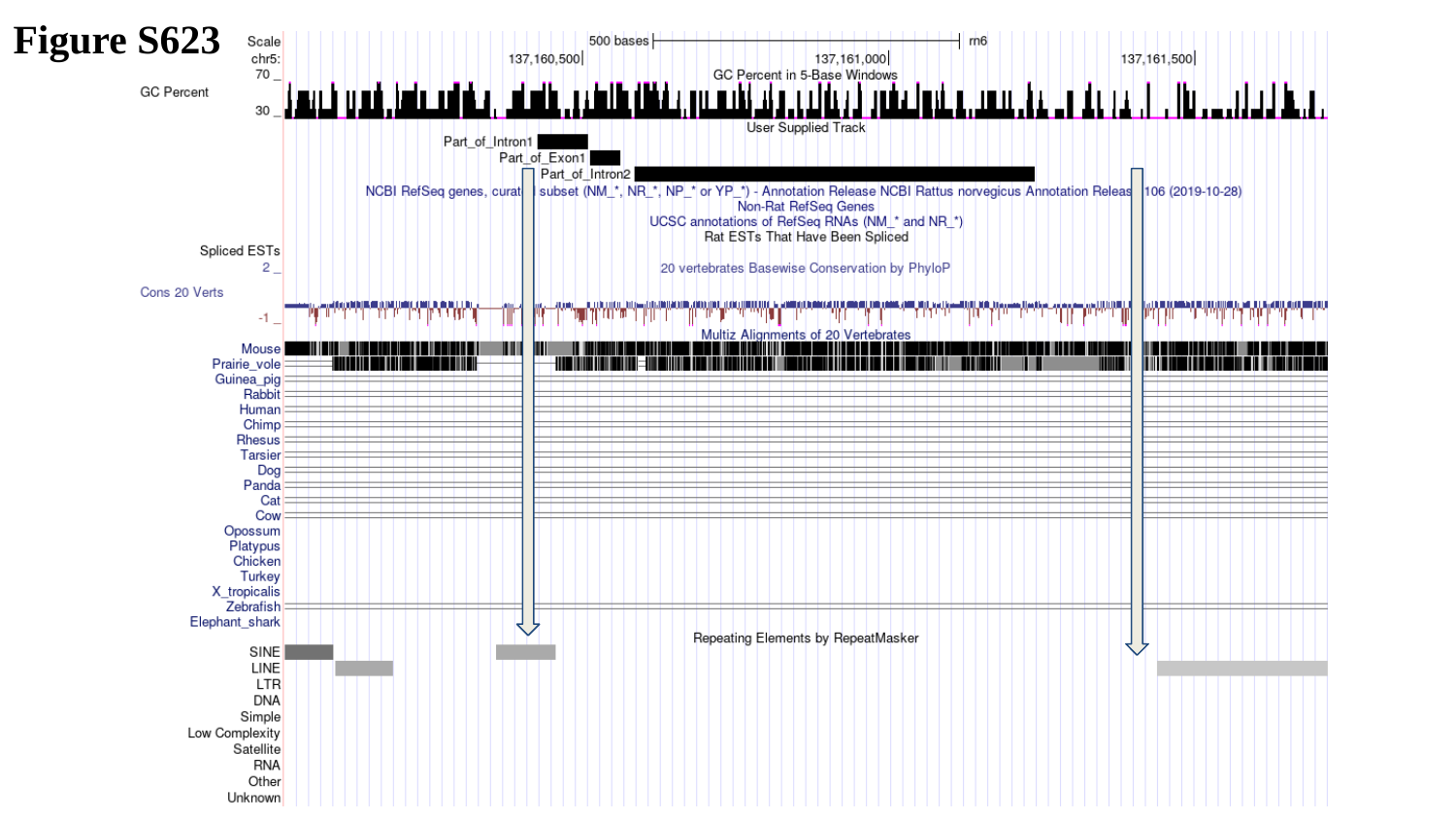

Figure S623

### Slide 624
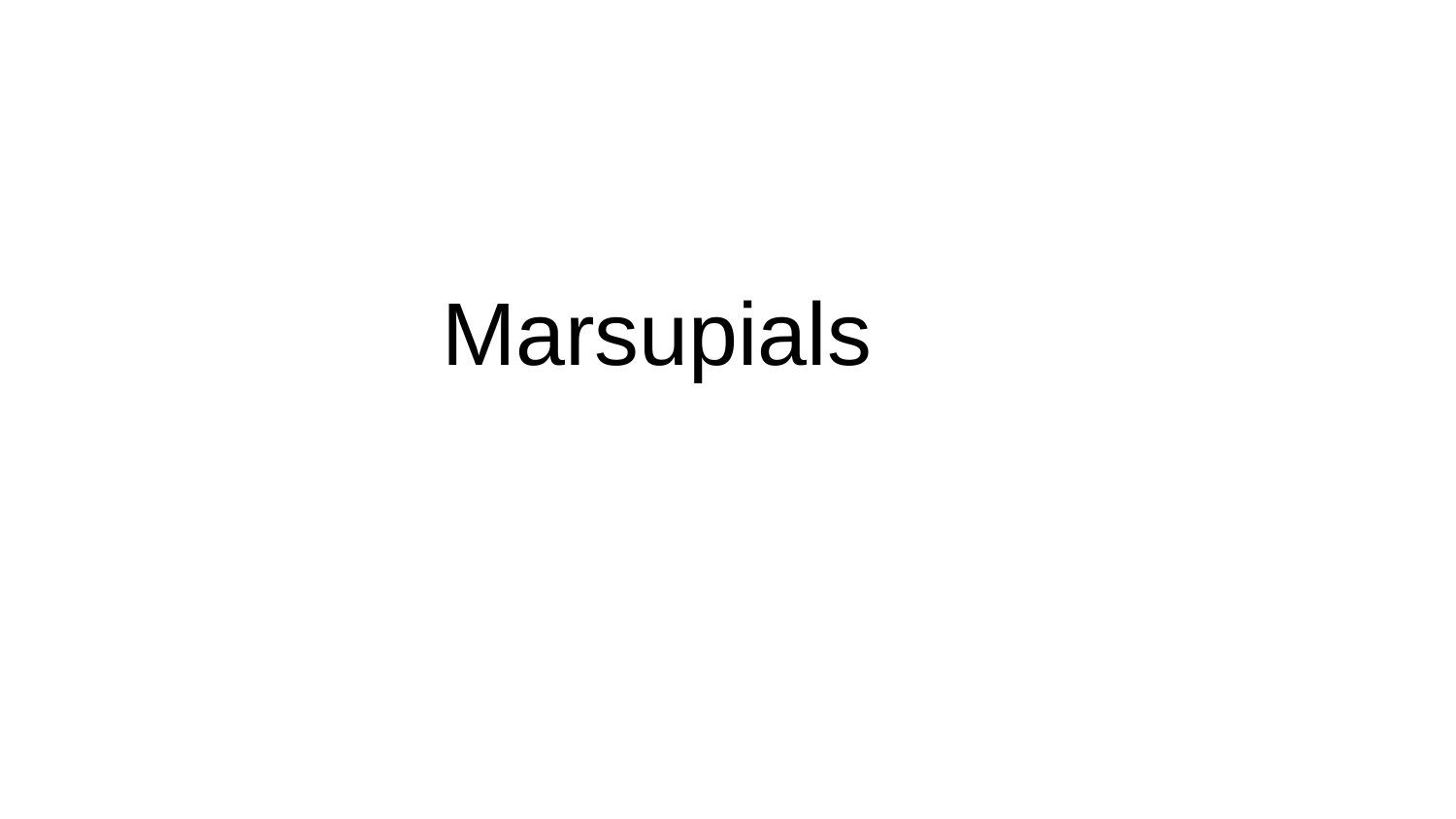

Marsupials

### Slide 625
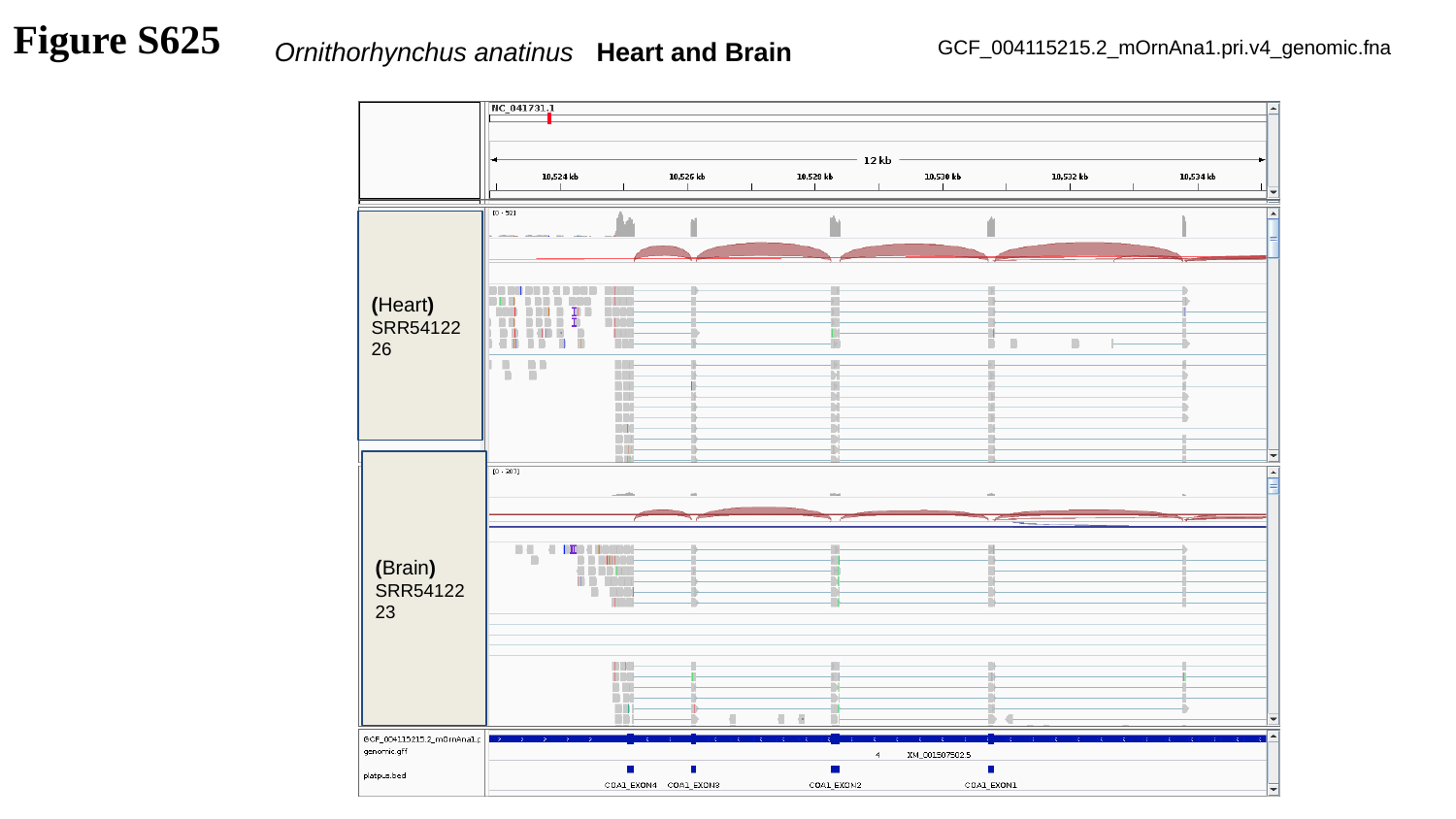

Figure S625
Ornithorhynchus anatinus
Heart and Brain
GCF_004115215.2_mOrnAna1.pri.v4_genomic.fna
(Heart)
SRR5412226
(Brain)
SRR5412223

### Slide 626
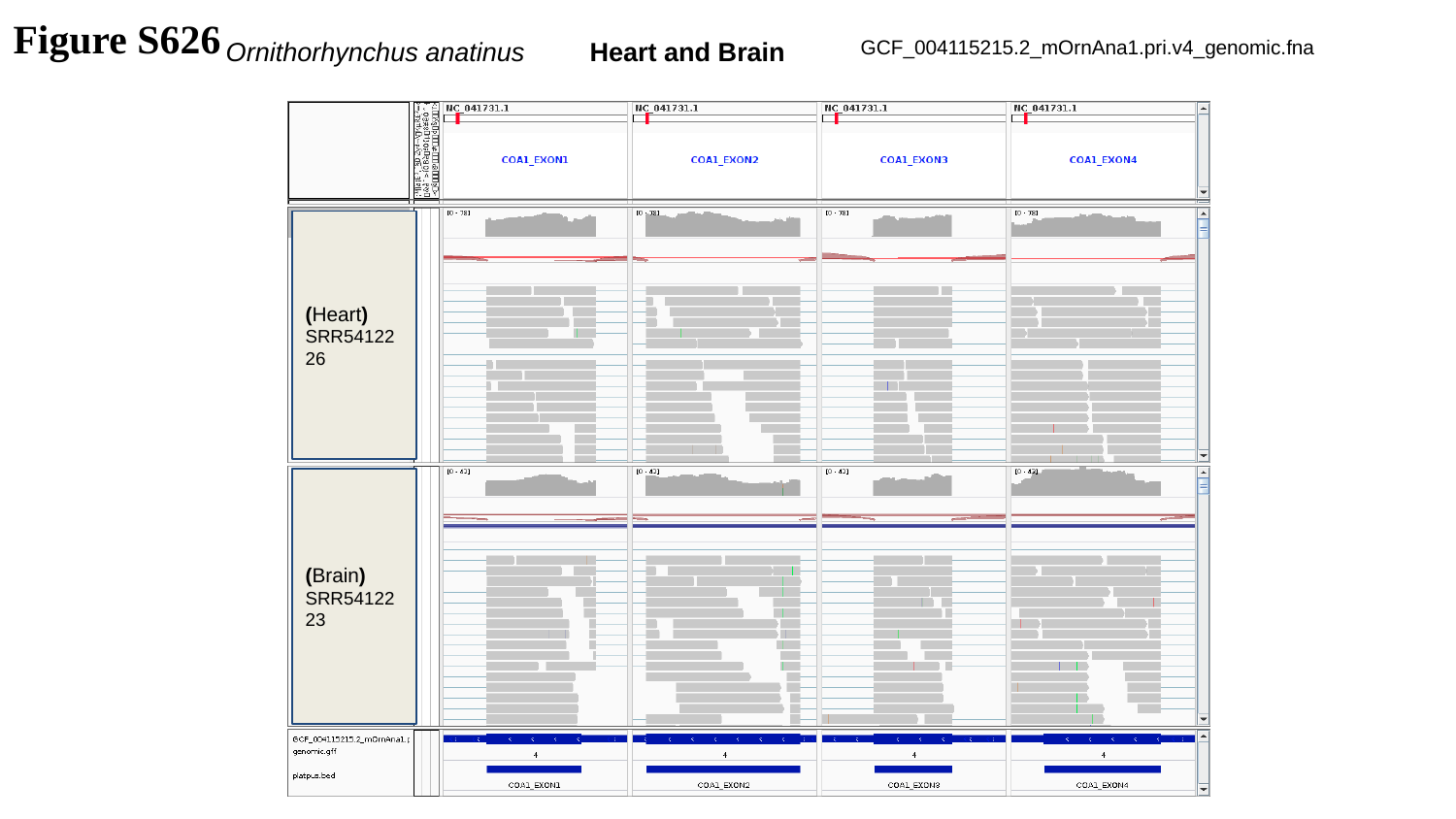

Figure S626
Ornithorhynchus anatinus
Heart and Brain
GCF_004115215.2_mOrnAna1.pri.v4_genomic.fna
(Heart)
SRR5412226
(Brain)
SRR5412223

### Slide 627
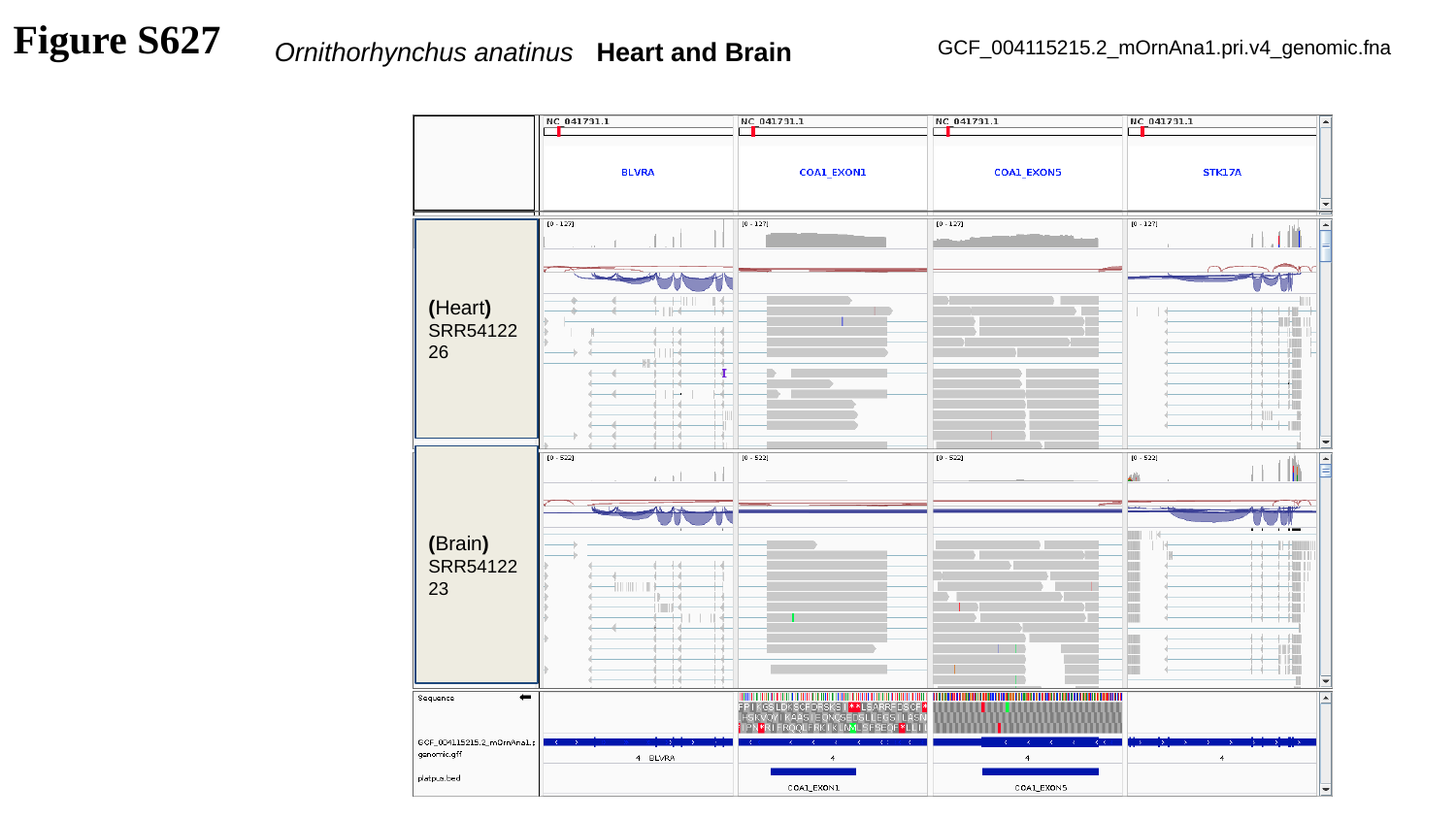

Figure S627
Ornithorhynchus anatinus
Heart and Brain
GCF_004115215.2_mOrnAna1.pri.v4_genomic.fna
(Heart)
SRR5412226
(Brain)
SRR5412223

### Slide 628
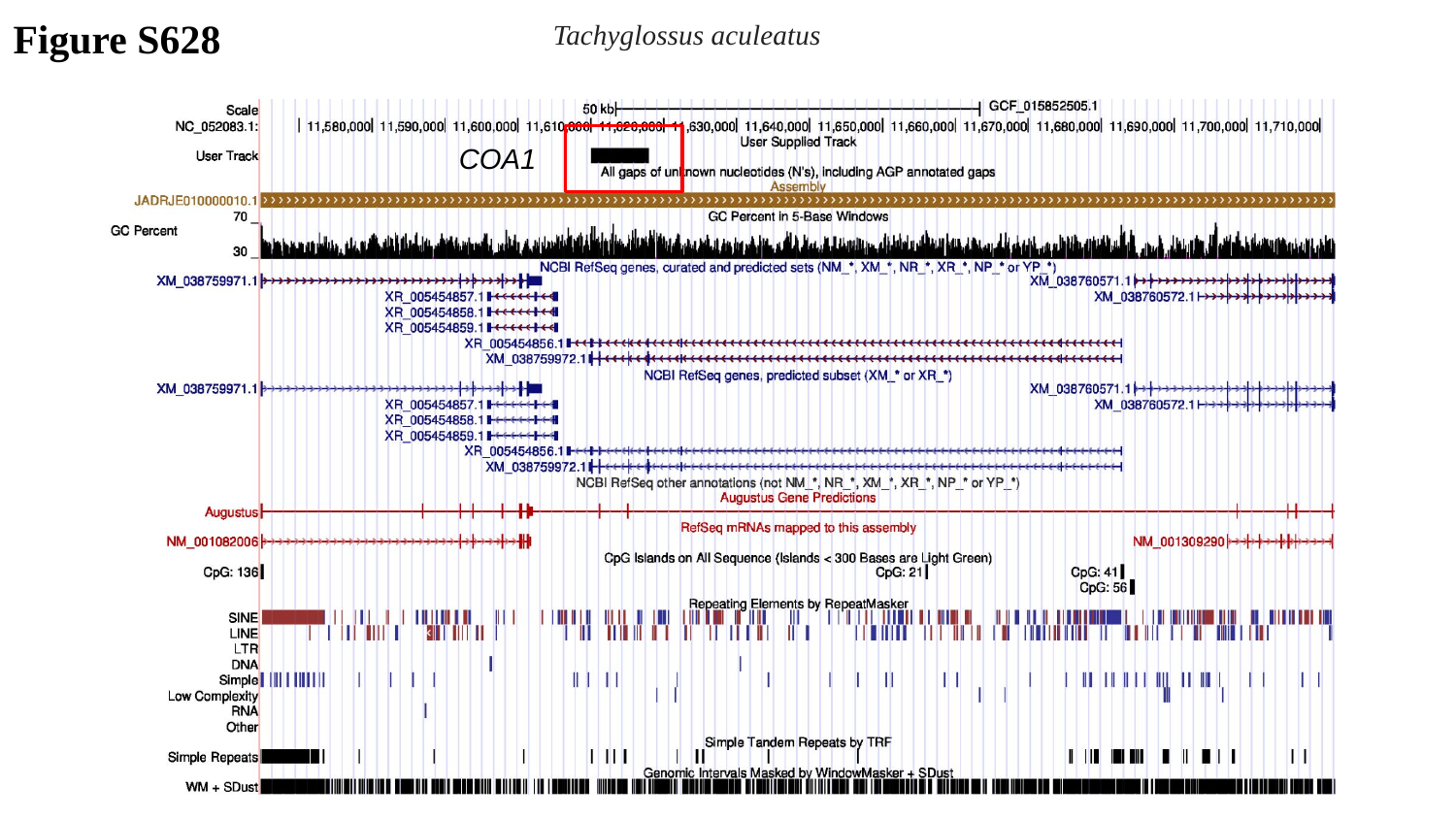

Figure S628
Tachyglossus aculeatus
COA1

### Slide 629
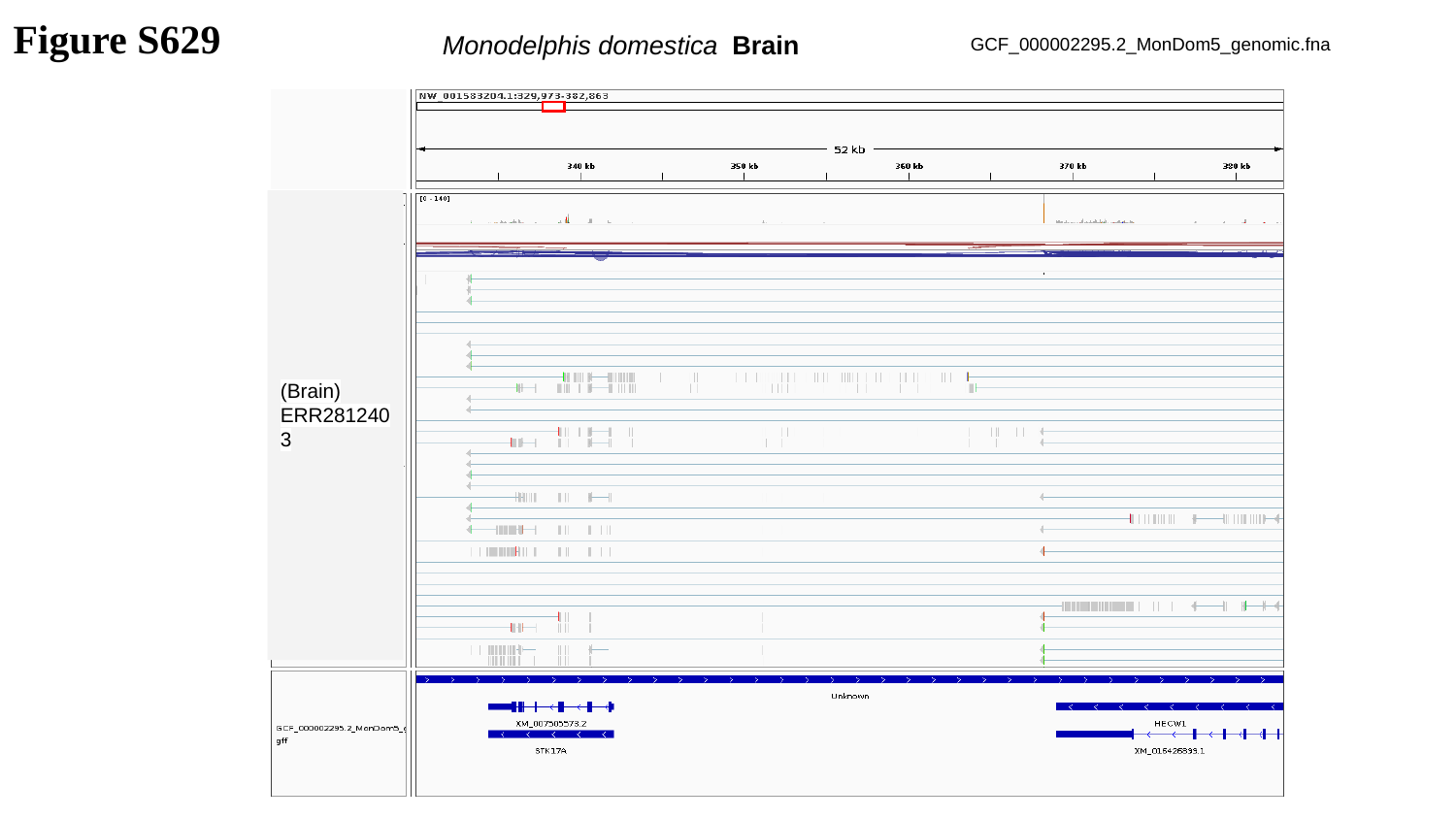

Figure S629
Monodelphis domestica Brain
GCF_000002295.2_MonDom5_genomic.fna
(Brain)
ERR2812403

### Slide 630
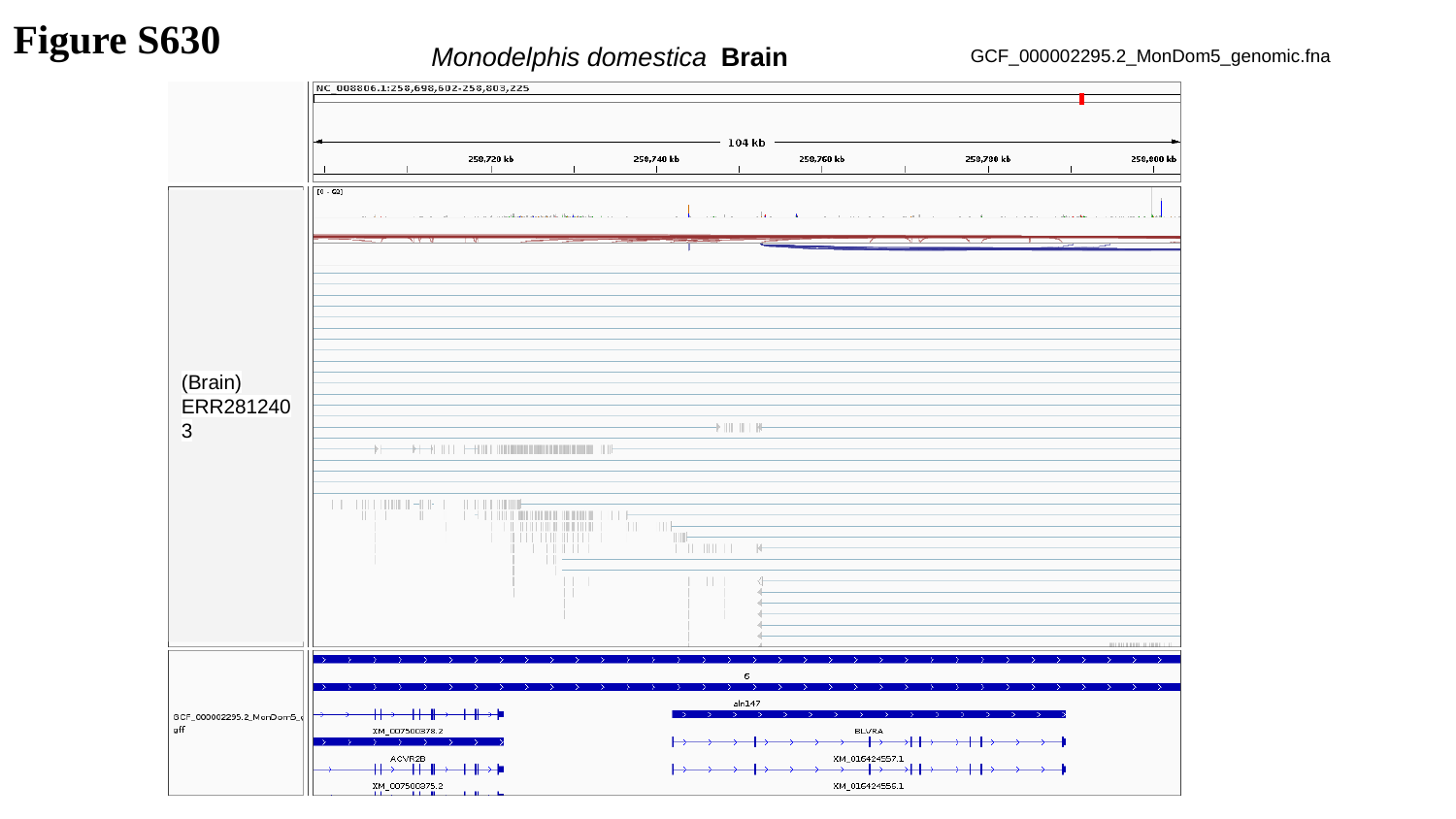

Figure S630
Monodelphis domestica Brain
GCF_000002295.2_MonDom5_genomic.fna
(Brain)
ERR2812403

### Slide 631
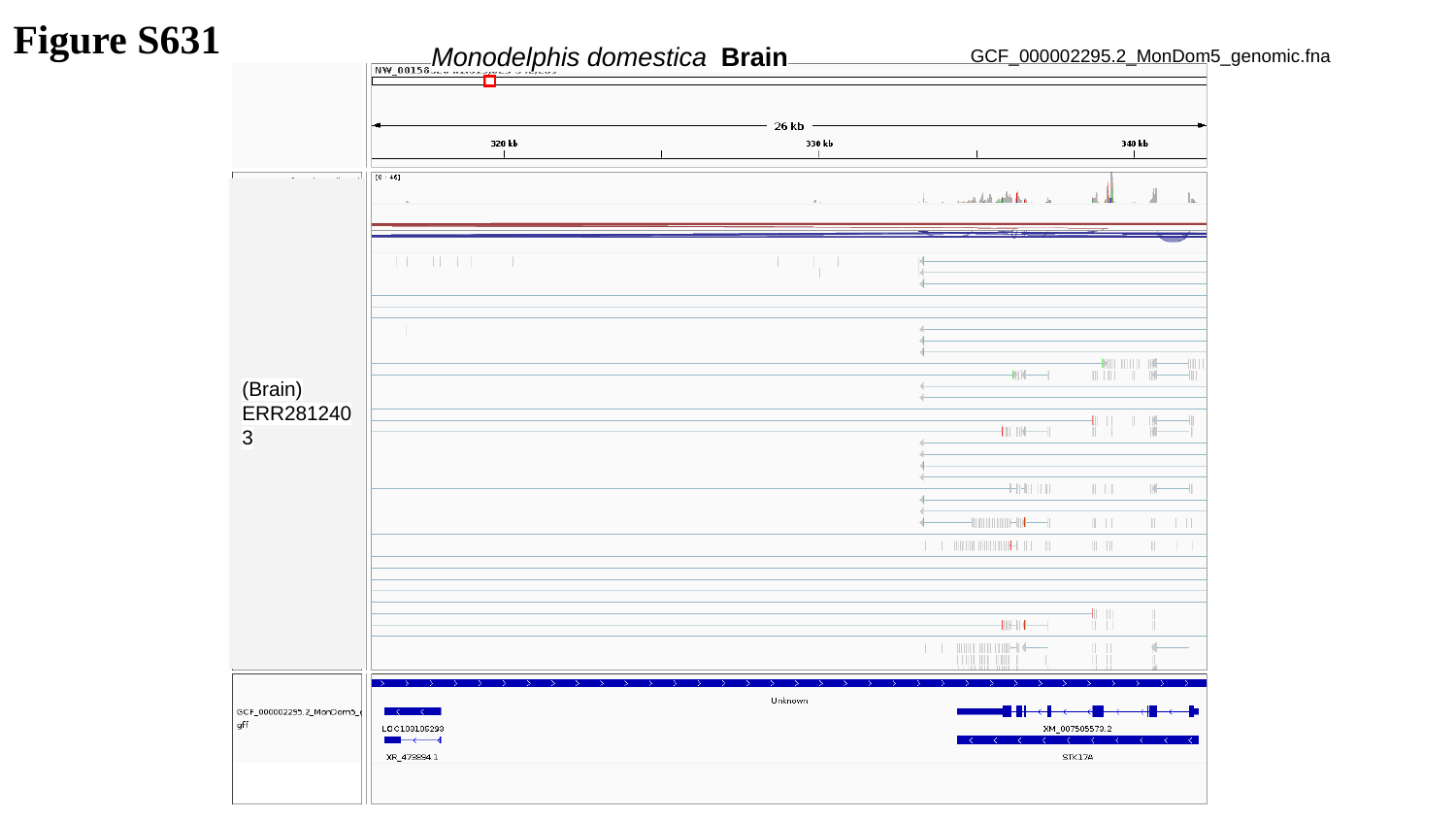

Figure S631
Monodelphis domestica Brain
GCF_000002295.2_MonDom5_genomic.fna
(Brain)
ERR2812403

### Slide 632
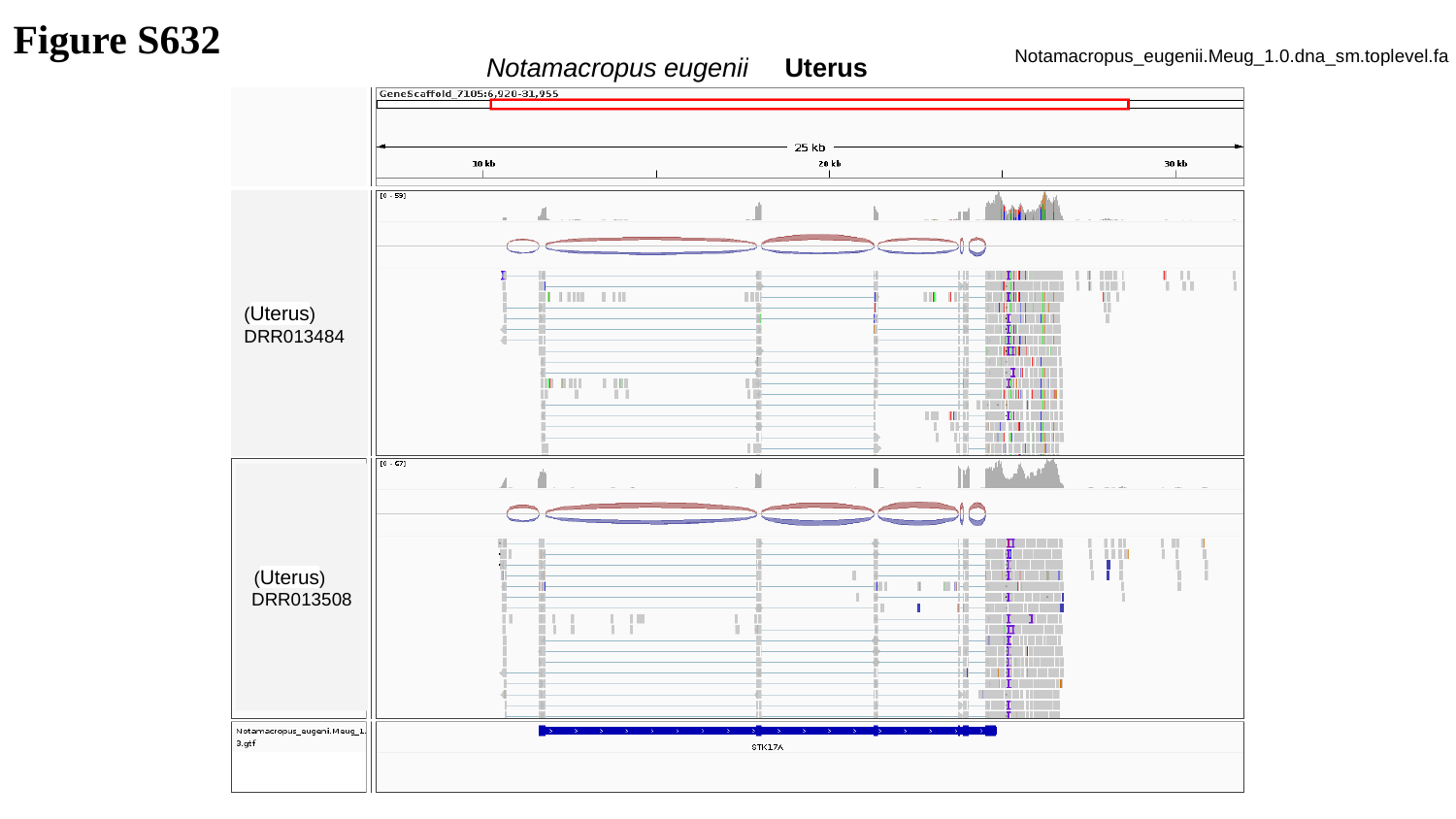

Figure S632
Notamacropus eugenii Uterus
Notamacropus_eugenii.Meug_1.0.dna_sm.toplevel.fa
(Uterus)
DRR013484
 (Uterus)
DRR013508

### Slide 633
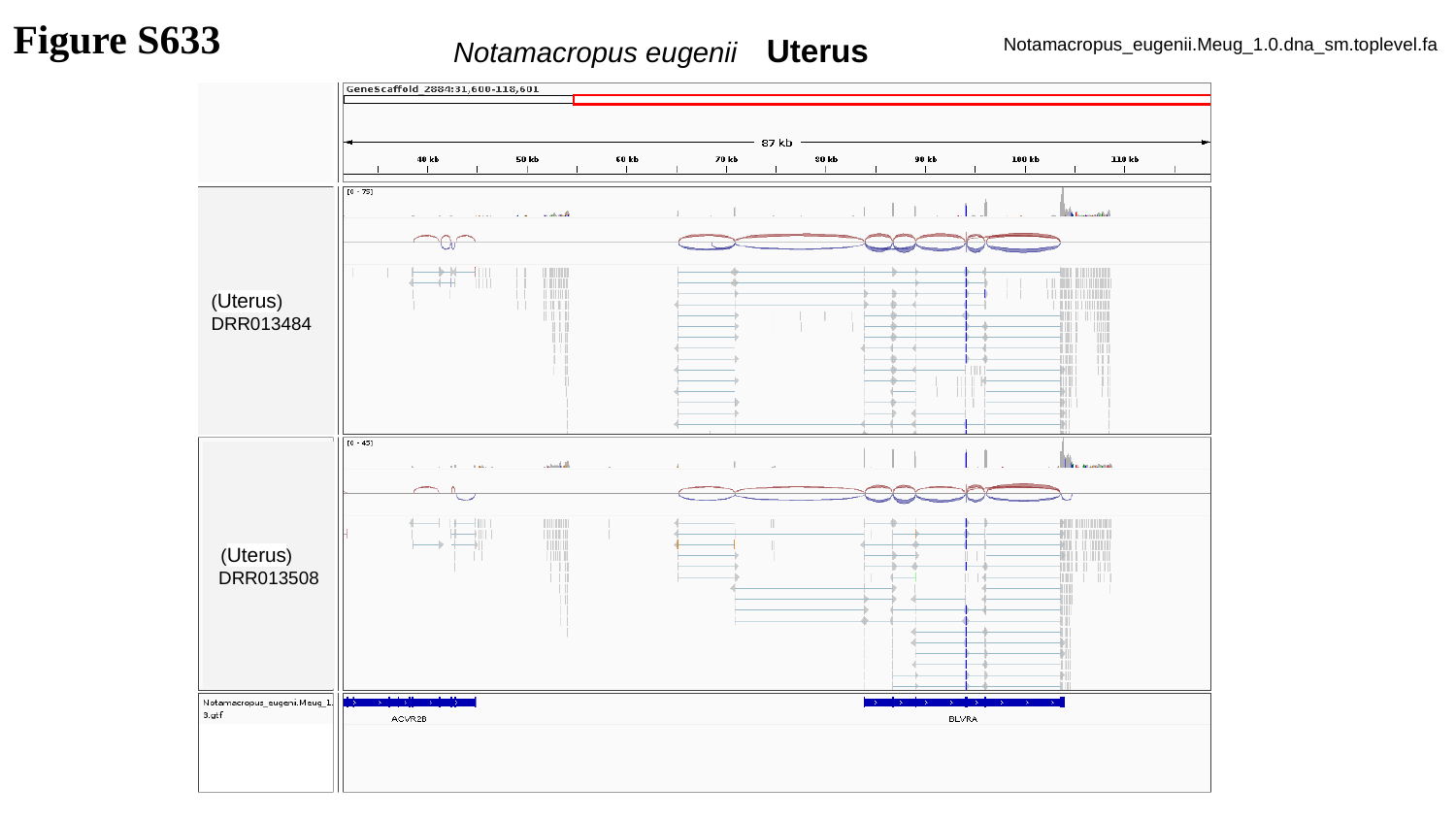

Figure S633
Notamacropus eugenii Uterus
Notamacropus_eugenii.Meug_1.0.dna_sm.toplevel.fa
(Uterus)
DRR013484
 (Uterus)
DRR013508

### Slide 634
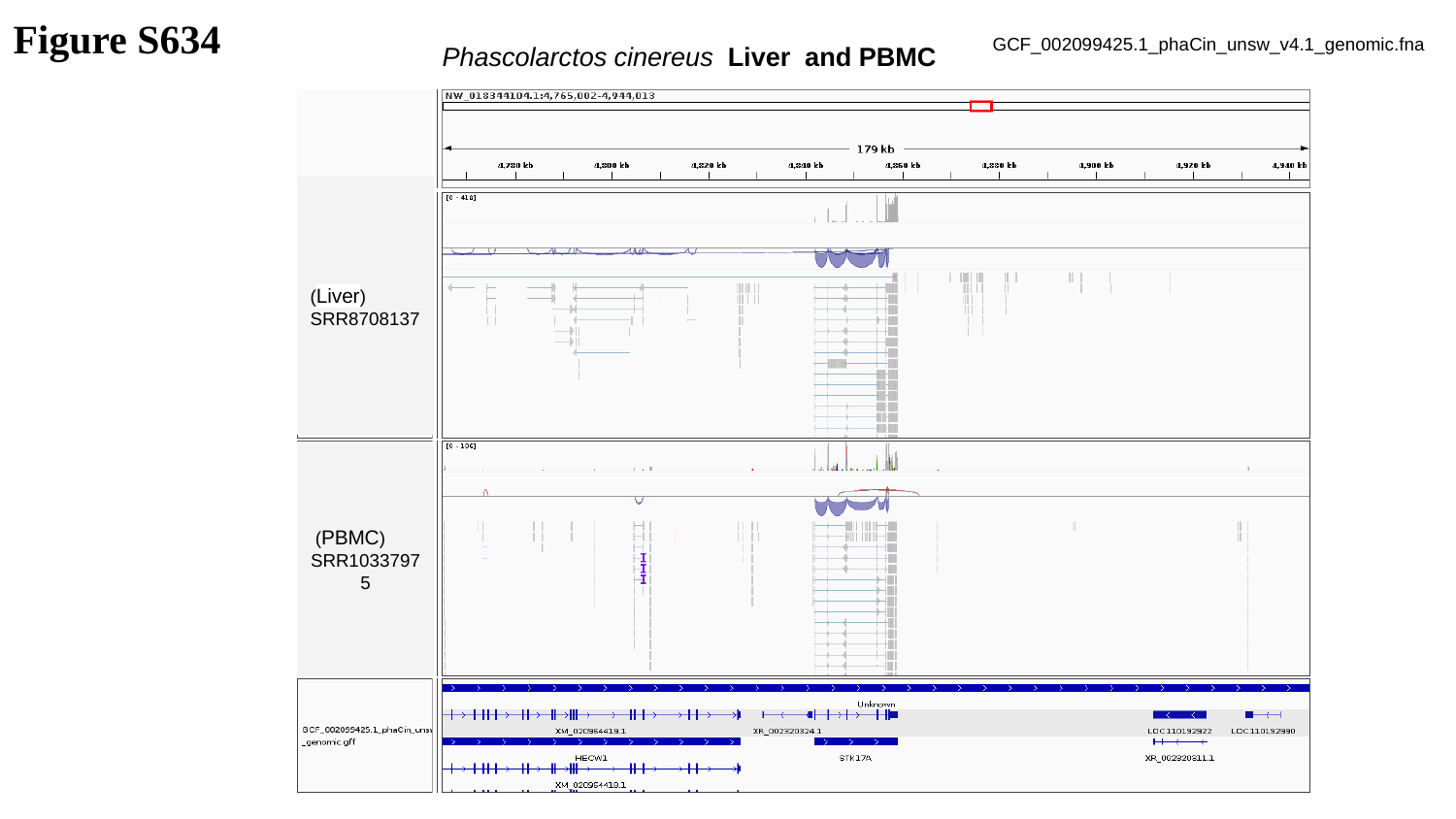

Figure S634
Phascolarctos cinereus Liver and PBMC
GCF_002099425.1_phaCin_unsw_v4.1_genomic.fna
(Liver)
SRR8708137
 (PBMC)
SRR10337975

### Slide 635
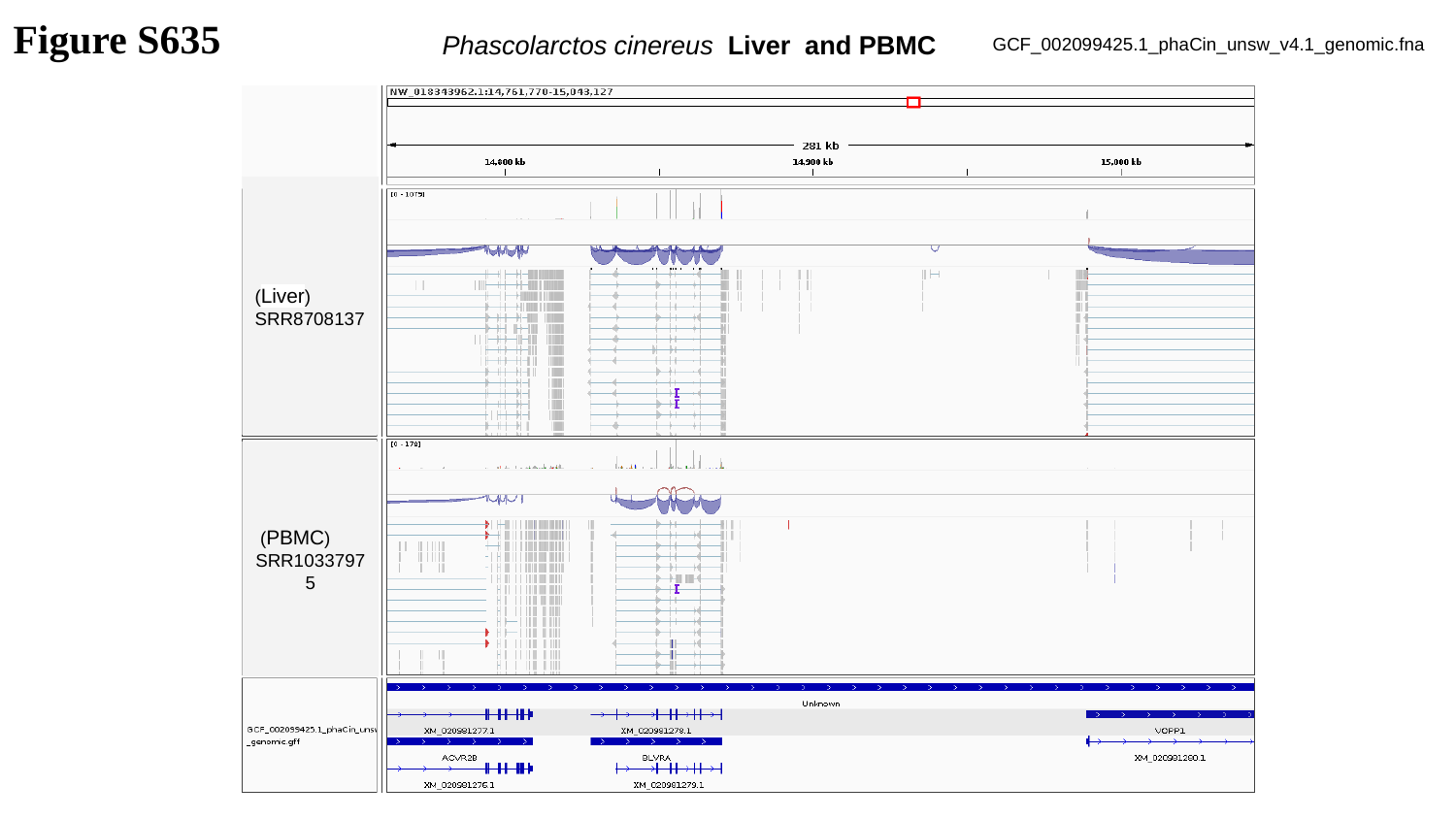

Figure S635
Phascolarctos cinereus Liver and PBMC
GCF_002099425.1_phaCin_unsw_v4.1_genomic.fna
(Liver)
SRR8708137
 (PBMC)
SRR10337975

### Slide 636
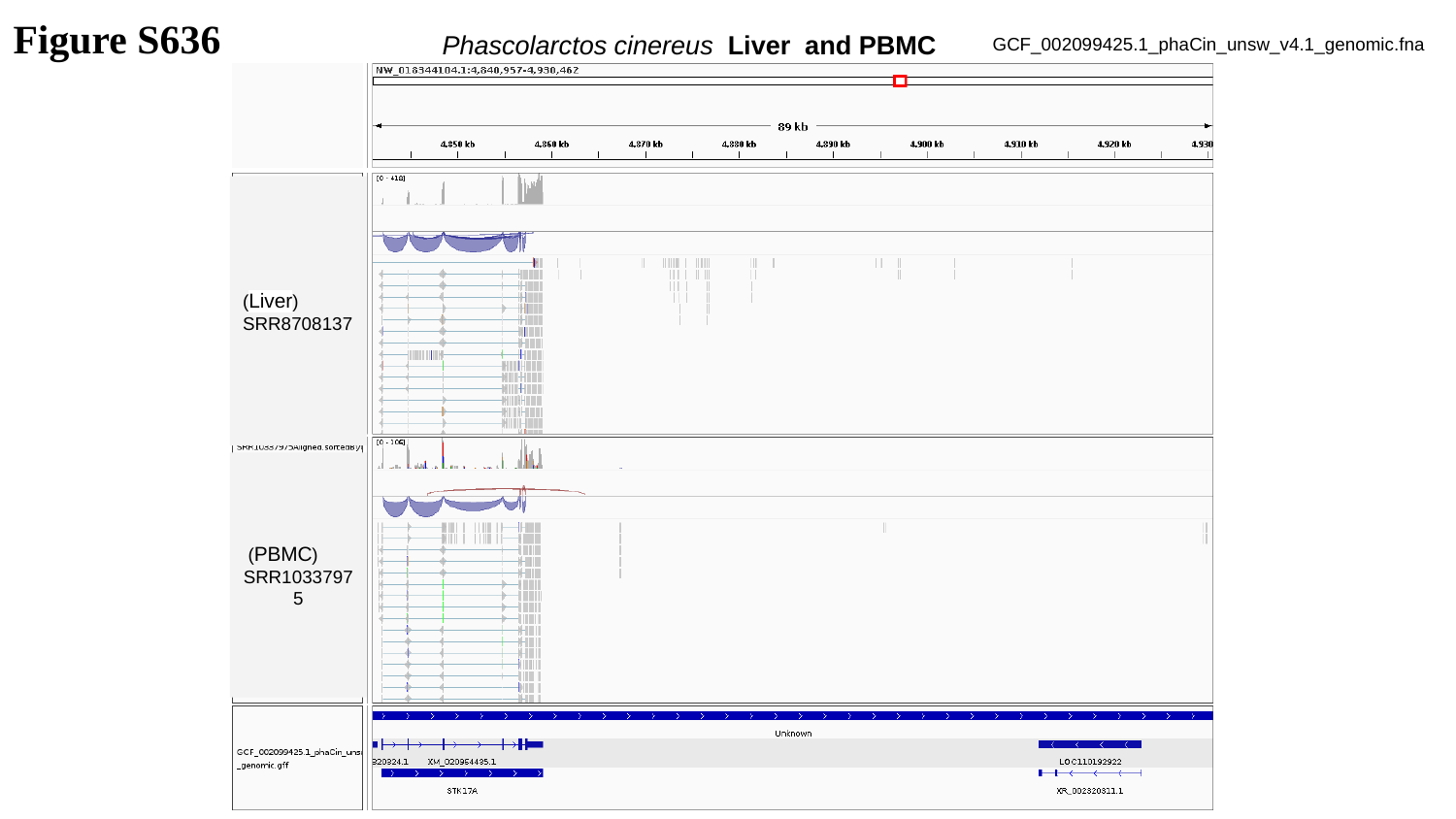

Figure S636
Phascolarctos cinereus Liver and PBMC
GCF_002099425.1_phaCin_unsw_v4.1_genomic.fna
(Liver)
SRR8708137
 (PBMC)
SRR10337975

### Slide 637
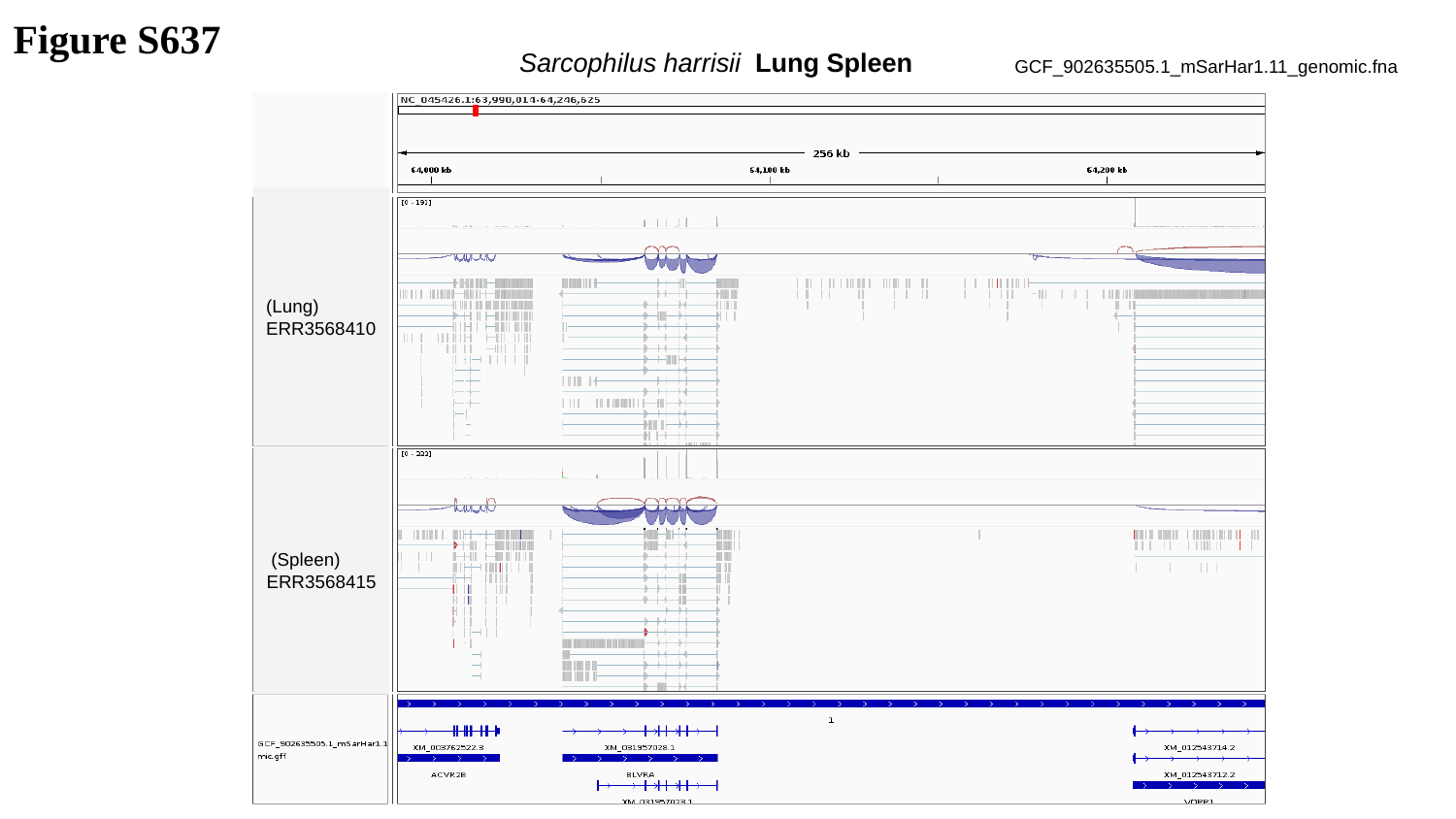

Figure S637
Sarcophilus harrisii Lung Spleen
GCF_902635505.1_mSarHar1.11_genomic.fna
(Lung)
ERR3568410
 (Spleen)
ERR3568415

### Slide 638
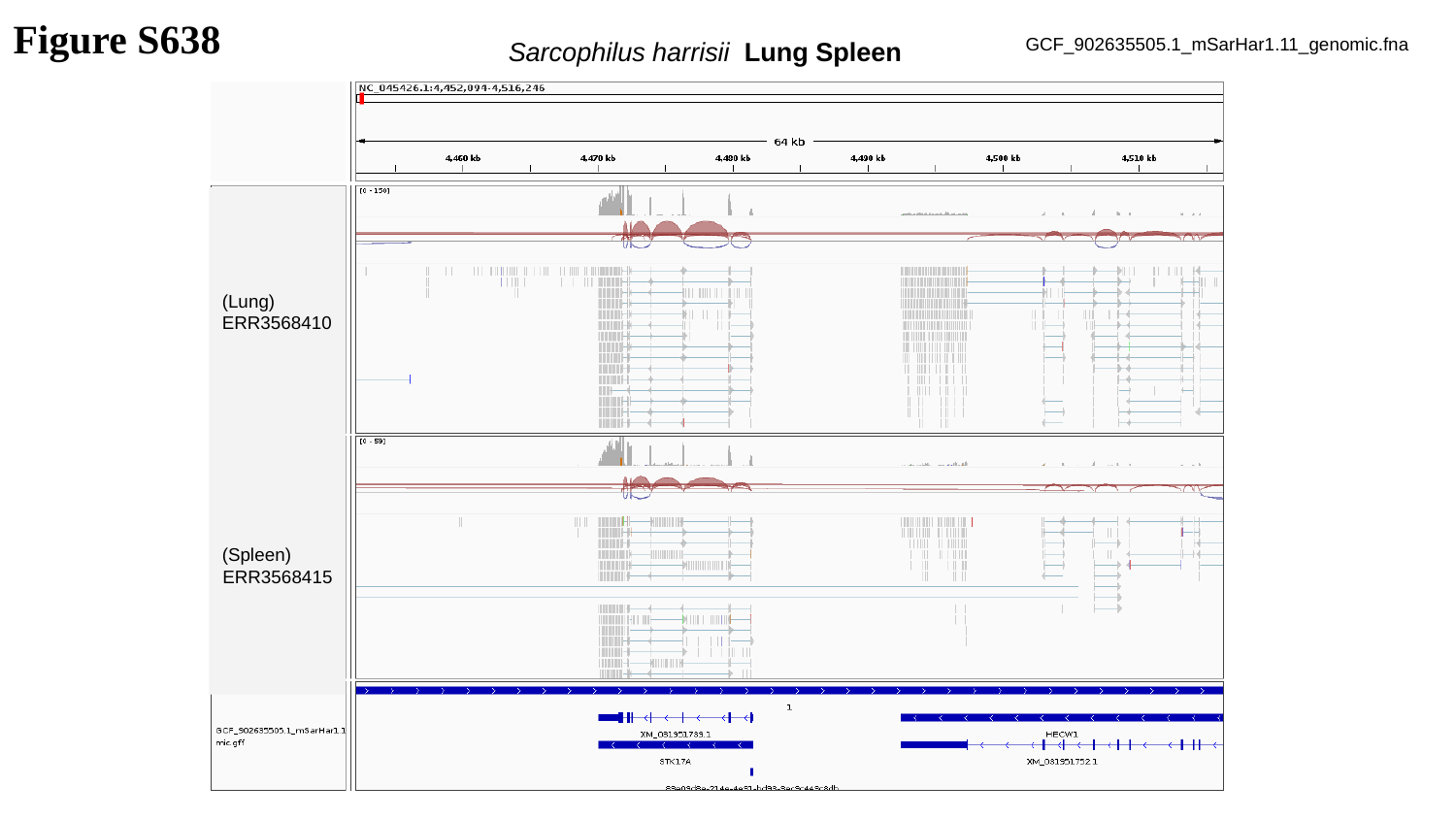

Figure S638
Sarcophilus harrisii Lung Spleen
GCF_902635505.1_mSarHar1.11_genomic.fna
(Lung)
ERR3568410
(Spleen)
ERR3568415

### Slide 639
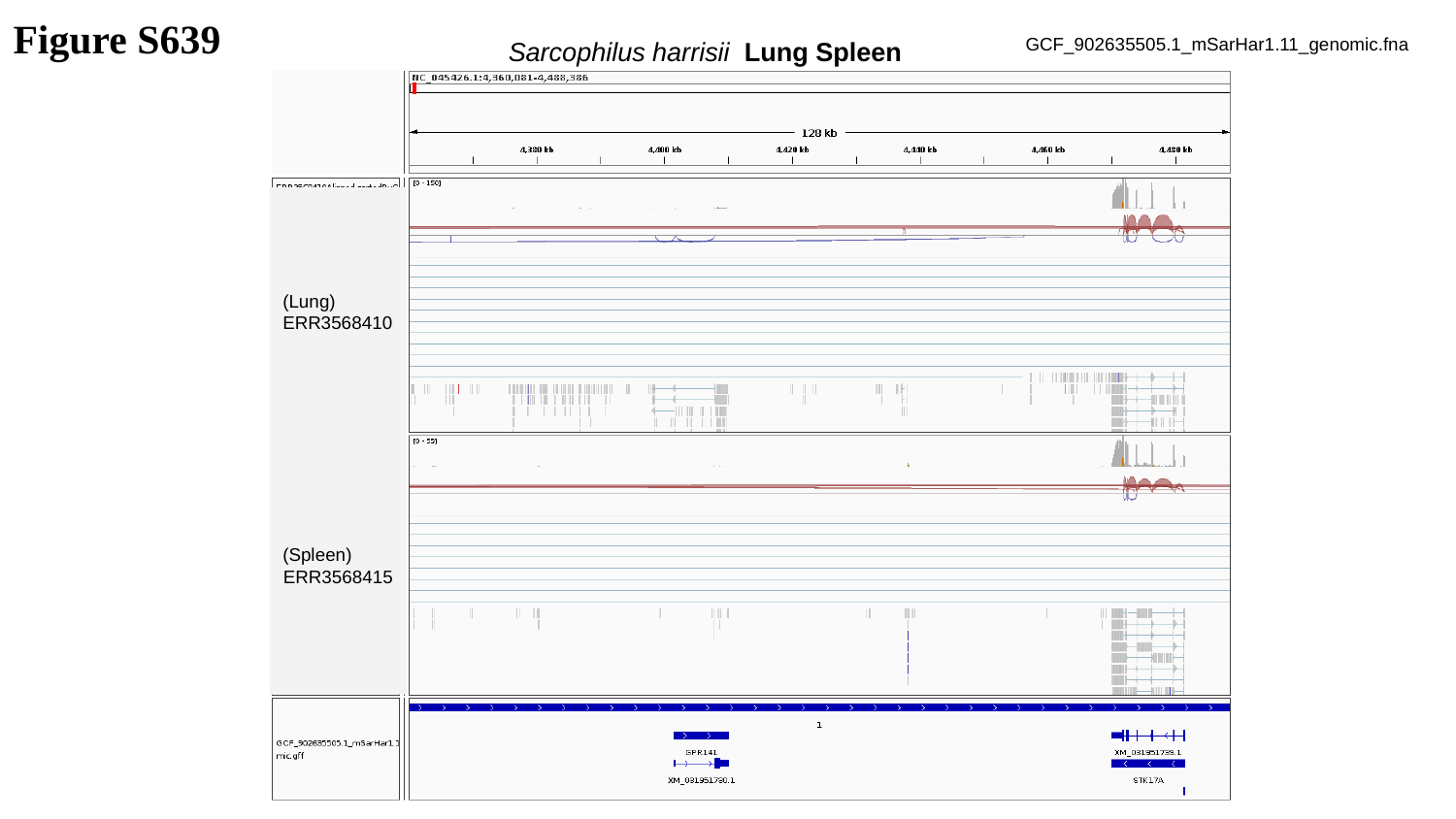

Figure S639
Sarcophilus harrisii Lung Spleen
GCF_902635505.1_mSarHar1.11_genomic.fna
(Lung)
ERR3568410
(Spleen)
ERR3568415

### Slide 640
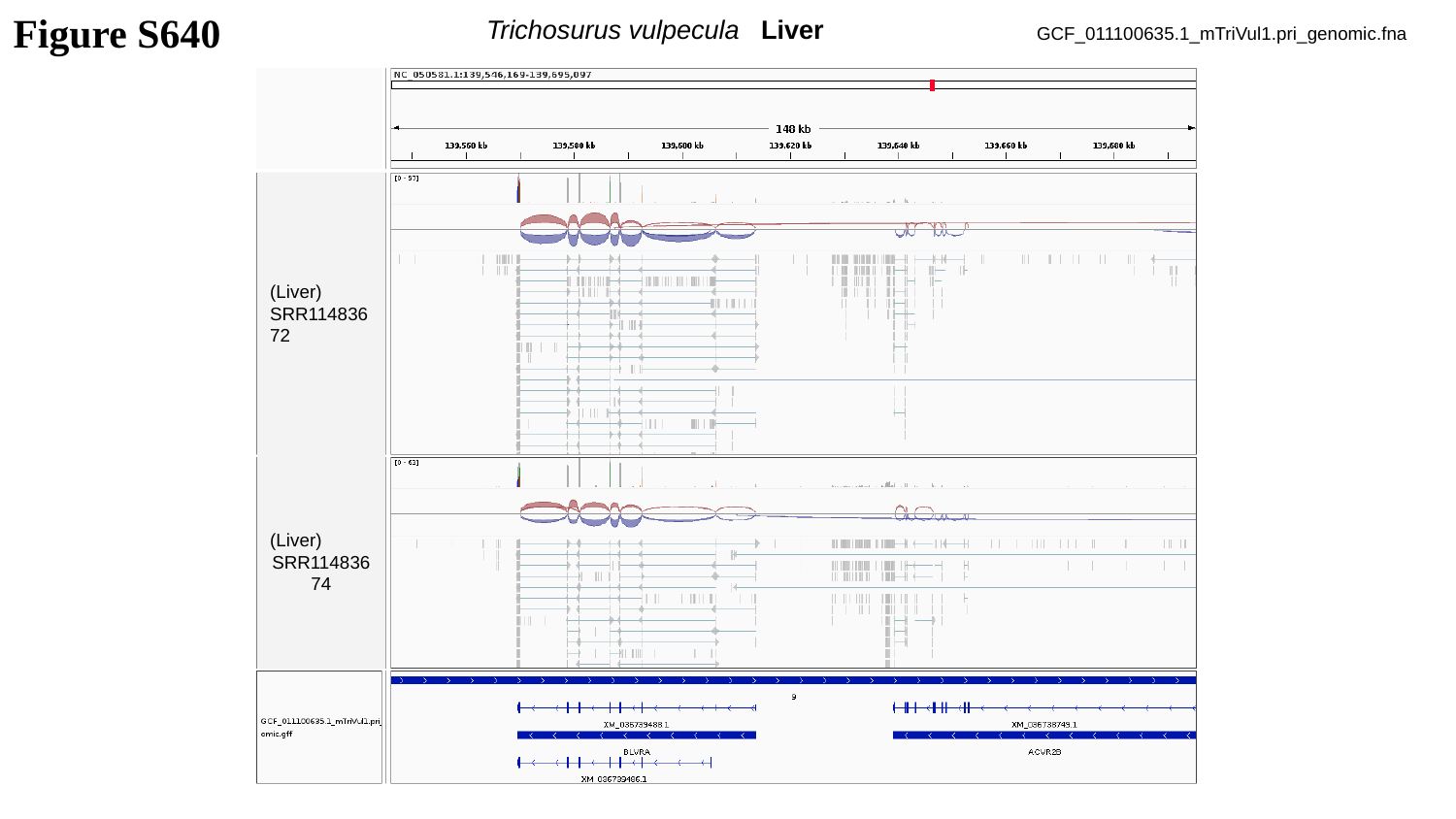

Trichosurus vulpecula Liver
Figure S640
GCF_011100635.1_mTriVul1.pri_genomic.fna
(Liver)
SRR11483672
(Liver)
SRR11483674

### Slide 641
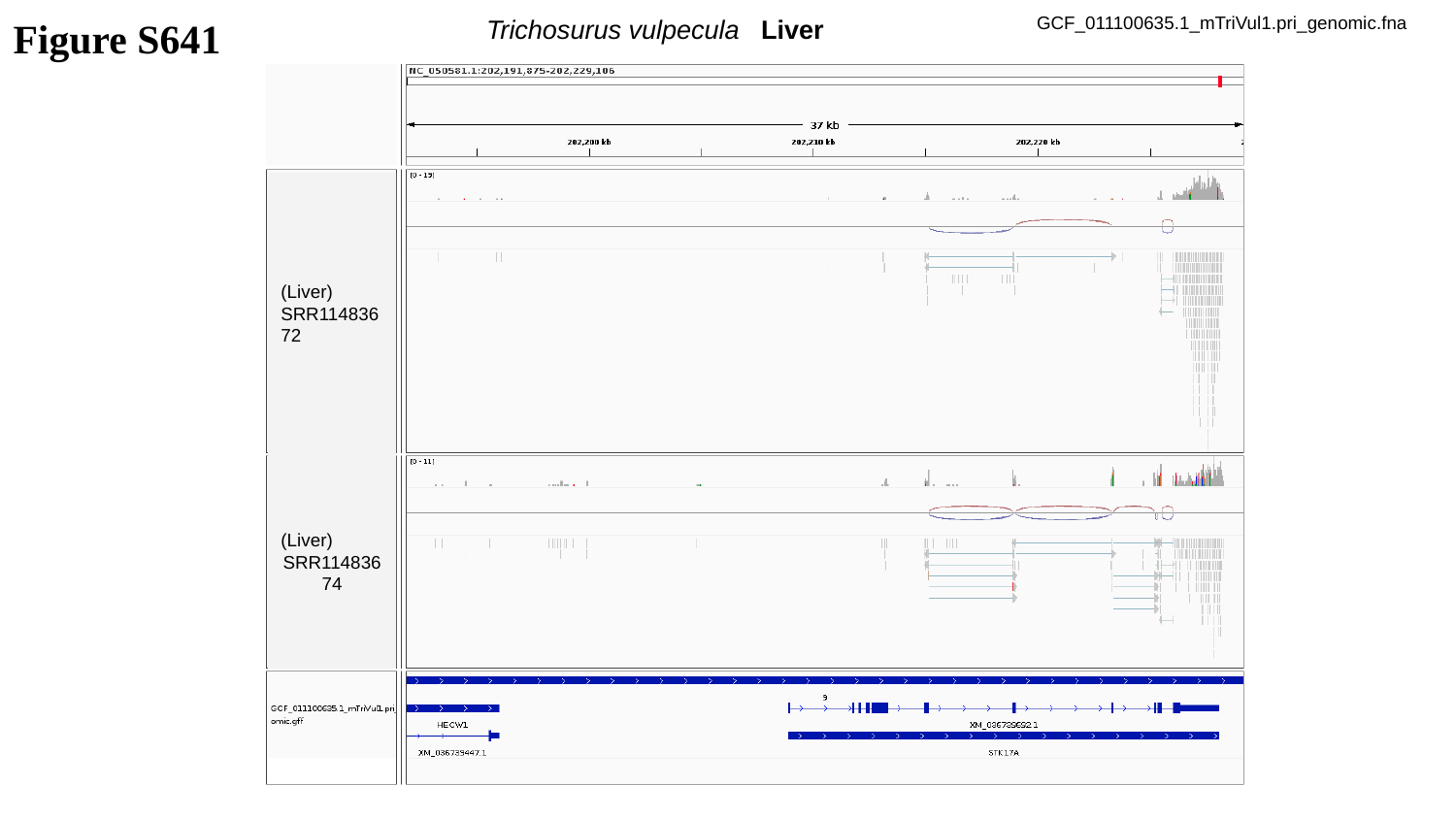

Figure S641
Trichosurus vulpecula Liver
GCF_011100635.1_mTriVul1.pri_genomic.fna
(Liver)
SRR11483672
(Liver)
SRR11483674

### Slide 642
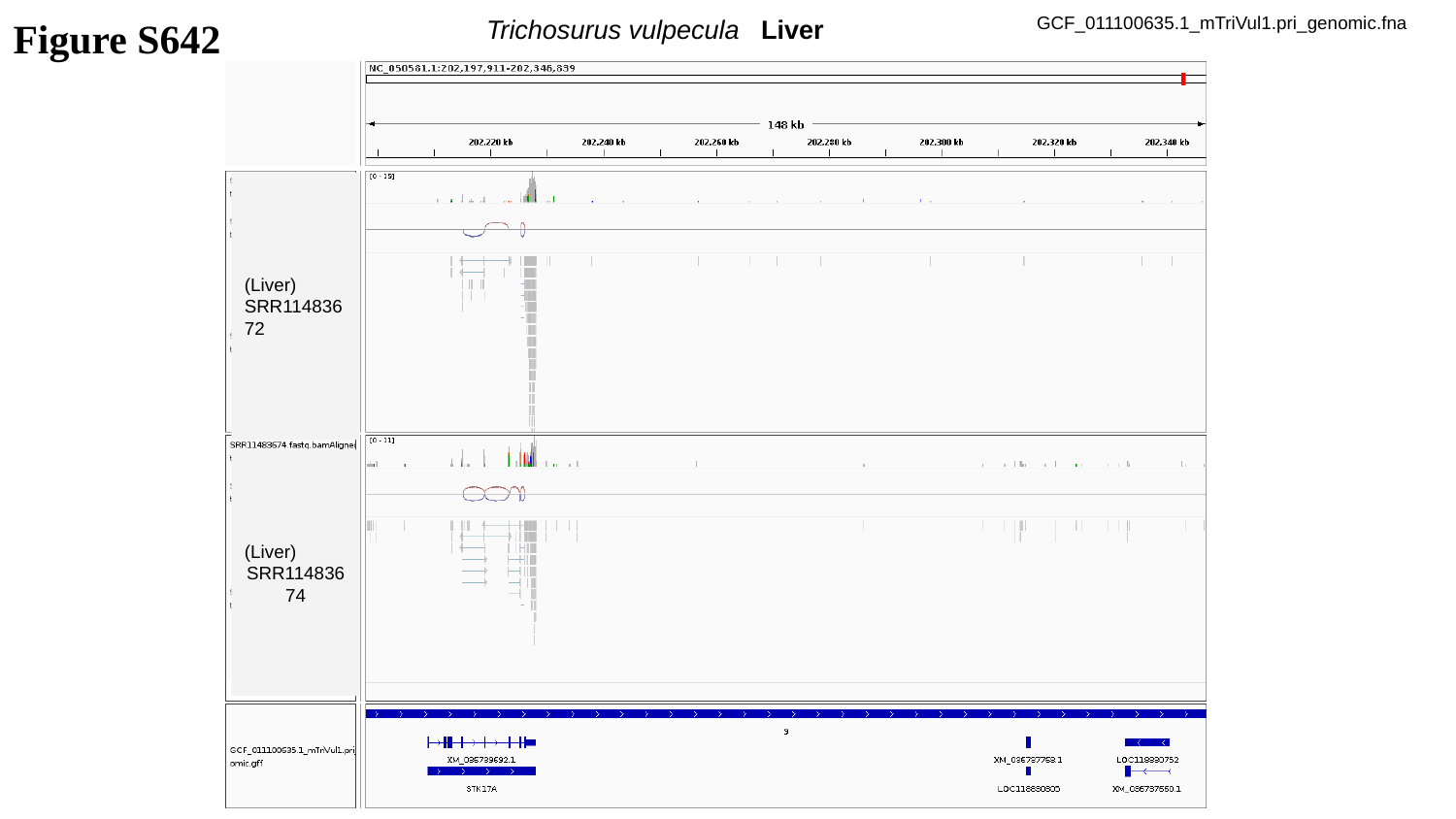

Figure S642
Trichosurus vulpecula Liver
GCF_011100635.1_mTriVul1.pri_genomic.fna
(Liver)
SRR11483672
(Liver)
SRR11483674

### Slide 643
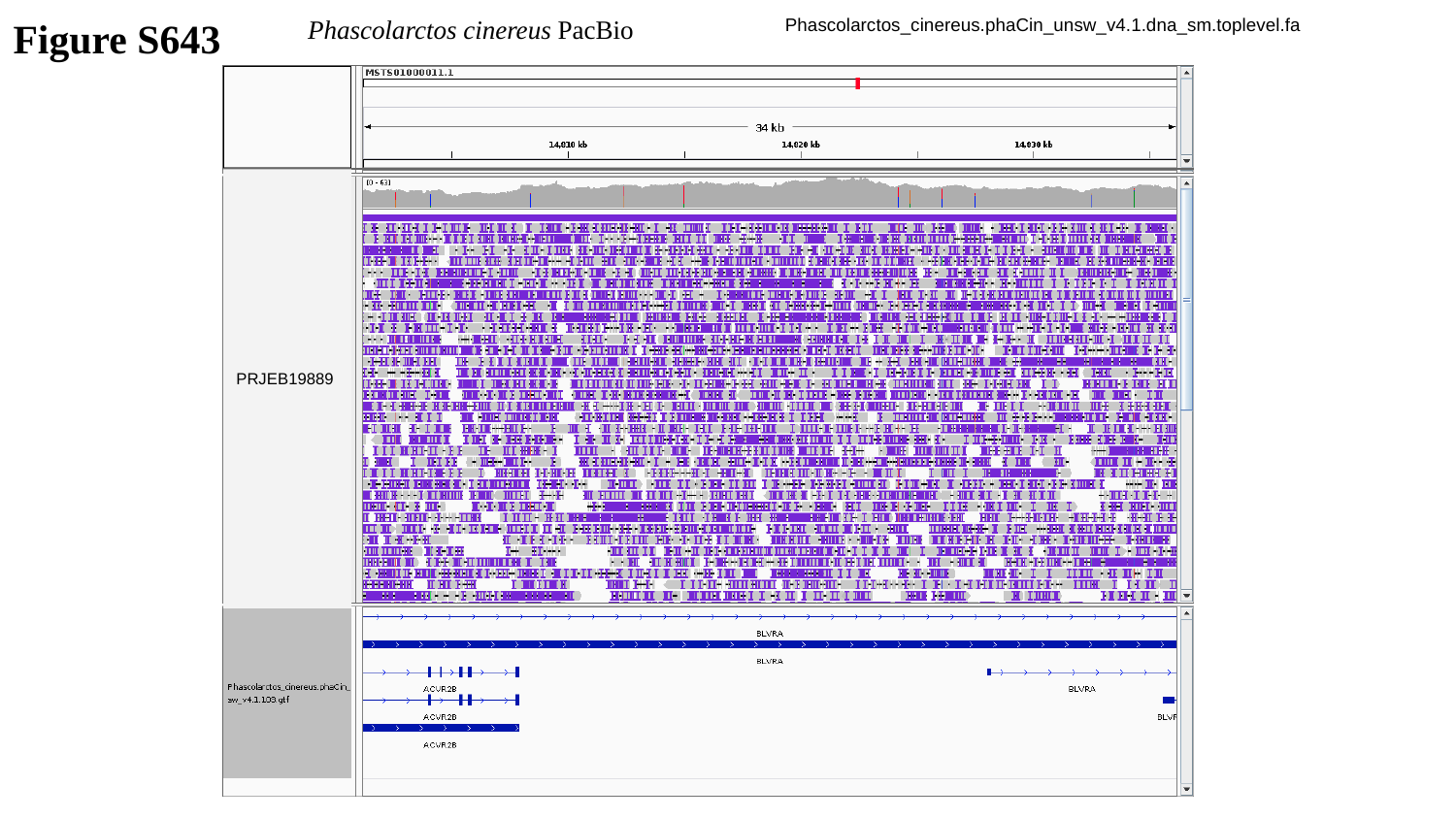

Figure S643
Phascolarctos cinereus PacBio
Phascolarctos_cinereus.phaCin_unsw_v4.1.dna_sm.toplevel.fa
PRJEB19889

### Slide 644
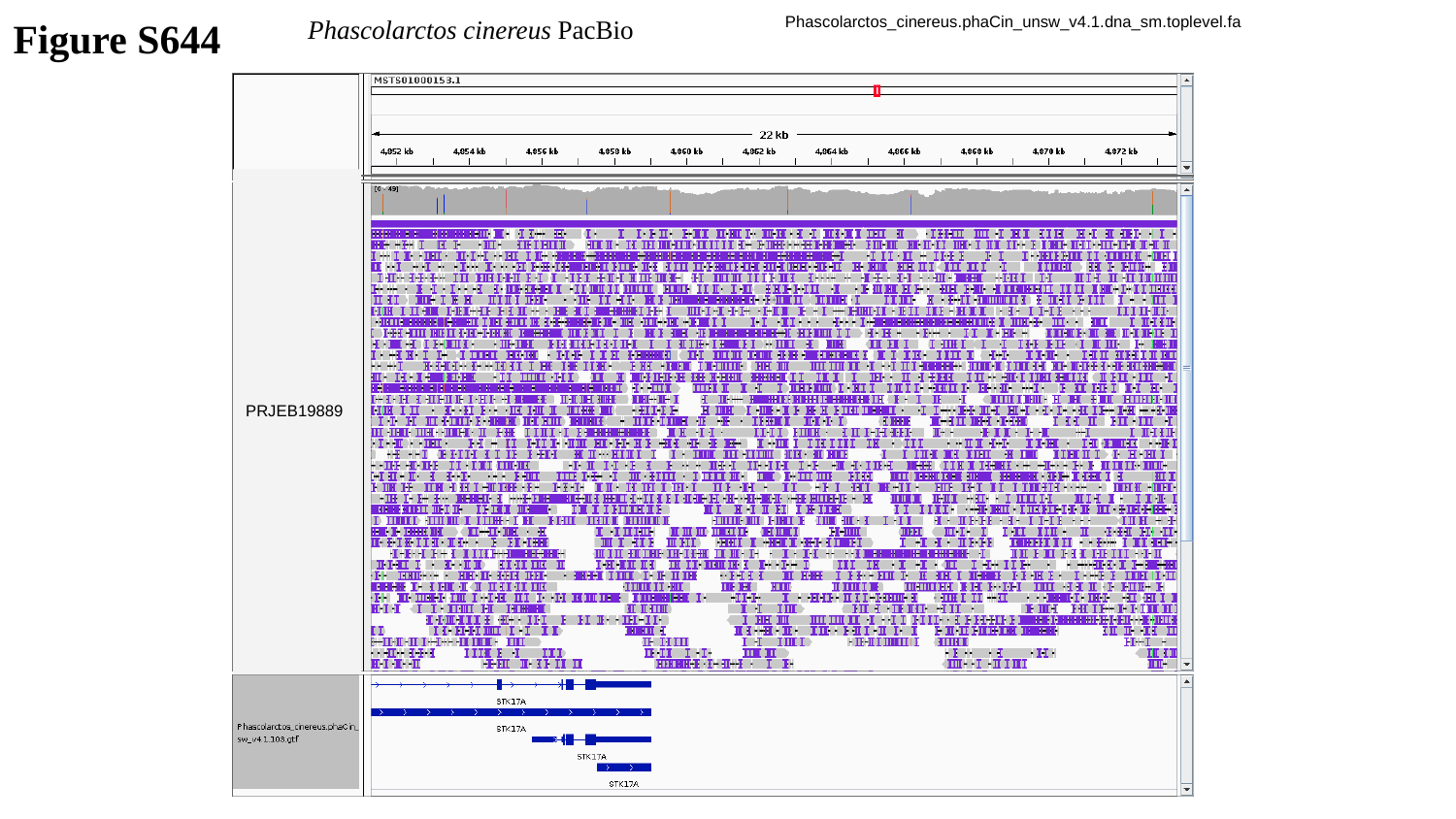

Figure S644
Phascolarctos cinereus PacBio
Phascolarctos_cinereus.phaCin_unsw_v4.1.dna_sm.toplevel.fa
PRJEB19889

### Slide 645
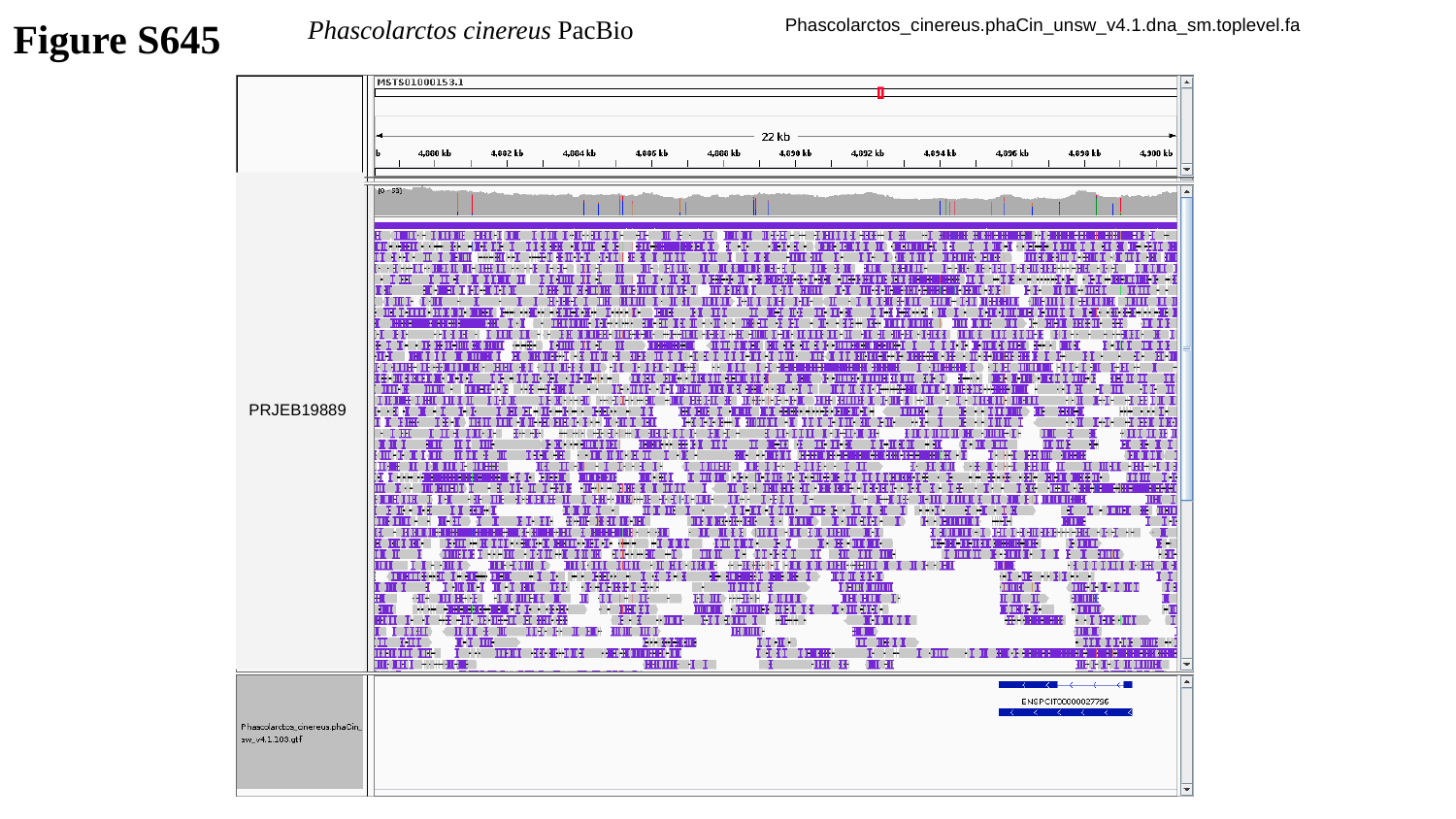

Figure S645
Phascolarctos cinereus PacBio
Phascolarctos_cinereus.phaCin_unsw_v4.1.dna_sm.toplevel.fa
PRJEB19889

### Slide 646
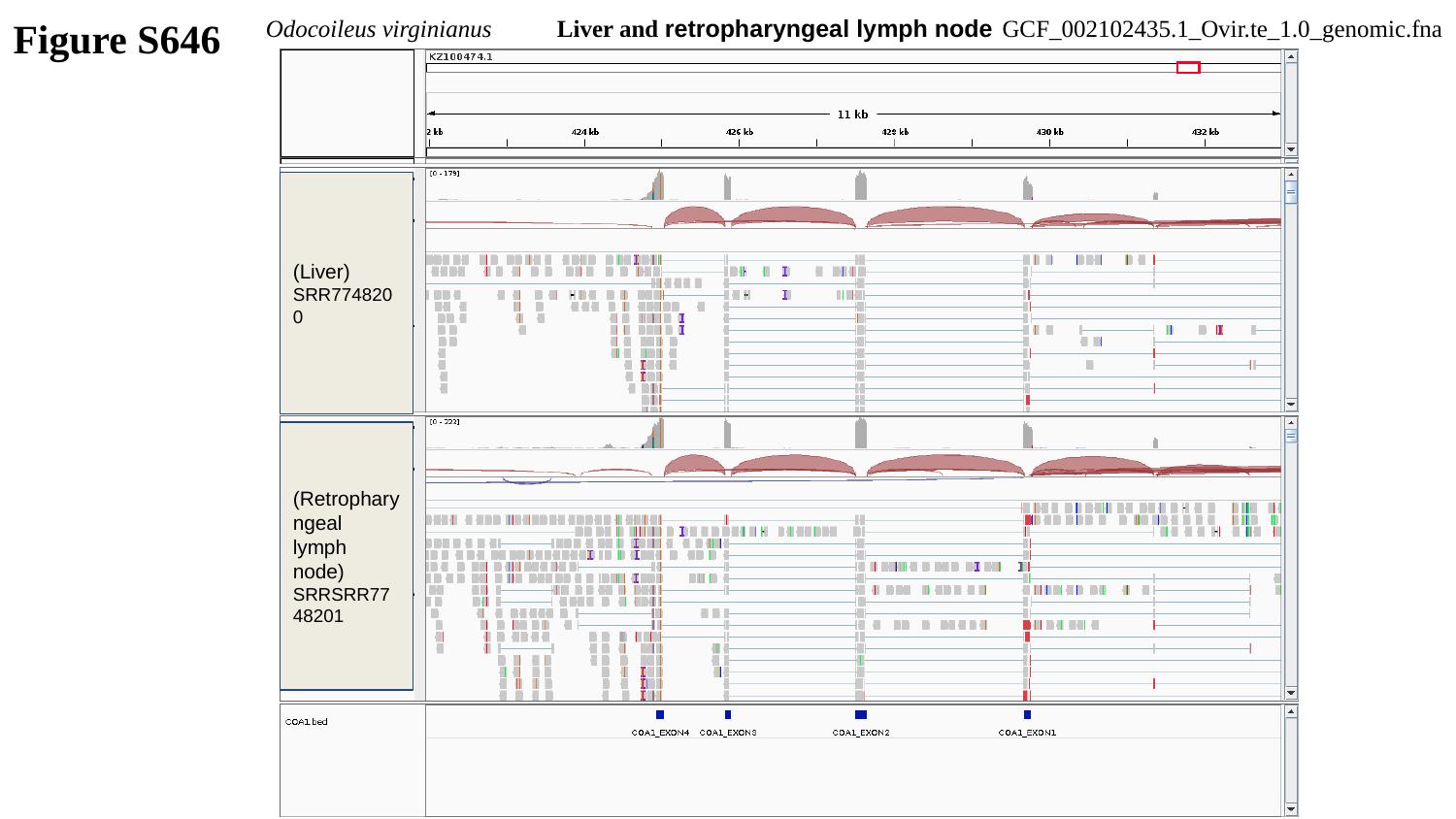

Figure S646
Odocoileus virginianus	Liver and retropharyngeal lymph node
GCF_002102435.1_Ovir.te_1.0_genomic.fna
(Liver)
SRR7748200
(Retropharyngeal lymph node)
SRRSRR7748201

### Slide 647
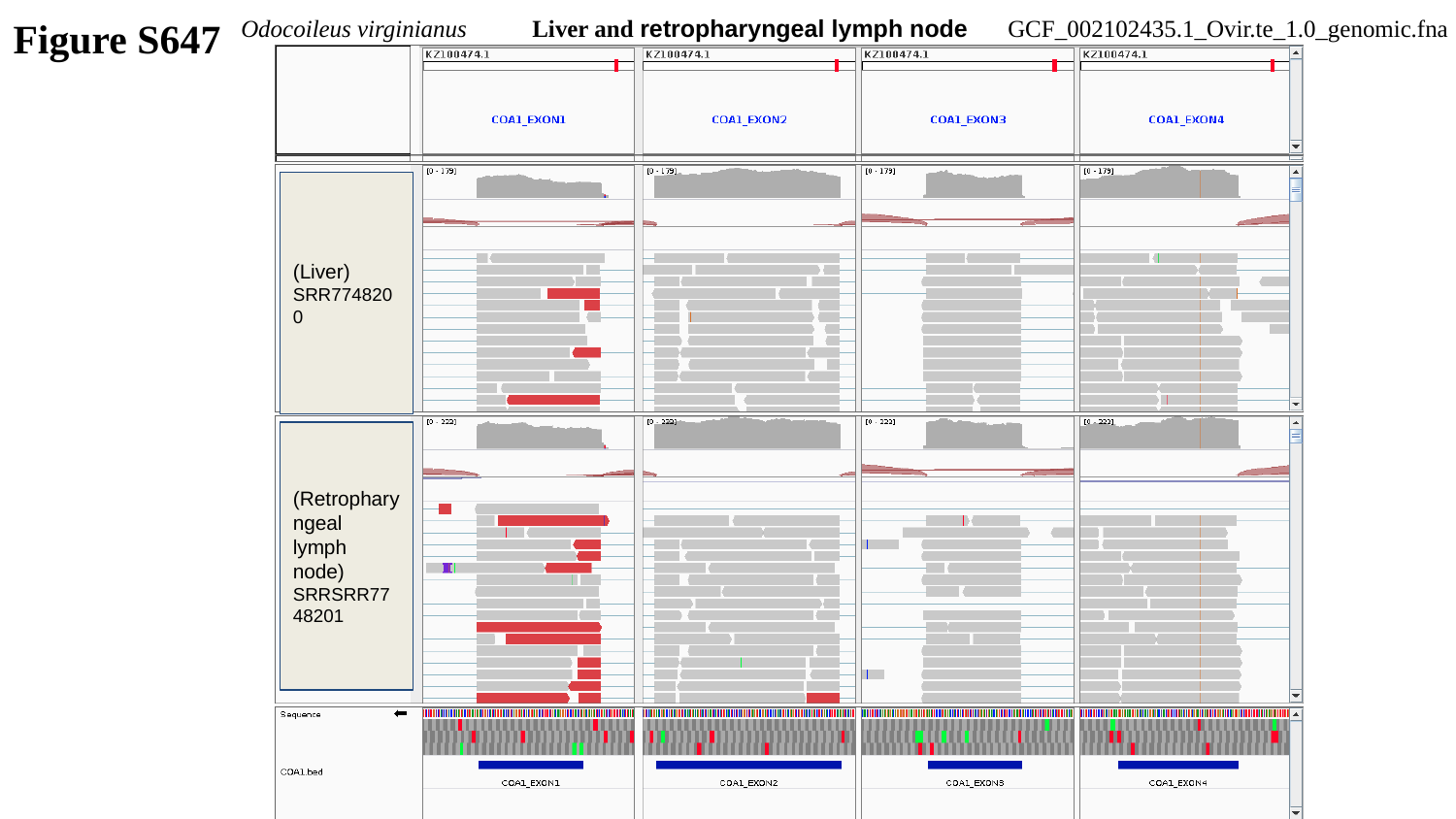

Figure S647
Odocoileus virginianus	Liver and retropharyngeal lymph node
GCF_002102435.1_Ovir.te_1.0_genomic.fna
(Liver)
SRR7748200
(Retropharyngeal lymph node)
SRRSRR7748201

### Slide 648
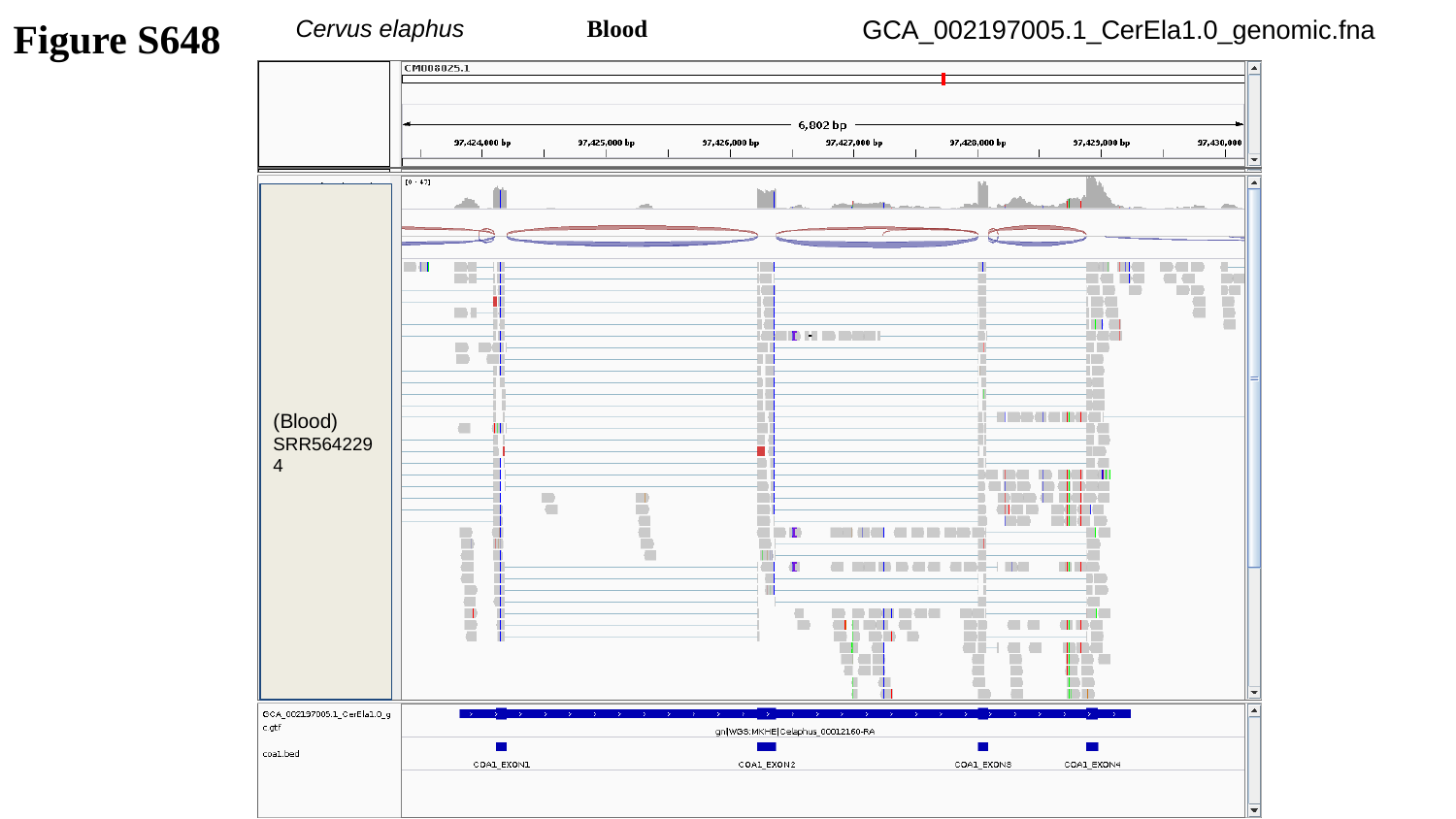

Figure S648
Cervus elaphus	Blood
GCA_002197005.1_CerEla1.0_genomic.fna
(Blood)
SRR5642294

### Slide 649
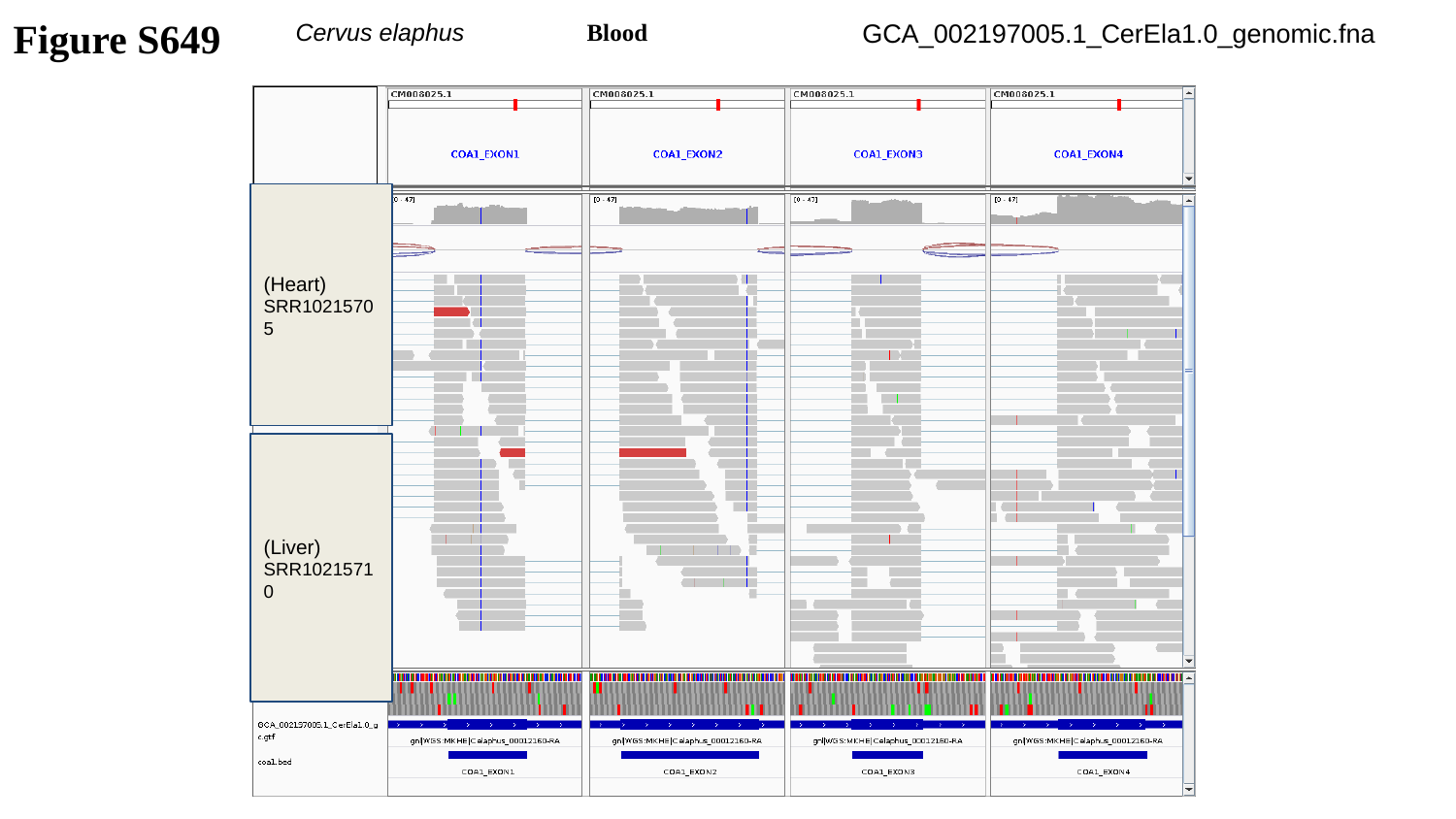

Figure S649
Cervus elaphus	Blood
GCA_002197005.1_CerEla1.0_genomic.fna
(Heart)
SRR10215705
(Liver)
SRR10215710

### Slide 650
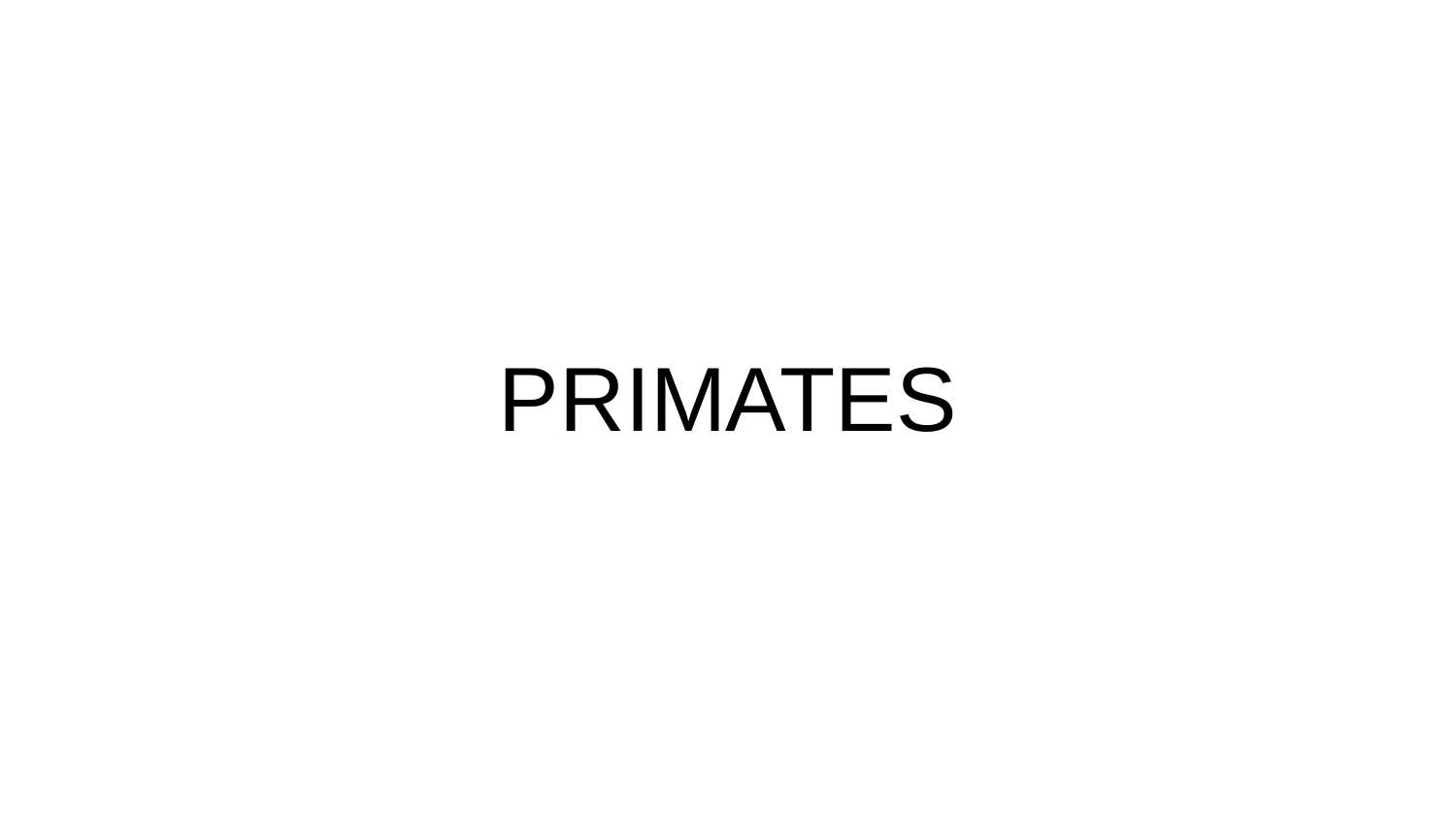

PRIMATES

### Slide 651
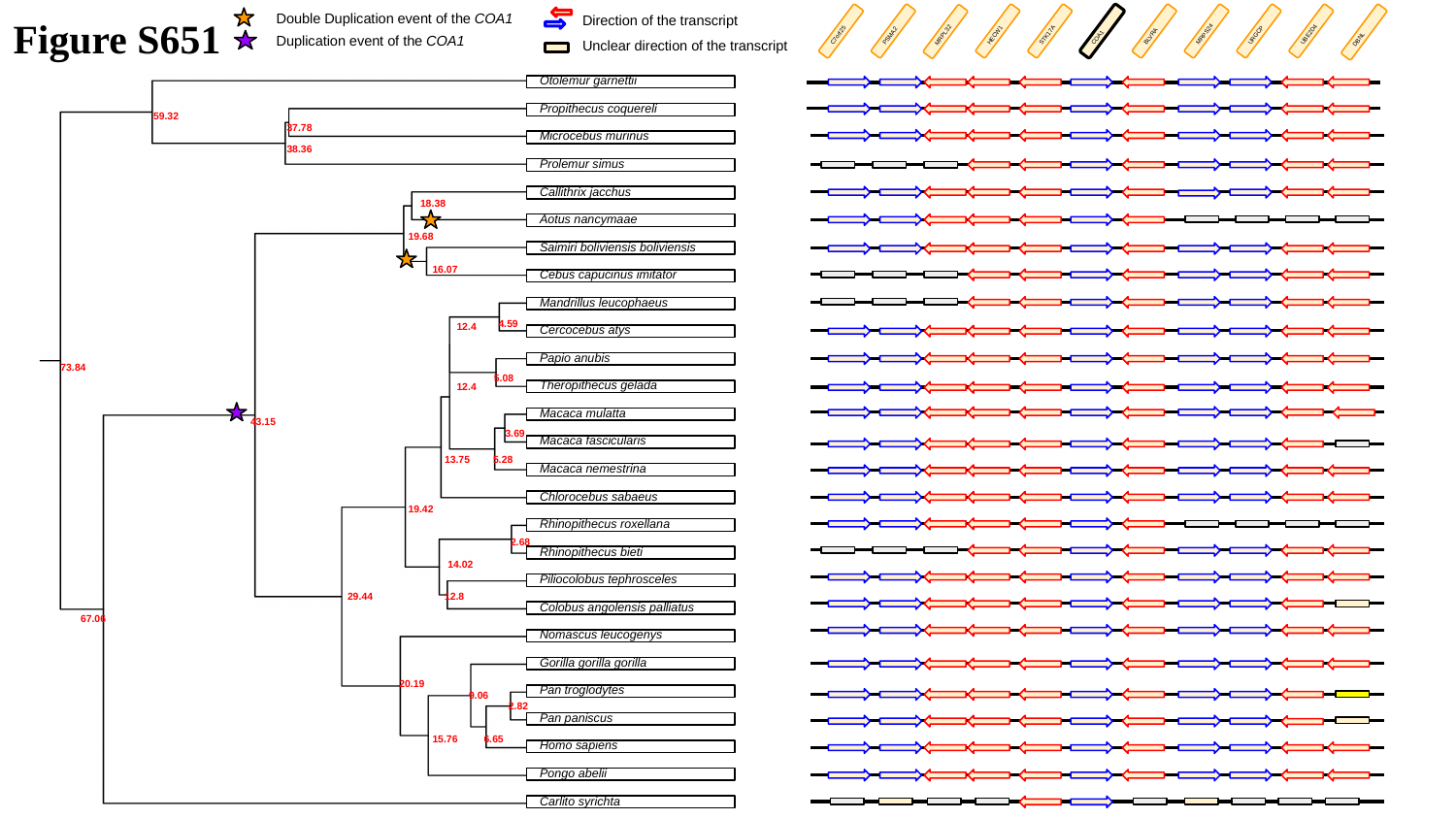

Figure S651
Direction of the transcript
Double Duplication event of the COA1
URGCP
UBE2D4
PSMA2
HECW1
STK17A
BLVRA
MRPS24
C7orf25
COA1
MRPL32
DBNL
Duplication event of the COA1
Unclear direction of the transcript
Otolemur garnettii
Propithecus coquereli
59.32
37.78
Microcebus murinus
38.36
Prolemur simus
Callithrix jacchus
18.38
Aotus nancymaae
19.68
Saimiri boliviensis boliviensis
16.07
Cebus capucinus imitator
Mandrillus leucophaeus
4.59
12.4
Cercocebus atys
Papio anubis
73.84
5.08
Theropithecus gelada
12.4
Macaca mulatta
43.15
3.69
Macaca fascicularis
13.75
5.28
Macaca nemestrina
Chlorocebus sabaeus
19.42
Rhinopithecus roxellana
2.68
Rhinopithecus bieti
14.02
Piliocolobus tephrosceles
29.44
12.8
Colobus angolensis palliatus
67.06
Nomascus leucogenys
Gorilla gorilla gorilla
20.19
Pan troglodytes
9.06
2.82
Pan paniscus
15.76
6.65
Homo sapiens
Pongo abelii
Carlito syrichta

### Slide 652
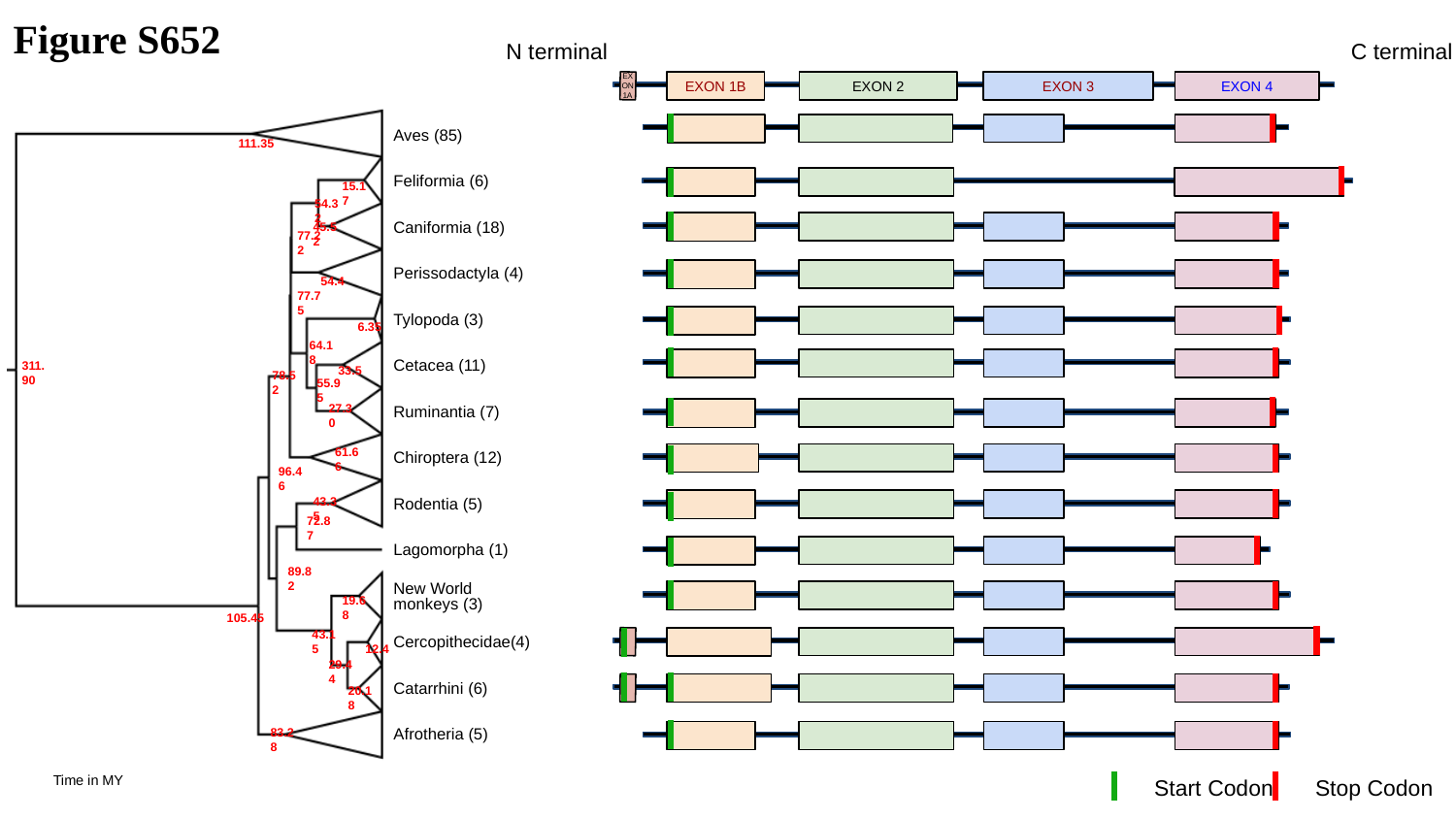

Figure S652
N terminal
C terminal
EXON 1A
EXON 1B
EXON 2
EXON 3
EXON 4
111.35
15.17
54.32
45.52
77.22
54.4
77.75
6.35
64.18
33.5
311.90
78.52
55.95
27.30
61.66
96.46
43.35
72.87
89.82
19.68
105.45
43.15
12.4
29.44
20.18
83.28
Aves (85)
Feliformia (6)
Caniformia (18)
Perissodactyla (4)
Tylopoda (3)
Cetacea (11)
Ruminantia (7)
Chiroptera (12)
Rodentia (5)
Lagomorpha (1)
New World monkeys (3)
Cercopithecidae(4)
Catarrhini (6)
Afrotheria (5)
Time in MY
Start Codon
Stop Codon

### Slide 653
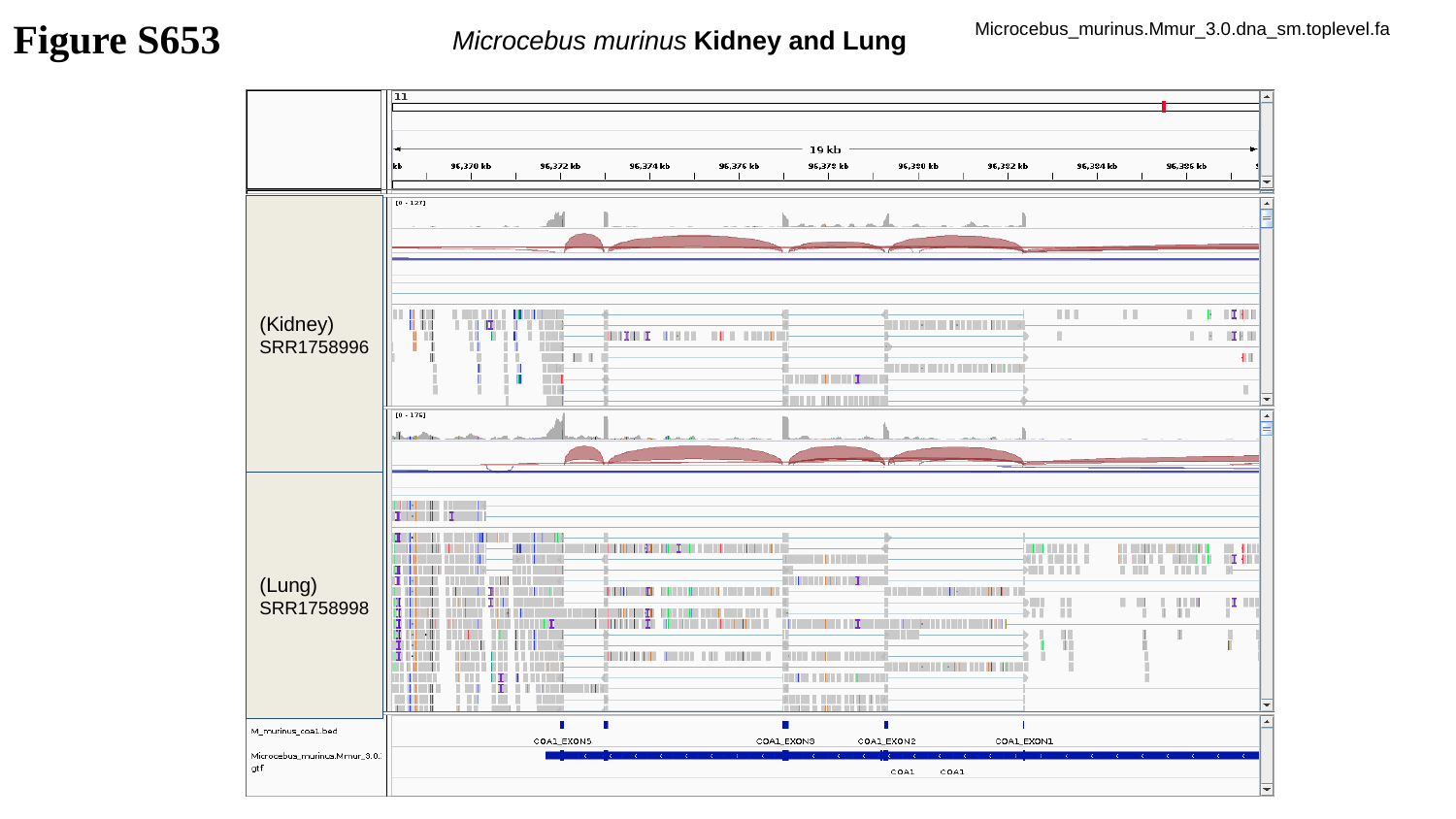

Figure S653
Microcebus_murinus.Mmur_3.0.dna_sm.toplevel.fa
Microcebus murinus Kidney and Lung
(Kidney)
SRR1758996
(Lung)
SRR1758998

### Slide 654
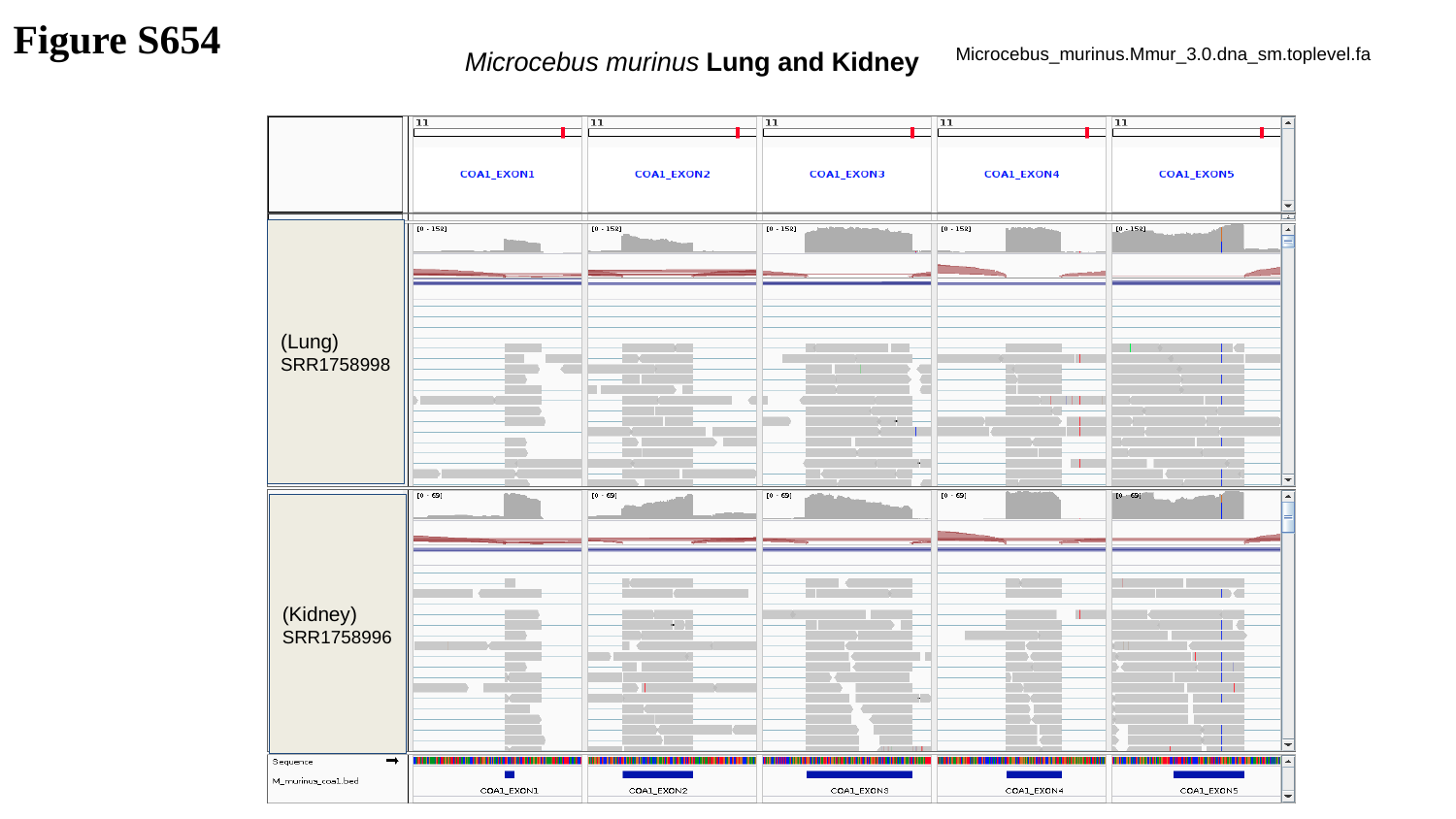

Figure S654
Microcebus murinus Lung and Kidney
Microcebus_murinus.Mmur_3.0.dna_sm.toplevel.fa
(Lung)
SRR1758998
(Kidney)
SRR1758996

### Slide 655
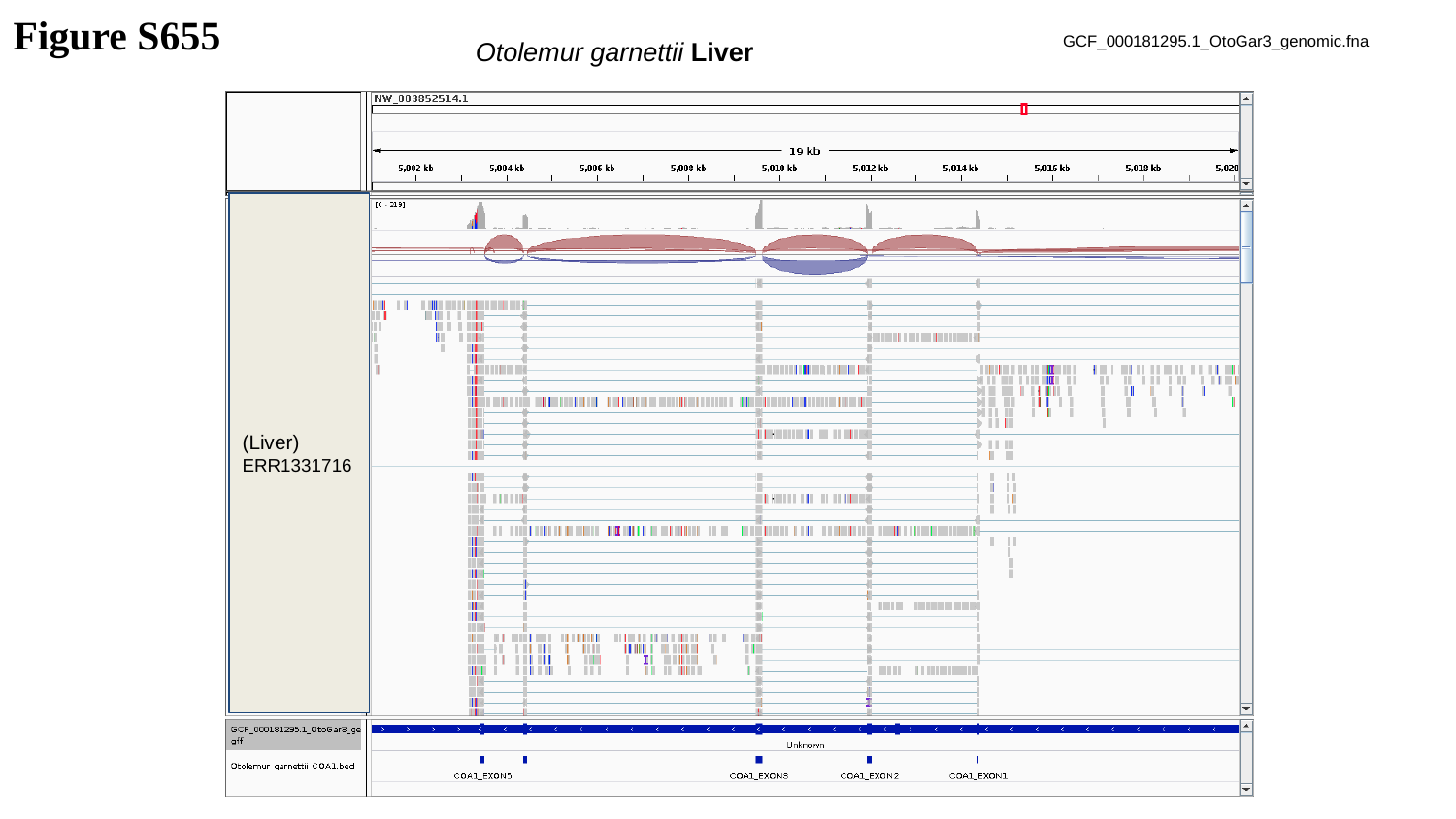

Figure S655
GCF_000181295.1_OtoGar3_genomic.fna
Otolemur garnettii Liver
(Liver)
ERR1331716

### Slide 656
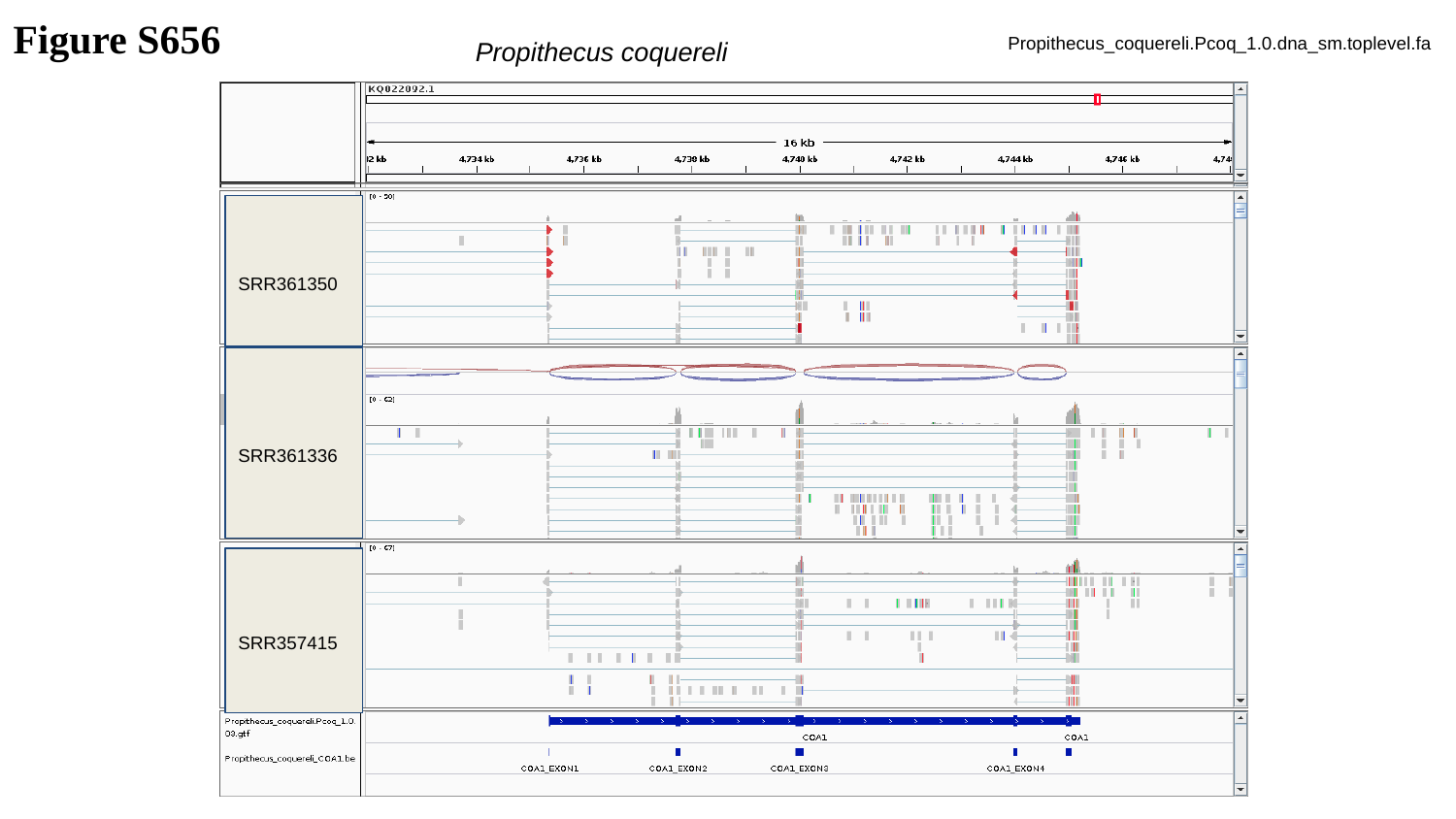

Figure S656
Propithecus_coquereli.Pcoq_1.0.dna_sm.toplevel.fa
Propithecus coquereli
SRR361350
SRR361336
SRR357415

### Slide 657
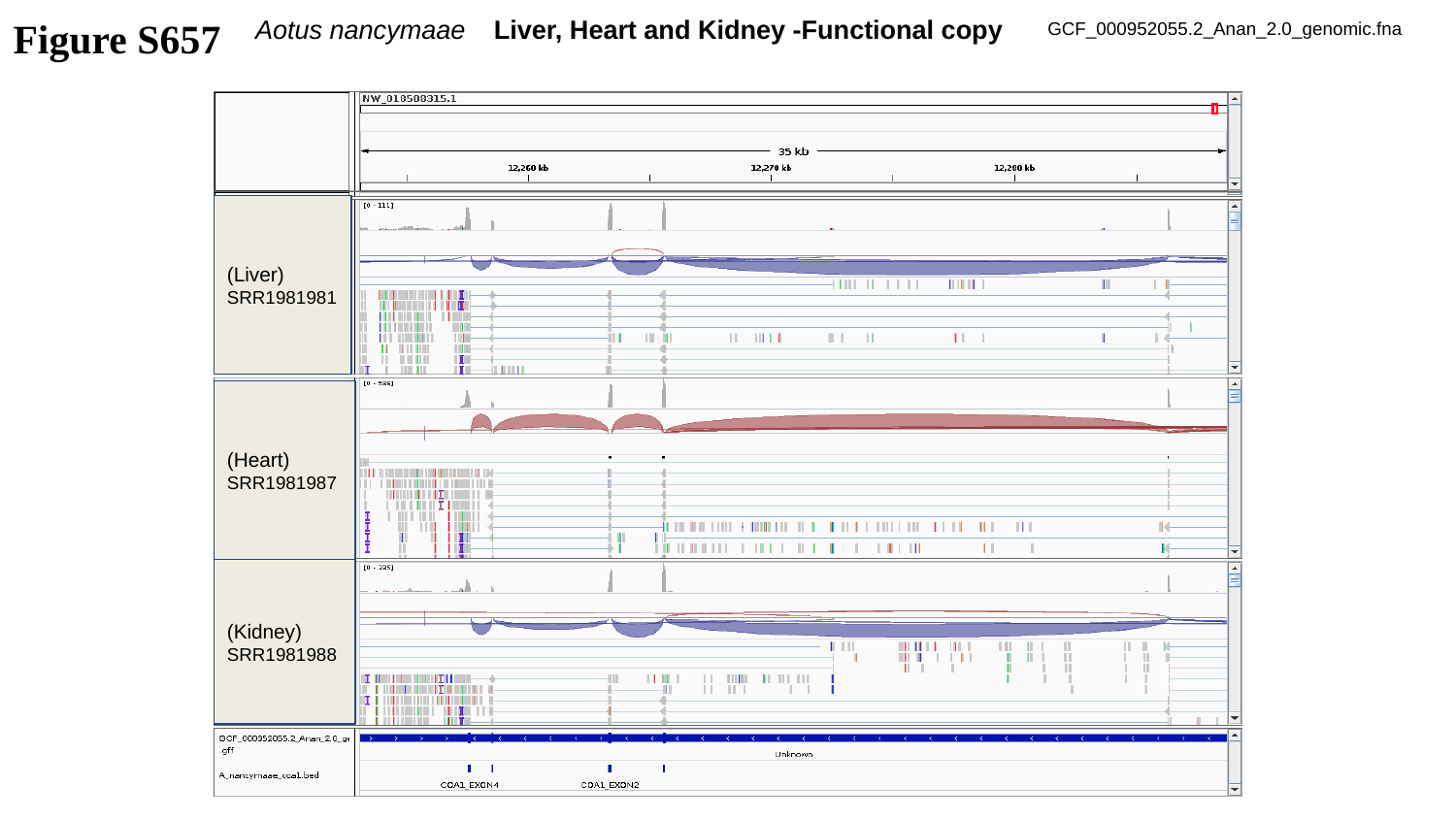

Figure S657
Aotus nancymaae Liver, Heart and Kidney -Functional copy
GCF_000952055.2_Anan_2.0_genomic.fna
(Liver)
SRR1981981
(Heart)
SRR1981987
(Kidney)
SRR1981988

### Slide 658
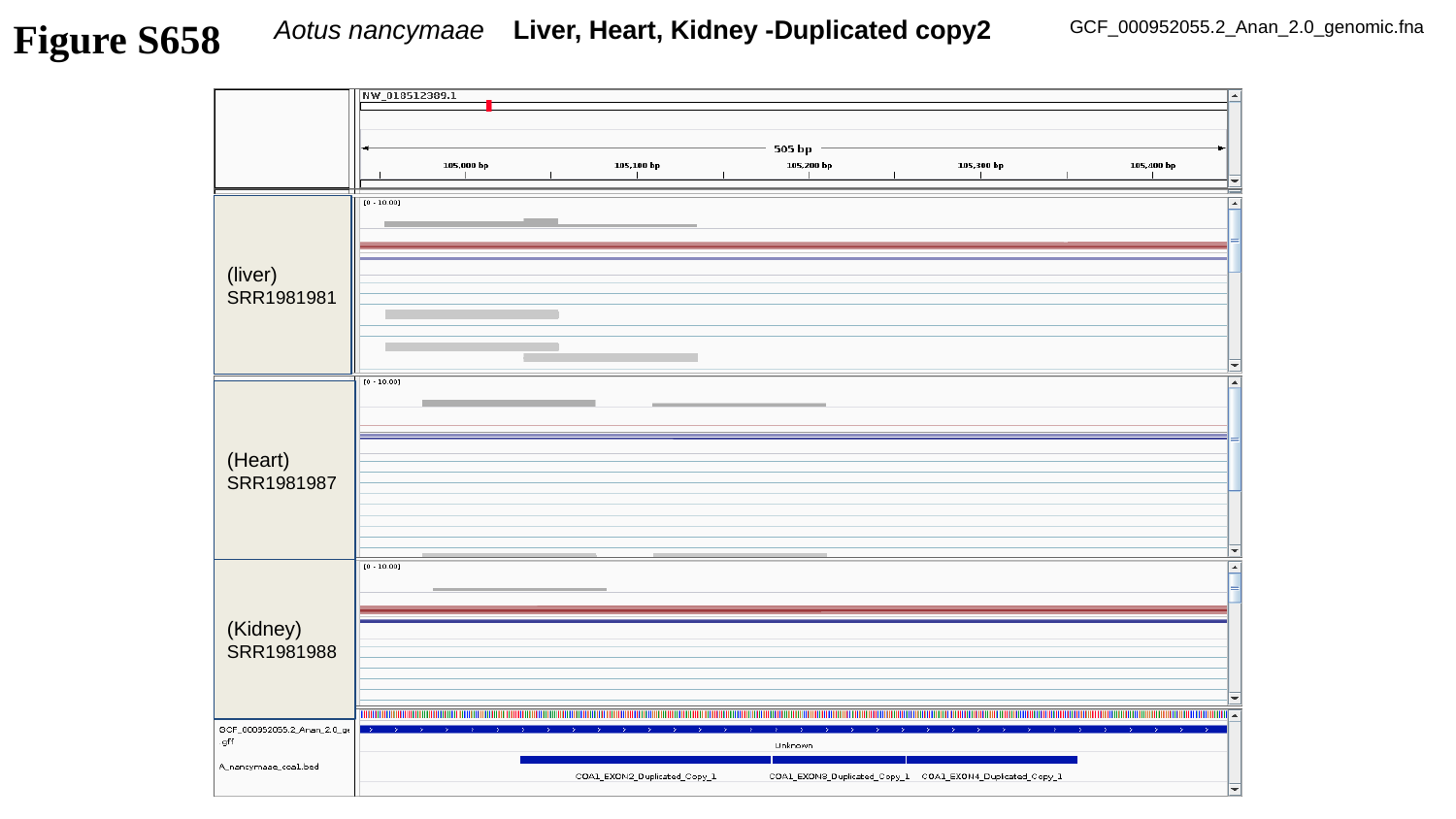

Figure S658
Aotus nancymaae Liver, Heart, Kidney -Duplicated copy2
GCF_000952055.2_Anan_2.0_genomic.fna
(liver)
SRR1981981
(Heart)
SRR1981987
(Kidney)
SRR1981988

### Slide 659
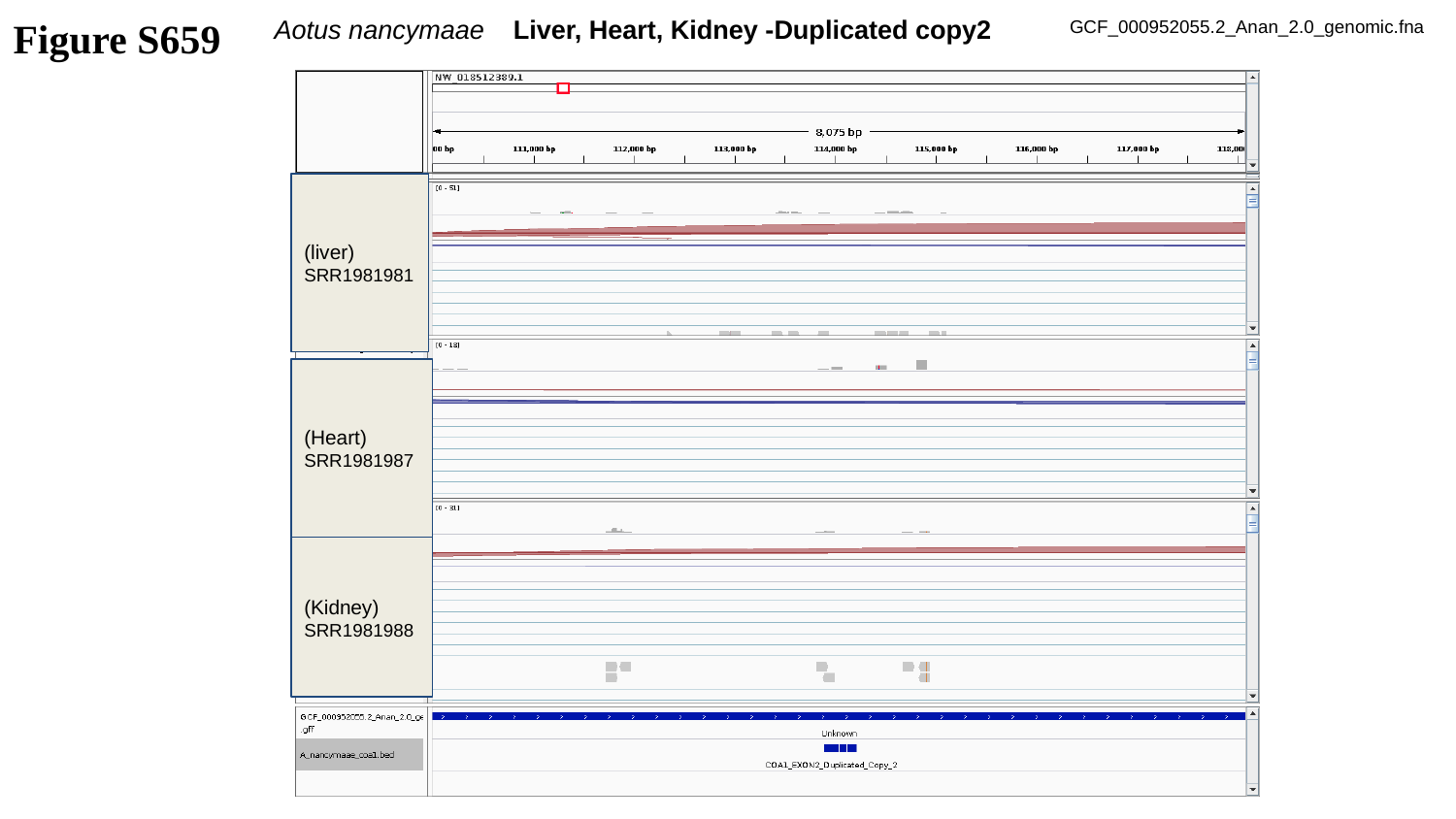

Figure S659
Aotus nancymaae Liver, Heart, Kidney -Duplicated copy2
GCF_000952055.2_Anan_2.0_genomic.fna
(liver)
SRR1981981
(Heart)
SRR1981987
(Kidney)
SRR1981988

### Slide 660
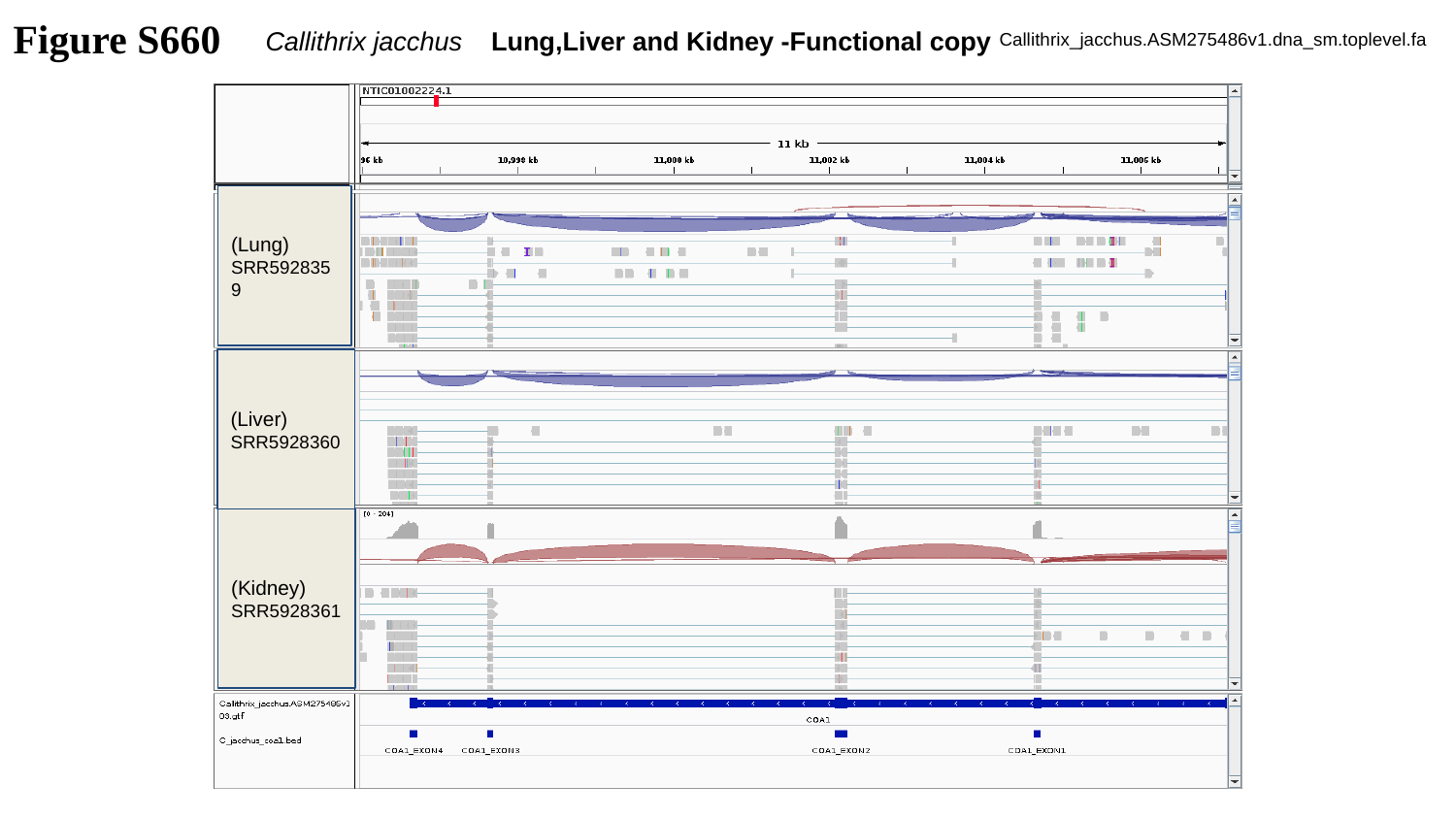

Figure S660
Callithrix jacchus Lung,Liver and Kidney -Functional copy
Callithrix_jacchus.ASM275486v1.dna_sm.toplevel.fa
(Lung)
SRR5928359
(Liver)
SRR5928360
(Kidney)
SRR5928361

### Slide 661
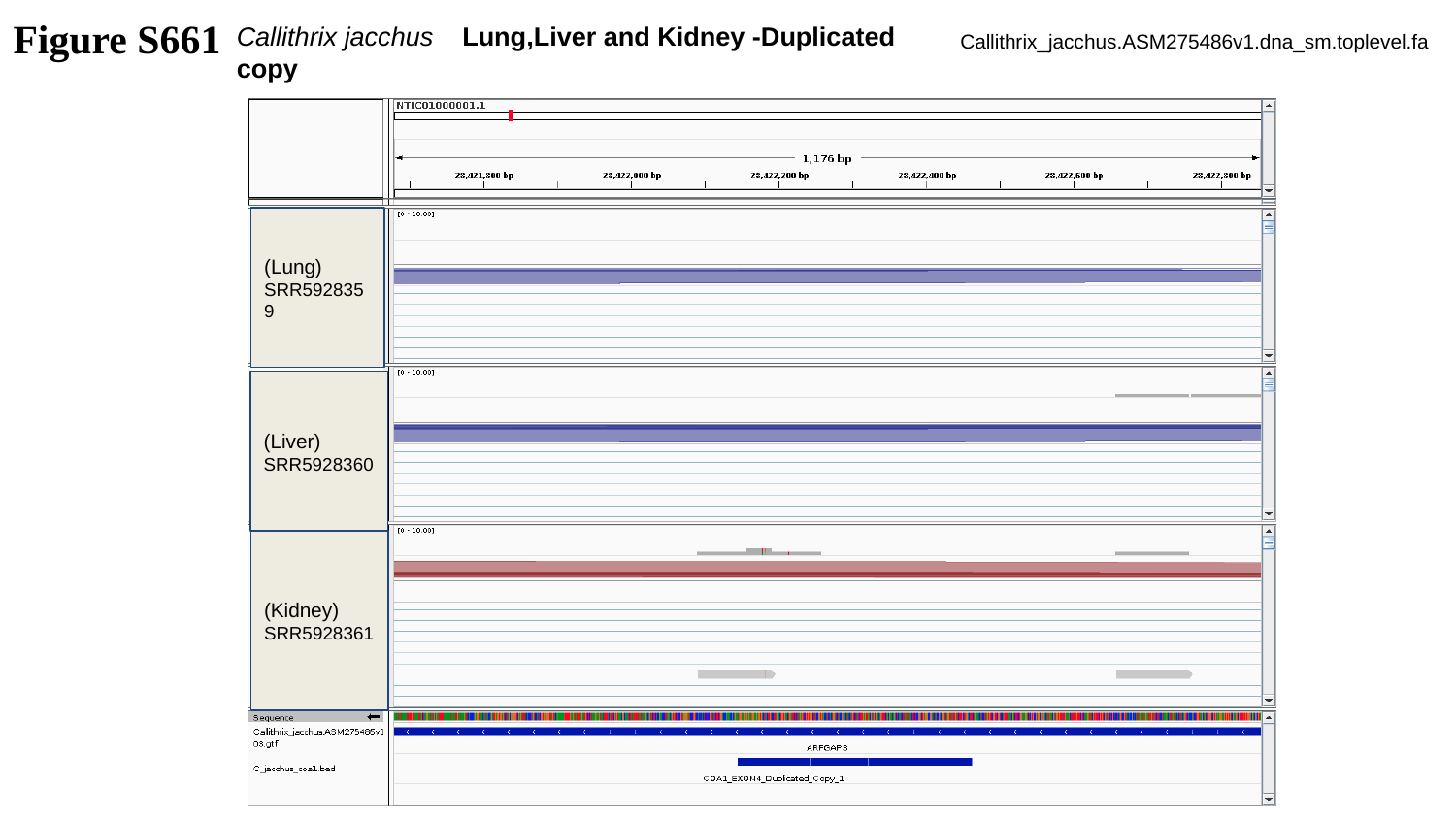

Figure S661
Callithrix jacchus Lung,Liver and Kidney -Duplicated copy
Callithrix_jacchus.ASM275486v1.dna_sm.toplevel.fa
(Lung)
SRR5928359
(Liver)
SRR5928360
(Kidney)
SRR5928361

### Slide 662
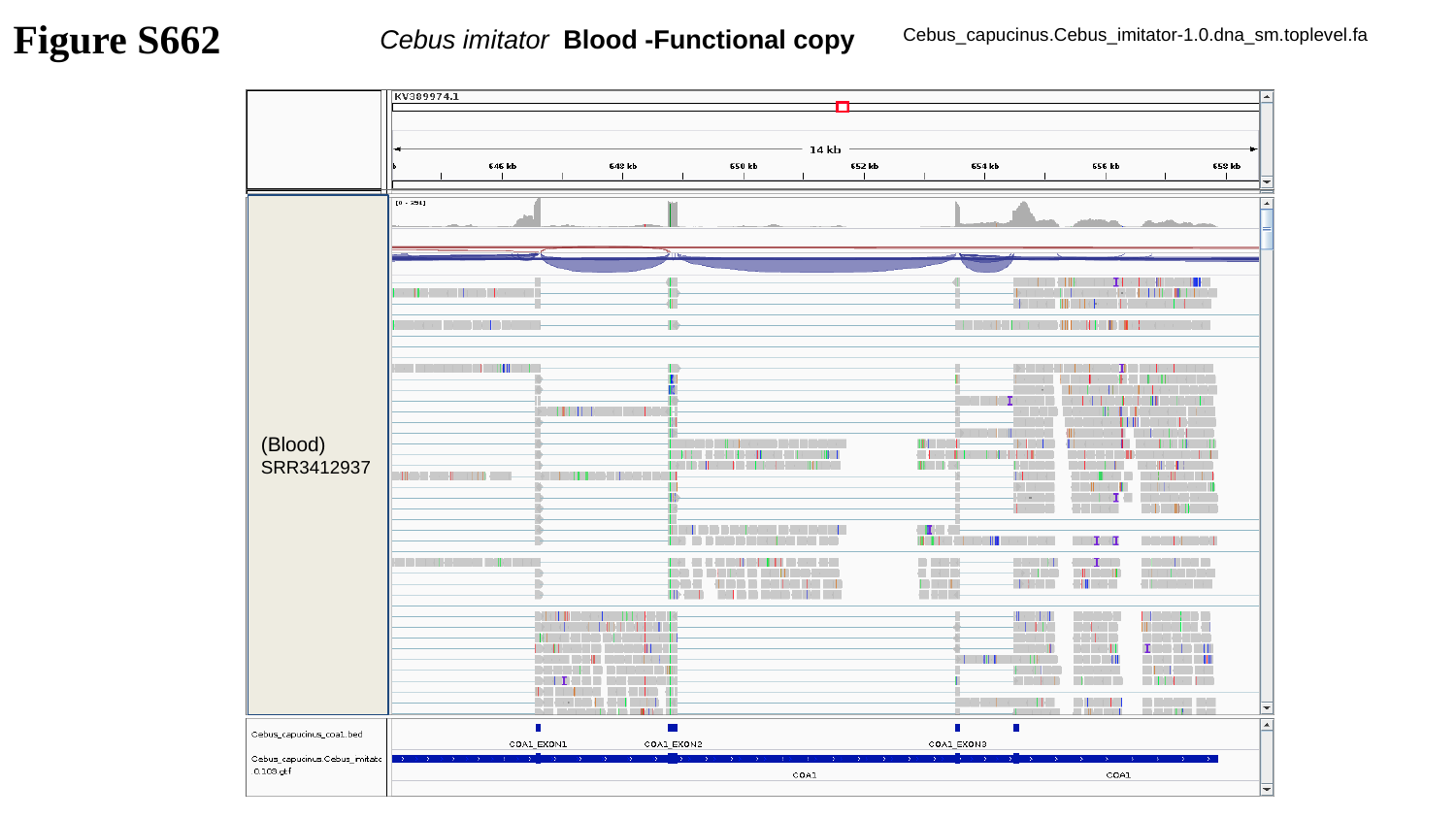

Figure S662
Cebus imitator Blood -Functional copy
Cebus_capucinus.Cebus_imitator-1.0.dna_sm.toplevel.fa
(Blood)
SRR3412937

### Slide 663
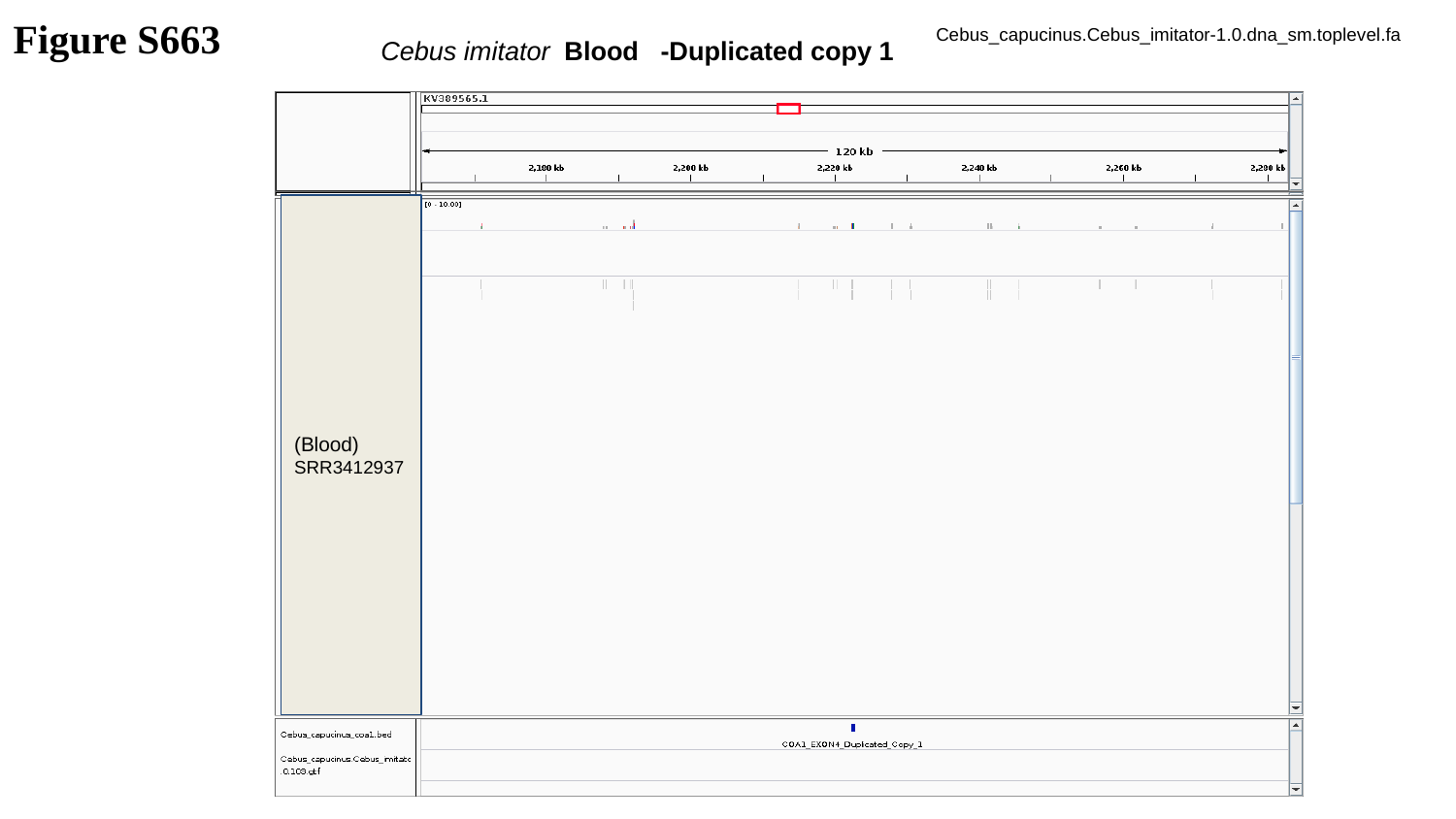

Figure S663
Cebus_capucinus.Cebus_imitator-1.0.dna_sm.toplevel.fa
Cebus imitator Blood -Duplicated copy 1
(Blood)
SRR3412937

### Slide 664
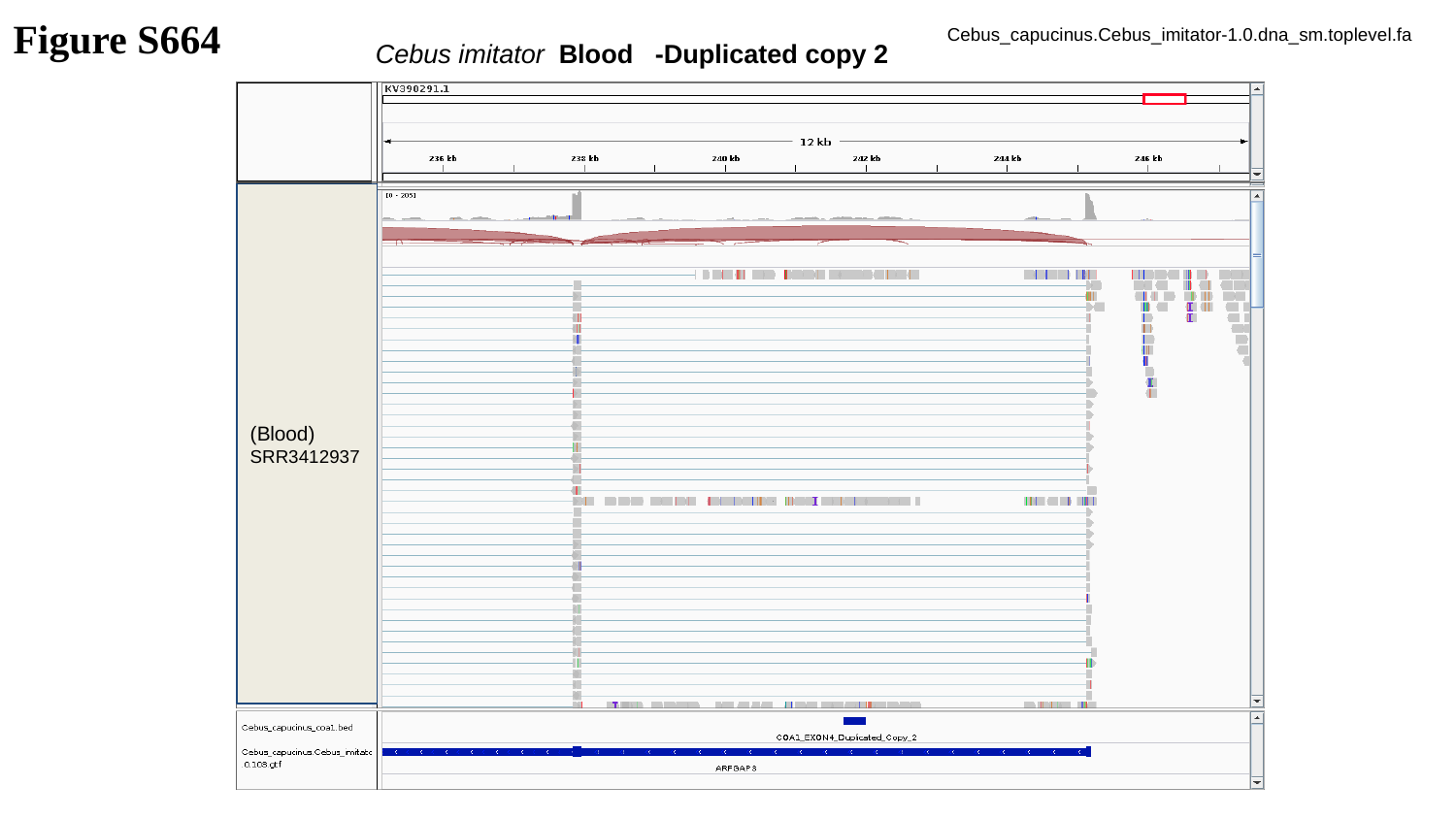

Figure S664
Cebus_capucinus.Cebus_imitator-1.0.dna_sm.toplevel.fa
Cebus imitator Blood -Duplicated copy 2
(Blood)
SRR3412937

### Slide 665
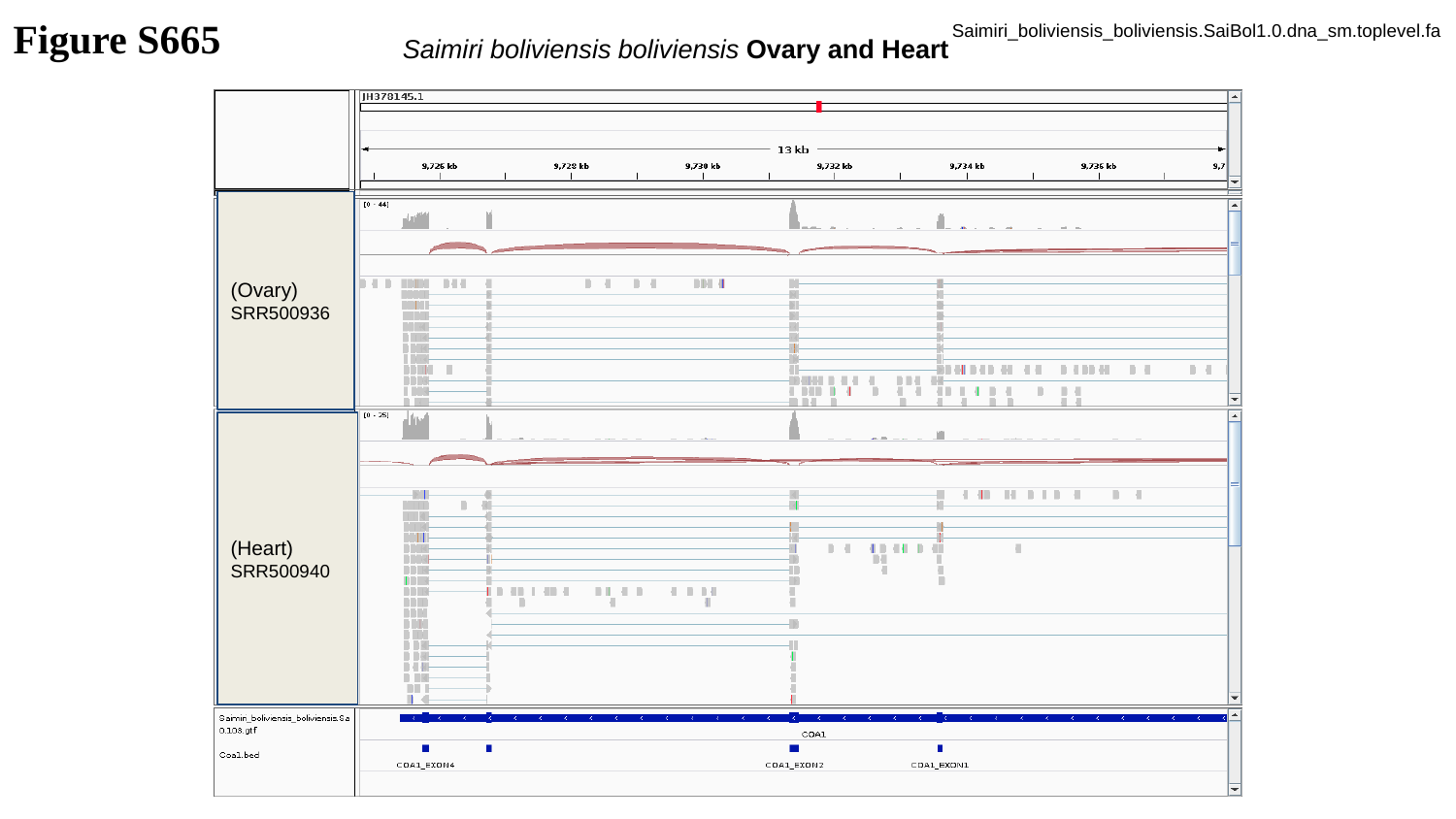

Figure S665
Saimiri_boliviensis_boliviensis.SaiBol1.0.dna_sm.toplevel.fa
Saimiri boliviensis boliviensis Ovary and Heart
(Ovary)
SRR500936
(Heart)
SRR500940

### Slide 666
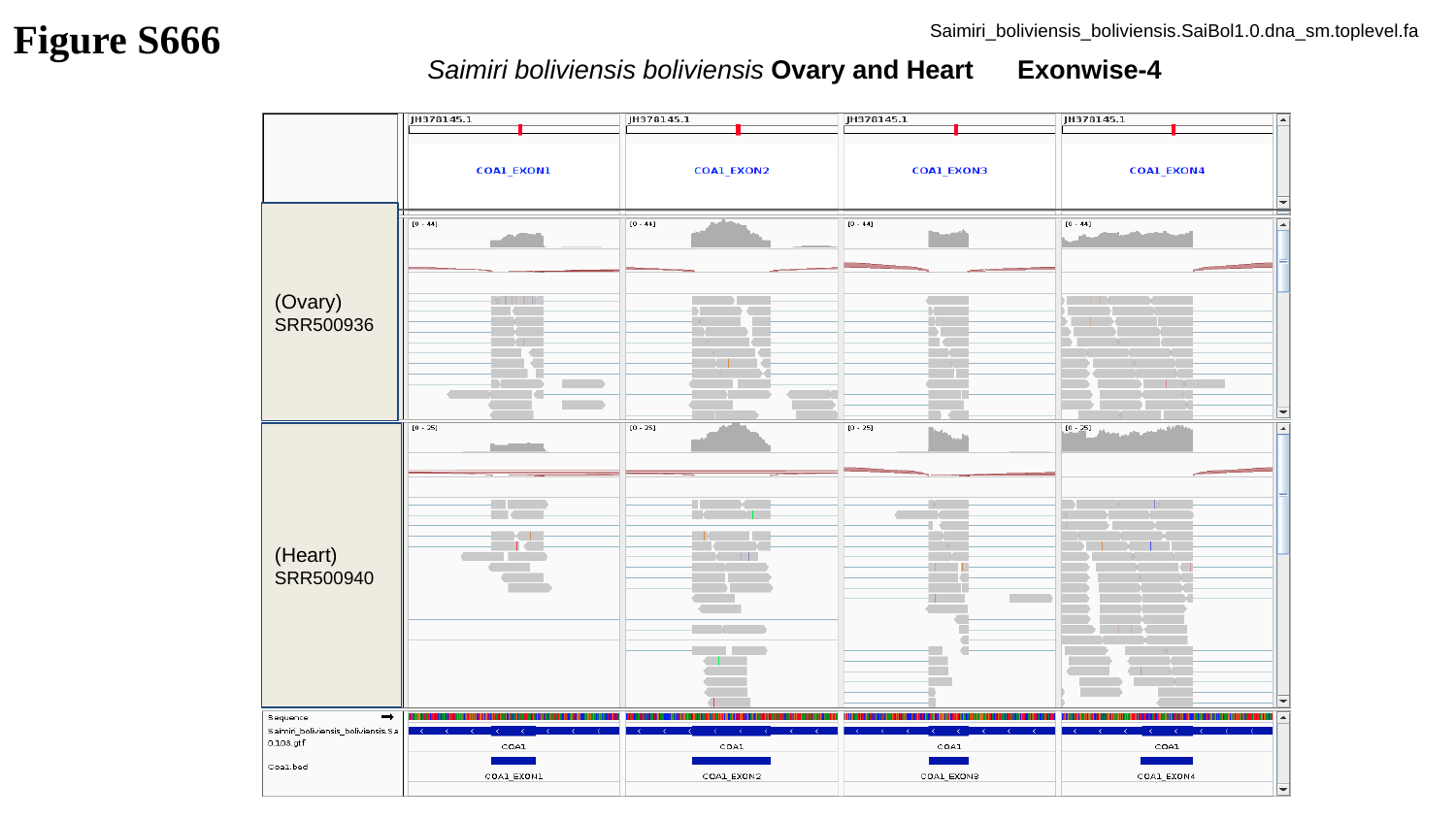

Figure S666
Saimiri_boliviensis_boliviensis.SaiBol1.0.dna_sm.toplevel.fa
Saimiri boliviensis boliviensis Ovary and Heart Exonwise-4
(Ovary)
SRR500936
(Heart)
SRR500940

### Slide 667
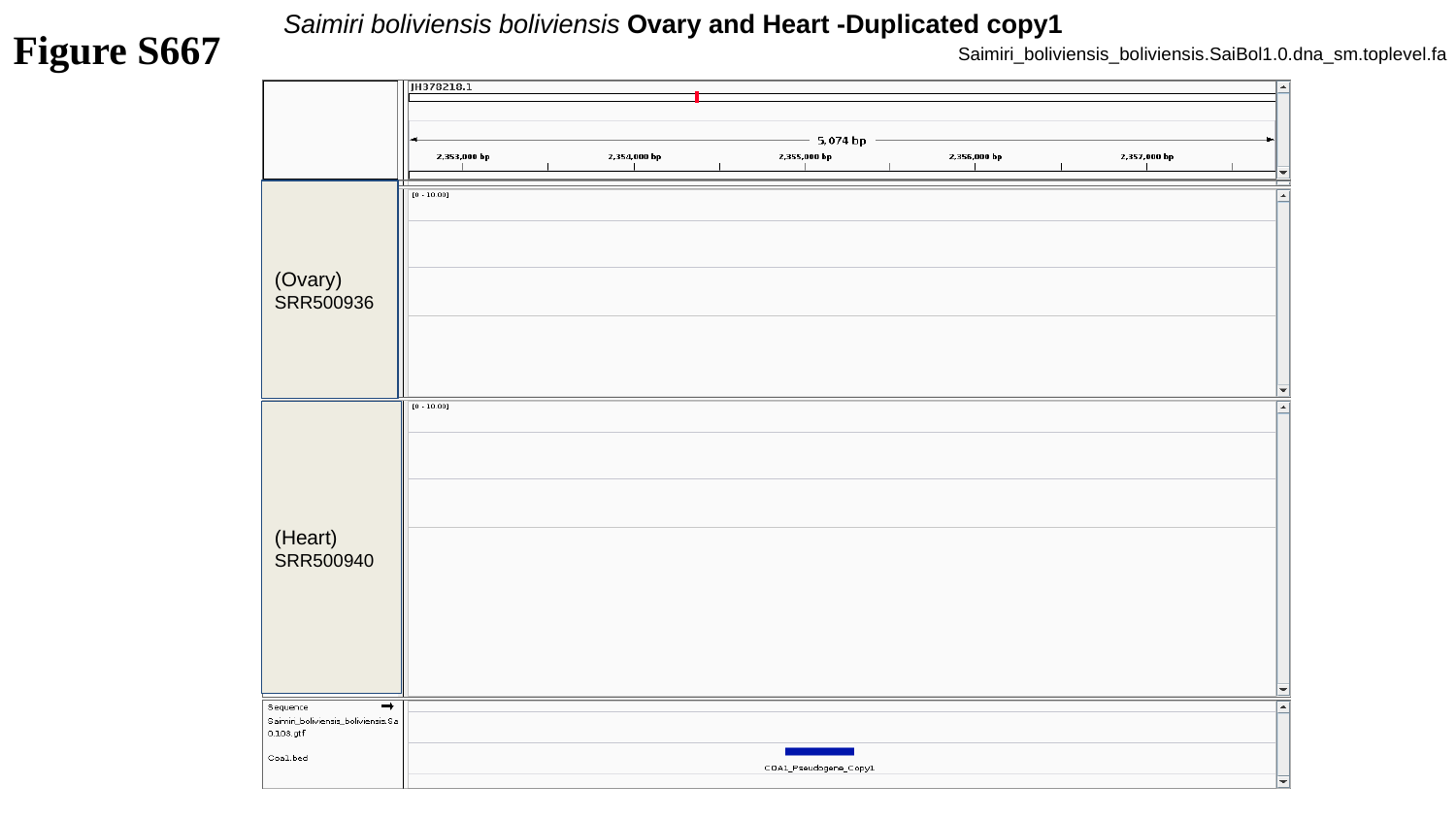

Saimiri boliviensis boliviensis Ovary and Heart -Duplicated copy1
Figure S667
Saimiri_boliviensis_boliviensis.SaiBol1.0.dna_sm.toplevel.fa
(Ovary)
SRR500936
(Heart)
SRR500940

### Slide 668
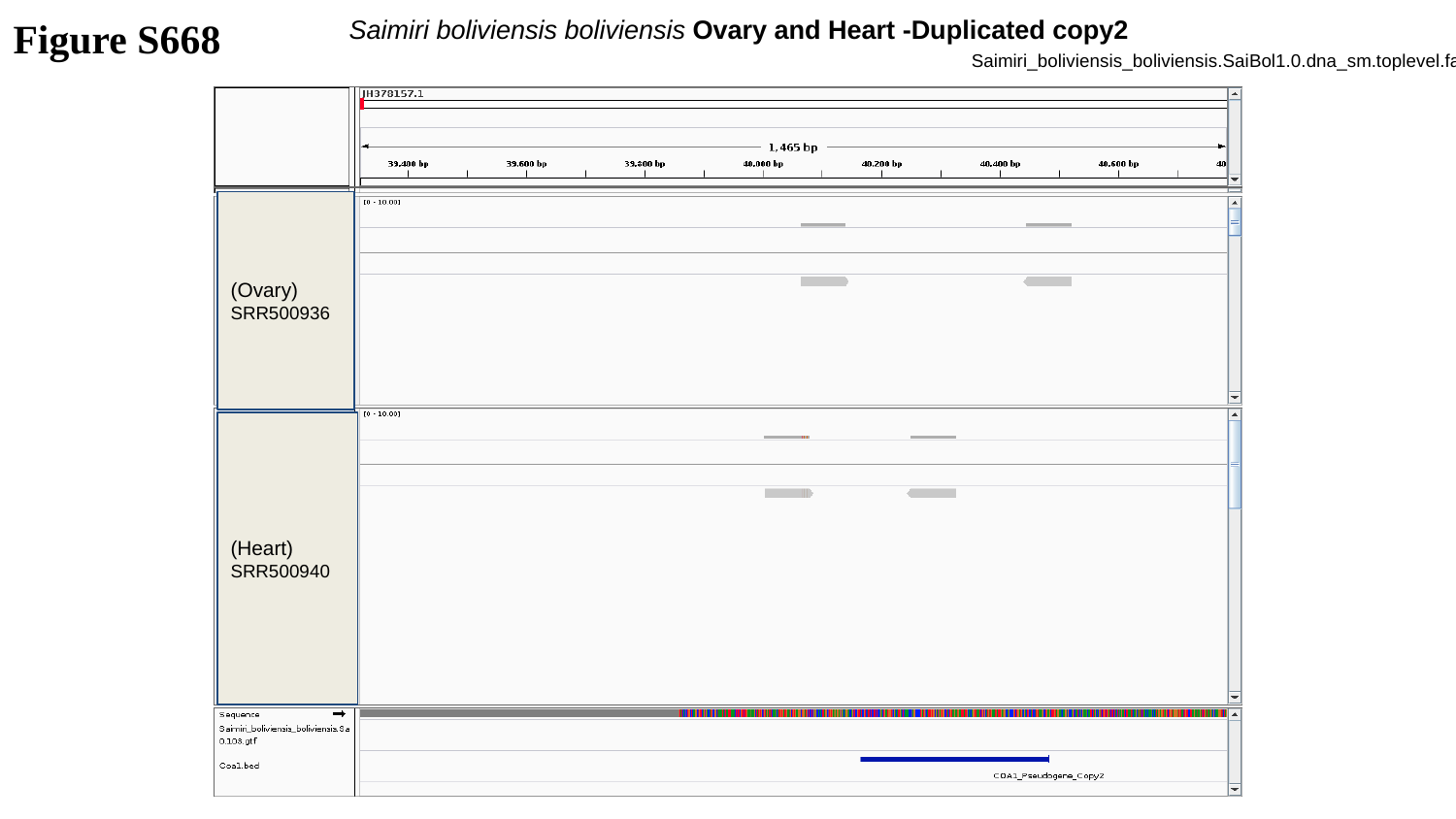

Figure S668
Saimiri boliviensis boliviensis Ovary and Heart -Duplicated copy2
Saimiri_boliviensis_boliviensis.SaiBol1.0.dna_sm.toplevel.fa
(Ovary)
SRR500936
(Heart)
SRR500940

### Slide 669
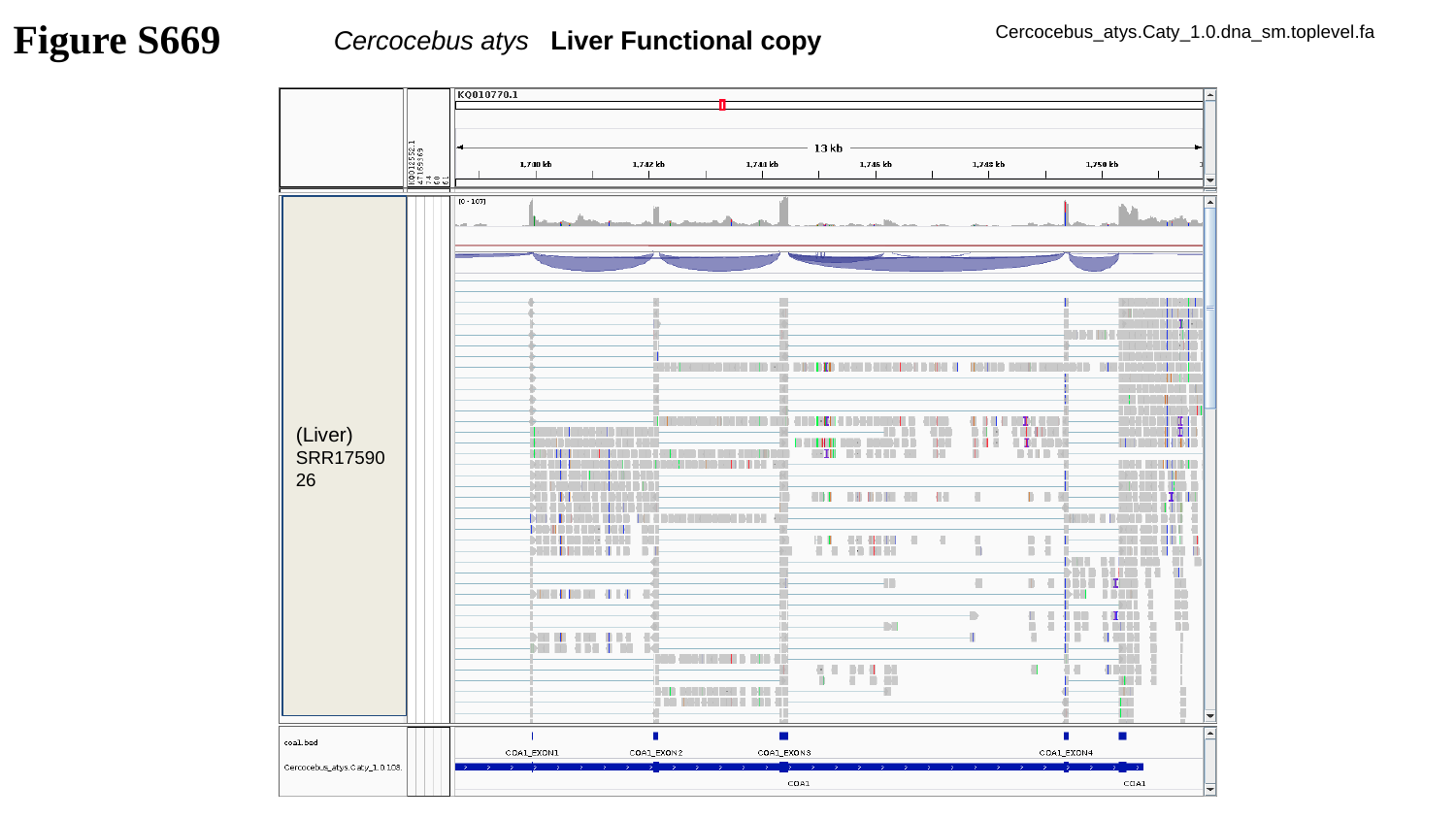

Figure S669
Cercocebus_atys.Caty_1.0.dna_sm.toplevel.fa
Cercocebus atys Liver Functional copy
(Liver)
SRR1759026

### Slide 670
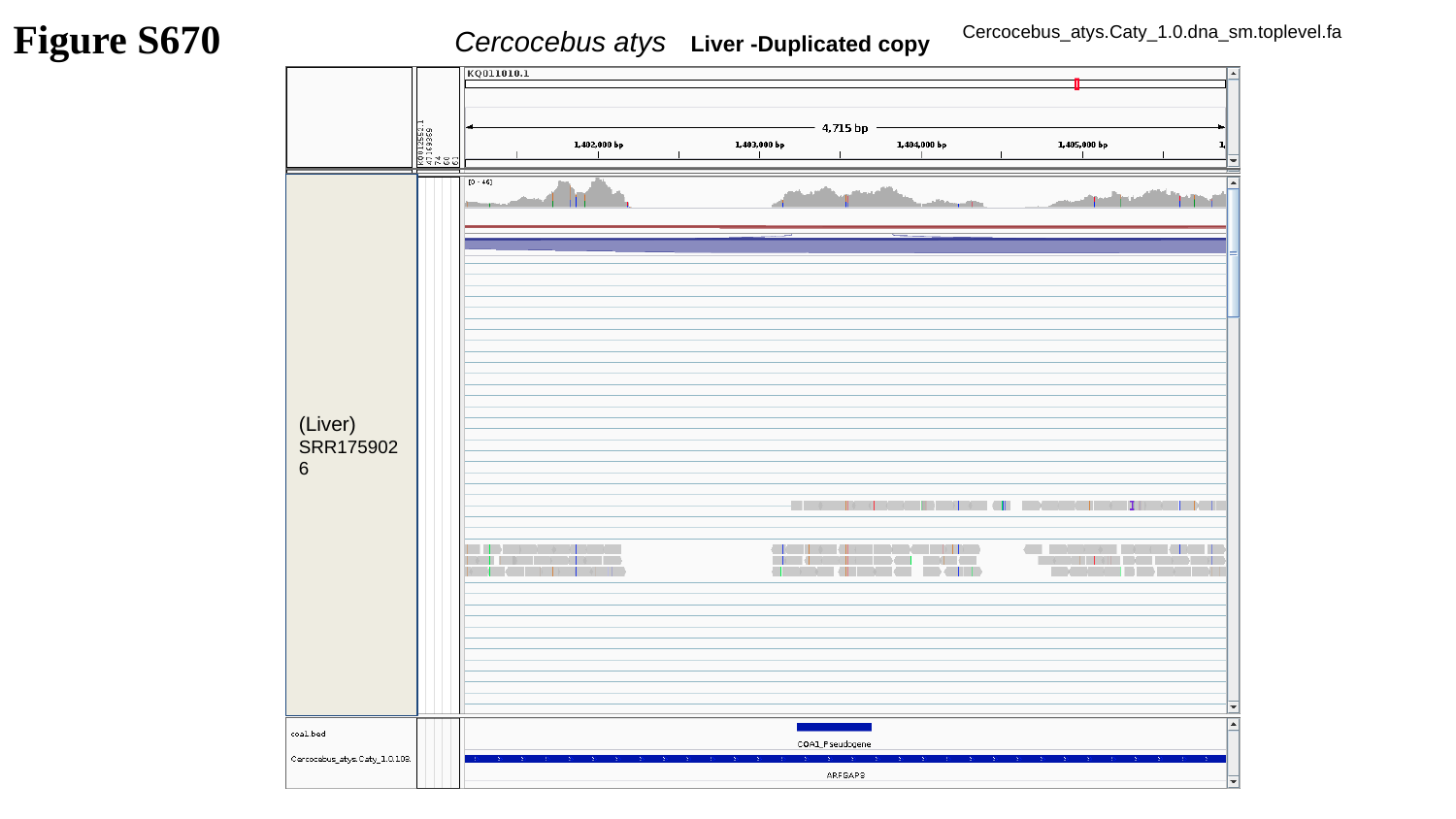

Figure S670
Cercocebus_atys.Caty_1.0.dna_sm.toplevel.fa
Cercocebus atys Liver -Duplicated copy
(Liver)
SRR1759026

### Slide 671
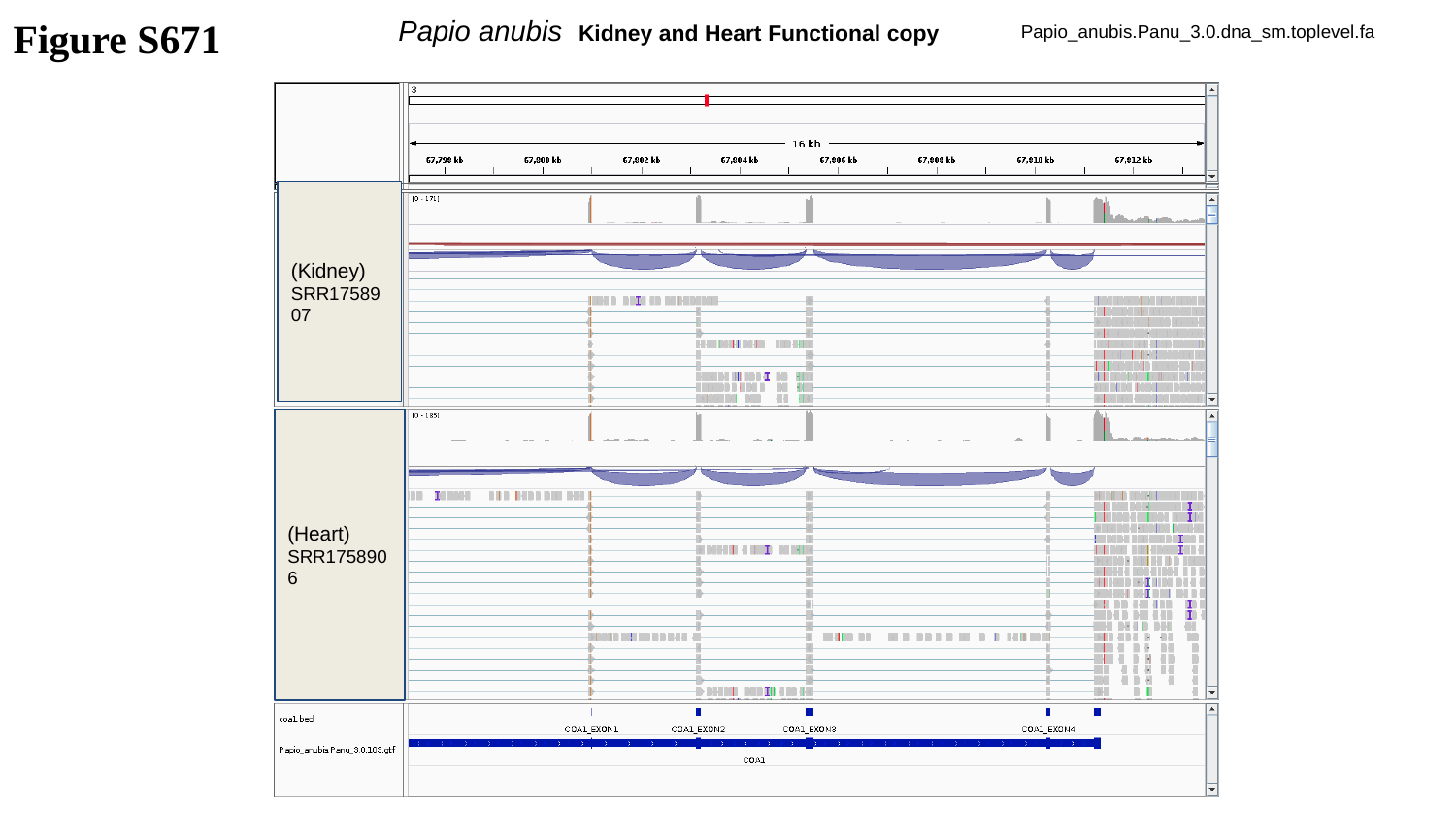

Figure S671
Papio anubis Kidney and Heart Functional copy
Papio_anubis.Panu_3.0.dna_sm.toplevel.fa
(Kidney)
SRR1758907
(Heart)
SRR1758906

### Slide 672
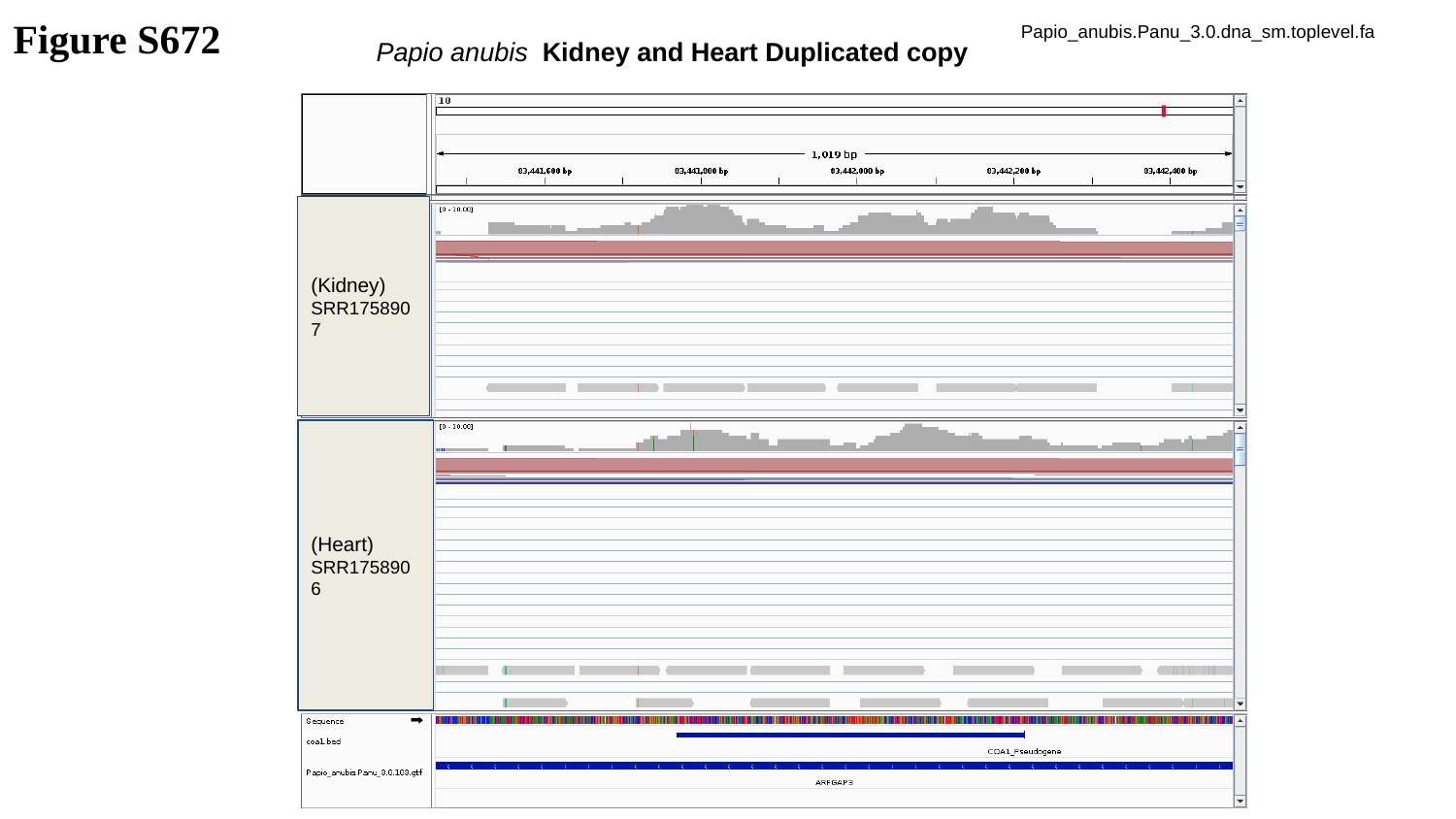

Figure S672
Papio_anubis.Panu_3.0.dna_sm.toplevel.fa
Papio anubis Kidney and Heart Duplicated copy
(Kidney)
SRR1758907
(Heart)
SRR1758906

### Slide 673
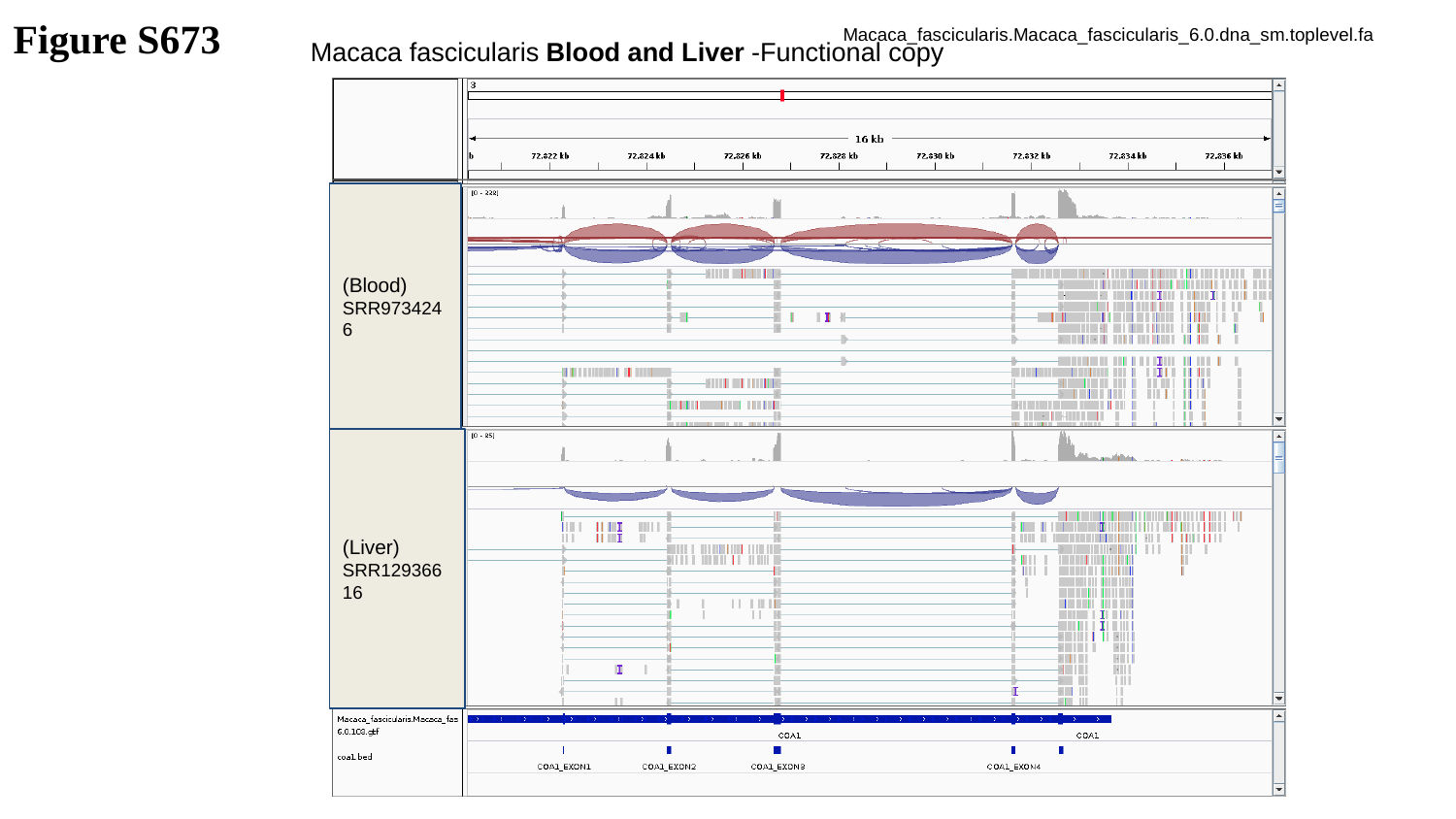

Figure S673
Macaca_fascicularis.Macaca_fascicularis_6.0.dna_sm.toplevel.fa
Macaca fascicularis Blood and Liver -Functional copy
(Blood)
SRR9734246
(Liver)
SRR12936616

### Slide 674
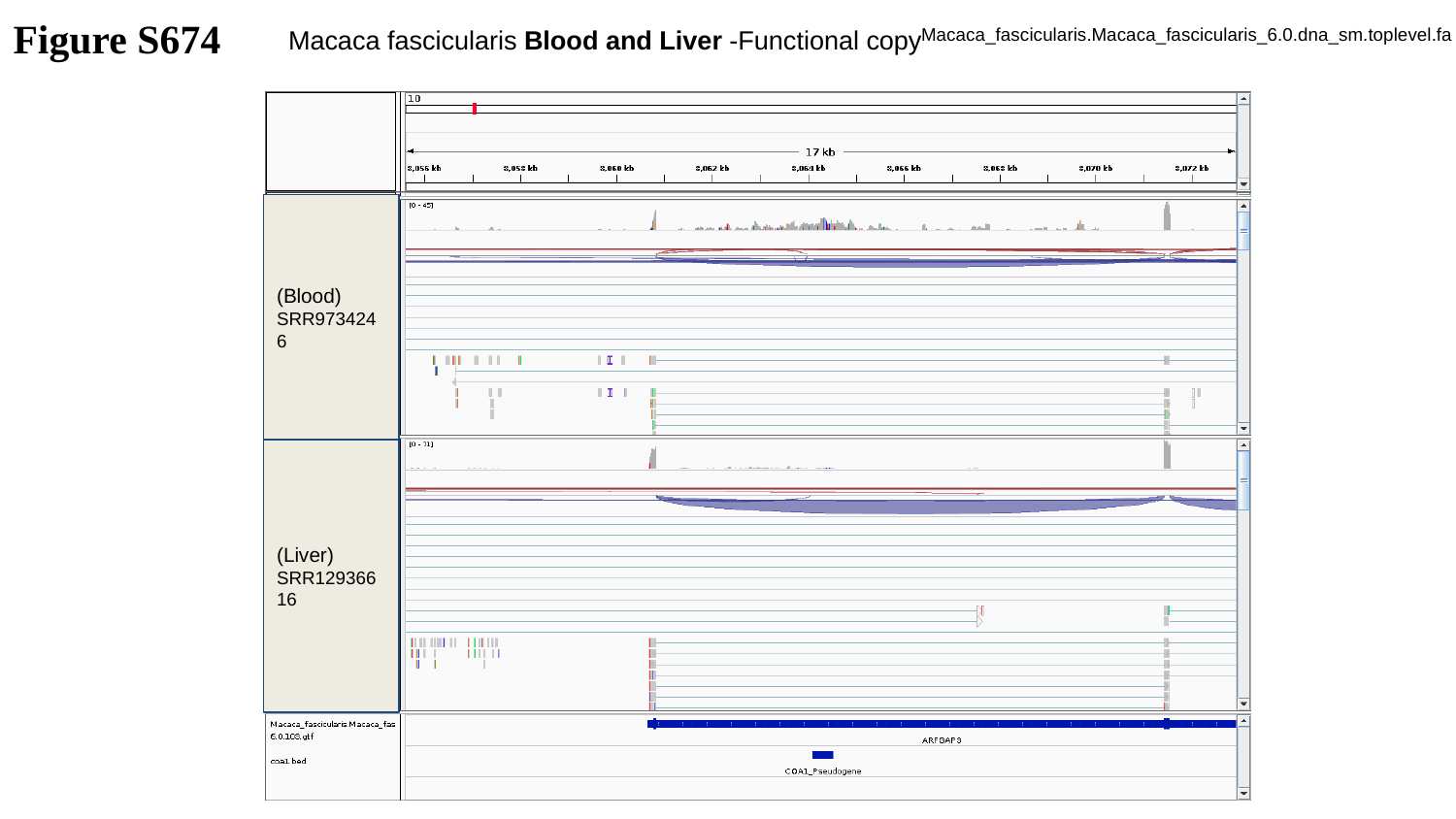

Figure S674
Macaca_fascicularis.Macaca_fascicularis_6.0.dna_sm.toplevel.fa
Macaca fascicularis Blood and Liver -Functional copy
(Blood)
SRR9734246
(Liver)
SRR12936616

### Slide 675
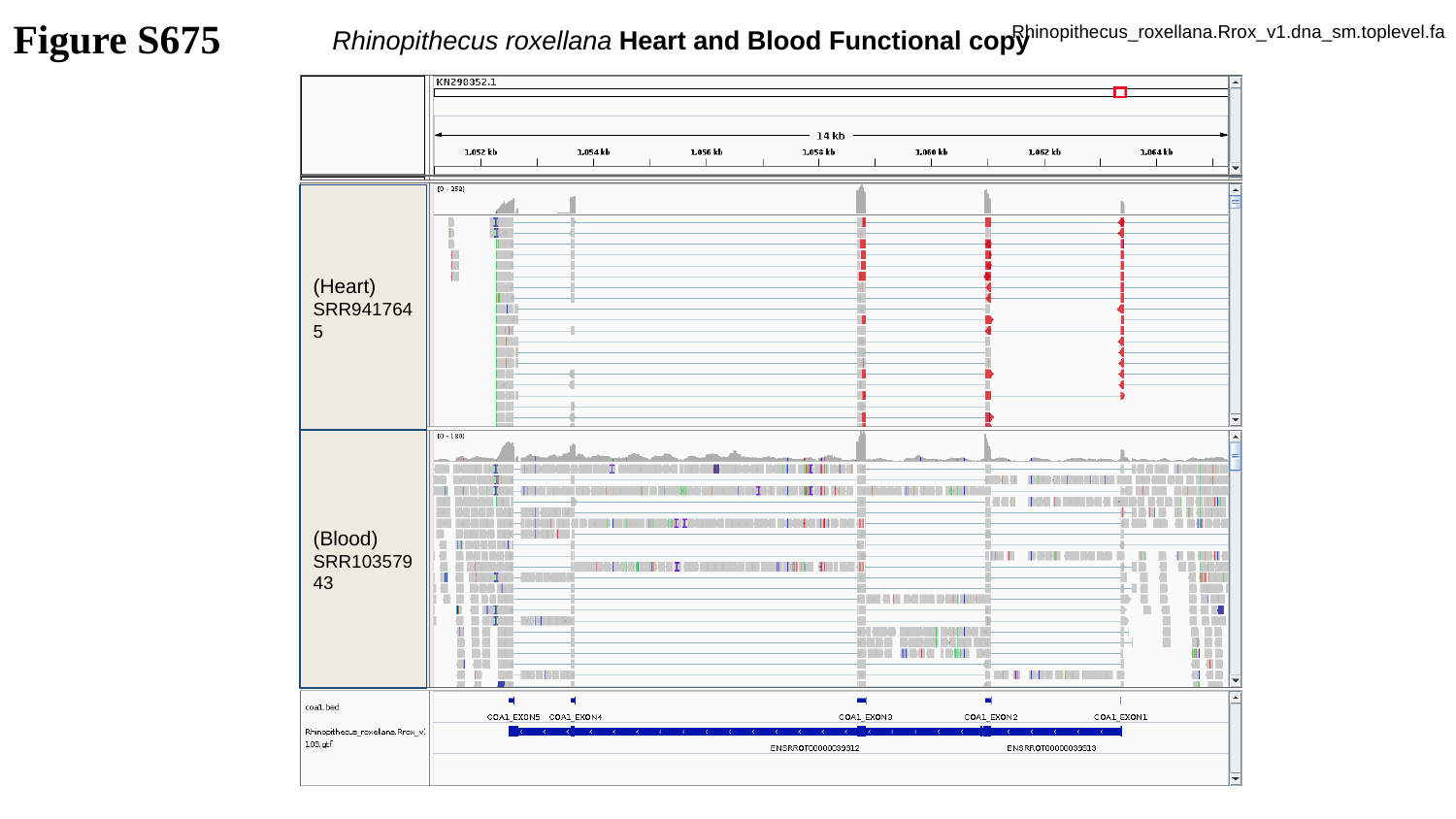

Figure S675
Rhinopithecus_roxellana.Rrox_v1.dna_sm.toplevel.fa
Rhinopithecus roxellana Heart and Blood Functional copy
(Heart)
SRR9417645
(Blood)
SRR10357943

### Slide 676
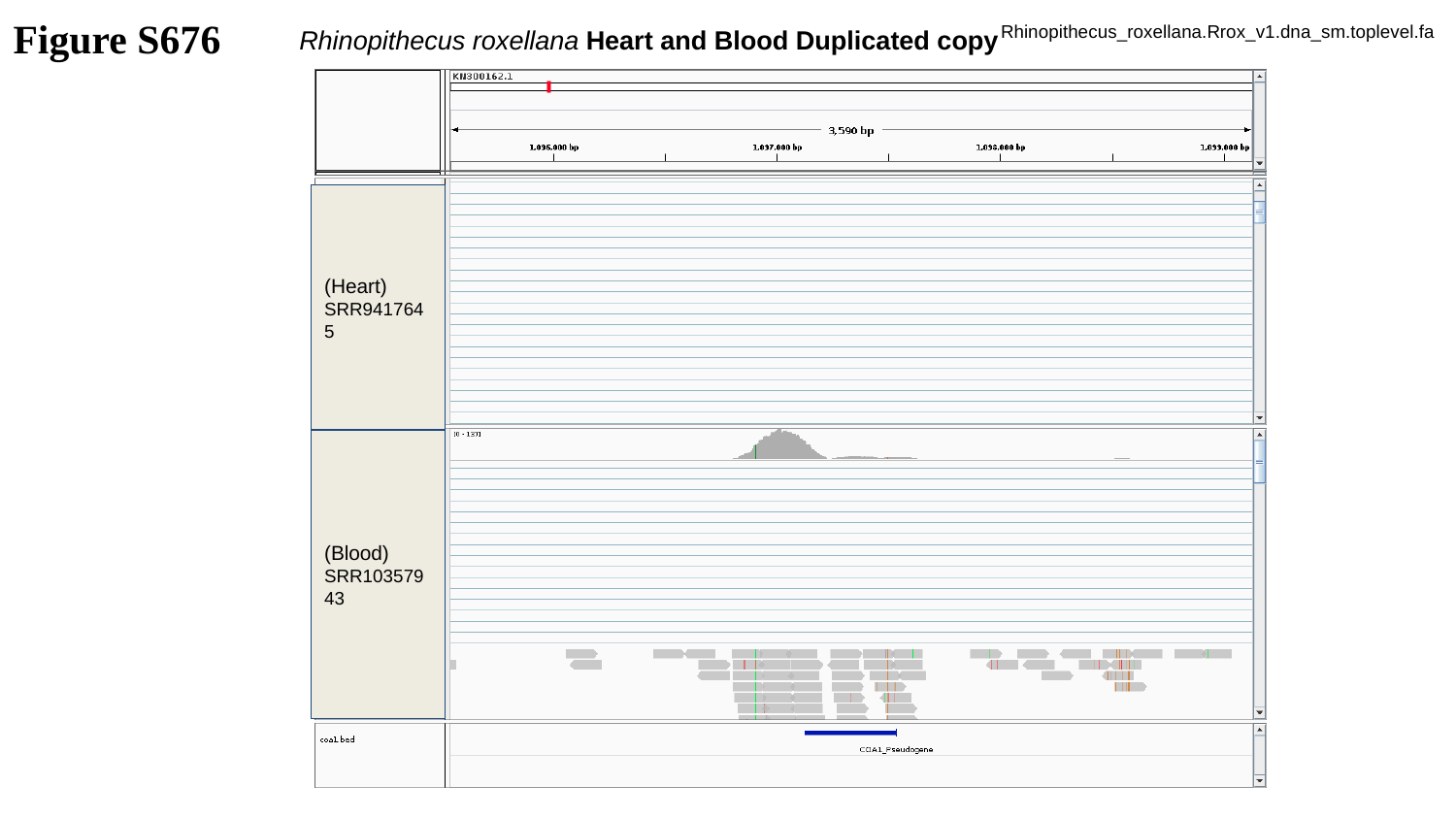

Figure S676
Rhinopithecus_roxellana.Rrox_v1.dna_sm.toplevel.fa
Rhinopithecus roxellana Heart and Blood Duplicated copy
(Heart)
SRR9417645
(Blood)
SRR10357943

### Slide 677
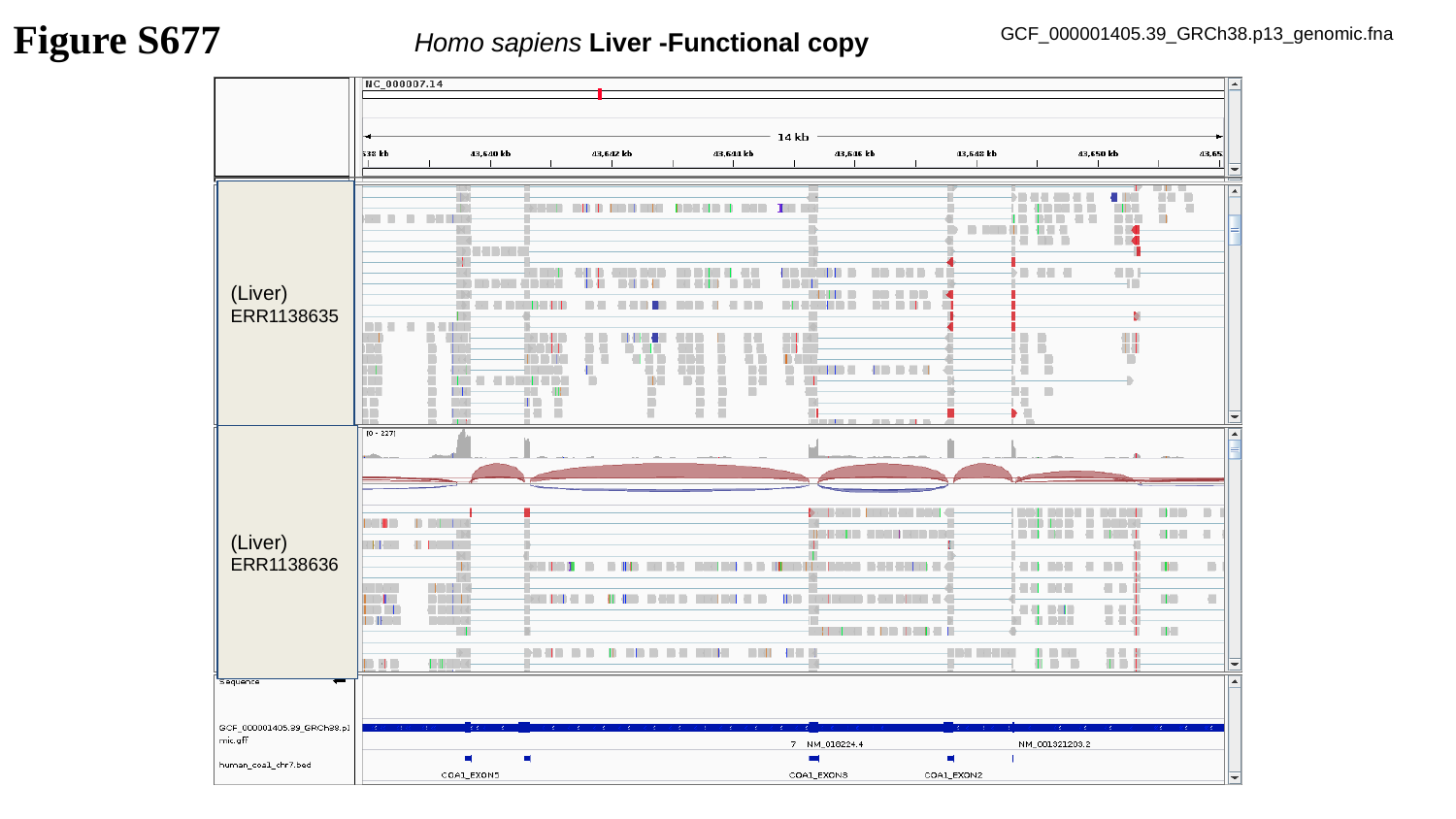

Figure S677
GCF_000001405.39_GRCh38.p13_genomic.fna
Homo sapiens Liver -Functional copy
(Liver)
ERR1138635
(Liver)
ERR1138636

### Slide 678
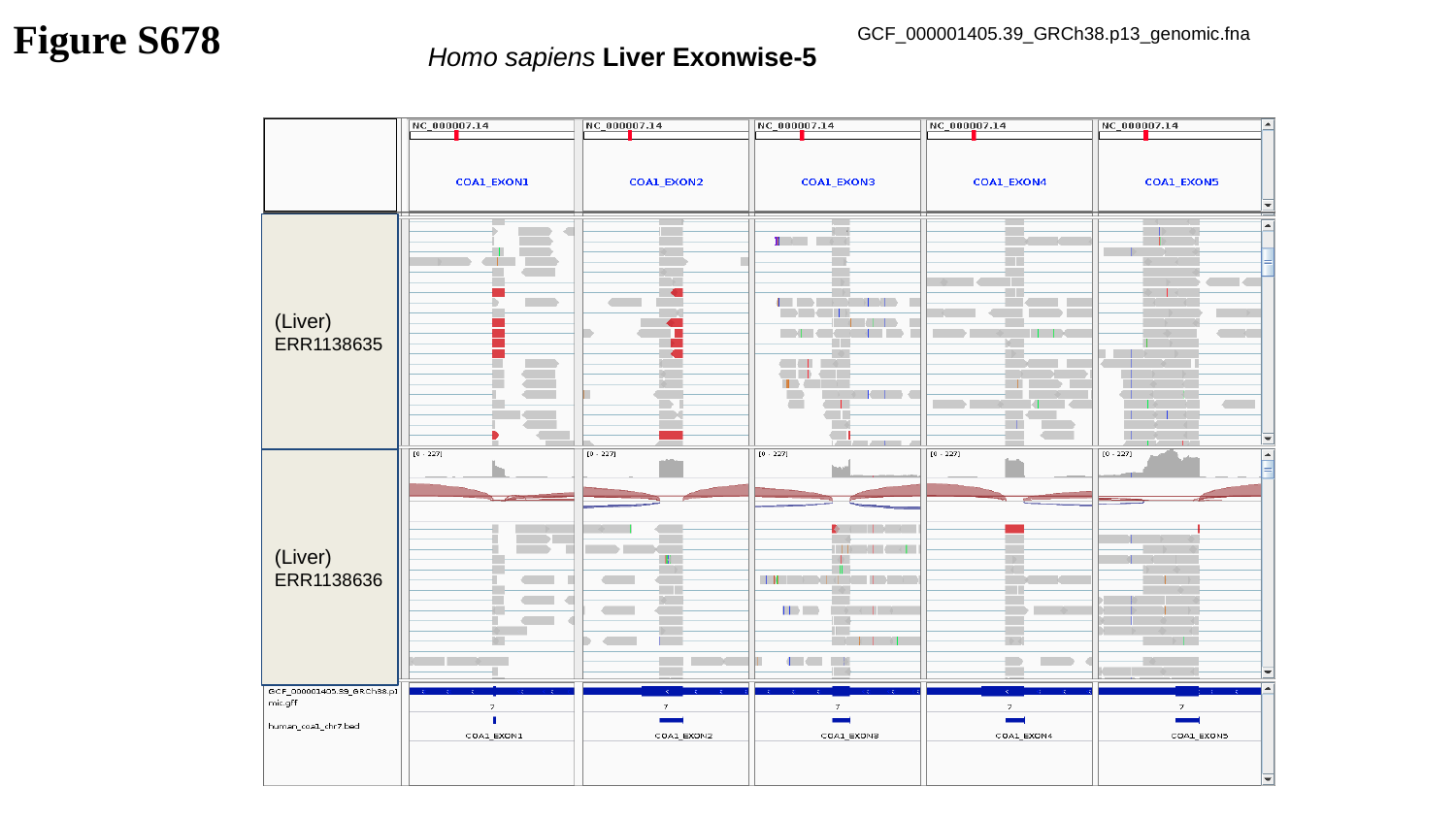

Figure S678
GCF_000001405.39_GRCh38.p13_genomic.fna
Homo sapiens Liver Exonwise-5
(Liver)
ERR1138635
(Liver)
ERR1138636

### Slide 679
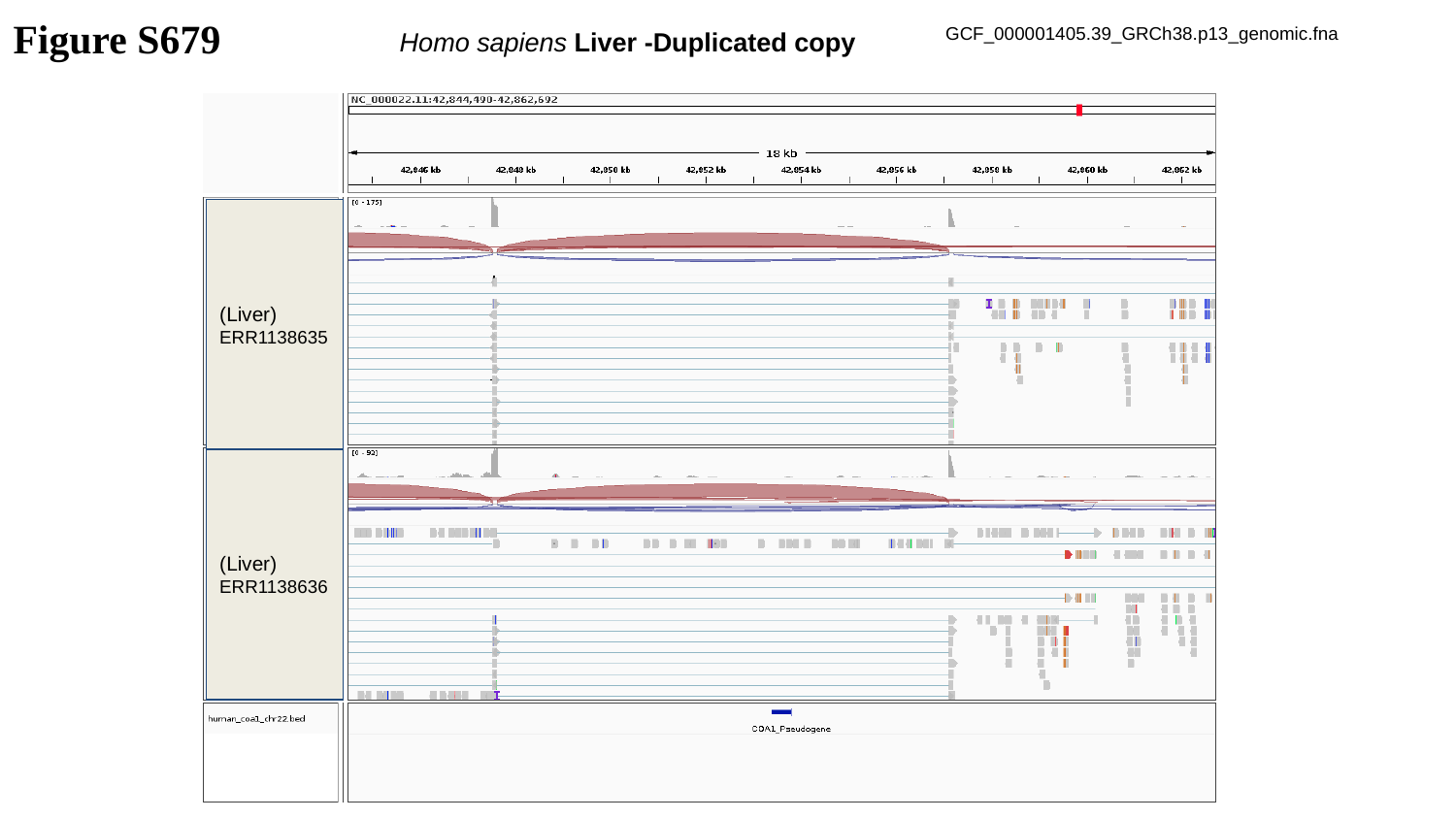

Figure S679
GCF_000001405.39_GRCh38.p13_genomic.fna
Homo sapiens Liver -Duplicated copy
(Liver)
ERR1138635
(Liver)
ERR1138636

### Slide 680
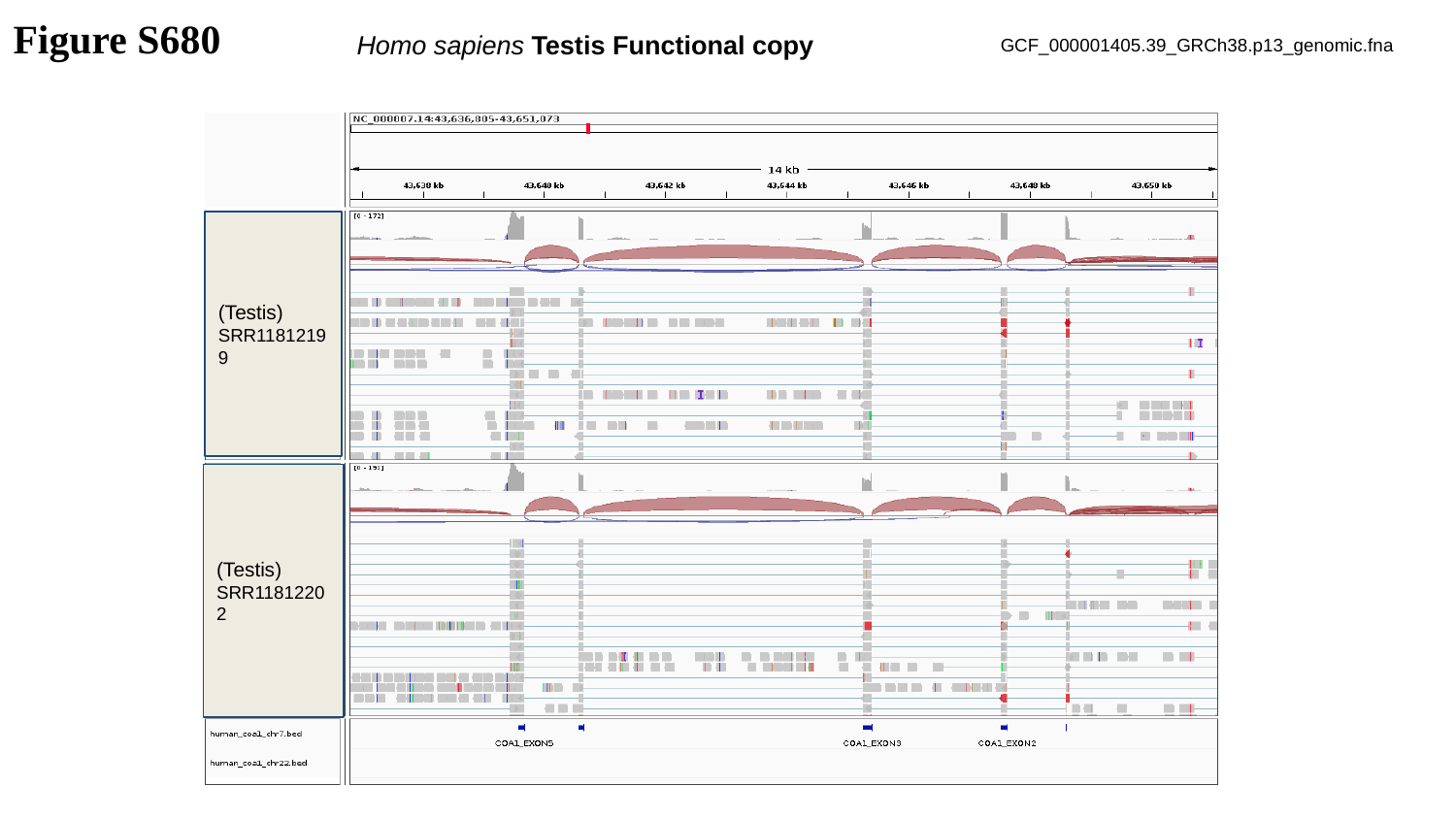

Figure S680
Homo sapiens Testis Functional copy
GCF_000001405.39_GRCh38.p13_genomic.fna
(Testis)
SRR11812199
(Testis)
SRR11812202

### Slide 681
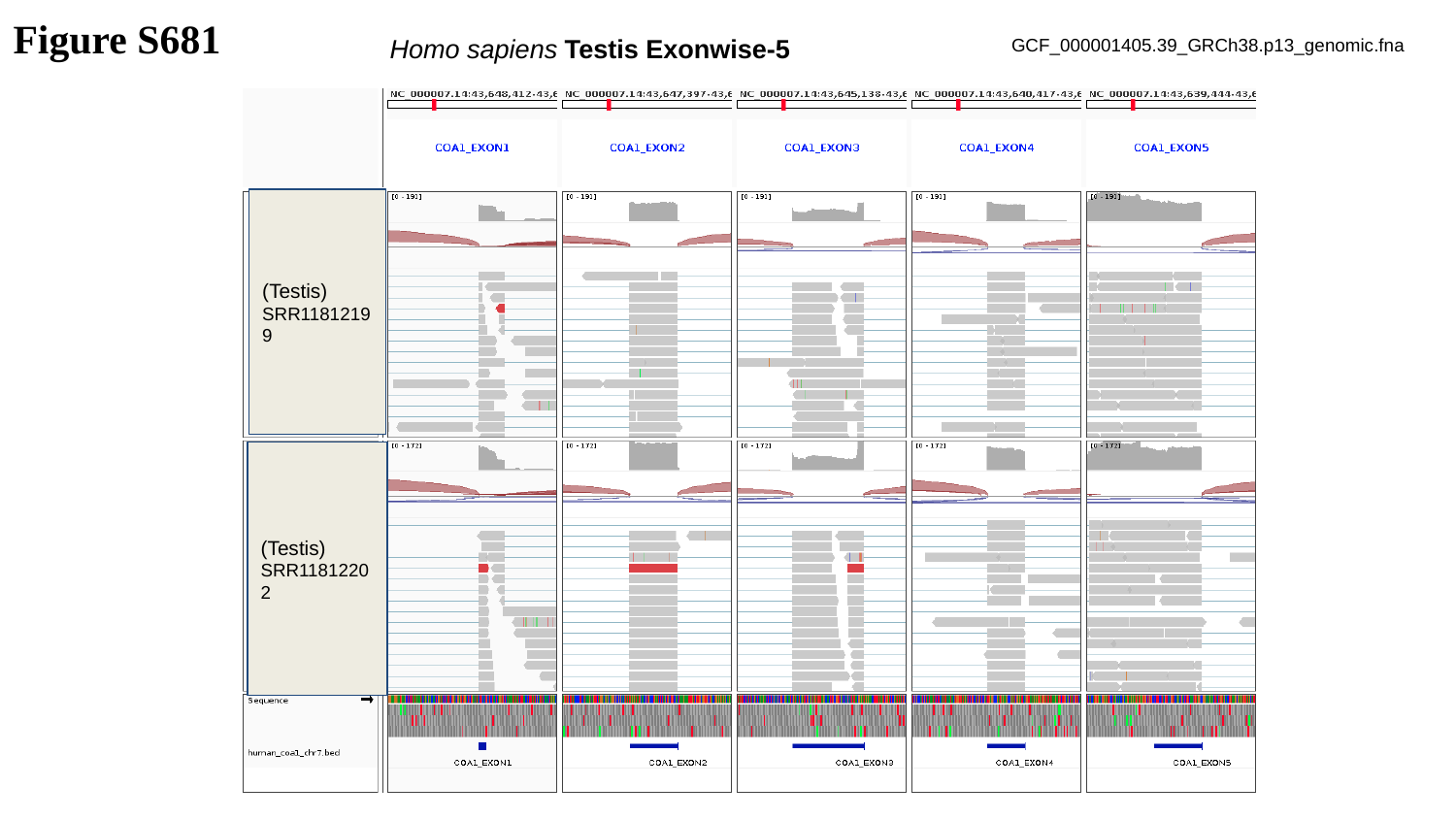

Figure S681
Homo sapiens Testis Exonwise-5
GCF_000001405.39_GRCh38.p13_genomic.fna
(Testis)
SRR11812199
(Testis)
SRR11812202

### Slide 682
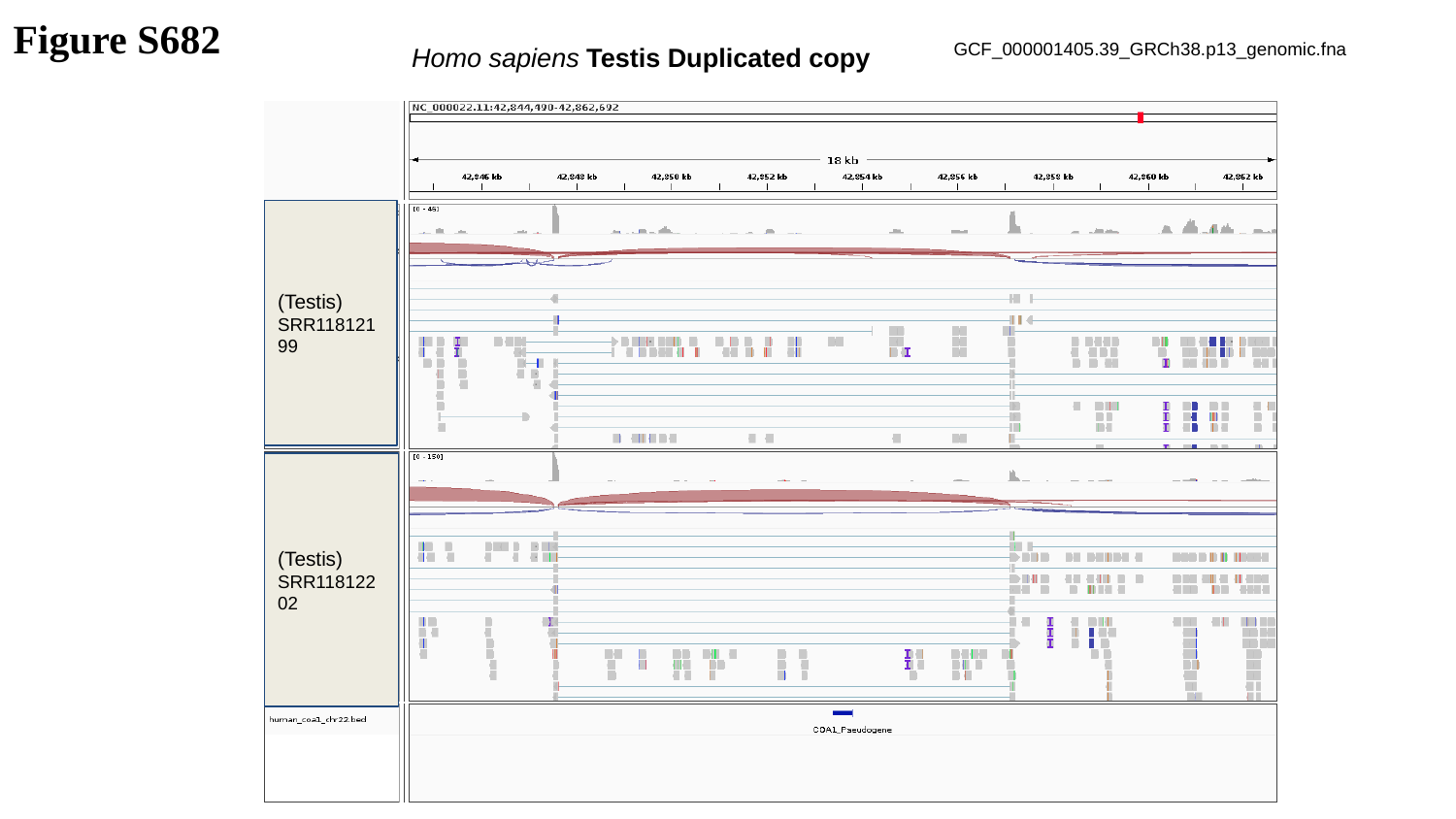

Figure S682
GCF_000001405.39_GRCh38.p13_genomic.fna
Homo sapiens Testis Duplicated copy
(Testis)
SRR11812199
(Testis)
SRR11812202

### Slide 683
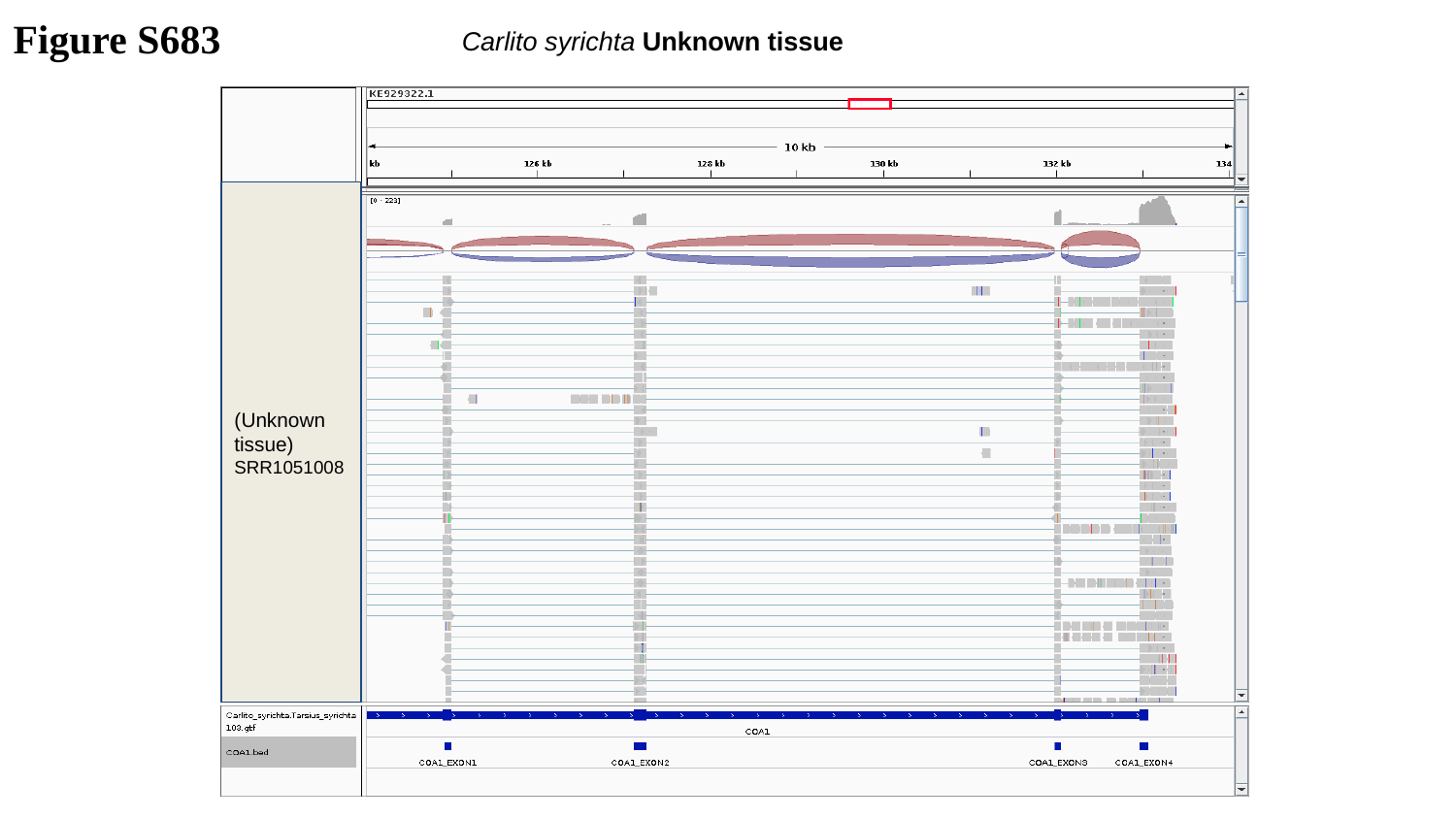

Figure S683
Carlito syrichta Unknown tissue
(Unknown tissue)
SRR1051008

### Slide 684
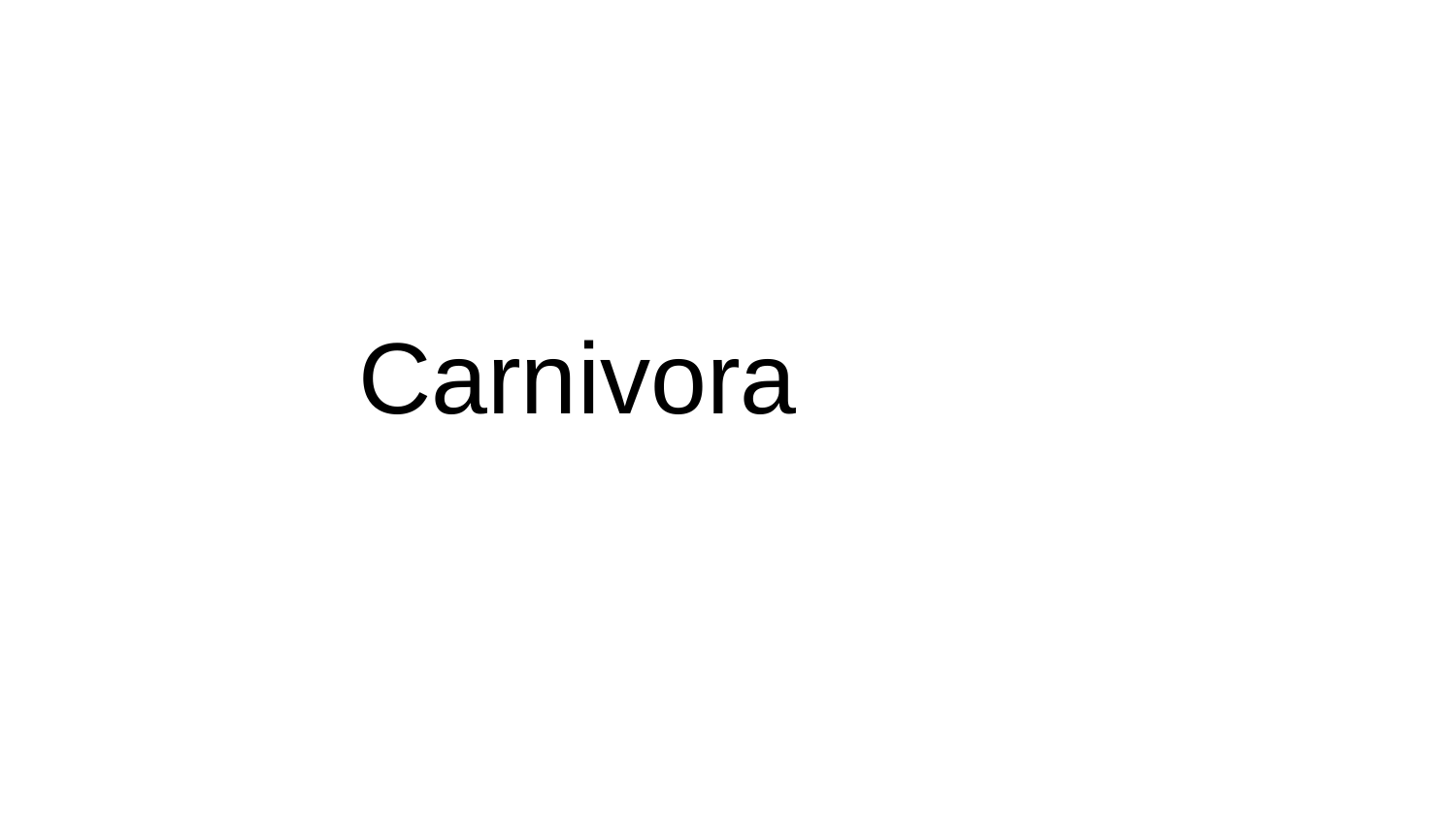

Carnivora

### Slide 685
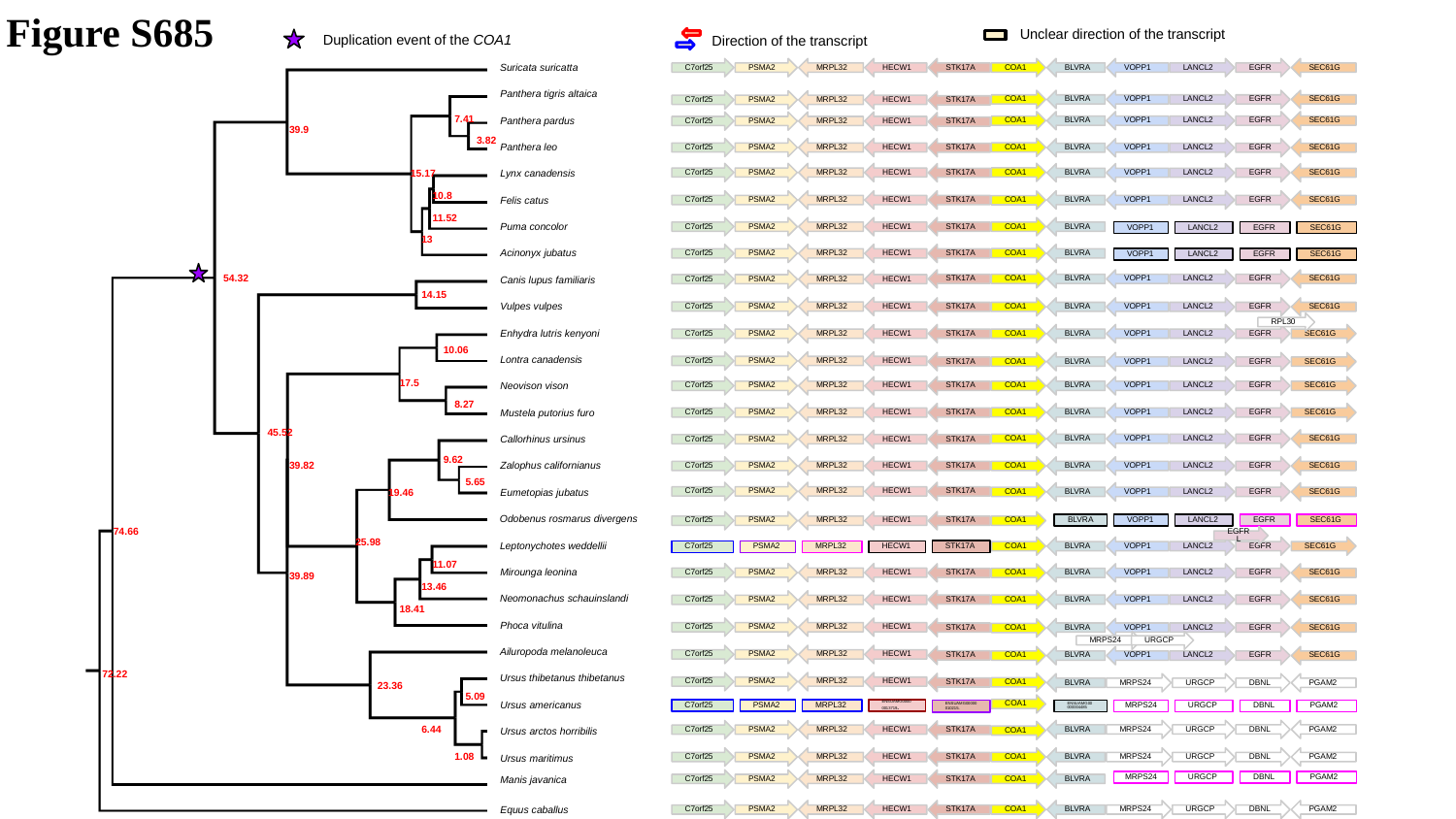

Figure S685
Unclear direction of the transcript
Direction of the transcript
Duplication event of the COA1
C7orf25
PSMA2
MRPL32
HECW1
STK17A
COA1
BLVRA
VOPP1
LANCL2
EGFR
SEC61G
Suricata suricatta
Panthera tigris altaica
COA1
BLVRA
VOPP1
LANCL2
EGFR
SEC61G
C7orf25
PSMA2
MRPL32
HECW1
STK17A
COA1
BLVRA
VOPP1
LANCL2
EGFR
SEC61G
7.41
C7orf25
PSMA2
MRPL32
HECW1
STK17A
Panthera pardus
39.9
3.82
COA1
C7orf25
PSMA2
MRPL32
HECW1
STK17A
BLVRA
VOPP1
LANCL2
EGFR
SEC61G
Panthera leo
COA1
C7orf25
PSMA2
MRPL32
HECW1
STK17A
BLVRA
VOPP1
LANCL2
EGFR
SEC61G
15.17
Lynx canadensis
10.8
C7orf25
PSMA2
MRPL32
HECW1
STK17A
BLVRA
VOPP1
LANCL2
EGFR
SEC61G
COA1
Felis catus
11.52
C7orf25
PSMA2
MRPL32
HECW1
STK17A
COA1
BLVRA
Puma concolor
VOPP1
LANCL2
EGFR
SEC61G
13
COA1
C7orf25
PSMA2
MRPL32
HECW1
STK17A
BLVRA
Acinonyx jubatus
VOPP1
LANCL2
EGFR
SEC61G
STK17A
COA1
BLVRA
VOPP1
LANCL2
EGFR
SEC61G
C7orf25
PSMA2
MRPL32
HECW1
54.32
Canis lupus familiaris
14.15
C7orf25
PSMA2
MRPL32
HECW1
STK17A
COA1
BLVRA
VOPP1
LANCL2
EGFR
SEC61G
Vulpes vulpes
RPL30
STK17A
COA1
BLVRA
VOPP1
LANCL2
EGFR
SEC61G
C7orf25
PSMA2
MRPL32
HECW1
Enhydra lutris kenyoni
10.06
C7orf25
PSMA2
MRPL32
HECW1
STK17A
COA1
BLVRA
VOPP1
LANCL2
EGFR
SEC61G
Lontra canadensis
17.5
C7orf25
PSMA2
MRPL32
HECW1
STK17A
COA1
BLVRA
VOPP1
LANCL2
EGFR
SEC61G
Neovison vison
8.27
COA1
C7orf25
PSMA2
MRPL32
HECW1
STK17A
BLVRA
VOPP1
LANCL2
EGFR
SEC61G
Mustela putorius furo
45.52
COA1
BLVRA
VOPP1
LANCL2
EGFR
SEC61G
C7orf25
PSMA2
MRPL32
HECW1
STK17A
Callorhinus ursinus
9.62
C7orf25
PSMA2
MRPL32
HECW1
STK17A
COA1
BLVRA
VOPP1
LANCL2
EGFR
SEC61G
39.82
Zalophus californianus
5.65
C7orf25
PSMA2
MRPL32
HECW1
STK17A
COA1
BLVRA
VOPP1
LANCL2
EGFR
SEC61G
19.46
Eumetopias jubatus
C7orf25
PSMA2
MRPL32
HECW1
STK17A
COA1
Odobenus rosmarus divergens
BLVRA
VOPP1
LANCL2
EGFR
SEC61G
74.66
EGFRL
25.98
COA1
BLVRA
VOPP1
LANCL2
EGFR
SEC61G
Leptonychotes weddellii
STK17A
C7orf25
PSMA2
MRPL32
HECW1
11.07
C7orf25
PSMA2
MRPL32
HECW1
STK17A
COA1
BLVRA
VOPP1
LANCL2
EGFR
SEC61G
Mirounga leonina
39.89
13.46
STK17A
COA1
BLVRA
VOPP1
LANCL2
EGFR
SEC61G
C7orf25
PSMA2
MRPL32
HECW1
Neomonachus schauinslandi
18.41
C7orf25
PSMA2
MRPL32
HECW1
STK17A
BLVRA
VOPP1
LANCL2
EGFR
SEC61G
COA1
Phoca vitulina
MRPS24
URGCP
C7orf25
PSMA2
MRPL32
HECW1
STK17A
COA1
BLVRA
VOPP1
LANCL2
EGFR
SEC61G
Ailuropoda melanoleuca
72.22
C7orf25
PSMA2
MRPL32
HECW1
Ursus thibetanus thibetanus
COA1
STK17A
BLVRA
MRPS24
URGCP
DBNL
PGAM2
23.36
5.09
COA1
Ursus americanus
C7orf25
PSMA2
MRPL32
ENSUAMG00000013719.
ENSUAMG00000004485
MRPS24
URGCP
DBNL
PGAM2
ENSUAMG00000010215.
BLVRA
MRPS24
URGCP
DBNL
PGAM2
C7orf25
PSMA2
MRPL32
HECW1
STK17A
COA1
6.44
Ursus arctos horribilis
C7orf25
PSMA2
MRPL32
HECW1
STK17A
COA1
BLVRA
MRPS24
URGCP
DBNL
PGAM2
1.08
Ursus maritimus
C7orf25
PSMA2
MRPL32
HECW1
STK17A
COA1
BLVRA
MRPS24
URGCP
DBNL
PGAM2
Manis javanica
C7orf25
PSMA2
MRPL32
HECW1
STK17A
COA1
BLVRA
MRPS24
URGCP
DBNL
PGAM2
Equus caballus

### Slide 686
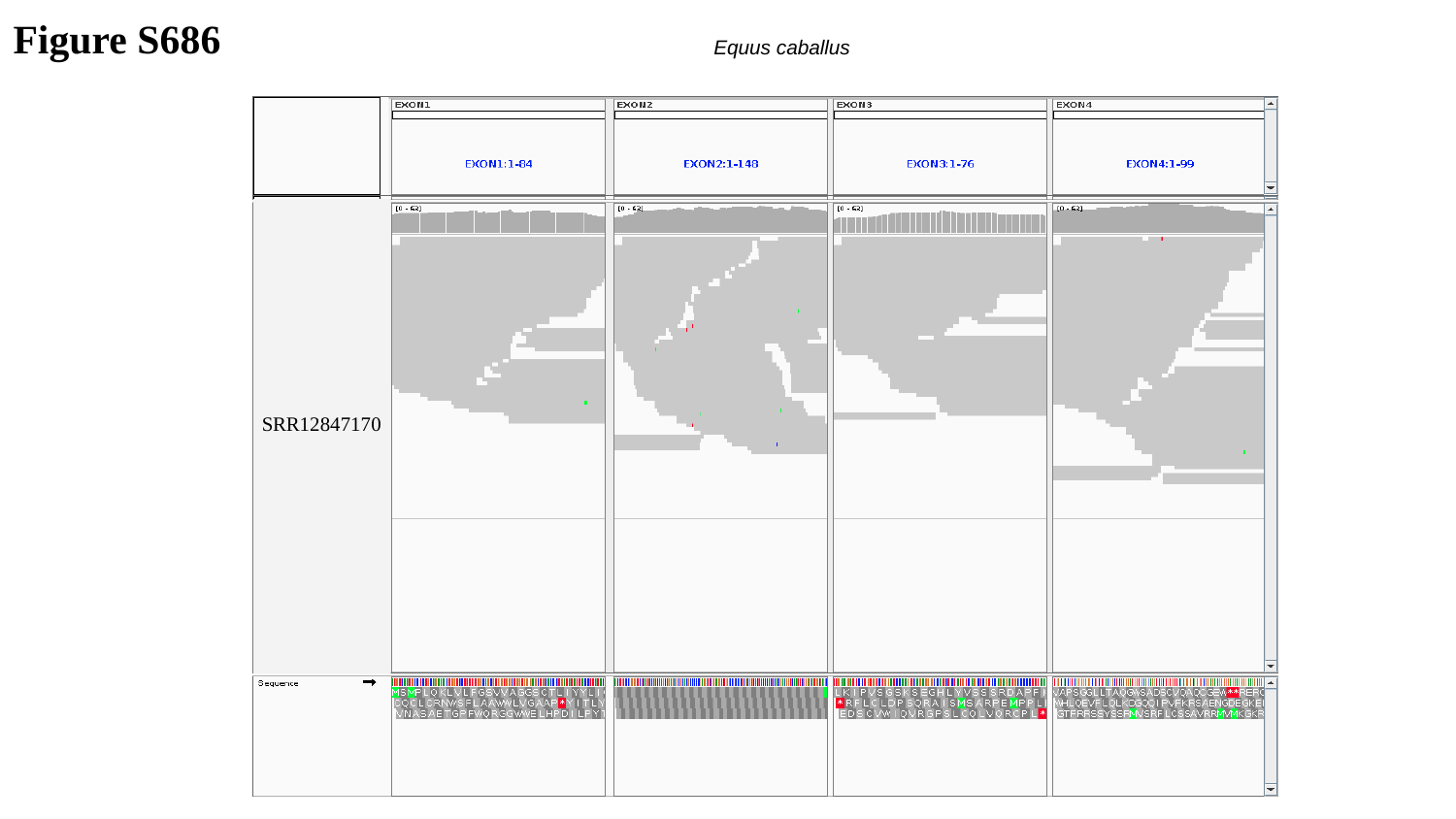

Figure S686
 Equus caballus
SRR12847170

### Slide 687
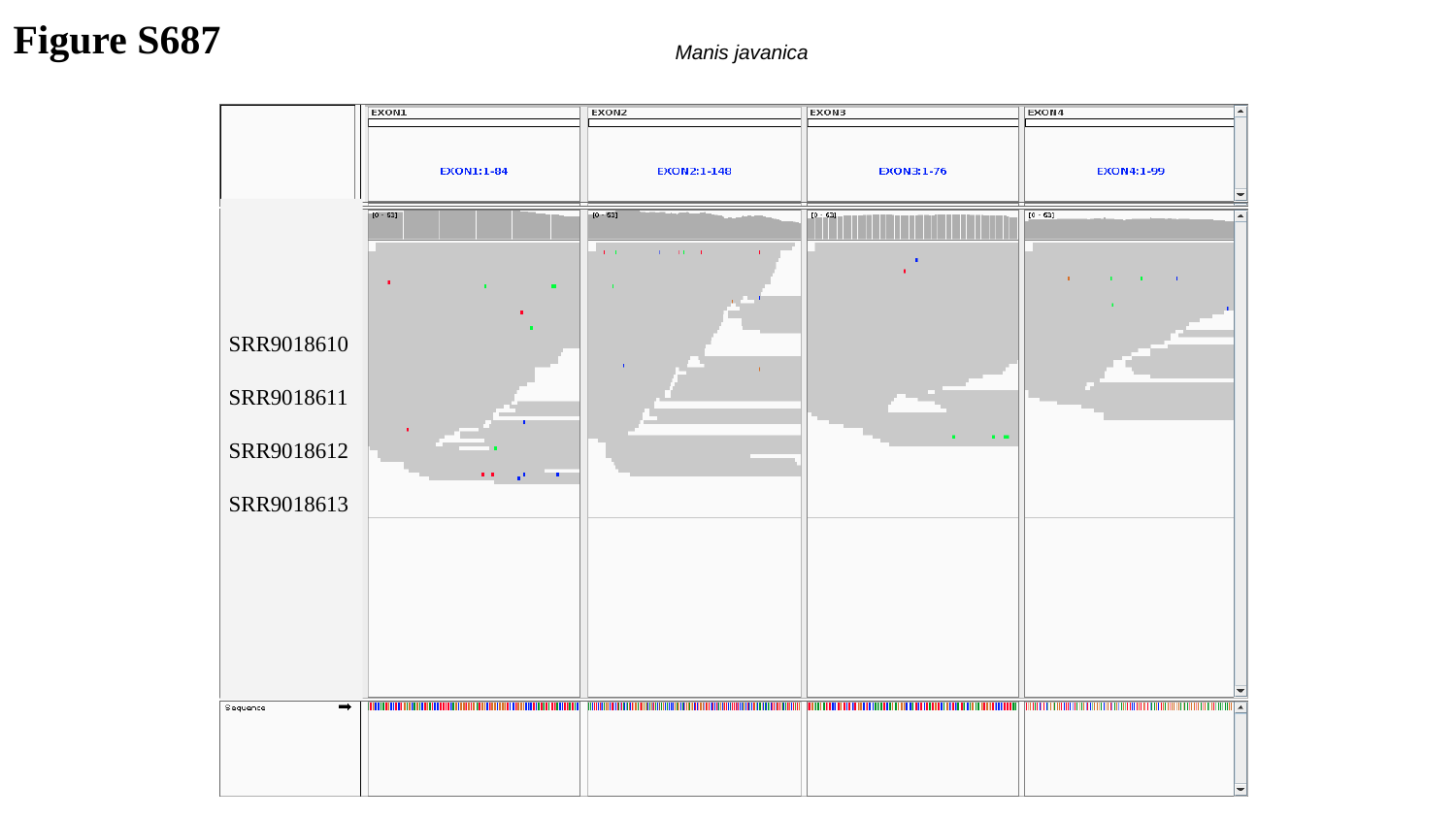

Figure S687
 Manis javanica
SRR9018610
SRR9018611
SRR9018612
SRR9018613

### Slide 688
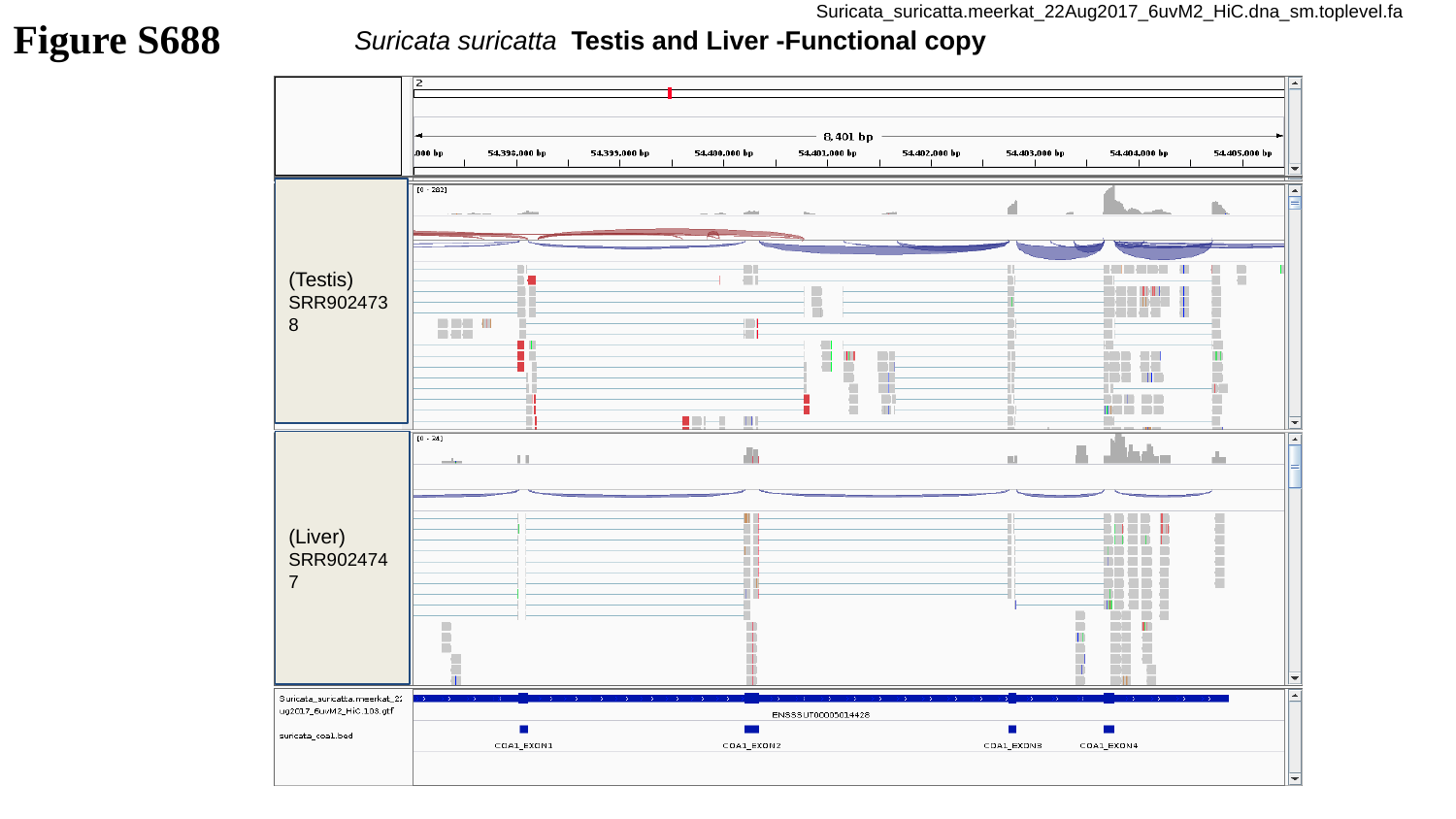

Figure S688
Suricata_suricatta.meerkat_22Aug2017_6uvM2_HiC.dna_sm.toplevel.fa
Suricata suricatta Testis and Liver -Functional copy
(Testis)
SRR9024738
(Liver)
SRR9024747

### Slide 689
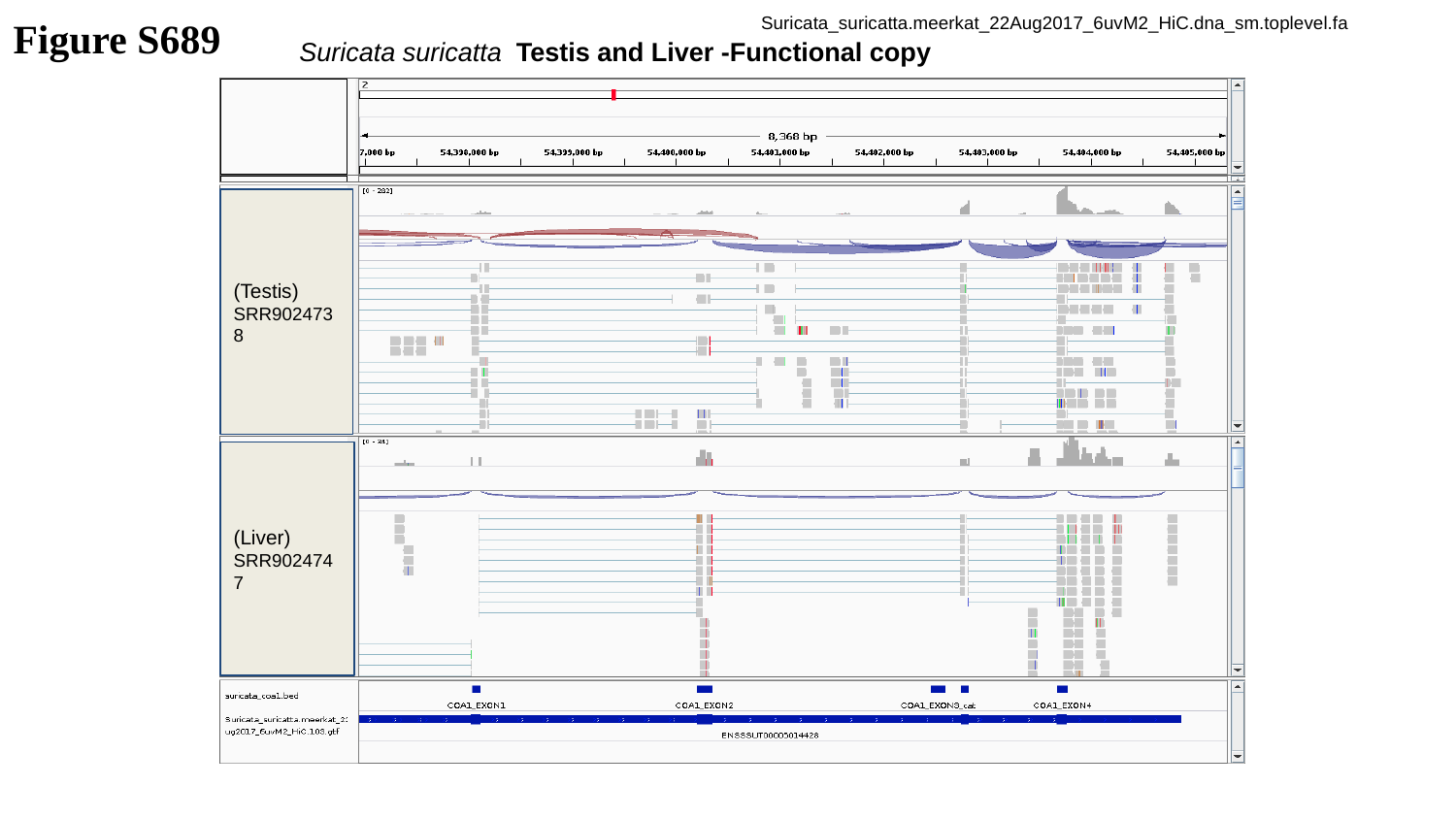

Figure S689
Suricata_suricatta.meerkat_22Aug2017_6uvM2_HiC.dna_sm.toplevel.fa
Suricata suricatta Testis and Liver -Functional copy
(Testis)
SRR9024738
(Liver)
SRR9024747

### Slide 690
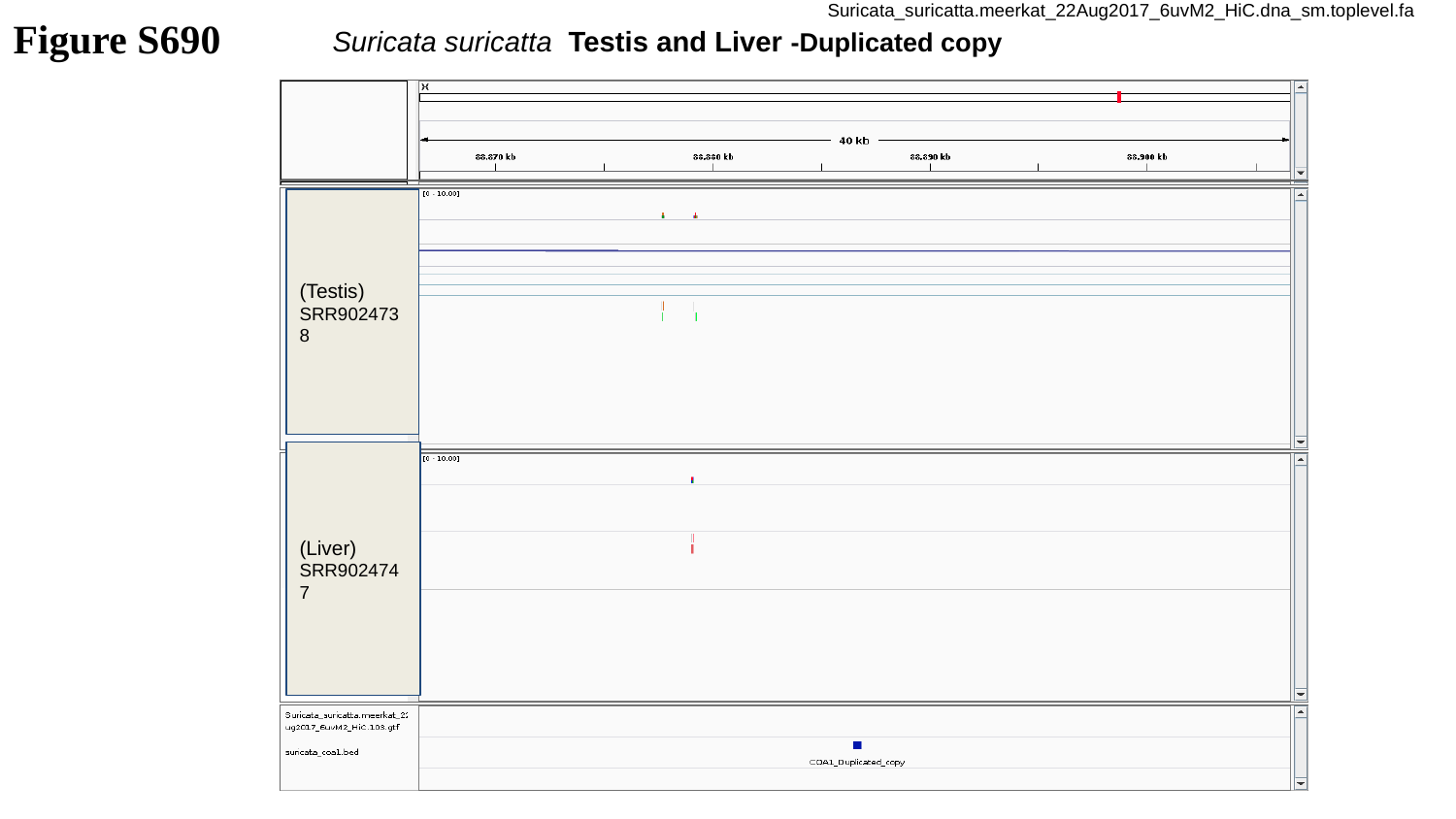

Figure S690
Suricata_suricatta.meerkat_22Aug2017_6uvM2_HiC.dna_sm.toplevel.fa
Suricata suricatta Testis and Liver -Duplicated copy
(Testis)
SRR9024738
(Liver)
SRR9024747

### Slide 691
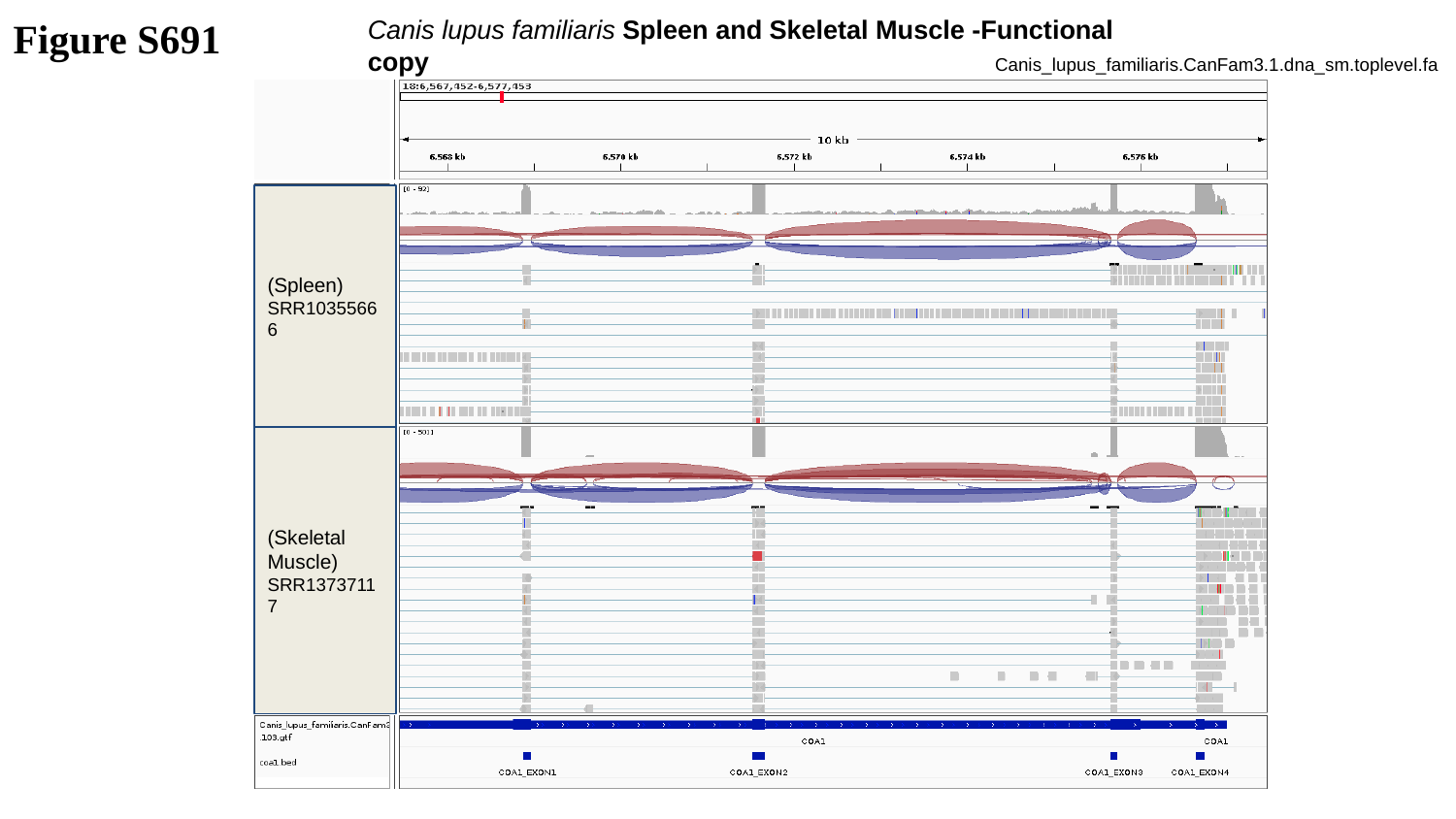

Figure S691
Canis lupus familiaris Spleen and Skeletal Muscle -Functional copy
Canis_lupus_familiaris.CanFam3.1.dna_sm.toplevel.fa
(Spleen)
SRR10355666
(Skeletal Muscle)
SRR13737117

### Slide 692
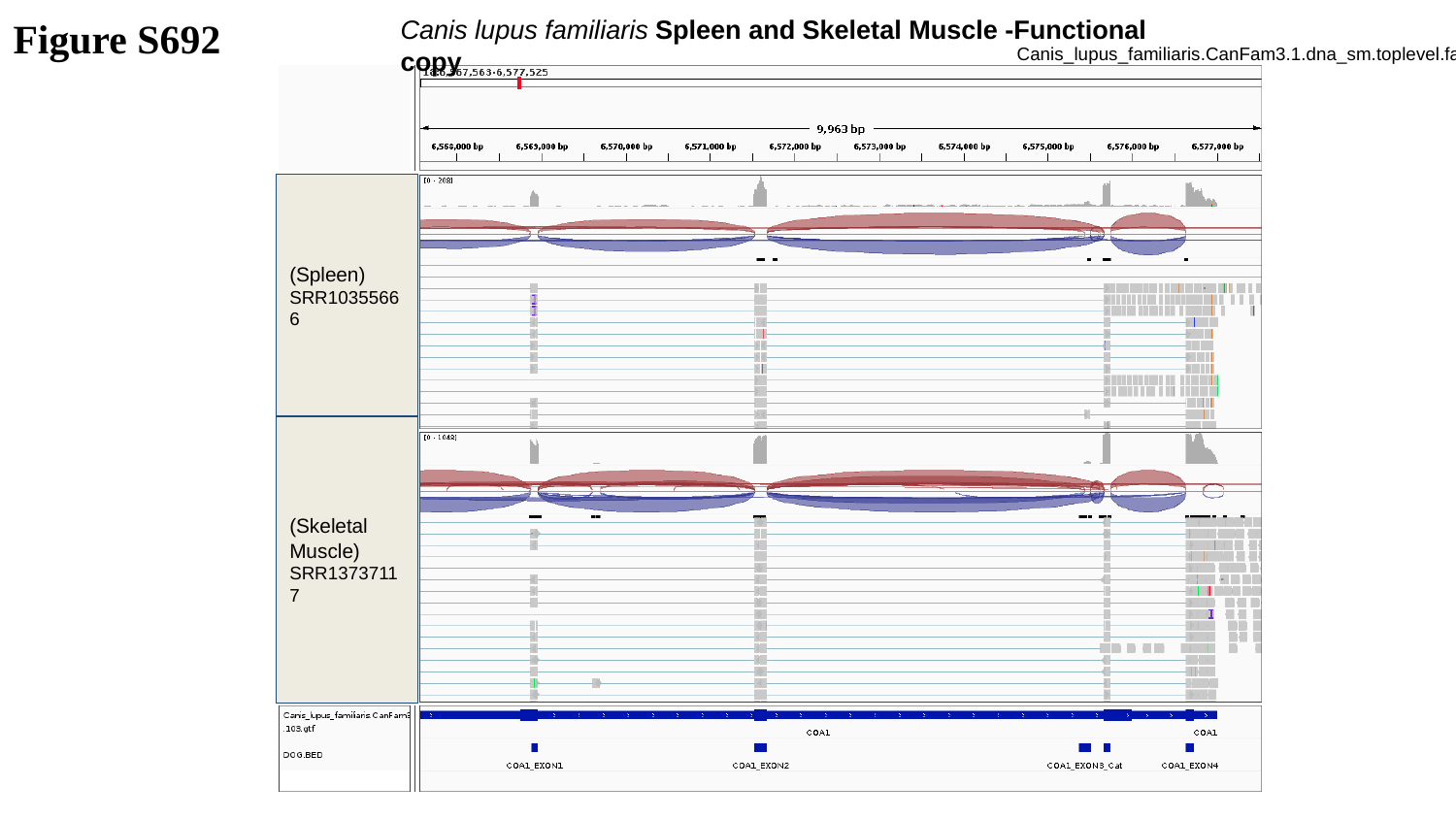

Figure S692
Canis lupus familiaris Spleen and Skeletal Muscle -Functional copy
Canis_lupus_familiaris.CanFam3.1.dna_sm.toplevel.fa
(Spleen)
SRR10355666
(Skeletal Muscle)
SRR13737117

### Slide 693
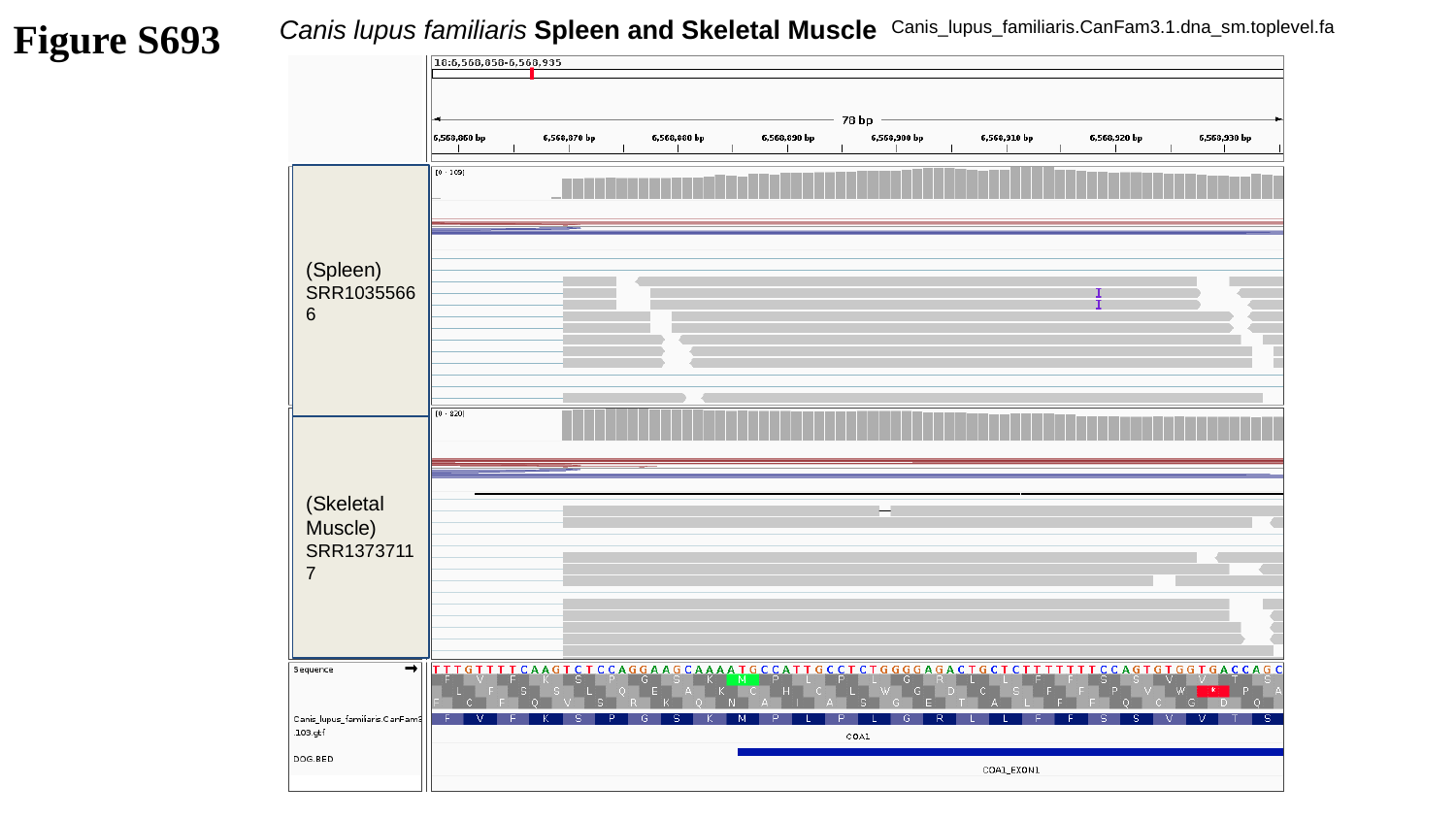

Figure S693
Canis lupus familiaris Spleen and Skeletal Muscle
Canis_lupus_familiaris.CanFam3.1.dna_sm.toplevel.fa
(Spleen)
SRR10355666
(Skeletal Muscle)
SRR13737117

### Slide 694
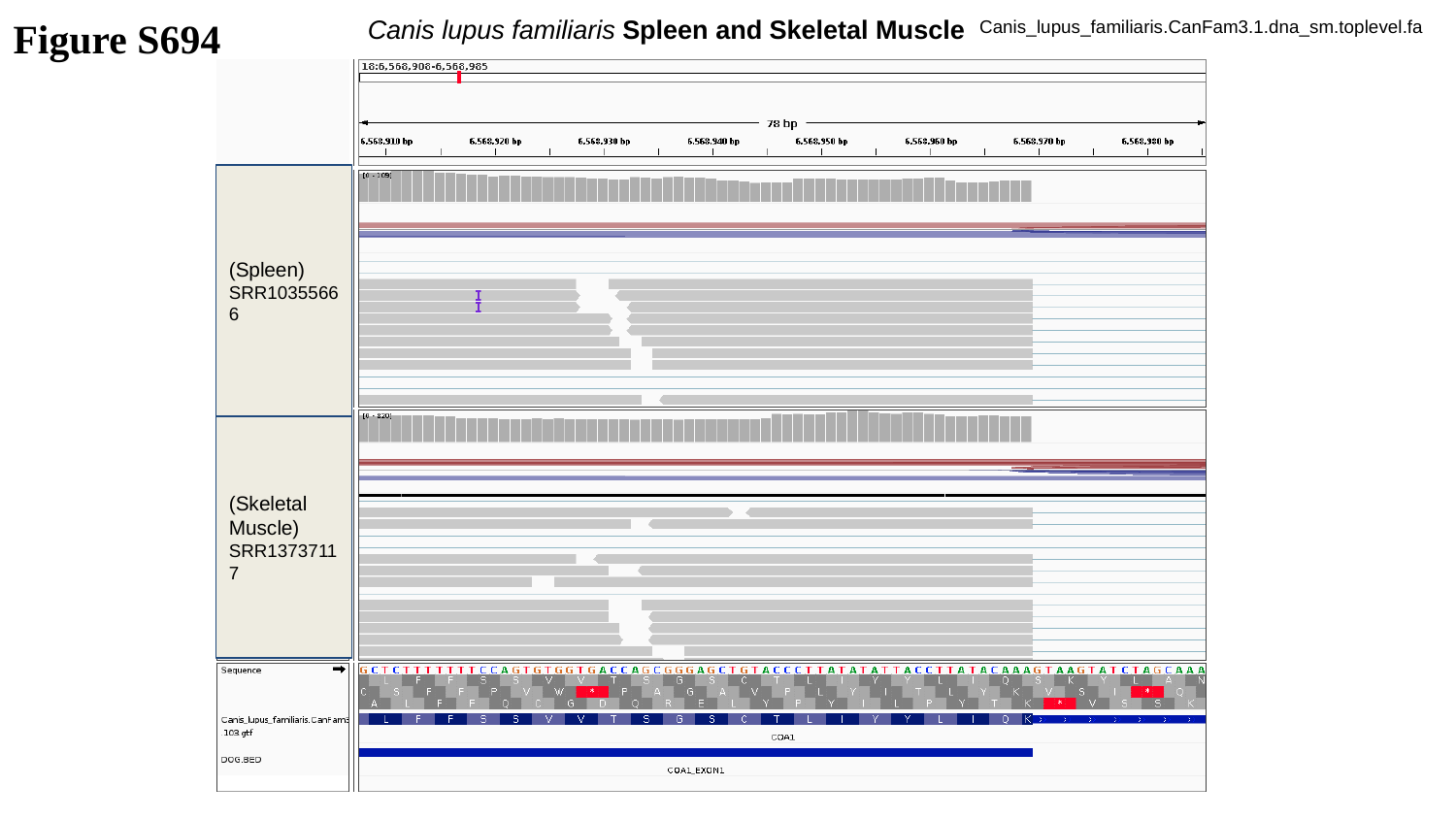

Figure S694
Canis lupus familiaris Spleen and Skeletal Muscle
Canis_lupus_familiaris.CanFam3.1.dna_sm.toplevel.fa
(Spleen)
SRR10355666
(Skeletal Muscle)
SRR13737117

### Slide 695
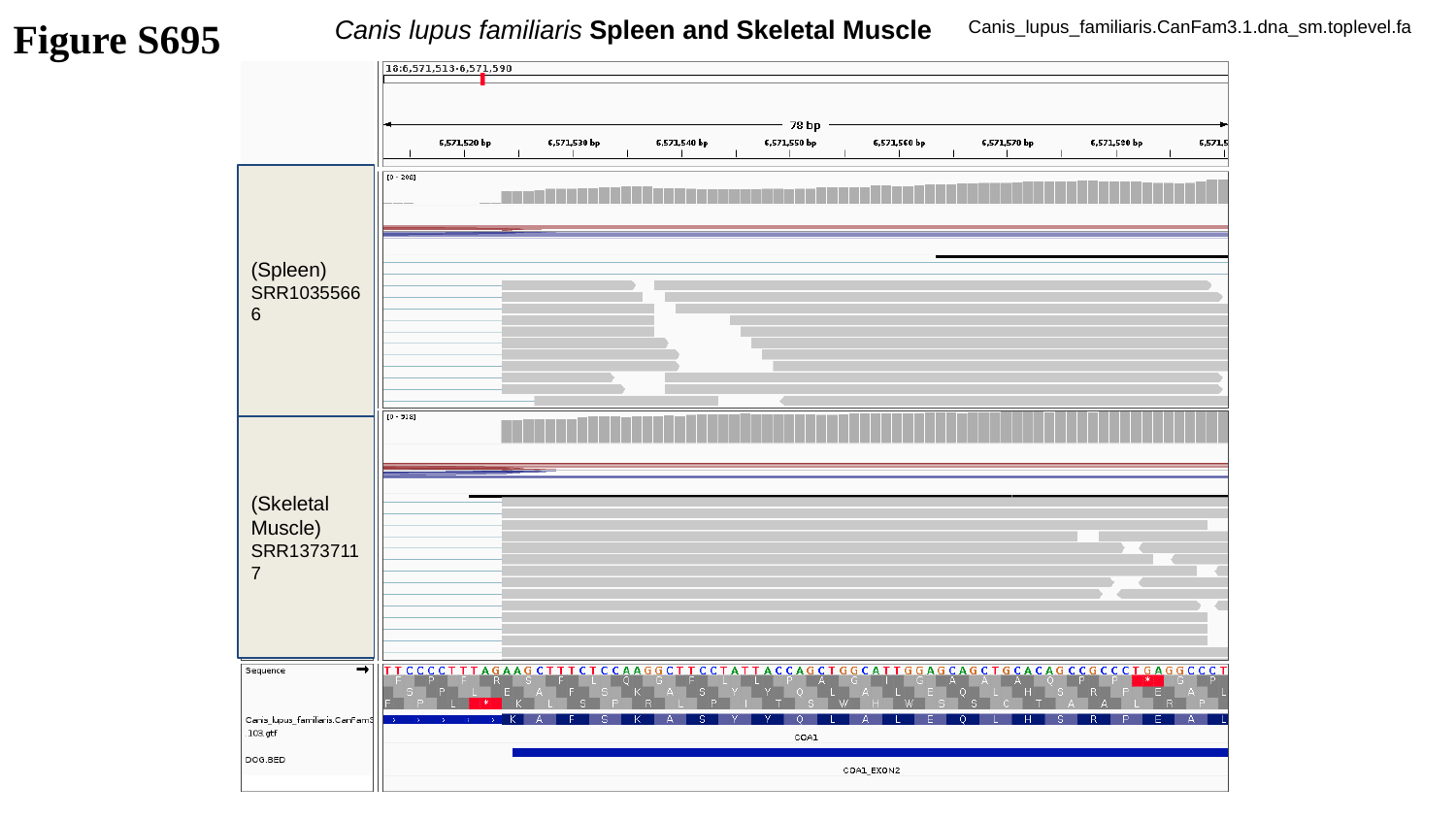

Figure S695
Canis lupus familiaris Spleen and Skeletal Muscle
Canis_lupus_familiaris.CanFam3.1.dna_sm.toplevel.fa
(Spleen)
SRR10355666
(Skeletal Muscle)
SRR13737117

### Slide 696
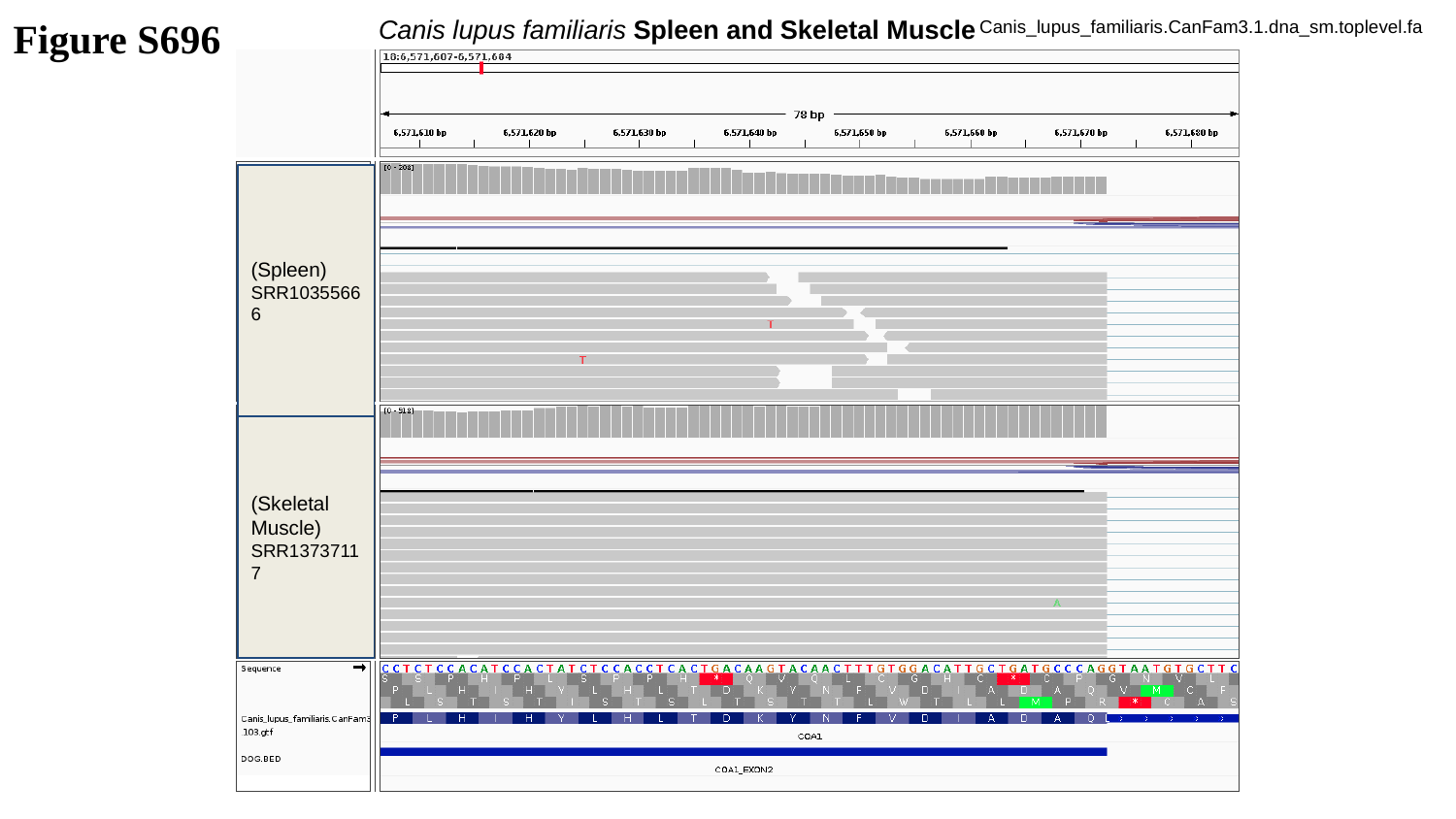

Figure S696
Canis lupus familiaris Spleen and Skeletal Muscle
Canis_lupus_familiaris.CanFam3.1.dna_sm.toplevel.fa
(Spleen)
SRR10355666
(Skeletal Muscle)
SRR13737117

### Slide 697
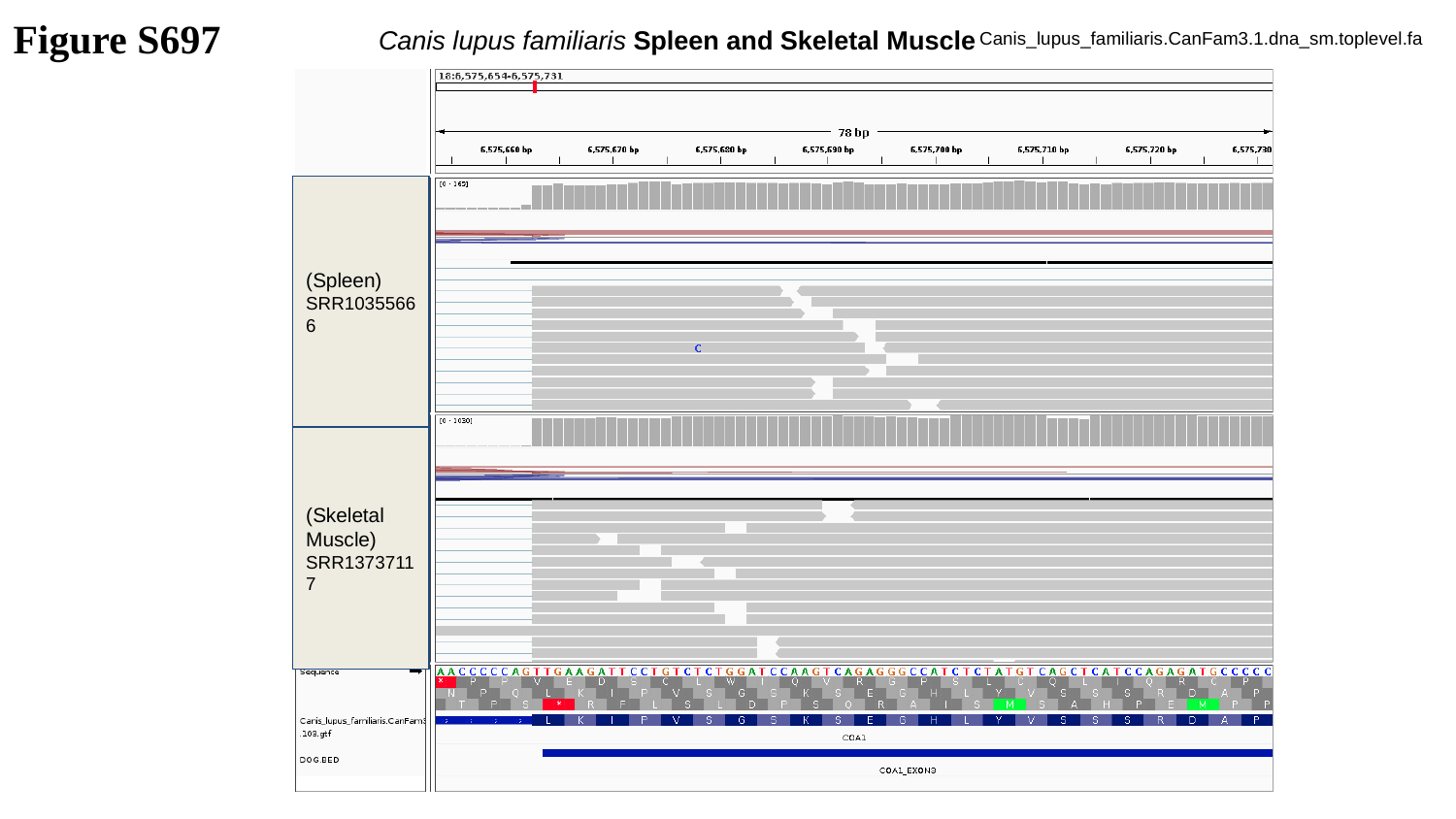

Figure S697
Canis lupus familiaris Spleen and Skeletal Muscle
Canis_lupus_familiaris.CanFam3.1.dna_sm.toplevel.fa
(Spleen)
SRR10355666
(Skeletal Muscle)
SRR13737117

### Slide 698
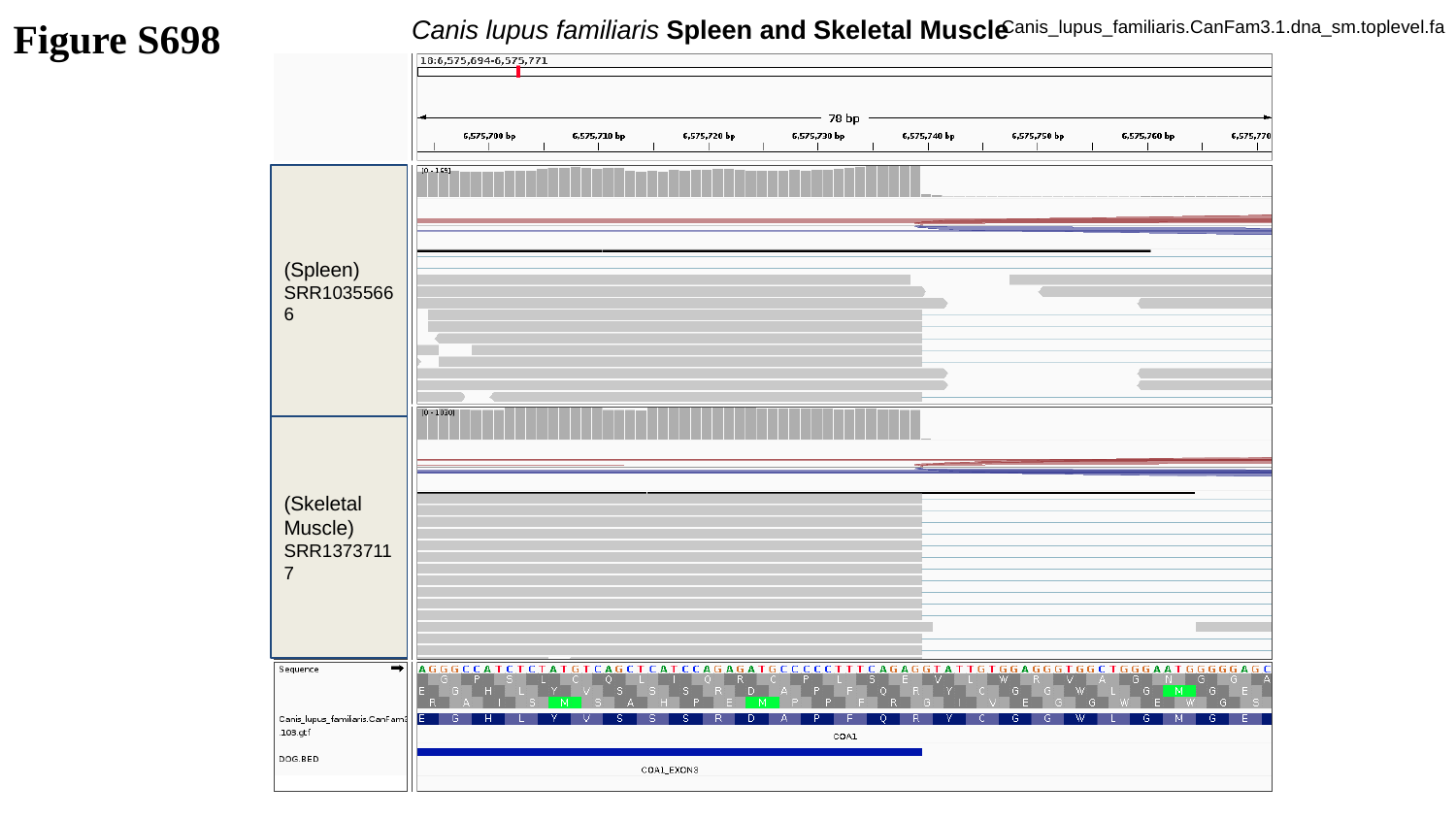

Figure S698
Canis lupus familiaris Spleen and Skeletal Muscle
Canis_lupus_familiaris.CanFam3.1.dna_sm.toplevel.fa
(Spleen)
SRR10355666
(Skeletal Muscle)
SRR13737117

### Slide 699
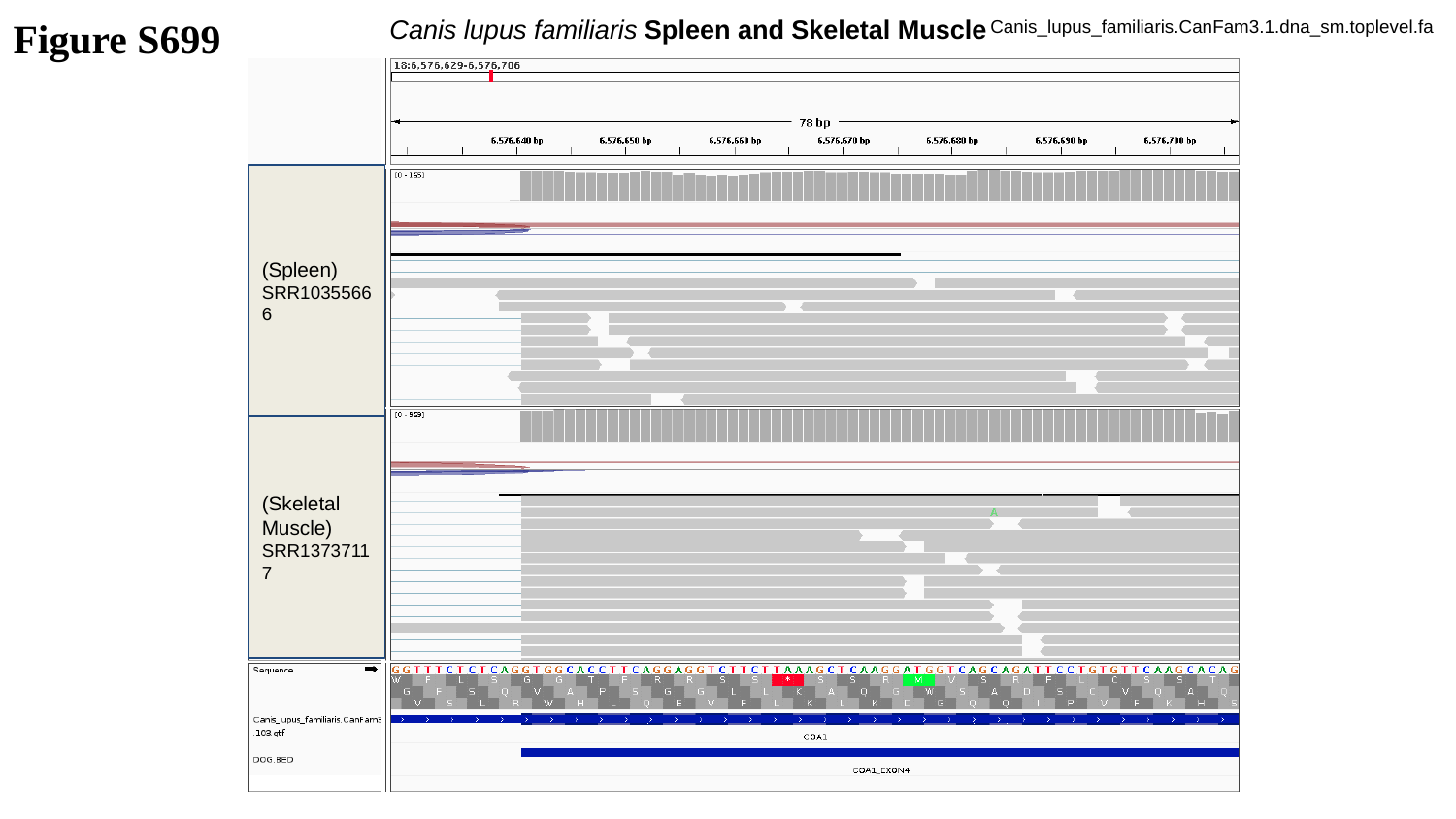

Figure S699
Canis lupus familiaris Spleen and Skeletal Muscle
Canis_lupus_familiaris.CanFam3.1.dna_sm.toplevel.fa
(Spleen)
SRR10355666
(Skeletal Muscle)
SRR13737117

### Slide 700
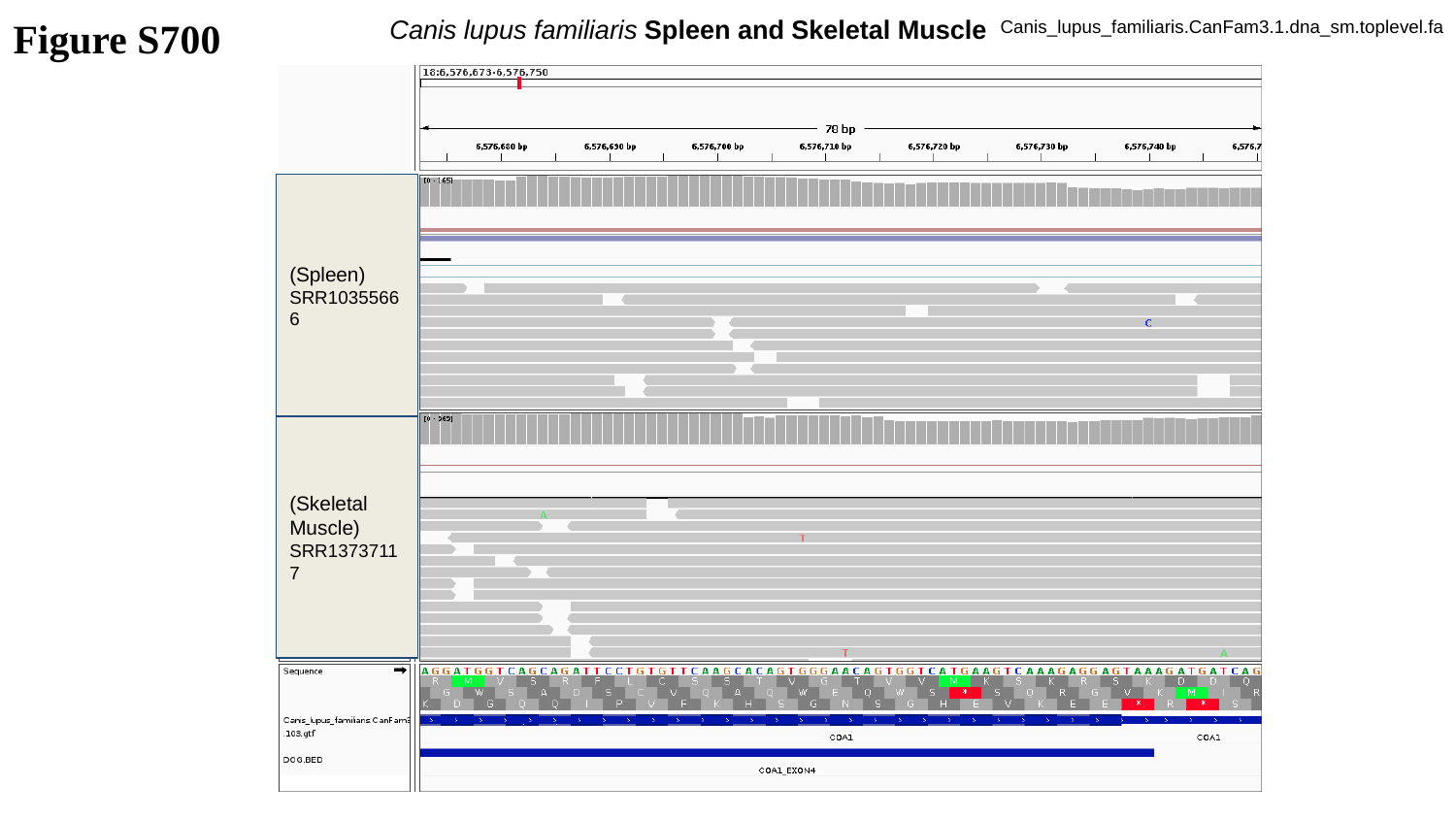

Figure S700
Canis lupus familiaris Spleen and Skeletal Muscle
Canis_lupus_familiaris.CanFam3.1.dna_sm.toplevel.fa
(Spleen)
SRR10355666
(Skeletal Muscle)
SRR13737117

### Slide 701
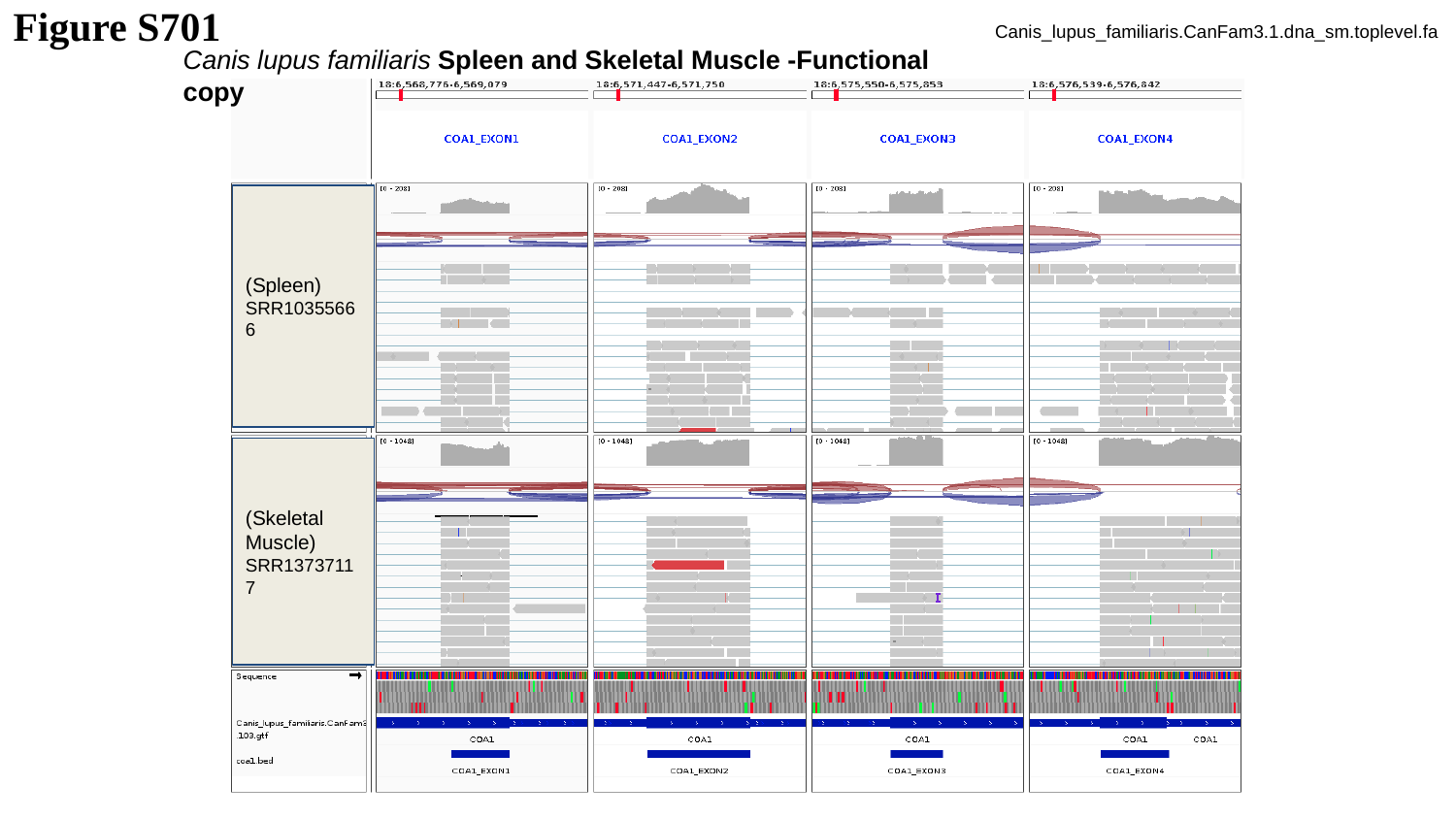

Figure S701
Canis_lupus_familiaris.CanFam3.1.dna_sm.toplevel.fa
Canis lupus familiaris Spleen and Skeletal Muscle -Functional copy
(Spleen)
SRR10355666
(Skeletal Muscle)
SRR13737117

### Slide 702
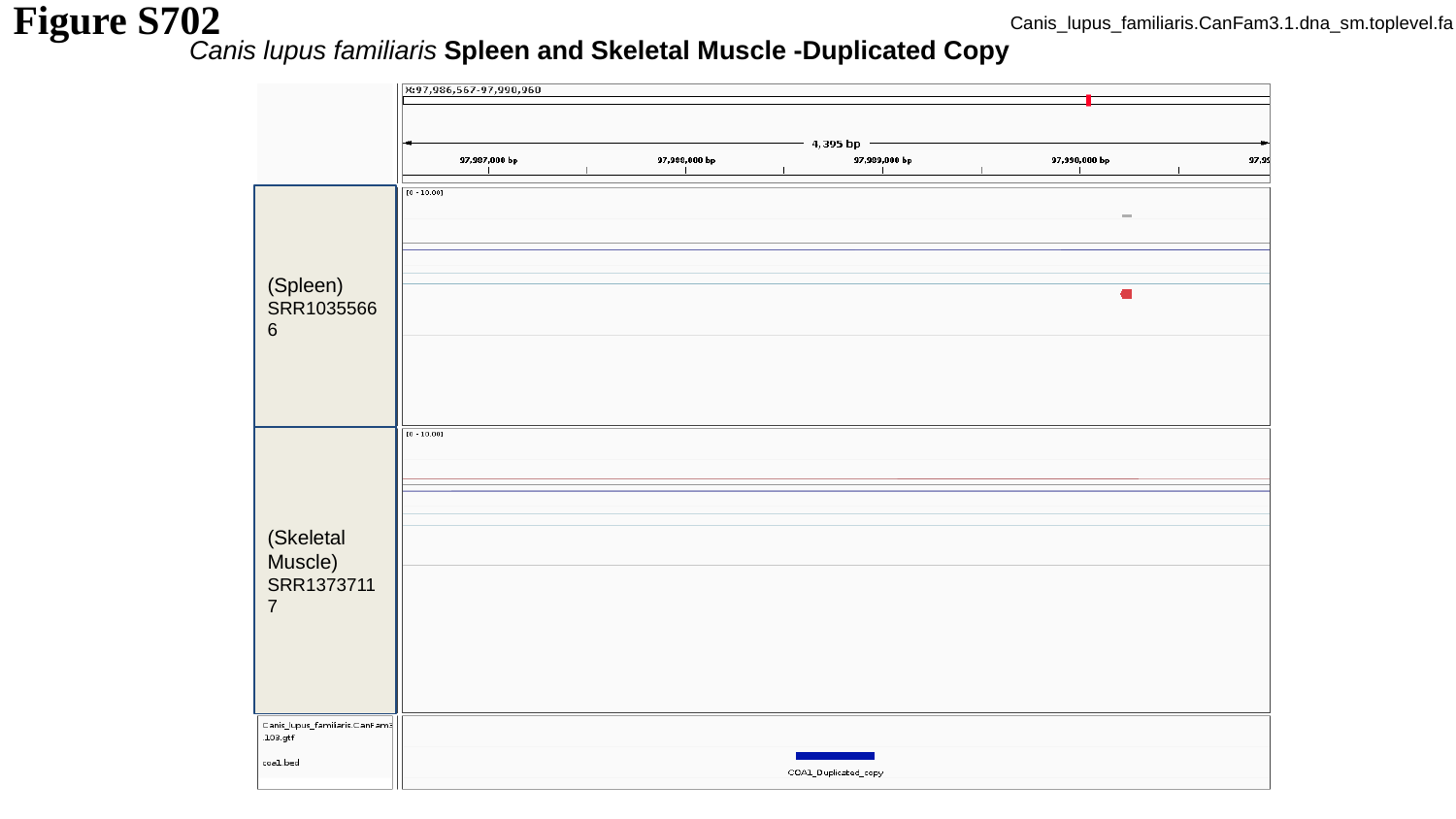

Canis_lupus_familiaris.CanFam3.1.dna_sm.toplevel.fa
Figure S702
Canis lupus familiaris Spleen and Skeletal Muscle -Duplicated Copy
(Spleen)
SRR10355666
(Skeletal Muscle)
SRR13737117

### Slide 703
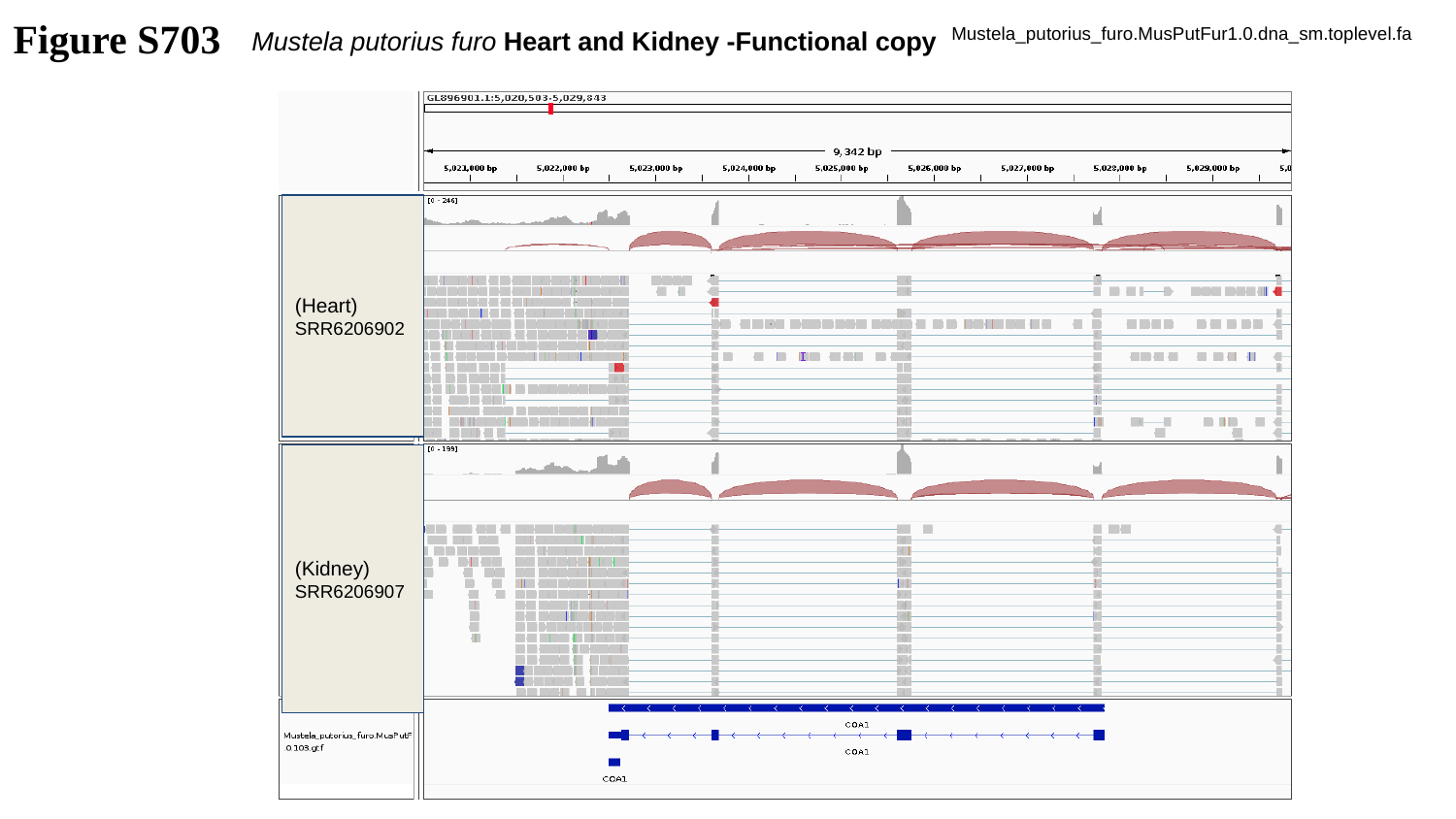

Figure S703
Mustela putorius furo Heart and Kidney -Functional copy
Mustela_putorius_furo.MusPutFur1.0.dna_sm.toplevel.fa
(Heart)
SRR6206902
(Kidney)
SRR6206907

### Slide 704
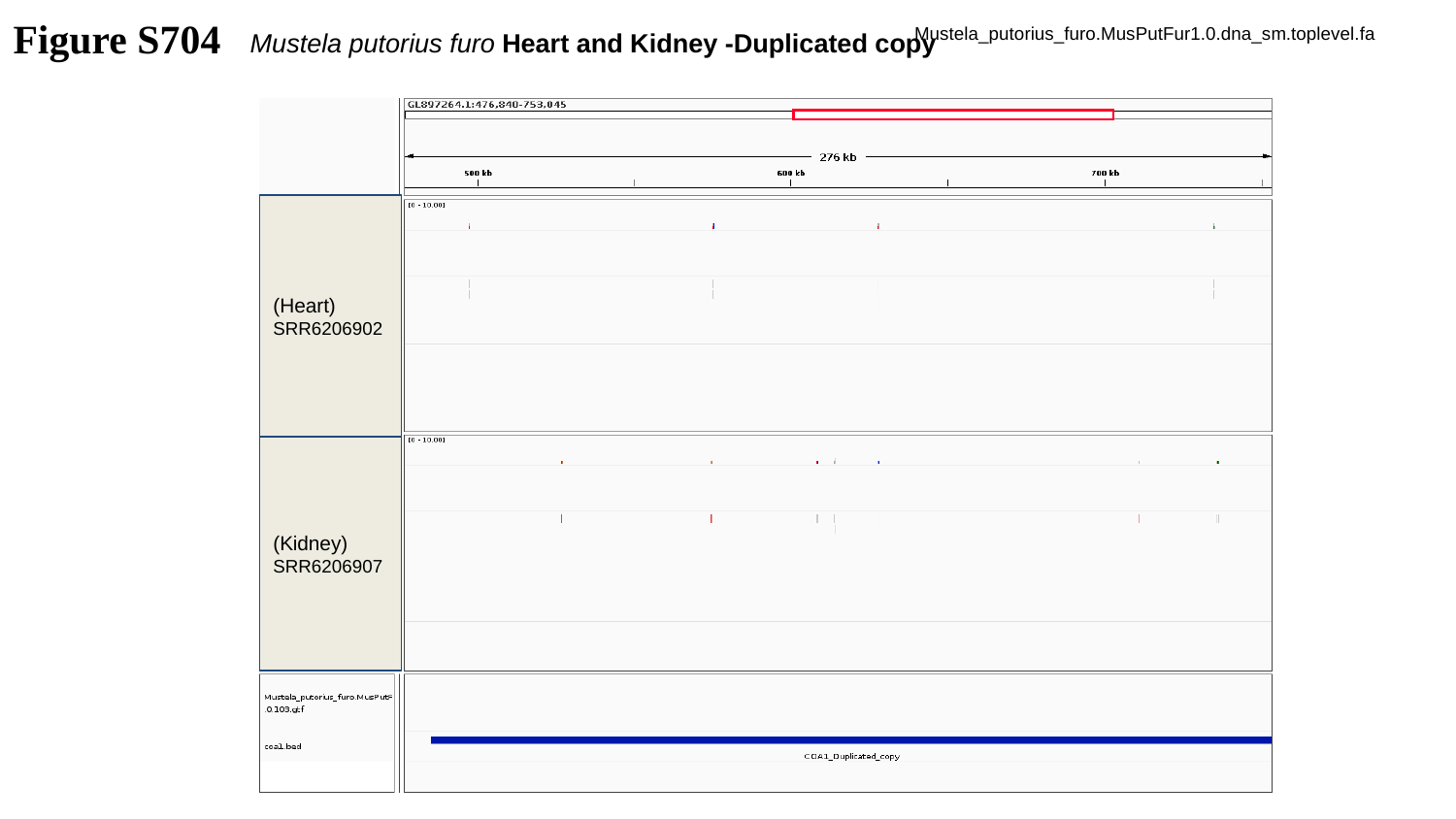

Figure S704
Mustela_putorius_furo.MusPutFur1.0.dna_sm.toplevel.fa
Mustela putorius furo Heart and Kidney -Duplicated copy
(Heart)
SRR6206902
(Kidney)
SRR6206907

### Slide 705
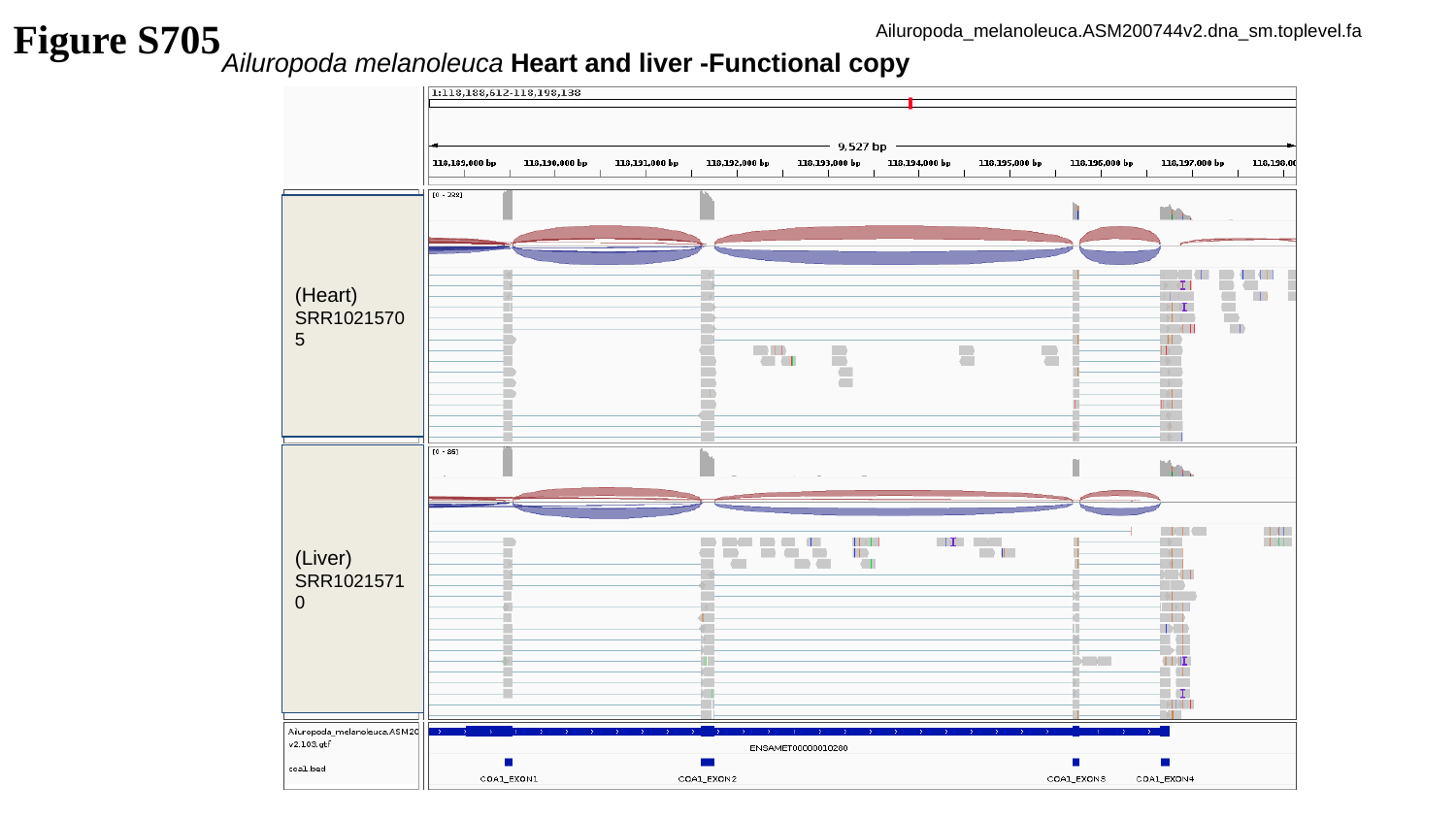

Figure S705
Ailuropoda_melanoleuca.ASM200744v2.dna_sm.toplevel.fa
Ailuropoda melanoleuca Heart and liver -Functional copy
(Heart)
SRR10215705
(Liver)
SRR10215710

### Slide 706
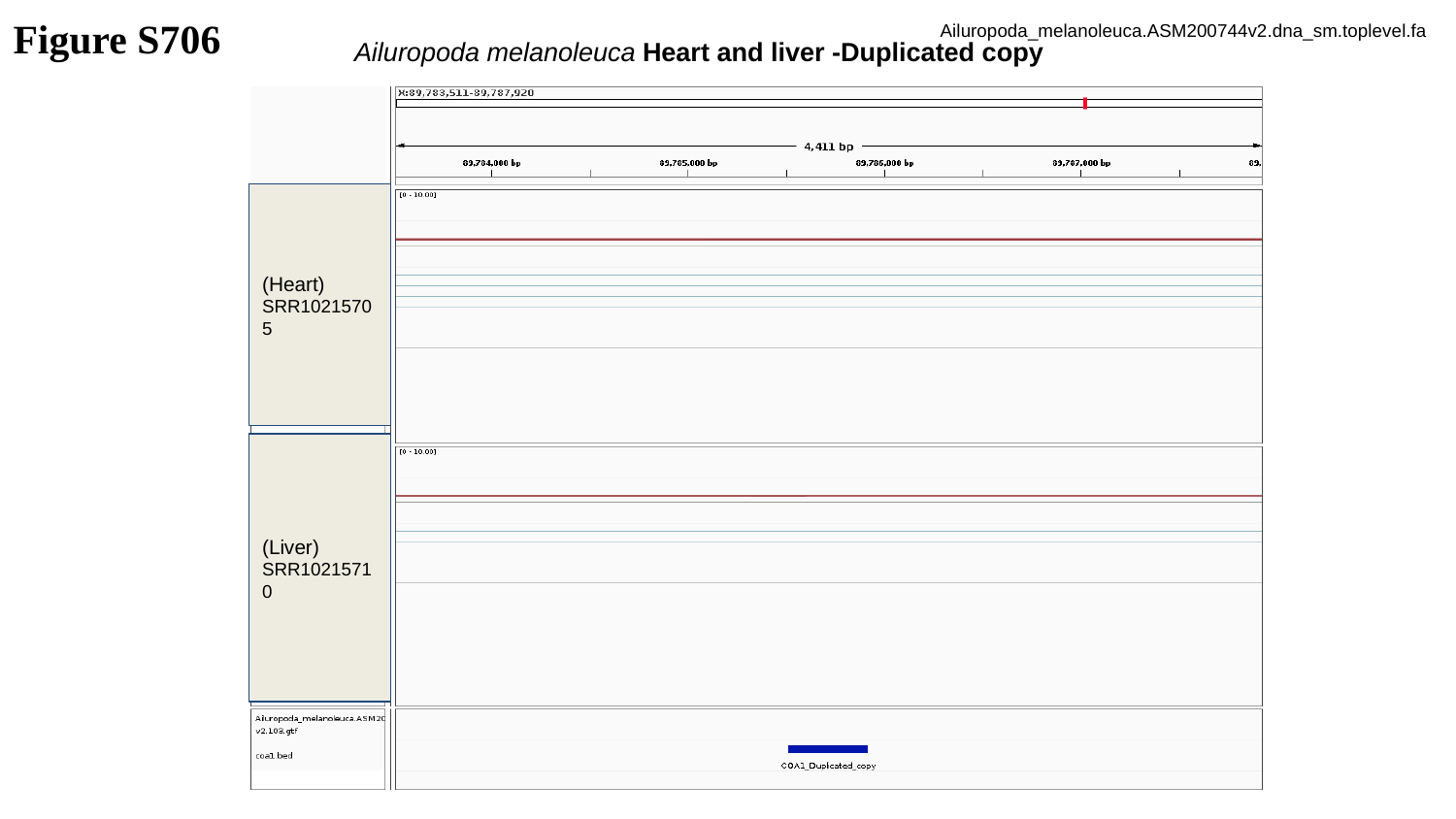

Figure S706
Ailuropoda_melanoleuca.ASM200744v2.dna_sm.toplevel.fa
Ailuropoda melanoleuca Heart and liver -Duplicated copy
(Heart)
SRR10215705
(Liver)
SRR10215710

### Slide 707
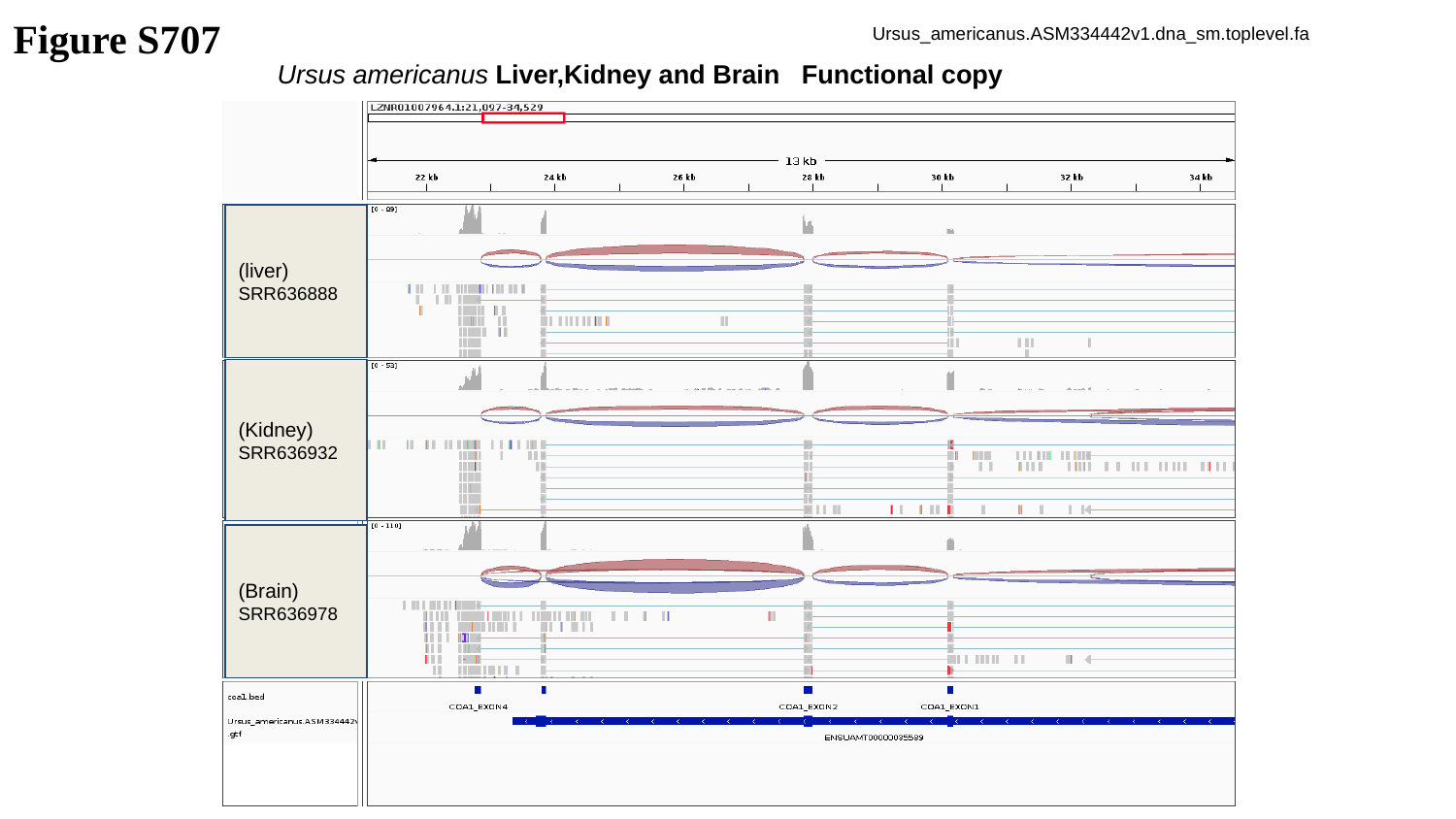

Figure S707
Ursus_americanus.ASM334442v1.dna_sm.toplevel.fa
Ursus americanus Liver,Kidney and Brain Functional copy
(liver)
SRR636888
(Kidney)
SRR636932
(Brain)
SRR636978

### Slide 708
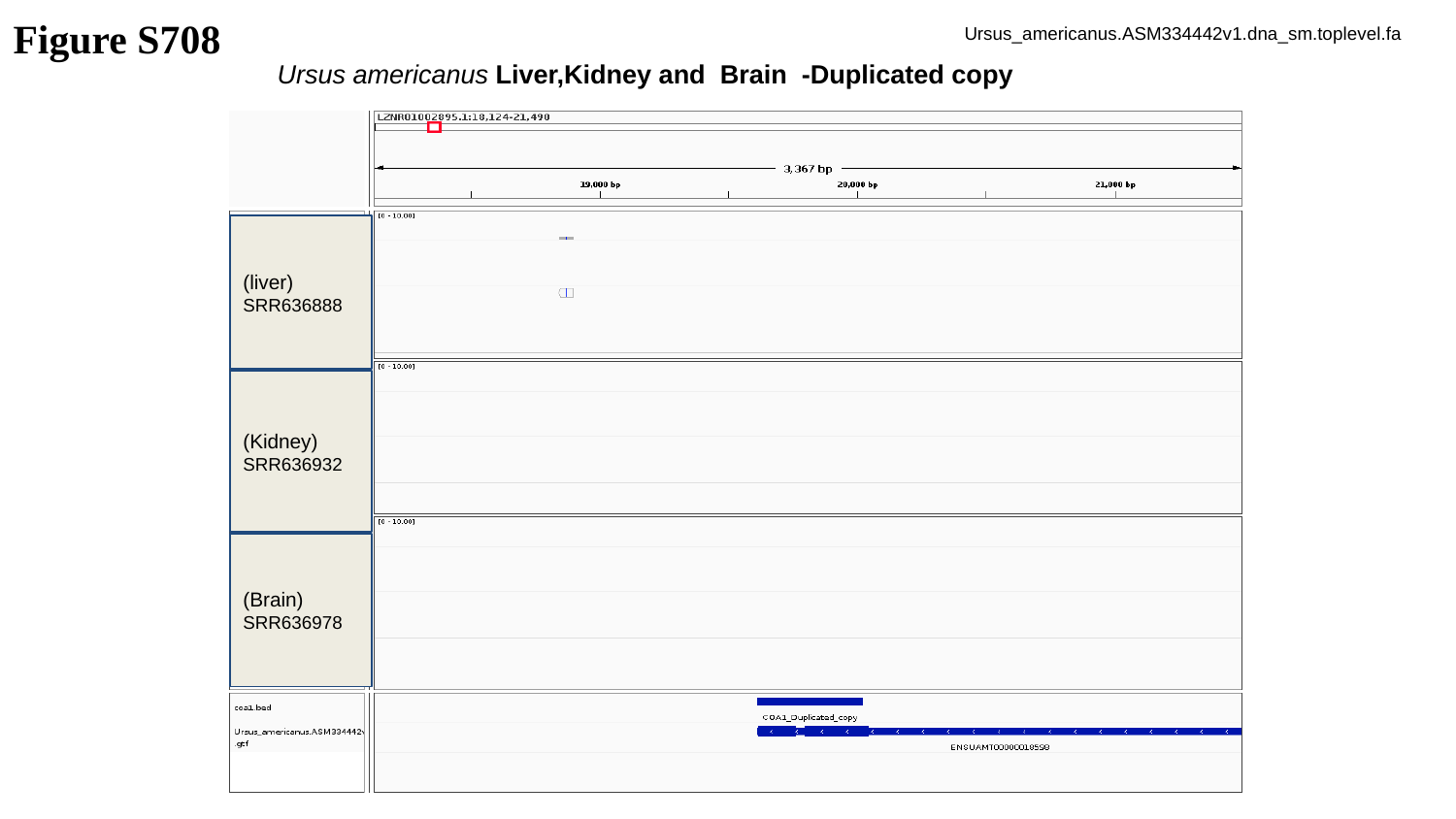

Figure S708
Ursus_americanus.ASM334442v1.dna_sm.toplevel.fa
Ursus americanus Liver,Kidney and Brain -Duplicated copy
(liver)
SRR636888
(Kidney)
SRR636932
(Brain)
SRR636978

### Slide 709
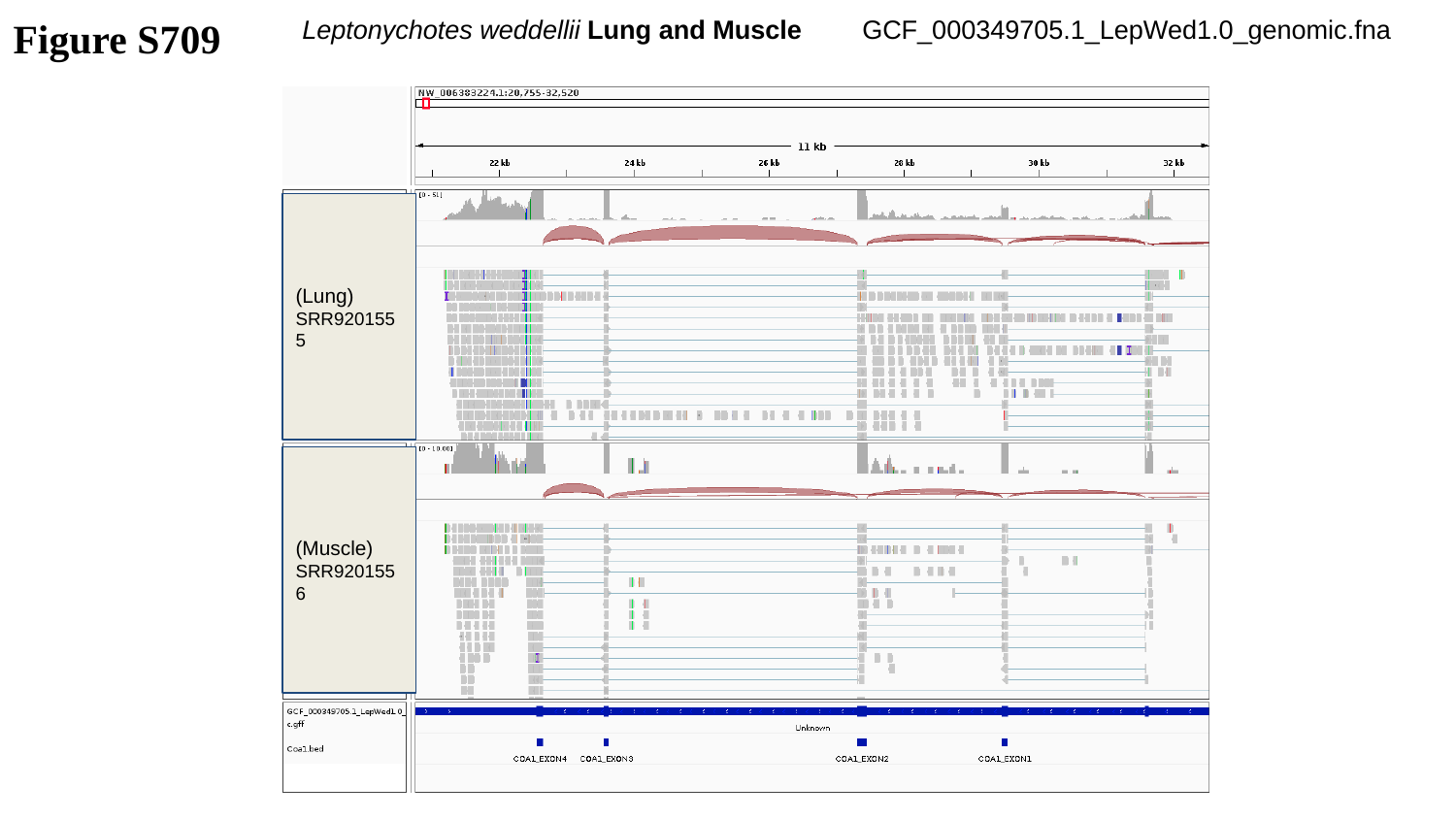

Figure S709
Leptonychotes weddellii Lung and Muscle
GCF_000349705.1_LepWed1.0_genomic.fna
(Lung)
SRR9201555
(Muscle)
SRR9201556

### Slide 710
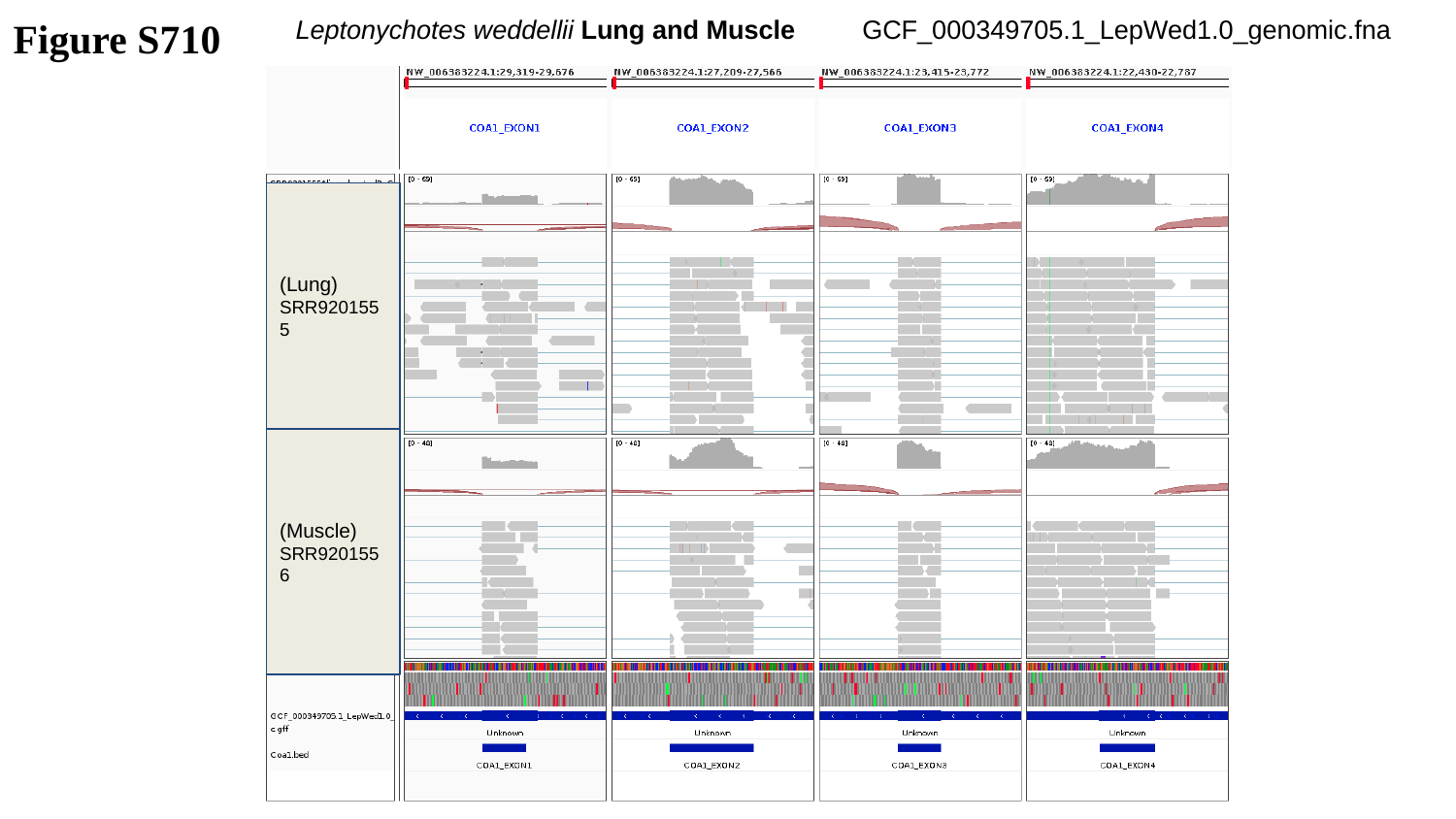

Figure S710
Leptonychotes weddellii Lung and Muscle
GCF_000349705.1_LepWed1.0_genomic.fna
(Lung)
SRR9201555
(Muscle)
SRR9201556

### Slide 711
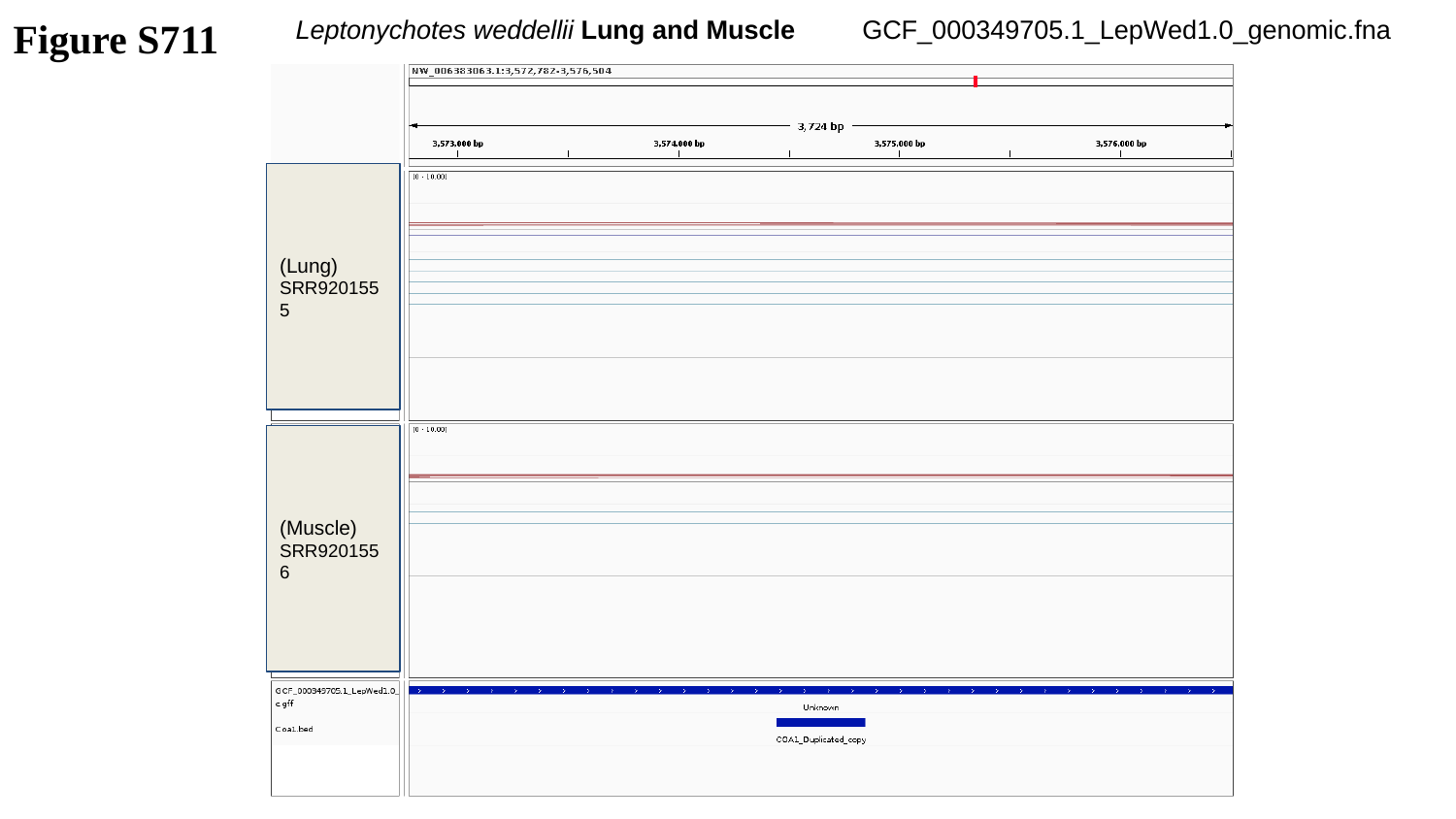

Figure S711
Leptonychotes weddellii Lung and Muscle
GCF_000349705.1_LepWed1.0_genomic.fna
(Lung)
SRR9201555
(Muscle)
SRR9201556

### Slide 712
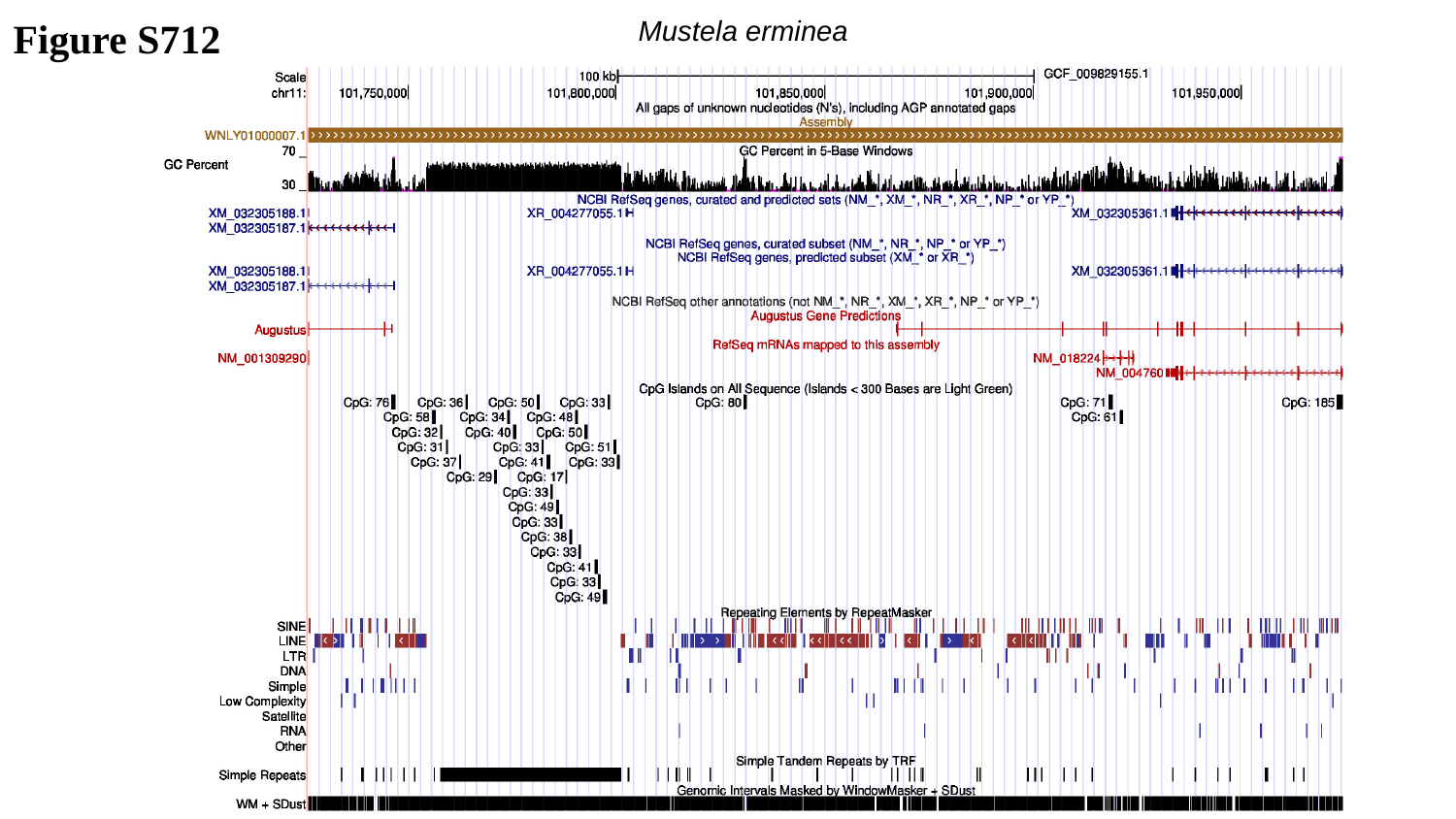

Figure S712
Mustela erminea

### Slide 713
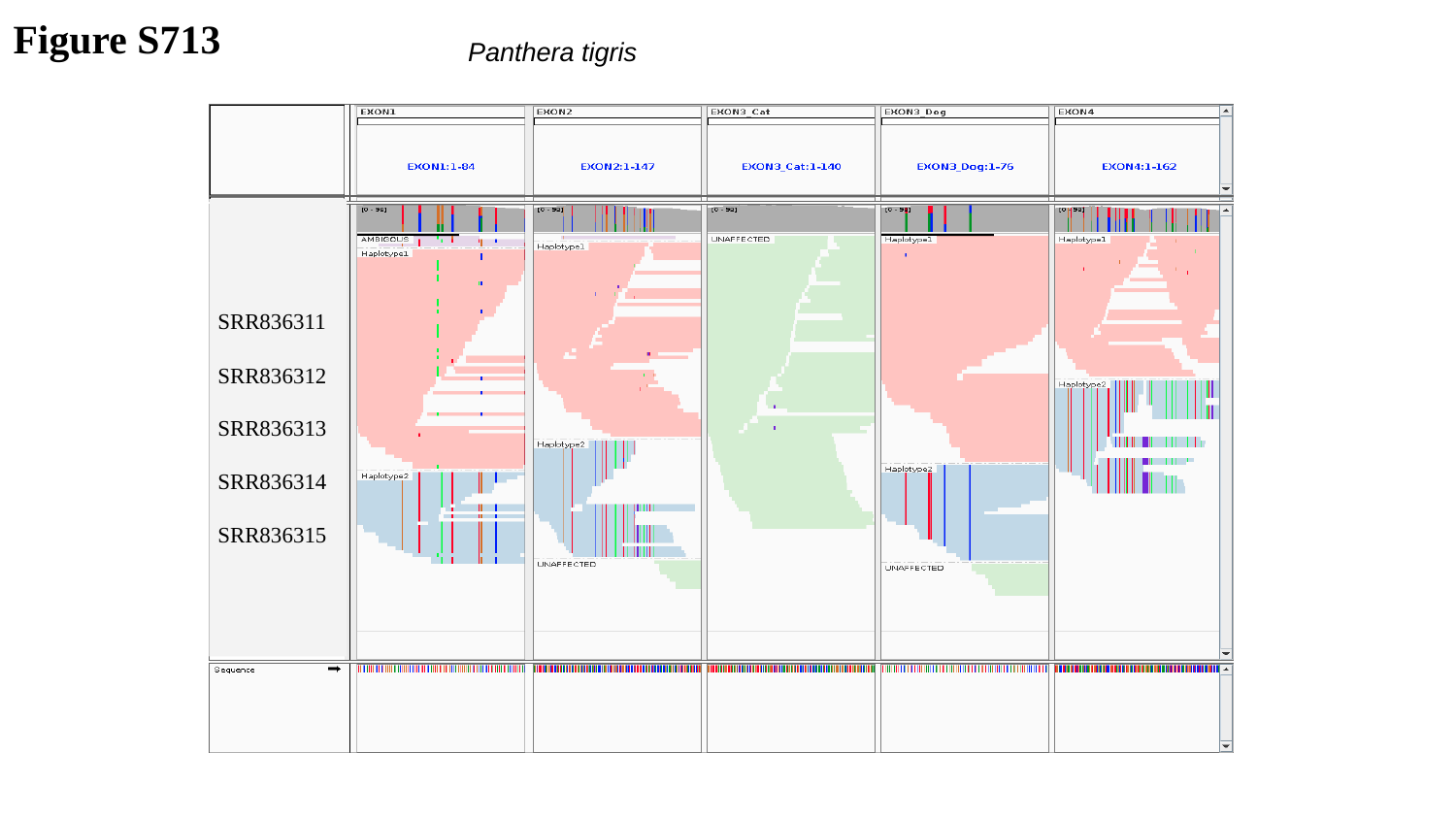

Figure S713
Panthera tigris
SRR836311
SRR836312
SRR836313
SRR836314
SRR836315

### Slide 714
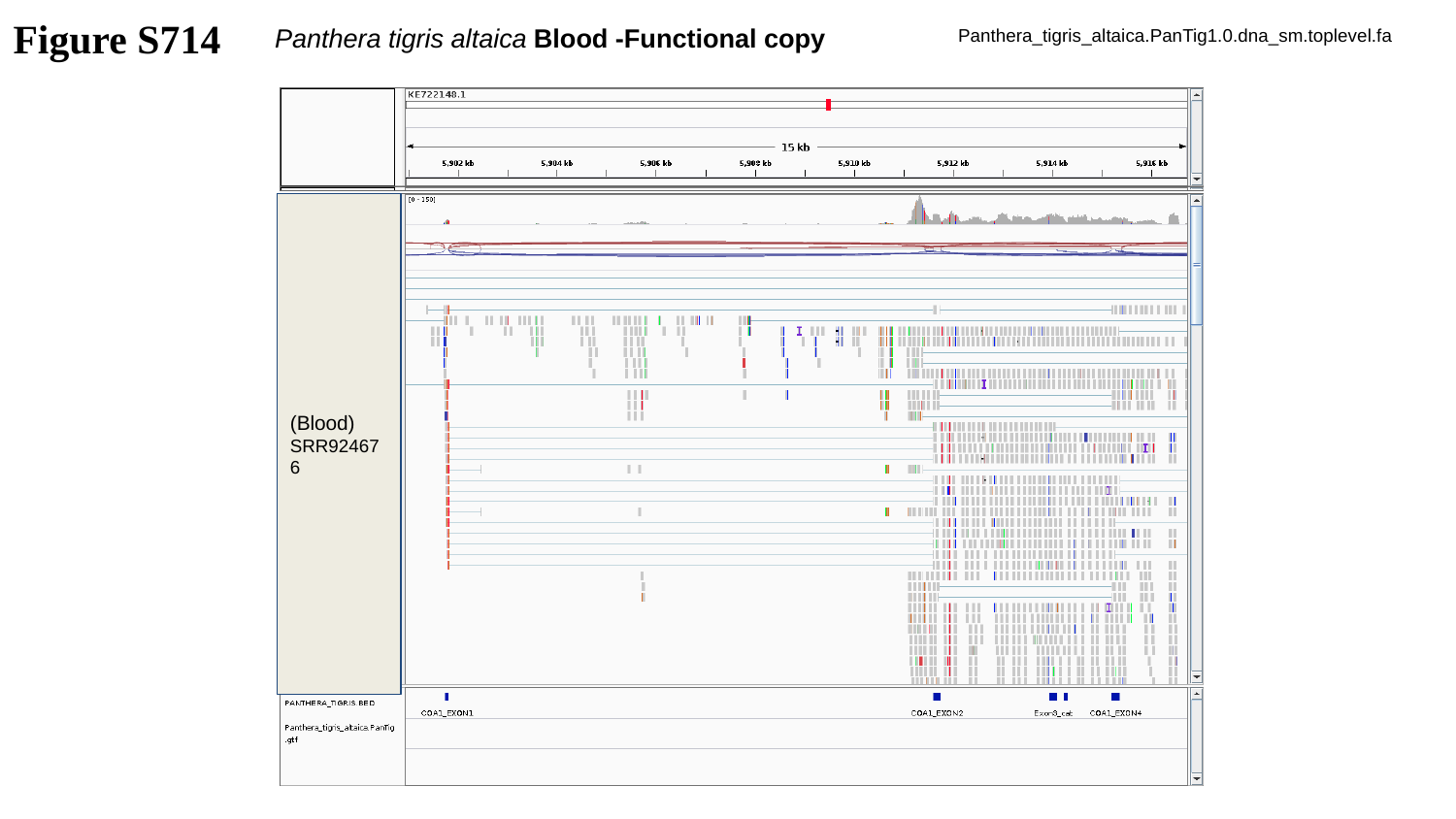

Figure S714
Panthera tigris altaica Blood -Functional copy
Panthera_tigris_altaica.PanTig1.0.dna_sm.toplevel.fa
(Blood)
SRR924676

### Slide 715
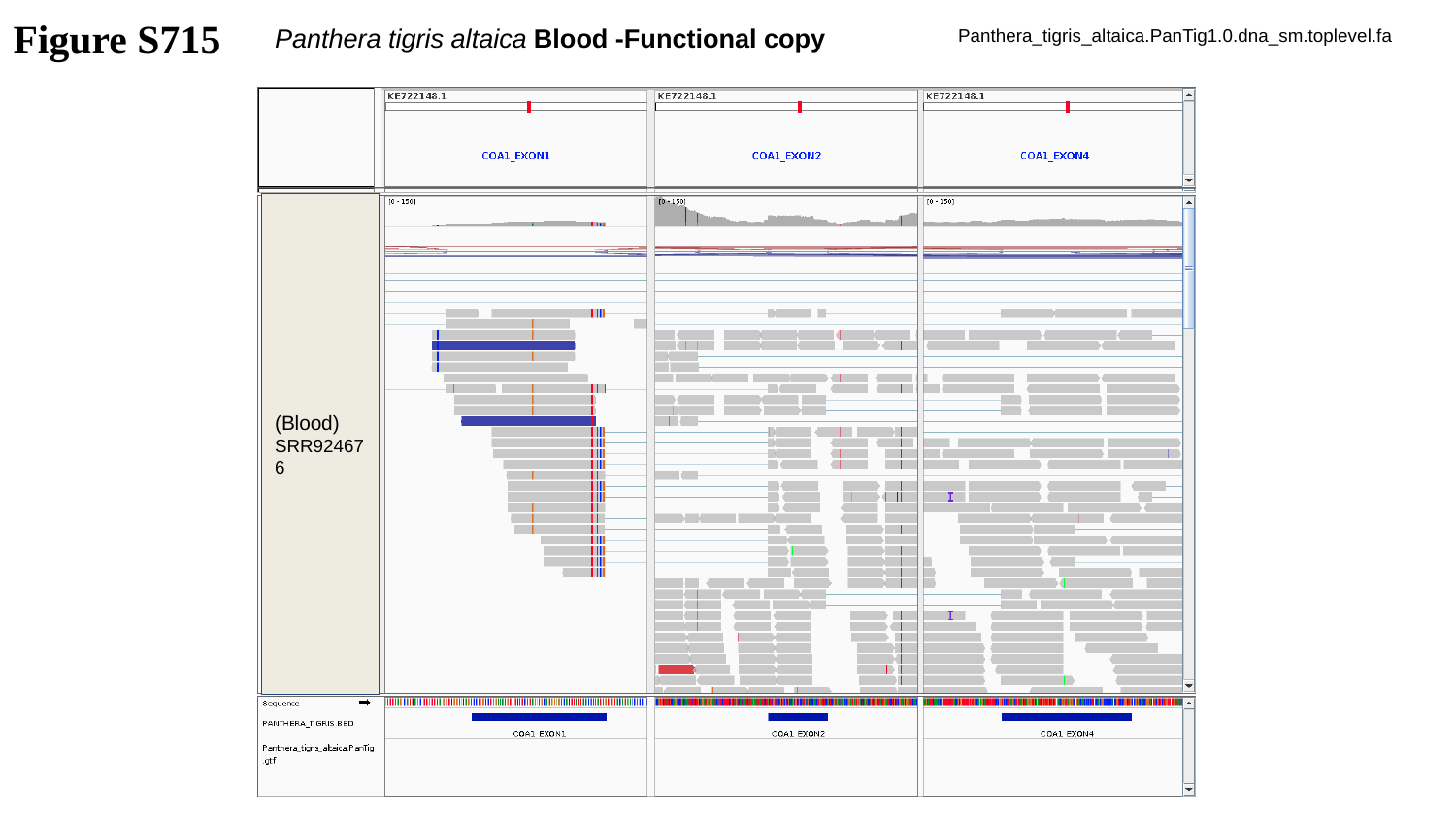

Figure S715
Panthera tigris altaica Blood -Functional copy
Panthera_tigris_altaica.PanTig1.0.dna_sm.toplevel.fa
(Blood)
SRR924676

### Slide 716
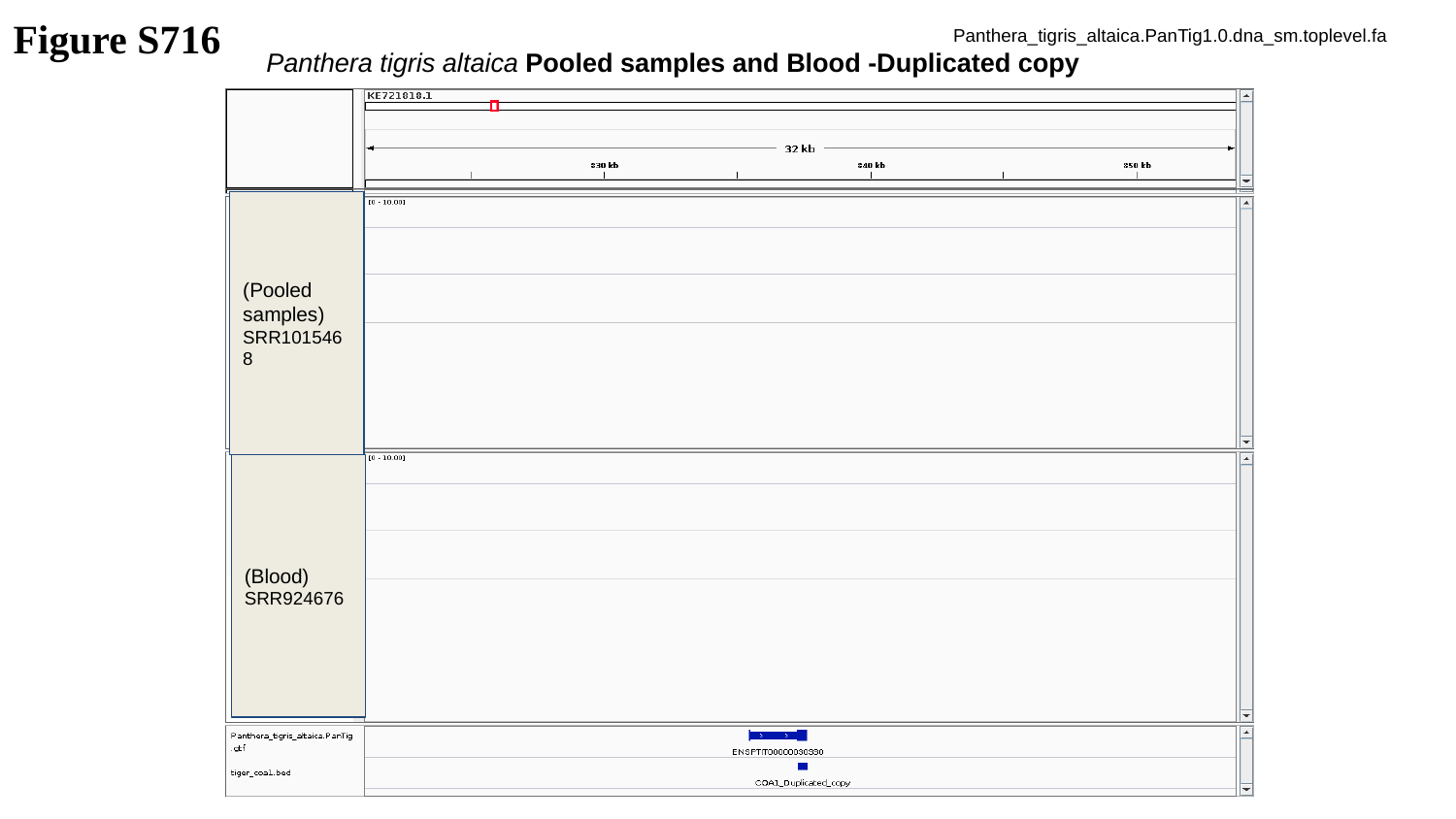

Figure S716
Panthera_tigris_altaica.PanTig1.0.dna_sm.toplevel.fa
Panthera tigris altaica Pooled samples and Blood -Duplicated copy
(Pooled samples)
SRR1015468
(Blood)
SRR924676

### Slide 717
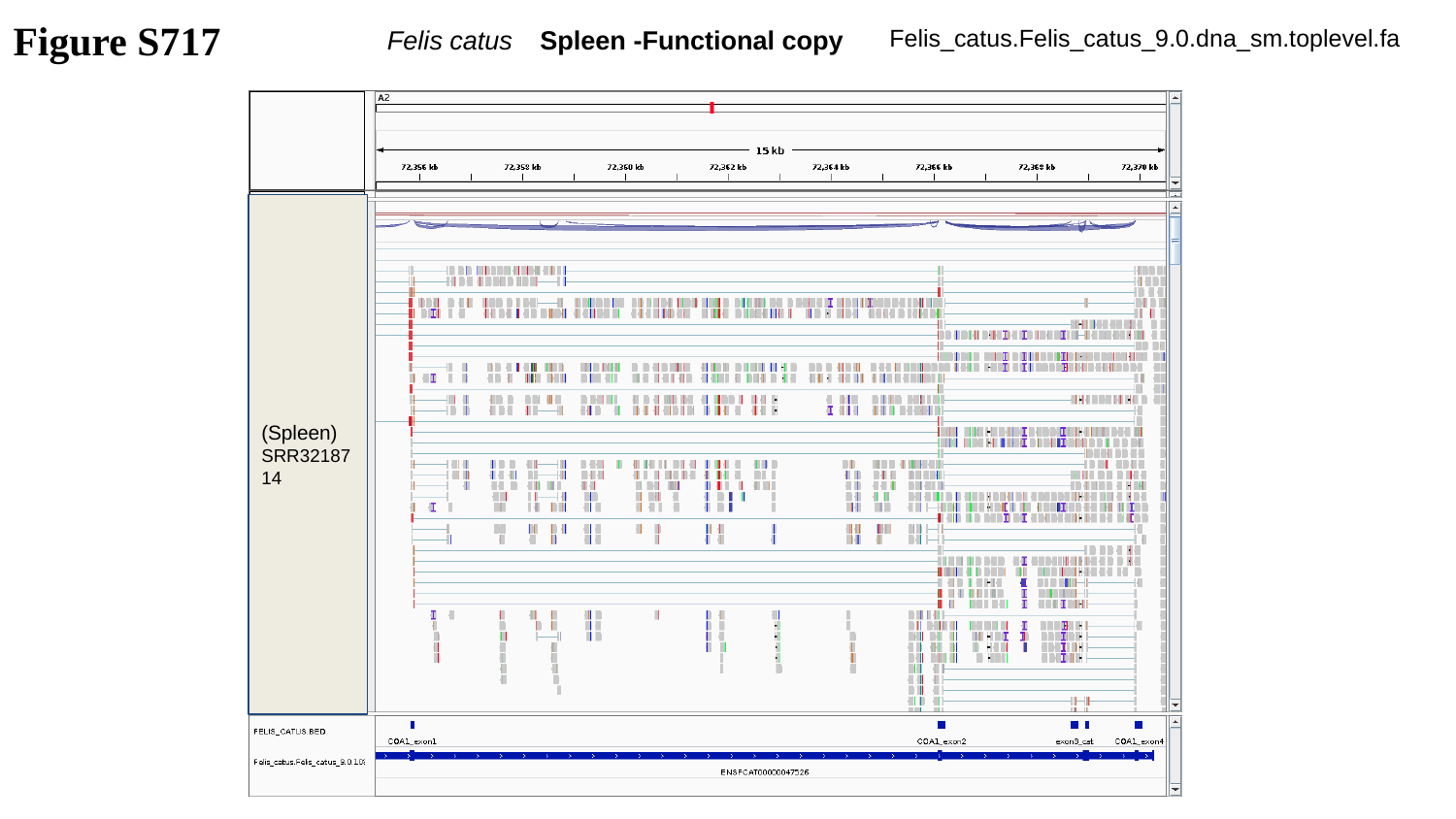

Figure S717
Felis_catus.Felis_catus_9.0.dna_sm.toplevel.fa
Felis catus	 Spleen -Functional copy
(Spleen)
SRR3218714
(Spleen)
SRR3218716

### Slide 718
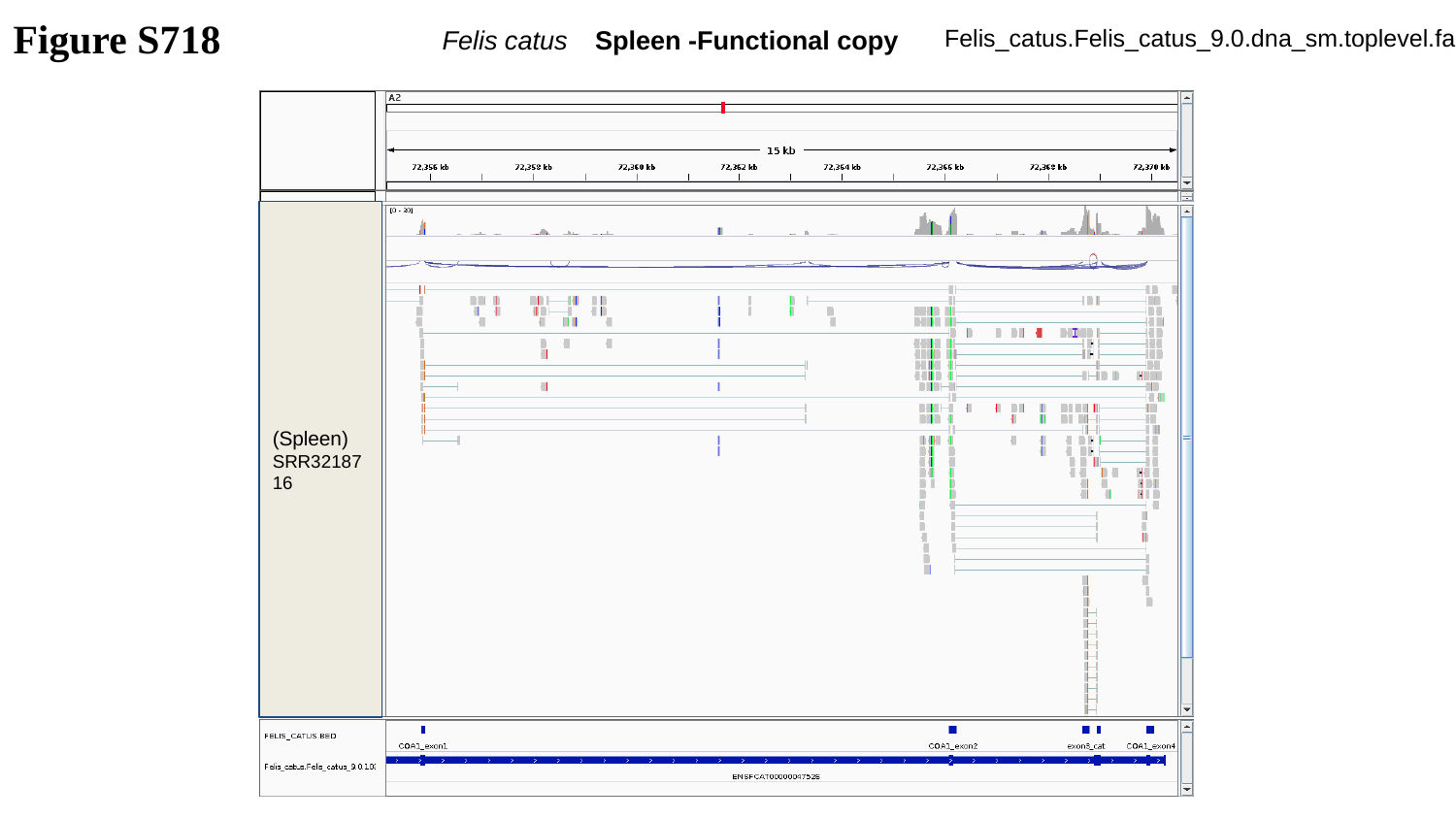

Figure S718
Felis_catus.Felis_catus_9.0.dna_sm.toplevel.fa
Felis catus	 Spleen -Functional copy
(Spleen)
SRR3218716

### Slide 719
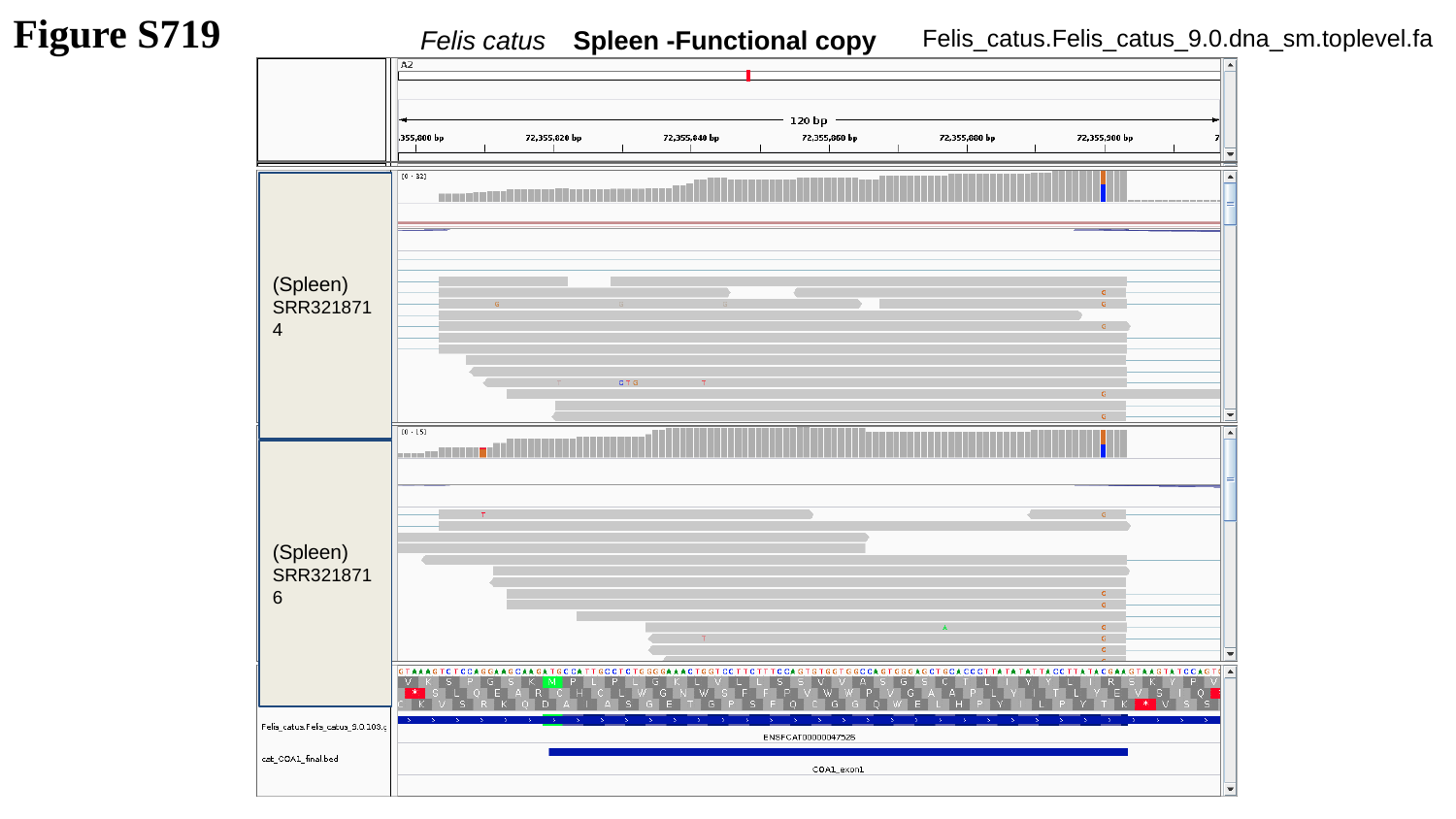

Figure S719
Felis_catus.Felis_catus_9.0.dna_sm.toplevel.fa
Felis catus	 Spleen -Functional copy
(Spleen)
SRR3218714
(Spleen)
SRR3218716

### Slide 720
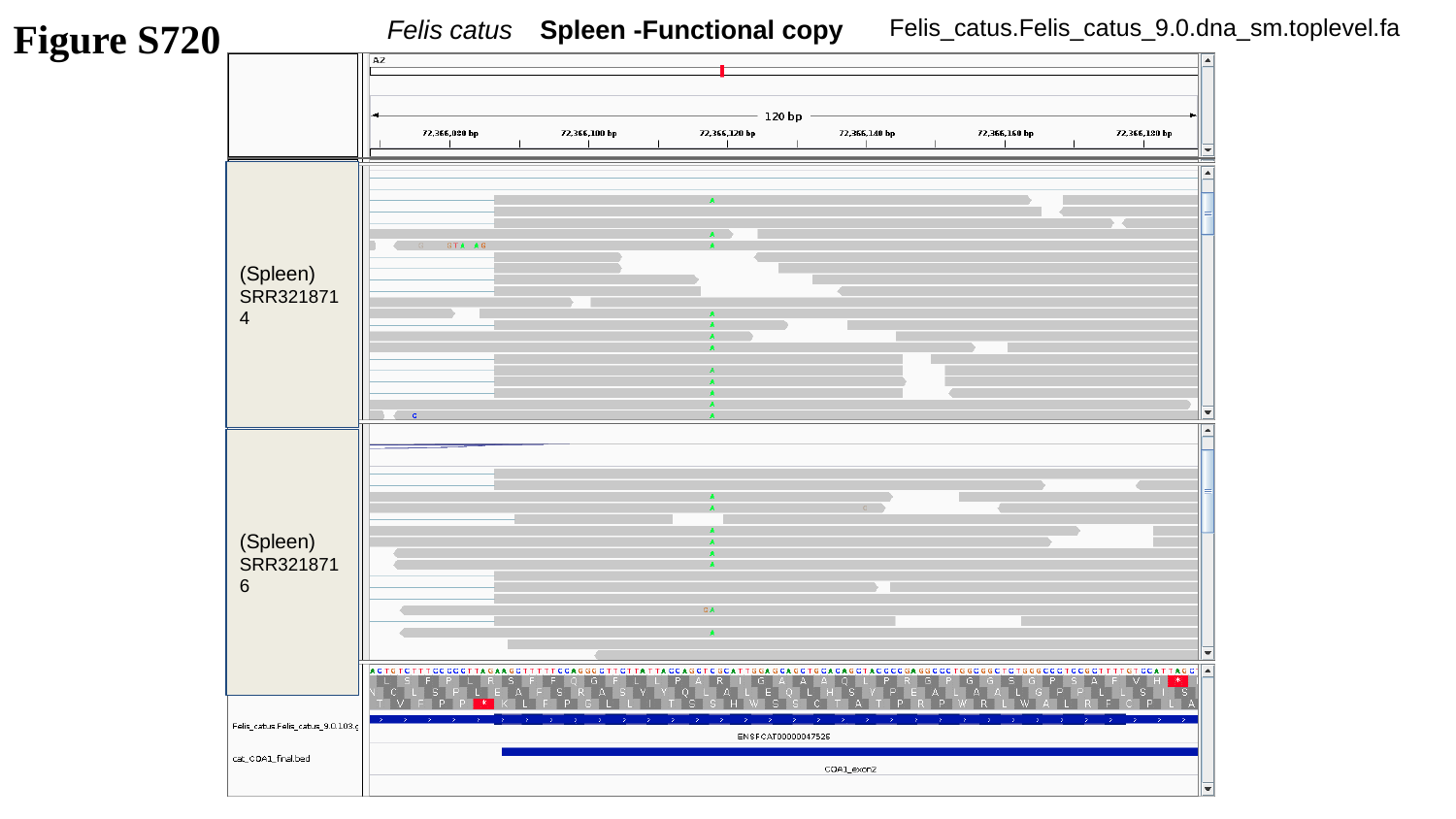

Figure S720
Felis catus	 Spleen -Functional copy
Felis_catus.Felis_catus_9.0.dna_sm.toplevel.fa
(Spleen)
SRR3218714
(Spleen)
SRR3218716

### Slide 721
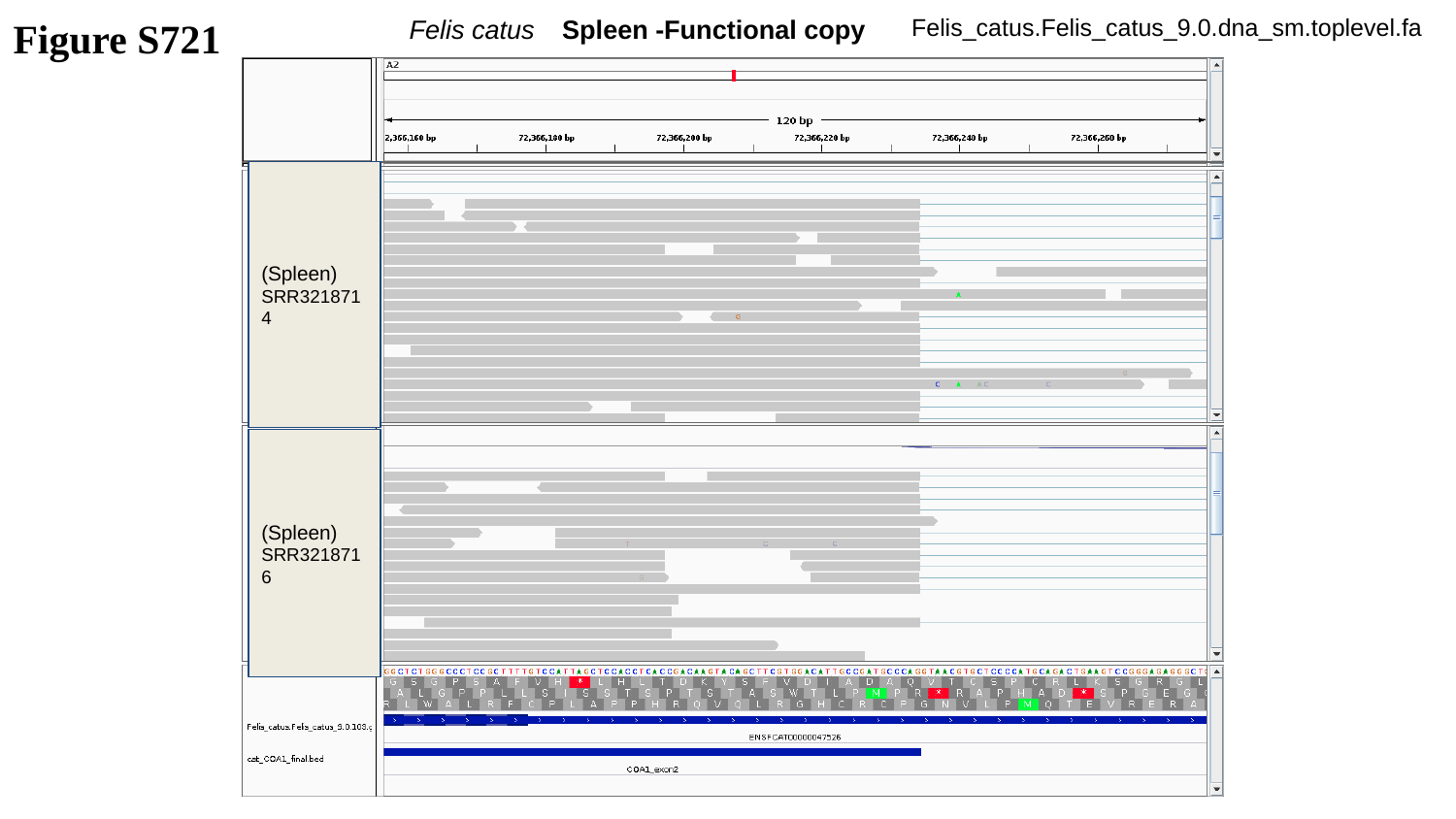

Figure S721
Felis catus	 Spleen -Functional copy
Felis_catus.Felis_catus_9.0.dna_sm.toplevel.fa
(Spleen)
SRR3218714
(Spleen)
SRR3218716

### Slide 722
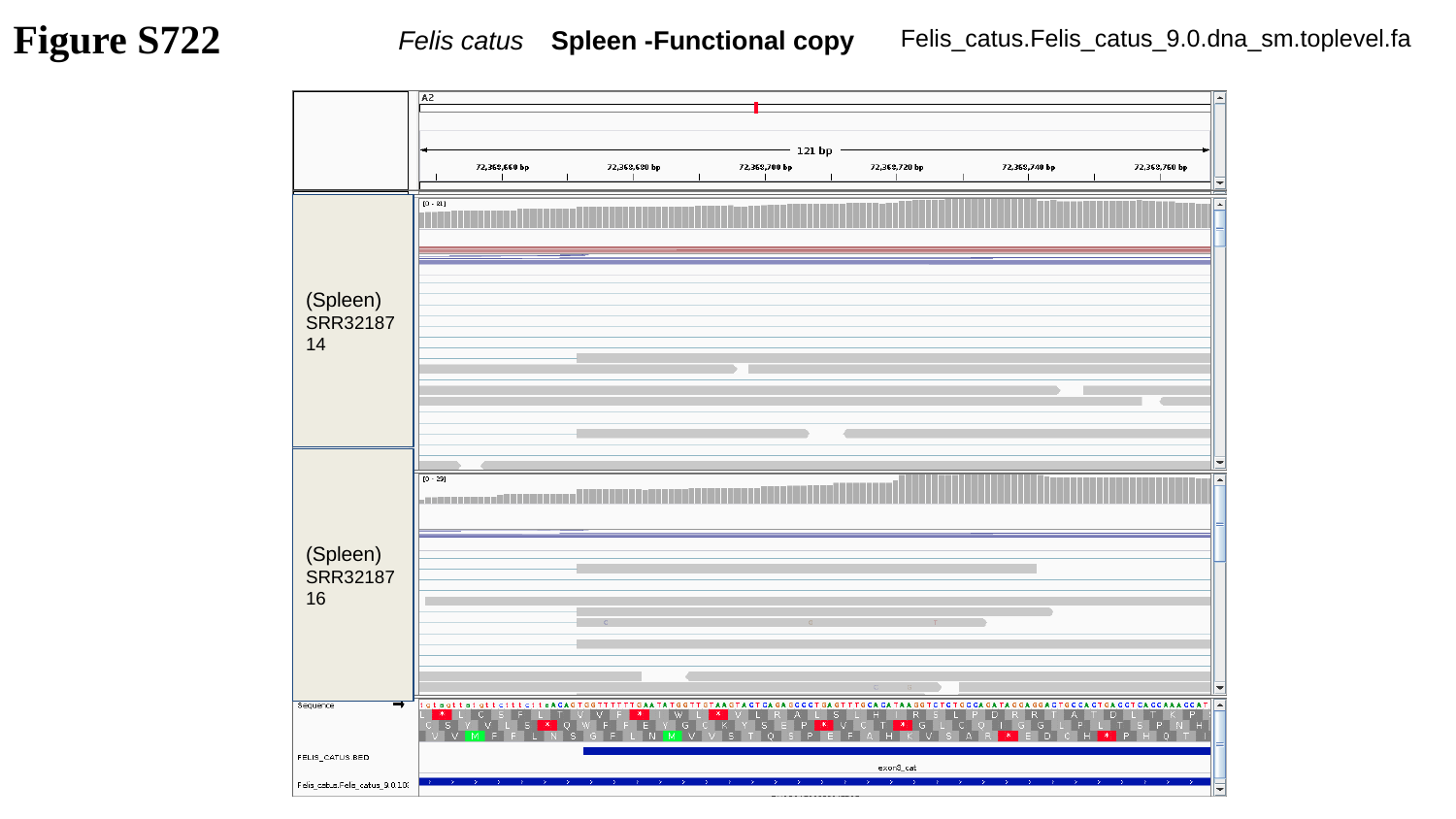

Figure S722
Felis_catus.Felis_catus_9.0.dna_sm.toplevel.fa
Felis catus	 Spleen -Functional copy
(Spleen)
SRR3218714
(Spleen)
SRR3218716

### Slide 723
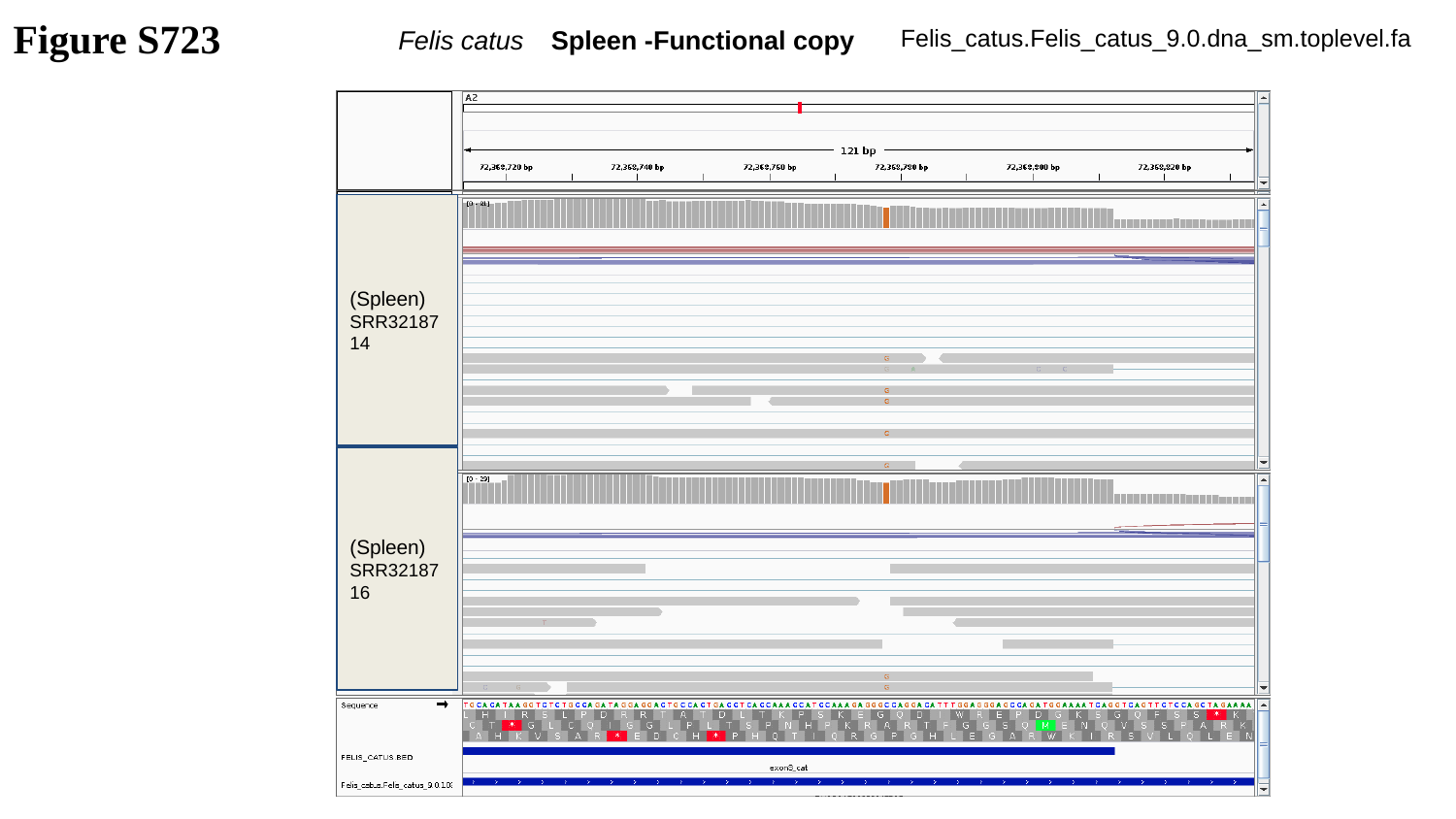

Figure S723
Felis_catus.Felis_catus_9.0.dna_sm.toplevel.fa
Felis catus	 Spleen -Functional copy
(Spleen)
SRR3218714
(Spleen)
SRR3218716

### Slide 724
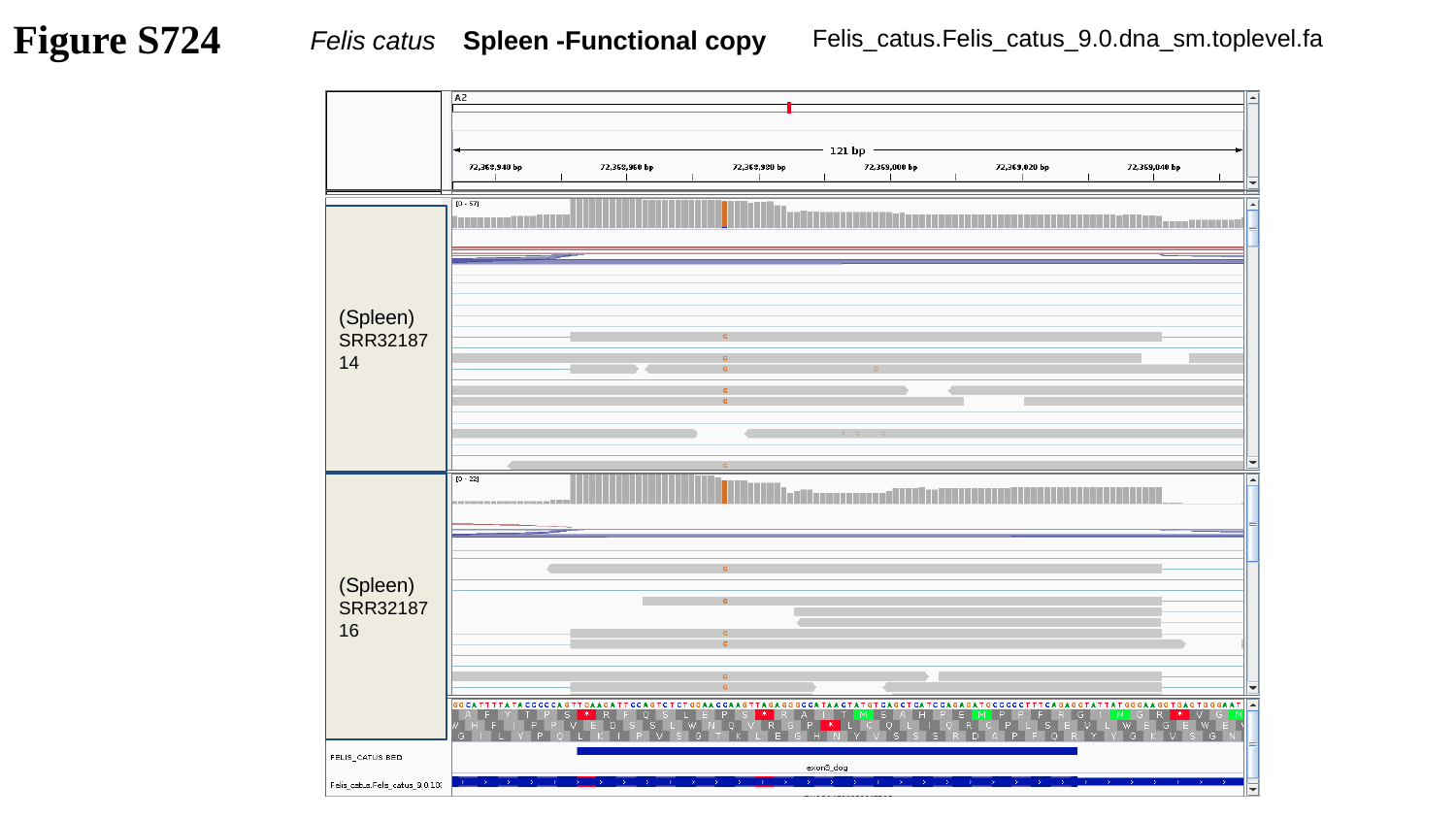

Figure S724
Felis_catus.Felis_catus_9.0.dna_sm.toplevel.fa
Felis catus	 Spleen -Functional copy
(Spleen)
SRR3218714
(Spleen)
SRR3218716

### Slide 725
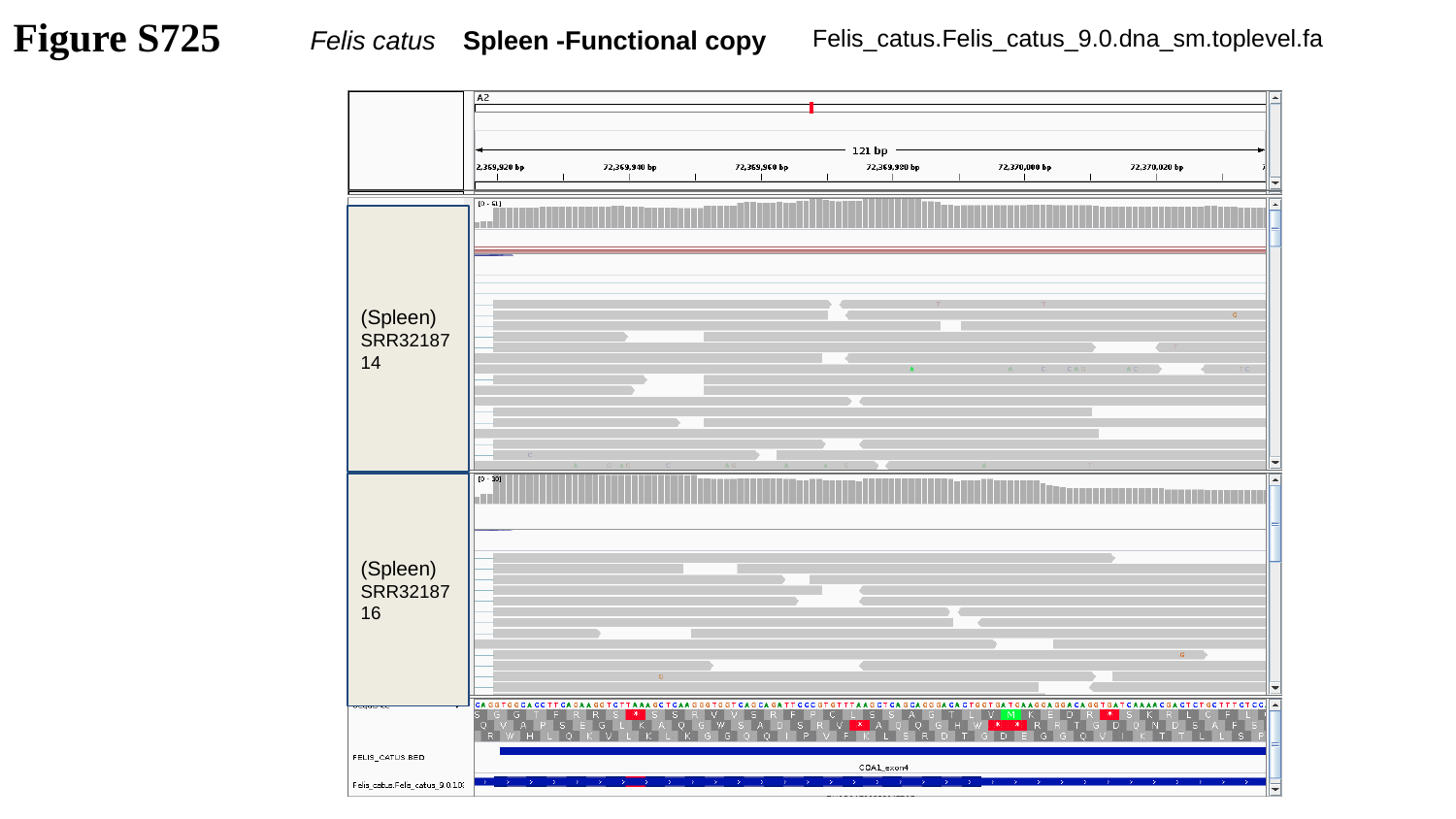

Figure S725
Felis_catus.Felis_catus_9.0.dna_sm.toplevel.fa
Felis catus	 Spleen -Functional copy
(Spleen)
SRR3218714
(Spleen)
SRR3218716

### Slide 726
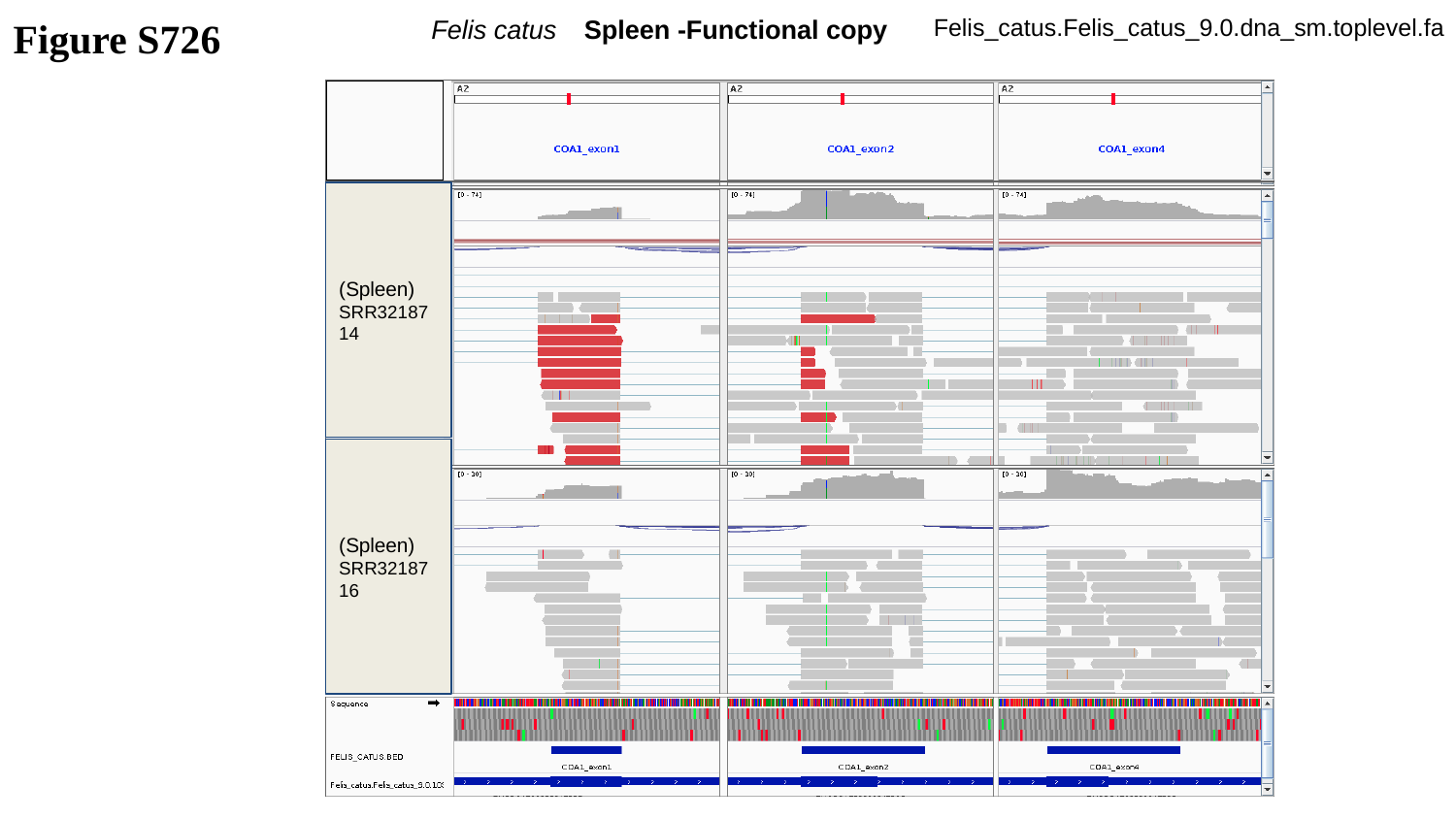

Figure S726
Felis catus	 Spleen -Functional copy
Felis_catus.Felis_catus_9.0.dna_sm.toplevel.fa
(Spleen)
SRR3218714
(Spleen)
SRR3218716

### Slide 727
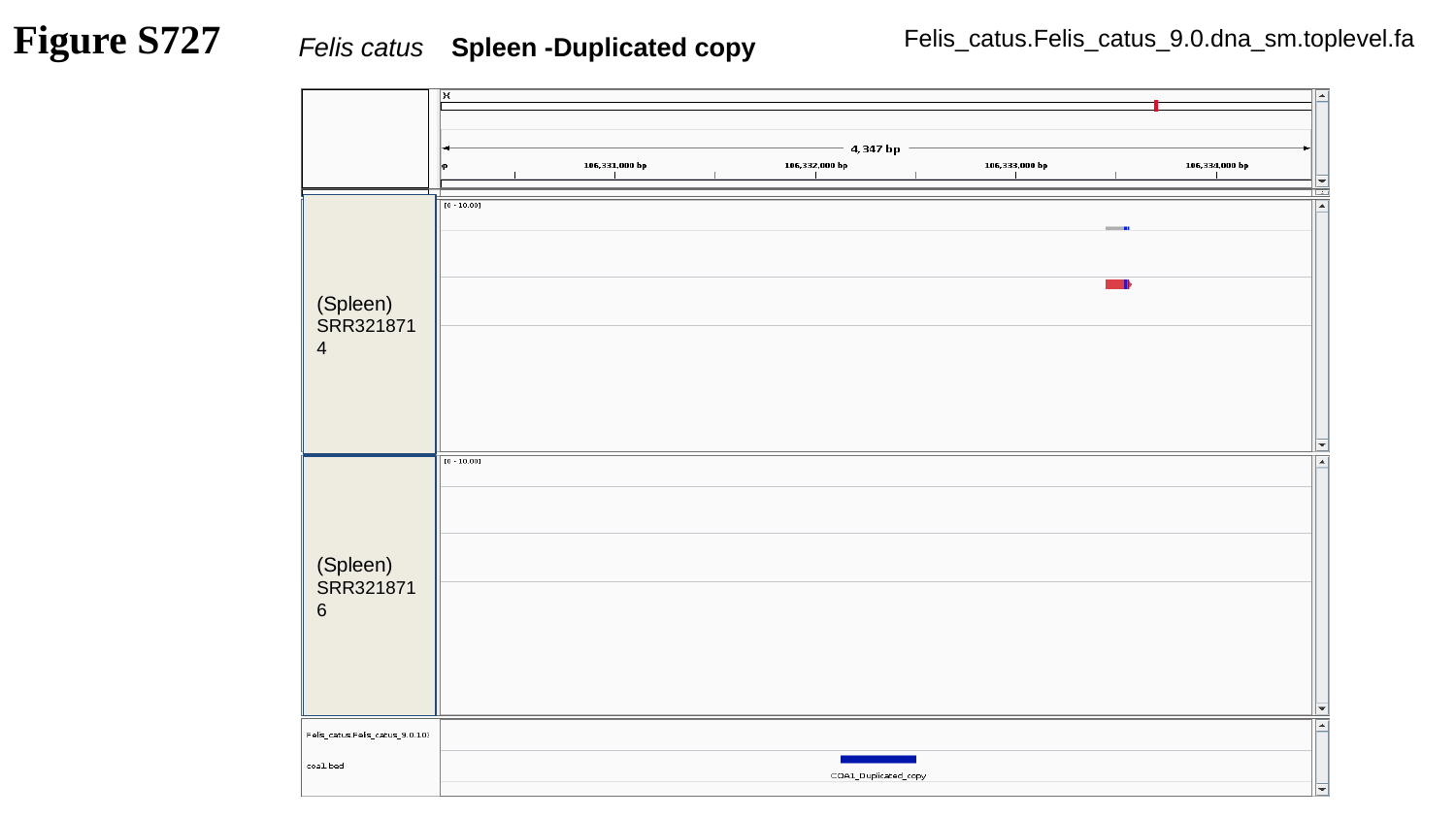

Figure S727
Felis_catus.Felis_catus_9.0.dna_sm.toplevel.fa
Felis catus	 Spleen -Duplicated copy
(Spleen)
SRR3218714
(Spleen)
SRR3218716

### Slide 728
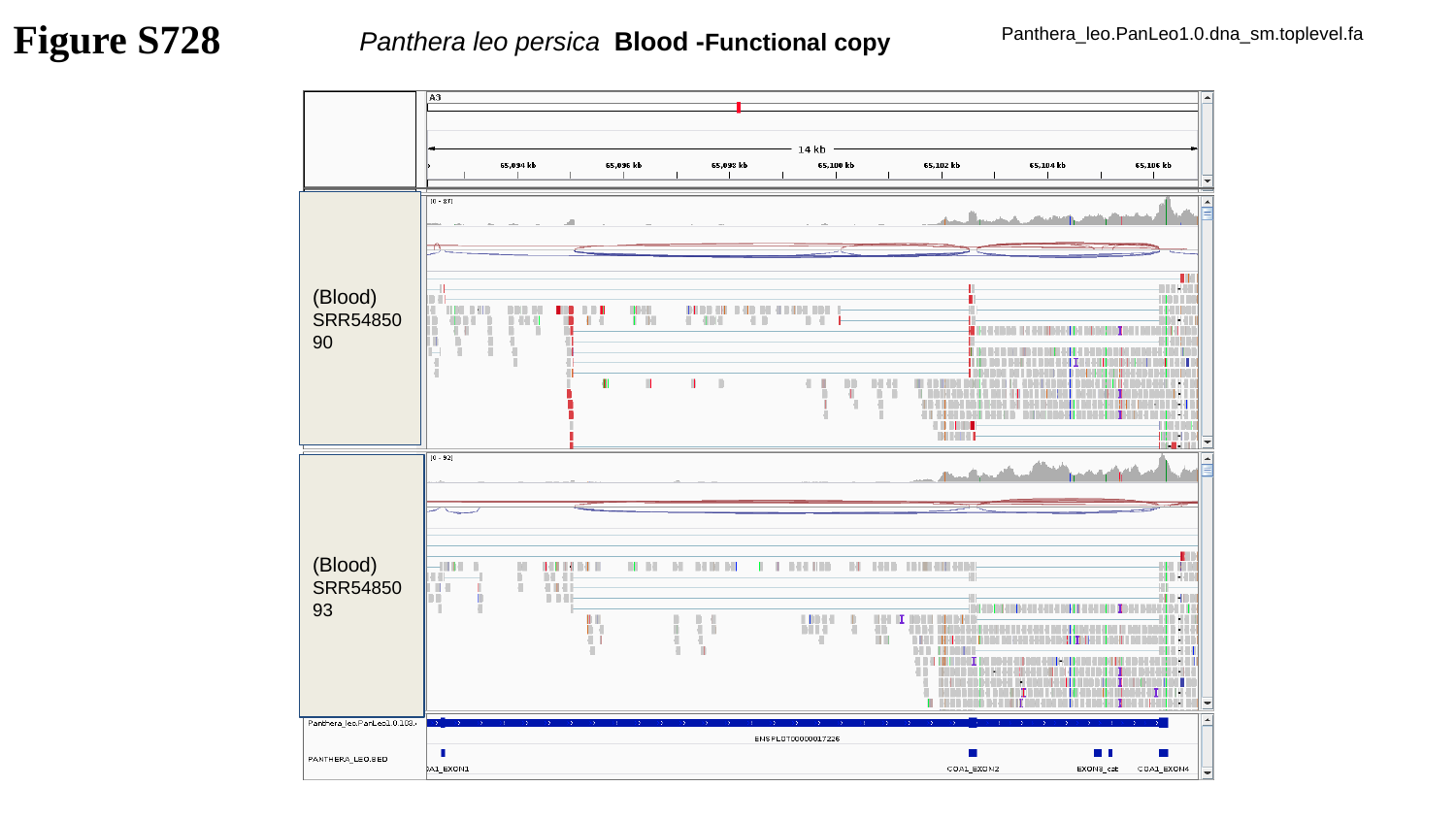

Figure S728
Panthera leo persica Blood -Functional copy
Panthera_leo.PanLeo1.0.dna_sm.toplevel.fa
(Blood)
SRR5485090
(Blood)
SRR5485093

### Slide 729
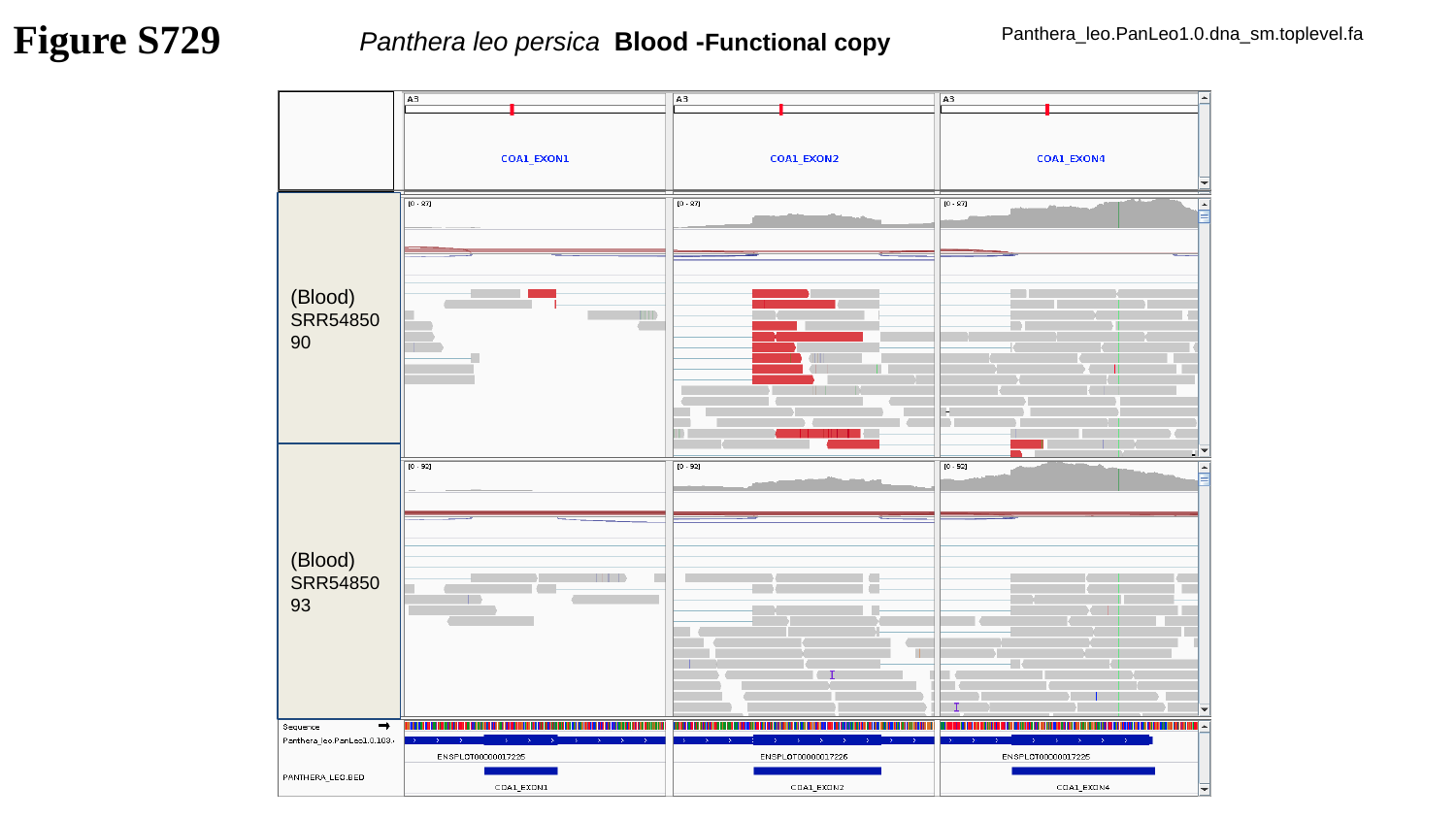

Figure S729
Panthera leo persica Blood -Functional copy
Panthera_leo.PanLeo1.0.dna_sm.toplevel.fa
(Blood)
SRR5485090
(Blood)
SRR5485093

### Slide 730
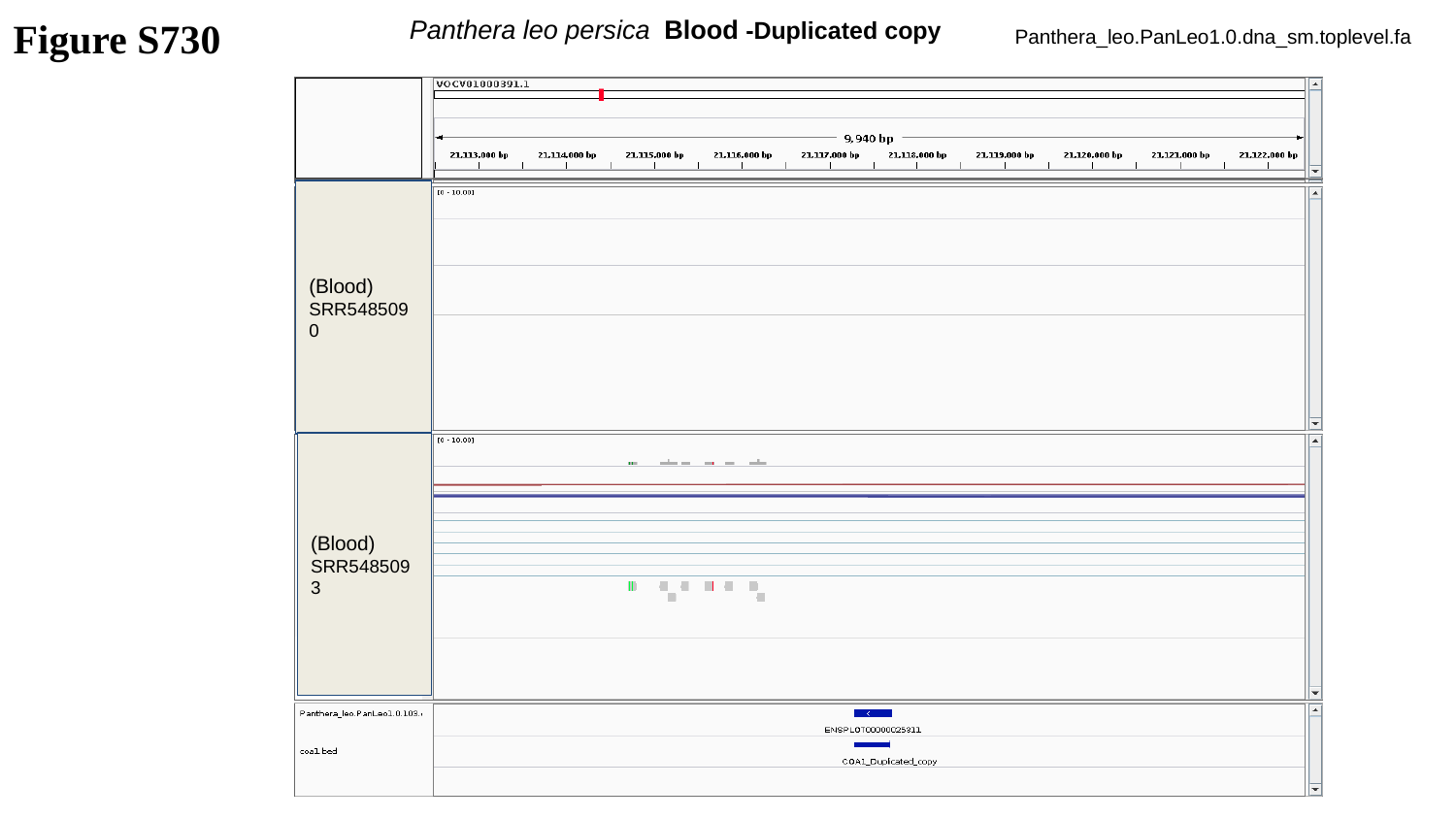

Figure S730
Panthera leo persica Blood -Duplicated copy
Panthera_leo.PanLeo1.0.dna_sm.toplevel.fa
(Blood)
SRR5485090
(Blood)
SRR5485093

### Slide 731
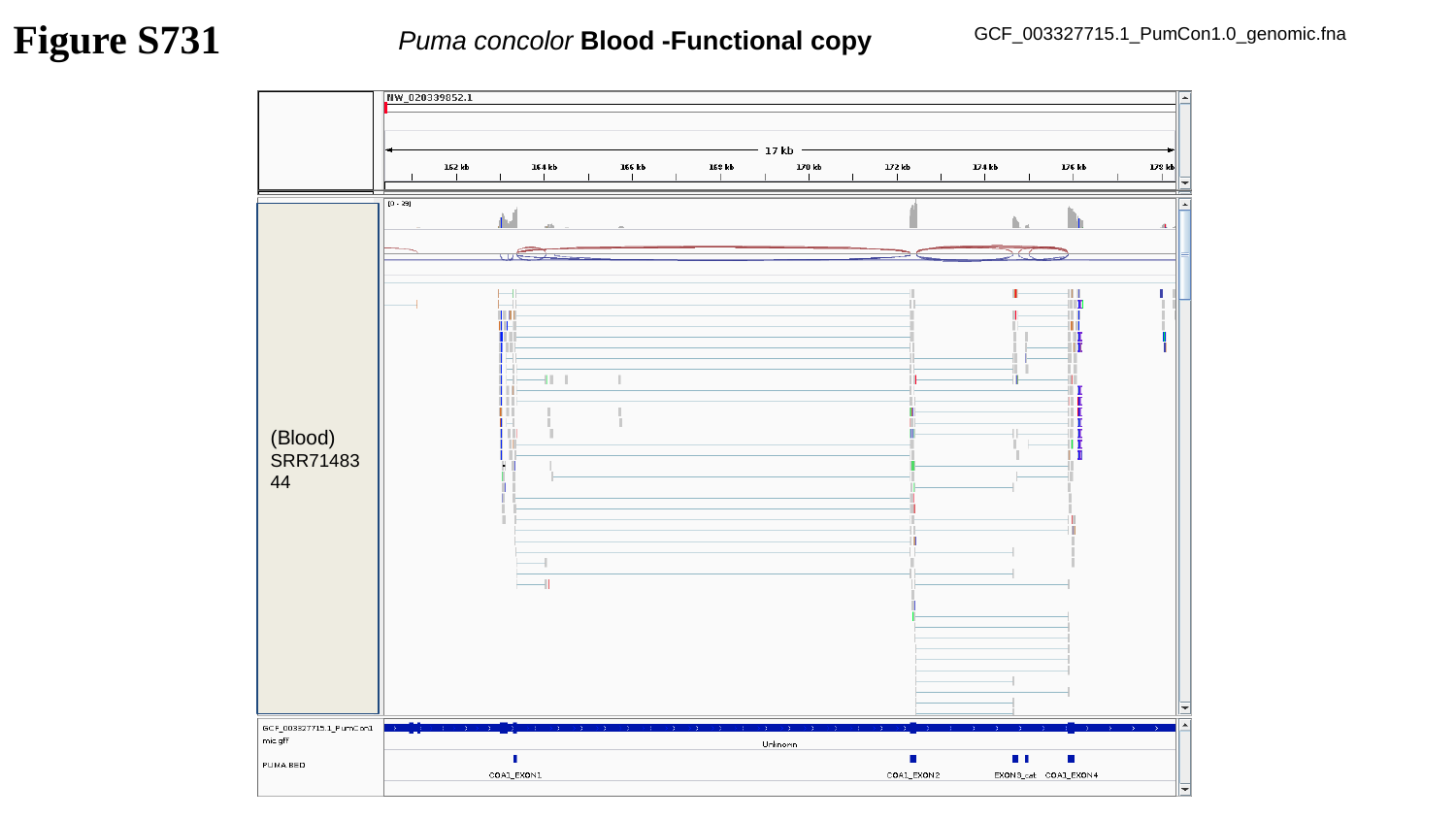

Figure S731
Puma concolor Blood -Functional copy
GCF_003327715.1_PumCon1.0_genomic.fna
(Blood)
SRR7148344

### Slide 732
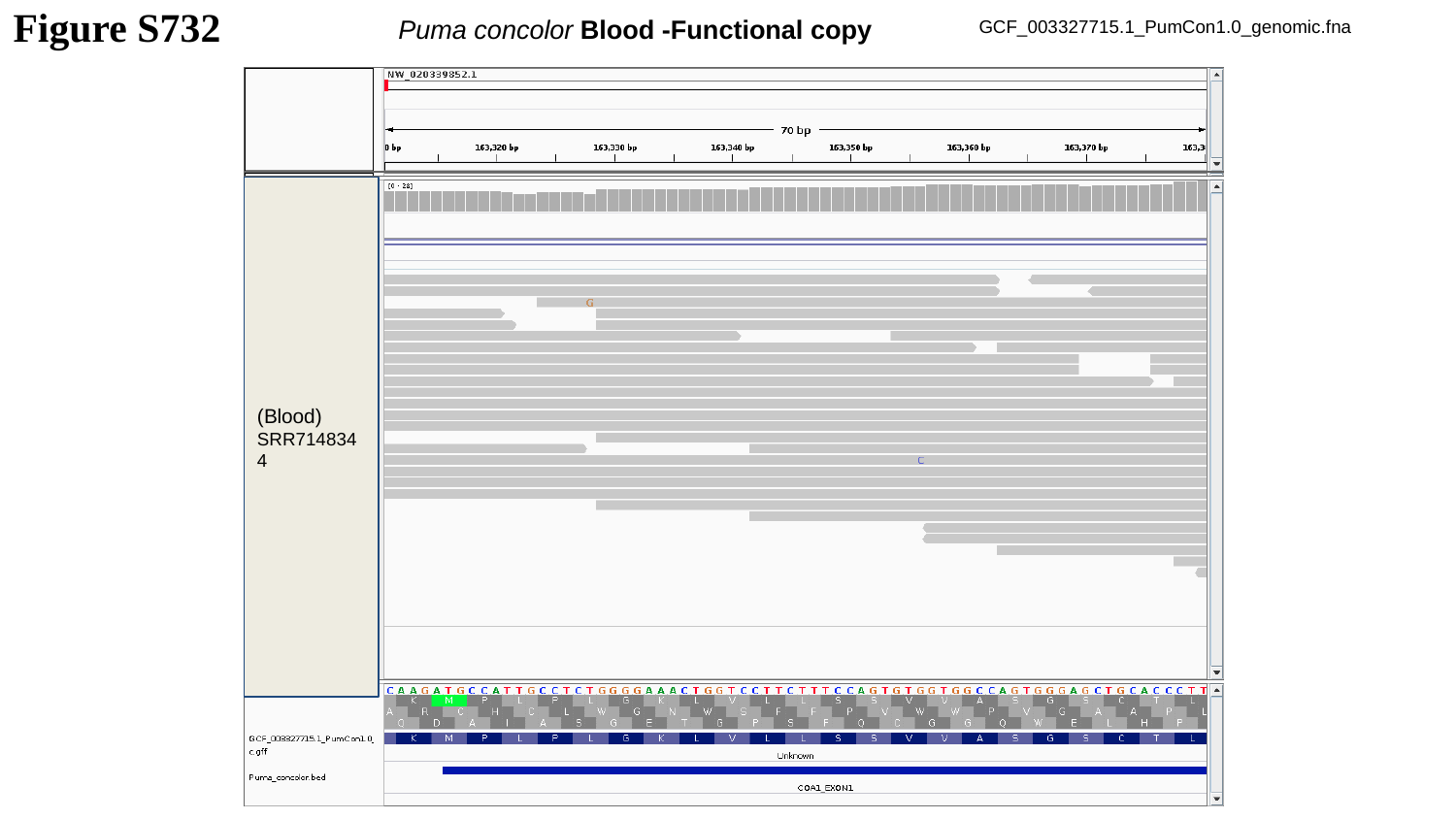

Puma concolor Blood -Functional copy
Figure S732
GCF_003327715.1_PumCon1.0_genomic.fna
(Blood)
SRR7148344

### Slide 733
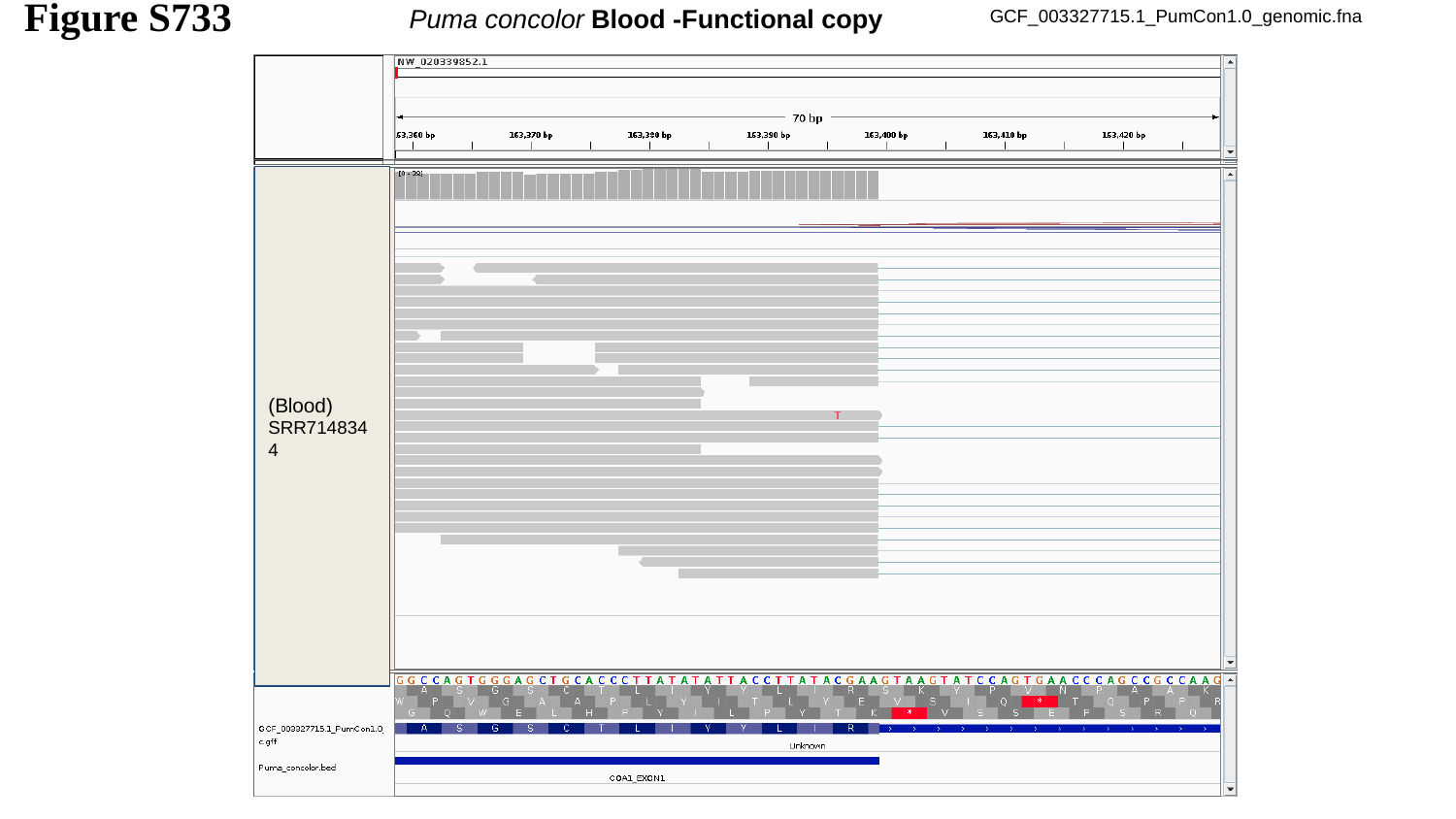

Figure S733
Puma concolor Blood -Functional copy
GCF_003327715.1_PumCon1.0_genomic.fna
(Blood)
SRR7148344

### Slide 734
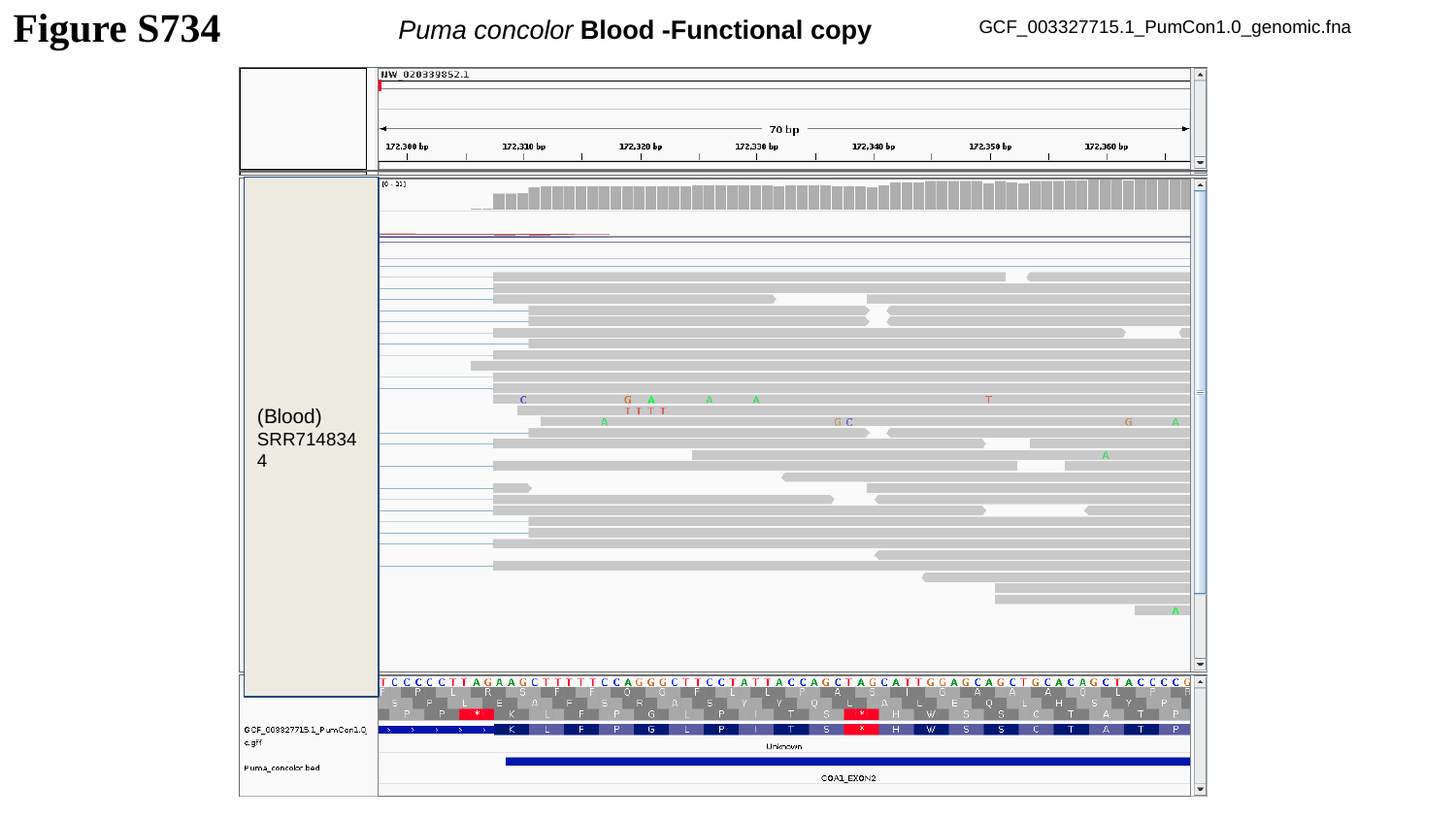

Puma concolor Blood -Functional copy
Figure S734
GCF_003327715.1_PumCon1.0_genomic.fna
(Blood)
SRR7148344

### Slide 735
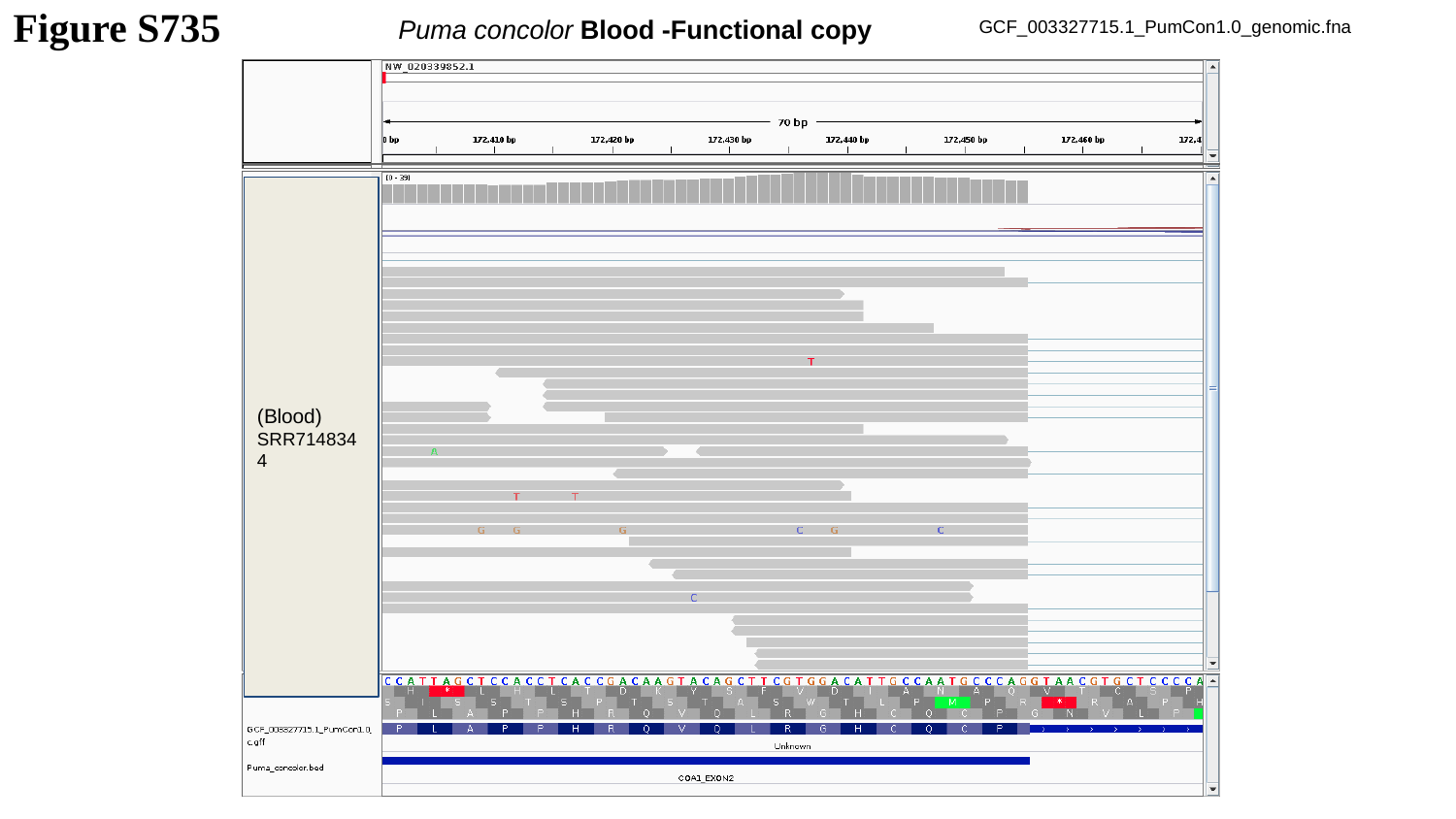

Puma concolor Blood -Functional copy
Figure S735
GCF_003327715.1_PumCon1.0_genomic.fna
(Blood)
SRR7148344

### Slide 736
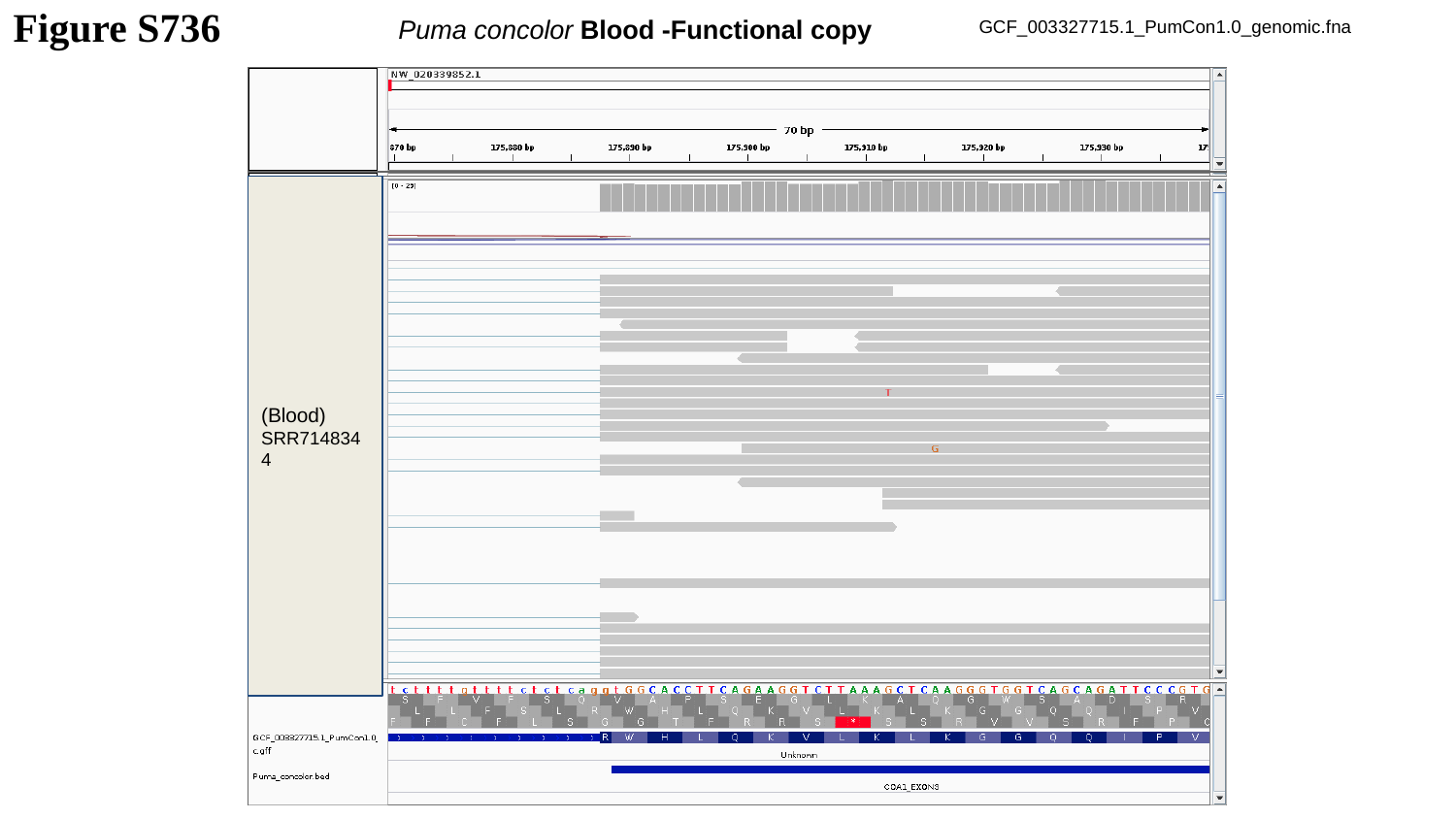

Puma concolor Blood -Functional copy
Figure S736
GCF_003327715.1_PumCon1.0_genomic.fna
(Blood)
SRR7148344

### Slide 737
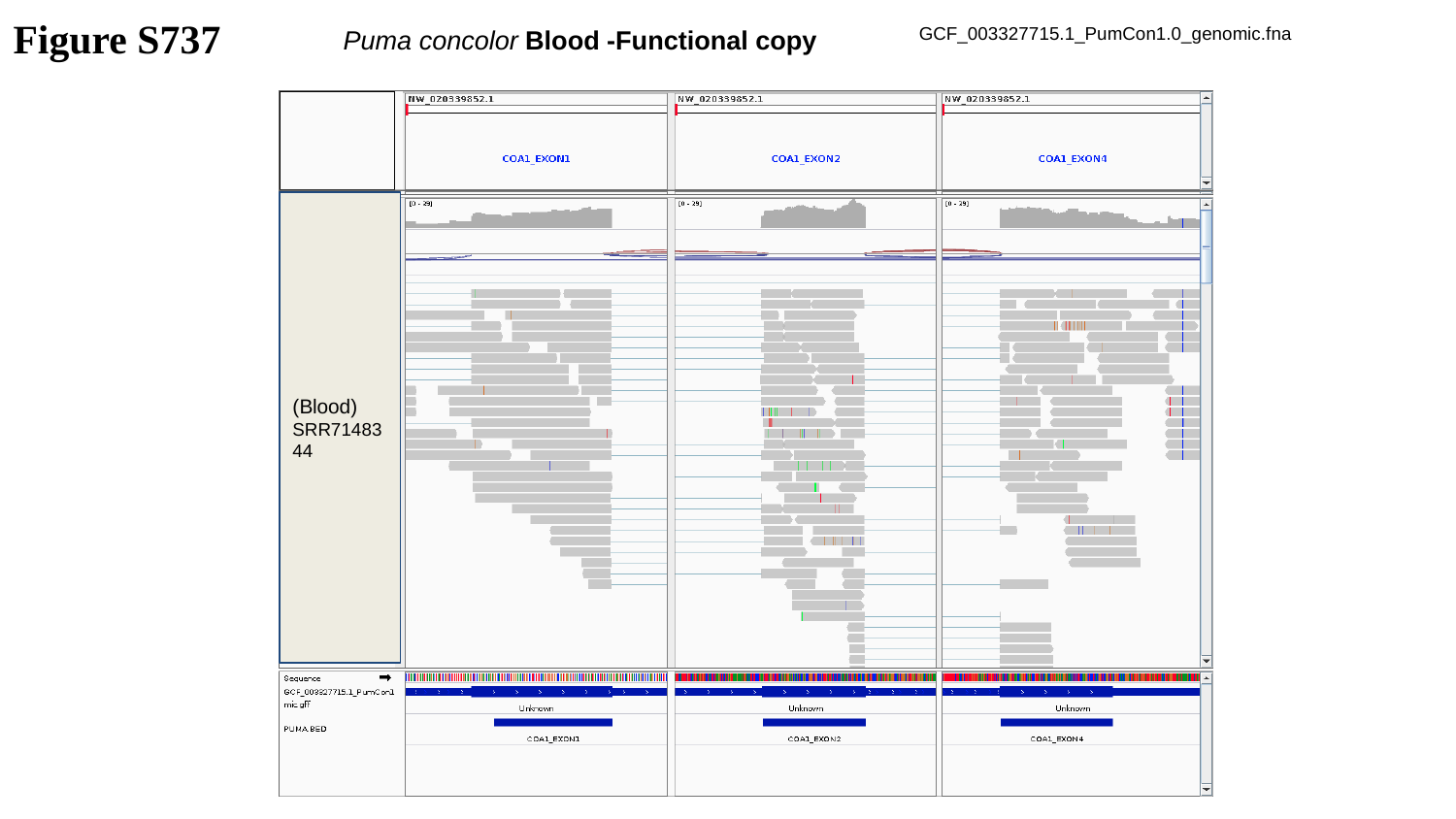

Figure S737
Puma concolor Blood -Functional copy
GCF_003327715.1_PumCon1.0_genomic.fna
(Blood)
SRR7148344

### Slide 738
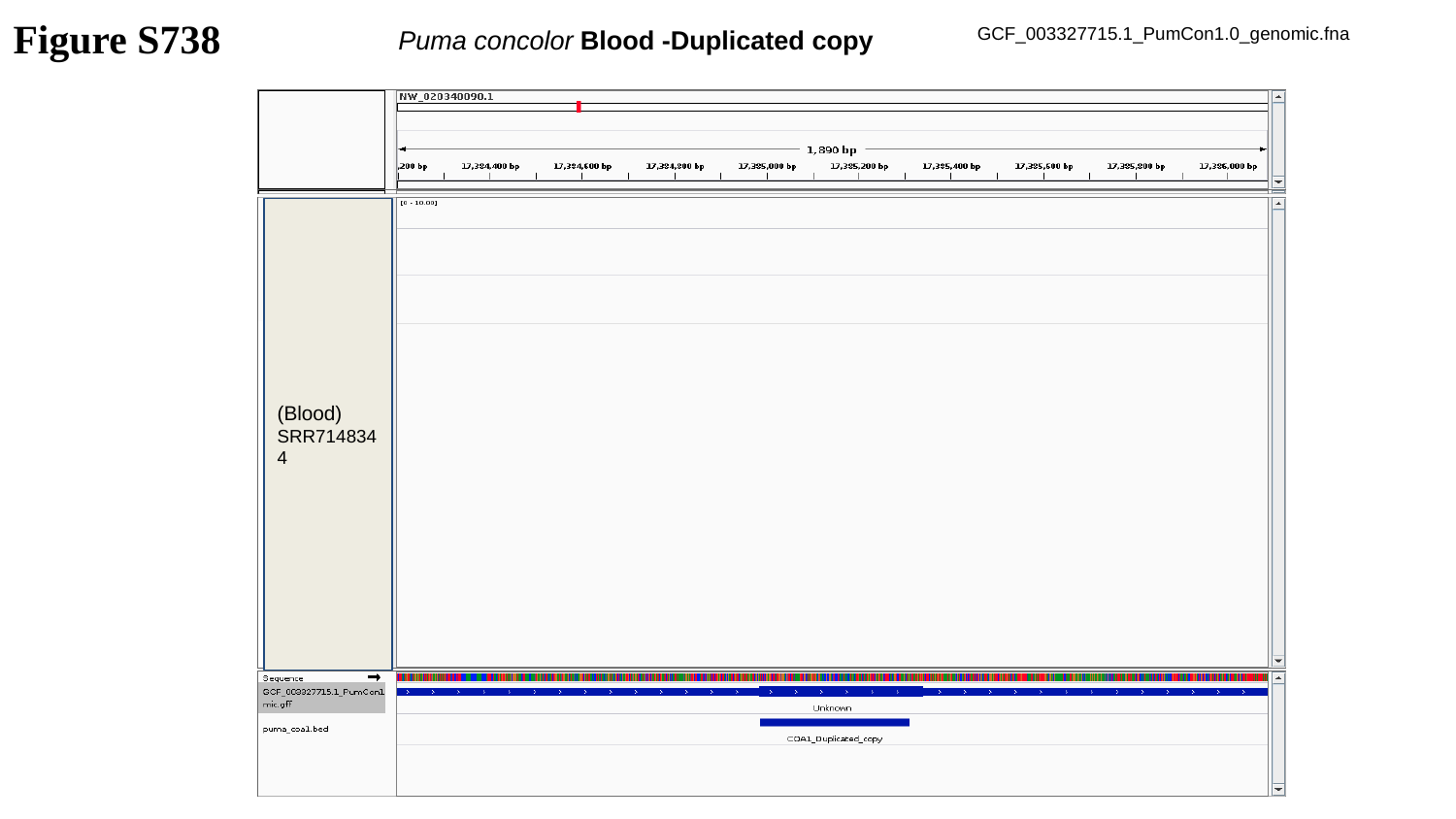

Figure S738
Puma concolor Blood -Duplicated copy
GCF_003327715.1_PumCon1.0_genomic.fna
(Blood)
SRR7148344

### Slide 739
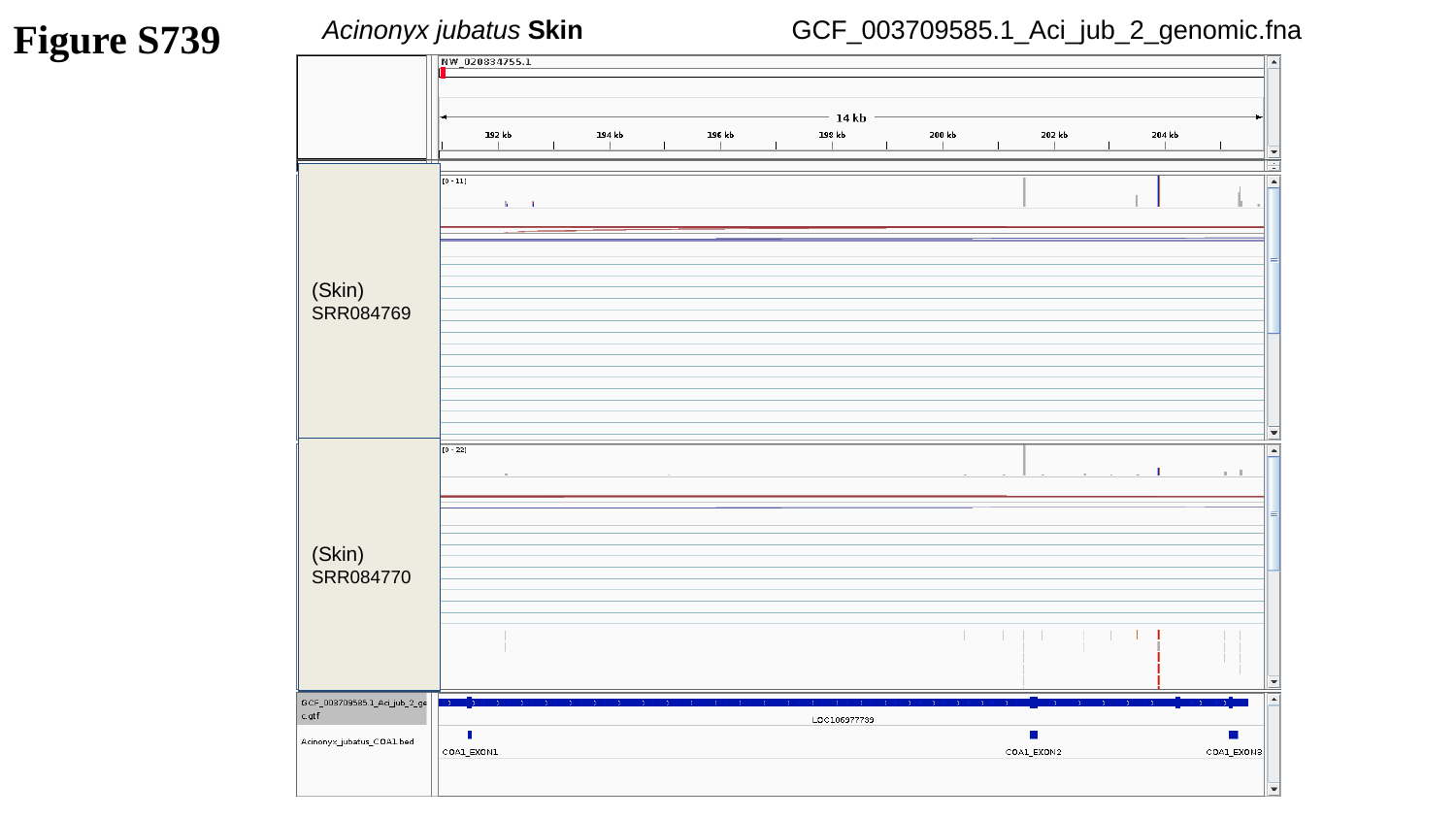

Figure S739
Acinonyx jubatus Skin
GCF_003709585.1_Aci_jub_2_genomic.fna
(Skin)
SRR084769
(Skin)
SRR084770

### Slide 740
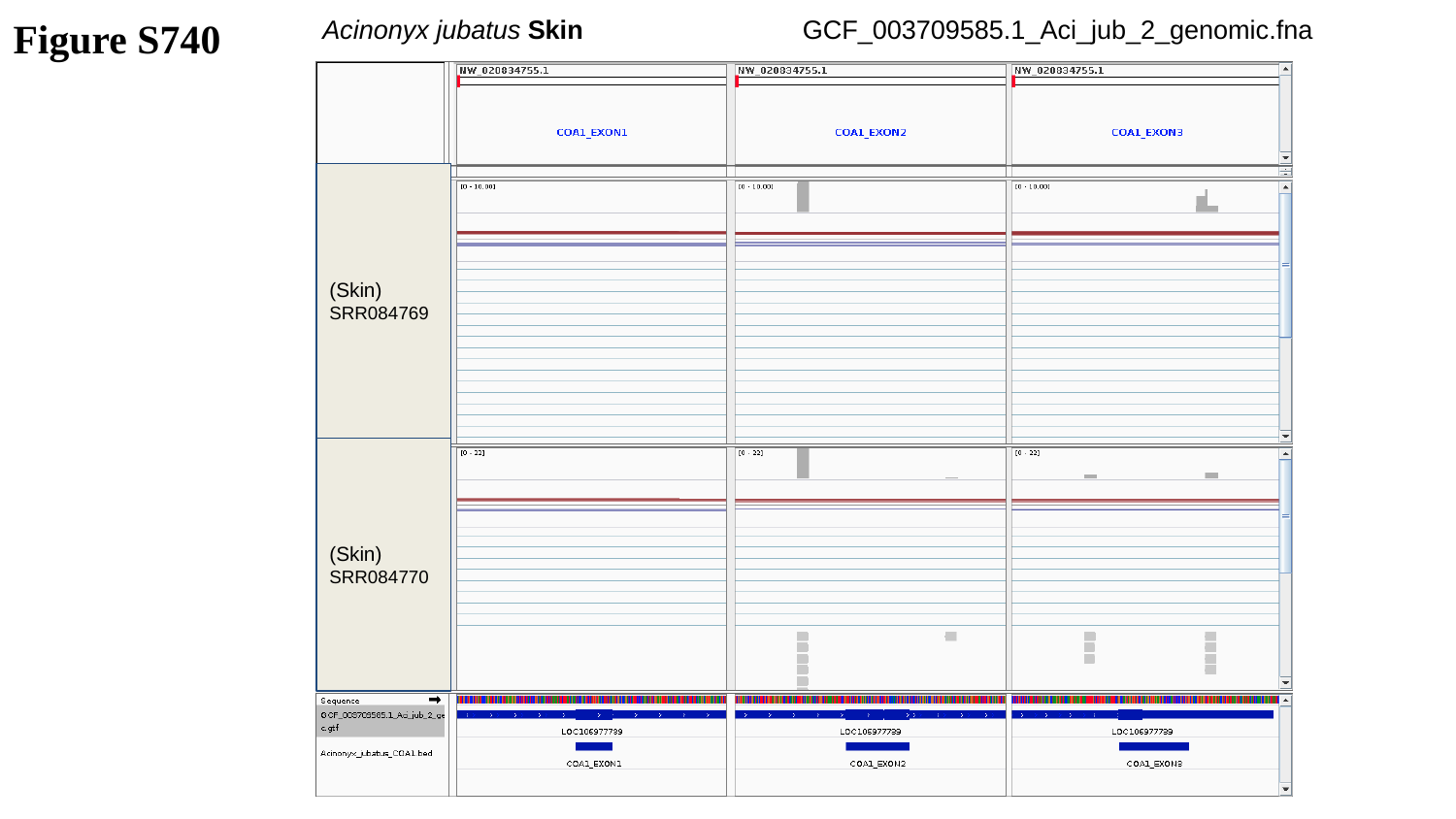

Figure S740
Acinonyx jubatus Skin
GCF_003709585.1_Aci_jub_2_genomic.fna
(Skin)
SRR084769
(Skin)
SRR084770

### Slide 741
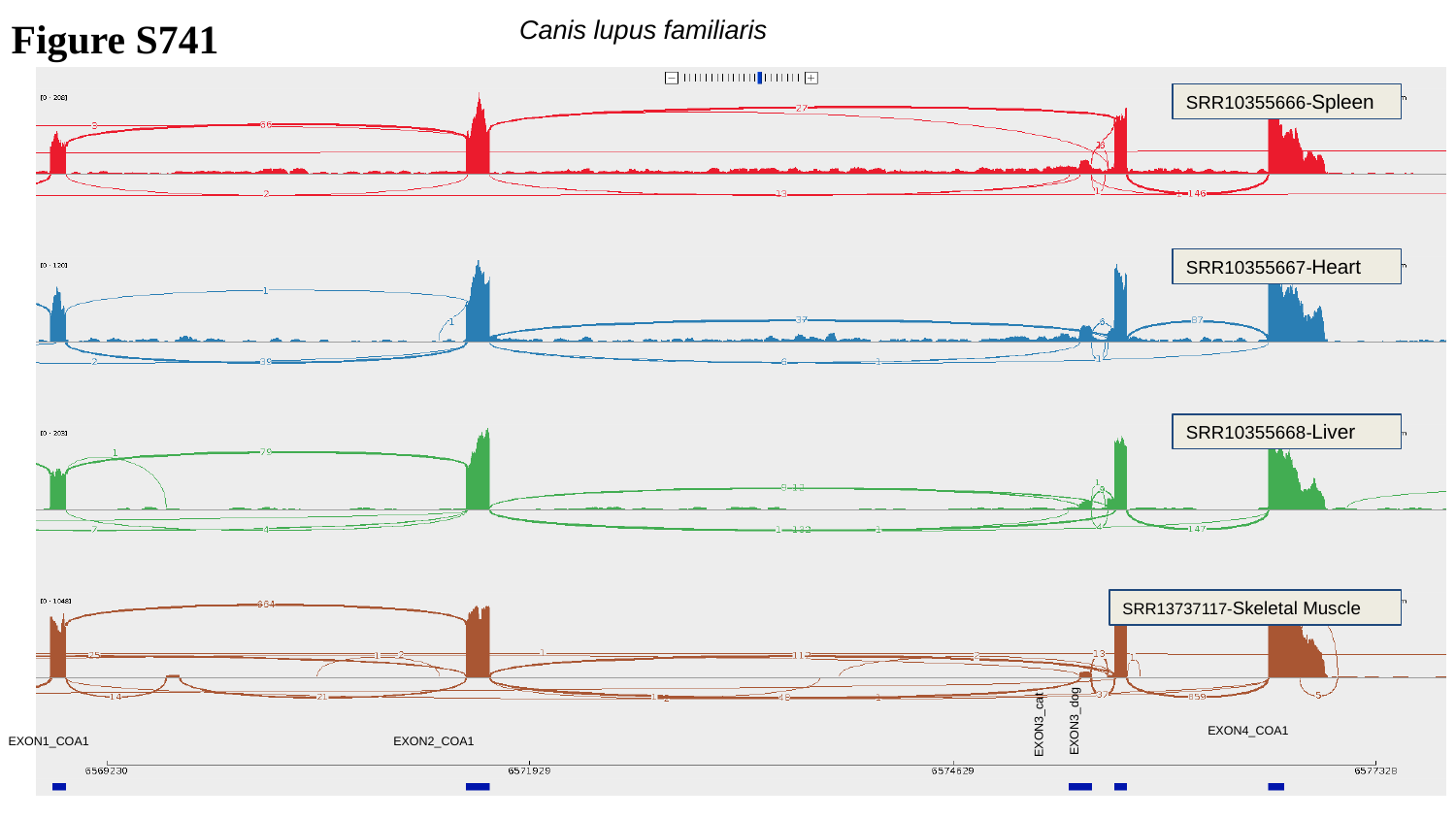

Figure S741
Canis lupus familiaris
SRR10355666-Spleen
SRR10355667-Heart
SRR10355668-Liver
SRR13737117-Skeletal Muscle
 EXON3_cat
EXON3_dog
EXON4_COA1
EXON1_COA1
EXON2_COA1

### Slide 742
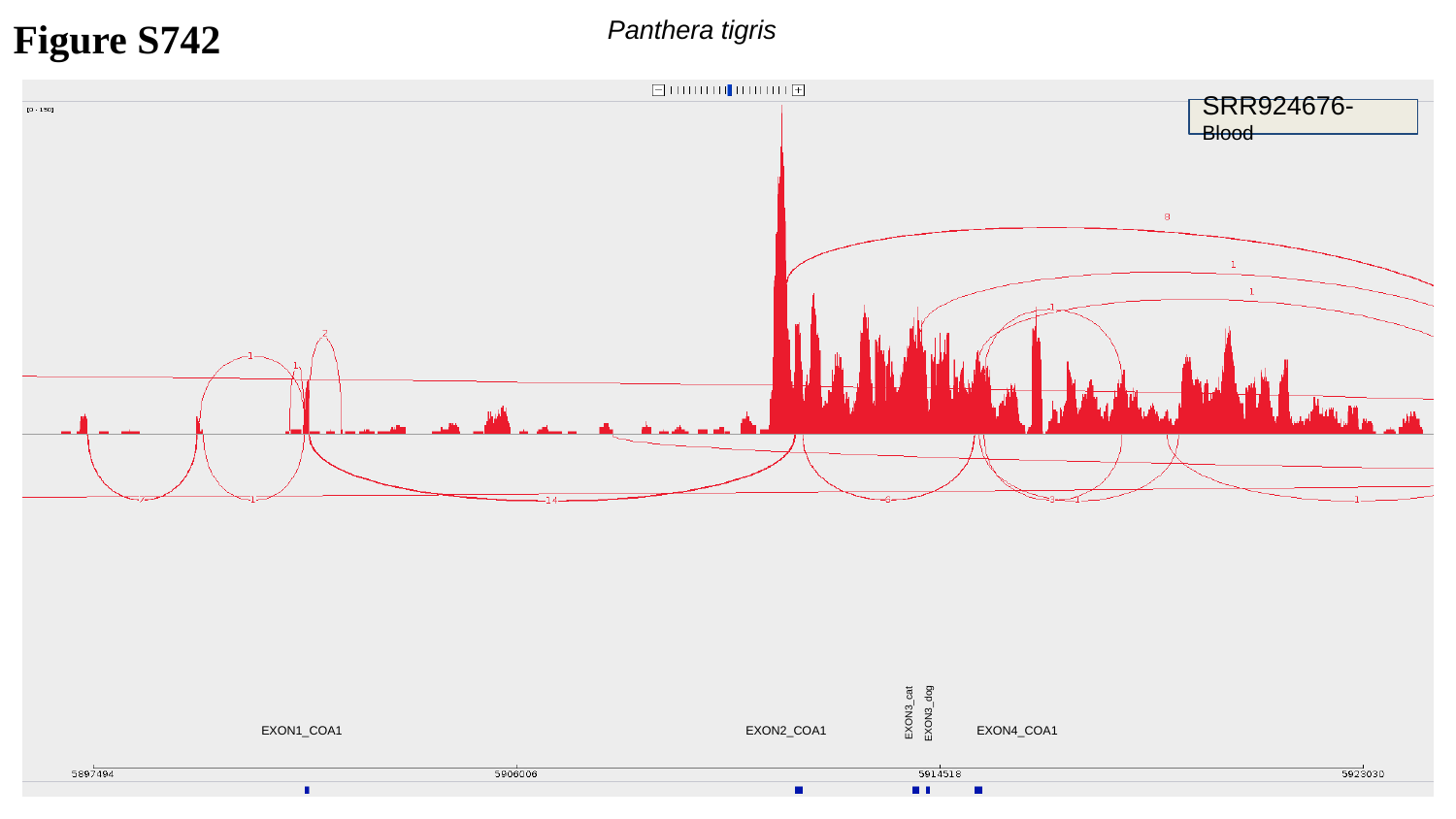

Figure S742
Panthera tigris
SRR924676-Blood
 EXON3_cat
 EXON3_dog
EXON1_COA1
EXON2_COA1
EXON4_COA1

### Slide 743
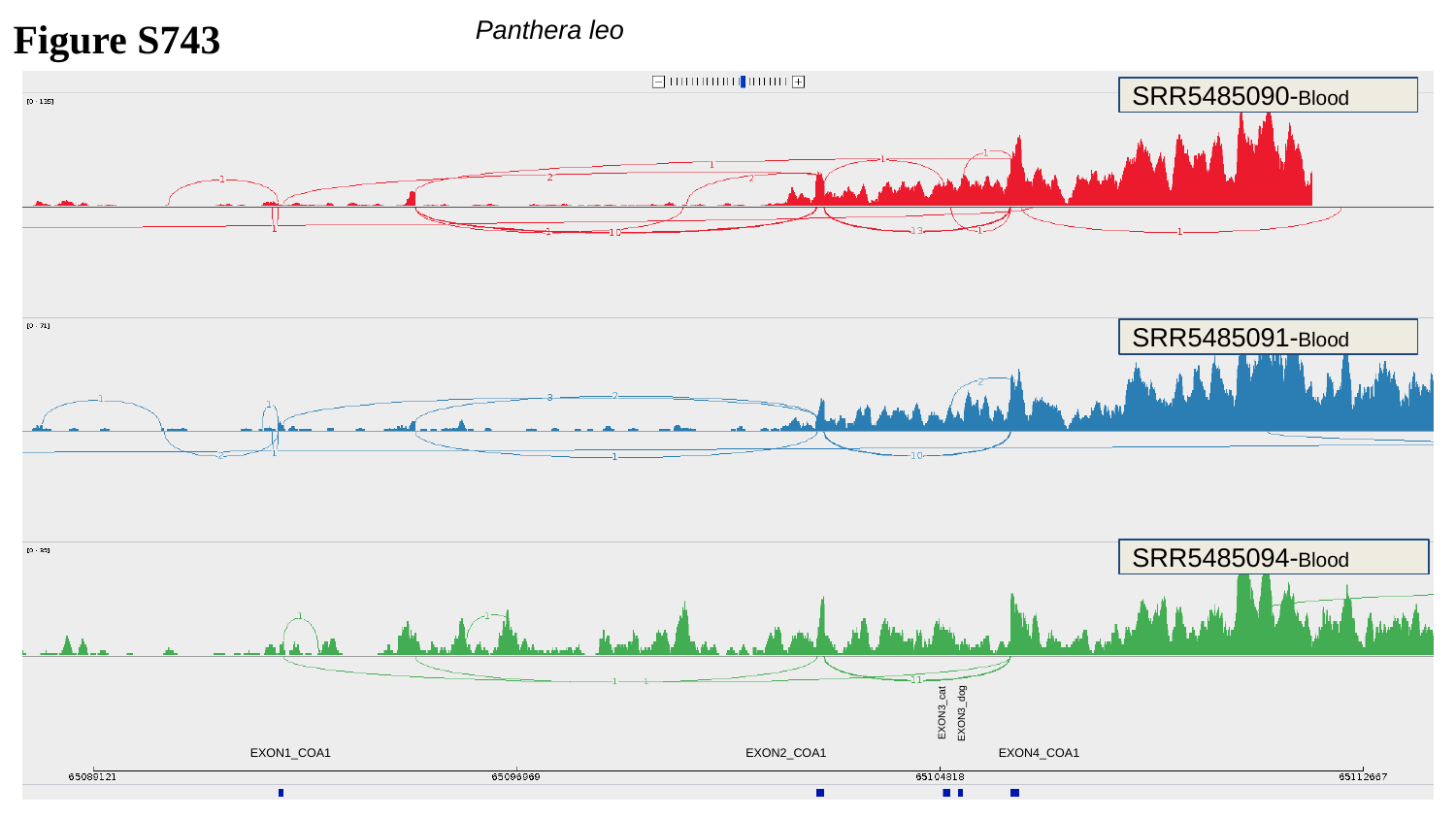

Figure S743
Panthera leo
SRR5485090-Blood
SRR5485091-Blood
SRR5485094-Blood
 EXON3_cat
 EXON3_dog
EXON1_COA1
EXON2_COA1
EXON4_COA1

### Slide 744
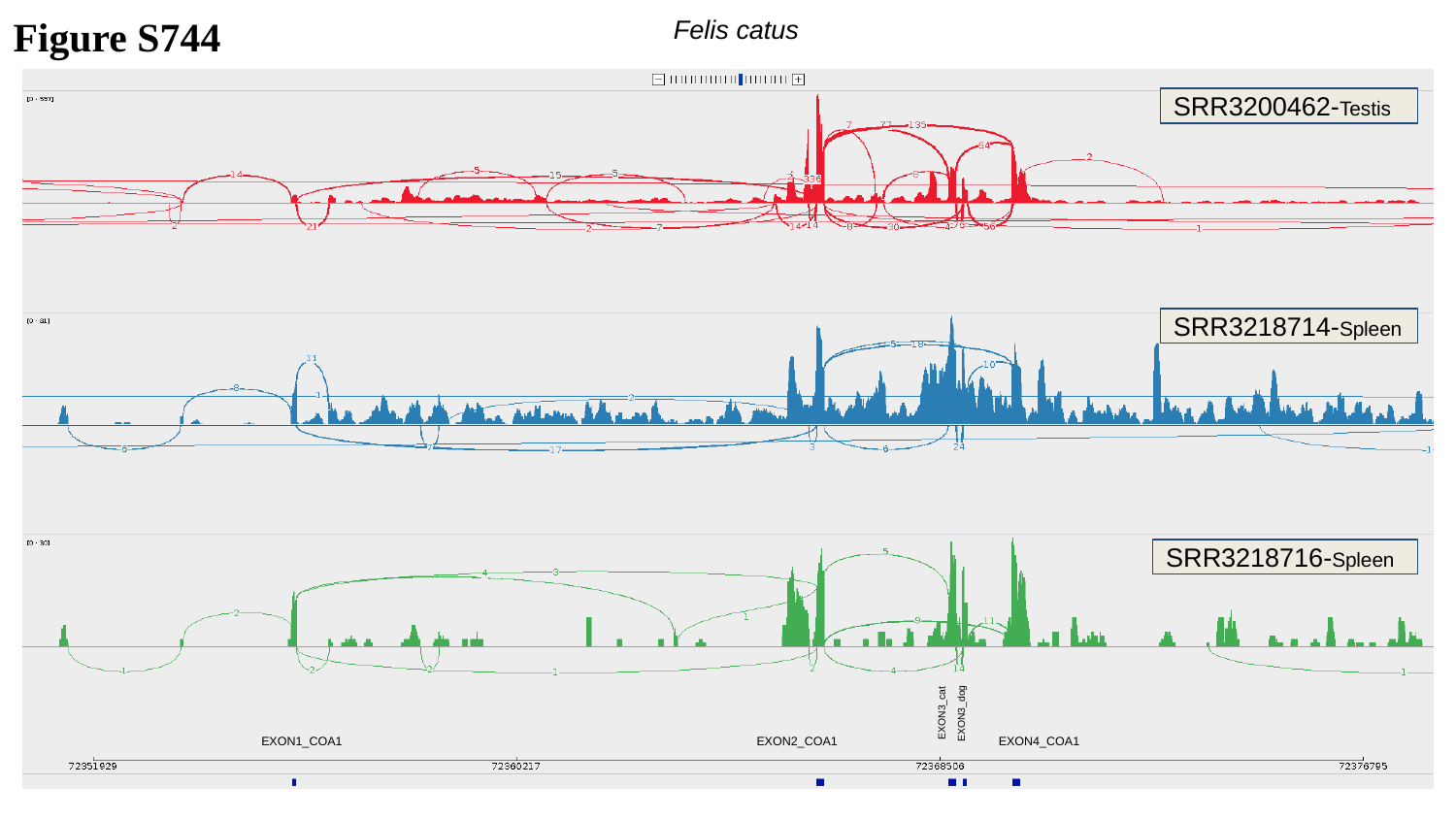

Felis catus
Figure S744
SRR3200462-Testis
SRR3218714-Spleen
SRR3218716-Spleen
 EXON3_cat
 EXON3_dog
EXON1_COA1
EXON2_COA1
EXON4_COA1

### Slide 745
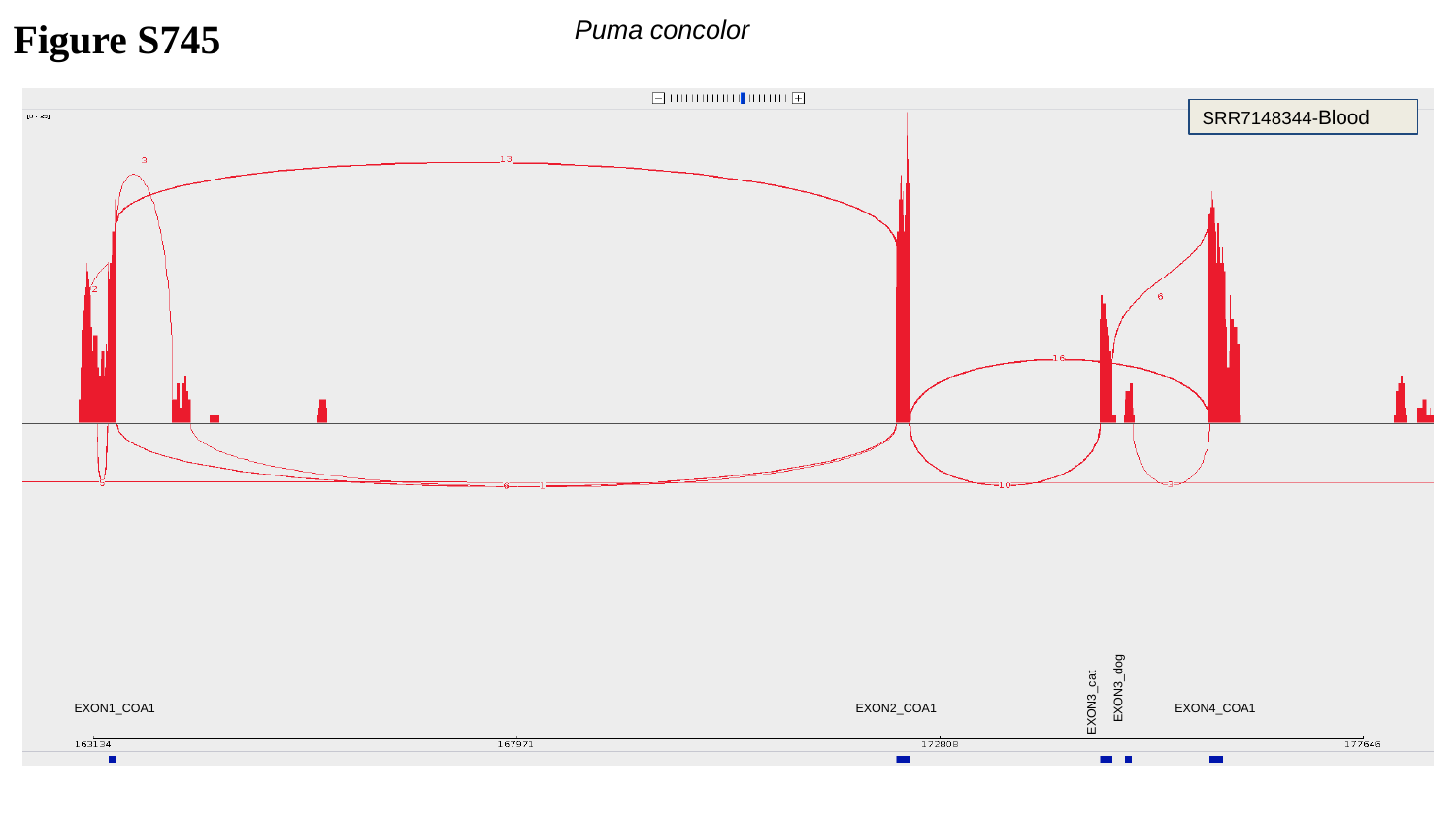

Figure S745
Puma concolor
SRR7148344-Blood
 EXON3_cat
EXON3_dog
EXON1_COA1
EXON2_COA1
EXON4_COA1

### Slide 746
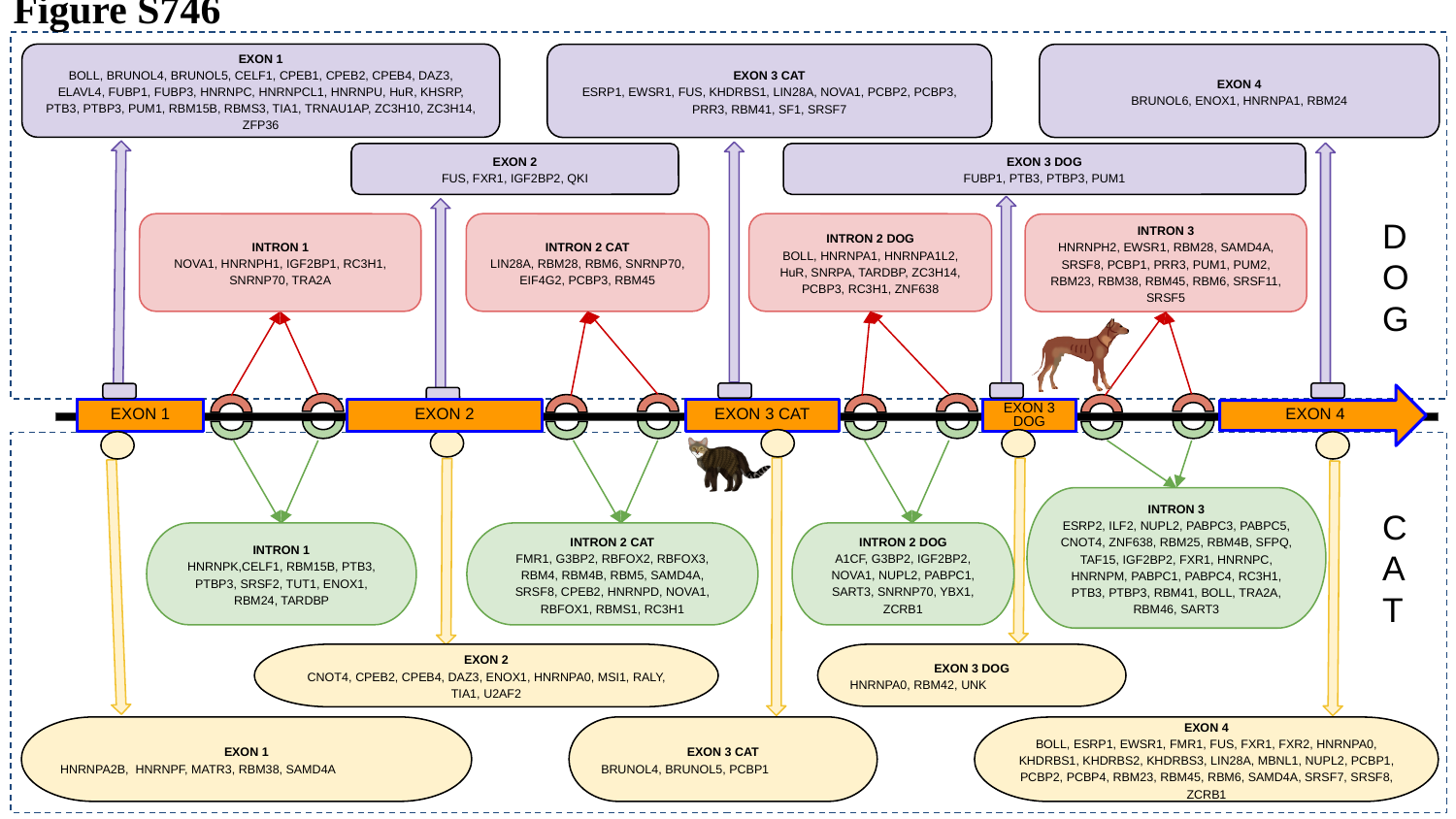

Figure S746
 EXON 1
BOLL, BRUNOL4, BRUNOL5, CELF1, CPEB1, CPEB2, CPEB4, DAZ3, ELAVL4, FUBP1, FUBP3, HNRNPC, HNRNPCL1, HNRNPU, HuR, KHSRP, PTB3, PTBP3, PUM1, RBM15B, RBMS3, TIA1, TRNAU1AP, ZC3H10, ZC3H14, ZFP36
EXON 3 CAT
ESRP1, EWSR1, FUS, KHDRBS1, LIN28A, NOVA1, PCBP2, PCBP3, PRR3, RBM41, SF1, SRSF7
EXON 4
BRUNOL6, ENOX1, HNRNPA1, RBM24
EXON 2
FUS, FXR1, IGF2BP2, QKI
EXON 3 DOG
FUBP1, PTB3, PTBP3, PUM1
D
O
G
INTRON 1
NOVA1, HNRNPH1, IGF2BP1, RC3H1, SNRNP70, TRA2A
INTRON 2 CAT
LIN28A, RBM28, RBM6, SNRNP70, EIF4G2, PCBP3, RBM45
INTRON 2 DOG
BOLL, HNRNPA1, HNRNPA1L2, HuR, SNRPA, TARDBP, ZC3H14, PCBP3, RC3H1, ZNF638
INTRON 3
HNRNPH2, EWSR1, RBM28, SAMD4A, SRSF8, PCBP1, PRR3, PUM1, PUM2, RBM23, RBM38, RBM45, RBM6, SRSF11, SRSF5
EXON 4
EXON 1
EXON 2
EXON 3 CAT
EXON 3 DOG
INTRON 3
ESRP2, ILF2, NUPL2, PABPC3, PABPC5, CNOT4, ZNF638, RBM25, RBM4B, SFPQ, TAF15, IGF2BP2, FXR1, HNRNPC, HNRNPM, PABPC1, PABPC4, RC3H1, PTB3, PTBP3, RBM41, BOLL, TRA2A, RBM46, SART3
C
A
T
INTRON 1
HNRNPK,CELF1, RBM15B, PTB3, PTBP3, SRSF2, TUT1, ENOX1, RBM24, TARDBP
INTRON 2 CAT
FMR1, G3BP2, RBFOX2, RBFOX3, RBM4, RBM4B, RBM5, SAMD4A, SRSF8, CPEB2, HNRNPD, NOVA1, RBFOX1, RBMS1, RC3H1
INTRON 2 DOG
A1CF, G3BP2, IGF2BP2, NOVA1, NUPL2, PABPC1, SART3, SNRNP70, YBX1, ZCRB1
EXON 2
CNOT4, CPEB2, CPEB4, DAZ3, ENOX1, HNRNPA0, MSI1, RALY, TIA1, U2AF2
EXON 3 DOG
HNRNPA0, RBM42, UNK
EXON 1
HNRNPA2B, HNRNPF, MATR3, RBM38, SAMD4A
EXON 3 CAT
BRUNOL4, BRUNOL5, PCBP1
EXON 4
BOLL, ESRP1, EWSR1, FMR1, FUS, FXR1, FXR2, HNRNPA0, KHDRBS1, KHDRBS2, KHDRBS3, LIN28A, MBNL1, NUPL2, PCBP1, PCBP2, PCBP4, RBM23, RBM45, RBM6, SAMD4A, SRSF7, SRSF8, ZCRB1

### Slide 747
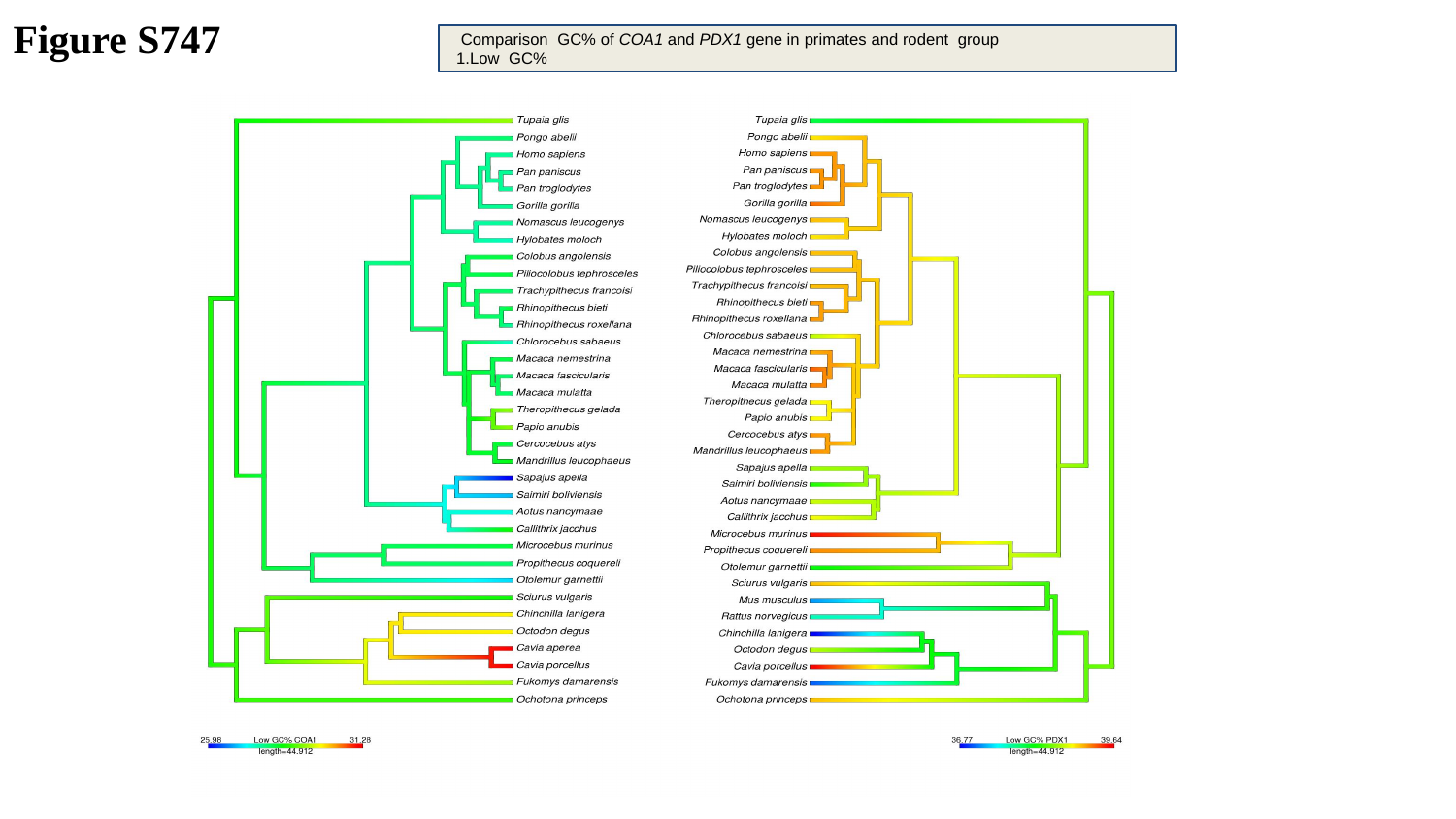

Figure S747
 Comparison GC% of COA1 and PDX1 gene in primates and rodent group
 1.Low GC%

### Slide 748
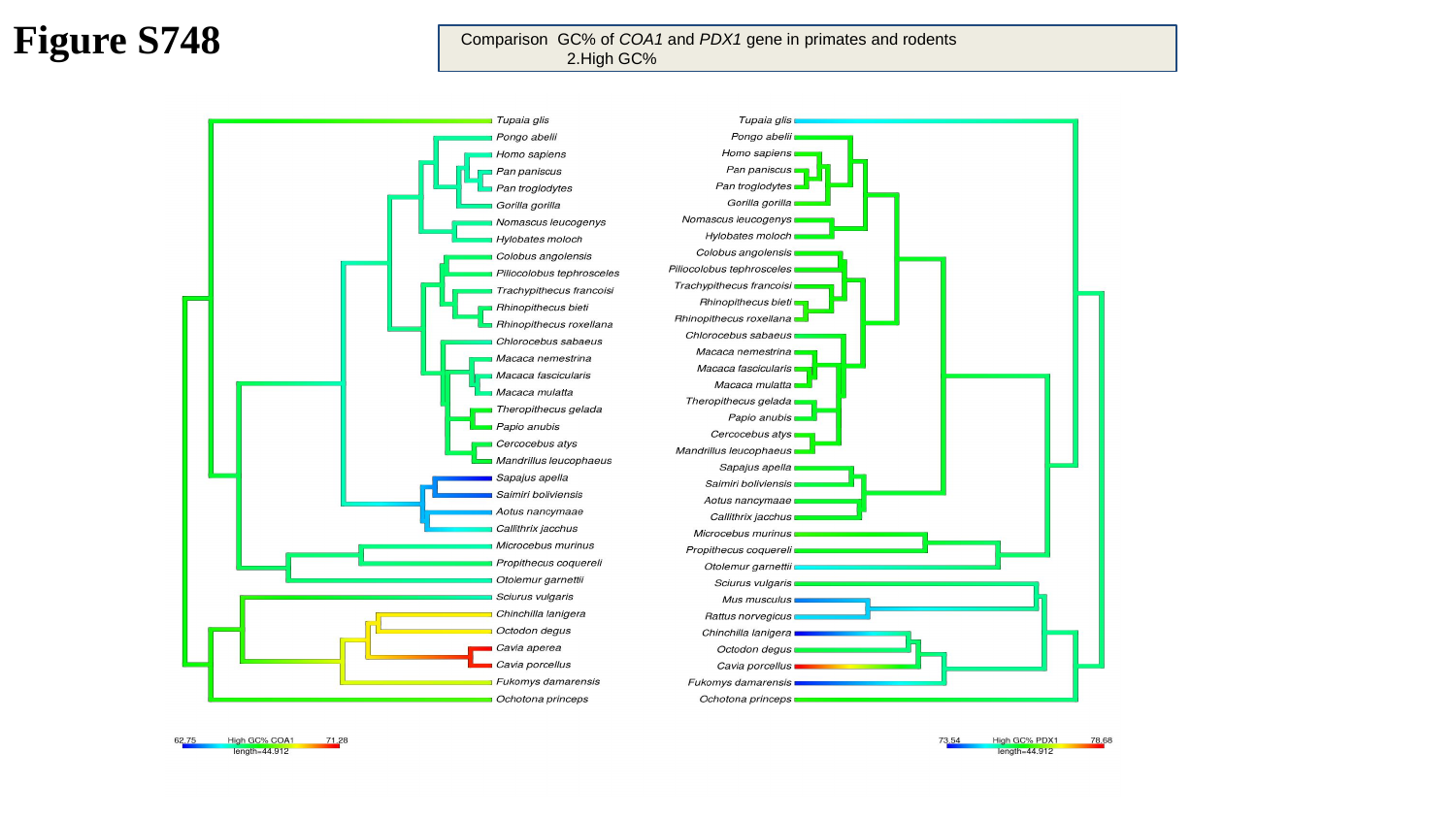

Figure S748
 Comparison GC% of COA1 and PDX1 gene in primates and rodents
 2.High GC%

### Slide 749
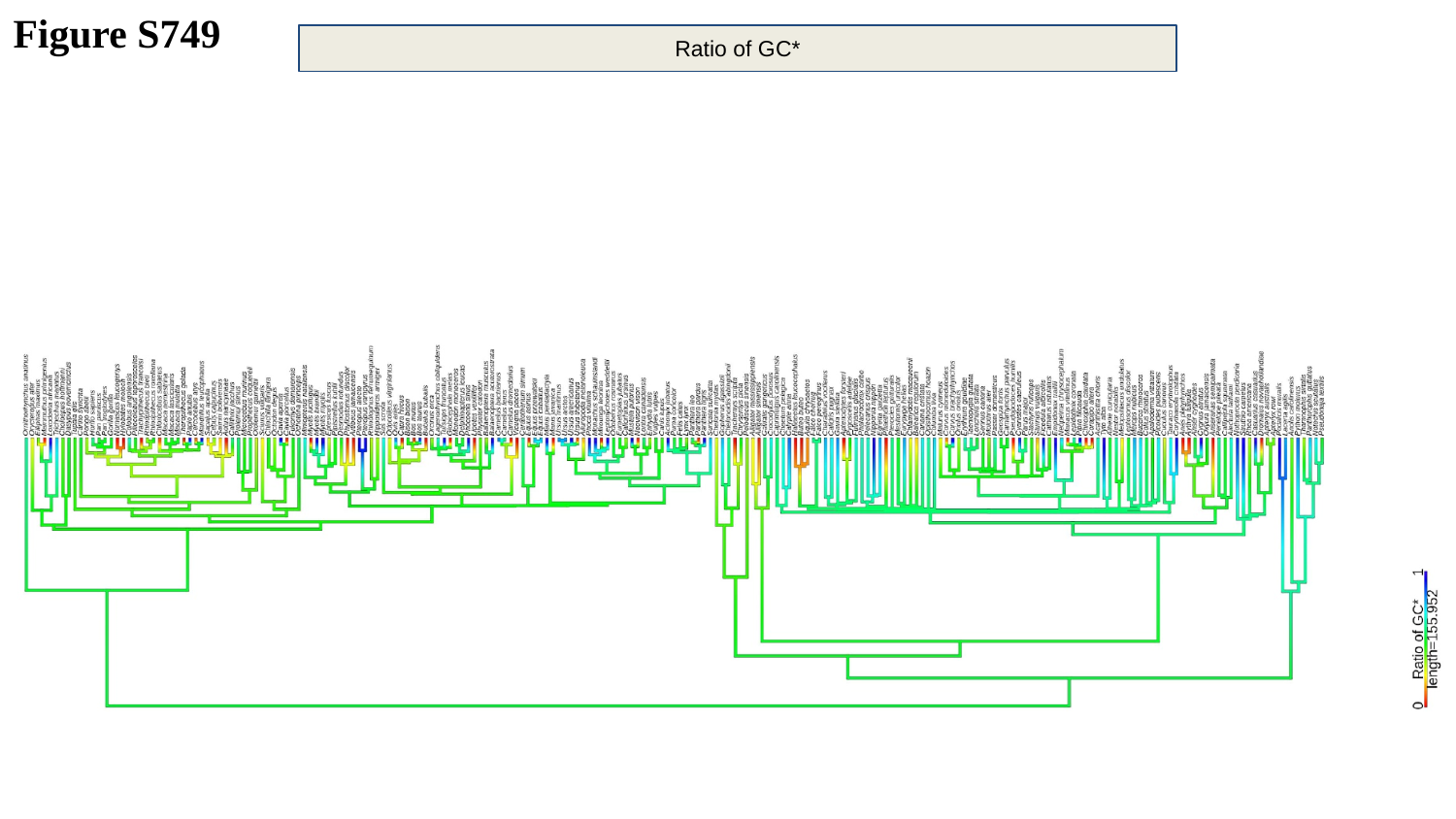

Figure S749
Ratio of GC*

### Slide 750
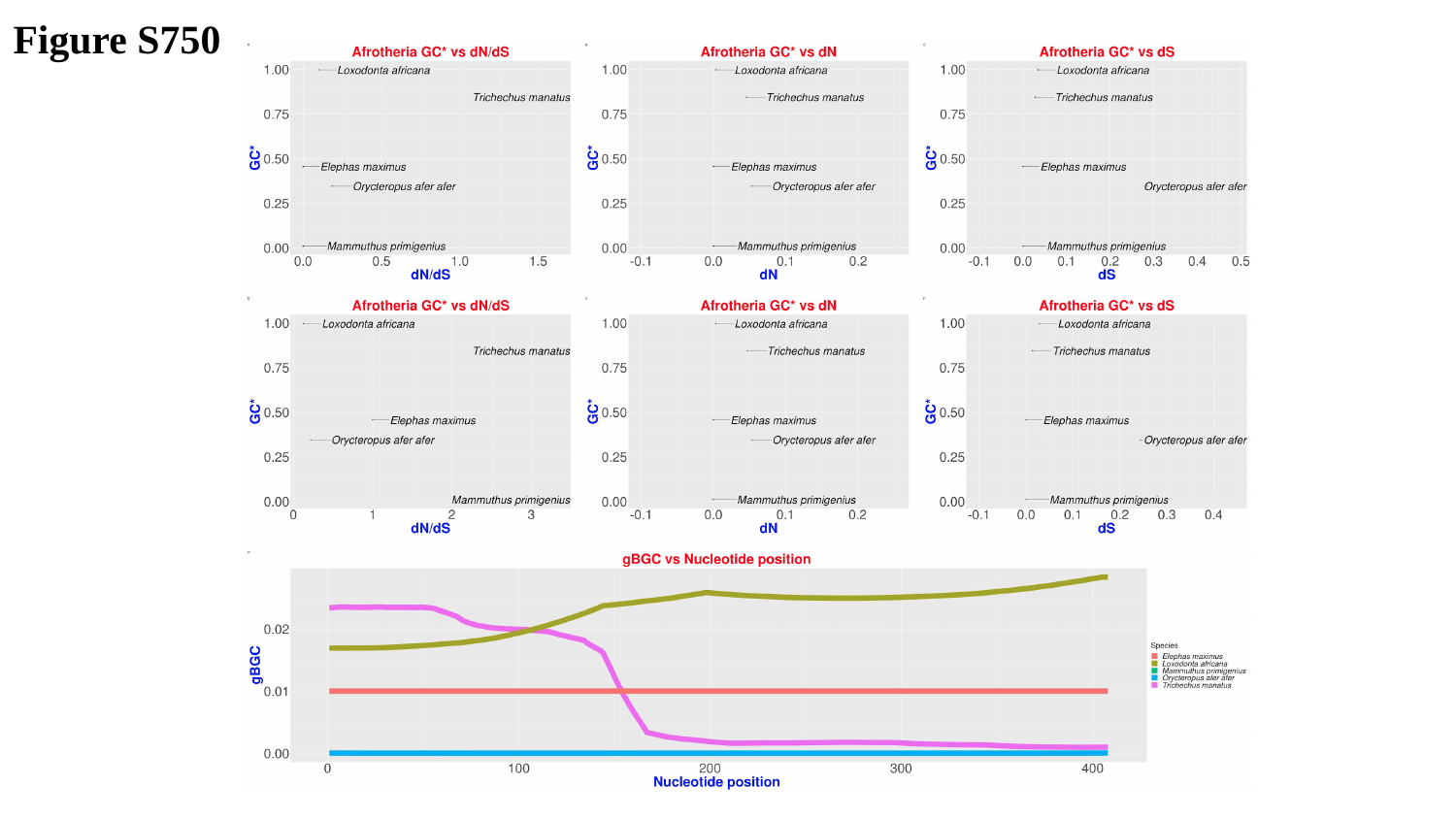

Figure S750

### Slide 751
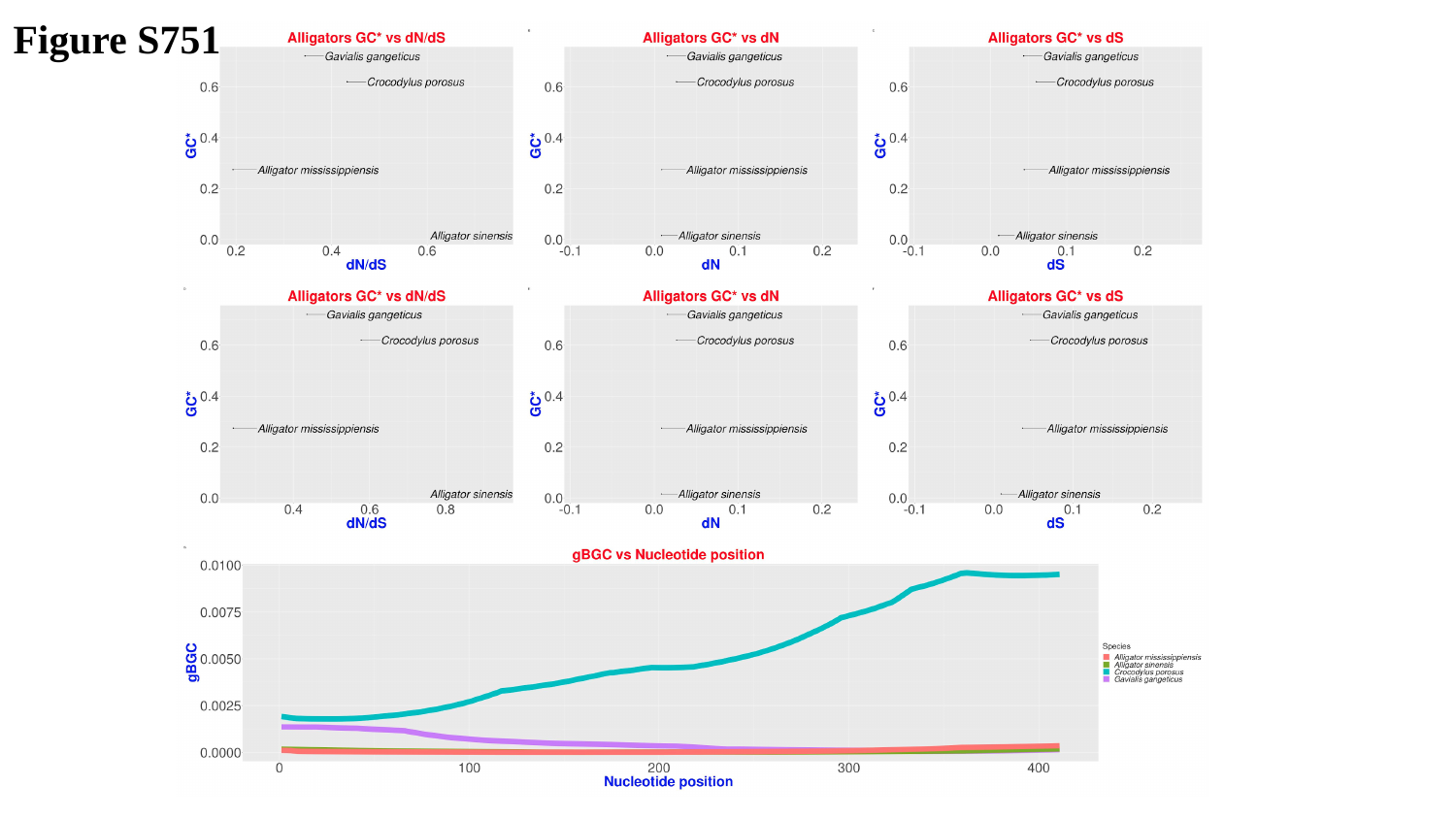

Figure S751

### Slide 752
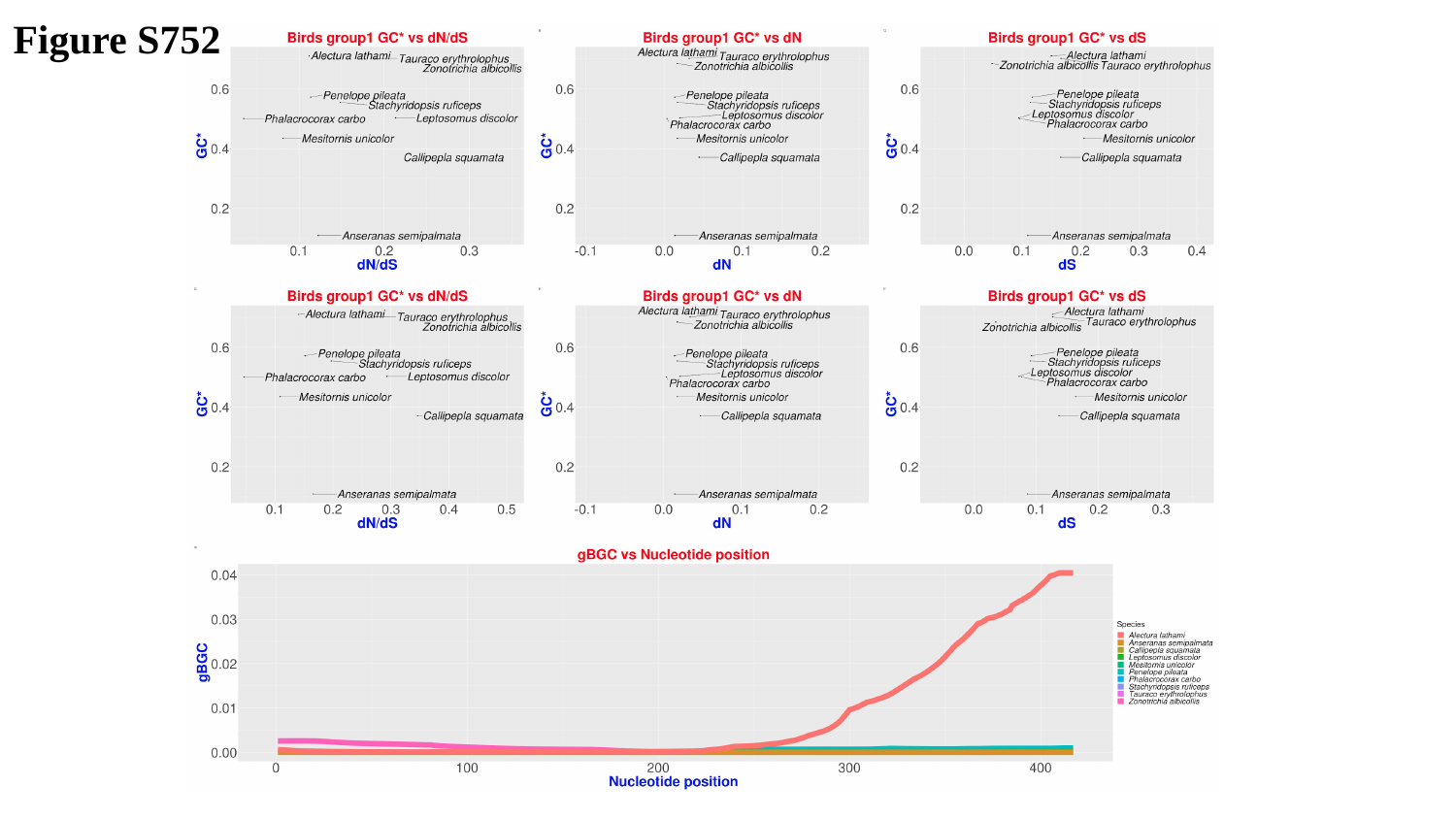

Figure S752

### Slide 753
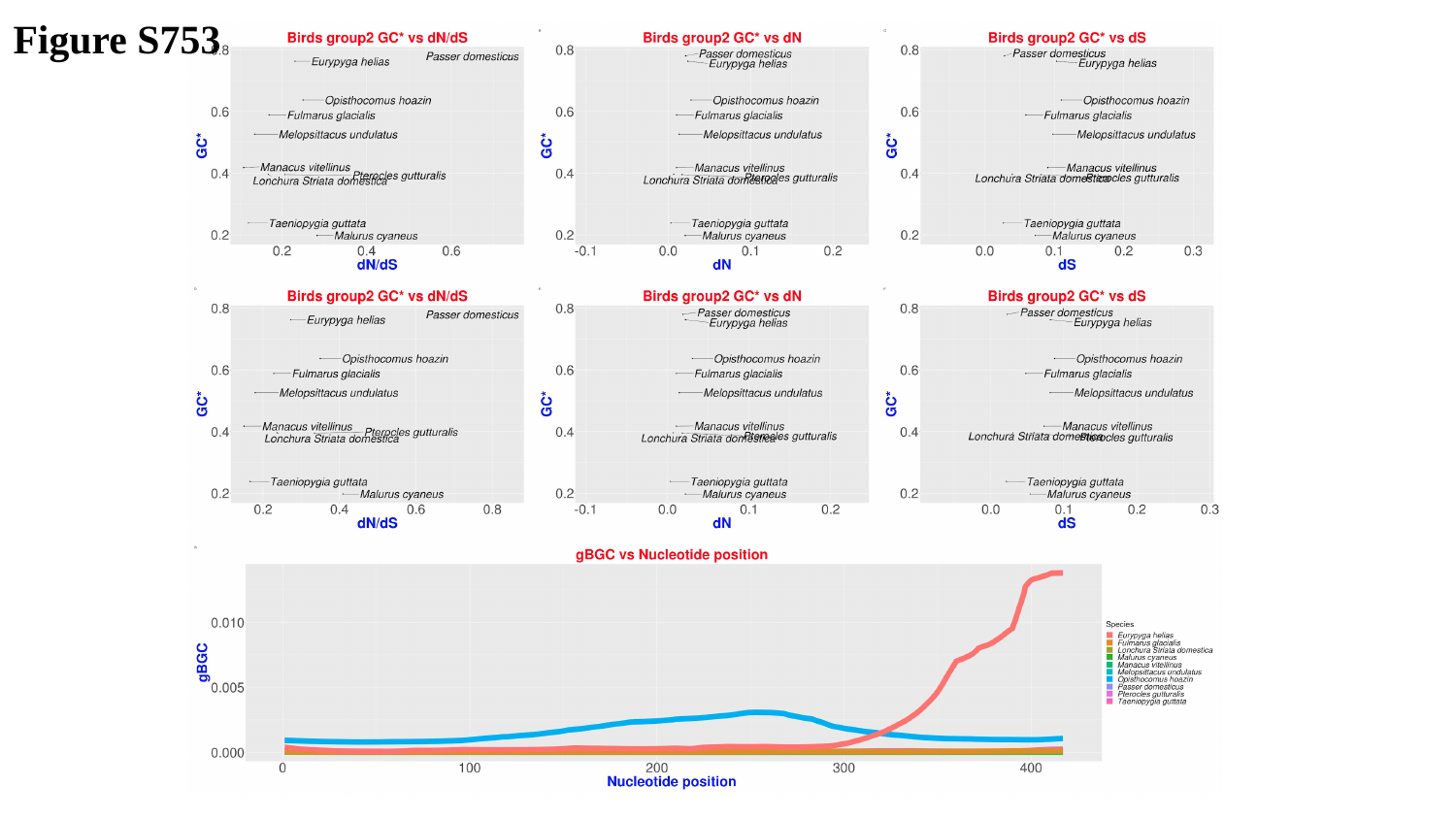

Figure S753

### Slide 754
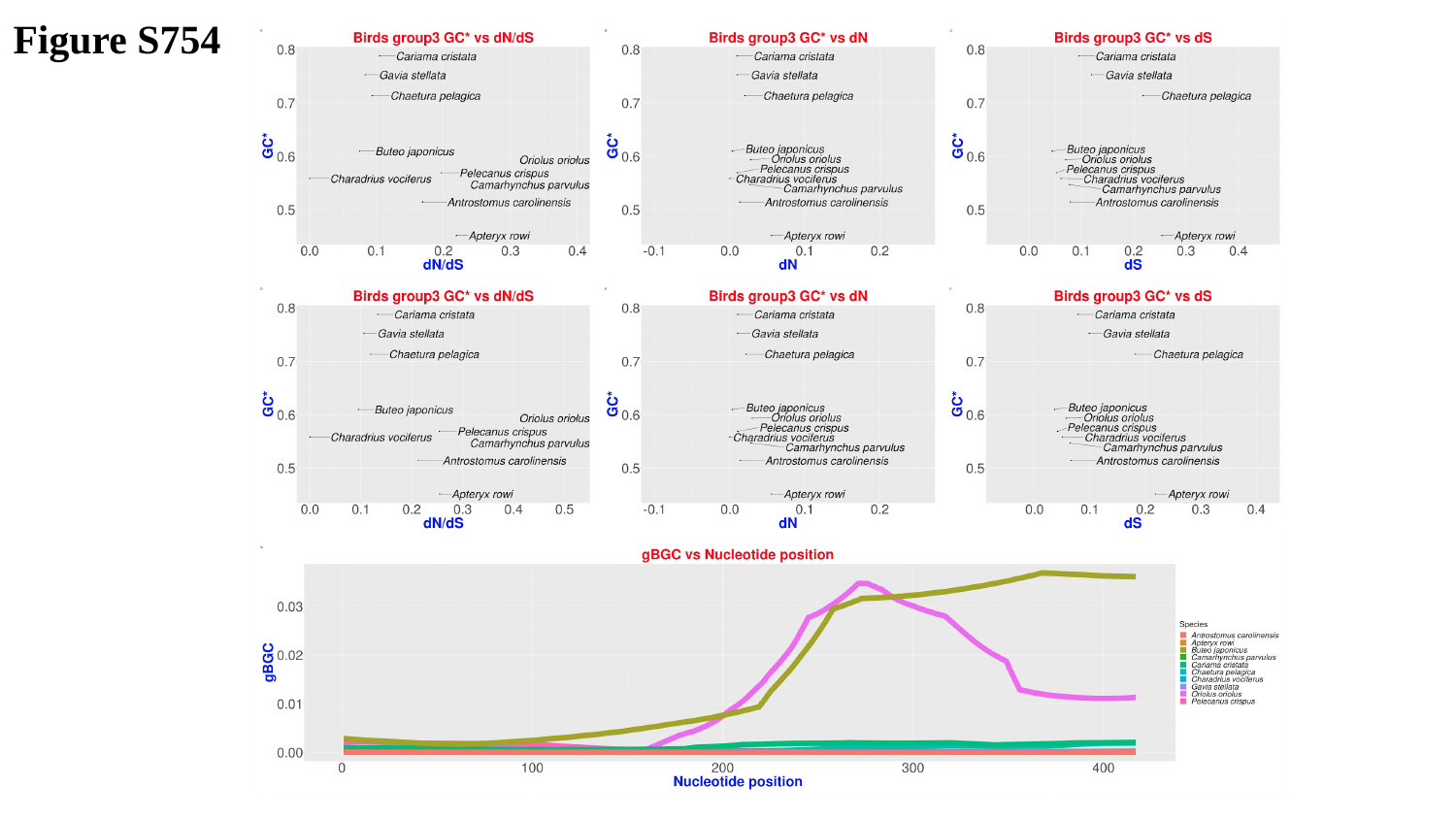

Figure S754

### Slide 755
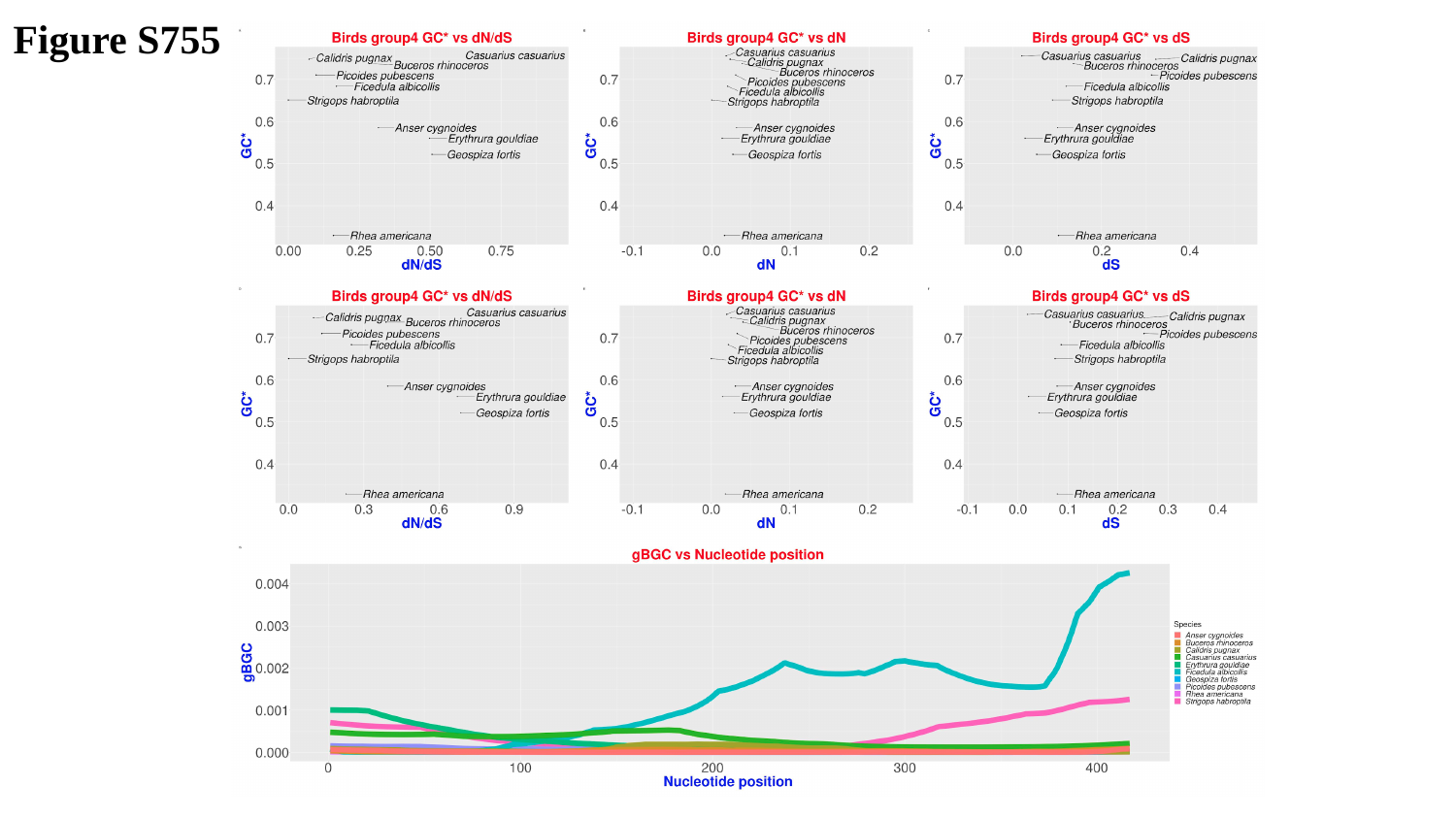

Figure S755

### Slide 756
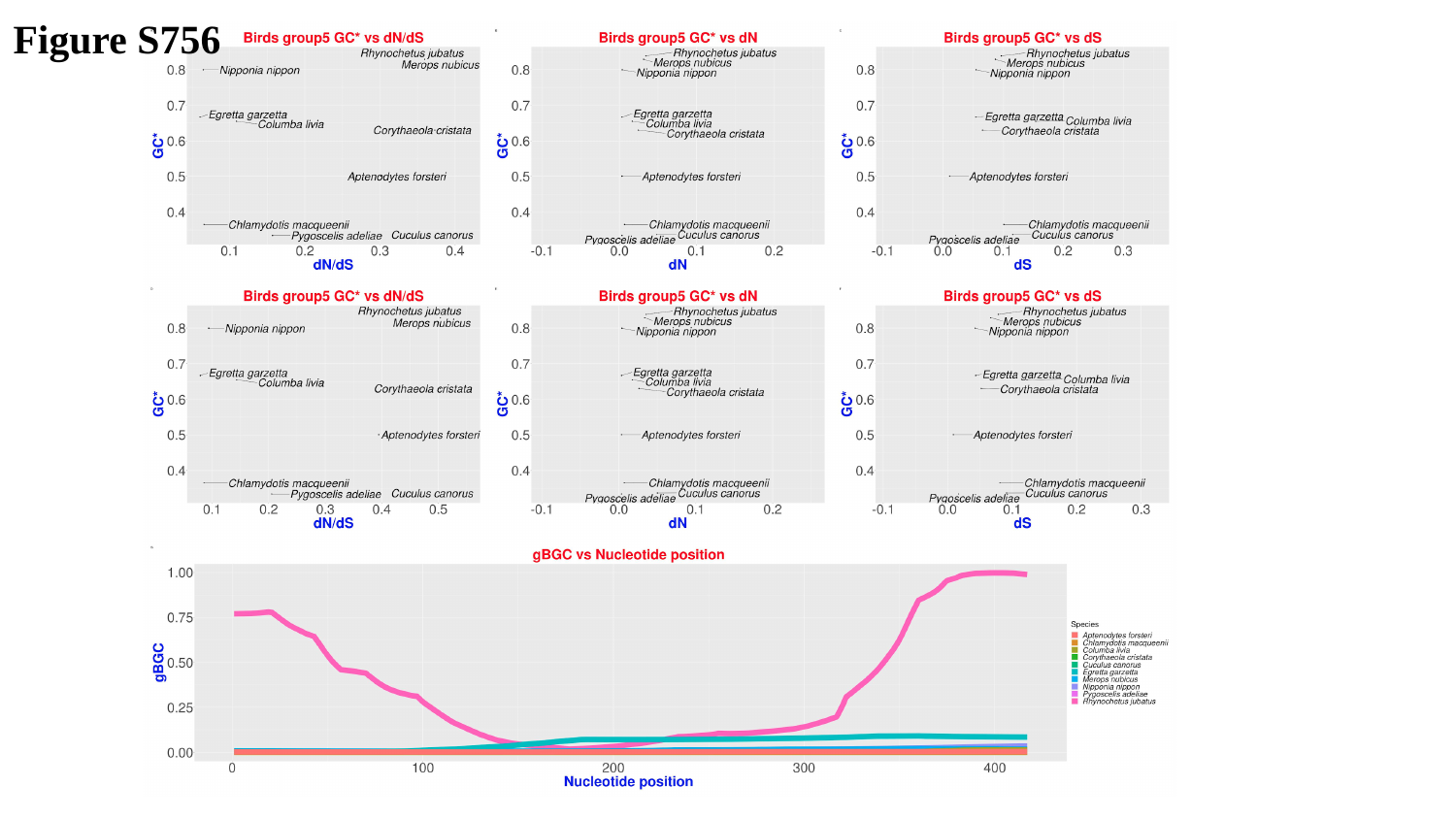

Figure S756

### Slide 757
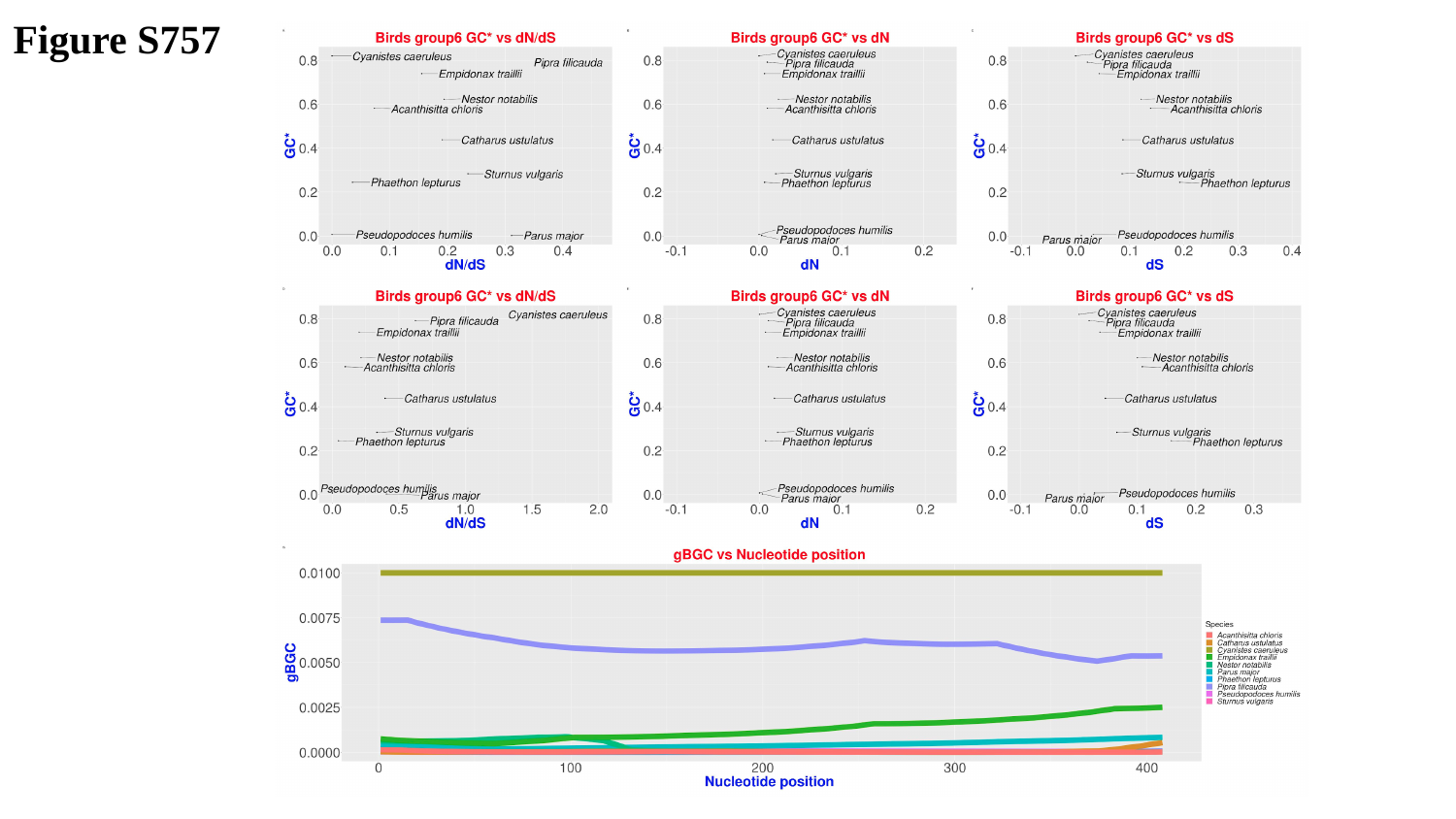

Figure S757

### Slide 758
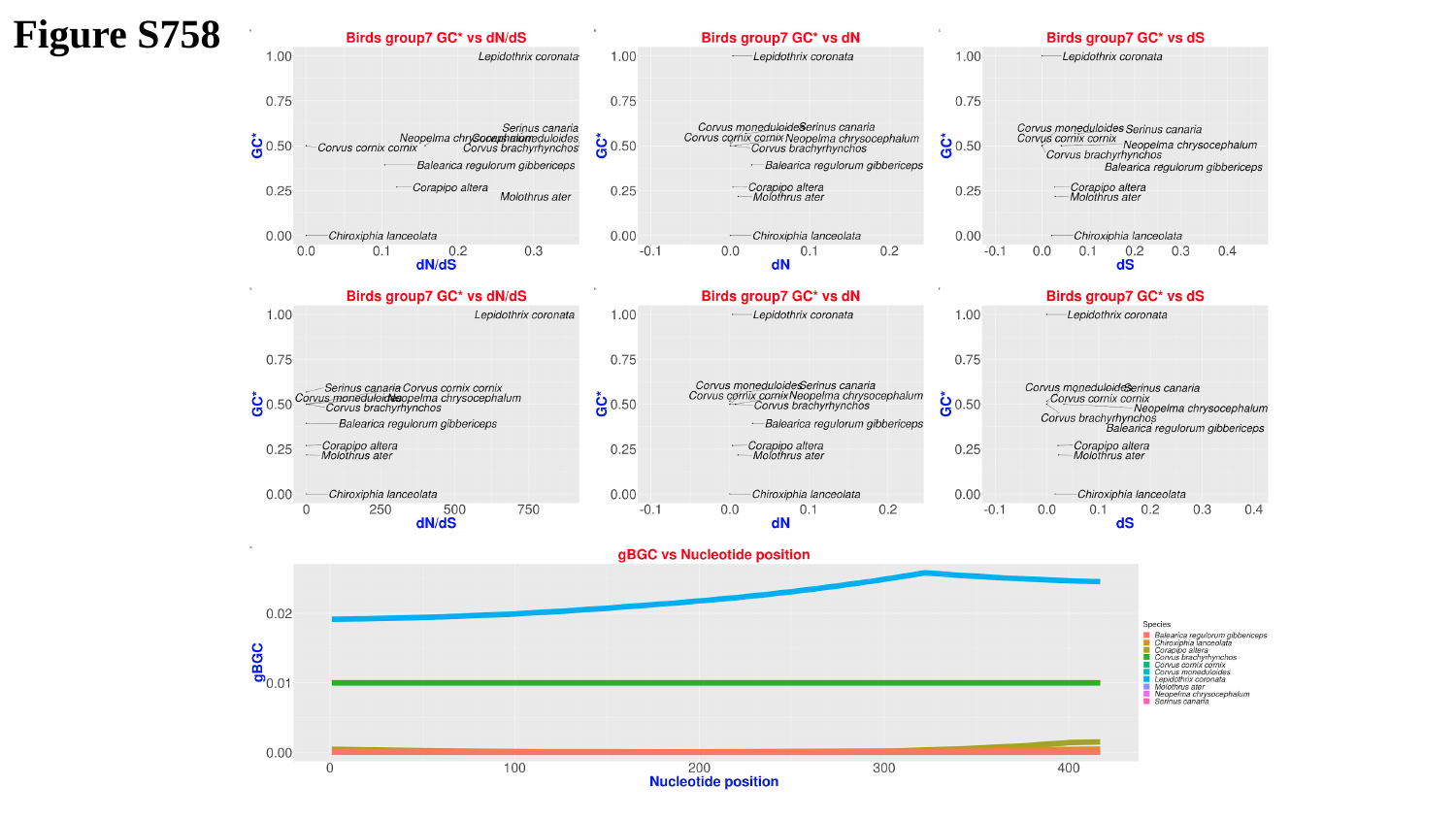

Figure S758

### Slide 759
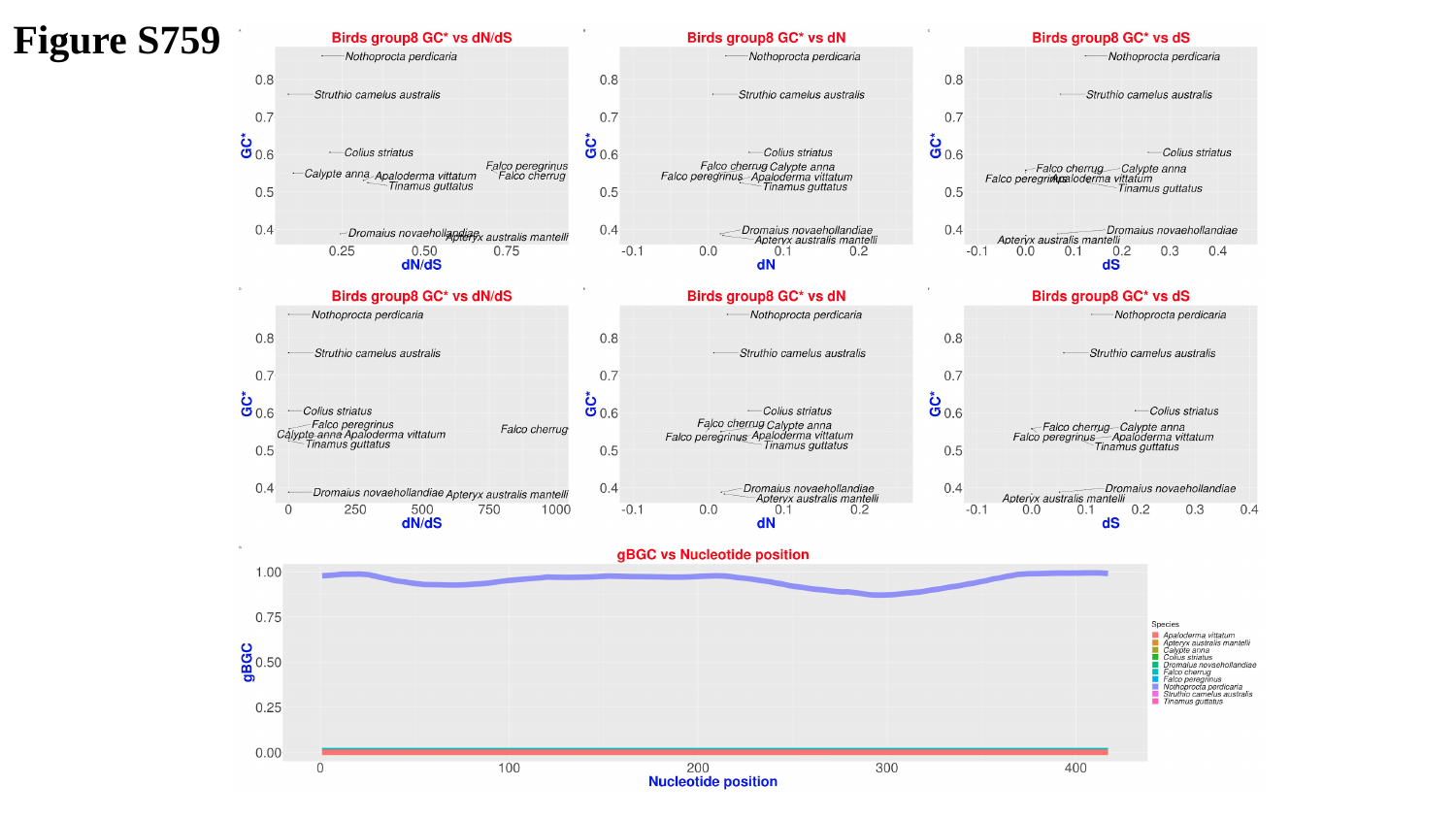

Figure S759

### Slide 760
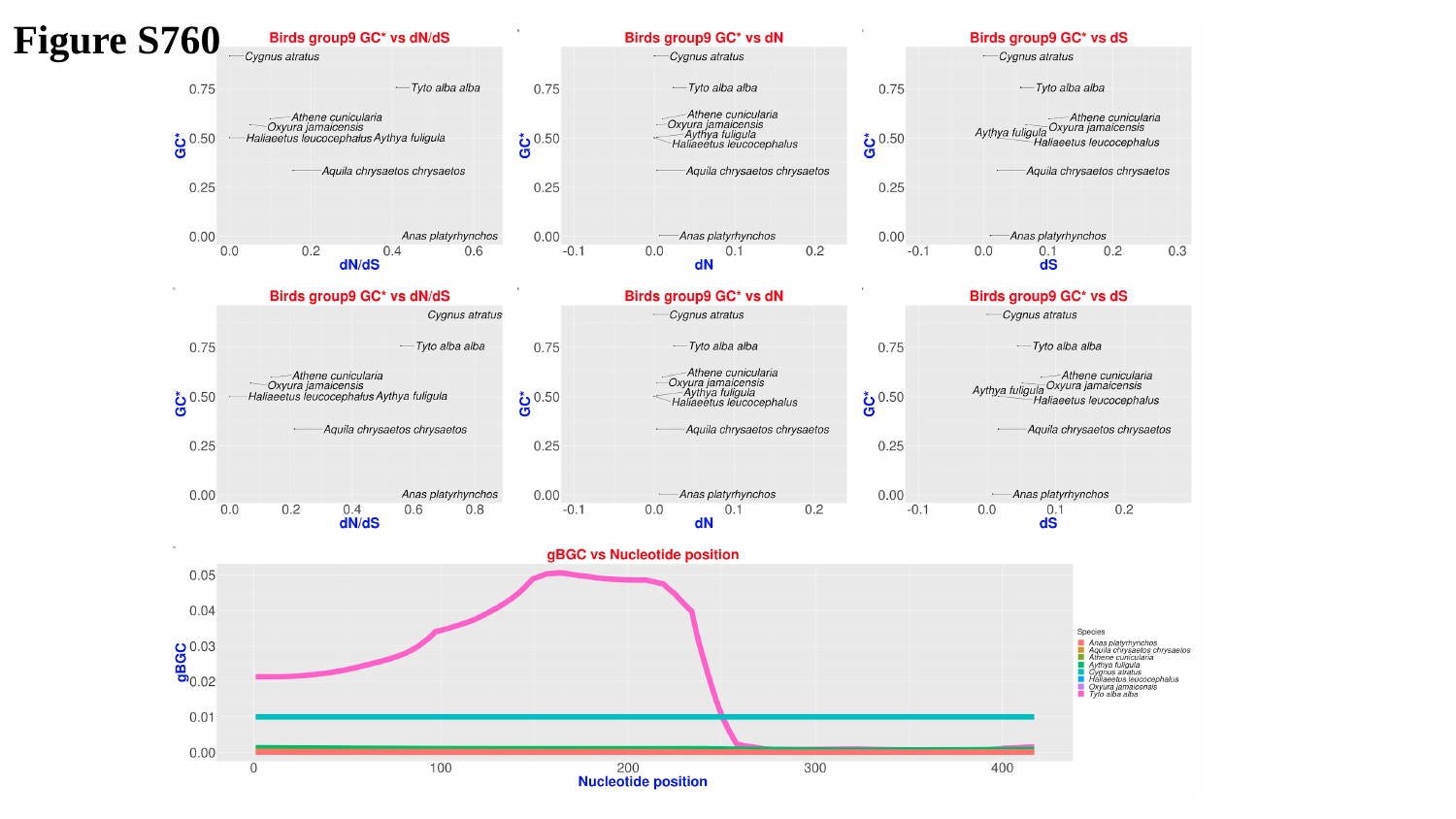

Figure S760

### Slide 761
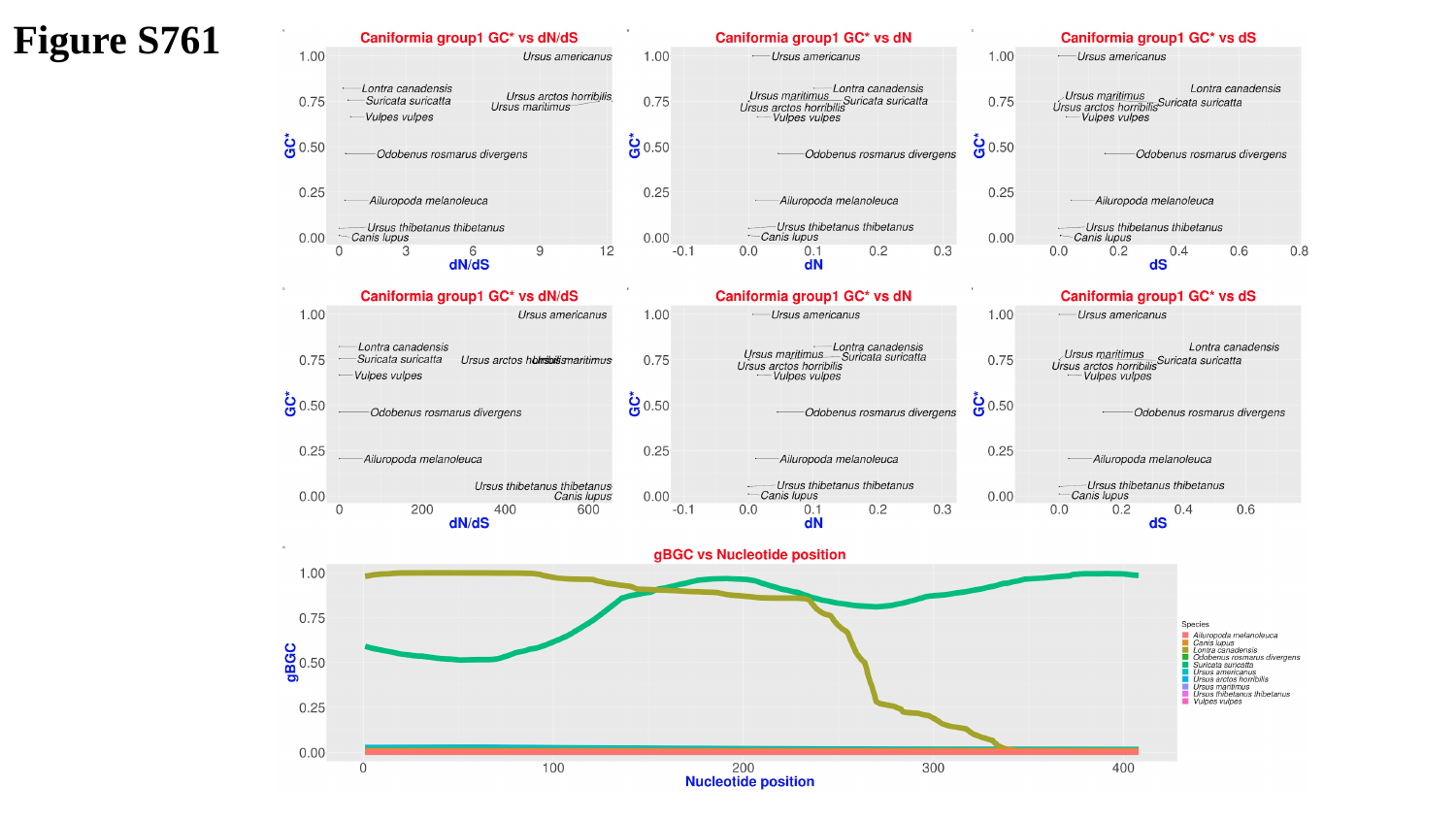

Figure S761

### Slide 762
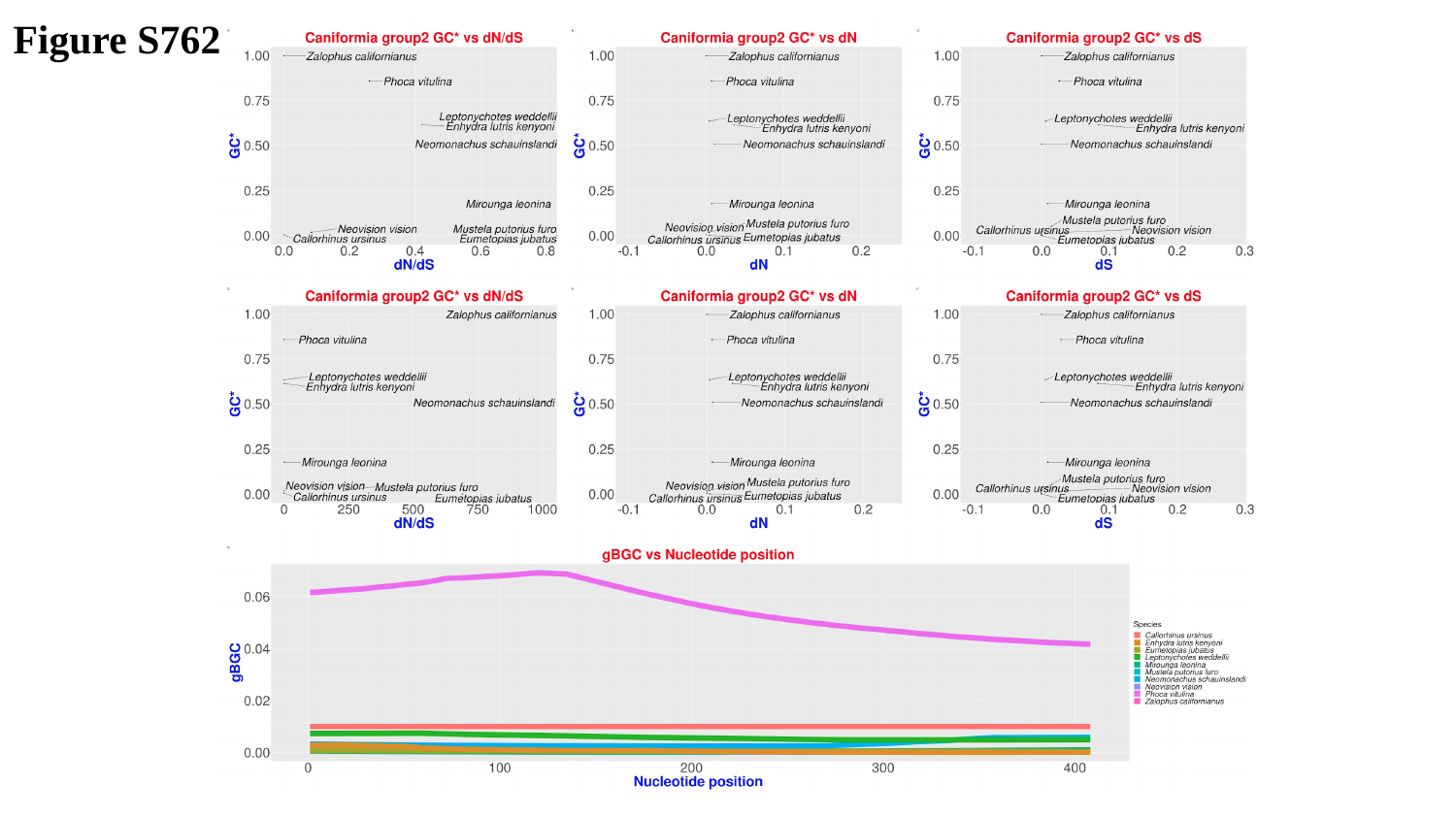

Figure S762

### Slide 763
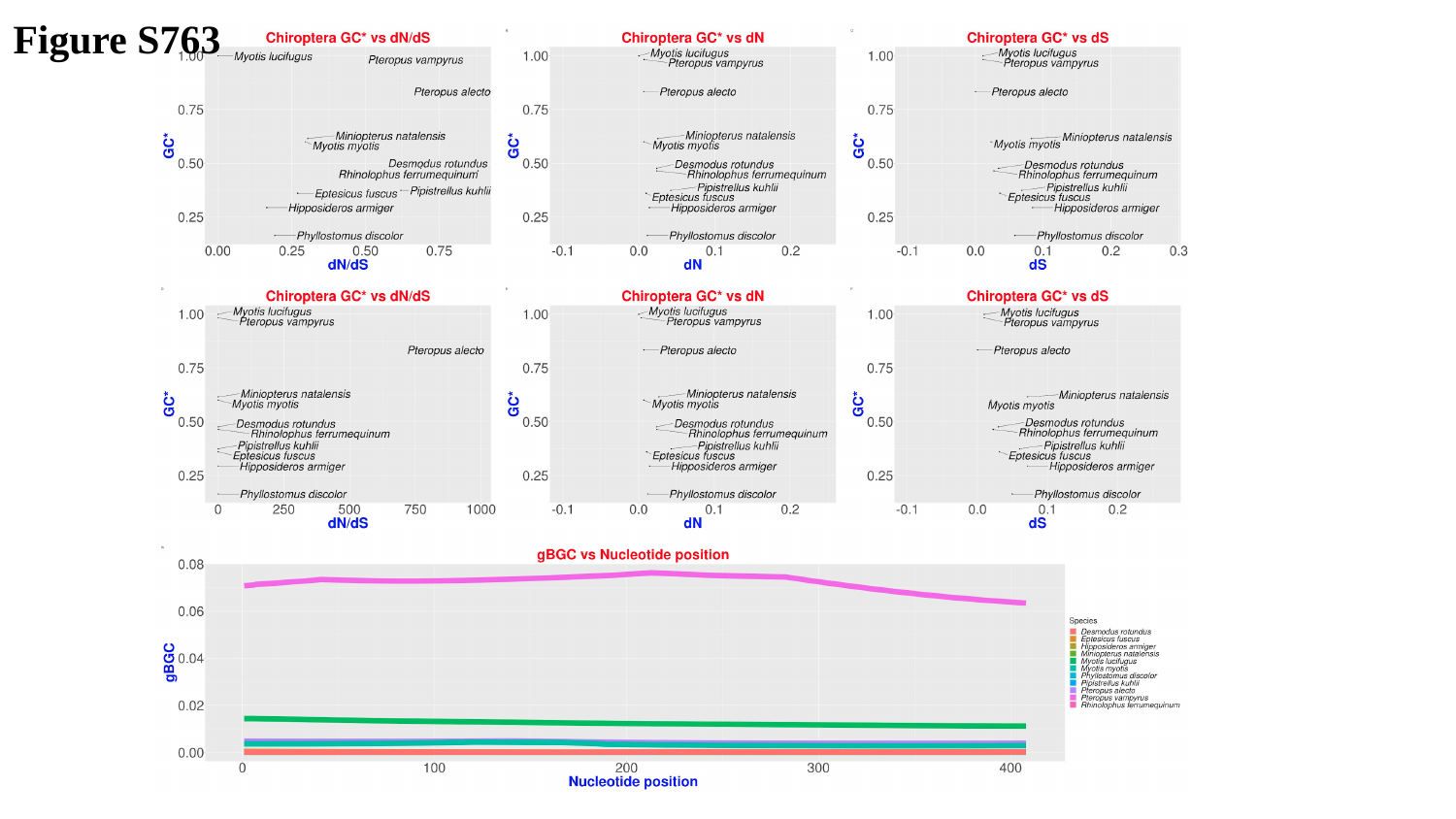

Figure S763

### Slide 764
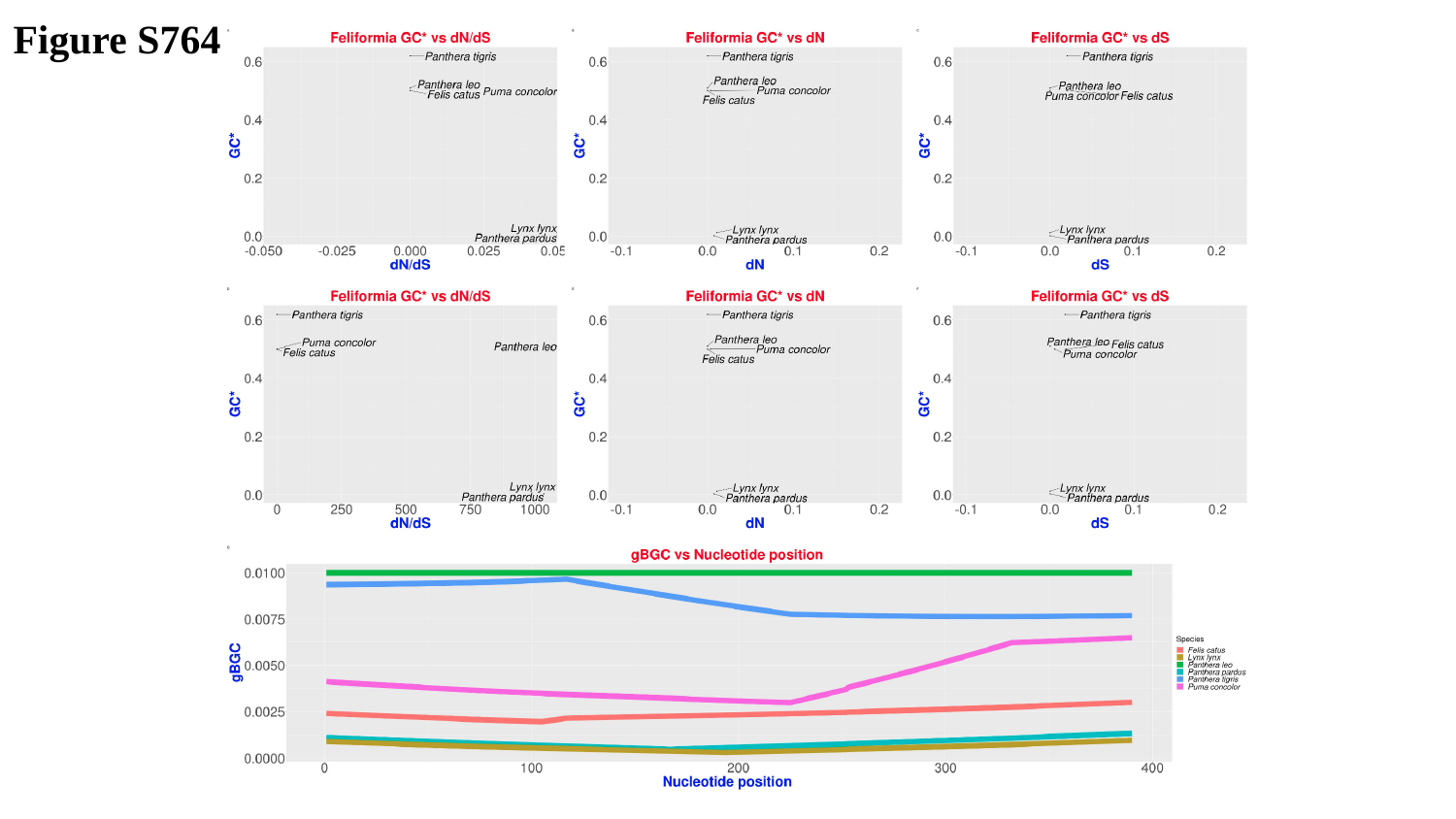

Figure S764

### Slide 765
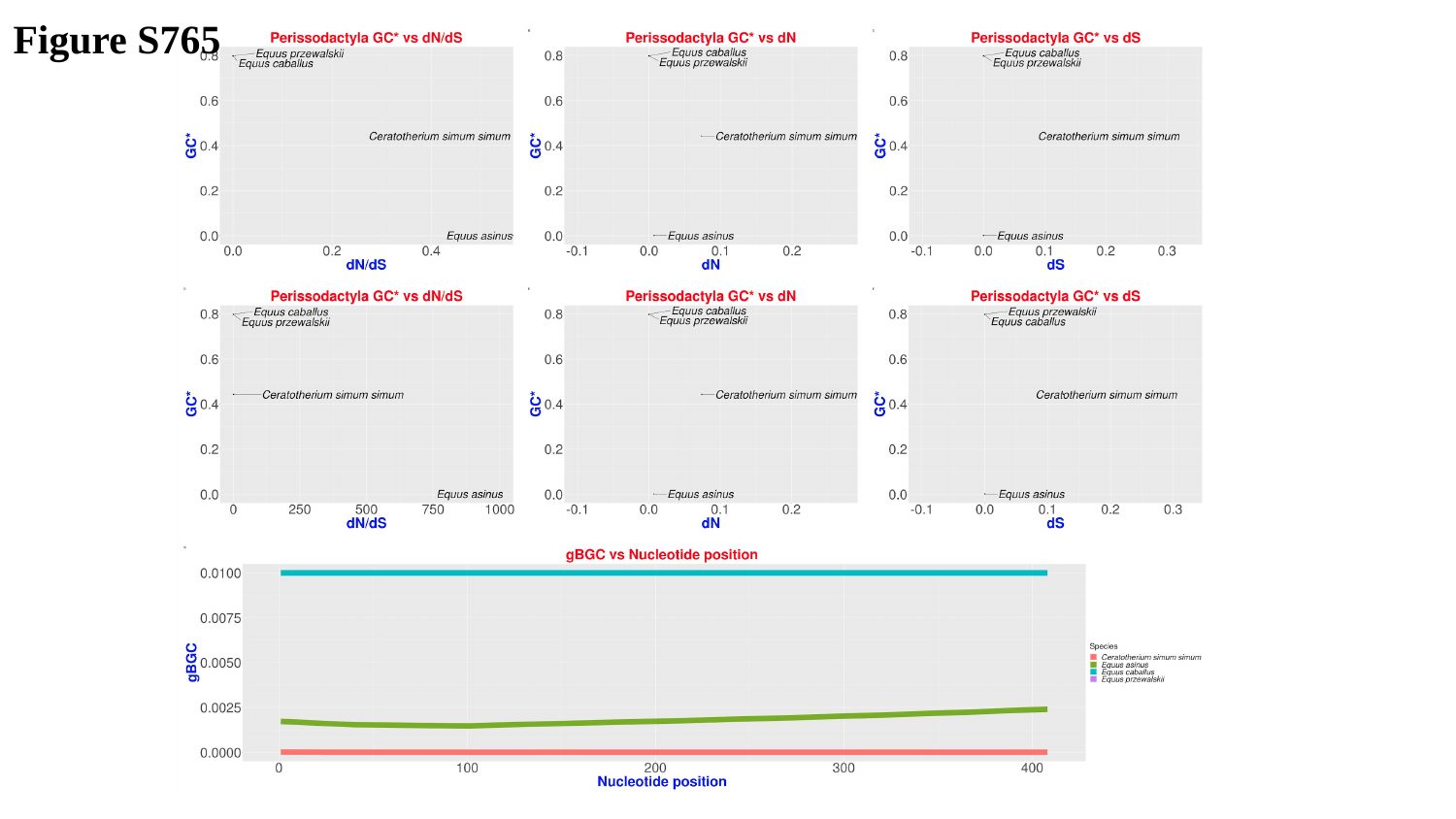

Figure S765

### Slide 766
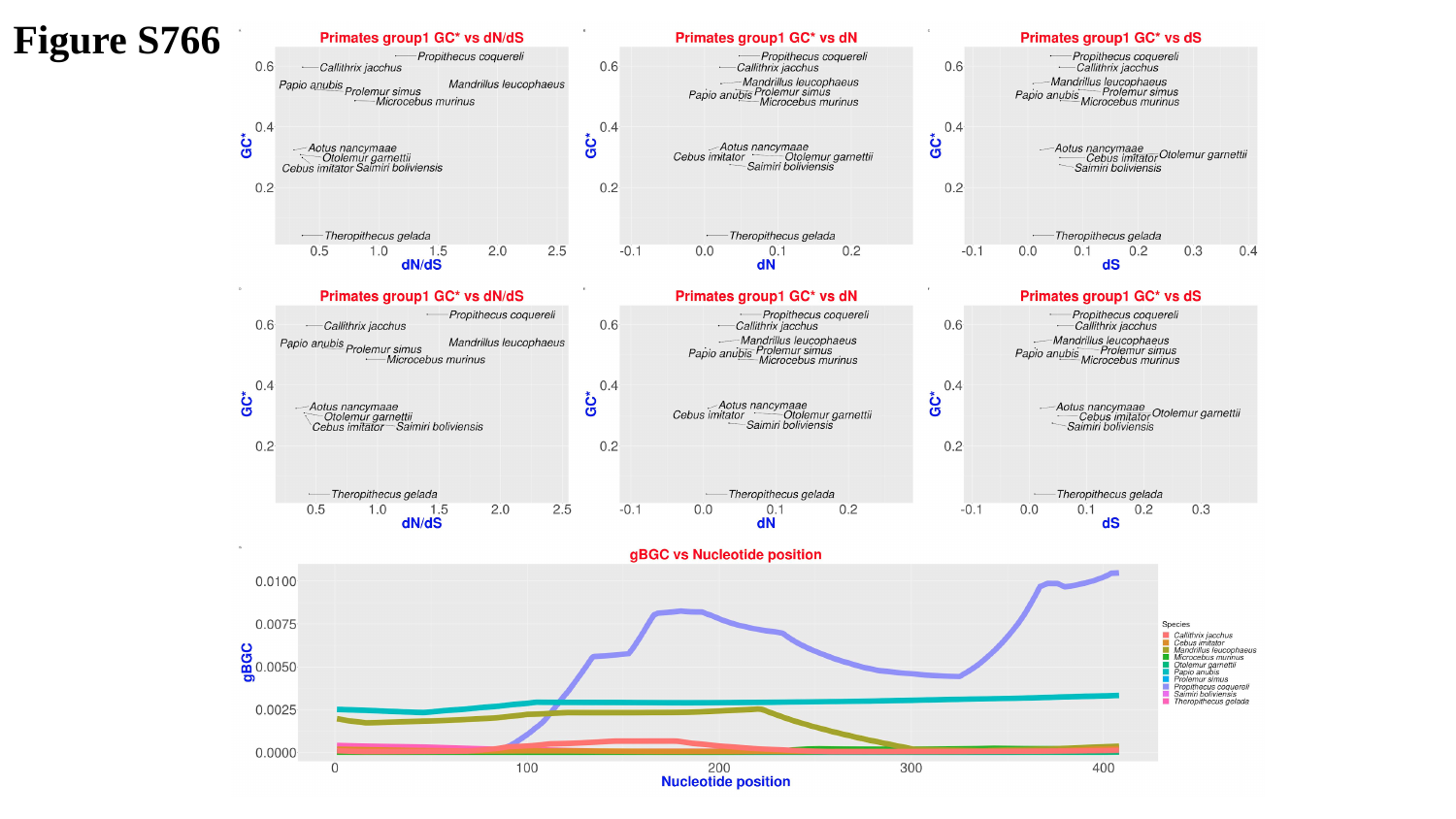

Figure S766

### Slide 767
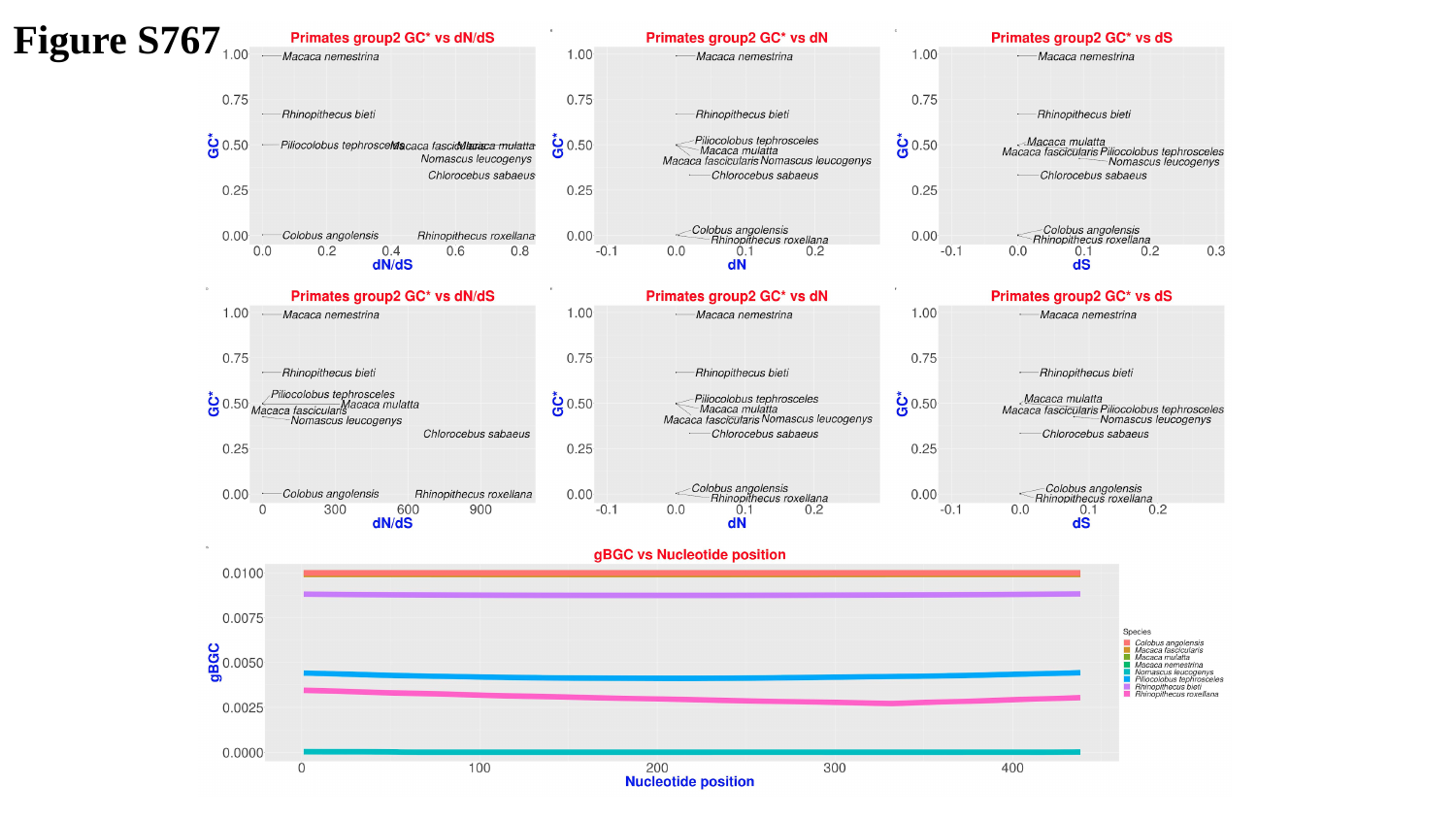

Figure S767

### Slide 768
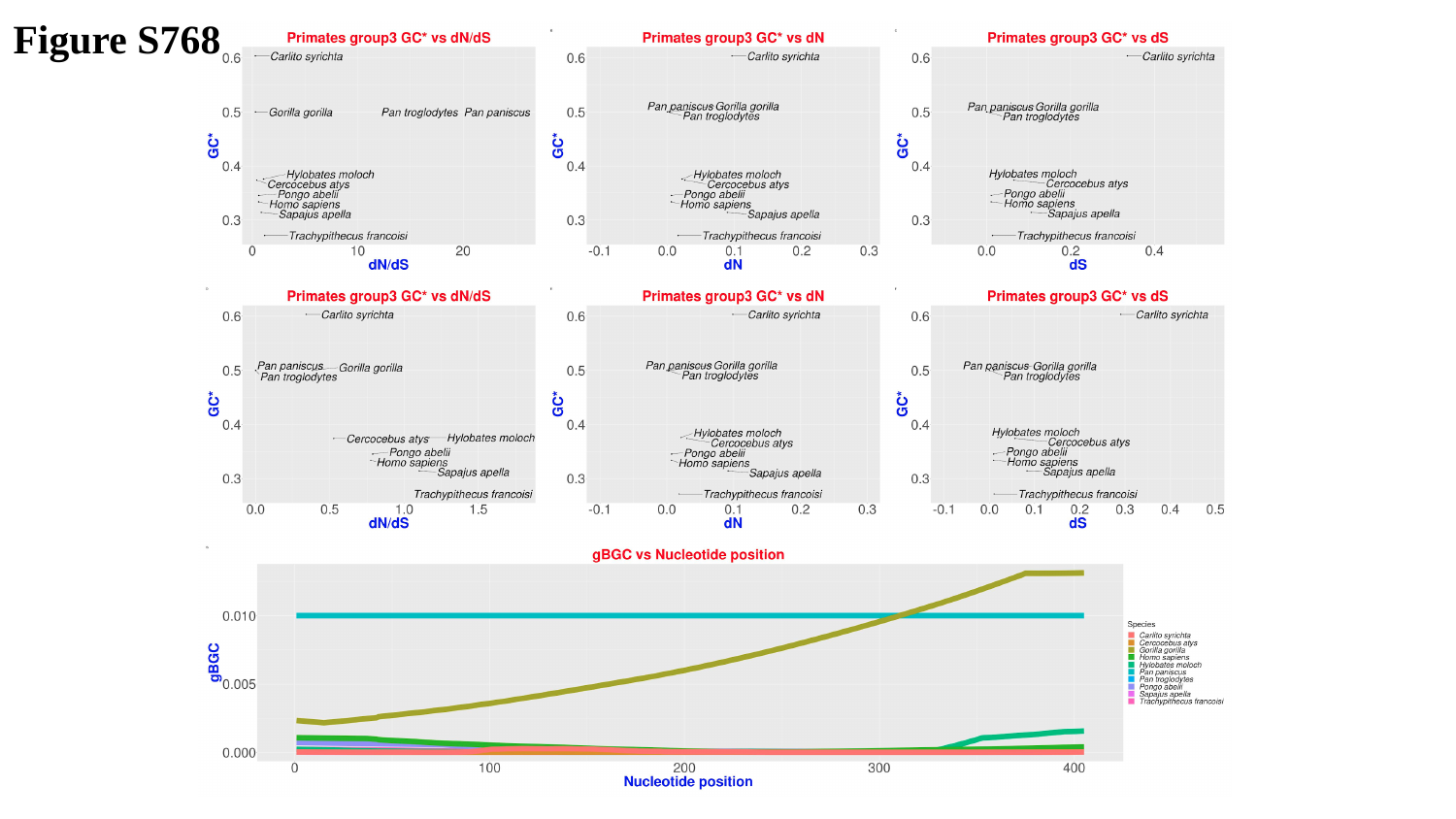

Figure S768

### Slide 769
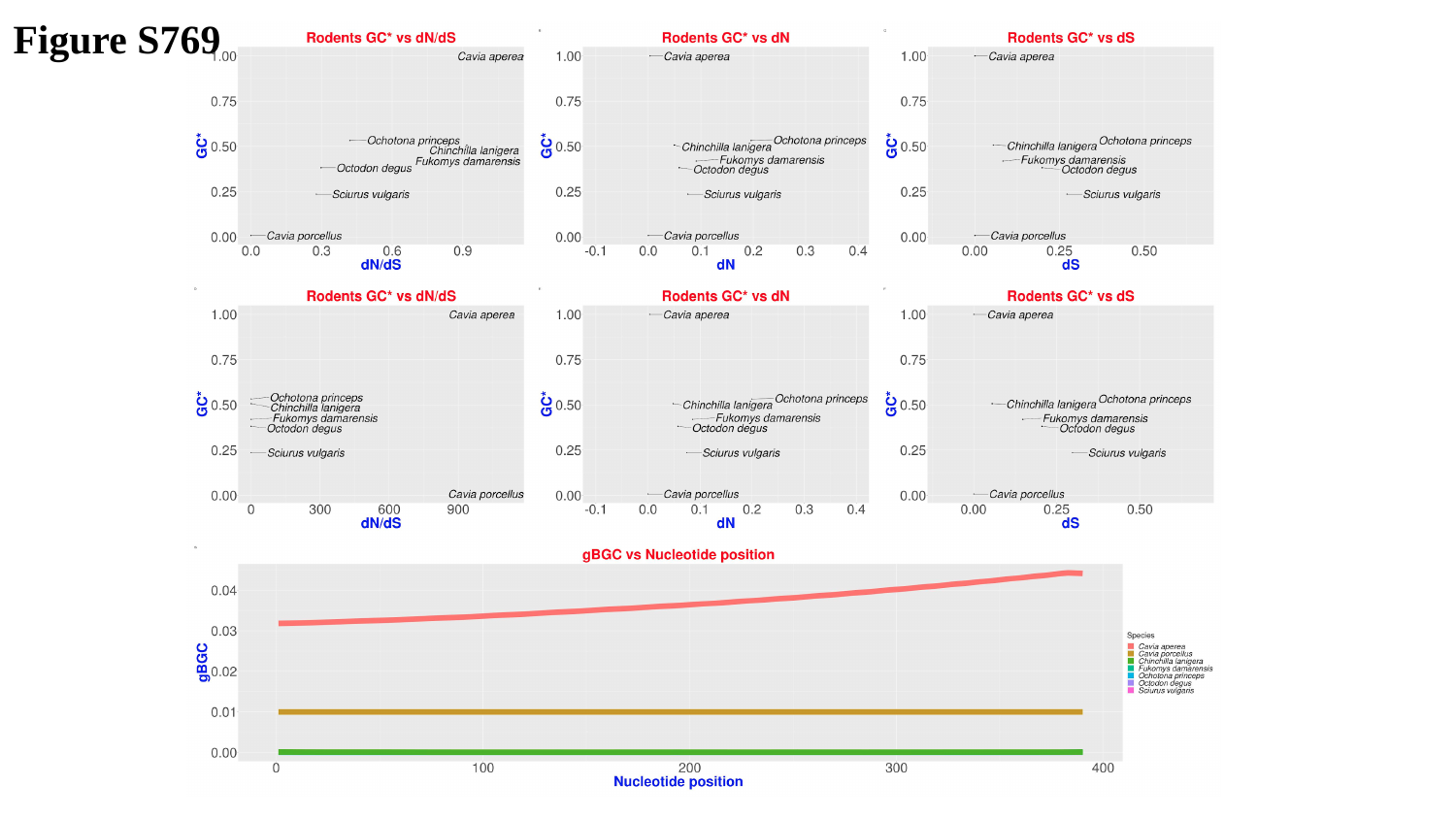

Figure S769

### Slide 770
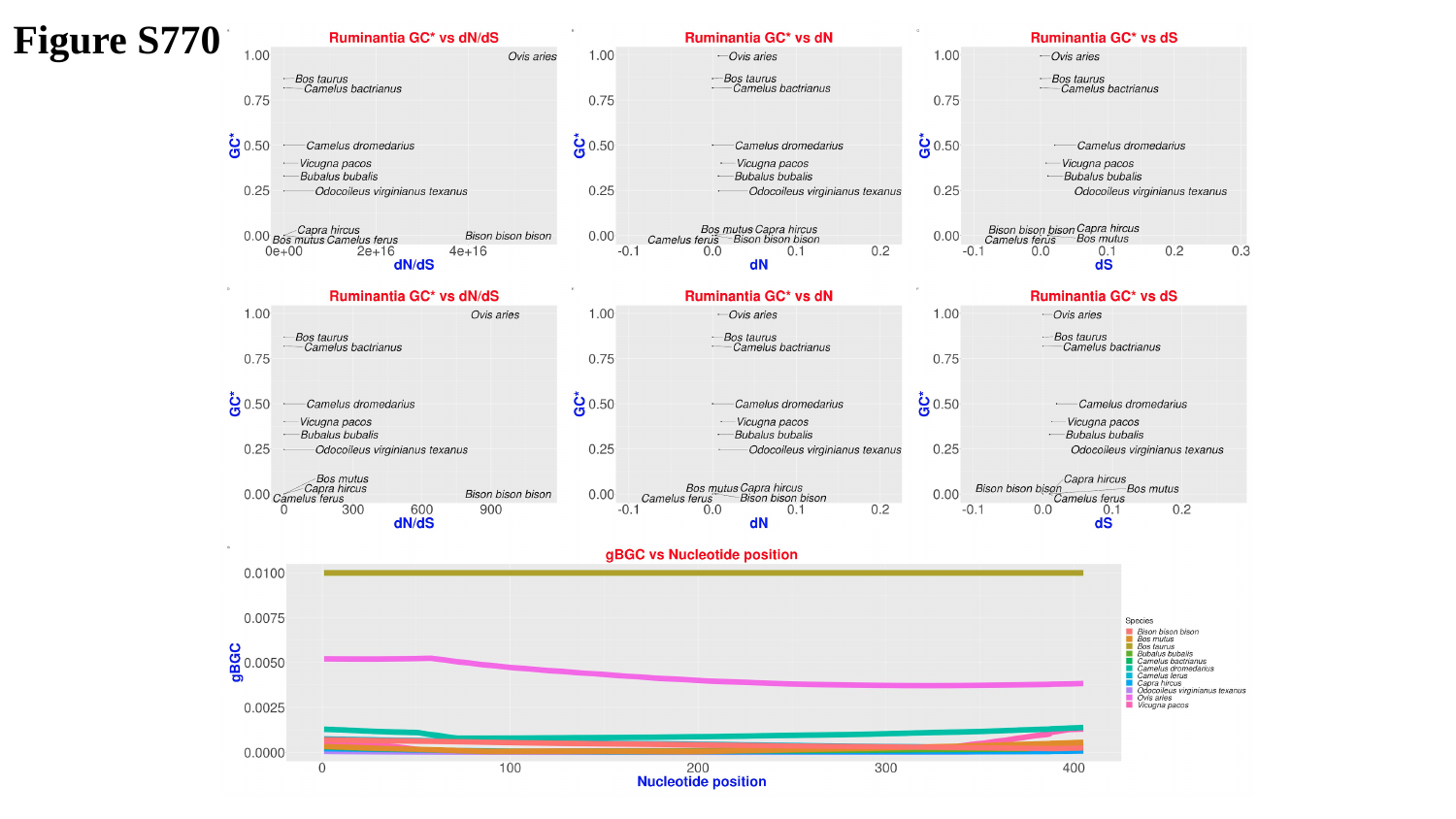

Figure S770

### Slide 771
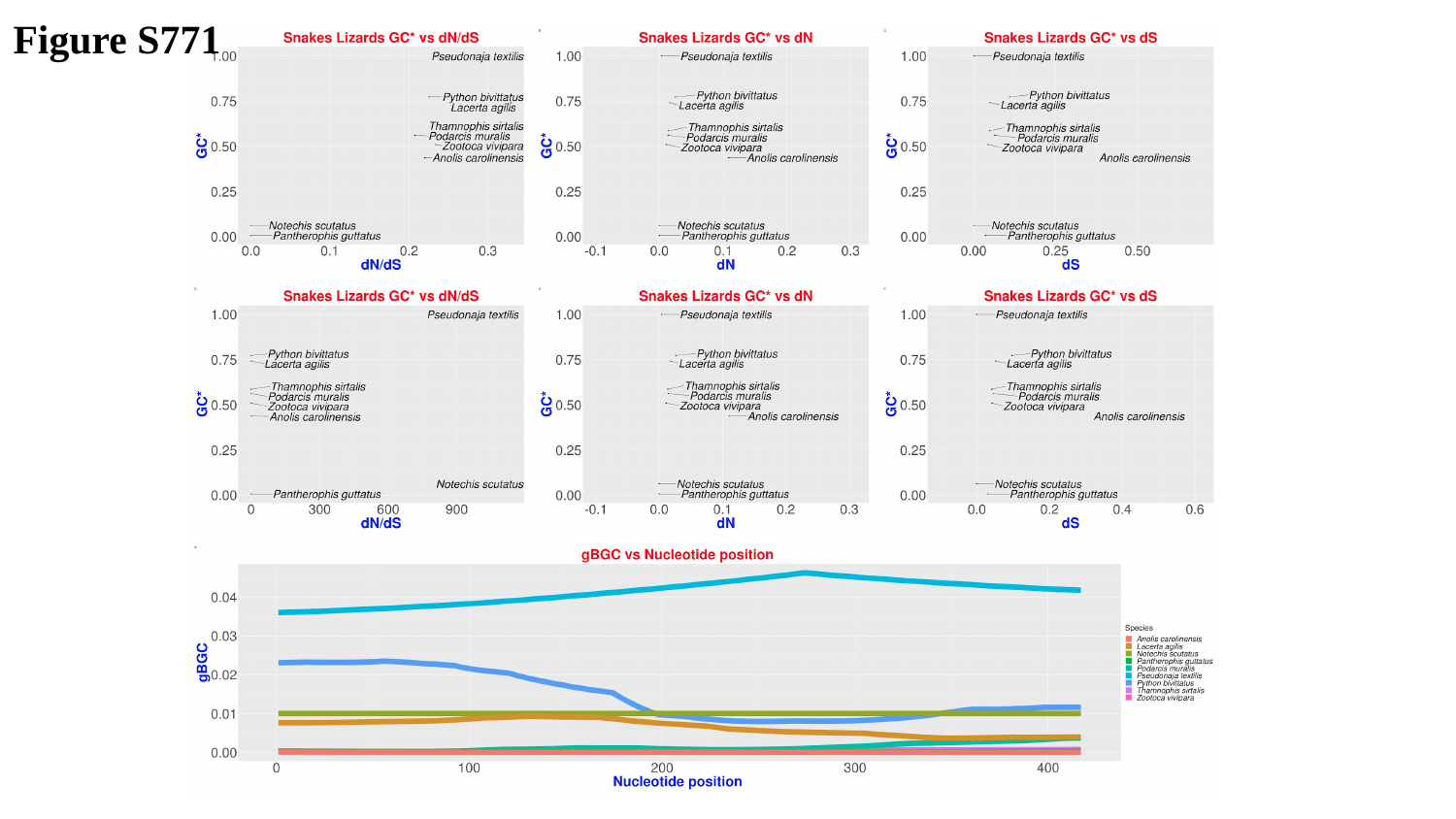

Figure S771

### Slide 772
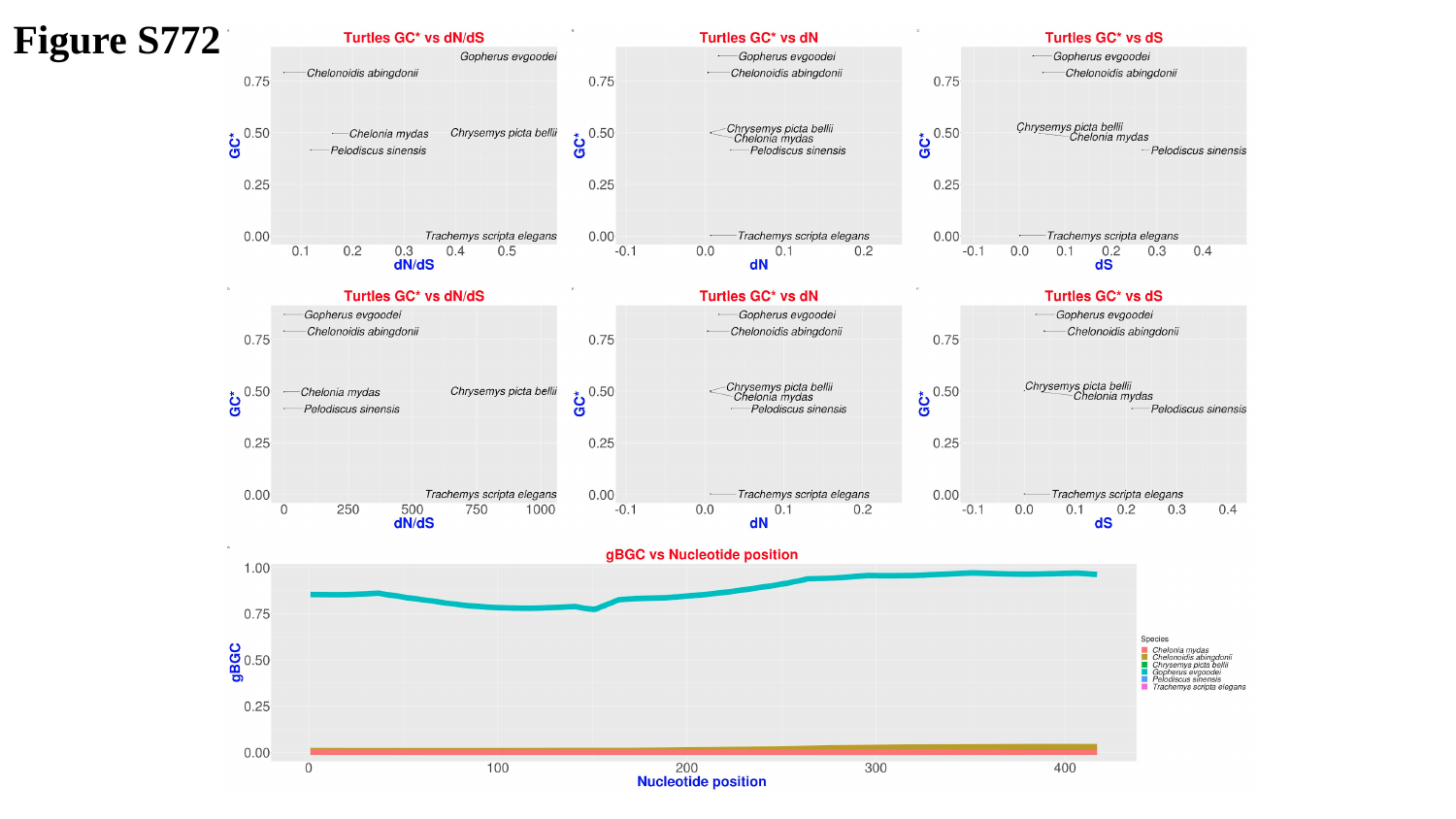

Figure S772

### Slide 773
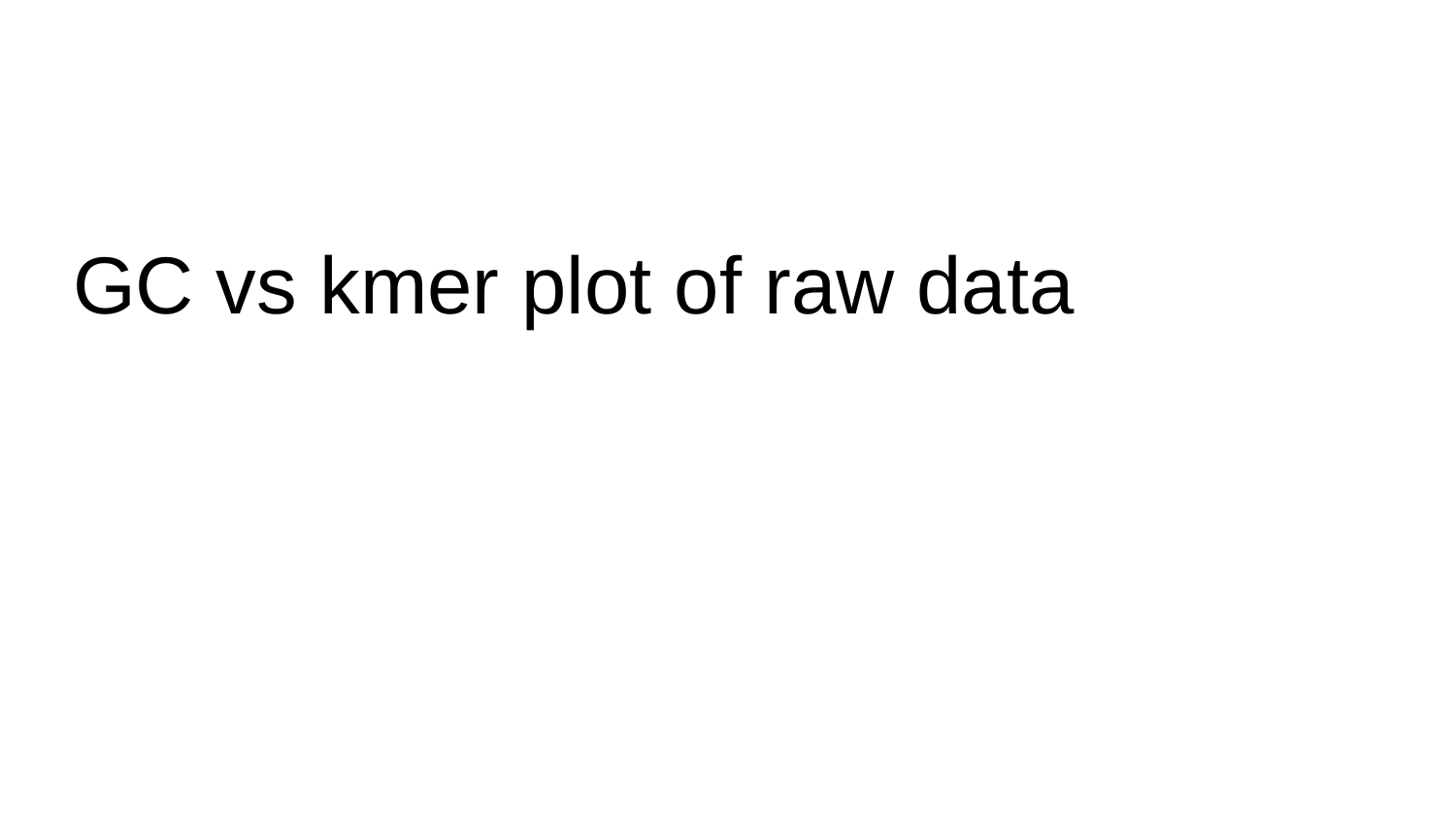

#### GC vs kmer plot of raw data

### Slide 774
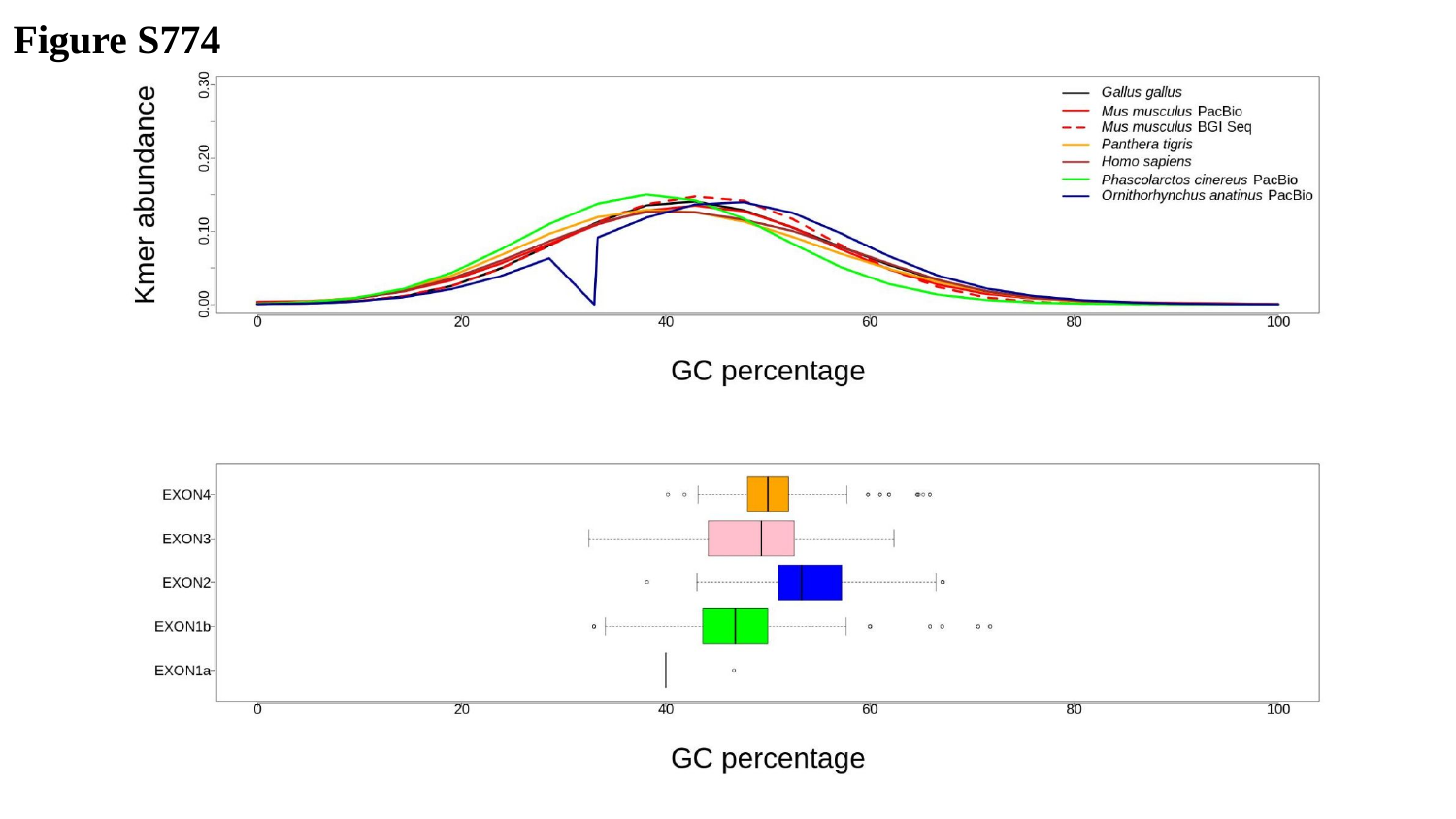

Figure S774

### Slide 775
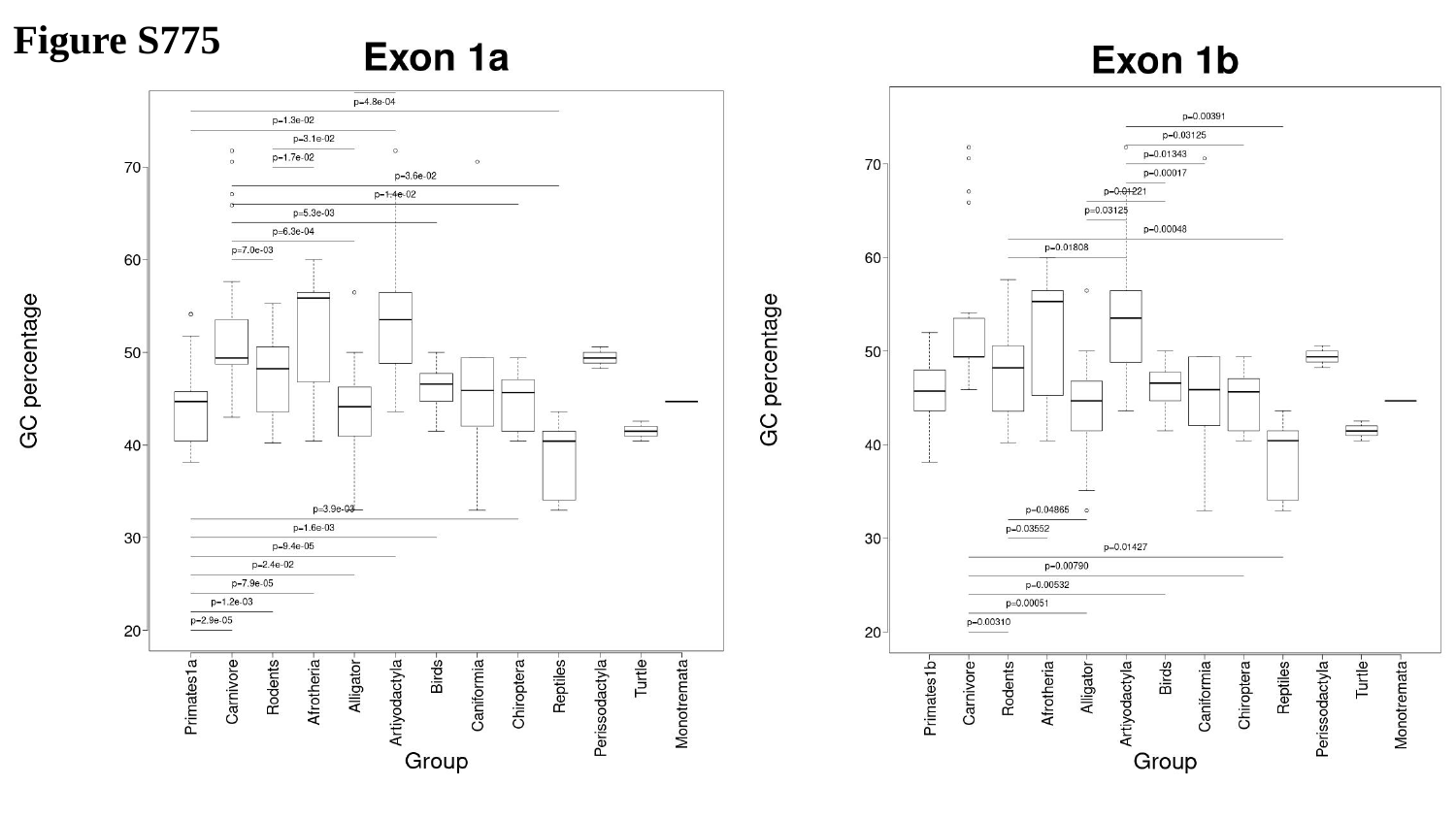

Figure S775

### Slide 776
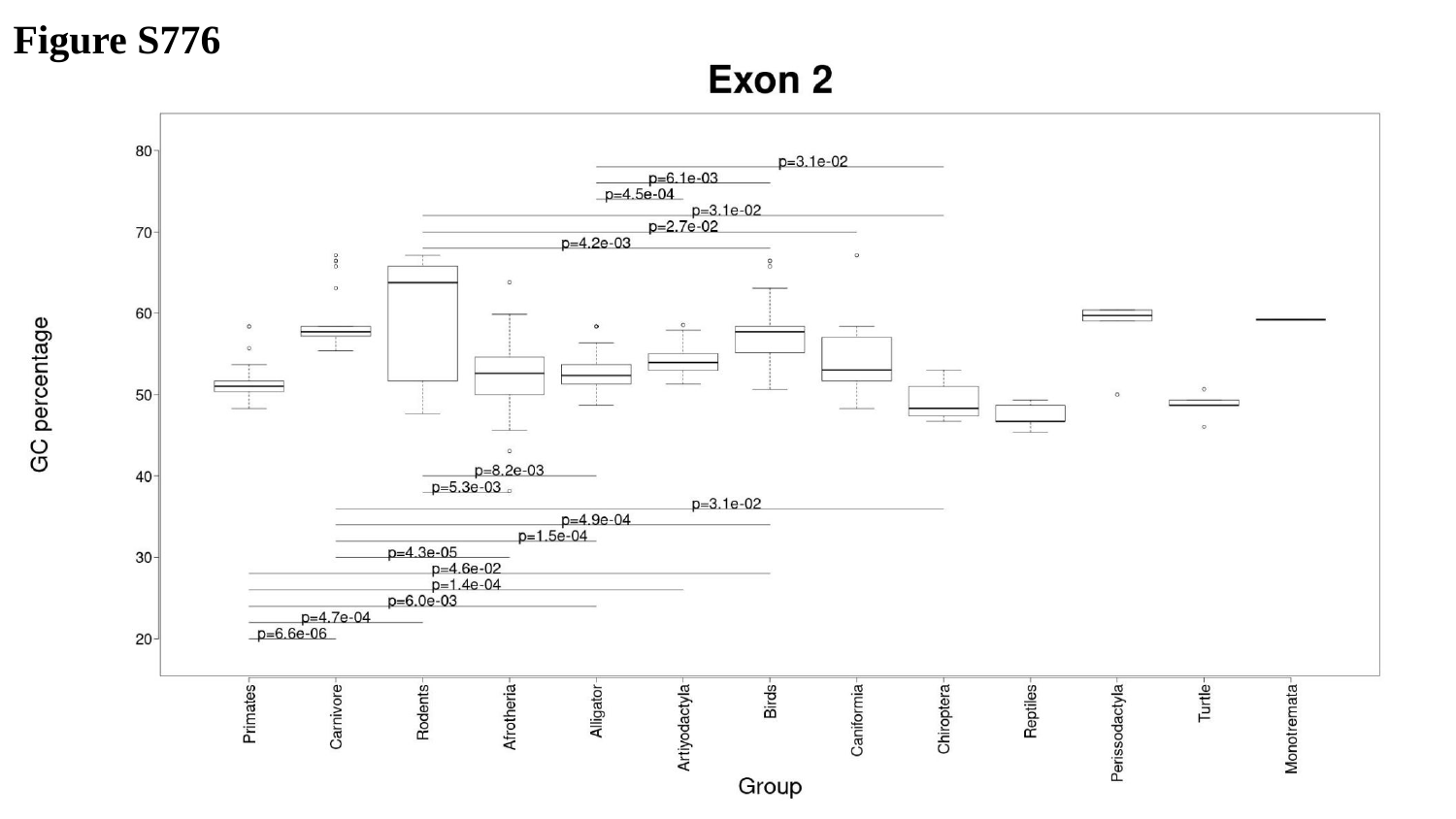

Figure S776

### Slide 777
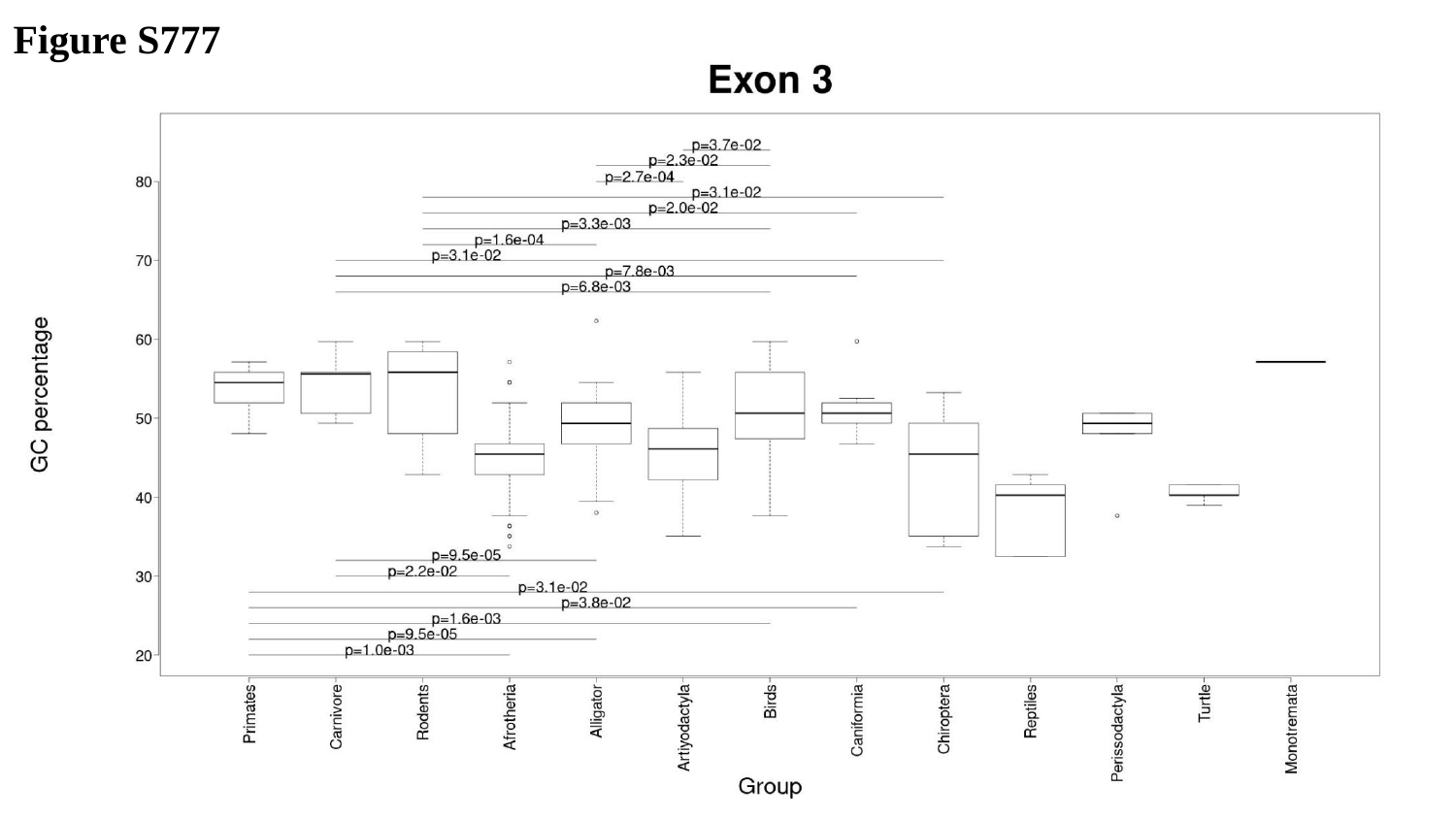

Figure S777

### Slide 778
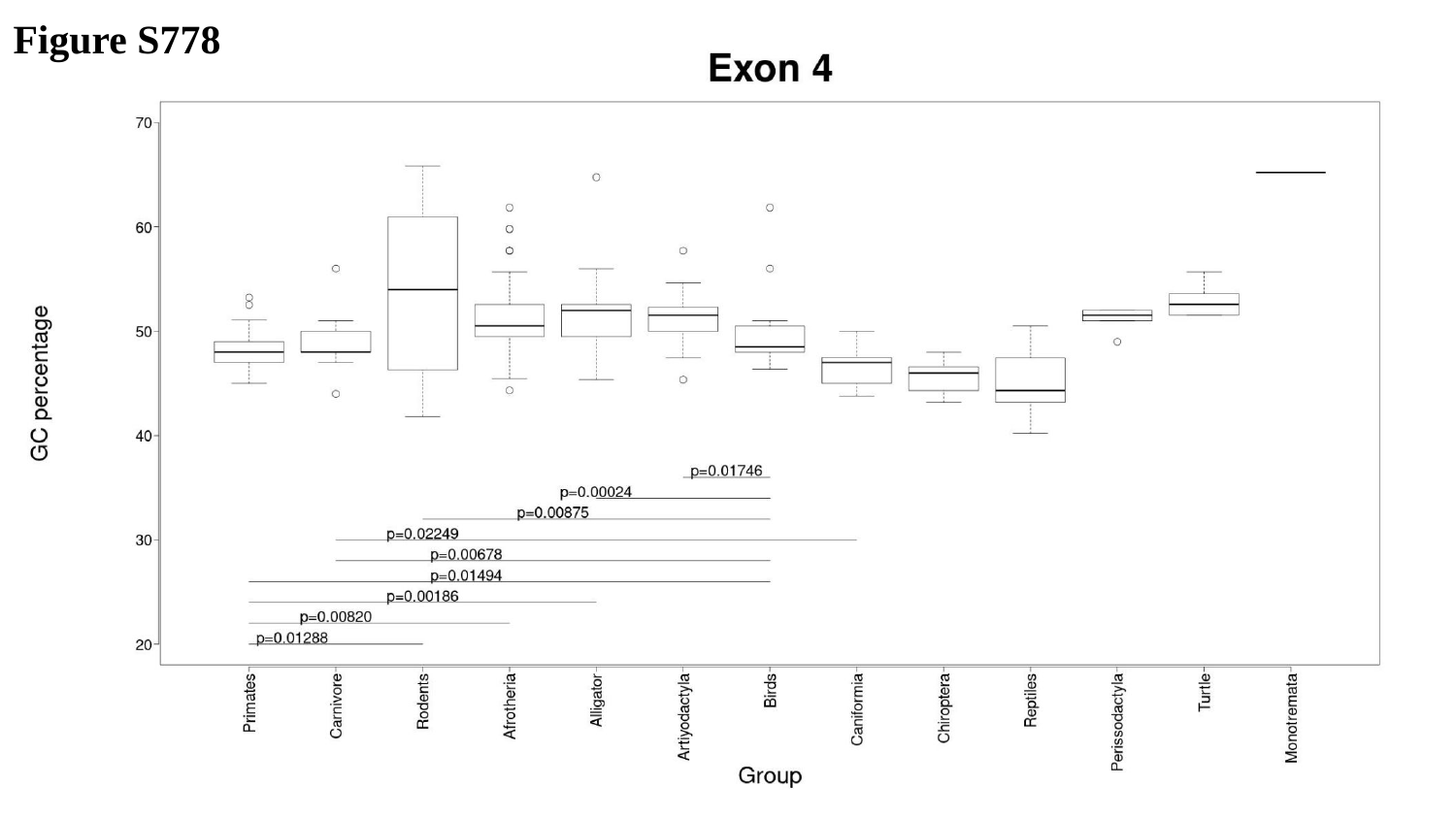

Figure S778

### Slide 779
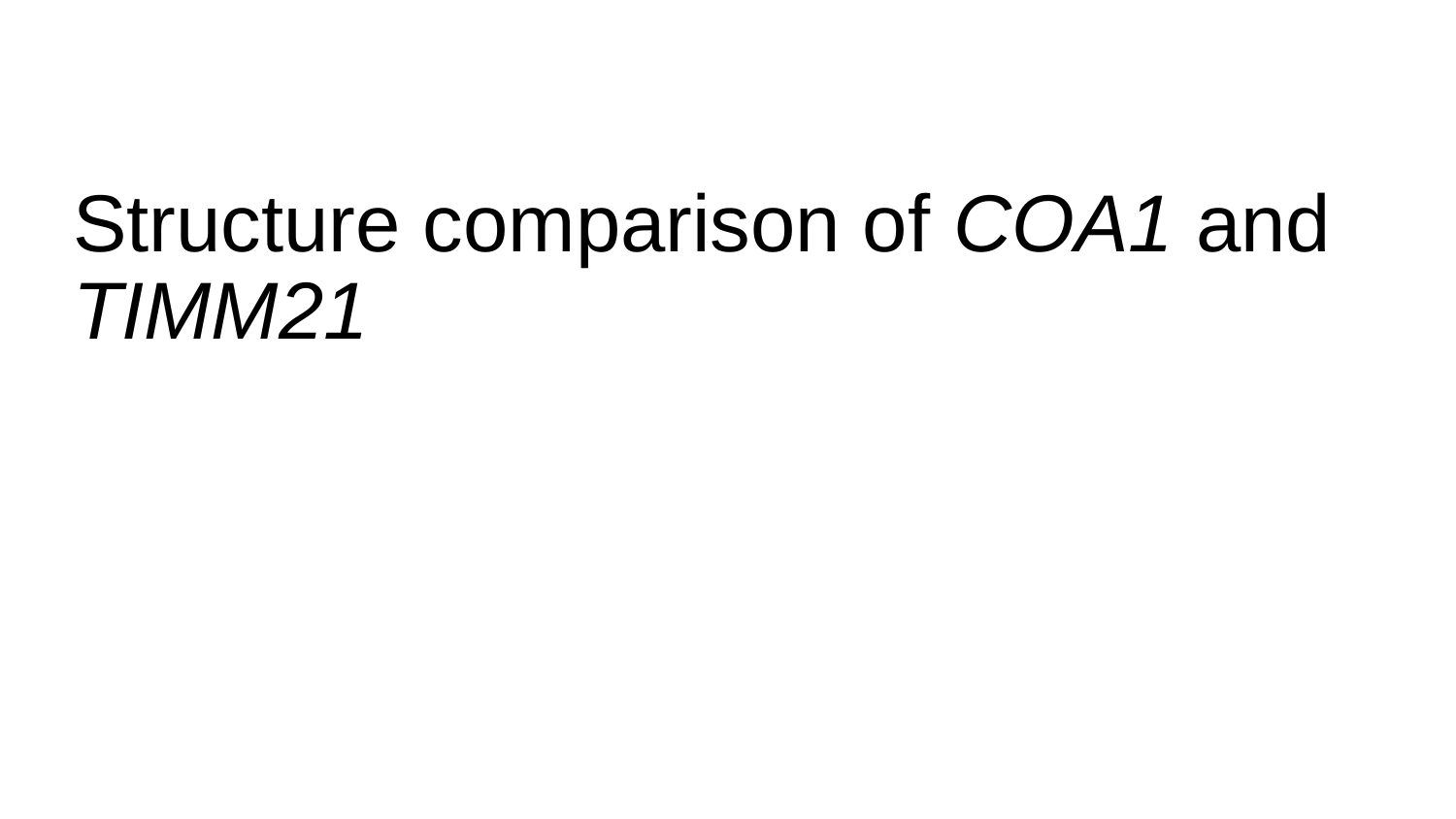

#### Structure comparison of COA1 and TIMM21

### Slide 780
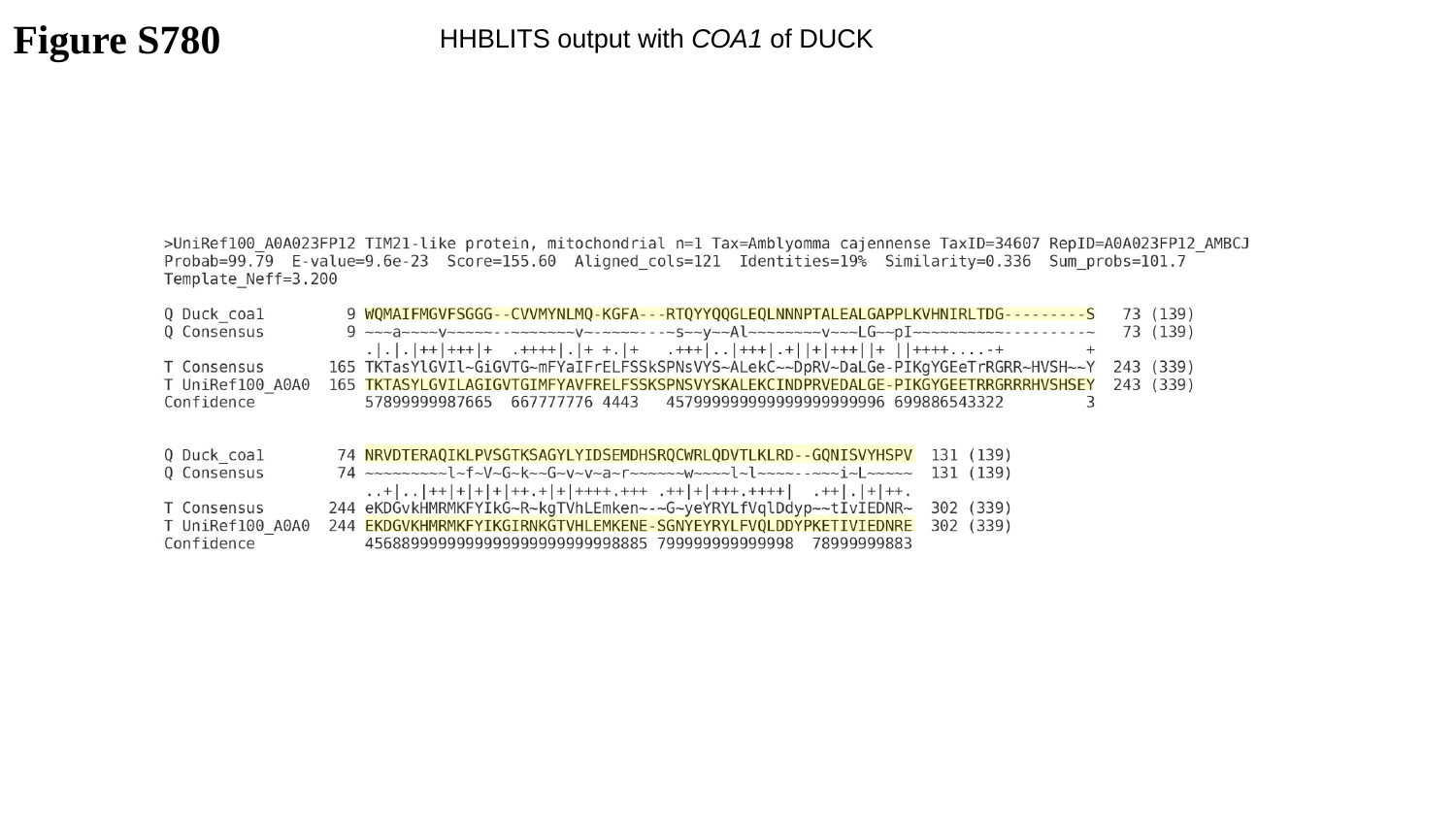

Figure S780
HHBLITS output with COA1 of DUCK

### Slide 781
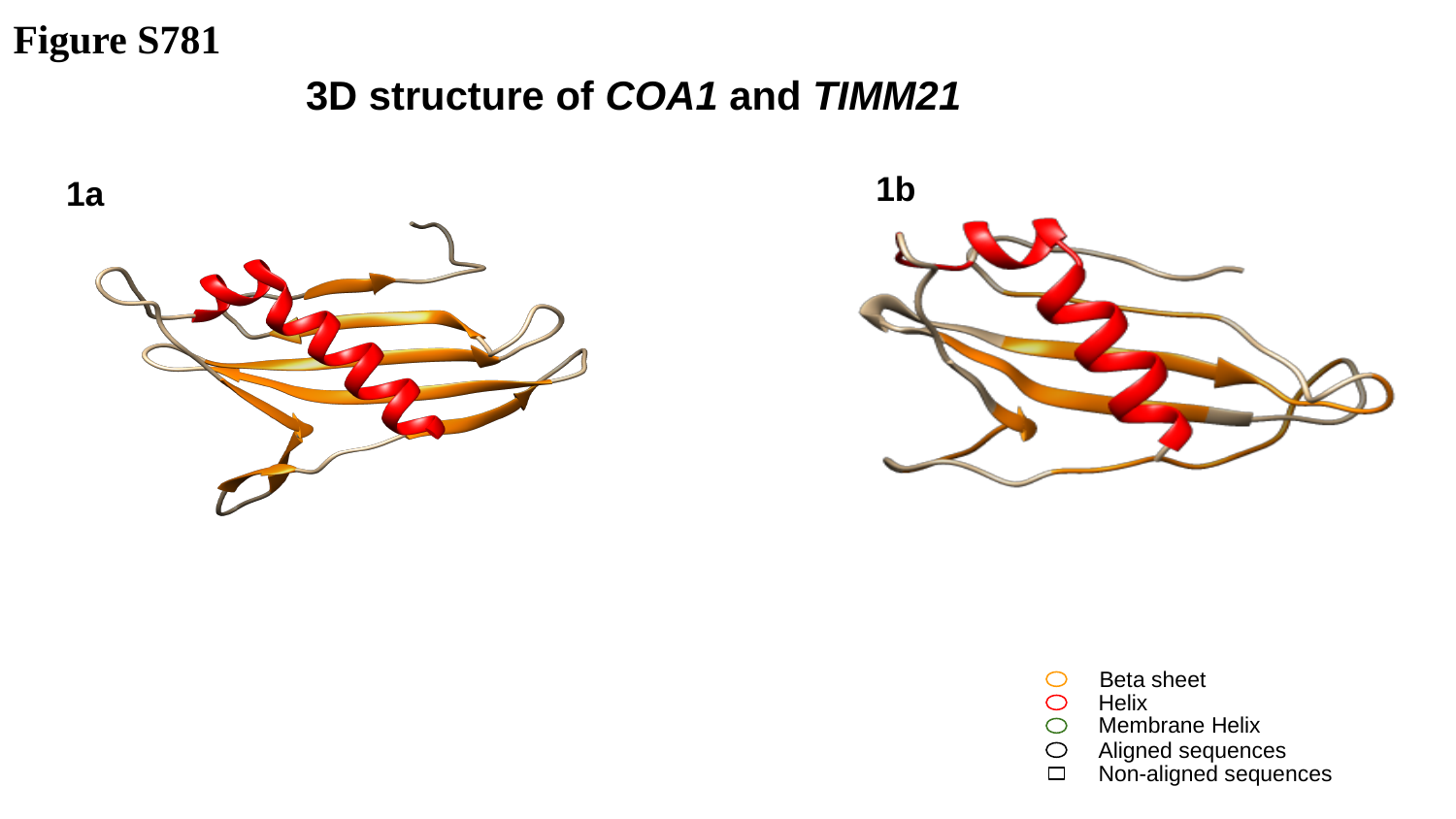

Figure S781
3D structure of COA1 and TIMM21
1b
1a
Beta sheet
Helix
Membrane Helix
Aligned sequences
Non-aligned sequences

### Slide 782
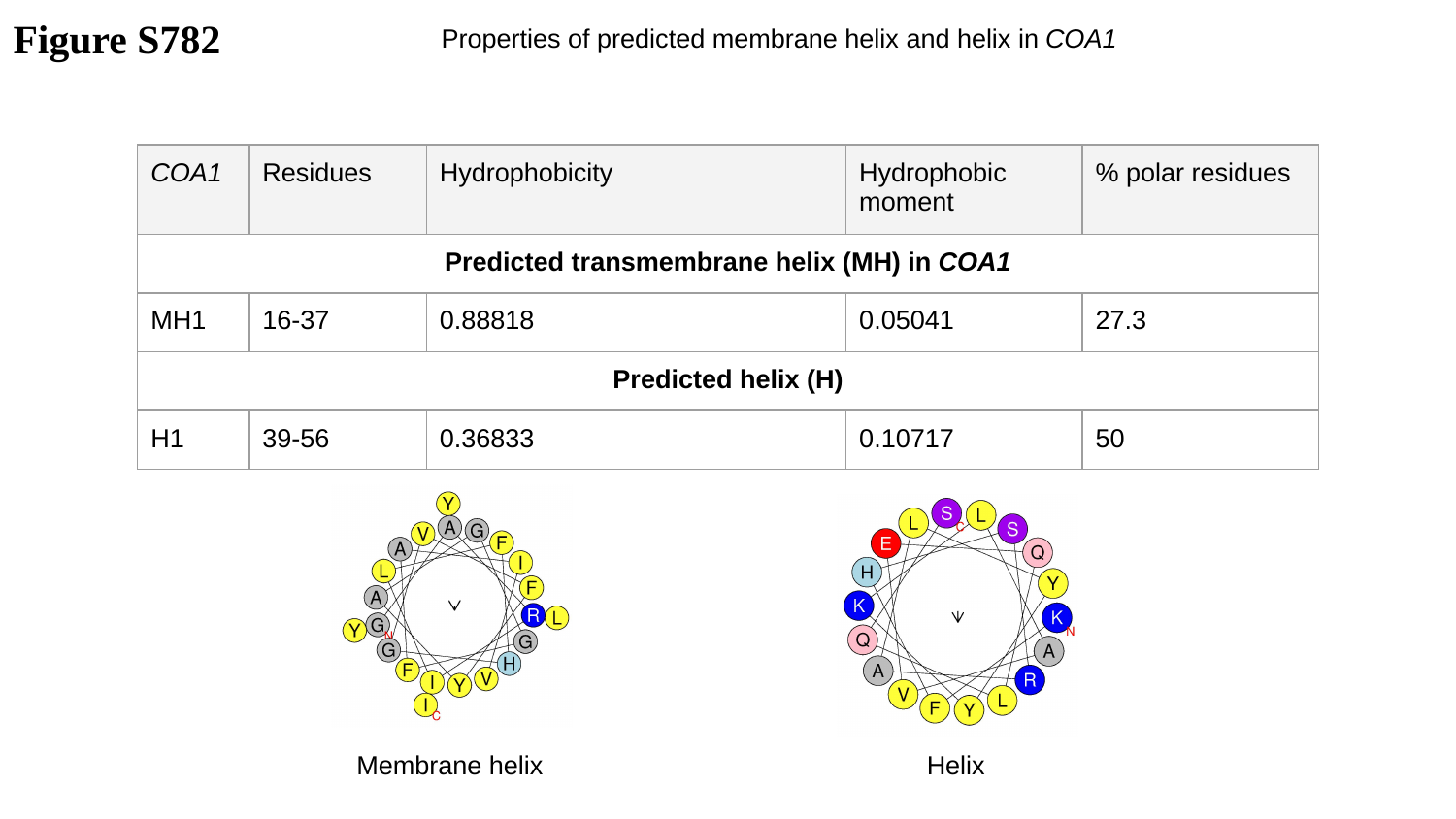

Figure S782
Properties of predicted membrane helix and helix in COA1
| COA1 | Residues | Hydrophobicity | Hydrophobic moment | % polar residues |
| --- | --- | --- | --- | --- |
| Predicted transmembrane helix (MH) in COA1 | | | | |
| MH1 | 16-37 | 0.88818 | 0.05041 | 27.3 |
| Predicted helix (H) | | | | |
| H1 | 39-56 | 0.36833 | 0.10717 | 50 |
Helix
Membrane helix

### Slide 783
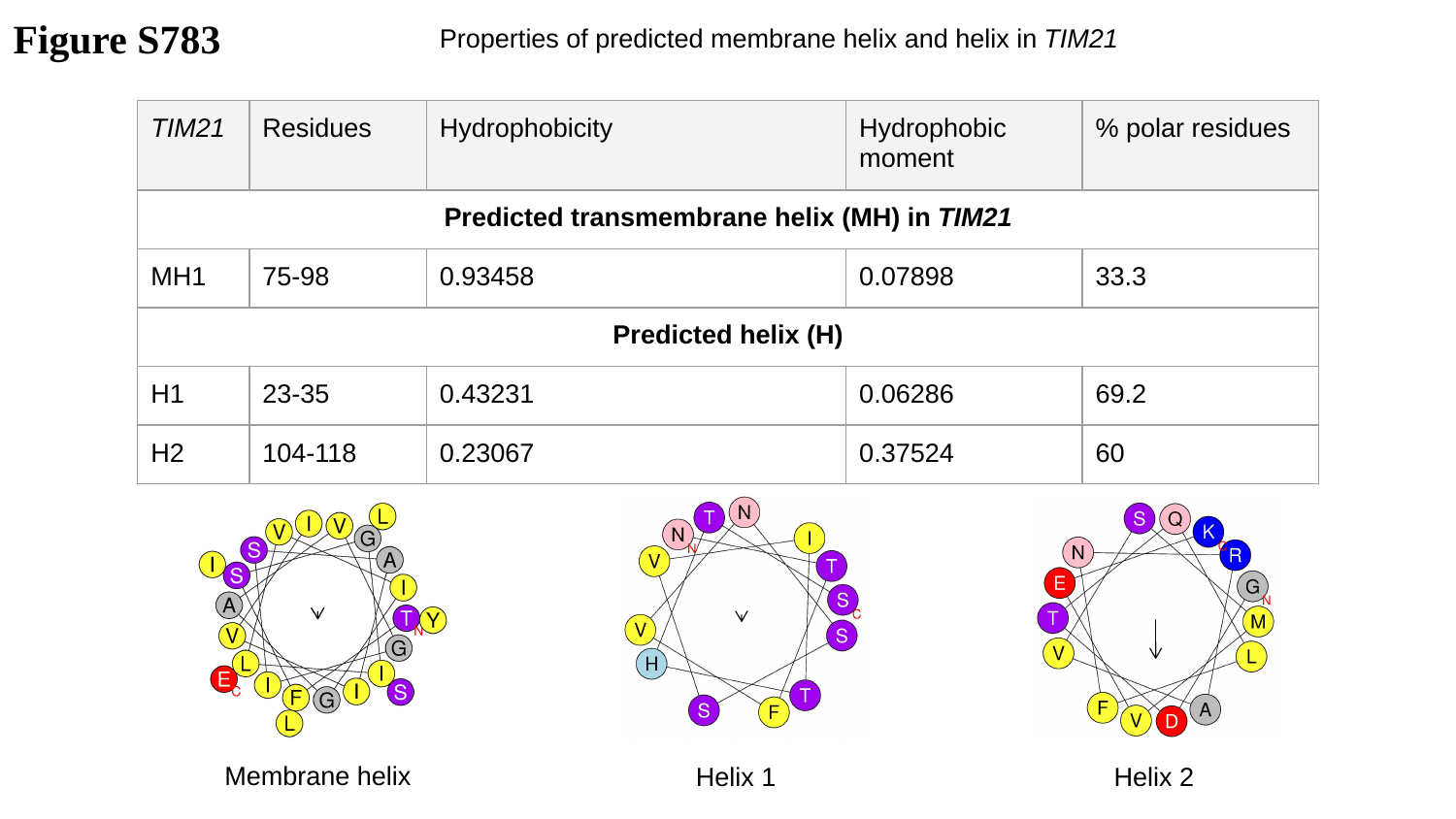

Figure S783
Properties of predicted membrane helix and helix in TIM21
| TIM21 | Residues | Hydrophobicity | Hydrophobic moment | % polar residues |
| --- | --- | --- | --- | --- |
| Predicted transmembrane helix (MH) in TIM21 | | | | |
| MH1 | 75-98 | 0.93458 | 0.07898 | 33.3 |
| Predicted helix (H) | | | | |
| H1 | 23-35 | 0.43231 | 0.06286 | 69.2 |
| H2 | 104-118 | 0.23067 | 0.37524 | 60 |
Membrane helix
Helix 1
Helix 2

### Slide 784
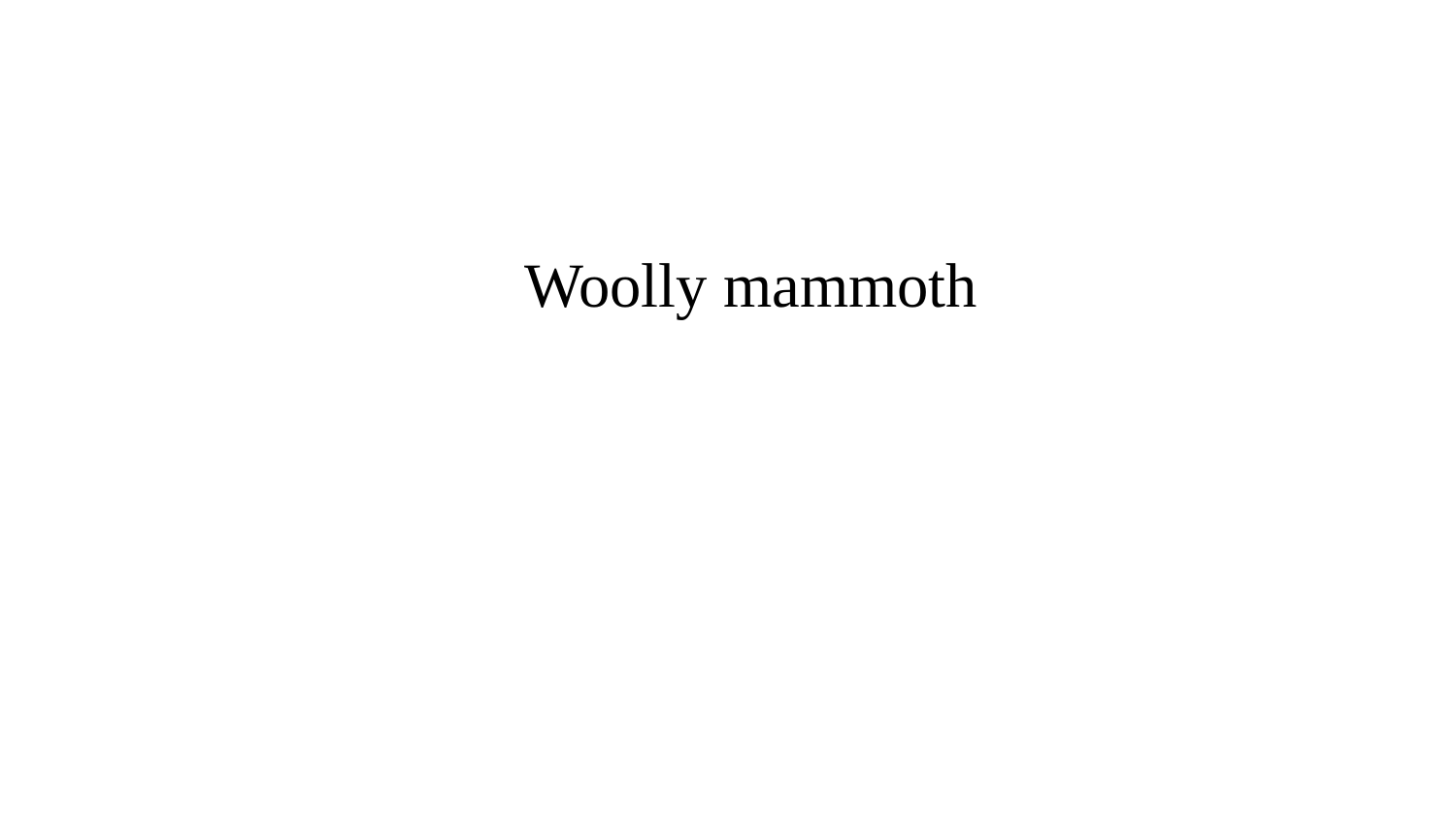

Woolly mammoth

### Slide 785
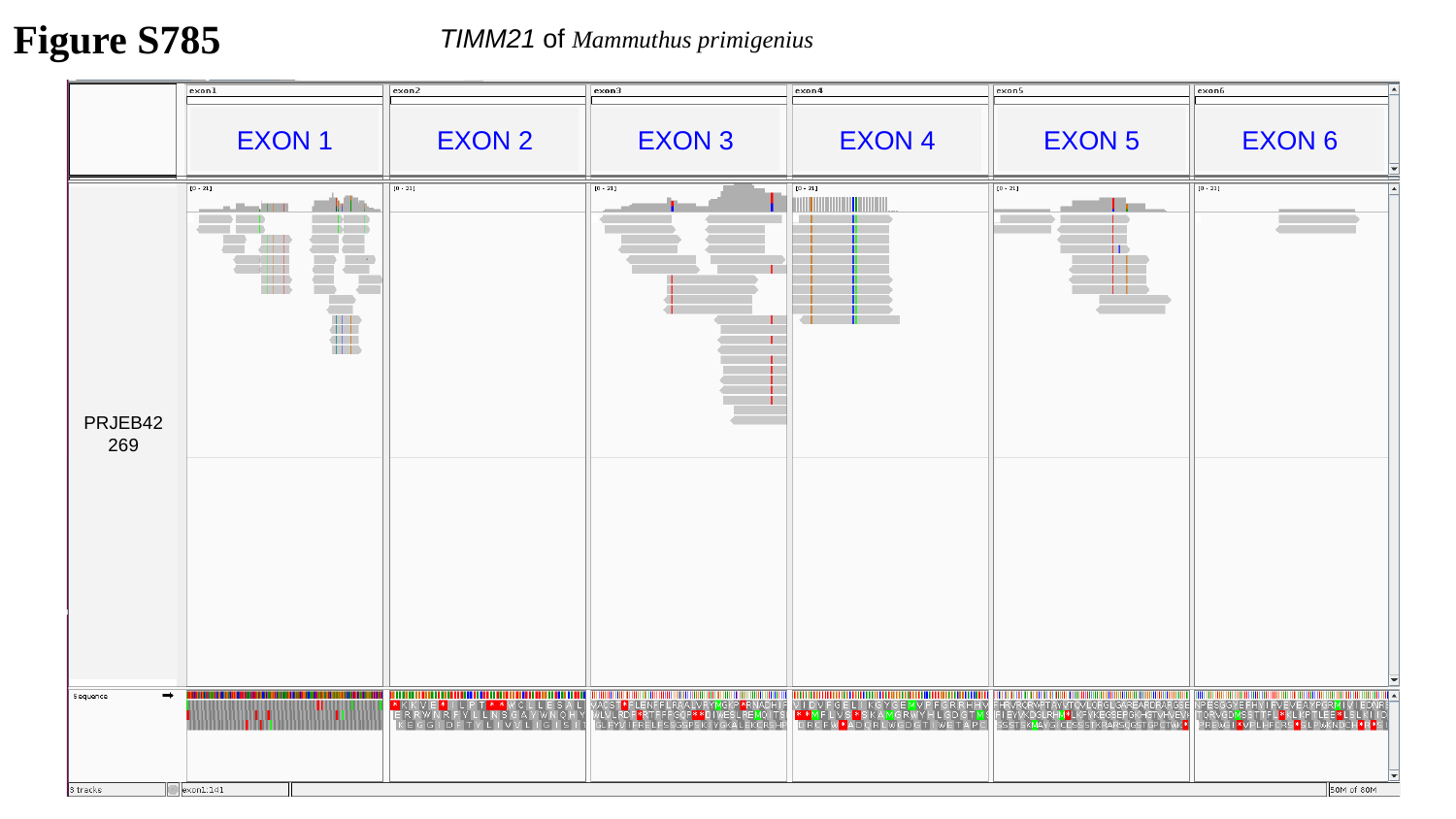

Figure S785
TIMM21 of Mammuthus primigenius
EXON 1
EXON 2
EXON 3
EXON 4
EXON 5
EXON 6
PRJEB42269

### Slide 786
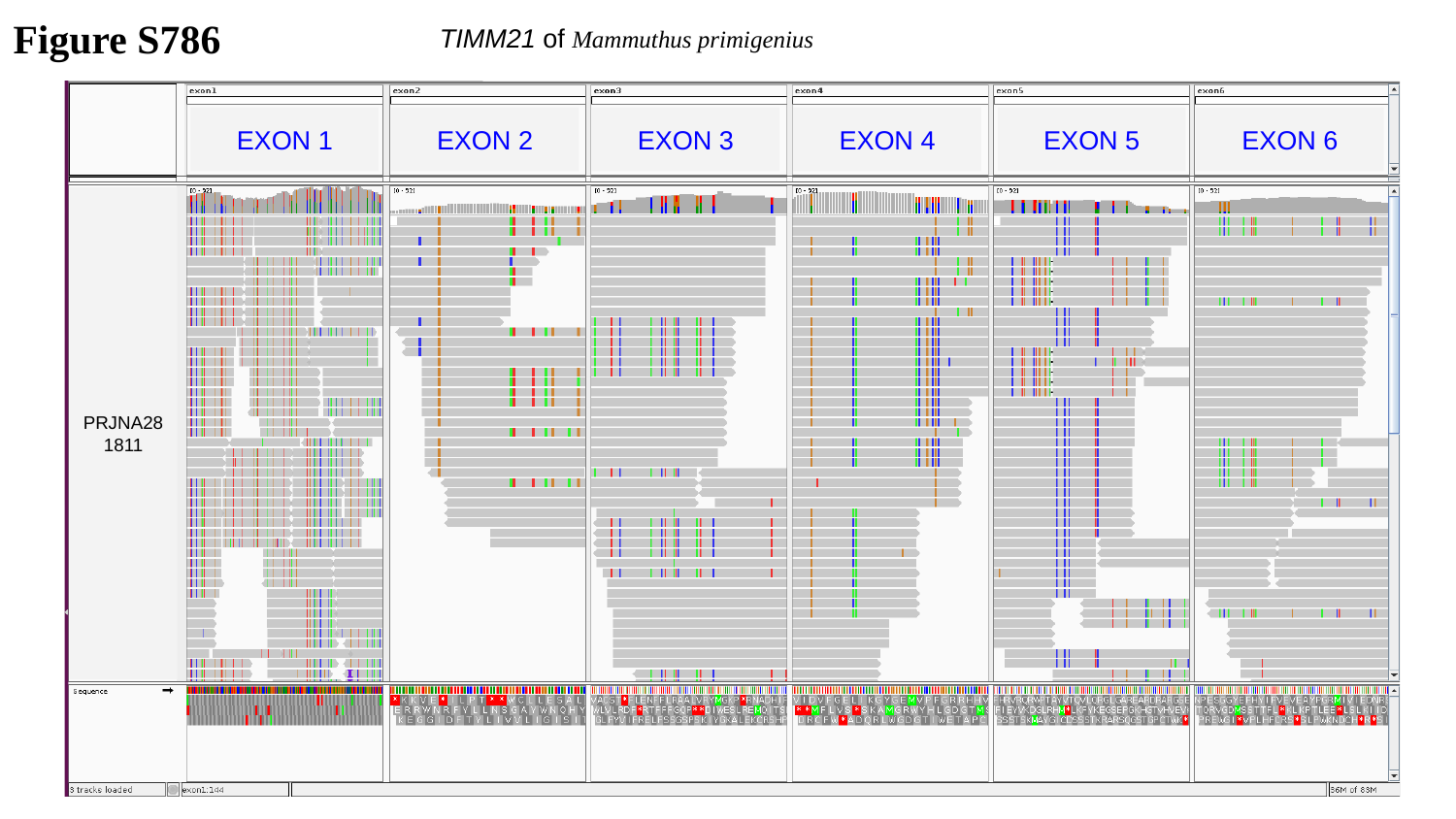

Figure S786
TIMM21 of Mammuthus primigenius
EXON 1
EXON 2
EXON 3
EXON 4
EXON 5
EXON 6
PRJNA281811

### Slide 787
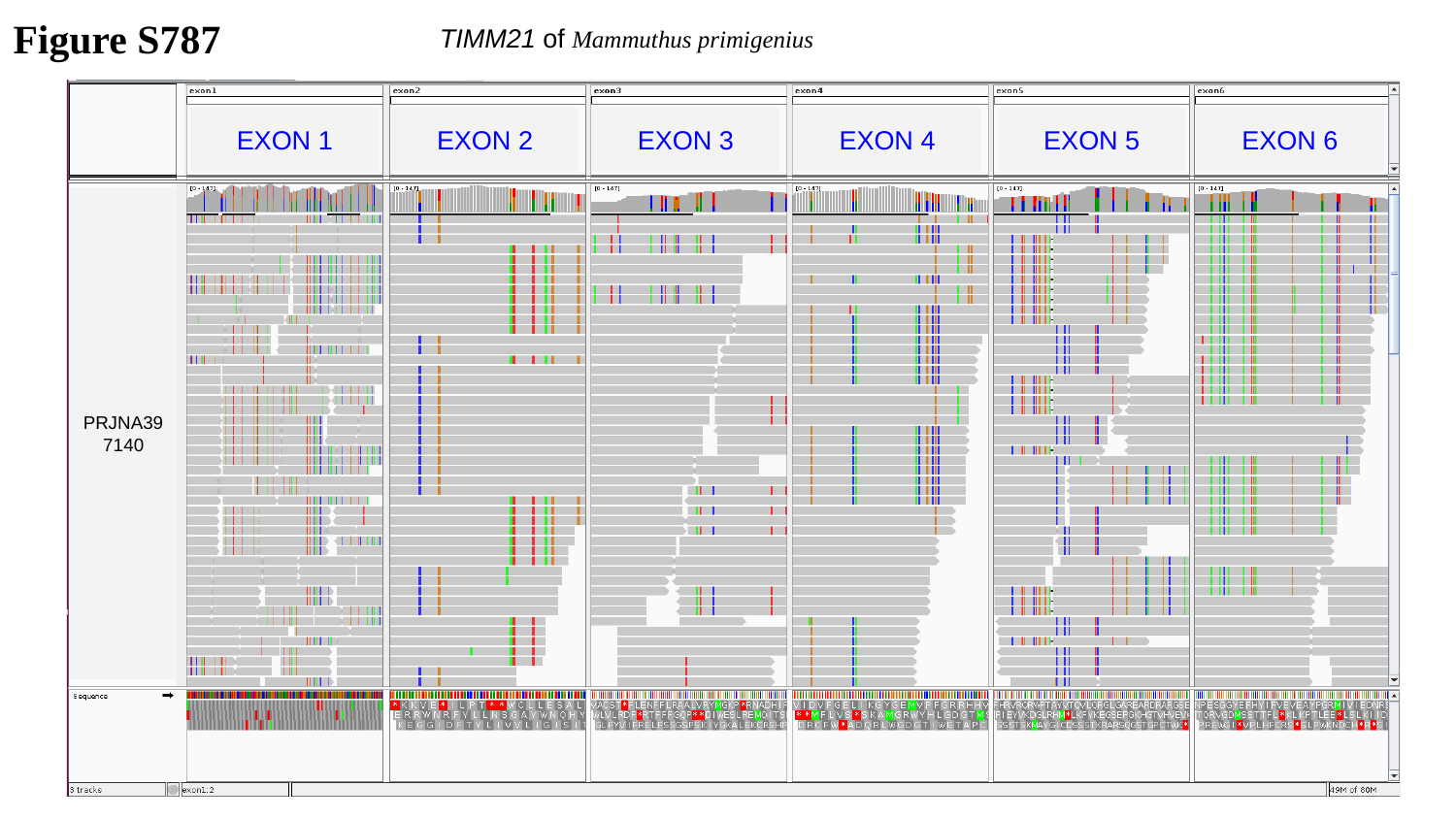

Figure S787
TIMM21 of Mammuthus primigenius
EXON 1
EXON 2
EXON 3
EXON 4
EXON 5
EXON 6
PRJNA397140

### Slide 788
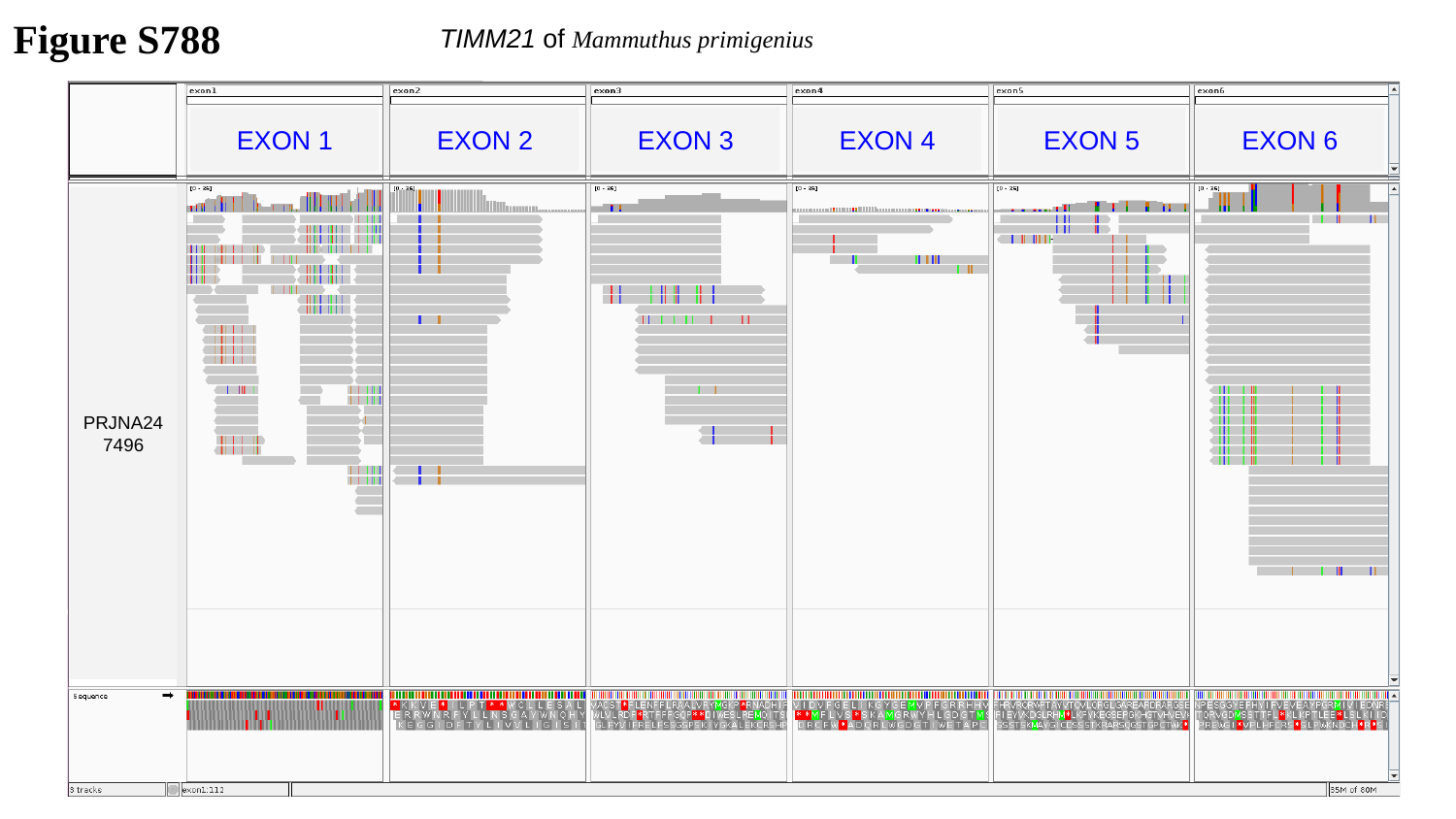

Figure S788
TIMM21 of Mammuthus primigenius
EXON 1
EXON 2
EXON 3
EXON 4
EXON 5
EXON 6
PRJNA247496

### Slide 789
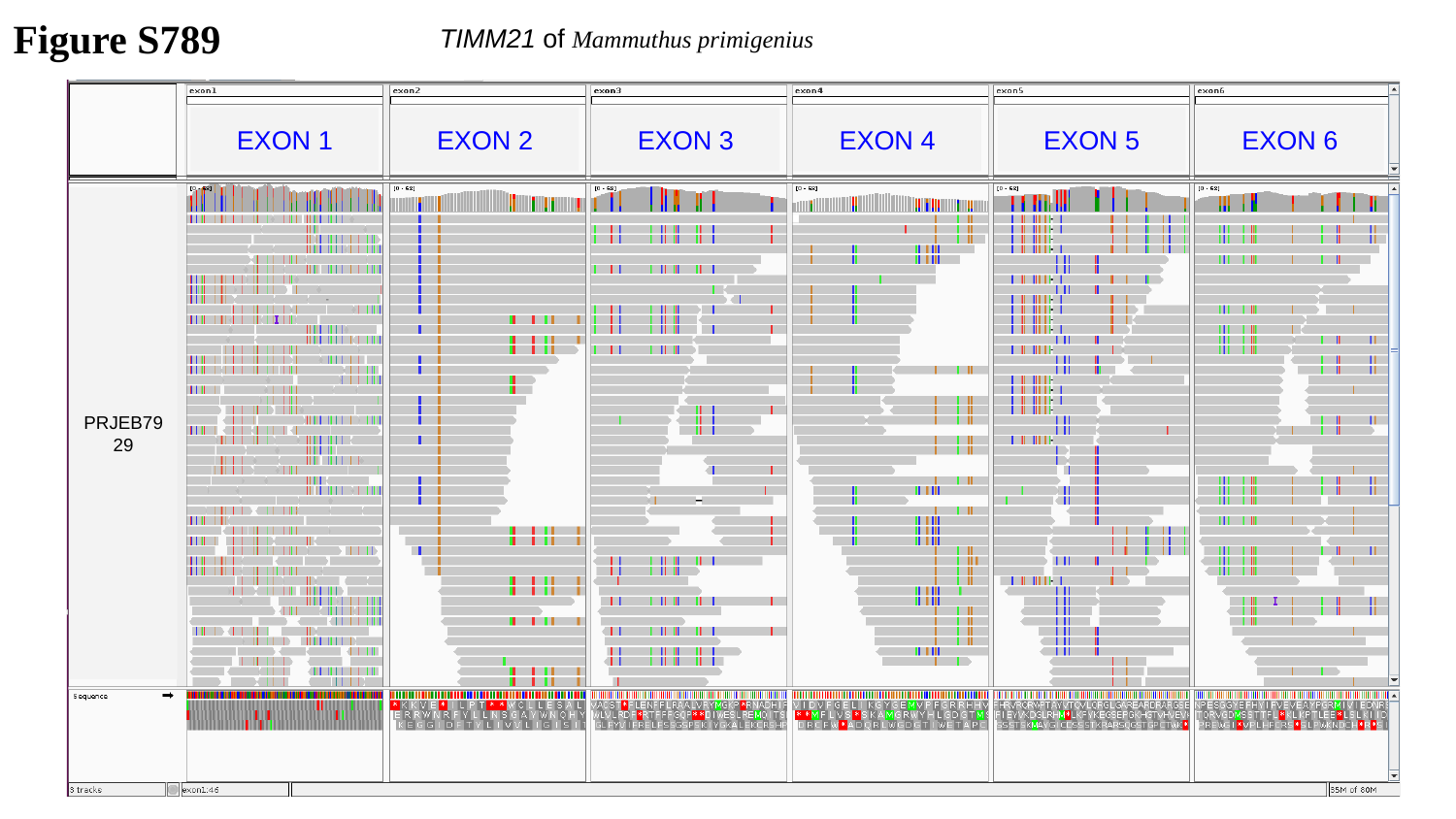

Figure S789
TIMM21 of Mammuthus primigenius
EXON 1
EXON 2
EXON 3
EXON 4
EXON 5
EXON 6
PRJEB7929

### Slide 790
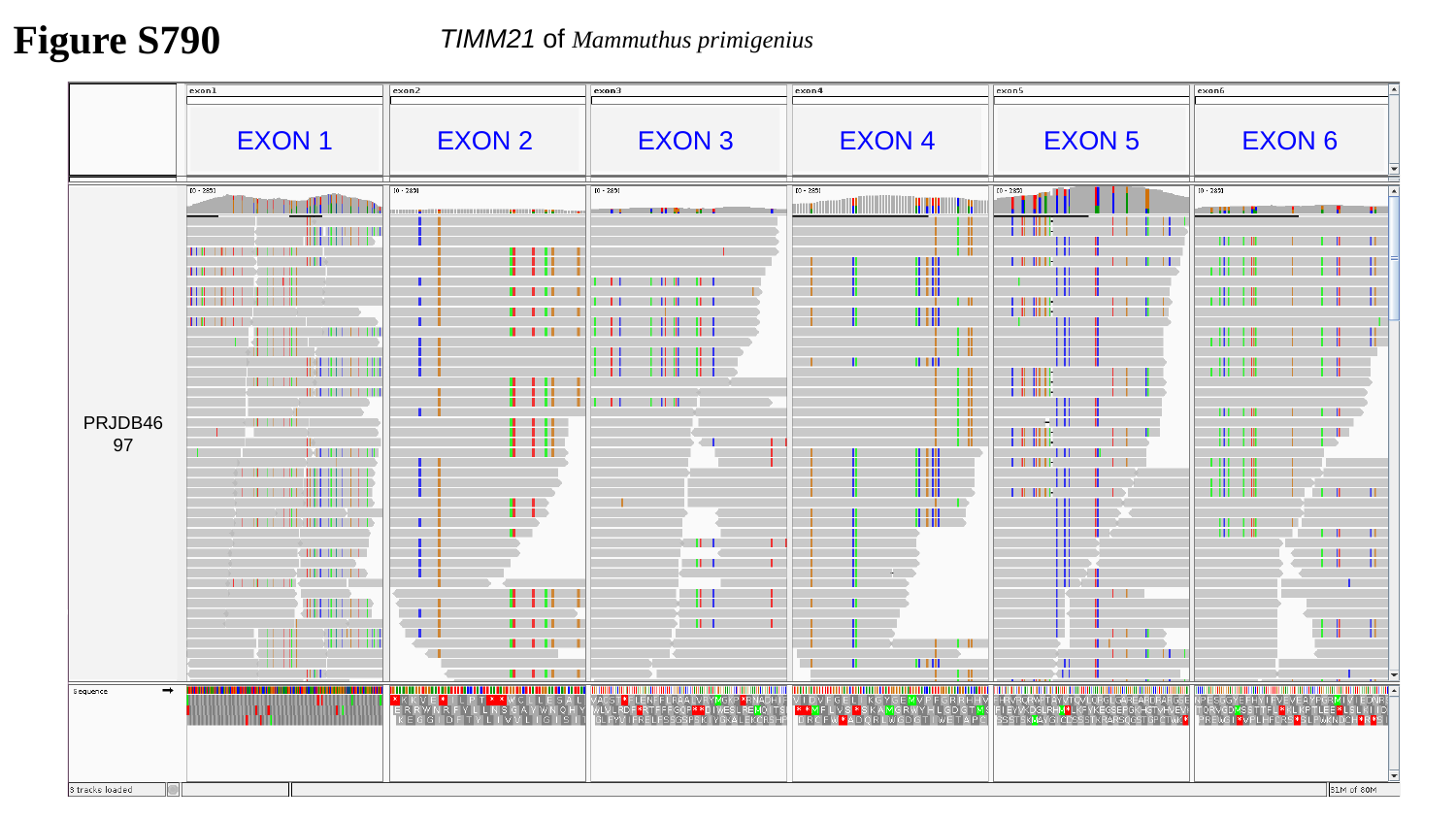

Figure S790
TIMM21 of Mammuthus primigenius
EXON 1
EXON 2
EXON 3
EXON 4
EXON 5
EXON 6
PRJDB4697

### Slide 791
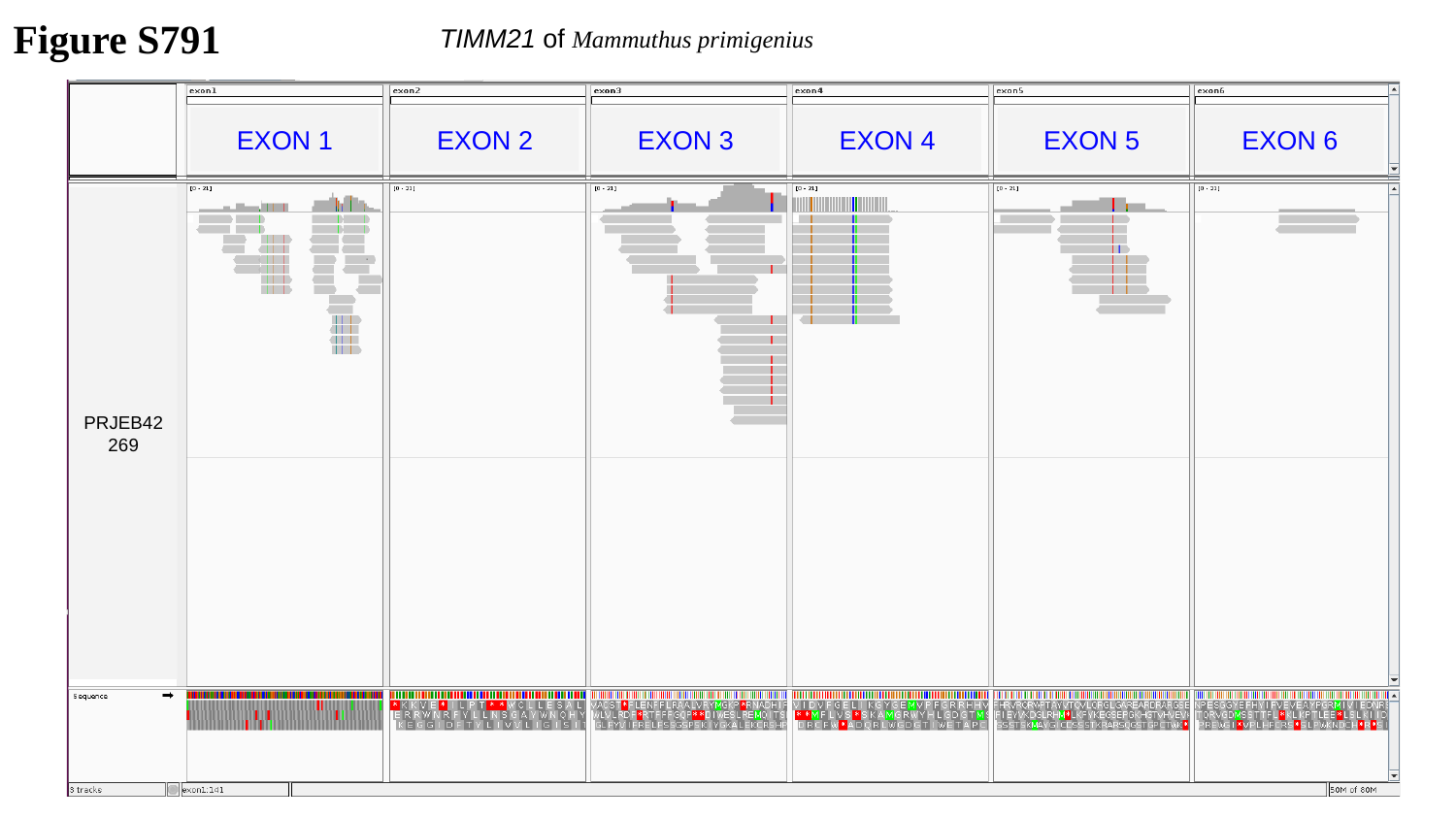

Figure S791
TIMM21 of Mammuthus primigenius
EXON 1
EXON 2
EXON 3
EXON 4
EXON 5
EXON 6
PRJEB42269

### Slide 792
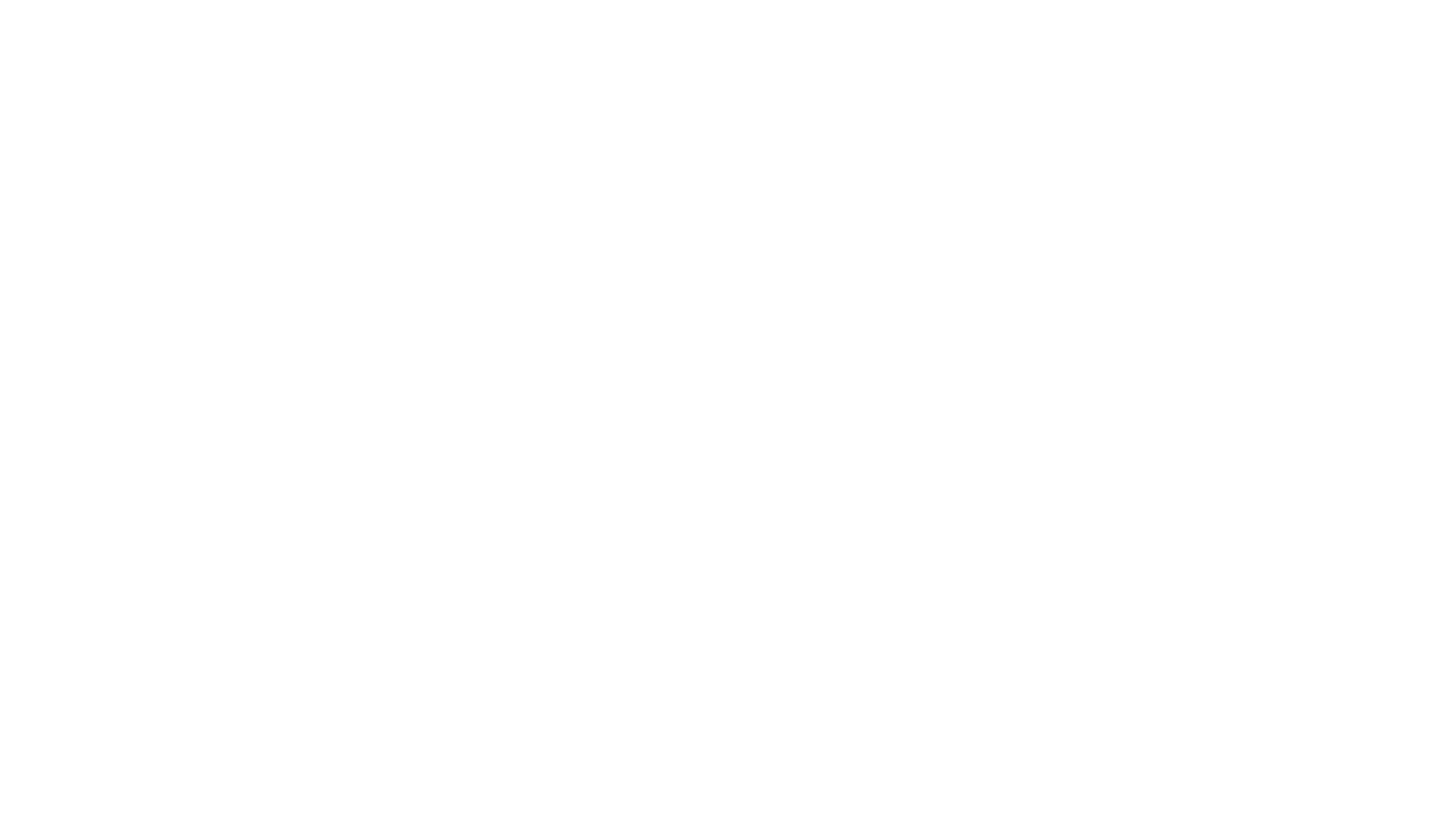
